# RNA m^6^A and 5hmC regulate monocyte and macrophage gene expression programs

**DOI:** 10.1101/2022.11.03.514952

**Authors:** Natalia Pinello, Renhua Song, Quintin Lee, Emilie Calonne, Kun-Long Duan, Emilie Wong, Jessica Tieng, Majid Mehravar, Bowen Rong, Fei Lan, Ben Roediger, Cheng-Jie Ma, Bi-Feng Yuan, John E J Rasko, Mark Larance, Dan Ye, François Fuks, Justin J. -L. Wong

## Abstract

**Background:** RNA modifications are essential for the establishment of cellular identity. Although increasing evidence indicates that RNA modifications regulate the innate immune response, their role in monocyte-to-macrophage differentiation and polarisation is unclear. To date, most studies have focused on m^6^A, while other RNA modifications, including 5hmC, remain poorly characterised. The interplay between different RNA modifications that may occur in specific cellular contexts remains similarly unexplored.

**Results:** We profiled m^6^A and 5hmC epitranscriptomes, transcriptomes, translatomes and proteomes of monocytes and macrophages at rest and pro- and anti-inflammatory states. We observed that decreased expression of m^6^A and 5hmC writers, METTL3 and TET-enzymes respectively, facilitated monocyte-to-macrophage differentiation. Despite a global trend of m^6^A and 5hmC loss during macrophage differentiation, enrichment of m^6^A and/or 5hmC on specific categories of transcripts essential for macrophage differentiation positively correlated with their expression and translation. m^6^A and 5hmC mark and are associated with the expression of transcripts with critical functions in pro- and anti-inflammatory macrophages. Notably, we also discovered the coexistence of m^6^A and 5hmC marking alternatively-spliced isoforms and/or opposing ends of the untranslated regions (UTR) of transcripts with key roles in macrophage biology. In specific examples, RNA 5hmC controls the decay of transcripts independently of m^6^A.

**Conclusions:** This study: i) uncovers m^6^A, 5hmC and their writer enzymes as regulators of monocyte and macrophage gene expression programs and ii) provides a comprehensive dataset to interrogate the role of RNA modifications in a plastic system. Altogether, this work sheds light on the role of RNA modifications as central regulators of effector cells in innate immunity.

## Background

RNA modifications modulate RNA metabolism and influence major biological processes, including splicing (1), RNA nuclear export (2), RNA decay (3) and translation (4). More than 180 RNA modifications have been identified so far mainly in tRNA and rRNA, and through recognition by effector RNA binding proteins, many of them are pivotal for cellular differentiation, maintenance of cellular identity, and function (5,6).

N-6 methyladenosine (m^6^A), the most abundant internal RNA modification on messenger RNAs (mRNAs), is widespread across mammalian transcriptomes (7,8). m^6^A deposition is catalysed by a methyltransferase (or ‘writer’) complex composed of a catalytic subunit formed by METTL3 and METTL14 (9,10) and several auxiliary proteins, including VIRMA, WTAP, CBLL1, ZC3H13 and RBM15 (11–14). Reversal of m^6^A methylation is performed by either of the two known m^6^A ‘eraser’ enzymes, FTO and ALKBH5 (15,16). RNA binding proteins known as m^6^A ‘readers’ recognise m^6^A-methylated RNAs to maintain key cellular processes and modulate response to environmental cues (17). m^6^A is now established as an essential player in physiological processes including development (18), maintenance of cellular identity (19) and immune response (20–22). Altered m^6^A patterns resulting from aberrant expression of m^6^A regulatory proteins have been linked to the development of human diseases, including cancer (23), cardiovascular disease (24) and autoimmune disorders (25).

5-hydroxymethylcytosine (5hmC) is one of the least studied RNA modifications. As with DNA, RNA 5hmC is generated by oxidation of 5mC by the Ten-eleven translocation proteins (TET1, TET2 and TET3) (26–28). In mammals, RNA 5hmC has been implicated in infection-induced myelopoiesis (29), degradation of transcripts derived from endogenous retroviruses (30) and stabilisation of pluripotency-promoting transcripts (27). In contrast to m^6^A, which has been extensively mapped across different tissues, cell types and pathological conditions (31,32), few studies have explored the roles of RNA 5hmC in normal physiology and disease states (27,28). Recent studies have revealed overlap between the functions of RNA m^6^A and 5hmC, including the regulation of RNA stability and translation (3,4,27,28). However, the interplay between m^6^A and 5hmC in the regulation of biological processes remains unknown.

Monocytes and macrophages are mononuclear phagocytic cells central to innate immunity (33). During inflammation, monocytes are recruited to sites of injury or infection where they differentiate into macrophages (34). Macrophages are remarkably plastic cells with roles in development, homeostasis, tissue repair and immunity (35). These cells rapidly alter their physiology in response to diverse environmental stimuli and give rise to morphologically and functionally diverse macrophage subpopulations with pro- and anti-inflammatory roles. Pro-inflammatory macrophages are the first line of defence against intracellular pathogens, whereas anti-inflammatory macrophages are crucial for wound healing and resolution of inflammation (36–38).

The roles of m^6^A methylation during monocyte differentiation into macrophages and subsequent polarisation of macrophages into pro- or anti-inflammatory cells remain elusive. Two recent studies reported the role of m^6^A in macrophage polarisation via exposure of METTL3-depleted mouse bone marrow-derived macrophages (BMDMs) to pro-inflammatory stimuli. However, one study reported that METTL3 depletion upregulates the expression of pro-inflammatory cytokines, including *TNF* and *IL-6* (39), whereas the other reported a decrease in these cytokines in METTL3-depleted BMDMs (40). Thus, additional and alternative experimental approaches are required to clarify the roles of METTL3 and m^6^A methylation in macrophage biology. Transcriptome-wide distribution of 5hmC, let alone its function in monocytes and macrophages, has not been explored.

Due to macrophage heterogeneity and the difficulty in capturing the complexity observed *in vivo*, generalised *in vitro* models of pro- and anti-inflammatory macrophages have been developed. A widely used approach is to use the monocytic cell line THP-1. Several studies by us and others have demonstrated that different populations of macrophages derived from this model exhibit similarity in both molecular and immunological signatures to those in primary human monocyte-derived macrophage populations (41–44), providing a suitable system to study RNA modifications in macrophage biology.

Here, we obtained transcriptome-wide RNA m^6^A and 5hmC distributions coupled with gene expression, polyribosome and proteomic profiles from THP-1 monocytic cells and THP-1-derived macrophages at rest and pro- and anti-inflammatory states (Fig. 1A). Through this multi-omics approach, we present data that serves as a reference set for studies on the regulatory roles of RNA m^6^A and 5hmC in shaping the transcriptomes and proteomes of these cells. Our work reveals METTL3 and TET proteins as key regulators of monocyte-to-macrophage differentiation, as well as several transcripts for which gene expression is regulated by m^6^A and/or 5hmC RNA modifications in a cellular context-specific manner. We report a positive association between RNA 5hmC and m^6^A with gene expression and translation. We observe the coexistence of m^6^A and 5hmC RNA that may regulate the expression, stability and alternative splicing of transcripts with crucial roles in macrophage biology. Altogether, our work provides new insights into the complex epitranscriptomic processes regulating effector cells of the innate immune system.

**Figure 1.**
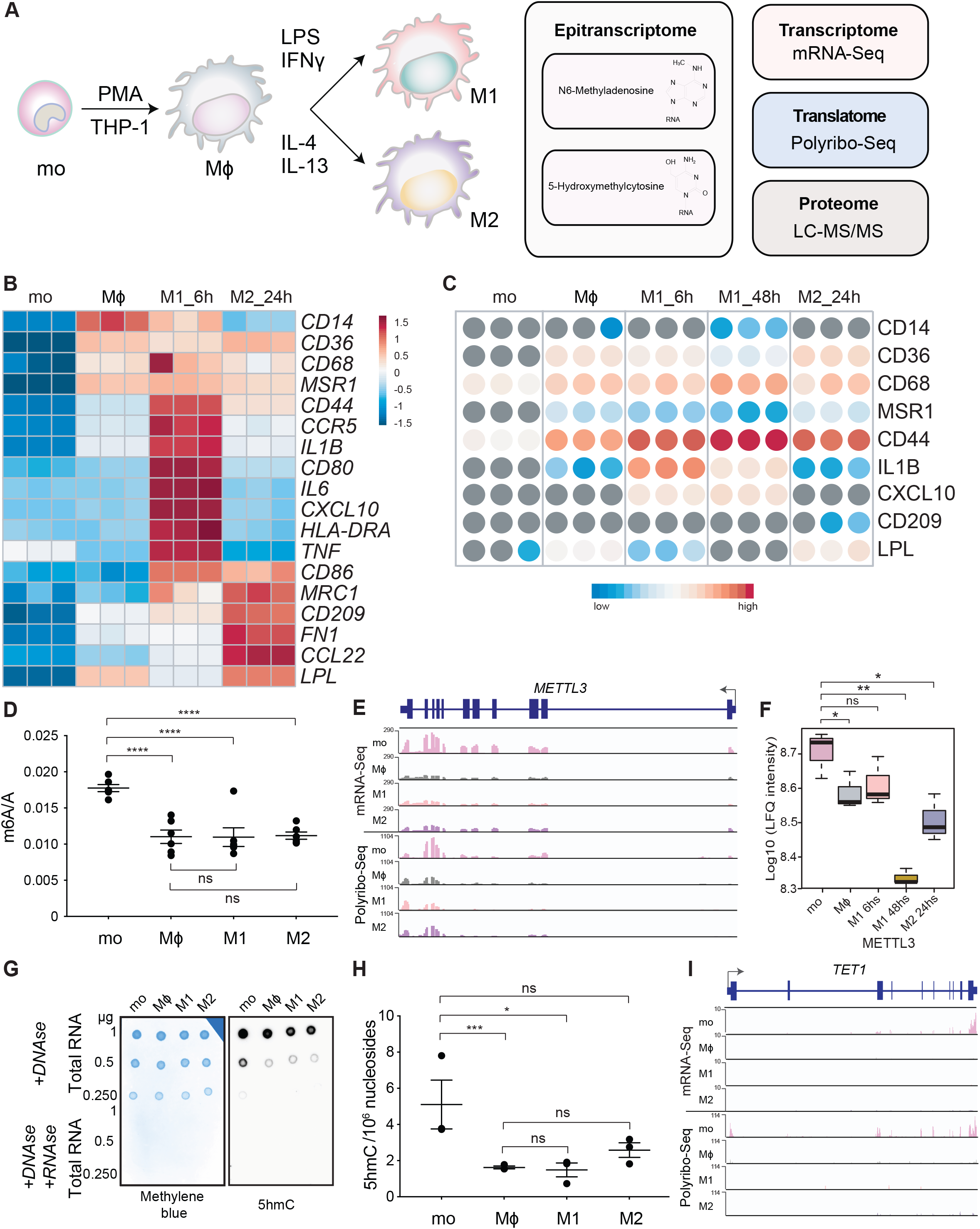
RNA m^6^A and 5hmC levels decrease during monocyte-to-macrophage differentiation. (**A**) Diagram detailing multi-omics approach to study m^6^A and 5hmC RNA modifications in the THP-1 macrophage differentiation and polarisation model. Heatmaps displaying (**B**) differential gene expression (z score) and (**C**) protein levels of monocytes (mo), resting-like (Mϕ), pro-inflammatory (M1) and anti-inflammatory macrophages (M2) based on established markers. h indicates hours of polarisation. (**D**) m^6^A/A on polyadenylated RNAs measured by LC-MS/MS. (**E**) *METTL3* coverage plots showing reads per kilobase per million reads (RPKM) for mRNA-Seq (top) and Polyribo-Seq (bottom) data from mo, Mϕ, M1 and M2. (**F**) METTL3 protein abundance as measured by LC-MS/MS. (**G**) Dot blot performed with total RNA from mo, Mϕ, M1 and M2 cells probed with anti-5hmC antibody (right) and methylene blue loading control (left). Samples were treated with DNase to eliminate the possibility of detecting 5hmC on DNA. Negative controls were treated with both DNase and RNase A. (**H**) 5hmC/10^6 nucleosides in total RNA measured by LC-MS/MS. (**I**) *TET1* coverage plots showing reads (RPKM) for mRNA-Seq (top) and Polyribo-Seq (bottom) data from mo, Mϕ, M1 and M2. All data are from at least three independent experiments. In **D** and **H**, bar plots show mean ± SEM. In **F**, box and whisker plots show median and 1.5 times the interquartile range. An unpaired two-tailed Student’s t-test was used to determine significance, denoted by ns, not significant; *, *p* <0.05; **, *p* <0.01; ***, *p* <0.001 and ****, *p* <0.0001.

## Results

### m^6^A and 5hmC levels on mRNA decrease during monocyte-to-macrophage differentiation

To study m^6^A and 5hmC RNA modifications during monocyte-to-macrophage differentiation and polarisation, we differentiated and polarised THP-1 monocytic cells into four cellular stages generated as follows: Untreated THP-1 cells (mo); Phorbol 12-myristate 13-acetate (PMA)-differentiated cells (Mϕ); PMA-differentiated cells plus lipopolysaccharide (LPS) and interferon-γ (IFNγ) (M1); and PMA-differentiated cells plus IL-4 and IL-13 (M2) (Fig. 1A).

We performed deep mRNA-Seq and liquid chromatography with tandem mass spectrometry (LC-MS/MS) in mo, Mϕ, M1 and M2. Differences in global gene and protein expression patterns were evident by Principal Component Analysis (PCA) of mRNA-Seq (Additional file 1: Fig. S1A) and LC-MS/MS (Additional file 1: Fig. S1B) data. For M1, LC-MS/MS was performed at 6 and 48 hours post-stimulation to capture dynamic proteomic changes associated with macrophage reprogramming in response to LPS and IFNγ stimulation (45). This analysis showed a clear separation of monocytes from macrophages and highlighted M1 as a highly distinct subset within our modelled macrophages. As expected, we observed clustering of technical replicates within each cellular stage. We further confirmed the identities of the four cell types via gene expression analysis (Fig. 1B) and proteomics profiling (Fig. 1C) of established cell type-specific markers (41,42,46–50). Overall, our data indicate that the THP-1 macrophage differentiation and polarisation model is robust in our hands. Importantly, these transcriptomic and proteomic profiles resemble, at least in part, those of human primary monocytes and macrophages (51). Therefore, we concluded this system was suitable to study molecular modulation of monocytes and macrophages’ functions *in vitro*.

To investigate changes in global m^6^A levels during monocyte-to-macrophage differentiation and polarisation, we quantified m^6^A on polyadenylated mRNAs from mo, Mϕ, M1 and M2 cells by LC-MS/MS. We found that m^6^A levels on mRNA decreased during the transition of monocytes to resting macrophages and remained unchanged when macrophages were treated with polarising stimuli (Fig. 1D). This observation was consistent with a significant decrease in *METTL3* expression revealed by mRNA-Seq and the relatively lower abundance of *METTL3* transcripts detected in the polysome fractions of macrophages compared to monocytes (Fig. 1E). METTL3 protein levels were also lower in macrophages compared to monocytes (Fig. 1F). Protein levels for other components of the m^6^A writers complex (METTL14, VIRMA, ZC3H13, WTAP and RBM15) were also diminished in macrophages compared to monocytes (Additional file 1: Fig. S1C and S1D).

To investigate global RNA 5hmC changes during macrophage differentiation and polarisation, we first performed dot blots. We found that total RNA 5hmC levels decreased in macrophages compared to monocytes (Fig. 1G). This decrease is likely due to changes in 5hmC levels on mRNA, as 5hmC signal was markedly increased on these RNA species compared to total, ribosomal (rRNA) or small (sRNA) fractions (Additional file 1: Fig. S1D). No 5hmC signal was detected on rRNA or sRNA by dot blot (Additional file 1: Fig. S1D). Higher global 5hmC levels in total RNA of monocytes compared to macrophages was further confirmed using LC-MS/MS (Fig. 1H). Similar to m^6^A, global 5hmC levels significantly decreased during monocyte-to-macrophage differentiation, and no significant difference was observed during polarisation. Based on mRNA-Seq data, *TET1* expression diminished during monocyte-to-macrophage differentiation and our polysome profiling data similarly inferred lower *TET1* translation in macrophages compared to monocytes (Fig. 1I). *TET2* and *TET3* mRNA enrichment showed no major difference in the polysome fraction of macrophages compared to monocytes (Additional file 1: Fig. S1E and S1F). Collectively, these data demonstrate that TET1, but not TET2 or TET3, expression correlated best with global RNA 5hmC levels. We therefore speculate that TET1 may be the primary mediator of RNA 5-hydroxymethylation during monocyte-to-macrophage differentiation.

In summary, global m^6^A and 5hmC RNA levels decrease during monocyte-to-macrophage differentiation and remain stable following exposure to pro- and anti-inflammatory stimuli. Diminished expression of m^6^A and 5hmC writers, including *METTL3* and *TET1* respectively, may explain the reduction in m^6^A and 5hmC levels detected on RNA during monocyte-to-macrophage differentiation.

### METTL3 regulates monocyte-to-macrophage differentiation and polarisation

To determine the role of METTL3 during monocyte-to-macrophage differentiation *in vitro*, we depleted *METTL3* in THP-1 cells using short hairpin RNAs (Fig. 2A). As expected, depletion of *METTL3* led to a significant decrease in global m^6^A levels on mRNA (Fig 2B). Following METTL3 depletion, THP-1 cells (mo) exhibited macrophage-like morphology, including adherence to the bottom of the culture flask, enlargement of the cell body and amoeboid appearance (Fig. 2C). Additionally, METTL3-depleted cells expressed higher levels of the macrophage-associated cell surface markers CD11b and CD44 (Fig. 2D). However, their expression levels were markedly lower (3- to 4-fold less in mean fluorescence intensity) than PMA-differentiated cells, indicating that the spontaneous differentiation into macrophages facilitated by METTL3 depletion was incomplete. We detected no significant difference in the expression of M1 (CD80 and CD38) or M2 (CD209) cell surface markers by flow cytometry (Additional file 2: Fig. S2A), ruling out the possibility that METTL3 depletion promotes spontaneous macrophage polarisation.

**Figure 2.**
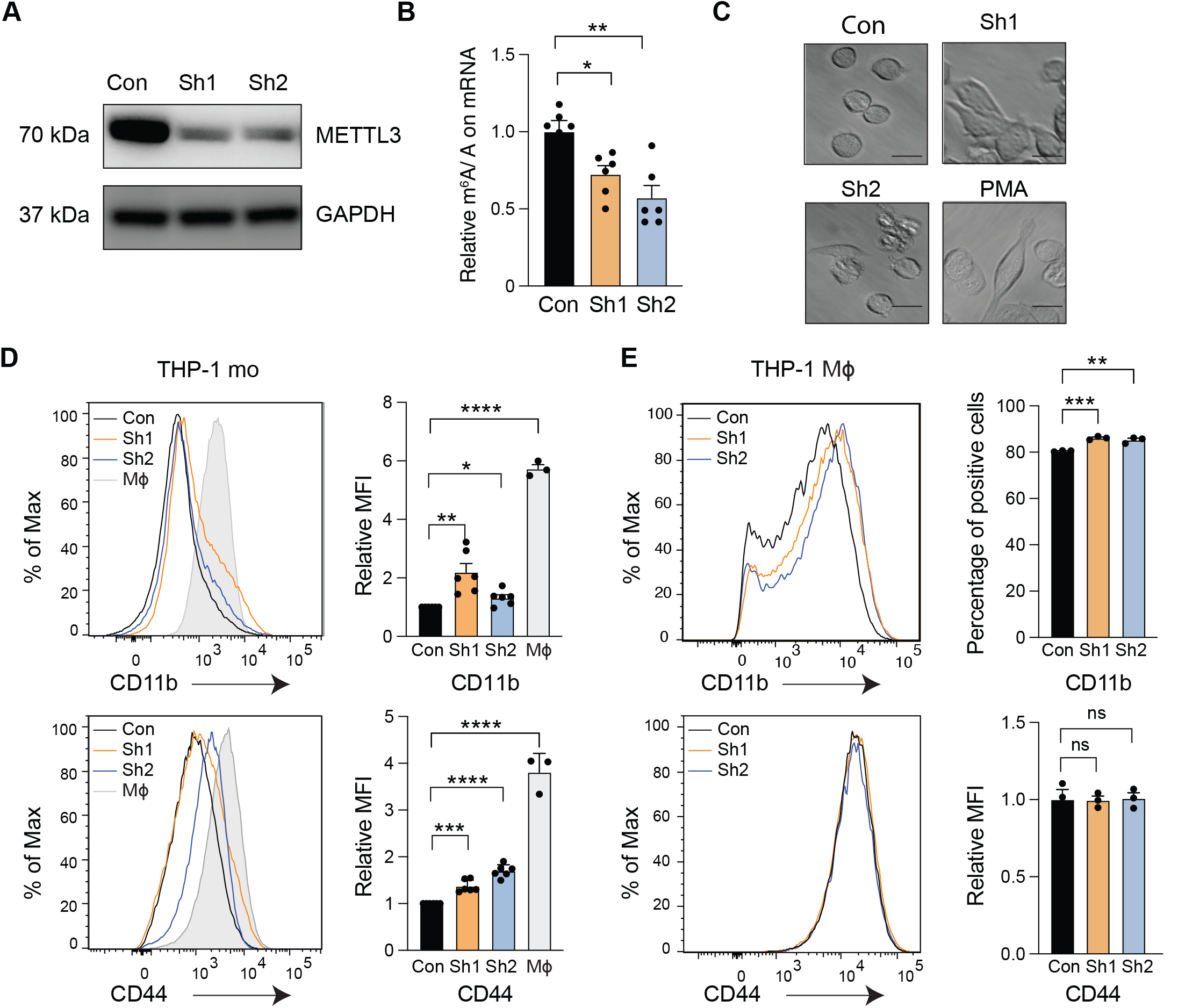
METTL3 depletion triggers monocyte-to-macrophage differentiation. (**A**) Western blot showing METTL3 and GAPDH protein levels following THP-1 transduction with control (Con) or METTL3-specific (Sh1 and Sh2) shRNAs. (**B**) Relative m^6^A/A in polyadenylated RNA from control and METTL3 depleted cells measured by LC-MS/MS. (**C**) Microscopy images showing morphological changes in THP-1 cells transduced with control and METTL3-targeting shRNAs or exposed to PMA (positive control). Flow cytometry profiles and quantification (Mean Fluorescence Intensity (MFI) or Percentage of Positive Cells) showing changes in the relative expression of the macrophage cell surface markers CD11b and CD44 following METTL3 depletion in (**D**) THP-1 monocytes (mo) or (**E**) THP-1-derived macrophages (Mϕ). Representative histograms from at least three independent experiments are shown. Bar plots show mean ± SEM. An unpaired two-tailed Student’s t-test was used to determine significance, denoted by ns, not significant; *, *p* <0.05; **, *p* <0.01; ***, *p* <0.001 and ****, *p* <0.0001.

Next, we explored whether METTL3 depletion influenced the trajectory of PMA-induced differentiation. In comparison with control cells, METTL3-depleted cells exhibited a slight increase in CD11b but no difference in surface CD44 (Fig. 2E) following PMA treatment, indicating that METTL3 depletion does not significantly enhance PMA-mediated differentiation. METTL3 depletion did not promote the polarisation of PMA-differentiated macrophages based on the lack of consistent increase in the levels of M1 (CD80 and CD38) and M2 (CD209) markers (Additional file 2: Fig. S2B).

We further explored the effect of METTL3-depletion on M1 and M2 polarisation triggered by specific stimuli. In METTL3-depleted macrophages stimulated with LPS and IFNγ, we observed a significant decrease in the levels of CD38 (Additional file 3: Fig S3A) and mRNA expression of other M1 markers, including *CXCL10, IL-6, CD80* and *TNF* measured by qRT-PCR (Additional file 3: Fig. S3B). METTL3-depleted macrophages stimulated with IL-4 and IL-13 exhibited decreased levels of CD209 (Additional file 3: Fig. S3C) and decreased mRNA expression of *CD209* and *CD206* (another M2 marker) compared to control cells (Additional file 3: Fig. S3D).

Our results suggest that METTL3 depletion in THP-1 monocytes stimulates transition towards a resting-like macrophage phenotype, and that METTL3 may facilitate M1 and M2 polarisation of THP-1-derived macrophages following exposure to pro- and anti-inflammatory stimuli, respectively.

### Changes in m^6^A levels are associated with altered transcription and translation of genes involved in monocyte-to-macrophage differentiation and polarisation

To determine transcripts that are modified by m^6^A in monocytes and macrophages, we performed m^6^A RNA immunoprecipitation sequencing (m^6^A-IP-Seq) in mo, Mϕ, M1 and M2 cells. We identified 8511–12425 m^6^A peaks (corresponding to 6118–7130 genes) in monocytes and macrophages (Additional file 4: Table S1), of which 4044 m^6^A peaks (corresponding to 4185 genes) were common to all four cell stages. 1866 m^6^A peaks (in 747 genes) were exclusive to monocytes and 651 m^6^A peaks (in 507 genes) were present in all types of macrophages (Fig. 3A). Overall m^6^A profiles obtained were consistent with previous studies (7,8), whereby half (~45%) of the m^6^A peaks identified in both monocytes and macrophages were present in the coding sequence (CDS) while approximately 30% were located in the 3’untranslated region (3’UTR) (Fig. 3B). Metagene profiles of all four cellular stages presented the characteristic m^6^A distribution along the mRNA transcript with a marked enrichment of this modification preceding the stop codon (Fig. 3C). As expected, sequence logo analysis revealed the canonical DRACH motif in all four data sets (Fig. 3C).

**Figure 3.**
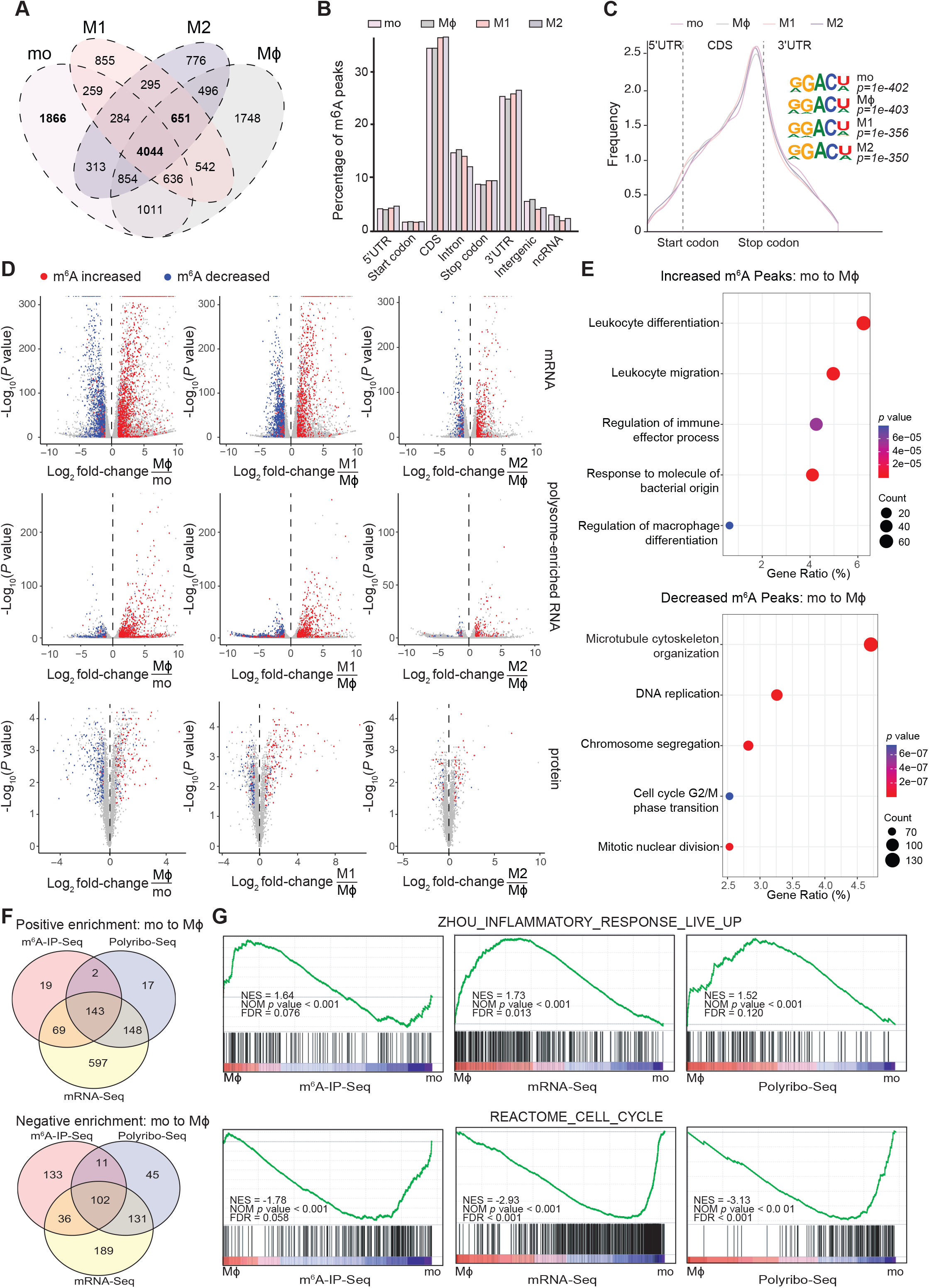
Changes in m^6^A levels are associated with altered transcription and translation of genes involved in monocyte-to-macrophage differentiation and polarisation. (**A**) Venn diagram showing the overlap of m^6^A peaks found in THP-1 monocytes (mo), resting-like (Mϕ), pro-inflammatory (M1) and anti-inflammatory macrophages (M2). (**B**) Percentage of m^6^A peaks identified by m^6^A-IP-Seq in defined transcript and genomic regions: coding sequence (CDS), 5’untranslated region (5’UTR), 3’untranslated region (3’UTR), start and stop codons, intergenic regions and non-coding RNA (ncRNA) in mo, Mϕ, M1 and M2. (**C**) Metagene profiles detailing m^6^A distribution along a normalised transcript and top enriched sequence motifs identified by m^6^A-IP-Seq in all cell types. (**D**) Volcano plots showing differential gene expression (mRNA, top), translation (polysome-enriched RNA, middle) and protein levels (bottom). Red and blue dots represent individual genes with increased and decreased m^6^A peaks respectively in Mϕ compared to mo, M1 compared to Mϕ and M2 compared to Mϕ. (**E**) Gene ontology analysis of increased (top) and decreased (bottom) m^6^A peaks during mo to Mϕ differentiation. (**F**) Venn diagram showing overlapping gene sets identified by Gene Set Enrichment Analysis (GSEA) on m^6^A-Seq, mRNA-Seq and polyribo-Seq data presenting positive (top) or negative (bottom) enrichment during mo to Mϕ differentiation. (**G**) GSEA signatures showing significantly positive (top) or negative (bottom) enrichment during mo to Mϕ differentiation.

By comparing m^6^A profiles to gene expression changes, we found that increased m^6^A methylation levels during macrophage differentiation and polarisation are predominantly associated with increased mRNA expression and *vice versa* (Fig. 3D). Similar positive associations were observed between transcriptome-wide m^6^A levels and protein translation as measured by polyribosome sequencing (Polyribo-Seq) and LC-MS/MS (Fig. 3D). These results support previous reports showing that m^6^A is coupled to transcription and/or promotes translational processes (52–55).

To identify a functional association between differential m^6^A modification and macrophage differentiation and/or polarisation, we performed Gene Ontology (GO) analysis on differential m^6^A peaks. While the majority of mRNAs presenting differential m^6^A methylation (n=6141) showed decreased m^6^A peaks in Mϕ compared to mo, ~30% of transcripts showed increased m^6^A peaks (n=1904) in Mϕ. mRNAs with increased m^6^A peaks in Mϕ compared to mo showed enrichment for functions associated with macrophage differentiation and immune response (Fig. 3E). Consistent with the cessation of cell proliferation typical of differentiated Mϕ (58), transcripts related to cell cycle, DNA replication and chromosome segregation were enriched in mRNAs with decreased m^6^A peaks (Fig. 3E).

GO analysis for transcripts showing increased m^6^A in M1 compared to Mϕ cells revealed a compelling enrichment for biological processes underlying pro-inflammatory responses, including the NF-kappa B signalling pathway, response to lipopolysaccharide and virus, pattern recognition and toll-like receptor signalling pathways, response to interferon gamma and tumour necrosis factor and regulation of pro-inflammatory cytokines (56–59) (Additional file 5: Fig. S4A). Transcripts associated with increased or decreased m^6^A during M2 polarisation were enriched in GO terms associated with homeostatic and anti-inflammatory functions, including tissue development, glucosaminoglycan metabolic processes, regulation of autophagy and vacuolar transport (60–63) (Additional file 5: Fig. S4B). These results indicate that m^6^A RNA methylation may have a central role in the regulation of critical pathways that control macrophage differentiation and polarisation.

Using gene set enrichment analysis (GSEA), we identified 143 overlapping gene sets representing transcripts with increased m^6^A, expression and translation levels in Mϕ compared to mo, and 102 overlapping gene sets showing the opposite pattern (Fig. 3F). In line with the GO analyses, the inflammatory response gene set was significantly enriched in Mϕ compared to mo, whereas the cell cycle regulation gene set showed significant negative enrichment (Fig. 3G). 153 gene sets representing transcripts with increased m^6^A, expression and translation levels were positively enriched in M1 compared to Mϕ, including sets associated with an inflammatory response induced by LPS and IFNγ (Additional file 5: Fig. S4C and S4D). Only two gene sets were enriched for transcripts with decreased m^6^A, expression and translation levels; one is also associated with downregulation of the inflammatory response (Additional file 5: Fig. S4C and S4D). When comparing M2 to Mϕ, we found no significantly enriched gene set in association with decreased m^6^A, gene expression or translation levels. However, 28 gene sets, including sets enriched for immune response and response to IL-4 stimulation genes, typical of M2 cells, were positively correlated with increased m^6^A, mRNA expression and translation levels (Additional file 5: Fig. S4E and S4F).

Collectively, our *in silico* analyses using multiple approaches indicate that m^6^A may regulate transcription and translation of transcripts that are directly involved in establishing the identity and function of macrophages.

### Modulation of m^6^A levels alters the expression and translation of critical genes that regulate macrophage differentiation and function

Coupling of m^6^A with transcriptional and translational changes occurs on critical genes involved in different states of macrophages (Fig. 3D, Fig. 4A and Additional file 6: Table S2), indicating that m^6^A may be a regulatory mechanism controlling their expression and function. Increased m^6^A on transcripts with essential roles in macrophage development, maintenance and phagocytic function, including *CSF1* (Colony Stimulating factor), *MSR1* (Macrophage Scavenger Receptor 1) and *CD36* (Scavenger Receptor Class B, Member 3), correlated positively with gene expression and translation levels in macrophages compared to monocytes (Fig. 4B, 4C and Additional file 7: Fig. S5A). In METTL3-depleted Mϕ macrophages (Fig. 2A), which demonstrated significantly lower global m^6^A levels on mRNA (Fig. 2B), we observed a significant reduction in both mRNA expression and protein levels of MSR1 (Fig. 4B) and CD36 (Fig. 4C), and mRNA expression of *CSF1* (Additional file 7: Fig. S5A).

**Figure 4.**
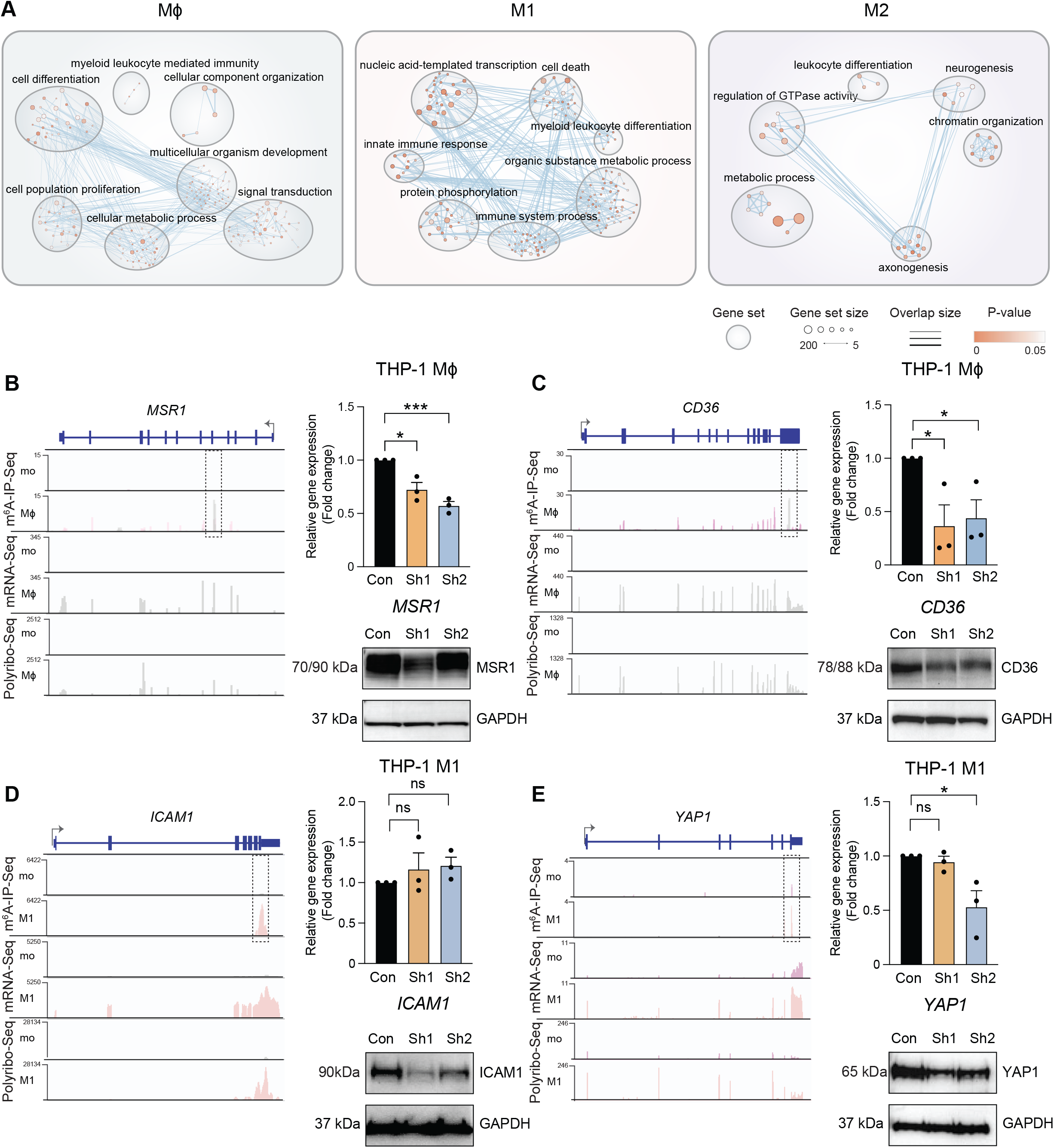
Modulation of m^6^A levels alters the expression and translation of critical genes that regulate macrophage differentiation and function. (**A**) Enrichment map of gene ontology terms representing genes with enriched m^6^A peaks in Mϕ, M1 and M2 populations. (**B-E**) Left: Coverage plots of m^6^A-IP-Seq (top), mRNA-Seq (middle) and Polyribo-Seq (bottom) for (**B**) *MSR1* and (**C**) *CD36* in Mϕ, and (**D**) *ICAM1* and (**E**) *YAP1* in M1. m^6^A-IP-Seq tracks show the overlay of input and IP data. An m^6^A peak is highlighted within a dotted box. m^6^A-IP-Seq coverage plots are displayed in BPM (bins per million reads, Bin size=1). mRNA-Seq and Polyribo-Seq coverage plots are displayed in RPKM (reads per kilobase per million reads). Right: Gene expression (top) and protein level (bottom) changes following METTL3 depletion in Mϕ or M1 cells. All data are from at least three independent experiments. Bar plots show mean ± SEM. An unpaired two-tailed Student’s t-test was used to determine significance, denoted by ns, not significant; *, *p* <0.05; *** and *p* <0.001. Con, non-targeting shRNA; Sh1 and Sh2, METTL3-targeting shRNAs.

In M1 pro-inflammatory-like macrophages, high levels of m^6^A were found in transcripts that are highly expressed and translated during macrophage response to infection including *ICAM1* (Intracellular Adhesion Molecule 1) and *YAP1* (Yes-Associated Protein 1) (Fig. 4D and E). Although METTL3-depleted M1 macrophages did not produce lower levels of *ICAM1* mRNA, ICAM1 protein levels decreased in these cells (Fig. 4D). The expression of *YAP1* mRNA decreased in METTL3-depleted M1 cells generated using one of the two shRNAs against METTL3 (sh2), however, YAP1 protein levels decreased in cells transduced with either shRNA (Fig 4E). In M2 macrophages, mRNA expression levels were significantly reduced following METTL3 depletion in the case of the scavenger receptor *CD163* (Additional file 7: Fig. S5B) and the anti-inflammatory factor, *COL6A2* (Collagen type VI alpha 2 chain) (Additional file 7: Fig. S5C). Both of these genes have established roles in M2 macrophage biology (64–67). In conclusion, we confirmed that METTL3 controls the expression of genes that are pivotal for macrophage functions.

### Inhibition of TET enzymes may facilitate monocyte-to-macrophage differentiation

To explore whether TET enzymes regulate monocyte-to-macrophage differentiation *in vitro*, we inhibited the catalytic activity of TETs by using the metabolite Itaconic Acid (ITA) to block their active sites, as reported in our recent study (68). We confirmed that inhibition of TETs’ activities led to a decrease in global RNA 5hmC levels in THP-1 monocytes (Fig. 5A). Following ITA treatment, THP-1 monocytes spontaneously expressed higher levels of the macrophage markers CD11b and CD44, although these changes were markedly lower than the levels achieved via PMA treatment (Fig. 5B). Negligible changes in the expression of M1 markers (CD80 and CD38) were observed (<1.2-fold), while no significant difference was detected in the levels of an M2 marker (CD209) by flow cytometry (Fig. 5C and 5D). When inhibition of TETs by ITA was performed together with PMA-induced differentiation, no difference in CD11b or CD44 levels was observed in THP-1 macrophages compared to control cells without ITA treatment (Additional file 8: Fig. S6A). There was no increase in the expression levels of CD80 and CD38 in macrophages stimulated with LPS and IFNγ following exposure to ITA compared to control (Additional file 8: Fig. S6B). In macrophages stimulated with IL-4 and IL-13, no significant change was observed in the expression levels of CD209 following ITA treatment (Additional file 8: Fig. S6C).

**Figure 5.**
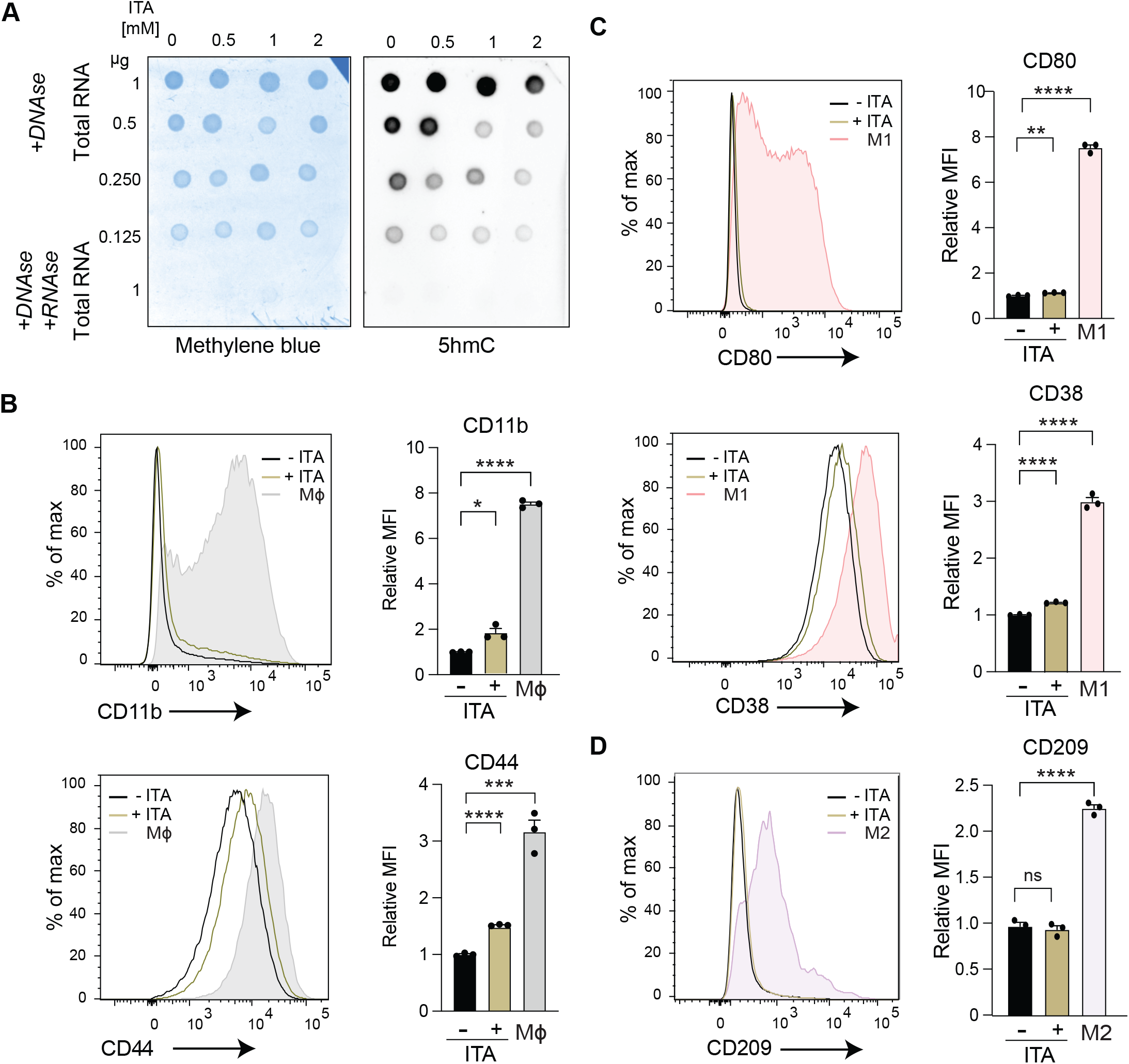
Inhibition of TET enzymes facilitates monocyte-to-macrophage differentiation. (**A**) Dot blot on total RNA from THP-1 mo treated with increasing concentrations of Itaconic Acid (ITA) probed with anti-5hmC antibody (right) and methylene blue loading control (left). Samples were treated with DNase to eliminate the possibility of detecting 5hmC on DNA. Negative controls were treated with both DNase and RNase A. Flow cytometry profiles and quantification (Mean Fluorescence Intensity (MFI)) showing changes in the relative expression of (**B**) Mϕ cell surface markers CD11b and CD44, (**C**) M1 markers CD80 and CD38 and (**D**) M2 marker CD209 following treatment of mo with 2mM Itaconic Acid (ITA) compared to controls. All data are from at least three independent experiments. Bar plots show mean ± SEM. An unpaired two-tailed Student’s t-test was used to determine significance, denoted by *, *p* <0.05; **, *p* <0.01; ***, *p* <0.001 and ****, *p* <0.0001.

Our data indicate that inhibition of TETs in THP-1 monocytes may trigger their differentiation into macrophages but, like METTL3 depletion, it is insufficient to facilitate complete differentiation and polarisation of macrophages. ITA-mediated inhibition of TETs does not accelerate PMA-mediated differentiation or promote M1 and M2 polarisation of THP-1-derived macrophages triggered by pro- and anti-inflammatory stimuli.

### Changes in RNA 5hmC levels are predominantly associated with the transcription and translation of transcripts that also bear m^6^A in a different region

Next, we examined the transcriptome-wide distribution of 5hmC in monocytes and macrophages using 5hmC RNA immunoprecipitation sequencing (5hmC-IP-Seq). Overall, we identified between 151–182 5hmC peaks for monocytes and macrophages corresponding to 128–161 genes (Additional file 9: Table S3), of which 31 5hmC peaks (corresponding to 24 genes) were common to monocytes and macrophages, 99 5hmC peaks (in 94 genes) were exclusive to monocytes and 26 5hmC peaks (in 25 genes) were shared by all macrophages (Fig. 6A). Despite a global decrease of RNA 5hmC in macrophages compared to monocytes (Fig. 1G and 1H), the number of increased and decreased RNA 5hmC peaks in Mϕ compared to mo was similar (Additional file 10: Fig. S7A). The number of differential 5hmC peaks detected during the transition of Mϕ to M1 or Mϕ to M2 was less than half of those observed in mo to Mϕ (Additional file 10: Fig. S7A). This finding is consistent with the observed lack of changes in global 5hmC levels between polarised and non-polarised macrophages (Fig. 1H).

**Figure 6.**
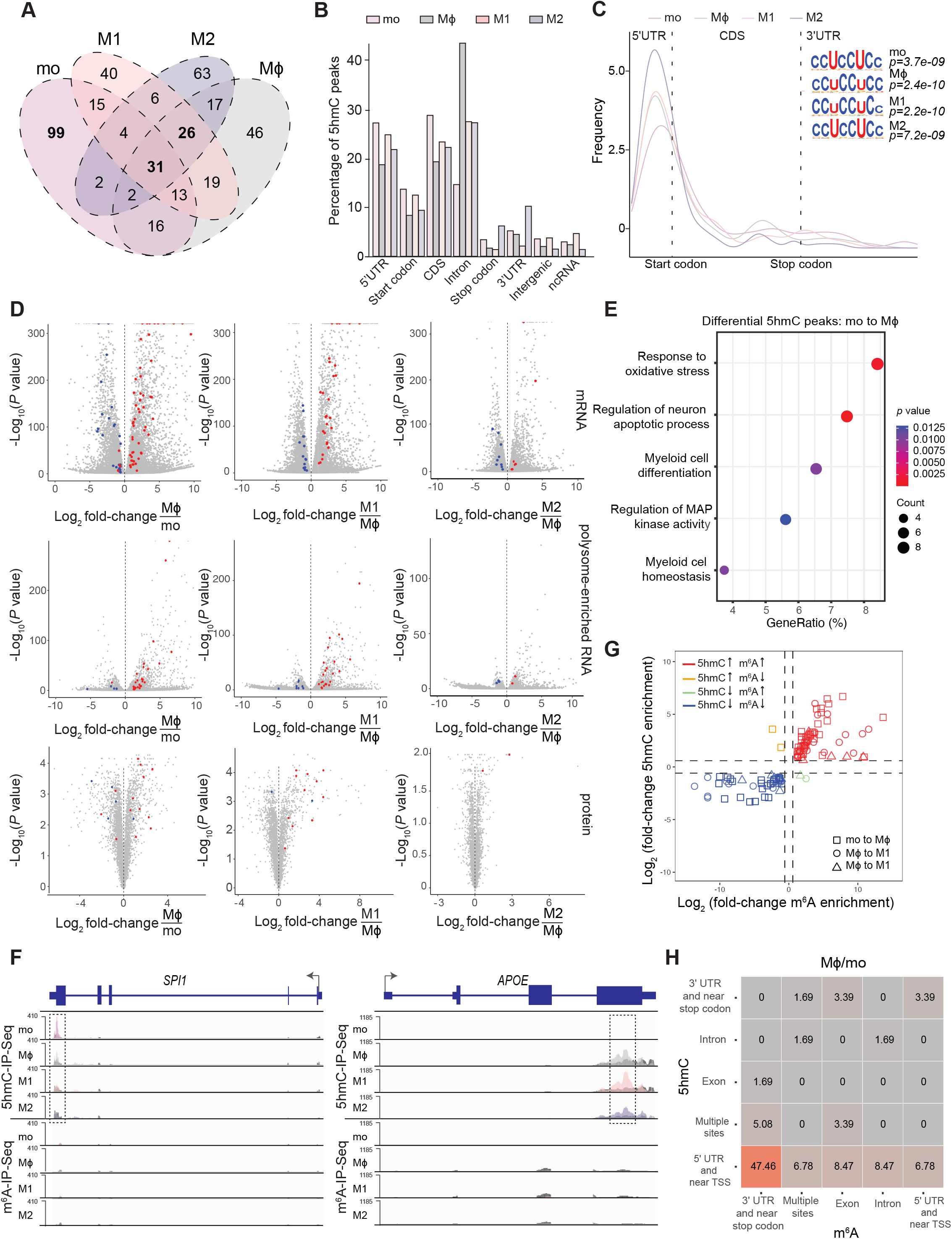
Changes in RNA 5hmC levels are predominantly associated with the transcription and translation of transcripts that also bear m^6^A in a different region. (**A**) Venn diagram showing the overlap of 5hmC peaks detected in mo, Mϕ, M1 and M2. (**B**) Percentage of 5hmC peaks identified by 5hmC-IP-Seq in defined transcript and genomic regions: coding sequence (CDS), 5’untranslated region (5’UTR), 3’untranslated region (3’UTR), start and stop codons, intergenic regions and non-coding RNA (ncRNA) found in mo, Mϕ, M1 and M2. (**C**) Metagene profiles detailing 5hmC distribution along a normalised transcript and top enriched sequence motifs identified by 5hmC-IP-Seq in all cell types. (**D**) Volcano plots showing differential gene expression (mRNA, top), translation (polysome-enriched RNA, middle) and protein (bottom) levels. Differential 5hmC peaks (red: increased, blue: decreased) are shown for Mϕ compared to mo, M1 compared to Mϕ and M2 compared to Mϕ. (**E**) Gene ontology analysis on differential 5hmC peaks during mo to Mϕ differentiation. (**F**) Coverage plots for 5hmC-IP-Seq (top) and m^6^A-IP-Seq (bottom) data from mo, Mϕ, M1 and M2 shown for *SPI1* and *APOE*. 5hmC- and m^6^A-IP-Seq tracks show the overlay of input and IP data. A 5hmC peak is highlighted within a dotted box. 5hmC and m^6^A-IP-Seq coverage plots are displayed in BPM (bins per million reads, Bin size=1). (**G**) The association between m^6^A and 5hmC enrichment on 5hmC-modified transcripts during macrophage differentiation and polarisation. (**H**) The proportion of differentially enriched 5hmC and m^6^A peaks that cooccurred in defined transcript regions (UTR, untranslated region; TSS, transcriptional start site, exon and intron) during monocyte to macrophage differentiation.

The majority of 5hmC peaks were found in the CDS, intronic and 5’UTR regions (~25% for each feature) (Fig. 6B). In contrast to m^6^A, 5hmC metagene profiles revealed strong enrichment of this modification towards the 5’UTR of mRNA transcripts (Fig. 6C). In addition to an enrichment of UCC-rich repeats reported previously in peaks containing RNA 5hmC (27,28), we also identified CCG- and CA-rich motifs in all four stages of monocyte-to-macrophage differentiation and polarisation (Fig. 6C and S7B). Notably, like m^6^A, we also observed a trend by which increased RNA 5hmC levels are associated with increased transcript expression and translation as mo differentiate into Mϕ and following polarisation of Mϕ into M1 macrophages (Fig. 6D). The opposite was true for transcripts with RNA 5hmC levels that decreased between these stages (Fig. 6D). The positive association between RNA 5hmC levels and transcription or translation during M2 polarisation was not as obvious due to fewer quantifiable transcripts and proteins in M2 macrophages (Fig. 6D).

GO analysis on transcripts with differential RNA 5hmC peaks between mo and Mϕ revealed enrichment of functions relevant to myeloid differentiation and homeostasis (Fig. 6E). Relevant genes within these GO categories include the macrophage transcription factor *SPI1/PU.1* (69), and *FOXO3* which regulates autophagy (70); an essential stimulus for monocyte-macrophage differentiation (Fig. 6F and S7C). *QKI* transcripts encoding a splicing factor that promotes monocyte differentiation into macrophages also exhibited increased RNA 5hmC peak in Mϕ (71) (Additional file 10: Fig. S7C). Functions associated with the regulation of MAP kinase activity and oxidative stress, which are pivotal for macrophage differentiation (72,73), were enriched (Fig. 6E). Significant enrichment of differential RNA 5hmC peaks was also observed in *MCL1*, a switch that modulates macrophage viability or apoptosis during bacterial clearance (74) (Additional file 10: Fig. S7C).

In line with the roles of M1 macrophages, differential RNA 5hmC peaks in M1 compared to Mϕ were enriched in functions associated with responses to reactive oxygen species, tumour necrosis factor and apoptosis involved in pro-inflammatory processes (59,72,74) (Additional file 10: Fig S7D). Key genes within these categories include *MCL1* and *APOE* (74,75) (Fig. 6F and Additional file 10: Fig. S7C). GO analysis was not performed on M2 macrophages given the limited number of differential RNA 5hmC peaks (n=30) in M2 compared to Mϕ. However, we observed enriched RNA 5hmC peaks in transcripts encoding proteins known to be expressed at high levels and/or secreted by M2. They include a classical M2 marker, FN1 (41), and a mesenchymal marker, Vimentin (VIM); both have been shown to be involved in the antiinflammatory process of wound healing (76,77) (Additional file 10: Fig S7C).

Given that m^6^A and 5hmC-modified mRNAs were both enriched for functions pertinent to different states of macrophages, we determined whether these modifications occurred on the same transcript or if the presence of one or the other was exclusive. Nearly two-thirds of the transcripts bearing 5hmC in monocytes or macrophages were also modified by m^6^A (Additional file 10: Fig. S7E). A significant association between 5hmC and m^6^A changes was also observed during monocyte-to-macrophage differentiation and polarisation (Fig. 6G, *p* = 2.2e-16, Fisher’s Exact Test). Transcripts bearing both 5hmC and m^6^A marks include some of the transcripts highlighted above: *FOXO3, MCL1* and *VIM* (Additional file 10: Fig. S7C). However, we did not detect m^6^A peaks in the other 5hmC-modified transcripts described herein (Fig. 6F and Additional file 10: Fig. S7C). In the vast majority of transcripts, RNA 5hmC peaks were found in the 5’ UTR and near transcriptional start sites, whereas m^6^A peaks were enriched in the 3’ UTR and near stop codons (Fig. 6H and Additional file 10: Fig. S7F). Notably, in some cases, such as *FAM20C* and *MFSD12*, we observed an enrichment of 5hmC and m^6^A peaks mapping to different alternative 5’ and 3’ isoforms in macrophages (Additional file 10: Fig. S7C). Overall, we have identified several key transcripts presenting differential 5hmC methylation in monocytes and macrophages. As with m^6^A, we observe a positive association between 5hmC, gene expression and translation levels, suggesting that 5hmC may also be regulating genes with functions relevant to monocytes and macrophages. Furthermore, we identify important macrophage regulators bearing both RNA modifications, located on opposite ends of the transcript. To the best of our knowledge, these results are the first to identify the presence of 5hmC in mRNAs harbouring m^6^A. This concurrence of modifications on the same transcript adds another dimension to the complex regulatory dynamics exhibited by RNA.

### Depletion of RNA 5hmC leads to increased half-life of critical transcripts in macrophages

While exploring our 5hmC-IP-Seq data, two genes were particularly noteworthy: *ARID1A* and *GRK2* (Fig. 7A and B). *ARID1A* (AT-Rich Interaction Domain 1A), a principal component of the BAF SWI/SNF chromatin remodelling complex, is required for maintenance of lineagespecific enhancers (78), essential for myeloid differentiation (79) and induces the antiviral interferon response in macrophages (80). *GRK2* (G Protein-Coupled Receptor Kinase 2), a central signalling node that modulates G protein-coupled receptors (GPCRs) and other cell signalling routes including the NF-KB and MAPK inflammatory pathways, is a critical regulator of chemotaxis (81,82) and myeloid-specific deficiency of *GRK2* results in excessive cytokine production in a sepsis model (83). Since both *ARID1A* and *GRK2* have important functions in myeloid cell fate, we reasoned that the expression of these genes must be tightly regulated and hypothesised that RNA 5hmC is involved in this process. Even though the role of RNA 5hmC is still unclear, studies have linked the presence of this mark with transcript stability (27,30). We therefore determined whether 5hmC regulates the stability of *ARID1A* and *GRK2* transcripts by measuring their decay rates in 5hmC-depleted Mϕ using ITA to block TETs’ activity. As a negative control *MYC* transcripts are not modified by 5hmC (Additional file 11: Fig. S8A) and thus, *MYC* RNA half-life was not affected by ITA treatment in Mϕ (Additional file 11: Fig. S8B). However, the half-life of *ARID1A* and *GRK2* was 4 to 6 times longer in 5hmC-depleted Mϕ (Fig. 7C and 7D). This finding is consistent with previous work performed in mESCs showing that RNA 5hmC reduces the stability of crucial pluripotency-promoting transcripts (27) and that 5hmC promotes *MERVL* retrotransposon destabilization (30). Given that *ARID1A* but not *GRK2* (Fig. 7A and B) harbours m^6^A, we also measured *ARID1A* RNA decay rate following METTL3 depletion in Mϕ (Fig. 7E) and found that METTL3 depletion does not affect *ARID1A* half-life. These data indicate that TET-mediated RNA 5-hydroxymethylation reduces the stability of *ARID1A* and *GRK2* in THP-1-derived macrophages independently of m^6^A.

**Figure 7.**
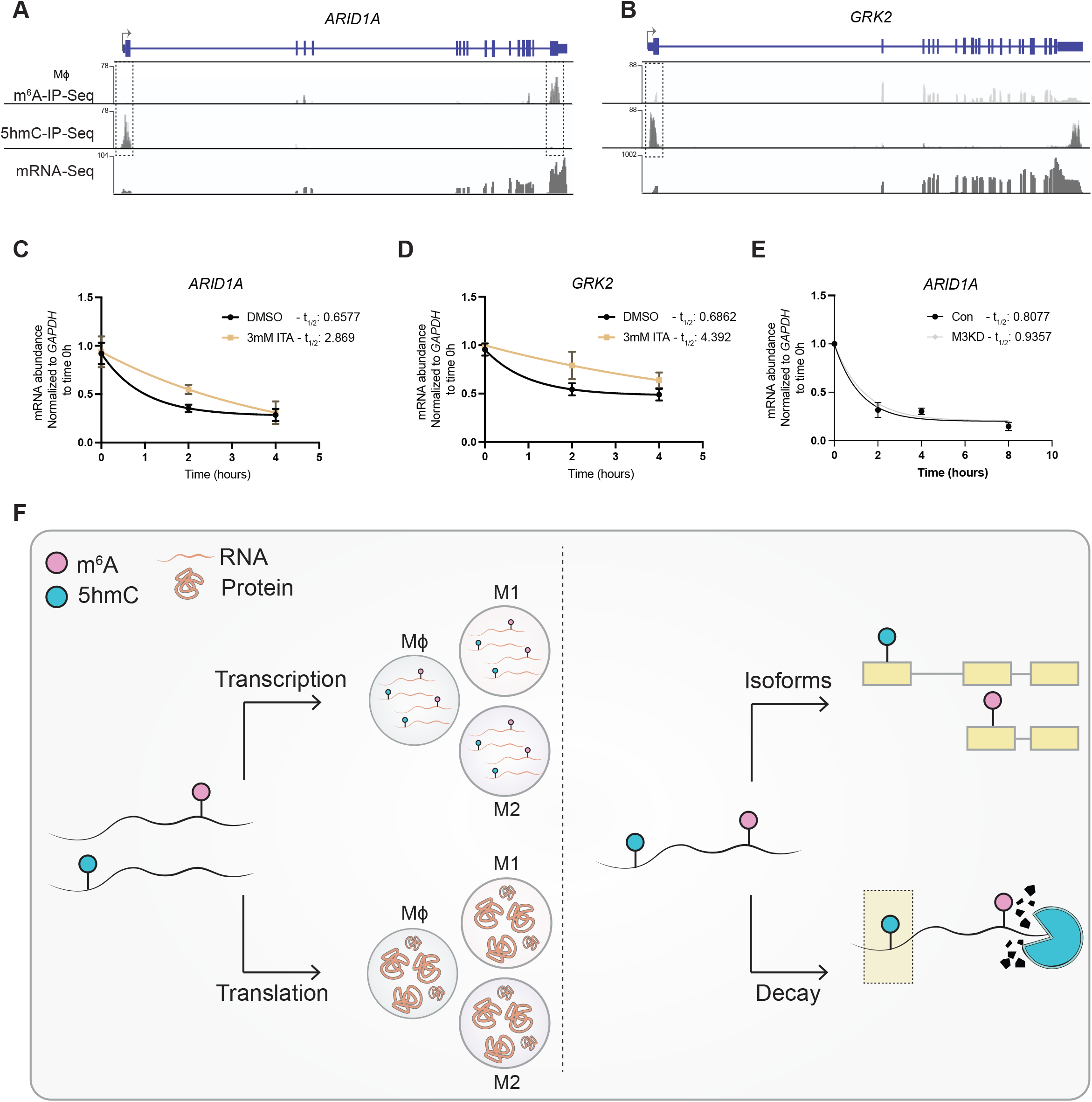
Depletion of RNA 5hmC levels leads to increased half-life of critical transcripts in macrophages. Coverage plots of m^6^A-IP-Seq (top), 5hmC-IP-Seq (middle) and mRNA-Seq (bottom) data for (**A**) *ARID1A* and (**B**) *GRK2* in Mϕ. (**C and D**) mRNA decay plots for ARID*1A* and *GRK2* in Itaconic Acid (ITA)-treated Mϕ. (**E**) mRNA decay plots for *ARID1A in* METTL3-depleted Mϕ. (**F**) Diagram summarising key findings. Left: m^6^A and 5hmC are associated with transcription and translation of key macrophage genes. Right: m^6^A and 5hmC can coexist on specific transcripts. In some cases, these modifications mark distinct alternative mRNA isoforms. Although m^6^A is known to regulate RNA decay, 5hmC can also facilitate the decay of specific mRNAs independently of m^6^A.

## Discussion

Following infection, sensing of danger signals leads to activation of the innate immune response and profound phenotypic changes drive monocyte-to-macrophage differentiation and polarisation (61,84–86). Due to the critical role of these immune cells, extensive research has been conducted to understand the cellular and molecular mechanisms controlling them in homeostasis and disease. However, little is known as to how RNA modifications and altered expression of RNA modification enzymes are involved in this process. To address this gap, we generated a multi-omics dataset that profiled the transcriptome, m^6^A and 5hmC epitranscriptomes, polyribosome-enriched mRNAs and proteome across four monocyte-macrophage states (Fig. 1A), and analysed m^6^A and 5hmC patterns in the context of monocyte-to-macrophage differentiation and polarisation.

Consistent with other reports (87), our results indicate that depletion of METTL3 triggers differentiation of myeloid cells as evidenced by increased number of cells exhibiting high expression of CD11b. Higher expression of CD44 that accompanied macrophage-like morphology confirms that METTL3-depleted THP-1 cells can differentiate toward macrophages (Fig. 2C and D). We also demonstrate for the first time that depletion of RNA 5hmC via inhibition of TETs activity exerts a similar effect in triggering monocyte differentiation (Fig. 5C). However, in comparison to an appropriate positive (i.e., PMA-treated) control, we show that either downregulation of METTL3 or inhibition of TET enzymes alone is insufficient to directly drive monocyte-to-macrophage differentiation.

Although there are conflicting reports on the role of METTL3 in polarising the pro-inflammatory macrophage phenotype in mice (39,40), our results indicate that METTL3 has a role in promoting the M1 pro-inflammatory phenotype in human cells *in vitro* (Additional file 3: Fig. S3A and B). In contrast to previous reports suggesting that METTL3 depletion promotes the polarisation of M2 anti-inflammatory macrophages in mice (39,40), our data indicate that METTL3 depletion does not promote the M2 phenotype (Additional file 3: Fig. S3C and D). This discrepancy may be due to differences between human and mouse macrophages, differences in the levels of METTL3 depletion and/or differences in stimulation conditions to model the anti-inflammatory phenotype. Future studies should aim to further characterise the role of METTL3 in the anti-inflammatory context.

Despite an overall decrease in RNA m^6^A and 5hmC levels in macrophages compared to monocytes (Fig. 1D to I), high levels of these modifications are present on a substantial number of transcripts. This observation implies that the global decrease in these RNA modification marks cannot be directly extrapolated to explain mechanisms acting at the level of specific transcripts. RNA modifications persist during cellular differentiation to maintain and/or enhance the levels of transcripts that are critical for this process. This result may also explain our observations concerning the incomplete monocyte-to-macrophage differentiation, as “forceful” METTL3 depletion or 5hmC inhibition may inadvertently remove RNA modifications that are required for the expression of specific transcripts to enhance this process. In addition, some transcripts bearing RNA modifications may not promote monocyte differentiation and macrophage polarisation per se but are important to regulate macrophage functions in a context-dependent manner. Herein, we have provided key examples of these genes, but many others remain to be identified in future studies.

Through an integrative approach, we have identified a positive association between RNA 5hmC, m^6^A, and transcriptional changes (Fig. 3D and 6D), coherent with previously proposed co-transcriptional deposition models where m^6^A (52,53) and 5hmC (27) are deposited as transcription proceeds. In line with our findings, m^6^A has been shown to promote translation in various contexts (88,89). Interestingly, while RNA 5hmC abundance is positively correlated with active mRNA translation in *D.melanogaster* (28), it did not seem to impact mRNA translation but instead reduced the stability of critical transcripts in mESCs (27). By focusing on *ARID1A* and *GRK2*, our results support the role of 5hmC in regulating transcript stability in mammalian cells (Fig 7C and E). Since the temporal expression order of inflammatory molecules is influenced by mRNA stability (90), 5hmC may be an additional regulatory layer that fine-tunes gene expression timing of key genes during the inflammatory response.

Another noteworthy observation arising from our study is that transcripts harbouring 5hmC often harbour m^6^A, frequently on opposite ends of the molecule (Fig. 6H and S7F). In some cases, 5hmC and m^6^A mark alternatively spliced isoforms (Additional file 10: Fig. S7D), indicating that they may regulate alternative splicing and/or functions of these specific isoforms. Recently, others have recognised the interplay between RNA modifications (91). For example, cooperation between m^6^A and 5mC has been reported to promote replication of murine leukemia virus (92) and enhance p21 translation in oxidative stress-induced cellular senescence (93). Since RNA 5hmC is a product of 5mC oxidation (94,95), it is tempting to speculate that an m^6^A-5mC-5hmC axis can act on specific transcripts under distinct cellular contexts. Future studies exploring the nature of the interaction between RNA modifications should aim to establish whether one modification prevails over others, and if they act synergistically or exclusively. Herein, we provide evidence that 5hmC can regulate mRNA decay independently of m^6^A (Fig. 7).

While there are studies investigating TET enzymes and DNA 5-hydroxymehtylation in macrophages ((96–98)), their roles in gene expression regulation remain unclear. To advance the current understanding of these enzymes in macrophage biology, their contribution to both RNA and DNA 5-hydroxymethylation must be considered. Future research should aim to establish the mechanisms underpinning TETs’ selectivity towards DNA and/or RNA. It is plausible that crosstalk between RNA-m^6^A and DNA-5hmC is also regulating macrophage biology. Notably, recent work has shown that RNA-m^6^A regulates TET-1-mediated DNA 5-hydroxymethylation to promote transcription and chromatin accessibility in cancer (99). Future work should aim to distinguish the importance of the RNA-m^6^A-RNA-5hmC axis and the RNA-m^6^A-DNA-5hmC axis to gene expression regulation in macrophage development and functions.

Altogether, our observations raise at least three important points; first, patterns/associations observed at the global level cannot be directly extrapolated to explain mechanisms acting at the transcript level. Second, these data reinforce a concept that has emerged in the m^6^A literature (100), namely that RNA modifications have context-specific functions, and this is likely to be dictated by context-specific expression patterns of writers, erasers and reader proteins. Third, a transcript’s fate could be determined by more than one RNA modification, several modifications could co-exist and perhaps interact, cooperate and/or cross-regulate in different biological contexts. From a clinical perspective, macrophage dysfunction has been associated with the development of multiple disorders including cancer, metabolic diseases, and autoimmune disorders. In recent years, macrophage reprogramming has emerged as a novel approach to therapy. By adding to the current understanding of how the epitranscriptome regulates macrophage plasticity, this work provides a novel perspective that may inform new strategies to reprogram macrophages for the treatment of human diseases.

## Conclusions

In summary, we identify m^6^A and 5hmC as regulators of transcription and translation of mRNAs that control differentiation, polarisation and functions of macrophages (Fig. 7F). Our study highlights the possible interplay between two distinct types of RNA modifications in dynamic cellular contexts. We propose the coexistence of m^6^A and 5hmC as a potential mechanism coordinating the execution of complex gene expression programs by facilitating RNA decay and alternative isoform expression (Fig. 7F). These regulatory roles of m^6^A and 5hmC may be essential for the establishment and maintenance of cellular identity and functional plasticity of macrophages. This comprehensive dataset provides new understanding of the molecular mechanisms governing cell development intrinsic to innate immunity.

## Methods

### Monocyte-to-macrophage differentiation and polarisation

THP-1 cells were maintained in RPMI medium (ThermoFisher Scientific) supplemented with 10% (v/v) fetal calf serum (Hyclone, GE Healthcare), 1% (v/v) non-essential amino acids (ThermoFisher Scientific), 1 mM sodium pyruvate (ThermoFisher Scientific) and 0.1 mg/ml penicillin and streptomycin (ThermoFisher Scientific). To generate THP-1-derived macrophages(Mϕ), THP-1 monocytes were stimulated with 100 nM PMA (Sigma) for 48 hours. M1 macrophages were generated by stimulating Mϕ with 1 μg/ml LPS (Sigma) and 20 ng/ml IFN-γ for 6 or 48 hours. M2 macrophages were generated by stimulating Mϕ with 20 ng/ml IL-4 (R&D Systems) and 20 ng/ml IL-13 (R&D system) for 24 hours.

### RNA extraction, cDNA synthesis and RT-qPCR

RNA extraction was performed using TRIzol Reagent (Invitrogen) and cDNA synthesis was performed using iScript gDNA Clear cDNA synthesis kit (Bio-Rad) following the manufacturer’s instructions. RT-qPCR was performed using SensiFAST SYBR No-ROX Kit (Bioline) and 0.3 μM of forward and reverse primers. Samples were amplified and analysed using the LightCycler 480 Instrument II (Roche), cycling conditions: 95°C for 3 min, followed by 40 cycles at 95°C for 10 s, annealing at 60°C for 30 s, and extension at 72°C for 20 s. Fold change was calculated using the ΔΔCT method. See Additional file 12: Table S4 for a list of the primers used in this work.

### m^6^A and 5hmC quantification

Sample preparation for m^6^A quantification was performed as described previously (12). In brief, mRNA was isolated using the Magnetic mRNA Isolation Kit (New England Biolabs), followed by rRNA depletion using the RiboMinus Eukaryote System v2 (Invitrogen) as per manufacturer’s instructions. 200 ng mRNA were digested with 1U nuclease P1 (Sigma) in 20 mL buffer containing 25 mM NaCl, 2.5 mM ZnCl2 at 37°C for 2 hours, followed by the addition of 1M NH4HCO3 (2 mL) and 1 U of alkaline phosphatase (Sigma). The solution was incubated at 37°C for 2 hours, centrifugated at 13000 g for 10 minutes at 37°C, and then 10 μL of the solution were injected into the mass spectrometer. Quantification was performed by comparison with a standard curve obtained from pure nucleoside standards. The m^6^A to A ratio was calculated based on obtained concentrations.

To quantify global RNA 5hmC levels, LC-MS/MS was performed as previously described (27). Briefly, 10X buffer (500 mM Tris-HCl, 100 mM NaCl, 10 mM MgCl2, 10 mM ZnSO_4_, pH 7.0), 180 U of S1 nuclease, 0.001 U of venom phosphodiesterase I, and 30 U of CAIP were added to 10 μg of total RNA. The mixture was incubated at 37°C for 4 hours. The resulting solution was then extracted with chloroform three times. The upper aqueous phase was collected and passed through a solid-phase extraction cartridge filled with 50 mg of sorbent of graphitised carbon black to remove the salts. The elution was then dried with nitrogen gas at 37°C for subsequent chemical labelling and LC–ESI-MS/MS analysis by an AB 3200 QTRAP mass spectrometer (Applied Biosystems).

### Western blotting

Protein lysates were generated by incubation with RIPA Buffer (Sigma) supplemented with 1% complete protease inhibitor cocktail (Sigma) on ice for 30 min. Protein was quantified using the Micro BCA Protein Assay kit (ThermoFisher Scientific) following the manufacturer’s instructions. Protein lysates were loaded onto 4-12% BIS-Tris Protein gels (ThermoFisher) for electrophoresis followed by transfer onto a PVDF membrane (Merck Millipore). Membrane was blocked with 5% (v/v) skim milk TBS- 0.1% Tween-20 for 1 hour at room temperature, incubated overnight with a 1:1000 dilution of rabbit anti-METTL3 (ab195352, Abcam), anti-MSR1 (ab271070, Abcam), anti-CD36 (ab133625, Abcam), anti ICAM1 (4915, Cell Signalling Technologies), anti-YAP1 (4912, Cell Signalling Techologies) or 1:5000 dilution of mouse anti-GAPDH (ab8245, Abcam) antibody, followed by incubation with a 1:5000 dilution of a donkey anti-rabbit IgG HRP (AP182P, Merck Millipore) or donkey anti-mouse IgG HRP (AP192P, Merck Millipore) antibodies. Protein detection was performed using SuperSignal West Pico PLUS (ThermoFisher) and imaged on a ChemiDoc Imaging System (BioRad).

### mRNA Sequencing

Paired-end mRNA-Seq was performed for all samples. 2 μg of DNAse treated total RNA was sent to Novogene for directional mRNA library preparation (mRNA enrichment) and sequencing using NovaSeq. 50 million paired-end 150 bp reads were obtained for each sample.

### Polyribo-Seq

Polyribo-Seq sample preparation and uHPLC Size Exclusion Chromatography (SEC) were performed as described by Yoshikawa et al., (101). Briefly, cell lysates in CHAPS buffer containing RNAse inhibitors were separated by size exclusion chromatography on a Thermo Dionex BioRs UHPLC and Agilent SEC-5 7.8 × 300 mm HPLC column with 2000Å pores and 5 mm particles. Polysome fractions were harvested in TRIzol LS reagent (10296028, Invitrogen) and RNA was extracted following the manufacturer’s protocol. RNA-Seq was then performed on RNA from the polysome-bound fraction.

### Proteomic Analysis

Sample preparation, LC-MS/MS and analysis of spectra was performed as described in Green et al.(42). Briefly, 20 μg of protein was denatured, reduced and alkylated followed by trypsin digestion at 37°C for 16 hours. Peptides were then purified using StageTips. Dried peptides were resuspended in 60 μL of 5% formic acid and stored at 4°C until analysed by LC-MS. Using a Thermo Fisher RSLCnano, peptides were separated using a 50 cm × 75 um C18 (Dr Maisch, Ammerbuch, Germany, 1.9 μm) column with a ~10 μm pulled tip, coupled online to a nanospray ESI source. Peptides were resolved over a gradient from 5% acetonitrile to 40% acetonitrile over 140 min with a flow rate of 300 nL min–1. Tandem mass spectrometry analysis was carried out on a Fusion Lumos mass spectrometer (Thermo Fisher) using HCD fragmentation. The data-dependent acquisition method used acquired MS/MS spectra of the top 20 most abundant ions at any one point during the gradient. Raw data were analysed using MaxQuant (38) (version 1.6.3.4). Peptide and protein level identification were set to a false discovery rate (FDR) of 1% using the human Uniprot database. Mass tolerance was set to 4.5 ppm for precursor ions and MS/MS mass tolerance was 20 ppm. Enzyme specificity was set to trypsin, with a maximum of 2 missed cleavages permitted. Deamidation of Asn and Gln, oxidation of Met, pyro-Glu and protein N-terminal acetylation were set as variable modifications. Carbamidomethyl on Cys was searched as a fixed modification.

### m^6^A-IP-Seq

m^6^A-IP-Seq was performed as previously described (102). Briefly, 5 μg of DNAse treated total RNA were subjected to fragmentation with ZnCl2 incubation at 70°C for 3min. Following precipitation, fragmented RNA size distribution was assessed using RNA 6000 Nano Bioanalyzer kit (Agilent). Approximately 500ng of sample were stored as input control. Fragmented RNA was subjected to two rounds of m^6^A immunoprecipitation for 2 hours each using an anti-m^6^A antibody (ABE572, Merck) previously conjugated to protein-A magnetic beads (Thermo Fisher Scientific) and of protein-G magnetic beads (Thermo Fisher Scientific) by incubation at 4°C for at least 6 hours. Following extensive washing, RNA was eluted from the beads using RLT buffer and RNeasy mini kit (Qiagen). RNA was quantified using RNA 6000 Pico Bioanalyzer kit (Agilent). To confirm m^6^A enrichment, cDNA was synthesized using the SensiFast cDNA synthesis kit (Bioline) and SETD7 and GAPDH levels measured by qPCR were used as positive and negative control respectively. Finally, library preparation was performed using the SMARTER Stranded Total RNA Seq kit v2-Pico Input Mammalian kit (Takara Bio) following the manufacturer’s instructions. Libraries were then sequenced using HiSeq2500 (Illumina) and a minimum of 20 million paired-end reads were obtained per sample. This experiment was performed in two replicates for each sample.

### 5hmC-IP-Seq

5hmC-IP-Seq was performed as described by Delatte, et al., (28) and Lan et al., (27). In brief, 1 mg of DNAse treated total RNA was chemically fragmented by incubating RNA in fragmentation buffer (10 mM Tris-HCl pH7, 100 mM ZnCl2) at 94°C for 40 seconds, the reaction was stopped by the addition of 50 mM EDTA. Fragmented RNA was ethanol precipitated and resuspended in nuclease-free water. Fragmentation efficiency was checked by running a Nano RNA Bioanalyzer chip (Agilent) on the Agilent 2100 Bioanalyzer. Prior to immunoprecipitation, fragmented RNA was denatured by incubation at 70°C for 5 minutes and then placed on ice. RNA was then incubated at 4°C overnight with or without 12.5 μg of anti-5hmC antibody (C15220001, Diagenode) in freshly prepared 1X Immunoprecipitation (IP) buffer (50 mM Tris-HCl pH 7.4, 750 mM NaCl and 0.5%Igepal CA-630, RNasin 400 U/ml and RVC 2 mM) supplemented with protease inhibitor (complete EDTA free, Roche). 60 μL of equilibrated Dynabeads Protein G (Invitrogen) were added to each sample and incubated for 2.5 hours at 4°C. After three washes with 1X IP buffer, samples were eluted by addition of 1 mL of TriPure Reagent (Roche) as per manufacturer’s instructions. Samples were analysed by deep sequencing. Libraries were prepared using the TruSeq ChIP Sample Prep Kit (Illumina) after reverse transcription of pulled-down RNA and synthesis of a second strand (NEBNext mRNA second strand synthesis module (NEB)). Briefly, 5 to 10 ng of dsDNA underwent 5’ and 3’ protruding end repair. Then, non-templated adenines were added to the 3’ ends of the blunted DNA fragments. This last step allows ligation of Illumina multiplex adapters. The DNA fragments were size selected to remove unligated adapters and to sequence fragments of 200-300 bp of length. The library was amplified through 18 cycles of PCR. DNA was quantified using Qubit, and DNA integrity was assessed by running a DNA Bioanalyzer chip (Agilent) on the Agilent 2100 Bioanalyzer. 6 pM of DNA library spiked with 1% PhiX viral DNA was clustered on cBot (Illumina) and then sequenced on a HiScanSQ module (Illumina). hmeRIP-Seq experiments were performed in triplicates for each condition.

### Generation of METTL3 knock-down cell lines

Stable gene knockdown of METTL3, in THP-1 cells was achieved by lentiviral transduction of the pLKO.1 containing specific short hairpin RNAs (Additional file 12: Table S4). The transduced cells were subject to selection by 7 days culture in 0.6 μg/mL puromycin. Knockdown was confirmed by western blotting.

### Enrichment of polyadenylated RNA, rRNA and sRNA

Polyadenylated RNA was enriched using the Poly(A)Purist MAG Kit (Invitrogen) as per manufacturer’s instructions. rRNA was separated from polyA RNA using GeneElute mRNA Miniprep kit (Sigma) and concentrated using RNA Clean and Concentrator kit (Zymo Research) and sRNA (including tRNA) was extracted using miRNeasy mini-Kit (Qiagen) and RNeasy MiniElute Cleanup Kit (Qiagen). RNA quality and purity of the fractions was performed using Small RNA, PICO and NANO RNA Bioanalyzer kits (Agilent).

### 5hmC Dot Blotting

To perform dot blot, RNA was treated with TURBO DNase (ThermoFisher) following the manufacturer’s protocol. As a control, RNA was treated with 1U RNase A (Qiagen) for 1 hour at 56°C. RNA was then loaded onto a Hybond-N+ membrane (GE Healthcare), allowed to air dry and crosslinked twice at 200000 μJ/cm^2^ UV. To control for RNA loading, the membrane was then incubated with 0.04% methylene blue in 0.5 M sodium acetate for 5 min, following rinsing with PBS 0.1% Tween-20, it was imaged using the colorimetric function of ChemiDoc Imaging System (BioRad). The membrane was then blocked in 3% w/v skim milk PBS-0.1% Tween-20 for 1 hr at room temperature, incubated in blocking buffer with 1:500 rat anti-5hmC (Diagenode) overnight at 4°C and followed by incubation with 1:1000 anti-rat IgG HRP (ab6734, Abcam) in blocking buffer. 5hmC detection was performed using SuperSignal West Femto Maximum Sensitivity Substrate (ThermoFisher) and imaged on a ChemiDoc Imaging System (BioRad).

### Flow Cytometry

Flow cytometry analysis was performed using the following cell surface markers: anti-CD11b conjugated to phycoerythrin (PE, BD Biosciences, clone ICRF44) and anti-CD44 allophycocyanin (APC, BD Biosciences, clone IM7) conjugated to staining to confirm the differentiation of THP-1 mo into Mϕ. Anti-CD38 conjugated to PE-Cy7 (BioLegend, clone HIT2) and anti-CD80 conjugated to V450 (BD Biosciences, L307.4) to confirm the polarisation into M1-like cells. Anti-CD209 conjugated to BV421 (BD Biosciences, DCN46) to confirm the polarisation into M2-like cells. Cell viability was determined by staining cells with 0.5 μg/ml DAPI (Invitrogen). Acquisition was performed using a BD FACSCanto™ II and a BD FACSFortessa™ II (BD Biosciences) and data were analysed using FlowJo software (BD Biosciences). To quantify shifts in fluorescence intensity in normally distributed uniform populations, Relative Median Fluorescence Intensity (MFI) was calculated. Percentage of positive populations was calculated for populations displaying bi-modal distributions.

### RNA decay assay

METTL3-depleted and control THP-1-derived macrophages were treated with 10 μg/ml actinomycin D (Sigma) for 2, 4 and 8 hours and harvested. For RNA 5hmC depletion assays, THP-1-derived macrophages were treated with DMSO or 3 mM Itaconic Acid (Sigma) for 10 hours prior to actinomycin D treatment. RNA was extracted using TRIzol. cDNA and qPCR were performed as described above. mRNA decay rate was estimated by non-linear regression curve fitting as previously described (103).

### Bioinformatic analysis

For m^6^A- and 5hmC-IP-Seq, raw reads were trimmed to remove adaptor sequences and low-quality reads using Trimmomatic (104) with the default setting. Read quality was assessed using FastQC. Clean reads were aligned to the human reference genome hg38 (ENSEMBL version 86) using the STAR aligner (105). To minimise the rate of false positives, only uniquely mapped reads were selected using samtools (106). Peaks enriched in immunoprecipitated over corresponding input samples were called using MACS2 (107). Peaks identified in both biological replicates were merged using the mergePeaks command in the HOMER software (108) and overlapping peaks were mapped to the RefSeq gene annotation using intersectBed from BEDTools (109). Enriched m^6^A and 5hmC motifs were identified using *de novo* motif search with the HOMER software (version 4.9.1). Motifs with the most significant *P*-values were visualised using WebLogo (110) The metagene profiles were plotted using the ‘Guitar’ (111) R package. DESeq2 (112) was used to identify m^6^A peak levels that were significantly different (1.5-fold) between two samples with a Benjamini-Hochberg correction (*p* <0.05). Gene ontology (GO) biological processes (BP) enrichment analysis was performed for the genes with significantly increased/decreased m^6^A peaks using the ‘clusterProfiler’ (113) package.

For mRNA- and Polyribo-Seq, Truseq3-PE adapter and poor-quality sequences were assessed using FastQC were trimmed by Trimmomatic (104) using the default settings. Trimmed reads were then aligned to the hg38 (ENSEMBL version 86) reference genome using STAR. FeatureCounts (114) was subsequently employed to convert aligned short reads into read counts for each sample. The data were then analysed using R and DESeq2 (112). Differentially enriched mRNAs undergoing translation in the polysomes were identified using Wald statistical test, with fold-change > 1.5 and *p* < 0.05 after Benjamini-Hochberg correction. Expressed genes were identified as those with RPKM greater than 1 for at least one group of samples. Differentially expressed genes between two groups were identified using Wald statistics, with fold-change > 1.5 and *p* < 0.05 after Benjamini-Hochberg correction.

### Statistical Analysis

Unless stated otherwise, statistical analyses were performed using GraphPad Prism 9.0. Statistical significance was determined using Student’s t-test unless indicated otherwise in figure legends and text. *p* < 0.05 was considered statistically significant. All error bars represent the standard error of the mean (SEM) from independent experiments (n>=3 unless specified otherwise).

## Declarations

### Ethics approval and consent to participate

Not applicable

### Consent for publication

Not applicable

### Availability of data and materials

All data have been deposited at Gene Expression Omnibus (GEO) repository, accession number GSE130011 (42), GSE213207 and GSE130011. RAW MS data have been deposited to the ProteomeXchange Consortium (http://proteomecentral.proteomexchange.org) via the PRIDE partner repository with the dataset identifier PXD017391 (42).

### Author’s contributions

N.P. performed most of the experiments with critical contributions from Q.L., E.W., J.T., M.M. and K.-L.D. R.S. performed all bioinformatic analyses. B.Rong and F.L. performed m^6^A quantification by LC-MS/MS. E.C. performed 5hmC-IP-Seq. C.M. and B.Y. performed 5hmC quantification by LC-MS/MS. M.L. performed size exclusion chromatography for polysome fractionation experiments, and LC-MS/MS for proteomic analysis. B.Roediger, J.E.J R., D.Y. and F.F. provided input on experimental design and resources. J.J.-L.W. supervised the overall project. N.P. and J.J.-L.W. conceived the study, designed the experiments and wrote the manuscript with contributions from all authors.

## Acknowledgments

We thank Renjing Liu, Ka Ka Ting and Paul Coleman for reagents, Chau-To Kwok, Stephanie Sun, Immanuel D. Green, Alex Wong and Stuart J. Cook for technical assistance and Sydney Cytometry and Sydney MS for providing the instrumentation used in this study.

