## Supplementary Figures for "RNA m^6^A and 5hmC regulate monocyte and macrophage gene expression programs"

**Supplementary information**

**Additional file 1: Figure S1. Expression of m$^6$A and 5hmC writers in monocytes and macrophages.** Principal Component Analysis (PCA) of (**A**) RNA-Seq and (**B**) LC-MS/MS data from mo, Mϕ, M1_6h, M1_48h and M2. (**C**) Protein levels for METTL14, VIRMA, ZC3H13, WTAP and RBM15 in mo, Mϕ, M1 and M2. (**D**) Dot blot on total RNA, polyadenylated RNA, rRNA and sRNA (including tRNA) from Mϕ cells detected with anti-5hmC antibody (bottom) or methylene blue loading control (top). Samples were treated with DNase to eliminate the possibility of detecting 5hmC on DNA. Negative controls were treated with both DNase and RNase A. (**E and F**) *TET2* and *TET3* coverage plots of reads per million mapped reads for mRNA-Seq (top) and Polyribo-Seq (bottom) data from mo, Mϕ, M1 and M2. All data are from at least three independent experiments and show mean ± SEM. An unpaired two-tailed Student's t-test was used to determine significance, denoted by ns, not significant; *, $p$ <0.05; **, $p$ <0.01 and ***, $p$ <0.001.

**Additional file 2: Figure S2. Expression of M1 and M2 markers in METTL3-depleted THP-1 cells without polarising stimuli.** Flow cytometry profiles and quantification (Mean Fluorescence Intensity (MFI) or Percentage of Positive Cells) after METTL3 depletion showing changes in the relative expression of M1 cell surface markers CD80 and CD38 and M2 cell surface marker CD209 in (**A**) THP-1 mo and (**B**) THP-1 Mϕ. All data are from at least three independent experiments. Bar plots show mean ± SEM. An unpaired two-tailed Student's t-test was used to determine significance, denoted by ns, not significant; *, $p$ <0.05; **, $p$ <0.01; ***, $p$ <0.001 and ****, $p$ <0.0001. Con, non-targeting shRNA; Sh1 and Sh2, METTL3-targeting shRNAs.

**Additional file 3: Figure S3. Expression of M1 and M2 markers in METTL3-depleted THP-1 cells treated with polarising stimuli.** (**A**) Flow cytometry profiles and quantification (Mean Fluorescence Intensity (MFI)) after METTL3 depletion showing changes in the relative expression of M1 cell surface marker CD38 in THP-1-derived M1 macrophages. (**B**) Changes in gene expression following METTL3 depletion measured by qRT-PCR of M1 markers *CXCL10, IL-6*, *CD80* and *TNF* in THP-1-derived M1 cells. (**C**) Flow cytometry analysis of M2 cell surface marker CD209 in THP-1-derived M2 macrophages. (**D**) Expression of M2 markers CD209 and CD206 measured in THP-1-derived M2 cells by qRT-PCR following METTL3 depletion. All data are from at least three independent experiments. Bar plots show mean ± SEM. An unpaired two-tailed Student's t-test was used to determine significance, denoted by ns, not significant; *, $p$ <0.05; **, $p$ <0.01; ***, $p$ <0.001 and ****, $p$ <0.0001. Con, non-targeting shRNA; Sh1 and Sh2, METTL3-targeting shRNAs.

**Additional file 5: Figure S4. Association between m$^6$A changes and enrichment of gene functions relevant to polarised macrophages.** Gene ontology analysis on increased (top) and decreased (bottom) m$^6$A peaks during (**A**) Mϕ to M1 and (**B**) Mϕ to M2 polarisation. (**C**) Venn diagram showing overlapping gene sets identified by Gene Set Enrichment Analysis (GSEA) on m$^6$A-IP-Seq, mRNA-Seq and Polyribo-Seq data presenting positive (top) or negative (bottom) enrichment during Mϕ to M1. (**D**) GSEA signatures showing significantly positive and negative enrichment during Mϕ to M1 polarisation. (**E**) Similar Venn diagram shown in (**C**) for Mϕ to M2 polarisation. (**F**) GSEA signature showing significantly positive enrichment during Mϕ to M2 polarisation.

**Additional file 7: Figure S5. Association between changes in m$^6$A levels and expression of genes that regulate macrophage differentiation and function.** Left: Coverage plots of m$^6$A-

IP-Seq (top), RNA-Seq (middle) and Polyribo-Seq (bottom) for (**A**) *CSF1* in Mϕ, and (**B and C**) *CD163* and *COL6A2* in M2. $m^6A$-IP-Seq tracks show the overlay of input and IP data. $m^6A$ peak is highlighted within a dotted box. $m^6A$-IP-Seq coverage plots are displayed in BPM (bins per million reads, Bin size=1). mRNA-Seq and Polyribo-Seq coverage plots are displayed in reads per kilobase per million reads (RPKM). Right: Changes in gene expression following METTL3 depletion for (**A**) *CSF1* in Mϕ, and (**B-C**) *CD163* and *COL6A2* in M2. All data are from at least three independent experiments. Bar plots show mean ± SEM. An unpaired two-tailed Student's t-test was used to determine significance, denoted by *, $p < 0.05$; ***, $p < 0.001$ and ****, $p < 0.0001$. Con, non-targeting shRNA; Sh1 and Sh2, METTL3-targeting shRNAs.

**Additional file 8: Figure S6. Expression of M1 and M2 markers following TETs inhibition in THP-1-derived macrophages.** Flow cytometry profiles and quantification (Mean Fluorescence Intensity (MFI) or Percentage of Positive Cells following treatment with 2mM Itaconic Acid (ITA) showing changes in the relative expression of (**A**) Mϕ cell surface markers CD11b and CD44 in Mϕ cells, (**B**) M1 cell surface markers CD80 and CD38 in M1 cells and (**C**) M2 cell surface marker CD209 in M2 cells. All data are from at least three independent experiments. Bar plots show mean ± SEM. An unpaired two-tailed Student's t-test was used to determine significance, denoted by ns, not significant and $p < 0.05$; **.

**Additional file 10: Figure S7. Coexistence of 5hmC and $m^6A$ in genes that regulate macrophage differentiation and polarisation** (**A**) Number of increased (red) and decreased (blue) 5hmC peaks during macrophage differentiation and polarisation. (**B**) Significantly enriched sequence motifs identified by 5hmC-IP -Seq in mo, Mϕ, M1 and M2. (**C**) 5hmC and $m^6A$ peaks in *QKI, FOXO3, MCL1, VIM, FAM20C* and *MSDF12* transcripts identified in 5hmC-IP-Seq (top) and $m^6A$-IP-Seq (bottom) data from mo, Mϕ, M1 and M2. 5hmC- and $m^6A$-

IP-Seq tracks show the overlay of input and IP data. $m^6A$ and 5hmC peaks are highlighted within dotted boxes. 5hmC- and $m^6A$-IP-Seq coverage plots are displayed in BPM (bins per million reads, Bin size=1). (**D**) GO analysis on differential 5hmC peaks during Mϕ to M1 polarisation. (**E**) Number and percentages of transcripts bearing both $m^6A$ and 5hmC in mo, Mϕ, M1 and M2. (**F**) The proportion of differentially enriched 5hmC and $m^6A$ peaks that co-occurred in defined transcript regions (UTR, untranslated region; TSS, transcriptional start site, exon and intron) during M1 and M2 macrophage polarisation.

**Additional file 11: Figure S8. Half-life of *MYC* mRNA following treatment with Itaconic Acid.** (**A)** Coverage plots of 5hmC-IP-Seq (top) and mRNA-Seq (bottom) data for *MYC* in Mϕ. (**B**) mRNA decay plot. for *MYC in* Itaconic Acid (ITA)-treated Mϕ.

#### Additional file 1: Figure S1

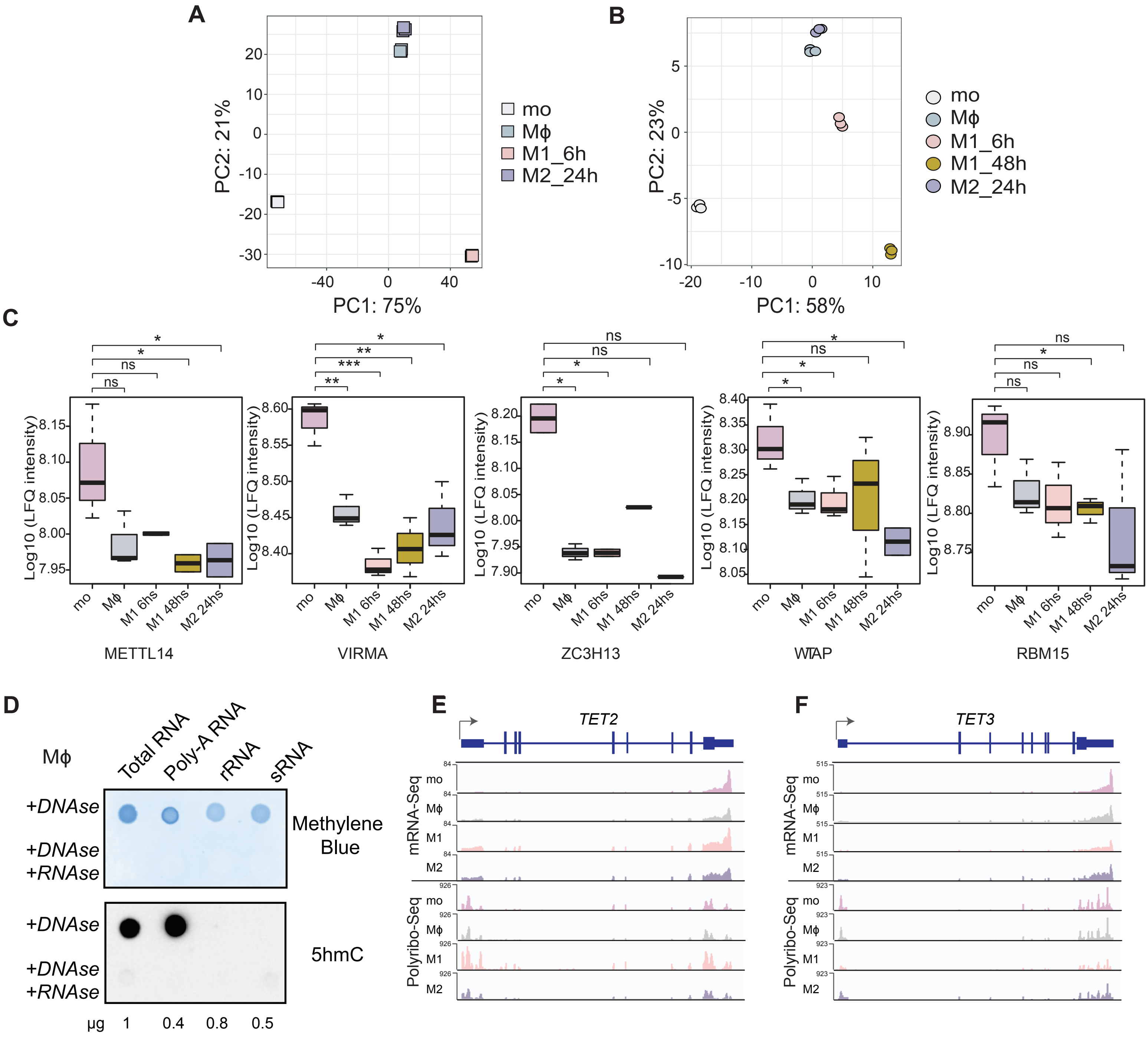

### Additional file 2: Figure S2

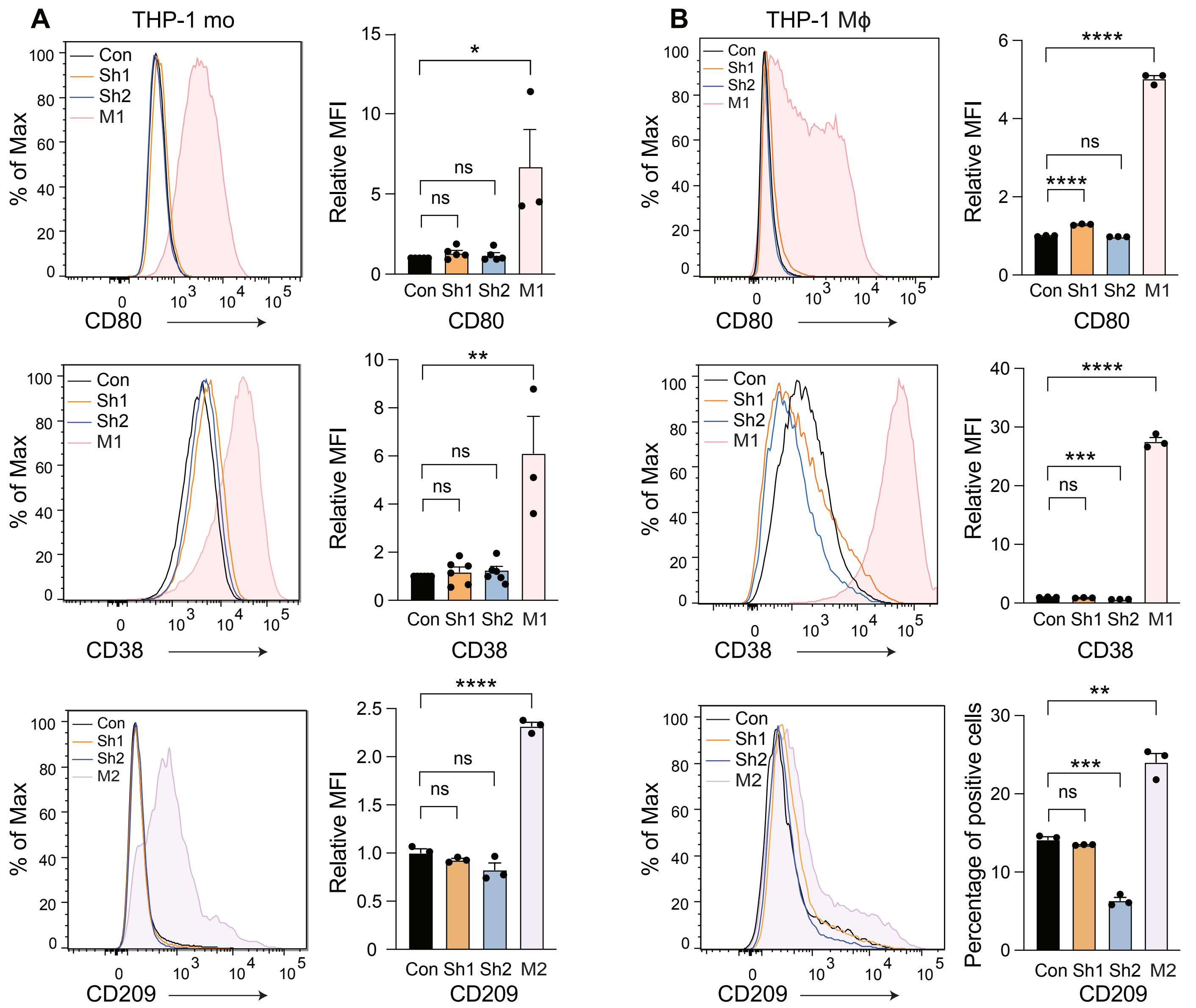

#### Additional file 3: Figure S3

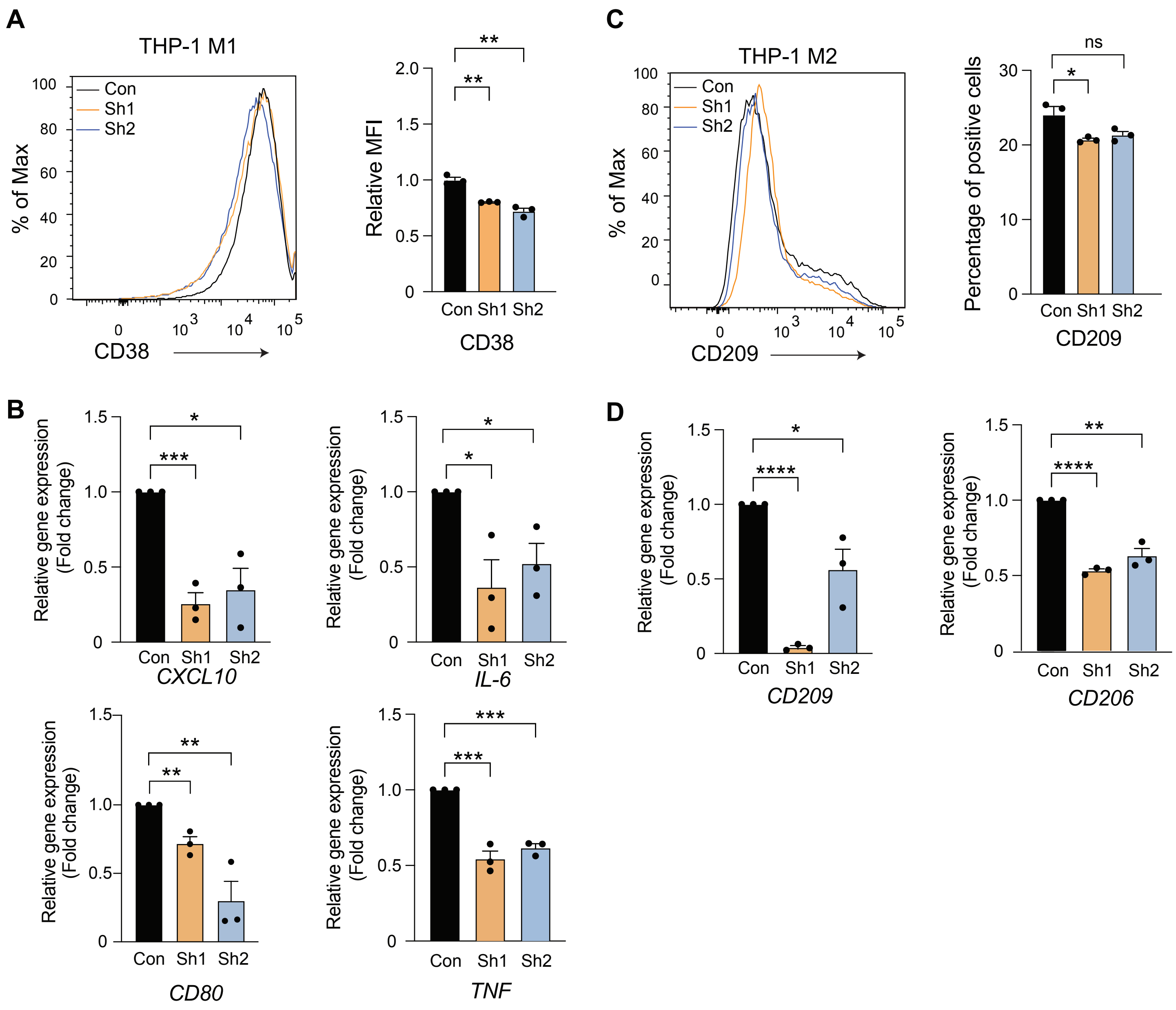

#### Additional file 5:Figure S4

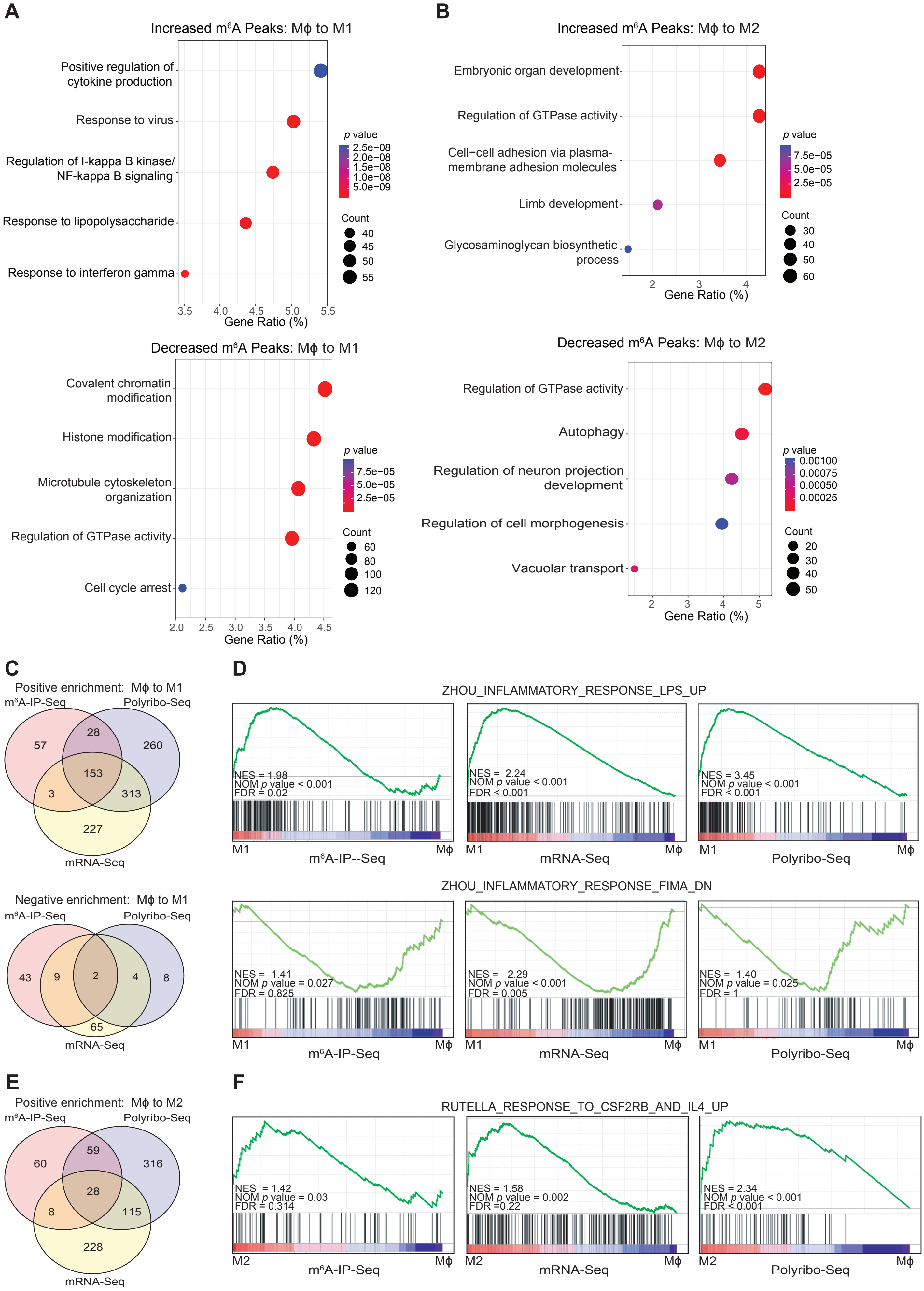

#### Additional file 7: Figure S5

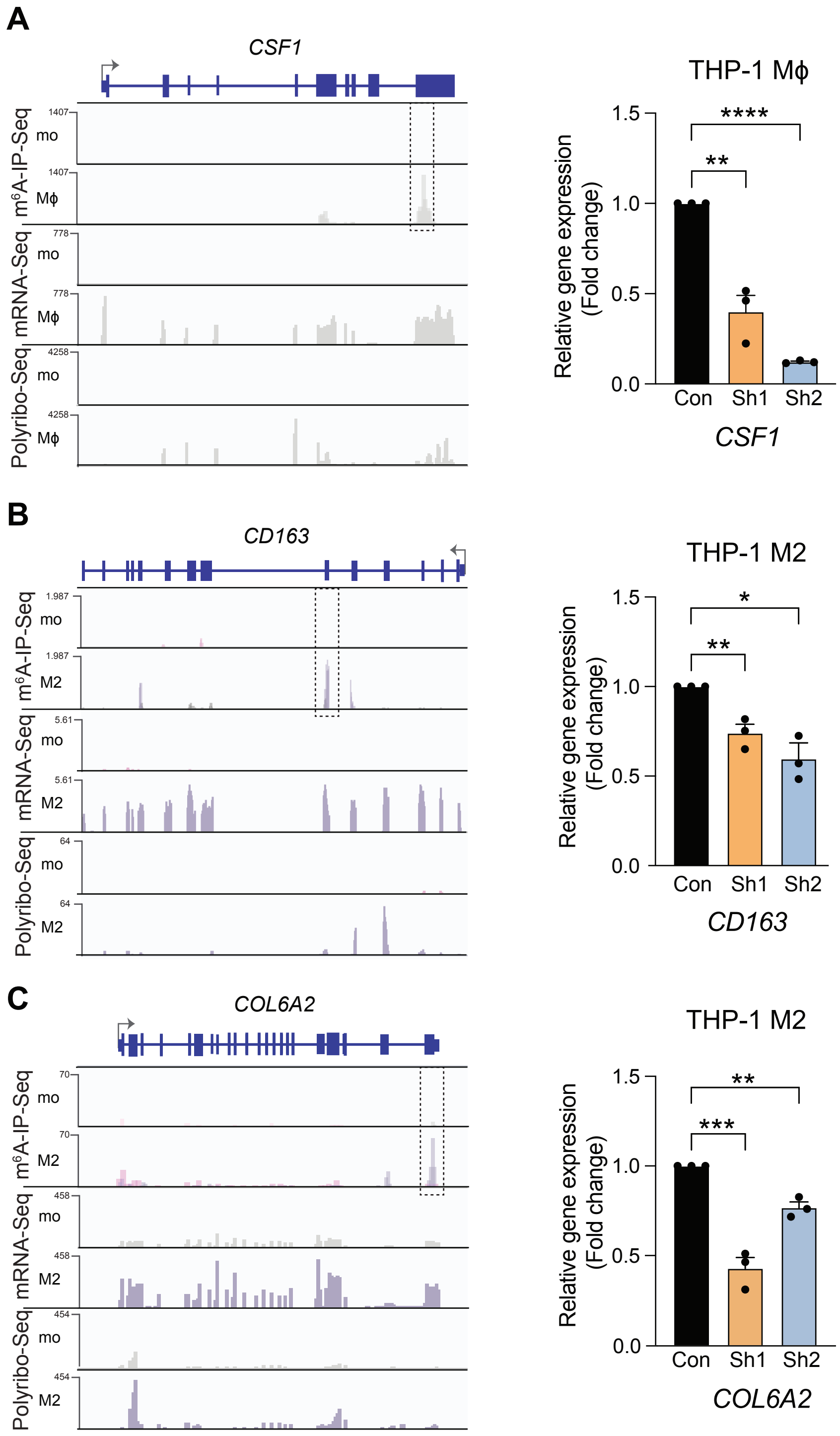

#### Additional file 8: Figure S6

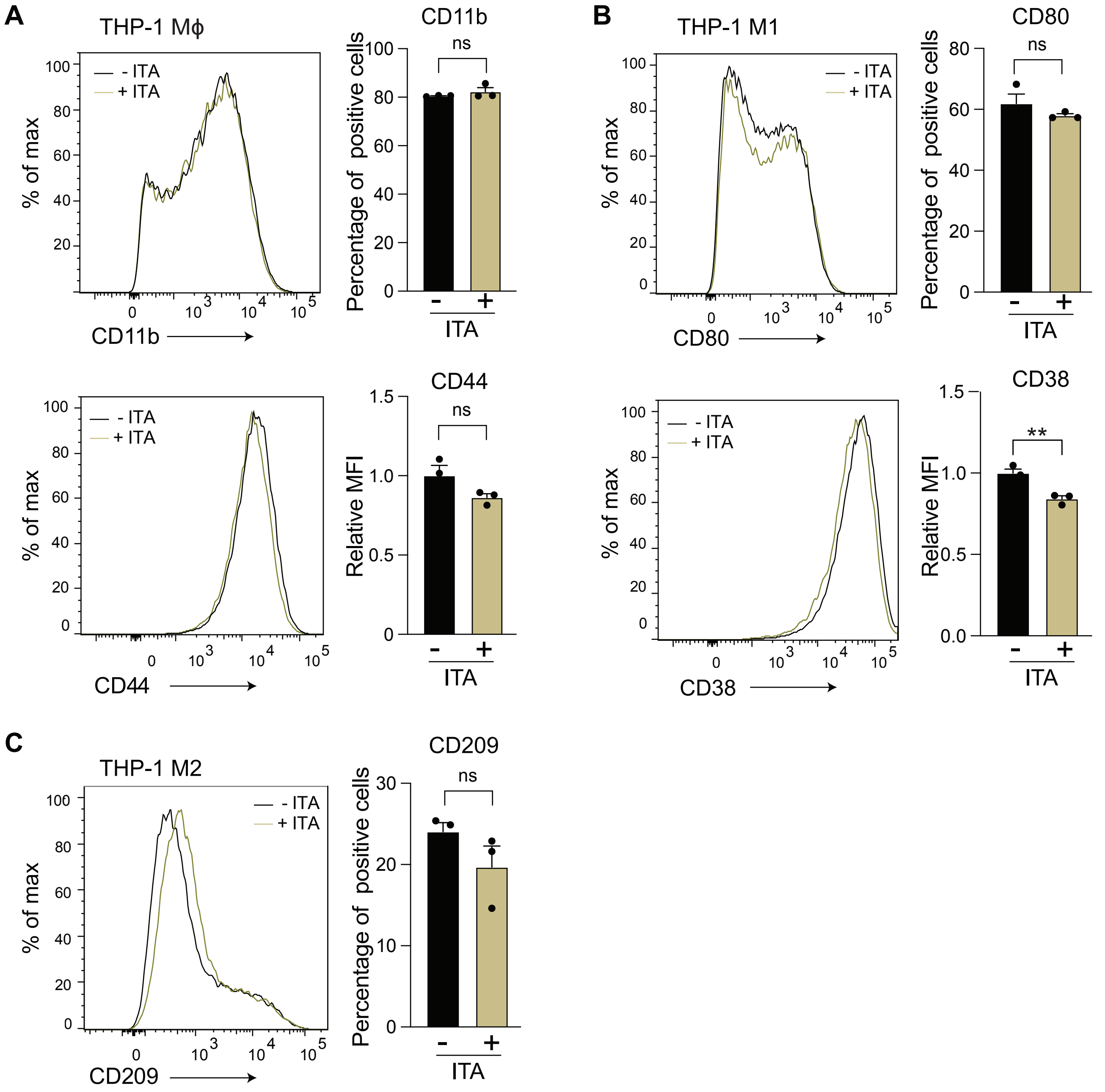

#### Additional file 10: Figure S7

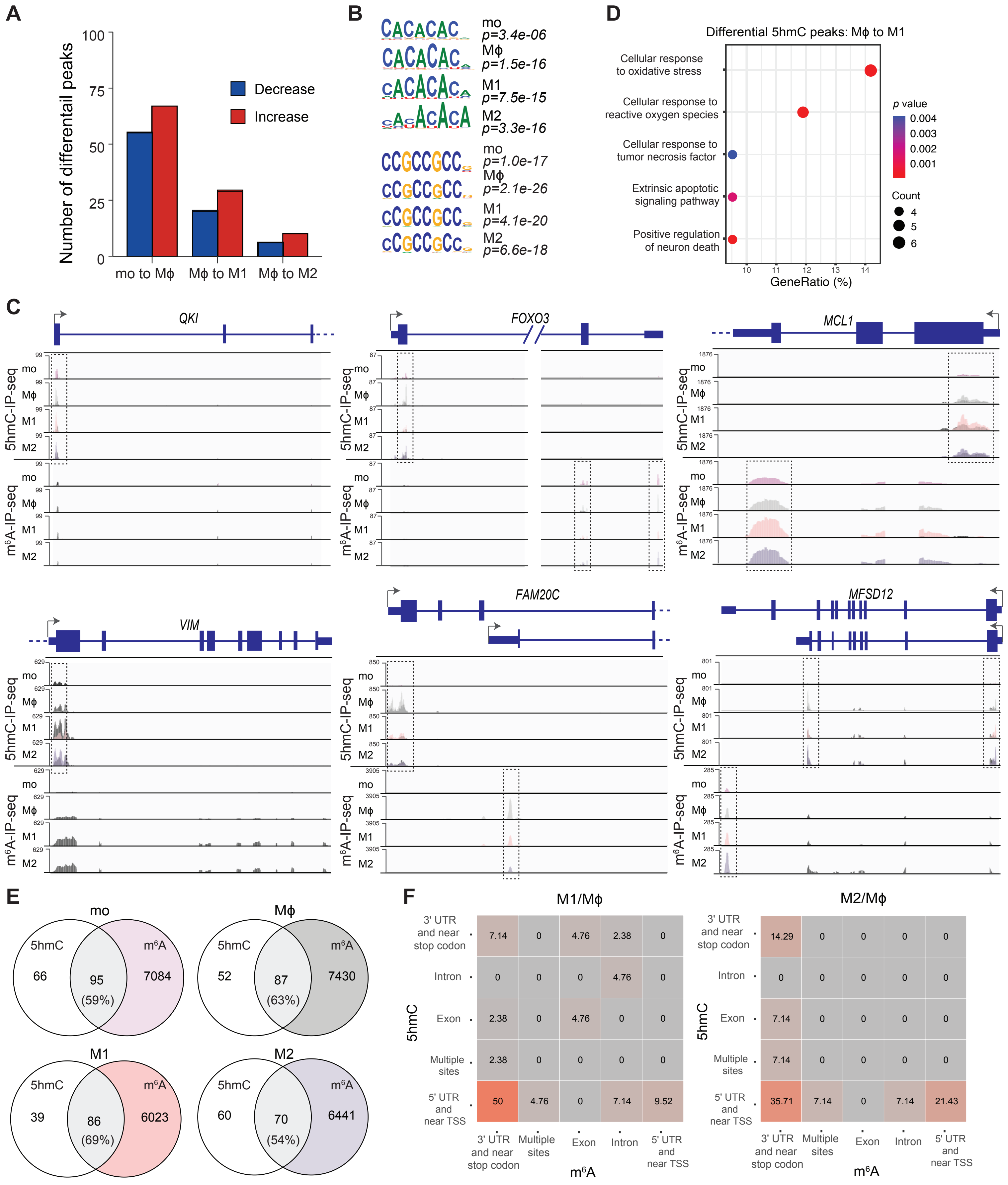

#### Additional file 11: Figure S8

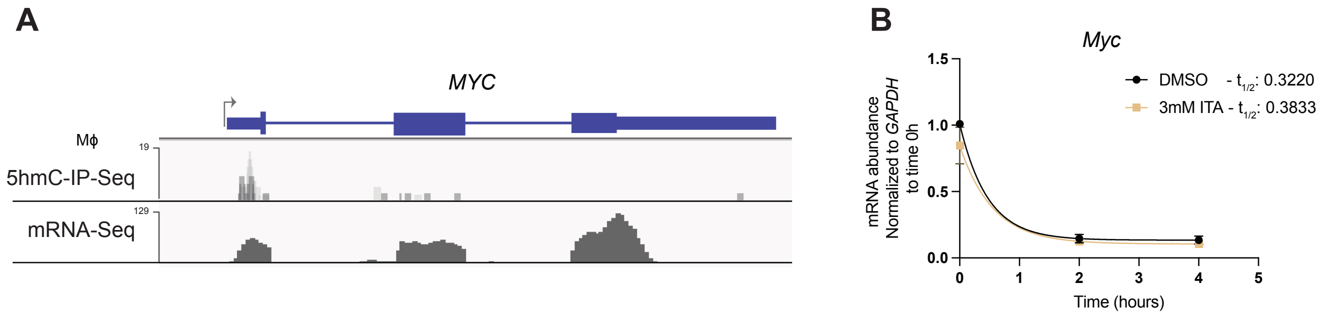
