## Supplementary Tables for "RNA m^6^A and 5hmC regulate monocyte and macrophage gene expression programs"

**Additional file 4: Table S1.** Lists of differentially enriched $m^6A$ peaks during macrophage differentiation and polarisation.

**Additional file 6: Table S2.** Lists of gene ontology terms representing genes with enriched $m^6A$ peaks in Mϕ, M1 and M2 populations used to generate enrichment maps in Fig. 4A.

**Additional file 9: Table S3.** Lists of RNA 5hmC-enriched peaks in monocytes and macrophages.

**Additional file 12: Table S4.** Oligonucleotide sequences.

**Additional file 4**
**Table S1: Lists of differentially enriched m6A peaks during macrophage differentiation and**

| Mϕ to mo | | | | |
|---|---|---|---|---|
| **chr** | **start** | **end** | **log2 fold change** | **Gene name** |
| 1 | 14539 | 14732 | -1.125339533 | WASH7P |
| 1 | 155659732 | 155659932 | -1.505965517 | YY1AP1 |
| 1 | 155659732 | 155659932 | -1.505965517 | RP11-29H23.5 |
| 22 | 31738499 | 31738718 | -1.775587866 | PRR14L |
| 22 | 31782959 | 31783179 | -5.948922728 | DEPDC5 |
| 22 | 31956966 | 31957162 | -2.620740771 | YWHAH |
| 22 | 32385197 | 32385359 | 2.407406497 | RP1-149A16.17 |
| 22 | 32541325 | 32541523 | -3.457589283 | SYN3 |
| 22 | 32546241 | 32546436 | -2.830544473 | SYN3 |
| 1 | 155660636 | 155660834 | -2.7540941 | YY1AP1 |
| 22 | 35347296 | 35347550 | 1.717398642 | TOM1 |
| 22 | 35424190 | 35424391 | -3.672686183 | MCM5 |
| 22 | 35551864 | 35552074 | 8.529207295 | RASD2 |
| 22 | 35667832 | 35668027 | 2.426500468 | APOL6 |
| 22 | 36227696 | 36227953 | 1.200364506 | APOL2 |
| 1 | 155689058 | 155689261 | -7.325862217 | DAP3 |
| 22 | 36236670 | 36236890 | 2.317685526 | APOL2 |
| 22 | 36265540 | 36265738 | 1.480949657 | APOL1 |
| 22 | 36282531 | 36282713 | -11.03674752 | MYH9 |
| 22 | 36330892 | 36331092 | -6.899235901 | MYH9 |
| 22 | 36490015 | 36490158 | -1.994361467 | FOXRED2 |
| 22 | 37029509 | 37029721 | -1.666590355 | MPST |
| 22 | 37182298 | 37182488 | 2.997546989 | C1QTNF6 |
| 22 | 37182298 | 37182488 | 2.997546989 | RP1-151B14.6 |
| 1 | 155738143 | 155738343 | -1.10057212 | DAP3 |
| 22 | 37314194 | 37314428 | 1.295594581 | CYTH4 |
| 22 | 37665716 | 37665917 | -1.536923622 | PDXP |
| 22 | 37665716 | 37665917 | -1.536923622 | SH3BP1 |
| 22 | 37734526 | 37734718 | -1.889553746 | TRIOBP |
| 22 | 37831455 | 37831645 | -2.510233154 | ANKRD54 |
| 1 | 155748247 | 155748450 | -1.153095379 | MSTO2P |
| 22 | 37940816 | 37941016 | 1.425323249 | MICALL1 |
| 22 | 38078115 | 38078310 | 2.966439548 | SLC16A8 |
| 22 | 38080102 | 38080355 | 1.758698146 | SLC16A8 |
| 22 | 38081711 | 38081913 | 1.121626904 | SLC16A8 |
| 22 | 38089654 | 38089847 | 3.906059688 | BAIAP2L2 |
| 22 | 38091962 | 38092151 | 2.141152438 | BAIAP2L2 |
| 22 | 38093639 | 38093808 | 2.085102103 | BAIAP2L2 |
| 1 | 155749784 | 155749961 | -1.32686022 | GON4L |
| 1 | 155749784 | 155749961 | -1.32686022 | MSTO2P |
| 22 | 38341039 | 38341149 | 3.25389732 | CSNK1E |
| 22 | 38341039 | 38341149 | 3.25389732 | RP3-449O17.1 |
| 22 | 38699898 | 38700105 | 1.137474351 | JOSD1 |
| 22 | 38741375 | 38741665 | -10.0530725 | SUN2 |
| 22 | 38741375 | 38741665 | -10.0530725 | RP3-508I15.14 |
| 22 | 38742668 | 38742944 | -9.161679141 | RP3-508I15.21 |
| 22 | 38742668 | 38742944 | -9.161679141 | SUN2 |
| 22 | 38742668 | 38742944 | -9.161679141 | RP3-508I15.14 |
| 1 | 155750401 | 155750624 | -1.654681677 | GON4L |

| M1 to Mϕ | | | | |
|---|---|---|---|---|
| **chr** | **start** | **end** | **log2 fold change** | **Gene name** |
| 1 | 912473 | 912664 | -5.157008352 | RP11-54O7.16 |
| 1 | 912473 | 912664 | -5.157008352 | RP11-54O7.1 |
| 22 | 31782959 | 31783179 | 6.106096186 | DEPDC5 |
| 22 | 31933732 | 31933908 | -2.566699563 | C22orf24 |
| 22 | 35347296 | 35347550 | -1.499685564 | TOM1 |
| 22 | 35551864 | 35552074 | -2.590948822 | RASD2 |
| 22 | 35659233 | 35659433 | 2.920553726 | APOL6 |
| 22 | 35667832 | 35668027 | 1.430813098 | APOL6 |
| 22 | 36226781 | 36226978 | 3.950888342 | APOL2 |
| 22 | 36227696 | 36227953 | 3.582451386 | APOL2 |
| 22 | 36236670 | 36236890 | 3.83915031 | APOL2 |
| 22 | 36265540 | 36265738 | 3.883800219 | APOL1 |
| 22 | 36266820 | 36267016 | 9.858496786 | APOL1 |
| 22 | 37029509 | 37029721 | -1.180868352 | MPST |
| 22 | 37225385 | 37225582 | -2.555746761 | RAC2 |
| 22 | 37287860 | 37288039 | -8.022666534 | CYTH4 |
| 22 | 37314194 | 37314428 | -1.321938493 | CYTH4 |
| 22 | 37314951 | 37315080 | -1.22202147 | CYTH4 |
| 22 | 37665716 | 37665917 | -1.499954877 | SH3BP1 |
| 22 | 37665716 | 37665917 | -1.499954877 | PDXP |
| 22 | 37757791 | 37758043 | 9.611116799 | TRIOBP |
| 22 | 37805818 | 37806021 | -2.213728968 | H1F0 |
| 22 | 37831455 | 37831645 | -1.240512657 | ANKRD54 |
| 22 | 38078115 | 38078310 | -1.175086284 | SLC16A8 |
| 22 | 38080102 | 38080355 | -1.184449511 | SLC16A8 |
| 22 | 38091962 | 38092151 | -1.659163994 | BAIAP2L2 |
| 22 | 38093639 | 38093808 | -4.099200549 | BAIAP2L2 |
| 22 | 38215038 | 38215237 | 2.343682319 | MAFF |
| 22 | 38341039 | 38341149 | -1.524181511 | RP3-449O17.1 |
| 22 | 38341039 | 38341149 | -1.524181511 | CSNK1E |
| 22 | 38673136 | 38673324 | -6.383988751 | RP3-508I15.9 |
| 22 | 38673136 | 38673324 | -6.383988751 | CBY1 |
| 22 | 38699898 | 38700105 | -1.300738191 | JOSD1 |
| 22 | 38739932 | 38740065 | 10.86749039 | SUN2 |
| 22 | 38739932 | 38740065 | 10.86749039 | RP3-508I15.14 |
| 22 | 38741375 | 38741665 | 13.30559167 | SUN2 |
| 22 | 38741375 | 38741665 | 13.30559167 | RP3-508I15.14 |
| 22 | 38742668 | 38742944 | 12.11307827 | RP3-508I15.14 |
| 22 | 38742668 | 38742944 | 12.11307827 | SUN2 |
| 22 | 38742668 | 38742944 | 12.11307827 | RP3-508I15.21 |
| 22 | 38785121 | 38785314 | 5.444371624 | SUN2 |
| 22 | 38785121 | 38785314 | 5.444371624 | DNAL4 |
| 22 | 39044232 | 39044421 | 1.696842256 | APOBEC3F |
| 22 | 39044232 | 39044421 | 1.696842256 | APOBEC3G |
| 22 | 39045240 | 39045438 | 2.484322338 | APOBEC3F |
| 22 | 39045240 | 39045438 | 2.484322338 | APOBEC3G |
| 22 | 39080257 | 39080549 | 2.159518195 | APOBEC3G |
| 22 | 39224879 | 39225079 | 2.674755807 | PDGFB |
| 22 | 39529220 | 39529418 | -7.712919375 | RPS19BP1 |

| M2 to Mϕ | | | | |
|---|---|---|---|---|
| **chr** | **start** | **end** | **log2 fold change** | **Gene name** |
| 22 | 32385197 | 32385359 | -7.247146547 | RP1-149A16.17 |
| 1 | 155660636 | 155660834 | -7.079225952 | YY1AP1 |
| 22 | 35424190 | 35424391 | 2.259417334 | MCM5 |
| 22 | 35551864 | 35552074 | 1.157726563 | RASD2 |
| 22 | 35667832 | 35668027 | -1.072901926 | APOL6 |
| 22 | 36266820 | 36267016 | 4.986489893 | APOL1 |
| 22 | 36282531 | 36282713 | 9.825830688 | MYH9 |
| 22 | 36330892 | 36331092 | 7.099069026 | MYH9 |
| 22 | 37287860 | 37288039 | -8.022666534 | CYTH4 |
| 22 | 37665716 | 37665917 | 1.290898272 | SH3BP1 |
| 22 | 37665716 | 37665917 | 1.290898272 | PDXP |
| 22 | 37757791 | 37758043 | 11.80182975 | TRIOBP |
| 22 | 38075412 | 38075610 | 1.815784134 | PICK1 |
| 22 | 38341039 | 38341149 | 1.227671138 | CSNK1E |
| 22 | 38341039 | 38341149 | 1.227671138 | RP3-449O17.1 |
| 22 | 38741375 | 38741665 | 9.970952217 | RP3-508I15.14 |
| 22 | 38741375 | 38741665 | 9.970952217 | SUN2 |
| 22 | 38742668 | 38742944 | 9.023886805 | RP3-508I15.21 |
| 22 | 38742668 | 38742944 | 9.023886805 | RP3-508I15.14 |
| 22 | 38742668 | 38742944 | 9.023886805 | SUN2 |
| 22 | 38785121 | 38785314 | 5.417235525 | SUN2 |
| 22 | 38785121 | 38785314 | 5.417235525 | DNAL4 |
| 22 | 38866865 | 38866989 | 11.77322023 | CBX6 |
| 22 | 39045240 | 39045438 | 1.200026536 | APOBEC3F |
| 22 | 39045240 | 39045438 | 1.200026536 | APOBEC3G |
| 22 | 39224879 | 39225079 | 3.293478519 | PDGFB |
| 22 | 39529220 | 39529418 | 2.063175004 | RPS19BP1 |
| 22 | 40405900 | 40406085 | 7.629135788 | SGSM3 |
| 22 | 40407938 | 40408129 | 7.013132033 | SGSM3 |
| 22 | 40418800 | 40418998 | 1.772499353 | MKL1 |
| 22 | 41125739 | 41125924 | -8.137607581 | EP300 |
| 22 | 41395913 | 41396131 | -1.077392223 | TEF |
| 22 | 41674095 | 41674321 | 1.031526939 | SNU13 |
| 1 | 155853395 | 155853566 | 1.866651114 | GON4L |
| 22 | 41694090 | 41694290 | 1.053787496 | C22orf46 |
| 22 | 41925158 | 41925384 | 1.036802782 | TNFRSF13C |
| 22 | 42059945 | 42060140 | 1.11589132 | NAGA |
| 22 | 42213210 | 42213410 | 1.44256189 | TCF20 |
| 22 | 42275143 | 42275351 | 1.022267425 | OGFRP1 |
| 22 | 42275143 | 42275351 | 1.022267425 | TCF20 |
| 22 | 44018531 | 44018758 | 5.428601736 | PARVB |
| 22 | 44185904 | 44186102 | 8.078930031 | PARVG |
| 22 | 44185904 | 44186102 | 8.078930031 | RP4-671O14.5 |
| 22 | 45240238 | 45240431 | 8.907641608 | KIAA0930 |
| 22 | 46040534 | 46040646 | 1.380024604 | LINC00899 |
| 22 | 46238294 | 46238413 | -5.944942085 | PPARA |
| 1 | 156076583 | 156076846 | 1.627426231 | MEX3A |
| 1 | 156177253 | 156177358 | 5.457939689 | SEMA4A |
| 22 | 49432901 | 49433082 | -1.321431952 | C22orf34 |

| | | | | |
|---|---|---|---|---|
| 22 | 38785121 | 38785314 | -6.450622016 | DNAL4 |
| 22 | 38785121 | 38785314 | -6.450622016 | SUN2 |
| 22 | 39224879 | 39225079 | 5.29248125 | PDGFB |
| 22 | 39430260 | 39430459 | -1.070191626 | TAB1 |
| 22 | 39488579 | 39488781 | 4.433595236 | MGAT3 |
| 22 | 39529220 | 39529418 | -1.692391544 | RPS19BP1 |
| 22 | 40405900 | 40406085 | -10.1734646 | SGSM3 |
| 22 | 40406916 | 40407115 | -9.163188409 | SGSM3 |
| 22 | 40407938 | 40408129 | -8.782841127 | SGSM3 |
| 22 | 40411416 | 40411718 | -2.004530372 | MKL1 |
| 22 | 40418800 | 40418998 | -2.175031474 | MKL1 |
| 22 | 40586890 | 40587073 | -2.006393953 | RPL4P6 |
| 22 | 40586890 | 40587073 | -2.006393953 | MKL1 |
| 22 | 40926292 | 40926493 | -1.08852846 | XPNPEP3 |
| 22 | 41230318 | 41230573 | -1.870534618 | CHADL |
| 22 | 41230318 | 41230573 | -1.870534618 | L3MBTL2 |
| 22 | 41246473 | 41246641 | -2.584443317 | RANGAP1 |
| 22 | 41436259 | 41436461 | -1.361436716 | TOB2 |
| 22 | 41460095 | 41460308 | -1.875273589 | PHF5A |
| 22 | 41528817 | 41529016 | -1.666936931 | POLR3H |
| 22 | 41528817 | 41529016 | -1.666936931 | ACO2 |
| 22 | 41531867 | 41532065 | -1.744143241 | POLR3H |
| 22 | 41534391 | 41534590 | -2.063898641 | POLR3H |
| 22 | 41600579 | 41600747 | -2.340878255 | DESI1 |
| 22 | 41674095 | 41674321 | -2.281071772 | SNU13 |
| 22 | 41676806 | 41677027 | -8.265386043 | SNU13 |
| 1 | 155853395 | 155853566 | -3.211216552 | GON4L |
| 22 | 41694090 | 41694290 | -1.953772774 | C22orf46 |
| 22 | 41695797 | 41695984 | -3.938769886 | C22orf46 |
| 22 | 41697307 | 41697521 | -1.277664788 | C22orf46 |
| 22 | 41925158 | 41925384 | -1.316390603 | TNFRSF13C |
| 22 | 42052721 | 42052917 | -2.412239779 | WBP2NL |
| 22 | 42056904 | 42057085 | -2.44155084 | WBP2NL |
| 22 | 42059945 | 42060140 | -1.6117169 | NAGA |
| 1 | 155900360 | 155900561 | 1.459781126 | RIT1 |
| 22 | 42213210 | 42213410 | -2.636394435 | TCF20 |
| 22 | 42692534 | 42692734 | 1.003008834 | A4GALT |
| 22 | 42871359 | 42871558 | 1.326445802 | PACSIN2 |
| 22 | 43132987 | 43133185 | -1.768517498 | MCAT |
| 22 | 44018531 | 44018758 | -8.011465628 | PARVB |
| 22 | 44185904 | 44186102 | -8.593965971 | PARVG |
| 22 | 44185904 | 44186102 | -8.593965971 | RP4-671O14.5 |
| 22 | 44727727 | 44727930 | -1.245357295 | PRR5-ARHGAP8 |
| 22 | 44727727 | 44727930 | -1.245357295 | PRR5 |
| 22 | 45170005 | 45170215 | 1.320120875 | NUP50 |
| 22 | 45178295 | 45178499 | -1.146203793 | NUP50 |
| 22 | 45184667 | 45184799 | -1.437250848 | NUP50 |
| 22 | 45185324 | 45185525 | -1.472630193 | NUP50 |
| 22 | 45194828 | 45195028 | -4.578977533 | KIAA0930 |
| 22 | 45340496 | 45340704 | -1.554199523 | FAM118A |
| 22 | 45422430 | 45422543 | -4.055596822 | RIBC2 |
| 22 | 45922516 | 45922644 | -3.771397371 | WNT7B |
| 22 | 46040534 | 46040646 | -3.098198606 | LINC00899 |
| 22 | 46044480 | 46044685 | -2.281844851 | LINC00899 |
| 22 | 40266041 | 40266240 | -1.119608832 | TNRC6B |
| 22 | 40323105 | 40323307 | -1.094163627 | TNRC6B |
| 22 | 40405900 | 40406085 | 8.005089202 | SGSM3 |
| 22 | 40407938 | 40408129 | 6.466476446 | SGSM3 |
| 22 | 40586890 | 40587073 | -5.817405525 | RPL4P6 |
| 22 | 40586890 | 40587073 | -5.817405525 | MKL1 |
| 22 | 41125739 | 41125924 | -1.693235957 | EP300 |
| 22 | 41395913 | 41396131 | -2.871073685 | TEF |
| 22 | 41397445 | 41397652 | -2.035647185 | TEF |
| 1 | 6620034 | 6620161 | -2.831005066 | PHF13 |
| 22 | 41697307 | 41697521 | -1.054006316 | C22orf46 |
| 22 | 42052721 | 42052917 | -6.272482216 | WBP2NL |
| 22 | 42056904 | 42057085 | -6.379319819 | WBP2NL |
| 22 | 42213210 | 42213410 | 1.069424034 | TCF20 |
| 22 | 42871359 | 42871558 | -1.93656669 | PACSIN2 |
| 22 | 43200430 | 43200664 | -1.161418535 | SCUBE1 |
| 22 | 43203536 | 43203868 | -1.291658199 | SCUBE1 |
| 22 | 44185904 | 44186102 | 6.011357287 | RP4-671O14.5 |
| 22 | 44185904 | 44186102 | 6.011357287 | PARVG |
| 22 | 44495636 | 44495817 | -2.161644472 | LDOC1L |
| 22 | 44496971 | 44497178 | -1.282662825 | LDOC1L |
| 22 | 44728864 | 44729062 | -2.835462983 | PRR5-ARHGAP8 |
| 22 | 44728864 | 44729062 | -2.835462983 | PRR5 |
| 22 | 44730196 | 44730359 | -1.932550429 | PRR5 |
| 22 | 44730196 | 44730359 | -1.932550429 | PRR5-ARHGAP8 |
| 22 | 45170005 | 45170215 | -1.532291744 | NUP50 |
| 22 | 45240238 | 45240431 | 9.428495397 | KIAA0930 |
| 1 | 155951139 | 155951340 | -1.83082998 | MIR6738 |
| 1 | 155951139 | 155951340 | -1.83082998 | ARHGEF2 |
| 22 | 46040534 | 46040646 | -4.269579406 | LINC00899 |
| 22 | 46044480 | 46044685 | -3.076564879 | LINC00899 |
| 22 | 46239386 | 46239635 | -1.020460128 | PPARA |
| 22 | 46262251 | 46262435 | -1.585912267 | PKDREJ |
| 1 | 156076583 | 156076846 | -6.934604125 | MEX3A |
| 22 | 46308412 | 46308655 | -7.436698024 | GTSE1 |
| 22 | 46331219 | 46331527 | -7.099606114 | TRMU |
| 22 | 46677471 | 46677675 | -2.484593321 | GRAMD4 |
| 22 | 46686235 | 46686431 | -3.671881561 | CERK |
| 1 | 156177253 | 156177358 | 5.866507661 | SEMA4A |
| 22 | 46762099 | 46762299 | -1.4450807 | CTA-29F11.1 |
| 22 | 49415090 | 49415278 | -3.664212803 | C22orf34 |
| 22 | 49415710 | 49415847 | -1.286161702 | C22orf34 |
| 22 | 49433881 | 49434047 | -2.170845675 | C22orf34 |
| 22 | 49435895 | 49436088 | -1.5 | C22orf34 |
| 22 | 49440585 | 49440784 | -4.159983289 | C22orf34 |
| 22 | 49459056 | 49459294 | -5.202216 | C22orf34 |
| 22 | 49460282 | 49460466 | -4.850219859 | C22orf34 |
| 22 | 49466125 | 49466312 | -6.560442921 | C22orf34 |
| 1 | 156200174 | 156200375 | 1.048704414 | SLC25A44 |
| 22 | 49593140 | 49593332 | -2.04541146 | C22orf34 |
| 22 | 49603465 | 49603663 | 1.536686593 | C22orf34 |
| 22 | 49607102 | 49607298 | -1.815628076 | C22orf34 |
| 22 | 49617326 | 49617428 | 5.357122759 | C22orf34 |
| 22 | 49900167 | 49900362 | 2.227526589 | ALG12 |
| 22 | 49435895 | 49436088 | -6.581824838 | C22orf34 |
| 22 | 49466125 | 49466312 | -1.022035123 | C22orf34 |
| 22 | 49543825 | 49543940 | -6.646160816 | C22orf34 |
| 22 | 49617326 | 49617428 | 3.350219859 | C22orf34 |
| 22 | 49885485 | 49885654 | -1.130002956 | ZBED4 |
| 22 | 49900167 | 49900362 | 2.652255521 | ALG12 |
| 22 | 50217525 | 50217686 | 1.014825308 | SELO |
| 22 | 50264521 | 50264672 | 1.48602443 | MAPK11 |
| 22 | 50483139 | 50483341 | -1.459597465 | ADM2 |
| 22 | 50503221 | 50503423 | 1.892514259 | LMF2 |
| 3 | 2099359 | 2099479 | -2.418250634 | CNTN4 |
| 3 | 2616930 | 2617042 | 5.75389732 | CNTN4 |
| 1 | 156642474 | 156642752 | 1.273021472 | RP11-284F21.10 |
| 1 | 156642474 | 156642752 | 1.273021472 | BCAN |
| 3 | 9747420 | 9747605 | 1.64547306 | BRPF1 |
| 3 | 9932550 | 9932765 | 1.204627136 | IL17RC |
| 3 | 9944467 | 9944667 | 1.216360451 | CRELD1 |
| 3 | 9949399 | 9949544 | 1.08926229 | PRRT3-AS1 |
| 3 | 9949399 | 9949544 | 1.08926229 | PRRT3 |
| 3 | 10005403 | 10005592 | 1.031640191 | EMC3-AS1 |
| 3 | 10005403 | 10005592 | 1.031640191 | EMC3 |
| 3 | 10068962 | 10069140 | 1.713132377 | FANCD2 |
| 3 | 10247993 | 10248181 | 1.155736154 | TATDN2 |
| 1 | 156723222 | 156723422 | 1.38020187 | ISG20L2 |
| 3 | 10332509 | 10332613 | 4.102176075 | ATP2B2 |
| 3 | 11203375 | 11203644 | -1.551179393 | HRH1 |
| 3 | 12935616 | 12935824 | 1.145377245 | IQSEC1 |
| 3 | 14125342 | 14125544 | 3.046539343 | TMEM43 |
| 3 | 14968521 | 14968809 | -6.13178132 | NR2C2 |
| 3 | 15043834 | 15044027 | 6.942657531 | NR2C2 |
| 3 | 15043834 | 15044027 | 6.942657531 | MRPS25 |
| 1 | 156728007 | 156728207 | 1.791314237 | ISG20L2 |
| 3 | 15052335 | 15052554 | 1.132853667 | MRPS25 |
| 3 | 15052335 | 15052554 | 1.132853667 | NR2C2 |
| 3 | 15256040 | 15256237 | -2.260821473 | SH3BP5-AS1 |
| 3 | 15256040 | 15256237 | -2.260821473 | SH3BP5 |
| 3 | 15272446 | 15272642 | 7.493298794 | SH3BP5 |
| 3 | 15797628 | 15797826 | -1.227574585 | ANKRD28 |
| 3 | 16319098 | 16319295 | 8.781255379 | OXNAD1 |
| 3 | 16319098 | 16319295 | 8.781255379 | RFTN1 |
| 3 | 16319353 | 16319504 | 8.014212056 | OXNAD1 |
| 3 | 16319353 | 16319504 | 8.014212056 | RFTN1 |
| 3 | 16319994 | 16320151 | 1.794537711 | RFTN1 |
| 3 | 16319994 | 16320151 | 1.794537711 | OXNAD1 |
| 3 | 17011389 | 17011595 | -1.035297575 | PLCL2 |
| 3 | 17485674 | 17485821 | -5.94208526 | TBC1D5 |
| 3 | 17618690 | 17618894 | -5.581824838 | TBC1D5 |
| 3 | 17654661 | 17654789 | 5.987386534 | TBC1D5 |
| 3 | 18444414 | 18444621 | 1.145960604 | TBC1D5 |
| 3 | 18444414 | 18444621 | 1.145960604 | SATB1 |
| 1 | 156914769 | 156914947 | 2.015312982 | PEAR1 |
| 3 | 20174686 | 20174871 | -1.341857026 | SGO1 |
| 3 | 20174686 | 20174871 | -1.341857026 | SGO1-AS1 |
| 3 | 21556254 | 21556406 | -3.946397883 | ZNF385D |

| | | | | |
|---|---|---|---|---|
| 22 | 46087293 | 46087430 | -4.367257647 | MIRLET7BHG |
| 22 | 46235324 | 46235527 | -1.231079257 | PPARA |
| 22 | 46238294 | 46238413 | -1.198914105 | PPARA |
| 22 | 46244724 | 46244910 | -2.503594074 | CDPF1 |
| 1 | 156076583 | 156076846 | -1.246847222 | MEX3A |
| 22 | 46308412 | 46308655 | -3.862657193 | GTSE1 |
| 22 | 46331219 | 46331527 | -4.007296351 | TRMU |
| 22 | 46362814 | 46363015 | 1.147055647 | CELSR1 |
| 22 | 46417031 | 46417172 | -1.667980577 | CELSR1 |
| 22 | 46596167 | 46596359 | -6.582139219 | GRAMD4 |
| 22 | 46597819 | 46598008 | -5.96831897 | GRAMD4 |
| 22 | 46686235 | 46686431 | -1.027929829 | CERK |
| 1 | 156177253 | 156177358 | -8.325736096 | SEMA4A |
| 22 | 46762099 | 46762299 | -2.395814811 | CTA-29F11.1 |
| 22 | 49415090 | 49415278 | 3.019459495 | C22orf34 |
| 22 | 49415710 | 49415847 | 3.360744923 | C22orf34 |
| 22 | 49435895 | 49436088 | 6.581824838 | C22orf34 |
| 22 | 49459056 | 49459294 | -1.055058846 | C22orf34 |
| 22 | 49460282 | 49460466 | -2.125422498 | C22orf34 |
| 1 | 156200174 | 156200375 | -1.605940548 | SLC25A44 |
| 22 | 49468882 | 49469052 | -1.278304196 | C22orf34 |
| 22 | 49543825 | 49543940 | -1.241129774 | C22orf34 |
| 22 | 49593140 | 49593332 | -1.644336267 | C22orf34 |
| 22 | 49603465 | 49603663 | -3.993106524 | C22orf34 |
| 22 | 49607102 | 49607298 | -2.9205253 | C22orf34 |
| 22 | 49609028 | 49609222 | -1.243877919 | C22orf34 |
| 22 | 49617326 | 49617428 | -8.570604542 | C22orf34 |
| 22 | 49823130 | 49823334 | -1.531783704 | BRD1 |
| 22 | 49883624 | 49883822 | -1.221656839 | ZBED4 |
| 22 | 49885485 | 49885654 | -1.949796082 | ZBED4 |
| 22 | 49886726 | 49886924 | -1.790866645 | ZBED4 |
| 22 | 49900167 | 49900362 | -3.756850641 | ALG12 |
| 22 | 49900970 | 49901162 | -1.43481201 | ALG12 |
| 22 | 50059999 | 50060195 | -4.213205147 | MLC1 |
| 22 | 50071151 | 50071318 | -5.401320317 | MLC1 |
| 22 | 50203872 | 50204070 | -1.121902452 | SELO |
| 22 | 50214933 | 50215133 | -1.44419367 | SELO |
| 22 | 50217525 | 50217686 | -2.626186543 | SELO |
| 1 | 156291612 | 156291820 | -2.145653577 | TMEM79 |
| 1 | 156291612 | 156291820 | -2.145653577 | GLMP |
| 22 | 50253183 | 50253355 | -2.004080515 | MAPK12 |
| 22 | 50264521 | 50264672 | -1.262964102 | MAPK11 |
| 22 | 50310452 | 50310655 | 6.954712178 | DENND6B |
| 22 | 50316284 | 50316541 | -2.742499534 | DENND6B |
| 22 | 50316284 | 50316541 | -2.742499534 | XX-C00717C00720L.1 |
| 22 | 50393939 | 50394118 | 1.106112821 | PPP6R2 |
| 22 | 50503221 | 50503423 | -2.869384461 | LMF2 |
| 1 | 156539116 | 156539313 | -1.259933736 | IQGAP3 |
| 22 | 50569486 | 50569689 | 1.844931772 | CHKB-CPT1B |
| 22 | 50569486 | 50569689 | 1.844931772 | CPT1B |
| 22 | 50571447 | 50571638 | 8.045345683 | CPT1B |
| 22 | 50571447 | 50571638 | 8.045345683 | CHKB-CPT1B |
| 22 | 50798853 | 50799047 | 1.700183966 | RPL23AP82 |
| 3 | 2099359 | 2099479 | -2.123330718 | CNTN4 |
| 22 | 49963095 | 49963307 | 1.378533622 | PIM3 |
| 22 | 50217525 | 50217686 | 1.276431052 | SELO |
| 1 | 6621563 | 6621876 | -1.634596215 | PHF13 |
| 1 | 156291612 | 156291820 | 1.530167832 | GLMP |
| 1 | 156291612 | 156291820 | 1.530167832 | TMEM79 |
| 22 | 50253183 | 50253355 | -1.464930768 | MAPK12 |
| 22 | 50483139 | 50483341 | -2.920929736 | ADM2 |
| 22 | 50523769 | 50523973 | 1.470716655 | CTA-384D8.36 |
| 22 | 50523769 | 50523973 | 1.470716655 | SCO2 |
| 1 | 156539116 | 156539313 | -3.023087316 | IQGAP3 |
| 22 | 50624956 | 50625162 | -1.078902295 | ARSA |
| 22 | 50798853 | 50799047 | -3.396342678 | RPL23AP82 |
| 3 | 2616930 | 2617042 | 4.775373393 | CNTN4 |
| 3 | 4303654 | 4303867 | -1.533941236 | SUMF1 |
| 3 | 4303654 | 4303867 | -1.533941236 | SETMAR |
| 3 | 4328791 | 4328917 | -6.274651684 | SUMF1 |
| 3 | 9371235 | 9371435 | -1.006123958 | THUMPD3-AS1 |
| 3 | 9371235 | 9371435 | -1.006123958 | THUMPD3 |
| 3 | 9734395 | 9734626 | -1.285034294 | BRPF1 |
| 3 | 9738915 | 9739116 | -1.237209961 | BRPF1 |
| 3 | 9835547 | 9835738 | -1.349261089 | TTLL3 |
| 3 | 9854727 | 9854902 | -1.08638484 | TTLL3 |
| 3 | 9932550 | 9932765 | -1.319022506 | IL17RC |
| 3 | 9944467 | 9944667 | -1.016973666 | CRELD1 |
| 3 | 9946003 | 9946209 | -1.169691306 | PRRT3 |
| 3 | 9949399 | 9949544 | -1.134695413 | PRRT3 |
| 3 | 9949399 | 9949544 | -1.134695413 | PRRT3-AS1 |
| 3 | 10005403 | 10005592 | -1.207939156 | EMC3 |
| 3 | 10005403 | 10005592 | -1.207939156 | EMC3-AS1 |
| 3 | 10007282 | 10007452 | -6.605335672 | EMC3 |
| 3 | 10007282 | 10007452 | -6.605335672 | AC022007.5 |
| 3 | 10142746 | 10142945 | -7.908541792 | VHL |
| 3 | 10328585 | 10328857 | -3.097713787 | ATP2B2 |
| 3 | 10332509 | 10332613 | -3.533044595 | ATP2B2 |
| 3 | 10377691 | 10377951 | -5.538507781 | ATP2B2 |
| 3 | 10449461 | 10449644 | -1.97944895 | ATP2B2 |
| 3 | 11193531 | 11193731 | -1.592530572 | HRH1 |
| 3 | 11203375 | 11203644 | -1.946409159 | HRH1 |
| 3 | 11204739 | 11204958 | -1.816770688 | HRH1 |
| 3 | 11206510 | 11206638 | -2.674001783 | HRH1 |
| 3 | 11259914 | 11260109 | -1.118113248 | HRH1 |
| 3 | 12503300 | 12503496 | -1.939308675 | TSEN2 |
| 3 | 12995238 | 12995414 | -7.306434249 | IQSEC1 |
| 3 | 14112989 | 14113187 | -1.14910964 | CHCHD4 |
| 3 | 14125342 | 14125544 | -6.493454312 | TMEM43 |
| 3 | 14158286 | 14158486 | -2.115992891 | XPC |
| 3 | 14158286 | 14158486 | -2.115992891 | AC093495.4 |
| 3 | 14968521 | 14968809 | -6.13178132 | NR2C2 |
| 3 | 15042782 | 15043022 | -1.104245557 | NR2C2 |
| 3 | 15042782 | 15043022 | -1.104245557 | MRPS25 |
| 3 | 15045656 | 15045854 | -6.738119953 | NR2C2 |
| 3 | 15045656 | 15045854 | -6.738119953 | MRPS25 |
| 3 | 15272446 | 15272642 | 8.954946542 | SH3BP5 |
| 1 | 156728770 | 156728994 | -1.082306154 | ISG20L2 |
| 3 | 21556254 | 21556406 | -3.946397883 | ZNF385D-AS1 |
| 3 | 21558913 | 21559024 | -2.548921464 | ZNF385D-AS1 |
| 3 | 21558913 | 21559024 | -2.548921464 | ZNF385D |
| 3 | 21636906 | 21637074 | -1.988639962 | ZNF385D |
| 3 | 21699079 | 21699263 | -6.139224729 | ZNF385D |
| 1 | 156937395 | 156937584 | -6.564641508 | ARHGEF11 |
| 3 | 21755252 | 21755429 | -1.289833529 | ZNF385D |
| 3 | 21837257 | 21837455 | -1.034518561 | ZNF385D |
| 3 | 21950537 | 21950746 | -1.826184565 | ZNF385D |
| 3 | 21950537 | 21950746 | -1.826184565 | ZNF385D-AS2 |
| 3 | 21981637 | 21981857 | -2.369955583 | ZNF385D |
| 3 | 21984115 | 21984288 | -2.538955244 | ZNF385D |
| 1 | 156938453 | 156938648 | -1.236253528 | ARHGEF11 |
| 3 | 21985978 | 21986176 | -6.99460854 | ZNF385D |
| 3 | 21986616 | 21986814 | -1.581234598 | ZNF385D |
| 3 | 21987037 | 21987237 | -1.268397609 | ZNF385D |
| 3 | 22048510 | 22048744 | -7.898577436 | ZNF385D |
| 3 | 22124968 | 22125148 | -8.494353437 | ZNF385D |
| 3 | 22126908 | 22127105 | -1.122249981 | ZNF385D |
| 3 | 22280036 | 22280228 | -6.949489602 | ZNF385D |
| 3 | 23544225 | 23544334 | -5.807354922 | UBE2E2 |
| 3 | 23962313 | 23962512 | -1.876444043 | NR1D2 |
| 3 | 26710036 | 26710165 | -5.083709073 | LRRC3B |
| 3 | 27633399 | 27633644 | 2.747214578 | AC098614.2 |
| 3 | 27717384 | 27717486 | 2.822170524 | EOMES |
| 3 | 28324105 | 28324306 | -1.03879656 | CMC1 |
| 3 | 28324105 | 28324306 | -1.03879656 | AZI2 |
| 3 | 32469469 | 32469569 | -5.198837316 | CMTM7 |
| 3 | 32570747 | 32570863 | 8.310439233 | DYNC1LI1 |
| 3 | 33798457 | 33798602 | 2.105255261 | PDCD6IP |
| 3 | 36831413 | 36831613 | -1.230602134 | TRANK1 |
| 3 | 36855638 | 36855806 | -1.156295115 | TRANK1 |
| 3 | 37282023 | 37282182 | -1.189941348 | GOLGA4 |
| 3 | 38411078 | 38411190 | -1.467153729 | XYLB |
| 1 | 158958218 | 158958476 | 2.168188069 | PYHIN1 |
| 3 | 39184213 | 39184412 | 4.11738898 | XIRP1 |
| 3 | 39185527 | 39185728 | 4.740601613 | XIRP1 |
| 3 | 39186682 | 39186876 | 3.952619368 | XIRP1 |
| 3 | 39187976 | 39188173 | 3.797975173 | XIRP1 |
| 3 | 39188843 | 39188950 | 3.790266364 | XIRP1 |
| 3 | 39265585 | 39265692 | 2.080386764 | CX3CR1 |
| 3 | 40324008 | 40324181 | 1.805717356 | ENTPD3-AS1 |
| 3 | 40517048 | 40517247 | 2.257013297 | ZNF620 |
| 3 | 41245072 | 41245251 | 7.062383768 | CTNNB1 |
| 3 | 42106114 | 42106378 | -1.813636653 | TRAK1 |
| 3 | 42638216 | 42638418 | -1.519532377 | RP4-613B23.1 |
| 3 | 42638216 | 42638418 | -1.519532377 | NKTR |
| 3 | 42659945 | 42660133 | 5.864172857 | ZBTB47 |
| 3 | 44917369 | 44917569 | 1.26182093 | ZDHHC3 |
| 3 | 45085541 | 45085731 | 3.18718098 | CDCP1 |
| 3 | 45224891 | 45225068 | 1.230664634 | TMEM158 |
| 3 | 45707590 | 45707824 | 2.095973797 | SACM1L |
| 1 | 159848980 | 159849154 | -5.211032383 | C1orf204 |
| 3 | 47380380 | 47380547 | 1.717429685 | RP11-708J19.1 |

| | | | | |
|---|---|---|---|---|
| 3 | 2616930 | 2617042 | -8.44686999 | CNTN4 |
| 1 | 156595361 | 156595562 | -1.840991676 | GPATCH4 |
| 3 | 4303654 | 4303867 | 1.174396248 | SUMF1 |
| 3 | 4303654 | 4303867 | 1.174396248 | SETMAR |
| 3 | 4983040 | 4983242 | 2.820819253 | BHLHE40 |
| 3 | 9371235 | 9371435 | -1.068358473 | THUMPD3 |
| 3 | 9371235 | 9371435 | -1.068358473 | THUMPD3-AS1 |
| 3 | 9384595 | 9384748 | -1.536518041 | THUMPD3 |
| 3 | 9384595 | 9384748 | -1.536518041 | THUMPD3-AS1 |
| 1 | 156642474 | 156642752 | 1.743792214 | RP11-284F21.10 |
| 1 | 156642474 | 156642752 | 1.743792214 | BCAN |
| 3 | 9747420 | 9747605 | -1.428818277 | BRPF1 |
| 3 | 9841527 | 9841739 | -1.106813665 | RPUSD3 |
| 3 | 9841527 | 9841739 | -1.106813665 | TTLL3 |
| 3 | 9854727 | 9854902 | 2.35449532 | TTLL3 |
| 3 | 9944467 | 9944667 | 1.165926895 | CRELD1 |
| 3 | 9946003 | 9946209 | 2.548633008 | PRRT3 |
| 3 | 9949399 | 9949544 | 2.50892603 | PRRT3 |
| 3 | 9949399 | 9949544 | 2.50892603 | PRRT3-AS1 |
| 3 | 10005403 | 10005592 | -2.384057092 | EMC3 |
| 3 | 10005403 | 10005592 | -2.384057092 | EMC3-AS1 |
| 3 | 10007282 | 10007452 | 1.514759543 | AC022007.5 |
| 3 | 10007282 | 10007452 | 1.514759543 | EMC3 |
| 3 | 10068962 | 10069140 | -5.097070619 | FANCD2 |
| 3 | 10142746 | 10142945 | -1.237935784 | VHL |
| 3 | 10260277 | 10260491 | -1.249718493 | TATDN2 |
| 3 | 10260277 | 10260491 | -1.249718493 | RP11-438J1.1 |
| 3 | 10279717 | 10279943 | -1.118096281 | TATDN2 |
| 3 | 10279717 | 10279943 | -1.118096281 | RP11-438J1.1 |
| 1 | 156723222 | 156723422 | -1.965133909 | ISG20L2 |
| 3 | 11193531 | 11193731 | 1.7313697 | HRH1 |
| 3 | 11203375 | 11203644 | 1.579560814 | HRH1 |
| 3 | 11204739 | 11204958 | 1.250757068 | HRH1 |
| 3 | 11206510 | 11206638 | 1.164056587 | HRH1 |
| 1 | 156724238 | 156724439 | -1.173489348 | ISG20L2 |
| 3 | 11259914 | 11260109 | 2.508984266 | HRH1 |
| 3 | 11260567 | 11260748 | 1.964559966 | HRH1 |
| 3 | 11468491 | 11468727 | 1.094474756 | ATG7 |
| 3 | 12503300 | 12503496 | -1.520473538 | TSEN2 |
| 3 | 12899068 | 12899357 | 1.304063344 | IQSEC1 |
| 3 | 13316887 | 13317090 | -2.547430451 | NUP210 |
| 3 | 14112989 | 14113187 | -1.528545902 | CHCHD4 |
| 1 | 156726986 | 156727189 | -1.141353305 | ISG20L2 |
| 3 | 14968521 | 14968809 | -1.545618852 | NR2C2 |
| 3 | 15043834 | 15044027 | -8.533423002 | MRPS25 |
| 3 | 15043834 | 15044027 | -8.533423002 | NR2C2 |
| 3 | 15045656 | 15045854 | -2.807458259 | NR2C2 |
| 3 | 15045656 | 15045854 | -2.807458259 | MRPS25 |
| 1 | 156728007 | 156728207 | -2.763675108 | ISG20L2 |
| 3 | 15052335 | 15052554 | -1.986970291 | MRPS25 |
| 3 | 15052335 | 15052554 | -1.986970291 | NR2C2 |
| 3 | 15256040 | 15256237 | 9.472773356 | SH3BP5-AS1 |
| 3 | 15256040 | 15256237 | 9.472773356 | SH3BP5 |
| 3 | 15670216 | 15670416 | 1.049328485 | ANKRD28 |
| 1 | 156728770 | 156728994 | -1.082306154 | RRNAD1 |
| 3 | 15670216 | 15670416 | -3.124727159 | ANKRD28 |
| 3 | 15797628 | 15797826 | -2.623026244 | ANKRD28 |
| 3 | 16319098 | 16319295 | 10.70856983 | RFTN1 |
| 3 | 16319098 | 16319295 | 10.70856983 | OXNAD1 |
| 3 | 16319353 | 16319504 | 10.4377067 | OXNAD1 |
| 3 | 16319353 | 16319504 | 10.4377067 | RFTN1 |
| 3 | 16319994 | 16320151 | 4.568981486 | RFTN1 |
| 3 | 16319994 | 16320151 | 4.568981486 | OXNAD1 |
| 3 | 17089866 | 17090052 | -1.238074507 | PLCL2 |
| 3 | 17490477 | 17490699 | -1.468204841 | TBC1D5 |
| 3 | 17618690 | 17618894 | -5.581824838 | TBC1D5 |
| 3 | 19484530 | 19484688 | 1.119732967 | KCNH8 |
| 3 | 20174686 | 20174871 | -2.032833459 | SGO1 |
| 3 | 20174686 | 20174871 | -2.032833459 | SGO1-AS1 |
| 3 | 21421151 | 21421350 | -1.6228311 | ZNF385D |
| 3 | 21556254 | 21556406 | -1.600816931 | ZNF385D-AS1 |
| 3 | 21556254 | 21556406 | -1.600816931 | ZNF385D |
| 3 | 21558913 | 21559024 | -1.807370577 | ZNF385D |
| 3 | 21558913 | 21559024 | -1.807370577 | ZNF385D-AS1 |
| 3 | 21636906 | 21637074 | -4.127060064 | ZNF385D |
| 3 | 21664771 | 21664974 | -1.143346716 | ZNF385D |
| 3 | 21683551 | 21683853 | -7.819458328 | ZNF385D |
| 3 | 21699079 | 21699263 | -6.139224729 | ZNF385D |
| 3 | 21727853 | 21728051 | -7.106741993 | ZNF385D |
| 3 | 21751132 | 21751328 | -2.12662762 | ZNF385D |
| 1 | 156937395 | 156937584 | -6.564641508 | ARHGEF11 |
| 3 | 21981637 | 21981857 | -1.332221428 | ZNF385D |
| 3 | 21984115 | 21984288 | -1.017848931 | ZNF385D |
| 3 | 21985978 | 21986176 | -1.217314114 | ZNF385D |
| 3 | 21987037 | 21987237 | -1.315761188 | ZNF385D |
| 3 | 22048510 | 22048744 | -1.381092004 | ZNF385D |
| 3 | 22124968 | 22125148 | -1.172876325 | ZNF385D |
| 3 | 22126908 | 22127105 | -1.656381594 | ZNF385D |
| 3 | 23269027 | 23269224 | 1.141670505 | UBE2E2 |
| 3 | 23962313 | 23962512 | -2.04978945 | NR1D2 |
| 3 | 23977281 | 23977447 | -2.249001747 | NR1D2 |
| 3 | 27633399 | 27633644 | -1.251854727 | AC098614.2 |
| 3 | 27717384 | 27717486 | 2.371944824 | EOMES |
| 3 | 30671862 | 30672086 | -1.327886816 | TGFBR2 |
| 3 | 32469469 | 32469569 | -5.198837316 | CMTM7 |
| 1 | 157124931 | 157125135 | 2.05888037 | ETV3 |
| 3 | 33798457 | 33798602 | 1.554987295 | PDCD6IP |
| 3 | 36992053 | 36992257 | -1.312765824 | EPM2AIP1 |
| 1 | 158184556 | 158184758 | -1.639581144 | CD1D |
| 3 | 38141411 | 38141617 | 2.197908082 | MYD88 |
| 3 | 38483819 | 38484061 | -1.006057288 | ACVR2B |
| 1 | 158958218 | 158958476 | 2.683274652 | PYHIN1 |
| 3 | 38540950 | 38541150 | -1.578701541 | EXOG |
| 3 | 39143677 | 39143881 | 1.565713667 | CSRNP1 |
| 3 | 39184213 | 39184412 | 6.004255523 | XIRP1 |
| 3 | 39185527 | 39185728 | 6.993659372 | XIRP1 |
| 3 | 39186682 | 39186876 | 6.636232067 | XIRP1 |
| 3 | 39187976 | 39188173 | 6.232812368 | XIRP1 |
| 3 | 48287054 | 48287194 | -1.378450104 | ZNF589 |
| 3 | 48460342 | 48460533 | 1.294264558 | ATRIP |
| 3 | 48561726 | 48561887 | 1.233361164 | UCN2 |
| 3 | 48561726 | 48561887 | 1.233361164 | PFKFB4 |
| 3 | 48562535 | 48562706 | 1.01739667 | UCN2 |
| 3 | 48681040 | 48681240 | 1.029033971 | NCKIPSD |
| 3 | 48989714 | 48989883 | 6.372416919 | RP13-131K19.2 |
| 3 | 49424527 | 49424699 | -1.214590719 | RP11-949J7.8 |
| 3 | 49424527 | 49424699 | -1.214590719 | NICN1 |
| 3 | 49642819 | 49643010 | 1.063674083 | BSN |
| 3 | 49653744 | 49653895 | 1.608227433 | BSN |
| 3 | 49655654 | 49655848 | 1.773655279 | BSN |
| 3 | 49657187 | 49657383 | 1.75056 | BSN |
| 3 | 49660802 | 49660998 | 1.662792309 | BSN |
| 3 | 49662072 | 49662225 | 1.599400672 | BSN |
| 3 | 49685279 | 49685478 | 2.870733493 | MST1 |
| 3 | 49686808 | 49686916 | 3.723382603 | MST1 |
| 3 | 49687198 | 49687397 | 6.190771476 | MST1 |
| 3 | 49721010 | 49721203 | 1.402295723 | RNF123 |
| 3 | 49721010 | 49721203 | 1.402295723 | GMPPB |
| 3 | 50294927 | 50295126 | 2.703408496 | HYAL3 |
| 3 | 50358985 | 50359185 | 1.153714263 | TMEM115 |
| 3 | 50358985 | 50359185 | 1.153714263 | XXcos-LUCA11.5 |
| 3 | 50607529 | 50607752 | 5.531494958 | CISH |
| 3 | 51652717 | 51652919 | -1.24633419 | RAD54L2 |
| 3 | 51653557 | 51653679 | -2.894247947 | RAD54L2 |
| 3 | 51957551 | 51957754 | 1.417437805 | PCBP4 |
| 3 | 52050765 | 52050961 | 9.667960112 | DUSP7 |
| 3 | 52054097 | 52054277 | 10.42520179 | DUSP7 |
| 3 | 56728546 | 56728766 | 1.085994749 | ARHGEF3 |
| 1 | 7791782 | 7791971 | -1.044255085 | PER3 |
| 1 | 160750025 | 160750223 | 5.079935668 | SLAMF7 |
| 3 | 57298137 | 57298342 | -1.743116353 | DNAH12 |
| 3 | 57912407 | 57912590 | -1.443358297 | SLMAP |
| 3 | 58123390 | 58123595 | 1.188157146 | FLNB |
| 3 | 58501468 | 58501664 | 1.292021295 | KCTD6 |
| 3 | 66380388 | 66380590 | 1.07899582 | LRIG1 |
| 3 | 66380388 | 66380590 | 1.07899582 | SLC25A26 |
| 3 | 68992206 | 68992393 | -2.345063672 | EOGT |
| 3 | 69417513 | 69417704 | -7.196862809 | FRMD4B |
| 3 | 71582843 | 71582990 | 8.253192535 | RP11-154H23.4 |
| 3 | 71582843 | 71582990 | 8.253192535 | FOXP1 |
| 3 | 72378400 | 72378599 | 1.068872468 | RYBP |
| 3 | 75484431 | 75484532 | -1.106673641 | ENPP7P2 |
| 3 | 77098076 | 77098217 | 7.558171981 | ROBO2 |
| 1 | 161048095 | 161048195 | 8.987599322 | ARHGAP30 |
| 3 | 82288606 | 82288785 | -3.660964047 | RP11-260O18.1 |
| 3 | 82402312 | 82402472 | -2.529807428 | RP11-260O18.1 |
| 3 | 85158372 | 85158499 | -5.61860498 | CADM2 |
| 1 | 161120694 | 161120877 | 1.259059935 | NIT1 |
| 3 | 97905025 | 97905219 | -5.114409345 | CRYBG3 |
| 3 | 99850080 | 99850245 | 1.230907135 | FILIP1L |
| 3 | 99850080 | 99850245 | 1.230907135 | CMSS1 |
| 3 | 101677016 | 101677234 | 1.211249625 | ZBTB11-AS1 |

| | | | | |
|---|---|---|---|---|
| 3 | 15797628 | 15797826 | 1.615550379 | ANKRD28 |
| 3 | 16319098 | 16319295 | -4.658706307 | OXNAD1 |
| 3 | 16319098 | 16319295 | -4.658706307 | RFTN1 |
| 3 | 16323790 | 16323910 | -8.275733976 | OXNAD1 |
| 3 | 16323790 | 16323910 | -8.275733976 | RFTN1 |
| 3 | 17089866 | 17090052 | -1.42979244 | PLCL2 |
| 3 | 17485674 | 17485821 | -1.473543618 | TBC1D5 |
| 3 | 17613156 | 17613356 | -2.067976893 | TBC1D5 |
| 3 | 17613872 | 17614040 | -5.486130924 | TBC1D5 |
| 3 | 17618690 | 17618894 | -1.753558602 | TBC1D5 |
| 3 | 18349196 | 18349399 | -1.785550978 | TBC1D5 |
| 3 | 18349196 | 18349399 | -1.785550978 | SATB1 |
| 3 | 18444414 | 18444621 | -1.711029318 | SATB1 |
| 3 | 18444414 | 18444621 | -1.711029318 | TBC1D5 |
| 3 | 19484530 | 19484688 | -3.821928095 | KCNH8 |
| 3 | 20174686 | 20174871 | -2.08900245 | SGO1-AS1 |
| 3 | 20174686 | 20174871 | -2.08900245 | SGO1 |
| 3 | 21421151 | 21421350 | 7.290247064 | ZNF385D |
| 3 | 21556254 | 21556406 | 6.399843181 | ZNF385D |
| 3 | 21556254 | 21556406 | 6.399843181 | ZNF385D-AS1 |
| 3 | 21558913 | 21559024 | 9.038366053 | ZNF385D-AS1 |
| 3 | 21558913 | 21559024 | 9.038366053 | ZNF385D |
| 3 | 21636906 | 21637074 | 6.477279923 | ZNF385D |
| 3 | 21664771 | 21664974 | -2.788315899 | ZNF385D |
| 3 | 21683551 | 21683853 | -1.998863717 | ZNF385D |
| 3 | 21699079 | 21699263 | -1.542141023 | ZNF385D |
| 1 | 156937395 | 156937584 | 6.564641508 | ARHGEF11 |
| 3 | 21755252 | 21755429 | 7.672269324 | ZNF385D |
| 3 | 21906674 | 21906811 | 8.553051412 | ZNF385D |
| 3 | 21949821 | 21950028 | 7.089998962 | ZNF385D-AS2 |
| 3 | 21949821 | 21950028 | 7.089998962 | ZNF385D |
| 3 | 21950537 | 21950746 | 5.178664723 | ZNF385D |
| 3 | 21950537 | 21950746 | 5.178664723 | ZNF385D-AS2 |
| 3 | 21980661 | 21980864 | 5.833588632 | ZNF385D |
| 3 | 21983450 | 21983651 | 5.578568185 | ZNF385D |
| 3 | 21984115 | 21984288 | 4.62591871 | ZNF385D |
| 1 | 156938453 | 156938648 | 1.246298452 | ARHGEF11 |
| 3 | 21986616 | 21986814 | 3.777965581 | ZNF385D |
| 3 | 22036982 | 22037178 | 8.334366332 | ZNF385D |
| 3 | 22124968 | 22125148 | 8.494353437 | ZNF385D |
| 3 | 22126908 | 22127105 | 5.645186681 | ZNF385D |
| 3 | 23269027 | 23269224 | -1.064932036 | UBE2E2 |
| 3 | 23352163 | 23352298 | 6.890167354 | UBE2E2 |
| 3 | 23364104 | 23364300 | 2.642947682 | UBE2E2 |
| 3 | 23962313 | 23962512 | 1.917948119 | NR1D2 |
| 3 | 23977281 | 23977447 | 8.236567475 | NR1D2 |
| 3 | 25794147 | 25794348 | -1.459108519 | OXSM |
| 3 | 27633399 | 27633644 | -1.803606437 | AC098614.2 |
| 1 | 157123313 | 157123511 | 1.471114287 | ETV3 |
| 3 | 32469469 | 32469569 | -1.572618157 | CMTM7 |
| 3 | 32753438 | 32753538 | -4.306521982 | CNOT10 |
| 3 | 32753438 | 32753538 | -4.306521982 | SUGT1P2 |
| 3 | 33093101 | 33093304 | -1.144860298 | GLB1 |
| 3 | 33093101 | 33093304 | -1.144860298 | TMPPE |
| 3 | 39188843 | 39188950 | 6.515546748 | XIRP1 |
| 3 | 39396827 | 39397029 | -1.054416603 | SLC25A38 |
| 3 | 40535342 | 40535530 | -1.299196573 | ZNF621 |
| 3 | 41245072 | 41245251 | 8.694345014 | CTNNB1 |
| 3 | 42106114 | 42106378 | -1.590823743 | TRAK1 |
| 3 | 42223363 | 42223562 | -1.561522329 | TRAK1 |
| 3 | 42536963 | 42537075 | -5.697747942 | VIPR1 |
| 3 | 42582480 | 42582669 | 4.766664866 | SS18L2 |
| 3 | 42582480 | 42582669 | 4.766664866 | SEC22C |
| 3 | 42638216 | 42638418 | -1.613158287 | NKTR |
| 3 | 42638216 | 42638418 | -1.613158287 | RP4-613B23.1 |
| 3 | 42659076 | 42659279 | -2.427057438 | ZBTB47 |
| 3 | 43303256 | 43303542 | -1.047912538 | SNRK |
| 3 | 44445677 | 44445917 | -2.592907225 | ZNF445 |
| 3 | 44446588 | 44446702 | -2.055931933 | ZNF445 |
| 3 | 44448162 | 44448353 | -1.879415463 | ZNF445 |
| 3 | 44455244 | 44455418 | -1.933854784 | ZNF445 |
| 1 | 159809898 | 159810097 | 5.361136599 | FCRL6 |
| 3 | 44569907 | 44570135 | -4.117737886 | ZKSCAN7 |
| 3 | 44569907 | 44570135 | -4.117737886 | RP11-944L7.4 |
| 3 | 44644832 | 44645057 | -1.05280591 | ZNF197 |
| 3 | 44644832 | 44645057 | -1.05280591 | RP11-944L7.4 |
| 3 | 45012908 | 45013079 | -2.002719737 | CLEC3B |
| 3 | 45012908 | 45013079 | -2.002719737 | EXOSC7 |
| 1 | 159835199 | 159835379 | 2.99057821 | C1orf204 |
| 1 | 159835199 | 159835379 | 2.99057821 | SLAMF8 |
| 3 | 45224891 | 45225068 | -1.049059069 | TMEM158 |
| 3 | 45595164 | 45595357 | 1.322361786 | LIMD1 |
| 3 | 45707590 | 45707824 | 1.267564046 | SACM1L |
| 1 | 159836080 | 159836278 | 3.092380362 | C1orf204 |
| 1 | 159836080 | 159836278 | 3.092380362 | SLAMF8 |
| 3 | 46203039 | 46203234 | 1.155358036 | CCR1 |
| 3 | 46203039 | 46203234 | 1.155358036 | CCR3 |
| 3 | 47122357 | 47122633 | -1.093836407 | SETD2 |
| 1 | 159848980 | 159849154 | 1.530434603 | C1orf204 |
| 3 | 47468711 | 47468895 | -3.025792497 | SCAP |
| 3 | 47495950 | 47496136 | -1.37259305 | ELP6 |
| 3 | 47731265 | 47731539 | -1.618362018 | SMARCC1 |
| 3 | 48269712 | 48269908 | -1.888803789 | ZNF589 |
| 3 | 48286434 | 48286664 | -6.774772019 | ZNF589 |
| 3 | 48290997 | 48291204 | -1.129993844 | ZNF589 |
| 3 | 48290997 | 48291204 | -1.129993844 | FCF1P2 |
| 3 | 48404083 | 48404272 | -1.525393748 | PLXNB1 |
| 3 | 48519541 | 48519757 | -1.905496693 | PFKFB4 |
| 3 | 48561726 | 48561887 | -1.044449955 | PFKFB4 |
| 3 | 48561726 | 48561887 | -1.044449955 | UCN2 |
| 3 | 48562535 | 48562706 | -5.370761583 | UCN2 |
| 3 | 48639844 | 48640021 | -1.417679645 | CELSR3 |
| 3 | 48659739 | 48659930 | -1.131820354 | CELSR3 |
| 1 | 160030952 | 160031092 | -1.117346953 | PIGM |
| 1 | 160030952 | 160031092 | -1.117346953 | KCNJ10 |
| 3 | 48688278 | 48688517 | -1.059897215 | IP6K2 |
| 3 | 48751434 | 48751636 | -1.047652969 | PRKAR2A |
| 3 | 48851566 | 48851767 | -2.748983562 | PRKAR2A-AS1 |
| 3 | 101677016 | 101677234 | 1.211249625 | ZBTB11 |
| 3 | 101712373 | 101712630 | 1.137406867 | PDCL3P4 |
| 3 | 101801330 | 101801522 | -5.9291848 | NXPE3 |
| 3 | 105367034 | 105367327 | 2.218237542 | ALCAM |
| 3 | 105574393 | 105574589 | -1.310931907 | ALCAM |
| 3 | 107773317 | 107773450 | -1.754049419 | BBX |
| 3 | 111133749 | 111133945 | 4.667360501 | NECTIN3 |
| 3 | 113954030 | 113954140 | 3.073949348 | ZDHHC23 |
| 3 | 119415953 | 119416154 | 1.139509452 | ARHGAP31 |
| 3 | 120094942 | 120095074 | 2.043470733 | RP11-18H7.1 |
| 3 | 120735627 | 120735809 | -6.36161715 | RABL3 |
| 3 | 121488011 | 121488207 | 5.613507097 | POLQ |
| 3 | 121696145 | 121696341 | -1.011974612 | GOLGB1 |
| 3 | 121697614 | 121697814 | -1.24165476 | GOLGB1 |
| 3 | 122396125 | 122396332 | -3.971257253 | FAM162A |
| 3 | 122728275 | 122728474 | -1.397524113 | PARP14 |
| 3 | 122839262 | 122839414 | -5.10715956 | DIRC2 |
| 1 | 161442688 | 161442876 | 2.026131596 | RP11-122G18.10 |
| 3 | 124796486 | 124796696 | 10.52205562 | ITGB5 |
| 3 | 125083333 | 125083528 | -1.207748957 | SLC12A8 |
| 1 | 161457562 | 161457735 | -5.283978038 | RP11-122G18.10 |
| 3 | 126269236 | 126269432 | -4.832667959 | RP11-71E19.1 |
| 3 | 126351992 | 126352116 | -1.227318249 | KLF15 |
| 3 | 126542824 | 126543050 | 1.438864667 | CHST13 |
| 3 | 126542824 | 126543050 | 1.438864667 | C3orf22 |
| 3 | 126989494 | 126989693 | 8.720539989 | PLXNA1 |
| 3 | 127672282 | 127672481 | 10.10613953 | PODXL2 |
| 3 | 128071006 | 128071206 | 10.48615512 | RUVBL1 |
| 3 | 128071006 | 128071206 | 10.48615512 | SEC61A1 |
| 3 | 128480786 | 128480987 | -1.234603904 | GATA2 |
| 3 | 128501872 | 128502036 | -1.917891548 | GATA2-AS1 |
| 3 | 128538512 | 128538747 | -1.026674385 | TMED10P2 |
| 3 | 128865188 | 128865303 | -1.390017158 | RP11-723O4.2 |
| 3 | 128880983 | 128881183 | 4.75389732 | ACAD9 |
| 3 | 129122158 | 129122306 | 8.114709844 | ISY1-RAB43 |
| 3 | 129122158 | 129122306 | 8.114709844 | RAB43 |
| 1 | 161749596 | 161749764 | 1.44051624 | DUSP12 |
| 3 | 129847537 | 129847738 | -1.352407732 | RP11-332M2.1 |
| 3 | 129847537 | 129847738 | -1.352407732 | TMCC1 |
| 3 | 131024525 | 131024635 | 2.012939316 | ASTE1 |
| 3 | 131024701 | 131024880 | 1.630370082 | ASTE1 |
| 3 | 131383249 | 131383458 | 1.261303121 | NUDT16 |
| 3 | 136370795 | 136370899 | -5.26569073 | STAG1 |
| 3 | 136819323 | 136819502 | 1.565198319 | SLC35G2 |
| 3 | 138295999 | 138296197 | -1.084421078 | ARMC8 |
| 3 | 138295999 | 138296197 | -1.084421078 | NME9 |
| 3 | 138347963 | 138348161 | -1.255949519 | MRAS |
| 3 | 138349669 | 138349870 | 2.538407799 | MRAS |
| 3 | 139406777 | 139406966 | -2.388194286 | RP11-319G6.1 |
| 3 | 139407484 | 139407654 | -1.777849789 | RP11-319G6.1 |
| 3 | 140973670 | 140973872 | -1.298523987 | SLC25A36 |
| 3 | 141443821 | 141444022 | -1.03441937 | ZBTB38 |
| 3 | 142135547 | 142135738 | -3.064641508 | TFDP2 |
| 1 | 162383847 | 162384011 | -2.432925286 | C1orf226 |

| | | | | |
|---|---|---|---|---|
| 3 | 33389819 | 33390025 | -1.119239112 | FBXL2 |
| 3 | 33389819 | 33390025 | -1.119239112 | UBP1 |
| 3 | 36831413 | 36831613 | 3.254175549 | TRANK1 |
| 3 | 36833460 | 36833664 | 1.955714573 | TRANK1 |
| 3 | 36855638 | 36855806 | 1.748788068 | TRANK1 |
| 3 | 36857298 | 36857505 | 2.621647841 | TRANK1 |
| 3 | 37142805 | 37143170 | -1.262799271 | LRRFIP2 |
| 1 | 158184556 | 158184758 | 1.548151095 | CD1D |
| 3 | 38411078 | 38411190 | -1.4473235 | XYLB |
| 3 | 38415158 | 38415352 | -3.166031923 | XYLB |
| 1 | 158958218 | 158958476 | -2.305440517 | PYHIN1 |
| 3 | 38524045 | 38524282 | -1.095264994 | EXOG |
| 3 | 39143677 | 39143881 | 3.283990918 | CSRNP1 |
| 3 | 39184213 | 39184412 | 6.483613129 | XIRP1 |
| 3 | 39185527 | 39185728 | 5.445512095 | XIRP1 |
| 3 | 39186682 | 39186876 | 7.462545387 | XIRP1 |
| 3 | 39187976 | 39188173 | 8.870114373 | XIRP1 |
| 3 | 39188843 | 39188950 | 6.065446135 | XIRP1 |
| 1 | 7772262 | 7772463 | -1.059671815 | VAMP3 |
| 3 | 39396827 | 39397029 | -1.684623784 | SLC25A38 |
| 3 | 40486802 | 40487010 | -1.241199301 | ZNF619 |
| 3 | 40517048 | 40517247 | -2.809977197 | ZNF620 |
| 3 | 41239222 | 41239418 | 2.064476179 | CTNNB1 |
| 3 | 42104137 | 42104333 | -6.399640811 | TRAK1 |
| 3 | 42106114 | 42106378 | -1.443274134 | TRAK1 |
| 3 | 42536963 | 42537075 | 5.071978558 | VIPR1 |
| 3 | 42582480 | 42582669 | -6.587774775 | SEC22C |
| 3 | 42582480 | 42582669 | -6.587774775 | SS18L2 |
| 3 | 42659945 | 42660133 | -8.557074503 | ZBTB47 |
| 3 | 43079405 | 43079592 | -1.636840133 | POMGNT2 |
| 3 | 44445677 | 44445917 | -2.482409314 | ZNF445 |
| 3 | 44446588 | 44446702 | -1.533431105 | ZNF445 |
| 3 | 44448162 | 44448353 | -1.272279515 | ZNF445 |
| 3 | 44455244 | 44455418 | -2.336601957 | ZNF445 |
| 1 | 159809898 | 159810097 | 5.803697796 | FCRL6 |
| 3 | 44500369 | 44500619 | -1.951414185 | ZNF852 |
| 3 | 44569907 | 44570135 | -1.225949956 | RP11-944L7.4 |
| 3 | 44569907 | 44570135 | -1.225949956 | ZKSCAN7 |
| 3 | 44709867 | 44710103 | -2.26569073 | SOCS5P3 |
| 3 | 44917369 | 44917569 | -2.552868454 | ZDHHC3 |
| 3 | 44926125 | 44926335 | -1.460589972 | ZDHHC3 |
| 3 | 45012908 | 45013079 | -2.101792674 | EXOSC7 |
| 3 | 45012908 | 45013079 | -2.101792674 | CLEC3B |
| 3 | 45085541 | 45085731 | 2.712213051 | CDCP1 |
| 1 | 159835199 | 159835379 | 6.026643661 | SLAMF8 |
| 1 | 159835199 | 159835379 | 6.026643661 | C1orf204 |
| 3 | 45224891 | 45225068 | 2.115642294 | TMEM158 |
| 3 | 45547605 | 45547803 | -1.460216104 | LARS2 |
| 3 | 45594879 | 45595072 | -1.199379414 | LIMD1 |
| 3 | 45595164 | 45595357 | -1.289697628 | LIMD1 |
| 3 | 45707590 | 45707824 | -2.742967902 | SACM1L |
| 1 | 159836080 | 159836278 | 5.363801025 | C1orf204 |
| 1 | 159836080 | 159836278 | 5.363801025 | SLAMF8 |
| 3 | 46358189 | 46358444 | -5.5130279 | CCR2 |
| 3 | 49011730 | 49011967 | -1.428466441 | WDR6 |
| 1 | 160031190 | 160031336 | -1.299527004 | KCNJ10 |
| 1 | 160031190 | 160031336 | -1.299527004 | PIGM |
| 3 | 49015166 | 49015430 | -1.322575297 | WDR6 |
| 3 | 49268696 | 49268894 | -3.115233272 | C3orf62 |
| 3 | 49271144 | 49271350 | -1.109980918 | C3orf62 |
| 3 | 49278164 | 49278369 | -1.15185173 | USP4 |
| 3 | 49317446 | 49317627 | -2.233811785 | USP4 |
| 3 | 49424527 | 49424699 | -1.077421401 | RP11-949J7.8 |
| 3 | 49424527 | 49424699 | -1.077421401 | NICN1 |
| 3 | 49531080 | 49531347 | -1.180663723 | DAG1 |
| 3 | 49653146 | 49653251 | 1.144284749 | BSN |
| 3 | 49662072 | 49662225 | -1.396124015 | BSN |
| 3 | 49685279 | 49685478 | 1.84044346 | MST1 |
| 3 | 49719270 | 49719482 | 1.157547325 | RNF123 |
| 3 | 49719270 | 49719482 | 1.157547325 | GMPPB |
| 3 | 49719270 | 49719482 | 1.157547325 | AMIGO3 |
| 3 | 49721010 | 49721203 | 1.587239384 | GMPPB |
| 3 | 49721010 | 49721203 | 1.587239384 | RNF123 |
| 3 | 49804613 | 49804718 | -3.859449886 | FAM212A |
| 1 | 160544696 | 160544900 | -2.124627626 | RP11-528G1.2 |
| 1 | 160544696 | 160544900 | -2.124627626 | CD84 |
| 3 | 49828926 | 49829100 | -2.210850615 | TRAIP |
| 3 | 50188297 | 50188487 | -1.464070525 | SEMA3F |
| 3 | 50297407 | 50297612 | -1.43605754 | NAT6 |
| 3 | 50297407 | 50297612 | -1.43605754 | HYAL3 |
| 3 | 50320026 | 50320224 | -1.521502304 | TUSC2 |
| 3 | 50320026 | 50320224 | -1.521502304 | HYAL2 |
| 1 | 160547134 | 160547336 | -1.030968808 | CD84 |
| 1 | 160547134 | 160547336 | -1.030968808 | RP11-528G1.2 |
| 3 | 50584315 | 50584525 | -1.259578842 | HEMK1 |
| 3 | 50607529 | 50607752 | 2.278952104 | CISH |
| 3 | 51381127 | 51381314 | -1.172508409 | DOCK3 |
| 3 | 51381918 | 51382101 | -1.140018589 | DOCK3 |
| 3 | 51393819 | 51394022 | -1.009631034 | RBM15B |
| 3 | 51653557 | 51653679 | 1.373212962 | RAD54L2 |
| 3 | 51703798 | 51703996 | -2.200625519 | TEX264 |
| 3 | 51712979 | 51713154 | -5.353679566 | GRM2 |
| 3 | 51715334 | 51715481 | -2.514643613 | GRM2 |
| 3 | 51957551 | 51957754 | -7.445132139 | PCBP4 |
| 3 | 52050765 | 52050961 | 7.103621578 | DUSP7 |
| 3 | 52075795 | 52075998 | -1.5798625 | POC1A |
| 1 | 160565498 | 160565705 | -1.099333968 | CD84 |
| 1 | 160565498 | 160565705 | -1.099333968 | RP11-528G1.2 |
| 3 | 52411113 | 52411310 | -1.331833066 | PHF7 |
| 3 | 52491988 | 52492231 | -1.527824603 | NISCH |
| 1 | 160610384 | 160610569 | 4.274082708 | SLAMF1 |
| 3 | 52752135 | 52752334 | -1.069816706 | NEK4 |
| 3 | 52807485 | 52807655 | 2.296591504 | ITIH3 |
| 3 | 52823777 | 52823971 | 1.578651297 | RP5-966M1.6 |
| 3 | 52823777 | 52823971 | 1.578651297 | ITIH4 |
| 3 | 52823777 | 52823971 | 1.578651297 | ITIH4-AS1 |
| 3 | 52883274 | 52883574 | 5.326422487 | TMEM110 |
| 3 | 52883274 | 52883574 | 5.326422487 | TMEM110-MUSTN1 |
| 3 | 142392762 | 142392952 | 5.157008352 | XRN1 |
| 3 | 142562415 | 142562540 | 1.371018292 | ATR |
| 3 | 146070653 | 146070859 | 2.896008685 | PLOD2 |
| 3 | 146070653 | 146070859 | 2.896008685 | RP11-274H2.2 |
| 3 | 149129687 | 149129855 | 6.668671574 | HPS3 |
| 3 | 149813141 | 149813330 | 2.037681037 | RNF13 |
| 3 | 149813141 | 149813330 | 2.037681037 | ANKUB1 |
| 1 | 162780235 | 162780415 | -1.937234559 | DDR2 |
| 3 | 151881518 | 151881712 | 1.142335581 | SUCNR1 |
| 3 | 151881518 | 151881712 | 1.142335581 | AADACL2-AS1 |
| 3 | 152268308 | 152268507 | 1.958729776 | MBNL1-AS1 |
| 3 | 152268308 | 152268507 | 1.958729776 | MBNL1 |
| 3 | 152835172 | 152835372 | 5.099477025 | P2RY1 |
| 3 | 152836466 | 152836667 | 3.516412779 | P2RY1 |
| 3 | 152839585 | 152839818 | 2.321707724 | P2RY1 |
| 3 | 160568250 | 160568446 | 1.129706396 | KRT8P12 |
| 3 | 160754695 | 160754819 | 2.597401152 | RP11-16N11.2 |
| 3 | 165474584 | 165474775 | -2.013229455 | LINC01322 |
| 3 | 167636916 | 167637091 | -1.999836039 | WDR49 |
| 3 | 170128792 | 170128990 | -1.195159433 | PHC3 |
| 3 | 170360322 | 170360529 | -1.133958059 | SKIL |
| 3 | 170390976 | 170391169 | -1.330985848 | SKIL |
| 3 | 170392155 | 170392360 | -1.042461837 | SKIL |
| 3 | 171602892 | 171603151 | -1.556094964 | PLD1 |
| 1 | 167704715 | 167704913 | -1.456416186 | RCSD1 |
| 3 | 172828958 | 172829092 | -1.179479413 | ATP5G1P4 |
| 3 | 175096858 | 175097050 | -1.923082183 | NAALADL2 |
| 3 | 175096858 | 175097050 | -1.923082183 | NAALADL2-AS3 |
| 3 | 179801305 | 179801501 | -6.344125155 | PEX5L |
| 3 | 183154112 | 183154311 | -1.784927804 | LAMP3 |
| 3 | 183491757 | 183491955 | -1.45125904 | KLHL6 |
| 3 | 183508302 | 183508403 | -1.792896653 | KLHL6 |
| 3 | 183650825 | 183651002 | -2.364519495 | KLHL24 |
| 3 | 183679021 | 183679219 | -1.661704593 | KLHL24 |
| 3 | 183742231 | 183742457 | -1.689039992 | YEATS2 |
| 3 | 184334832 | 184335035 | 1.284066685 | EIF4G1 |
| 1 | 920545 | 920739 | 2.352368442 | RP11-54O7.3 |
| 3 | 185482740 | 185482911 | 1.310424907 | MAP3K13 |
| 3 | 185482740 | 185482911 | 1.310424907 | TMEM41A |
| 3 | 186148015 | 186148207 | 1.241037181 | DGKG |
| 1 | 169255393 | 169255580 | -1.659053192 | NME7 |
| 3 | 187729437 | 187729632 | 7.270608821 | BCL6 |
| 3 | 187729437 | 187729632 | 7.270608821 | RP11-211G3.3 |
| 3 | 188153285 | 188153475 | -1.545091889 | LPP |
| 3 | 188153285 | 188153475 | -1.545091889 | LPP-AS2 |
| 3 | 189118141 | 189118329 | -6.191852146 | TPRG1 |
| 3 | 189284892 | 189285041 | -1.000151783 | TPRG1 |
| 1 | 169871599 | 169871835 | -2.239023648 | SCYL3 |
| 3 | 194487876 | 194488010 | 1.579956697 | LINC00884 |
| 3 | 194487876 | 194488010 | 1.579956697 | ATP13A3 |
| 3 | 194637760 | 194637893 | 5.451187557 | AC046143.3 |
| 1 | 170552014 | 170552243 | -1.424929825 | GORAB |
| 3 | 194687517 | 194687719 | 1.664025478 | FAM43A |
| 3 | 195069517 | 195069724 | 1.285346588 | XXYLT1 |

| | | | | |
|---|---|---|---|---|
| 3 | 46359753 | 46359948 | -7.77119841 | CCR2 |
| 3 | 47016990 | 47017155 | -1.070210554 | SETD2 |
| 1 | 159848980 | 159849154 | 5.211032383 | C1orf204 |
| 3 | 47225807 | 47225939 | -1.665245967 | KIF9-AS1 |
| 3 | 47380380 | 47380547 | -2.193007299 | RP11-708J19.1 |
| 3 | 47413831 | 47413991 | -1.590916102 | SCAP |
| 3 | 47468711 | 47468895 | -1.838760225 | SCAP |
| 3 | 47495950 | 47496136 | -2.673355074 | ELP6 |
| 3 | 47577365 | 47577487 | -3.192970691 | CSPG5 |
| 1 | 159866737 | 159866936 | 2.493467983 | RP11-190A12.7 |
| 3 | 48158741 | 48158947 | -4.061541399 | CDC25A |
| 3 | 48269712 | 48269908 | -2.414952961 | ZNF589 |
| 1 | 159980942 | 159981140 | -7.833112001 | LINC01133 |
| 3 | 48432614 | 48432814 | -2.897871512 | CCDC51 |
| 3 | 48460342 | 48460533 | -2.872676999 | ATRIP |
| 3 | 48561726 | 48561887 | 7.664845621 | UCN2 |
| 3 | 48561726 | 48561887 | 7.664845621 | PFKFB4 |
| 3 | 48562535 | 48562706 | 8.616688131 | UCN2 |
| 1 | 160030952 | 160031092 | -1.104118365 | KCNJ10 |
| 1 | 160030952 | 160031092 | -1.104118365 | PIGM |
| 3 | 48681040 | 48681240 | -2.48300802 | NCKIPSD |
| 3 | 48917969 | 48918168 | 1.393426669 | ARIH2OS |
| 3 | 48989714 | 48989883 | -8.38739353 | RP13-131K19.2 |
| 3 | 49011730 | 49011967 | -1.141112957 | WDR6 |
| 3 | 49013625 | 49013811 | -1.958500441 | WDR6 |
| 1 | 160031190 | 160031336 | -1.188871119 | PIGM |
| 1 | 160031190 | 160031336 | -1.188871119 | KCNJ10 |
| 3 | 49015166 | 49015430 | -1.404063708 | WDR6 |
| 3 | 49021305 | 49021498 | -3.483308801 | DALRD3 |
| 3 | 49021305 | 49021498 | -3.483308801 | NDUFAF3 |
| 3 | 49163035 | 49163185 | -1.985788248 | CCDC71 |
| 3 | 49268696 | 49268894 | 2.058023316 | C3orf62 |
| 3 | 49271144 | 49271350 | 1.173480209 | C3orf62 |
| 3 | 49653146 | 49653251 | -1.443414411 | BSN |
| 3 | 49685279 | 49685478 | -4.39395128 | MST1 |
| 3 | 49686808 | 49686916 | -3.970597616 | MST1 |
| 3 | 49687198 | 49687397 | -8.141942082 | MST1 |
| 3 | 49721010 | 49721203 | -2.877206509 | RNF123 |
| 3 | 49721010 | 49721203 | -2.877206509 | GMPPB |
| 3 | 49721477 | 49721670 | -1.945194573 | GMPPB |
| 3 | 49721477 | 49721670 | -1.945194573 | RNF123 |
| 3 | 49804613 | 49804718 | -1.124266183 | FAM212A |
| 3 | 49828926 | 49829100 | -3.213538123 | TRAIP |
| 3 | 50188297 | 50188487 | -1.87108056 | SEMA3F |
| 3 | 50294927 | 50295126 | -6.228237492 | HYAL3 |
| 3 | 50297407 | 50297612 | -2.511112137 | NAT6 |
| 3 | 50297407 | 50297612 | -2.511112137 | HYAL3 |
| 3 | 50582437 | 50582643 | -2.764778658 | HEMK1 |
| 1 | 160547134 | 160547336 | 1.703960722 | RP11-528G1.2 |
| 1 | 160547134 | 160547336 | 1.703960722 | CD84 |
| 3 | 50584315 | 50584525 | -1.766836636 | HEMK1 |
| 3 | 50607529 | 50607752 | -1.673980308 | CISH |
| 3 | 51381127 | 51381314 | 7.479978333 | DOCK3 |
| 1 | 160547984 | 160548232 | 2.137861908 | CD84 |
| 3 | 53064610 | 53064809 | -5.658140766 | SERBP1P3 |
| 3 | 53064610 | 53064809 | -5.658140766 | RP11-894J14.5 |
| 3 | 53287410 | 53287610 | 1.399267126 | DCP1A |
| 3 | 53868702 | 53868971 | -1.126537674 | ACTR8 |
| 3 | 56563756 | 56563991 | -2.330403129 | CCDC66 |
| 3 | 56617171 | 56617295 | -1.337503212 | CCDC66 |
| 3 | 56623526 | 56623732 | -1.213291142 | FAM208A |
| 3 | 56633245 | 56633404 | -1.323095989 | FAM208A |
| 3 | 56633740 | 56633940 | -1.447364453 | FAM208A |
| 1 | 7791782 | 7791971 | -5.714180086 | PER3 |
| 3 | 57628166 | 57628367 | -1.035906947 | DENND6A |
| 3 | 57912407 | 57912590 | -2.977735955 | SLMAP |
| 1 | 160753807 | 160754008 | 4.276083229 | SLAMF7 |
| 3 | 58500996 | 58501121 | -2.345705598 | KCTD6 |
| 3 | 58501468 | 58501664 | -1.269315064 | KCTD6 |
| 3 | 63999377 | 63999574 | 1.231255442 | ATXN7 |
| 3 | 66380388 | 66380590 | 1.525362599 | SLC25A26 |
| 3 | 66380388 | 66380590 | 1.525362599 | LRIG1 |
| 3 | 67007982 | 67008181 | 1.897856405 | KBTBD8 |
| 1 | 160998133 | 160998317 | -1.236981414 | RP11-544M22.13 |
| 1 | 160998133 | 160998317 | -1.236981414 | F11R |
| 3 | 69171535 | 69171734 | -2.250915706 | FRMD4B |
| 3 | 69176589 | 69176785 | -2.186538966 | FRMD4B |
| 3 | 69181017 | 69181222 | -2.467967886 | FRMD4B |
| 3 | 69964885 | 69965074 | -1.825012572 | MITF |
| 3 | 70014133 | 70014261 | -5.245926548 | SAMMSON |
| 3 | 70958837 | 70958988 | -1.044270062 | FOXP1 |
| 1 | 161021536 | 161021703 | -5.207342618 | RP11-544M22.13 |
| 3 | 71582843 | 71582990 | 6.400854179 | FOXP1 |
| 3 | 71582843 | 71582990 | 6.400854179 | RP11-154H23.4 |
| 3 | 75415547 | 75415731 | -3.495742518 | ALG1L6P |
| 3 | 75418643 | 75418859 | -1.196921639 | ALG1L6P |
| 3 | 75422270 | 75422472 | -1.640753996 | FAM86DP |
| 3 | 75425735 | 75425937 | -1.860490604 | FAM86DP |
| 3 | 75453633 | 75453806 | -2.796340636 | ENPP7P2 |
| 3 | 75453633 | 75453806 | -2.796340636 | RP11-803B1.8 |
| 3 | 75737795 | 75738061 | -1.092962366 | ZNF717 |
| 3 | 78938771 | 78938908 | 2.165038746 | ROBO1 |
| 3 | 79018544 | 79018741 | 2.709752726 | ROBO1 |
| 3 | 81524400 | 81524598 | -1.757504008 | GBE1 |
| 3 | 82287309 | 82287584 | -1.475625935 | RP11-260O18.1 |
| 1 | 161048095 | 161048195 | 6.361830472 | ARHGAP30 |
| 3 | 82288606 | 82288785 | -2.702195128 | RP11-260O18.1 |
| 3 | 85290690 | 85290872 | -1.944137459 | CADM2 |
| 3 | 85295476 | 85295665 | -2.794357318 | CADM2 |
| 3 | 88099758 | 88099956 | -1.086502197 | CGGBP1 |
| 3 | 88099758 | 88099956 | -1.086502197 | ZNF654 |
| 3 | 94061110 | 94061260 | -1.658132888 | DHFR2 |
| 3 | 96987594 | 96987788 | 3.153644198 | EPHA6 |
| 3 | 97877471 | 97877658 | 1.426899203 | CRYBG3 |
| 3 | 97905025 | 97905219 | -5.114409345 | CRYBG3 |
| 3 | 99850080 | 99850245 | -4.390679857 | CMSS1 |
| 3 | 99850080 | 99850245 | -4.390679857 | FILIP1L |
| 3 | 100351666 | 100351858 | -6.425874521 | NIT2 |
| 3 | 195180180 | 195180341 | 2.657241641 | XXYLT1 |
| 3 | 195618941 | 195619100 | -3.069349222 | MUC20P1 |
| 3 | 195868311 | 195868440 | 8.588446924 | TNK2 |
| 3 | 196051875 | 196052094 | -1.219344878 | TFRC |
| 3 | 196219864 | 196220045 | -1.766664866 | PCYT1A |
| 3 | 196219864 | 196220045 | -1.766664866 | SLC51A |
| 1 | 171535309 | 171535509 | 5.31467831 | PRRC2C |
| 3 | 196356665 | 196356860 | -1.148387309 | UBXN7 |
| 3 | 196487494 | 196487700 | -1.675273481 | RNF168 |
| 3 | 196488617 | 196488818 | -2.128072698 | RNF168 |
| 3 | 196496379 | 196496563 | -1.263735003 | RNF168 |
| 1 | 8004893 | 8005048 | 2.309910629 | ERRFI1 |
| 3 | 197001954 | 197002093 | 1.613687654 | MFI2-AS1 |
| 3 | 197001954 | 197002093 | 1.613687654 | MELTF |
| 3 | 197512020 | 197512210 | 1.747698558 | BDH1 |
| 3 | 198058666 | 198058841 | -1.143788295 | ANKRD18DP |
| 4 | 372987 | 373105 | -5.844562202 | ZNF141 |
| 4 | 537909 | 538106 | -5.680971887 | PIGG |
| 4 | 675305 | 675409 | 6.679050854 | MYL5 |
| 4 | 687892 | 688051 | 1.279844022 | MFSD7 |
| 4 | 688732 | 688954 | 2.396413141 | MFSD7 |
| 4 | 857760 | 857923 | -1.829549679 | GAK |
| 4 | 879302 | 879499 | -1.34474407 | GAK |
| 4 | 880359 | 880535 | 6.020829576 | GAK |
| 4 | 989167 | 989311 | 2.715138774 | SLC26A1 |
| 4 | 989167 | 989311 | 2.715138774 | IDUA |
| 1 | 172570964 | 172571158 | 4.480290606 | SUCO |
| 4 | 990214 | 990362 | 1.962827821 | SLC26A1 |
| 4 | 990214 | 990362 | 1.962827821 | IDUA |
| 4 | 1010402 | 1010588 | -1.04438543 | FGFRL1 |
| 4 | 1574179 | 1574375 | -6.222507423 | RP11-1398P2.1 |
| 4 | 1978869 | 1979068 | 1.288318578 | WHSC1 |
| 4 | 2063888 | 2064090 | 4.945324257 | NAT8L |
| 4 | 2304618 | 2304736 | 2.011681635 | RP11-478C1.7 |
| 4 | 2304618 | 2304736 | 2.011681635 | ZFYVE28 |
| 1 | 173041121 | 173041321 | 2.534921333 | TNFSF18 |
| 1 | 173041121 | 173041321 | 2.534921333 | RP1-15D23.2 |
| 4 | 2731795 | 2731996 | 1.094933997 | FAM193A |
| 4 | 2940591 | 2940794 | 2.422949537 | NOP14-AS1 |
| 4 | 2940591 | 2940794 | 2.422949537 | NOP14 |
| 4 | 2941466 | 2941663 | 1.855960274 | NOP14 |
| 4 | 2941466 | 2941663 | 1.855960274 | NOP14-AS1 |
| 1 | 173185841 | 173186022 | 1.197116938 | TNFSF4 |
| 4 | 3374447 | 3374625 | 1.461972129 | RGS12 |
| 4 | 3375029 | 3375194 | 3.342633703 | RGS12 |
| 4 | 3487793 | 3487956 | 1.303328786 | DOK7 |
| 4 | 3512761 | 3512963 | 1.114679785 | LRPAP1 |
| 4 | 3768005 | 3768205 | 1.436850965 | ADRA2C |
| 4 | 3947495 | 3947682 | 2.414056741 | AC226119.5 |
| 4 | 3947495 | 3947682 | 2.414056741 | FAM86EP |
| 1 | 8013362 | 8013506 | 2.786679228 | ERRFI1 |
| 1 | 173186477 | 173186678 | 1.177573278 | TNFSF4 |
| 4 | 6640930 | 6641131 | 10.48982494 | MRFAP1 |
| 1 | 173728040 | 173728142 | -5.975642357 | KLHL20 |

| | | | | |
|---|---|---|---|---|
| 1 | 160547984 | 160548232 | 2.137861908 | RP11-528G1.2 |
| 3 | 51653557 | 51653679 | -1.444691939 | RAD54L2 |
| 3 | 51712979 | 51713154 | 5.353679566 | GRM2 |
| 3 | 52050765 | 52050961 | -7.438282454 | DUSP7 |
| 3 | 52075795 | 52075998 | -2.025150189 | POC1A |
| 1 | 160565498 | 160565705 | 1.737430399 | RP11-528G1.2 |
| 1 | 160565498 | 160565705 | 1.737430399 | CD84 |
| 3 | 52277993 | 52278191 | 8.175952721 | WDR82 |
| 3 | 52393414 | 52393598 | -1.60116188 | DNAH1 |
| 3 | 52400461 | 52400643 | 2.098004134 | DNAH1 |
| 3 | 52411113 | 52411310 | -2.367850767 | PHF7 |
| 3 | 52435676 | 52435866 | -2.855531951 | SEMA3G |
| 3 | 52609576 | 52609775 | -1.082696126 | PBRM1 |
| 3 | 52807485 | 52807655 | 4.225605556 | ITIH3 |
| 3 | 52823777 | 52823971 | 1.131067758 | ITIH4-AS1 |
| 3 | 52823777 | 52823971 | 1.131067758 | ITIH4 |
| 3 | 52823777 | 52823971 | 1.131067758 | RP5-966M1.6 |
| 3 | 52883274 | 52883574 | -7.105792077 | TMEM110 |
| 3 | 52883274 | 52883574 | -7.105792077 | TMEM110-MUSTN1 |
| 3 | 52904902 | 52905102 | -1.187371093 | SFMBT1 |
| 3 | 56563756 | 56563991 | -1.931734158 | CCDC66 |
| 3 | 56728546 | 56728766 | 2.854614701 | ARHGEF3 |
| 1 | 7791782 | 7791971 | 5.714180086 | PER3 |
| 3 | 56729340 | 56729541 | 3.779879322 | ARHGEF3 |
| 3 | 57557108 | 57557365 | -1.143191503 | PDE12 |
| 3 | 57559504 | 57559717 | -1.421925118 | PDE12 |
| 1 | 160753807 | 160754008 | 10.21094612 | SLAMF7 |
| 3 | 62203473 | 62203783 | 1.856909768 | PTPRG |
| 3 | 63999377 | 63999574 | -1.040435365 | ATXN7 |
| 3 | 68992206 | 68992393 | -1.356402876 | EOGT |
| 3 | 68993369 | 68993541 | -1.820459453 | EOGT |
| 1 | 160998133 | 160998317 | 1.561017686 | RP11-544M22.13 |
| 1 | 160998133 | 160998317 | 1.561017686 | F11R |
| 3 | 69417513 | 69417704 | 2.150484238 | FRMD4B |
| 3 | 69964885 | 69965074 | 3.141242039 | MITF |
| 3 | 71579966 | 71580163 | -2.429859696 | FOXP1 |
| 3 | 72750557 | 72750763 | -1.728721459 | SHQ1 |
| 1 | 161048095 | 161048195 | -9.605822969 | ARHGAP30 |
| 3 | 82416859 | 82416991 | -2.011923371 | RP11-260O18.1 |
| 3 | 85290690 | 85290872 | 2.196158711 | CADM2 |
| 3 | 85295476 | 85295665 | 7.040283866 | CADM2 |
| 3 | 96987594 | 96987788 | -3.421457192 | EPHA6 |
| 1 | 161120694 | 161120877 | -1.738830375 | NIT1 |
| 3 | 97871886 | 97872087 | -1.276020867 | RP11-389G6.4 |
| 3 | 97871886 | 97872087 | -1.276020867 | CRYBG3 |
| 3 | 97876633 | 97876835 | -1.051364709 | CRYBG3 |
| 3 | 97877471 | 97877658 | -1.995548367 | CRYBG3 |
| 3 | 97943194 | 97943392 | -1.062116616 | MINA |
| 3 | 97943194 | 97943392 | -1.062116616 | CRYBG3 |
| 3 | 98799488 | 98799679 | -1.369220162 | ST3GAL6 |
| 3 | 98799488 | 98799679 | -1.369220162 | DCBLD2 |
| 3 | 99850080 | 99850245 | -3.501637566 | CMSS1 |
| 3 | 99850080 | 99850245 | -3.501637566 | FILIP1L |
| 3 | 99850593 | 99850787 | -1.494308097 | FILIP1L |
| 3 | 101243772 | 101243968 | -2.396925329 | IMPG2 |
| 3 | 101712373 | 101712630 | -1.322913746 | PDCL3P4 |
| 3 | 101758033 | 101758218 | -2.132622505 | CEP97 |
| 3 | 101765081 | 101765260 | -2.360422599 | CEP97 |
| 1 | 161191175 | 161191370 | -5.449444269 | ADAMTS4 |
| 3 | 101821827 | 101822024 | -1.083854412 | NXPE3 |
| 3 | 105574393 | 105574589 | -2.132170082 | ALCAM |
| 1 | 7839777 | 7840052 | -3.923637157 | PER3 |
| 3 | 105702163 | 105702351 | -1.560109306 | CBLB |
| 3 | 108644388 | 108644489 | -1.0206156 | DZIP3 |
| 3 | 111133749 | 111133945 | 2.921379127 | NECTIN3 |
| 1 | 161230264 | 161230455 | -1.233549602 | TOMM40L |
| 1 | 161230264 | 161230455 | -1.233549602 | NR1I3 |
| 3 | 113291373 | 113291573 | -1.444347275 | AC026348.1 |
| 3 | 113291373 | 113291573 | -1.444347275 | CFAP44 |
| 3 | 113413383 | 113413544 | -5.504214311 | AC026348.1 |
| 3 | 113413383 | 113413544 | -5.504214311 | CFAP44-AS1 |
| 3 | 113413383 | 113413544 | -5.504214311 | CFAP44 |
| 3 | 113453630 | 113453828 | -6.054915327 | SPICE1 |
| 3 | 113628425 | 113628624 | 2.090998917 | SIDT1 |
| 1 | 161400265 | 161400466 | -1.248348427 | RP11-122G18.12 |
| 3 | 113954030 | 113954140 | 3.032065169 | ZDHHC23 |
| 3 | 119415953 | 119416154 | 3.165637562 | ARHGAP31 |
| 3 | 119587518 | 119587733 | 1.542989804 | ADPRH |
| 3 | 120094942 | 120095074 | 2.46093874 | RP11-18H7.1 |
| 3 | 120733546 | 120733737 | -6.531360384 | RABL3 |
| 1 | 161400924 | 161401178 | -1.625509612 | RP11-122G18.12 |
| 3 | 120735627 | 120735809 | -6.36161715 | RABL3 |
| 3 | 121419270 | 121419465 | -6.149604009 | STXBP5L |
| 3 | 121488782 | 121488982 | -2.211036829 | POLQ |
| 3 | 121664395 | 121664567 | -6.374934714 | GOLGB1 |
| 3 | 121697614 | 121697814 | -1.108397273 | GOLGB1 |
| 3 | 122396125 | 122396332 | 2.306516633 | FAM162A |
| 3 | 122555429 | 122555641 | 1.109358618 | PARP9 |
| 3 | 122569594 | 122569802 | 1.04324408 | DTX3L |
| 3 | 122699687 | 122699887 | 2.254879888 | PARP14 |
| 3 | 122700609 | 122700812 | 1.996815718 | PARP14 |
| 3 | 122718461 | 122718665 | 1.822370967 | PARP14 |
| 3 | 122740567 | 122740768 | -1.802481053 | HSPBAP1 |
| 3 | 122839262 | 122839414 | -5.10715956 | DIRC2 |
| 3 | 123946952 | 123947155 | -1.324574039 | CCDC14 |
| 3 | 123958853 | 123959090 | -1.366267503 | CCDC14 |
| 1 | 161442688 | 161442876 | -4.83974005 | RP11-122G18.10 |
| 3 | 124796486 | 124796696 | 9.352362172 | ITGB5 |
| 3 | 125233669 | 125233875 | -1.201950981 | ZNF148 |
| 1 | 161457562 | 161457735 | -2.761780978 | RP11-122G18.10 |
| 3 | 125594831 | 125595033 | -1.015165402 | OSBPL11 |
| 3 | 126351992 | 126352116 | -2.331706215 | KLF15 |
| 3 | 126451652 | 126451903 | -1.41929889 | ZXDC |
| 3 | 126459666 | 126459873 | -1.947318814 | ZXDC |
| 3 | 126461799 | 126462000 | -1.21339911 | ZXDC |
| 3 | 126542824 | 126543050 | -2.137240466 | CHST13 |
| 3 | 126542824 | 126543050 | -2.137240466 | C3orf22 |
| 3 | 126989494 | 126989693 | 8.689807638 | PLXNA1 |
| 4 | 6967177 | 6967359 | 7.086370009 | TBC1D14 |
| 4 | 7042398 | 7042571 | 1.291186996 | TADA2B |
| 4 | 7042398 | 7042571 | 1.291186996 | RP11-367J11.2 |
| 4 | 7042398 | 7042571 | 1.291186996 | CCDC96 |
| 4 | 7717284 | 7717476 | -1.249487413 | SORCS2 |
| 4 | 8383926 | 8384121 | 1.395685555 | ACOX3 |
| 4 | 8386045 | 8386245 | -1.107997783 | ACOX3 |
| 4 | 8387565 | 8387728 | 2.244764022 | ACOX3 |
| 4 | 10019427 | 10019620 | -5.239890132 | SLC2A9 |
| 4 | 13544757 | 13544877 | 6.87545815 | NKX3-2 |
| 4 | 14535460 | 14535648 | -6.856639552 | LINC00504 |
| 4 | 15003000 | 15003165 | 9.631847792 | CPEB2 |
| 4 | 15356967 | 15357285 | -1.322763718 | RP11-665G4.1 |
| 4 | 15356967 | 15357285 | -1.322763718 | C1QTNF7 |
| 4 | 15612336 | 15612536 | -1.594003838 | FBXL5 |
| 4 | 15631795 | 15632103 | -1.115417735 | FBXL5 |
| 4 | 15821610 | 15821809 | -2.903677461 | CD38 |
| 4 | 16360779 | 16360896 | -2.229715809 | ZEB2P1 |
| 4 | 17646375 | 17646492 | -1.476690608 | FAM184B |
| 4 | 17860582 | 17860749 | -3.890679857 | LCORL |
| 1 | 173938682 | 173938862 | -8.607415675 | RC3H1 |
| 4 | 20253590 | 20253729 | 5.064641508 | SLIT2 |
| 4 | 20619205 | 20619394 | 1.909112504 | SLIT2 |
| 4 | 20742494 | 20742695 | -1.236049015 | KCNIP4 |
| 4 | 20742494 | 20742695 | -1.236049015 | PACRGL |
| 4 | 20744259 | 20744532 | -1.906658904 | KCNIP4 |
| 4 | 20744259 | 20744532 | -1.906658904 | PACRGL |
| 4 | 24807412 | 24807553 | -7.328659225 | CCDC149 |
| 1 | 8014000 | 8014231 | 2.324977061 | ERRFI1 |
| 4 | 24808369 | 24808506 | -1.115372909 | CCDC149 |
| 4 | 24909840 | 24910020 | -1.863321561 | CCDC149 |
| 4 | 24910227 | 24910478 | -1.106463151 | CCDC149 |
| 4 | 27022615 | 27022813 | 1.298092684 | STIM2 |
| 4 | 30720986 | 30721185 | 1.60322657 | PCDH7 |
| 4 | 30786921 | 30787121 | -5.934411277 | PCDH7 |
| 4 | 30786921 | 30787121 | -5.934411277 | RP11-619J20.1 |
| 1 | 174437616 | 174437813 | -2.960785245 | RABGAP1L |
| 4 | 38697031 | 38697258 | -1.578741511 | KLF3 |
| 4 | 38790552 | 38790747 | -1.284689303 | TLR1 |
| 4 | 38796539 | 38796753 | -1.279833354 | TLR1 |
| 4 | 38798165 | 38798285 | -1.732362959 | TLR1 |
| 1 | 174489065 | 174489281 | 1.012807987 | RABGAP1L |
| 4 | 39125411 | 39125614 | -1.419996683 | RP11-360F5.1 |
| 4 | 39125411 | 39125614 | -1.419996683 | KLHL5 |
| 1 | 174561827 | 174562026 | 7.70025629 | RABGAP1L |
| 4 | 41982539 | 41982707 | -1.860201097 | DCAF4L1 |
| 4 | 48716183 | 48716336 | -5.099836172 | FRYL |
| 4 | 53096004 | 53096201 | -1.173769369 | SCFD2 |
| 1 | 174729505 | 174729665 | -2.213389254 | RABGAP1L |
| 4 | 53473457 | 53473586 | -1.262763193 | RP11-231C18.3 |
| 4 | 53473457 | 53473586 | -1.262763193 | LNX1 |
| 1 | 174989759 | 174989944 | -3.199085887 | RABGAP1L |
| 4 | 56976630 | 56976825 | 3.9412799 | NOA1 |
| 4 | 66002189 | 66002390 | -6.149604009 | RP11-25H12.1 |

| | | | | |
|---|---|---|---|---|
| 3 | 99850593 | 99850787 | -1.494308097 | CMSS1 |
| 3 | 100574324 | 100574478 | -3.532101921 | TMEM45A |
| 3 | 101565183 | 101565383 | -1.271185025 | TRMT10C |
| 3 | 101651311 | 101651622 | -1.117436134 | ZBTB11 |
| 3 | 101677016 | 101677234 | -1.398635687 | ZBTB11-AS1 |
| 3 | 101677016 | 101677234 | -1.398635687 | ZBTB11 |
| 3 | 101712373 | 101712630 | 2.119910181 | PDCL3P4 |
| 3 | 101758033 | 101758218 | -2.48885359 | CEP97 |
| 3 | 101765081 | 101765260 | -2.238320454 | CEP97 |
| 1 | 161191175 | 161191370 | 5.075213652 | ADAMTS4 |
| 3 | 101801330 | 101801522 | -3.128575822 | NXPE3 |
| 3 | 101821827 | 101822024 | -3.081290378 | NXPE3 |
| 3 | 105405375 | 105405573 | 2.406162908 | ALCAM |
| 3 | 105574393 | 105574589 | 2.906033641 | ALCAM |
| 1 | 7839777 | 7840052 | 1.549360373 | PER3 |
| 3 | 105576494 | 105576698 | 2.44302867 | ALCAM |
| 3 | 105658951 | 105659152 | 3.202385527 | CBLB |
| 3 | 105702163 | 105702351 | 4.354123708 | CBLB |
| 3 | 105727530 | 105727735 | 3.46094629 | CBLB |
| 3 | 112580637 | 112580838 | -1.139568997 | SLC35A5 |
| 3 | 113291373 | 113291573 | -1.495199265 | CFAP44 |
| 3 | 113291373 | 113291573 | -1.495199265 | AC026348.1 |
| 3 | 113413383 | 113413544 | -1.709900338 | CFAP44 |
| 3 | 113413383 | 113413544 | -1.709900338 | AC026348.1 |
| 3 | 113413383 | 113413544 | -1.709900338 | CFAP44-AS1 |
| 3 | 113453630 | 113453828 | -2.547704478 | SPICE1 |
| 3 | 113628425 | 113628624 | -2.037541315 | SIDT1 |
| 3 | 113954030 | 113954140 | -6.262554613 | ZDHHC23 |
| 3 | 114085992 | 114086192 | -2.607133252 | QTRT2 |
| 3 | 119415953 | 119416154 | 5.269842769 | ARHGAP31 |
| 3 | 120733546 | 120733737 | 1.566731179 | RABL3 |
| 3 | 120735627 | 120735809 | 6.36161715 | RABL3 |
| 3 | 121488011 | 121488207 | -9.896025757 | POLQ |
| 3 | 121488782 | 121488982 | -3.682604716 | POLQ |
| 3 | 121664395 | 121664567 | -1.900709134 | GOLGB1 |
| 1 | 161421933 | 161422127 | -1.353912941 | RP11-122G18.10 |
| 1 | 161421933 | 161422127 | -1.353912941 | RP11-122G18.12 |
| 3 | 122396125 | 122396332 | -2.124889155 | FAM162A |
| 3 | 122414607 | 122414884 | -1.325057551 | WDR5B |
| 3 | 122415164 | 122415357 | -1.441750392 | WDR5B |
| 3 | 122426532 | 122426731 | -1.522437738 | KPNA1 |
| 3 | 122426532 | 122426731 | -1.522437738 | RP11-299J3.8 |
| 3 | 122528371 | 122528573 | 4.515645168 | PARP9 |
| 3 | 122536986 | 122537182 | 3.626607672 | PARP9 |
| 1 | 161432461 | 161432640 | -1.232881934 | RP11-122G18.10 |
| 3 | 122540580 | 122540751 | 3.916938051 | PARP9 |
| 3 | 122555429 | 122555641 | 3.253999382 | PARP9 |
| 3 | 122569594 | 122569802 | 1.251362925 | DTX3L |
| 3 | 122699687 | 122699887 | 1.702281426 | PARP14 |
| 3 | 122700609 | 122700812 | 2.65932424 | PARP14 |
| 3 | 122718461 | 122718665 | 2.777411884 | PARP14 |
| 3 | 122728275 | 122728474 | 4.26797268 | PARP14 |
| 3 | 122839262 | 122839414 | -1.452310976 | DIRC2 |
| 3 | 123271833 | 123272045 | -1.193636718 | SEC22A |
| 3 | 127621723 | 127622011 | -1.34573488 | MCM2 |
| 3 | 127672282 | 127672481 | 5.108372929 | PODXL2 |
| 1 | 7841533 | 7841684 | -4.920388962 | PER3 |
| 3 | 128071006 | 128071206 | 10.21610119 | SEC61A1 |
| 3 | 128071006 | 128071206 | 10.21610119 | RUVBL1 |
| 3 | 128153107 | 128153307 | -1.071616442 | RUVBL1 |
| 3 | 128341471 | 128341674 | -1.234712544 | EEFSEC |
| 3 | 128480786 | 128480987 | -1.778504819 | GATA2 |
| 3 | 128501872 | 128502036 | -2.016691528 | GATA2-AS1 |
| 3 | 128538512 | 128538747 | -2.764944914 | TMED10P2 |
| 3 | 128880983 | 128881183 | 6.311140328 | ACAD9 |
| 3 | 129122474 | 129122672 | 1.145777223 | ISY1-RAB43 |
| 3 | 129122474 | 129122672 | 1.145777223 | RAB43 |
| 3 | 129436992 | 129437242 | -2.007278673 | MBD4 |
| 3 | 129556137 | 129556403 | -1.438684928 | PLXND1 |
| 3 | 129650968 | 129651133 | -3.187860456 | TMCC1 |
| 3 | 129655022 | 129655226 | -2.104790095 | TMCC1 |
| 3 | 129670426 | 129670627 | -1.739938548 | TMCC1 |
| 3 | 129828221 | 129828446 | -1.846876223 | TMCC1 |
| 3 | 129847537 | 129847738 | -1.144500071 | TMCC1 |
| 3 | 129847537 | 129847738 | -1.144500071 | RP11-332M2.1 |
| 3 | 129905711 | 129905878 | -3.252823087 | TMCC1-AS1 |
| 3 | 131024525 | 131024635 | -4.39395128 | ASTE1 |
| 3 | 131024701 | 131024880 | -1.326912179 | ASTE1 |
| 3 | 132660070 | 132660262 | -1.324583994 | NPHP3-ACAD11 |
| 3 | 132660070 | 132660262 | -1.324583994 | ACAD11 |
| 3 | 132660070 | 132660262 | -1.324583994 | UBA5 |
| 3 | 134854140 | 134854307 | 6.257357027 | EPHB1 |
| 3 | 136151539 | 136151741 | -1.054778594 | MSL2 |
| 3 | 136370795 | 136370899 | -5.26569073 | STAG1 |
| 3 | 136819323 | 136819502 | -5.169368191 | SLC35G2 |
| 3 | 136855199 | 136855352 | -2.536082573 | SLC35G2 |
| 3 | 136855199 | 136855352 | -2.536082573 | NCK1-AS1 |
| 3 | 136861793 | 136861939 | -2.277649395 | NCK1-AS1 |
| 3 | 136945761 | 136945967 | -1.477314118 | NCK1 |
| 3 | 136948238 | 136948443 | -1.599632324 | NCK1 |
| 3 | 136948238 | 136948443 | -1.599632324 | IL20RB |
| 3 | 138295999 | 138296197 | -1.019105441 | ARMC8 |
| 3 | 138295999 | 138296197 | -1.019105441 | NME9 |
| 3 | 138349669 | 138349870 | 3.369508985 | MRAS |
| 3 | 138655212 | 138655413 | -1.585725428 | PIK3CB |
| 3 | 139406777 | 139406966 | -1.017278611 | RP11-319G6.1 |
| 3 | 139407484 | 139407654 | -6.671801069 | RP11-319G6.1 |
| 3 | 139494272 | 139494479 | -1.780498636 | RP11-319G6.1 |
| 3 | 139494272 | 139494479 | -1.780498636 | ACTG1P1 |
| 3 | 140973670 | 140973872 | -1.692482378 | SLC25A36 |
| 3 | 140976370 | 140976576 | -1.235083447 | SLC25A36 |
| 3 | 141292335 | 141292504 | -2.767531783 | PXYLP1 |
| 3 | 141292335 | 141292504 | -2.767531783 | RP11-438D8.2 |
| 3 | 141292585 | 141292762 | -1.464658972 | RP11-438D8.2 |
| 3 | 141292585 | 141292762 | -1.464658972 | PXYLP1 |
| 3 | 141443821 | 141444022 | -1.315998108 | ZBTB38 |
| 3 | 141445730 | 141445899 | -1.342899921 | ZBTB38 |
| 1 | 162383847 | 162384011 | -1.38948656 | C1orf226 |
| 4 | 66023964 | 66024156 | -1.504086966 | RP11-25H12.1 |
| 4 | 66111116 | 66111252 | -1.231485988 | RP11-25H12.1 |
| 4 | 67514204 | 67514408 | -1.120553474 | CENPC |
| 1 | 8358288 | 8358482 | 2.138572246 | RERE |
| 1 | 175160519 | 175160718 | 3.339010588 | KIAA0040 |
| 4 | 70799209 | 70799359 | 3.120520564 | RUFY3 |
| 4 | 73098209 | 73098408 | -2.960532913 | ANKRD17 |
| 4 | 74961363 | 74961560 | 5.294357318 | RP11-44F21.3 |
| 4 | 74961363 | 74961560 | 5.294357318 | PARM1 |
| 4 | 78795474 | 78795589 | -6.991184873 | BMP2K |
| 4 | 78796072 | 78796281 | -8.264989757 | BMP2K |
| 1 | 176125239 | 176125478 | 1.912350143 | RFWD2 |
| 4 | 79905858 | 79906060 | 1.053773449 | ANTXR2 |
| 4 | 79907069 | 79907271 | 1.157491355 | ANTXR2 |
| 4 | 80072750 | 80072949 | 10.70338588 | ANTXR2 |
| 4 | 80201932 | 80202131 | 2.51599025 | PRDM8 |
| 4 | 80203406 | 80203662 | 1.169669729 | PRDM8 |
| 4 | 82429747 | 82429948 | 1.829158874 | HNRNPDL |
| 4 | 83283996 | 83284200 | 2.444144806 | COQ2 |
| 4 | 84672702 | 84672832 | -1.321553534 | WDFY3 |
| 4 | 87089834 | 87090031 | 1.711840797 | AFF1 |
| 4 | 87125982 | 87126181 | -1.623304329 | AFF1 |
| 4 | 87662198 | 87662391 | 4.125293216 | DMP1 |
| 4 | 87662198 | 87662391 | 4.125293216 | RP11-742B18.1 |
| 4 | 87811618 | 87811811 | -7.465276277 | IBSP |
| 4 | 87982532 | 87982747 | 15.03566447 | SPP1 |
| 4 | 88696701 | 88696909 | 1.586797576 | NAP1L5 |
| 4 | 88696701 | 88696909 | 1.586797576 | HERC3 |
| 4 | 94289485 | 94289590 | -1.077495927 | SMARCAD1 |
| 4 | 95549021 | 95549220 | 3.002232376 | RP11-710C12.1 |
| 4 | 95549021 | 95549220 | 3.002232376 | UNC5C |
| 4 | 99651265 | 99651462 | 6.07476772 | RP11-766F14.2 |
| 1 | 178916457 | 178916656 | 1.354541906 | RALGPS2 |
| 4 | 100187930 | 100188143 | -2.132503441 | RP11-15B17.1 |
| 4 | 100187930 | 100188143 | -2.132503441 | DDIT4L |
| 4 | 102570759 | 102570950 | 4.804589369 | NFKB1 |
| 4 | 102839714 | 102839877 | 1.026105891 | RP11-10L12.4 |
| 4 | 102839714 | 102839877 | 1.026105891 | UBE2D3 |
| 4 | 105473284 | 105473443 | 5.721471748 | PPA2 |
| 4 | 106316191 | 106316340 | 6.851735523 | AIMP1 |
| 4 | 106316191 | 106316340 | 6.851735523 | TBCK |
| 4 | 106320699 | 106320903 | 7.338529458 | AIMP1 |
| 4 | 106320699 | 106320903 | 7.338529458 | TBCK |
| 4 | 106923850 | 106924086 | 1.603414872 | DKK2 |
| 4 | 107034419 | 107034699 | 3.12240229 | DKK2 |
| 4 | 107035984 | 107036193 | 2.135741365 | DKK2 |
| 4 | 107909610 | 107909797 | 3.114409345 | SGMS2 |
| 4 | 107909610 | 107909797 | 3.114409345 | RP11-286E11.1 |
| 4 | 107996845 | 107997072 | -7.554119284 | HADH |
| 4 | 110618093 | 110618282 | 4.29248125 | PITX2 |
| 4 | 112343221 | 112343395 | -6.769276357 | ALPK1 |
| 4 | 113761280 | 113761477 | 1.476857421 | CAMK2D |
| 4 | 118782389 | 118782502 | 4.975642357 | SEC24D |
| 4 | 119027013 | 119027211 | -2.05173683 | SYNPO2 |

| | | | | |
|---|---|---|---|---|
| 3 | 123914989 | 123915193 | -2.006657927 | CCDC14 |
| 3 | 123946952 | 123947155 | -1.915114162 | CCDC14 |
| 3 | 124737592 | 124737793 | -1.481763275 | UMPS |
| 1 | 161442688 | 161442876 | -1.133713087 | RP11-122G18.10 |
| 3 | 124969441 | 124969641 | 1.508654918 | HEG1 |
| 3 | 124970582 | 124970779 | 2.40003527 | HEG1 |
| 3 | 125012776 | 125012992 | 2.140705335 | HEG1 |
| 3 | 125013754 | 125013952 | 1.823194232 | HEG1 |
| 3 | 125083333 | 125083528 | 8.411615194 | SLC12A8 |
| 3 | 125107731 | 125107973 | 6.635043568 | SLC12A8 |
| 3 | 126269236 | 126269432 | -3.743157456 | RP11-71E19.1 |
| 3 | 126351992 | 126352116 | -3.318767382 | KLF15 |
| 3 | 126451652 | 126451903 | -1.397702405 | ZXDC |
| 3 | 127621723 | 127622011 | -3.177185404 | MCM2 |
| 1 | 7841533 | 7841684 | 2.300231797 | PER3 |
| 3 | 128071006 | 128071206 | -11.84282525 | SEC61A1 |
| 3 | 128071006 | 128071206 | -11.84282525 | RUVBL1 |
| 3 | 128496870 | 128497067 | -2.27927071 | GATA2-AS1 |
| 3 | 128501872 | 128502036 | -1.260385926 | GATA2-AS1 |
| 1 | 161525967 | 161526168 | 2.548923662 | HSPA6 |
| 1 | 161525967 | 161526168 | 2.548923662 | RP11-25K21.6 |
| 3 | 128538512 | 128538747 | 2.410746693 | TMED10P2 |
| 3 | 128865188 | 128865303 | 1.897340546 | RP11-723O4.2 |
| 3 | 128870627 | 128870825 | 1.245453855 | RP11-723O4.2 |
| 3 | 128880983 | 128881183 | -8.446887762 | ACAD9 |
| 3 | 128912685 | 128912907 | -1.051969776 | RP11-723O4.6 |
| 3 | 128912685 | 128912907 | -1.051969776 | KIAA1257 |
| 3 | 128912685 | 128912907 | -1.051969776 | ACAD9 |
| 3 | 128923361 | 128923585 | -1.582955969 | RP11-723O4.6 |
| 3 | 128923361 | 128923585 | -1.582955969 | KIAA1257 |
| 3 | 129122474 | 129122672 | 2.177647708 | ISY1-RAB43 |
| 3 | 129122474 | 129122672 | 2.177647708 | RAB43 |
| 3 | 129390429 | 129390636 | -1.614209059 | RPL32P3 |
| 3 | 129390429 | 129390636 | -1.614209059 | RP11-529F4.1 |
| 3 | 129434041 | 129434228 | -2.681042611 | MBD4 |
| 3 | 129556137 | 129556403 | 1.74241007 | PLXND1 |
| 1 | 161749596 | 161749764 | 1.00318075 | DUSP12 |
| 3 | 129905711 | 129905878 | -2.784305148 | TMCC1-AS1 |
| 3 | 130744742 | 130744942 | -1.162400233 | PIK3R4 |
| 3 | 131024701 | 131024880 | -2.561973007 | ASTE1 |
| 3 | 131383249 | 131383458 | -1.707533954 | NUDT16 |
| 1 | 161828880 | 161829040 | -4.277705411 | ATF6 |
| 3 | 132660070 | 132660262 | 1.887522303 | UBA5 |
| 3 | 132660070 | 132660262 | 1.887522303 | NPHP3-ACAD11 |
| 3 | 132660070 | 132660262 | 1.887522303 | ACAD11 |
| 3 | 134556191 | 134556390 | -1.56140738 | CEP63 |
| 3 | 136001580 | 136001772 | 1.693257274 | PPP2R3A |
| 3 | 136819323 | 136819502 | 5.169368191 | SLC35G2 |
| 3 | 136855199 | 136855352 | 2.645077874 | SLC35G2 |
| 3 | 136855199 | 136855352 | 2.645077874 | NCK1-AS1 |
| 3 | 138161977 | 138162173 | -1.490006555 | DBR1 |
| 3 | 138349669 | 138349870 | 4.538407799 | MRAS |
| 3 | 138403629 | 138403792 | 1.195008223 | MRAS |
| 3 | 138404318 | 138404507 | 1.64317603 | MRAS |
| 3 | 142392762 | 142392952 | 5.805512399 | XRN1 |
| 3 | 143121989 | 143122195 | 2.720907928 | CHST2 |
| 3 | 143985509 | 143985689 | -2.156405405 | C3orf58 |
| 3 | 146070653 | 146070859 | 2.353392846 | RP11-274H2.2 |
| 3 | 146070653 | 146070859 | 2.353392846 | PLOD2 |
| 3 | 146462637 | 146462748 | 4.175101274 | PLSCR2 |
| 3 | 149813141 | 149813330 | 3.116221516 | ANKUB1 |
| 3 | 149813141 | 149813330 | 3.116221516 | RNF13 |
| 3 | 149960761 | 149960965 | -1.645003102 | RNF13 |
| 3 | 149960761 | 149960965 | -1.645003102 | ANKUB1 |
| 3 | 151881518 | 151881712 | -1.538608647 | AADACL2-AS1 |
| 3 | 151881518 | 151881712 | -1.538608647 | SUCNR1 |
| 3 | 152835172 | 152835372 | -4.178776002 | P2RY1 |
| 3 | 152839585 | 152839818 | -1.932344365 | P2RY1 |
| 3 | 155763676 | 155763894 | -1.845292356 | C3orf33 |
| 3 | 155937620 | 155937820 | -1.641586026 | GMPS |
| 3 | 157443028 | 157443186 | 5.235067513 | PTX3 |
| 3 | 157443028 | 157443186 | 5.235067513 | VEPH1 |
| 3 | 158238800 | 158238997 | 6.178501046 | RSRC1 |
| 1 | 166859415 | 166859616 | -4.060523894 | TADA1 |
| 3 | 158691287 | 158691481 | -1.270794381 | GFM1 |
| 3 | 160226594 | 160226793 | -2.324370211 | C3orf80 |
| 3 | 160438640 | 160438857 | -2.310464791 | RP11-432B6.3 |
| 3 | 160438640 | 160438857 | -2.310464791 | TRIM59 |
| 3 | 160550663 | 160550862 | 5.582453463 | KPNA4 |
| 3 | 160568250 | 160568446 | -1.489331412 | KRT8P12 |
| 3 | 160754695 | 160754819 | -4.672147954 | RP11-16N11.2 |
| 3 | 161044818 | 161044933 | -2.032065169 | PPM1L |
| 3 | 161047890 | 161048081 | -1.116177394 | PPM1L |
| 3 | 161069029 | 161069175 | -1.731918511 | PPM1L |
| 3 | 165474584 | 165474775 | -2.032506075 | LINC01322 |
| 3 | 167523566 | 167523763 | -2.203408496 | WDR49 |
| 3 | 167524439 | 167524622 | -4.646160816 | WDR49 |
| 3 | 167549628 | 167549922 | -2.139487975 | WDR49 |
| 3 | 170128792 | 170128990 | -8.567418697 | PHC3 |
| 3 | 171602892 | 171603151 | -3.322205483 | PLD1 |
| 3 | 171872661 | 171872853 | -5.206813965 | TMEM212 |
| 3 | 172506419 | 172506633 | 6.130442341 | TNFSF10 |
| 3 | 172828958 | 172829092 | -5.793888758 | ATP5G1P4 |
| 3 | 177025246 | 177025440 | -2.429254054 | TBL1XR1 |
| 3 | 179333766 | 179333960 | -1.488002651 | ZNF639 |
| 3 | 183154112 | 183154311 | 5.484961464 | LAMP3 |
| 3 | 183491757 | 183491955 | -1.014026459 | KLHL6 |
| 3 | 183508302 | 183508403 | -1.44395646 | KLHL6 |
| 3 | 183679021 | 183679219 | -1.047556188 | KLHL24 |
| 1 | 168242629 | 168242832 | 1.623642391 | SFT2D2 |
| 3 | 183811047 | 183811249 | 1.010822454 | YEATS2 |
| 3 | 183816623 | 183816807 | -2.511106759 | MAP6D1 |
| 3 | 184344692 | 184344890 | -1.376680016 | FAM131A |
| 3 | 184711332 | 184711618 | -1.850194906 | MAGEF1 |
| 3 | 185193255 | 185193443 | -1.830488189 | EHHADH |
| 1 | 169131363 | 169131559 | 3.244263635 | ATP1B1 |
| 3 | 185470254 | 185470414 | -6.037070731 | MAP3K13 |
| 3 | 185482391 | 185482535 | 1.611692061 | TMEM41A |
| 4 | 119030075 | 119030292 | -1.485380149 | SYNPO2 |
| 4 | 119057529 | 119057726 | -1.880687073 | SYNPO2 |
| 4 | 119482968 | 119483073 | -6.126923742 | RP11-33B1.1 |
| 4 | 122271219 | 122271456 | -1.000227689 | KIAA1109 |
| 4 | 125317033 | 125317293 | 6.620544689 | FAT4 |
| 4 | 127809355 | 127809549 | -5.114409345 | HSPA4L |
| 4 | 127809355 | 127809549 | -5.114409345 | RP11-50D9.4 |
| 4 | 127832617 | 127832781 | 1.273826143 | HSPA4L |
| 4 | 128872038 | 128872250 | 1.590620728 | JADE1 |
| 4 | 128872038 | 128872250 | 1.590620728 | SCLT1 |
| 4 | 129069685 | 129069883 | -4.821928095 | SCLT1 |
| 1 | 180196456 | 180196756 | 2.40627385 | QSOX1 |
| 4 | 139115759 | 139115942 | -6.369257266 | ELF2 |
| 4 | 139597456 | 139597700 | 1.129870352 | SETD7 |
| 4 | 139600946 | 139601111 | 4.497176718 | SETD7 |
| 4 | 139889616 | 139889756 | 1.442754463 | MAML3 |
| 4 | 139890605 | 139890805 | 1.007696814 | MAML3 |
| 4 | 145033130 | 145033297 | -4.953445298 | ANAPC10 |
| 4 | 145557836 | 145558026 | -1.66922771 | SMAD1 |
| 4 | 145760740 | 145760981 | 2.983063531 | C4orf51 |
| 4 | 145760740 | 145760981 | 2.983063531 | ZNF827 |
| 4 | 146640842 | 146641043 | 1.389558693 | POU4F2 |
| 4 | 147634277 | 147634484 | 1.503574167 | TMEM184C |
| 4 | 148072140 | 148072355 | 1.507479794 | ARHGAP10 |
| 4 | 150828276 | 150828450 | 1.645475102 | LRBA |
| 4 | 152346995 | 152347188 | -6.476643534 | FBXW7 |
| 4 | 152626961 | 152627162 | -1.224135852 | TMEM154 |
| 4 | 153334931 | 153335131 | -1.111258605 | TRIM2 |
| 4 | 153703349 | 153703543 | -1.400269849 | TLR2 |
| 4 | 153704986 | 153705198 | -1.425576792 | TLR2 |
| 4 | 155710962 | 155711147 | -1.22704681 | GUCY1A3 |
| 4 | 156762829 | 156763012 | -1.195430366 | PDGFC |
| 4 | 158568163 | 158568360 | 1.12465686 | RXFP1 |
| 4 | 158868882 | 158869089 | -1.221405287 | FNIP2 |
| 4 | 158880143 | 158880429 | -1.649587587 | FNIP2 |
| 4 | 158881254 | 158881454 | -3.023603276 | FNIP2 |
| 4 | 158904635 | 158904859 | -1.004135045 | FNIP2 |
| 4 | 158904635 | 158904859 | -1.004135045 | C4orf45 |
| 4 | 159353625 | 159353826 | -1.03410777 | RAPGEF2 |
| 4 | 168512001 | 168512197 | -1.834339332 | PALLD |
| 4 | 168512001 | 168512197 | -1.834339332 | DDX60L |
| 4 | 168952793 | 168952896 | -4.616809838 | CBR4 |
| 1 | 9037438 | 9037630 | -2.463824578 | SLC2A5 |
| 1 | 181089798 | 181089997 | 3.134553555 | IER5 |
| 4 | 182793464 | 182793664 | 2.019084356 | TENM3 |
| 4 | 182800328 | 182800527 | 4.506288425 | TENM3 |
| 4 | 182890894 | 182891094 | 4.506470653 | DCTD |
| 4 | 184009358 | 184009475 | -1.527120953 | STOX2 |
| 1 | 182585904 | 182586131 | -1.325835809 | RNASEL |
| 4 | 186192409 | 186192515 | -3.921175172 | CYP4V2 |
| 4 | 186210424 | 186210587 | -1.448329555 | KLKB1 |
| 4 | 186210424 | 186210587 | -1.448329555 | CYP4V2 |
| 1 | 182599061 | 182599262 | -1.227340091 | RGS16 |
| 4 | 189982653 | 189982796 | 1.204420617 | TUBB7P |

| | | | | |
|---|---|---|---|---|
| 3 | 138655212 | 138655413 | -1.794539992 | PIK3CB |
| 3 | 139407484 | 139407654 | 6.671801069 | RP11-319G6.1 |
| 3 | 139494272 | 139494479 | 2.67904193 | RP11-319G6.1 |
| 3 | 139494272 | 139494479 | 2.67904193 | ACTG1P1 |
| 1 | 162383279 | 162383588 | 1.82785815 | C1orf226 |
| 1 | 162383279 | 162383588 | 1.82785815 | RP11-565P22.6 |
| 3 | 141292335 | 141292504 | 3.625147292 | PXYLP1 |
| 3 | 141292335 | 141292504 | 3.625147292 | RP11-438D8.2 |
| 3 | 141292585 | 141292762 | 3.018386501 | RP11-438D8.2 |
| 3 | 141292585 | 141292762 | 3.018386501 | PXYLP1 |
| 3 | 141443821 | 141444022 | 1.153825105 | ZBTB38 |
| 3 | 142135547 | 142135738 | -4.135197726 | TFDP2 |
| 1 | 162383847 | 162384011 | 2.43791733 | C1orf226 |
| 3 | 142141262 | 142141417 | -6.077092605 | TFDP2 |
| 3 | 142562415 | 142562540 | -2.657643892 | ATR |
| 3 | 143121989 | 143122195 | 2.153402576 | CHST2 |
| 3 | 143985509 | 143985689 | -1.264091802 | C3orf58 |
| 3 | 143989670 | 143989870 | -1.629088755 | C3orf58 |
| 3 | 146070653 | 146070859 | -4.200191517 | PLOD2 |
| 3 | 146070653 | 146070859 | -4.200191517 | RP11-274H2.2 |
| 3 | 149960761 | 149960965 | 1.54168077 | RNF13 |
| 3 | 149960761 | 149960965 | 1.54168077 | ANKUB1 |
| 3 | 150410967 | 150411077 | -1.290486609 | TSC22D2 |
| 3 | 150458406 | 150458613 | 1.290584864 | TSC22D2 |
| 1 | 162780235 | 162780415 | 1.861233012 | DDR2 |
| 3 | 151881518 | 151881712 | 3.031055201 | SUCNR1 |
| 3 | 151881518 | 151881712 | 3.031055201 | AADACL2-AS1 |
| 3 | 152268308 | 152268507 | -1.171733048 | MBNL1-AS1 |
| 3 | 152268308 | 152268507 | -1.171733048 | MBNL1 |
| 3 | 152299843 | 152300056 | -1.52424997 | MBNL1 |
| 3 | 152464921 | 152465152 | -1.471958082 | MBNL1 |
| 3 | 152464921 | 152465152 | -1.471958082 | RP11-362A9.3 |
| 1 | 166849676 | 166849875 | -1.376870883 | POGK |
| 3 | 152836466 | 152836667 | -1.932547424 | P2RY1 |
| 3 | 153162912 | 153163116 | 1.106466397 | RAP2B |
| 3 | 155937620 | 155937820 | -3.178704364 | GMPS |
| 3 | 156678291 | 156678492 | 1.227339939 | TIPARP |
| 1 | 166852670 | 166852872 | -1.601369329 | POGK |
| 3 | 156693884 | 156694091 | 1.538808393 | TIPARP |
| 3 | 156849718 | 156849896 | -1.543731421 | LEKR1 |
| 3 | 157443028 | 157443186 | -5.925347962 | VEPH1 |
| 3 | 157443028 | 157443186 | -5.925347962 | PTX3 |
| 3 | 158238800 | 158238997 | -5.163214744 | RSRC1 |
| 3 | 160226594 | 160226793 | 1.134713949 | C3orf80 |
| 3 | 160438640 | 160438857 | -2.271622093 | RP11-432B6.3 |
| 3 | 160438640 | 160438857 | -2.271622093 | TRIM59 |
| 3 | 160754695 | 160754819 | -1.073778594 | RP11-16N11.2 |
| 3 | 161047890 | 161048081 | 1.514007188 | PPM1L |
| 3 | 161069029 | 161069175 | 1.629212682 | PPM1L |
| 3 | 165474584 | 165474775 | 3.985951373 | LINC01322 |
| 3 | 167523566 | 167523763 | -4.352850748 | WDR49 |
| 3 | 167524439 | 167524622 | -4.888663711 | WDR49 |
| 3 | 167549628 | 167549922 | -4.249069805 | WDR49 |
| 3 | 167627046 | 167627233 | -2.910462531 | WDR49 |
| 3 | 185482391 | 185482535 | 1.611692061 | MAP3K13 |
| 3 | 185482740 | 185482911 | 2.213993947 | TMEM41A |
| 3 | 185482740 | 185482911 | 2.213993947 | MAP3K13 |
| 3 | 187042859 | 187043095 | -1.48113645 | ST6GAL1 |
| 3 | 187075762 | 187075961 | -1.537723008 | ST6GAL1 |
| 3 | 187077889 | 187078090 | -1.767922959 | ST6GAL1 |
| 3 | 187371012 | 187371205 | 2.010263988 | RTP4 |
| 3 | 188153285 | 188153475 | -1.666859762 | LPP |
| 3 | 188153285 | 188153475 | -1.666859762 | LPP-AS2 |
| 3 | 188576537 | 188576734 | 7.090925534 | LPP |
| 3 | 189284892 | 189285041 | -2.653866045 | TPRG1 |
| 3 | 194138163 | 194138363 | -1.486444006 | HES1 |
| 3 | 194587825 | 194588026 | -1.028511361 | TMEM44-AS1 |
| 3 | 194587825 | 194588026 | -1.028511361 | TMEM44 |
| 3 | 194629380 | 194629615 | -1.859836405 | TMEM44 |
| 3 | 194637760 | 194637893 | 5.505963033 | AC046143.3 |
| 3 | 195069517 | 195069724 | -1.035605741 | XXYLT1 |
| 3 | 195618941 | 195619100 | -1.884732317 | MUC20P1 |
| 3 | 195868311 | 195868440 | 6.303318272 | TNK2 |
| 1 | 171535309 | 171535509 | 7.471257253 | PRRC2C |
| 3 | 196356665 | 196356860 | -1.944427581 | UBXN7 |
| 3 | 196472073 | 196472275 | -2.644585299 | RNF168 |
| 3 | 196487494 | 196487700 | -3.111241406 | RNF168 |
| 3 | 196488617 | 196488818 | -2.981567016 | RNF168 |
| 3 | 196496379 | 196496563 | -3.114454022 | RNF168 |
| 3 | 196502915 | 196503161 | -2.251988998 | RNF168 |
| 3 | 196554321 | 196554514 | -1.324831125 | WDR53 |
| 3 | 196584204 | 196584404 | -1.294950703 | FBXO45 |
| 3 | 196649220 | 196649436 | -8.43089483 | NRROS |
| 3 | 196649220 | 196649436 | -8.43089483 | PIGX |
| 3 | 196655707 | 196655905 | -8.954149395 | PIGX |
| 3 | 196655707 | 196655905 | -8.954149395 | NRROS |
| 3 | 196659975 | 196660174 | -1.316536257 | PIGX |
| 3 | 196659975 | 196660174 | -1.316536257 | NRROS |
| 3 | 196661310 | 196661505 | -2.424492127 | NRROS |
| 3 | 196661310 | 196661505 | -2.424492127 | PIGX |
| 3 | 196830417 | 196830635 | -1.356333335 | PAK2 |
| 3 | 196910909 | 196911218 | 1.12043305 | SENP5 |
| 3 | 196912209 | 196912347 | 1.998335991 | SENP5 |
| 3 | 196943216 | 196943417 | -1.183238588 | NCBP2-AS2 |
| 3 | 196947060 | 196947261 | -1.728429875 | PIGZ |
| 3 | 196947976 | 196948146 | -1.494159097 | PIGZ |
| 3 | 196963014 | 196963204 | -2.198220509 | PIGZ |
| 3 | 197512020 | 197512210 | -1.165178358 | BDH1 |
| 3 | 197674374 | 197674573 | 1.355711058 | RUBCN |
| 4 | 160935 | 161036 | -1.236405324 | ZNF718 |
| 4 | 161836 | 162035 | -1.978152045 | ZNF718 |
| 4 | 254030 | 254173 | -5.626923742 | ZNF876P |
| 4 | 373840 | 374034 | -3.332441838 | ZNF141 |
| 4 | 375087 | 375240 | -2.40650267 | ZNF141 |
| 4 | 423421 | 423671 | -1.864674673 | RP11-2H3.7 |
| 4 | 426300 | 426479 | -3.001317425 | ABCA11P |
| 4 | 426300 | 426479 | -3.001317425 | RP11-2H3.7 |
| 4 | 426300 | 426479 | -3.001317425 | ZNF721 |
| 1 | 182599995 | 182600205 | -1.631599065 | RGS16 |
| 5 | 199052 | 199250 | -1.817289188 | CCDC127 |
| 5 | 322194 | 322411 | 2.264556785 | PDCD6 |
| 5 | 322194 | 322411 | 2.264556785 | AHRR |
| 5 | 388720 | 388926 | 7.915579385 | AHRR |
| 5 | 389622 | 389827 | 2.068026441 | AHRR |
| 5 | 401986 | 402190 | 7.236789462 | AHRR |
| 5 | 443575 | 443755 | 2.443850305 | EXOC3 |
| 5 | 476378 | 476618 | 1.780514966 | SLC9A3 |
| 5 | 476378 | 476618 | 1.780514966 | CTD-2228K2.7 |
| 5 | 662414 | 662602 | 8.546997594 | TPPP |
| 5 | 662414 | 662602 | 8.546997594 | CEP72 |
| 1 | 9103981 | 9104179 | 1.477626809 | GPR157 |
| 5 | 840100 | 840284 | 6.44807158 | ZDHHC11 |
| 5 | 1051710 | 1051922 | 1.205492898 | SLC12A7 |
| 5 | 1052183 | 1052382 | -1.79248125 | SLC12A7 |
| 5 | 1054509 | 1054610 | 3.298183632 | SLC12A7 |
| 5 | 1056360 | 1056556 | 2.170348949 | SLC12A7 |
| 5 | 1060706 | 1060898 | 2.90763006 | SLC12A7 |
| 5 | 1064523 | 1064717 | -4.93518236 | SLC12A7 |
| 1 | 184432238 | 184432406 | -4.178776002 | C1orf21 |
| 5 | 1610346 | 1610546 | -1.091740499 | RP11-43F13.1 |
| 5 | 1610346 | 1610546 | -1.091740499 | CTD-2012J19.3 |
| 5 | 1611777 | 1611978 | -1.553303523 | RP11-43F13.1 |
| 5 | 1806718 | 1806920 | -7.870707158 | NDUFS6 |
| 5 | 5479596 | 5479769 | 6.220564311 | ICE1 |
| 5 | 10649505 | 10649694 | 10.67695892 | ANKRD33B |
| 5 | 10649896 | 10650097 | 9.631630015 | ANKRD33B |
| 5 | 10657083 | 10657259 | -2.000417326 | ANKRD33B |
| 5 | 10680170 | 10680374 | 1.009636423 | DAP |
| 5 | 14610389 | 14610652 | -1.012589507 | FAM105A |
| 5 | 14808975 | 14809158 | -5.324628089 | ANKH |
| 5 | 16465402 | 16465593 | 1.250816276 | ZNF622 |
| 5 | 16474761 | 16474997 | -1.327978839 | FAM134B |
| 5 | 16666267 | 16666466 | -1.785468277 | MYO10 |
| 5 | 16701144 | 16701346 | -1.470349712 | MYO10 |
| 5 | 17713737 | 17713936 | -1.210731884 | RP11-454P21.1 |
| 5 | 18707721 | 18707907 | 5.587462841 | CTD-2023M8.1 |
| 5 | 20304850 | 20305070 | 1.014457044 | CDH18 |
| 5 | 20304850 | 20305070 | 1.014457044 | RP11-420O16.2 |
| 5 | 21473591 | 21473790 | 1.654004849 | GUSBP1 |
| 5 | 21474142 | 21474317 | 1.61326402 | GUSBP1 |
| 5 | 21580558 | 21580756 | -1.536954615 | GUSBP1 |
| 5 | 25190316 | 25190502 | 1.745926548 | RP11-549K20.1 |
| 5 | 31496743 | 31496931 | 6.155732258 | DROSHA |
| 5 | 34050259 | 34050464 | -2.17636786 | C1QTNF3-AMACR |
| 5 | 34823182 | 34823417 | -1.22335512 | RAI14 |
| 5 | 36686069 | 36686272 | -1.318719978 | CTD-2353F22.1 |
| 5 | 36686069 | 36686272 | -1.318719978 | SLC1A3 |
| 1 | 192558118 | 192558237 | 1.138420103 | RP5-1011O1.2 |
| 5 | 36877198 | 36877359 | 7.794591484 | NIPBL |
| 5 | 37519283 | 37519492 | 1.348161805 | WDR70 |
| 5 | 37524670 | 37524821 | -7.096339049 | WDR70 |
| 5 | 38942124 | 38942336 | -1.902674113 | OSMR |

| | | | | |
|---|---|---|---|---|
| 3 | 167636916 | 167637091 | -3.619483578 | WDR49 |
| 3 | 167651498 | 167651603 | -2.827811823 | WDR49 |
| 3 | 169767852 | 169768042 | -2.634882987 | ACTRT3 |
| 3 | 170128792 | 170128990 | -3.411999243 | PHC3 |
| 1 | 167697285 | 167697485 | -3.000546042 | RCSD1 |
| 3 | 170360322 | 170360529 | 1.912716969 | SKIL |
| 3 | 170390976 | 170391169 | 2.445095582 | SKIL |
| 3 | 170392155 | 170392360 | 2.128010849 | SKIL |
| 3 | 170418438 | 170418614 | 2.242713414 | RP11-469J4.3 |
| 1 | 167704715 | 167704913 | -2.31831083 | RCSD1 |
| 3 | 172828958 | 172829092 | -1.171479911 | ATP5G1P4 |
| 3 | 179259077 | 179259302 | -2.23764574 | LRRFIP1P1 |
| 3 | 179259077 | 179259302 | -2.23764574 | KCNMB3 |
| 3 | 179333382 | 179333575 | -1.358841278 | ZNF639 |
| 3 | 179333766 | 179333960 | -1.595735306 | ZNF639 |
| 3 | 179784012 | 179784201 | -2.462136337 | USP13 |
| 3 | 183491757 | 183491955 | 4.500578168 | KLHL6 |
| 3 | 183508302 | 183508403 | 7.67367227 | KLHL6 |
| 3 | 183522732 | 183522897 | 5.892317423 | KLHL6 |
| 3 | 183650825 | 183651002 | 2.887640011 | KLHL24 |
| 3 | 183679021 | 183679219 | 3.334461031 | KLHL24 |
| 3 | 183811047 | 183811249 | -1.534530897 | YEATS2 |
| 3 | 183816623 | 183816807 | -1.499400447 | MAP6D1 |
| 3 | 184321715 | 184321915 | -1.363300497 | EIF4G1 |
| 3 | 184334832 | 184335035 | -1.409655714 | EIF4G1 |
| 3 | 185193255 | 185193443 | -1.586128171 | EHHADH |
| 1 | 920545 | 920739 | 6.177117733 | RP11-54O7.3 |
| 1 | 169131363 | 169131559 | 5.527641218 | ATP1B1 |
| 3 | 185470254 | 185470414 | 6.037070731 | MAP3K13 |
| 3 | 185482391 | 185482535 | 1.10433666 | TMEM41A |
| 3 | 185482391 | 185482535 | 1.10433666 | MAP3K13 |
| 3 | 185482740 | 185482911 | -1.683402373 | TMEM41A |
| 3 | 185482740 | 185482911 | -1.683402373 | MAP3K13 |
| 3 | 186148015 | 186148207 | -4.098028457 | DGKG |
| 3 | 186554356 | 186554556 | -1.222103115 | TBCCD1 |
| 1 | 169255393 | 169255580 | 6.440174404 | NME7 |
| 3 | 186826748 | 186826958 | -2.347201279 | RP11-573D15.1 |
| 3 | 187039028 | 187039224 | -3.060951743 | ST6GAL1 |
| 3 | 187042859 | 187043095 | 1.70146204 | ST6GAL1 |
| 3 | 187075762 | 187075961 | 2.752427654 | ST6GAL1 |
| 3 | 187077889 | 187078090 | 2.026844303 | ST6GAL1 |
| 3 | 187729437 | 187729632 | -5.257357027 | BCL6 |
| 3 | 187729437 | 187729632 | -5.257357027 | RP11-211G3.3 |
| 3 | 187738129 | 187738237 | 3.396124015 | BCL6 |
| 3 | 187739457 | 187739630 | 3.967518451 | BCL6 |
| 1 | 169793360 | 169793559 | -2.52849205 | C1orf112 |
| 1 | 169793360 | 169793559 | -2.52849205 | METTL18 |
| 3 | 188153285 | 188153475 | -1.365300729 | LPP |
| 3 | 188153285 | 188153475 | -1.365300729 | LPP-AS2 |
| 3 | 189049732 | 189049990 | 1.65632858 | TPRG1 |
| 3 | 189052688 | 189052887 | 1.094235192 | TPRG1 |
| 3 | 189118141 | 189118329 | 6.191852146 | TPRG1 |
| 1 | 169871599 | 169871835 | -1.861039133 | SCYL3 |
| 3 | 191375136 | 191375432 | 1.92968532 | CCDC50 |
| 4 | 426564 | 426712 | -2.992290349 | ABCA11P |
| 4 | 426564 | 426712 | -2.992290349 | RP11-2H3.7 |
| 4 | 426564 | 426712 | -2.992290349 | ZNF721 |
| 4 | 442554 | 442753 | -2.201374612 | ABCA11P |
| 4 | 442554 | 442753 | -2.201374612 | RP11-2H3.7 |
| 4 | 442554 | 442753 | -2.201374612 | ZNF721 |
| 4 | 443298 | 443497 | -1.494444923 | RP11-2H3.7 |
| 4 | 443298 | 443497 | -1.494444923 | ZNF721 |
| 4 | 443298 | 443497 | -1.494444923 | ABCA11P |
| 4 | 443973 | 444173 | -1.821461077 | ZNF721 |
| 4 | 443973 | 444173 | -1.821461077 | RP11-2H3.7 |
| 4 | 443973 | 444173 | -1.821461077 | ABCA11P |
| 4 | 497373 | 497549 | -1.089209898 | ZNF721 |
| 4 | 577958 | 578157 | -5.953445298 | RP11-1263C18.2 |
| 4 | 674970 | 675131 | -1.824356938 | MYL5 |
| 4 | 675305 | 675409 | 6.515505954 | MYL5 |
| 4 | 687892 | 688051 | -2.071152255 | MFSD7 |
| 4 | 762020 | 762218 | -1.33329675 | PCGF3 |
| 4 | 879302 | 879499 | -1.809409597 | GAK |
| 4 | 880359 | 880535 | 6.481086516 | GAK |
| 4 | 959723 | 959916 | -1.423162471 | DGKQ |
| 4 | 989167 | 989311 | -5.218355771 | IDUA |
| 4 | 989167 | 989311 | -5.218355771 | SLC26A1 |
| 1 | 172570964 | 172571158 | 3.616330378 | SUCO |
| 4 | 990214 | 990362 | -6.715552899 | SLC26A1 |
| 4 | 990214 | 990362 | -6.715552899 | IDUA |
| 4 | 1175237 | 1175409 | -2.078059601 | SPON2 |
| 4 | 1235504 | 1235713 | -1.570810831 | CTBP1 |
| 4 | 1250555 | 1250745 | -1.849575429 | CTBP1-AS2 |
| 4 | 1251547 | 1251747 | -1.246217943 | CTBP1-AS2 |
| 4 | 1385805 | 1385992 | -1.28822932 | UVSSA |
| 4 | 1728305 | 1728502 | -1.826960118 | TACC3 |
| 4 | 1978744 | 1978849 | -1.266175512 | WHSC1 |
| 4 | 1983281 | 1983476 | -2.135962357 | NELFA |
| 4 | 2744808 | 2744994 | 1.482768609 | TNIP2 |
| 4 | 2829755 | 2829979 | 1.272542124 | SH3BP2 |
| 4 | 2833812 | 2834006 | 1.684638988 | SH3BP2 |
| 4 | 3316324 | 3316505 | -5.689959909 | RGS12 |
| 1 | 173185841 | 173186022 | -1.058449646 | TNFSF4 |
| 4 | 3375029 | 3375194 | -4.482892142 | RGS12 |
| 4 | 3378226 | 3378477 | -1.211372919 | RGS12 |
| 4 | 3487793 | 3487956 | 1.457098892 | DOK7 |
| 4 | 3768005 | 3768205 | -1.97376629 | ADRA2C |
| 4 | 3942718 | 3942862 | -4.414944181 | FAM86EP |
| 4 | 3942718 | 3942862 | -4.414944181 | AC226119.5 |
| 4 | 4236755 | 4236917 | -2.200292519 | TMEM128 |
| 1 | 173186477 | 173186678 | -1.770284191 | TNFSF4 |
| 4 | 4302419 | 4302622 | -1.093639445 | ZBTB49 |
| 4 | 6642197 | 6642388 | -10.30634878 | MRFAP1 |
| 4 | 6675272 | 6675491 | -3.229472696 | AC093323.3 |
| 1 | 173728040 | 173728142 | -5.975642357 | KLHL20 |
| 4 | 6709393 | 6709595 | -1.439073886 | MRFAP1L1 |
| 4 | 6717050 | 6717212 | -1.627635497 | BLOC1S4 |
| 4 | 6861400 | 6861570 | -1.907008653 | KIAA0232 |
| 5 | 38942124 | 38942336 | -1.902674113 | RICTOR |
| 5 | 39005378 | 39005502 | -7.789952577 | RICTOR |
| 5 | 39153535 | 39153727 | -5.321477112 | FYB |
| 5 | 40680047 | 40680257 | 8.983041468 | PTGER4 |
| 5 | 40680047 | 40680257 | 8.983041468 | TTC33 |
| 5 | 40721129 | 40721316 | 6.161386588 | TTC33 |
| 5 | 40852238 | 40852440 | -1.719928627 | CARD6 |
| 5 | 41742815 | 41743060 | 4.39395128 | OXCT1 |
| 1 | 193218751 | 193219040 | -1.203962599 | CDC73 |
| 5 | 43043069 | 43043258 | 1.173294006 | CTD-2201E18.3 |
| 5 | 43043069 | 43043258 | 1.173294006 | AC025171.1 |
| 5 | 43043069 | 43043258 | 1.173294006 | ANXA2R |
| 5 | 43051707 | 43051901 | -1.212996054 | CTD-2201E18.3 |
| 5 | 43061705 | 43061887 | -2.322422564 | CTD-2035E11.5 |
| 5 | 43061705 | 43061887 | -2.322422564 | CTD-2201E18.3 |
| 5 | 43136764 | 43136940 | -7.288005937 | ZNF131 |
| 5 | 43161647 | 43161779 | -7.891959523 | ZNF131 |
| 5 | 43245742 | 43245922 | 1.734384742 | NIM1K |
| 5 | 43298655 | 43298861 | -1.219390708 | HMGCS1 |
| 1 | 925028 | 925204 | 4.888555888 | SAMD11 |
| 1 | 9265044 | 9265241 | 1.094090954 | H6PD |
| 1 | 197101343 | 197101538 | -1.472776108 | ASPM |
| 5 | 43381809 | 43382006 | -1.434197302 | CCL28 |
| 5 | 43446335 | 43446507 | -6.794415866 | TMEM267 |
| 1 | 197104402 | 197104591 | -5.22766361 | ASPM |
| 5 | 53982505 | 53982680 | -2.925227382 | ARL15 |
| 5 | 58974271 | 58974428 | 1.252368913 | PDE4D |
| 5 | 60121801 | 60121986 | 2.111196211 | PDE4D |
| 5 | 61146752 | 61146951 | -6.741781944 | NDUFAF2 |
| 5 | 61347415 | 61347562 | -1.73472492 | RP11-631M6.3 |
| 5 | 61347415 | 61347562 | -1.73472492 | ZSWIM6 |
| 5 | 62545443 | 62545630 | -4.52219706 | IPO11 |
| 5 | 64845766 | 64845994 | -1.496384215 | CWC27 |
| 5 | 66078403 | 66078626 | -1.257633759 | ERBIN |
| 5 | 67163753 | 67163952 | 5.345290028 | MAST4 |
| 5 | 67164527 | 67164755 | 1.055231074 | MAST4 |
| 5 | 67166708 | 67166878 | 1.157731058 | MAST4 |
| 5 | 67182554 | 67182754 | 2.203188454 | CD180 |
| 5 | 67183189 | 67183383 | 2.51108025 | CD180 |
| 5 | 67183615 | 67183791 | 1.694136252 | CD180 |
| 5 | 67183999 | 67184185 | 1.671378587 | CD180 |
| 1 | 200553551 | 200553736 | -3.413675915 | KIF14 |
| 5 | 71074138 | 71074299 | -5.844125155 | RP11-195E2.1 |
| 1 | 200617887 | 200618090 | -1.029841537 | KIF14 |
| 5 | 72443879 | 72444105 | 3.003879748 | ZNF366 |
| 5 | 72461108 | 72461241 | 2.614577421 | ZNF366 |
| 5 | 72817092 | 72817236 | 8.704877771 | TNPO1 |
| 5 | 74635630 | 74635831 | 1.785808058 | ENC1 |
| 5 | 76732254 | 76732427 | 1.592514028 | F2R |
| 5 | 76734384 | 76734599 | 1.185484799 | F2R |
| 5 | 77076987 | 77077182 | -1.701546142 | ZBED3 |
| 5 | 77630877 | 77631068 | 1.549767837 | OTP |
| 5 | 78170006 | 78170327 | -5.998766174 | AP3B1 |
| 5 | 78170733 | 78170960 | -6.152460835 | AP3B1 |

| | | | | |
|---|---|---|---|---|
| 3 | 194138163 | 194138363 | 5.986291361 | HES1 |
| 3 | 194487876 | 194488010 | -1.784003686 | ATP13A3 |
| 3 | 194487876 | 194488010 | -1.784003686 | LINC00884 |
| 3 | 194637760 | 194637893 | -7.588865484 | AC046143.3 |
| 3 | 194642003 | 194642185 | -1.543567825 | LSG1 |
| 3 | 194642003 | 194642185 | -1.543567825 | AC046143.3 |
| 3 | 194652876 | 194653094 | -1.271566293 | LSG1 |
| 3 | 194687517 | 194687719 | -1.726463412 | FAM43A |
| 3 | 195069517 | 195069724 | -1.437113466 | XXYLT1 |
| 3 | 195180180 | 195180341 | -4.308004102 | XXYLT1 |
| 3 | 195726222 | 195726429 | 8.303578451 | LINC00969 |
| 3 | 195726222 | 195726429 | 8.303578451 | MUC20 |
| 3 | 196220170 | 196220337 | -7.988639962 | SLC51A |
| 3 | 196220170 | 196220337 | -7.988639962 | PCYT1A |
| 1 | 171535309 | 171535509 | -8.636424223 | PRRC2C |
| 3 | 196287281 | 196287483 | 1.711305926 | PCYT1A |
| 3 | 196287281 | 196287483 | 1.711305926 | RP11-447L10.1 |
| 3 | 196322314 | 196322513 | 12.80734313 | TM4SF19-AS1 |
| 3 | 196322314 | 196322513 | 12.80734313 | TM4SF19 |
| 3 | 196322314 | 196322513 | 12.80734313 | TM4SF19-TCTEX1D2 |
| 3 | 196356665 | 196356860 | 1.181093917 | UBXN7 |
| 3 | 196561148 | 196561299 | -1.627214687 | WDR53 |
| 3 | 196566981 | 196567182 | -2.034483159 | WDR53 |
| 3 | 196649220 | 196649436 | 8.43089483 | PIGX |
| 3 | 196649220 | 196649436 | 8.43089483 | NRROS |
| 3 | 196655707 | 196655905 | -1.017317584 | NRROS |
| 3 | 196655707 | 196655905 | -1.017317584 | PIGX |
| 1 | 171541846 | 171542083 | -1.17601759 | PRRC2C |
| 3 | 196910909 | 196911218 | -1.067461243 | SENP5 |
| 3 | 196912209 | 196912347 | -1.129482892 | SENP5 |
| 3 | 196943216 | 196943417 | -1.011678638 | NCBP2-AS2 |
| 1 | 8004893 | 8005048 | -3.729270175 | ERRFI1 |
| 3 | 197001954 | 197002093 | 3.66450902 | MFI2-AS1 |
| 3 | 197001954 | 197002093 | 3.66450902 | MELTF |
| 3 | 197512020 | 197512210 | -4.511218451 | BDH1 |
| 3 | 198038521 | 198038673 | -3.310113425 | LMLN |
| 3 | 198038521 | 198038673 | -3.310113425 | LMLN-AS1 |
| 3 | 198058666 | 198058841 | -1.638564171 | ANKRD18DP |
| 1 | 171783863 | 171784063 | -1.592533677 | METTL13 |
| 4 | 160935 | 161036 | -2.028881652 | ZNF718 |
| 4 | 161836 | 162035 | -1.310109371 | ZNF718 |
| 4 | 372987 | 373105 | -2.75571471 | ZNF141 |
| 4 | 373840 | 374034 | -1.399056235 | ZNF141 |
| 4 | 375087 | 375240 | -2.215669656 | ZNF141 |
| 4 | 426300 | 426479 | -1.279562695 | ZNF721 |
| 4 | 426300 | 426479 | -1.279562695 | ABCA11P |
| 4 | 426300 | 426479 | -1.279562695 | RP11-2H3.7 |
| 4 | 426564 | 426712 | -1.570713674 | ZNF721 |
| 4 | 426564 | 426712 | -1.570713674 | RP11-2H3.7 |
| 4 | 426564 | 426712 | -1.570713674 | ABCA11P |
| 4 | 481544 | 481730 | -1.648135802 | ZNF721 |
| 4 | 523516 | 523710 | -1.397929642 | PIGG |
| 1 | 171796597 | 171796796 | -1.286865761 | METTL13 |
| 4 | 537909 | 538106 | -1.913380483 | PIGG |
| 4 | 6862879 | 6863079 | -1.152007035 | KIAA0232 |
| 4 | 6863596 | 6863772 | -1.50962896 | KIAA0232 |
| 4 | 6923839 | 6924045 | -2.011515119 | TBC1D14 |
| 4 | 7030359 | 7030565 | -2.349892839 | TBC1D14 |
| 4 | 7030359 | 7030565 | -2.349892839 | RP11-367J11.2 |
| 4 | 7041363 | 7041562 | -1.169136412 | CCDC96 |
| 4 | 7041363 | 7041562 | -1.169136412 | RP11-367J11.2 |
| 4 | 7042398 | 7042571 | -1.521401748 | RP11-367J11.2 |
| 4 | 7042398 | 7042571 | -1.521401748 | CCDC96 |
| 4 | 7042398 | 7042571 | -1.521401748 | TADA2B |
| 4 | 7671490 | 7671708 | -7.613958362 | SORCS2 |
| 4 | 7717284 | 7717476 | -6.929087655 | SORCS2 |
| 4 | 7737335 | 7737532 | -2.647665904 | SORCS2 |
| 4 | 8227334 | 8227534 | -1.610593007 | SH3TC1 |
| 4 | 8227854 | 8228004 | -1.272062038 | SH3TC1 |
| 4 | 8229790 | 8230024 | -1.592600278 | SH3TC1 |
| 4 | 8232927 | 8233127 | -1.753473076 | SH3TC1 |
| 4 | 8383926 | 8384121 | 1.321223998 | ACOX3 |
| 4 | 8386045 | 8386245 | -1.147848277 | ACOX3 |
| 4 | 8469357 | 8469553 | 1.601699356 | TRMT44 |
| 4 | 9825703 | 9825899 | -1.486207084 | SLC2A9 |
| 4 | 9877853 | 9878010 | 5.877443751 | SLC2A9 |
| 4 | 10442901 | 10443100 | -1.979452263 | ZNF518B |
| 4 | 10446249 | 10446449 | -1.990237143 | ZNF518B |
| 4 | 13544757 | 13544877 | 7.02882667 | NKX3-2 |
| 4 | 13614449 | 13614646 | -1.520798132 | BOD1L1 |
| 4 | 14473185 | 14473384 | -1.674643044 | LINC00504 |
| 4 | 14474130 | 14474367 | -1.595691315 | LINC00504 |
| 4 | 14535460 | 14535648 | -6.856639552 | LINC00504 |
| 4 | 14660634 | 14660819 | -1.247109891 | LINC00504 |
| 4 | 14862088 | 14862321 | -1.41902586 | LINC00504 |
| 4 | 15614124 | 15614318 | -6.418813967 | FBXL5 |
| 4 | 15820401 | 15820583 | 9.719420287 | CD38 |
| 4 | 15821610 | 15821809 | 5.807145408 | CD38 |
| 4 | 15823030 | 15823307 | 8.957107882 | CD38 |
| 4 | 15846956 | 15847135 | 7.127956875 | CD38 |
| 4 | 16227071 | 16227297 | 2.457005726 | TAPT1-AS1 |
| 4 | 16227071 | 16227297 | 2.457005726 | TAPT1 |
| 4 | 16283762 | 16283947 | -2.73170026 | TAPT1-AS1 |
| 4 | 16360779 | 16360896 | -1.540459998 | ZEB2P1 |
| 4 | 17646375 | 17646492 | -1.60158962 | FAM184B |
| 4 | 17843712 | 17843906 | -1.726698316 | LCORL |
| 4 | 17843712 | 17843906 | -1.726698316 | NCAPG |
| 4 | 17850358 | 17850555 | -6.265934388 | LCORL |
| 4 | 17860582 | 17860749 | -3.890679857 | LCORL |
| 1 | 173938682 | 173938862 | -8.607415675 | RC3H1 |
| 4 | 17874063 | 17874250 | -2.013497286 | LCORL |
| 4 | 20253590 | 20253729 | 8.044859715 | SLIT2 |
| 4 | 20619205 | 20619394 | 3.981203324 | SLIT2 |
| 4 | 22388048 | 22388250 | -1.650809012 | ADGRA3 |
| 1 | 173992899 | 173993103 | -7.544207797 | RC3H1 |
| 4 | 24909840 | 24910020 | -2.382058914 | CCDC149 |
| 4 | 25123964 | 25124165 | -2.499909736 | SEPSECS |
| 4 | 27022615 | 27022813 | -1.866052398 | STIM2 |
| 5 | 79456943 | 79457138 | 6.171315149 | HOMER1 |
| 5 | 80429654 | 80429909 | -1.657228507 | ZFYVE16 |
| 5 | 80430329 | 80430521 | -7.331501044 | ZFYVE16 |
| 5 | 80432971 | 80433169 | -7.898729443 | ZFYVE16 |
| 5 | 81239101 | 81239252 | -1.545849917 | CKMT2-AS1 |
| 5 | 81239101 | 81239252 | -1.545849917 | CKMT2 |
| 5 | 82001568 | 82001719 | -5.363960227 | ATG10 |
| 5 | 83522027 | 83522210 | 3.550782026 | VCAN |
| 5 | 83537429 | 83537602 | 3.570677925 | VCAN |
| 5 | 83537429 | 83537602 | 3.570677925 | VCAN-AS1 |
| 5 | 83580928 | 83581177 | 3.724725557 | VCAN-AS1 |
| 5 | 83580928 | 83581177 | 3.724725557 | VCAN |
| 1 | 200971457 | 200971656 | -1.939406914 | KIF21B |
| 5 | 87355731 | 87355884 | 8.958639415 | RASA1 |
| 5 | 88190446 | 88190649 | -6.180971887 | TMEM161B |
| 5 | 88335733 | 88335876 | -6.136606405 | TMEM161B-AS1 |
| 5 | 88350647 | 88350810 | -1.240079862 | TMEM161B-AS1 |
| 1 | 201206410 | 201206556 | -6.408491812 | IGFN1 |
| 5 | 88797290 | 88797490 | -7.164197219 | MEF2C |
| 5 | 88944465 | 88944663 | -1.068251 | MEF2C-AS1 |
| 5 | 89423730 | 89423854 | 2.842609491 | MEF2C-AS1 |
| 5 | 90461409 | 90461614 | 1.274509212 | MBLAC2 |
| 5 | 90473807 | 90473933 | 2.082135446 | MBLAC2 |
| 5 | 90473807 | 90473933 | 2.082135446 | POLR3G |
| 5 | 93587818 | 93587970 | -1.61195389 | NR2F1 |
| 5 | 93814998 | 93815169 | -5.806203897 | FAM172A |
| 1 | 201520320 | 201520525 | 1.174063857 | RP11-134G8.7 |
| 1 | 201520320 | 201520525 | 1.174063857 | RP11-134G8.5 |
| 1 | 201520320 | 201520525 | 1.174063857 | RP11-134G8.6 |
| 5 | 95620230 | 95620418 | 1.970058167 | GPR150 |
| 5 | 96880095 | 96880286 | -7.479502876 | CTD-2260A17.2 |
| 5 | 96880095 | 96880286 | -7.479502876 | ERAP2 |
| 5 | 98779677 | 98779873 | 1.454076612 | RGMB |
| 5 | 98793371 | 98793547 | 1.603186986 | RGMB |
| 5 | 98793371 | 98793547 | 1.603186986 | AC008522.1 |
| 5 | 103276078 | 103276242 | 1.03279779 | C5orf30 |
| 5 | 103559276 | 103559465 | -1.062395484 | NUDT12 |
| 5 | 108208401 | 108208526 | -6.44208526 | FBXL17 |
| 5 | 108211083 | 108211236 | -5.999295215 | FBXL17 |
| 5 | 111484244 | 111484438 | -1.621294568 | CAMK4 |
| 5 | 112891727 | 112891930 | -2.086470593 | REEP5 |
| 5 | 112891727 | 112891930 | -2.086470593 | CTC-487M23.8 |
| 5 | 112891727 | 112891930 | -2.086470593 | ZRSR1 |
| 5 | 116268627 | 116268854 | -1.326373973 | COMMD10 |
| 5 | 116446324 | 116446523 | 6.903677461 | SEMA6A |
| 5 | 119149573 | 119149771 | -1.036264611 | DMXL1 |
| 5 | 120686264 | 120686482 | 1.351213672 | PRR16 |
| 5 | 121998130 | 121998235 | 1.721985563 | SRFBP1 |
| 5 | 122078111 | 122078268 | 10.9043254 | LOX |
| 5 | 122152018 | 122152192 | 5.483189637 | CTC-441N14.2 |
| 5 | 122152018 | 122152192 | 5.483189637 | CTC-441N14.4 |
| 5 | 122152018 | 122152192 | 5.483189637 | ZNF474 |
| 5 | 122451247 | 122451426 | 3.640281055 | MGC32805 |
| 5 | 122451247 | 122451426 | 3.640281055 | SNCAIP |

| | | | | |
|---|---|---|---|---|
| 4 | 577958 | 578157 | 5.953445298 | RP11-1263C18.2 |
| 4 | 674970 | 675131 | 2.49179542 | MYL5 |
| 4 | 675305 | 675409 | -5.941130465 | MYL5 |
| 4 | 857760 | 857923 | 1.147617255 | GAK |
| 4 | 879302 | 879499 | 2.405901606 | GAK |
| 4 | 880359 | 880535 | -7.717672353 | GAK |
| 4 | 989167 | 989311 | -2.504992044 | IDUA |
| 4 | 989167 | 989311 | -2.504992044 | SLC26A1 |
| 1 | 172570964 | 172571158 | -2.816134108 | SUCO |
| 4 | 990214 | 990362 | -1.838525539 | IDUA |
| 4 | 990214 | 990362 | -1.838525539 | SLC26A1 |
| 4 | 990433 | 990565 | -1.548901363 | SLC26A1 |
| 4 | 990433 | 990565 | -1.548901363 | IDUA |
| 4 | 991264 | 991461 | -2.513345392 | IDUA |
| 4 | 991264 | 991461 | -2.513345392 | SLC26A1 |
| 4 | 992402 | 992599 | -2.888200371 | SLC26A1 |
| 4 | 992402 | 992599 | -2.888200371 | IDUA |
| 4 | 1010402 | 1010588 | 4.00199316 | FGFRL1 |
| 4 | 1175237 | 1175409 | -1.934351929 | SPON2 |
| 4 | 1196051 | 1196239 | -3.530242451 | SPON2 |
| 4 | 1235504 | 1235713 | -1.587421528 | CTBP1 |
| 4 | 1250555 | 1250745 | -1.343218072 | CTBP1-AS2 |
| 4 | 1251547 | 1251747 | -1.155233827 | CTBP1-AS2 |
| 4 | 1728305 | 1728502 | -3.489571223 | TACC3 |
| 4 | 1729355 | 1729554 | -9.181057528 | TACC3 |
| 4 | 1813993 | 1814196 | -1.617551881 | LETM1 |
| 4 | 1900887 | 1901087 | -2.418780115 | WHSC1 |
| 4 | 1978744 | 1978849 | -2.084262312 | WHSC1 |
| 4 | 1978869 | 1979068 | -2.667641337 | WHSC1 |
| 4 | 2240259 | 2240584 | -1.396962942 | POLN |
| 4 | 2240259 | 2240584 | -1.396962942 | HAUS3 |
| 4 | 2240721 | 2240903 | -1.309855436 | HAUS3 |
| 4 | 2240721 | 2240903 | -1.309855436 | POLN |
| 4 | 2744808 | 2744994 | -1.504389957 | TNIP2 |
| 4 | 2936123 | 2936238 | -1.141107833 | NOP14-AS1 |
| 4 | 3316324 | 3316505 | -2.918081442 | RGS12 |
| 1 | 173185841 | 173186022 | 4.493632006 | TNFSF4 |
| 4 | 3374447 | 3374625 | -2.300666387 | RGS12 |
| 4 | 3375029 | 3375194 | -3.361929432 | RGS12 |
| 4 | 3378226 | 3378477 | -1.301234528 | RGS12 |
| 4 | 3942718 | 3942862 | 1.331178938 | FAM86EP |
| 4 | 3942718 | 3942862 | 1.331178938 | AC226119.5 |
| 4 | 3947495 | 3947682 | -1.740095089 | AC226119.5 |
| 4 | 3947495 | 3947682 | -1.740095089 | FAM86EP |
| 1 | 173186477 | 173186678 | 2.940677752 | TNFSF4 |
| 4 | 4550862 | 4550995 | -4.126332716 | STX18-AS1 |
| 4 | 6707912 | 6708114 | -1.01745056 | MRFAP1L1 |
| 4 | 6967177 | 6967359 | -7.525944328 | TBC1D14 |
| 4 | 7041363 | 7041562 | -1.445321095 | RP11-367J11.2 |
| 4 | 7041363 | 7041562 | -1.445321095 | CCDC96 |
| 4 | 7042398 | 7042571 | -2.556559323 | CCDC96 |
| 4 | 7042398 | 7042571 | -2.556559323 | RP11-367J11.2 |
| 4 | 7042398 | 7042571 | -2.556559323 | TADA2B |
| 4 | 7650480 | 7650689 | -1.412036159 | SORCS2 |
| 4 | 30723203 | 30723417 | -1.320537813 | PCDH7 |
| 4 | 36068017 | 36068218 | 4.172790221 | ARAP2 |
| 4 | 37590743 | 37590914 | 3.897207933 | C4orf19 |
| 4 | 37590743 | 37590914 | 3.897207933 | RELL1 |
| 1 | 174437616 | 174437813 | -7.190501055 | RABGAP1L |
| 4 | 38825457 | 38825578 | -7.247427792 | TLR1 |
| 4 | 38825457 | 38825578 | -7.247427792 | TLR6 |
| 4 | 39124855 | 39125083 | -2.059650319 | KLHL5 |
| 4 | 39124855 | 39125083 | -2.059650319 | RP11-360F5.1 |
| 4 | 39125411 | 39125614 | -1.056443293 | KLHL5 |
| 4 | 39125411 | 39125614 | -1.056443293 | RP11-360F5.1 |
| 4 | 39534382 | 39534597 | -1.807661928 | UGDH-AS1 |
| 4 | 39770217 | 39770322 | 6.982892142 | ZBTB12BP |
| 4 | 39770217 | 39770322 | 6.982892142 | UBE2K |
| 4 | 39825225 | 39825424 | -1.575922357 | PDS5A |
| 4 | 39837966 | 39838164 | -1.586418528 | PDS5A |
| 1 | 174561827 | 174562026 | 5.825525846 | RABGAP1L |
| 4 | 40243562 | 40243933 | 2.18877269 | RHOH |
| 4 | 40425800 | 40426002 | -1.341176239 | RBM47 |
| 4 | 42086124 | 42086315 | -1.346035697 | SLC30A9 |
| 4 | 46928399 | 46928574 | -4.183592406 | GABRA4 |
| 1 | 174562111 | 174562293 | -1.27359447 | RABGAP1L |
| 4 | 48421978 | 48422200 | -2.724790722 | SLAIN2 |
| 4 | 48488670 | 48488866 | -4.982892142 | SLC10A4 |
| 4 | 48716183 | 48716336 | -1.903677461 | FRYL |
| 4 | 53096004 | 53096201 | -6.706813965 | SCFD2 |
| 1 | 174729505 | 174729665 | -8.271561235 | RABGAP1L |
| 4 | 53473457 | 53473586 | -2.060442029 | RP11-231C18.3 |
| 4 | 53473457 | 53473586 | -2.060442029 | LNX1 |
| 4 | 55370301 | 55370513 | -1.981843118 | SRD5A3-AS1 |
| 4 | 55370301 | 55370513 | -1.981843118 | SRD5A3 |
| 4 | 55373690 | 55373850 | -3.204621669 | SRD5A3-AS1 |
| 4 | 55386922 | 55387092 | -1.766017071 | SRD5A3-AS1 |
| 1 | 174989759 | 174989944 | -2.252039227 | RABGAP1L |
| 4 | 56345217 | 56345391 | -1.836212671 | AASDH |
| 4 | 56349304 | 56349529 | -1.128662386 | AASDH |
| 4 | 56387594 | 56387750 | -3.469299728 | RP11-646I6.5 |
| 4 | 56910712 | 56910933 | -1.430164641 | REST |
| 4 | 56930085 | 56930320 | -1.711810964 | REST |
| 4 | 56931573 | 56931771 | -1.435894332 | REST |
| 4 | 56977292 | 56977416 | -1.367484488 | NOA1 |
| 4 | 66002189 | 66002390 | -6.149604009 | RP11-25H12.1 |
| 4 | 66023964 | 66024156 | -1.135651011 | RP11-25H12.1 |
| 4 | 66110411 | 66110611 | -2.803544257 | RP11-25H12.1 |
| 4 | 66111116 | 66111252 | -6.726892528 | RP11-25H12.1 |
| 1 | 175159696 | 175159894 | 8.122033232 | KIAA0040 |
| 4 | 67514204 | 67514408 | -1.711449809 | CENPC |
| 4 | 67814614 | 67814813 | -3.485068927 | UBA6-AS1 |
| 4 | 68337225 | 68337423 | -1.957082485 | YTHDC1 |
| 4 | 70688732 | 70688914 | -1.347834642 | UTP3 |
| 1 | 8358288 | 8358482 | 2.639743806 | RERE |
| 1 | 175160519 | 175160718 | 5.477746483 | KIAA0040 |
| 4 | 70799209 | 70799359 | 2.259387454 | RUFY3 |
| 4 | 73091873 | 73092072 | -1.041664958 | ANKRD17 |
| 5 | 123037180 | 123037331 | 5.971257253 | RP11-359P5.1 |
| 5 | 124646987 | 124647088 | -1.139655425 | ZNF608 |
| 5 | 124647826 | 124648021 | 2.097403295 | ZNF608 |
| 5 | 126892594 | 126892780 | -6.558821551 | Mar-03 |
| 5 | 126904187 | 126904350 | -1.734742642 | Mar-03 |
| 5 | 126904717 | 126904838 | -4.972721918 | Mar-03 |
| 5 | 127054715 | 127054912 | -3.721471748 | C5orf63 |
| 5 | 128322295 | 128322456 | -6.074079386 | FBN2 |
| 5 | 131723970 | 131724094 | -6.369890305 | CTC-432M15.3 |
| 5 | 131723970 | 131724094 | -6.369890305 | FNIP1 |
| 5 | 131724732 | 131724931 | -2.506661041 | FNIP1 |
| 5 | 131724732 | 131724931 | -2.506661041 | CTC-432M15.3 |
| 5 | 132294485 | 132294606 | 2.866044132 | P4HA2 |
| 5 | 132294485 | 132294606 | 2.866044132 | SLC22A4 |
| 5 | 132294953 | 132295132 | 3.096805085 | P4HA2 |
| 5 | 132294953 | 132295132 | 3.096805085 | SLC22A4 |
| 5 | 132349302 | 132349442 | 1.267353865 | AC034220.3 |
| 1 | 9270578 | 9270775 | 1.230879664 | H6PD |
| 5 | 134286251 | 134286359 | 3.438675207 | CTD-2410N18.4 |
| 5 | 134286251 | 134286359 | 3.438675207 | CDKL3 |
| 5 | 134741119 | 134741300 | -1.488757709 | CAMLG |
| 5 | 135028486 | 135028688 | 1.171853002 | PITX1 |
| 5 | 135031384 | 135031583 | 9.506379096 | PITX1 |
| 5 | 135033291 | 135033530 | 2.025401302 | PITX1 |
| 5 | 135033291 | 135033530 | 2.025401302 | C5orf66 |
| 5 | 135321643 | 135321849 | 1.01264701 | C5orf66 |
| 5 | 135322280 | 135322471 | -1.063744868 | C5orf66 |
| 5 | 139009296 | 139009434 | 7.38517709 | SIL1 |
| 1 | 203064544 | 203064715 | -1.213712509 | PPFIA4 |
| 5 | 139325341 | 139325523 | 7.044435518 | MATR3 |
| 5 | 139393817 | 139393932 | 1.077797881 | PROB1 |
| 5 | 139394492 | 139394640 | 8.397664723 | PROB1 |
| 5 | 139475649 | 139475835 | -2.852565707 | TMEM173 |
| 5 | 139680816 | 139681015 | 1.372419112 | CXXC5 |
| 5 | 139809524 | 139809695 | -2.019446737 | PSD2 |
| 5 | 140109215 | 140109404 | -5.26080022 | PURA |
| 1 | 203179036 | 203179237 | 12.34355396 | CHI3L1 |
| 5 | 140631938 | 140632129 | -2.239811214 | CD14 |
| 5 | 140967232 | 140967402 | -1.0870147 | PCDHA4 |
| 5 | 140967232 | 140967402 | -1.0870147 | PCDHAC1 |
| 5 | 140967232 | 140967402 | -1.0870147 | PCDHA6 |
| 5 | 140967232 | 140967402 | -1.0870147 | PCDHA8 |
| 5 | 140967232 | 140967402 | -1.0870147 | PCDHA2 |
| 5 | 140967232 | 140967402 | -1.0870147 | PCDHA10 |
| 5 | 140967232 | 140967402 | -1.0870147 | PCDHA13 |
| 5 | 140967232 | 140967402 | -1.0870147 | PCDHA11 |
| 5 | 140967232 | 140967402 | -1.0870147 | PCDHAC2 |
| 5 | 140967232 | 140967402 | -1.0870147 | PCDHA5 |
| 5 | 140967232 | 140967402 | -1.0870147 | PCDHA7 |
| 5 | 140967232 | 140967402 | -1.0870147 | PCDHA1 |
| 5 | 140967232 | 140967402 | -1.0870147 | PCDHA9 |
| 5 | 140967232 | 140967402 | -1.0870147 | PCDHA12 |
| 5 | 140967232 | 140967402 | -1.0870147 | PCDHA3 |
| 5 | 140968426 | 140968679 | -2.033926094 | PCDHA5 |

| | | | | |
|---|---|---|---|---|
| 4 | 7671490 | 7671708 | -1.349768594 | SORCS2 |
| 4 | 7717284 | 7717476 | 6.929087655 | SORCS2 |
| 4 | 7737335 | 7737532 | 3.442383892 | SORCS2 |
| 4 | 7763449 | 7763681 | 2.324482023 | AFAP1-AS1 |
| 4 | 7763449 | 7763681 | 2.324482023 | AFAP1 |
| 4 | 8227334 | 8227534 | 4.550217366 | SH3TC1 |
| 4 | 8227854 | 8228004 | 4.242269952 | SH3TC1 |
| 4 | 8228157 | 8228355 | 5.072491861 | SH3TC1 |
| 4 | 8229790 | 8230024 | 3.035655055 | SH3TC1 |
| 4 | 8232927 | 8233127 | 4.330342115 | SH3TC1 |
| 4 | 8383926 | 8384121 | -3.398530435 | ACOX3 |
| 4 | 8387008 | 8387209 | -1.658264327 | ACOX3 |
| 4 | 8387565 | 8387728 | -3.605467993 | ACOX3 |
| 1 | 173800303 | 173800527 | -2.412766496 | CENPL |
| 4 | 8398926 | 8399056 | 5.049016041 | ACOX3 |
| 4 | 8468081 | 8468282 | -1.03943029 | TRMT44 |
| 4 | 8469357 | 8469553 | -2.554548906 | TRMT44 |
| 4 | 9798631 | 9798814 | -2.501213369 | SLC2A9 |
| 4 | 9825703 | 9825899 | -2.789258596 | SLC2A9 |
| 4 | 9877853 | 9878010 | -8.915703109 | SLC2A9 |
| 4 | 9880136 | 9880341 | -2.807740379 | SLC2A9 |
| 4 | 9982245 | 9982399 | -7.082139219 | SLC2A9 |
| 4 | 9993118 | 9993318 | -7.342237346 | SLC2A9 |
| 1 | 173803078 | 173803366 | -2.03952383 | CENPL |
| 4 | 10019427 | 10019620 | -3.493207858 | SLC2A9 |
| 4 | 13544757 | 13544877 | -6.346743479 | NKX3-2 |
| 1 | 173824898 | 173825117 | -2.143144348 | DARS2 |
| 4 | 14471358 | 14471561 | 1.940657446 | LINC00504 |
| 4 | 14473185 | 14473384 | 2.553282218 | LINC00504 |
| 4 | 14474130 | 14474367 | 1.730563522 | LINC00504 |
| 4 | 14535460 | 14535648 | 6.856639552 | LINC00504 |
| 4 | 14660634 | 14660819 | 1.105159795 | LINC00504 |
| 4 | 14862088 | 14862321 | 1.939934127 | LINC00504 |
| 4 | 15003000 | 15003165 | -7.037154371 | CPEB2 |
| 4 | 15066280 | 15066483 | 3.129905989 | CPEB2 |
| 4 | 15066280 | 15066483 | 3.129905989 | RP11-665G4.1 |
| 1 | 173870764 | 173870986 | -1.271725904 | ZBTB37 |
| 4 | 15614124 | 15614318 | 1.40072716 | FBXL5 |
| 4 | 15631795 | 15632103 | -1.160780225 | FBXL5 |
| 4 | 15820401 | 15820583 | -7.082139219 | CD38 |
| 4 | 15821610 | 15821809 | -4.376429239 | CD38 |
| 4 | 15823030 | 15823307 | -7.309106707 | CD38 |
| 4 | 15846956 | 15847135 | -4.488639962 | CD38 |
| 4 | 16227071 | 16227297 | -1.303815378 | TAPT1 |
| 4 | 16227071 | 16227297 | -1.303815378 | TAPT1-AS1 |
| 4 | 16284356 | 16284555 | 2.827675914 | TAPT1-AS1 |
| 4 | 17646375 | 17646492 | 2.401801394 | FAM184B |
| 4 | 17804305 | 17804499 | -1.358492199 | DCAF16 |
| 4 | 17843712 | 17843906 | -3.293809944 | NCAPG |
| 4 | 17843712 | 17843906 | -3.293809944 | LCORL |
| 4 | 17850358 | 17850555 | -2.791729709 | LCORL |
| 4 | 17860582 | 17860749 | -3.194047792 | LCORL |
| 4 | 17861791 | 17861970 | -2.138322719 | LCORL |
| 1 | 173938682 | 173938862 | 8.607415675 | RC3H1 |
| 4 | 73098209 | 73098408 | -2.385744735 | ANKRD17 |
| 4 | 74961363 | 74961560 | 7.611020079 | PARM1 |
| 4 | 74961363 | 74961560 | 7.611020079 | RP11-44F21.3 |
| 4 | 76034563 | 76034757 | 11.67176881 | ART3 |
| 4 | 76034563 | 76034757 | 11.67176881 | CXCL11 |
| 4 | 77034746 | 77034925 | -6.183161107 | Sep-11 |
| 4 | 78911557 | 78911759 | -1.026245935 | BMP2K |
| 4 | 78911557 | 78911759 | -1.026245935 | PAQR3 |
| 4 | 81427546 | 81427747 | 1.008033218 | RASGEF1B |
| 4 | 82429747 | 82429948 | -1.099339362 | HNRNPDL |
| 4 | 82894320 | 82894521 | -1.814498545 | THAP9-AS1 |
| 4 | 82894320 | 82894521 | -1.814498545 | SEC31A |
| 4 | 83283996 | 83284200 | 2.13787359 | COQ2 |
| 1 | 178411327 | 178411483 | 5.33974005 | RASAL2 |
| 4 | 84809974 | 84810171 | -1.282460116 | WDFY3 |
| 4 | 84809974 | 84810171 | -1.282460116 | WDFY3-AS1 |
| 4 | 85995376 | 85995573 | 4.53303279 | ARHGAP24 |
| 4 | 86701426 | 86701621 | -5.927693809 | PTPN13 |
| 4 | 86913262 | 86913436 | -3.79248125 | RP11-397E7.4 |
| 4 | 86913262 | 86913436 | -3.79248125 | C4orf36 |
| 4 | 86988978 | 86989181 | 1.014338662 | AFF1 |
| 1 | 178442798 | 178443005 | -1.211143103 | RASAL2 |
| 4 | 87089834 | 87090031 | 1.719129363 | AFF1 |
| 4 | 87125982 | 87126181 | -6.70826683 | AFF1 |
| 4 | 87811618 | 87811811 | -1.719349729 | IBSP |
| 4 | 87982532 | 87982747 | 14.27216899 | SPP1 |
| 4 | 89113663 | 89113765 | -5.798094878 | TIGD2 |
| 4 | 89114230 | 89114448 | -2.049023708 | TIGD2 |
| 4 | 89243071 | 89243270 | -1.542762521 | GPRIN3 |
| 4 | 89247988 | 89248182 | -1.288219355 | GPRIN3 |
| 4 | 89249277 | 89249511 | -1.535179794 | GPRIN3 |
| 4 | 94289485 | 94289590 | -2.147652134 | SMARCAD1 |
| 4 | 95546840 | 95547036 | 7.669524838 | UNC5C |
| 4 | 95549021 | 95549220 | 6.428506626 | RP11-710C12.1 |
| 4 | 95549021 | 95549220 | 6.428506626 | UNC5C |
| 4 | 98325345 | 98325657 | -1.29248125 | RAP1GDS1 |
| 4 | 98471883 | 98472057 | -1.54587569 | TSPAN5 |
| 4 | 98893697 | 98893902 | -1.553121468 | EIF4E |
| 1 | 178916457 | 178916656 | 4.441973932 | RALGPS2 |
| 4 | 99868201 | 99868388 | 2.865676648 | DAPP1 |
| 4 | 100187930 | 100188143 | -3.38528556 | DDIT4L |
| 4 | 100187930 | 100188143 | -3.38528556 | RP11-15B17.1 |
| 4 | 102571275 | 102571509 | 2.672558417 | NFKB1 |
| 4 | 102631966 | 102632222 | -1.319884521 | MANBA |
| 4 | 102701166 | 102701339 | -1.32107073 | MANBA |
| 4 | 105473284 | 105473443 | 4.573602462 | PPA2 |
| 4 | 106071758 | 106071876 | -1.686247991 | TBCK |
| 1 | 8361262 | 8361462 | 1.659510587 | RERE |
| 1 | 179107767 | 179107961 | 1.212827949 | ABL2 |
| 4 | 106320699 | 106320903 | 7.851627367 | TBCK |
| 4 | 106320699 | 106320903 | 7.851627367 | AIMP1 |
| 4 | 106923850 | 106924086 | 2.225426266 | DKK2 |
| 4 | 107034419 | 107034699 | 3.390589773 | DKK2 |
| 4 | 107035984 | 107036193 | 2.426967736 | DKK2 |
| 5 | 140968426 | 140968679 | -2.033926094 | PCDHA1 |
| 5 | 140968426 | 140968679 | -2.033926094 | PCDHA9 |
| 5 | 140968426 | 140968679 | -2.033926094 | PCDHA6 |
| 5 | 140968426 | 140968679 | -2.033926094 | PCDHA2 |
| 5 | 140968426 | 140968679 | -2.033926094 | PCDHA13 |
| 5 | 140968426 | 140968679 | -2.033926094 | PCDHA11 |
| 5 | 140968426 | 140968679 | -2.033926094 | PCDHA10 |
| 5 | 140968426 | 140968679 | -2.033926094 | PCDHA8 |
| 5 | 140968426 | 140968679 | -2.033926094 | PCDHAC1 |
| 5 | 140968426 | 140968679 | -2.033926094 | PCDHA7 |
| 5 | 140968426 | 140968679 | -2.033926094 | PCDHA12 |
| 5 | 140968426 | 140968679 | -2.033926094 | PCDHAC2 |
| 5 | 140968426 | 140968679 | -2.033926094 | PCDHA3 |
| 5 | 140968426 | 140968679 | -2.033926094 | PCDHA4 |
| 5 | 140969027 | 140969230 | -3.229715809 | PCDHA8 |
| 5 | 140969027 | 140969230 | -3.229715809 | PCDHA5 |
| 5 | 140969027 | 140969230 | -3.229715809 | PCDHA13 |
| 5 | 140969027 | 140969230 | -3.229715809 | PCDHA10 |
| 5 | 140969027 | 140969230 | -3.229715809 | PCDHA2 |
| 5 | 140969027 | 140969230 | -3.229715809 | PCDHA7 |
| 5 | 140969027 | 140969230 | -3.229715809 | PCDHA4 |
| 5 | 140969027 | 140969230 | -3.229715809 | PCDHA9 |
| 5 | 140969027 | 140969230 | -3.229715809 | PCDHA11 |
| 5 | 140969027 | 140969230 | -3.229715809 | PCDHA3 |
| 5 | 140969027 | 140969230 | -3.229715809 | PCDHAC1 |
| 5 | 140969027 | 140969230 | -3.229715809 | PCDHAC2 |
| 5 | 140969027 | 140969230 | -3.229715809 | PCDHA12 |
| 5 | 140969027 | 140969230 | -3.229715809 | PCDHA6 |
| 5 | 140969027 | 140969230 | -3.229715809 | PCDHA1 |
| 1 | 203305619 | 203305810 | 1.696892933 | BTG2 |
| 5 | 141122150 | 141122346 | -1.195025408 | AC005754.7 |
| 5 | 141122150 | 141122346 | -1.195025408 | PCDHB4 |
| 1 | 203307287 | 203307504 | 1.544922353 | BTG2 |
| 5 | 141157246 | 141157397 | -1.409938268 | CH17-140K24.2 |
| 5 | 141157246 | 141157397 | -1.409938268 | PCDHB17P |
| 5 | 141157246 | 141157397 | -1.409938268 | AC005754.7 |
| 5 | 141187211 | 141187408 | -6.524924275 | PCDHB9 |
| 5 | 141187211 | 141187408 | -6.524924275 | CH17-140K24.2 |
| 5 | 141187211 | 141187408 | -6.524924275 | CH17-140K24.5 |
| 5 | 141187211 | 141187408 | -6.524924275 | CH17-140K24.7 |
| 5 | 141189487 | 141189648 | 1.60771983 | CH17-140K24.2 |
| 5 | 141189487 | 141189648 | 1.60771983 | PCDHB9 |
| 5 | 141189487 | 141189648 | 1.60771983 | CH17-140K24.7 |
| 5 | 141208881 | 141209009 | -1.916445007 | CH17-140K24.2 |
| 5 | 141208881 | 141209009 | -1.916445007 | PCDHB12 |
| 5 | 141210034 | 141210194 | -1.943314887 | PCDHB12 |
| 5 | 141210034 | 141210194 | -1.943314887 | CH17-140K24.2 |
| 5 | 141340734 | 141340933 | 2.122805337 | PCDHGA1 |
| 5 | 141340734 | 141340933 | 2.122805337 | PCDHGA2 |
| 5 | 141414952 | 141415154 | 2.459890204 | PCDHGA3 |
| 5 | 141414952 | 141415154 | 2.459890204 | PCDHGA9 |
| 5 | 141414952 | 141415154 | 2.459890204 | PCDHGA8 |
| 5 | 141414952 | 141415154 | 2.459890204 | PCDHGA7 |
| 5 | 141414952 | 141415154 | 2.459890204 | PCDHGB1 |

| | | | | |
|---|---|---|---|---|
| 4 | 20253590 | 20253729 | -3.988639962 | SLIT2 |
| 4 | 22388048 | 22388250 | -2.215793869 | ADGRA3 |
| 4 | 24808519 | 24808716 | 1.315839817 | CCDC149 |
| 4 | 24909840 | 24910020 | 1.725739748 | CCDC149 |
| 4 | 25201574 | 25201719 | -1.076894791 | RP11-717K11.2 |
| 4 | 25929229 | 25929460 | -1.355265996 | SMIM20 |
| 4 | 26181027 | 26181215 | -2.473708402 | RBPJ |
| 4 | 26320668 | 26320859 | -6.905084969 | RBPJ |
| 4 | 30720986 | 30721185 | 2.789370131 | PCDH7 |
| 4 | 30723203 | 30723417 | 2.914892874 | PCDH7 |
| 4 | 30724029 | 30724226 | 2.605872589 | PCDH7 |
| 4 | 30786921 | 30787121 | 4.434411277 | PCDH7 |
| 4 | 30786921 | 30787121 | 4.434411277 | RP11-619J20.1 |
| 4 | 37590743 | 37590914 | -8.425770829 | RELL1 |
| 4 | 37590743 | 37590914 | -8.425770829 | C4orf19 |
| 4 | 38018194 | 38018397 | -2.444255274 | TBC1D1 |
| 1 | 174437616 | 174437813 | 2.182096911 | RABGAP1L |
| 4 | 38665184 | 38665356 | 7.537154371 | KLF3 |
| 4 | 38697031 | 38697258 | 1.449504844 | KLF3 |
| 4 | 38790552 | 38790747 | 2.708926257 | TLR1 |
| 4 | 38796539 | 38796753 | 4.201399681 | TLR1 |
| 4 | 38798165 | 38798285 | 7.505810189 | TLR1 |
| 4 | 38827181 | 38827374 | 2.680651172 | TLR1 |
| 4 | 38827181 | 38827374 | 2.680651172 | TLR6 |
| 4 | 39124855 | 39125083 | 2.282088227 | KLHL5 |
| 4 | 39124855 | 39125083 | 2.282088227 | RP11-360F5.1 |
| 1 | 174489065 | 174489281 | 6.495051982 | RABGAP1L |
| 4 | 39125411 | 39125614 | 2.075298196 | KLHL5 |
| 4 | 39125411 | 39125614 | 2.075298196 | RP11-360F5.1 |
| 4 | 39308805 | 39308998 | -1.523200801 | RFC1 |
| 4 | 39532693 | 39532894 | -1.529110474 | UGDH-AS1 |
| 4 | 39770217 | 39770322 | -5.767394606 | UBE2K |
| 4 | 39770217 | 39770322 | -5.767394606 | ZBTB12BP |
| 4 | 40120379 | 40120576 | -1.007073916 | N4BP2 |
| 4 | 40425800 | 40426002 | 2.979046615 | RBM47 |
| 4 | 46928399 | 46928574 | 1.506113958 | GABRA4 |
| 4 | 48136168 | 48136382 | -1.844151543 | TEC |
| 1 | 174562111 | 174562293 | 6.9291848 | RABGAP1L |
| 4 | 48488670 | 48488866 | -3.736491603 | SLC10A4 |
| 4 | 48716183 | 48716336 | 5.099836172 | FRYL |
| 4 | 52661158 | 52661346 | -1.56509178 | USP46-AS1 |
| 4 | 53096004 | 53096201 | 2.510655253 | SCFD2 |
| 1 | 174729505 | 174729665 | 2.082955969 | RABGAP1L |
| 4 | 53365202 | 53365414 | -1.342236586 | SCFD2 |
| 4 | 53473457 | 53473586 | 2.760710994 | RP11-231C18.3 |
| 4 | 53473457 | 53473586 | 2.760710994 | LNX1 |
| 4 | 55386922 | 55387092 | 1.588769093 | SRD5A3-AS1 |
| 1 | 174989759 | 174989944 | 4.788747737 | RABGAP1L |
| 4 | 56345217 | 56345391 | -1.232539091 | AASDH |
| 4 | 56387594 | 56387750 | -1.919190909 | RP11-646I6.5 |
| 4 | 56976630 | 56976825 | -2.872555155 | NOA1 |
| 4 | 56977292 | 56977416 | -1.170950768 | NOA1 |
| 4 | 66111116 | 66111252 | 1.22548502 | RP11-25H12.1 |
| 4 | 67514204 | 67514408 | -1.066280909 | CENPC |
| 4 | 107895439 | 107895639 | -2.947647648 | SGMS2 |
| 4 | 107895439 | 107895639 | -2.947647648 | RP11-286E11.1 |
| 1 | 179814382 | 179814563 | -2.029446845 | FAM163A |
| 4 | 107996865 | 107997072 | -7.554119284 | HADH |
| 4 | 109714280 | 109714481 | -1.362018685 | PLA2G12A |
| 4 | 110618093 | 110618282 | 5.27344723 | PITX2 |
| 4 | 112276350 | 112276474 | 1.783765128 | TIFA |
| 4 | 112277990 | 112278226 | 1.974600079 | TIFA |
| 4 | 112357908 | 112358107 | 1.793505165 | ALPK1 |
| 4 | 112357908 | 112358107 | 1.793505165 | RTEL1P1 |
| 4 | 112359713 | 112359818 | 2.209805496 | ALPK1 |
| 4 | 112359713 | 112359818 | 2.209805496 | RTEL1P1 |
| 4 | 113489477 | 113489616 | 1.241851159 | CAMK2D |
| 4 | 118281848 | 118282046 | -1.413051984 | PRSS12 |
| 4 | 118630506 | 118630690 | -1.674550237 | RP11-384K6.6 |
| 4 | 118780966 | 118781100 | 4.649604009 | SEC24D |
| 4 | 119027013 | 119027211 | -2.342160026 | SYNPO2 |
| 4 | 119030075 | 119030292 | -1.666785663 | SYNPO2 |
| 4 | 119057529 | 119057726 | -1.896915179 | SYNPO2 |
| 4 | 119271451 | 119271626 | -2.822661636 | USP53 |
| 4 | 119292534 | 119292721 | -1.614593913 | USP53 |
| 4 | 119482968 | 119483073 | -6.126923742 | RP11-33B1.1 |
| 4 | 122271219 | 122271456 | -1.044504398 | KIAA1109 |
| 4 | 122743738 | 122743950 | -1.368676936 | BBS12 |
| 1 | 179996776 | 179996982 | -1.022983003 | CEP350 |
| 4 | 122892861 | 122893009 | -5.762760405 | NUDT6 |
| 4 | 122892861 | 122893009 | -5.762760405 | FGF2 |
| 4 | 123151096 | 123151249 | -1.68128504 | SPATA5 |
| 4 | 123401696 | 123401895 | -1.482844762 | SPRY1 |
| 4 | 124669172 | 124669407 | -1.203967285 | ANKRD50 |
| 4 | 124710168 | 124710366 | -1.210469633 | ANKRD50 |
| 4 | 125317033 | 125317293 | 5.082453463 | FAT4 |
| 4 | 125321247 | 125321420 | 1.636294784 | FAT4 |
| 4 | 127643686 | 127643886 | -1.815755646 | INTU |
| 4 | 127706657 | 127706852 | -1.297289019 | INTU |
| 4 | 127809355 | 127809549 | -5.114409345 | HSPA4L |
| 4 | 127809355 | 127809549 | -5.114409345 | RP11-50D9.4 |
| 4 | 127832617 | 127832781 | -2.379008197 | HSPA4L |
| 4 | 127885833 | 127886029 | -2.394780637 | PLK4 |
| 4 | 127886327 | 127886429 | -1.058655188 | PLK4 |
| 4 | 127889839 | 127890038 | -2.08942865 | PLK4 |
| 4 | 127920555 | 127920749 | -1.461676343 | MFSD8 |
| 4 | 128271268 | 128271462 | -1.051269566 | PGRMC2 |
| 4 | 128862017 | 128862219 | -1.133613617 | JADE1 |
| 4 | 128872038 | 128872250 | -2.344240249 | SCLT1 |
| 4 | 128872038 | 128872250 | -2.344240249 | JADE1 |
| 4 | 129068559 | 129068839 | 1.140785179 | SCLT1 |
| 4 | 129069685 | 129069883 | 1.084962501 | SCLT1 |
| 4 | 129070817 | 129070946 | 2.297559808 | SCLT1 |
| 4 | 129072934 | 129073156 | 3.119649352 | SCLT1 |
| 4 | 131379926 | 131380106 | -3.008459743 | RP11-314N14.1 |
| 1 | 8656143 | 8656341 | -1.709034414 | RERE |
| 4 | 134200224 | 134200525 | -1.472286229 | PABPC4L |
| 4 | 137591064 | 137591270 | -2.193266435 | RP11-714L20.1 |
| 5 | 141414952 | 141415154 | 2.459890204 | PCDHGA4 |
| 5 | 141414952 | 141415154 | 2.459890204 | PCDHGB4 |
| 5 | 141414952 | 141415154 | 2.459890204 | PCDHGB6 |
| 5 | 141414952 | 141415154 | 2.459890204 | PCDHGA10 |
| 5 | 141414952 | 141415154 | 2.459890204 | PCDHGA6 |
| 5 | 141414952 | 141415154 | 2.459890204 | PCDHGA1 |
| 5 | 141414952 | 141415154 | 2.459890204 | PCDHGB2 |
| 5 | 141414952 | 141415154 | 2.459890204 | PCDHGB5 |
| 5 | 141414952 | 141415154 | 2.459890204 | PCDHGA2 |
| 5 | 141414952 | 141415154 | 2.459890204 | PCDHGB3 |
| 5 | 141414952 | 141415154 | 2.459890204 | PCDHGA5 |
| 5 | 141485539 | 141485736 | 10.43533362 | PCDHGB4 |
| 5 | 141485539 | 141485736 | 10.43533362 | PCDHGA2 |
| 5 | 141485539 | 141485736 | 10.43533362 | PCDHGA7 |
| 5 | 141485539 | 141485736 | 10.43533362 | PCDHGA8 |
| 5 | 141485539 | 141485736 | 10.43533362 | PCDHGA5 |
| 5 | 141485539 | 141485736 | 10.43533362 | PCDHGB3 |
| 5 | 141485539 | 141485736 | 10.43533362 | PCDHGB5 |
| 5 | 141485539 | 141485736 | 10.43533362 | PCDHGA1 |
| 5 | 141485539 | 141485736 | 10.43533362 | PCDHGB7 |
| 5 | 141485539 | 141485736 | 10.43533362 | PCDHGA3 |
| 5 | 141485539 | 141485736 | 10.43533362 | PCDHGB1 |
| 5 | 141485539 | 141485736 | 10.43533362 | PCDHGA10 |
| 5 | 141485539 | 141485736 | 10.43533362 | PCDHGA4 |
| 5 | 141485539 | 141485736 | 10.43533362 | PCDHGA9 |
| 5 | 141485539 | 141485736 | 10.43533362 | PCDHGA6 |
| 5 | 141485539 | 141485736 | 10.43533362 | PCDHGC3 |
| 5 | 141485539 | 141485736 | 10.43533362 | PCDHGB6 |
| 5 | 141485539 | 141485736 | 10.43533362 | PCDHGA11 |
| 5 | 141485539 | 141485736 | 10.43533362 | PCDHGA12 |
| 5 | 141485539 | 141485736 | 10.43533362 | PCDHGB2 |
| 5 | 141485539 | 141485736 | 10.43533362 | PCDHGC4 |
| 5 | 141488883 | 141489085 | 10.46862799 | PCDHGA2 |
| 5 | 141488883 | 141489085 | 10.46862799 | PCDHGA7 |
| 5 | 141488883 | 141489085 | 10.46862799 | PCDHGA6 |
| 5 | 141488883 | 141489085 | 10.46862799 | PCDHGA8 |
| 5 | 141488883 | 141489085 | 10.46862799 | PCDHGB1 |
| 5 | 141488883 | 141489085 | 10.46862799 | PCDHGC3 |
| 5 | 141488883 | 141489085 | 10.46862799 | PCDHGB3 |
| 5 | 141488883 | 141489085 | 10.46862799 | PCDHGA9 |
| 5 | 141488883 | 141489085 | 10.46862799 | PCDHGA3 |
| 5 | 141488883 | 141489085 | 10.46862799 | PCDHGB7 |
| 5 | 141488883 | 141489085 | 10.46862799 | PCDHGA1 |
| 5 | 141488883 | 141489085 | 10.46862799 | PCDHGA10 |
| 5 | 141488883 | 141489085 | 10.46862799 | PCDHGA11 |
| 5 | 141488883 | 141489085 | 10.46862799 | PCDHGA4 |
| 5 | 141488883 | 141489085 | 10.46862799 | PCDHGC4 |
| 5 | 141488883 | 141489085 | 10.46862799 | PCDHGA12 |
| 5 | 141488883 | 141489085 | 10.46862799 | PCDHGB5 |
| 5 | 141488883 | 141489085 | 10.46862799 | PCDHGB6 |
| 5 | 141488883 | 141489085 | 10.46862799 | PCDHGB2 |
| 5 | 141488883 | 141489085 | 10.46862799 | PCDHGB4 |
| 5 | 141488883 | 141489085 | 10.46862799 | PCDHGA5 |
| 5 | 141490123 | 141490274 | 10.39650315 | PCDHGA9 |

| | | | | |
|---|---|---|---|---|
| 4 | 67615172 | 67615349 | -6.81785906 | UBA6 |
| 4 | 67720530 | 67720832 | -1.639953988 | UBA6-AS1 |
| 4 | 67720530 | 67720832 | -1.639953988 | RP11-453E17.3 |
| 4 | 70688732 | 70688914 | -1.752364462 | UTP3 |
| 4 | 70799209 | 70799359 | -1.427805045 | RUFY3 |
| 4 | 73098209 | 73098408 | -1.627698598 | ANKRD17 |
| 4 | 75645157 | 75645357 | -1.173788608 | G3BP2 |
| 4 | 75962834 | 75963091 | -1.696158711 | SDAD1 |
| 4 | 77034746 | 77034925 | -2.262137287 | Sep-11 |
| 4 | 77890399 | 77890558 | -5.864172857 | MRPL1 |
| 1 | 176125239 | 176125478 | 5.385744735 | RFWD2 |
| 4 | 79905858 | 79906060 | 1.921279102 | ANTXR2 |
| 4 | 79907069 | 79907271 | 1.605438615 | ANTXR2 |
| 4 | 81427546 | 81427747 | 11.89018297 | RASGEF1B |
| 4 | 82429747 | 82429948 | -1.58063007 | HNRNPDL |
| 4 | 82894320 | 82894521 | -1.066272274 | THAP9-AS1 |
| 4 | 82894320 | 82894521 | -1.066272274 | SEC31A |
| 1 | 178134585 | 178134786 | 3.163396925 | RASAL2 |
| 4 | 82927996 | 82928196 | -1.387327584 | LIN54 |
| 4 | 82984369 | 82984612 | -2.126080074 | LIN54 |
| 4 | 83246491 | 83246689 | -1.520313645 | RP11-576N17.5 |
| 4 | 83283416 | 83283623 | -3.212086734 | COQ2 |
| 4 | 83283996 | 83284200 | -4.697818168 | COQ2 |
| 4 | 83453671 | 83453872 | -1.140093764 | HELQ |
| 4 | 84672702 | 84672832 | 1.071104416 | WDFY3 |
| 4 | 85995376 | 85995573 | -1.221617196 | ARHGAP24 |
| 4 | 86913262 | 86913436 | -3.462282553 | RP11-397E7.4 |
| 4 | 86913262 | 86913436 | -3.462282553 | C4orf36 |
| 4 | 86988978 | 86989181 | -2.308437527 | AFF1 |
| 4 | 86989737 | 86989872 | -2.00311231 | AFF1 |
| 4 | 87047273 | 87047482 | -2.056996311 | AFF1 |
| 4 | 87083943 | 87084148 | -2.440475802 | AFF1 |
| 1 | 178442798 | 178443005 | 1.7762074 | RASAL2 |
| 4 | 87089834 | 87090031 | -4.362207733 | AFF1 |
| 4 | 87114943 | 87115144 | -1.758185222 | AFF1 |
| 4 | 87124879 | 87125078 | -1.356640903 | AFF1 |
| 4 | 87125982 | 87126181 | -1.64643665 | AFF1 |
| 4 | 87135722 | 87135948 | -1.784697433 | AFF1 |
| 4 | 87161165 | 87161313 | -1.4100044 | KLHL8 |
| 4 | 87185417 | 87185671 | -1.37970858 | KLHL8 |
| 4 | 87662198 | 87662391 | 7.443086436 | RP11-742B18.1 |
| 4 | 87662198 | 87662391 | 7.443086436 | DMP1 |
| 4 | 88262574 | 88262736 | 1.662786748 | PPM1K |
| 4 | 88521581 | 88521739 | -1.851587264 | PYURF |
| 4 | 88521581 | 88521739 | -1.851587264 | HERC3 |
| 4 | 88521581 | 88521739 | -1.851587264 | PIGY |
| 4 | 88696701 | 88696909 | -2.443598586 | HERC3 |
| 4 | 88696701 | 88696909 | -2.443598586 | NAP1L5 |
| 4 | 89113663 | 89113765 | -2.044881764 | TIGD2 |
| 4 | 89243071 | 89243270 | 11.10085961 | GPRIN3 |
| 4 | 89247988 | 89248182 | 9.51921929 | GPRIN3 |
| 4 | 89249277 | 89249511 | 9.07546366 | GPRIN3 |
| 1 | 178473217 | 178473408 | 2.007827759 | RASAL2 |
| 4 | 95169129 | 95169342 | -3.400780041 | UNC5C |
| 4 | 137594434 | 137594669 | -1.750820924 | RP11-714L20.1 |
| 4 | 137595281 | 137595453 | -3.322275327 | RP11-714L20.1 |
| 4 | 139044868 | 139045069 | 2.003265914 | NOCT |
| 4 | 139044868 | 139045069 | 2.003265914 | ELF2 |
| 4 | 139115759 | 139115942 | -1.991813515 | ELF2 |
| 4 | 139300711 | 139300939 | 1.521460096 | NDUFC1 |
| 4 | 139511541 | 139511742 | -1.231654172 | SETD7 |
| 4 | 139718719 | 139719021 | -4.263561146 | MAML3 |
| 4 | 139718719 | 139719021 | -4.263561146 | MGST2 |
| 4 | 139719701 | 139719860 | -4.765039651 | MGST2 |
| 4 | 139719701 | 139719860 | -4.765039651 | MAML3 |
| 4 | 139889616 | 139889756 | -3.656945388 | MAML3 |
| 4 | 139890605 | 139890805 | -1.71393077 | MAML3 |
| 4 | 140153821 | 140154021 | -2.805644489 | MAML3 |
| 4 | 140622268 | 140622470 | 1.512840454 | TBC1D9 |
| 4 | 145033130 | 145033297 | -4.953445298 | ANAPC10 |
| 4 | 145557836 | 145558026 | -6.884453986 | SMAD1 |
| 4 | 146323173 | 146323328 | 1.794539305 | SLC10A7 |
| 4 | 146640842 | 146641043 | 2.906106194 | POU4F2 |
| 4 | 147542579 | 147542721 | -6.447408882 | EDNRA |
| 4 | 150256719 | 150256923 | -2.528467065 | DCLK2 |
| 4 | 150837531 | 150837767 | -1.731915299 | LRBA |
| 4 | 152346995 | 152347188 | -6.476643534 | FBXW7 |
| 4 | 152350079 | 152350261 | -1.599104953 | FBXW7 |
| 1 | 180950250 | 180950450 | -7.808059019 | KIAA1614 |
| 1 | 180950250 | 180950450 | -7.808059019 | RP11-46A10.5 |
| 1 | 180950250 | 180950450 | -7.808059019 | KIAA1614-AS1 |
| 4 | 152626961 | 152627162 | -2.558486662 | TMEM154 |
| 4 | 152627920 | 152628149 | -1.588176238 | TMEM154 |
| 4 | 153295555 | 153295756 | -2.065661138 | TRIM2 |
| 4 | 153334931 | 153335131 | -3.538426341 | TRIM2 |
| 4 | 153704986 | 153705198 | 1.030322823 | TLR2 |
| 4 | 155710962 | 155711147 | 2.115502776 | GUCY1A3 |
| 4 | 156762829 | 156763012 | -3.321116465 | PDGFC |
| 1 | 180976338 | 180976537 | -1.210025194 | RP11-46A10.5 |
| 1 | 180976338 | 180976537 | -1.210025194 | STX6 |
| 4 | 156971090 | 156971319 | -2.751689446 | PDGFC |
| 4 | 158170788 | 158170977 | -2.330787148 | FAM198B |
| 4 | 158170788 | 158170977 | -2.330787148 | RP11-597D13.9 |
| 4 | 158544200 | 158544393 | 3.228620131 | RXFP1 |
| 4 | 158556013 | 158556336 | 2.59742268 | RXFP1 |
| 4 | 158568163 | 158568360 | 4.127605924 | RXFP1 |
| 4 | 158651929 | 158652129 | 2.549840124 | RXFP1 |
| 4 | 158847115 | 158847306 | 7.601521028 | FNIP2 |
| 4 | 158881254 | 158881454 | -3.291903016 | FNIP2 |
| 4 | 159358052 | 159358250 | -7.628251784 | RAPGEF2 |
| 4 | 163325376 | 163325501 | -4.249087295 | NPY1R |
| 4 | 165341763 | 165341979 | -1.609377418 | MSMO1 |
| 4 | 168504488 | 168504669 | 2.104148796 | DDX60L |
| 4 | 168504488 | 168504669 | 2.104148796 | PALLD |
| 4 | 168947511 | 168947705 | -2.309285447 | CBR4 |
| 4 | 169096422 | 169096640 | -1.036778586 | SH3RF1 |
| 1 | 181089798 | 181089997 | 2.267603052 | IER5 |
| 4 | 169719909 | 169720111 | -1.746881343 | CLCN3 |
| 5 | 141490123 | 141490274 | 10.39650315 | PCDHGA12 |
| 5 | 141490123 | 141490274 | 10.39650315 | PCDHGA10 |
| 5 | 141490123 | 141490274 | 10.39650315 | PCDHGA2 |
| 5 | 141490123 | 141490274 | 10.39650315 | PCDHGA4 |
| 5 | 141490123 | 141490274 | 10.39650315 | PCDHGB6 |
| 5 | 141490123 | 141490274 | 10.39650315 | PCDHGA6 |
| 5 | 141490123 | 141490274 | 10.39650315 | PCDHGB2 |
| 5 | 141490123 | 141490274 | 10.39650315 | PCDHGB4 |
| 5 | 141490123 | 141490274 | 10.39650315 | PCDHGA11 |
| 5 | 141490123 | 141490274 | 10.39650315 | PCDHGB3 |
| 5 | 141490123 | 141490274 | 10.39650315 | PCDHGA7 |
| 5 | 141490123 | 141490274 | 10.39650315 | PCDHGB1 |
| 5 | 141490123 | 141490274 | 10.39650315 | PCDHGC4 |
| 5 | 141490123 | 141490274 | 10.39650315 | PCDHGB7 |
| 5 | 141490123 | 141490274 | 10.39650315 | PCDHGA3 |
| 5 | 141490123 | 141490274 | 10.39650315 | PCDHGB5 |
| 5 | 141490123 | 141490274 | 10.39650315 | PCDHGA5 |
| 5 | 141490123 | 141490274 | 10.39650315 | PCDHGA8 |
| 5 | 141490123 | 141490274 | 10.39650315 | PCDHGA1 |
| 5 | 141490123 | 141490274 | 10.39650315 | PCDHGC5 |
| 5 | 141490123 | 141490274 | 10.39650315 | PCDHGC3 |
| 5 | 141490719 | 141490905 | 10.64678073 | PCDHGA1 |
| 5 | 141490719 | 141490905 | 10.64678073 | PCDHGC5 |
| 5 | 141490719 | 141490905 | 10.64678073 | PCDHGA2 |
| 5 | 141490719 | 141490905 | 10.64678073 | PCDHGA12 |
| 5 | 141490719 | 141490905 | 10.64678073 | PCDHGA9 |
| 5 | 141490719 | 141490905 | 10.64678073 | PCDHGA8 |
| 5 | 141490719 | 141490905 | 10.64678073 | PCDHGA3 |
| 5 | 141490719 | 141490905 | 10.64678073 | PCDHGA11 |
| 5 | 141490719 | 141490905 | 10.64678073 | PCDHGA7 |
| 5 | 141490719 | 141490905 | 10.64678073 | PCDHGA5 |
| 5 | 141490719 | 141490905 | 10.64678073 | PCDHGB3 |
| 5 | 141490719 | 141490905 | 10.64678073 | PCDHGB1 |
| 5 | 141490719 | 141490905 | 10.64678073 | PCDHGC3 |
| 5 | 141490719 | 141490905 | 10.64678073 | PCDHGA4 |
| 5 | 141490719 | 141490905 | 10.64678073 | PCDHGB2 |
| 5 | 141490719 | 141490905 | 10.64678073 | PCDHGB5 |
| 5 | 141490719 | 141490905 | 10.64678073 | PCDHGA10 |
| 5 | 141490719 | 141490905 | 10.64678073 | PCDHGC4 |
| 5 | 141490719 | 141490905 | 10.64678073 | PCDHGB7 |
| 5 | 141490719 | 141490905 | 10.64678073 | PCDHGB6 |
| 5 | 141490719 | 141490905 | 10.64678073 | PCDHGA6 |
| 5 | 141490719 | 141490905 | 10.64678073 | PCDHGB4 |
| 5 | 142995001 | 142995149 | -3.974183616 | ARHGAP26 |
| 5 | 143203663 | 143203829 | 7.219265989 | ARHGAP26 |
| 5 | 143405674 | 143405872 | 1.083500803 | NR3C1 |
| 5 | 148827443 | 148827629 | -1.571120635 | ADRB2 |
| 5 | 148827892 | 148828092 | -2.240375599 | ADRB2 |
| 5 | 149499124 | 149499312 | 8.230979493 | CSNK1A1 |
| 5 | 149499124 | 149499312 | 8.230979493 | CTB-89H12.4 |
| 5 | 149501291 | 149501487 | 8.26862437 | CSNK1A1 |
| 5 | 149501291 | 149501487 | 8.26862437 | CTB-89H12.4 |
| 5 | 149833261 | 149833456 | 1.010444335 | PPARGC1B |
| 5 | 149981325 | 149981599 | 1.054032916 | SLC26A2 |

| | | | | |
|---|---|---|---|---|
| 4 | 95549021 | 95549220 | -4.218876036 | RP11-710C12.1 |
| 4 | 95549021 | 95549220 | -4.218876036 | UNC5C |
| 4 | 95549838 | 95550037 | -2.143464037 | RP11-710C12.1 |
| 4 | 98471883 | 98472057 | -1.407930079 | TSPAN5 |
| 1 | 178916457 | 178916656 | -1.16171506 | RALGPS2 |
| 4 | 99868201 | 99868388 | -1.020400978 | DAPP1 |
| 4 | 102570759 | 102570950 | -5.215226276 | NFKB1 |
| 4 | 102697771 | 102697970 | 1.377719669 | MANBA |
| 4 | 102701166 | 102701339 | 1.241008163 | MANBA |
| 1 | 179095179 | 179095377 | -1.432329861 | TOR3A |
| 4 | 102839714 | 102839877 | -1.242202001 | RP11-10L12.4 |
| 4 | 102839714 | 102839877 | -1.242202001 | UBE2D3 |
| 4 | 105473284 | 105473443 | -6.986579607 | PPA2 |
| 1 | 179107767 | 179107961 | 1.041864244 | ABL2 |
| 4 | 106163151 | 106163438 | -1.029887228 | TBCK |
| 4 | 106316191 | 106316340 | -6.073602462 | AIMP1 |
| 4 | 106316191 | 106316340 | -6.073602462 | TBCK |
| 4 | 106320699 | 106320903 | -8.531117955 | AIMP1 |
| 4 | 106320699 | 106320903 | -8.531117955 | TBCK |
| 4 | 107034419 | 107034699 | 2.650945171 | DKK2 |
| 4 | 107035984 | 107036193 | 1.930026173 | DKK2 |
| 4 | 107895439 | 107895639 | 1.774001202 | SGMS2 |
| 4 | 107895439 | 107895639 | 1.774001202 | RP11-286E11.1 |
| 4 | 107909610 | 107909797 | -7.1386437 | SGMS2 |
| 4 | 107909610 | 107909797 | -7.1386437 | RP11-286E11.1 |
| 4 | 107950451 | 107950637 | -7.248551679 | CYP2U1 |
| 4 | 107950451 | 107950637 | -7.248551679 | RP11-286E11.1 |
| 1 | 179814382 | 179814563 | -3.601437388 | FAM163A |
| 4 | 107996845 | 107997072 | -1.779521396 | HADH |
| 4 | 108048546 | 108048674 | 1.716675403 | LEF1 |
| 4 | 110618093 | 110618282 | -6.999647694 | PITX2 |
| 4 | 112276350 | 112276474 | -4.458361833 | TIFA |
| 4 | 112277990 | 112278226 | -1.886329483 | TIFA |
| 4 | 112357908 | 112358107 | -1.745926548 | ALPK1 |
| 4 | 112357908 | 112358107 | -1.745926548 | RTEL1P1 |
| 4 | 112359713 | 112359818 | -2.444908541 | ALPK1 |
| 4 | 112359713 | 112359818 | -2.444908541 | RTEL1P1 |
| 4 | 112587440 | 112587609 | -3.287328645 | ZGRF1 |
| 4 | 112618099 | 112618343 | -2.980942934 | ZGRF1 |
| 4 | 114623181 | 114623383 | -1.007307037 | UGT8 |
| 4 | 118281848 | 118282046 | -3.354244008 | PRSS12 |
| 4 | 118636472 | 118636597 | -2.350767233 | CICP16 |
| 4 | 118778034 | 118778234 | 7.013175265 | SEC24D |
| 4 | 118780966 | 118781100 | -5.533044595 | SEC24D |
| 4 | 118782389 | 118782502 | -7.805079876 | SEC24D |
| 4 | 119027013 | 119027211 | -3.922674544 | SYNPO2 |
| 4 | 119030075 | 119030292 | -3.008579691 | SYNPO2 |
| 4 | 119033047 | 119033200 | -4.74934623 | SYNPO2 |
| 4 | 119057529 | 119057726 | -3.616209299 | SYNPO2 |
| 4 | 119271451 | 119271626 | 1.051435534 | USP53 |
| 4 | 119292534 | 119292721 | 1.510624968 | USP53 |
| 4 | 119486543 | 119486683 | -1.835854872 | RP11-33B1.1 |
| 1 | 179996776 | 179996982 | -1.027040133 | CEP350 |
| 4 | 122892191 | 122892316 | 1.761373552 | NUDT6 |
| 4 | 173321639 | 173321836 | -1.183215346 | GALNT7 |
| 4 | 174283549 | 174283734 | -5.218355771 | FBXO8 |
| 4 | 174283549 | 174283734 | -5.218355771 | CEP44 |
| 4 | 174307002 | 174307196 | -1.345717452 | CEP44 |
| 4 | 176683717 | 176683971 | 1.976536344 | VEGFC |
| 4 | 176683717 | 176683971 | 1.976536344 | RP11-313E19.2 |
| 4 | 176687289 | 176687486 | 1.657917502 | RP11-313E19.2 |
| 4 | 176687289 | 176687486 | 1.657917502 | VEGFC |
| 4 | 177599043 | 177599242 | -1.030083047 | RP11-130F10.1 |
| 4 | 182800328 | 182800527 | 1.804500756 | TENM3 |
| 1 | 181798709 | 181799009 | -1.439588194 | CACNA1E |
| 4 | 183097573 | 183097770 | -6.654311723 | WWC2-AS2 |
| 4 | 183510665 | 183510865 | -1.312935741 | ING2 |
| 4 | 184011105 | 184011296 | -1.812245432 | STOX2 |
| 4 | 184388426 | 184388671 | 1.351192865 | IRF2 |
| 1 | 182585904 | 182586131 | -1.520055107 | RNASEL |
| 4 | 184672675 | 184672825 | -1.127079734 | PRIMPOL |
| 4 | 186210424 | 186210587 | -7.754763804 | CYP4V2 |
| 4 | 186210424 | 186210587 | -7.754763804 | KLKB1 |
| 1 | 182586559 | 182586747 | -1.486181931 | RNASEL |
| 4 | 186600049 | 186600242 | -1.365894571 | FAT1 |
| 4 | 186621317 | 186621518 | -1.033055327 | FAT1 |
| 1 | 182599061 | 182599262 | 1.053244896 | RGS16 |
| 4 | 188632139 | 188632239 | -2.453445298 | LINC01060 |
| 5 | 199052 | 199250 | -5.328473095 | CCDC127 |
| 5 | 346452 | 346555 | -2.43253521 | PDCD6 |
| 5 | 346452 | 346555 | -2.43253521 | AHRR |
| 5 | 388720 | 388926 | 8.137916755 | AHRR |
| 1 | 182616020 | 182616192 | 1.427539612 | RP11-317P15.4 |
| 5 | 389622 | 389827 | 1.130367783 | AHRR |
| 5 | 401986 | 402190 | 8.161395389 | AHRR |
| 5 | 430699 | 430908 | -1.509939497 | AHRR |
| 5 | 431837 | 431959 | -1.489072108 | AHRR |
| 5 | 432669 | 432839 | -1.643911256 | AHRR |
| 5 | 435531 | 435736 | -1.049133602 | AHRR |
| 5 | 436842 | 437042 | -1.013131264 | AHRR |
| 5 | 437627 | 437828 | -1.230482625 | AHRR |
| 5 | 442693 | 442836 | -8.292018813 | EXOC3-AS1 |
| 5 | 453526 | 453733 | -1.580567349 | EXOC3 |
| 5 | 471476 | 471675 | -1.78494592 | EXOC3 |
| 5 | 471476 | 471675 | -1.78494592 | PP7080 |
| 5 | 473983 | 474187 | -1.151833061 | CTD-2228K2.7 |
| 5 | 473983 | 474187 | -1.151833061 | SLC9A3 |
| 5 | 477744 | 477925 | -1.008472833 | CTD-2228K2.7 |
| 5 | 477744 | 477925 | -1.008472833 | SLC9A3 |
| 5 | 640793 | 641005 | -1.13572277 | CEP72 |
| 5 | 747976 | 748170 | -2.003213135 | ZDHHC11B |
| 1 | 9103981 | 9104179 | 3.460548544 | GPR157 |
| 1 | 183926591 | 183926874 | 5.685149355 | RGL1 |
| 5 | 840100 | 840284 | 6.614409345 | ZDHHC11 |
| 5 | 990181 | 990369 | -2.792909986 | RP11-43F13.3 |
| 5 | 1017611 | 1017794 | -7.448308111 | NKD2 |
| 1 | 184431455 | 184431714 | 1.324815187 | C1orf21 |
| 5 | 1051371 | 1051568 | 7.267098238 | SLC12A7 |
| 5 | 150053572 | 150053769 | 11.03809709 | CSF1R |
| 5 | 150650449 | 150650648 | 7.333001016 | SYNPO |
| 5 | 169823532 | 169823728 | -4.482892142 | DOCK2 |
| 5 | 169825281 | 169825430 | -4.857122759 | DOCK2 |
| 5 | 170233173 | 170233331 | -3.668396722 | C5orf58 |
| 5 | 170240060 | 170240267 | -1.286141683 | C5orf58 |
| 5 | 170242721 | 170242921 | -3.913226094 | C5orf58 |
| 5 | 170243510 | 170243747 | -1.139770752 | C5orf58 |
| 1 | 204222076 | 204222296 | 2.608813649 | PLEKHA6 |
| 5 | 170271692 | 170271873 | -7.890436463 | LCP2 |
| 5 | 172043699 | 172043897 | -1.113998769 | STK10 |
| 5 | 172254415 | 172254588 | -1.432631291 | UBTD2 |
| 5 | 172254415 | 172254588 | -1.432631291 | KLF3P1 |
| 5 | 172770910 | 172771109 | 12.96914788 | DUSP1 |
| 5 | 172770910 | 172771109 | 12.96914788 | RP11-779O18.3 |
| 5 | 173317817 | 173318023 | -1.083581234 | STC2 |
| 5 | 175682975 | 175683182 | 1.358224403 | HRH2 |
| 5 | 176598555 | 176598749 | 1.078727478 | GPRIN1 |
| 5 | 177135601 | 177135812 | 1.228081747 | NSD1 |
| 5 | 177477947 | 177478142 | 1.145801779 | RP11-1334A24.5 |
| 5 | 178121604 | 178121801 | 2.302452268 | N4BP3 |
| 5 | 178366059 | 178366253 | -4.398830763 | COL23A1 |
| 5 | 178394741 | 178394889 | 5.038407799 | COL23A1 |
| 5 | 178508041 | 178508169 | -1.019678331 | COL23A1 |
| 5 | 178514091 | 178514285 | -5.93633744 | COL23A1 |
| 5 | 178883173 | 178883346 | -1.941761591 | ZNF354B |
| 5 | 179079527 | 179079727 | -1.20314897 | ZNF354C |
| 5 | 179111633 | 179111836 | -1.294339473 | ADAMTS2 |
| 5 | 179575335 | 179575535 | -3.127847198 | RUFY1 |
| 5 | 179606934 | 179607144 | -2.341447144 | RUFY1 |
| 5 | 179768751 | 179768953 | -1.992297654 | MAML1 |
| 5 | 179774683 | 179774882 | 1.674739766 | MAML1 |
| 5 | 179883848 | 179884051 | 2.454624452 | TBC1D9B |
| 5 | 180792000 | 180792200 | 1.633777659 | MGAT1 |
| 1 | 205211527 | 205211699 | 4.99858974 | DSTYK |
| 6 | 326876 | 327013 | 6.968226955 | DUSP22 |
| 6 | 1394811 | 1394992 | 1.51775959 | FOXF2 |
| 6 | 1610739 | 1610924 | 1.988709352 | FOXC1 |
| 6 | 1612029 | 1612223 | 1.548941353 | FOXC1 |
| 6 | 2291487 | 2291691 | -3.243092173 | GMDS-AS1 |
| 6 | 2292258 | 2292454 | 1.069668341 | GMDS-AS1 |
| 1 | 205616444 | 205616643 | -1.023343904 | ELK4 |
| 6 | 2841184 | 2841383 | 6.269579406 | SERPINB1 |
| 6 | 3003754 | 3003939 | -6.118903737 | NQO2 |
| 6 | 3105638 | 3105854 | -1.284661144 | RIPK1 |
| 6 | 3722875 | 3723063 | 4.380775616 | PXDC1 |
| 6 | 5045259 | 5045372 | 3.126143485 | RP11-428J1.4 |
| 6 | 5086463 | 5086635 | 3.111620996 | PPP1R3G |
| 6 | 5998339 | 5998549 | 1.882890317 | NRN1 |
| 6 | 6685464 | 6685664 | 2.785231466 | RP1-80N2.4 |
| 6 | 6700348 | 6700548 | -4.067238083 | RP1-80N2.2 |
| 6 | 6795364 | 6795556 | -3.488639962 | BTF3P7 |
| 6 | 7246517 | 7246738 | 2.275108453 | RREB1 |
| 6 | 7726649 | 7726820 | 6.136256576 | BMP6 |

| | | | | | | | | | | | | | | |
|---|---|---|---|---|---|---|---|---|---|---|---|---|---|---|
| 4 | 122892191 | 122892316 | 1.761373552 | FGF2 | 5 | 1051710 | 1051922 | 2.250705962 | SLC12A7 | 1 | 205769463 | 205769672 | 1.098359754 | RAB29 |
| 4 | 122892861 | 122893009 | -1.875301798 | NUDT6 | 5 | 1052183 | 1052382 | 6.206053346 | SLC12A7 | 6 | 11351786 | 11351987 | -1.945406569 | NEDD9 |
| 4 | 122892861 | 122893009 | -1.875301798 | FGF2 | 5 | 1060706 | 1060898 | 2.103225439 | SLC12A7 | 6 | 12120248 | 12120469 | -1.129833324 | HIVEP1 |
| 4 | 122934230 | 122934441 | -1.993453153 | SPATA5 | 1 | 184432238 | 184432406 | 3.122805337 | C1orf21 | 6 | 12121148 | 12121343 | -1.639716689 | HIVEP1 |
| 4 | 123151096 | 123151249 | -1.871182791 | SPATA5 | 5 | 1475730 | 1475934 | 1.006300018 | LPCAT1 | 6 | 12122555 | 12122722 | -2.117189108 | HIVEP1 |
| 1 | 180020676 | 180020871 | -2.704329441 | CEP350 | 5 | 1491188 | 1491383 | 4.740899716 | LPCAT1 | 6 | 12125518 | 12125642 | -1.069541301 | HIVEP1 |
| 4 | 125317033 | 125317293 | -6.827541723 | FAT4 | 5 | 1798608 | 1798799 | -1.314644937 | MRPL36 | 6 | 12142364 | 12142471 | -1.161844276 | HIVEP1 |
| 4 | 125321247 | 125321420 | -3.050738826 | FAT4 | 5 | 1806718 | 1806920 | -7.870707158 | NDUFS6 | 6 | 12161580 | 12161809 | -1.72751061 | HIVEP1 |
| 4 | 127643686 | 127643886 | -1.344405235 | INTU | 5 | 5462757 | 5462965 | 1.200134888 | ICE1 | 1 | 205810790 | 205810988 | 1.29038059 | SLC41A1 |
| 4 | 127706657 | 127706852 | -1.975348163 | INTU | 5 | 5479596 | 5479769 | 5.945891852 | ICE1 | 6 | 13384993 | 13385173 | 1.878365024 | GFOD1 |
| 4 | 127809355 | 127809549 | -3.006284337 | HSPA4L | 5 | 6665102 | 6665214 | 3.007939374 | SRD5A1 | 6 | 13486734 | 13486933 | 2.228726985 | GFOD1 |
| 4 | 127809355 | 127809549 | -3.006284337 | RP11-50D9.4 | 5 | 6754765 | 6754981 | 1.820492644 | PAPD7 | 6 | 13486734 | 13486933 | 2.228726985 | GFOD1-AS1 |
| 1 | 180093769 | 180093905 | -1.356467275 | CEP350 | 5 | 10227367 | 10227543 | -1.623024853 | FAM173B | 1 | 205979360 | 205979471 | 7.20436389 | RAB7B |
| 4 | 127832617 | 127832781 | -3.247389759 | HSPA4L | 5 | 10649505 | 10649694 | 10.87065099 | ANKRD33B | 6 | 15520707 | 15520899 | -1.11850012 | JARID2 |
| 4 | 127885833 | 127886029 | -3.189554716 | PLK4 | 5 | 10649896 | 10650097 | 11.57436249 | ANKRD33B | 6 | 15523936 | 15524132 | -1.892275396 | DTNBP1 |
| 4 | 127886327 | 127886429 | -4.205004881 | PLK4 | 5 | 10652756 | 10652955 | 3.194554776 | RP11-54F2.1 | 6 | 16144921 | 16145119 | -1.019014558 | MYLIP |
| 4 | 127889839 | 127890038 | -3.655910633 | PLK4 | 5 | 10652756 | 10652955 | 3.194554776 | ANKRD33B | 6 | 16146690 | 16146888 | -1.041686145 | MYLIP |
| 4 | 128519039 | 128519252 | -1.254984506 | RP11-420A23.1 | 5 | 10654899 | 10655098 | 2.918110014 | ANKRD33B | 6 | 17707041 | 17707236 | 1.122653407 | RP11-500C11.3 |
| 1 | 180111181 | 180111383 | -1.78224104 | CEP350 | 5 | 10654899 | 10655098 | 2.918110014 | RP11-54F2.1 | 6 | 18465267 | 18465467 | 3.175338517 | RNF144B |
| 4 | 128862017 | 128862219 | -1.96118586 | JADE1 | 5 | 10656250 | 10656432 | 3.000756355 | ANKRD33B | 6 | 21030063 | 21030246 | 1.066927873 | CDKAL1 |
| 4 | 128872038 | 128872250 | -1.703871898 | SCLT1 | 5 | 10657083 | 10657259 | 2.321378156 | ANKRD33B | 6 | 25138165 | 25138357 | 1.672744469 | CMAHP |
| 4 | 128872038 | 128872250 | -1.703871898 | JADE1 | 5 | 10725160 | 10725347 | -6.687519716 | DAP | 6 | 25140398 | 25140597 | 1.377189263 | CMAHP |
| 4 | 129068559 | 129068839 | -1.720995076 | SCLT1 | 5 | 14610389 | 14610652 | -1.497272604 | FAM105A | 6 | 25140398 | 25140597 | 1.377189263 | RP3-425P12.4 |
| 4 | 129069685 | 129069883 | -2.508960954 | SCLT1 | 5 | 14710798 | 14710997 | -2.884463032 | ANKH | 6 | 26215911 | 26216111 | 2.142131774 | HIST1H2BG |
| 4 | 129070817 | 129070946 | -2.48022765 | SCLT1 | 5 | 17144741 | 17144933 | -2.362930139 | BASP1 | 6 | 26423425 | 26423689 | -1.326904832 | BTN2A3P |
| 4 | 129071577 | 129071764 | -2.232512818 | SCLT1 | 5 | 17144741 | 17144933 | -2.362930139 | AC091878.1 | 6 | 26430749 | 26430959 | -1.575169588 | BTN2A3P |
| 4 | 129072934 | 129073156 | -3.248212913 | SCLT1 | 5 | 17275634 | 17275835 | 1.530319308 | BASP1 | 1 | 206463949 | 206464164 | -2.086518387 | SRGAP2 |
| 4 | 131379926 | 131380106 | 1.796820745 | RP11-314N14.1 | 5 | 17713737 | 17713936 | 2.572668899 | RP11-454P21.1 | 6 | 28148482 | 28148672 | -2.587726484 | ZKSCAN8 |
| 4 | 131475334 | 131475572 | 1.597067348 | RP11-314N14.1 | 5 | 18707721 | 18707907 | 7.401863763 | CTD-2023M8.1 | 6 | 28166516 | 28166680 | -4.94208526 | ZNF192P1 |
| 1 | 180196456 | 180196756 | -1.476533416 | QSOX1 | 5 | 21534194 | 21534394 | -2.35207835 | GUSBP1 | 6 | 28365408 | 28365578 | 1.080768608 | ZKSCAN3 |
| 4 | 134200224 | 134200525 | -1.614697226 | PABPC4L | 5 | 21580558 | 21580756 | -1.748425964 | GUSBP1 | 1 | 206588219 | 206588395 | 1.191109963 | RASSF5 |
| 4 | 138407045 | 138407240 | -5.374096425 | LINC00499 | 5 | 21585761 | 21585969 | -1.730243329 | GUSBP1 | 1 | 206588219 | 206588395 | 1.191109963 | EIF2D |
| 4 | 139044868 | 139045069 | -2.601595439 | ELF2 | 5 | 31496743 | 31496931 | 7.85443417 | DROSHA | 6 | 28398104 | 28398280 | -1.072595868 | ZSCAN12 |
| 4 | 139044868 | 139045069 | -2.601595439 | NOCT | 5 | 31548712 | 31548894 | 1.407099652 | C5orf22 | 6 | 29003959 | 29004094 | -5.499295215 | ZNF311 |
| 4 | 139114928 | 139115129 | -1.549011134 | ELF2 | 5 | 32070504 | 32070641 | -5.207342618 | PDZD2 | 6 | 30186303 | 30186509 | -1.323606974 | TRIM26 |
| 4 | 139114928 | 139115129 | -1.549011134 | PPP1R14BP3 | 5 | 32099447 | 32099721 | -1.748779897 | PDZD2 | 6 | 30198838 | 30199043 | -9.289847636 | TRIM26 |
| 4 | 139300711 | 139300939 | -2.937148964 | NDUFC1 | 5 | 32126378 | 32126578 | -1.454452023 | GOLPH3 | 6 | 30278657 | 30278844 | -7.156335139 | HLA-L |
| 4 | 139446295 | 139446420 | 1.830079001 | RP11-83A24.2 | 5 | 32229723 | 32229923 | -1.262395874 | MTMR12 | 6 | 30278657 | 30278844 | -7.156335139 | HCG17 |
| 1 | 180502221 | 180502421 | -1.596249826 | ACBD6 | 5 | 33439612 | 33439810 | -1.040531987 | CTD-2203K17.1 | 6 | 30287450 | 30287603 | -1.286480687 | HLA-L |
| 4 | 139597456 | 139597700 | -3.133096104 | SETD7 | 5 | 34050259 | 34050464 | -1.122504619 | C1QTNF3-AMACR | 6 | 30287450 | 30287603 | -1.286480687 | HCG18 |
| 4 | 139600946 | 139601111 | -6.828715054 | SETD7 | 5 | 34051778 | 34051972 | -2.538310641 | C1QTNF3-AMACR | 6 | 30287450 | 30287603 | -1.286480687 | HCG17 |
| 4 | 139718719 | 139719021 | 1.429585213 | MGST2 | 5 | 34823182 | 34823417 | -2.056930635 | RAI14 | 6 | 30291639 | 30291894 | 1.110006663 | HLA-L |
| 4 | 139718719 | 139719021 | 1.429585213 | MAML3 | 5 | 34823946 | 34824142 | -1.861063921 | RAI14 | 6 | 30291639 | 30291894 | 1.110006663 | HCG18 |
| 4 | 139719701 | 139719860 | 2.227360854 | MAML3 | 5 | 34954526 | 34954723 | -1.502380665 | DNAJC21 | 6 | 30291639 | 30291894 | 1.110006663 | HCG17 |
| 4 | 139719701 | 139719860 | 2.227360854 | MGST2 | 5 | 34958497 | 34958696 | -1.648648615 | DNAJC21 | 6 | 30713022 | 30713150 | 1.768888835 | MDC1 |
| 4 | 139889616 | 139889756 | 1.462314956 | MAML3 | 5 | 35065282 | 35065482 | 3.663318452 | PRLR | 6 | 30713022 | 30713150 | 1.768888835 | MDC1-AS1 |
| 4 | 140622268 | 140622470 | 4.664715378 | TBC1D9 | 5 | 35876316 | 35876546 | 1.988551554 | IL7R | 1 | 206960782 | 206960980 | -2.129008166 | FCAMR |
| 4 | 143438245 | 143438447 | -1.072119739 | GAB1 | 5 | 35877596 | 35877814 | 2.697304667 | IL7R | 6 | 30743798 | 30743998 | -1.710200503 | IER3 |
| 4 | 145033130 | 145033297 | -1.861915783 | ANAPC10 | 5 | 36194936 | 36195135 | -1.591019523 | NADK2 | 1 | 207049396 | 207049603 | -1.093021495 | YOD1 |
| 4 | 146323173 | 146323328 | -2.243632913 | SLC10A7 | 1 | 192558118 | 192558237 | -1.698390205 | RP5-1011O1.2 | 1 | 207049396 | 207049603 | -1.093021495 | PFKFB2 |
| 4 | 146640842 | 146641043 | -5.083659592 | POU4F2 | 5 | 36877198 | 36877359 | 7.807699872 | NIPBL | 6 | 31577561 | 31577782 | -2.255028846 | TNF |
| 4 | 147636880 | 147637079 | 1.456351263 | TMEM184C | 1 | 192579237 | 192579434 | -1.195412958 | RGS1 | 6 | 31581298 | 31581484 | -6.073602462 | LTB |
| 1 | 180935687 | 180935872 | -1.224382404 | KIAA1614 | 5 | 37815885 | 37816047 | 2.746378662 | GDNF | 6 | 31629515 | 31629715 | 2.597523431 | PRRC2A |
| 4 | 147850786 | 147850984 | 8.026092965 | ARHGAP10 | 5 | 37815885 | 37816047 | 2.746378662 | GDNF-AS1 | 6 | 31811153 | 31811362 | 1.348287246 | HSPA1L |

| | | | | |
|---|---|---|---|---|
| 4 | 148072140 | 148072355 | 2.352528173 | ARHGAP10 |
| 4 | 150837531 | 150837767 | -1.146270197 | LRBA |
| 4 | 150852152 | 150852511 | -1.322839887 | LRBA |
| 1 | 180950250 | 180950450 | 2.244706783 | RP11-46A10.5 |
| 1 | 180950250 | 180950450 | 2.244706783 | KIAA1614 |
| 1 | 180950250 | 180950450 | 2.244706783 | KIAA1614-AS1 |
| 4 | 153295555 | 153295756 | 4.998544581 | TRIM2 |
| 4 | 153334931 | 153335131 | 7.663891342 | TRIM2 |
| 4 | 153603908 | 153604107 | -1.764999019 | KIAA0922 |
| 4 | 155710962 | 155711147 | 6.680492108 | GUCY1A3 |
| 4 | 156762829 | 156763012 | 2.37502861 | PDGFC |
| 1 | 180976338 | 180976537 | 1.072385912 | RP11-46A10.5 |
| 1 | 180976338 | 180976537 | 1.072385912 | STX6 |
| 4 | 158556013 | 158556336 | -1.208667941 | RXFP1 |
| 4 | 158568163 | 158568360 | -1.95014393 | RXFP1 |
| 4 | 158651929 | 158652129 | -1.404912557 | RXFP1 |
| 4 | 158868882 | 158869089 | 3.698795732 | FNIP2 |
| 4 | 158870191 | 158870390 | 5.183272037 | FNIP2 |
| 4 | 158880143 | 158880429 | 5.739601535 | FNIP2 |
| 4 | 158881254 | 158881454 | 8.045800336 | FNIP2 |
| 4 | 158904635 | 158904859 | 4.362858167 | FNIP2 |
| 4 | 158904635 | 158904859 | 4.362858167 | C4orf45 |
| 4 | 159358052 | 159358250 | 1.354563502 | RAPGEF2 |
| 4 | 165322097 | 165322298 | 1.250739043 | KLHL2 |
| 4 | 165341763 | 165341979 | 1.663304561 | MSMO1 |
| 4 | 168504488 | 168504669 | 4.538407799 | PALLD |
| 4 | 168504488 | 168504669 | 4.538407799 | DDX60L |
| 4 | 168512001 | 168512197 | 8.115460544 | PALLD |
| 4 | 168512001 | 168512197 | 8.115460544 | DDX60L |
| 4 | 168952793 | 168952896 | -1.227315316 | CBR4 |
| 4 | 169096422 | 169096640 | 1.278959232 | SH3RF1 |
| 1 | 9037438 | 9037630 | -2.977883443 | SLC2A5 |
| 4 | 173321639 | 173321836 | -1.092515947 | GALNT7 |
| 4 | 174283549 | 174283734 | -2.54913 | FBXO8 |
| 4 | 174283549 | 174283734 | -2.54913 | CEP44 |
| 4 | 176683717 | 176683971 | 3.093540112 | RP11-313E19.2 |
| 4 | 176683717 | 176683971 | 3.093540112 | VEGFC |
| 4 | 176687289 | 176687486 | 4.311994523 | RP11-313E19.2 |
| 4 | 176687289 | 176687486 | 4.311994523 | VEGFC |
| 4 | 177353389 | 177353589 | -2.415927578 | RP11-376O6.2 |
| 4 | 177353389 | 177353589 | -2.415927578 | NEIL3 |
| 4 | 177599043 | 177599242 | -1.242645276 | RP11-130F10.1 |
| 4 | 182793464 | 182793664 | -1.619798474 | TENM3 |
| 4 | 182800328 | 182800527 | -2.750145397 | TENM3 |
| 4 | 182890894 | 182891094 | -5.215947363 | DCTD |
| 1 | 181798709 | 181799009 | 1.771716885 | CACNA1E |
| 4 | 183183375 | 183183575 | 1.122068687 | WWC2 |
| 4 | 184009358 | 184009475 | -1.6118173 | STOX2 |
| 4 | 184011105 | 184011296 | -1.63581263 | STOX2 |
| 4 | 184388426 | 184388671 | 1.57239226 | IRF2 |
| 4 | 184475120 | 184475315 | 1.366990868 | RP11-326I11.3 |
| 4 | 184672675 | 184672825 | -2.077689206 | PRIMPOL |
| 4 | 184674898 | 184675092 | -5.198805861 | PRIMPOL |
| 4 | 185190877 | 185191073 | -1.542615782 | CFAP97 |

| | | | | |
|---|---|---|---|---|
| 5 | 39005378 | 39005502 | -2.256571611 | RICTOR |
| 1 | 192811632 | 192811832 | -2.478413057 | RGS2 |
| 5 | 39153535 | 39153727 | 1.781801158 | FYB |
| 5 | 39202029 | 39202230 | 2.610547029 | FYB |
| 5 | 39202621 | 39202820 | 2.561135612 | FYB |
| 5 | 39372405 | 39372606 | -2.316967697 | DAB2 |
| 5 | 39372405 | 39372606 | -2.316967697 | C9 |
| 5 | 39381539 | 39381741 | -1.349774382 | DAB2 |
| 5 | 39381539 | 39381741 | -1.349774382 | C9 |
| 5 | 39382718 | 39382924 | -1.858615807 | DAB2 |
| 5 | 39382718 | 39382924 | -1.858615807 | C9 |
| 5 | 40691929 | 40692105 | 1.734339972 | PTGER4 |
| 5 | 40691929 | 40692105 | 1.734339972 | TTC33 |
| 5 | 40692263 | 40692477 | 1.877221362 | PTGER4 |
| 5 | 40692263 | 40692477 | 1.877221362 | TTC33 |
| 1 | 192993561 | 192993740 | -1.938392057 | ZNF101P2 |
| 5 | 40721129 | 40721316 | 6.178225985 | TTC33 |
| 5 | 40827701 | 40827901 | -1.304373378 | RPL37 |
| 5 | 40843340 | 40843536 | 1.550736001 | CARD6 |
| 5 | 40852906 | 40853109 | 1.376244069 | CARD6 |
| 5 | 40853793 | 40853995 | 1.625404294 | CARD6 |
| 5 | 41742815 | 41743060 | 7.289628501 | OXCT1 |
| 1 | 193218751 | 193219040 | -1.494462015 | CDC73 |
| 5 | 43017858 | 43018058 | 1.08791749 | CTD-2201E18.3 |
| 5 | 43039692 | 43039942 | 3.401367622 | ANXA2R |
| 5 | 43039692 | 43039942 | 3.401367622 | CTD-2201E18.3 |
| 5 | 43042828 | 43043026 | 2.157113512 | AC025171.1 |
| 5 | 43042828 | 43043026 | 2.157113512 | CTD-2201E18.3 |
| 5 | 43042828 | 43043026 | 2.157113512 | ANXA2R |
| 5 | 43043069 | 43043258 | 2.551675503 | ANXA2R |
| 5 | 43043069 | 43043258 | 2.551675503 | CTD-2201E18.3 |
| 5 | 43043069 | 43043258 | 2.551675503 | AC025171.1 |
| 5 | 43044590 | 43044792 | 2.011960457 | AC025171.1 |
| 5 | 43044590 | 43044792 | 2.011960457 | CTD-2201E18.3 |
| 5 | 43055313 | 43055513 | 1.182271137 | CTD-2201E18.3 |
| 5 | 43056995 | 43057207 | 1.481401007 | CTD-2201E18.3 |
| 5 | 43057990 | 43058114 | 1.479964175 | CTD-2201E18.3 |
| 5 | 43059369 | 43059597 | 1.261343771 | CTD-2201E18.3 |
| 5 | 43060023 | 43060312 | 1.088708973 | CTD-2201E18.3 |
| 5 | 43136764 | 43136940 | -7.288005937 | ZNF131 |
| 1 | 925028 | 925204 | -3.488639962 | SAMD11 |
| 1 | 197101343 | 197101538 | -2.111640253 | ASPM |
| 5 | 43381809 | 43382006 | -2.089124933 | CCL28 |
| 5 | 43446335 | 43446507 | -1.045988978 | TMEM267 |
| 5 | 43453832 | 43454034 | -2.892151137 | TMEM267 |
| 5 | 43476262 | 43476472 | -2.241358012 | TMEM267 |
| 5 | 43506078 | 43506295 | -2.250743339 | RP11-159F24.3 |
| 5 | 43506078 | 43506295 | -2.250743339 | C5orf34 |
| 5 | 43522472 | 43522692 | -1.029682455 | RP11-159F24.5 |
| 5 | 43575404 | 43575684 | -2.06938445 | NNT-AS1 |
| 5 | 43579080 | 43579279 | -1.021202783 | NNT-AS1 |
| 5 | 44814915 | 44815130 | -9.346763897 | MRPS30 |
| 5 | 53982505 | 53982680 | -2.658979842 | ARL15 |
| 5 | 54519393 | 54519600 | -1.558924674 | SNX18 |

| | | | | |
|---|---|---|---|---|
| 1 | 207337086 | 207337285 | 7.688469661 | CD55 |
| 6 | 31828539 | 31828711 | -1.836212671 | HSPA1B |
| 6 | 32096916 | 32097053 | 1.015265536 | TNXB |
| 1 | 208216837 | 208217030 | 1.650160819 | PLXNA2 |
| 6 | 32838031 | 32838180 | 7.973544488 | XXbac-BPG246D15.9 |
| 6 | 32838031 | 32838180 | 7.973544488 | TAP2 |
| 6 | 33194485 | 33194684 | 1.380917861 | RXRB |
| 6 | 33203798 | 33203998 | 1.652780961 | SLC39A7 |
| 6 | 33404831 | 33404953 | -1.290704868 | KIFC1 |
| 6 | 34263427 | 34263614 | 1.022456211 | RP11-513I15.6 |
| 6 | 34263427 | 34263614 | 1.022456211 | RPL35P2 |
| 6 | 34270331 | 34270520 | 2.6139463 | RP11-513I15.6 |
| 6 | 35455936 | 35456148 | -1.298834604 | FANCE |
| 6 | 36085713 | 36085939 | -2.082094316 | MAPK14 |
| 6 | 36088042 | 36088254 | -1.760028412 | MAPK14 |
| 6 | 36363281 | 36363446 | -2.660964047 | ETV7 |
| 6 | 36366386 | 36366580 | -3.745926548 | ETV7 |
| 6 | 36684293 | 36684495 | 3.336031042 | CDKN1A |
| 6 | 36721593 | 36721770 | -1.426113843 | RAB44 |
| 6 | 36721973 | 36722218 | -1.410789099 | RAB44 |
| 1 | 211576199 | 211576399 | 1.176714491 | SLC30A1 |
| 6 | 37696505 | 37696698 | 5.304589369 | MDGA1 |
| 6 | 41098771 | 41098966 | 1.536308102 | NFYA |
| 6 | 41598977 | 41599175 | 1.720293734 | FOXP4 |
| 6 | 41796546 | 41796725 | -2.36310908 | USP49 |
| 6 | 42978343 | 42978764 | 1.317938113 | PEX6 |
| 6 | 43348439 | 43348644 | -1.024771584 | ZNF318 |
| 6 | 43454798 | 43455002 | 2.98919951 | DLK2 |
| 6 | 43671593 | 43671795 | 9.33507915 | MRPS18A |
| 6 | 43671593 | 43671795 | 9.33507915 | RSPH9 |
| 6 | 43770491 | 43770710 | 1.5820844 | RP1-261G23.7 |
| 6 | 43770491 | 43770710 | 1.5820844 | VEGFA |
| 6 | 44002882 | 44003082 | 1.503184937 | RP5-1120P11.1 |
| 6 | 44002882 | 44003082 | 1.503184937 | C6orf223 |
| 6 | 44938088 | 44938219 | -1.612550854 | SUPT3H |
| 6 | 44938088 | 44938219 | -1.612550854 | AL161905.1 |
| 6 | 45421763 | 45421937 | 1.777121659 | RP1-244F24.1 |
| 6 | 45421763 | 45421937 | 1.777121659 | RUNX2 |
| 6 | 45422208 | 45422460 | 1.953111696 | RUNX2 |
| 6 | 47284074 | 47284274 | 1.362756228 | TNFRSF21 |
| 6 | 47286218 | 47286439 | 1.064651938 | TNFRSF21 |
| 6 | 52404912 | 52405119 | -1.069775676 | EFHC1 |
| 6 | 52404912 | 52405119 | -1.069775676 | PAQR8 |
| 6 | 53498692 | 53498881 | -1.21490372 | GCLC |
| 6 | 54940776 | 54940974 | -2.019264114 | FAM83B |
| 1 | 944187 | 944287 | 1.424659796 | NOC2L |
| 1 | 944187 | 944287 | 1.424659796 | SAMD11 |
| 1 | 212858282 | 212858468 | 8.194000231 | FLVCR1 |
| 6 | 56618571 | 56618715 | 2.014428543 | DST |
| 6 | 57122446 | 57122599 | -2.974447218 | ZNF451 |
| 6 | 57122446 | 57122599 | -2.974447218 | RP11-203B9.4 |
| 6 | 57949697 | 57949933 | 1.240635104 | LINC00680 |
| 6 | 57949697 | 57949933 | 1.240635104 | GUSBP1 |
| 6 | 68764364 | 68764557 | -1.730448514 | ADGRB3 |

| | | | | |
|---|---|---|---|---|
| 1 | 182599061 | 182599262 | 3.644807699 | RGS16 |
| 4 | 188632139 | 188632239 | -4.041251952 | LINC01060 |
| 4 | 189982653 | 189982796 | -1.236800696 | TUBB7P |
| 1 | 182599995 | 182600205 | 5.573262225 | RGS16 |
| 5 | 199052 | 199250 | 1.021118173 | CCDC127 |
| 5 | 388720 | 388926 | -10.35289887 | AHRR |
| 1 | 182616020 | 182616192 | 1.161416161 | RP11-317P15.4 |
| 5 | 389622 | 389827 | -3.740345966 | AHRR |
| 5 | 393316 | 393516 | -9.544901087 | AHRR |
| 5 | 398371 | 398535 | -10.70268501 | AHRR |
| 5 | 401986 | 402190 | -10.29188067 | AHRR |
| 5 | 453526 | 453733 | -1.274335005 | EXOC3 |
| 5 | 471476 | 471675 | -1.053564329 | EXOC3 |
| 5 | 471476 | 471675 | -1.053564329 | PP7080 |
| 5 | 476378 | 476618 | 1.196589953 | CTD-2228K2.7 |
| 5 | 476378 | 476618 | 1.196589953 | SLC9A3 |
| 5 | 640793 | 641005 | -2.037206602 | CEP72 |
| 5 | 642418 | 642619 | -2.69187821 | CEP72 |
| 5 | 643116 | 643318 | -2.633004984 | CEP72 |
| 5 | 665670 | 665882 | -2.822733959 | TPPP |
| 5 | 665670 | 665882 | -2.822733959 | CEP72 |
| 5 | 667595 | 667795 | -2.224960374 | TPPP |
| 5 | 675840 | 676104 | -1.634603056 | CTD-2589H19.6 |
| 5 | 675840 | 676104 | -1.634603056 | TPPP |
| 5 | 685950 | 686315 | 2.669533156 | TPPP |
| 5 | 747976 | 748170 | -1.97227307 | ZDHHC11B |
| 5 | 750042 | 750174 | -2.19537761 | ZDHHC11B |
| 1 | 9103981 | 9104179 | -2.480000966 | GPR157 |
| 5 | 840100 | 840284 | -7.595529607 | ZDHHC11 |
| 5 | 863251 | 863476 | -1.451326972 | BRD9 |
| 5 | 864123 | 864361 | -2.367661782 | BRD9 |
| 5 | 990181 | 990369 | -2.228053038 | RP11-43F13.3 |
| 5 | 1017611 | 1017794 | -2.255794554 | NKD2 |
| 5 | 1038373 | 1038573 | -2.690649935 | NKD2 |
| 5 | 1051371 | 1051568 | -4.379944092 | SLC12A7 |
| 5 | 1051710 | 1051922 | -2.15155211 | SLC12A7 |
| 5 | 1052183 | 1052382 | -2.029446845 | SLC12A7 |
| 5 | 1056360 | 1056556 | -3.816468757 | SLC12A7 |
| 5 | 1060706 | 1060898 | -3.324697187 | SLC12A7 |
| 5 | 1064523 | 1064717 | -2.430105778 | SLC12A7 |
| 5 | 1262045 | 1262245 | -9.10710975 | TERT |
| 5 | 1318236 | 1318451 | -2.013866649 | CLPTM1L |
| 5 | 1461917 | 1462111 | -1.996790867 | LPCAT1 |
| 5 | 1463548 | 1463746 | -1.095400898 | LPCAT1 |
| 5 | 1468677 | 1468877 | -7.832001381 | LPCAT1 |
| 5 | 1475730 | 1475934 | -2.282525979 | LPCAT1 |
| 5 | 1491188 | 1491383 | -7.416445007 | LPCAT1 |
| 5 | 1798608 | 1798799 | -1.365163301 | MRPL36 |
| 5 | 1806718 | 1806920 | 7.870707158 | NDUFS6 |
| 5 | 5460624 | 5460879 | -1.02106247 | ICE1 |
| 1 | 184795603 | 184795802 | 1.023915443 | FAM129A |
| 5 | 5462757 | 5462965 | -1.183164367 | ICE1 |
| 5 | 5464640 | 5464886 | -1.421745848 | ICE1 |
| 5 | 5479596 | 5479769 | -6.413472825 | ICE1 |
| 5 | 54543096 | 54543297 | -1.939492619 | SNX18 |
| 5 | 55917150 | 55917273 | 5.851185575 | IL31RA |
| 5 | 55941120 | 55941320 | 1.335487678 | CTD-2031P19.5 |
| 5 | 55941120 | 55941320 | 1.335487678 | IL6ST |
| 5 | 57213927 | 57214121 | 1.196720733 | GPBP1 |
| 5 | 58454294 | 58454508 | -1.085104961 | PLK2 |
| 5 | 58495022 | 58495223 | -2.036204248 | GAPT |
| 5 | 61146752 | 61146951 | -2.884659185 | NDUFAF2 |
| 1 | 198211749 | 198211991 | 1.273471309 | NEK7 |
| 5 | 62545443 | 62545630 | -1.671977201 | IPO11 |
| 5 | 64226986 | 64227184 | -1.652427291 | RNF180 |
| 5 | 64709800 | 64709994 | 1.431679529 | FAM159B |
| 5 | 64845127 | 64845333 | -1.497101486 | CWC27 |
| 5 | 64845766 | 64845994 | -3.768544473 | CWC27 |
| 5 | 65470963 | 65471161 | -1.522901845 | ADAMTS6 |
| 1 | 198218657 | 198218834 | 4.200439718 | NEK7 |
| 5 | 66053714 | 66053924 | -1.049440565 | ERBIN |
| 5 | 66054363 | 66054550 | -1.30224107 | ERBIN |
| 1 | 200407180 | 200407386 | -2.337260872 | ZNF281 |
| 1 | 200407180 | 200407386 | -2.337260872 | RP11-469A15.2 |
| 5 | 66078403 | 66078626 | -1.221869572 | ERBIN |
| 5 | 66170612 | 66170830 | -1.100685027 | SREK1 |
| 5 | 67163753 | 67163952 | 6.527146704 | MAST4 |
| 5 | 67164527 | 67164755 | 2.924950051 | MAST4 |
| 5 | 67165997 | 67166205 | 2.446021466 | MAST4 |
| 1 | 200409102 | 200409285 | -2.22376697 | ZNF281 |
| 1 | 200409102 | 200409285 | -2.22376697 | RP11-469A15.2 |
| 5 | 67166708 | 67166878 | 2.686017371 | MAST4 |
| 5 | 67182554 | 67182754 | -2.51641838 | CD180 |
| 5 | 67183189 | 67183383 | -2.143785524 | CD180 |
| 5 | 67183615 | 67183791 | -4.070627829 | CD180 |
| 5 | 68297967 | 68298175 | -1.078557791 | PIK3R1 |
| 5 | 69133587 | 69133824 | -2.950423822 | CTC-498J12.3 |
| 5 | 69187867 | 69188110 | -2.132195722 | CTC-351M12.1 |
| 1 | 200553551 | 200553736 | -7.135147663 | KIF14 |
| 5 | 69189088 | 69189291 | -3.616650054 | CTC-351M12.1 |
| 5 | 69313458 | 69313712 | -1.796891101 | CFL1P5 |
| 5 | 69313458 | 69313712 | -1.796891101 | CCDC125 |
| 5 | 69414097 | 69414296 | -1.40531726 | RAD17 |
| 5 | 69553589 | 69553762 | -2.31871496 | OCLN |
| 5 | 70120516 | 70120702 | -5.001407508 | RP11-1319K7.1 |
| 5 | 71074138 | 71074299 | -2.239398472 | RP11-195E2.1 |
| 1 | 200617887 | 200618090 | -2.870753557 | KIF14 |
| 5 | 72220768 | 72220966 | -1.250807898 | MRPS27 |
| 5 | 72461108 | 72461241 | -1.05700813 | ZNF366 |
| 5 | 74781556 | 74781758 | -5.475642357 | FAM169A |
| 5 | 75028817 | 75028999 | 4.761360816 | GCNT4 |
| 5 | 76732254 | 76732427 | -1.882435795 | F2R |
| 5 | 76734384 | 76734599 | -2.139600183 | F2R |
| 5 | 77076987 | 77077182 | -3.990069789 | ZBED3 |
| 5 | 78170006 | 78170327 | 1.119614658 | AP3B1 |
| 5 | 78510037 | 78510327 | 1.027832909 | LHFPL2 |
| 5 | 79314520 | 79314720 | -2.062279525 | JMY |
| 5 | 79316012 | 79316212 | -1.835494903 | JMY |
| 6 | 70758059 | 70758249 | -5.836212671 | SMAP1 |
| 1 | 214383388 | 214383585 | 1.186070623 | PTPN14 |
| 6 | 73425362 | 73425565 | -1.306698859 | MB21D1 |
| 6 | 73695151 | 73695271 | 1.617388073 | RP11-553A21.3 |
| 6 | 79032421 | 79032659 | -7.237613389 | PHIP |
| 6 | 79201626 | 79201802 | -6.948260811 | HMGN3 |
| 6 | 79487088 | 79487286 | 1.322261864 | LCA5 |
| 1 | 214644603 | 214644802 | -1.022099923 | CENPF |
| 1 | 214645651 | 214645847 | -1.518624503 | CENPF |
| 6 | 83195783 | 83195983 | -1.057326083 | RWDD2A |
| 6 | 84762919 | 84763102 | 7.305281752 | TBX18 |
| 6 | 88143892 | 88144085 | 2.070904241 | CNR1 |
| 6 | 88144442 | 88144639 | 4.955321365 | CNR1 |
| 6 | 89283055 | 89283250 | 2.041884179 | GABRR2 |
| 6 | 89295482 | 89295681 | -1.889933101 | GABRR2 |
| 6 | 89630874 | 89631083 | -1.37191878 | ANKRD6 |
| 6 | 89630874 | 89631083 | -1.37191878 | LYRM2 |
| 6 | 89932406 | 89932627 | -1.660868329 | BACH2 |
| 6 | 89950916 | 89951113 | -2.412214218 | RP3-512E2.2 |
| 6 | 89950916 | 89951113 | -2.412214218 | BACH2 |
| 6 | 89951527 | 89951706 | -7.351951787 | BACH2 |
| 6 | 89951527 | 89951706 | -7.351951787 | RP3-512E2.2 |
| 6 | 93243236 | 93243524 | -1.20245873 | EPHA7 |
| 6 | 96487436 | 96487624 | -7.855039977 | UFL1-AS1 |
| 6 | 96494619 | 96494832 | -1.287557357 | UFL1-AS1 |
| 6 | 96497390 | 96497592 | -1.429820391 | UFL1-AS1 |
| 6 | 96500131 | 96500343 | -1.339625048 | UFL1-AS1 |
| 6 | 97260429 | 97260628 | 3.475051331 | MMS22L |
| 6 | 98835802 | 98836027 | -1.704042369 | POU3F2 |
| 6 | 99614729 | 99614863 | -5.497176718 | PRDM13 |
| 6 | 100740763 | 100740946 | -5.836212671 | ASCC3 |
| 6 | 102068583 | 102068757 | 1.403242783 | GRIK2 |
| 6 | 106512155 | 106512303 | 4.208926257 | AIM1 |
| 6 | 107070141 | 107070269 | -6.236852875 | BEND3 |
| 6 | 107635106 | 107635343 | 1.323382076 | SOBP |
| 6 | 107887987 | 107888187 | -8.903501329 | SEC63 |
| 1 | 219197327 | 219197526 | -1.260858856 | LYPLAL1 |
| 6 | 108416317 | 108416520 | -1.361579897 | LACE1 |
| 6 | 108514691 | 108514891 | -7.022026266 | LACE1 |
| 6 | 108683204 | 108683424 | -1.094312176 | FOXO3 |
| 6 | 108882967 | 108883075 | -2.162879978 | ARMC2 |
| 6 | 109094016 | 109094217 | -1.70373338 | SESN1 |
| 6 | 109133569 | 109133673 | -4.89395128 | CEP57L1 |
| 6 | 109136141 | 109136423 | -1.752663192 | CEP57L1 |
| 6 | 109162712 | 109162918 | -1.410568572 | CEP57L1 |
| 6 | 109415193 | 109415384 | -1.051110228 | PPIL6 |
| 6 | 110144456 | 110144649 | -3.599836172 | WASF1 |
| 6 | 111372692 | 111372996 | -1.087760037 | REV3L |
| 6 | 111375944 | 111376138 | -1.094714754 | REV3L |
| 6 | 113860518 | 113860705 | -1.120497346 | MARCKS |
| 6 | 113995466 | 113995650 | -4.112000837 | RP3-399L15.3 |
| 6 | 113995466 | 113995650 | -4.112000837 | HDAC2 |
| 1 | 220267100 | 220267295 | -1.093128279 | AURKAPS1 |
| 1 | 220267100 | 220267295 | -1.093128279 | RAB3GAP2 |

| | | | | |
|---|---|---|---|---|
| 5 | 6599748 | 6599947 | -1.055315805 | NSUN2 |
| 5 | 7867281 | 7867504 | -1.673319119 | MTRR |
| 5 | 7867281 | 7867504 | -1.673319119 | FASTKD3 |
| 1 | 185174465 | 185174666 | 2.263176985 | SWT1 |
| 5 | 10649896 | 10650097 | -8.186865902 | ANKRD33B |
| 5 | 10652756 | 10652955 | -1.161215617 | RP11-54F2.1 |
| 5 | 10652756 | 10652955 | -1.161215617 | ANKRD33B |
| 5 | 10654899 | 10655098 | -1.037409476 | ANKRD33B |
| 5 | 10654899 | 10655098 | -1.037409476 | RP11-54F2.1 |
| 5 | 10657083 | 10657259 | -1.312807015 | ANKRD33B |
| 1 | 185180244 | 185180440 | 2.948843131 | SWT1 |
| 5 | 14290859 | 14291058 | 1.047100861 | TRIO |
| 5 | 14508231 | 14508416 | 3.764606812 | TRIO |
| 5 | 14610389 | 14610652 | -3.878023245 | FAM105A |
| 5 | 14710798 | 14710997 | 3.237812386 | ANKH |
| 1 | 185236541 | 185236804 | 4.857046983 | SWT1 |
| 5 | 17144741 | 17144933 | -2.611744139 | BASP1 |
| 5 | 17144741 | 17144933 | -2.611744139 | AC091878.1 |
| 5 | 17713737 | 17713936 | -4.875236259 | RP11-454P21.1 |
| 5 | 18707721 | 18707907 | -4.02219706 | CTD-2023M8.1 |
| 5 | 20304850 | 20305070 | -2.430027182 | CDH18 |
| 5 | 20304850 | 20305070 | -2.430027182 | RP11-420O16.2 |
| 5 | 21470599 | 21470842 | -1.215863295 | GUSBP1 |
| 5 | 21473591 | 21473790 | -2.796087715 | GUSBP1 |
| 5 | 21474142 | 21474317 | -2.481151505 | GUSBP1 |
| 5 | 25190316 | 25190502 | -3.148266942 | RP11-549K20.1 |
| 5 | 31496743 | 31496931 | -8.214490832 | DROSHA |
| 5 | 31548712 | 31548894 | -2.478640413 | C5orf22 |
| 5 | 33435648 | 33435835 | -1.801158034 | CTD-2203K17.1 |
| 5 | 33439612 | 33439810 | -1.493405488 | CTD-2203K17.1 |
| 5 | 34050259 | 34050464 | -2.51672549 | C1QTNF3-AMACR |
| 5 | 34051778 | 34051972 | -2.858099292 | C1QTNF3-AMACR |
| 5 | 34823182 | 34823417 | 1.52635824 | RAI14 |
| 5 | 34823946 | 34824142 | 2.052335762 | RAI14 |
| 5 | 34950573 | 34950771 | -1.542142908 | DNAJC21 |
| 5 | 34958497 | 34958696 | -1.577835074 | DNAJC21 |
| 5 | 35065282 | 35065482 | -4.942786596 | PRLR |
| 5 | 36686069 | 36686272 | 6.974394404 | CTD-2353F22.1 |
| 5 | 36686069 | 36686272 | 6.974394404 | SLC1A3 |
| 5 | 36877198 | 36877359 | -8.179866202 | NIPBL |
| 5 | 36885411 | 36885613 | -1.280336335 | NIPBL |
| 5 | 36885411 | 36885613 | -1.280336335 | KRT18P31 |
| 1 | 192579237 | 192579434 | 15.26462439 | RGS1 |
| 5 | 37168894 | 37169095 | -1.151009956 | C5orf42 |
| 5 | 37286583 | 37286796 | -2.088327278 | CTD-2124B8.2 |
| 5 | 37446724 | 37446921 | -1.640581435 | WDR70 |
| 5 | 37527462 | 37527661 | -1.004468328 | WDR70 |
| 5 | 38942124 | 38942336 | 1.554998732 | OSMR |
| 5 | 38942124 | 38942336 | 1.554998732 | RICTOR |
| 5 | 39005378 | 39005502 | 7.789952577 | RICTOR |
| 1 | 192811632 | 192811832 | 4.088733947 | RGS2 |
| 5 | 39153535 | 39153727 | 2.006798802 | FYB |
| 5 | 39372405 | 39372606 | 4.017216271 | C9 |
| 5 | 39372405 | 39372606 | 4.017216271 | DAB2 |

| | | | | |
|---|---|---|---|---|
| 5 | 79321611 | 79321814 | -8.807297414 | JMY |
| 1 | 200848996 | 200849194 | -1.101814469 | CAMSAP2 |
| 5 | 79512861 | 79513064 | 2.398994268 | HOMER1 |
| 5 | 79738098 | 79738299 | -2.015058998 | CMYA5 |
| 5 | 80143328 | 80143505 | -1.520582688 | CTC-458I2.2 |
| 5 | 80143328 | 80143505 | -1.520582688 | SERINC5 |
| 5 | 80429654 | 80429909 | -2.599427764 | ZFYVE16 |
| 5 | 80430329 | 80430521 | -7.331501044 | ZFYVE16 |
| 5 | 80432971 | 80433169 | -7.898729443 | ZFYVE16 |
| 1 | 200849784 | 200849970 | -1.692188838 | CAMSAP2 |
| 5 | 81239101 | 81239252 | -5.791776465 | CKMT2-AS1 |
| 5 | 81239101 | 81239252 | -5.791776465 | CKMT2 |
| 5 | 81304830 | 81305046 | -1.194283077 | ZCCHC9 |
| 5 | 81699310 | 81699499 | -1.856599303 | SSBP2 |
| 5 | 82318370 | 82318605 | -2.705899922 | ATP6AP1L |
| 5 | 83522027 | 83522210 | -1.076001547 | VCAN |
| 5 | 83537429 | 83537602 | -4.667695177 | VCAN-AS1 |
| 5 | 83537429 | 83537602 | -4.667695177 | VCAN |
| 5 | 83580928 | 83581177 | 1.16461951 | VCAN |
| 5 | 83580928 | 83581177 | 1.16461951 | VCAN-AS1 |
| 1 | 200971457 | 200971656 | -2.661324921 | KIF21B |
| 5 | 87305411 | 87305580 | -1.148769767 | RASA1 |
| 5 | 88190446 | 88190649 | -6.180971887 | TMEM161B |
| 5 | 88335733 | 88335876 | -2.374825427 | TMEM161B-AS1 |
| 5 | 88350647 | 88350810 | -2.692548764 | TMEM161B-AS1 |
| 5 | 88667038 | 88667237 | -2.824670864 | LINC00461 |
| 5 | 88797290 | 88797490 | -1.880219181 | MEF2C |
| 5 | 88944465 | 88944663 | -6.81467831 | MEF2C-AS1 |
| 5 | 90461409 | 90461614 | -2.158127382 | MBLAC2 |
| 5 | 90473807 | 90473933 | -3.190188905 | POLR3G |
| 5 | 90473807 | 90473933 | -3.190188905 | MBLAC2 |
| 1 | 201514871 | 201515070 | -2.386828299 | RP11-134G8.7 |
| 5 | 91476482 | 91476611 | 1.334777724 | RAB5CP2 |
| 5 | 93814998 | 93815169 | -5.806203897 | FAM172A |
| 5 | 93953138 | 93953297 | 4.554262228 | FAM172A |
| 5 | 94695149 | 94695415 | -1.494953766 | SLF1 |
| 5 | 95556347 | 95556459 | -3.196158711 | ARSK |
| 1 | 201520320 | 201520525 | -2.263759798 | RP11-134G8.7 |
| 1 | 201520320 | 201520525 | -2.263759798 | RP11-134G8.6 |
| 1 | 201520320 | 201520525 | -2.263759798 | RP11-134G8.5 |
| 5 | 95620230 | 95620418 | 1.142910653 | GPR150 |
| 5 | 96880095 | 96880286 | 1.079947388 | ERAP2 |
| 5 | 96880095 | 96880286 | 1.079947388 | CTD-2260A17.2 |
| 5 | 96917552 | 96917747 | 1.747019905 | ERAP2 |
| 5 | 96917552 | 96917747 | 1.747019905 | CTD-2260A17.2 |
| 5 | 98779677 | 98779873 | -2.283727891 | RGMB |
| 5 | 102982495 | 102982707 | -1.522336631 | PAM |
| 5 | 103129413 | 103129604 | -1.630800455 | PPIP5K2 |
| 5 | 103559276 | 103559465 | -7.358624003 | NUDT12 |
| 5 | 108208401 | 108208526 | -3.120157165 | FBXL17 |
| 5 | 108211083 | 108211236 | -2.38488587 | FBXL17 |
| 5 | 110566518 | 110566641 | -4.421175172 | TMEM232 |
| 5 | 112837762 | 112837969 | -1.101896822 | APC |
| 5 | 112837762 | 112837969 | -1.101896822 | CTC-554D6.1 |

| | | | | |
|---|---|---|---|---|
| 6 | 116739829 | 116740024 | -1.667651386 | KPNA5 |
| 6 | 119306951 | 119307152 | 1.493924861 | MAN1A1 |
| 1 | 220831246 | 220831441 | -6.891601445 | LINC01352 |
| 6 | 119348542 | 119348747 | 1.604832908 | MAN1A1 |
| 6 | 121447700 | 121447920 | 1.541349285 | GJA1 |
| 6 | 124962755 | 124962976 | 1.966650595 | RNF217-AS1 |
| 6 | 124962755 | 124962976 | 1.966650595 | RNF217 |
| 1 | 220861858 | 220861965 | -2.379319819 | HLX-AS1 |
| 6 | 125889366 | 125889561 | -1.0584993 | NCOA7 |
| 1 | 220878827 | 220879033 | 1.377772513 | HLX-AS1 |
| 6 | 126644097 | 126644256 | -5.383264454 | PRELID1P1 |
| 6 | 127363340 | 127363539 | 1.322264309 | RPL5P18 |
| 6 | 127446871 | 127447055 | -1.69682666 | RP3-403A15.5 |
| 6 | 127446871 | 127447055 | -1.69682666 | KIAA0408 |
| 6 | 127516445 | 127516682 | 1.546872613 | SOGA3 |
| 6 | 127516445 | 127516682 | 1.546872613 | RP3-403A15.5 |
| 6 | 129409069 | 129409260 | 2.082890257 | LAMA2 |
| 6 | 130440835 | 130440946 | 2.103946426 | TMEM200A |
| 6 | 136192088 | 136192288 | 1.770241834 | PDE7B |
| 6 | 136192088 | 136192288 | 1.770241834 | RP13-143G15.4 |
| 6 | 136636389 | 136636573 | -6.771092744 | MAP3K5 |
| 6 | 136636389 | 136636573 | -6.771092744 | RP3-325F22.3 |
| 6 | 136786163 | 136786302 | -2.921222475 | MAP3K5 |
| 6 | 137856385 | 137856597 | -1.307978142 | RP11-356I2.4 |
| 6 | 137879966 | 137880166 | -1.482701598 | TNFAIP3 |
| 6 | 138415278 | 138415473 | -7.13795312 | HEBP2 |
| 6 | 138422348 | 138422518 | -1.233079135 | NHSL1 |
| 6 | 138432354 | 138432567 | -1.089258379 | NHSL1 |
| 6 | 138433187 | 138433382 | -1.159429647 | NHSL1 |
| 6 | 138650303 | 138650445 | -1.736243886 | NHSL1 |
| 6 | 138650303 | 138650445 | -1.736243886 | RP11-390P2.2 |
| 6 | 138730109 | 138730287 | -3.157157455 | GVQW2 |
| 6 | 139372606 | 139372805 | 2.37797848 | CITED2 |
| 6 | 139373132 | 139373318 | 1.045885254 | CITED2 |
| 6 | 139373606 | 139373743 | 14.04230237 | CITED2 |
| 6 | 142211643 | 142211821 | -4.448922728 | VTA1 |
| 6 | 142770430 | 142770624 | -1.597417505 | HIVEP2 |
| 1 | 222627988 | 222628189 | -1.429460163 | MIA3 |
| 6 | 143948212 | 143948385 | 2.655617759 | PLAGL1 |
| 6 | 147254250 | 147254364 | -3.439113419 | STXBP5 |
| 6 | 147303511 | 147303701 | 5.850653231 | STXBP5 |
| 6 | 147407808 | 147407991 | -1.512928225 | YAP1P1 |
| 6 | 149317227 | 149317380 | 8.658551711 | TAB2 |
| 1 | 223002478 | 223002661 | 1.776455734 | DISP1 |
| 6 | 149969021 | 149969220 | -1.40931566 | ULBP1 |
| 6 | 149972392 | 149972534 | -3.813504433 | ULBP1 |
| 6 | 150935273 | 150935466 | -1.116207329 | MTHFD1L |
| 6 | 150935273 | 150935466 | -1.116207329 | RP1-99E18.2 |
| 6 | 151091160 | 151091284 | -2.018820216 | MTHFD1L |
| 6 | 151091160 | 151091284 | -2.018820216 | RP1-292B18.4 |
| 6 | 154222752 | 154222940 | 1.278771459 | OPRM1 |
| 6 | 154222752 | 154222940 | 1.278771459 | IPCEF1 |
| 6 | 154860621 | 154860801 | 6.757295392 | TIAM2 |
| 6 | 156778035 | 156778268 | 10.84293067 | RP11-230C9.2 |

| | | | | |
|---|---|---|---|---|
| 5 | 39381539 | 39381741 | 4.472010359 | DAB2 |
| 5 | 39381539 | 39381741 | 4.472010359 | C9 |
| 5 | 39382718 | 39382924 | 4.760877674 | DAB2 |
| 5 | 39382718 | 39382924 | 4.760877674 | C9 |
| 5 | 40680047 | 40680257 | -7.928686795 | PTGER4 |
| 5 | 40680047 | 40680257 | -7.928686795 | TTC33 |
| 5 | 40681136 | 40681335 | -8.858054702 | PTGER4 |
| 5 | 40681136 | 40681335 | -8.858054702 | TTC33 |
| 5 | 40721129 | 40721316 | -6.502198026 | TTC33 |
| 5 | 40843340 | 40843536 | -1.088396866 | CARD6 |
| 5 | 41742815 | 41743060 | -7.033380966 | OXCT1 |
| 5 | 41743498 | 41743699 | -1.165789132 | OXCT1 |
| 5 | 41925909 | 41926103 | -1.619445998 | FBXO4 |
| 1 | 193218751 | 193219040 | 1.069566921 | CDC73 |
| 5 | 43017858 | 43018058 | -1.462328446 | CTD-2201E18.3 |
| 5 | 43034309 | 43034508 | -1.866168134 | CTD-2035E11.4 |
| 5 | 43034309 | 43034508 | -1.866168134 | CTD-2201E18.3 |
| 5 | 43039692 | 43039942 | -1.271289325 | CTD-2201E18.3 |
| 5 | 43039692 | 43039942 | -1.271289325 | ANXA2R |
| 5 | 43042828 | 43043026 | -1.151509908 | CTD-2201E18.3 |
| 5 | 43042828 | 43043026 | -1.151509908 | ANXA2R |
| 5 | 43042828 | 43043026 | -1.151509908 | AC025171.1 |
| 5 | 43043069 | 43043258 | -1.222939315 | CTD-2201E18.3 |
| 5 | 43043069 | 43043258 | -1.222939315 | ANXA2R |
| 5 | 43043069 | 43043258 | -1.222939315 | AC025171.1 |
| 5 | 43044590 | 43044792 | -1.668466278 | AC025171.1 |
| 5 | 43044590 | 43044792 | -1.668466278 | CTD-2201E18.3 |
| 5 | 43054013 | 43054213 | -1.249904197 | CTD-2201E18.3 |
| 5 | 43057990 | 43058114 | -1.117443997 | CTD-2201E18.3 |
| 5 | 43071143 | 43071346 | -1.553295186 | ZNF131 |
| 5 | 43136764 | 43136940 | -1.2996665 | ZNF131 |
| 5 | 43161647 | 43161779 | -1.193785266 | ZNF131 |
| 5 | 43174765 | 43174968 | -1.153878652 | ZNF131 |
| 5 | 43245742 | 43245922 | -2.757849919 | NIM1K |
| 1 | 925028 | 925204 | 3.488639962 | SAMD11 |
| 1 | 197101343 | 197101538 | -3.125934364 | ASPM |
| 5 | 43496224 | 43496402 | -5.819579898 | RP11-159F24.3 |
| 5 | 43496224 | 43496402 | -5.819579898 | EEF1A1P19 |
| 5 | 43496224 | 43496402 | -5.819579898 | C5orf34 |
| 5 | 43506078 | 43506295 | -2.387440409 | RP11-159F24.3 |
| 5 | 43506078 | 43506295 | -2.387440409 | C5orf34 |
| 1 | 197104402 | 197104591 | -4.512966102 | ASPM |
| 5 | 50794967 | 50795201 | -1.025618012 | PARP8 |
| 5 | 53982505 | 53982680 | 1.329355826 | ARL15 |
| 5 | 55231389 | 55231543 | -2.125603305 | CCNO |
| 5 | 55283309 | 55283653 | -1.509089176 | DHX29 |
| 5 | 55283309 | 55283653 | -1.509089176 | RP11-506H20.1 |
| 5 | 55917150 | 55917273 | -2.39106695 | IL31RA |
| 5 | 56881803 | 56882092 | -1.439955591 | MAP3K1 |
| 5 | 58495022 | 58495223 | -2.934507543 | GAPT |
| 5 | 58974271 | 58974428 | -1.930554343 | PDE4D |
| 5 | 59894267 | 59894465 | -3.232456814 | PDE4D |
| 5 | 60121801 | 60121986 | -5.403677461 | PDE4D |
| 5 | 60122947 | 60123083 | -7.974274874 | PDE4D |
| 5 | 112841579 | 112841769 | -1.436923998 | CTC-554D6.1 |
| 5 | 112841579 | 112841769 | -1.436923998 | APC |
| 5 | 114482271 | 114482401 | -3.309015101 | KCNN2 |
| 5 | 115126490 | 115126690 | 1.935184053 | TRIM36 |
| 5 | 115525285 | 115525486 | 1.452235122 | FEM1C |
| 5 | 115543415 | 115543616 | 1.314371127 | FEM1C |
| 5 | 115580678 | 115580877 | 1.666273851 | TMED7-TICAM2 |
| 5 | 115580678 | 115580877 | 1.666273851 | TICAM2 |
| 5 | 116168052 | 116168269 | -1.751449824 | COMMD10 |
| 5 | 116268627 | 116268854 | -1.359794263 | COMMD10 |
| 5 | 116446324 | 116446523 | 9.363401775 | SEMA6A |
| 5 | 119393316 | 119393596 | 2.325838565 | TNFAIP8 |
| 5 | 120631437 | 120631593 | -1.367393874 | PRR16 |
| 1 | 201877769 | 201877967 | -1.437881919 | IPO9 |
| 5 | 121998130 | 121998235 | 2.153556913 | SRFBP1 |
| 5 | 122078111 | 122078268 | 3.953445298 | LOX |
| 5 | 123037180 | 123037331 | 4.820122468 | RP11-359P5.1 |
| 5 | 123087426 | 123087598 | -1.382285028 | AC106786.1 |
| 1 | 201878951 | 201879147 | -2.134857548 | IPO9 |
| 5 | 123346588 | 123346803 | -2.288358241 | CEP120 |
| 5 | 123401854 | 123402005 | -3.034120431 | KRT8P33 |
| 5 | 123401854 | 123402005 | -3.034120431 | CEP120 |
| 5 | 123552947 | 123553117 | -1.388882558 | CSNK1G3 |
| 5 | 124647826 | 124648021 | 2.152904215 | ZNF608 |
| 5 | 126608225 | 126608431 | -1.877075909 | PHAX |
| 5 | 126904187 | 126904350 | -6.31467831 | Mar-03 |
| 5 | 126904717 | 126904838 | -2.276563207 | Mar-03 |
| 5 | 127551725 | 127551943 | -1.272805048 | PRRC1 |
| 5 | 128322295 | 128322456 | -6.074079386 | FBN2 |
| 5 | 131429100 | 131429301 | -1.278737665 | RAPGEF6 |
| 5 | 131429100 | 131429301 | -1.278737665 | CTC-432M15.3 |
| 5 | 131430998 | 131431198 | -1.163924524 | CTC-432M15.3 |
| 5 | 131430998 | 131431198 | -1.163924524 | RAPGEF6 |
| 5 | 131461840 | 131461997 | -1.630766748 | CTC-432M15.3 |
| 5 | 131461840 | 131461997 | -1.630766748 | RAPGEF6 |
| 5 | 131672330 | 131672536 | -1.076977265 | FNIP1 |
| 5 | 131672330 | 131672536 | -1.076977265 | CTC-432M15.3 |
| 1 | 202147817 | 202148014 | -1.203779535 | PTPN7 |
| 5 | 131724732 | 131724931 | -2.590248236 | CTC-432M15.3 |
| 5 | 131724732 | 131724931 | -2.590248236 | FNIP1 |
| 5 | 132294485 | 132294606 | 1.600816931 | SLC22A4 |
| 5 | 132294485 | 132294606 | 1.600816931 | P4HA2 |
| 5 | 132294953 | 132295132 | 1.29248125 | SLC22A4 |
| 5 | 132294953 | 132295132 | 1.29248125 | P4HA2 |
| 5 | 132349302 | 132349442 | -2.26569073 | AC034220.3 |
| 5 | 132470118 | 132470308 | 3.842748009 | C5orf56 |
| 5 | 132470118 | 132470308 | 3.842748009 | AC116366.6 |
| 1 | 202728951 | 202729153 | -2.845410533 | KDM5B |
| 5 | 134197645 | 134197748 | -1.205108703 | CTD-2410N18.5 |
| 5 | 134197645 | 134197748 | -1.205108703 | PPP2CA |
| 5 | 134331521 | 134331767 | -2.260029767 | CDKL3 |
| 5 | 134331521 | 134331767 | -2.260029767 | CTD-2410N18.4 |
| 5 | 134336907 | 134337009 | -5.518086806 | CTD-2410N18.4 |
| 5 | 134336907 | 134337009 | -5.518086806 | CDKL3 |
| 6 | 156778035 | 156778268 | 10.84293067 | ARID1B |
| 6 | 156778677 | 156778865 | 9.175661587 | ARID1B |
| 6 | 157292726 | 157292928 | 1.071539363 | TMEM242 |
| 6 | 157673099 | 157673299 | 1.150301386 | ZDHHC14 |
| 6 | 158096037 | 158096205 | 1.714534768 | SYNJ2 |
| 6 | 158197020 | 158197147 | 1.450002105 | GTF2H5 |
| 6 | 158614827 | 158614943 | -5.991853096 | TMEM181 |
| 6 | 158614827 | 158614943 | -5.991853096 | TATDN2P2 |
| 6 | 159047908 | 159048106 | 1.170760616 | RP1-111C20.4 |
| 6 | 160036524 | 160036709 | -5.241403978 | IGF2R |
| 6 | 160992178 | 160992366 | 2.178892429 | MAP3K4 |
| 6 | 160992178 | 160992366 | 2.178892429 | RP3-428L16.2 |
| 6 | 161137406 | 161137619 | 2.534622953 | AGPAT4 |
| 6 | 161175788 | 161175931 | -7.087618825 | AGPAT4 |
| 6 | 161176410 | 161176610 | -1.839729483 | AGPAT4 |
| 6 | 161193188 | 161193382 | -1.693884062 | AGPAT4 |
| 6 | 166467669 | 166467860 | 5.254887502 | RPS6KA2 |
| 6 | 169237802 | 169238003 | -3.095795448 | THBS2 |
| 6 | 169237802 | 169238003 | -3.095795448 | XXyac-YX65C7_A.2 |
| 6 | 169694796 | 169694992 | -1.167312941 | WDR27 |
| 6 | 169723271 | 169723467 | -7.899261214 | PHF10 |
| 6 | 170276514 | 170276695 | -5.269579406 | RP5-894D12.5 |
| 6 | 170276514 | 170276695 | -5.269579406 | LINC01624 |
| 6 | 170394549 | 170394739 | 2.537192262 | FAM120B |
| 7 | 80463 | 80669 | -1.195940466 | AC093627.8 |
| 7 | 80463 | 80669 | -1.195940466 | AC093627.7 |
| 1 | 224739326 | 224739526 | -1.52840219 | CNIH3 |
| 1 | 224772698 | 224772896 | -1.095853448 | RP11-449J1.1 |
| 7 | 196089 | 196376 | -1.978940505 | FAM20C |
| 7 | 198195 | 198396 | -1.442395674 | FAM20C |
| 7 | 199827 | 200045 | -1.748463082 | FAM20C |
| 7 | 203645 | 203843 | 8.8575453 | FAM20C |
| 7 | 205146 | 205388 | 9.545573046 | FAM20C |
| 7 | 218660 | 218944 | 1.419831594 | FAM20C |
| 1 | 224938043 | 224938194 | -5.475642357 | DNAH14 |
| 7 | 242891 | 243101 | -1.81486699 | FAM20C |
| 7 | 244990 | 245192 | -1.263872314 | FAM20C |
| 7 | 250822 | 251025 | 6.569455859 | FAM20C |
| 1 | 224977737 | 224977920 | -1.887782425 | DNAH14 |
| 7 | 550199 | 550361 | 1.375628126 | PRKAR1B |
| 7 | 751857 | 752072 | -1.69487174 | DNAAF5 |
| 7 | 881797 | 882039 | -6.377443751 | GET4 |
| 7 | 881797 | 882039 | -6.377443751 | SUN1 |
| 7 | 881797 | 882039 | -6.377443751 | RP11-449P15.2 |
| 7 | 884839 | 884959 | -3.745926548 | RP11-449P15.2 |
| 7 | 884839 | 884959 | -3.745926548 | GET4 |
| 7 | 884839 | 884959 | -3.745926548 | SUN1 |
| 7 | 895059 | 895246 | -6.142701109 | SUN1 |
| 7 | 895059 | 895246 | -6.142701109 | GET4 |
| 7 | 898780 | 898979 | 1.262021708 | COX19 |
| 7 | 898780 | 898979 | 1.262021708 | ADAP1 |
| 7 | 907541 | 907740 | 6.037070731 | ADAP1 |
| 7 | 907541 | 907740 | 6.037070731 | COX19 |
| 7 | 932345 | 932554 | 1.383516189 | COX19 |

| | | | | |
|---|---|---|---|---|
| 5 | 60892025 | 60892232 | -1.321151244 | ERCC8 |
| 5 | 60892025 | 60892232 | -1.321151244 | GNL3LP1 |
| 5 | 60892416 | 60892602 | -2.092061169 | ERCC8 |
| 5 | 60892416 | 60892602 | -2.092061169 | GNL3LP1 |
| 1 | 198211749 | 198211991 | -1.35042765 | NEK7 |
| 5 | 61543921 | 61544121 | 1.182564656 | ZSWIM6 |
| 5 | 62545443 | 62545630 | -2.512863413 | IPO11 |
| 5 | 64226986 | 64227184 | -1.164770639 | RNF180 |
| 5 | 64845127 | 64845333 | -1.066747902 | CWC27 |
| 5 | 65548707 | 65548879 | -7.615160358 | CENPK |
| 5 | 65629840 | 65630037 | -1.063766617 | TRAPPC13 |
| 5 | 65629840 | 65630037 | -1.063766617 | CTC-534A2.2 |
| 1 | 200407180 | 200407386 | 1.222848269 | RP11-469A15.2 |
| 1 | 200407180 | 200407386 | 1.222848269 | ZNF281 |
| 5 | 66078403 | 66078626 | 1.205340315 | ERBIN |
| 5 | 66178944 | 66179147 | -1.087074327 | SREK1 |
| 5 | 67164527 | 67164755 | 1.560145284 | MAST4 |
| 5 | 67165997 | 67166205 | 2.764788799 | MAST4 |
| 1 | 200409102 | 200409285 | 1.396390989 | ZNF281 |
| 1 | 200409102 | 200409285 | 1.396390989 | RP11-469A15.2 |
| 5 | 67166708 | 67166878 | 1.786610964 | MAST4 |
| 5 | 67183615 | 67183791 | 1.561666875 | CD180 |
| 5 | 67183999 | 67184185 | 7.048974905 | CD180 |
| 5 | 69187867 | 69188110 | -1.577823733 | CTC-351M12.1 |
| 1 | 200553551 | 200553736 | -2.597017519 | KIF14 |
| 5 | 69313458 | 69313712 | -1.002756566 | CFL1P5 |
| 5 | 69313458 | 69313712 | -1.002756566 | CCDC125 |
| 5 | 69414097 | 69414296 | -1.831079868 | RAD17 |
| 5 | 69553589 | 69553762 | -3.02595889 | OCLN |
| 5 | 69632528 | 69632721 | -8.111152182 | RP11-848G14.5 |
| 5 | 70120516 | 70120702 | -1.944389423 | RP11-1319K7.1 |
| 1 | 200617887 | 200618090 | -3.118363329 | KIF14 |
| 5 | 71510715 | 71511013 | -1.228536443 | BDP1 |
| 5 | 72220768 | 72220966 | -1.609005513 | MRPS27 |
| 5 | 72443879 | 72444105 | 3.019860955 | ZNF366 |
| 5 | 72461108 | 72461241 | 5.680971887 | ZNF366 |
| 5 | 73580038 | 73580250 | -1.572774628 | UTP15 |
| 5 | 74635630 | 74635831 | 3.460211425 | ENC1 |
| 5 | 76732254 | 76732427 | 1.728721533 | F2R |
| 5 | 76734384 | 76734599 | 1.357265168 | F2R |
| 5 | 77630877 | 77631068 | -5.359006474 | OTP |
| 5 | 77775645 | 77775844 | -8.797599597 | TBCA |
| 5 | 78175086 | 78175283 | -2.018358856 | AP3B1 |
| 5 | 78510037 | 78510327 | 1.16577225 | LHFPL2 |
| 5 | 79314520 | 79314720 | 1.589273077 | JMY |
| 5 | 79316012 | 79316212 | 1.267724084 | JMY |
| 5 | 79321611 | 79321814 | 1.334849895 | JMY |
| 1 | 200848996 | 200849194 | 4.079804234 | CAMSAP2 |
| 5 | 79456943 | 79457138 | -4.75389732 | HOMER1 |
| 5 | 79512861 | 79513064 | 1.719235783 | HOMER1 |
| 5 | 79738098 | 79738299 | -1.656843916 | CMYA5 |
| 5 | 80429654 | 80429909 | 2.609110426 | ZFYVE16 |
| 5 | 80430329 | 80430521 | 7.331501044 | ZFYVE16 |
| 5 | 80432971 | 80433169 | 4.521285692 | ZFYVE16 |
| 5 | 134661230 | 134661389 | 1.124952133 | SEC24A |
| 5 | 134741119 | 134741300 | -2.077806637 | CAMLG |
| 5 | 134857240 | 134857469 | -1.808548273 | C5orf24 |
| 5 | 135028486 | 135028688 | 1.199890627 | PITX1 |
| 5 | 135031384 | 135031583 | 9.65851637 | PITX1 |
| 5 | 135033291 | 135033530 | 2.407698852 | C5orf66 |
| 5 | 135033291 | 135033530 | 2.407698852 | PITX1 |
| 5 | 135321643 | 135321849 | 1.025854309 | C5orf66 |
| 5 | 135322280 | 135322471 | -6.118007096 | C5orf66 |
| 5 | 138164864 | 138165079 | -1.104928979 | BRD8 |
| 5 | 138391223 | 138391432 | -1.952976718 | KDM3B |
| 5 | 138391957 | 138392166 | -1.16264168 | KDM3B |
| 5 | 138393055 | 138393249 | -1.732907618 | KDM3B |
| 5 | 138467094 | 138467252 | -1.316934925 | EGR1 |
| 1 | 203064544 | 203064715 | -5.754392581 | PPFIA4 |
| 5 | 139399240 | 139399471 | -2.665128227 | SPATA24 |
| 5 | 139682863 | 139683059 | -1.828431999 | CXXC5 |
| 5 | 139809524 | 139809695 | -1.768393953 | PSD2 |
| 5 | 140109215 | 140109404 | -5.26080022 | PURA |
| 1 | 203109535 | 203109752 | -7.100525907 | ADORA1 |
| 5 | 140111602 | 140111796 | -1.541494682 | PURA |
| 5 | 140115022 | 140115224 | -1.016249419 | PURA |
| 5 | 140453969 | 140454118 | 3.656441478 | ANKHD1-EIF4EBP3 |
| 5 | 140453969 | 140454118 | 3.656441478 | ANKHD1 |
| 5 | 140631938 | 140632129 | -1.146827425 | CD14 |
| 1 | 203305619 | 203305810 | 3.505594388 | BTG2 |
| 5 | 141100544 | 141100721 | -4.811133598 | PCDHB3 |
| 5 | 141100544 | 141100721 | -4.811133598 | AC005754.7 |
| 5 | 141101608 | 141101807 | -1.287567744 | AC005754.7 |
| 5 | 141101608 | 141101807 | -1.287567744 | PCDHB3 |
| 5 | 141122150 | 141122346 | -1.855989455 | AC005754.7 |
| 5 | 141122150 | 141122346 | -1.855989455 | PCDHB4 |
| 5 | 141122722 | 141122897 | -3.052808777 | AC005754.7 |
| 5 | 141122722 | 141122897 | -3.052808777 | PCDHB4 |
| 5 | 141123197 | 141123436 | -1.288250461 | AC005754.7 |
| 5 | 141123197 | 141123436 | -1.288250461 | PCDHB4 |
| 5 | 141124133 | 141124325 | -4.093428534 | AC005754.7 |
| 5 | 141124133 | 141124325 | -4.093428534 | PCDHB4 |
| 5 | 141135812 | 141135965 | -2.190654687 | PCDHB5 |
| 5 | 141135812 | 141135965 | -2.190654687 | AC005754.7 |
| 5 | 141136531 | 141136871 | -1.25889982 | CH17-140K24.2 |
| 5 | 141136531 | 141136871 | -1.25889982 | AC005754.7 |
| 5 | 141136531 | 141136871 | -1.25889982 | PCDHB5 |
| 5 | 141137627 | 141137844 | -1.658127583 | PCDHB5 |
| 5 | 141137627 | 141137844 | -1.658127583 | AC005754.7 |
| 5 | 141137627 | 141137844 | -1.658127583 | CH17-140K24.2 |
| 5 | 141150167 | 141150433 | -2.299099164 | AC005754.7 |
| 5 | 141150167 | 141150433 | -2.299099164 | PCDHB6 |
| 5 | 141150167 | 141150433 | -2.299099164 | CH17-140K24.2 |
| 1 | 203307287 | 203307504 | 3.023214553 | BTG2 |
| 5 | 141151233 | 141151577 | -1.007547283 | CH17-140K24.2 |
| 5 | 141151233 | 141151577 | -1.007547283 | PCDHB6 |
| 5 | 141151233 | 141151577 | -1.007547283 | AC005754.7 |
| 5 | 141157246 | 141157397 | -2.204449442 | CH17-140K24.2 |
| 7 | 932345 | 932554 | 1.383516189 | ADAP1 |
| 7 | 1026297 | 1026476 | 9.926167216 | C7orf50 |
| 7 | 1045849 | 1046051 | 3.43219405 | GPR146 |
| 7 | 1045849 | 1046051 | 3.43219405 | C7orf50 |
| 7 | 1047942 | 1048131 | 7.933735903 | C7orf50 |
| 7 | 1047942 | 1048131 | 7.933735903 | GPR146 |
| 7 | 1050865 | 1051051 | 6.241403978 | GPR146 |
| 7 | 1050865 | 1051051 | 6.241403978 | C7orf50 |
| 1 | 225497567 | 225497767 | 2.81167657 | ENAH |
| 7 | 1058339 | 1058618 | 3.363175696 | C7orf50 |
| 7 | 1058339 | 1058618 | 3.363175696 | RP11-449P15.1 |
| 7 | 1058339 | 1058618 | 3.363175696 | GPR146 |
| 7 | 1434416 | 1434653 | 1.899455589 | MICALL2 |
| 7 | 1585082 | 1585289 | 2.925917899 | PSMG3-AS1 |
| 7 | 1586419 | 1586618 | 1.995092993 | PSMG3-AS1 |
| 7 | 1587066 | 1587258 | 3.164649199 | PSMG3-AS1 |
| 7 | 1587772 | 1587972 | 1.37114053 | PSMG3-AS1 |
| 7 | 1841797 | 1841985 | 1.726586314 | MAD1L1 |
| 7 | 1841797 | 1841985 | 1.726586314 | AC110781.3 |
| 7 | 1842326 | 1842509 | 6.999691741 | AC110781.3 |
| 7 | 1842326 | 1842509 | 6.999691741 | MAD1L1 |
| 7 | 1845297 | 1845484 | 5.569455859 | AC110781.3 |
| 7 | 1845297 | 1845484 | 5.569455859 | MAD1L1 |
| 7 | 1968309 | 1968508 | 8.07101363 | MAD1L1 |
| 7 | 2110051 | 2110266 | 2.287269431 | MAD1L1 |
| 1 | 225846738 | 225846938 | -2.513881067 | TMEM63A |
| 7 | 2254621 | 2254817 | 1.375886105 | SNX8 |
| 7 | 2354256 | 2354435 | 2.176994652 | SNX8 |
| 7 | 2354256 | 2354435 | 2.176994652 | EIF3B |
| 7 | 2432701 | 2432907 | 8.609768621 | CHST12 |
| 7 | 2443508 | 2443694 | -1.875985009 | CHST12 |
| 7 | 2446138 | 2446351 | -2.504086966 | CHST12 |
| 7 | 2538422 | 2538594 | 1.003521732 | BRAT1 |
| 7 | 2662085 | 2662289 | 1.050187254 | TTYH3 |
| 7 | 2721957 | 2722153 | -1.713085121 | AMZ1 |
| 7 | 4267833 | 4267999 | -5.52117169 | SDK1 |
| 7 | 4797555 | 4797765 | -1.108380277 | RADIL |
| 7 | 4877676 | 4877884 | 3.908473101 | RADIL |
| 7 | 5010018 | 5010124 | -6.310339021 | RBAK-RBAKDN |
| 7 | 5010018 | 5010124 | -6.310339021 | RNF216P1 |
| 7 | 5067063 | 5067228 | -1.095617754 | RBAK |
| 7 | 5067063 | 5067228 | -1.095617754 | RBAK-RBAKDN |
| 7 | 5071676 | 5071795 | -1.874679685 | RBAK-RBAKDN |
| 7 | 5313182 | 5313385 | 5.04646311 | TNRC18 |
| 7 | 5370682 | 5370931 | 1.297435725 | TNRC18 |
| 7 | 5377970 | 5378172 | 1.551227632 | TNRC18 |
| 1 | 226309759 | 226309879 | 1.759691307 | LIN9 |
| 7 | 5387952 | 5388155 | 2.44365712 | TNRC18 |
| 7 | 5390565 | 5390743 | -1.30436147 | TNRC18 |
| 7 | 5429106 | 5429306 | -2.248936647 | RP11-1275H24.3 |
| 7 | 5481567 | 5481767 | 1.339563271 | FBXL18 |
| 7 | 5495495 | 5495606 | 2.06935424 | FBXL18 |
| 7 | 5500396 | 5500678 | 1.77718709 | FBXL18 |
| 7 | 5501177 | 5501373 | 1.330839359 | FBXL18 |

| | | | | |
|---|---|---|---|---|
| 1 | 200849784 | 200849970 | 2.589778942 | CAMSAP2 |
| 5 | 81238670 | 81238846 | -3.126103802 | CKMT2-AS1 |
| 5 | 81238670 | 81238846 | -3.126103802 | CKMT2 |
| 5 | 81699310 | 81699499 | 2.179857889 | SSBP2 |
| 5 | 82001568 | 82001719 | -1.15866364 | ATG10 |
| 5 | 82084976 | 82085159 | -1.599227728 | ATG10 |
| 5 | 87268654 | 87268863 | 2.003422458 | RASA1 |
| 1 | 200971457 | 200971656 | 1.190631258 | KIF21B |
| 5 | 87305411 | 87305580 | 2.454399936 | RASA1 |
| 5 | 87306174 | 87306368 | 3.264055464 | RASA1 |
| 5 | 87355731 | 87355884 | -5.658140766 | RASA1 |
| 5 | 88190446 | 88190649 | -1.593589533 | TMEM161B |
| 5 | 88667038 | 88667237 | -1.728750796 | LINC00461 |
| 5 | 88794676 | 88794876 | -1.695368252 | MEF2C |
| 5 | 88797290 | 88797490 | -1.511410442 | MEF2C |
| 5 | 88798467 | 88798607 | -6.547368753 | MEF2C |
| 5 | 90461409 | 90461614 | -1.963886982 | MBLAC2 |
| 5 | 90473807 | 90473933 | -2.335340383 | MBLAC2 |
| 5 | 90473807 | 90473933 | -2.335340383 | POLR3G |
| 5 | 90519022 | 90519224 | -1.432003744 | LYSMD3 |
| 5 | 91370761 | 91370981 | 1.122818969 | ARRDC3 |
| 5 | 93506046 | 93506220 | 1.246888987 | NR2F1-AS1 |
| 5 | 94695149 | 94695415 | -3.208876219 | SLF1 |
| 5 | 95556347 | 95556459 | -2.380617304 | ARSK |
| 1 | 201520320 | 201520525 | -2.059393076 | RP11-134G8.7 |
| 1 | 201520320 | 201520525 | -2.059393076 | RP11-134G8.5 |
| 1 | 201520320 | 201520525 | -2.059393076 | RP11-134G8.6 |
| 5 | 95620230 | 95620418 | -2.395865862 | GPR150 |
| 5 | 95898356 | 95898593 | 2.924111437 | ELL2 |
| 5 | 98779677 | 98779873 | 2.539208851 | RGMB |
| 5 | 98793371 | 98793547 | 2.046042199 | RGMB |
| 5 | 98793371 | 98793547 | 2.046042199 | AC008522.1 |
| 1 | 201623147 | 201623484 | 1.557530055 | NAV1 |
| 5 | 103129413 | 103129604 | -1.694062896 | PPIP5K2 |
| 5 | 103276078 | 103276242 | -1.892328018 | C5orf30 |
| 5 | 103559276 | 103559465 | -1.502432878 | NUDT12 |
| 1 | 201782585 | 201782788 | 2.850219859 | IPO9-AS1 |
| 1 | 201782585 | 201782788 | 2.850219859 | NAV1 |
| 5 | 108208401 | 108208526 | 1.903677461 | FBXL17 |
| 5 | 110566518 | 110566641 | 1.226858984 | TMEM232 |
| 5 | 111484244 | 111484438 | 3.256271478 | CAMK4 |
| 5 | 111499885 | 111500083 | 1.117729476 | STARD4 |
| 1 | 201820054 | 201820265 | 1.592298747 | NAV1 |
| 1 | 201820054 | 201820265 | 1.592298747 | IPO9-AS1 |
| 5 | 112984179 | 112984388 | -1.129208321 | DCP2 |
| 5 | 114404562 | 114404761 | -2.313603859 | KCNN2 |
| 5 | 114482271 | 114482401 | -1.408551164 | KCNN2 |
| 5 | 115126490 | 115126690 | -1.87717705 | TRIM36 |
| 5 | 115580678 | 115580877 | 1.001922716 | TMED7-TICAM2 |
| 5 | 115580678 | 115580877 | 1.001922716 | TICAM2 |
| 1 | 201876108 | 201876320 | -1.872592107 | IPO9 |
| 5 | 116446324 | 116446523 | -8.371996431 | SEMA6A |
| 5 | 120686264 | 120686482 | 9.102896625 | PRR16 |
| 1 | 201877769 | 201877967 | -1.783558051 | IPO9 |
| 5 | 141157246 | 141157397 | -2.204449442 | AC005754.7 |
| 5 | 141157246 | 141157397 | -2.204449442 | PCDHB17P |
| 5 | 141173795 | 141173992 | -2 | AC005754.7 |
| 5 | 141173795 | 141173992 | -2 | CH17-140K24.2 |
| 5 | 141173795 | 141173992 | -2 | PCDHB7 |
| 5 | 141178382 | 141178591 | -1.927541749 | CH17-140K24.2 |
| 5 | 141178382 | 141178591 | -1.927541749 | PCDHB8 |
| 5 | 141178382 | 141178591 | -1.927541749 | CH17-140K24.4 |
| 5 | 141179163 | 141179478 | -7.312740275 | CH17-140K24.2 |
| 5 | 141179163 | 141179478 | -7.312740275 | CH17-140K24.4 |
| 5 | 141179163 | 141179478 | -7.312740275 | PCDHB8 |
| 5 | 141188447 | 141188636 | 1.330452666 | CH17-140K24.2 |
| 5 | 141188447 | 141188636 | 1.330452666 | CH17-140K24.5 |
| 5 | 141188447 | 141188636 | 1.330452666 | CH17-140K24.7 |
| 5 | 141188447 | 141188636 | 1.330452666 | PCDHB9 |
| 5 | 141189487 | 141189648 | 3.24500115 | CH17-140K24.7 |
| 5 | 141189487 | 141189648 | 3.24500115 | CH17-140K24.2 |
| 5 | 141189487 | 141189648 | 3.24500115 | PCDHB9 |
| 1 | 9356281 | 9356483 | 1.058337851 | SPSB1 |
| 5 | 141199772 | 141199961 | -6.720434584 | CH17-140K24.2 |
| 5 | 141199772 | 141199961 | -6.720434584 | PCDHB11 |
| 5 | 141199772 | 141199961 | -6.720434584 | CH17-140K24.7 |
| 5 | 141208881 | 141209009 | -1.644284749 | CH17-140K24.2 |
| 5 | 141208881 | 141209009 | -1.644284749 | PCDHB12 |
| 5 | 141210034 | 141210194 | -1.162375443 | CH17-140K24.2 |
| 5 | 141210034 | 141210194 | -1.162375443 | PCDHB12 |
| 5 | 141215235 | 141215462 | -1.02070135 | PCDHB13 |
| 5 | 141215235 | 141215462 | -1.02070135 | CH17-140K24.2 |
| 5 | 141421580 | 141421743 | -1.913320986 | PCDHGB5 |
| 5 | 141421580 | 141421743 | -1.913320986 | PCDHGA7 |
| 5 | 141421580 | 141421743 | -1.913320986 | PCDHGA8 |
| 5 | 141421580 | 141421743 | -1.913320986 | PCDHGA2 |
| 5 | 141421580 | 141421743 | -1.913320986 | PCDHGB2 |
| 5 | 141421580 | 141421743 | -1.913320986 | PCDHGA4 |
| 5 | 141421580 | 141421743 | -1.913320986 | PCDHGB1 |
| 5 | 141421580 | 141421743 | -1.913320986 | PCDHGA5 |
| 5 | 141421580 | 141421743 | -1.913320986 | PCDHGB7 |
| 5 | 141421580 | 141421743 | -1.913320986 | PCDHGA10 |
| 5 | 141421580 | 141421743 | -1.913320986 | PCDHGB6 |
| 5 | 141421580 | 141421743 | -1.913320986 | PCDHGB4 |
| 5 | 141421580 | 141421743 | -1.913320986 | PCDHGB3 |
| 5 | 141421580 | 141421743 | -1.913320986 | PCDHGA3 |
| 5 | 141421580 | 141421743 | -1.913320986 | PCDHGA6 |
| 5 | 141421580 | 141421743 | -1.913320986 | PCDHGA9 |
| 5 | 141421580 | 141421743 | -1.913320986 | PCDHGA11 |
| 5 | 141421580 | 141421743 | -1.913320986 | PCDHGA1 |
| 5 | 141432497 | 141432655 | -2.114613954 | PCDHGB7 |
| 5 | 141432497 | 141432655 | -2.114613954 | PCDHGB5 |
| 5 | 141432497 | 141432655 | -2.114613954 | PCDHGA5 |
| 5 | 141432497 | 141432655 | -2.114613954 | PCDHGB2 |
| 5 | 141432497 | 141432655 | -2.114613954 | PCDHGA12 |
| 5 | 141432497 | 141432655 | -2.114613954 | PCDHGB1 |
| 5 | 141432497 | 141432655 | -2.114613954 | PCDHGA9 |
| 5 | 141432497 | 141432655 | -2.114613954 | PCDHGA6 |
| 7 | 5527630 | 5527831 | 1.422004099 | ACTB |
| 7 | 5847305 | 5847431 | -4.75389732 | ZNF815P |
| 7 | 5973211 | 5973384 | 1.549870076 | PMS2 |
| 7 | 6653196 | 6653327 | 9.456235644 | ZNF316 |
| 7 | 6676041 | 6676234 | -1.16018379 | AC073343.13 |
| 7 | 6676041 | 6676234 | -1.16018379 | RP11-611L7.2 |
| 1 | 10639761 | 10639957 | 1.381294955 | CASZ1 |
| 1 | 10639761 | 10639957 | 1.381294955 | RP4-734G22.3 |
| 7 | 6794141 | 6794331 | -1.463824578 | CCZ1B |
| 7 | 6794141 | 6794331 | -1.463824578 | RSPH10B2 |
| 7 | 7720986 | 7721127 | 6.062060656 | RP11-505D17.1 |
| 7 | 7720986 | 7721127 | 6.062060656 | UMAD1 |
| 7 | 11348814 | 11348972 | -2.107286458 | AC004538.3 |
| 7 | 11351495 | 11351691 | -1.479731852 | AC004538.3 |
| 7 | 12370089 | 12370277 | -2.523990391 | VWDE |
| 7 | 12491330 | 12491491 | 1.752574959 | AC005281.1 |
| 7 | 12491330 | 12491491 | 1.752574959 | TAS2R2P |
| 7 | 12499192 | 12499377 | -7.511097201 | AC005281.1 |
| 7 | 22882053 | 22882237 | -4.86904613 | RPL12P10 |
| 7 | 24882819 | 24883007 | 1.196349873 | OSBPL3 |
| 7 | 25135997 | 25136149 | 1.043344867 | C7orf31 |
| 1 | 226928022 | 226928220 | -2.396994719 | COQ8A |
| 7 | 26163915 | 26164123 | -2.531373267 | NFE2L3 |
| 7 | 26164642 | 26164833 | -4.981204412 | NFE2L3 |
| 7 | 26165713 | 26165881 | -6.831556654 | NFE2L3 |
| 7 | 27176020 | 27176222 | 4.418682562 | HOXA10 |
| 7 | 27176020 | 27176222 | 4.418682562 | HOXA10-HOXA9 |
| 7 | 27204924 | 27205127 | 1.360580958 | HOTTIP |
| 7 | 27840972 | 27841073 | -2.327451598 | TAX1BP1 |
| 7 | 27840972 | 27841073 | -2.327451598 | JAZF1 |
| 7 | 30403665 | 30403878 | -2.974901199 | GS1-114I9.3 |
| 7 | 30403665 | 30403878 | -2.974901199 | LINC01176 |
| 7 | 30535648 | 30535845 | -4.836212671 | RP4-777O23.3 |
| 7 | 30535648 | 30535845 | -4.836212671 | AC005154.6 |
| 7 | 30633714 | 30633914 | -1.209678822 | GARS |
| 7 | 32689040 | 32689159 | 4.261780978 | DPY19L1P1 |
| 7 | 35697220 | 35697443 | 1.158847171 | RP11-379H18.1 |
| 1 | 10653750 | 10653963 | 2.816978822 | RP4-734G22.3 |
| 1 | 10653750 | 10653963 | 2.816978822 | CASZ1 |
| 1 | 227286247 | 227286421 | -1.745162813 | CDC42BPA |
| 7 | 38203671 | 38203871 | -5.938642067 | STARD3NL |
| 7 | 38327365 | 38327554 | 5.346743479 | RP11-121A8.1 |
| 7 | 38354613 | 38354812 | 6.394359166 | TRG-AS1 |
| 7 | 39950229 | 39950433 | 10.48963723 | CDK13 |
| 7 | 39951466 | 39951694 | 10.47056967 | CDK13 |
| 7 | 41690207 | 41690410 | 1.573396354 | INHBA |
| 7 | 41691341 | 41691525 | -3.63496088 | INHBA |
| 7 | 43609179 | 43609382 | -6.718746005 | COA1 |
| 7 | 43609179 | 43609382 | -6.718746005 | STK17A |
| 7 | 43944473 | 43944642 | -1.465185103 | POLR2J4 |
| 7 | 43944473 | 43944642 | -1.465185103 | UBE2D4 |
| 7 | 44006622 | 44006818 | 1.879735662 | POLR2J4 |
| 7 | 44006622 | 44006818 | 1.879735662 | AC004951.6 |
| 7 | 44006622 | 44006818 | 1.879735662 | SPDYE1 |

| | | | | |
|---|---|---|---|---|
| 5 | 122078111 | 122078268 | -4.729715809 | LOX |
| 5 | 122152018 | 122152192 | -1.418971621 | CTC-441N14.4 |
| 5 | 122152018 | 122152192 | -1.418971621 | CTC-441N14.2 |
| 5 | 122152018 | 122152192 | -1.418971621 | ZNF474 |
| 5 | 122451247 | 122451426 | 1.105808975 | SNCAIP |
| 5 | 122451247 | 122451426 | 1.105808975 | MGC32805 |
| 5 | 123037180 | 123037331 | -7.147202681 | RP11-359P5.1 |
| 5 | 123087426 | 123087598 | -2.707415079 | AC106786.1 |
| 1 | 201878951 | 201879147 | -2.186507541 | IPO9 |
| 5 | 123346588 | 123346803 | -1.060553457 | CEP120 |
| 5 | 123401854 | 123402005 | -1.162144437 | CEP120 |
| 5 | 123401854 | 123402005 | -1.162144437 | KRT8P33 |
| 5 | 123552947 | 123553117 | 1.231371754 | CSNK1G3 |
| 5 | 124744776 | 124744972 | -1.575173439 | ZNF608 |
| 5 | 126890638 | 126890828 | -1.26586024 | MARCH3 |
| 5 | 127054715 | 127054912 | -3.212553679 | C5orf63 |
| 5 | 130184692 | 130184823 | 2.625764875 | CHSY3 |
| 5 | 130185277 | 130185471 | 2.292081457 | CHSY3 |
| 5 | 131427031 | 131427249 | -1.006635262 | RAPGEF6 |
| 5 | 131427031 | 131427249 | -1.006635262 | CTC-432M15.3 |
| 5 | 131429100 | 131429301 | -1.15112246 | RAPGEF6 |
| 5 | 131429100 | 131429301 | -1.15112246 | CTC-432M15.3 |
| 5 | 131430998 | 131431198 | -1.16644248 | CTC-432M15.3 |
| 5 | 131430998 | 131431198 | -1.16644248 | RAPGEF6 |
| 5 | 131723970 | 131724094 | 6.369890305 | FNIP1 |
| 5 | 131723970 | 131724094 | 6.369890305 | CTC-432M15.3 |
| 1 | 202147817 | 202148014 | -1.537095143 | PTPN7 |
| 5 | 131727526 | 131727720 | 1.244194254 | FNIP1 |
| 5 | 131727526 | 131727720 | 1.244194254 | CTC-432M15.3 |
| 5 | 132294485 | 132294606 | -3.408214522 | SLC22A4 |
| 5 | 132294485 | 132294606 | -3.408214522 | P4HA2 |
| 5 | 132294953 | 132295132 | -2.042750662 | SLC22A4 |
| 5 | 132294953 | 132295132 | -2.042750662 | P4HA2 |
| 5 | 132349302 | 132349442 | 1.97320948 | AC034220.3 |
| 5 | 132470118 | 132470308 | -1.256612218 | AC116366.6 |
| 5 | 132470118 | 132470308 | -1.256612218 | C5orf56 |
| 5 | 132881054 | 132881252 | 1.601589073 | AFF4 |
| 5 | 134286251 | 134286359 | -1.114053036 | CDKL3 |
| 5 | 134286251 | 134286359 | -1.114053036 | CTD-2410N18.4 |
| 5 | 134578706 | 134578982 | -1.057513568 | JADE2 |
| 1 | 202853021 | 202853169 | -5.187315993 | RP11-480I12.5 |
| 5 | 135028486 | 135028688 | -1.597526921 | PITX1 |
| 5 | 135033291 | 135033530 | -2.235475003 | PITX1 |
| 5 | 135033291 | 135033530 | -2.235475003 | C5orf66 |
| 5 | 135444307 | 135444516 | -1.263703748 | TIFAB |
| 5 | 135444307 | 135444516 | -1.263703748 | DCANP1 |
| 5 | 135446722 | 135446916 | -1.631135122 | DCANP1 |
| 5 | 135446722 | 135446916 | -1.631135122 | TIFAB |
| 5 | 135449269 | 135449495 | -1.49152088 | TIFAB |
| 5 | 137753771 | 137753966 | -1.31848871 | HNRNPA0 |
| 5 | 137939504 | 137939798 | -1.249543287 | PKD2L2 |
| 5 | 137939504 | 137939798 | -1.249543287 | FAM13B |
| 5 | 138164864 | 138165079 | -1.360336491 | BRD8 |
| 1 | 202880949 | 202881167 | -1.308311368 | RABIF |
| 5 | 141432497 | 141432655 | -2.114613954 | PCDHGA1 |
| 5 | 141432497 | 141432655 | -2.114613954 | PCDHGA11 |
| 5 | 141432497 | 141432655 | -2.114613954 | PCDHGB6 |
| 5 | 141432497 | 141432655 | -2.114613954 | PCDHGA4 |
| 5 | 141432497 | 141432655 | -2.114613954 | PCDHGB3 |
| 5 | 141432497 | 141432655 | -2.114613954 | PCDHGA10 |
| 5 | 141432497 | 141432655 | -2.114613954 | PCDHGA2 |
| 5 | 141432497 | 141432655 | -2.114613954 | PCDHGB4 |
| 5 | 141432497 | 141432655 | -2.114613954 | PCDHGA3 |
| 5 | 141432497 | 141432655 | -2.114613954 | PCDHGA7 |
| 5 | 141432497 | 141432655 | -2.114613954 | PCDHGA8 |
| 5 | 141475982 | 141476185 | -1.623236924 | PCDHGA1 |
| 5 | 141475982 | 141476185 | -1.623236924 | PCDHGB3 |
| 5 | 141475982 | 141476185 | -1.623236924 | PCDHGB5 |
| 5 | 141475982 | 141476185 | -1.623236924 | PCDHGA7 |
| 5 | 141475982 | 141476185 | -1.623236924 | PCDHGA2 |
| 5 | 141475982 | 141476185 | -1.623236924 | PCDHGB7 |
| 5 | 141475982 | 141476185 | -1.623236924 | PCDHGA10 |
| 5 | 141475982 | 141476185 | -1.623236924 | PCDHGC3 |
| 5 | 141475982 | 141476185 | -1.623236924 | PCDHGA12 |
| 5 | 141475982 | 141476185 | -1.623236924 | PCDHGA8 |
| 5 | 141475982 | 141476185 | -1.623236924 | PCDHGA11 |
| 5 | 141475982 | 141476185 | -1.623236924 | PCDHGB1 |
| 5 | 141475982 | 141476185 | -1.623236924 | PCDHGB4 |
| 5 | 141475982 | 141476185 | -1.623236924 | PCDHGA6 |
| 5 | 141475982 | 141476185 | -1.623236924 | PCDHGA3 |
| 5 | 141475982 | 141476185 | -1.623236924 | PCDHGA4 |
| 5 | 141475982 | 141476185 | -1.623236924 | PCDHGA9 |
| 5 | 141475982 | 141476185 | -1.623236924 | PCDHGA5 |
| 5 | 141475982 | 141476185 | -1.623236924 | PCDHGB6 |
| 5 | 141475982 | 141476185 | -1.623236924 | PCDHGB2 |
| 5 | 141477595 | 141477767 | -1.52656201 | PCDHGB6 |
| 5 | 141477595 | 141477767 | -1.52656201 | PCDHGA5 |
| 5 | 141477595 | 141477767 | -1.52656201 | PCDHGA3 |
| 5 | 141477595 | 141477767 | -1.52656201 | PCDHGA2 |
| 5 | 141477595 | 141477767 | -1.52656201 | PCDHGA11 |
| 5 | 141477595 | 141477767 | -1.52656201 | PCDHGA7 |
| 5 | 141477595 | 141477767 | -1.52656201 | PCDHGB3 |
| 5 | 141477595 | 141477767 | -1.52656201 | PCDHGB1 |
| 5 | 141477595 | 141477767 | -1.52656201 | PCDHGB5 |
| 5 | 141477595 | 141477767 | -1.52656201 | PCDHGA9 |
| 5 | 141477595 | 141477767 | -1.52656201 | PCDHGB4 |
| 5 | 141477595 | 141477767 | -1.52656201 | PCDHGA10 |
| 5 | 141477595 | 141477767 | -1.52656201 | PCDHGA12 |
| 5 | 141477595 | 141477767 | -1.52656201 | PCDHGA1 |
| 5 | 141477595 | 141477767 | -1.52656201 | PCDHGA4 |
| 5 | 141477595 | 141477767 | -1.52656201 | PCDHGC3 |
| 5 | 141477595 | 141477767 | -1.52656201 | PCDHGB7 |
| 5 | 141477595 | 141477767 | -1.52656201 | PCDHGA8 |
| 5 | 141477595 | 141477767 | -1.52656201 | PCDHGB2 |
| 5 | 141477595 | 141477767 | -1.52656201 | PCDHGA6 |
| 5 | 141478245 | 141478373 | -1.406978088 | PCDHGB1 |
| 5 | 141478245 | 141478373 | -1.406978088 | PCDHGA2 |
| 5 | 141478245 | 141478373 | -1.406978088 | PCDHGB5 |
| 7 | 44006622 | 44006818 | 1.879735662 | RP5-1165K10.2 |
| 7 | 44060919 | 44061117 | 2.448855513 | DBNL |
| 1 | 227669098 | 227669338 | -1.132854996 | ZNF678 |
| 7 | 45046633 | 45046818 | -2.170393561 | CCM2 |
| 7 | 45913477 | 45913680 | 1.556374373 | IGFBP3 |
| 7 | 47303249 | 47303392 | 10.7459422 | TNS3 |
| 7 | 50592704 | 50592910 | -1.408842259 | GRB10 |
| 1 | 227701827 | 227702030 | -1.430174005 | ZNF847P |
| 7 | 55543949 | 55544149 | -5.885744735 | VOPP1 |
| 1 | 227732451 | 227732665 | 1.14927518 | SNAP47 |
| 1 | 227732451 | 227732665 | 1.14927518 | JMJD4 |
| 1 | 227734720 | 227734927 | 1.372952807 | SNAP47 |
| 1 | 227734720 | 227734927 | 1.372952807 | JMJD4 |
| 7 | 64710006 | 64710201 | 1.19473669 | ZNF107 |
| 7 | 65200603 | 65200794 | -1.169018256 | INTS4P1 |
| 7 | 65214850 | 65215050 | -1.791191978 | INTS4P1 |
| 7 | 66359307 | 66359480 | -1.014099469 | TPST1 |
| 7 | 66653560 | 66653771 | 1.618936924 | KCTD7 |
| 7 | 66682427 | 66682593 | 8.193234394 | RABGEF1 |
| 7 | 66682427 | 66682593 | 8.193234394 | KCTD7 |
| 7 | 72913073 | 72913275 | 1.401597757 | POM121 |
| 7 | 73573475 | 73573634 | 6.652460835 | TBL2 |
| 7 | 73682750 | 73682874 | 1.080067786 | DNAJC30 |
| 7 | 74883136 | 74883247 | 6.695084478 | STAG3L2 |
| 7 | 75415778 | 75415971 | -1.337044196 | NSUN5P1 |
| 7 | 75501100 | 75501218 | -6.614784949 | SPDYE5 |
| 7 | 76072809 | 76072937 | 3.904964433 | AC005077.14 |
| 1 | 228212760 | 228212948 | 2.879666704 | OBSCN |
| 1 | 228212760 | 228212948 | 2.879666704 | C1orf145 |
| 7 | 76482733 | 76482932 | 1.3182004 | DTX2 |
| 7 | 76980907 | 76981112 | 1.249603885 | DTX2P1 |
| 7 | 76980907 | 76981112 | 1.249603885 | DTX2P1-UPK3BP1-PMS2P11 |
| 7 | 77196728 | 77196881 | 3.009463528 | CCDC146 |
| 7 | 77196728 | 77196881 | 3.009463528 | RP11-467H10.2 |
| 7 | 77196728 | 77196881 | 3.009463528 | FGL2 |
| 1 | 228274489 | 228274698 | 1.421743368 | OBSCN |
| 1 | 228274489 | 228274698 | 1.421743368 | RP5-1139B12.3 |
| 7 | 77199482 | 77199682 | 2.798889981 | CCDC146 |
| 7 | 77199482 | 77199682 | 2.798889981 | FGL2 |
| 7 | 77696090 | 77696269 | 2.333175414 | APTR |
| 7 | 78018898 | 78019099 | -1.025593675 | MAGI2 |
| 1 | 228396561 | 228396757 | 2.042414737 | RP11-245P10.8 |
| 1 | 228396561 | 228396757 | 2.042414737 | TRIM11 |
| 7 | 83989715 | 83989903 | -1.305717356 | SEMA3A |
| 7 | 84999592 | 84999793 | 1.982351074 | SEMA3D |
| 1 | 228410535 | 228410687 | 1.036828521 | TRIM17 |
| 7 | 91175499 | 91175695 | -6.736789462 | CDK14 |
| 7 | 91265751 | 91265975 | 2.094088353 | FZD1 |
| 7 | 91266673 | 91266860 | 1.227238437 | FZD1 |
| 7 | 92833614 | 92833812 | 7.700172898 | CDK6 |
| 7 | 96274393 | 96274547 | -1.340191033 | SLC25A13 |
| 7 | 97938440 | 97938632 | -2.580863239 | AC079781.8 |
| 7 | 98207243 | 98207414 | 4.267637688 | LMTK2 |
| 7 | 98302485 | 98302746 | -1.214282316 | BAIAP2L1 |

| | | | | |
|---|---|---|---|---|
| 5 | 138386303 | 138386505 | -1.778291304 | KDM3B |
| 5 | 138391223 | 138391432 | -1.419496991 | KDM3B |
| 5 | 138391957 | 138392166 | -1.431242757 | KDM3B |
| 5 | 138393055 | 138393249 | -1.439715364 | KDM3B |
| 5 | 138467094 | 138467252 | 2.371133551 | EGR1 |
| 5 | 138467950 | 138468140 | 1.853364787 | EGR1 |
| 5 | 138947085 | 138947286 | 1.200798844 | SIL1 |
| 5 | 139009296 | 139009434 | -7.540658495 | SIL1 |
| 1 | 203064544 | 203064715 | 1.886285584 | PPFIA4 |
| 5 | 139325341 | 139325523 | -7.30453164 | MATR3 |
| 5 | 139393817 | 139393932 | -2.776225546 | PROB1 |
| 5 | 139399240 | 139399471 | -1.846333067 | SPATA24 |
| 5 | 139475649 | 139475835 | -3.062525251 | TMEM173 |
| 5 | 140109215 | 140109404 | -1.860340255 | PURA |
| 5 | 140333839 | 140334039 | 3.400180627 | HBEGF |
| 5 | 140401005 | 140401210 | -1.405289411 | CTC-329D1.2 |
| 5 | 140453969 | 140454118 | -4.329551982 | ANKHD1-EIF4EBP3 |
| 5 | 140453969 | 140454118 | -4.329551982 | ANKHD1 |
| 5 | 140456725 | 140456987 | -5.684798673 | ANKHD1-EIF4EBP3 |
| 5 | 140456725 | 140456987 | -5.684798673 | ANKHD1 |
| 5 | 140457271 | 140457432 | -6.808044657 | ANKHD1-EIF4EBP3 |
| 5 | 140457271 | 140457432 | -6.808044657 | ANKHD1 |
| 1 | 203179036 | 203179237 | -12.77999734 | CHI3L1 |
| 5 | 140526145 | 140526311 | -1.718361562 | ANKHD1-EIF4EBP3 |
| 5 | 140526145 | 140526311 | -1.718361562 | ANKHD1 |
| 5 | 140529175 | 140529373 | -1.250469857 | ANKHD1-EIF4EBP3 |
| 5 | 140529175 | 140529373 | -1.250469857 | ANKHD1 |
| 5 | 140550167 | 140550388 | -1.426146256 | SRA1 |
| 5 | 140631938 | 140632129 | 4.160131243 | CD14 |
| 5 | 140670810 | 140670988 | -1.756347321 | WDR55 |
| 5 | 140670810 | 140670988 | -1.756347321 | DND1 |
| 5 | 140698691 | 140698893 | -1.175704002 | HARS2 |
| 5 | 140698691 | 140698893 | -1.175704002 | ZMAT2 |
| 5 | 140969027 | 140969230 | -1.243069723 | PCDHA1 |
| 5 | 140969027 | 140969230 | -1.243069723 | PCDHAC2 |
| 5 | 140969027 | 140969230 | -1.243069723 | PCDHA2 |
| 5 | 140969027 | 140969230 | -1.243069723 | PCDHA9 |
| 5 | 140969027 | 140969230 | -1.243069723 | PCDHA5 |
| 5 | 140969027 | 140969230 | -1.243069723 | PCDHAC1 |
| 5 | 140969027 | 140969230 | -1.243069723 | PCDHA4 |
| 5 | 140969027 | 140969230 | -1.243069723 | PCDHA13 |
| 5 | 140969027 | 140969230 | -1.243069723 | PCDHA12 |
| 5 | 140969027 | 140969230 | -1.243069723 | PCDHA3 |
| 5 | 140969027 | 140969230 | -1.243069723 | PCDHA8 |
| 5 | 140969027 | 140969230 | -1.243069723 | PCDHA10 |
| 5 | 140969027 | 140969230 | -1.243069723 | PCDHA6 |
| 5 | 140969027 | 140969230 | -1.243069723 | PCDHA7 |
| 5 | 140969027 | 140969230 | -1.243069723 | PCDHA11 |
| 1 | 203305619 | 203305810 | 2.537095064 | BTG2 |
| 1 | 203307287 | 203307504 | 3.030246392 | BTG2 |
| 5 | 141151233 | 141151577 | -1.286394185 | CH17-140K24.2 |
| 5 | 141151233 | 141151577 | -1.286394185 | AC005754.7 |
| 5 | 141151233 | 141151577 | -1.286394185 | PCDHB6 |
| 5 | 141157246 | 141157397 | 1.277745691 | CH17-140K24.2 |
| 5 | 141478245 | 141478373 | -1.406978088 | PCDHGA1 |
| 5 | 141478245 | 141478373 | -1.406978088 | PCDHGA8 |
| 5 | 141478245 | 141478373 | -1.406978088 | PCDHGB3 |
| 5 | 141478245 | 141478373 | -1.406978088 | PCDHGC3 |
| 5 | 141478245 | 141478373 | -1.406978088 | PCDHGB4 |
| 5 | 141478245 | 141478373 | -1.406978088 | PCDHGB2 |
| 5 | 141478245 | 141478373 | -1.406978088 | PCDHGA3 |
| 5 | 141478245 | 141478373 | -1.406978088 | PCDHGA4 |
| 5 | 141478245 | 141478373 | -1.406978088 | PCDHGA10 |
| 5 | 141478245 | 141478373 | -1.406978088 | PCDHGA9 |
| 5 | 141478245 | 141478373 | -1.406978088 | PCDHGB7 |
| 5 | 141478245 | 141478373 | -1.406978088 | PCDHGA5 |
| 5 | 141478245 | 141478373 | -1.406978088 | PCDHGA12 |
| 5 | 141478245 | 141478373 | -1.406978088 | PCDHGB6 |
| 5 | 141478245 | 141478373 | -1.406978088 | PCDHGA11 |
| 5 | 141478245 | 141478373 | -1.406978088 | PCDHGA6 |
| 5 | 141478245 | 141478373 | -1.406978088 | PCDHGA7 |
| 5 | 141485539 | 141485736 | 9.216295417 | PCDHGA10 |
| 5 | 141485539 | 141485736 | 9.216295417 | PCDHGA11 |
| 5 | 141485539 | 141485736 | 9.216295417 | PCDHGB6 |
| 5 | 141485539 | 141485736 | 9.216295417 | PCDHGB4 |
| 5 | 141485539 | 141485736 | 9.216295417 | PCDHGC3 |
| 5 | 141485539 | 141485736 | 9.216295417 | PCDHGA9 |
| 5 | 141485539 | 141485736 | 9.216295417 | PCDHGA5 |
| 5 | 141485539 | 141485736 | 9.216295417 | PCDHGA12 |
| 5 | 141485539 | 141485736 | 9.216295417 | PCDHGA2 |
| 5 | 141485539 | 141485736 | 9.216295417 | PCDHGA3 |
| 5 | 141485539 | 141485736 | 9.216295417 | PCDHGA4 |
| 5 | 141485539 | 141485736 | 9.216295417 | PCDHGB1 |
| 5 | 141485539 | 141485736 | 9.216295417 | PCDHGA8 |
| 5 | 141485539 | 141485736 | 9.216295417 | PCDHGB5 |
| 5 | 141485539 | 141485736 | 9.216295417 | PCDHGA6 |
| 5 | 141485539 | 141485736 | 9.216295417 | PCDHGB7 |
| 5 | 141485539 | 141485736 | 9.216295417 | PCDHGB2 |
| 5 | 141485539 | 141485736 | 9.216295417 | PCDHGB3 |
| 5 | 141485539 | 141485736 | 9.216295417 | PCDHGC4 |
| 5 | 141485539 | 141485736 | 9.216295417 | PCDHGA1 |
| 5 | 141485539 | 141485736 | 9.216295417 | PCDHGA7 |
| 5 | 141488883 | 141489085 | 9.39357346 | PCDHGA3 |
| 5 | 141488883 | 141489085 | 9.39357346 | PCDHGB6 |
| 5 | 141488883 | 141489085 | 9.39357346 | PCDHGA5 |
| 5 | 141488883 | 141489085 | 9.39357346 | PCDHGA8 |
| 5 | 141488883 | 141489085 | 9.39357346 | PCDHGB7 |
| 5 | 141488883 | 141489085 | 9.39357346 | PCDHGB5 |
| 5 | 141488883 | 141489085 | 9.39357346 | PCDHGB4 |
| 5 | 141488883 | 141489085 | 9.39357346 | PCDHGA2 |
| 5 | 141488883 | 141489085 | 9.39357346 | PCDHGA4 |
| 5 | 141488883 | 141489085 | 9.39357346 | PCDHGB2 |
| 5 | 141488883 | 141489085 | 9.39357346 | PCDHGC3 |
| 5 | 141488883 | 141489085 | 9.39357346 | PCDHGB3 |
| 5 | 141488883 | 141489085 | 9.39357346 | PCDHGA11 |
| 5 | 141488883 | 141489085 | 9.39357346 | PCDHGA6 |
| 5 | 141488883 | 141489085 | 9.39357346 | PCDHGB1 |
| 5 | 141488883 | 141489085 | 9.39357346 | PCDHGA7 |
| 7 | 98302485 | 98302746 | -1.214282316 | BRI3 |
| 7 | 98307390 | 98307620 | -1.049211201 | BRI3 |
| 7 | 98307390 | 98307620 | -1.049211201 | BAIAP2L1 |
| 7 | 99012222 | 99012419 | 1.308836493 | TRRAP |
| 7 | 99028683 | 99028883 | -1.082495973 | SMURF1 |
| 7 | 99028683 | 99028883 | -1.082495973 | AC004893.11 |
| 7 | 99486967 | 99487151 | 1.162730147 | ZNF394 |
| 7 | 99486967 | 99487151 | 1.162730147 | ZNF789 |
| 7 | 100120071 | 100120191 | 6.873571999 | RP11-506M12.1 |
| 7 | 100120071 | 100120191 | 6.873571999 | CNPY4 |
| 7 | 100319303 | 100319503 | -1.070182141 | SPDYE3 |
| 7 | 100374178 | 100374383 | -1.168714594 | PILRA |
| 7 | 100865861 | 100866025 | 3.001206603 | SLC12A9 |
| 7 | 101091018 | 101091220 | 1.204738987 | TRIM56 |
| 7 | 102227355 | 102227521 | -1.392285763 | CUX1 |
| 1 | 229268511 | 229268710 | 1.122447796 | RP5-1061H20.4 |
| 7 | 102248858 | 102249068 | 3.115440817 | CUX1 |
| 7 | 102396010 | 102396210 | 1.386486381 | PRKRIP1 |
| 7 | 102592691 | 102592862 | 6.239890132 | RASA4 |
| 7 | 102592691 | 102592862 | 6.239890132 | RP11-514P8.6 |
| 7 | 102668615 | 102668803 | -4.725605556 | POLR2J2 |
| 7 | 102668615 | 102668803 | -4.725605556 | RP11-577H5.5 |
| 7 | 102782742 | 102782940 | -4.038407799 | FAM185A |
| 7 | 103109504 | 103109660 | -1.386082747 | NAPEPLD |
| 7 | 103112316 | 103112489 | -6.478551021 | NAPEPLD |
| 7 | 105607853 | 105607992 | -6.028995861 | ATXN7L1 |
| 7 | 105712524 | 105712728 | -6.598224222 | ATXN7L1 |
| 7 | 106659819 | 106660175 | 2.481228368 | CTB-111H14.1 |
| 7 | 106659819 | 106660175 | 2.481228368 | CCDC71L |
| 7 | 107629053 | 107629263 | 1.262262377 | BCAP29 |
| 7 | 107629053 | 107629263 | 1.262262377 | WBP1LP2 |
| 7 | 107758236 | 107758434 | -1.058559969 | CBLL1 |
| 7 | 111420573 | 111420820 | -1.150451296 | IMMP2L |
| 7 | 114358570 | 114358771 | -1.587259841 | FOXP2 |
| 7 | 114689779 | 114689980 | -1.696591741 | FOXP2 |
| 7 | 114690758 | 114690958 | -1.470330498 | FOXP2 |
| 7 | 120974797 | 120974983 | -8.095163133 | ING3 |
| 7 | 128028720 | 128028919 | 9.323883787 | LRRC4 |
| 7 | 128028720 | 128028919 | 9.323883787 | SND1 |
| 7 | 128029352 | 128029551 | 2.425773263 | LRRC4 |
| 7 | 128029352 | 128029551 | 2.425773263 | SND1 |
| 1 | 229635191 | 229635393 | 1.068919063 | URB2 |
| 7 | 128525735 | 128525857 | -6.162793273 | RP11-212P7.2 |
| 7 | 128767362 | 128767558 | -1.150161405 | CALU |
| 7 | 128768965 | 128769167 | -1.007128476 | CALU |
| 7 | 128870476 | 128870676 | -2.170272542 | KCP |
| 7 | 128870476 | 128870676 | -2.170272542 | RP11-309L24.4 |
| 7 | 129056170 | 129056395 | 1.044429102 | TPI1P2 |
| 7 | 129485580 | 129485779 | -6.825862217 | STRIP2 |
| 1 | 229636705 | 229636900 | 1.402400414 | URB2 |
| 7 | 129724688 | 129724792 | -3.858477402 | NRF1 |
| 7 | 129755057 | 129755230 | -6.533380966 | NRF1 |
| 7 | 130440552 | 130440746 | -2.293828664 | CEP41 |
| 7 | 131107979 | 131108162 | 4.40024775 | LINC-PINT |

| | | | | |
|---|---|---|---|---|
| 5 | 141157246 | 141157397 | 1.277745691 | AC005754.7 |
| 5 | 141157246 | 141157397 | 1.277745691 | PCDHB17P |
| 5 | 141189487 | 141189648 | -3.140206944 | PCDHB9 |
| 5 | 141189487 | 141189648 | -3.140206944 | CH17-140K24.2 |
| 5 | 141189487 | 141189648 | -3.140206944 | CH17-140K24.7 |
| 5 | 141192231 | 141192451 | 1.205974264 | CH17-140K24.7 |
| 5 | 141192231 | 141192451 | 1.205974264 | CH17-140K24.2 |
| 5 | 141192231 | 141192451 | 1.205974264 | PCDHB10 |
| 5 | 141192231 | 141192451 | 1.205974264 | CH17-140K24.6 |
| 1 | 9356281 | 9356483 | 5.556463277 | SPSB1 |
| 5 | 141199772 | 141199961 | 2.009402201 | PCDHB11 |
| 5 | 141199772 | 141199961 | 2.009402201 | CH17-140K24.7 |
| 5 | 141199772 | 141199961 | 2.009402201 | CH17-140K24.2 |
| 5 | 141340734 | 141340933 | 1.275098541 | PCDHGA2 |
| 5 | 141340734 | 141340933 | 1.275098541 | PCDHGA1 |
| 5 | 141414952 | 141415154 | 1.136709564 | PCDHGB2 |
| 5 | 141414952 | 141415154 | 1.136709564 | PCDHGA2 |
| 5 | 141414952 | 141415154 | 1.136709564 | PCDHGA1 |
| 5 | 141414952 | 141415154 | 1.136709564 | PCDHGA3 |
| 5 | 141414952 | 141415154 | 1.136709564 | PCDHGA4 |
| 5 | 141414952 | 141415154 | 1.136709564 | PCDHGA8 |
| 5 | 141414952 | 141415154 | 1.136709564 | PCDHGA7 |
| 5 | 141414952 | 141415154 | 1.136709564 | PCDHGB1 |
| 5 | 141414952 | 141415154 | 1.136709564 | PCDHGB4 |
| 5 | 141414952 | 141415154 | 1.136709564 | PCDHGA5 |
| 5 | 141414952 | 141415154 | 1.136709564 | PCDHGB5 |
| 5 | 141414952 | 141415154 | 1.136709564 | PCDHGA10 |
| 5 | 141414952 | 141415154 | 1.136709564 | PCDHGA6 |
| 5 | 141414952 | 141415154 | 1.136709564 | PCDHGA9 |
| 5 | 141414952 | 141415154 | 1.136709564 | PCDHGB6 |
| 5 | 141414952 | 141415154 | 1.136709564 | PCDHGB3 |
| 5 | 141421580 | 141421743 | 1.827783394 | PCDHGB3 |
| 5 | 141421580 | 141421743 | 1.827783394 | PCDHGA6 |
| 5 | 141421580 | 141421743 | 1.827783394 | PCDHGA8 |
| 5 | 141421580 | 141421743 | 1.827783394 | PCDHGA5 |
| 5 | 141421580 | 141421743 | 1.827783394 | PCDHGA4 |
| 5 | 141421580 | 141421743 | 1.827783394 | PCDHGA9 |
| 5 | 141421580 | 141421743 | 1.827783394 | PCDHGA2 |
| 5 | 141421580 | 141421743 | 1.827783394 | PCDHGA10 |
| 5 | 141421580 | 141421743 | 1.827783394 | PCDHGB2 |
| 5 | 141421580 | 141421743 | 1.827783394 | PCDHGB1 |
| 5 | 141421580 | 141421743 | 1.827783394 | PCDHGA1 |
| 5 | 141421580 | 141421743 | 1.827783394 | PCDHGB6 |
| 5 | 141421580 | 141421743 | 1.827783394 | PCDHGB4 |
| 5 | 141421580 | 141421743 | 1.827783394 | PCDHGA7 |
| 5 | 141421580 | 141421743 | 1.827783394 | PCDHGA11 |
| 5 | 141421580 | 141421743 | 1.827783394 | PCDHGA3 |
| 5 | 141421580 | 141421743 | 1.827783394 | PCDHGB7 |
| 5 | 141421580 | 141421743 | 1.827783394 | PCDHGB5 |
| 5 | 141422997 | 141423195 | 1.438685471 | PCDHGA1 |
| 5 | 141422997 | 141423195 | 1.438685471 | PCDHGA9 |
| 5 | 141422997 | 141423195 | 1.438685471 | PCDHGB6 |
| 5 | 141422997 | 141423195 | 1.438685471 | PCDHGA3 |
| 5 | 141422997 | 141423195 | 1.438685471 | PCDHGA10 |
| 5 | 141488883 | 141489085 | 9.39357346 | PCDHGA1 |
| 5 | 141488883 | 141489085 | 9.39357346 | PCDHGA12 |
| 5 | 141488883 | 141489085 | 9.39357346 | PCDHGC4 |
| 5 | 141488883 | 141489085 | 9.39357346 | PCDHGA9 |
| 5 | 141488883 | 141489085 | 9.39357346 | PCDHGA10 |
| 5 | 141490123 | 141490274 | 9.154738177 | PCDHGB3 |
| 5 | 141490123 | 141490274 | 9.154738177 | PCDHGA6 |
| 5 | 141490123 | 141490274 | 9.154738177 | PCDHGB6 |
| 5 | 141490123 | 141490274 | 9.154738177 | PCDHGA1 |
| 5 | 141490123 | 141490274 | 9.154738177 | PCDHGA9 |
| 5 | 141490123 | 141490274 | 9.154738177 | PCDHGA7 |
| 5 | 141490123 | 141490274 | 9.154738177 | PCDHGA10 |
| 5 | 141490123 | 141490274 | 9.154738177 | PCDHGB7 |
| 5 | 141490123 | 141490274 | 9.154738177 | PCDHGC5 |
| 5 | 141490123 | 141490274 | 9.154738177 | PCDHGB2 |
| 5 | 141490123 | 141490274 | 9.154738177 | PCDHGC3 |
| 5 | 141490123 | 141490274 | 9.154738177 | PCDHGB5 |
| 5 | 141490123 | 141490274 | 9.154738177 | PCDHGA2 |
| 5 | 141490123 | 141490274 | 9.154738177 | PCDHGA12 |
| 5 | 141490123 | 141490274 | 9.154738177 | PCDHGA3 |
| 5 | 141490123 | 141490274 | 9.154738177 | PCDHGA4 |
| 5 | 141490123 | 141490274 | 9.154738177 | PCDHGC4 |
| 5 | 141490123 | 141490274 | 9.154738177 | PCDHGA11 |
| 5 | 141490123 | 141490274 | 9.154738177 | PCDHGA5 |
| 5 | 141490123 | 141490274 | 9.154738177 | PCDHGA8 |
| 5 | 141490123 | 141490274 | 9.154738177 | PCDHGB1 |
| 5 | 141490123 | 141490274 | 9.154738177 | PCDHGB4 |
| 5 | 141490719 | 141490905 | 9.231818061 | PCDHGB3 |
| 5 | 141490719 | 141490905 | 9.231818061 | PCDHGA5 |
| 5 | 141490719 | 141490905 | 9.231818061 | PCDHGA11 |
| 5 | 141490719 | 141490905 | 9.231818061 | PCDHGC4 |
| 5 | 141490719 | 141490905 | 9.231818061 | PCDHGA1 |
| 5 | 141490719 | 141490905 | 9.231818061 | PCDHGB1 |
| 5 | 141490719 | 141490905 | 9.231818061 | PCDHGC3 |
| 5 | 141490719 | 141490905 | 9.231818061 | PCDHGB5 |
| 5 | 141490719 | 141490905 | 9.231818061 | PCDHGA6 |
| 5 | 141490719 | 141490905 | 9.231818061 | PCDHGB4 |
| 5 | 141490719 | 141490905 | 9.231818061 | PCDHGB6 |
| 5 | 141490719 | 141490905 | 9.231818061 | PCDHGB2 |
| 5 | 141490719 | 141490905 | 9.231818061 | PCDHGA7 |
| 5 | 141490719 | 141490905 | 9.231818061 | PCDHGA3 |
| 5 | 141490719 | 141490905 | 9.231818061 | PCDHGA4 |
| 5 | 141490719 | 141490905 | 9.231818061 | PCDHGB7 |
| 5 | 141490719 | 141490905 | 9.231818061 | PCDHGA10 |
| 5 | 141490719 | 141490905 | 9.231818061 | PCDHGA2 |
| 5 | 141490719 | 141490905 | 9.231818061 | PCDHGC5 |
| 5 | 141490719 | 141490905 | 9.231818061 | PCDHGA8 |
| 5 | 141490719 | 141490905 | 9.231818061 | PCDHGA9 |
| 5 | 141490719 | 141490905 | 9.231818061 | PCDHGA12 |
| 5 | 141505236 | 141505437 | 8.135859168 | PCDHGA4 |
| 5 | 141505236 | 141505437 | 8.135859168 | PCDHGB5 |
| 5 | 141505236 | 141505437 | 8.135859168 | PCDHGB1 |
| 5 | 141505236 | 141505437 | 8.135859168 | PCDHGA3 |
| 5 | 141505236 | 141505437 | 8.135859168 | PCDHGB2 |
| 1 | 229637939 | 229638155 | 1.939454092 | URB2 |
| 7 | 132367934 | 132368133 | 2.169925001 | AC011625.1 |
| 7 | 132367934 | 132368133 | 2.169925001 | PLXNA4 |
| 7 | 134994257 | 134994454 | -6.824178791 | AGBL3 |
| 7 | 135165458 | 135165658 | -1.365096474 | TMEM140 |
| 7 | 135165458 | 135165658 | -1.365096474 | C7orf49 |
| 1 | 230323109 | 230323310 | -1.2115958 | PGBD5 |
| 7 | 137880406 | 137880572 | -1.624069715 | CREB3L2 |
| 7 | 138585011 | 138585202 | -1.181749624 | TRIM24 |
| 7 | 139678248 | 139678431 | -1.840548224 | HIPK2 |
| 7 | 140177537 | 140177688 | 2.759850541 | JHDM1D-AS1 |
| 7 | 140695434 | 140695624 | 1.100528334 | NDUFB2-AS1 |
| 7 | 140695434 | 140695624 | 1.100528334 | ADCK2 |
| 7 | 140695434 | 140695624 | 1.100528334 | NDUFB2 |
| 7 | 140726205 | 140726398 | -1.277742226 | BRAF |
| 7 | 143269234 | 143269470 | 1.545023842 | GSTK1 |
| 7 | 143269234 | 143269470 | 1.545023842 | AC073342.12 |
| 7 | 144215295 | 144215506 | -1.357417577 | RP4-545C24.1 |
| 7 | 144286096 | 144286292 | -2.31634525 | ARHGEF34P |
| 7 | 144286096 | 144286292 | -2.31634525 | OR2A1-AS1 |
| 7 | 144336589 | 144336790 | -1.147092799 | OR2A1-AS1 |
| 7 | 144753787 | 144753907 | -4.027641218 | TPK1 |
| 7 | 149282069 | 149282250 | 8.819529011 | ZNF783 |
| 7 | 149432832 | 149433031 | -1.80871325 | ZNF777 |
| 7 | 149861813 | 149862013 | 1.147727942 | ZNF862 |
| 7 | 149861813 | 149862013 | 1.147727942 | RP4-751H13.7 |
| 7 | 150720325 | 150720560 | -1.33530768 | GIMAP1 |
| 7 | 150720325 | 150720560 | -1.33530768 | GIMAP1-GIMAP5 |
| 7 | 151122778 | 151122955 | 6.508404144 | AGAP3 |
| 7 | 151130993 | 151131190 | 1.139416669 | AGAP3 |
| 1 | 11295110 | 11295307 | 1.478729241 | UBIAD1 |
| 7 | 151378953 | 151379132 | -3.947408882 | WDR86 |
| 7 | 151859747 | 151859919 | -6.183161107 | PRKAG2 |
| 7 | 152181708 | 152181835 | -1.248735045 | KMT2C |
| 7 | 152182259 | 152182414 | -1.316045686 | KMT2C |
| 1 | 231947448 | 231947590 | -1.561198316 | DISC1 |
| 7 | 155458155 | 155458376 | 2.365564819 | EN2 |
| 7 | 155775040 | 155775273 | -1.052905774 | RBM33 |
| 1 | 232036767 | 232036935 | -1.452743277 | DISC1 |
| 7 | 157002171 | 157002301 | -4.642701109 | MNX1 |
| 7 | 157110167 | 157110373 | -8.954090746 | RP4-639J15.3 |
| 7 | 157241208 | 157241413 | -1.264130989 | UBE3C |
| 7 | 157498506 | 157498669 | -3.321928095 | AC006372.4 |
| 7 | 157546453 | 157546685 | -1.145170621 | PTPRN2 |
| 7 | 157586688 | 157586880 | 1.75389732 | PTPRN2 |
| 7 | 157587697 | 157587840 | 1.002194313 | PTPRN2 |
| 7 | 157588361 | 157588588 | 2.427177466 | PTPRN2 |
| 7 | 157598555 | 157598746 | 1.374230617 | PTPRN2 |
| 7 | 157628510 | 157628667 | 1.640716658 | PTPRN2 |
| 7 | 157651231 | 157651423 | 1.45892831 | PTPRN2 |
| 7 | 157673949 | 157674134 | 7.959245296 | PTPRN2 |
| 7 | 157860199 | 157860407 | -1.213059752 | PTPRN2 |
| 7 | 157860199 | 157860407 | -1.213059752 | AC011899.9 |
| 7 | 157862854 | 157863057 | -1.219757229 | AC011899.9 |

| | | | | |
|---|---|---|---|---|
| 5 | 141422997 | 141423195 | 1.438685471 | PCDHGB1 |
| 5 | 141422997 | 141423195 | 1.438685471 | PCDHGA8 |
| 5 | 141422997 | 141423195 | 1.438685471 | PCDHGB4 |
| 5 | 141422997 | 141423195 | 1.438685471 | PCDHGB7 |
| 5 | 141422997 | 141423195 | 1.438685471 | PCDHGA2 |
| 5 | 141422997 | 141423195 | 1.438685471 | PCDHGA11 |
| 5 | 141422997 | 141423195 | 1.438685471 | PCDHGB3 |
| 5 | 141422997 | 141423195 | 1.438685471 | PCDHGA7 |
| 5 | 141422997 | 141423195 | 1.438685471 | PCDHGA6 |
| 5 | 141422997 | 141423195 | 1.438685471 | PCDHGA5 |
| 5 | 141422997 | 141423195 | 1.438685471 | PCDHGA4 |
| 5 | 141422997 | 141423195 | 1.438685471 | PCDHGB2 |
| 5 | 141422997 | 141423195 | 1.438685471 | PCDHGB5 |
| 5 | 141432497 | 141432655 | 3.472600545 | PCDHGB7 |
| 5 | 141432497 | 141432655 | 3.472600545 | PCDHGA11 |
| 5 | 141432497 | 141432655 | 3.472600545 | PCDHGB2 |
| 5 | 141432497 | 141432655 | 3.472600545 | PCDHGB3 |
| 5 | 141432497 | 141432655 | 3.472600545 | PCDHGA3 |
| 5 | 141432497 | 141432655 | 3.472600545 | PCDHGA5 |
| 5 | 141432497 | 141432655 | 3.472600545 | PCDHGA10 |
| 5 | 141432497 | 141432655 | 3.472600545 | PCDHGA8 |
| 5 | 141432497 | 141432655 | 3.472600545 | PCDHGB5 |
| 5 | 141432497 | 141432655 | 3.472600545 | PCDHGA1 |
| 5 | 141432497 | 141432655 | 3.472600545 | PCDHGA2 |
| 5 | 141432497 | 141432655 | 3.472600545 | PCDHGB6 |
| 5 | 141432497 | 141432655 | 3.472600545 | PCDHGA12 |
| 5 | 141432497 | 141432655 | 3.472600545 | PCDHGB1 |
| 5 | 141432497 | 141432655 | 3.472600545 | PCDHGA7 |
| 5 | 141432497 | 141432655 | 3.472600545 | PCDHGB4 |
| 5 | 141432497 | 141432655 | 3.472600545 | PCDHGA4 |
| 5 | 141432497 | 141432655 | 3.472600545 | PCDHGA6 |
| 5 | 141432497 | 141432655 | 3.472600545 | PCDHGA9 |
| 5 | 141475982 | 141476185 | 1.760771584 | PCDHGA9 |
| 5 | 141475982 | 141476185 | 1.760771584 | PCDHGB5 |
| 5 | 141475982 | 141476185 | 1.760771584 | PCDHGB6 |
| 5 | 141475982 | 141476185 | 1.760771584 | PCDHGB3 |
| 5 | 141475982 | 141476185 | 1.760771584 | PCDHGA11 |
| 5 | 141475982 | 141476185 | 1.760771584 | PCDHGB2 |
| 5 | 141475982 | 141476185 | 1.760771584 | PCDHGA8 |
| 5 | 141475982 | 141476185 | 1.760771584 | PCDHGA10 |
| 5 | 141475982 | 141476185 | 1.760771584 | PCDHGB7 |
| 5 | 141475982 | 141476185 | 1.760771584 | PCDHGB4 |
| 5 | 141475982 | 141476185 | 1.760771584 | PCDHGA6 |
| 5 | 141475982 | 141476185 | 1.760771584 | PCDHGA7 |
| 5 | 141475982 | 141476185 | 1.760771584 | PCDHGA2 |
| 5 | 141475982 | 141476185 | 1.760771584 | PCDHGA4 |
| 5 | 141475982 | 141476185 | 1.760771584 | PCDHGC3 |
| 5 | 141475982 | 141476185 | 1.760771584 | PCDHGA3 |
| 5 | 141475982 | 141476185 | 1.760771584 | PCDHGB1 |
| 5 | 141475982 | 141476185 | 1.760771584 | PCDHGA12 |
| 5 | 141475982 | 141476185 | 1.760771584 | PCDHGA5 |
| 5 | 141475982 | 141476185 | 1.760771584 | PCDHGA1 |
| 5 | 141477595 | 141477767 | 1.206457518 | PCDHGB4 |
| 5 | 141477595 | 141477767 | 1.206457518 | PCDHGB3 |
| 5 | 141505236 | 141505437 | 8.135859168 | PCDHGB6 |
| 5 | 141505236 | 141505437 | 8.135859168 | PCDHGC5 |
| 5 | 141505236 | 141505437 | 8.135859168 | PCDHGA9 |
| 5 | 141505236 | 141505437 | 8.135859168 | PCDHGA8 |
| 5 | 141505236 | 141505437 | 8.135859168 | PCDHGB7 |
| 5 | 141505236 | 141505437 | 8.135859168 | PCDHGA2 |
| 5 | 141505236 | 141505437 | 8.135859168 | PCDHGC4 |
| 5 | 141505236 | 141505437 | 8.135859168 | PCDHGA6 |
| 5 | 141505236 | 141505437 | 8.135859168 | PCDHGA5 |
| 5 | 141505236 | 141505437 | 8.135859168 | PCDHGA10 |
| 5 | 141505236 | 141505437 | 8.135859168 | PCDHGB3 |
| 5 | 141505236 | 141505437 | 8.135859168 | PCDHGC3 |
| 5 | 141505236 | 141505437 | 8.135859168 | PCDHGB4 |
| 5 | 141505236 | 141505437 | 8.135859168 | PCDHGA7 |
| 5 | 141505236 | 141505437 | 8.135859168 | PCDHGA12 |
| 5 | 141505236 | 141505437 | 8.135859168 | PCDHGA1 |
| 5 | 141505236 | 141505437 | 8.135859168 | PCDHGA11 |
| 5 | 141653731 | 141653929 | -2.688721282 | ARAP3 |
| 5 | 141863288 | 141863511 | -2.932001109 | PCDH1 |
| 5 | 141864402 | 141864609 | -2.807501285 | PCDH1 |
| 5 | 141945241 | 141945437 | -4.718355771 | PCDH12 |
| 5 | 142314439 | 142314628 | -1.352229842 | SPRY4 |
| 5 | 142995001 | 142995149 | -3.974183616 | ARHGAP26 |
| 5 | 143103546 | 143103739 | -1.12027125 | ARHGAP26 |
| 5 | 143203663 | 143203829 | 5.090576128 | ARHGAP26 |
| 5 | 143222428 | 143222625 | -1.355913073 | ARHGAP26 |
| 5 | 143223757 | 143223925 | -1.252683228 | ARHGAP26 |
| 5 | 143405674 | 143405872 | 2.095671993 | NR3C1 |
| 5 | 146510784 | 146510925 | -1.61536749 | TCERG1 |
| 5 | 148827443 | 148827629 | -3.050006517 | ADRB2 |
| 5 | 148827892 | 148828092 | -2.228149077 | ADRB2 |
| 5 | 149350990 | 149351216 | -3.106356304 | GRPEL2-AS1 |
| 5 | 149350990 | 149351216 | -3.106356304 | GRPEL2 |
| 5 | 149368494 | 149368700 | -1.213526988 | PCYOX1L |
| 5 | 149493317 | 149493514 | -2.091096437 | CSNK1A1 |
| 5 | 149499124 | 149499312 | 7.898577436 | CTB-89H12.4 |
| 5 | 149499124 | 149499312 | 7.898577436 | CSNK1A1 |
| 5 | 149501291 | 149501487 | 7.563513522 | CTB-89H12.4 |
| 5 | 149501291 | 149501487 | 7.563513522 | CSNK1A1 |
| 5 | 149977615 | 149977812 | -1.125877296 | SLC26A2 |
| 5 | 149980430 | 149980715 | -1.389106974 | SLC26A2 |
| 5 | 150053572 | 150053769 | 6.851302801 | CSF1R |
| 5 | 150520928 | 150521082 | -7.426800982 | NDST1 |
| 5 | 150650449 | 150650648 | 4.215226276 | SYNPO |
| 5 | 150779091 | 150779253 | 3.761780978 | SMIM3 |
| 5 | 151030416 | 151030624 | 2.49356424 | TNIP1 |
| 5 | 154417070 | 154417247 | -2.596885872 | GALNT10 |
| 5 | 154417070 | 154417247 | -2.596885872 | SAP30L-AS1 |
| 5 | 154802131 | 154802296 | -1.370731522 | LARP1 |
| 5 | 157086788 | 157086987 | -1.207144964 | HAVCR2 |
| 5 | 157106717 | 157106900 | -1.186318629 | CTB-120L21.1 |
| 5 | 157106717 | 157106900 | -1.186318629 | HAVCR2 |
| 5 | 157138748 | 157138989 | -1.362589702 | HAVCR2 |
| 5 | 157138748 | 157138989 | -1.362589702 | MED7 |
| 7 | 157862854 | 157863057 | -1.219757229 | PTPRN2 |
| 7 | 157868433 | 157868632 | -1.816450627 | PTPRN2 |
| 7 | 157868433 | 157868632 | -1.816450627 | RP11-452C13.1 |
| 7 | 158791142 | 158791330 | -1.358246062 | ESYT2 |
| 7 | 158876438 | 158876630 | -2.520041059 | WDR60 |
| 8 | 245290 | 245501 | -1.052754524 | ZNF596 |
| 8 | 710331 | 710546 | 6.218355771 | ERICH1 |
| 8 | 1761479 | 1761691 | -1.088726706 | CLN8 |
| 8 | 1948182 | 1948453 | -3.740899716 | ARHGEF10 |
| 8 | 1948182 | 1948453 | -3.740899716 | RP11-439C15.6 |
| 8 | 2000542 | 2000747 | -1.691773772 | KBTBD11 |
| 8 | 2006525 | 2006744 | -1.156706574 | KBTBD11 |
| 8 | 6405905 | 6406037 | 1.309211291 | RP11-115C21.2 |
| 1 | 232976556 | 232976709 | -1.131623526 | NTPCR |
| 8 | 8319000 | 8319149 | 4.945462622 | SGK223 |
| 8 | 8319210 | 8319404 | 1.673600112 | SGK223 |
| 8 | 8327803 | 8328009 | 1.416291141 | SGK223 |
| 8 | 8376405 | 8376583 | 1.016200565 | SGK223 |
| 8 | 8377503 | 8377703 | 1.562873819 | SGK223 |
| 8 | 8381517 | 8381714 | 3.158767864 | SGK223 |
| 8 | 8703230 | 8703426 | 1.845793561 | CLDN23 |
| 8 | 9032831 | 9033029 | -1.053616612 | ERI1 |
| 8 | 9661615 | 9661745 | -5.683161107 | TNKS |
| 8 | 11107905 | 11108084 | 1.602122537 | XKR6 |
| 8 | 11107905 | 11108084 | 1.602122537 | AF131215.2 |
| 8 | 11108769 | 11108906 | 2.347241131 | AF131215.2 |
| 8 | 11108769 | 11108906 | 2.347241131 | XKR6 |
| 8 | 11119314 | 11119488 | -5.595221009 | XKR6 |
| 8 | 11844826 | 11845025 | 10.46177328 | CTSB |
| 8 | 16131712 | 16131843 | -1.409713877 | MSR1 |
| 8 | 17754020 | 17754211 | 1.176596995 | MTUS1 |
| 1 | 233345294 | 233345480 | -6.500704097 | RP5-862P8.2 |
| 8 | 17910302 | 17910489 | 2.993544028 | FGL1 |
| 8 | 18020952 | 18021095 | 6.998236744 | PCM1 |
| 8 | 19395498 | 19395675 | -1.739301314 | SH2D4A |
| 8 | 20246739 | 20246947 | 1.352018588 | LZTS1 |
| 8 | 20249196 | 20249389 | 2.755331891 | LZTS1 |
| 8 | 20253259 | 20253394 | 1.43431412 | LZTS1 |
| 8 | 22211761 | 22211954 | 1.179022155 | BMP1 |
| 8 | 23007121 | 23007319 | -1.077506186 | RP11-875O11.1 |
| 8 | 23007121 | 23007319 | -1.077506186 | RHOBTB2 |
| 8 | 23017554 | 23017749 | -1.223115476 | RP11-875O11.1 |
| 8 | 23017554 | 23017749 | -1.223115476 | RHOBTB2 |
| 8 | 23018249 | 23018449 | -1.354703372 | RP11-875O11.1 |
| 8 | 23018249 | 23018449 | -1.354703372 | RHOBTB2 |
| 8 | 23022679 | 23022879 | -1.294087724 | TNFRSF10B |
| 8 | 23246477 | 23246676 | 1.028791439 | CHMP7 |
| 8 | 23291251 | 23291443 | -6.72766361 | R3HCC1 |
| 8 | 23571539 | 23571741 | -1.429960966 | SLC25A37 |
| 8 | 26324646 | 26324900 | -2.689582202 | PPP2R2A |
| 8 | 26379636 | 26379803 | -6.15983606 | SDAD1P1 |
| 8 | 27776516 | 27776701 | 2.277294426 | ESCO2 |
| 8 | 28502830 | 28503079 | 1.059035414 | FZD3 |
| 8 | 28562747 | 28562948 | 1.085526075 | FZD3 |

| | | | | |
|---|---|---|---|---|
| 5 | 141477595 | 141477767 | 1.206457518 | PCDHGC3 |
| 5 | 141477595 | 141477767 | 1.206457518 | PCDHGA10 |
| 5 | 141477595 | 141477767 | 1.206457518 | PCDHGA1 |
| 5 | 141477595 | 141477767 | 1.206457518 | PCDHGB6 |
| 5 | 141477595 | 141477767 | 1.206457518 | PCDHGB7 |
| 5 | 141477595 | 141477767 | 1.206457518 | PCDHGA7 |
| 5 | 141477595 | 141477767 | 1.206457518 | PCDHGA8 |
| 5 | 141477595 | 141477767 | 1.206457518 | PCDHGB2 |
| 5 | 141477595 | 141477767 | 1.206457518 | PCDHGA12 |
| 5 | 141477595 | 141477767 | 1.206457518 | PCDHGA6 |
| 5 | 141477595 | 141477767 | 1.206457518 | PCDHGA9 |
| 5 | 141477595 | 141477767 | 1.206457518 | PCDHGA4 |
| 5 | 141477595 | 141477767 | 1.206457518 | PCDHGA5 |
| 5 | 141477595 | 141477767 | 1.206457518 | PCDHGB1 |
| 5 | 141477595 | 141477767 | 1.206457518 | PCDHGA3 |
| 5 | 141477595 | 141477767 | 1.206457518 | PCDHGB5 |
| 5 | 141477595 | 141477767 | 1.206457518 | PCDHGA11 |
| 5 | 141477595 | 141477767 | 1.206457518 | PCDHGA2 |
| 5 | 141478245 | 141478373 | 2.184927772 | PCDHGB5 |
| 5 | 141478245 | 141478373 | 2.184927772 | PCDHGA11 |
| 5 | 141478245 | 141478373 | 2.184927772 | PCDHGB2 |
| 5 | 141478245 | 141478373 | 2.184927772 | PCDHGB7 |
| 5 | 141478245 | 141478373 | 2.184927772 | PCDHGA5 |
| 5 | 141478245 | 141478373 | 2.184927772 | PCDHGA9 |
| 5 | 141478245 | 141478373 | 2.184927772 | PCDHGB4 |
| 5 | 141478245 | 141478373 | 2.184927772 | PCDHGA7 |
| 5 | 141478245 | 141478373 | 2.184927772 | PCDHGA10 |
| 5 | 141478245 | 141478373 | 2.184927772 | PCDHGA4 |
| 5 | 141478245 | 141478373 | 2.184927772 | PCDHGB3 |
| 5 | 141478245 | 141478373 | 2.184927772 | PCDHGA1 |
| 5 | 141478245 | 141478373 | 2.184927772 | PCDHGA8 |
| 5 | 141478245 | 141478373 | 2.184927772 | PCDHGB1 |
| 5 | 141478245 | 141478373 | 2.184927772 | PCDHGA12 |
| 5 | 141478245 | 141478373 | 2.184927772 | PCDHGB6 |
| 5 | 141478245 | 141478373 | 2.184927772 | PCDHGA2 |
| 5 | 141478245 | 141478373 | 2.184927772 | PCDHGA3 |
| 5 | 141478245 | 141478373 | 2.184927772 | PCDHGA6 |
| 5 | 141478245 | 141478373 | 2.184927772 | PCDHGC3 |
| 5 | 141485539 | 141485736 | -8.812882955 | PCDHGA1 |
| 5 | 141485539 | 141485736 | -8.812882955 | PCDHGA10 |
| 5 | 141485539 | 141485736 | -8.812882955 | PCDHGB5 |
| 5 | 141485539 | 141485736 | -8.812882955 | PCDHGB1 |
| 5 | 141485539 | 141485736 | -8.812882955 | PCDHGB3 |
| 5 | 141485539 | 141485736 | -8.812882955 | PCDHGA9 |
| 5 | 141485539 | 141485736 | -8.812882955 | PCDHGA8 |
| 5 | 141485539 | 141485736 | -8.812882955 | PCDHGB4 |
| 5 | 141485539 | 141485736 | -8.812882955 | PCDHGB2 |
| 5 | 141485539 | 141485736 | -8.812882955 | PCDHGC4 |
| 5 | 141485539 | 141485736 | -8.812882955 | PCDHGA5 |
| 5 | 141485539 | 141485736 | -8.812882955 | PCDHGA7 |
| 5 | 141485539 | 141485736 | -8.812882955 | PCDHGB6 |
| 5 | 141485539 | 141485736 | -8.812882955 | PCDHGA12 |
| 5 | 141485539 | 141485736 | -8.812882955 | PCDHGA6 |
| 5 | 141485539 | 141485736 | -8.812882955 | PCDHGA4 |
| 5 | 162153203 | 162153347 | -1.919062505 | GABRG2 |
| 5 | 168566108 | 168566338 | -1.128765403 | PANK3 |
| 5 | 168568770 | 168569030 | -1.533584722 | PANK3 |
| 5 | 169823532 | 169823728 | -4.482892142 | DOCK2 |
| 5 | 169825281 | 169825430 | -4.857122759 | DOCK2 |
| 5 | 170242721 | 170242921 | -1.245192622 | C5orf58 |
| 1 | 9367340 | 9367552 | 1.404803936 | SPSB1 |
| 1 | 204222076 | 204222296 | 3.922531156 | PLEKHA6 |
| 5 | 170271692 | 170271873 | 1.385094707 | LCP2 |
| 5 | 170979940 | 170980102 | -2.548184197 | RANBP17 |
| 5 | 172043699 | 172043897 | -1.53481971 | STK10 |
| 5 | 172044704 | 172044906 | -1.03555922 | STK10 |
| 5 | 172257745 | 172257915 | -5.309651478 | UBTD2 |
| 5 | 172334320 | 172334520 | -1.48941701 | SH3PXD2B |
| 5 | 172337975 | 172338155 | -1.739749229 | SH3PXD2B |
| 5 | 172770910 | 172771109 | 10.82192237 | RP11-779O18.3 |
| 5 | 172770910 | 172771109 | 10.82192237 | DUSP1 |
| 5 | 173317817 | 173318023 | -4.721453648 | STC2 |
| 5 | 175682975 | 175683182 | -1.537393226 | HRH2 |
| 5 | 175684024 | 175684196 | -3.497176718 | HRH2 |
| 5 | 175707968 | 175708076 | -3.780734953 | HRH2 |
| 5 | 176303914 | 176304106 | 3.488639962 | SIMC1 |
| 5 | 176507277 | 176507404 | -1.651976883 | FAF2 |
| 5 | 176598028 | 176598224 | -1.03899246 | GPRIN1 |
| 5 | 177209744 | 177209945 | -2.0418815 | NSD1 |
| 5 | 177210955 | 177211135 | -8.179900513 | NSD1 |
| 5 | 177211703 | 177211959 | -2.144234234 | NSD1 |
| 1 | 204469242 | 204469458 | -1.386591946 | PIK3C2B |
| 5 | 177294105 | 177294302 | -2.409321825 | NSD1 |
| 5 | 177477947 | 177478142 | -2.003681694 | RP11-1334A24.5 |
| 5 | 177503560 | 177503827 | -3.110438035 | RP11-1334A24.6 |
| 5 | 177503560 | 177503827 | -3.110438035 | DOK3 |
| 5 | 177615553 | 177615731 | -5.724316284 | RP11-1277A3.1 |
| 5 | 177617897 | 177618082 | -2.036481636 | RP11-1277A3.1 |
| 5 | 177618989 | 177619186 | -1.867122813 | RP11-1277A3.1 |
| 5 | 177618989 | 177619186 | -1.867122813 | RP11-1277A3.2 |
| 5 | 178207618 | 178207816 | 6.957379419 | HNRNPAB |
| 5 | 178313436 | 178313618 | -5.764227055 | COL23A1 |
| 1 | 205083707 | 205083942 | -1.932968782 | TMEM81 |
| 5 | 178366059 | 178366253 | -4.398830763 | COL23A1 |
| 5 | 178507261 | 178507545 | -2.934275525 | COL23A1 |
| 5 | 178508041 | 178508169 | -7.740631683 | COL23A1 |
| 5 | 178508773 | 178508969 | -1.635377721 | COL23A1 |
| 5 | 178514091 | 178514285 | -5.93633744 | COL23A1 |
| 5 | 178883173 | 178883346 | -2.669385931 | ZNF354B |
| 5 | 178883886 | 178884087 | -1.315748811 | ZNF354B |
| 1 | 205147458 | 205147567 | -1.756688731 | DSTYK |
| 5 | 179079527 | 179079727 | -2.654085231 | ZNF354C |
| 5 | 179083043 | 179083162 | -2.264791449 | ZNF354C |
| 5 | 179083043 | 179083162 | -2.264791449 | RP11-281O15.7 |
| 5 | 179113296 | 179113483 | 1.286181232 | ADAMTS2 |
| 5 | 179192815 | 179192965 | 6.498766174 | ADAMTS2 |
| 5 | 179193547 | 179193746 | 5.619799265 | ADAMTS2 |
| 5 | 179575335 | 179575535 | -6.770548308 | RUFY1 |
| 8 | 29350188 | 29350325 | 3.881196868 | DUSP4 |
| 8 | 30156017 | 30156204 | 5.618007096 | RP11-51J9.5 |
| 8 | 30156017 | 30156204 | 5.618007096 | LEPROTL1 |
| 8 | 30565804 | 30566076 | -2.180021734 | RBPMS |
| 8 | 30596694 | 30596896 | -4.122968389 | GTF2E2 |
| 8 | 30624037 | 30624208 | -6.272120003 | GTF2E2 |
| 8 | 32763972 | 32764172 | 3.482715914 | NRG1 |
| 8 | 35952129 | 35952324 | 2.29248125 | RP11-473J6.1 |
| 8 | 35956125 | 35956297 | -1.030700272 | RP11-473J6.1 |
| 8 | 37842435 | 37842633 | -2.992446554 | ADGRA2 |
| 8 | 37859971 | 37860155 | 1.037762576 | RP11-205M5.3 |
| 8 | 37859971 | 37860155 | 1.037762576 | RAB11FIP1 |
| 8 | 38275620 | 38275809 | -7.223799929 | WHSC1L1 |
| 8 | 41621003 | 41621269 | 1.385588935 | RP11-360L9.8 |
| 8 | 41621003 | 41621269 | 1.385588935 | GPAT4 |
| 8 | 41655501 | 41655683 | 3.599968275 | ANK1 |
| 8 | 42025756 | 42025887 | -1.341700212 | KAT6A |
| 8 | 42542126 | 42542287 | -6.084649348 | SMIM19 |
| 8 | 42908593 | 42908800 | -1.277730681 | HOOK3 |
| 1 | 235450236 | 235450371 | -1.612344731 | B3GALNT2 |
| 8 | 47468624 | 47468814 | -7.102323778 | SPIDR |
| 8 | 47623364 | 47623628 | 1.41034478 | SPIDR |
| 8 | 47626248 | 47626370 | -6.273567766 | SPIDR |
| 1 | 11805564 | 11805754 | 1.023324896 | MTHFR |
| 1 | 235662804 | 235662982 | -2.338535255 | LYST |
| 8 | 47737218 | 47737404 | 13.29669727 | CEBPD |
| 8 | 49176835 | 49177064 | -1.61546111 | RP11-10H3.1 |
| 8 | 49178172 | 49178323 | -1.204375475 | RP11-10H3.1 |
| 8 | 49200478 | 49200671 | -2.995879194 | RP11-10H3.1 |
| 1 | 235738003 | 235738197 | -1.135182668 | LDHAP2 |
| 1 | 235738003 | 235738197 | -1.135182668 | LYST |
| 8 | 51860946 | 51861150 | -1.129340014 | PCMTD1 |
| 8 | 52656154 | 52656358 | -1.17352552 | RB1CC1 |
| 8 | 52714786 | 52714980 | 2.27321153 | RB1CC1 |
| 8 | 53780804 | 53781022 | -1.154776101 | ATP6V1H |
| 8 | 53781425 | 53781614 | -1.522863065 | ATP6V1H |
| 8 | 53784307 | 53784499 | -1.663673322 | ATP6V1H |
| 8 | 53936820 | 53937017 | -2.79370699 | RGS20 |
| 8 | 57545475 | 57545688 | -1.164147399 | RP11-246K15.1 |
| 8 | 57572649 | 57572751 | -2.962396455 | RP11-246K15.1 |
| 8 | 57714972 | 57715177 | -1.203806371 | RP11-388G22.1 |
| 8 | 58146130 | 58146328 | 1.639446705 | FAM110B |
| 8 | 58146948 | 58147260 | 1.534156767 | FAM110B |
| 1 | 235806031 | 235806256 | -1.217212118 | LYST |
| 8 | 60465067 | 60465219 | 5.793888758 | LINC01301 |
| 8 | 60760872 | 60761082 | -3.332041687 | CHD7 |
| 1 | 235808863 | 235809071 | -1.340003504 | LYST |
| 8 | 60860818 | 60860996 | -1.091346149 | CHD7 |
| 8 | 61579995 | 61580186 | 4.730739724 | ASPH |
| 8 | 61579995 | 61580186 | 4.730739724 | AC090094.1 |
| 8 | 63167983 | 63168167 | 1.510363491 | YTHDF3-AS1 |
| 1 | 235809998 | 235810196 | -1.15729413 | LYST |
| 8 | 67313471 | 67313622 | -4.330889049 | ARFGEF1 |
| 8 | 70113033 | 70113245 | -1.376151857 | NCOA2 |

| | | | | |
|---|---|---|---|---|
| 5 | 141485539 | 141485736 | -8.812882955 | PCDHGC3 |
| 5 | 141485539 | 141485736 | -8.812882955 | PCDHGB7 |
| 5 | 141485539 | 141485736 | -8.812882955 | PCDHGA11 |
| 5 | 141485539 | 141485736 | -8.812882955 | PCDHGA3 |
| 5 | 141485539 | 141485736 | -8.812882955 | PCDHGA2 |
| 5 | 141488883 | 141489085 | -8.786557299 | PCDHGC3 |
| 5 | 141488883 | 141489085 | -8.786557299 | PCDHGB2 |
| 5 | 141488883 | 141489085 | -8.786557299 | PCDHGA3 |
| 5 | 141488883 | 141489085 | -8.786557299 | PCDHGA11 |
| 5 | 141488883 | 141489085 | -8.786557299 | PCDHGA8 |
| 5 | 141488883 | 141489085 | -8.786557299 | PCDHGA10 |
| 5 | 141488883 | 141489085 | -8.786557299 | PCDHGB5 |
| 5 | 141488883 | 141489085 | -8.786557299 | PCDHGB3 |
| 5 | 141488883 | 141489085 | -8.786557299 | PCDHGA9 |
| 5 | 141488883 | 141489085 | -8.786557299 | PCDHGB4 |
| 5 | 141488883 | 141489085 | -8.786557299 | PCDHGA6 |
| 5 | 141488883 | 141489085 | -8.786557299 | PCDHGA12 |
| 5 | 141488883 | 141489085 | -8.786557299 | PCDHGC4 |
| 5 | 141488883 | 141489085 | -8.786557299 | PCDHGA7 |
| 5 | 141488883 | 141489085 | -8.786557299 | PCDHGB6 |
| 5 | 141488883 | 141489085 | -8.786557299 | PCDHGA4 |
| 5 | 141488883 | 141489085 | -8.786557299 | PCDHGB1 |
| 5 | 141488883 | 141489085 | -8.786557299 | PCDHGB7 |
| 5 | 141488883 | 141489085 | -8.786557299 | PCDHGA2 |
| 5 | 141488883 | 141489085 | -8.786557299 | PCDHGA1 |
| 5 | 141488883 | 141489085 | -8.786557299 | PCDHGA5 |
| 5 | 141490123 | 141490274 | -7.862709758 | PCDHGC3 |
| 5 | 141490123 | 141490274 | -7.862709758 | PCDHGA10 |
| 5 | 141490123 | 141490274 | -7.862709758 | PCDHGA11 |
| 5 | 141490123 | 141490274 | -7.862709758 | PCDHGB4 |
| 5 | 141490123 | 141490274 | -7.862709758 | PCDHGB7 |
| 5 | 141490123 | 141490274 | -7.862709758 | PCDHGB1 |
| 5 | 141490123 | 141490274 | -7.862709758 | PCDHGB5 |
| 5 | 141490123 | 141490274 | -7.862709758 | PCDHGA5 |
| 5 | 141490123 | 141490274 | -7.862709758 | PCDHGA8 |
| 5 | 141490123 | 141490274 | -7.862709758 | PCDHGA3 |
| 5 | 141490123 | 141490274 | -7.862709758 | PCDHGB6 |
| 5 | 141490123 | 141490274 | -7.862709758 | PCDHGC4 |
| 5 | 141490123 | 141490274 | -7.862709758 | PCDHGA12 |
| 5 | 141490123 | 141490274 | -7.862709758 | PCDHGA6 |
| 5 | 141490123 | 141490274 | -7.862709758 | PCDHGA2 |
| 5 | 141490123 | 141490274 | -7.862709758 | PCDHGA1 |
| 5 | 141490123 | 141490274 | -7.862709758 | PCDHGA9 |
| 5 | 141490123 | 141490274 | -7.862709758 | PCDHGB3 |
| 5 | 141490123 | 141490274 | -7.862709758 | PCDHGB2 |
| 5 | 141490123 | 141490274 | -7.862709758 | PCDHGA4 |
| 5 | 141490123 | 141490274 | -7.862709758 | PCDHGA7 |
| 5 | 141490123 | 141490274 | -7.862709758 | PCDHGC5 |
| 5 | 141490719 | 141490905 | -8.53523869 | PCDHGA9 |
| 5 | 141490719 | 141490905 | -8.53523869 | PCDHGA12 |
| 5 | 141490719 | 141490905 | -8.53523869 | PCDHGA5 |
| 5 | 141490719 | 141490905 | -8.53523869 | PCDHGB1 |
| 5 | 141490719 | 141490905 | -8.53523869 | PCDHGC5 |
| 5 | 141490719 | 141490905 | -8.53523869 | PCDHGB7 |

| | | | | |
|---|---|---|---|---|
| 1 | 205169385 | 205169589 | -1.196385865 | DSTYK |
| 5 | 179768751 | 179768953 | -7.120991575 | MAML1 |
| 5 | 179883848 | 179884051 | 2.297296255 | TBC1D9B |
| 5 | 179912831 | 179912970 | -6.281121212 | RNF130 |
| 1 | 205187526 | 205187707 | -1.152709287 | DSTYK |
| 5 | 180789790 | 180789990 | -1.239938487 | MGAT1 |
| 5 | 180808752 | 180808953 | -1.112931918 | MGAT1 |
| 5 | 180809590 | 180809715 | -7.079935668 | MGAT1 |
| 5 | 180834408 | 180834607 | -1.074190392 | LINC00847 |
| 1 | 205211527 | 205211699 | 5.36735481 | DSTYK |
| 5 | 180851197 | 180851398 | -1.262100933 | ZFP62 |
| 5 | 181257132 | 181257319 | -1.784652953 | CTC-338M12.4 |
| 5 | 181257132 | 181257319 | -1.784652953 | TRIM52 |
| 6 | 326876 | 327013 | 7.36310908 | DUSP22 |
| 6 | 407744 | 407929 | 3.534850748 | IRF4 |
| 6 | 1394811 | 1394992 | -4.196158711 | FOXF2 |
| 6 | 1610739 | 1610924 | 1.381785418 | FOXC1 |
| 6 | 1612029 | 1612223 | 1.263943695 | FOXC1 |
| 6 | 2288998 | 2289111 | -1.750455547 | GMDS-AS1 |
| 6 | 2666977 | 2667087 | -1.011981622 | MYLK4 |
| 6 | 2841184 | 2841383 | 10.51792376 | SERPINB1 |
| 6 | 3003754 | 3003939 | 2.246298161 | NQO2 |
| 6 | 3020030 | 3020232 | -1.750571248 | NQO2 |
| 6 | 3023513 | 3023656 | -7.464814142 | HTATSF1P2 |
| 6 | 3023513 | 3023656 | -7.464814142 | NQO2 |
| 6 | 3090219 | 3090415 | 2.122936212 | RIPK1 |
| 6 | 3094007 | 3094212 | 1.834536842 | RIPK1 |
| 6 | 3722875 | 3723063 | 8.842336648 | PXDC1 |
| 6 | 5998339 | 5998549 | 2.011297014 | NRN1 |
| 6 | 6685464 | 6685664 | -3.321928095 | RP1-80N2.4 |
| 6 | 6700348 | 6700548 | -2.768054057 | RP1-80N2.2 |
| 6 | 7230125 | 7230317 | -7.311997884 | RREB1 |
| 6 | 7231610 | 7231760 | -1.277433426 | RREB1 |
| 6 | 7726649 | 7726820 | 3.401096608 | BMP6 |
| 1 | 205663231 | 205663429 | -2.473624167 | SLC45A3 |
| 6 | 7975112 | 7975328 | -1.466872148 | BLOC1S5-TXNDC5 |
| 6 | 8015612 | 8015851 | -1.322354815 | BLOC1S5-TXNDC5 |
| 6 | 8015612 | 8015851 | -1.322354815 | BLOC1S5 |
| 6 | 8015612 | 8015851 | -1.322354815 | EEF1E1-BLOC1S5 |
| 6 | 10994757 | 10994959 | -2.921229848 | ELOVL2 |
| 1 | 205769463 | 205769672 | 1.402004368 | RAB29 |
| 6 | 11351786 | 11351987 | -1.465578777 | NEDD9 |
| 6 | 11575417 | 11575621 | -1.154298831 | TMEM170B |
| 6 | 12122555 | 12122722 | -2.593446413 | HIVEP1 |
| 6 | 13287435 | 13287650 | -1.208600871 | RP1-257A7.4 |
| 6 | 13287435 | 13287650 | -1.208600871 | TBC1D7 |
| 6 | 13287435 | 13287650 | -1.208600871 | PHACTR1 |
| 6 | 13364728 | 13364932 | -1.242714873 | GFOD1 |
| 6 | 13604820 | 13604980 | -5.804589369 | SIRT5 |
| 6 | 14135576 | 14135781 | 1.28244794 | CD83 |
| 1 | 205978340 | 205978541 | 1.542668219 | RAB7B |
| 6 | 15244540 | 15244740 | -1.743015254 | RP11-560J1.2 |
| 1 | 205979360 | 205979471 | 6.220953352 | RAB7B |
| 6 | 15500901 | 15501132 | -1.759253799 | JARID2 |

| | | | | |
|---|---|---|---|---|
| 8 | 70585270 | 70585454 | 4 | TRAM1 |
| 1 | 235962407 | 235962616 | -1.59829448 | RP4-580N22.2 |
| 1 | 11920200 | 11920407 | 1.304775665 | KIAA2013 |
| 8 | 74236861 | 74237061 | 2.823695271 | JPH1 |
| 8 | 74314920 | 74315129 | 1.64766513 | JPH1 |
| 1 | 236195324 | 236195444 | 5.440939354 | GPR137B |
| 8 | 75444470 | 75444671 | -6.035060472 | HNF4G |
| 8 | 80486289 | 80486486 | 8.676138122 | ZBTB10 |
| 8 | 80487185 | 80487368 | 11.06087172 | ZBTB10 |
| 8 | 80975603 | 80975805 | -1.262278024 | PAG1 |
| 8 | 81279987 | 81280303 | 2.034708867 | RP11-363E6.3 |
| 8 | 89703075 | 89703256 | 1.953445298 | RP11-37B2.1 |
| 8 | 94432295 | 94432483 | -5.657574781 | FSBP |
| 8 | 94432295 | 94432483 | -5.657574781 | RAD54B |
| 8 | 94511412 | 94511609 | 1.218201768 | KIAA1429 |
| 8 | 95247521 | 95247693 | 1.224223469 | C8orf37 |
| 8 | 95247521 | 95247693 | 1.224223469 | C8orf37-AS1 |
| 8 | 96493834 | 96494034 | 9.258842382 | SDC2 |
| 1 | 236820114 | 236820292 | -6.828380012 | RPSAP21 |
| 1 | 236820114 | 236820292 | -6.828380012 | MTR |
| 8 | 96742215 | 96742429 | -1.511836414 | CPQ |
| 8 | 96794951 | 96795149 | 6.916840374 | CPQ |
| 8 | 96930670 | 96930770 | 5.054915327 | CPQ |
| 8 | 96955852 | 96956040 | -7.164898169 | CPQ |
| 8 | 96974133 | 96974304 | 1.029446845 | CPQ |
| 8 | 96976967 | 96977141 | -1.19818497 | CPQ |
| 8 | 98694701 | 98694984 | -2.888659274 | STK3 |
| 8 | 98942657 | 98942816 | 4.26569073 | STK3 |
| 8 | 98949310 | 98949511 | 1.561269206 | OSR2 |
| 8 | 99205066 | 99205269 | -1.320389052 | VPS13B |
| 8 | 99293547 | 99293730 | -5.663214744 | VPS13B |
| 8 | 99425369 | 99425510 | 1.763272907 | VPS13B |
| 8 | 99741015 | 99741183 | -1.309271152 | VPS13B |
| 1 | 11925300 | 11925499 | 7.061413997 | KIAA2013 |
| 8 | 100310458 | 100310717 | 1.894764875 | RNF19A |
| 8 | 100312144 | 100312390 | 1.333820673 | RNF19A |
| 8 | 100336178 | 100336428 | 1.946621453 | RNF19A |
| 8 | 100344322 | 100344578 | 1.139905992 | KB-1991G8.1 |
| 8 | 100346500 | 100346703 | 1.378173205 | KB-1991G8.1 |
| 8 | 100510007 | 100510217 | -1.218702656 | ANKRD46 |
| 8 | 100550532 | 100550703 | 1.087022418 | ANKRD46 |
| 1 | 237077341 | 237077540 | -5.302739759 | RYR2 |
| 8 | 101205381 | 101205577 | 11.34653884 | ZNF706 |
| 8 | 101719500 | 101719687 | 2.968821358 | NCALD |
| 8 | 101825111 | 101825317 | 2.052061472 | NCALD |
| 8 | 102651667 | 102651867 | 1.004487968 | KLF10 |
| 8 | 102809055 | 102809239 | 1.085392312 | KB-1732A1.1 |
| 8 | 103267033 | 103267227 | -2.327744066 | BAALC-AS1 |
| 8 | 103885737 | 103885937 | 1.177668422 | RIMS2 |
| 1 | 238535838 | 238536028 | -2.9129853 | RP11-177F15.1 |
| 8 | 104627095 | 104627244 | 1.95166453 | ZFPM2 |
| 8 | 105792065 | 105792263 | -1.698443373 | ZFPM2-AS1 |
| 8 | 105792065 | 105792263 | -1.698443373 | ZFPM2 |
| 8 | 105792857 | 105793057 | -1.309523696 | ZFPM2 |

| | | | | |
|---|---|---|---|---|
| 5 | 141490719 | 141490905 | -8.53523869 | PCDHGB5 |
| 5 | 141490719 | 141490905 | -8.53523869 | PCDHGA8 |
| 5 | 141490719 | 141490905 | -8.53523869 | PCDHGC3 |
| 5 | 141490719 | 141490905 | -8.53523869 | PCDHGA10 |
| 5 | 141490719 | 141490905 | -8.53523869 | PCDHGA11 |
| 5 | 141490719 | 141490905 | -8.53523869 | PCDHGC4 |
| 5 | 141490719 | 141490905 | -8.53523869 | PCDHGB4 |
| 5 | 141490719 | 141490905 | -8.53523869 | PCDHGA1 |
| 5 | 141490719 | 141490905 | -8.53523869 | PCDHGA4 |
| 5 | 141490719 | 141490905 | -8.53523869 | PCDHGB2 |
| 5 | 141490719 | 141490905 | -8.53523869 | PCDHGA6 |
| 5 | 141490719 | 141490905 | -8.53523869 | PCDHGA7 |
| 5 | 141490719 | 141490905 | -8.53523869 | PCDHGB3 |
| 5 | 141490719 | 141490905 | -8.53523869 | PCDHGA3 |
| 5 | 141490719 | 141490905 | -8.53523869 | PCDHGA2 |
| 5 | 141490719 | 141490905 | -8.53523869 | PCDHGB6 |
| 1 | 203683076 | 203683279 | -2.025982733 | ATP2B4 |
| 5 | 141494635 | 141494843 | 1.855163824 | PCDHGA8 |
| 5 | 141494635 | 141494843 | 1.855163824 | PCDHGB2 |
| 5 | 141494635 | 141494843 | 1.855163824 | PCDHGA7 |
| 5 | 141494635 | 141494843 | 1.855163824 | PCDHGA12 |
| 5 | 141494635 | 141494843 | 1.855163824 | PCDHGA10 |
| 5 | 141494635 | 141494843 | 1.855163824 | PCDHGA11 |
| 5 | 141494635 | 141494843 | 1.855163824 | PCDHGB6 |
| 5 | 141494635 | 141494843 | 1.855163824 | PCDHGB3 |
| 5 | 141494635 | 141494843 | 1.855163824 | PCDHGA3 |
| 5 | 141494635 | 141494843 | 1.855163824 | PCDHGA9 |
| 5 | 141494635 | 141494843 | 1.855163824 | PCDHGA1 |
| 5 | 141494635 | 141494843 | 1.855163824 | PCDHGA5 |
| 5 | 141494635 | 141494843 | 1.855163824 | PCDHGA2 |
| 5 | 141494635 | 141494843 | 1.855163824 | PCDHGB1 |
| 5 | 141494635 | 141494843 | 1.855163824 | PCDHGB4 |
| 5 | 141494635 | 141494843 | 1.855163824 | PCDHGC3 |
| 5 | 141494635 | 141494843 | 1.855163824 | PCDHGC4 |
| 5 | 141494635 | 141494843 | 1.855163824 | PCDHGA6 |
| 5 | 141494635 | 141494843 | 1.855163824 | PCDHGA4 |
| 5 | 141494635 | 141494843 | 1.855163824 | PCDHGB5 |
| 5 | 141494635 | 141494843 | 1.855163824 | PCDHGB7 |
| 5 | 141494635 | 141494843 | 1.855163824 | PCDHGC5 |
| 5 | 141945241 | 141945437 | -3.770071709 | PCDH12 |
| 5 | 141956573 | 141956734 | -4.344562202 | PCDH12 |
| 5 | 142314439 | 142314628 | 1.260159707 | SPRY4 |
| 5 | 142995001 | 142995149 | 3.974183616 | ARHGAP26 |
| 5 | 143103546 | 143103739 | 2.218962749 | ARHGAP26 |
| 1 | 203739789 | 203740080 | -1.774403377 | ATP2B4 |
| 5 | 143202919 | 143203105 | 1.16387983 | ARHGAP26 |
| 5 | 143222428 | 143222625 | 1.681037703 | ARHGAP26 |
| 5 | 143223757 | 143223925 | 1.512233808 | ARHGAP26 |
| 5 | 143353288 | 143353494 | 1.520277049 | NR3C1 |
| 5 | 143400010 | 143400209 | 1.015601288 | NR3C1 |
| 5 | 143405674 | 143405872 | 6.135147663 | NR3C1 |
| 5 | 146510784 | 146510925 | -2.626732392 | TCERG1 |
| 5 | 148827443 | 148827629 | 2.505824113 | ADRB2 |
| 5 | 148827892 | 148828092 | 2.285252925 | ADRB2 |
| 6 | 15520707 | 15520899 | -2.00806696 | JARID2 |
| 6 | 15522917 | 15523108 | -1.030133152 | DTNBP1 |
| 6 | 15523936 | 15524132 | -7.666927081 | DTNBP1 |
| 6 | 15525490 | 15525672 | -8.083767876 | DTNBP1 |
| 6 | 16144921 | 16145119 | -1.398467394 | MYLIP |
| 6 | 16146690 | 16146888 | -1.097598904 | MYLIP |
| 1 | 206203164 | 206203365 | 2.070321496 | SRGAP2 |
| 1 | 206203164 | 206203365 | 2.070321496 | FAM72A |
| 6 | 16306505 | 16306713 | -1.295768219 | ATXN1 |
| 6 | 17615936 | 17616136 | -1.654407366 | NUP153 |
| 6 | 17707041 | 17707236 | 2.327832857 | RP11-500C11.3 |
| 6 | 18465267 | 18465467 | 6.934182749 | RNF144B |
| 6 | 21594036 | 21594237 | -1.590605961 | SOX4 |
| 6 | 21594036 | 21594237 | -1.590605961 | RP11-204E9.3 |
| 6 | 21594725 | 21594910 | -2.376566519 | SOX4 |
| 6 | 21594725 | 21594910 | -2.376566519 | RP11-204E9.3 |
| 6 | 21596617 | 21596815 | -1.353106066 | SOX4 |
| 6 | 25966520 | 25966720 | 1.076027586 | TRIM38 |
| 6 | 26215911 | 26216111 | 1.650805634 | HIST1H2BG |
| 1 | 206463949 | 206464164 | -1.087906929 | SRGAP2 |
| 6 | 27406202 | 27406402 | -2.359623099 | RP1-153G14.4 |
| 6 | 27451360 | 27451560 | -1.32653364 | ZNF184 |
| 6 | 27453011 | 27453221 | -1.025059124 | ZNF184 |
| 6 | 28092545 | 28092737 | -7.623277527 | ZSCAN12P1 |
| 6 | 28148482 | 28148672 | -1.794800917 | ZKSCAN8 |
| 6 | 28166516 | 28166680 | -4.94208526 | ZNF192P1 |
| 6 | 28227190 | 28227404 | -1.904958946 | ZSCAN9 |
| 6 | 28227904 | 28228104 | -1.114510873 | ZSCAN9 |
| 6 | 28232529 | 28232677 | -1.782844244 | ZSCAN9 |
| 6 | 28245374 | 28245573 | -2.584443646 | ZKSCAN4 |
| 6 | 28276225 | 28276443 | -2.088974617 | ZSCAN26 |
| 6 | 28276225 | 28276443 | -2.088974617 | RP5-874C20.8 |
| 6 | 28300826 | 28301021 | -2.084976246 | PGBD1 |
| 6 | 28365837 | 28366087 | -1.750322294 | ZKSCAN3 |
| 6 | 28380337 | 28380541 | -1.76769712 | ZSCAN12 |
| 6 | 28382286 | 28382459 | -2.203362863 | ZSCAN12 |
| 1 | 206588219 | 206588395 | 1.817591817 | EIF2D |
| 1 | 206588219 | 206588395 | 1.817591817 | RASSF5 |
| 6 | 28391293 | 28391510 | -1.899624936 | ZSCAN12 |
| 6 | 28398104 | 28398280 | -1.676625934 | ZSCAN12 |
| 6 | 28995524 | 28995723 | -1.285515796 | ZNF311 |
| 6 | 29553810 | 29554008 | 7.367751569 | OR2I1P |
| 6 | 29555944 | 29556140 | 16.60224498 | UBD |
| 6 | 29555944 | 29556140 | 16.60224498 | GABBR1 |
| 6 | 30103404 | 30103585 | 9.313268051 | TRIM31 |
| 6 | 30198838 | 30199043 | -9.289847636 | TRIM26 |
| 6 | 30278657 | 30278844 | -1.265655282 | HLA-L |
| 6 | 30278657 | 30278844 | -1.265655282 | HCG17 |
| 6 | 30287450 | 30287603 | -1.429420426 | HCG17 |
| 6 | 30287450 | 30287603 | -1.429420426 | HLA-L |
| 6 | 30287450 | 30287603 | -1.429420426 | HCG18 |
| 6 | 30298872 | 30299071 | -1.247965165 | HCG17 |
| 6 | 30298872 | 30299071 | -1.247965165 | HCG18 |
| 6 | 30300567 | 30300754 | -1.257184006 | HCG18 |
| 8 | 105792857 | 105793057 | -1.309523696 | ZFPM2-AS1 |
| 1 | 239908098 | 239908278 | -1.548423981 | CHRM3 |
| 8 | 106657455 | 106657600 | 3.698997717 | RP11-649G15.2 |
| 8 | 106657455 | 106657600 | 3.698997717 | OXR1 |
| 1 | 239909023 | 239909222 | -1.925989743 | CHRM3 |
| 1 | 241593806 | 241594069 | 1.367264304 | OPN3 |
| 1 | 241593806 | 241594069 | 1.367264304 | KMO |
| 8 | 120538538 | 120538740 | -2.168975984 | SNTB1 |
| 8 | 120538538 | 120538740 | -2.168975984 | MTBP |
| 8 | 120573594 | 120573781 | -1.129101249 | SNTB1 |
| 8 | 122877624 | 122877761 | 4.715226276 | ZHX2 |
| 8 | 124568045 | 124568183 | 7.369890305 | MTSS1 |
| 8 | 124568045 | 124568183 | 7.369890305 | NDUFB9 |
| 8 | 124832400 | 124832571 | -1.949335044 | RP11-1082L8.3 |
| 8 | 124976462 | 124976650 | 1.810744188 | ZNF572 |
| 8 | 127890673 | 127890852 | -1.180513957 | PVT1_3 |
| 8 | 127890673 | 127890852 | -1.180513957 | PVT1 |
| 8 | 128020382 | 128020572 | 2.625859546 | PVT1 |
| 1 | 242376974 | 242377148 | 1.453445298 | PLD5 |
| 1 | 242376974 | 242377148 | 1.453445298 | RP11-513D4.1 |
| 8 | 129548322 | 129548523 | -4.074064939 | CCDC26 |
| 8 | 129628671 | 129628867 | -1.27344723 | CCDC26 |
| 8 | 131962287 | 131962426 | 6.215226276 | EFR3A |
| 8 | 132123505 | 132123676 | 1.977098155 | KCNQ3 |
| 8 | 132129452 | 132129655 | 1.316850404 | KCNQ3 |
| 8 | 133037906 | 133038232 | 2.529393085 | TG |
| 8 | 133037906 | 133038232 | 2.529393085 | SLA |
| 8 | 134602164 | 134602269 | -5.642701109 | ZFAT |
| 8 | 139742282 | 139742467 | 6.151033836 | TRAPPC9 |
| 8 | 139796981 | 139797180 | -1.16107653 | TRAPPC9 |
| 8 | 139989440 | 139989554 | 4.425874521 | TRAPPC9 |
| 8 | 140659424 | 140659559 | -2.066699563 | PTK2 |
| 1 | 243185914 | 243186113 | -1.869777568 | CEP170 |
| 8 | 141129013 | 141129237 | 6.04637857 | DENND3 |
| 8 | 141129013 | 141129237 | 6.04637857 | RP11-809O17.1 |
| 8 | 141129525 | 141129696 | 5.96868691 | DENND3 |
| 8 | 141129525 | 141129696 | 5.96868691 | RP11-809O17.1 |
| 8 | 141173171 | 141173359 | -1.00736179 | DENND3 |
| 8 | 141252383 | 141252581 | 6.40024775 | SLC45A4 |
| 8 | 141252383 | 141252581 | 6.40024775 | RP11-10J21.4 |
| 8 | 142271015 | 142271216 | -1.002694391 | TSNARE1 |
| 8 | 142273138 | 142273345 | -1.186242774 | TSNARE1 |
| 1 | 243303055 | 243303239 | -1.041109167 | SDCCAG8 |
| 8 | 143008964 | 143009151 | 2.175248624 | RP11-273G15.2 |
| 8 | 143250467 | 143250661 | 1.058563504 | RP13-582O9.6 |
| 8 | 143250467 | 143250661 | 1.058563504 | ZFP41 |
| 8 | 143295910 | 143296098 | 2.550909957 | ZNF696 |
| 8 | 143560248 | 143560447 | 1.162766242 | GSDMD |
| 8 | 143573580 | 143573777 | 1.167967476 | RP11-661A12.9 |
| 8 | 143587928 | 143588087 | 8.124139845 | EEF1D |
| 8 | 143589780 | 143589979 | 2.053785397 | EEF1D |
| 8 | 143596800 | 143597035 | 1.157131062 | EEF1D |
| 8 | 143599505 | 143599703 | 1.796505363 | EEF1D |
| 8 | 143599505 | 143599703 | 1.796505363 | TIGD5 |

| | | | | |
|---|---|---|---|---|
| 5 | 149981325 | 149981599 | -1.721435689 | SLC26A2 |
| 5 | 150053572 | 150053769 | -5.746677561 | CSF1R |
| 5 | 150520928 | 150521082 | 7.426800982 | NDST1 |
| 5 | 150553225 | 150553438 | 3.10860495 | NDST1 |
| 5 | 150779091 | 150779253 | -8.000923987 | SMIM3 |
| 5 | 150793756 | 150793940 | 5.256863798 | SMIM3 |
| 5 | 151030416 | 151030624 | -1.729186037 | TNIP1 |
| 5 | 151746282 | 151746476 | -1.685375819 | ATOX1 |
| 5 | 154802131 | 154802296 | -1.063872191 | LARP1 |
| 5 | 154814658 | 154814867 | -2.208930026 | LARP1 |
| 5 | 154815623 | 154815822 | -1.959256477 | LARP1 |
| 5 | 154887963 | 154888158 | -2.289742297 | GEMIN5 |
| 5 | 154891336 | 154891535 | -2.615108531 | GEMIN5 |
| 5 | 157787676 | 157787878 | -1.480755778 | CLINT1 |
| 5 | 160078018 | 160078219 | -1.618813287 | PWWP2A |
| 5 | 160092920 | 160093121 | -1.757255051 | PWWP2A |
| 5 | 160093604 | 160093804 | -1.371506923 | PWWP2A |
| 5 | 162153203 | 162153347 | 2.427611164 | GABRG2 |
| 5 | 169825281 | 169825430 | -2.710731884 | DOCK2 |
| 5 | 170233173 | 170233331 | 3.733038231 | C5orf58 |
| 5 | 170240060 | 170240267 | 3.085872719 | C5orf58 |
| 5 | 170242721 | 170242921 | 2.514772027 | C5orf58 |
| 5 | 170243510 | 170243747 | 3.042527539 | C5orf58 |
| 1 | 9367340 | 9367552 | 5.29721744 | SPSB1 |
| 5 | 170244795 | 170244977 | 2.565109764 | C5orf58 |
| 5 | 170248582 | 170248772 | 1.384498635 | LCP2 |
| 5 | 170248582 | 170248772 | 1.384498635 | C5orf58 |
| 5 | 170979940 | 170980102 | -2.106987657 | RANBP17 |
| 5 | 172254415 | 172254588 | -1.376760942 | UBTD2 |
| 5 | 172254415 | 172254588 | -1.376760942 | KLF3P1 |
| 5 | 172257745 | 172257915 | -1.872826212 | UBTD2 |
| 5 | 172334320 | 172334520 | 2.588010988 | SH3PXD2B |
| 5 | 172337975 | 172338155 | 1.922749933 | SH3PXD2B |
| 5 | 172768596 | 172768799 | 5.470018948 | RP11-779O18.3 |
| 5 | 172768596 | 172768799 | 5.470018948 | DUSP1 |
| 5 | 173034541 | 173034760 | -1.428401745 | ATP6V0E1 |
| 5 | 173090788 | 173090997 | 1.656934377 | CREBRF |
| 5 | 173607957 | 173608160 | -1.219621918 | BOD1 |
| 5 | 173888602 | 173888809 | 1.726138231 | CPEB4 |
| 5 | 175682975 | 175683182 | -2.456685302 | HRH2 |
| 5 | 175684024 | 175684196 | -2.535039218 | HRH2 |
| 5 | 175684924 | 175685106 | -1.954112202 | HRH2 |
| 5 | 175685831 | 175686000 | -1.738545692 | HRH2 |
| 5 | 175707968 | 175708076 | -2.375432598 | HRH2 |
| 5 | 176289681 | 176289879 | -2.634516576 | SIMC1 |
| 5 | 176303914 | 176304106 | -8.546270187 | SIMC1 |
| 5 | 176506838 | 176507042 | -1.251148441 | FAF2 |
| 5 | 177135601 | 177135812 | -2.242383137 | NSD1 |
| 5 | 177477947 | 177478142 | -2.695158749 | RP11-1334A24.5 |
| 5 | 177613111 | 177613280 | 1.342296511 | RP11-1277A3.1 |
| 5 | 177615553 | 177615731 | -2.008020402 | RP11-1277A3.1 |
| 5 | 177617897 | 177618082 | -2.205940409 | RP11-1277A3.1 |
| 5 | 177618989 | 177619186 | -3.579386366 | RP11-1277A3.1 |
| 5 | 177618989 | 177619186 | -3.579386366 | RP11-1277A3.2 |
| 6 | 30300567 | 30300754 | -1.257184006 | HCG17 |
| 6 | 30561335 | 30561532 | -1.628709903 | PRR3 |
| 6 | 30562409 | 30562609 | -2.290352932 | PRR3 |
| 1 | 206960782 | 206960980 | 4.660163437 | FCAMR |
| 6 | 31159226 | 31159424 | -1.707581162 | TCF19 |
| 6 | 31394498 | 31394697 | 5.927628914 | XXbac-BPG181B23.7 |
| 6 | 31410679 | 31410828 | 1.075289867 | HCP5 |
| 6 | 31410679 | 31410828 | 1.075289867 | MICA |
| 1 | 207049396 | 207049603 | -1.506085402 | YOD1 |
| 1 | 207049396 | 207049603 | -1.506085402 | PFKFB2 |
| 6 | 31577561 | 31577782 | 7.754068508 | TNF |
| 6 | 31629515 | 31629715 | -6.658352896 | PRRC2A |
| 6 | 31811153 | 31811362 | 1.340131276 | HSPA1L |
| 6 | 31817392 | 31817623 | 2.189300147 | HSPA1A |
| 6 | 31828539 | 31828711 | 3.574915061 | HSPA1B |
| 6 | 31829608 | 31829813 | 1.272383834 | HSPA1B |
| 6 | 32129227 | 32129427 | -1.026747013 | FKBPL |
| 6 | 32828779 | 32828973 | 2.613282342 | XXbac-BPG246D15.9 |
| 6 | 32828779 | 32828973 | 2.613282342 | TAP2 |
| 6 | 32838031 | 32838180 | 6.202570732 | TAP2 |
| 6 | 32838031 | 32838180 | 6.202570732 | XXbac-BPG246D15.9 |
| 6 | 32845337 | 32845528 | 3.060731533 | PSMB9 |
| 6 | 32845337 | 32845528 | 3.060731533 | TAP1 |
| 6 | 32845337 | 32845528 | 3.060731533 | PSMB8-AS1 |
| 6 | 32845785 | 32845980 | -11.45778726 | PSMB8-AS1 |
| 6 | 32845785 | 32845980 | -11.45778726 | TAP1 |
| 6 | 32845785 | 32845980 | -11.45778726 | PSMB9 |
| 1 | 209675902 | 209676104 | 1.802759057 | G0S2 |
| 1 | 209675902 | 209676104 | 1.802759057 | RP1-28O10.1 |
| 6 | 33277786 | 33277981 | -2.13579821 | B3GALT4 |
| 6 | 33278524 | 33278708 | -1.730029371 | B3GALT4 |
| 6 | 33305088 | 33305300 | 1.079300903 | TAPBP |
| 6 | 33320980 | 33321231 | 1.27055072 | DAXX |
| 6 | 33391409 | 33391558 | -1.034048758 | KIFC1 |
| 6 | 33403826 | 33404025 | -2.433846299 | KIFC1 |
| 6 | 33404831 | 33404953 | -8.474616988 | KIFC1 |
| 6 | 33452037 | 33452217 | -1.123528977 | SYNGAP1 |
| 6 | 33452037 | 33452217 | -1.123528977 | XXbac-BPG294E21.9 |
| 6 | 33456395 | 33456606 | -1.178458184 | ZBTB9 |
| 6 | 33456395 | 33456606 | -1.178458184 | SYNGAP1 |
| 6 | 34270331 | 34270520 | 1.020441528 | RP11-513I15.6 |
| 6 | 34279540 | 34279740 | -2.142750848 | NUDT3 |
| 6 | 34279540 | 34279740 | -2.142750848 | RP11-513I15.6 |
| 6 | 34280149 | 34280345 | -1.099069706 | RP11-513I15.6 |
| 6 | 34280149 | 34280345 | -1.099069706 | NUDT3 |
| 6 | 34859080 | 34859306 | 1.116941218 | UHRF1BP1 |
| 1 | 209851181 | 209851319 | -1.356670248 | DIEXF |
| 6 | 34878361 | 34878599 | -1.101892041 | UHRF1BP1 |
| 6 | 34878361 | 34878599 | -1.101892041 | TAF11 |
| 1 | 211313639 | 211313872 | -2.041353319 | RCOR3 |
| 6 | 35455936 | 35456148 | -2.171498857 | FANCE |
| 6 | 35575665 | 35575861 | 1.507284464 | FKBP5 |
| 6 | 35869582 | 35869787 | -1.398489645 | SRPK1 |
| 6 | 36084566 | 36084772 | -1.443413291 | MAPK14 |
| 8 | 143726048 | 143726248 | 4.117836296 | FAM83H |
| 1 | 1014125 | 1014325 | 6.711886243 | ISG15 |
| 1 | 243685247 | 243685380 | -5.488639962 | AKT3 |
| 8 | 143744738 | 143744856 | 1.053628355 | FAM83H-AS1 |
| 8 | 143770406 | 143770587 | 6.221212597 | RP11-429J17.7 |
| 8 | 143770406 | 143770587 | 6.221212597 | FAM83H-AS1 |
| 8 | 143916432 | 143916629 | 8.822301352 | PLEC |
| 8 | 143922948 | 143923148 | 12.9272785 | PLEC |
| 1 | 243730449 | 243730649 | 1.10128549 | RP11-370K11.1 |
| 1 | 243730449 | 243730649 | 1.10128549 | AKT3 |
| 8 | 144080353 | 144080538 | 1.253961346 | EXOSC4 |
| 8 | 144263475 | 144263696 | 1.244764152 | BOP1 |
| 8 | 144268114 | 144268314 | 4.72204634 | SCX |
| 8 | 144268114 | 144268314 | 4.72204634 | BOP1 |
| 8 | 144292687 | 144292886 | 4.408491812 | HSF1 |
| 8 | 144298636 | 144298828 | -6.933735903 | HSF1 |
| 8 | 144310522 | 144310709 | 7.954219767 | HSF1 |
| 8 | 144415123 | 144415303 | 3.231068041 | SLC39A4 |
| 8 | 144496536 | 144496734 | 1.10879995 | CTD-2517M22.14 |
| 8 | 144496536 | 144496734 | 1.10879995 | PPP1R16A |
| 8 | 144509120 | 144509318 | 3.484151164 | MFSD3 |
| 8 | 144523099 | 144523213 | 3.630199766 | LRRC24 |
| 8 | 144523099 | 144523213 | 3.630199766 | LRRC14 |
| 8 | 144547291 | 144547489 | 6.846525648 | ARHGAP39 |
| 1 | 244401416 | 244401571 | -3.583782864 | RP11-518L10.5 |
| 8 | 144548068 | 144548249 | 5.149604009 | ARHGAP39 |
| 8 | 144742692 | 144742890 | -1.139736076 | ZNF251 |
| 8 | 144808311 | 144808560 | 1.612037952 | ZNF517 |
| 9 | 116629 | 116821 | 1.020875993 | FOXD4 |
| 9 | 306981 | 307165 | -7.124037905 | DOCK8 |
| 9 | 1047282 | 1047473 | -3.375493032 | LINC01230 |
| 1 | 244656943 | 244657125 | -7.91585155 | DESI2 |
| 9 | 2655098 | 2655234 | -1.154195071 | VLDLR |
| 9 | 3530417 | 3530641 | -1.100439013 | RFX3-AS1 |
| 9 | 3675231 | 3675353 | 1.22712569 | RFX3-AS1 |
| 9 | 4662718 | 4662835 | 2.428882341 | SPATA6L |
| 9 | 4662718 | 4662835 | 2.428882341 | PLPP6 |
| 9 | 4726475 | 4726635 | -1.098250327 | AK3 |
| 1 | 244701321 | 244701454 | 1.346482299 | DESI2 |
| 9 | 5113478 | 5113674 | 1.167569923 | IGHEP2 |
| 9 | 5113478 | 5113674 | 1.167569923 | JAK2 |
| 9 | 5133836 | 5134036 | -1.086907959 | INSL6 |
| 9 | 5138350 | 5138451 | -1.423421606 | INSL6 |
| 9 | 5179712 | 5179911 | -3.903677461 | INSL6 |
| 9 | 5467826 | 5468026 | 2.50050311 | CD274 |
| 9 | 5569975 | 5570179 | 1.460687077 | PDCD1LG2 |
| 9 | 5897637 | 5897833 | -3.714867753 | KIAA2026 |
| 9 | 5897637 | 5897833 | -3.714867753 | MLANA |
| 1 | 245010214 | 245010367 | -3.123963757 | EFCAB2 |
| 9 | 5988462 | 5988626 | 6.28788779 | KIAA2026 |
| 9 | 6007543 | 6007739 | 9.060655456 | KIAA2026 |
| 9 | 6433887 | 6434085 | -1.118228049 | RP11-307L3.4 |
| 9 | 6433887 | 6434085 | -1.118228049 | UHRF2 |
| 9 | 6460497 | 6460668 | -8.668758688 | UHRF2 |

| | | | | |
|---|---|---|---|---|
| 5 | 177632521 | 177632759 | -2.441896994 | RP11-1277A3.2 |
| 5 | 178121604 | 178121801 | 4.060730277 | N4BP3 |
| 5 | 178148212 | 178148411 | -1.170245152 | RMND5B |
| 5 | 178237925 | 178238124 | -1.503824715 | COL23A1 |
| 5 | 178238227 | 178238402 | -1.728669614 | COL23A1 |
| 5 | 178287248 | 178287443 | -7.666804808 | COL23A1 |
| 5 | 178313436 | 178313618 | -3.950196324 | COL23A1 |
| 5 | 178350216 | 178350404 | -9.304185172 | COL23A1 |
| 5 | 178366059 | 178366253 | -4.243093237 | COL23A1 |
| 5 | 178394741 | 178394889 | -9.035799265 | COL23A1 |
| 5 | 178507261 | 178507545 | -3.871077921 | COL23A1 |
| 5 | 178508041 | 178508169 | -3.948252987 | COL23A1 |
| 5 | 178508773 | 178508969 | -4.66897531 | COL23A1 |
| 5 | 178514091 | 178514285 | -3.940416216 | COL23A1 |
| 5 | 179192815 | 179192965 | -6.813310826 | ADAMTS2 |
| 5 | 179193547 | 179193746 | -5.954946542 | ADAMTS2 |
| 5 | 179575335 | 179575535 | -1.143507535 | RUFY1 |
| 5 | 179606934 | 179607144 | -2.715591328 | RUFY1 |
| 5 | 179765630 | 179765932 | -1.072956675 | MAML1 |
| 5 | 179774683 | 179774882 | -1.814227137 | MAML1 |
| 5 | 179834758 | 179834948 | -7.948355408 | SQSTM1 |
| 5 | 179883848 | 179884051 | -2.826445575 | TBC1D9B |
| 5 | 180574119 | 180574321 | -1.507773871 | CNOT6 |
| 5 | 180830521 | 180830717 | -1.332011903 | HEIH |
| 6 | 326876 | 327013 | -4.488639962 | DUSP22 |
| 6 | 407744 | 407929 | 4.5 | IRF4 |
| 6 | 1394811 | 1394992 | -3.094000019 | FOXF2 |
| 6 | 1610739 | 1610924 | -1.640816592 | FOXC1 |
| 6 | 2272871 | 2273073 | -1.500236643 | GMDS-AS1 |
| 6 | 2288998 | 2289111 | -1.429542521 | GMDS-AS1 |
| 6 | 2291487 | 2291691 | -1.616904183 | GMDS-AS1 |
| 6 | 2292258 | 2292454 | -2.128693921 | GMDS-AS1 |
| 6 | 2301831 | 2301978 | -1.226537879 | GMDS-AS1 |
| 6 | 2306378 | 2306577 | -1.860411083 | GMDS-AS1 |
| 6 | 2841184 | 2841383 | -9.456958931 | SERPINB1 |
| 6 | 3003754 | 3003939 | -1.365699575 | NQO2 |
| 6 | 3020030 | 3020232 | 2.063506103 | NQO2 |
| 6 | 3023513 | 3023656 | 1.088623818 | HTATSF1P2 |
| 6 | 3023513 | 3023656 | 1.088623818 | NQO2 |
| 6 | 3025791 | 3025993 | 2.427849657 | NQO2 |
| 6 | 5045259 | 5045372 | -3.672325495 | RP11-428J1.4 |
| 6 | 5084827 | 5085019 | -3.585485475 | PPP1R3G |
| 6 | 5086463 | 5086635 | -3.355502861 | PPP1R3G |
| 6 | 5261826 | 5262019 | -7.263921749 | FARS2 |
| 6 | 5368851 | 5369055 | -1.508880574 | FARS2 |
| 6 | 5998339 | 5998549 | -2.249780637 | NRN1 |
| 6 | 6685464 | 6685664 | -2.594516912 | RP1-80N2.4 |
| 6 | 6795364 | 6795556 | -5.281613271 | BTF3P7 |
| 6 | 7229113 | 7229312 | -1.001021365 | RREB1 |
| 6 | 7231610 | 7231760 | -1.137500914 | RREB1 |
| 6 | 7246517 | 7246738 | -1.726790414 | RREB1 |
| 6 | 7342607 | 7342801 | -8.015915204 | SSR1 |
| 6 | 7342607 | 7342801 | -8.015915204 | CAGE1 |
| 1 | 205663231 | 205663429 | 2.804545386 | SLC45A3 |
| 6 | 36088042 | 36088254 | -1.510134888 | MAPK14 |
| 6 | 36139284 | 36139451 | -2.070181589 | MAPK13 |
| 6 | 36158392 | 36158593 | -1.047881501 | RP1-179N16.6 |
| 6 | 36363281 | 36363446 | 6.061737399 | ETV7 |
| 6 | 36366386 | 36366580 | 8.089952116 | ETV7 |
| 6 | 36386864 | 36387007 | 7.595953714 | ETV7 |
| 6 | 36386864 | 36387007 | 7.595953714 | RP1-50J22.4 |
| 6 | 36684293 | 36684495 | -8.868128288 | CDKN1A |
| 6 | 36721593 | 36721770 | -1.130695226 | RAB44 |
| 6 | 36721973 | 36722218 | -1.788826307 | RAB44 |
| 6 | 37174077 | 37174274 | 1.837384481 | PIM1 |
| 6 | 37212347 | 37212545 | 3.374537998 | TMEM217 |
| 6 | 37218505 | 37218699 | 3.650483849 | TMEM217 |
| 6 | 37637193 | 37637398 | 2.319901069 | MDGA1 |
| 6 | 37696505 | 37696698 | 7.574436873 | MDGA1 |
| 6 | 38087117 | 38087314 | -7.505832026 | ZFAND3 |
| 6 | 39105500 | 39105724 | -1.204510362 | SAYSD1 |
| 6 | 41158734 | 41158937 | -5.614823626 | TREM2 |
| 6 | 41598977 | 41599175 | 1.14508071 | FOXP4 |
| 6 | 42228350 | 42228550 | -3.701569039 | TRERF1 |
| 6 | 42268327 | 42268529 | -2.577554502 | TRERF1 |
| 6 | 42269421 | 42269618 | -2.826203232 | TRERF1 |
| 6 | 42829599 | 42829810 | -2.370912585 | GLTSCR1L |
| 6 | 42864915 | 42865032 | -2.056516727 | GLTSCR1L |
| 6 | 42865085 | 42865282 | -2.046741633 | GLTSCR1L |
| 6 | 42938987 | 42939188 | -1.134930711 | CNPY3 |
| 6 | 43337555 | 43337800 | -2.520835153 | ZNF318 |
| 6 | 43339988 | 43340191 | -2.439728683 | ZNF318 |
| 6 | 43348439 | 43348644 | -2.638379715 | ZNF318 |
| 6 | 43355809 | 43356010 | -2.528580124 | ZNF318 |
| 6 | 43357413 | 43357631 | -2.880091613 | ZNF318 |
| 1 | 212625555 | 212625748 | 1.303503583 | FAM71A |
| 1 | 212625555 | 212625748 | 1.303503583 | RP11-338C15.5 |
| 6 | 43454798 | 43455002 | 1.366025062 | DLK2 |
| 6 | 43625233 | 43625437 | 2.496802571 | GTPBP2 |
| 6 | 43671593 | 43671795 | 8.559206956 | MRPS18A |
| 6 | 43671593 | 43671795 | 8.559206956 | RSPH9 |
| 6 | 43785162 | 43785434 | -1.466865862 | VEGFA |
| 6 | 43996657 | 43996844 | -5.642701109 | RP5-1120P11.1 |
| 6 | 44002882 | 44003082 | -1.532796422 | RP5-1120P11.1 |
| 6 | 44002882 | 44003082 | -1.532796422 | C6orf223 |
| 6 | 44257802 | 44258002 | 1.022015933 | SLC35B2 |
| 6 | 44258792 | 44258992 | 1.139912644 | NFKBIE |
| 6 | 44299707 | 44299901 | -1.225813068 | RP11-444E17.6 |
| 6 | 44299707 | 44299901 | -1.225813068 | AARS2 |
| 6 | 44299707 | 44299901 | -1.225813068 | TMEM151B |
| 6 | 44938088 | 44938219 | -5.528995861 | SUPT3H |
| 6 | 44938088 | 44938219 | -5.528995861 | AL161905.1 |
| 6 | 44940834 | 44941016 | -1.782001441 | SUPT3H |
| 6 | 44940834 | 44941016 | -1.782001441 | AL161905.1 |
| 6 | 45421763 | 45421937 | -5.086838568 | RP1-244F24.1 |
| 6 | 45421763 | 45421937 | -5.086838568 | RUNX2 |
| 6 | 46139975 | 46140290 | 1.082051881 | ENPP4 |
| 6 | 46140905 | 46141076 | 1.21917323 | ENPP4 |
| 9 | 6835202 | 6835413 | 1.091923689 | ACTG1P14 |
| 9 | 6835202 | 6835413 | 1.091923689 | RP11-146B14.1 |
| 9 | 6835202 | 6835413 | 1.091923689 | AL513412.1 |
| 9 | 6835202 | 6835413 | 1.091923689 | KDM4C |
| 1 | 245091241 | 245091453 | -1.801176319 | EFCAB2 |
| 9 | 20919895 | 20920071 | -5.283978038 | FOCAD |
| 9 | 20937514 | 20937714 | 1.900382085 | FOCAD |
| 9 | 21005350 | 21005504 | -2.707442974 | HACD4 |
| 9 | 21022190 | 21022377 | -1.787435016 | HACD4 |
| 9 | 21022917 | 21023107 | -1.377706997 | HACD4 |
| 9 | 27330304 | 27330489 | 3.453445298 | MOB3B |
| 9 | 27455274 | 27455600 | 1.828999111 | MOB3B |
| 1 | 245672071 | 245672239 | -5.330889049 | KIF26B |
| 9 | 32431347 | 32431547 | -1.741388171 | ACO1 |
| 9 | 32446367 | 32446565 | -1.284947945 | ACO1 |
| 9 | 33602325 | 33602477 | -1.220939173 | CYP4F26P |
| 9 | 34214669 | 34214919 | -7.469605941 | UBAP1 |
| 1 | 245687312 | 245687481 | -1.437234559 | KIF26B |
| 9 | 34367376 | 34367586 | 4.065605284 | KIAA1161 |
| 9 | 34370741 | 34370845 | 5.391106664 | KIAA1161 |
| 9 | 34400582 | 34400798 | 1.08299649 | FAM219A |
| 9 | 34610826 | 34611025 | 1.41686115 | RPP25L |
| 9 | 34635466 | 34635670 | 1.201690847 | SIGMAR1 |
| 9 | 35649668 | 35649829 | 3.76128115 | SIT1 |
| 9 | 36170527 | 36170729 | -1.016462597 | CCIN |
| 9 | 36839531 | 36839722 | 1.102679889 | PAX5 |
| 9 | 37503228 | 37503428 | -1.509884288 | POLR1E |
| 9 | 37765322 | 37765468 | -1.107864346 | TRMT10B |
| 9 | 37765322 | 37765468 | -1.107864346 | RP11-613M10.9 |
| 9 | 37766008 | 37766135 | -5.238373102 | TRMT10B |
| 9 | 37766008 | 37766135 | -5.238373102 | RP11-613M10.9 |
| 9 | 38068817 | 38068991 | 10.79647644 | SHB |
| 9 | 38396117 | 38396315 | 1.412891086 | ALDH1B1 |
| 9 | 38397096 | 38397289 | 1.651259692 | ALDH1B1 |
| 1 | 246254191 | 246254375 | -6.491051153 | SMYD3 |
| 9 | 38553897 | 38554095 | 1.390638741 | RP11-392E22.11 |
| 9 | 38553897 | 38554095 | 1.390638741 | RP11-392E22.9 |
| 9 | 38566467 | 38566704 | 1.235869819 | RP11-392E22.11 |
| 9 | 38566467 | 38566704 | 1.235869819 | RP11-392E22.9 |
| 9 | 38566467 | 38566704 | 1.235869819 | SNX18P3 |
| 9 | 38567530 | 38567698 | 1.68639872 | RP11-392E22.11 |
| 9 | 38567530 | 38567698 | 1.68639872 | RP11-392E22.9 |
| 9 | 38567530 | 38567698 | 1.68639872 | SNX18P3 |
| 9 | 38568712 | 38568919 | 1.505311823 | RP11-392E22.9 |
| 9 | 38595678 | 38595878 | 1.053661813 | ANKRD18A |
| 9 | 38620410 | 38620577 | 3.417288195 | ANKRD18A |
| 9 | 38620410 | 38620577 | 3.417288195 | FAM201A |
| 9 | 38622192 | 38622509 | 1.910858551 | FAM201A |
| 1 | 246257207 | 246257392 | -7.501407508 | SMYD3 |
| 9 | 41023412 | 41023559 | -2.776667366 | FRG1HP |
| 9 | 41023412 | 41023559 | -2.776667366 | PGM5P2 |
| 9 | 41062124 | 41062289 | -6.09722454 | RP11-87H9.5 |
| 9 | 41062124 | 41062289 | -6.09722454 | PGM5P2 |
| 9 | 41062124 | 41062289 | -6.09722454 | FRG1HP |

| | | | | | | | | | | | | | | |
|---|---|---|---|---|---|---|---|---|---|---|---|---|---|---|
| 6 | 11185231 | 11185433 | 3.107628944 | NEDD9 | 6 | 47232767 | 47233011 | -1.497583408 | TNFRSF21 | 1 | 246271045 | 246271174 | -2.044653271 | SMYD3 |
| 6 | 11185231 | 11185433 | 3.107628944 | RP3-510L9.1 | 6 | 47284074 | 47284274 | -1.148987301 | TNFRSF21 | 9 | 62812986 | 62813246 | 1.362659042 | LINC01410 |
| 6 | 11190299 | 11190500 | 2.662810798 | NEDD9 | 6 | 47286218 | 47286439 | -1.260723368 | TNFRSF21 | 9 | 63818002 | 63818170 | 2.527002857 | DUX4L50 |
| 6 | 11190299 | 11190500 | 2.662810798 | RP3-510L9.1 | 1 | 212726574 | 212726751 | -7.184662351 | NSL1 | 9 | 65291878 | 65292076 | -1.217771569 | RP11-15J10.1 |
| 6 | 11351786 | 11351987 | 1.413978295 | NEDD9 | 6 | 51410058 | 51410288 | 1.794445551 | RP3-437C15.1 | 9 | 65730219 | 65730371 | 2.147878709 | CBWD5 |
| 6 | 11548543 | 11548658 | -1.107721353 | TMEM170B | 6 | 52502363 | 52502563 | -1.359385276 | EFHC1 | 9 | 66919568 | 66919739 | 6.464629204 | ZNF658 |
| 6 | 12120248 | 12120469 | 1.096837796 | HIVEP1 | 6 | 52502363 | 52502563 | -1.359385276 | TRAM2 | 9 | 68780279 | 68780458 | 7.675072492 | FAM122A |
| 6 | 12121148 | 12121343 | 1.128299164 | HIVEP1 | 6 | 52667336 | 52667528 | -3.827573675 | RP1-152L7.5 | 9 | 68780279 | 68780458 | 7.675072492 | PIP5K1B |
| 6 | 12122555 | 12122722 | 1.695567079 | HIVEP1 | 6 | 53552248 | 53552438 | -1.271405133 | GCLC | 9 | 68782493 | 68782637 | -1.289862926 | PIP5K1B |
| 6 | 12161580 | 12161809 | 1.304003727 | HIVEP1 | 1 | 944187 | 944287 | -2.188675786 | NOC2L | 9 | 68782493 | 68782637 | -1.289862926 | FAM122A |
| 1 | 205810790 | 205810988 | -1.31994821 | SLC41A1 | 1 | 944187 | 944287 | -2.188675786 | SAMD11 | 9 | 69254360 | 69254557 | 2.117092517 | TJP2 |
| 6 | 13287435 | 13287650 | -2.574796262 | PHACTR1 | 6 | 56552310 | 56552554 | -1.159940133 | DST | 9 | 70413093 | 70413320 | 1.192656196 | KLF9 |
| 6 | 13287435 | 13287650 | -2.574796262 | RP1-257A7.4 | 1 | 213001870 | 213002124 | -1.317572684 | ANGEL2 | 9 | 71745276 | 71745475 | 1.444759277 | TMEM2 |
| 6 | 13287435 | 13287650 | -2.574796262 | TBC1D7 | 6 | 57147591 | 57147798 | -1.405147071 | RP11-203B9.4 | 9 | 76503410 | 76503606 | 1.125393716 | GCNT1 |
| 6 | 13364728 | 13364932 | 1.774752213 | GFOD1 | 6 | 57147591 | 57147798 | -1.405147071 | ZNF451 | 9 | 77373591 | 77373783 | -6.254392581 | VPS13A |
| 6 | 13486734 | 13486933 | 1.678020756 | GFOD1 | 6 | 63576392 | 63576599 | -1.143343199 | PTP4A1 | 9 | 77414149 | 77414289 | -2.289891131 | VPS13A |
| 6 | 13486734 | 13486933 | 1.678020756 | GFOD1-AS1 | 6 | 63684756 | 63684957 | -1.445713994 | PHF3 | 9 | 77721352 | 77721554 | -1.126747018 | GNAQ |
| 6 | 13604820 | 13604980 | -1.534270452 | SIRT5 | 6 | 63685462 | 63685670 | -1.538992788 | PHF3 | 9 | 78329180 | 78329364 | -2.074185693 | PSAT1 |
| 6 | 14135576 | 14135781 | 5.051511829 | CD83 | 6 | 63711605 | 63711801 | -1.937889456 | PHF3 | 9 | 81633996 | 81634190 | 1.014420416 | TLE1 |
| 1 | 205978340 | 205978541 | 2.005440482 | RAB7B | 6 | 63713322 | 63713544 | -1.419014436 | PHF3 | 9 | 81689158 | 81689372 | 2.747018788 | TLE1 |
| 6 | 14910615 | 14910772 | -2.701964871 | RP11-146I2.1 | 6 | 70758059 | 70758249 | -5.836212671 | SMAP1 | 9 | 86048147 | 86048266 | -4.649604009 | GOLM1 |
| 1 | 9613127 | 9613388 | -1.729073495 | TMEM201 | 1 | 214383388 | 214383585 | -3.08043539 | PTPN14 | 9 | 86048897 | 86049094 | 6.079778014 | GOLM1 |
| 1 | 205979360 | 205979471 | -5.056219753 | RAB7B | 6 | 71319058 | 71319260 | 1.058352037 | RP3-331H24.5 | 9 | 87706709 | 87706931 | -1.24016874 | DAPK1 |
| 6 | 15500901 | 15501132 | 1.173275909 | JARID2 | 6 | 71319058 | 71319260 | 1.058352037 | RP11-154D6.1 | 1 | 246987138 | 246987338 | -7.090925534 | ZNF670-ZNF695 |
| 6 | 15520707 | 15520899 | 1.604479572 | JARID2 | 6 | 72831169 | 72831478 | 3.218702656 | KCNQ5 | 1 | 246987138 | 246987338 | -7.090925534 | ZNF695 |
| 6 | 15525490 | 15525672 | 8.083767876 | DTNBP1 | 6 | 73266941 | 73267085 | -4.281502777 | RP11-398K22.12 | 9 | 88996363 | 88996559 | -5.551643904 | S1PR3 |
| 6 | 16144921 | 16145119 | -1.237310098 | MYLIP | 6 | 73266941 | 73267085 | -4.281502777 | KHDC1 | 9 | 89001619 | 89001814 | -2.611583511 | S1PR3 |
| 6 | 16146690 | 16146888 | -1.495004288 | MYLIP | 6 | 73290899 | 73291091 | -5.564641508 | KHDC1 | 9 | 89361460 | 89361656 | 2.976481355 | SEMA4D |
| 1 | 206203164 | 206203365 | 1.041722073 | SRGAP2 | 6 | 73290899 | 73291091 | -5.564641508 | RP11-398K22.12 | 9 | 89606251 | 89606423 | 1.306488438 | GADD45G |
| 1 | 206203164 | 206203365 | 1.041722073 | FAM72A | 6 | 73290899 | 73291091 | -5.564641508 | RPSAP41 | 9 | 91723334 | 91723507 | -1.365378809 | ROR2 |
| 6 | 16306505 | 16306713 | 1.276412709 | ATXN1 | 6 | 73299258 | 73299467 | -2.976755879 | RP11-398K22.12 | 9 | 92083217 | 92083359 | -6.530347966 | SPTLC1 |
| 6 | 16762630 | 16762829 | 2.687682092 | RP1-151F17.1 | 6 | 73299258 | 73299467 | -2.976755879 | KHDC1 | 9 | 92226237 | 92226383 | 6.014988673 | IARS |
| 6 | 17615936 | 17616136 | -1.437230981 | NUP153 | 6 | 73425362 | 73425565 | 1.785498072 | MB21D1 | 9 | 92614118 | 92614318 | 1.967781493 | IPPK |
| 6 | 17628923 | 17629174 | -2.101727456 | NUP153 | 1 | 214640235 | 214640540 | -7.320912986 | CENPF | 9 | 92614118 | 92614318 | 1.967781493 | CENPP |
| 6 | 17637316 | 17637517 | -1.09424779 | NUP153 | 6 | 78927561 | 78927682 | 1.860793747 | IRAK1BP1 | 9 | 92631987 | 92632162 | 3.83974005 | IPPK |
| 6 | 17707041 | 17707236 | -1.02215175 | RP11-500C11.3 | 6 | 79201626 | 79201802 | -6.948260811 | HMGN3 | 9 | 92832229 | 92832396 | -5.612603718 | ANKRD19P |
| 6 | 17764311 | 17764516 | 1.709013257 | KIF13A | 1 | 214642140 | 214642340 | -5.149604009 | CENPF | 9 | 92847235 | 92847499 | -1.006247308 | ANKRD19P |
| 6 | 18465267 | 18465467 | -2.95548013 | RNF144B | 6 | 81548044 | 81548244 | -1.805280043 | FAM46A | 9 | 92847235 | 92847499 | -1.006247308 | ZNF484 |
| 6 | 20402008 | 20402159 | -4.186495091 | E2F3 | 6 | 81748859 | 81749087 | 1.158294373 | FAM46A | 9 | 92848247 | 92848440 | -2.745926548 | ANKRD19P |
| 6 | 21594036 | 21594237 | 1.897845076 | RP11-204E9.3 | 1 | 214644603 | 214644802 | -2.562078697 | CENPF | 9 | 92848247 | 92848440 | -2.745926548 | ZNF484 |
| 6 | 21594036 | 21594237 | 1.897845076 | SOX4 | 6 | 83129051 | 83129252 | 1.591580023 | DOPEY1 | 9 | 93035619 | 93035754 | -2.095576556 | FGD3 |
| 6 | 21594725 | 21594910 | 1.745409827 | RP11-204E9.3 | 6 | 83137191 | 83137394 | 2.162810114 | DOPEY1 | 9 | 93678379 | 93678548 | -1.143870566 | PHF2 |
| 6 | 21594725 | 21594910 | 1.745409827 | SOX4 | 6 | 83137950 | 83138304 | 1.763171307 | DOPEY1 | 9 | 93951931 | 93952136 | 1.803228464 | BARX1 |
| 6 | 21596617 | 21596815 | 1.915171289 | SOX4 | 6 | 83138683 | 83138976 | 3.669466013 | DOPEY1 | 9 | 93959071 | 93959248 | 4.380775616 | BARX1-AS1 |
| 6 | 25140398 | 25140597 | -2.419263244 | CMAHP | 6 | 83166320 | 83166462 | -3.223311921 | DOPEY1 | 9 | 94772907 | 94773089 | -5.018086806 | C9orf3 |
| 6 | 25140398 | 25140597 | -2.419263244 | RP3-425P12.4 | 6 | 83166320 | 83166462 | -3.223311921 | PGM3 | 9 | 95508597 | 95508796 | 5.916445007 | PTCH1 |
| 6 | 26215911 | 26216111 | 7.181108908 | HIST1H2BG | 1 | 214645651 | 214645847 | -3.133423842 | CENPF | 9 | 96388164 | 96388272 | -2.845059211 | ZNF367 |
| 6 | 26636490 | 26636642 | -2.033805885 | ZNF322 | 6 | 83965424 | 83965603 | -6.618903737 | CYB5R4 | 9 | 96926350 | 96926546 | 1.943762635 | MFSD14C |
| 6 | 26636754 | 26636929 | -1.459383443 | ZNF322 | 6 | 85450551 | 85450691 | -1.128169877 | NT5E | 1 | 247135015 | 247135212 | -7.093173207 | ZNF124 |
| 6 | 26637277 | 26637483 | -1.008525645 | ZNF322 | 6 | 85509881 | 85510095 | -1.059001881 | SNX14 | 9 | 98015266 | 98015466 | -6.429476155 | ANP32B |
| 6 | 27406202 | 27406402 | 4.15210435 | RP1-153G14.4 | 1 | 214646981 | 214647186 | -2.426825579 | CENPF | 9 | 98213191 | 98213393 | -1.513581165 | TBC1D2 |
| 6 | 28166516 | 28166680 | -2.09977905 | ZNF192P1 | 6 | 85660039 | 85660239 | -1.246758726 | PKMP3 | 9 | 98220868 | 98221053 | -1.471400348 | TBC1D2 |
| 1 | 206587013 | 206587221 | 1.129843639 | RASSF5 | 6 | 87257731 | 87257921 | -1.523967239 | ZNF292 | 9 | 98736249 | 98736436 | 1.352342956 | ANKS6 |
| 1 | 206587013 | 206587221 | 1.129843639 | EIF2D | 6 | 87511448 | 87511629 | -2.374575924 | SLC35A1 | 9 | 98790068 | 98790168 | 1.503881251 | ANKS6 |

| | | | | |
|---|---|---|---|---|
| 6 | 28245374 | 28245573 | -1.133557393 | ZKSCAN4 |
| 6 | 28300826 | 28301021 | -1.07254668 | PGBD1 |
| 6 | 28326646 | 28326843 | -2.450696582 | ZSCAN31 |
| 6 | 28380337 | 28380541 | -1.393762523 | ZSCAN12 |
| 6 | 28382286 | 28382459 | -1.178753964 | ZSCAN12 |
| 1 | 206588219 | 206588395 | -1.007430847 | RASSF5 |
| 1 | 206588219 | 206588395 | -1.007430847 | EIF2D |
| 6 | 28390732 | 28390901 | -2.853184015 | ZSCAN12 |
| 6 | 28391293 | 28391510 | -1.837105428 | ZSCAN12 |
| 6 | 28904342 | 28904541 | -1.762143272 | TRIM27 |
| 6 | 29003959 | 29004094 | -1.659693696 | ZNF311 |
| 6 | 29738021 | 29738208 | -6.301697265 | HLA-F-AS1 |
| 6 | 29738021 | 29738208 | -6.301697265 | HLA-F |
| 1 | 206647658 | 206647860 | 4.754827241 | DYRK3 |
| 6 | 30006455 | 30006640 | -6.867778012 | HLA-J |
| 6 | 30006455 | 30006640 | -6.867778012 | ZNRD1ASP |
| 6 | 30006455 | 30006640 | -6.867778012 | HCG4P3 |
| 6 | 30278657 | 30278844 | 1.42738822 | HLA-L |
| 6 | 30278657 | 30278844 | 1.42738822 | HCG17 |
| 6 | 30291639 | 30291894 | -1.511693573 | HCG18 |
| 6 | 30291639 | 30291894 | -1.511693573 | HLA-L |
| 6 | 30291639 | 30291894 | -1.511693573 | HCG17 |
| 6 | 30294479 | 30294644 | -1.268730391 | HCG18 |
| 6 | 30294479 | 30294644 | -1.268730391 | HCG17 |
| 6 | 30562409 | 30562609 | -1.145848484 | PRR3 |
| 6 | 30684947 | 30685156 | 1.777022636 | PPP1R18 |
| 6 | 30712242 | 30712428 | -2.107091024 | MDC1 |
| 6 | 30712242 | 30712428 | -2.107091024 | MDC1-AS1 |
| 6 | 30713022 | 30713150 | -3.354761554 | MDC1 |
| 6 | 30713022 | 30713150 | -3.354761554 | MDC1-AS1 |
| 6 | 30723915 | 30724115 | -12.5198998 | TUBB |
| 1 | 9726928 | 9727166 | -2.649572325 | PIK3CD |
| 6 | 30743798 | 30743998 | 9.451939311 | IER3 |
| 6 | 31159226 | 31159424 | -3.287827579 | TCF19 |
| 6 | 31162728 | 31162928 | -3.937637674 | TCF19 |
| 6 | 31197017 | 31197158 | -1.40112847 | XXbac-BPG299F13.17 |
| 6 | 31394498 | 31394697 | -7.787740079 | XXbac-BPG181B23.7 |
| 6 | 31399656 | 31399805 | -1.744139454 | MICA |
| 6 | 31558436 | 31558642 | -1.176618942 | NFKBIL1 |
| 6 | 31577561 | 31577782 | -3.229354022 | TNF |
| 6 | 31581298 | 31581484 | -1.785851245 | LTB |
| 6 | 31627863 | 31628045 | -8.375928918 | PRRC2A |
| 6 | 31629515 | 31629715 | -4.13665505 | PRRC2A |
| 6 | 31632565 | 31632783 | -1.31048802 | PRRC2A |
| 6 | 31659629 | 31659760 | -1.16451116 | C6orf47 |
| 6 | 31659629 | 31659760 | -1.16451116 | C6orf47-AS1 |
| 6 | 31662389 | 31662579 | -2.070038734 | GPANK1 |
| 6 | 31817392 | 31817623 | -1.462632022 | HSPA1A |
| 6 | 31828539 | 31828711 | -1.871129947 | HSPA1B |
| 6 | 31899968 | 31900207 | -2.074412091 | ZBTB12 |
| 6 | 31899968 | 31900207 | -2.074412091 | C2 |
| 6 | 32096916 | 32097053 | 3.058683812 | TNXB |
| 6 | 32129227 | 32129427 | -1.791517279 | FKBPL |
| 6 | 32838031 | 32838180 | -6.346743479 | XXbac-BPG246D15.9 |

| | | | | |
|---|---|---|---|---|
| 6 | 87511448 | 87511629 | -2.374575924 | RP3-382I10.7 |
| 1 | 214656970 | 214657158 | -3.757849919 | CENPF |
| 6 | 88143892 | 88144085 | -1.165518587 | CNR1 |
| 6 | 89162077 | 89162297 | -1.337644408 | PM20D2 |
| 6 | 89283055 | 89283250 | 4.657624015 | GABRR2 |
| 1 | 215620073 | 215620274 | -1.160680737 | KCTD3 |
| 6 | 89637065 | 89637267 | -1.116320367 | LYRM2 |
| 6 | 89643909 | 89644065 | -1.322316595 | MDN1 |
| 6 | 89864058 | 89864250 | -2.312624102 | CASP8AP2 |
| 6 | 89932406 | 89932627 | -1.544375673 | BACH2 |
| 6 | 89949377 | 89949483 | -6.462035093 | BACH2 |
| 6 | 89950916 | 89951113 | -1.290597835 | RP3-512E2.2 |
| 6 | 89950916 | 89951113 | -1.290597835 | BACH2 |
| 6 | 93243236 | 93243524 | 1.47662587 | EPHA7 |
| 6 | 93410893 | 93411115 | 1.82118588 | EPHA7 |
| 1 | 217614155 | 217614354 | -1.144830947 | GPATCH2 |
| 6 | 96487436 | 96487624 | -2.589349247 | UFL1-AS1 |
| 6 | 96497390 | 96497592 | -2.239490346 | UFL1-AS1 |
| 6 | 96500131 | 96500343 | -1.664112667 | UFL1-AS1 |
| 6 | 96799618 | 96799807 | -1.443719116 | GPR63 |
| 6 | 98835802 | 98836027 | 2.798793895 | POU3F2 |
| 6 | 99446084 | 99446284 | -1.262688343 | USP45 |
| 6 | 99614729 | 99614863 | -1.73539574 | PRDM13 |
| 6 | 100584456 | 100584645 | 1.364135059 | ASCC3 |
| 6 | 100740763 | 100740946 | -5.836212671 | ASCC3 |
| 6 | 100758489 | 100758686 | 1.545848097 | ASCC3 |
| 1 | 218441358 | 218441522 | -2.64520145 | TGFB2 |
| 6 | 100882093 | 100882294 | 1.132530365 | RP3-467N11.1 |
| 6 | 105314674 | 105314870 | 2.035053694 | PREP |
| 6 | 106104942 | 106105134 | 1.794223656 | ATG5 |
| 6 | 106104942 | 106105134 | 1.794223656 | PRDM1 |
| 6 | 106107077 | 106107260 | 1.912645661 | ATG5 |
| 6 | 106107077 | 106107260 | 1.912645661 | PRDM1 |
| 6 | 107069083 | 107069290 | -1.249479389 | BEND3 |
| 6 | 107165060 | 107165251 | -2.5 | PDSS2 |
| 6 | 107168240 | 107168445 | -1.615652951 | PDSS2 |
| 6 | 107363807 | 107363985 | -4.447408882 | PDSS2 |
| 6 | 107635106 | 107635343 | -2.234839441 | SOBP |
| 6 | 107640700 | 107640899 | -1.780462868 | SOBP |
| 6 | 107641279 | 107641550 | -3.235295546 | SOBP |
| 6 | 107646029 | 107646223 | -1.312584412 | SOBP |
| 6 | 107658264 | 107658506 | -1.947004652 | SOBP |
| 6 | 107887987 | 107888187 | -1.637475591 | SEC63 |
| 1 | 219197327 | 219197526 | -2.678992373 | LYPLAL1 |
| 6 | 107959500 | 107959693 | 1.024428083 | RP1-191J18.66 |
| 6 | 108514691 | 108514891 | -2.024849547 | LACE1 |
| 6 | 108683204 | 108683424 | -1.136124566 | FOXO3 |
| 6 | 109094016 | 109094217 | -2.13055095 | SESN1 |
| 6 | 109133569 | 109133673 | 1.117406007 | CEP57L1 |
| 6 | 109162712 | 109162918 | -1.954384615 | CEP57L1 |
| 6 | 109175496 | 109175640 | -1.473505147 | CCDC162P |
| 6 | 109383034 | 109383309 | -1.507094274 | RP11-425D10.10 |
| 6 | 109415193 | 109415384 | -3.816864876 | PPIL6 |
| 6 | 109481339 | 109481532 | -1.216964783 | ZBTB24 |

| | | | | |
|---|---|---|---|---|
| 9 | 98790203 | 98790402 | 2.170300497 | ANKS6 |
| 9 | 99149365 | 99149552 | -1.509574782 | TGFBR1 |
| 9 | 99828185 | 99828384 | 1.040691238 | NR4A3 |
| 9 | 100179414 | 100179543 | -5.416445007 | INVS |
| 9 | 100182033 | 100182232 | -2.390381718 | INVS |
| 9 | 100317505 | 100317681 | -3.033941236 | TEX10 |
| 9 | 104784212 | 104784425 | -1.446031427 | ABCA1 |
| 1 | 247184053 | 247184248 | -1.546829554 | RP11-488L18.3 |
| 9 | 111026591 | 111026776 | -2.113419163 | LPAR1 |
| 1 | 247185048 | 247185301 | -1.051206857 | RP11-488L18.3 |
| 9 | 111661245 | 111661373 | 2.156017921 | DNAJC25-GNG10 |
| 9 | 111661245 | 111661373 | 2.156017921 | GNG10 |
| 9 | 113541487 | 113541674 | 2.35907224 | RGS3 |
| 9 | 113543683 | 113543867 | 2.439100607 | RGS3 |
| 9 | 113584309 | 113584502 | 1.84071267 | RP11-168K11.2 |
| 9 | 113584309 | 113584502 | 1.84071267 | RGS3 |
| 9 | 114053917 | 114054114 | -1.081694696 | ZNF618 |
| 9 | 114269724 | 114269927 | 3.933484889 | COL27A1 |
| 9 | 114377002 | 114377199 | -1.532263128 | AKNA |
| 9 | 114789159 | 114789358 | -2.369654133 | TNFSF15 |
| 9 | 114790232 | 114790434 | -2.18324662 | TNFSF15 |
| 9 | 120902797 | 120902951 | 1.9486445 | TRAF1 |
| 9 | 120904609 | 120904762 | 2.129905414 | TRAF1 |
| 9 | 120904788 | 120905009 | 2.488257881 | TRAF1 |
| 9 | 121100607 | 121100801 | -5.459431619 | CNTRL |
| 9 | 121205366 | 121205555 | 1.88862915 | RAB14 |
| 9 | 122820341 | 122820563 | -1.655519084 | PDCL |
| 1 | 247423080 | 247423275 | -2.053995451 | NLRP3 |
| 9 | 123103037 | 123103240 | -1.439299632 | RABGAP1 |
| 9 | 123371035 | 123371231 | -6.136606405 | CRB2 |
| 9 | 123382023 | 123382187 | 7.941655758 | DENND1A |
| 9 | 123416587 | 123416744 | 6.866984305 | DENND1A |
| 9 | 124032571 | 124032768 | 1.880852206 | LHX2 |
| 9 | 124344587 | 124344787 | 7.842319528 | NEK6 |
| 9 | 125150476 | 125150604 | 4.54637857 | PPP6C |
| 1 | 247451642 | 247451832 | -2.146715538 | OR2B11 |
| 9 | 127506810 | 127507023 | 1.880011875 | FAM129B |
| 9 | 128063191 | 128063401 | 1.207592898 | NAIF1 |
| 9 | 128107581 | 128107852 | 1.262276579 | SLC25A25 |
| 9 | 128193112 | 128193311 | 1.474167178 | CIZ1 |
| 9 | 128193112 | 128193311 | 1.474167178 | DNM1 |
| 9 | 128323190 | 128323428 | 1.137140635 | COQ4 |
| 9 | 128333630 | 128333804 | 1.079384692 | COQ4 |
| 9 | 128633696 | 128633877 | 1.782325339 | WDR34 |
| 9 | 129176763 | 129176962 | 16.02150762 | IER5L |
| 9 | 129176763 | 129176962 | 16.02150762 | RP11-247A12.8 |
| 9 | 129176763 | 129176962 | 16.02150762 | RP11-247A12.2 |
| 9 | 129177179 | 129177347 | 2.502317165 | IER5L |
| 9 | 129177179 | 129177347 | 2.502317165 | RP11-247A12.8 |
| 9 | 129177179 | 129177347 | 2.502317165 | RP11-247A12.2 |
| 1 | 248812084 | 248812288 | 2.975188247 | SH3BP5L |
| 9 | 130701569 | 130701739 | -5.766664866 | EXOSC2 |
| 1 | 248825151 | 248825342 | 2.480208243 | SH3BP5L |
| 9 | 131474792 | 131474996 | -7.503469818 | PRRC2B |

| | | | | |
|---|---|---|---|---|
| 6 | 32838031 | 32838180 | -6.346743479 | TAP2 |
| 6 | 32850485 | 32850648 | -8.553817988 | PSMB9 |
| 6 | 32850485 | 32850648 | -8.553817988 | TAP1 |
| 1 | 209675902 | 209676104 | 3.197721635 | G0S2 |
| 1 | 209675902 | 209676104 | 3.197721635 | RP1-28O10.1 |
| 6 | 33194485 | 33194684 | -1.752704279 | RXRB |
| 6 | 33203798 | 33203998 | -2.036642743 | SLC39A7 |
| 6 | 33320980 | 33321231 | -1.796697368 | DAXX |
| 6 | 33391409 | 33391558 | -1.727418522 | KIFC1 |
| 6 | 33403826 | 33404025 | -2.931273679 | KIFC1 |
| 6 | 33404831 | 33404953 | -3.143744432 | KIFC1 |
| 6 | 33405428 | 33405616 | -4.020405027 | KIFC1 |
| 1 | 209783056 | 209783270 | -1.89671772 | C1orf74 |
| 6 | 34279540 | 34279740 | 2.325366574 | NUDT3 |
| 6 | 34279540 | 34279740 | 2.325366574 | RP11-513I15.6 |
| 6 | 34280149 | 34280345 | 3.409454726 | NUDT3 |
| 6 | 34280149 | 34280345 | 3.409454726 | RP11-513I15.6 |
| 6 | 35058908 | 35059065 | -3.53599929 | ANKS1A |
| 6 | 35295474 | 35295673 | -1.035912249 | ZNF76 |
| 6 | 35373961 | 35374155 | -7.25042094 | PPARD |
| 6 | 35425982 | 35426150 | 2.443442881 | PPARD |
| 6 | 35426560 | 35426798 | 2.618117233 | PPARD |
| 6 | 35455936 | 35456148 | 1.480335229 | FANCE |
| 6 | 35833742 | 35833945 | -2.84034266 | SRPK1 |
| 6 | 35833742 | 35833945 | -2.84034266 | LHFPL5 |
| 6 | 35869582 | 35869787 | -1.749974154 | SRPK1 |
| 6 | 36084566 | 36084772 | -1.41001098 | MAPK14 |
| 6 | 36085713 | 36085939 | -2.146119563 | MAPK14 |
| 6 | 36088042 | 36088254 | -1.184685189 | MAPK14 |
| 6 | 36089172 | 36089377 | -2.136446702 | MAPK14 |
| 6 | 36139284 | 36139451 | 1.015421708 | MAPK13 |
| 1 | 9930118 | 9930314 | -1.348985731 | LZIC |
| 6 | 36487797 | 36487995 | -2.144074869 | KCTD20 |
| 6 | 36495736 | 36495946 | -1.509033767 | STK38 |
| 6 | 36721593 | 36721770 | -4.578972379 | RAB44 |
| 6 | 36721973 | 36722218 | -4.534265207 | RAB44 |
| 6 | 37174077 | 37174274 | 3.106839103 | PIM1 |
| 6 | 37212347 | 37212545 | 4.229410323 | TMEM217 |
| 6 | 37368928 | 37369135 | -1.015766161 | RNF8 |
| 6 | 37637193 | 37637398 | 3.983740288 | MDGA1 |
| 6 | 38087117 | 38087314 | 7.505832026 | ZFAND3 |
| 6 | 38173879 | 38174085 | -1.221531347 | BTBD9 |
| 1 | 211941872 | 211942070 | -1.643428862 | INTS7 |
| 6 | 39190924 | 39191152 | -3.340797349 | KCNK5 |
| 6 | 41098771 | 41098966 | -2.611360212 | NFYA |
| 6 | 41158734 | 41158937 | 1.729940892 | TREM2 |
| 1 | 212100765 | 212100914 | -9.23661505 | DTL |
| 6 | 41684292 | 41684492 | 2.681032397 | TFEB |
| 6 | 41787152 | 41787320 | -1.032635236 | PRICKLE4 |
| 6 | 41796546 | 41796725 | 1.11959104 | USP49 |
| 6 | 41906888 | 41907098 | -2.211012616 | MED20 |
| 6 | 41932563 | 41932762 | -2.099852307 | BYSL |
| 6 | 42228350 | 42228550 | 1.110957337 | TRERF1 |
| 6 | 42745192 | 42745393 | -1.480811718 | TBCC |
| 6 | 110614346 | 110614546 | -1.683973704 | CDK19 |
| 6 | 110858371 | 110858501 | -1.646135123 | CNN2P9 |
| 6 | 110858539 | 110858765 | -1.643349018 | CNN2P9 |
| 1 | 219610610 | 219610809 | -1.970605334 | ZC3H11B |
| 6 | 110874813 | 110875023 | -1.666600991 | AMD1 |
| 6 | 111367608 | 111367813 | -1.64201951 | REV3L |
| 6 | 111372692 | 111372996 | -2.225577258 | REV3L |
| 6 | 111373613 | 111373818 | -1.38712964 | REV3L |
| 6 | 111510984 | 111511097 | -1.869787223 | TRAF3IP2-AS1 |
| 6 | 111591494 | 111591708 | 1.281336292 | TRAF3IP2 |
| 6 | 111591494 | 111591708 | 1.281336292 | TRAF3IP2-AS1 |
| 6 | 113860518 | 113860705 | 2.265851841 | MARCKS |
| 6 | 113868499 | 113868617 | 1.103768902 | LINC01268 |
| 6 | 113995466 | 113995650 | -4.112000837 | HDAC2 |
| 6 | 113995466 | 113995650 | -4.112000837 | RP3-399L15.3 |
| 6 | 116252896 | 116253095 | -1.658531862 | TSPYL4 |
| 6 | 116435686 | 116435874 | 1.663180631 | DSE |
| 6 | 116436719 | 116436924 | 1.282297823 | DSE |
| 6 | 116666752 | 116666951 | 1.109877813 | ZUFSP |
| 6 | 117272532 | 117272730 | 11.55411194 | VGLL2 |
| 6 | 117548225 | 117548420 | 1.060386486 | DCBLD1 |
| 6 | 117548225 | 117548420 | 1.060386486 | RP1-179P9.3 |
| 6 | 117699696 | 117699878 | -7.216303572 | NUS1 |
| 1 | 220813517 | 220813713 | -2.279713704 | Mar-01 |
| 6 | 117997994 | 117998165 | 4.077409055 | SLC35F1 |
| 6 | 118049994 | 118050097 | 5.146160816 | SLC35F1 |
| 6 | 118834886 | 118835047 | -1.295883839 | MCM9 |
| 6 | 119306951 | 119307152 | 1.614024329 | MAN1A1 |
| 1 | 220831246 | 220831441 | -6.891601445 | LINC01352 |
| 6 | 119348542 | 119348747 | 1.74169582 | MAN1A1 |
| 6 | 121100072 | 121100239 | -2.45876892 | TBC1D32 |
| 6 | 124963551 | 124963757 | -2.142744834 | RNF217 |
| 1 | 220861858 | 220861965 | -3.501876067 | HLX-AS1 |
| 6 | 125296590 | 125296911 | -7.213328939 | HDDC2 |
| 6 | 125889366 | 125889561 | -1.961120686 | NCOA7 |
| 6 | 125889613 | 125889809 | -1.623491188 | NCOA7 |
| 6 | 125928734 | 125928934 | 1.30475451 | NCOA7 |
| 6 | 126644097 | 126644256 | -2.318622946 | PRELID1P1 |
| 6 | 127363340 | 127363539 | -5.372416919 | RPL5P18 |
| 6 | 127446871 | 127447055 | -3.170221791 | RP3-403A15.5 |
| 6 | 127446871 | 127447055 | -3.170221791 | KIAA0408 |
| 6 | 127518804 | 127518956 | -1.854535426 | SOGA3 |
| 6 | 127518804 | 127518956 | -1.854535426 | RP3-403A15.5 |
| 6 | 129589345 | 129589548 | -1.881458855 | ARHGAP18 |
| 6 | 130139570 | 130139774 | -1.575152403 | RP11-73O6.3 |
| 6 | 130139570 | 130139774 | -1.575152403 | L3MBTL3 |
| 6 | 130441543 | 130441704 | -3.890679857 | TMEM200A |
| 6 | 131949202 | 131949391 | 2.964012404 | CTGF |
| 6 | 135465950 | 135466149 | -1.175619691 | AHI1 |
| 6 | 136192088 | 136192288 | -5.015381127 | RP13-143G15.4 |
| 6 | 136192088 | 136192288 | -5.015381127 | PDE7B |
| 6 | 136786163 | 136786302 | -9.033223312 | MAP3K5 |
| 6 | 136788485 | 136788852 | -3.40616226 | MAP3K5 |
| 6 | 137198243 | 137198434 | -2.576378107 | IFNGR1 |
| 9 | 131552423 | 131552606 | -7.664898169 | RP11-40A7.2 |
| 9 | 131621136 | 131621330 | 6.877443751 | RAPGEF1 |
| 1 | 248847259 | 248847474 | 2.452742344 | ZNF672 |
| 9 | 132905688 | 132905893 | -1.454282978 | TSC1 |
| 9 | 133464443 | 133464585 | 3.769579406 | CACFD1 |
| 9 | 133471615 | 133471830 | 1.649534871 | SLC2A6 |
| 9 | 135468293 | 135468442 | -5.852600189 | PPP1R26-AS1 |
| 10 | 150782 | 150954 | -6.611246526 | ZMYND11 |
| 9 | 136357799 | 136358114 | 2.414265966 | GPSM1 |
| 9 | 136378320 | 136378520 | 1.538106264 | SNAPC4 |
| 9 | 136423225 | 136423470 | 1.078334384 | PMPCA |
| 9 | 136438882 | 136439085 | 1.688659115 | INPP5E |
| 9 | 136614588 | 136614788 | 1.542295996 | LINC01451 |
| 9 | 136673495 | 136673637 | 8.266512744 | AGPAT2 |
| 9 | 136721215 | 136721422 | 1.574220637 | SNHG7 |
| 9 | 136721215 | 136721422 | 1.574220637 | FAM69B |
| 9 | 136802973 | 136803167 | 1.698152176 | RP11-216L13.18 |
| 9 | 136802973 | 136803167 | 1.698152176 | RP11-216L13.19 |
| 9 | 136802973 | 136803167 | 1.698152176 | CCDC183 |
| 9 | 136839991 | 136840280 | 7.139224729 | RABL6 |
| 9 | 136840681 | 136840881 | 2.180964505 | RABL6 |
| 9 | 136940769 | 136940932 | 2.849733297 | FBXW5 |
| 9 | 136941593 | 136941792 | 1.257466813 | FBXW5 |
| 9 | 136942106 | 136942302 | 10.52339981 | FBXW5 |
| 9 | 137062844 | 137063117 | 1.087447514 | SAPCD2 |
| 9 | 137064107 | 137064277 | 1.850774742 | SAPCD2 |
| 9 | 137064107 | 137064277 | 1.850774742 | RP11-229P13.23 |
| 9 | 137085603 | 137085740 | 1.288760303 | MAN1B1-AS1 |
| 9 | 137232832 | 137232940 | 1.075402911 | SLC34A3 |
| 9 | 137279800 | 137280002 | 4.013526125 | TOR4A |
| 9 | 137488203 | 137488401 | -1.044513311 | PNPLA7 |
| 9 | 137612155 | 137612256 | -6.26116816 | ARRDC1 |
| 9 | 137612155 | 137612256 | -6.26116816 | RP11-48C7.8 |
| 10 | 3122435 | 3122638 | 10.95583862 | PFKP |
| 9 | 137728232 | 137728446 | 9.813916542 | EHMT1 |
| X | 310152 | 310282 | -2.07609171 | GTPBP6 |
| X | 313480 | 313654 | -8.224106559 | GTPBP6 |
| X | 350431 | 350614 | 1.351086343 | PPP2R3B |
| X | 1190725 | 1190921 | 3.264812882 | CRLF2 |
| X | 1309551 | 1309755 | -1.473887719 | CSF2RA |
| X | 1389344 | 1389543 | 3.323408252 | SLC25A6 |
| X | 1463169 | 1463370 | -1.804671565 | P2RY8 |
| X | 1742581 | 1742818 | -1.677906135 | RP13-297E16.4 |
| 10 | 4826051 | 4826228 | -3.549767837 | AKR1E2 |
| X | 9719104 | 9719247 | 1.680693631 | TBL1X |
| X | 9932253 | 9932427 | -6.38975709 | SHROOM2 |
| X | 9947994 | 9948111 | -1.273743898 | SHROOM2 |
| X | 10117452 | 10117641 | 2.661281596 | WWC3 |
| X | 10122446 | 10122635 | 2.979071993 | WWC3 |
| X | 11586420 | 11586562 | -3.341038605 | ARHGAP6 |
| X | 12920577 | 12920722 | 2.599782246 | TLR8 |
| X | 12921726 | 12921930 | 3.936752214 | TLR8 |
| X | 15321667 | 15321797 | -2.621469899 | PIGA |
| X | 15697329 | 15697439 | 6.268123608 | CA5BP1 |

| | | | | |
|---|---|---|---|---|
| 6 | 42886906 | 42887107 | -1.135046996 | RPL7L1 |
| 6 | 42938987 | 42939188 | -1.462517561 | CNPY3 |
| 6 | 42978343 | 42978764 | -1.952945644 | PEX6 |
| 1 | 212619396 | 212619588 | 10.04589523 | ATF3 |
| 6 | 43029160 | 43029358 | -1.272269079 | RRP36 |
| 6 | 43339988 | 43340191 | -1.461360962 | ZNF318 |
| 6 | 43355809 | 43356010 | -1.26744917 | ZNF318 |
| 6 | 43357413 | 43357631 | -1.038755185 | ZNF318 |
| 6 | 43454798 | 43455002 | -1.605918284 | DLK2 |
| 6 | 43505390 | 43505616 | -1.654607988 | TJAP1 |
| 6 | 43533599 | 43533793 | 1.745423867 | XPO5 |
| 6 | 43614132 | 43614282 | -2.528238842 | GTPBP2 |
| 6 | 43614132 | 43614282 | -2.528238842 | POLH |
| 6 | 43671593 | 43671795 | -9.821434818 | MRPS18A |
| 6 | 43671593 | 43671795 | -9.821434818 | RSPH9 |
| 6 | 43770491 | 43770710 | 1.802237791 | RP1-261G23.7 |
| 6 | 43770491 | 43770710 | 1.802237791 | VEGFA |
| 6 | 43785162 | 43785434 | 1.784741588 | VEGFA |
| 6 | 43996657 | 43996844 | 1.689255812 | RP5-1120P11.1 |
| 6 | 44113899 | 44114099 | -1.158133918 | MRPL14 |
| 6 | 44257802 | 44258002 | 1.106818246 | SLC35B2 |
| 6 | 44299707 | 44299901 | -1.826066532 | TMEM151B |
| 6 | 44299707 | 44299901 | -1.826066532 | RP11-444E17.6 |
| 6 | 44299707 | 44299901 | -1.826066532 | AARS2 |
| 6 | 44300297 | 44300503 | -1.633409644 | AARS2 |
| 6 | 44300297 | 44300503 | -1.633409644 | TMEM151B |
| 6 | 44300297 | 44300503 | -1.633409644 | RP11-444E17.6 |
| 6 | 44938088 | 44938219 | 5.528995861 | SUPT3H |
| 6 | 44938088 | 44938219 | 5.528995861 | AL161905.1 |
| 6 | 45421763 | 45421937 | -3.99631654 | RP1-244F24.1 |
| 6 | 45421763 | 45421937 | -3.99631654 | RUNX2 |
| 6 | 45422208 | 45422460 | -3.116612847 | RUNX2 |
| 6 | 45547656 | 45547857 | -1.106475112 | RUNX2 |
| 6 | 47232767 | 47233011 | 4.091662389 | TNFRSF21 |
| 6 | 47284074 | 47284274 | 3.867802579 | TNFRSF21 |
| 6 | 47286218 | 47286439 | 4.03294673 | TNFRSF21 |
| 1 | 212726574 | 212726751 | -1.36751417 | NSL1 |
| 6 | 52264295 | 52264503 | -3.738626007 | MCM3 |
| 6 | 52404375 | 52404574 | -2.873686245 | EFHC1 |
| 6 | 52404375 | 52404574 | -2.873686245 | PAQR8 |
| 6 | 52404912 | 52405119 | -3.259905429 | EFHC1 |
| 6 | 52404912 | 52405119 | -3.259905429 | PAQR8 |
| 6 | 52502363 | 52502563 | 1.728644739 | TRAM2 |
| 6 | 52502363 | 52502563 | 1.728644739 | EFHC1 |
| 6 | 53498692 | 53498881 | 4.340814303 | GCLC |
| 1 | 944187 | 944287 | 9.102297512 | NOC2L |
| 1 | 944187 | 944287 | 9.102297512 | SAMD11 |
| 1 | 9982478 | 9982677 | -1.017839195 | NMNAT1 |
| 6 | 56459050 | 56459267 | -2.239995547 | DST |
| 6 | 56541275 | 56541466 | -2.406515976 | DST |
| 6 | 56552310 | 56552554 | -1.556664375 | DST |
| 6 | 56555489 | 56555696 | -2.119639721 | DST |
| 6 | 57052876 | 57053083 | -1.267276746 | KIAA1586 |
| 6 | 57556562 | 57556756 | -1.838138834 | PRIM2 |
| 6 | 137824172 | 137824359 | 1.44783167 | RP11-356I2.4 |
| 6 | 137879004 | 137879204 | 3.625647776 | TNFAIP3 |
| 6 | 137879966 | 137880166 | 3.349936652 | TNFAIP3 |
| 6 | 137881547 | 137881748 | 3.466074689 | TNFAIP3 |
| 6 | 138415278 | 138415473 | -2.509259198 | HEBP2 |
| 6 | 138418778 | 138418892 | -3.251191882 | HEBP2 |
| 6 | 138420000 | 138420154 | -2.387970985 | HEBP2 |
| 6 | 138422348 | 138422518 | -6.192324431 | NHSL1 |
| 6 | 138424142 | 138424344 | -2.089870319 | NHSL1 |
| 6 | 138430966 | 138431161 | -3.523951453 | NHSL1 |
| 6 | 138432354 | 138432567 | -3.65723792 | NHSL1 |
| 6 | 138433187 | 138433382 | -3.10780531 | NHSL1 |
| 6 | 139373606 | 139373743 | 10.77344205 | CITED2 |
| 6 | 142753245 | 142753508 | 4.281395724 | HIVEP2 |
| 6 | 142753245 | 142753508 | 4.281395724 | RP1-67K17.3 |
| 6 | 142760203 | 142760409 | 4.182548637 | HIVEP2 |
| 6 | 142769806 | 142769999 | 5.543779586 | HIVEP2 |
| 6 | 142770430 | 142770624 | 4.590504637 | HIVEP2 |
| 6 | 142772648 | 142772845 | 3.244091158 | HIVEP2 |
| 6 | 142773861 | 142774085 | 3.903959725 | HIVEP2 |
| 6 | 142774627 | 142774828 | 3.999404117 | HIVEP2 |
| 6 | 142950107 | 142950295 | 5.381638084 | RP11-439L18.1 |
| 6 | 142950107 | 142950295 | 5.381638084 | RP11-439L18.2 |
| 1 | 222627988 | 222628189 | -1.404029682 | MIA3 |
| 6 | 143490913 | 143491115 | 1.445706629 | RP1-20N2.8 |
| 6 | 143490913 | 143491115 | 1.445706629 | VDAC1P8 |
| 6 | 143942219 | 143942433 | 1.485540082 | PLAGL1 |
| 6 | 143948212 | 143948385 | 3.54333132 | PLAGL1 |
| 6 | 144187076 | 144187310 | 3.111794414 | STX11 |
| 6 | 145627023 | 145627234 | -1.150418219 | EPM2A |
| 6 | 145844187 | 145844312 | -6.703036053 | RP11-545I5.3 |
| 6 | 145855742 | 145855888 | -1.25086432 | RP11-545I5.3 |
| 6 | 145871843 | 145872041 | -2.015101826 | RP11-545I5.3 |
| 6 | 145871843 | 145872041 | -2.015101826 | SHPRH |
| 6 | 145872794 | 145872993 | -1.278903775 | RP11-545I5.3 |
| 6 | 145872794 | 145872993 | -1.278903775 | SHPRH |
| 6 | 146844580 | 146844754 | -1.061628471 | STXBP5-AS1 |
| 6 | 147346956 | 147347151 | -1.092151649 | STXBP5 |
| 6 | 147407808 | 147407991 | -1.043547708 | YAP1P1 |
| 6 | 149027437 | 149027747 | 1.381485751 | RP11-162J8.3 |
| 6 | 149027437 | 149027747 | 1.381485751 | UST |
| 6 | 149317227 | 149317380 | 6.41585155 | TAB2 |
| 6 | 149591172 | 149591289 | -1.014904893 | GINM1 |
| 6 | 149858221 | 149858419 | -2.79248125 | LRP11 |
| 6 | 149969021 | 149969220 | -8.228080936 | ULBP1 |
| 6 | 149972392 | 149972534 | -3.569535409 | ULBP1 |
| 6 | 150935273 | 150935466 | -2.669684271 | MTHFD1L |
| 6 | 150935273 | 150935466 | -2.669684271 | RP1-99E18.2 |
| 6 | 151365682 | 151365879 | -1.445085368 | ZBTB2 |
| 6 | 151414568 | 151414769 | -1.434833034 | RMND1 |
| 1 | 223005538 | 223005736 | -1.848773403 | DISP1 |
| 6 | 151445433 | 151445602 | -1.358603317 | RMND1 |
| 6 | 152122210 | 152122398 | -1.556276911 | ESR1 |
| 6 | 152122210 | 152122398 | -1.556276911 | SYNE1 |
| X | 15697329 | 15697439 | 6.268123608 | CA5B |
| X | 16597179 | 16597400 | 5.748426889 | CTPS2 |
| X | 19779786 | 19779891 | 1.886294752 | SH3KBP1 |
| X | 19780413 | 19780601 | 6.001407508 | SH3KBP1 |
| X | 30559834 | 30560061 | 1.029881402 | CXorf21 |
| X | 37727816 | 37727973 | -7.161034488 | XK |
| X | 37727816 | 37727973 | -7.161034488 | RP5-972B16.2 |
| X | 38796931 | 38797097 | 2.51796627 | RP4-646N3.1 |
| X | 40074875 | 40075006 | -1.337907966 | BCOR |
| X | 46625358 | 46625493 | 4.783978038 | SLC9A7 |
| X | 46648011 | 46648220 | 1.062621172 | SLC9A7 |
| X | 48957841 | 48958039 | 1.844085565 | OTUD5 |
| X | 53145577 | 53145724 | 1.328038674 | KANTR |
| X | 53147160 | 53147360 | 1.017257162 | KANTR |
| X | 53169039 | 53169228 | -5.329551982 | RP11-258C19.7 |
| 10 | 5953049 | 5953245 | 3.52219706 | IL15RA |
| X | 54173945 | 54174132 | 2.780648519 | FAM120C |
| X | 64190483 | 64190608 | -4.416445007 | AMER1 |
| X | 64311504 | 64311701 | -1.103225439 | MTMR8 |
| 1 | 1055055 | 1055216 | 1.219114638 | RP11-54O7.14 |
| 1 | 1055055 | 1055216 | 1.219114638 | AGRN |
| X | 64381330 | 64381535 | -1.159351582 | MTMR8 |
| X | 64381539 | 64381699 | -1.121978294 | MTMR8 |
| X | 66021887 | 66022083 | -2.762261604 | VSIG4 |
| X | 68717309 | 68717503 | -6.9376827 | STARD8 |
| 10 | 7197233 | 7197418 | 2.650613326 | SFMBT2 |
| X | 71096404 | 71096608 | -1.523112392 | FOXO4 |
| X | 71183471 | 71183652 | 2.674687187 | RP5-1091N2.9 |
| X | 71492069 | 71492205 | 1.984605743 | TAF1 |
| X | 71492069 | 71492205 | 1.984605743 | INGX |
| 10 | 7411383 | 7411501 | 4.245926548 | SFMBT2 |
| X | 72208481 | 72208678 | -2.74920727 | PIN4 |
| X | 72208481 | 72208678 | -2.74920727 | ERCC6L |
| 10 | 7412296 | 7412473 | 1.688729198 | RP5-1031D4.3 |
| X | 74095528 | 74095688 | -1.270066752 | FTX |
| X | 74097210 | 74097362 | -2.181130094 | FTX |
| X | 74193104 | 74193282 | -1.932247151 | FTX |
| X | 75741148 | 75741303 | 1.07850168 | TTC3P1 |
| X | 75744278 | 75744435 | -1.422414814 | TTC3P1 |
| X | 75745119 | 75745222 | -6.47244752 | TTC3P1 |
| X | 77843748 | 77843932 | 6.277534196 | MAGT1 |
| X | 77846837 | 77847094 | 7.383988751 | MAGT1 |
| X | 80571860 | 80572059 | -5.408491812 | HK2P1 |
| 10 | 11462636 | 11462841 | -1.183490473 | USP6NL |
| X | 92022227 | 92022411 | -6.834719369 | PCDH11X |
| X | 103609272 | 103609475 | 1.204130368 | TCEAL3 |
| X | 103630108 | 103630359 | 1.380241686 | TCEAL1 |
| X | 103825526 | 103825722 | 1.927577476 | RAB9B |
| X | 109624043 | 109624241 | 3.405776914 | KCNE5 |
| X | 111755236 | 111755415 | -2.452728006 | ALG13 |
| X | 113700055 | 113700249 | -1.817180139 | XACT |
| X | 113820181 | 113820330 | -1.743628168 | XACT |
| X | 113823081 | 113823237 | -1.04512183 | XACT |
| X | 113825461 | 113825646 | -6.83941009 | XACT |

| | | | | |
|---|---|---|---|---|
| 6 | 57946208 | 57946399 | -2.302329151 | GUSBP1 |
| 6 | 57946208 | 57946399 | -2.302329151 | LINC00680 |
| 6 | 57946706 | 57946920 | -2.95070689 | GUSBP1 |
| 6 | 57946706 | 57946920 | -2.95070689 | LINC00680 |
| 6 | 57949697 | 57949933 | -2.04717369 | GUSBP1 |
| 6 | 57949697 | 57949933 | -2.04717369 | LINC00680 |
| 6 | 63713322 | 63713544 | -1.186804116 | PHF3 |
| 6 | 70757342 | 70757534 | -5.629283017 | SMAP1 |
| 1 | 214383388 | 214383585 | 4.508343595 | PTPN14 |
| 6 | 71318434 | 71318570 | 2.723764168 | RP3-331H24.5 |
| 6 | 71318434 | 71318570 | 2.723764168 | RP11-154D6.1 |
| 6 | 71319058 | 71319260 | 5.503163039 | RP3-331H24.5 |
| 6 | 71319058 | 71319260 | 5.503163039 | RP11-154D6.1 |
| 6 | 72831169 | 72831478 | -5.497176718 | KCNQ5 |
| 6 | 73195072 | 73195276 | -2.264222532 | KCNQ5 |
| 6 | 73266941 | 73267085 | -2.155411511 | KHDC1 |
| 6 | 73266941 | 73267085 | -2.155411511 | RP11-398K22.12 |
| 6 | 73290899 | 73291091 | -1.759200125 | RP11-398K22.12 |
| 6 | 73290899 | 73291091 | -1.759200125 | RPSAP41 |
| 6 | 73290899 | 73291091 | -1.759200125 | KHDC1 |
| 1 | 214640235 | 214640540 | -4.198466736 | CENPF |
| 6 | 73620588 | 73620785 | 6.674087434 | SLC17A5 |
| 6 | 73695151 | 73695271 | -2.536624491 | RP11-553A21.3 |
| 6 | 75681842 | 75682045 | -2.294116055 | SENP6 |
| 6 | 78927561 | 78927682 | 4.903677461 | IRAK1BP1 |
| 6 | 78940890 | 78941089 | -1.122915945 | PHIP |
| 6 | 78940890 | 78941089 | -1.122915945 | IRAK1BP1 |
| 6 | 79201626 | 79201802 | -1.135605059 | HMGN3 |
| 6 | 79487088 | 79487286 | -1.684886086 | LCA5 |
| 6 | 79487656 | 79487855 | -1.200130353 | LCA5 |
| 6 | 79513300 | 79513502 | -1.920676296 | LCA5 |
| 1 | 214642140 | 214642340 | -4.744448467 | CENPF |
| 6 | 81748859 | 81749087 | -1.249186761 | FAM46A |
| 1 | 214644603 | 214644802 | -3.240483927 | CENPF |
| 6 | 81764293 | 81764422 | 2.111885945 | RP5-991C6.2 |
| 6 | 83011556 | 83011723 | -2.183374808 | UBE3D |
| 6 | 83137191 | 83137394 | -1.328625772 | DOPEY1 |
| 6 | 83137950 | 83138304 | -1.077010876 | DOPEY1 |
| 6 | 83138683 | 83138976 | -2.438753394 | DOPEY1 |
| 6 | 83166320 | 83166462 | 2.082112056 | PGM3 |
| 6 | 83166320 | 83166462 | 2.082112056 | DOPEY1 |
| 1 | 214645651 | 214645847 | -3.581046522 | CENPF |
| 6 | 84736973 | 84737177 | -1.879852695 | TBX18 |
| 1 | 214646981 | 214647186 | -3.661115878 | CENPF |
| 6 | 85660039 | 85660239 | 1.87123535 | PKMP3 |
| 6 | 87254954 | 87255150 | -1.147411257 | ZNF292 |
| 6 | 87256446 | 87256682 | -1.036001099 | ZNF292 |
| 6 | 87259627 | 87259867 | -1.602559022 | ZNF292 |
| 6 | 87260232 | 87260434 | -1.105445322 | ZNF292 |
| 6 | 87261324 | 87261694 | -1.457113619 | ZNF292 |
| 1 | 214656970 | 214657158 | -6.119481308 | CENPF |
| 6 | 87700559 | 87700682 | -7.495051982 | AKIRIN2 |
| 6 | 88143892 | 88144085 | 2.353038302 | CNR1 |
| 6 | 88275803 | 88276000 | -1.861285938 | ACTBP8 |

| | | | | |
|---|---|---|---|---|
| 6 | 152975271 | 152975472 | -1.511299076 | FBXO5 |
| 6 | 154159265 | 154159465 | -3.2266441 | OPRM1 |
| 6 | 154159265 | 154159465 | -3.2266441 | IPCEF1 |
| 6 | 154222752 | 154222940 | -1.839713984 | IPCEF1 |
| 6 | 154222752 | 154222940 | -1.839713984 | OPRM1 |
| 6 | 154860621 | 154860801 | 6.575825415 | TIAM2 |
| 6 | 155129489 | 155129735 | -1.089061077 | TIAM2 |
| 6 | 155256678 | 155256877 | -1.308018675 | TIAM2 |
| 6 | 155256678 | 155256877 | -1.308018675 | RP11-477D19.2 |
| 6 | 156778035 | 156778268 | 9.226007981 | ARID1B |
| 6 | 156778035 | 156778268 | 9.226007981 | RP11-230C9.2 |
| 6 | 157200889 | 157201143 | -1.091482112 | ARID1B |
| 6 | 157207305 | 157207505 | -1.336404807 | ARID1B |
| 6 | 157292726 | 157292928 | -3.267320458 | TMEM242 |
| 6 | 157882797 | 157882993 | -8.16614068 | SNX9 |
| 6 | 158111011 | 158111180 | -1.784755138 | SERAC1 |
| 6 | 158503622 | 158503820 | -8.817022633 | TULP4 |
| 6 | 158631189 | 158631322 | 1.22750439 | TMEM181 |
| 6 | 158677150 | 158677334 | 4.774410954 | SYTL3 |
| 6 | 158977572 | 158977783 | -1.10243685 | RSPH3 |
| 6 | 158990918 | 158991113 | -6.189959909 | RP1-111C20.4 |
| 6 | 158990918 | 158991113 | -6.189959909 | RSPH3 |
| 6 | 159035978 | 159036183 | 1.498099421 | TAGAP |
| 6 | 159035978 | 159036183 | 1.498099421 | RP1-111C20.4 |
| 6 | 159748214 | 159748415 | 1.986781182 | WTAP |
| 6 | 159748214 | 159748415 | 1.986781182 | SOD2 |
| 6 | 159755239 | 159755439 | 1.167439985 | SOD2 |
| 6 | 159755239 | 159755439 | 1.167439985 | WTAP |
| 6 | 160052969 | 160053183 | 1.387379811 | IGF2R |
| 6 | 160092741 | 160092935 | 8.141761243 | IGF2R |
| 6 | 160093787 | 160093955 | 1.65757703 | RP11-288H12.3 |
| 6 | 160093787 | 160093955 | 1.65757703 | IGF2R |
| 6 | 160991327 | 160991472 | -1.670715796 | RP3-428L16.2 |
| 6 | 161048961 | 161049159 | -1.459732155 | MAP3K4 |
| 6 | 161049697 | 161049887 | -1.292310674 | MAP3K4 |
| 6 | 161070503 | 161070699 | -1.685853572 | MAP3K4 |
| 6 | 161131804 | 161132002 | -2.741557621 | AGPAT4 |
| 6 | 161135031 | 161135212 | -1.982432816 | AGPAT4 |
| 6 | 161137406 | 161137619 | 1.685822931 | AGPAT4 |
| 6 | 161139762 | 161139959 | 1.273236953 | AGPAT4 |
| 6 | 161175788 | 161175931 | -7.087618825 | AGPAT4 |
| 6 | 161176410 | 161176610 | -2.879814657 | AGPAT4 |
| 6 | 166374307 | 166374502 | -1.39094137 | MPC1 |
| 6 | 166467669 | 166467860 | 4.04637857 | RPS6KA2 |
| 6 | 166533224 | 166533397 | -5.892317423 | RPS6KA2 |
| 6 | 167969794 | 167969946 | -2.100816931 | AFDN |
| 6 | 169214362 | 169214539 | -2.316134108 | XXyac-YX65C7_A.2 |
| 6 | 169216164 | 169216397 | -2.567661357 | THBS2 |
| 6 | 169216164 | 169216397 | -2.567661357 | XXyac-YX65C7_A.2 |
| 6 | 169237802 | 169238003 | -2.650607693 | THBS2 |
| 6 | 169237802 | 169238003 | -2.650607693 | XXyac-YX65C7_A.2 |
| 6 | 169559081 | 169559276 | -2.426821921 | WDR27 |
| 6 | 169624822 | 169625020 | -1.261113373 | WDR27 |
| 6 | 169625744 | 169625868 | -6.128693921 | WDR27 |

| | | | | |
|---|---|---|---|---|
| X | 113861627 | 113861812 | -7.02117169 | XACT |
| 10 | 12121177 | 12121436 | -1.308227524 | DHTKD1 |
| X | 118392694 | 118392904 | 1.230929178 | WDR44 |
| X | 118768538 | 118768724 | 4.628693921 | IL13RA1 |
| X | 118975042 | 118975241 | 9.63048962 | LONRF3 |
| X | 118998211 | 118998376 | -2.588326821 | LONRF3 |
| X | 119822713 | 119822878 | -5.310568057 | UPF3B |
| X | 119871377 | 119871576 | -1.08303401 | RNF113A |
| X | 119871377 | 119871576 | -1.08303401 | NDUFA1 |
| X | 126164318 | 126164454 | -5.810339021 | DCAF12L2 |
| X | 126551839 | 126552069 | 7.10958426 | DCAF12L1 |
| 10 | 13646359 | 13646569 | 1.049781836 | FRMD4A |
| 10 | 13646359 | 13646569 | 1.049781836 | RP11-295P9.13 |
| 10 | 13646359 | 13646569 | 1.049781836 | RP11-295P9.3 |
| X | 130056105 | 130056230 | 1.375415544 | BCORL1 |
| X | 132218436 | 132218627 | 2.288824156 | RAP2C-AS1 |
| X | 132218436 | 132218627 | 2.288824156 | RAP2C |
| 10 | 13657294 | 13657502 | 1.201831126 | FRMD4A |
| 10 | 13657294 | 13657502 | 1.201831126 | RP11-295P9.3 |
| X | 135293965 | 135294177 | -1.028169862 | ZNF75D |
| X | 135360247 | 135360464 | -1.106558287 | ZNF449 |
| 10 | 13707253 | 13707452 | 1.162937896 | FRMD4A |
| X | 149533086 | 149533298 | 1.124547607 | AF011889.5 |
| X | 149533086 | 149533298 | 1.124547607 | LINC00893 |
| X | 149546747 | 149546847 | 2.928990498 | CXorf40A |
| X | 153347836 | 153348053 | 1.166281019 | ZNF275 |
| 10 | 14567803 | 14567998 | 1.66130429 | FAM107B |
| X | 153459257 | 153459435 | -5.357122759 | HAUS7 |
| X | 153459257 | 153459435 | -5.357122759 | TREX2 |
| X | 153580309 | 153580515 | -1.539092244 | ATP2B3 |
| X | 153724911 | 153725145 | 2.086073121 | ABCD1 |
| X | 153954478 | 153954684 | 1.776482273 | HCFC1 |
| X | 153971091 | 153971209 | 11.74412945 | HCFC1 |
| X | 153982872 | 153983099 | 1.052719732 | TMEM187 |
| 10 | 14842567 | 14842736 | 7.554915327 | HSPA14 |
| X | 154460319 | 154460614 | 2.146507701 | PLXNA3 |
| X | 155383145 | 155383251 | -2.065338126 | F8A2 |
| X | 155383145 | 155383251 | -2.065338126 | MIR1184-2 |
| X | 155457370 | 155457570 | -1.004992044 | F8A3 |
| X | 155457370 | 155457570 | -1.004992044 | MIR1184-3 |
| X | 155776411 | 155776570 | 1.277294426 | SPRY3 |
| Y | 2924675 | 2924858 | 1.130646149 | RPS4Y1 |
| Y | 3069859 | 3070050 | -2.83053274 | LINC00278 |
| Y | 5098094 | 5098294 | 2.02416054 | PCDH11Y |
| Y | 14622121 | 14622300 | 1.569177461 | NLGN4Y |
| Y | 14622344 | 14622489 | 7.778971143 | NLGN4Y |
| Y | 14840725 | 14840935 | 1.093598603 | NLGN4Y |
| Y | 19008567 | 19008777 | -1.130109641 | TTTY14 |
| 10 | 17216229 | 17216416 | 1.988238097 | VIM-AS1 |
| 10 | 22252539 | 22252730 | 3.194971406 | RP11-573G6.4 |
| 10 | 22436293 | 22436513 | 1.424319061 | SPAG6 |
| 10 | 27170141 | 27170336 | 1.133447334 | MASTL |
| 10 | 29467416 | 29467617 | 1.565976548 | SVIL |
| 10 | 29467416 | 29467617 | 1.565976548 | SVIL-AS1 |

| | | | | |
|---|---|---|---|---|
| 6 | 89162077 | 89162297 | -2.005217781 | PM20D2 |
| 6 | 89257371 | 89257639 | 4.414861368 | GABRR2 |
| 1 | 215620073 | 215620274 | -1.937187135 | KCTD3 |
| 6 | 89295482 | 89295681 | 1.735961063 | GABRR2 |
| 6 | 89630874 | 89631083 | 1.438811189 | LYRM2 |
| 6 | 89630874 | 89631083 | 1.438811189 | ANKRD6 |
| 6 | 89637065 | 89637267 | -1.071229962 | LYRM2 |
| 6 | 89643909 | 89644065 | -1.230945269 | MDN1 |
| 6 | 89692568 | 89692782 | -1.153374048 | MDN1 |
| 6 | 89862418 | 89862621 | -2.298162382 | CASP8AP2 |
| 6 | 89864058 | 89864250 | -2.408979846 | CASP8AP2 |
| 6 | 89866112 | 89866329 | -2.406237027 | CASP8AP2 |
| 6 | 89868338 | 89868550 | -2.521347095 | CASP8AP2 |
| 6 | 89932406 | 89932627 | 3.956343149 | BACH2 |
| 6 | 89949377 | 89949483 | 6.462035093 | BACH2 |
| 6 | 89950916 | 89951113 | 7.743103267 | BACH2 |
| 6 | 89950916 | 89951113 | 7.743103267 | RP3-512E2.2 |
| 6 | 95586586 | 95586791 | -1.547566872 | MANEA |
| 6 | 96494619 | 96494832 | -1.547109986 | UFL1-AS1 |
| 6 | 96798705 | 96798848 | -3.82926897 | GPR63 |
| 6 | 96799618 | 96799807 | -2.0069029 | GPR63 |
| 6 | 97186621 | 97186820 | -3.091365292 | MMS22L |
| 6 | 97228957 | 97229185 | -3.389021027 | MMS22L |
| 6 | 97260429 | 97260628 | -6.811792472 | MMS22L |
| 6 | 98835802 | 98836027 | -1.929430966 | POU3F2 |
| 6 | 98926581 | 98926813 | -1.150066112 | FBXL4 |
| 6 | 99446084 | 99446284 | -1.045023148 | USP45 |
| 6 | 99614729 | 99614863 | -2.655533551 | PRDM13 |
| 6 | 100390457 | 100390615 | -1.865185342 | SIM1 |
| 6 | 100584456 | 100584645 | -1.715332847 | ASCC3 |
| 6 | 100740763 | 100740946 | 5.836212671 | ASCC3 |
| 1 | 218441358 | 218441522 | 2.813718944 | TGFB2 |
| 6 | 100848271 | 100848501 | -1.208987327 | ASCC3 |
| 6 | 100859077 | 100859374 | -1.606669007 | ASCC3 |
| 6 | 100882093 | 100882294 | -1.169217154 | RP3-467N11.1 |
| 6 | 105278118 | 105278318 | -1.443616249 | PREP |
| 6 | 105314674 | 105314870 | -4.237810428 | PREP |
| 6 | 106088146 | 106088348 | 2.737041019 | ATG5 |
| 6 | 106088146 | 106088348 | 2.737041019 | PRDM1 |
| 6 | 106104942 | 106105134 | 2.927100206 | PRDM1 |
| 6 | 106104942 | 106105134 | 2.927100206 | ATG5 |
| 6 | 106107077 | 106107260 | 3.031887369 | PRDM1 |
| 6 | 106107077 | 106107260 | 3.031887369 | ATG5 |
| 6 | 106512155 | 106512303 | -6.169368191 | AIM1 |
| 6 | 106629168 | 106629330 | -1.265977626 | RTN4IP1 |
| 6 | 107070141 | 107070269 | -1.06478643 | BEND3 |
| 6 | 107363807 | 107363985 | -3.549016041 | PDSS2 |
| 6 | 107404091 | 107404302 | -1.968814258 | PDSS2 |
| 6 | 107640700 | 107640899 | -1.235925767 | SOBP |
| 6 | 107658984 | 107659265 | 1.518628969 | SOBP |
| 1 | 219197327 | 219197526 | 1.148361815 | LYPLAL1 |
| 6 | 107957493 | 107957647 | -8.306059889 | SEC63 |
| 6 | 107957493 | 107957647 | -8.306059889 | RP1-191J18.66 |
| 6 | 107959500 | 107959693 | -2.443795961 | RP1-191J18.66 |

| | | | | |
|---|---|---|---|---|
| 6 | 169626582 | 169626787 | -1.645541064 | WDR27 |
| 6 | 169655051 | 169655179 | -1.355867624 | WDR27 |
| 1 | 10461561 | 10461758 | -1.810678128 | DFFA |
| 1 | 224192413 | 224192645 | -1.283288393 | DEGS1 |
| 6 | 169657127 | 169657360 | -1.6323375 | WDR27 |
| 6 | 169664148 | 169664398 | -1.854198562 | WDR27 |
| 6 | 169665240 | 169665421 | -2.071516214 | WDR27 |
| 6 | 169667861 | 169668051 | -2.489272106 | WDR27 |
| 6 | 169694796 | 169694992 | -2.242967603 | WDR27 |
| 6 | 169710245 | 169710433 | 5.169925001 | PHF10 |
| 1 | 224389646 | 224389846 | -1.115023977 | WDR26 |
| 1 | 224598134 | 224598323 | -3.771326699 | CNIH3 |
| 6 | 170162399 | 170162599 | -1.955674854 | RP11-302L19.1 |
| 6 | 170271383 | 170271509 | -1.850219859 | RP5-894D12.5 |
| 6 | 170304718 | 170304913 | -2.690699373 | FAM120B |
| 6 | 170304718 | 170304913 | -2.690699373 | DLL1 |
| 6 | 170317480 | 170317675 | -1.370166031 | FAM120B |
| 6 | 170581880 | 170582071 | 7.148350581 | PDCD2 |
| 7 | 80463 | 80669 | -1.220893337 | AC093627.8 |
| 7 | 80463 | 80669 | -1.220893337 | AC093627.7 |
| 7 | 82640 | 82818 | -1.225070443 | AC093627.7 |
| 1 | 224739326 | 224739526 | -1.90743135 | CNIH3 |
| 7 | 137337 | 137514 | -3.363055979 | AC093627.9 |
| 7 | 145460 | 145654 | -1.385380446 | AC093627.9 |
| 7 | 147649 | 147798 | -2.088993637 | AC093627.9 |
| 7 | 153934 | 154100 | -1.528215208 | AC093627.10 |
| 7 | 163339 | 163533 | -2.093963841 | RP11-90P13.1 |
| 1 | 224772698 | 224772896 | -3.487989102 | RP11-449J1.1 |
| 7 | 196089 | 196376 | -1.267741228 | FAM20C |
| 7 | 197197 | 197481 | -11.76398867 | FAM20C |
| 7 | 203645 | 203843 | 9.084076514 | FAM20C |
| 7 | 205146 | 205388 | 7.857444717 | FAM20C |
| 7 | 214884 | 215095 | 1.576019808 | FAM20C |
| 7 | 218660 | 218944 | 2.616753665 | FAM20C |
| 7 | 242891 | 243101 | -1.203598373 | FAM20C |
| 7 | 250822 | 251025 | 7.406289222 | FAM20C |
| 1 | 224977737 | 224977920 | -2.803383257 | DNAH14 |
| 7 | 605167 | 605348 | 2.0559235 | AC147651.4 |
| 7 | 605167 | 605348 | 2.0559235 | PRKAR1B |
| 7 | 751857 | 752072 | -1.595902051 | DNAAF5 |
| 7 | 873301 | 873458 | -4.249175304 | SUN1 |
| 7 | 881797 | 882039 | -6.377443751 | SUN1 |
| 7 | 881797 | 882039 | -6.377443751 | RP11-449P15.2 |
| 7 | 881797 | 882039 | -6.377443751 | GET4 |
| 7 | 884839 | 884959 | -3.745926548 | RP11-449P15.2 |
| 7 | 884839 | 884959 | -3.745926548 | GET4 |
| 7 | 884839 | 884959 | -3.745926548 | SUN1 |
| 7 | 895059 | 895246 | -6.142701109 | SUN1 |
| 7 | 895059 | 895246 | -6.142701109 | GET4 |
| 7 | 898780 | 898979 | -6.774410954 | COX19 |
| 7 | 898780 | 898979 | -6.774410954 | ADAP1 |
| 7 | 907541 | 907740 | 6.477279923 | ADAP1 |
| 7 | 907541 | 907740 | 6.477279923 | COX19 |
| 7 | 1026297 | 1026476 | 7.65226479 | C7orf50 |

| | | | | |
|---|---|---|---|---|
| 10 | 30844934 | 30845140 | -1.099215715 | ZNF438 |
| 10 | 30849196 | 30849412 | -1.28772288 | ZNF438 |
| 10 | 30928688 | 30928869 | 1.466442902 | ZNF438 |
| 10 | 31318404 | 31318609 | 1.27137236 | ZEB1 |
| 10 | 31318404 | 31318609 | 1.27137236 | ZEB1-AS1 |
| 10 | 31408847 | 31408971 | -1.323154877 | ZEB1 |
| 10 | 31455125 | 31455336 | -1.910262189 | ZEB1 |
| 10 | 31455985 | 31456140 | -6.037739575 | ZEB1 |
| 10 | 31474673 | 31474827 | 5.062060656 | ZEB1 |
| 10 | 31475938 | 31476042 | 4.86735481 | ZEB1 |
| 10 | 31891650 | 31891821 | -1.668061851 | ARHGAP12 |
| 10 | 32269035 | 32269234 | -1.106250836 | RP11-166N17.1 |
| 10 | 32269035 | 32269234 | -1.106250836 | EPC1 |
| 10 | 37829136 | 37829357 | 1.544780057 | AL135791.1 |
| 10 | 37829136 | 37829357 | 1.544780057 | ZNF248 |
| 10 | 38064031 | 38064292 | 1.338010583 | ZNF33A |
| 1 | 15734152 | 15734326 | 1.429146784 | PLEKHM2 |
| 1 | 15734152 | 15734326 | 1.429146784 | RP11-288I21.1 |
| 10 | 42337214 | 42337340 | -6.772964885 | RP11-313J2.1 |
| 1 | 15739858 | 15739967 | -1.460282766 | SLC25A34 |
| 10 | 42534747 | 42534960 | -5.980000966 | ZNF37BP |
| 10 | 42831029 | 42831220 | 1.172725199 | BMS1 |
| 1 | 15836078 | 15836362 | 1.064041698 | FLJ37453 |
| 10 | 45004787 | 45004972 | -1.771571163 | ZNF22 |
| 10 | 45004787 | 45004972 | -1.771571163 | CEP164P1 |
| 10 | 45429231 | 45429397 | 3.014873672 | ALOX5 |
| 10 | 45458352 | 45458554 | -1.327396853 | Mar-08 |
| 10 | 46785827 | 46786112 | 1.301052645 | GLUD1P7 |
| 10 | 46888179 | 46888375 | 1.097097977 | FRMPD2B |
| 10 | 48450640 | 48450825 | 1.053839423 | ARHGAP22 |
| 10 | 48574732 | 48574869 | -4.63339327 | ARHGAP22 |
| 10 | 48823365 | 48823530 | -4.769579406 | WDFY4 |
| 10 | 49515725 | 49515948 | -1.463903088 | PGBD3 |
| 10 | 49515725 | 49515948 | -1.463903088 | ERCC6 |
| 10 | 49969830 | 49969970 | -2.289862926 | PARG |
| 10 | 49969830 | 49969970 | -2.289862926 | TIMM23B |
| 10 | 52730633 | 52730816 | -1.275305498 | RP11-556E13.1 |
| 10 | 52731337 | 52731535 | -1.027441765 | RP11-556E13.1 |
| 10 | 52734233 | 52734422 | -1.018130393 | RP11-556E13.1 |
| 10 | 58828875 | 58829050 | 1.505269482 | BICC1 |
| 10 | 59654177 | 59654377 | 1.39958581 | SLC16A9 |
| 10 | 62053058 | 62053267 | 1.953491925 | ARID5B |
| 10 | 62197969 | 62198191 | 1.142409273 | RTKN2 |
| 10 | 63465660 | 63465862 | 2.922948754 | JMJD1C-AS1 |
| 10 | 63465660 | 63465862 | 2.922948754 | JMJD1C |
| 10 | 65653141 | 65653302 | -1.184258827 | LINC01515 |
| 1 | 16231785 | 16231958 | 1.76759229 | RSG1 |
| 10 | 69123588 | 69123734 | 2.627907711 | VPS26A |
| 10 | 70098540 | 70098738 | -6.909291089 | AIFM2 |
| 10 | 70098540 | 70098738 | -6.909291089 | H2AFY2 |
| 10 | 70108414 | 70108627 | -1.105262055 | H2AFY2 |
| 10 | 70108414 | 70108627 | -1.105262055 | AIFM2 |
| 10 | 70566729 | 70566950 | 1.731206789 | PALD1 |
| 10 | 70760662 | 70760861 | 1.788802607 | ADAMTS14 |

| | | | | | | | | | | | | | | |
|---|---|---|---|---|---|---|---|---|---|---|---|---|---|---|
| 6 | 108411310 | 108411557 | -1.382299441 | LACE1 | 7 | 1026648 | 1026848 | -3.61820528 | C7orf50 | 10 | 70877237 | 70877433 | 1.561975071 | SGPL1 |
| 1 | 10304397 | 10304598 | 1.287137156 | KIF1B | 7 | 1045849 | 1046051 | 3.609123901 | GPR146 | 10 | 71810198 | 71810383 | -1.69712391 | CDH23 |
| 6 | 108907799 | 108907993 | -2.430754544 | ARMC2 | 7 | 1045849 | 1046051 | 3.609123901 | C7orf50 | 10 | 72008533 | 72008732 | 2.420595366 | CHST3 |
| 6 | 108907799 | 108907993 | -2.430754544 | RP11-249L21.4 | 7 | 1047942 | 1048131 | 7.350165669 | GPR146 | 10 | 72354677 | 72354821 | -6.252037057 | DNAJB12 |
| 6 | 109133569 | 109133673 | -1.817343937 | CEP57L1 | 7 | 1047942 | 1048131 | 7.350165669 | C7orf50 | 10 | 72516378 | 72516623 | 1.302431029 | MICU1 |
| 6 | 109135398 | 109135653 | -2.195448374 | CEP57L1 | 7 | 1050865 | 1051051 | 6.496292172 | C7orf50 | 10 | 73166846 | 73166988 | -7.085431975 | ECD |
| 6 | 109368722 | 109368927 | 2.116282818 | CD164 | 7 | 1050865 | 1051051 | 6.496292172 | GPR146 | 10 | 73498455 | 73498643 | 4.027641218 | USP54 |
| 6 | 109383034 | 109383309 | 2.460672185 | RP11-425D10.10 | 7 | 1051451 | 1051623 | 3.29248125 | GPR146 | 10 | 73498455 | 73498643 | 4.027641218 | PPP3CB-AS1 |
| 6 | 109415193 | 109415384 | 2.505578895 | PPIL6 | 7 | 1051451 | 1051623 | 3.29248125 | C7orf50 | 1 | 16462092 | 16462364 | -1.087587733 | RP4-798A10.2 |
| 6 | 109465995 | 109466193 | -1.246078013 | ZBTB24 | 7 | 1054846 | 1055046 | 6.590576128 | GPR146 | 10 | 73516582 | 73516782 | 1.021400352 | PPP3CB-AS1 |
| 6 | 110144456 | 110144649 | -1.73460632 | WASF1 | 7 | 1054846 | 1055046 | 6.590576128 | C7orf50 | 10 | 73516582 | 73516782 | 1.021400352 | USP54 |
| 6 | 110442410 | 110442610 | -1.435587929 | SLC22A16 | 1 | 10629690 | 10629897 | -1.210698578 | PEX14 | 10 | 73647436 | 73647687 | 3.447613839 | SYNPO2L |
| 1 | 219610610 | 219610809 | 1.839088131 | ZC3H11B | 1 | 225497567 | 225497767 | -1.507820975 | ENAH | 10 | 73648515 | 73648688 | 2.335185076 | SYNPO2L |
| 6 | 111375944 | 111376138 | -1.075182004 | REV3L | 7 | 1058339 | 1058618 | 3.340109438 | GPR146 | 10 | 73674707 | 73674808 | -1.527067855 | RP11-464F9.1 |
| 1 | 220263231 | 220263530 | -7.732017576 | RAB3GAP2 | 7 | 1058339 | 1058618 | 3.340109438 | RP11-449P15.1 | 10 | 73674707 | 73674808 | -1.527067855 | RP11-464F9.21 |
| 6 | 111601296 | 111601472 | -2.306458281 | TRAF3IP2 | 7 | 1058339 | 1058618 | 3.340109438 | C7orf50 | 10 | 73674707 | 73674808 | -1.527067855 | AGAP5 |
| 6 | 111601296 | 111601472 | -2.306458281 | C6orf3 | 7 | 1066901 | 1067113 | -1.406934115 | C7orf50 | 10 | 73675160 | 73675348 | 1.287045922 | RP11-464F9.1 |
| 6 | 113860518 | 113860705 | 1.845685513 | MARCKS | 7 | 1164452 | 1164595 | -1.587932886 | AC091729.9 | 10 | 73675160 | 73675348 | 1.287045922 | AGAP5 |
| 6 | 113868499 | 113868617 | -1.643956971 | LINC01268 | 7 | 1587772 | 1587972 | -1.231303807 | PSMG3-AS1 | 10 | 73730321 | 73730528 | 2.437347952 | RP11-464F9.1 |
| 6 | 113995466 | 113995650 | -4.115847389 | RP3-399L15.3 | 7 | 1747043 | 1747249 | -2.742480353 | ELFN1 | 10 | 73730321 | 73730528 | 2.437347952 | BMS1P4 |
| 6 | 113995466 | 113995650 | -4.115847389 | HDAC2 | 7 | 1841797 | 1841985 | -2.307354922 | AC110781.3 | 10 | 73731095 | 73731266 | 3.7667161 | GLUD1P3 |
| 1 | 220267100 | 220267295 | -1.017691984 | RAB3GAP2 | 7 | 1841797 | 1841985 | -2.307354922 | MAD1L1 | 10 | 73732121 | 73732317 | 2.681562297 | GLUD1P3 |
| 1 | 220267100 | 220267295 | -1.017691984 | AURKAPS1 | 7 | 1842572 | 1842765 | -5.374096425 | MAD1L1 | 10 | 73732121 | 73732317 | 2.681562297 | DUSP8P5 |
| 6 | 116435686 | 116435874 | 3.873047192 | DSE | 7 | 1842572 | 1842765 | -5.374096425 | AC110781.3 | 10 | 73733435 | 73733623 | 10.05840993 | GLUD1P3 |
| 6 | 116436719 | 116436924 | 3.270719325 | DSE | 7 | 2110051 | 2110266 | -7.387496454 | MAD1L1 | 10 | 73733435 | 73733623 | 10.05840993 | DUSP8P5 |
| 6 | 116666752 | 116666951 | -1.322469971 | ZUFSP | 7 | 2132566 | 2132768 | -1.459637532 | MAD1L1 | 1 | 16563988 | 16564183 | 4.834442492 | NBPF1 |
| 1 | 220813517 | 220813713 | -2.092897884 | Mar-01 | 7 | 2133559 | 2133767 | -8.079068143 | MAD1L1 | 10 | 73916583 | 73916789 | 2.148422502 | PLAU |
| 6 | 117997994 | 117998165 | -7.139696471 | SLC35F1 | 7 | 2289508 | 2289689 | -1.203587691 | SNX8 | 10 | 73916583 | 73916789 | 2.148422502 | C10orf55 |
| 6 | 118049994 | 118050097 | -5.950056031 | SLC35F1 | 7 | 2432701 | 2432907 | 9.645630325 | CHST12 | 10 | 74352387 | 74352603 | 6.400753143 | ADK |
| 6 | 118834886 | 118835047 | 5.365659516 | MCM9 | 7 | 2443508 | 2443694 | -5.766664866 | CHST12 | 10 | 74843005 | 74843204 | -1.07729921 | KAT6B |
| 6 | 119024320 | 119024499 | -2.644476301 | FAM184A | 7 | 2538017 | 2538216 | -8.544642475 | BRAT1 | 10 | 75028771 | 75028969 | -2.58662268 | KAT6B |
| 6 | 119024320 | 119024499 | -2.644476301 | RP11-351A11.1 | 7 | 2538422 | 2538594 | -3.041939539 | BRAT1 | 10 | 75398674 | 75398879 | 2.878334585 | ZNF503 |
| 6 | 119306951 | 119307152 | 1.644424252 | MAN1A1 | 7 | 2613697 | 2613906 | -1.597356872 | IQCE | 10 | 75398674 | 75398879 | 2.878334585 | RP11-399K21.11 |
| 1 | 220831246 | 220831441 | 3.014157694 | LINC01352 | 7 | 2718379 | 2718555 | -2.215708621 | AMZ1 | 10 | 75399839 | 75400033 | 10.61047047 | ZNF503 |
| 6 | 119348542 | 119348747 | 1.221389371 | MAN1A1 | 7 | 2721957 | 2722153 | -2.114946795 | AMZ1 | 10 | 75399839 | 75400033 | 10.61047047 | RP11-399K21.11 |
| 6 | 121100072 | 121100239 | 1.359114516 | TBC1D32 | 7 | 2723890 | 2723999 | -2.80966627 | AMZ1 | 10 | 75401962 | 75402091 | 1.749811014 | ZNF503-AS2 |
| 1 | 220861858 | 220861965 | 6.379319819 | HLX-AS1 | 7 | 2728762 | 2728945 | -1.713994619 | AMZ1 | 10 | 75580892 | 75581094 | -1.027219831 | C10orf11 |
| 6 | 125889366 | 125889561 | 1.644404654 | NCOA7 | 7 | 2728762 | 2728945 | -1.713994619 | GNA12 | 1 | 1082669 | 1082824 | 3.262373087 | C1orf159 |
| 6 | 125889613 | 125889809 | 1.531473481 | NCOA7 | 7 | 2729347 | 2729545 | -8.718144306 | GNA12 | 10 | 76179385 | 76179588 | -1.253747111 | C10orf11 |
| 6 | 125928734 | 125928934 | 2.034796946 | NCOA7 | 7 | 2729347 | 2729545 | -8.718144306 | AMZ1 | 10 | 76222129 | 76222297 | -5.807354922 | C10orf11 |
| 6 | 126644097 | 126644256 | -1.882669933 | PRELID1P1 | 7 | 2731262 | 2731463 | -1.363188836 | GNA12 | 10 | 76475350 | 76475511 | -5.114409345 | C10orf11 |
| 6 | 127363340 | 127363539 | -2.076671194 | RPL5P18 | 7 | 2731262 | 2731463 | -1.363188836 | AMZ1 | 10 | 77780949 | 77781109 | -7.005089202 | RP13-39P12.2 |
| 6 | 127446871 | 127447055 | 1.756917284 | KIAA0408 | 7 | 3991034 | 3991234 | 1.157787263 | SDK1 | 10 | 78054505 | 78054724 | -1.665118549 | RPS24 |
| 6 | 127446871 | 127447055 | 1.756917284 | RP3-403A15.5 | 7 | 4267833 | 4267999 | -1.032531728 | SDK1 | 10 | 79315278 | 79315471 | -1.075352141 | ZMIZ1 |
| 6 | 127516445 | 127516682 | 1.17988078 | RP3-403A15.5 | 7 | 4762434 | 4762633 | -1.092708583 | FOXK1 | 10 | 79353843 | 79354041 | 11.01027976 | PPIF |
| 6 | 127516445 | 127516682 | 1.17988078 | SOGA3 | 7 | 4764029 | 4764228 | -1.616246674 | FOXK1 | 10 | 79701170 | 79701383 | 1.15505133 | NUTM2B-AS1 |
| 6 | 129589345 | 129589548 | 1.086683969 | ARHGAP18 | 7 | 4766770 | 4766971 | -1.405501361 | FOXK1 | 10 | 79711698 | 79711873 | -6.226506677 | NUTM2B-AS1 |
| 6 | 130139570 | 130139774 | -1.444296336 | L3MBTL3 | 7 | 5067063 | 5067228 | -1.284229793 | RBAK | 10 | 79711698 | 79711873 | -6.226506677 | NUTM2B |
| 6 | 130139570 | 130139774 | -1.444296336 | RP11-73O6.3 | 7 | 5067063 | 5067228 | -1.284229793 | RBAK-RBAKDN | 1 | 17068851 | 17069073 | -3.425348779 | PADI2 |
| 6 | 130265198 | 130265380 | -7.968088911 | SAMD3 | 1 | 226152421 | 226152626 | -1.07457557 | ACBD3 | 10 | 79958364 | 79958534 | -2.150570247 | SFTPD |
| 6 | 130440835 | 130440946 | -4.926462393 | TMEM200A | 1 | 226152421 | 226152626 | -1.07457557 | RP11-275I14.4 | 1 | 17115541 | 17115727 | -5.938833787 | PADI2 |
| 6 | 130441543 | 130441704 | -4.393298181 | TMEM200A | 7 | 5071676 | 5071795 | -6.136460663 | RBAK-RBAKDN | 10 | 86709672 | 86709779 | 2.377686038 | LDB3 |
| 6 | 131281842 | 131282042 | -2.811890875 | AKAP7 | 7 | 5230818 | 5231054 | -1.28850734 | WIPI2 | 1 | 17407853 | 17408053 | -1.021818133 | RCC2 |
| 6 | 131949202 | 131949391 | -2.138897027 | CTGF | 7 | 5232457 | 5232658 | -6.623370299 | WIPI2 | 10 | 87009549 | 87009692 | -1.215563433 | RP11-96C23.14 |

| | | | | |
|---|---|---|---|---|
| 1 | 10377434 | 10377543 | -6.248426889 | KIF1B |
| 6 | 132352885 | 132353084 | -1.612422438 | MOXD1 |
| 6 | 135465950 | 135466149 | -1.022785753 | AHI1 |
| 6 | 136139149 | 136139343 | 5.623963757 | RP13-143G15.4 |
| 6 | 136139149 | 136139343 | 5.623963757 | PDE7B |
| 6 | 136192088 | 136192288 | -3.304852378 | RP13-143G15.4 |
| 6 | 136192088 | 136192288 | -3.304852378 | PDE7B |
| 6 | 136276039 | 136276239 | -1.224948196 | BCLAF1 |
| 6 | 137198243 | 137198434 | 1.490232501 | IFNGR1 |
| 6 | 137879004 | 137879204 | 1.223251393 | TNFAIP3 |
| 6 | 137879966 | 137880166 | 1.176746214 | TNFAIP3 |
| 1 | 221702402 | 221702647 | 1.31272249 | DUSP10 |
| 6 | 138415278 | 138415473 | -1.494927648 | HEBP2 |
| 6 | 138418778 | 138418892 | -1.512181333 | HEBP2 |
| 6 | 138420000 | 138420154 | -1.760956279 | HEBP2 |
| 6 | 138650303 | 138650445 | 5.397207933 | NHSL1 |
| 6 | 138650303 | 138650445 | 5.397207933 | RP11-390P2.2 |
| 1 | 221739055 | 221739257 | 1.729796374 | DUSP10 |
| 6 | 138730109 | 138730287 | -1.877443751 | GVQW2 |
| 6 | 139372606 | 139372805 | -3.422024137 | CITED2 |
| 6 | 139373132 | 139373318 | -1.136878034 | CITED2 |
| 6 | 139373606 | 139373743 | -12.17957477 | CITED2 |
| 6 | 142211643 | 142211821 | -1.473822297 | VTA1 |
| 6 | 142753245 | 142753508 | -1.720037777 | RP1-67K17.3 |
| 6 | 142753245 | 142753508 | -1.720037777 | HIVEP2 |
| 6 | 142760203 | 142760409 | -2.045074355 | HIVEP2 |
| 6 | 142769806 | 142769999 | -5.251525519 | HIVEP2 |
| 6 | 142770430 | 142770624 | -2.88832291 | HIVEP2 |
| 6 | 142772648 | 142772845 | -2.103573923 | HIVEP2 |
| 6 | 142773861 | 142774085 | -2.012088543 | HIVEP2 |
| 6 | 142774627 | 142774828 | -1.791822341 | HIVEP2 |
| 6 | 143338143 | 143338305 | -4.011405565 | AIG1 |
| 6 | 143343204 | 143343400 | -4.567480514 | RP1-95L4.4 |
| 6 | 143948212 | 143948385 | -3.043377176 | PLAGL1 |
| 6 | 144095029 | 144095223 | -2.128471794 | SF3B5 |
| 1 | 222628889 | 222629067 | -1.559287611 | MIA3 |
| 6 | 145871843 | 145872041 | 1.540630751 | RP11-545I5.3 |
| 6 | 145871843 | 145872041 | 1.540630751 | SHPRH |
| 6 | 145908785 | 145908985 | -1.362036401 | SHPRH |
| 1 | 222629854 | 222630052 | -1.122487237 | MIA3 |
| 6 | 147303511 | 147303701 | -7.579423166 | STXBP5 |
| 6 | 147407808 | 147407991 | 3.531197488 | YAP1P1 |
| 6 | 149027437 | 149027747 | 3.74383265 | UST |
| 6 | 149027437 | 149027747 | 3.74383265 | RP11-162J8.3 |
| 1 | 222665379 | 222665578 | -1.467475781 | MIA3 |
| 6 | 149858221 | 149858419 | -6.37196576 | LRP11 |
| 6 | 149862393 | 149862583 | -7.557257532 | LRP11 |
| 6 | 149946398 | 149946550 | 3.068546804 | ULBP2 |
| 6 | 149969021 | 149969220 | 1.674799466 | ULBP1 |
| 6 | 149972392 | 149972534 | 2.711159223 | ULBP1 |
| 6 | 151091160 | 151091284 | -1.608808312 | MTHFD1L |
| 6 | 151091160 | 151091284 | -1.608808312 | RP1-292B18.4 |
| 6 | 151414568 | 151414769 | 1.098573235 | RMND1 |
| 1 | 10378236 | 10378414 | 1.018544659 | KIF1B |
| 7 | 5233208 | 5233421 | -2.532073044 | WIPI2 |
| 7 | 5351997 | 5352192 | -1.761921115 | TNRC18 |
| 7 | 5357061 | 5357287 | -1.806278429 | TNRC18 |
| 1 | 226309759 | 226309879 | -1.36268881 | LIN9 |
| 7 | 5390565 | 5390743 | -3.168500171 | TNRC18 |
| 7 | 5429106 | 5429306 | -3.332446282 | RP11-1275H24.3 |
| 7 | 5495495 | 5495606 | -4.204695468 | FBXL18 |
| 7 | 5741195 | 5741308 | -6.235837607 | RNF216 |
| 7 | 5973211 | 5973384 | -5.231251136 | PMS2 |
| 7 | 5987139 | 5987424 | -1.019674847 | PMS2 |
| 7 | 6154906 | 6155124 | 1.141419906 | USP42 |
| 7 | 6156710 | 6156957 | 1.362970333 | USP42 |
| 7 | 6164812 | 6165012 | -1.109144115 | CYTH3 |
| 7 | 6409558 | 6409866 | -1.182657338 | DAGLB |
| 7 | 6652216 | 6652425 | -2.006598391 | ZNF316 |
| 7 | 6653196 | 6653327 | 5.178776002 | ZNF316 |
| 7 | 6676041 | 6676234 | -1.542518052 | RP11-611L7.2 |
| 7 | 6676041 | 6676234 | -1.542518052 | AC073343.13 |
| 7 | 6691893 | 6692090 | -1.461997541 | AC073343.13 |
| 7 | 6691893 | 6692090 | -1.461997541 | ZNF12 |
| 1 | 10639761 | 10639957 | 2.75528854 | CASZ1 |
| 1 | 10639761 | 10639957 | 2.75528854 | RP4-734G22.3 |
| 7 | 6751855 | 6752052 | -5.382435795 | PMS2CL |
| 7 | 6794141 | 6794331 | -2.087185453 | RSPH10B2 |
| 7 | 6794141 | 6794331 | -2.087185453 | CCZ1B |
| 7 | 7607174 | 7607373 | -1.008836757 | MIOS |
| 7 | 7946155 | 7946315 | -2.025640252 | UMAD1 |
| 7 | 7946155 | 7946315 | -2.025640252 | RP11-505D17.1 |
| 1 | 226728998 | 226729229 | -1.790087338 | ITPKB |
| 7 | 8086472 | 8086672 | -1.56637839 | GLCCI1 |
| 7 | 11348814 | 11348972 | -2.842809357 | AC004538.3 |
| 7 | 11351495 | 11351691 | -3.217992573 | AC004538.3 |
| 7 | 11375544 | 11375849 | -1.347225824 | AC004538.3 |
| 7 | 11375544 | 11375849 | -1.347225824 | THSD7A |
| 7 | 12219489 | 12219614 | -2.179312564 | TMEM106B |
| 7 | 12370089 | 12370277 | -1.707856283 | VWDE |
| 1 | 226735797 | 226736004 | -2.667214066 | ITPKB |
| 7 | 12499192 | 12499377 | -3.806401733 | AC005281.1 |
| 7 | 12502607 | 12502727 | -5.934411277 | AC005281.1 |
| 7 | 12688550 | 12688752 | -1.474497578 | ARL4A |
| 7 | 17793994 | 17794165 | -1.203559028 | SNX13 |
| 7 | 18188280 | 18188482 | -1.707564334 | HDAC9 |
| 1 | 226895609 | 226895919 | -2.100127884 | PSEN2 |
| 7 | 18666190 | 18666448 | -1.893903184 | HDAC9 |
| 7 | 20331220 | 20331416 | 9.486231902 | ITGB8 |
| 7 | 20331220 | 20331416 | 9.486231902 | CTA-293F17.1 |
| 7 | 21429787 | 21429958 | -1.29383316 | SP4 |
| 1 | 226926988 | 226927232 | -2.077451568 | COQ8A |
| 7 | 22882053 | 22882237 | -4.86904613 | RPL12P10 |
| 7 | 22945911 | 22946113 | 1.54850738 | FAM126A |
| 7 | 23200467 | 23200671 | -1.177365082 | NUPL2 |
| 7 | 23611336 | 23611499 | -1.71692467 | CCDC126 |
| 7 | 23642819 | 23643019 | -1.712416923 | CCDC126 |
| 1 | 226928022 | 226928220 | -1.931388331 | COQ8A |
| 10 | 87009549 | 87009692 | -1.215563433 | AGAP11 |
| 10 | 87009549 | 87009692 | -1.215563433 | RP11-96C23.13 |
| 10 | 87009549 | 87009692 | -1.215563433 | RP11-96C23.11 |
| 10 | 87213986 | 87214198 | 1.719372247 | NUTM2A-AS1 |
| 10 | 87366168 | 87366348 | -1.5069029 | NUTM2D |
| 10 | 87505240 | 87505364 | 9.066726656 | MINPP1 |
| 10 | 87862731 | 87862915 | 1.192269278 | KLLN |
| 10 | 88321819 | 88321919 | -4.350219859 | RNLS |
| 10 | 89584285 | 89584464 | 1.20128376 | PANK1 |
| 10 | 89765356 | 89765476 | 3.142701109 | KIF20B |
| 10 | 91259791 | 91259988 | 1.276638238 | PCGF5 |
| 10 | 94532952 | 94533149 | -1.549044198 | TBC1D12 |
| 10 | 94532952 | 94533149 | -1.549044198 | HELLS |
| 10 | 95663563 | 95663756 | 1.179567616 | TCTN3 |
| 1 | 789218 | 789408 | 1.061851973 | RP5-857K21.4 |
| 1 | 789218 | 789408 | 1.061851973 | RP11-206L10.9 |
| 1 | 1083551 | 1083758 | 1.221786705 | C1orf159 |
| 10 | 96983664 | 96983831 | -1.13463835 | LCOR |
| 10 | 97309555 | 97309789 | -2.321928095 | RPL12P27 |
| 10 | 97570323 | 97570593 | 1.079451095 | UBTD1 |
| 10 | 97675443 | 97675603 | 8.64691542 | PI4K2A |
| 10 | 97714635 | 97714853 | 1.43712439 | MARVELD1 |
| 10 | 98223078 | 98223249 | -2.14596596 | R3HCC1L |
| 1 | 19657127 | 19657327 | 1.179665853 | MINOS1-NBL1 |
| 1 | 19657127 | 19657327 | 1.179665853 | NBL1 |
| 10 | 99876858 | 99877057 | 1.41705196 | DNMBP |
| 10 | 99929748 | 99929997 | 1.063666788 | DNMBP |
| 10 | 99929748 | 99929997 | 1.063666788 | DNMBP-AS1 |
| 10 | 99956354 | 99956477 | 1.23977513 | DNMBP |
| 10 | 99956354 | 99956477 | 1.23977513 | DNMBP-AS1 |
| 10 | 100382630 | 100382867 | 1.389448614 | OLMALINC |
| 10 | 100924594 | 100924753 | 1.705635638 | SLF2 |
| 10 | 100987749 | 100987984 | 1.015959546 | C10orf2 |
| 10 | 101006713 | 101006886 | 10.34231182 | LZTS2 |
| 10 | 101060653 | 101060763 | 2.269579406 | RP11-108L7.15 |
| 10 | 101067708 | 101067910 | 1.398181522 | KAZALD1 |
| 10 | 101136889 | 101137116 | 1.138802218 | TLX1 |
| 10 | 101798062 | 101798262 | 6.663425387 | MGEA5 |
| 10 | 102643119 | 102643317 | 2.864128236 | RP11-47A8.5 |
| 10 | 102644511 | 102644707 | 8.690298173 | TRIM8 |
| 10 | 102927245 | 102927445 | -1.427039827 | CNNM2 |
| 10 | 103445408 | 103445620 | 1.216437846 | PDCD11 |
| 1 | 20618573 | 20618772 | -4.353679566 | CDA |
| 10 | 103677052 | 103677242 | -1.210999001 | SH3PXD2A |
| 10 | 112595912 | 112596104 | -5.762760405 | VTI1A |
| 10 | 113886611 | 113886917 | -1.807018739 | NHLRC2 |
| 1 | 1087171 | 1087322 | -2.044087906 | C1orf159 |
| 10 | 114125112 | 114125306 | -1.876680016 | CCDC186 |
| 10 | 114861541 | 114861712 | -1.769398854 | FAM160B1 |
| 1 | 20832556 | 20832788 | -1.023892707 | EIF4G3 |
| 10 | 119037213 | 119037413 | -1.890340975 | EIF3A |
| 10 | 122971416 | 122971564 | 7.158776968 | PSTK |
| 10 | 123135960 | 123136110 | 6.671315149 | HMX3 |
| 10 | 123137053 | 123137237 | 9.740150551 | HMX3 |

| | | | | |
|---|---|---|---|---|
| 6 | 151445433 | 151445602 | -1.56822467 | RMND1 |
| 6 | 152975271 | 152975472 | -3.259618011 | FBXO5 |
| 6 | 152983351 | 152983515 | -2.793888758 | FBXO5 |
| 6 | 152983351 | 152983515 | -2.793888758 | RP1-101K10.6 |
| 6 | 154222752 | 154222940 | 2.238897054 | IPCEF1 |
| 6 | 154222752 | 154222940 | 2.238897054 | OPRM1 |
| 6 | 154860621 | 154860801 | -8.127025276 | TIAM2 |
| 6 | 155129489 | 155129735 | -1.279920075 | TIAM2 |
| 6 | 155257782 | 155257981 | -1.431189946 | TFB1M |
| 6 | 156778035 | 156778268 | -9.572218344 | ARID1B |
| 6 | 156778035 | 156778268 | -9.572218344 | RP11-230C9.2 |
| 6 | 157292726 | 157292928 | -1.293749336 | TMEM242 |
| 6 | 157882797 | 157882993 | 8.16614068 | SNX9 |
| 6 | 158086588 | 158086748 | -7.289952577 | SYNJ2 |
| 6 | 158111011 | 158111180 | 1.378121783 | SERAC1 |
| 6 | 158197020 | 158197147 | -2.647966443 | GTF2H5 |
| 6 | 158502026 | 158502127 | -1.273234491 | TULP4 |
| 6 | 158507126 | 158507324 | 1.034585623 | TULP4 |
| 1 | 223952092 | 223952247 | -1.884506849 | GTF2IP20 |
| 6 | 158609584 | 158609841 | -1.334888463 | MIR7161 |
| 6 | 158609584 | 158609841 | -1.334888463 | TATDN2P2 |
| 6 | 158609584 | 158609841 | -1.334888463 | TMEM181 |
| 6 | 158614827 | 158614943 | 5.991853096 | TMEM181 |
| 6 | 158614827 | 158614943 | 5.991853096 | TATDN2P2 |
| 6 | 158631189 | 158631322 | 1.26368012 | TMEM181 |
| 6 | 158977572 | 158977783 | 1.173861472 | RSPH3 |
| 6 | 159047908 | 159048106 | -1.418471137 | RP1-111C20.4 |
| 6 | 159726554 | 159726755 | -1.67332314 | SOD2 |
| 6 | 159726554 | 159726755 | -1.67332314 | WTAP |
| 6 | 160036524 | 160036709 | 5.241403978 | IGF2R |
| 6 | 160092741 | 160092935 | -7.292950725 | IGF2R |
| 6 | 160991327 | 160991472 | 4.553827339 | RP3-428L16.2 |
| 6 | 160992178 | 160992366 | 9.053332584 | RP3-428L16.2 |
| 6 | 160992178 | 160992366 | 9.053332584 | MAP3K4 |
| 6 | 161131804 | 161132002 | 1.872080548 | AGPAT4 |
| 6 | 161133511 | 161133711 | 1.962045823 | AGPAT4 |
| 6 | 161135031 | 161135212 | 1.407013236 | AGPAT4 |
| 6 | 161136470 | 161136674 | 2.270063847 | AGPAT4 |
| 6 | 161139762 | 161139959 | 1.693800173 | AGPAT4 |
| 6 | 161154954 | 161155154 | 1.510746594 | AGPAT4 |
| 6 | 161172982 | 161173087 | 6.963333315 | AGPAT4 |
| 6 | 161175788 | 161175931 | 7.087618825 | AGPAT4 |
| 6 | 161176410 | 161176610 | 3.400724745 | AGPAT4 |
| 6 | 161193188 | 161193382 | 3.44130254 | AGPAT4 |
| 6 | 161221929 | 161222122 | 1.756722947 | AGPAT4 |
| 6 | 166533224 | 166533397 | 4.307354922 | RPS6KA2 |
| 6 | 167951917 | 167952100 | 1.135487876 | AFDN |
| 6 | 167962329 | 167962504 | 1.704402773 | AFDN |
| 6 | 167969794 | 167969946 | 7.076459288 | AFDN |
| 6 | 169214362 | 169214539 | -1.355091298 | XXyac-YX65C7_A.2 |
| 6 | 169237802 | 169238003 | 2.72660924 | THBS2 |
| 6 | 169237802 | 169238003 | 2.72660924 | XXyac-YX65C7_A.2 |
| 6 | 169559081 | 169559276 | -1.362436935 | WDR27 |
| 6 | 169624822 | 169625020 | -1.105159795 | WDR27 |
| 7 | 26164642 | 26164833 | -1.452738325 | NFE2L3 |
| 7 | 26165713 | 26165881 | -1.759865047 | NFE2L3 |
| 7 | 26185764 | 26185986 | 2.318026942 | NFE2L3 |
| 7 | 27128839 | 27129025 | -2.24539361 | RP1-170O19.22 |
| 7 | 27128839 | 27129025 | -2.24539361 | HOXA4 |
| 7 | 27128839 | 27129025 | -2.24539361 | HOXA-AS2 |
| 7 | 27128839 | 27129025 | -2.24539361 | HOXA3 |
| 1 | 226929032 | 226929249 | -1.880171706 | COQ8A |
| 7 | 27189694 | 27189943 | -1.523361933 | RP1-170O19.14 |
| 1 | 226929835 | 226929974 | -1.984907146 | COQ8A |
| 7 | 27194867 | 27195064 | -1.042864937 | HOXA13 |
| 7 | 27198028 | 27198214 | -1.355206621 | HOXA13 |
| 7 | 27204924 | 27205127 | -6.469054663 | HOTTIP |
| 7 | 27540233 | 27540452 | -1.294438775 | HIBADH |
| 7 | 30452474 | 30452674 | 1.102043009 | NOD1 |
| 7 | 30534298 | 30534503 | -2.398591303 | AC005154.6 |
| 7 | 30534298 | 30534503 | -2.398591303 | RP4-777O23.3 |
| 7 | 30535648 | 30535845 | 2.132014284 | RP4-777O23.3 |
| 7 | 30535648 | 30535845 | 2.132014284 | AC005154.6 |
| 7 | 30554474 | 30554592 | -7.386878686 | AC005154.6 |
| 7 | 30633714 | 30633914 | -1.670186695 | GARS |
| 7 | 32689040 | 32689159 | 4.906890596 | DPY19L1P1 |
| 7 | 32704030 | 32704155 | -5.218355771 | DPY19L1P1 |
| 7 | 32826190 | 32826389 | -1.583020783 | DPY19L1P2 |
| 7 | 35202323 | 35202531 | 3.201612555 | TBX20 |
| 7 | 35673255 | 35673454 | 1.063376516 | HERPUD2 |
| 7 | 36154762 | 36154963 | -1.102029207 | EEPD1 |
| 1 | 10653750 | 10653963 | 2.465781797 | RP4-734G22.3 |
| 1 | 10653750 | 10653963 | 2.465781797 | CASZ1 |
| 1 | 227286247 | 227286421 | -5.602285572 | CDC42BPA |
| 7 | 36299285 | 36299492 | -1.060746673 | EEPD1 |
| 7 | 37950297 | 37950506 | -2.431524211 | EPDR1 |
| 7 | 37950297 | 37950506 | -2.431524211 | SFRP4 |
| 7 | 38203671 | 38203871 | -5.938642067 | STARD3NL |
| 7 | 38329391 | 38329591 | -2.882604987 | RP11-121A8.1 |
| 7 | 39631906 | 39632101 | 8.450622016 | RALA |
| 7 | 39950229 | 39950433 | 8.31527577 | CDK13 |
| 7 | 40545342 | 40545556 | 2.807157536 | AC004988.1 |
| 7 | 40545342 | 40545556 | 2.807157536 | SUGCT |
| 7 | 41690207 | 41690410 | 3.493656502 | INHBA |
| 7 | 43609179 | 43609382 | -6.718746005 | COA1 |
| 7 | 43609179 | 43609382 | -6.718746005 | STK17A |
| 7 | 43627871 | 43628048 | -1.472182651 | COA1 |
| 7 | 43628755 | 43628954 | -2.152860402 | COA1 |
| 7 | 43944473 | 43944642 | -1.770902459 | UBE2D4 |
| 7 | 43944473 | 43944642 | -1.770902459 | POLR2J4 |
| 7 | 43948251 | 43948431 | -6.218355771 | UBE2D4 |
| 7 | 43948251 | 43948431 | -6.218355771 | POLR2J4 |
| 1 | 227654325 | 227654526 | -1.155244779 | ZNF678 |
| 7 | 44060919 | 44061117 | 2.830057387 | DBNL |
| 7 | 44382425 | 44382659 | -1.198356926 | NUDCD3 |
| 7 | 44769614 | 44769817 | 1.78458318 | ZMIZ2 |
| 1 | 227669098 | 227669338 | -1.790812491 | ZNF678 |
| 7 | 44973375 | 44973572 | 3.792660134 | MYO1G |
| 10 | 125000992 | 125001118 | 5.243417511 | CTBP2 |
| 10 | 125000992 | 125001118 | 5.243417511 | RP11-59C5.3 |
| 10 | 125842483 | 125842651 | 2.127607213 | BCCIP |
| 10 | 125842483 | 125842651 | 2.127607213 | DHX32 |
| 10 | 125880825 | 125881012 | -1.711156082 | DHX32 |
| 10 | 127935744 | 127935866 | 2.018329235 | PTPRE |
| 10 | 127935744 | 127935866 | 2.018329235 | RP11-4C20.4 |
| 10 | 129837744 | 129837867 | -4.584962501 | RP11-234G16.4 |
| 10 | 129837744 | 129837867 | -4.584962501 | EBF3 |
| 10 | 131911674 | 131911875 | -7.80453164 | PPP2R2D |
| 10 | 131970407 | 131970577 | -2.254123767 | BNIP3 |
| 10 | 131996776 | 131997043 | -1.265257358 | RP11-45A17.3 |
| 10 | 132192389 | 132192549 | -1.105448391 | DPYSL4 |
| 10 | 132348083 | 132348284 | -1.313305405 | LRRC27 |
| 10 | 132405524 | 132405698 | 2.231065479 | PWWP2B |
| 10 | 132416931 | 132417124 | 1.525886825 | PWWP2B |
| 10 | 132439046 | 132439235 | -6.765995289 | RP11-432J24.3 |
| 10 | 132522435 | 132522554 | 2.60715956 | LINC01165 |
| 10 | 132771555 | 132771765 | 6.458033025 | INPP5A |
| 1 | 1088576 | 1088766 | -6.821928095 | C1orf159 |
| 10 | 132772322 | 132772520 | 4.079935668 | INPP5A |
| 10 | 133420064 | 133420251 | 1.590023065 | MTG1 |
| 10 | 133420064 | 133420251 | 1.590023065 | RP11-108K14.8 |
| 10 | 133476833 | 133476989 | -4.303665157 | SCART1 |
| 1 | 22913692 | 22913794 | -1.520738318 | EPHB2 |
| 11 | 459272 | 459497 | -2.97320948 | PTDSS2 |
| 11 | 529039 | 529242 | 1.808901995 | RP13-46H24.1 |
| 11 | 532370 | 532576 | 1.25617924 | HRAS |
| 11 | 532939 | 533150 | 1.331494517 | HRAS |
| 11 | 536254 | 536396 | 1.858852682 | HRAS |
| 11 | 538578 | 538702 | 2.678277219 | LRRC56 |
| 11 | 546576 | 546760 | 2.274660599 | LRRC56 |
| 11 | 562270 | 562441 | 2.02046928 | RASSF7 |
| 11 | 566293 | 566488 | 3.453445298 | MIR210HG |
| 11 | 567272 | 567472 | 1.126529288 | MIR210HG |
| 11 | 798897 | 799107 | 1.088606324 | PANO1 |
| 11 | 805422 | 805616 | 2.242400261 | PIDD1 |
| 11 | 869094 | 869300 | 1.360777137 | CHID1 |
| 11 | 1010728 | 1010849 | 1.011183907 | AP2A2 |
| 11 | 1453421 | 1453607 | -6.281121212 | BRSK2 |
| 11 | 2313029 | 2313221 | 1.19487174 | TSPAN32 |
| 11 | 2317620 | 2317940 | 1.064883937 | TSPAN32 |
| 11 | 2328882 | 2329076 | -1.867875639 | CD81-AS1 |
| 11 | 2403020 | 2403214 | 1.458130595 | TSSC4 |
| 11 | 2698468 | 2698632 | -5.237866715 | KCNQ1 |
| 11 | 2698468 | 2698632 | -5.237866715 | KCNQ1OT1 |
| 11 | 3133668 | 3133830 | 5.294357318 | OSBPL5 |
| 11 | 3855226 | 3855478 | 3.458126815 | STIM1 |
| 11 | 3855226 | 3855478 | 3.458126815 | AC090587.5 |
| 11 | 3855226 | 3855478 | 3.458126815 | AC090587.4 |
| 11 | 3856113 | 3856278 | 1.798573312 | STIM1 |
| 1 | 1090210 | 1090350 | -2.257281793 | C1orf159 |
| 11 | 5714902 | 5715103 | -3.008034696 | TRIM22 |
| 11 | 5714902 | 5715103 | -3.008034696 | TRIM5 |

| | | | | |
|---|---|---|---|---|
| 6 | 169625744 | 169625868 | -1.84844968 | WDR27 |
| 6 | 169626582 | 169626787 | -2.183097546 | WDR27 |
| 6 | 169655051 | 169655179 | -1.954885789 | WDR27 |
| 6 | 169657127 | 169657360 | -1.532261565 | WDR27 |
| 6 | 169665240 | 169665421 | 1.162221846 | WDR27 |
| 6 | 169694796 | 169694992 | -1.393635262 | WDR27 |
| 6 | 169703003 | 169703223 | -1.29352111 | C6orf120 |
| 6 | 169725129 | 169725325 | -1.915721565 | RP1-266L20.2 |
| 6 | 169725129 | 169725325 | -1.915721565 | PHF10 |
| 1 | 224389646 | 224389846 | 1.023406785 | WDR26 |
| 6 | 170162399 | 170162599 | -2.256564907 | RP11-302L19.1 |
| 6 | 170271383 | 170271509 | -2.703366879 | RP5-894D12.5 |
| 6 | 170272937 | 170273060 | -5.402756788 | LINC01624 |
| 6 | 170272937 | 170273060 | -5.402756788 | RP5-894D12.5 |
| 6 | 170276514 | 170276695 | -2.810671468 | LINC01624 |
| 6 | 170276514 | 170276695 | -2.810671468 | RP5-894D12.5 |
| 1 | 224616473 | 224616702 | 3.668332974 | CNIH3 |
| 6 | 170304718 | 170304913 | -1.472140742 | DLL1 |
| 6 | 170304718 | 170304913 | -1.472140742 | FAM120B |
| 6 | 170581880 | 170582071 | -8.866063834 | PDCD2 |
| 7 | 80463 | 80669 | 4.508676013 | AC093627.8 |
| 7 | 80463 | 80669 | 4.508676013 | AC093627.7 |
| 7 | 82640 | 82818 | 5.449300694 | AC093627.7 |
| 7 | 145460 | 145654 | 8.128546572 | AC093627.9 |
| 7 | 153934 | 154100 | -1.454521227 | AC093627.10 |
| 7 | 163339 | 163533 | 2.589560893 | RP11-90P13.1 |
| 7 | 163830 | 164025 | -1.472949488 | RP11-90P13.1 |
| 7 | 196089 | 196376 | 4.415766583 | FAM20C |
| 7 | 197197 | 197481 | 2.631716968 | FAM20C |
| 7 | 198195 | 198396 | 3.963067264 | FAM20C |
| 7 | 199827 | 200045 | 5.028676657 | FAM20C |
| 7 | 203645 | 203843 | -6.796403695 | FAM20C |
| 7 | 214884 | 215095 | 2.427439899 | FAM20C |
| 7 | 218660 | 218944 | 1.388898135 | FAM20C |
| 1 | 224938043 | 224938194 | 5.475642357 | DNAH14 |
| 7 | 242891 | 243101 | 2.734764241 | FAM20C |
| 7 | 244990 | 245192 | 4.773268621 | FAM20C |
| 7 | 259903 | 260089 | 3.610293976 | FAM20C |
| 7 | 498295 | 498497 | 5.269310099 | PDGFA |
| 7 | 550199 | 550361 | -1.434680029 | PRKAR1B |
| 7 | 605167 | 605348 | -2.449487694 | PRKAR1B |
| 7 | 605167 | 605348 | -2.449487694 | AC147651.4 |
| 7 | 611028 | 611221 | 1.181067209 | PRKAR1B |
| 7 | 764613 | 764864 | -1.849106949 | DNAAF5 |
| 7 | 780307 | 780486 | -4.789091984 | DNAAF5 |
| 7 | 782871 | 783061 | -1.658866767 | DNAAF5 |
| 7 | 785664 | 785864 | -1.199550671 | DNAAF5 |
| 7 | 881797 | 882039 | -3.60171224 | SUN1 |
| 7 | 881797 | 882039 | -3.60171224 | GET4 |
| 7 | 881797 | 882039 | -3.60171224 | RP11-449P15.2 |
| 7 | 884839 | 884959 | -4.635614782 | SUN1 |
| 7 | 884839 | 884959 | -4.635614782 | GET4 |
| 7 | 884839 | 884959 | -4.635614782 | RP11-449P15.2 |
| 7 | 887940 | 888117 | -8.005231272 | GET4 |
| 7 | 45714071 | 45714269 | -1.462141389 | ADCY1 |
| 7 | 45714456 | 45714571 | -1.582470895 | ADCY1 |
| 7 | 45717038 | 45717237 | -1.234595872 | ADCY1 |
| 7 | 47275425 | 47275643 | -1.403480923 | TNS3 |
| 7 | 47303249 | 47303392 | 7.956958931 | TNS3 |
| 7 | 50592704 | 50592910 | -3.148484773 | GRB10 |
| 1 | 227701827 | 227702030 | -7.418813967 | ZNF847P |
| 7 | 53762895 | 53763082 | -1.064641508 | LINC01446 |
| 7 | 55431211 | 55431428 | 1.06332659 | LANCL2 |
| 7 | 55543949 | 55544149 | 2.428933575 | VOPP1 |
| 7 | 64334124 | 64334325 | -2.912575096 | ZNF736 |
| 7 | 64348745 | 64348889 | -1.986782513 | ZNF736 |
| 7 | 64500803 | 64500909 | -1.596739865 | HNRNPCP7 |
| 7 | 64521158 | 64521318 | -1.388220915 | ZNF680 |
| 7 | 64709224 | 64709435 | 1.957349403 | ZNF107 |
| 7 | 64832938 | 64833076 | -2.137161273 | ZNF138 |
| 7 | 64889982 | 64890144 | -1.080232336 | ZNF273 |
| 7 | 64889982 | 64890144 | -1.080232336 | RP11-797H7.5 |
| 7 | 64942400 | 64942601 | -3.519347844 | RP11-460N20.7 |
| 7 | 64976975 | 64977176 | -1.270802482 | ZNF117 |
| 1 | 10948088 | 10948287 | -3.604218799 | C1orf127 |
| 7 | 64979139 | 64979337 | -1.355048192 | ZNF117 |
| 7 | 64993932 | 64994083 | -1.478607567 | ZNF117 |
| 7 | 64993932 | 64994083 | -1.478607567 | ERV3-1 |
| 7 | 65200603 | 65200794 | -6.53890856 | INTS4P1 |
| 7 | 65214850 | 65215050 | -5.855833487 | INTS4P1 |
| 7 | 65398604 | 65398810 | -1.470137338 | ZNF92 |
| 7 | 65720768 | 65720950 | -1.824851646 | RP11-479O9.4 |
| 7 | 65954176 | 65954420 | -1.220348067 | VKORC1L1 |
| 7 | 65957379 | 65957561 | -3.233993514 | VKORC1L1 |
| 7 | 65958962 | 65959160 | -2.195276848 | VKORC1L1 |
| 7 | 66240436 | 66240701 | -2.006372861 | TPST1 |
| 7 | 66359307 | 66359480 | -3.146087761 | TPST1 |
| 7 | 66379775 | 66379986 | -2.83505023 | TPST1 |
| 7 | 66379775 | 66379986 | -2.83505023 | LINC00174 |
| 7 | 66682427 | 66682593 | 5.862256927 | KCTD7 |
| 7 | 66682427 | 66682593 | 5.862256927 | RABGEF1 |
| 7 | 72575329 | 72575579 | -1.498612808 | TYW1B |
| 1 | 228104384 | 228104595 | -1.139588304 | C1orf35 |
| 7 | 73442255 | 73442454 | -1.270618171 | BAZ1B |
| 7 | 73570464 | 73570645 | -1.506773077 | TBL2 |
| 7 | 73682750 | 73682874 | -1.601046827 | DNAJC30 |
| 7 | 73736187 | 73736431 | -1.903988022 | ABHD11 |
| 7 | 74376174 | 74376352 | -1.347631124 | CLIP2 |
| 7 | 74386351 | 74386573 | -1.251948415 | CLIP2 |
| 7 | 74403841 | 74404056 | -1.409098717 | CLIP2 |
| 1 | 228175246 | 228175451 | 1.089751989 | IBA57 |
| 7 | 75415778 | 75415971 | -1.315310181 | NSUN5P1 |
| 7 | 75501100 | 75501218 | -2.093955373 | SPDYE5 |
| 7 | 75536538 | 75536794 | -1.735707169 | HIP1 |
| 7 | 76389440 | 76389630 | -2.260215802 | SSC4D |
| 1 | 228212760 | 228212948 | -3.696158711 | OBSCN |
| 1 | 228212760 | 228212948 | -3.696158711 | C1orf145 |
| 7 | 76592335 | 76592552 | -1.41595705 | AC004980.7 |
| 11 | 5715752 | 5715934 | -6.515850601 | TRIM5 |
| 11 | 5715752 | 5715934 | -6.515850601 | TRIM22 |
| 11 | 6390458 | 6390661 | 9.783777537 | SMPD1 |
| 11 | 6391713 | 6391907 | 1.163296398 | SMPD1 |
| 11 | 6557275 | 6557475 | -1.127854122 | DNHD1 |
| 11 | 6567119 | 6567228 | -7.47891378 | DNHD1 |
| 11 | 7653344 | 7653590 | 1.105228909 | PPFIBP2 |
| 11 | 8000243 | 8000406 | 2.314408627 | EIF3F |
| 11 | 8730376 | 8730575 | -2.028700072 | ST5 |
| 11 | 9139538 | 9139749 | -1.609908957 | DENND5A |
| 11 | 9280746 | 9280944 | -7.60360251 | TMEM41B |
| 11 | 10170152 | 10170269 | 4.874096425 | SBF2 |
| 11 | 10778795 | 10778991 | 1.077230942 | CTR9 |
| 1 | 23845560 | 23845758 | -1.32760585 | FUCA1 |
| 11 | 14606153 | 14606337 | 1.674682639 | PSMA1 |
| 11 | 14643900 | 14644095 | 8.408766663 | PDE3B |
| 11 | 14666321 | 14666520 | -1.021762644 | PDE3B |
| 11 | 14717964 | 14718158 | -1.024737346 | PDE3B |
| 11 | 14779861 | 14780010 | -7.519373778 | PDE3B |
| 11 | 17168888 | 17169105 | 1.015554821 | PIK3C2A |
| 11 | 17353070 | 17353264 | 3.393298181 | NCR3LG1 |
| 11 | 17353678 | 17353884 | 5.041889942 | NCR3LG1 |
| 11 | 17381147 | 17381362 | 2.231061055 | RP1-239B22.5 |
| 11 | 17386694 | 17386879 | 3.808004102 | KCNJ11 |
| 11 | 17387963 | 17388142 | 1.693394293 | KCNJ11 |
| 1 | 24902097 | 24902296 | 11.0332312 | RUNX3 |
| 11 | 22224571 | 22224727 | -6.372416919 | ANO5 |
| 11 | 22280056 | 22280191 | -4.75259564 | ANO5 |
| 11 | 27658359 | 27658531 | 3.433000519 | BDNF-AS |
| 11 | 27658359 | 27658531 | 3.433000519 | BDNF |
| 11 | 28159004 | 28159204 | -2.12520581 | METTL15 |
| 1 | 25253665 | 25253774 | -6.526284025 | RSRP1 |
| 11 | 32441056 | 32441233 | -1.722521784 | WT1-AS |
| 11 | 33237091 | 33237240 | -4.990069789 | RP1-89D4.1 |
| 11 | 33353148 | 33353334 | -1.196139478 | HIPK3 |
| 11 | 33710148 | 33710346 | 1.986032408 | CD59 |
| 11 | 34159358 | 34159534 | -6.488103045 | ABTB2 |
| 11 | 35137798 | 35137952 | 3.953445298 | RP4-607I7.1 |
| 11 | 35229128 | 35229326 | 1.410599055 | CD44 |
| 11 | 43325521 | 43325707 | -6.324178791 | API5 |
| 11 | 44244335 | 44244481 | 1.123464852 | EXT2 |
| 11 | 45804454 | 45804647 | 3.822829216 | SLC35C1 |
| 11 | 46238413 | 46238575 | 1.143626507 | CTD-2589M5.5 |
| 11 | 46379990 | 46380195 | 1.51547825 | DGKZ |
| 11 | 46603819 | 46603982 | 1.076237837 | HARBI1 |
| 11 | 46616086 | 46616262 | -1.555999403 | HARBI1 |
| 11 | 47268624 | 47268806 | -1.327270126 | NR1H3 |
| 11 | 47354987 | 47355195 | 10.74829891 | SPI1 |
| 11 | 47355361 | 47355569 | 10.23290077 | SPI1 |
| 1 | 25617636 | 25617834 | 2.956745503 | MAN1C1 |
| 1 | 25783782 | 25783980 | 1.03944907 | MAN1C1 |
| 1 | 1093104 | 1093319 | 1.11251054 | C1orf159 |
| 11 | 61744013 | 61744219 | 1.116167429 | DAGLA |
| 11 | 61786454 | 61786761 | -1.074606255 | TMEM258 |

| | | | | |
|---|---|---|---|---|
| 7 | 887940 | 888117 | -8.005231272 | SUN1 |
| 7 | 888682 | 888871 | -8.449825973 | GET4 |
| 7 | 888682 | 888871 | -8.449825973 | SUN1 |
| 7 | 893580 | 893761 | -8.464906592 | SUN1 |
| 7 | 893580 | 893761 | -8.464906592 | GET4 |
| 7 | 895059 | 895246 | -3.694276027 | GET4 |
| 7 | 895059 | 895246 | -3.694276027 | SUN1 |
| 7 | 898780 | 898979 | -2.58539041 | COX19 |
| 7 | 898780 | 898979 | -2.58539041 | ADAP1 |
| 7 | 907541 | 907740 | -9.278713153 | ADAP1 |
| 7 | 907541 | 907740 | -9.278713153 | COX19 |
| 7 | 932345 | 932554 | -3.443759282 | ADAP1 |
| 7 | 932345 | 932554 | -3.443759282 | COX19 |
| 1 | 224985350 | 224985502 | -1.021663716 | DNAH14 |
| 7 | 1026297 | 1026476 | -11.29870213 | C7orf50 |
| 7 | 1026648 | 1026848 | -2.980393335 | C7orf50 |
| 7 | 1045849 | 1046051 | -4.246434308 | C7orf50 |
| 7 | 1045849 | 1046051 | -4.246434308 | GPR146 |
| 7 | 1047942 | 1048131 | -9.345459369 | GPR146 |
| 7 | 1047942 | 1048131 | -9.345459369 | C7orf50 |
| 7 | 1050865 | 1051051 | -8.074814027 | C7orf50 |
| 7 | 1050865 | 1051051 | -8.074814027 | GPR146 |
| 7 | 1051451 | 1051623 | -5.924507037 | GPR146 |
| 7 | 1051451 | 1051623 | -5.924507037 | C7orf50 |
| 7 | 1054846 | 1055046 | -4.72766361 | C7orf50 |
| 7 | 1054846 | 1055046 | -4.72766361 | GPR146 |
| 1 | 225497567 | 225497767 | 1.636706829 | ENAH |
| 7 | 1058339 | 1058618 | -4.260863156 | RP11-449P15.1 |
| 7 | 1058339 | 1058618 | -4.260863156 | GPR146 |
| 7 | 1058339 | 1058618 | -4.260863156 | C7orf50 |
| 7 | 1164452 | 1164595 | -1.284211543 | AC091729.9 |
| 7 | 1587772 | 1587972 | 1.143225329 | PSMG3-AS1 |
| 7 | 1747043 | 1747249 | 5.234596695 | ELFN1 |
| 7 | 1841797 | 1841985 | -3.53664271 | AC110781.3 |
| 7 | 1841797 | 1841985 | -3.53664271 | MAD1L1 |
| 7 | 1842326 | 1842509 | -7.313424882 | AC110781.3 |
| 7 | 1842326 | 1842509 | -7.313424882 | MAD1L1 |
| 7 | 1842572 | 1842765 | -3.099836172 | MAD1L1 |
| 7 | 1842572 | 1842765 | -3.099836172 | AC110781.3 |
| 7 | 1845297 | 1845484 | -7.01628118 | AC110781.3 |
| 7 | 1845297 | 1845484 | -7.01628118 | MAD1L1 |
| 7 | 1968309 | 1968508 | -8.907979309 | MAD1L1 |
| 7 | 2110051 | 2110266 | -1.142872618 | MAD1L1 |
| 7 | 2132566 | 2132768 | -1.407317194 | MAD1L1 |
| 1 | 225846738 | 225846938 | 1.618889079 | TMEM63A |
| 7 | 2133559 | 2133767 | -1.007916788 | MAD1L1 |
| 7 | 2234796 | 2234995 | -2.289434414 | MRM2 |
| 7 | 2254621 | 2254817 | -1.556318832 | SNX8 |
| 7 | 2432701 | 2432907 | -8.941951068 | CHST12 |
| 7 | 2433642 | 2433844 | -1.406162771 | CHST12 |
| 7 | 2434844 | 2435007 | -5.962019624 | CHST12 |
| 7 | 2443508 | 2443694 | -1.981886812 | CHST12 |
| 1 | 225985337 | 225985584 | 1.341644595 | SDE2 |
| 7 | 2446138 | 2446351 | -2.604887513 | CHST12 |
| 7 | 76595429 | 76595643 | -1.011920933 | AC004980.7 |
| 7 | 76626265 | 76626431 | -1.182040025 | AC004980.7 |
| 7 | 76626265 | 76626431 | -1.182040025 | POMZP3 |
| 1 | 228274489 | 228274698 | -7.741466986 | OBSCN |
| 1 | 228274489 | 228274698 | -7.741466986 | RP5-1139B12.3 |
| 7 | 77696090 | 77696269 | 3.323215067 | APTR |
| 7 | 77944763 | 77945014 | -1.958075467 | PHTF2 |
| 7 | 78018898 | 78019099 | -1.51410294 | MAGI2 |
| 7 | 80744967 | 80745189 | 1.075277257 | SEMA3C |
| 7 | 82914852 | 82914965 | -1.360100196 | PCLO |
| 7 | 82915347 | 82915458 | -2.450601225 | PCLO |
| 7 | 82916227 | 82916440 | -4.025582539 | PCLO |
| 7 | 82950172 | 82950357 | -3.181384634 | PCLO |
| 7 | 82953322 | 82953520 | -2.291676848 | PCLO |
| 7 | 82954153 | 82954330 | -2.057468674 | PCLO |
| 7 | 83134469 | 83134755 | -1.60496968 | PCLO |
| 7 | 83135209 | 83135417 | -1.558687939 | PCLO |
| 7 | 83155139 | 83155347 | -1.888119325 | PCLO |
| 7 | 86880292 | 86880471 | -2.078482406 | KIAA1324L |
| 7 | 87907462 | 87907667 | -1.493718585 | DBF4 |
| 7 | 88007799 | 88007998 | 2.111020536 | ADAM22 |
| 7 | 90358718 | 90358898 | -1.305248796 | GTPBP10 |
| 7 | 90412674 | 90412900 | -1.019459495 | CLDN12 |
| 7 | 90412674 | 90412900 | -1.019459495 | CTB-13L3.1 |
| 7 | 91175499 | 91175695 | -2.356013846 | CDK14 |
| 7 | 91178882 | 91179076 | -2 | CDK14 |
| 7 | 91265751 | 91265975 | 1.137508735 | FZD1 |
| 7 | 91266673 | 91266860 | 1.165090384 | FZD1 |
| 7 | 92001028 | 92001241 | -1.411351833 | AKAP9 |
| 7 | 92079565 | 92079758 | -1.279833648 | AKAP9 |
| 7 | 92083225 | 92083375 | -1.743707444 | AKAP9 |
| 7 | 92096913 | 92097114 | -1.175817414 | AKAP9 |
| 7 | 92517483 | 92517690 | -4.625780654 | PEX1 |
| 7 | 92833614 | 92833812 | 7.532181928 | CDK6 |
| 7 | 93100451 | 93100634 | 1.63996806 | SAMD9 |
| 7 | 93104996 | 93105198 | 1.292222383 | SAMD9 |
| 7 | 93105826 | 93106026 | 1.301307779 | SAMD9 |
| 7 | 93132585 | 93132790 | 2.48022414 | SAMD9L |
| 7 | 93133320 | 93133474 | 2.163484276 | SAMD9L |
| 7 | 93133979 | 93134085 | 2.355332615 | SAMD9L |
| 7 | 93135659 | 93135835 | 2.191375775 | SAMD9L |
| 7 | 96664264 | 96664456 | -5.526284025 | SHFM1 |
| 7 | 97938440 | 97938632 | -1.324876248 | AC079781.8 |
| 7 | 98194051 | 98194262 | 1.393337394 | LMTK2 |
| 7 | 98207243 | 98207414 | 7.052790591 | LMTK2 |
| 1 | 228687438 | 228687633 | 6.5915075 | FTH1P2 |
| 7 | 98302485 | 98302746 | -2.6120753 | BAIAP2L1 |
| 7 | 98302485 | 98302746 | -2.6120753 | BRI3 |
| 7 | 98307390 | 98307620 | -2.46430255 | BAIAP2L1 |
| 7 | 98307390 | 98307620 | -2.46430255 | BRI3 |
| 7 | 98309554 | 98309751 | -4.320224794 | BAIAP2L1 |
| 7 | 98309554 | 98309751 | -4.320224794 | BRI3 |
| 7 | 99028683 | 99028883 | -1.599302258 | SMURF1 |
| 7 | 99028683 | 99028883 | -1.599302258 | AC004893.11 |
| 11 | 61786454 | 61786761 | -1.074606255 | MYRF |
| 11 | 61898107 | 61898307 | -1.354736006 | RAB3IL1 |
| 11 | 61962573 | 61962790 | -1.14863772 | FTH1 |
| 11 | 61962573 | 61962790 | -1.14863772 | BEST1 |
| 1 | 25835197 | 25835404 | -1.190061063 | AUNIP |
| 1 | 25862843 | 25862956 | -4.281121212 | PAQR7 |
| 1 | 25862843 | 25862956 | -4.281121212 | RP1-125I3.2 |
| 11 | 62602205 | 62602489 | 1.95048346 | EML3 |
| 11 | 62672055 | 62672285 | 1.074554845 | UQCC3 |
| 11 | 62672055 | 62672285 | 1.074554845 | LBHD1 |
| 11 | 62786416 | 62786606 | 2.458819918 | TMEM223 |
| 11 | 62786416 | 62786606 | 2.458819918 | TAF6L |
| 11 | 62786416 | 62786606 | 2.458819918 | RP11-727F15.12 |
| 11 | 62787223 | 62787407 | 3.376616992 | TMEM223 |
| 11 | 62787223 | 62787407 | 3.376616992 | TAF6L |
| 11 | 63719306 | 63719494 | 6.789127371 | RTN3 |
| 11 | 64118260 | 64118460 | 1.160770756 | MACROD1 |
| 11 | 64118260 | 64118460 | 1.160770756 | FLRT1 |
| 11 | 64118260 | 64118460 | 1.160770756 | RP11-21A7A.3 |
| 11 | 64765735 | 64765956 | 1.010319206 | SF1 |
| 11 | 64854212 | 64854379 | 1.557358365 | EHD1 |
| 11 | 65116584 | 65116704 | 1.466385997 | ZNHIT2 |
| 11 | 65242162 | 65242349 | -6.344125155 | SLC22A20 |
| 11 | 65321329 | 65321575 | 2.410828292 | CDC42EP2 |
| 11 | 65424373 | 65424612 | 11.82775466 | NEAT1 |
| 11 | 65499486 | 65499627 | -13.67087184 | MALAT1 |
| 11 | 65618432 | 65618607 | 1.385023704 | PCNX3 |
| 11 | 65787204 | 65787411 | 2.120231534 | OVOL1 |
| 11 | 65891381 | 65891593 | 1.685489664 | CCDC85B |
| 11 | 65899626 | 65899794 | 5.36310908 | FOSL1 |
| 11 | 66002120 | 66002304 | 7.089332426 | EIF1AD |
| 11 | 66002120 | 66002304 | 7.089332426 | BANF1 |
| 11 | 66677468 | 66677663 | 1.151176748 | RBM4B |
| 11 | 66686168 | 66686273 | 1.236800696 | SPTBN2 |
| 11 | 66700706 | 66700887 | 6.279210357 | SPTBN2 |
| 1 | 26692273 | 26692461 | 1.43301025 | RP5-968P14.2 |
| 11 | 66848962 | 66849161 | -8.056993142 | PC |
| 11 | 66857890 | 66858084 | 1.146398312 | PC |
| 11 | 66857890 | 66858084 | 1.146398312 | LRFN4 |
| 11 | 66858686 | 66858881 | 1.924581699 | PC |
| 11 | 66858686 | 66858881 | 1.924581699 | LRFN4 |
| 11 | 66859520 | 66859725 | 1.266872741 | PC |
| 11 | 66859520 | 66859725 | 1.266872741 | LRFN4 |
| 11 | 66860011 | 66860210 | 9.710041789 | PC |
| 11 | 66860011 | 66860210 | 9.710041789 | LRFN4 |
| 11 | 67290612 | 67290792 | -2.120552651 | ANKRD13D |
| 11 | 67333036 | 67333201 | -4.099836172 | RAD9A |
| 11 | 67365189 | 67365345 | 2.430229957 | AP003419.11 |
| 11 | 67365189 | 67365345 | 2.430229957 | RAD9A |
| 11 | 67365189 | 67365345 | 2.430229957 | CLCF1 |
| 11 | 67365415 | 67365539 | 2.009038736 | AP003419.11 |
| 11 | 67365415 | 67365539 | 2.009038736 | CLCF1 |
| 11 | 67365415 | 67365539 | 2.009038736 | RAD9A |
| 11 | 67420483 | 67420677 | 1.691236171 | PPP1CA |

| | | | | |
|---|---|---|---|---|
| 7 | 2446981 | 2447155 | -1.799914405 | CHST12 |
| 7 | 2538017 | 2538216 | -1.321497558 | BRAT1 |
| 7 | 2560139 | 2560302 | -1.26477469 | IQCE |
| 7 | 2613697 | 2613906 | 1.080026442 | IQCE |
| 7 | 2718379 | 2718555 | 2.59216219 | AMZ1 |
| 7 | 2721957 | 2722153 | 3.876727773 | AMZ1 |
| 1 | 225988000 | 225988200 | 1.145607931 | SDE2 |
| 7 | 2723890 | 2723999 | 1.940601976 | AMZ1 |
| 7 | 2729347 | 2729545 | 1.139411623 | AMZ1 |
| 7 | 2729347 | 2729545 | 1.139411623 | GNA12 |
| 7 | 2731262 | 2731463 | 1.158589352 | GNA12 |
| 7 | 2731262 | 2731463 | 1.158589352 | AMZ1 |
| 7 | 3991034 | 3991234 | -1.479715358 | SDK1 |
| 7 | 4267833 | 4267999 | -2.335185842 | SDK1 |
| 7 | 4797555 | 4797765 | 1.85328291 | RADIL |
| 7 | 4877676 | 4877884 | -1.22944241 | RADIL |
| 7 | 5010018 | 5010124 | 3.72537652 | RNF216P1 |
| 7 | 5010018 | 5010124 | 3.72537652 | RBAK-RBAKDN |
| 7 | 5064030 | 5064229 | -1.391485382 | RBAK-RBAKDN |
| 7 | 5064030 | 5064229 | -1.391485382 | RBAK |
| 7 | 5232457 | 5232658 | -1.801515497 | WIPI2 |
| 7 | 5233208 | 5233421 | -1.034095723 | WIPI2 |
| 7 | 5313182 | 5313385 | -2.568807756 | TNRC18 |
| 7 | 5370682 | 5370931 | -1.02002015 | TNRC18 |
| 7 | 5377970 | 5378172 | -1.95579712 | TNRC18 |
| 1 | 226309759 | 226309879 | -1.937713668 | LIN9 |
| 7 | 5495495 | 5495606 | -2.999469402 | FBXL18 |
| 7 | 5500396 | 5500678 | -2.455258996 | FBXL18 |
| 7 | 5501177 | 5501373 | -1.605621774 | FBXL18 |
| 7 | 5527630 | 5527831 | -1.922102064 | ACTB |
| 7 | 5741195 | 5741308 | -2.702606695 | RNF216 |
| 7 | 5847305 | 5847431 | -2.221562856 | ZNF815P |
| 7 | 5973211 | 5973384 | -2.792353431 | PMS2 |
| 7 | 5987139 | 5987424 | -1.621567286 | PMS2 |
| 7 | 6164812 | 6165012 | 1.209770064 | CYTH3 |
| 7 | 6330975 | 6331184 | -1.381426559 | FAM220A |
| 7 | 6409558 | 6409866 | -1.84584811 | DAGLB |
| 7 | 6690505 | 6690813 | -1.062222009 | ZNF12 |
| 7 | 6690505 | 6690813 | -1.062222009 | AC073343.13 |
| 1 | 10639761 | 10639957 | 5.866507661 | RP4-734G22.3 |
| 1 | 10639761 | 10639957 | 5.866507661 | CASZ1 |
| 7 | 7720986 | 7721127 | -6.836709752 | RP11-505D17.1 |
| 7 | 7720986 | 7721127 | -6.836709752 | UMAD1 |
| 7 | 7725931 | 7726058 | -7.239890132 | RP11-505D17.1 |
| 7 | 7725931 | 7726058 | -7.239890132 | UMAD1 |
| 7 | 7855900 | 7856056 | 6.127514285 | RP11-505D17.1 |
| 7 | 7855900 | 7856056 | 6.127514285 | UMAD1 |
| 1 | 226728998 | 226729229 | 1.229859176 | ITPKB |
| 7 | 11348814 | 11348972 | 6.422745025 | AC004538.3 |
| 7 | 11351495 | 11351691 | 3.7256584 | AC004538.3 |
| 7 | 11375544 | 11375849 | -1.210421315 | AC004538.3 |
| 7 | 11375544 | 11375849 | -1.210421315 | THSD7A |
| 7 | 11636738 | 11636938 | -1.532550012 | THSD7A |
| 7 | 12219489 | 12219614 | 2.696635635 | TMEM106B |
| 7 | 99486967 | 99487151 | -1.271467155 | ZNF789 |
| 7 | 99486967 | 99487151 | -1.271467155 | ZNF394 |
| 7 | 99525942 | 99526161 | -1.024030459 | ZKSCAN5 |
| 7 | 99531667 | 99531883 | -1.293745664 | ZKSCAN5 |
| 7 | 99547159 | 99547289 | -2.166098215 | FAM200A |
| 7 | 99573054 | 99573253 | -1.808670664 | ZNF655 |
| 7 | 99573054 | 99573253 | -1.808670664 | GS1-259H13.13 |
| 1 | 228743550 | 228743747 | 2.784387202 | RHOU |
| 7 | 100033279 | 100033478 | -1.881346528 | ZKSCAN1 |
| 7 | 100064085 | 100064288 | -3.918751396 | ZSCAN21 |
| 7 | 100064085 | 100064288 | -3.918751396 | ZNF3 |
| 7 | 100071914 | 100072127 | -1.035160267 | ZNF3 |
| 7 | 100211327 | 100211528 | -2.020334315 | STAG3 |
| 7 | 100211327 | 100211528 | -2.020334315 | GATS |
| 7 | 100319303 | 100319503 | -3.779645087 | SPDYE3 |
| 7 | 100865861 | 100866025 | -1.595341781 | SLC12A9 |
| 7 | 101088358 | 101088557 | 1.294422346 | TRIM56 |
| 7 | 101091018 | 101091220 | 1.672052573 | TRIM56 |
| 7 | 102197008 | 102197210 | -1.263672943 | CUX1 |
| 7 | 102227355 | 102227521 | -7.074079386 | CUX1 |
| 1 | 229268511 | 229268710 | -2.279815314 | RP5-1061H20.4 |
| 7 | 102559861 | 102560145 | 5.271032271 | POLR2J3 |
| 7 | 102559861 | 102560145 | 5.271032271 | SPDYE2 |
| 7 | 102559861 | 102560145 | 5.271032271 | RP11-514P8.7 |
| 7 | 102592691 | 102592862 | 6.272964885 | RASA4 |
| 7 | 102592691 | 102592862 | 6.272964885 | RP11-514P8.6 |
| 7 | 102668615 | 102668803 | -4.725605556 | RP11-577H5.5 |
| 7 | 102668615 | 102668803 | -4.725605556 | POLR2J2 |
| 1 | 229269260 | 229269509 | -1.635216814 | RP5-1061H20.4 |
| 7 | 102934350 | 102934506 | -6.256863798 | FBXL13 |
| 7 | 102934350 | 102934506 | -6.256863798 | LRRC17 |
| 7 | 103109504 | 103109660 | -5.23456651 | NAPEPLD |
| 7 | 103112316 | 103112489 | -3.257079273 | NAPEPLD |
| 7 | 104950206 | 104950406 | 1.709348115 | RP11-325F22.2 |
| 7 | 104950206 | 104950406 | 1.709348115 | LINC01004 |
| 7 | 105101862 | 105102068 | -1.99541694 | KMT2E |
| 7 | 105107630 | 105107829 | -1.465571104 | KMT2E |
| 1 | 229325212 | 229325412 | -1.560454738 | CCSAP |
| 7 | 105142389 | 105142588 | -1.509293545 | SRPK2 |
| 7 | 105607853 | 105607992 | -3.736514611 | ATXN7L1 |
| 7 | 105610541 | 105610722 | -3.537695004 | ATXN7L1 |
| 7 | 107629053 | 107629263 | 2.091268981 | BCAP29 |
| 7 | 107629053 | 107629263 | 2.091268981 | WBP1LP2 |
| 7 | 107758236 | 107758434 | -1.65457416 | CBLL1 |
| 7 | 112343940 | 112344127 | 2.861951237 | AC004112.4 |
| 7 | 114689779 | 114689980 | -1.091297901 | FOXP2 |
| 7 | 114690758 | 114690958 | -1.256798962 | FOXP2 |
| 7 | 115940311 | 115940519 | 1.306418231 | TFEC |
| 7 | 120273843 | 120274041 | -3.584962501 | KCND2 |
| 7 | 120274372 | 120274491 | -3.807354922 | KCND2 |
| 7 | 128029352 | 128029551 | -7.302508224 | SND1 |
| 7 | 128029352 | 128029551 | -7.302508224 | LRRC4 |
| 7 | 128457424 | 128457621 | 1.325517209 | HILPDA |
| 7 | 128457424 | 128457621 | 1.325517209 | RP11-155G14.6 |
| 11 | 67420483 | 67420677 | 1.691236171 | CARNS1 |
| 1 | 26696907 | 26697019 | 8.821099902 | ARID1A |
| 11 | 67991356 | 67991604 | 2.227831138 | UNC93B1 |
| 11 | 68052150 | 68052347 | 5.767394606 | RP11-802E16.3 |
| 1 | 26731190 | 26731339 | -8.709156915 | ARID1A |
| 11 | 68386414 | 68386530 | 2.637199862 | LRP5 |
| 11 | 68980292 | 68980450 | -1.106212619 | MRGPRD |
| 11 | 69161933 | 69162048 | -5.72869044 | TPCN2 |
| 11 | 69161933 | 69162048 | -5.72869044 | RP11-554A11.8 |
| 11 | 69367511 | 69367701 | -2.18121232 | MYEOV |
| 11 | 71800330 | 71800529 | 1.341703952 | ALG1L9P |
| 11 | 71800330 | 71800529 | 1.341703952 | FAM86C1 |
| 11 | 72013169 | 72013365 | 1.666038336 | NUMA1 |
| 11 | 72015511 | 72015696 | -2.020946719 | NUMA1 |
| 11 | 72015511 | 72015696 | -2.020946719 | RP11-849H4.4 |
| 11 | 72293306 | 72293601 | 1.107981969 | CLPB |
| 11 | 72434167 | 72434362 | -5.769336977 | CLPB |
| 11 | 72838104 | 72838304 | 5.849569313 | FCHSD2 |
| 11 | 72838104 | 72838304 | 5.849569313 | ATG16L2 |
| 11 | 73309464 | 73309686 | 1.685282406 | ARHGEF17 |
| 11 | 73310951 | 73311072 | 1.325694931 | ARHGEF17 |
| 11 | 73353126 | 73353248 | 7.466568684 | ARHGEF17 |
| 11 | 73367613 | 73367813 | 1.181344238 | ARHGEF17 |
| 11 | 74128962 | 74129140 | 1.370733493 | C2CD3 |
| 11 | 74497759 | 74497926 | -1.582200477 | AP001372.2 |
| 11 | 74497759 | 74497926 | -1.582200477 | POLD3 |
| 11 | 74818866 | 74819064 | -4.55033117 | RNF169 |
| 11 | 74818866 | 74819064 | -4.55033117 | XRRA1 |
| 11 | 74877761 | 74877930 | 5.797628741 | XRRA1 |
| 11 | 74877761 | 74877930 | 5.797628741 | RP11-147I3.1 |
| 11 | 74878464 | 74878663 | 5.038407799 | XRRA1 |
| 11 | 74878464 | 74878663 | 5.038407799 | RP11-147I3.1 |
| 11 | 75571800 | 75572002 | 1.429809787 | SERPINH1 |
| 1 | 1232976 | 1233163 | 8.851211491 | B3GALT6 |
| 11 | 76211235 | 76211434 | -4.872416919 | RP11-619A14.3 |
| 11 | 77103530 | 77103700 | 1.144224267 | CAPN5 |
| 11 | 77734664 | 77734864 | 1.37886115 | FTH1P16 |
| 11 | 77734664 | 77734864 | 1.37886115 | RSF1 |
| 11 | 79194509 | 79194628 | -2.53846232 | TENM4 |
| 11 | 82931807 | 82931967 | 2.12740695 | PRCP |
| 11 | 82931807 | 82931967 | 2.12740695 | DDIAS |
| 11 | 83286085 | 83286210 | 6.189959909 | RP11-727A23.10 |
| 11 | 83286085 | 83286210 | 6.189959909 | CCDC90B |
| 11 | 86808389 | 86808595 | 3.229336425 | PRSS23 |
| 11 | 88294014 | 88294214 | 1.192900961 | CTSC |
| 11 | 88326250 | 88326467 | 1.215627641 | CTSC |
| 1 | 27551186 | 27551384 | 1.643116296 | AHDC1 |
| 11 | 93156881 | 93157077 | -1.69630861 | SLC36A4 |
| 11 | 93178033 | 93178230 | -6.714114649 | SLC36A4 |
| 11 | 94549325 | 94549525 | -1.260517647 | RP11-867G2.8 |
| 11 | 94549325 | 94549525 | -1.260517647 | FUT4 |
| 11 | 94549325 | 94549525 | -1.260517647 | PIWIL4 |
| 11 | 94566397 | 94566594 | -1.983940076 | RP11-867G2.8 |
| 11 | 94566397 | 94566594 | -1.983940076 | PIWIL4 |

| | | | | |
|---|---|---|---|---|
| 7 | 12370089 | 12370277 | -1.149385187 | VWDE |
| 1 | 226735797 | 226736004 | 1.876536396 | ITPKB |
| 7 | 15612356 | 15612571 | -3.38151559 | MEOX2 |
| 7 | 18188280 | 18188482 | -1.938974446 | HDAC9 |
| 1 | 226895609 | 226895919 | 2.389575369 | PSEN2 |
| 7 | 18666190 | 18666448 | 3.35403142 | HDAC9 |
| 7 | 18988033 | 18988229 | -6.530094639 | HDAC9 |
| 7 | 18988861 | 18988996 | -6.018086806 | HDAC9 |
| 7 | 19116024 | 19116224 | -2.744445353 | TWIST1 |
| 7 | 19116734 | 19116971 | -2.785114569 | TWIST1 |
| 7 | 19698428 | 19698630 | -1.502654106 | TWISTNB |
| 7 | 21429787 | 21429958 | -1.154755615 | SP4 |
| 1 | 226926988 | 226927232 | 3.638969701 | COQ8A |
| 7 | 21511043 | 21511242 | -1.573445356 | SP4 |
| 7 | 22869286 | 22869487 | -1.442778928 | AC005682.6 |
| 7 | 22882053 | 22882237 | -1.743783539 | RPL12P10 |
| 7 | 22945911 | 22946113 | -1.560426402 | FAM126A |
| 7 | 25135997 | 25136149 | -2.605959657 | C7orf31 |
| 7 | 25136341 | 25136550 | -1.63374724 | C7orf31 |
| 1 | 226928022 | 226928220 | 7.339079979 | COQ8A |
| 7 | 26163915 | 26164123 | 7.72268528 | NFE2L3 |
| 7 | 26164642 | 26164833 | 8.24298539 | NFE2L3 |
| 7 | 26165713 | 26165881 | 6.831556654 | NFE2L3 |
| 7 | 26765449 | 26765655 | -1.755183128 | SKAP2 |
| 7 | 27143549 | 27143730 | -3.055225022 | HOXA5 |
| 7 | 27143549 | 27143730 | -3.055225022 | HOXA3 |
| 7 | 27143549 | 27143730 | -3.055225022 | HOXA-AS3 |
| 7 | 27143549 | 27143730 | -3.055225022 | RP1-170O19.22 |
| 1 | 226929032 | 226929249 | 2.084519693 | COQ8A |
| 7 | 27145084 | 27145264 | -7.014988673 | HOXA3 |
| 7 | 27145084 | 27145264 | -7.014988673 | RP1-170O19.22 |
| 7 | 27145084 | 27145264 | -7.014988673 | HOXA-AS3 |
| 7 | 27150606 | 27150790 | -2.77344723 | RP1-170O19.22 |
| 7 | 27150606 | 27150790 | -2.77344723 | HOXA3 |
| 7 | 27150606 | 27150790 | -2.77344723 | HOXA-AS3 |
| 7 | 27150606 | 27150790 | -2.77344723 | RP1-170O19.23 |
| 7 | 27171486 | 27171691 | -1.194429386 | HOXA9 |
| 7 | 27171486 | 27171691 | -1.194429386 | HOXA10-AS |
| 7 | 27171486 | 27171691 | -1.194429386 | HOXA10-HOXA9 |
| 7 | 27171486 | 27171691 | -1.194429386 | HOXA10 |
| 7 | 27172011 | 27172145 | -1.248456647 | HOXA10 |
| 7 | 27172011 | 27172145 | -1.248456647 | HOXA9 |
| 7 | 27172011 | 27172145 | -1.248456647 | HOXA10-HOXA9 |
| 7 | 27176020 | 27176222 | -3.869702796 | HOXA10 |
| 7 | 27176020 | 27176222 | -3.869702796 | HOXA10-HOXA9 |
| 7 | 27182754 | 27182915 | -1.606117644 | HOXA11 |
| 7 | 27189694 | 27189943 | -2.669201372 | RP1-170O19.14 |
| 7 | 27191774 | 27191883 | -2.668648514 | RP1-170O19.14 |
| 7 | 27193300 | 27193459 | -2.478351583 | RP1-170O19.14 |
| 1 | 226929835 | 226929974 | 3.257442032 | COQ8A |
| 7 | 27194867 | 27195064 | -3.923558796 | HOXA13 |
| 7 | 27198028 | 27198214 | -2.604994902 | HOXA13 |
| 7 | 27204924 | 27205127 | -3.028652263 | HOTTIP |
| 7 | 27840972 | 27841073 | 2.773263517 | TAX1BP1 |
| 7 | 128457424 | 128457621 | 1.325517209 | RP11-212P7.3 |
| 7 | 128767362 | 128767558 | -1.202476718 | CALU |
| 7 | 128768965 | 128769167 | -1.097363101 | CALU |
| 7 | 128810120 | 128810291 | -4.087462841 | CCDC136 |
| 7 | 128867864 | 128867997 | 3.352258152 | KCP |
| 7 | 128867864 | 128867997 | 3.352258152 | RP11-309L24.4 |
| 7 | 128948755 | 128948921 | 2.291051694 | IRF5 |
| 7 | 128954975 | 128955173 | -1.002863169 | TNPO3 |
| 7 | 129485580 | 129485779 | -3.475642357 | STRIP2 |
| 7 | 129606935 | 129607259 | -3.167390253 | RP11-448A19.1 |
| 1 | 229636705 | 229636900 | 1.493089651 | URB2 |
| 7 | 129724688 | 129724792 | -1.398732086 | NRF1 |
| 7 | 129755057 | 129755230 | -6.533380966 | NRF1 |
| 7 | 130440552 | 130440746 | -5.936529774 | CEP41 |
| 7 | 130867667 | 130867866 | 4.289701488 | AC016831.7 |
| 7 | 130881583 | 130881783 | 2.564738762 | MIR29A |
| 7 | 130881583 | 130881783 | 2.564738762 | AC016831.7 |
| 7 | 131108443 | 131108601 | -1.602139478 | LINC-PINT |
| 1 | 229637939 | 229638155 | 1.041343248 | URB2 |
| 7 | 132128289 | 132128471 | -1.969611508 | PLXNA4 |
| 7 | 133034544 | 133034748 | -7.513088798 | CHCHD3 |
| 7 | 133034544 | 133034748 | -7.513088798 | RP11-371A22.1 |
| 7 | 134294697 | 134294896 | -1.649955801 | SLC35B4 |
| 7 | 134994257 | 134994454 | -1.769916563 | AGBL3 |
| 7 | 135085485 | 135085685 | -3.047294704 | AGBL3 |
| 7 | 135164506 | 135164702 | -8.069035884 | TMEM140 |
| 7 | 135164506 | 135164702 | -8.069035884 | C7orf49 |
| 7 | 135165458 | 135165658 | 3.287281658 | TMEM140 |
| 7 | 135165458 | 135165658 | 3.287281658 | C7orf49 |
| 7 | 135166385 | 135166645 | -1.948277584 | C7orf49 |
| 7 | 135167815 | 135167921 | 2.272897432 | C7orf49 |
| 7 | 135168458 | 135168623 | 2.83690886 | RP11-134L10.1 |
| 7 | 135168458 | 135168623 | 2.83690886 | C7orf49 |
| 7 | 135168850 | 135169053 | -1.356730171 | C7orf49 |
| 7 | 135168850 | 135169053 | -1.356730171 | RP11-134L10.1 |
| 7 | 135169903 | 135170103 | 1.769018754 | C7orf49 |
| 1 | 230323109 | 230323310 | -2.442536308 | PGBD5 |
| 7 | 135185900 | 135186113 | -2.177629938 | WDR91 |
| 7 | 137880406 | 137880572 | -1.170382855 | CREB3L2 |
| 7 | 138585011 | 138585202 | -1.198428922 | TRIM24 |
| 7 | 139079657 | 139079858 | 2.027849482 | ZC3HAV1 |
| 1 | 230706102 | 230706317 | 6.485951373 | AGT |
| 7 | 139678248 | 139678431 | -2.178340571 | HIPK2 |
| 7 | 140177537 | 140177688 | 2.607672808 | JHDM1D-AS1 |
| 7 | 140673885 | 140674086 | -1.542832877 | ADCK2 |
| 7 | 140673885 | 140674086 | -1.542832877 | DENND2A |
| 7 | 140695434 | 140695624 | -1.001501965 | NDUFB2 |
| 7 | 140695434 | 140695624 | -1.001501965 | ADCK2 |
| 7 | 140695434 | 140695624 | -1.001501965 | NDUFB2-AS1 |
| 7 | 140695878 | 140696056 | -1.462361944 | NDUFB2-AS1 |
| 7 | 140695878 | 140696056 | -1.462361944 | ADCK2 |
| 7 | 140695878 | 140696056 | -1.462361944 | NDUFB2 |
| 7 | 140726205 | 140726398 | -1.354852686 | BRAF |
| 7 | 143269234 | 143269470 | 1.337424147 | AC073342.12 |
| 11 | 95811131 | 95811312 | -6.036233804 | CEP57 |
| 11 | 95978883 | 95979061 | -1.152779572 | MAML2 |
| 1 | 27881763 | 27881960 | -1.050568889 | THEMIS2 |
| 11 | 102324986 | 102325168 | 3.082946624 | BIRC3 |
| 1 | 27882738 | 27882938 | -1.487284143 | THEMIS2 |
| 11 | 108104176 | 108104369 | -1.380590694 | CUL5 |
| 11 | 108206663 | 108206832 | -1.430453905 | NPAT |
| 11 | 108207003 | 108207240 | -1.020825415 | NPAT |
| 11 | 108513237 | 108513356 | -2.151059807 | EXPH5 |
| 11 | 108778478 | 108778677 | 4.732579115 | DDX10 |
| 11 | 110102293 | 110102531 | -2.843094616 | ZC3H12C |
| 11 | 110136967 | 110137135 | -1.103074279 | ZC3H12C |
| 11 | 110189224 | 110189423 | -2.416840374 | RDX |
| 11 | 110233352 | 110233503 | -1.762325826 | RDX |
| 11 | 110579514 | 110579707 | -2.446542398 | ARHGAP20 |
| 11 | 113225547 | 113225685 | -5.955321365 | NCAM1 |
| 11 | 113226165 | 113226323 | 6.025264453 | NCAM1 |
| 11 | 113227829 | 113228035 | -2.61610464 | NCAM1 |
| 11 | 116849197 | 116849394 | 1.124253853 | APOA1-AS |
| 11 | 116849197 | 116849394 | 1.124253853 | SIK3 |
| 11 | 117315812 | 117316011 | 2.809299971 | CEP164 |
| 11 | 117315812 | 117316011 | 2.809299971 | BACE1 |
| 1 | 28473495 | 28473694 | -1.19895623 | PHACTR4 |
| 11 | 117995577 | 117995774 | 1.027655242 | IL10RA |
| 11 | 117998863 | 117999058 | 1.664539668 | IL10RA |
| 11 | 118396210 | 118396403 | -1.461192751 | UBE4A |
| 11 | 118525274 | 118525430 | -1.422668884 | KMT2A |
| 11 | 118525274 | 118525430 | -1.422668884 | RP11-770J1.3 |
| 1 | 28496589 | 28496790 | -1.328887868 | PHACTR4 |
| 11 | 118908488 | 118908678 | -1.029183662 | BCL9L |
| 11 | 118996806 | 118996991 | 1.891553688 | RP11-110I1.12 |
| 11 | 119094849 | 119095039 | 10.30813262 | H2AFX |
| 11 | 119356382 | 119356498 | 10.41956421 | USP2 |
| 11 | 119372875 | 119373023 | 5.596146407 | RP11-334E6.3 |
| 11 | 119372875 | 119373023 | 5.596146407 | USP2 |
| 11 | 119664740 | 119664867 | -5.693739433 | NECTIN1 |
| 11 | 120407575 | 120407776 | 8.31120468 | ARHGEF12 |
| 11 | 122903976 | 122904175 | -1.252945465 | C11orf63 |
| 11 | 122934751 | 122934949 | -1.560066289 | C11orf63 |
| 11 | 123729712 | 123729912 | -1.893429946 | ZNF202 |
| 11 | 124746704 | 124746860 | 11.01099541 | NRGN |
| 11 | 124747325 | 124747520 | 1.618958123 | VSIG2 |
| 11 | 124748892 | 124749194 | 1.938971048 | VSIG2 |
| 11 | 125680571 | 125680716 | -4.25389732 | ACRV1 |
| 11 | 125960877 | 125961060 | -1.21770836 | CDON |
| 11 | 126277812 | 126277976 | 1.486504665 | FOXRED1 |
| 11 | 128972604 | 128972805 | -1.261790029 | ARHGAP32 |
| 11 | 129236434 | 129236628 | -2.850829363 | ARHGAP32 |
| 11 | 130108002 | 130108193 | -8.897528351 | APLP2 |
| 11 | 130108652 | 130108850 | -8.013132033 | APLP2 |
| 11 | 130405691 | 130405900 | 1.272341931 | ADAMTS8 |
| 1 | 29193345 | 29193555 | 1.141563732 | MECR |
| 11 | 130449028 | 130449222 | 1.182232708 | ADAMTS15 |
| 12 | 370862 | 371061 | -6.812626111 | KDM5A |

| | | | | |
|---|---|---|---|---|
| 7 | 27840972 | 27841073 | 2.773263517 | JAZF1 |
| 1 | 226994030 | 226994290 | 1.104204258 | CDC42BPA |
| 7 | 30452474 | 30452674 | -1.223520925 | NOD1 |
| 7 | 30535648 | 30535845 | -2.621727018 | RP4-777O23.3 |
| 7 | 30535648 | 30535845 | -2.621727018 | AC005154.6 |
| 7 | 30552526 | 30552679 | -1.268357804 | AC005154.6 |
| 7 | 30558401 | 30558605 | -1.318183998 | AC005154.6 |
| 7 | 32689040 | 32689159 | -6.502461339 | DPY19L1P1 |
| 7 | 32704030 | 32704155 | -1.356597158 | DPY19L1P1 |
| 7 | 32753728 | 32753927 | -1.04342926 | DPY19L1P1 |
| 7 | 32869437 | 32869626 | -1.066172145 | KBTBD2 |
| 7 | 35202323 | 35202531 | -2.289253739 | TBX20 |
| 1 | 227286247 | 227286421 | -1.392855066 | CDC42BPA |
| 7 | 36855453 | 36855652 | -1.095215133 | ELMO1 |
| 7 | 37950297 | 37950506 | -2.021325341 | EPDR1 |
| 7 | 37950297 | 37950506 | -2.021325341 | SFRP4 |
| 7 | 38203671 | 38203871 | -1.283994711 | STARD3NL |
| 7 | 38204383 | 38204575 | -5.238879133 | STARD3NL |
| 7 | 38327365 | 38327554 | -6.462035093 | RP11-121A8.1 |
| 7 | 38329391 | 38329591 | -2.858449133 | RP11-121A8.1 |
| 7 | 38342361 | 38342552 | -4.794892442 | TRG-AS1 |
| 7 | 38354613 | 38354812 | -9.576654785 | TRG-AS1 |
| 1 | 227317768 | 227317912 | -1.047837312 | CDC42BPA |
| 7 | 39951466 | 39951694 | -8.22612062 | CDK13 |
| 7 | 41690207 | 41690410 | 9.06266076 | INHBA |
| 7 | 41700110 | 41700303 | 8.166035762 | INHBA-AS1 |
| 7 | 41700110 | 41700303 | 8.166035762 | INHBA |
| 7 | 43609179 | 43609382 | -1.407956159 | STK17A |
| 7 | 43609179 | 43609382 | -1.407956159 | COA1 |
| 7 | 43627871 | 43628048 | -1.609382069 | COA1 |
| 7 | 43628755 | 43628954 | -1.667548014 | COA1 |
| 7 | 43631037 | 43631217 | -2.706432918 | COA1 |
| 7 | 43878990 | 43879152 | -1.128436174 | URGCP-MRPS24 |
| 7 | 43878990 | 43879152 | -1.128436174 | URGCP |
| 7 | 44006622 | 44006818 | -2.34535599 | RP5-1165K10.2 |
| 7 | 44006622 | 44006818 | -2.34535599 | AC004951.6 |
| 7 | 44006622 | 44006818 | -2.34535599 | SPDYE1 |
| 7 | 44006622 | 44006818 | -2.34535599 | POLR2J4 |
| 7 | 44060919 | 44061117 | -2.125157983 | DBNL |
| 7 | 44068755 | 44069111 | 1.538319384 | DBNL |
| 7 | 44565840 | 44566040 | -1.413035874 | DDX56 |
| 7 | 44645201 | 44645421 | -1.081494675 | OGDH |
| 7 | 44973375 | 44973572 | -6.079873737 | MYO1G |
| 7 | 45715045 | 45715241 | 1.103390337 | ADCY1 |
| 7 | 45717038 | 45717237 | 1.0460843 | ADCY1 |
| 7 | 45814774 | 45814974 | -1.346008233 | GTF2IP13 |
| 7 | 45913477 | 45913680 | 9.363853889 | IGFBP3 |
| 7 | 47275425 | 47275643 | 4.09520073 | TNS3 |
| 7 | 47277565 | 47277764 | 3.849626869 | TNS3 |
| 7 | 47368960 | 47369161 | 1.693710014 | TNS3 |
| 7 | 50400408 | 50400629 | -1.129064643 | IKZF1 |
| 7 | 50445321 | 50445497 | -3.57814697 | FIGNL1 |
| 7 | 50446111 | 50446348 | -2.960594742 | FIGNL1 |
| 7 | 50446967 | 50447147 | -3.459361314 | FIGNL1 |
| 7 | 143269234 | 143269470 | 1.337424147 | GSTK1 |
| 7 | 143289293 | 143289506 | -1.854529243 | CASP2 |
| 7 | 143356657 | 143356858 | -2.19600762 | FAM131B |
| 7 | 143379891 | 143380107 | -1.4452516 | AC093673.5 |
| 7 | 143876101 | 143876338 | -1.453749254 | TCAF1 |
| 7 | 144214123 | 144214356 | -1.193594757 | RP4-545C24.1 |
| 7 | 144215295 | 144215506 | -1.030603704 | RP4-545C24.1 |
| 7 | 144692618 | 144692807 | -2.620729436 | TPK1 |
| 7 | 144753787 | 144753907 | -4.027641218 | TPK1 |
| 7 | 148697876 | 148698063 | 2.089061958 | RP5-1136G13.2 |
| 7 | 148697876 | 148698063 | 2.089061958 | CUL1 |
| 7 | 149070367 | 149070578 | -1.105944163 | ZNF786 |
| 7 | 149072120 | 149072284 | -1.053005329 | ZNF786 |
| 7 | 149103062 | 149103258 | -1.262617229 | ZNF425 |
| 7 | 149145612 | 149145811 | -4.053457602 | ZNF398 |
| 7 | 149153991 | 149154201 | -1.680050575 | ZNF398 |
| 7 | 149178917 | 149179133 | -2.044269603 | ZNF398 |
| 7 | 149182032 | 149182301 | -1.612846133 | ZNF398 |
| 7 | 149266386 | 149266584 | -1.486563788 | ZNF783 |
| 7 | 149297022 | 149297189 | -1.752340407 | RP4-800G7.2 |
| 7 | 149297022 | 149297189 | -1.752340407 | ZNF783 |
| 7 | 149432832 | 149433031 | -1.561732238 | ZNF777 |
| 7 | 149733429 | 149733616 | -1.475872416 | KRBA1 |
| 7 | 150467077 | 150467230 | 2.76432097 | GIMAP8 |
| 7 | 150477229 | 150477547 | 2.293159179 | GIMAP8 |
| 7 | 150692390 | 150692593 | 1.230676187 | GIMAP2 |
| 7 | 150720325 | 150720560 | -1.808226533 | GIMAP1-GIMAP5 |
| 7 | 150720325 | 150720560 | -1.808226533 | GIMAP1 |
| 7 | 151122778 | 151122955 | 7.054915327 | AGAP3 |
| 7 | 151143918 | 151144118 | -1.599142897 | AGAP3 |
| 7 | 151378953 | 151379132 | 4.243784924 | WDR86 |
| 7 | 152136749 | 152136948 | -1.45451964 | KMT2C |
| 7 | 152162426 | 152162599 | -1.25540769 | KMT2C |
| 7 | 152163379 | 152163588 | -2.333491921 | KMT2C |
| 7 | 152177664 | 152177863 | -1.004666206 | KMT2C |
| 7 | 152182259 | 152182414 | -3.113717945 | KMT2C |
| 7 | 152646892 | 152647074 | -2.556104752 | XRCC2 |
| 7 | 152647390 | 152647631 | -1.860565063 | XRCC2 |
| 7 | 155775040 | 155775273 | -1.610113714 | RBM33 |
| 1 | 232036767 | 232036935 | -1.472684673 | DISC1 |
| 7 | 156639796 | 156639980 | -1.935876674 | LINC01006 |
| 7 | 157002171 | 157002301 | -4.642701109 | MNX1 |
| 7 | 157005144 | 157005268 | -6.125149209 | MNX1 |
| 7 | 157009323 | 157009502 | -2.14005396 | MNX1 |
| 7 | 157241208 | 157241413 | -1.194853309 | UBE3C |
| 7 | 157498506 | 157498669 | -3.321928095 | AC006372.4 |
| 7 | 157546453 | 157546685 | -1.470378402 | PTPRN2 |
| 7 | 157583133 | 157583239 | -4.005613628 | PTPRN2 |
| 7 | 157586688 | 157586880 | -3.79248125 | PTPRN2 |
| 7 | 157587697 | 157587840 | -1.053099702 | PTPRN2 |
| 7 | 157588361 | 157588588 | -2.136509247 | PTPRN2 |
| 7 | 157590814 | 157591013 | -6.419404993 | PTPRN2 |
| 7 | 157605917 | 157606105 | -6.04273623 | PTPRN2 |
| 7 | 157628510 | 157628667 | -4.196158711 | PTPRN2 |
| 12 | 373612 | 373761 | -7.126295761 | KDM5A |
| 12 | 374354 | 374549 | -1.637727232 | KDM5A |
| 12 | 753652 | 753877 | 1.675885614 | WNK1 |
| 12 | 917855 | 918042 | -7.074913245 | RAD52 |
| 12 | 975822 | 976021 | 1.975295953 | RAD52 |
| 12 | 975822 | 976021 | 1.975295953 | RP11-359B12.2 |
| 12 | 1533211 | 1533378 | 6.471532104 | WNT5B |
| 12 | 1646271 | 1646466 | 1.862903473 | WNT5B |
| 12 | 1792948 | 1793116 | -1.743118819 | CACNA2D4 |
| 12 | 3040379 | 3040584 | 1.699848477 | TEAD4 |
| 12 | 4352407 | 4352606 | 1.410211712 | TIGAR |
| 12 | 4520400 | 4520598 | 5.384753646 | C12orf4 |
| 12 | 4523499 | 4523685 | 4.749834953 | C12orf4 |
| 12 | 6018814 | 6019013 | -2.912945024 | VWF |
| 12 | 6329278 | 6329480 | 9.894918043 | TNFRSF1A |
| 12 | 6557135 | 6557334 | 1.271841888 | NOP2 |
| 12 | 6844971 | 6845276 | -1.912404271 | CDCA3 |
| 12 | 6844971 | 6845276 | -1.912404271 | GNB3 |
| 12 | 6872344 | 6872545 | 1.076663019 | SPSB2 |
| 12 | 6937257 | 6937442 | 9.558895098 | ATN1 |
| 12 | 6975761 | 6975929 | 1.084040651 | EMG1 |
| 12 | 6975761 | 6975929 | 1.084040651 | RP3-461F17.3 |
| 12 | 7096400 | 7096692 | 1.175225693 | C1RL |
| 12 | 7099512 | 7099721 | 1.136836174 | C1RL |
| 12 | 8059954 | 8060130 | -5.664617871 | C3AR1 |
| 12 | 8221590 | 8221753 | 1.614409345 | FAM90A1 |
| 12 | 8235494 | 8235598 | -1.030049006 | RP11-266K4.9 |
| 12 | 8235494 | 8235598 | -1.030049006 | FAM86FP |
| 12 | 8242461 | 8242777 | 3.894834743 | FAM86FP |
| 12 | 8242461 | 8242777 | 3.894834743 | RP11-266K4.9 |
| 12 | 8260756 | 8260917 | 5.353679566 | ENPP7P5 |
| 12 | 9753204 | 9753402 | 1.251393174 | CD69 |
| 12 | 10159368 | 10159565 | 2.125654324 | OLR1 |
| 12 | 10372998 | 10373195 | -2.829105741 | KLRC4-KLRK1 |
| 12 | 10372998 | 10373195 | -2.829105741 | KLRK1 |
| 12 | 10372998 | 10373195 | -2.829105741 | RP11-277P12.20 |
| 12 | 10847257 | 10847447 | -1.195934167 | PRR4 |
| 12 | 10847257 | 10847447 | -1.195934167 | PRH1 |
| 12 | 10847257 | 10847447 | -1.195934167 | PRH1-PRR4 |
| 12 | 11220100 | 11220238 | -7.200406371 | RP11-711K1.8 |
| 12 | 11379317 | 11379491 | -1.392993622 | PRB1 |
| 12 | 11387866 | 11388047 | -1.487719836 | PRB1 |
| 12 | 11513987 | 11514185 | -1.833648717 | DDX55P1 |
| 12 | 11720628 | 11720816 | -1.103397783 | ETV6 |
| 12 | 11824840 | 11824942 | 4.756863798 | ETV6 |
| 12 | 12165222 | 12165387 | -7.096069336 | BCL2L14 |
| 12 | 12165222 | 12165387 | -7.096069336 | LRP6 |
| 12 | 12465722 | 12465920 | 1.02428853 | BORCS5 |
| 12 | 12477336 | 12477536 | -1.000043377 | DUSP16 |
| 12 | 12774030 | 12774217 | -3.696158711 | APOLD1 |
| 12 | 12788787 | 12788986 | 1.98937044 | APOLD1 |
| 12 | 18598471 | 18598665 | -1.254777549 | PIK3C2G |
| 12 | 19274690 | 19274887 | 2.216626902 | PLEKHA5 |
| 12 | 19283407 | 19283607 | 1.775542648 | PLEKHA5 |

| | | | | | | | | | | | | | | |
|---|---|---|---|---|---|---|---|---|---|---|---|---|---|---|
| 1 | 227701827 | 227702030 | 1.389141736 | ZNF847P | 1 | 232514834 | 232515030 | -4.292604826 | SIPA1L2 | 12 | 20369052 | 20369203 | 1.975989243 | RP11-284H19.1 |
| 7 | 53762895 | 53763082 | -2.405876023 | LINC01446 | 7 | 157857806 | 157858002 | -3.92050133 | AC011899.9 | 12 | 21515736 | 21515860 | -1.032612408 | GOLT1B |
| 7 | 55543949 | 55544149 | 2.215819733 | VOPP1 | 7 | 157857806 | 157858002 | -3.92050133 | PTPRN2 | 12 | 23820795 | 23820931 | -5.093676037 | SOX5 |
| 1 | 227732451 | 227732665 | -2.341929796 | SNAP47 | 7 | 157860199 | 157860407 | -3.429539456 | PTPRN2 | 12 | 23823031 | 23823184 | -1.117533126 | SOX5 |
| 1 | 227732451 | 227732665 | -2.341929796 | JMJD4 | 7 | 157860199 | 157860407 | -3.429539456 | AC011899.9 | 12 | 24584079 | 24584225 | 1.921693827 | LINC00477 |
| 7 | 64500803 | 64500909 | -2.489520191 | HNRNPCP7 | 7 | 157862854 | 157863057 | -3.153957647 | AC011899.9 | 12 | 24584079 | 24584225 | 1.921693827 | KNOP1P1 |
| 7 | 64521158 | 64521318 | -1.925492382 | ZNF680 | 7 | 157862854 | 157863057 | -3.153957647 | PTPRN2 | 12 | 25958107 | 25958304 | -2.166811714 | RASSF8-AS1 |
| 7 | 64708077 | 64708277 | -2.054597589 | ZNF107 | 1 | 232805914 | 232806151 | -1.695004754 | MAP10 | 12 | 26122758 | 26123090 | 1.969822599 | SSPN |
| 7 | 64709224 | 64709435 | -2.706849818 | ZNF107 | 7 | 157868433 | 157868632 | -7.369732097 | PTPRN2 | 12 | 26122758 | 26123090 | 1.969822599 | BHLHE41 |
| 1 | 227734720 | 227734927 | -1.655039506 | SNAP47 | 7 | 157868433 | 157868632 | -7.369732097 | RP11-452C13.1 | 12 | 27160462 | 27160657 | 5.164898169 | RP11-582E3.2 |
| 1 | 227734720 | 227734927 | -1.655039506 | JMJD4 | 7 | 158876438 | 158876630 | -6.834719369 | WDR60 | 12 | 27405173 | 27405327 | 1.166645224 | ARNTL2 |
| 7 | 64710006 | 64710201 | -2.699635092 | ZNF107 | 8 | 245290 | 245501 | -1.29230371 | ZNF596 | 12 | 27405173 | 27405327 | 1.166645224 | ARNTL2-AS1 |
| 7 | 64832938 | 64833076 | -1.21004594 | ZNF138 | 1 | 232806936 | 232807120 | -3.08510677 | MAP10 | 12 | 27406787 | 27406990 | 7.18223834 | ARNTL2 |
| 7 | 64889982 | 64890144 | -2.546876727 | ZNF273 | 8 | 642978 | 643099 | -5.957939689 | ERICH1 | 12 | 27406787 | 27406990 | 7.18223834 | ARNTL2-AS1 |
| 7 | 64889982 | 64890144 | -2.546876727 | RP11-797H7.5 | 8 | 1761479 | 1761691 | -2.291240943 | CLN8 | 12 | 27410293 | 27410496 | -5.316043206 | ARNTL2 |
| 7 | 64928743 | 64928939 | -2.257897031 | ZNF273 | 8 | 1770922 | 1771136 | -1.493986838 | CLN8 | 12 | 27410293 | 27410496 | -5.316043206 | ARNTL2-AS1 |
| 7 | 64942400 | 64942601 | 3.266558873 | RP11-460N20.7 | 8 | 1770922 | 1771136 | -1.493986838 | RP11-439C15.6 | 12 | 27595376 | 27595567 | 3.335481524 | PPFIBP1 |
| 1 | 10948088 | 10948287 | -1.688062397 | C1orf127 | 8 | 1780362 | 1780566 | -1.75388163 | RP11-439C15.6 | 12 | 27702594 | 27702793 | 3.678922603 | RP11-1060J15.5 |
| 1 | 227747992 | 227748191 | -1.288742795 | SNAP47 | 8 | 1780362 | 1780566 | -1.75388163 | CLN8 | 12 | 27702594 | 27702793 | 3.678922603 | RP11-1060J15.4 |
| 7 | 65200603 | 65200794 | 6.53890856 | INTS4P1 | 1 | 11295782 | 11295909 | -2.658706307 | UBIAD1 | 12 | 28275782 | 28276008 | -2.037173671 | CCDC91 |
| 7 | 65214850 | 65215050 | 2.402388189 | INTS4P1 | 8 | 2000542 | 2000747 | -3.233234975 | KBTBD11 | 12 | 29326554 | 29326762 | -9.224628741 | FAR2 |
| 7 | 65399754 | 65399989 | -1.328331282 | ZNF92 | 8 | 2003183 | 2003384 | -2.871356086 | KBTBD11 | 12 | 29327848 | 29327980 | 2.190137603 | FAR2 |
| 7 | 65720768 | 65720950 | -2.013673033 | RP11-479O9.4 | 8 | 2006525 | 2006744 | -3.97652375 | KBTBD11 | 12 | 29328437 | 29328637 | 1.438914065 | FAR2 |
| 7 | 65721339 | 65721538 | -2.351695036 | RP11-479O9.4 | 8 | 6405905 | 6406037 | -3.542518052 | RP11-115C21.2 | 1 | 32826514 | 32826727 | -1.030755781 | S100PBP |
| 7 | 65958962 | 65959160 | -2.052912022 | VKORC1L1 | 8 | 6618978 | 6619151 | -6.378278161 | MCPH1-AS1 | 12 | 29406506 | 29406704 | -1.247908681 | OVCH1-AS1 |
| 7 | 66240436 | 66240701 | 4.330225857 | TPST1 | 8 | 6618978 | 6619151 | -6.378278161 | MCPH1 | 12 | 29410081 | 29410280 | -1.901399193 | OVCH1-AS1 |
| 7 | 66359307 | 66359480 | 4.290765863 | TPST1 | 8 | 6757388 | 6757581 | -1.137050742 | AGPAT5 | 12 | 29441514 | 29441649 | -2.934012562 | OVCH1-AS1 |
| 7 | 67238443 | 67238643 | -1.130375846 | TYW1 | 8 | 6758468 | 6758679 | -1.257302748 | AGPAT5 | 12 | 29441514 | 29441649 | -2.934012562 | OVCH1 |
| 7 | 72913073 | 72913275 | -3.590052084 | POM121 | 1 | 232976556 | 232976709 | -6.495583753 | NTPCR | 12 | 29442144 | 29442334 | -1.77077261 | OVCH1-AS1 |
| 7 | 72946342 | 72946542 | -1.075088124 | POM121 | 8 | 8235608 | 8235806 | -4.623963757 | FAM86B3P | 12 | 29442144 | 29442334 | -1.77077261 | OVCH1 |
| 1 | 228104384 | 228104595 | -2.924403104 | C1orf35 | 8 | 8236087 | 8236285 | -3.676100553 | FAM86B3P | 12 | 29449950 | 29450145 | -2.517636825 | OVCH1 |
| 7 | 73435519 | 73435714 | -1.119975396 | FZD9 | 8 | 8236087 | 8236285 | -3.676100553 | ALG1L13P | 12 | 29449950 | 29450145 | -2.517636825 | OVCH1-AS1 |
| 7 | 73573475 | 73573634 | -8.171618887 | TBL2 | 8 | 8319210 | 8319404 | -2.12510116 | SGK223 | 12 | 29465259 | 29465495 | -2.977089361 | OVCH1 |
| 7 | 73682750 | 73682874 | -1.152480011 | DNAJC30 | 8 | 8327803 | 8328009 | -2.262081522 | SGK223 | 12 | 29465259 | 29465495 | -2.977089361 | OVCH1-AS1 |
| 7 | 73736187 | 73736431 | -1.603380734 | ABHD11 | 8 | 8339545 | 8339734 | -3.823986149 | SGK223 | 12 | 29584585 | 29584776 | 5.421175172 | TMTC1 |
| 7 | 74338415 | 74338613 | 7.746302152 | CLIP2 | 8 | 8376405 | 8376583 | -2.622628593 | SGK223 | 12 | 31387425 | 31387630 | -1.400724484 | DENND5B |
| 7 | 74376174 | 74376352 | 3.277292005 | CLIP2 | 8 | 8377503 | 8377703 | -2.319558021 | SGK223 | 12 | 31551053 | 31551199 | -1.385783122 | DENND5B |
| 7 | 74380633 | 74380836 | 2.570976068 | CLIP2 | 1 | 232983449 | 232983592 | -1.427805045 | NTPCR | 12 | 31553177 | 31553376 | -2.867989905 | DENND5B |
| 7 | 74386351 | 74386573 | 2.790681952 | CLIP2 | 1 | 232983449 | 232983592 | -1.427805045 | PCNX2 | 12 | 31981166 | 31981359 | -1.280223839 | KIAA1551 |
| 7 | 74403841 | 74404056 | 3.631984985 | CLIP2 | 8 | 8381517 | 8381714 | -6.345435505 | SGK223 | 12 | 31982778 | 31982971 | -1.366863247 | KIAA1551 |
| 7 | 74883136 | 74883247 | -7.387444994 | STAG3L2 | 8 | 9661615 | 9661745 | -2.453445298 | TNKS | 12 | 31983661 | 31983861 | -1.248874819 | KIAA1551 |
| 7 | 74907100 | 74907355 | -1.219499664 | SPDYE12P | 8 | 9777441 | 9777644 | -1.307538344 | TNKS | 12 | 31985559 | 31985859 | -1.279039895 | KIAA1551 |
| 1 | 228175246 | 228175451 | -2.152769818 | IBA57 | 1 | 233258262 | 233258611 | -1.839290819 | PCNX2 | 12 | 32377653 | 32377852 | -1.363557442 | BICD1 |
| 7 | 75536538 | 75536794 | -1.664569838 | HIP1 | 8 | 11107905 | 11108084 | 2.49781078 | AF131215.2 | 12 | 38320344 | 38320540 | -7.579225952 | ALG10B |
| 7 | 76329593 | 76329781 | -1.112845551 | YWHAG | 8 | 11107905 | 11108084 | 2.49781078 | XKR6 | 12 | 42225394 | 42225593 | -1.914870718 | YAF2 |
| 7 | 76389440 | 76389630 | 1.446231762 | SSC4D | 8 | 11108769 | 11108906 | 1.718702656 | AF131215.2 | 12 | 43955064 | 43955260 | -2.245181739 | TMEM117 |
| 7 | 76980907 | 76981112 | -1.01242711 | DTX2P1-UPK3BP1-PMS2P11 | 8 | 11108769 | 11108906 | 1.718702656 | XKR6 | 1 | 1313234 | 1313497 | 1.526812534 | CPSF3L |
| 7 | 76980907 | 76981112 | -1.01242711 | DTX2P1 | 8 | 11118862 | 11119058 | 1.352145451 | XKR6 | 12 | 45726971 | 45727153 | 1.399353555 | RP11-352M15.2 |
| 7 | 77196728 | 77196881 | 8.927165722 | CCDC146 | 8 | 11119314 | 11119488 | 1.180092269 | XKR6 | 12 | 45823090 | 45823222 | -5.874934714 | ARID2 |
| 7 | 77196728 | 77196881 | 8.927165722 | RP11-467H10.2 | 1 | 233344667 | 233344868 | -3.440913869 | RP5-862P8.2 | 12 | 45974716 | 45974933 | -9.353005145 | SCAF11 |
| 7 | 77196728 | 77196881 | 8.927165722 | FGL2 | 8 | 12542574 | 12542774 | -4.953445298 | RP11-419I17.1 | 12 | 46399887 | 46400061 | -1.072013613 | RP11-96H19.1 |
| 1 | 228274489 | 228274698 | -1.455081578 | RP5-1139B12.3 | 8 | 12565457 | 12565614 | -1.058242871 | RP11-419I17.1 | 12 | 46643600 | 46643708 | 4.474183616 | RP11-96H19.1 |
| 1 | 228274489 | 228274698 | -1.455081578 | OBSCN | 8 | 12722995 | 12723251 | 1.690572097 | LONRF1 | 12 | 46643600 | 46643708 | 4.474183616 | RP11-446N19.1 |
| 7 | 80676512 | 80676750 | 5.620879788 | CD36 | 8 | 17645692 | 17645893 | -2.855824846 | MTUS1 | 12 | 47844316 | 47844513 | 1.23687879 | VDR |

| | | | | |
|---|---|---|---|---|
| 7 | 80744967 | 80745189 | 6.456251725 | SEMA3C |
| 7 | 82914852 | 82914965 | 1.515622735 | PCLO |
| 7 | 82916227 | 82916440 | 1.464690463 | PCLO |
| 7 | 82953322 | 82953520 | 1.141444625 | PCLO |
| 7 | 82966378 | 82966576 | 1.24318814 | PCLO |
| 1 | 228396561 | 228396757 | -2.654291409 | TRIM11 |
| 1 | 228396561 | 228396757 | -2.654291409 | RP11-245P10.8 |
| 7 | 83134469 | 83134755 | 1.358467563 | PCLO |
| 7 | 83135209 | 83135417 | 1.361168317 | PCLO |
| 7 | 83961468 | 83961669 | -1.867072836 | SEMA3A |
| 7 | 84194551 | 84194767 | -2.258734539 | SEMA3A |
| 7 | 84999592 | 84999793 | 4.202408786 | SEMA3D |
| 7 | 87907462 | 87907667 | -1.265922668 | DBF4 |
| 1 | 11023312 | 11023558 | -1.551298934 | TARDBP |
| 1 | 11023312 | 11023558 | -1.551298934 | RP4-635E18.9 |
| 1 | 228410535 | 228410687 | -3.413909512 | TRIM17 |
| 7 | 90412674 | 90412900 | -1.520807438 | CTB-13L3.1 |
| 7 | 90412674 | 90412900 | -1.520807438 | CLDN12 |
| 7 | 91175499 | 91175695 | 1.810914942 | CDK14 |
| 7 | 91873489 | 91873705 | -1.901460099 | MTERF1 |
| 7 | 92002647 | 92002921 | -1.093235655 | AKAP9 |
| 1 | 228494209 | 228494407 | -1.702525279 | RNF187 |
| 7 | 92491232 | 92491431 | 2.822288679 | PEX1 |
| 7 | 92491232 | 92491431 | 2.822288679 | AC007566.10 |
| 1 | 228651295 | 228651492 | -4.0110169 | DUSP5P1 |
| 7 | 92776931 | 92777116 | -8.311426293 | CDK6 |
| 7 | 92833614 | 92833812 | -9.264444507 | CDK6 |
| 7 | 93100451 | 93100634 | 1.933319994 | SAMD9 |
| 7 | 93102626 | 93102828 | 1.919488211 | SAMD9 |
| 7 | 93103841 | 93104033 | 1.114974886 | SAMD9 |
| 7 | 93104996 | 93105198 | 2.67984615 | SAMD9 |
| 7 | 93105826 | 93106026 | 2.841572913 | SAMD9 |
| 7 | 93132585 | 93132790 | 3.873677897 | SAMD9L |
| 7 | 93133320 | 93133474 | 4.130716087 | SAMD9L |
| 1 | 228652335 | 228652488 | -4.221666989 | SNORA51 |
| 7 | 93133979 | 93134085 | 3.932673322 | SAMD9L |
| 7 | 93135659 | 93135835 | 3.97814276 | SAMD9L |
| 7 | 93344025 | 93344130 | -5.321477112 | VPS50 |
| 7 | 98194051 | 98194262 | -1.169134533 | LMTK2 |
| 7 | 98207243 | 98207414 | -5.690501055 | LMTK2 |
| 1 | 228687438 | 228687633 | -9.625953835 | FTH1P2 |
| 7 | 98302485 | 98302746 | 3.493380112 | BAIAP2L1 |
| 7 | 98302485 | 98302746 | 3.493380112 | BRI3 |
| 7 | 98307390 | 98307620 | 3.190322879 | BRI3 |
| 7 | 98307390 | 98307620 | 3.190322879 | BAIAP2L1 |
| 7 | 98309554 | 98309751 | 7.684185021 | BAIAP2L1 |
| 7 | 98309554 | 98309751 | 7.684185021 | BRI3 |
| 7 | 99012222 | 99012419 | -1.310012728 | TRRAP |
| 7 | 99419814 | 99420021 | -1.728687048 | ATP5J2-PTCD1 |
| 7 | 99419814 | 99420021 | -1.728687048 | PTCD1 |
| 7 | 99425372 | 99425558 | -1.987441865 | ATP5J2-PTCD1 |
| 7 | 99425372 | 99425558 | -1.987441865 | PTCD1 |
| 7 | 99486967 | 99487151 | -3.320366173 | ZNF789 |
| 7 | 99486967 | 99487151 | -3.320366173 | ZNF394 |
| 8 | 17754020 | 17754211 | -2.786678223 | MTUS1 |
| 8 | 17754998 | 17755104 | -3.322564297 | MTUS1 |
| 1 | 233345294 | 233345480 | -2.978507038 | RP5-862P8.2 |
| 8 | 17989005 | 17989225 | -2.218665163 | PCM1 |
| 8 | 18528321 | 18528573 | 1.01086002 | PSD3 |
| 1 | 233379412 | 233379611 | -1.682164219 | RP5-862P8.2 |
| 8 | 19319400 | 19319597 | -7.50779464 | SH2D4A |
| 8 | 19395498 | 19395675 | -6.428990498 | SH2D4A |
| 8 | 19405737 | 19405937 | -1.744643366 | CSGALNACT1 |
| 8 | 20249196 | 20249389 | 1.35491655 | LZTS1 |
| 8 | 21909259 | 21909519 | -1.538959517 | DOK2 |
| 8 | 22107279 | 22107497 | -1.048315051 | NUDT18 |
| 8 | 22211761 | 22211954 | 1.473211203 | BMP1 |
| 1 | 233382426 | 233382632 | -1.306358203 | RP5-862P8.2 |
| 8 | 22603736 | 22603936 | -1.499668422 | C8orf58 |
| 8 | 22613997 | 22614220 | -1.736188172 | RP11-582J16.5 |
| 8 | 22613997 | 22614220 | -1.736188172 | CCAR2 |
| 8 | 22690905 | 22691100 | -1.29903409 | RP11-459E5.1 |
| 8 | 22690905 | 22691100 | -1.29903409 | EGR3 |
| 8 | 23007121 | 23007319 | -1.941024775 | RHOBTB2 |
| 8 | 23007121 | 23007319 | -1.941024775 | RP11-875O11.1 |
| 8 | 23017554 | 23017749 | -2.575737159 | RP11-875O11.1 |
| 8 | 23017554 | 23017749 | -2.575737159 | RHOBTB2 |
| 8 | 23018249 | 23018449 | -2.45261657 | RHOBTB2 |
| 8 | 23018249 | 23018449 | -2.45261657 | RP11-875O11.1 |
| 8 | 23246477 | 23246676 | -8.108306626 | CHMP7 |
| 8 | 23291251 | 23291443 | -6.72766361 | R3HCC1 |
| 8 | 24340885 | 24341082 | 7.062827811 | RP11-624C23.1 |
| 8 | 24340885 | 24341082 | 7.062827811 | ADAM28 |
| 1 | 234607364 | 234607569 | -1.43420954 | RP4-781K5.2 |
| 1 | 234607364 | 234607569 | -1.43420954 | IRF2BP2 |
| 8 | 25411627 | 25411827 | -1.648147303 | DOCK5 |
| 8 | 25507309 | 25507527 | -1.598371563 | CDCA2 |
| 8 | 26324646 | 26324900 | -7.859507204 | PPP2R2A |
| 8 | 26370569 | 26370770 | 1.270630358 | PPP2R2A |
| 8 | 26379636 | 26379803 | -1.297579134 | SDAD1P1 |
| 8 | 27776516 | 27776701 | 1.75389732 | ESCO2 |
| 8 | 28348546 | 28348749 | -3.0309998 | ZNF395 |
| 8 | 28348546 | 28348749 | -3.0309998 | FBXO16 |
| 8 | 28502830 | 28503079 | -1.951639671 | FZD3 |
| 8 | 28562747 | 28562948 | -2.684307794 | FZD3 |
| 8 | 28731068 | 28731269 | -1.060018063 | EXTL3 |
| 8 | 28750940 | 28751215 | -1.248264064 | EXTL3 |
| 8 | 30156017 | 30156204 | 4.430543453 | LEPROTL1 |
| 8 | 30156017 | 30156204 | 4.430543453 | RP11-51J9.5 |
| 8 | 30624037 | 30624208 | 1.64038422 | GTF2E2 |
| 8 | 30744142 | 30744338 | 2.326394657 | UBXN8 |
| 8 | 32763972 | 32764172 | 5.935324077 | NRG1 |
| 8 | 33389088 | 33389310 | -1.56484322 | FUT10 |
| 8 | 33453285 | 33453479 | -1.554902005 | FUT10 |
| 8 | 37698501 | 37698718 | -3.694147295 | ZNF703 |
| 8 | 37842435 | 37842633 | -1.690781163 | ADGRA2 |
| 8 | 37862826 | 37863050 | -2.150006847 | RAB11FIP1 |
| 8 | 37874809 | 37875011 | -1.320110378 | RAB11FIP1 |
| 12 | 48815211 | 48815408 | 1.055487056 | CACNB3 |
| 12 | 48981544 | 48981715 | 2.708995126 | WNT1 |
| 1 | 33494846 | 33494988 | -1.797129365 | ZSCAN20 |
| 12 | 49233311 | 49233520 | 3.330982785 | RP11-977B10.2 |
| 12 | 49233311 | 49233520 | 3.330982785 | TUBA1C |
| 12 | 49330219 | 49330413 | -3.717760723 | TROAP |
| 12 | 49389924 | 49390121 | -1.524083641 | SPATS2 |
| 12 | 49389924 | 49390121 | -1.524083641 | RP11-161H23.10 |
| 12 | 49642519 | 49642767 | 1.048519355 | PRPF40B |
| 12 | 49642519 | 49642767 | 1.048519355 | FMNL3 |
| 12 | 52054678 | 52054829 | 2.155590253 | NR4A1 |
| 12 | 53500889 | 53501068 | 3.13339327 | RP11-793H13.11 |
| 12 | 53500889 | 53501068 | 3.13339327 | TARBP2 |
| 12 | 54181534 | 54181734 | 1.499766403 | SMUG1 |
| 12 | 54292345 | 54292544 | 1.264739084 | NFE2 |
| 12 | 54292345 | 54292544 | 1.264739084 | RP11-968A15.8 |
| 12 | 54362565 | 54362768 | -1.162655245 | RP11-753H16.5 |
| 12 | 54362565 | 54362768 | -1.162655245 | GPR84 |
| 12 | 54362565 | 54362768 | -1.162655245 | RP11-753H16.3 |
| 12 | 54363011 | 54363227 | -1.208640424 | RP11-753H16.5 |
| 12 | 54363011 | 54363227 | -1.208640424 | RP11-753H16.3 |
| 12 | 54363011 | 54363227 | -1.208640424 | GPR84 |
| 1 | 1314475 | 1314612 | 1.362542534 | CPSF3L |
| 12 | 55752758 | 55753115 | -1.424598576 | GDF11 |
| 12 | 55752758 | 55753115 | -1.424598576 | SARNP |
| 12 | 57091496 | 57091696 | 1.117633613 | NAB2 |
| 12 | 57456344 | 57456550 | 2.73776501 | INHBE |
| 12 | 57516952 | 57517084 | 6.638062203 | DDIT3 |
| 12 | 57516952 | 57517084 | 6.638062203 | MARS |
| 12 | 57726324 | 57726534 | 1.272649504 | AGAP2 |
| 12 | 57726324 | 57726534 | 1.272649504 | AGAP2-AS1 |
| 12 | 62466903 | 62467103 | 8.134225311 | MON2 |
| 12 | 62511746 | 62511942 | -2.271678615 | MON2 |
| 12 | 62560499 | 62560698 | -2.287888572 | MON2 |
| 12 | 62648366 | 62648569 | -1.32829163 | PPM1H |
| 12 | 64143226 | 64143329 | 1.667580774 | SRGAP1 |
| 1 | 35560423 | 35560625 | 1.425926801 | NCDN |
| 12 | 64153130 | 64153320 | -6.19333543 | SRGAP1 |
| 12 | 64451718 | 64451897 | 7.565908389 | RP11-114F3.4 |
| 12 | 64451718 | 64451897 | 7.565908389 | TBK1 |
| 12 | 64876429 | 64876711 | 1.084604601 | TBC1D30 |
| 12 | 65464216 | 65464415 | -4.660964047 | MSRB3 |
| 12 | 68764739 | 68764939 | 1.581627665 | SLC35E3 |
| 12 | 69284865 | 69285064 | -6.861937109 | C1GALT1P1 |
| 12 | 70183386 | 70183534 | -2.634519386 | LINC01481 |
| 12 | 70183386 | 70183534 | -2.634519386 | RP11-320P7.2 |
| 12 | 70235412 | 70235607 | -1.435062945 | LINC01481 |
| 12 | 70556387 | 70556579 | -1.009901433 | PTPRB |
| 1 | 1315006 | 1315210 | 1.305169163 | CPSF3L |
| 1 | 35720766 | 35720966 | -3.857122759 | CLSPN |
| 12 | 71663121 | 71663308 | 1.111512003 | ZFC3H1 |
| 12 | 71663121 | 71663308 | 1.111512003 | THAP2 |
| 12 | 74537880 | 74538090 | 1.44664838 | ATXN7L3B |
| 12 | 75360216 | 75360392 | -5.827318014 | CAPS2 |

| | | | | | | | | | | | | | | |
|---|---|---|---|---|---|---|---|---|---|---|---|---|---|---|
| 7 | 99499923 | 99500078 | -1.688905064 | ZNF789 | 8 | 37877316 | 37877512 | -8.311697855 | RAB11FIP1 | 12 | 75360216 | 75360392 | -5.827318014 | GLIPR1L1 |
| 7 | 99499923 | 99500078 | -1.688905064 | ZNF394 | 8 | 38210249 | 38210505 | -1.215585315 | BAG4 | 12 | 78119517 | 78119715 | -1.9037611 | NAV3 |
| 7 | 99892519 | 99892723 | 1.017245442 | TRIM4 | 8 | 38275620 | 38275809 | -7.223799929 | WHSC1L1 | 12 | 79775856 | 79776040 | -1.058830905 | PPP1R12A |
| 7 | 99900944 | 99901141 | -2.073700261 | TRIM4 | 8 | 38329624 | 38329818 | -1.222930894 | WHSC1L1 | 12 | 79775856 | 79776040 | -1.058830905 | RP11-530C5.1 |
| 7 | 100106954 | 100107159 | -1.026619703 | TAF6 | 8 | 38392711 | 38392910 | -1.207586192 | LETM2 | 1 | 35764360 | 35764525 | -1.197488112 | CLSPN |
| 7 | 100106954 | 100107159 | -1.026619703 | AP4M1 | 8 | 39927983 | 39928169 | 13.72483337 | IDO1 | 12 | 89525562 | 89525742 | 1.225229952 | RP11-734K2.4 |
| 7 | 100120071 | 100120191 | -6.268123608 | RP11-506M12.1 | 8 | 39927983 | 39928169 | 13.72483337 | RP11-44K6.3 | 12 | 89525562 | 89525742 | 1.225229952 | POC1B |
| 7 | 100120071 | 100120191 | -6.268123608 | CNPY4 | 8 | 41828485 | 41828606 | -5.959990298 | RP11-930P14.2 | 12 | 89525562 | 89525742 | 1.225229952 | POC1B-GALNT4 |
| 7 | 100211327 | 100211528 | 8.52734455 | GATS | 8 | 41828485 | 41828606 | -5.959990298 | ANK1 | 12 | 89542108 | 89542334 | 1.573498254 | RP11-734K2.4 |
| 7 | 100211327 | 100211528 | 8.52734455 | STAG3 | 8 | 41831962 | 41832113 | -5.434411277 | ANK1 | 12 | 93398628 | 93398830 | -1.139237008 | NUDT4 |
| 7 | 100319303 | 100319503 | -1.297953771 | SPDYE3 | 8 | 41937466 | 41937664 | -9.078456304 | KAT6A | 12 | 94288575 | 94288745 | -6.59722454 | PLXNC1 |
| 7 | 100358814 | 100359015 | -1.157490803 | PILRB | 8 | 42025756 | 42025887 | -1.578035857 | KAT6A | 1 | 35855946 | 35856141 | -1.751503295 | AGO4 |
| 7 | 100358814 | 100359015 | -1.157490803 | STAG3L5P-PVRIG2P-PILRB | 8 | 42169298 | 42169446 | -5.699371846 | AP3M2 | 12 | 96249076 | 96249271 | -6.790541432 | ELK3 |
| 7 | 100367357 | 100367582 | -1.579436412 | STAG3L5P-PVRIG2P-PILRB | 8 | 42297433 | 42297610 | 6.493454312 | IKBKB | 12 | 96259659 | 96259857 | -2.344036649 | ELK3 |
| 7 | 100367357 | 100367582 | -1.579436412 | PILRB | 8 | 42297896 | 42298083 | 6.283978038 | IKBKB | 12 | 98547891 | 98548112 | -1.228319466 | TMPO |
| 7 | 100367357 | 100367582 | -1.579436412 | PILRA | 8 | 42437187 | 42437387 | -1.064288442 | SLC20A2 | 12 | 100586445 | 100586638 | -2.208745761 | GAS2L3 |
| 1 | 229258835 | 229258948 | -1.424774134 | RP5-1061H20.4 | 8 | 42472459 | 42472669 | -1.673954119 | SLC20A2 | 12 | 101764481 | 101764644 | -1.224984199 | GNPTAB |
| 7 | 100430700 | 100430912 | -1.676153974 | MEPCE | 8 | 42542126 | 42542287 | -6.084649348 | SMIM19 | 12 | 101921990 | 101922177 | -1.819036919 | DRAM1 |
| 7 | 100865861 | 100866025 | -3.095159274 | SLC12A9 | 8 | 43167022 | 43167225 | -1.923905681 | HGSNAT | 12 | 104304216 | 104304413 | -7.734773216 | TXNRD1 |
| 7 | 101091018 | 101091220 | -2.205602819 | TRIM56 | 1 | 235450236 | 235450371 | -1.980423843 | B3GALNT2 | 12 | 104304216 | 104304413 | -7.734773216 | EID3 |
| 1 | 11083890 | 11084090 | -1.717266436 | EXOSC10 | 8 | 47313488 | 47313704 | 1.291879877 | SPIDR | 1 | 36090215 | 36090382 | -1.035538631 | ADPRHL2 |
| 1 | 229268511 | 229268710 | -1.122253833 | RP5-1061H20.4 | 8 | 47468624 | 47468814 | -3.125225623 | SPIDR | 12 | 104348360 | 104348554 | 9.514996759 | TXNRD1 |
| 7 | 102348473 | 102348674 | -1.510241973 | SPDYE6 | 8 | 47623364 | 47623628 | -5.821928095 | SPIDR | 12 | 105024379 | 105024579 | -2.172632455 | ALDH1L2 |
| 7 | 102396010 | 102396210 | -1.883542467 | PRKRIP1 | 8 | 47626248 | 47626370 | -6.273567766 | SPIDR | 12 | 105024379 | 105024579 | -2.172632455 | C12orf45 |
| 7 | 102457297 | 102457492 | -1.523509338 | ALKBH4 | 8 | 47627067 | 47627230 | 4.183161107 | SPIDR | 12 | 105026643 | 105026842 | -2.169989854 | ALDH1L2 |
| 7 | 102559861 | 102560145 | -6.513434603 | POLR2J3 | 1 | 235662804 | 235662982 | -1.763661695 | LYST | 12 | 105026643 | 105026842 | -2.169989854 | C12orf45 |
| 7 | 102559861 | 102560145 | -6.513434603 | SPDYE2 | 8 | 47737218 | 47737404 | 12.49908865 | CEBPD | 12 | 106346429 | 106346628 | -1.475200169 | TCP11L2 |
| 7 | 102559861 | 102560145 | -6.513434603 | RP11-514P8.7 | 8 | 49176835 | 49177064 | -1.84509186 | RP11-10H3.1 | 1 | 36096988 | 36097186 | -2.109113429 | COL8A2 |
| 7 | 102659056 | 102659237 | -1.798439639 | RP11-577H5.5 | 8 | 49178172 | 49178323 | -1.196422387 | RP11-10H3.1 | 12 | 108291520 | 108291718 | 2.354475609 | CMKLR1 |
| 7 | 102659056 | 102659237 | -1.798439639 | SPDYE2B | 8 | 49200478 | 49200671 | -1.977373908 | RP11-10H3.1 | 12 | 109053735 | 109053919 | -3.660964047 | USP30-AS1 |
| 7 | 102668615 | 102668803 | -1.34464958 | RP11-577H5.5 | 1 | 235738003 | 235738197 | -1.518448475 | LDHAP2 | 12 | 109053735 | 109053919 | -3.660964047 | USP30 |
| 7 | 102668615 | 102668803 | -1.34464958 | POLR2J2 | 1 | 235738003 | 235738197 | -1.518448475 | LYST | 1 | 36097652 | 36097839 | -3.127372599 | COL8A2 |
| 1 | 229269260 | 229269509 | -1.042167805 | RP5-1061H20.4 | 8 | 51820426 | 51820642 | -1.085024377 | PCMTD1 | 12 | 109267599 | 109267794 | 2.793982494 | ACACB |
| 7 | 102782742 | 102782940 | -2.499248205 | FAM185A | 8 | 51860946 | 51861150 | -1.306453106 | PCMTD1 | 12 | 109769303 | 109769469 | 2.626868197 | FAM222A-AS1 |
| 7 | 102934350 | 102934506 | -1.291740774 | LRRC17 | 8 | 51946899 | 51947097 | -2.311219357 | RP11-110G21.1 | 12 | 109769303 | 109769469 | 2.626868197 | FAM222A |
| 7 | 102934350 | 102934506 | -1.291740774 | FBXL13 | 8 | 52656154 | 52656358 | -1.550849407 | RB1CC1 | 12 | 110429953 | 110430144 | -6.643567593 | RP11-478C19.2 |
| 7 | 103109504 | 103109660 | 2.741645541 | NAPEPLD | 8 | 52657213 | 52657415 | -1.126441103 | RB1CC1 | 12 | 110429953 | 110430144 | -6.643567593 | HMGA1P3 |
| 7 | 104950206 | 104950406 | 6.277534196 | LINC01004 | 8 | 53781942 | 53782139 | 7.283086494 | ATP6V1H | 12 | 110502478 | 110502710 | 1.670335278 | RAD9B |
| 7 | 104950206 | 104950406 | 6.277534196 | RP11-325F22.2 | 1 | 235738293 | 235738492 | -1.30708608 | LDHAP2 | 12 | 110502478 | 110502710 | 1.670335278 | RP3-424M6.4 |
| 7 | 105360119 | 105360317 | -1.643605945 | SRPK2 | 1 | 235738293 | 235738492 | -1.30708608 | LYST | 12 | 111347738 | 111347863 | -2.037685572 | CUX2 |
| 7 | 105508266 | 105508512 | -2.42040395 | PUS7 | 8 | 53783632 | 53783806 | 6.369573855 | ATP6V1H | 12 | 111813528 | 111813709 | -2.321928095 | ALDH2 |
| 7 | 105712524 | 105712728 | -1.252632062 | ATXN7L1 | 8 | 53934252 | 53934456 | -1.964546115 | RGS20 | 12 | 112140809 | 112140975 | 2.899915159 | TRAFD1 |
| 1 | 229518171 | 229518370 | -2.185709611 | ABCB10 | 8 | 56010263 | 56010507 | 1.715718457 | LYN | 12 | 112505740 | 112505950 | 1.492134828 | PTPN11 |
| 7 | 107629053 | 107629263 | -2.836807053 | BCAP29 | 1 | 235740457 | 235740653 | -1.961081726 | LYST | 12 | 113296082 | 113296231 | -1.314706329 | TPCN1 |
| 7 | 107629053 | 107629263 | -2.836807053 | WBP1LP2 | 8 | 56161428 | 56161591 | -1.058863434 | PLAG1 | 12 | 114671792 | 114672047 | 2.0853735 | TBX3 |
| 7 | 108573099 | 108573295 | 1.699689907 | DNAJB9 | 8 | 56966062 | 56966237 | -1.169282654 | IMPAD1 | 12 | 114683522 | 114683705 | 2.346511123 | TBX3 |
| 7 | 111420573 | 111420820 | -1.095475252 | IMMP2L | 1 | 235759304 | 235759516 | -1.39079135 | LYST | 12 | 115960933 | 115961180 | -1.129949744 | MED13L |
| 7 | 111421749 | 111421905 | -1.194201186 | IMMP2L | 8 | 57545475 | 57545688 | -3.481057964 | RP11-246K15.1 | 12 | 115991708 | 115991911 | -1.004041679 | MED13L |
| 1 | 229594352 | 229594549 | -1.097790266 | TAF5L | 8 | 57546710 | 57546974 | -1.918157134 | RP11-246K15.1 | 12 | 116008819 | 116009020 | -1.055495066 | MED13L |
| 7 | 116797959 | 116798156 | 6.999647694 | MET | 8 | 57568226 | 57568416 | -2.084652976 | RP11-246K15.1 | 12 | 116237586 | 116237800 | -1.235638925 | MED13L |
| 7 | 120273843 | 120274041 | -3.04637857 | KCND2 | 8 | 57569236 | 57569471 | -2.978161978 | RP11-246K15.1 | 12 | 118152310 | 118152517 | -1.322764839 | TAOK3 |
| 7 | 120274372 | 120274491 | -1.875806185 | KCND2 | 8 | 57572649 | 57572751 | -2.887824735 | RP11-246K15.1 | 12 | 118160180 | 118160419 | -1.08769063 | TAOK3 |
| 7 | 127347452 | 127347653 | 1.055196209 | ZNF800 | 8 | 57697364 | 57697565 | -7.978641728 | RP11-388G22.1 | 12 | 120095553 | 120095684 | 1.566075829 | RAB35 |
| 7 | 128028720 | 128028919 | -7.927263767 | LRRC4 | 1 | 235806031 | 235806256 | -1.508120091 | LYST | 12 | 120127535 | 120127676 | 1.803149833 | GCN1 |

| | | | | |
|---|---|---|---|---|
| 7 | 128028720 | 128028919 | -7.927263767 | SND1 |
| 1 | 229635191 | 229635393 | -2.171638628 | URB2 |
| 7 | 128457424 | 128457621 | -3.083579192 | RP11-155G14.6 |
| 7 | 128457424 | 128457621 | -3.083579192 | RP11-212P7.3 |
| 7 | 128457424 | 128457621 | -3.083579192 | HILPDA |
| 7 | 128656339 | 128656545 | -1.465674827 | CICP14 |
| 7 | 128767362 | 128767558 | 1.946936705 | CALU |
| 1 | 229636288 | 229636488 | -2.242430266 | URB2 |
| 7 | 128768965 | 128769167 | 1.665346966 | CALU |
| 7 | 128810120 | 128810291 | -1.954276822 | CCDC136 |
| 7 | 128870476 | 128870676 | 2.636102314 | KCP |
| 7 | 128870476 | 128870676 | 2.636102314 | RP11-309L24.4 |
| 7 | 128948755 | 128948921 | -1.466273329 | IRF5 |
| 7 | 128954975 | 128955173 | -1.116452389 | TNPO3 |
| 7 | 129056170 | 129056395 | -1.271035203 | TPI1P2 |
| 1 | 229636705 | 229636900 | -2.952538872 | URB2 |
| 7 | 130440552 | 130440746 | -2.202803846 | CEP41 |
| 7 | 130646044 | 130646242 | -1.508593013 | COPG2 |
| 7 | 131108443 | 131108601 | 3.443051498 | LINC-PINT |
| 1 | 229637939 | 229638155 | -2.578075632 | URB2 |
| 7 | 132128289 | 132128471 | 1.433636587 | PLXNA4 |
| 7 | 132367934 | 132368133 | -6.704428943 | AC011625.1 |
| 7 | 132367934 | 132368133 | -6.704428943 | PLXNA4 |
| 7 | 133034544 | 133034748 | -1.442144735 | RP11-371A22.1 |
| 7 | 133034544 | 133034748 | -1.442144735 | CHCHD3 |
| 7 | 134661429 | 134661615 | -1.363698443 | BPGM |
| 7 | 134994257 | 134994454 | 2.477435312 | AGBL3 |
| 7 | 135085485 | 135085685 | 2.211948517 | AGBL3 |
| 1 | 11107087 | 11107291 | -1.254484151 | MTOR |
| 7 | 135164506 | 135164702 | -1.508017155 | TMEM140 |
| 7 | 135164506 | 135164702 | -1.508017155 | C7orf49 |
| 7 | 135165458 | 135165658 | 10.49794291 | TMEM140 |
| 7 | 135165458 | 135165658 | 10.49794291 | C7orf49 |
| 7 | 135166385 | 135166645 | -1.072545126 | C7orf49 |
| 7 | 135169083 | 135169200 | -1.025043133 | C7orf49 |
| 7 | 135169083 | 135169200 | -1.025043133 | RP11-134L10.1 |
| 1 | 230323109 | 230323310 | 2.701268965 | PGBD5 |
| 7 | 135185900 | 135186113 | -1.664584574 | WDR91 |
| 7 | 137880406 | 137880572 | 1.318579356 | CREB3L2 |
| 7 | 138837526 | 138837715 | -2.156184062 | KIAA1549 |
| 7 | 138837526 | 138837715 | -2.156184062 | TMEM213 |
| 1 | 230706102 | 230706317 | -9.091749848 | AGT |
| 7 | 139572619 | 139572820 | 1.769372331 | HIPK2 |
| 7 | 139678248 | 139678431 | 2.024870521 | HIPK2 |
| 7 | 139716098 | 139716303 | 1.27547125 | HIPK2 |
| 7 | 140090879 | 140091080 | 5.570313849 | KDM7A |
| 1 | 230710089 | 230710288 | -2.577727871 | AGT |
| 7 | 140726205 | 140726398 | 1.413507873 | BRAF |
| 7 | 143269234 | 143269470 | -2.278815642 | AC073342.12 |
| 7 | 143269234 | 143269470 | -2.278815642 | GSTK1 |
| 7 | 143289293 | 143289506 | -1.287344103 | CASP2 |
| 7 | 143356657 | 143356858 | 4.518365559 | FAM131B |
| 7 | 143598206 | 143598406 | -2.383571092 | TCAF1P1 |
| 7 | 143876101 | 143876338 | -1.851446972 | TCAF1 |
| 8 | 60465067 | 60465219 | 2.79248125 | LINC01301 |
| 8 | 60741528 | 60741786 | -1.543707262 | CHD7 |
| 8 | 60742613 | 60742807 | -1.613120874 | CHD7 |
| 8 | 60760872 | 60761082 | -2.248465762 | CHD7 |
| 8 | 60781134 | 60781336 | -1.631803621 | CHD7 |
| 8 | 60853179 | 60853384 | -1.884655104 | CHD7 |
| 8 | 60856587 | 60856793 | -2.053288347 | CHD7 |
| 1 | 235808863 | 235809071 | -1.598634456 | LYST |
| 8 | 60860818 | 60860996 | -2.87807958 | CHD7 |
| 8 | 60865650 | 60865850 | -1.726653191 | CHD7 |
| 8 | 61579995 | 61580186 | 5.745926548 | ASPH |
| 8 | 61579995 | 61580186 | 5.745926548 | AC090094.1 |
| 8 | 64581828 | 64582054 | 2.922847204 | RP11-21C4.1 |
| 8 | 64581828 | 64582054 | 2.922847204 | BHLHE22 |
| 8 | 64801204 | 64801385 | -5.22766361 | RP11-1D12.2 |
| 1 | 235809998 | 235810196 | -1.972771462 | LYST |
| 8 | 66429852 | 66430086 | -1.046352302 | RRS1 |
| 8 | 67192871 | 67192973 | -2.716938969 | ARFGEF1 |
| 8 | 67192871 | 67192973 | -2.716938969 | CSPP1 |
| 8 | 67313471 | 67313622 | -4.330889049 | ARFGEF1 |
| 1 | 235962407 | 235962616 | -1.926039785 | RP4-580N22.2 |
| 8 | 71842628 | 71842775 | 1.870821265 | MSC |
| 8 | 71842628 | 71842775 | 1.870821265 | MSC-AS1 |
| 8 | 71879880 | 71880119 | 1.204038968 | MSC-AS1 |
| 8 | 71966701 | 71966887 | -7.554262228 | MSC-AS1 |
| 8 | 73055664 | 73055798 | -2.017323071 | RP11-531A24.5 |
| 8 | 73056589 | 73056769 | -1.341404912 | RP11-531A24.5 |
| 8 | 73057939 | 73058138 | -1.944546206 | RP11-531A24.5 |
| 8 | 73060970 | 73061169 | -1.592548182 | RP11-531A24.5 |
| 8 | 73787710 | 73787876 | -1.900224786 | UBE2W |
| 8 | 73787710 | 73787876 | -1.900224786 | RP11-463D19.2 |
| 8 | 73982146 | 73982350 | -1.282092071 | TMEM70 |
| 8 | 73982146 | 73982350 | -1.282092071 | RPS20P21 |
| 8 | 74236861 | 74237061 | -3.863960227 | JPH1 |
| 1 | 236195324 | 236195444 | 5.964814142 | GPR137B |
| 1 | 236482545 | 236482688 | 6.306434249 | EDARADD |
| 8 | 80486289 | 80486486 | 6.749175304 | ZBTB10 |
| 8 | 80487185 | 80487368 | 8.13084547 | ZBTB10 |
| 8 | 80975603 | 80975805 | -1.680495932 | PAG1 |
| 8 | 80976257 | 80976706 | -1.102097384 | PAG1 |
| 1 | 236483294 | 236483402 | 7.373178429 | ENO1P1 |
| 1 | 236483294 | 236483402 | 7.373178429 | EDARADD |
| 8 | 80985095 | 80985300 | -1.002981151 | PAG1 |
| 8 | 81279987 | 81280303 | 1.393462931 | RP11-363E6.3 |
| 8 | 81694603 | 81694851 | -1.915303059 | SLC10A5 |
| 8 | 84615937 | 84616147 | 1.682623198 | RALYL |
| 8 | 86481893 | 86482080 | -3.351086343 | RMDN1 |
| 8 | 86481893 | 86482080 | -3.351086343 | NTAN1P2 |
| 8 | 89687774 | 89687962 | 1.83774227 | RP11-37B2.1 |
| 8 | 89703075 | 89703256 | 1.629950078 | RP11-37B2.1 |
| 8 | 89757141 | 89757341 | 1.897120233 | RP11-37B2.1 |
| 8 | 89790086 | 89790288 | 1.748411008 | RIPK2 |
| 8 | 89953381 | 89953584 | 1.271610861 | NBN |
| 8 | 90005491 | 90005666 | -5.422745025 | DECR1 |
| 12 | 120303834 | 120304149 | 1.290208157 | SIRT4 |
| 12 | 120460167 | 120460311 | 1.022222453 | GATC |
| 12 | 120491129 | 120491326 | 1.414293897 | NRAV |
| 12 | 120491129 | 120491326 | 1.414293897 | DYNLL1 |
| 1 | 37482961 | 37483163 | 1.072581223 | ZC3H12A |
| 12 | 121020476 | 121020679 | -1.900739585 | OASL |
| 12 | 121185992 | 121186191 | -2.789391272 | P2RX7 |
| 12 | 121779242 | 121779463 | 1.890710034 | RHOF |
| 12 | 121779242 | 121779463 | 1.890710034 | TMEM120B |
| 12 | 122019712 | 122019859 | 1.913741064 | BCL7A |
| 12 | 122019712 | 122019859 | 1.913741064 | RP11-87C12.5 |
| 1 | 1318944 | 1319161 | 2.298427549 | CPSF3L |
| 12 | 122021136 | 122021282 | 1.772608161 | BCL7A |
| 12 | 122061443 | 122061658 | 1.004235038 | BCL7A |
| 12 | 122142314 | 122142512 | 1.026295543 | MLXIP |
| 12 | 122377784 | 122377995 | -1.180540353 | CLIP1 |
| 12 | 122701684 | 122701929 | -3.137536576 | HCAR2 |
| 12 | 122701684 | 122701929 | -3.137536576 | RP11-324E6.6 |
| 12 | 122850859 | 122851031 | -6.469605941 | HIP1R |
| 12 | 122960031 | 122960150 | 1.227576991 | ABCB9 |
| 12 | 122980181 | 122980326 | 1.181610117 | ABCB9 |
| 12 | 122980181 | 122980326 | 1.181610117 | ARL6IP4 |
| 12 | 122980181 | 122980326 | 1.181610117 | RP11-197N18.2 |
| 12 | 124324844 | 124325043 | 2.832384658 | NCOR2 |
| 12 | 124372202 | 124372522 | 9.982138702 | NCOR2 |
| 12 | 130973528 | 130973737 | 2.272160258 | ADGRD1 |
| 12 | 131089092 | 131089296 | 1.883419881 | ADGRD1 |
| 12 | 131943449 | 131943646 | 5.149604009 | PUS1 |
| 1 | 37990006 | 37990197 | -1.289065557 | SF3A3 |
| 12 | 132130438 | 132130655 | 1.298512059 | EP400NL |
| 12 | 132131238 | 132131433 | -1.006143187 | EP400NL |
| 1 | 38004596 | 38004840 | 5.101617085 | FHL3 |
| 12 | 132553114 | 132553299 | 6.372206705 | RP13-554M15.7 |
| 12 | 132553114 | 132553299 | 6.372206705 | FBRSL1 |
| 12 | 132623999 | 132624180 | 4.102825599 | POLE |
| 12 | 132773031 | 132773231 | 1.125848855 | GOLGA3 |
| 1 | 1327094 | 1327239 | 6.213656422 | CPTP |
| 13 | 19292813 | 19293046 | -2.67883564 | ANKRD26P3 |
| 13 | 19346161 | 19346361 | -6.215226276 | LINC00421 |
| 13 | 19564368 | 19564515 | -4.708926257 | MRPS31P2 |
| 13 | 19564368 | 19564515 | -4.708926257 | CASC4P1 |
| 13 | 20608076 | 20608249 | 1.027247355 | SLC35E1P1 |
| 13 | 20608076 | 20608249 | 1.027247355 | IFT88 |
| 13 | 20722933 | 20723068 | 1.096851192 | IL17D |
| 13 | 20773527 | 20773722 | 4.113492766 | EEF1AKMT1 |
| 13 | 20981570 | 20981750 | 1.219281213 | LATS2 |
| 13 | 20988090 | 20988287 | 1.910652859 | LATS2 |
| 13 | 20988541 | 20988713 | 8.550269576 | LATS2 |
| 13 | 23324548 | 23324743 | 1.031367878 | SGCG |
| 13 | 23335479 | 23335679 | 1.201567108 | SACS |
| 13 | 27430790 | 27431005 | -1.95888266 | GTF3A |
| 13 | 28302569 | 28302765 | 2.026691451 | FLT1 |
| 13 | 28389863 | 28390064 | 1.208646413 | FLT1 |
| 1 | 39358574 | 39358774 | -1.022901845 | MACF1 |

| | | | | |
|---|---|---|---|---|
| 7 | 144187511 | 144187897 | -1.087154085 | ARHGEF35 |
| 1 | 231000179 | 231000370 | -1.012759607 | ARV1 |
| 7 | 144365178 | 144365397 | -1.131360182 | ARHGEF5 |
| 7 | 144648261 | 144648471 | -1.053444567 | EEF1A1P10 |
| 7 | 144648261 | 144648471 | -1.053444567 | TPK1 |
| 7 | 144753787 | 144753907 | 4.027641218 | TPK1 |
| 7 | 148697876 | 148698063 | -1.317962239 | CUL1 |
| 7 | 148697876 | 148698063 | -1.317962239 | RP5-1136G13.2 |
| 7 | 148800596 | 148800840 | -1.073822404 | CUL1 |
| 7 | 149003793 | 149003992 | -1.844031894 | PDIA4 |
| 7 | 149145612 | 149145811 | -1.828633701 | ZNF398 |
| 1 | 231267817 | 231268007 | -1.747273806 | GNPAT |
| 7 | 149224576 | 149224939 | -1.754964937 | ZNF282 |
| 7 | 149225829 | 149226024 | -2.082603226 | ZNF282 |
| 7 | 149255005 | 149255153 | -1.108165301 | ZNF212 |
| 7 | 149297022 | 149297189 | 2.163302291 | ZNF783 |
| 7 | 149297022 | 149297189 | 2.163302291 | RP4-800G7.2 |
| 7 | 149733429 | 149733616 | -2.372611347 | KRBA1 |
| 7 | 149850037 | 149850221 | 2.872151376 | ZNF862 |
| 7 | 150372835 | 150373034 | -1.038983856 | ZNF775 |
| 7 | 150372835 | 150373034 | -1.038983856 | REPIN1 |
| 7 | 150396525 | 150396747 | -1.380688689 | ZNF775 |
| 7 | 150398070 | 150398234 | -1.563160808 | ZNF775 |
| 7 | 150477229 | 150477547 | 3.174364077 | GIMAP8 |
| 7 | 150720325 | 150720560 | 2.764199854 | GIMAP1-GIMAP5 |
| 7 | 150720325 | 150720560 | 2.764199854 | GIMAP1 |
| 7 | 151046211 | 151046424 | 1.137941175 | ABCB8 |
| 7 | 151104764 | 151104950 | 3.457939689 | AGAP3 |
| 7 | 151213317 | 151213496 | -2.848024047 | ABCF2 |
| 1 | 11295110 | 11295307 | -2.361029254 | UBIAD1 |
| 1 | 231421091 | 231421345 | -2.482865016 | EGLN1 |
| 7 | 151378953 | 151379132 | 3.947408882 | WDR86 |
| 7 | 151558559 | 151558737 | 1.27419961 | PRKAG2 |
| 7 | 151559124 | 151559331 | 1.548543388 | PRKAG2 |
| 7 | 151859747 | 151859919 | -1.495803867 | PRKAG2 |
| 7 | 152646892 | 152647074 | -3.799933893 | XRCC2 |
| 7 | 152647390 | 152647631 | -3.589010753 | XRCC2 |
| 7 | 152648805 | 152649014 | -4.225991625 | XRCC2 |
| 7 | 154950182 | 154950383 | 1.38343541 | PAXIP1 |
| 7 | 154968960 | 154969170 | -2.077622618 | PAXIP1 |
| 7 | 154975792 | 154975988 | -2.295287879 | PAXIP1 |
| 7 | 156950342 | 156950604 | -2.744113194 | NOM1 |
| 7 | 156959924 | 156960103 | -1.764415276 | NOM1 |
| 7 | 156996710 | 156996871 | -7.181108908 | MNX1 |
| 7 | 157002171 | 157002301 | -1.979332123 | MNX1 |
| 7 | 157005144 | 157005268 | -2.964129728 | MNX1 |
| 7 | 157009323 | 157009502 | -4.898461174 | MNX1 |
| 1 | 232465147 | 232465346 | 3.56592848 | SIPA1L2 |
| 7 | 157110167 | 157110373 | 4.257932035 | RP4-639J15.3 |
| 7 | 157498506 | 157498669 | -5.636361542 | AC006372.4 |
| 7 | 157539548 | 157539856 | -2.22196946 | PTPRN2 |
| 7 | 157546453 | 157546685 | -1.746863951 | PTPRN2 |
| 7 | 157583133 | 157583239 | -3.56587864 | PTPRN2 |
| 1 | 232514130 | 232514333 | 2.297218725 | SIPA1L2 |

| | | | | |
|---|---|---|---|---|
| 8 | 91069308 | 91069499 | -1.128581818 | OTUD6B-AS1 |
| 1 | 236547451 | 236547649 | -1.147144707 | LGALS8 |
| 1 | 236547451 | 236547649 | -1.147144707 | RP11-385F5.4 |
| 8 | 91069532 | 91069645 | -1.374797343 | OTUD6B-AS1 |
| 8 | 93922417 | 93922664 | 2.528293481 | PDP1 |
| 8 | 94432295 | 94432483 | -5.657574781 | RAD54B |
| 8 | 94432295 | 94432483 | -5.657574781 | FSBP |
| 8 | 94511412 | 94511609 | -2.275521346 | KIAA1429 |
| 8 | 94930579 | 94930815 | -1.477284692 | TP53INP1 |
| 8 | 94930579 | 94930815 | -1.477284692 | NDUFAF6 |
| 8 | 95247521 | 95247693 | -6.832667959 | C8orf37 |
| 8 | 95247521 | 95247693 | -6.832667959 | C8orf37-AS1 |
| 8 | 96333532 | 96333734 | -1.094917223 | PTDSS1 |
| 1 | 236820114 | 236820292 | -1.558800606 | MTR |
| 1 | 236820114 | 236820292 | -1.558800606 | RPSAP21 |
| 8 | 96742215 | 96742429 | -6.405787694 | CPQ |
| 8 | 96930670 | 96930770 | 5.33974005 | CPQ |
| 8 | 96955852 | 96956040 | -4.418971621 | CPQ |
| 8 | 96972109 | 96972303 | -7.110264741 | CPQ |
| 8 | 96974133 | 96974304 | -4.988639962 | CPQ |
| 8 | 96976967 | 96977141 | -7.142556594 | CPQ |
| 8 | 97092642 | 97092834 | -4.114890083 | CPQ |
| 8 | 97146872 | 97147087 | -6.204287351 | CPQ |
| 8 | 97724699 | 97724899 | -1.160617734 | MTDH |
| 8 | 98501149 | 98501337 | -1.09053278 | STK3 |
| 8 | 99066462 | 99066658 | 7.873558884 | VPS13B |
| 1 | 236899066 | 236899261 | -1.269591247 | MTR |
| 8 | 99293547 | 99293730 | -5.663214744 | VPS13B |
| 8 | 99425369 | 99425510 | 2.611885945 | VPS13B |
| 8 | 99741015 | 99741183 | -1.753784759 | VPS13B |
| 8 | 100258984 | 100259192 | 1.7371232 | RNF19A |
| 8 | 100258984 | 100259192 | 1.7371232 | SPAG1 |
| 8 | 100287598 | 100287793 | 1.896793534 | RNF19A |
| 1 | 236903528 | 236903727 | -3.714193092 | MTR |
| 8 | 100293796 | 100294149 | 2.448651174 | RNF19A |
| 8 | 100310458 | 100310717 | 1.281151763 | RNF19A |
| 8 | 100312144 | 100312390 | 1.705148718 | RNF19A |
| 8 | 100344322 | 100344578 | 1.390155108 | KB-1991G8.1 |
| 8 | 100346500 | 100346703 | 1.856548102 | KB-1991G8.1 |
| 8 | 100549927 | 100550125 | 1.325190465 | ANKRD46 |
| 8 | 101177869 | 101178073 | -1.705435554 | ZNF706 |
| 8 | 101719500 | 101719687 | -3.575430724 | NCALD |
| 8 | 101825111 | 101825317 | -7.50098994 | NCALD |
| 8 | 102650204 | 102650407 | -1.368978214 | KLF10 |
| 8 | 102651667 | 102651867 | -1.208844883 | KLF10 |
| 8 | 103072724 | 103072931 | -1.295120017 | ATP6V1C1 |
| 8 | 103132702 | 103132856 | 2.307779119 | LINC01181 |
| 8 | 103134868 | 103135076 | 2.645293936 | BAALC-AS2 |
| 8 | 103267033 | 103267227 | -1.563726349 | BAALC-AS1 |
| 8 | 103331271 | 103331464 | -1.506990014 | FZD6 |
| 1 | 238499412 | 238499654 | -6.987476137 | RP11-177F15.1 |
| 8 | 103885737 | 103885937 | 1.335849171 | RIMS2 |
| 8 | 104359840 | 104360047 | -1.049528808 | DPYS |
| 8 | 104380963 | 104381162 | 2.862906026 | DPYS |

| | | | | |
|---|---|---|---|---|
| 13 | 30658029 | 30658254 | -1.264915839 | USPL1 |
| 13 | 32332971 | 32333208 | 1.129470484 | BRCA2 |
| 13 | 32403002 | 32403203 | -1.542920702 | N4BP2L1 |
| 13 | 33105510 | 33105729 | -1.528289248 | STARD13 |
| 13 | 33129453 | 33129668 | -1.560488209 | STARD13 |
| 13 | 33947734 | 33947915 | -5.310568057 | RFC3 |
| 13 | 36819591 | 36819788 | 8.668257638 | RFXAP |
| 1 | 1328427 | 1328623 | 1.654869712 | CPTP |
| 13 | 40865558 | 40865729 | -3.375213927 | TPTE2P5 |
| 13 | 41043285 | 41043419 | -2.162853993 | ELF1 |
| 13 | 41046052 | 41046257 | -1.732395267 | ELF1 |
| 13 | 41371217 | 41371418 | -2.344501546 | NAA16 |
| 13 | 41618343 | 41618539 | 4.999295215 | VWA8 |
| 13 | 42040017 | 42040137 | 1.145470201 | DGKH |
| 13 | 42041456 | 42041608 | 2.001957525 | DGKH |
| 13 | 42048079 | 42048277 | -6.310568057 | DGKH |
| 13 | 42301891 | 42302085 | -2.020513634 | AKAP11 |
| 13 | 43971247 | 43971450 | 1.317586973 | RP11-65B7.2 |
| 13 | 43971247 | 43971450 | 1.317586973 | LINC00284 |
| 13 | 44434483 | 44434788 | 1.788485102 | TSC22D1 |
| 13 | 46750694 | 46750895 | -1.778735662 | LRCH1 |
| 13 | 48395405 | 48395612 | -2.243945212 | RB1 |
| 13 | 48395405 | 48395612 | -2.243945212 | LPAR6 |
| 13 | 48396446 | 48396634 | -4.011883687 | LPAR6 |
| 13 | 48396446 | 48396634 | -4.011883687 | RB1 |
| 13 | 48397192 | 48397392 | -2.43686567 | LPAR6 |
| 13 | 48397192 | 48397392 | -2.43686567 | RB1 |
| 13 | 48418527 | 48418683 | -1.147853898 | RB1 |
| 13 | 48418527 | 48418683 | -1.147853898 | LPAR6 |
| 13 | 49274119 | 49274272 | 4.794357318 | CDADC1 |
| 1 | 39524933 | 39525090 | -6.775313278 | BMP8A |
| 1 | 39524933 | 39525090 | -6.775313278 | RP11-69E11.4 |
| 13 | 50807750 | 50807943 | -7.174087434 | DLEU7-AS1 |
| 13 | 50807750 | 50807943 | -7.174087434 | DLEU7 |
| 13 | 50834010 | 50834178 | -5.925874521 | DLEU7-AS1 |
| 13 | 50834010 | 50834178 | -5.925874521 | DLEU7 |
| 13 | 50867621 | 50867780 | -4.089954545 | RNASEH2B-AS1 |
| 13 | 50906523 | 50906664 | -1.118007096 | RNASEH2B-AS1 |
| 13 | 57634846 | 57634997 | 4.767637688 | PCDH17 |
| 13 | 57725439 | 57725612 | 3.54637857 | PCDH17 |
| 13 | 61413235 | 61413428 | 3.318154611 | RP11-310K10.1 |
| 13 | 61413235 | 61413428 | 3.318154611 | PCDH20 |
| 13 | 71681840 | 71682045 | 7.803925211 | DACH1 |
| 1 | 1333701 | 1333879 | 7.940054807 | TAS1R3 |
| 13 | 73062278 | 73062492 | 1.583247548 | KLF5 |
| 13 | 73846143 | 73846354 | 1.218820597 | KLF12 |
| 13 | 76884861 | 76885063 | 1.00695586 | KCTD12 |
| 13 | 76991728 | 76991929 | 1.176292464 | CLN5 |
| 13 | 78600866 | 78601051 | 1.120754706 | POU4F1 |
| 13 | 78600866 | 78601051 | 1.120754706 | RNF219-AS1 |
| 13 | 78601441 | 78601646 | 1.753719198 | RNF219-AS1 |
| 13 | 78601441 | 78601646 | 1.753719198 | POU4F1 |
| 13 | 78602400 | 78602589 | 6.563513522 | POU4F1 |
| 13 | 78602400 | 78602589 | 6.563513522 | RNF219-AS1 |

| | | | | |
|---|---|---|---|---|
| 7 | 157586688 | 157586880 | -4.020957888 | PTPRN2 |
| 7 | 157587697 | 157587840 | -1.912414944 | PTPRN2 |
| 7 | 157588361 | 157588588 | -3.832027784 | PTPRN2 |
| 7 | 157590106 | 157590260 | -7.081982046 | PTPRN2 |
| 7 | 157590814 | 157591013 | -2.112318882 | PTPRN2 |
| 7 | 157598555 | 157598746 | -4.394496356 | PTPRN2 |
| 7 | 157605917 | 157606105 | -1.621741451 | PTPRN2 |
| 7 | 157610328 | 157610518 | -2.987604177 | PTPRN2 |
| 7 | 157628510 | 157628667 | -2.540694163 | PTPRN2 |
| 7 | 157650104 | 157650303 | -9.400209831 | PTPRN2 |
| 1 | 232514834 | 232515030 | 7.916568583 | SIPA1L2 |
| 7 | 157651231 | 157651423 | -2.797668853 | PTPRN2 |
| 7 | 157651837 | 157651995 | -2.929989169 | PTPRN2 |
| 7 | 157653095 | 157653328 | -2.572365478 | PTPRN2 |
| 7 | 157857806 | 157858002 | -2.26021602 | AC011899.9 |
| 7 | 157857806 | 157858002 | -2.26021602 | PTPRN2 |
| 7 | 157862067 | 157862228 | -3.100816931 | PTPRN2 |
| 7 | 157862067 | 157862228 | -3.100816931 | AC011899.9 |
| 7 | 157862854 | 157863057 | -1.252464655 | PTPRN2 |
| 7 | 157862854 | 157863057 | -1.252464655 | AC011899.9 |
| 7 | 157863798 | 157863940 | -1.945142647 | PTPRN2 |
| 7 | 157863798 | 157863940 | -1.945142647 | AC011899.9 |
| 7 | 157867585 | 157867778 | -1.833469901 | PTPRN2 |
| 1 | 232805914 | 232806151 | -1.670415656 | MAP10 |
| 7 | 158791142 | 158791330 | 3.272840181 | ESYT2 |
| 1 | 232806936 | 232807120 | -1.78827434 | MAP10 |
| 8 | 673456 | 673668 | -1.16107659 | ERICH1 |
| 8 | 710331 | 710546 | -9.430505579 | ERICH1 |
| 8 | 714953 | 715133 | -9.94841748 | ERICH1 |
| 8 | 1761479 | 1761691 | 1.826522661 | CLN8 |
| 8 | 1770922 | 1771136 | 1.676801188 | RP11-439C15.6 |
| 8 | 1770922 | 1771136 | 1.676801188 | CLN8 |
| 8 | 1780362 | 1780566 | 1.239788791 | CLN8 |
| 8 | 1780362 | 1780566 | 1.239788791 | RP11-439C15.6 |
| 8 | 1944606 | 1944731 | -2.634355885 | RP11-439C15.6 |
| 8 | 1944606 | 1944731 | -2.634355885 | ARHGEF10 |
| 1 | 11295782 | 11295909 | -1.769389725 | UBIAD1 |
| 1 | 232807779 | 232807910 | -1.668869948 | MAP10 |
| 8 | 1948182 | 1948453 | -3.058526364 | ARHGEF10 |
| 8 | 1948182 | 1948453 | -3.058526364 | RP11-439C15.6 |
| 8 | 2000542 | 2000747 | -1.170873054 | KBTBD11 |
| 8 | 2003183 | 2003384 | -1.643293529 | KBTBD11 |
| 8 | 2006525 | 2006744 | -1.511893126 | KBTBD11 |
| 8 | 6405905 | 6406037 | -1.892680822 | RP11-115C21.2 |
| 8 | 6445238 | 6445482 | -1.696173437 | MCPH1 |
| 8 | 6618978 | 6619151 | 6.378278161 | MCPH1-AS1 |
| 8 | 6618978 | 6619151 | 6.378278161 | MCPH1 |
| 1 | 232976556 | 232976709 | 6.495583753 | NTPCR |
| 8 | 8235608 | 8235806 | -2.538558672 | FAM86B3P |
| 8 | 8236087 | 8236285 | -1.836914123 | FAM86B3P |
| 8 | 8236087 | 8236285 | -1.836914123 | ALG1L13P |
| 8 | 8323717 | 8323864 | -3.169504056 | SGK223 |
| 8 | 8339545 | 8339734 | 5.257110455 | SGK223 |
| 8 | 8376405 | 8376583 | 1.282141679 | SGK223 |
| 1 | 238535838 | 238536028 | -6.142701109 | RP11-177F15.1 |
| 8 | 105789362 | 105789563 | -1.420671315 | ZFPM2 |
| 8 | 105789362 | 105789563 | -1.420671315 | ZFPM2-AS1 |
| 8 | 105792065 | 105792263 | -1.316949756 | ZFPM2 |
| 8 | 105792065 | 105792263 | -1.316949756 | ZFPM2-AS1 |
| 8 | 105792857 | 105793057 | -1.407651537 | ZFPM2 |
| 8 | 105792857 | 105793057 | -1.407651537 | ZFPM2-AS1 |
| 8 | 105794430 | 105794648 | -1.604620187 | ZFPM2 |
| 8 | 105794430 | 105794648 | -1.604620187 | ZFPM2-AS1 |
| 8 | 105796901 | 105797071 | -2.182330796 | ZFPM2 |
| 8 | 105796901 | 105797071 | -2.182330796 | ZFPM2-AS1 |
| 8 | 106272387 | 106272651 | -1.488411007 | RP11-395G23.3 |
| 1 | 239908098 | 239908278 | -2.737982492 | CHRM3 |
| 8 | 106565950 | 106566120 | -1.704338667 | RP11-649G15.2 |
| 8 | 106565950 | 106566120 | -1.704338667 | OXR1 |
| 8 | 106590629 | 106590828 | -2.613085661 | OXR1 |
| 8 | 106590629 | 106590828 | -2.613085661 | RP11-649G15.2 |
| 8 | 106657455 | 106657600 | 3.021212733 | OXR1 |
| 8 | 106657455 | 106657600 | 3.021212733 | RP11-649G15.2 |
| 1 | 239909023 | 239909222 | -3.65562526 | CHRM3 |
| 8 | 109334042 | 109334230 | 8.319875253 | NUDCD1 |
| 1 | 241593806 | 241594069 | -2.361879494 | KMO |
| 1 | 241593806 | 241594069 | -2.361879494 | OPN3 |
| 8 | 116875665 | 116875839 | -3.118246309 | RAD21-AS1 |
| 8 | 120477216 | 120477382 | -6.013952998 | MTBP |
| 8 | 120538538 | 120538740 | -1.749137749 | MTBP |
| 8 | 120538538 | 120538740 | -1.749137749 | SNTB1 |
| 1 | 241634115 | 241634321 | -1.769842656 | CHML |
| 1 | 241634115 | 241634321 | -1.769842656 | OPN3 |
| 8 | 122877624 | 122877761 | 6.909590589 | ZHX2 |
| 8 | 122951436 | 122951640 | -1.338651336 | ZHX2 |
| 8 | 122953646 | 122953846 | -1.348914452 | ZHX2 |
| 8 | 123122991 | 123123195 | -3.54411018 | TBC1D31 |
| 8 | 123122991 | 123123195 | -3.54411018 | HMGB1P19 |
| 8 | 123250201 | 123250333 | -1.024416592 | ZHX1-C8orf76 |
| 8 | 123250201 | 123250333 | -1.024416592 | ZHX1 |
| 1 | 241635521 | 241635690 | -2.511614937 | OPN3 |
| 1 | 241635521 | 241635690 | -2.511614937 | CHML |
| 8 | 124451274 | 124451496 | -1.882455778 | TRMT12 |
| 8 | 124452529 | 124452731 | -1.129922346 | TRMT12 |
| 8 | 124487262 | 124487492 | -1.557178517 | RNF139 |
| 8 | 124553103 | 124553316 | -1.225793574 | MTSS1 |
| 8 | 124553103 | 124553316 | -1.225793574 | NDUFB9 |
| 8 | 124832400 | 124832571 | -3.594088353 | RP11-1082L8.3 |
| 1 | 241878761 | 241878951 | -4.962406252 | EXO1 |
| 8 | 124999190 | 124999423 | -1.369136285 | SQLE |
| 8 | 125027353 | 125027549 | -1.54898758 | KIAA0196 |
| 8 | 127890673 | 127890852 | -1.6730176 | PVT1 |
| 8 | 127890673 | 127890852 | -1.6730176 | PVT1_3 |
| 8 | 129548322 | 129548523 | -3.22384508 | CCDC26 |
| 1 | 243126562 | 243126750 | -1.316075925 | CEP170 |
| 8 | 132123505 | 132123676 | 1.245926548 | KCNQ3 |
| 8 | 132988851 | 132989019 | 2.668802692 | TG |
| 8 | 133458573 | 133458768 | -1.510555009 | ST3GAL1 |
| 13 | 80340733 | 80340856 | 4.057220335 | SPRY2 |
| 13 | 99255320 | 99255502 | -6.20918986 | GPR18 |
| 13 | 99255320 | 99255502 | -6.20918986 | UBAC2 |
| 13 | 99295643 | 99295972 | 2.993229076 | UBAC2 |
| 13 | 99295643 | 99295972 | 2.993229076 | GPR183 |
| 1 | 39770956 | 39771152 | 7.070095352 | BMP8B |
| 1 | 39770956 | 39771152 | 7.070095352 | OXCT2 |
| 13 | 99982584 | 99982839 | 1.04059379 | ZIC2 |
| 13 | 100358000 | 100358185 | -6.526284025 | PCCA |
| 13 | 100540327 | 100540538 | -1.40638532 | RP11-151A6.4 |
| 13 | 100540327 | 100540538 | -1.40638532 | GGACT |
| 13 | 102772992 | 102773165 | 6.054588852 | TEX30 |
| 13 | 106492813 | 106492993 | 1.985468767 | EFNB2 |
| 13 | 109782253 | 109782446 | 11.2885837 | IRS2 |
| 13 | 109784244 | 109784442 | 12.82787978 | IRS2 |
| 13 | 109785219 | 109785420 | 11.41269797 | IRS2 |
| 13 | 110614133 | 110614303 | 2.307354922 | RP11-90L1.8 |
| 13 | 110659909 | 110660185 | 1.11774668 | CARS2 |
| 13 | 110879752 | 110879908 | -1.103022394 | ANKRD10 |
| 1 | 39869548 | 39869745 | 4.42648146 | TRIT1 |
| 13 | 112687308 | 112687487 | 1.465661428 | RP11-88E10.5 |
| 13 | 112807226 | 112807434 | 1.960698083 | ATP11A |
| 1 | 39869852 | 39870046 | 11.59574704 | TRIT1 |
| 13 | 113252438 | 113252597 | 7.677124691 | CUL4A |
| 13 | 113403433 | 113403645 | 2.040971649 | ADPRHL1 |
| 13 | 113405638 | 113405835 | 1.83149158 | ADPRHL1 |
| 13 | 113406400 | 113406674 | 1.907283859 | ADPRHL1 |
| 13 | 113407792 | 113407967 | 7.700639768 | ADPRHL1 |
| 13 | 113413737 | 113413931 | 1.134682748 | ADPRHL1 |
| 13 | 113441748 | 113441877 | -1.564821748 | ADPRHL1 |
| 13 | 113453467 | 113453626 | 1.258694094 | ADPRHL1 |
| 13 | 113498233 | 113498422 | -1.123980195 | TMCO3 |
| 13 | 113816679 | 113816812 | 1.78967596 | GAS6-AS1 |
| 13 | 113816679 | 113816812 | 1.78967596 | TMEM255B |
| 1 | 39901403 | 39901540 | 2.937529967 | MYCL |
| 13 | 113817848 | 113818102 | 2.341319216 | GAS6-AS1 |
| 13 | 113818813 | 113818947 | -5.372416919 | GAS6-AS1 |
| 13 | 113832019 | 113832212 | -5.919995535 | GAS6 |
| 13 | 113832019 | 113832212 | -5.919995535 | GAS6-AS1 |
| 13 | 113842829 | 113843020 | -6.965091458 | GAS6 |
| 13 | 113842829 | 113843020 | -6.965091458 | GAS6-AS1 |
| 13 | 113844092 | 113844420 | -1.727695227 | GAS6 |
| 13 | 113844092 | 113844420 | -1.727695227 | GAS6-AS1 |
| 13 | 113844754 | 113844858 | -7.221471748 | GAS6-AS1 |
| 13 | 113844754 | 113844858 | -7.221471748 | GAS6 |
| 1 | 39962665 | 39962862 | 1.297444385 | RP3-342P20.2 |
| 1 | 39962665 | 39962862 | 1.297444385 | MFSD2A |
| 13 | 113954447 | 113954642 | -1.158630285 | ABC13-47488600E17.1 |
| 13 | 113956509 | 113956703 | -1.803348413 | ABC13-47488600E17.1 |
| 13 | 114058727 | 114058909 | 3.142701109 | RASA3 |
| 13 | 114059932 | 114060119 | 1.646594282 | RASA3 |
| 1 | 39963123 | 39963323 | 9.198469598 | MFSD2A |
| 1 | 39963123 | 39963323 | 9.198469598 | RP3-342P20.2 |
| 13 | 114062758 | 114062909 | 5.877861077 | RASA3 |

| | | | | |
|---|---|---|---|---|
| 8 | 8377503 | 8377703 | 1.587304388 | SGK223 |
| 1 | 232983449 | 232983592 | -1.954962328 | NTPCR |
| 1 | 232983449 | 232983592 | -1.954962328 | PCNX2 |
| 8 | 8381517 | 8381714 | 6.345435505 | SGK223 |
| 8 | 8703230 | 8703426 | 1.631934529 | CLDN23 |
| 1 | 233258262 | 233258611 | 1.18626808 | PCNX2 |
| 8 | 10765441 | 10765642 | -2.075882108 | CTD-2135J3.3 |
| 8 | 10765441 | 10765642 | -2.075882108 | PINX1 |
| 8 | 10765441 | 10765642 | -2.075882108 | CTD-2135J3.4 |
| 8 | 11030224 | 11030402 | -6.681862059 | XKR6 |
| 8 | 11107905 | 11108084 | -2.418742294 | XKR6 |
| 8 | 11107905 | 11108084 | -2.418742294 | AF131215.2 |
| 8 | 11118862 | 11119058 | -1.322716782 | XKR6 |
| 8 | 11183380 | 11183557 | 1.297103172 | XKR6 |
| 8 | 11844826 | 11845025 | -7.267789573 | CTSB |
| 1 | 233344667 | 233344868 | -1.81101556 | RP5-862P8.2 |
| 8 | 17645692 | 17645893 | 4.696585751 | MTUS1 |
| 8 | 17754020 | 17754211 | 5.148629003 | MTUS1 |
| 8 | 17754998 | 17755104 | 4.60451474 | MTUS1 |
| 8 | 17989005 | 17989225 | 1.376778539 | PCM1 |
| 8 | 18020952 | 18021095 | -6.366083713 | PCM1 |
| 8 | 18528321 | 18528573 | 3.874469118 | PSD3 |
| 8 | 18872261 | 18872442 | 1.479332773 | PSD3 |
| 1 | 233379412 | 233379611 | -1.714389058 | RP5-862P8.2 |
| 8 | 19319400 | 19319597 | -2.683178529 | SH2D4A |
| 8 | 19395498 | 19395675 | -3.089308281 | SH2D4A |
| 8 | 19405737 | 19405937 | 1.208306505 | CSGALNACT1 |
| 8 | 20246739 | 20246947 | 1.056425191 | LZTS1 |
| 8 | 20253259 | 20253394 | 1.818121503 | LZTS1 |
| 8 | 21909259 | 21909519 | -2.836901139 | DOK2 |
| 8 | 22107279 | 22107497 | -1.694148676 | NUDT18 |
| 8 | 22107730 | 22108011 | -1.59934036 | NUDT18 |
| 1 | 233382426 | 233382632 | -1.868418037 | RP5-862P8.2 |
| 8 | 22419631 | 22419855 | -1.19805276 | SLC39A14 |
| 8 | 22613997 | 22614220 | 1.528107974 | RP11-582J16.5 |
| 8 | 22613997 | 22614220 | 1.528107974 | CCAR2 |
| 8 | 22635297 | 22635494 | 1.344779549 | BIN3 |
| 8 | 22690080 | 22690304 | 3.308341395 | EGR3 |
| 8 | 22690080 | 22690304 | 3.308341395 | RP11-459E5.1 |
| 8 | 22690905 | 22691100 | 3.93598021 | RP11-459E5.1 |
| 8 | 22690905 | 22691100 | 3.93598021 | EGR3 |
| 8 | 23018249 | 23018449 | 1.02210026 | RHOBTB2 |
| 8 | 23018249 | 23018449 | 1.02210026 | RP11-875O11.1 |
| 8 | 23022679 | 23022879 | 1.036853424 | TNFRSF10B |
| 8 | 23116731 | 23116894 | 1.918551133 | TNFRSF10C |
| 8 | 23137702 | 23137906 | 2.235229642 | TNFRSF10D |
| 8 | 23425001 | 23425223 | -1.773553191 | ENTPD4 |
| 8 | 23425001 | 23425223 | -1.773553191 | LOXL2 |
| 8 | 23432524 | 23432867 | -1.157253741 | ENTPD4 |
| 8 | 23482557 | 23482732 | -2.783787934 | CTC-756D1.2 |
| 8 | 23566666 | 23566867 | -1.417444344 | SLC25A37 |
| 1 | 11790324 | 11790527 | 1.452097863 | MTHFR |
| 1 | 234607364 | 234607569 | -1.114941088 | IRF2BP2 |
| 1 | 234607364 | 234607569 | -1.114941088 | RP4-781K5.2 |

| | | | | |
|---|---|---|---|---|
| 8 | 133459359 | 133459560 | -1.578238284 | ST3GAL1 |
| 8 | 133476307 | 133476527 | -1.355122605 | ST3GAL1 |
| 8 | 134573644 | 134573775 | 3.336212671 | ZFAT |
| 8 | 139742282 | 139742467 | 5.426654778 | TRAPPC9 |
| 8 | 139775788 | 139775971 | 1.436355479 | TRAPPC9 |
| 8 | 139796981 | 139797180 | -6.823448085 | TRAPPC9 |
| 8 | 139851817 | 139851927 | -1.877443751 | TRAPPC9 |
| 1 | 243165775 | 243165975 | -1.12967717 | RP11-261C10.4 |
| 1 | 243165775 | 243165975 | -1.12967717 | CEP170 |
| 8 | 140097412 | 140097606 | -1.669400957 | TRAPPC9 |
| 8 | 140097412 | 140097606 | -1.669400957 | PEG13 |
| 8 | 140530650 | 140530824 | -8.166035762 | AGO2 |
| 8 | 140636749 | 140636961 | 1.26672557 | ERICD |
| 8 | 140659424 | 140659559 | -2.066699563 | PTK2 |
| 1 | 243185914 | 243186113 | -1.234358039 | CEP170 |
| 8 | 141127984 | 141128183 | 2.722821495 | RP11-809O17.1 |
| 8 | 141127984 | 141128183 | 2.722821495 | DENND3 |
| 8 | 141129013 | 141129237 | 8.806851574 | RP11-809O17.1 |
| 8 | 141129013 | 141129237 | 8.806851574 | DENND3 |
| 8 | 141129525 | 141129696 | 9.150006435 | DENND3 |
| 8 | 141129525 | 141129696 | 9.150006435 | RP11-809O17.1 |
| 8 | 141193974 | 141194167 | -1.014636598 | DENND3 |
| 1 | 243191164 | 243191369 | -1.404661179 | CEP170 |
| 8 | 142271015 | 142271216 | -1.71772835 | TSNARE1 |
| 8 | 142271602 | 142271718 | -1.264402661 | TSNARE1 |
| 8 | 142273138 | 142273345 | -1.847677283 | TSNARE1 |
| 8 | 142308438 | 142308611 | 1.813391338 | TSNARE1 |
| 8 | 142663665 | 142663895 | -1.517530952 | JRK |
| 1 | 243303055 | 243303239 | -6.934652329 | SDCCAG8 |
| 8 | 142664981 | 142665178 | -1.330874431 | JRK |
| 8 | 142666084 | 142666311 | -1.089124083 | JRK |
| 8 | 143008964 | 143009151 | 3.031504899 | RP11-273G15.2 |
| 1 | 243318113 | 243318318 | -1.399916373 | SDCCAG8 |
| 8 | 143306303 | 143306466 | -1.137566942 | TOP1MT |
| 8 | 143367845 | 143367994 | -3.857634171 | RHPN1-AS1 |
| 8 | 143544496 | 143544663 | -1.199543586 | RP11-661A12.5 |
| 8 | 143559764 | 143559964 | 1.930182767 | GSDMD |
| 8 | 143560248 | 143560447 | 2.153450564 | GSDMD |
| 8 | 143587928 | 143588087 | 7.946366266 | EEF1D |
| 8 | 143589780 | 143589979 | 1.910905871 | EEF1D |
| 8 | 143596800 | 143597035 | 2.044560215 | EEF1D |
| 1 | 243685247 | 243685380 | -2.196158711 | AKT3 |
| 8 | 143744738 | 143744856 | -2.068751762 | FAM83H-AS1 |
| 8 | 143922948 | 143923148 | 10.13063428 | PLEC |
| 8 | 143983681 | 143983917 | 1.927335275 | PARP10 |
| 1 | 243730449 | 243730649 | -7.747334056 | RP11-370K11.1 |
| 1 | 243730449 | 243730649 | -7.747334056 | AKT3 |
| 8 | 144102666 | 144102839 | -1.043584174 | SHARPIN |
| 8 | 144107200 | 144107401 | -1.225705162 | MAF1 |
| 8 | 144107200 | 144107401 | -1.225705162 | SHARPIN |
| 8 | 144139683 | 144139905 | -1.131222029 | HGH1 |
| 8 | 144263475 | 144263696 | 1.161970392 | BOP1 |
| 8 | 144310522 | 144310709 | 6.409291089 | HSF1 |
| 8 | 144494208 | 144494449 | -1.355925802 | PPP1R16A |

| | | | | |
|---|---|---|---|---|
| 13 | 114116230 | 114116383 | 6.861403766 | RASA3 |
| 14 | 19300320 | 19300517 | 1.59013979 | AL589743.1 |
| 14 | 19300320 | 19300517 | 1.59013979 | LINC01296 |
| 14 | 19302682 | 19302838 | 1.036738506 | LINC01296 |
| 14 | 19302682 | 19302838 | 1.036738506 | BMS1P17 |
| 14 | 19302682 | 19302838 | 1.036738506 | AL589743.1 |
| 14 | 19316403 | 19316587 | -1.185984389 | BMS1P17 |
| 14 | 19316403 | 19316587 | -1.185984389 | LINC01296 |
| 14 | 20801695 | 20801869 | 7.099499581 | RNASE1 |
| 14 | 20983269 | 20983467 | 1.319148332 | RP11-84C10.1 |
| 14 | 21081896 | 21082080 | 1.301870465 | ARHGEF40 |
| 14 | 21384482 | 21384666 | 1.491772488 | RP11-524O1.4 |
| 14 | 22845956 | 22846158 | 1.319224441 | MMP14 |
| 14 | 23095827 | 23096089 | 1.427479704 | C14orf119 |
| 14 | 23320487 | 23320686 | 1.071311754 | BCL2L2-PABPN1 |
| 14 | 24213015 | 24213217 | -1.230847976 | CHMP4A |
| 14 | 24213015 | 24213217 | -1.230847976 | RP11-468E2.1 |
| 14 | 24213015 | 24213217 | -1.230847976 | AL136419.6 |
| 14 | 24317682 | 24317825 | 6.008404144 | LTB4R |
| 14 | 24431815 | 24432044 | 1.010234621 | KHNYN |
| 14 | 30666292 | 30666424 | -4.527641218 | SCFD1 |
| 14 | 31026847 | 31027017 | -5.810568057 | AP4S1 |
| 14 | 31127902 | 31128080 | -1.196947932 | HECTD1 |
| 14 | 34518747 | 34518871 | -5.647023157 | EAPP |
| 14 | 37591318 | 37591546 | 1.658682564 | FOXA1 |
| 14 | 41607789 | 41608031 | 6.868465186 | LRFN5 |
| 14 | 41886792 | 41886931 | 1.144753309 | LRFN5 |
| 14 | 49622154 | 49622355 | 11.71404333 | RP11-649E7.5 |
| 14 | 49622154 | 49622355 | 11.71404333 | MGAT2 |
| 14 | 50329969 | 50330186 | -1.261828289 | ATP5S |
| 14 | 50329969 | 50330186 | -1.261828289 | CDKL1 |
| 14 | 50757300 | 50757433 | 1.192747017 | NIN |
| 1 | 40473208 | 40473425 | 2.165243713 | RP11-656D10.7 |
| 14 | 51066332 | 51066454 | 2.141410468 | TRIM9 |
| 14 | 53428711 | 53428889 | 2.470130367 | AL163953.3 |
| 14 | 53950559 | 53950758 | 1.616343584 | BMP4 |
| 14 | 54405575 | 54405725 | -5.727149647 | CDKN3 |
| 14 | 54567734 | 54567887 | 1.75792983 | SAMD4A |
| 14 | 54736892 | 54737104 | 10.36929437 | SAMD4A |
| 1 | 40486959 | 40487161 | -1.684905158 | ZFP69 |
| 14 | 54748660 | 54748856 | 1.455530216 | SAMD4A |
| 1 | 40495532 | 40495772 | 1.591220995 | RP11-656D10.3 |
| 1 | 40495532 | 40495772 | 1.591220995 | ZFP69 |
| 14 | 56296859 | 56297060 | -1.804389241 | PELI2 |
| 14 | 57391289 | 57391557 | 2.379158092 | NAA30 |
| 1 | 1354281 | 1354460 | 10.18269774 | MXRA8 |
| 14 | 58521660 | 58521854 | 6.566972457 | HNRNPCP1 |
| 14 | 58521660 | 58521854 | 6.566972457 | KIAA0586 |
| 14 | 59727069 | 59727239 | 8.302906129 | RTN1 |
| 14 | 60248782 | 60248981 | 1.835587562 | PPM1A |
| 14 | 61012438 | 61012645 | -1.658167573 | SLC38A6 |
| 14 | 61013153 | 61013351 | -1.457639143 | SLC38A6 |
| 14 | 61014856 | 61015043 | -2.02371573 | SLC38A6 |
| 14 | 61061478 | 61061609 | -3.443571474 | SLC38A6 |

| | | | | |
|---|---|---|---|---|
| 8 | 25411627 | 25411827 | 1.051857536 | DOCK5 |
| 8 | 25506868 | 25507053 | -3.034584582 | CDCA2 |
| 8 | 25507309 | 25507527 | -2.573364242 | CDCA2 |
| 8 | 26324646 | 26324900 | 7.859507204 | PPP2R2A |
| 8 | 27776516 | 27776701 | -5.534787818 | ESCO2 |
| 8 | 27812387 | 27812566 | -3.250604174 | PBK |
| 8 | 27812387 | 27812566 | -3.250604174 | ESCO2 |
| 8 | 28348546 | 28348749 | -1.982604961 | ZNF395 |
| 8 | 28348546 | 28348749 | -1.982604961 | FBXO16 |
| 8 | 28502830 | 28503079 | -1.398666883 | FZD3 |
| 8 | 28562747 | 28562948 | -1.05609247 | FZD3 |
| 8 | 29336750 | 29336948 | 4.772602985 | DUSP4 |
| 8 | 29337333 | 29337510 | 5.233177299 | DUSP4 |
| 8 | 29350188 | 29350325 | 5.11860498 | DUSP4 |
| 8 | 30156017 | 30156204 | -7.0691359 | RP11-51J9.5 |
| 8 | 30156017 | 30156204 | -7.0691359 | LEPROTL1 |
| 8 | 30596694 | 30596896 | 1.308952778 | GTF2E2 |
| 8 | 30744142 | 30744338 | 2.665458439 | UBXN8 |
| 8 | 30837825 | 30838013 | -9.993148799 | TEX15 |
| 8 | 32763972 | 32764172 | 6.970156799 | NRG1 |
| 8 | 35952129 | 35952324 | -4.865222203 | RP11-473J6.1 |
| 8 | 35956125 | 35956297 | -2.39395128 | RP11-473J6.1 |
| 8 | 37698501 | 37698718 | 2.850007262 | ZNF703 |
| 8 | 37842435 | 37842633 | -1.00180335 | ADGRA2 |
| 8 | 37859971 | 37860155 | -1.158611988 | RP11-205M5.3 |
| 8 | 37859971 | 37860155 | -1.158611988 | RAB11FIP1 |
| 8 | 38392711 | 38392910 | -1.679589745 | LETM2 |
| 8 | 38820111 | 38820309 | -1.342242462 | TACC1 |
| 8 | 41655501 | 41655683 | -1.808486448 | ANK1 |
| 8 | 41831962 | 41832113 | -1.522968142 | ANK1 |
| 8 | 41937466 | 41937664 | 1.089950534 | KAT6A |
| 8 | 42169298 | 42169446 | -2.675287912 | AP3M2 |
| 8 | 42297433 | 42297610 | -6.367778012 | IKBKB |
| 8 | 42297896 | 42298083 | -8.278447588 | IKBKB |
| 1 | 235329377 | 235329576 | -1.893547807 | GGPS1 |
| 8 | 42542126 | 42542287 | -1.42207742 | SMIM19 |
| 8 | 47313488 | 47313704 | -2.890679857 | SPIDR |
| 8 | 47469438 | 47469626 | -1.568104378 | SPIDR |
| 8 | 47489839 | 47490062 | -1.424215629 | SPIDR |
| 8 | 47490637 | 47490825 | -1.361139363 | SPIDR |
| 8 | 47623364 | 47623628 | -1.973657746 | SPIDR |
| 1 | 11805564 | 11805754 | 1.439949013 | MTHFR |
| 1 | 235662804 | 235662982 | 1.499668812 | LYST |
| 8 | 49176835 | 49177064 | 2.157031728 | RP11-10H3.1 |
| 8 | 49178172 | 49178323 | 2.095679605 | RP11-10H3.1 |
| 8 | 49200478 | 49200671 | 2.097019381 | RP11-10H3.1 |
| 8 | 49200893 | 49201069 | 4.189448454 | RP11-10H3.1 |
| 8 | 49201695 | 49201827 | 3.800037249 | RP11-10H3.1 |
| 1 | 235738003 | 235738197 | -1.549792137 | LDHAP2 |
| 1 | 235738003 | 235738197 | -1.549792137 | LYST |
| 8 | 51820426 | 51820642 | 1.247762257 | PCMTD1 |
| 8 | 51860946 | 51861150 | 1.049326297 | PCMTD1 |
| 8 | 52714786 | 52714980 | 1.103499987 | RB1CC1 |
| 8 | 53780804 | 53781022 | 2.677925592 | ATP6V1H |
| 8 | 144509120 | 144509318 | -5.119799265 | MFSD3 |
| 1 | 244401416 | 244401571 | -7.357230094 | RP11-518L10.5 |
| 8 | 144548068 | 144548249 | 4.554262228 | ARHGAP39 |
| 8 | 144722051 | 144722250 | -2.200186644 | ZNF251 |
| 8 | 144723051 | 144723250 | -2.213286793 | ZNF251 |
| 8 | 144742692 | 144742890 | -1.215629882 | ZNF251 |
| 8 | 144773467 | 144773745 | -1.571988275 | ZNF34 |
| 8 | 144808311 | 144808560 | -5.511183907 | ZNF517 |
| 8 | 144810368 | 144810566 | -1.940158998 | ZNF517 |
| 8 | 144841416 | 144841628 | -1.259821351 | COMMD5 |
| 8 | 144841416 | 144841628 | -1.259821351 | ZNF7 |
| 8 | 144881294 | 144881493 | -1.227189195 | ZNF250 |
| 8 | 144978076 | 144978226 | -3.584962501 | ZNF252P |
| 9 | 112919 | 113117 | 2.229736573 | LINC01388 |
| 9 | 116629 | 116821 | 2.967322244 | FOXD4 |
| 9 | 306981 | 307165 | 1.02527085 | DOCK8 |
| 9 | 1047282 | 1047473 | -1.290530532 | LINC01230 |
| 1 | 244656943 | 244657125 | -2.023534127 | DESI2 |
| 9 | 1056691 | 1056950 | -2.383743769 | DMRT2 |
| 9 | 2655098 | 2655234 | -3.094912279 | VLDLR |
| 9 | 3530417 | 3530641 | -1.45093626 | RFX3-AS1 |
| 9 | 3674465 | 3674622 | -6.299491486 | RFX3-AS1 |
| 9 | 4662718 | 4662835 | -5.922745025 | SPATA6L |
| 9 | 4662718 | 4662835 | -5.922745025 | PLPP6 |
| 9 | 4663305 | 4663467 | -2.343310159 | PLPP6 |
| 9 | 4663305 | 4663467 | -2.343310159 | SPATA6L |
| 1 | 244701321 | 244701454 | 1.04219477 | DESI2 |
| 9 | 5034015 | 5034153 | 10.00399679 | JAK2 |
| 9 | 5035641 | 5035837 | 8.590299396 | JAK2 |
| 9 | 5037409 | 5037605 | 9.785942967 | JAK2 |
| 9 | 5106607 | 5106804 | 4.700586061 | JAK2 |
| 9 | 5111066 | 5111178 | 9.724840329 | TCF3P1 |
| 9 | 5111066 | 5111178 | 9.724840329 | JAK2 |
| 9 | 5113478 | 5113674 | 5.149707895 | JAK2 |
| 9 | 5113478 | 5113674 | 5.149707895 | IGHEP2 |
| 9 | 5114887 | 5115066 | 5.759254418 | JAK2 |
| 9 | 5133836 | 5134036 | 2.303127926 | INSL6 |
| 9 | 5135545 | 5135748 | 2.313175613 | INSL6 |
| 9 | 5136535 | 5136719 | 3.890117822 | INSL6 |
| 9 | 5137824 | 5138009 | 2.901783356 | INSL6 |
| 9 | 5138350 | 5138451 | 3.5025175 | INSL6 |
| 9 | 5154994 | 5155225 | 3.405794896 | INSL6 |
| 9 | 5168511 | 5168705 | 2.309240911 | INSL6 |
| 9 | 5169562 | 5169803 | 2.277489662 | INSL6 |
| 9 | 5172239 | 5172437 | 2.885133785 | INSL6 |
| 9 | 5172841 | 5173040 | 3.835198989 | INSL6 |
| 9 | 5173327 | 5173514 | 3.500310716 | INSL6 |
| 9 | 5179712 | 5179911 | 3.702418725 | INSL6 |
| 9 | 5467826 | 5468026 | 7.091007067 | CD274 |
| 9 | 5539361 | 5539537 | 1.131517203 | PDCD1LG2 |
| 9 | 5779964 | 5780190 | 1.419944985 | ERMP1 |
| 9 | 5896417 | 5896642 | -1.847230433 | KIAA2026 |
| 9 | 5896417 | 5896642 | -1.847230433 | MLANA |
| 9 | 5897637 | 5897833 | -2.714867753 | KIAA2026 |
| 14 | 64469693 | 64469889 | 3.252000106 | AKAP5 |
| 14 | 64469693 | 64469889 | 3.252000106 | ZBTB25 |
| 14 | 64542411 | 64542574 | -5.148458103 | HSPA2 |
| 14 | 64600908 | 64601108 | -1.600816931 | RP11-973N13.2 |
| 14 | 64741131 | 64741288 | -5.836654538 | PLEKHG3 |
| 14 | 65411798 | 65411999 | 2.548058982 | FUT8 |
| 14 | 65411798 | 65411999 | 2.548058982 | FUT8-AS1 |
| 14 | 65633099 | 65633233 | -6.509013133 | FUT8 |
| 14 | 67328524 | 67328681 | -7.666962012 | ATP6V1D |
| 14 | 67328524 | 67328681 | -7.666962012 | MPP5 |
| 14 | 67729109 | 67729304 | -2.339484836 | RDH12 |
| 14 | 67729109 | 67729304 | -2.339484836 | ZFYVE26 |
| 14 | 67976649 | 67976824 | -1.656830239 | RAD51B |
| 1 | 40838479 | 40838709 | -1.390935544 | KCNQ4 |
| 14 | 70730351 | 70730550 | -1.513236106 | MAP3K9 |
| 14 | 70943261 | 70943462 | -1.089742251 | PCNX1 |
| 14 | 71060987 | 71061225 | -1.802959234 | PCNX1 |
| 14 | 71092596 | 71092797 | -1.578273708 | PCNX1 |
| 14 | 71588402 | 71588580 | -1.677297142 | SIPA1L1 |
| 14 | 72671063 | 72671182 | 1.224784953 | DPF3 |
| 14 | 73058402 | 73058534 | 2.48012001 | RP11-109N23.4 |
| 14 | 73058402 | 73058534 | 2.48012001 | RBM25 |
| 14 | 73239507 | 73239713 | 1.74849858 | PAPLN |
| 1 | 41132575 | 41132723 | -4.751912869 | SCMH1 |
| 14 | 73522726 | 73522925 | -5.652460835 | RP3-414A15.2 |
| 14 | 73522726 | 73522925 | -5.652460835 | HEATR4 |
| 14 | 73721377 | 73721553 | -1.41923714 | ELMSAN1 |
| 14 | 74500295 | 74500487 | -1.397145908 | LTBP2 |
| 14 | 74840726 | 74840949 | -6.46721402 | YLPM1 |
| 14 | 74906591 | 74906781 | 2.347576418 | RPS6KL1 |
| 14 | 75046924 | 75047129 | -1.420283279 | MLH3 |
| 14 | 75071057 | 75071249 | 5.020829576 | ZC2HC1C |
| 14 | 75546665 | 75546859 | 2.091416243 | BATF |
| 14 | 75958898 | 75959049 | 3.009426702 | IFT43 |
| 14 | 75958898 | 75959049 | 3.009426702 | TGFB3 |
| 14 | 76206317 | 76206525 | -1.002683447 | GPATCH2L |
| 14 | 76762098 | 76762200 | 9.46405097 | VASH1 |
| 14 | 76775983 | 76776181 | 11.65246912 | VASH1 |
| 14 | 76775983 | 76776181 | 11.65246912 | RP11-488C13.6 |
| 14 | 77025365 | 77025564 | -1.123463592 | IRF2BPL |
| 14 | 80706659 | 80706858 | 1.077409055 | CEP128 |
| 14 | 81270764 | 81270957 | -1.189160465 | STON2 |
| 14 | 81276940 | 81277146 | -1.049898802 | STON2 |
| 1 | 41543186 | 41543363 | -7.174709732 | HIVEP3 |
| 1 | 41543186 | 41543363 | -7.174709732 | RP11-486B10.4 |
| 14 | 85562749 | 85562916 | -2.388276698 | FLRT2 |
| 14 | 88010651 | 88010791 | 1.989468621 | GPR65 |
| 14 | 88010651 | 88010791 | 1.989468621 | RP11-300J18.2 |
| 14 | 89215554 | 89215725 | 3.830870687 | FOXN3 |
| 14 | 91233560 | 91233766 | 1.873056148 | GPR68 |
| 14 | 91258344 | 91258556 | 1.527122971 | CTD-2547L24.4 |
| 1 | 41549558 | 41549758 | -1.058899959 | HIVEP3 |
| 14 | 92161680 | 92161879 | -1.131824225 | CPSF2 |
| 14 | 92941491 | 92941666 | 3.77595024 | ITPK1 |

| | | | | |
|---|---|---|---|---|
| 8 | 53781425 | 53781614 | 2.663105104 | ATP6V1H |
| 1 | 235738293 | 235738492 | -1.614040485 | LYST |
| 1 | 235738293 | 235738492 | -1.614040485 | LDHAP2 |
| 8 | 53784307 | 53784499 | 2.899767776 | ATP6V1H |
| 8 | 53785696 | 53785893 | 3.779104049 | ATP6V1H |
| 8 | 53935132 | 53935330 | -5.635147663 | RGS20 |
| 8 | 53938126 | 53938348 | -1.174308537 | RGS20 |
| 8 | 55786451 | 55786653 | -1.345748539 | TGS1 |
| 1 | 235740457 | 235740653 | -1.476312949 | LYST |
| 8 | 56161428 | 56161591 | -1.313477208 | PLAG1 |
| 1 | 235759304 | 235759516 | -1.53434313 | LYST |
| 8 | 57546710 | 57546974 | -2.370448342 | RP11-246K15.1 |
| 8 | 57568226 | 57568416 | -1.736418184 | RP11-246K15.1 |
| 8 | 58146130 | 58146328 | 7.189993742 | FAM110B |
| 8 | 58146948 | 58147260 | 2.859093936 | FAM110B |
| 1 | 235806031 | 235806256 | -1.677418754 | LYST |
| 8 | 58561291 | 58561475 | 1.323260244 | SDCBP |
| 8 | 60465067 | 60465219 | -3.584962501 | LINC01301 |
| 8 | 60760872 | 60761082 | 1.206203956 | CHD7 |
| 1 | 235808863 | 235809071 | -1.439408361 | LYST |
| 8 | 61578234 | 61578448 | 1.693313517 | ASPH |
| 8 | 61578234 | 61578448 | 1.693313517 | KRT8P3 |
| 8 | 61578234 | 61578448 | 1.693313517 | AC090094.1 |
| 8 | 61579995 | 61580186 | -4.754887502 | ASPH |
| 8 | 61579995 | 61580186 | -4.754887502 | AC090094.1 |
| 8 | 61607029 | 61607226 | 1.827967443 | ASPH |
| 8 | 61607029 | 61607226 | 1.827967443 | AC090094.1 |
| 8 | 64801204 | 64801385 | -6.63873312 | RP11-1D12.2 |
| 1 | 235809998 | 235810196 | -1.249001011 | LYST |
| 8 | 65604295 | 65604501 | -1.299538676 | ARMC1 |
| 8 | 65719140 | 65719466 | -1.056152963 | PDE7A |
| 8 | 65719140 | 65719466 | -1.056152963 | MTFR1 |
| 8 | 66429852 | 66430086 | -2.255537978 | RRS1 |
| 8 | 71842628 | 71842775 | 2.193575283 | MSC |
| 8 | 71842628 | 71842775 | 2.193575283 | MSC-AS1 |
| 8 | 71966701 | 71966887 | 1.887405015 | MSC-AS1 |
| 8 | 72033573 | 72033760 | 1.427072337 | TRPA1 |
| 8 | 72033573 | 72033760 | 1.427072337 | MSC-AS1 |
| 8 | 72052603 | 72052801 | 3.638551288 | TRPA1 |
| 8 | 72052603 | 72052801 | 3.638551288 | MSC-AS1 |
| 8 | 73055664 | 73055798 | -1.173087821 | RP11-531A24.5 |
| 8 | 73056589 | 73056769 | -1.355673299 | RP11-531A24.5 |
| 1 | 11920200 | 11920407 | -2.564556594 | KIAA2013 |
| 8 | 73787710 | 73787876 | -1.411726005 | UBE2W |
| 8 | 73787710 | 73787876 | -1.411726005 | RP11-463D19.2 |
| 8 | 74259464 | 74259643 | -1.132924109 | JPH1 |
| 8 | 74314920 | 74315129 | -1.850293684 | JPH1 |
| 8 | 80033441 | 80033658 | -2.406470326 | RP11-26J3.4 |
| 8 | 80975603 | 80975805 | 1.97106856 | PAG1 |
| 8 | 80976257 | 80976706 | 1.60670952 | PAG1 |
| 1 | 236483294 | 236483402 | -4.673256867 | ENO1P1 |
| 1 | 236483294 | 236483402 | -4.673256867 | EDARADD |
| 8 | 80985095 | 80985300 | 1.464409464 | PAG1 |
| 8 | 86481893 | 86482080 | -1.747675596 | RMDN1 |
| 9 | 5897637 | 5897833 | -2.714867753 | MLANA |
| 9 | 5911077 | 5911275 | -1.301215311 | KIAA2026 |
| 9 | 5919919 | 5920103 | -1.060410792 | KIAA2026 |
| 9 | 5921271 | 5921469 | -1.239892631 | KIAA2026 |
| 9 | 5923069 | 5923306 | -1.053557511 | KIAA2026 |
| 9 | 5988462 | 5988626 | 4.332667959 | KIAA2026 |
| 9 | 6460497 | 6460668 | -1.781828501 | UHRF2 |
| 9 | 12775342 | 12775543 | -1.318420844 | LURAP1L-AS1 |
| 9 | 12775342 | 12775543 | -1.318420844 | LURAP1L |
| 9 | 12821536 | 12821702 | -2.804404621 | LURAP1L |
| 1 | 245091241 | 245091453 | -1.836273113 | EFCAB2 |
| 9 | 19372100 | 19372298 | -1.445003003 | DENND4C |
| 9 | 20937514 | 20937714 | -5.079935668 | FOCAD |
| 9 | 21005350 | 21005504 | -7.315815903 | HACD4 |
| 9 | 21022190 | 21022377 | -6.798618923 | HACD4 |
| 9 | 21022917 | 21023107 | -6.75389732 | HACD4 |
| 1 | 245123230 | 245123417 | -2.4129853 | EFCAB2 |
| 9 | 27330304 | 27330489 | 4.42264701 | MOB3B |
| 9 | 27455274 | 27455600 | 2.471085471 | MOB3B |
| 1 | 245672071 | 245672239 | -5.330889049 | KIF26B |
| 9 | 32431347 | 32431547 | 3.659492292 | ACO1 |
| 9 | 32444532 | 32444758 | 3.232628675 | ACO1 |
| 9 | 32446367 | 32446565 | 3.328382719 | ACO1 |
| 9 | 32448674 | 32448867 | 2.191525391 | ACO1 |
| 9 | 32449747 | 32449930 | 8.860015223 | ACO1 |
| 9 | 32457123 | 32457325 | 2.617007149 | DDX58 |
| 9 | 32460594 | 32460783 | -6.658140766 | DDX58 |
| 1 | 245680643 | 245680745 | -5.251912869 | KIF26B |
| 9 | 32979288 | 32979488 | -1.253797769 | APTX |
| 9 | 33038678 | 33038883 | 1.601611929 | DNAJA1 |
| 9 | 33104237 | 33104430 | -6.030347966 | B4GALT1 |
| 1 | 245686296 | 245686464 | -1.898233303 | KIF26B |
| 9 | 33917230 | 33917427 | -1.073589745 | UBE2R2 |
| 9 | 34214669 | 34214919 | -7.469605941 | UBAP1 |
| 9 | 34249062 | 34249253 | 5.981989894 | UBAP1 |
| 9 | 34257550 | 34257683 | -1.034770467 | KIF24 |
| 9 | 34370741 | 34370845 | 1.488639962 | KIAA1161 |
| 9 | 34372835 | 34373089 | -6.473818469 | KIAA1161 |
| 9 | 34635466 | 34635670 | -1.000715435 | SIGMAR1 |
| 9 | 34680941 | 34681133 | -2.683430921 | RP11-195F19.9 |
| 1 | 245702616 | 245702820 | -3.584962501 | KIF26B |
| 9 | 35538064 | 35538256 | -1.183589542 | RUSC2 |
| 9 | 35546876 | 35547069 | -1.255014182 | RUSC2 |
| 9 | 35547610 | 35547813 | -1.396642081 | RUSC2 |
| 9 | 35555290 | 35555480 | -1.324509869 | RUSC2 |
| 9 | 35609682 | 35609885 | -1.535185575 | CD72 |
| 9 | 35609682 | 35609885 | -1.535185575 | TESK1 |
| 9 | 35649668 | 35649829 | -4.336212671 | SIT1 |
| 9 | 35660862 | 35661061 | -1.55851215 | CCDC107 |
| 9 | 35660862 | 35661061 | -1.55851215 | ARHGEF39 |
| 9 | 35661688 | 35661885 | -5.043731421 | ARHGEF39 |
| 1 | 246147122 | 246147330 | -2.393091816 | SMYD3 |
| 9 | 36170527 | 36170729 | -1.997777942 | CCIN |
| 9 | 36217304 | 36217528 | -1.254031662 | GNE |
| 1 | 41552654 | 41552852 | -1.520162477 | HIVEP3 |
| 14 | 95417753 | 95417896 | -1.236085365 | SYNE3 |
| 1 | 41555806 | 41556004 | -1.001819988 | HIVEP3 |
| 14 | 95543945 | 95544154 | 1.485011663 | GLRX5 |
| 14 | 96354378 | 96354588 | -1.132402817 | ATG2B |
| 14 | 96501731 | 96501886 | 5.781121212 | RP11-872J21.3 |
| 14 | 96501731 | 96501886 | 5.781121212 | PAPOLA |
| 14 | 99708816 | 99708963 | -3.172147954 | CYP46A1 |
| 14 | 100329186 | 100329348 | -2.033677634 | SLC25A47 |
| 14 | 100380915 | 100381105 | -1.558937077 | WDR25 |
| 14 | 100455211 | 100455379 | -4.805512399 | WDR25 |
| 14 | 101925331 | 101925478 | 1.058576813 | PPP2R5C |
| 14 | 101948756 | 101948911 | 1.817915842 | RP11-1017G21.5 |
| 14 | 102834058 | 102834234 | -6.110491426 | TRAF3 |
| 14 | 103412270 | 103412427 | 5.680971887 | MARK3 |
| 14 | 103412270 | 103412427 | 5.680971887 | RPL10AP1 |
| 14 | 103534709 | 103534912 | 2.29664543 | TRMT61A |
| 14 | 103590504 | 103590710 | 1.412836807 | KLC1 |
| 14 | 103590504 | 103590710 | 1.412836807 | APOPT1 |
| 14 | 103590504 | 103590710 | 1.412836807 | RP11-73M18.2 |
| 14 | 103705063 | 103705263 | 4.206689519 | KLC1 |
| 14 | 103705063 | 103705263 | 4.206689519 | XRCC3 |
| 14 | 103713418 | 103713598 | -1.626994633 | XRCC3 |
| 14 | 103713418 | 103713598 | -1.626994633 | KLC1 |
| 14 | 103734789 | 103734989 | 1.232576125 | PPP1R13B |
| 14 | 104704130 | 104704319 | 7.647023157 | INF2 |
| 14 | 104719716 | 104719917 | 1.395745314 | INF2 |
| 14 | 104884253 | 104884380 | 3.565101721 | CEP170B |
| 14 | 104886986 | 104887185 | 1.314512463 | CEP170B |
| 14 | 104889474 | 104889645 | 8.197114105 | CEP170B |
| 14 | 105050822 | 105051022 | -1.812977419 | GPR132 |
| 14 | 105054037 | 105054225 | -1.352177384 | GPR132 |
| 1 | 42559335 | 42559541 | 1.12050405 | CCDC30 |
| 14 | 105142119 | 105142317 | 1.239546765 | JAG2 |
| 14 | 105247368 | 105247545 | 1.084394195 | BRF1 |
| 14 | 105371268 | 105371467 | 1.253404091 | PACS2 |
| 14 | 105371268 | 105371467 | 1.253404091 | RP11-521B24.4 |
| 14 | 105375118 | 105375335 | 6.376190323 | PACS2 |
| 1 | 42653931 | 42654130 | -1.495193288 | CCDC30 |
| 14 | 105401393 | 105401575 | -1.002683255 | TEX22 |
| 14 | 105416831 | 105416959 | -1.689255812 | RP11-521B24.3 |
| 14 | 105416831 | 105416959 | -1.689255812 | TEX22 |
| 14 | 105442660 | 105442858 | 1.916123104 | TEX22 |
| 14 | 105442660 | 105442858 | 1.916123104 | MTA1 |
| 14 | 105498930 | 105499141 | 1.818811024 | C14orf80 |
| 15 | 21294182 | 21294371 | -2.44180677 | RP11-32B5.1 |
| 15 | 25709780 | 25709981 | -6.459338463 | RP11-345J18.2 |
| 15 | 25709780 | 25709981 | -6.459338463 | ATP10A |
| 15 | 25754139 | 25754307 | 5.892317423 | ATP10A |
| 15 | 28155038 | 28155231 | -1.275001448 | HERC2 |
| 15 | 28158214 | 28158325 | -4.827318014 | HERC2 |
| 15 | 28702565 | 28702759 | -1.502500341 | GOLGA8M |
| 15 | 28788564 | 28788702 | -6.165318412 | RP11-578F21.10 |
| 15 | 28789582 | 28789786 | -1.507251449 | PDCD6IPP2 |

| | | | | |
|---|---|---|---|---|
| 8 | 86481893 | 86482080 | -1.747675596 | NTAN1P2 |
| 1 | 236543255 | 236543449 | -8.141399429 | RP11-385F5.4 |
| 1 | 236543255 | 236543449 | -8.141399429 | LGALS8 |
| 8 | 89703075 | 89703256 | -2.980580129 | RP11-37B2.1 |
| 8 | 90005491 | 90005666 | -1.088872253 | DECR1 |
| 8 | 91069308 | 91069499 | -1.092802933 | OTUD6B-AS1 |
| 1 | 236547451 | 236547649 | -1.115686126 | RP11-385F5.4 |
| 1 | 236547451 | 236547649 | -1.115686126 | LGALS8 |
| 8 | 94432295 | 94432483 | -2.271112014 | FSBP |
| 8 | 94432295 | 94432483 | -2.271112014 | RAD54B |
| 8 | 94526605 | 94526805 | -1.086419172 | KIAA1429 |
| 8 | 94930579 | 94930815 | 4.352104404 | NDUFAF6 |
| 8 | 94930579 | 94930815 | 4.352104404 | TP53INP1 |
| 8 | 95247521 | 95247693 | -1.711783083 | C8orf37 |
| 8 | 95247521 | 95247693 | -1.711783083 | C8orf37-AS1 |
| 8 | 96333532 | 96333734 | -1.196799122 | PTDSS1 |
| 8 | 96493834 | 96494034 | -7.124927085 | SDC2 |
| 8 | 96930670 | 96930770 | -4.987923984 | CPQ |
| 8 | 96955852 | 96956040 | 1.961795666 | CPQ |
| 8 | 96972109 | 96972303 | 7.110264741 | CPQ |
| 8 | 96974133 | 96974304 | 4.988639962 | CPQ |
| 8 | 96976967 | 96977141 | 7.142556594 | CPQ |
| 8 | 97092642 | 97092834 | 1.266272626 | CPQ |
| 8 | 97144231 | 97144356 | 2.714927519 | CPQ |
| 8 | 97144231 | 97144356 | 2.714927519 | KB-1958F4.1 |
| 8 | 97146872 | 97147087 | 2.136186135 | CPQ |
| 8 | 98157713 | 98158025 | -2.50034207 | POP1 |
| 8 | 98694701 | 98694984 | 1.200471335 | STK3 |
| 8 | 98942657 | 98942816 | -7.382047034 | STK3 |
| 8 | 98949310 | 98949511 | -1.626561261 | OSR2 |
| 8 | 99066462 | 99066658 | -7.427750588 | VPS13B |
| 1 | 236899066 | 236899261 | -1.604624261 | MTR |
| 8 | 99205066 | 99205269 | 2.542710694 | VPS13B |
| 8 | 99336356 | 99336534 | -5.703633882 | VPS13B |
| 8 | 100140878 | 100140992 | -4.652214608 | FBXO43 |
| 1 | 11925300 | 11925499 | -8.963136413 | KIAA2013 |
| 1 | 236903528 | 236903727 | -1.213734455 | MTR |
| 8 | 100293796 | 100294149 | 1.565961246 | RNF19A |
| 8 | 100310458 | 100310717 | -1.353965766 | RNF19A |
| 8 | 100312144 | 100312390 | -1.222002372 | RNF19A |
| 8 | 100336178 | 100336428 | -2.161373036 | RNF19A |
| 8 | 100549927 | 100550125 | -2.213036027 | ANKRD46 |
| 8 | 100550532 | 100550703 | -2.906534287 | ANKRD46 |
| 1 | 237077341 | 237077540 | 5.302739759 | RYR2 |
| 8 | 101177869 | 101178073 | 1.444508478 | ZNF706 |
| 8 | 101719500 | 101719687 | -1.008278349 | NCALD |
| 8 | 102650204 | 102650407 | 3.114308301 | KLF10 |
| 8 | 102651667 | 102651867 | 2.455510383 | KLF10 |
| 8 | 103132702 | 103132856 | 2.008536757 | LINC01181 |
| 8 | 103227940 | 103228138 | 3.473642592 | BAALC |
| 8 | 103227940 | 103228138 | 3.473642592 | BAALC-AS1 |
| 8 | 103324541 | 103324709 | -1.104610985 | FZD6 |
| 8 | 104252251 | 104252430 | 2.357122759 | RIMS2 |
| 8 | 104490894 | 104491094 | 2.780318711 | LRP12 |
| 9 | 36217304 | 36217528 | -1.254031662 | CLTA |
| 9 | 36246124 | 36246329 | -1.032154181 | CLTA |
| 9 | 36246124 | 36246329 | -1.032154181 | GNE |
| 9 | 36293490 | 36293708 | -1.685884417 | CLTA |
| 1 | 246239415 | 246239581 | -6.3488096 | SMYD3 |
| 9 | 36413981 | 36414104 | 7.004214311 | RNF38 |
| 9 | 36839531 | 36839722 | 4.266388325 | PAX5 |
| 9 | 36840212 | 36840430 | 3.875078568 | PAX5 |
| 9 | 36900143 | 36900339 | 9.864172857 | PAX5 |
| 9 | 37027863 | 37028058 | 6.876190323 | PAX5 |
| 9 | 37030569 | 37030758 | 6.769165207 | PAX5 |
| 9 | 37126348 | 37126544 | -1.149039864 | ZCCHC7 |
| 1 | 246240071 | 246240310 | -3.093263484 | SMYD3 |
| 9 | 37503228 | 37503428 | -3.043602827 | POLR1E |
| 9 | 37537512 | 37537702 | -1.346075782 | FBXO10 |
| 9 | 37537512 | 37537702 | -1.346075782 | RP11-613M10.8 |
| 9 | 37588810 | 37589008 | -3.290894514 | RP11-613M10.9 |
| 9 | 37588810 | 37589008 | -3.290894514 | TOMM5 |
| 9 | 37588810 | 37589008 | -3.290894514 | RP11-613M10.8 |
| 9 | 37765322 | 37765468 | -6.330889049 | RP11-613M10.9 |
| 9 | 37765322 | 37765468 | -6.330889049 | TRMT10B |
| 9 | 37766008 | 37766135 | -5.238373102 | RP11-613M10.9 |
| 9 | 37766008 | 37766135 | -5.238373102 | TRMT10B |
| 9 | 37887996 | 37888211 | -1.539755203 | RP11-613M10.9 |
| 9 | 37887996 | 37888211 | -1.539755203 | SLC25A51 |
| 9 | 38396117 | 38396315 | -2.216724685 | ALDH1B1 |
| 9 | 38397096 | 38397289 | -1.698340613 | ALDH1B1 |
| 9 | 38397938 | 38398135 | -7.741766199 | ALDH1B1 |
| 1 | 246254191 | 246254375 | -2.074606146 | SMYD3 |
| 9 | 38553897 | 38554095 | -5.951187557 | RP11-392E22.9 |
| 9 | 38553897 | 38554095 | -5.951187557 | RP11-392E22.11 |
| 9 | 38620410 | 38620577 | 1.168934819 | ANKRD18A |
| 9 | 38620410 | 38620577 | 1.168934819 | FAM201A |
| 9 | 39446877 | 39447060 | -1.493987005 | ZNF658B |
| 1 | 246257207 | 246257392 | -1.20705019 | SMYD3 |
| 9 | 41023412 | 41023559 | -1.958208369 | PGM5P2 |
| 9 | 41023412 | 41023559 | -1.958208369 | FRG1HP |
| 1 | 246271045 | 246271174 | -6.598915499 | SMYD3 |
| 9 | 41135725 | 41135916 | -1.0726242 | FRG1HP |
| 9 | 41135725 | 41135916 | -1.0726242 | CBWD7 |
| 9 | 41480221 | 41480421 | -1.787920851 | RP11-187C18.3 |
| 1 | 246591663 | 246591848 | -1.286278178 | CNST |
| 9 | 62812986 | 62813246 | 1.895521133 | LINC01410 |
| 9 | 65291878 | 65292076 | 1.25682427 | RP11-15J10.1 |
| 9 | 65293863 | 65294041 | 1.605872589 | RP11-15J10.1 |
| 9 | 65730219 | 65730371 | 3.297814296 | CBWD5 |
| 9 | 66918214 | 66918381 | -1.455460574 | ZNF658 |
| 9 | 66919568 | 66919739 | 3.377443751 | ZNF658 |
| 9 | 67718526 | 67718733 | -7.461152824 | FAM27E3 |
| 9 | 70413093 | 70413320 | 1.306712357 | KLF9 |
| 9 | 71745276 | 71745475 | 1.275162521 | TMEM2 |
| 9 | 72333625 | 72333816 | -1.06344286 | LINC01504 |
| 1 | 246850616 | 246850838 | -1.578587736 | AHCTF1 |
| 9 | 75087809 | 75087967 | -3.566807719 | NMRK1 |
| 15 | 28789582 | 28789786 | -1.507251449 | RP11-578F21.10 |
| 15 | 28858219 | 28858424 | -1.053891102 | PDCD6IPP2 |
| 15 | 28858219 | 28858424 | -1.053891102 | WI2-2334D6.1 |
| 15 | 30626318 | 30626515 | 1.692507645 | ARHGAP11B |
| 15 | 30741707 | 30741894 | -5.462406252 | ARHGAP11B |
| 15 | 30904781 | 30904934 | -1.331634963 | FAN1 |
| 15 | 32499795 | 32499994 | -2.132305075 | RP11-632K20.2 |
| 15 | 32615702 | 32615905 | 8.605506857 | ARHGAP11A |
| 15 | 34287976 | 34288117 | -7.41277755 | SLC12A6 |
| 15 | 35092377 | 35092570 | -1.487744945 | RP11-323I15.2 |
| 15 | 35298106 | 35298305 | -3.083625434 | RBM17P4 |
| 15 | 35298106 | 35298305 | -3.083625434 | DPH6 |
| 15 | 38351104 | 38351282 | 2.309469688 | SPRED1 |
| 1 | 43363696 | 43363888 | 3.45111102 | ELOVL1 |
| 15 | 39595718 | 39595906 | 4.010890869 | THBS1 |
| 15 | 40089458 | 40089665 | 1.289062335 | BMF |
| 15 | 40282763 | 40282961 | 11.15225588 | PLCB2 |
| 15 | 40288771 | 40288976 | 1.607066087 | PLCB2 |
| 15 | 40368673 | 40368821 | 1.078115297 | DISP2 |
| 15 | 40369742 | 40369884 | 1.793846147 | DISP2 |
| 15 | 40471510 | 40471708 | 1.822688185 | CHST14 |
| 15 | 40521147 | 40521272 | 5.79902625 | RP11-111A22.1 |
| 15 | 40621206 | 40621406 | -6.343687842 | KNL1 |
| 1 | 43413534 | 43413731 | 7.129577384 | SZT2 |
| 15 | 40856967 | 40857198 | 1.609819825 | SPINT1 |
| 15 | 40872914 | 40873104 | 3.35696087 | RHOV |
| 15 | 40873168 | 40873336 | 4.810964879 | RHOV |
| 15 | 41713199 | 41713357 | -1.163571367 | MGA |
| 15 | 41825130 | 41825330 | 1.872877859 | RP11-23P13.4 |
| 15 | 41825130 | 41825330 | 1.872877859 | MAPKBP1 |
| 15 | 41900450 | 41900647 | 1.569044638 | EHD4 |
| 15 | 41908471 | 41908650 | 5.457939689 | EHD4 |
| 15 | 41908471 | 41908650 | 5.457939689 | CTD-2382E5.6 |
| 15 | 42249995 | 42250154 | 1.924981764 | TMEM87A |
| 15 | 42273655 | 42273851 | 8.785179876 | GANC |
| 15 | 42273655 | 42273851 | 8.785179876 | TMEM87A |
| 15 | 42299886 | 42300039 | -4.26569073 | GANC |
| 15 | 42304930 | 42305055 | -3.169925001 | GANC |
| 15 | 42691704 | 42691904 | 3.11786353 | STARD9 |
| 15 | 42745855 | 42746020 | -1.064046975 | TTBK2 |
| 1 | 43691005 | 43691187 | 3.02219706 | KDM4A |
| 15 | 43176623 | 43176760 | -1.951114609 | EPB42 |
| 15 | 43176623 | 43176760 | -1.951114609 | TMEM62 |
| 15 | 43328936 | 43329288 | 1.18450425 | LCMT2 |
| 15 | 43523096 | 43523296 | 6.631487851 | MAP1A |
| 15 | 43526380 | 43526559 | 5.856544524 | MAP1A |
| 15 | 43771155 | 43771304 | 7.894613978 | PDIA3 |
| 15 | 44711123 | 44711313 | 1.651039803 | PATL2 |
| 15 | 45200794 | 45200908 | -6.768123608 | SHF |
| 15 | 47771518 | 47771717 | -1.28257701 | SEMA6D |
| 15 | 48825693 | 48825833 | 2.511999449 | SHC4 |
| 1 | 43912870 | 43913068 | 4.803665157 | ST3GAL3 |
| 15 | 49936762 | 49936935 | -2.261422455 | ATP8B4 |
| 15 | 49998720 | 49998897 | -1.414273552 | ATP8B4 |

| | | | | |
|---|---|---|---|---|
| 8 | 104497715 | 104497912 | 2.718834957 | LRP12 |
| 8 | 105789362 | 105789563 | 2.242092095 | ZFPM2 |
| 8 | 105789362 | 105789563 | 2.242092095 | ZFPM2-AS1 |
| 8 | 105792065 | 105792263 | 3.11920166 | ZFPM2-AS1 |
| 8 | 105792065 | 105792263 | 3.11920166 | ZFPM2 |
| 8 | 105792857 | 105793057 | 2.589114624 | ZFPM2 |
| 8 | 105792857 | 105793057 | 2.589114624 | ZFPM2-AS1 |
| 8 | 105794430 | 105794648 | 1.178068289 | ZFPM2-AS1 |
| 8 | 105794430 | 105794648 | 1.178068289 | ZFPM2 |
| 8 | 105796901 | 105797071 | 2.587418517 | ZFPM2-AS1 |
| 8 | 105796901 | 105797071 | 2.587418517 | ZFPM2 |
| 8 | 105844419 | 105844624 | 3.675614633 | ZFPM2-AS1 |
| 8 | 106272387 | 106272651 | 1.607007994 | RP11-395G23.3 |
| 8 | 106565950 | 106566120 | 3.579670125 | OXR1 |
| 8 | 106565950 | 106566120 | 3.579670125 | RP11-649G15.2 |
| 8 | 106590629 | 106590828 | -1.990097216 | OXR1 |
| 8 | 106590629 | 106590828 | -1.990097216 | RP11-649G15.2 |
| 8 | 107251523 | 107251691 | 6.288478332 | ANGPT1 |
| 1 | 239909023 | 239909222 | 7.400308416 | CHRM3 |
| 8 | 109343981 | 109344156 | -1.432318856 | ENY2 |
| 8 | 115414058 | 115414261 | 1.251717046 | TRPS1 |
| 8 | 115587098 | 115587300 | 1.048749955 | TRPS1 |
| 8 | 115604444 | 115604569 | 1.013673405 | TRPS1 |
| 8 | 116771542 | 116771749 | -1.464531823 | UTP23 |
| 8 | 118110168 | 118110383 | 1.481238679 | EXT1 |
| 8 | 119832579 | 119832788 | -1.306797379 | TAF2 |
| 8 | 120477216 | 120477382 | -2.423105429 | MTBP |
| 8 | 120538538 | 120538740 | 1.176472916 | SNTB1 |
| 8 | 120538538 | 120538740 | 1.176472916 | MTBP |
| 8 | 120644487 | 120644636 | -6.756617057 | SNTB1 |
| 1 | 241634115 | 241634321 | -1.701937909 | OPN3 |
| 1 | 241634115 | 241634321 | -1.701937909 | CHML |
| 8 | 122877624 | 122877761 | -7.762025959 | ZHX2 |
| 8 | 122952136 | 122952278 | -1.651337838 | ZHX2 |
| 8 | 123122991 | 123123195 | -1.196789536 | HMGB1P19 |
| 8 | 123122991 | 123123195 | -1.196789536 | TBC1D31 |
| 8 | 123503165 | 123503364 | 7.673126606 | FBXO32 |
| 1 | 241635521 | 241635690 | -1.47511417 | OPN3 |
| 1 | 241635521 | 241635690 | -1.47511417 | CHML |
| 8 | 124240660 | 124240793 | -2.82688034 | RP11-37N22.1 |
| 8 | 124452529 | 124452731 | -1.052672839 | TRMT12 |
| 8 | 124553103 | 124553316 | 4.467208755 | MTSS1 |
| 8 | 124553103 | 124553316 | 4.467208755 | NDUFB9 |
| 8 | 124568045 | 124568183 | -5.673256867 | NDUFB9 |
| 8 | 124568045 | 124568183 | -5.673256867 | MTSS1 |
| 1 | 11975161 | 11975361 | -1.49756041 | PLOD1 |
| 1 | 241878761 | 241878951 | -4.599573603 | EXO1 |
| 8 | 124976462 | 124976650 | -4.419117117 | ZNF572 |
| 8 | 125433340 | 125433541 | 3.390993877 | TRIB1 |
| 8 | 125436187 | 125436415 | 4.021501783 | TRIB1 |
| 8 | 127740784 | 127741007 | -2.386186957 | MYC |
| 8 | 127890673 | 127890852 | -1.70816762 | PVT1_3 |
| 8 | 127890673 | 127890852 | -1.70816762 | PVT1 |
| 8 | 128020382 | 128020572 | -5.009993891 | PVT1 |

| | | | | |
|---|---|---|---|---|
| 9 | 76612733 | 76612866 | -6.657008352 | PRUNE2 |
| 9 | 77373591 | 77373783 | -6.254392581 | VPS13A |
| 9 | 77414149 | 77414289 | -6.059470536 | VPS13A |
| 9 | 78271357 | 78271488 | -6.369257266 | CEP78 |
| 9 | 78329180 | 78329364 | -3.49483682 | PSAT1 |
| 9 | 81633996 | 81634190 | -1.965664508 | TLE1 |
| 9 | 83903493 | 83903735 | -1.25236436 | KIF27 |
| 9 | 84001023 | 84001210 | -2.522701322 | RMI1 |
| 9 | 85632938 | 85633208 | -1.096760126 | AGTPBP1 |
| 9 | 86027714 | 86027915 | 1.199767999 | GOLM1 |
| 9 | 86048897 | 86049094 | 9.048851482 | GOLM1 |
| 1 | 246986719 | 246986891 | -2.260688162 | ZNF695 |
| 1 | 246986719 | 246986891 | -2.260688162 | ZNF670-ZNF695 |
| 9 | 87706709 | 87706931 | -1.240850281 | DAPK1 |
| 9 | 87921168 | 87921393 | -1.350837153 | SPATA31C1 |
| 9 | 87922709 | 87922842 | -2.674043321 | SPATA31C1 |
| 9 | 87923037 | 87923236 | -2.295250342 | SPATA31C1 |
| 9 | 88468295 | 88468487 | -2.168859799 | SPIN1 |
| 9 | 88475222 | 88475417 | -2.282336484 | SPIN1 |
| 9 | 89001619 | 89001814 | 1.425678381 | S1PR3 |
| 9 | 89002302 | 89002509 | 2.849317327 | S1PR3 |
| 9 | 89361460 | 89361656 | 3.294482216 | SEMA4D |
| 1 | 247038035 | 247038243 | -1.716218093 | ZNF670-ZNF695 |
| 1 | 247038035 | 247038243 | -1.716218093 | ZNF670 |
| 9 | 91229442 | 91229602 | -6.452317311 | AUH |
| 9 | 92549034 | 92549153 | -5.850653231 | CENPP |
| 9 | 92618865 | 92619065 | -2.29248125 | CENPP |
| 9 | 92618865 | 92619065 | -2.29248125 | IPPK |
| 9 | 92631987 | 92632162 | 5.52219706 | IPPK |
| 9 | 92719331 | 92719526 | -1.592941923 | BICD2 |
| 9 | 92832229 | 92832396 | -1.384940108 | ANKRD19P |
| 9 | 92836858 | 92837060 | -1.351102774 | EEF1DP2 |
| 9 | 92836858 | 92837060 | -1.351102774 | ANKRD19P |
| 9 | 92846270 | 92846469 | -1.19418134 | ZNF484 |
| 9 | 92846270 | 92846469 | -1.19418134 | ANKRD19P |
| 1 | 247100696 | 247100900 | -2.127657961 | ZNF669 |
| 9 | 92847235 | 92847499 | -1.216698992 | ANKRD19P |
| 9 | 92847235 | 92847499 | -1.216698992 | ZNF484 |
| 9 | 92848247 | 92848440 | -6.911384987 | ZNF484 |
| 9 | 92848247 | 92848440 | -6.911384987 | ANKRD19P |
| 9 | 93035619 | 93035754 | -4.27994848 | FGD3 |
| 9 | 93084842 | 93085042 | -2.006763495 | SUSD3 |
| 9 | 93121781 | 93121999 | 1.889463475 | NINJ1 |
| 9 | 93452925 | 93453025 | 6.772482216 | FAM120A |
| 9 | 93452925 | 93453025 | 6.772482216 | FAM120AOS |
| 9 | 93678379 | 93678548 | -1.278196674 | PHF2 |
| 9 | 94038000 | 94038126 | 1.438872125 | PTPDC1 |
| 9 | 94038000 | 94038126 | 1.438872125 | CYCSP24 |
| 9 | 94098294 | 94098494 | -1.030834527 | PTPDC1 |
| 9 | 94300052 | 94300253 | -1.292699728 | ZNF169 |
| 1 | 247133118 | 247133317 | -1.311856964 | ZNF124 |
| 9 | 94772907 | 94773089 | 1.660001609 | C9orf3 |
| 9 | 95447265 | 95447497 | -2.067891846 | PTCH1 |
| 9 | 96320992 | 96321215 | -1.129688981 | SLC35D2 |

| | | | | |
|---|---|---|---|---|
| 15 | 50000425 | 50000621 | -1.866147994 | ATP8B4 |
| 1 | 43913723 | 43913918 | 5.452317311 | ST3GAL3 |
| 15 | 50944970 | 50945164 | 1.427438295 | AP4E1 |
| 15 | 50944970 | 50945164 | 1.427438295 | DCAF13P3 |
| 15 | 51057983 | 51058183 | 1.40950095 | TNFAIP8L3 |
| 15 | 51057983 | 51058183 | 1.40950095 | RP11-108K3.1 |
| 15 | 51318654 | 51318807 | 1.012833947 | RP11-108K3.2 |
| 15 | 51318654 | 51318807 | 1.012833947 | CYP19A1 |
| 15 | 51384662 | 51384810 | 2.588864959 | GLDN |
| 15 | 51499379 | 51499578 | -2.996623447 | DMXL2 |
| 15 | 51549732 | 51549927 | -6.051643904 | DMXL2 |
| 15 | 52381130 | 52381316 | -3.325152514 | MYO5A |
| 15 | 52476078 | 52476275 | -6.818538829 | MYO5A |
| 15 | 55348472 | 55348671 | -3.625538769 | PIGB |
| 15 | 55348472 | 55348671 | -3.625538769 | CCPG1 |
| 15 | 55360419 | 55360642 | -1.839774695 | CCPG1 |
| 15 | 55360419 | 55360642 | -1.839774695 | DYX1C1-CCPG1 |
| 1 | 43926544 | 43926733 | 5.431318679 | ST3GAL3 |
| 15 | 55915731 | 55915930 | -2.867860678 | NEDD4 |
| 15 | 56123495 | 56123682 | -6.812854422 | RFX7 |
| 15 | 56399778 | 56399972 | 1.731738339 | TEX9 |
| 15 | 56403382 | 56403555 | 6.644577177 | TEX9 |
| 15 | 56820564 | 56820670 | -1.381480401 | ZNF280D |
| 15 | 56917101 | 56917257 | 2.526296237 | ZNF280D |
| 15 | 56917619 | 56917829 | 1.100906345 | ZNF280D |
| 15 | 56954392 | 56954606 | -2.573591562 | TCF12 |
| 15 | 56955097 | 56955281 | -2.737469301 | TCF12 |
| 15 | 57293449 | 57293607 | -1.041569022 | TCF12 |
| 15 | 57300580 | 57300784 | -2.567655951 | LINC00926 |
| 15 | 57646388 | 57646496 | 4.212141475 | MYZAP |
| 15 | 57646388 | 57646496 | 4.212141475 | GCOM1 |
| 15 | 57977254 | 57977374 | 2.680584863 | ALDH1A2 |
| 1 | 43947053 | 43947248 | 1.607478981 | IPO13 |
| 15 | 58110962 | 58111092 | 7.23634542 | ALDH1A2 |
| 15 | 58128518 | 58128641 | 6.584727649 | ALDH1A2 |
| 15 | 58131174 | 58131368 | 2.297648855 | ALDH1A2 |
| 15 | 58138318 | 58138440 | 9.27446174 | ALDH1A2 |
| 15 | 58138318 | 58138440 | 9.27446174 | AQP9 |
| 15 | 58184072 | 58184279 | 2.953623088 | ALDH1A2 |
| 15 | 58184072 | 58184279 | 2.953623088 | AQP9 |
| 15 | 58316144 | 58316338 | 1.151684427 | ALDH1A2 |
| 15 | 58847342 | 58847540 | -1.132089179 | FAM63B |
| 15 | 58851654 | 58851844 | -1.140831241 | FAM63B |
| 15 | 58987782 | 58987960 | 1.178928976 | RNF111 |
| 15 | 59125935 | 59126088 | -1.103226711 | RP11-59H7.3 |
| 15 | 59127576 | 59127736 | -1.162627457 | RP11-59H7.3 |
| 15 | 59128483 | 59128685 | -1.660527294 | RP11-59H7.3 |
| 15 | 59190397 | 59190597 | 7.040283866 | MYO1E |
| 15 | 59372661 | 59372788 | 8.346437135 | MYO1E |
| 15 | 59372661 | 59372788 | 8.346437135 | FAM81A |
| 15 | 59372805 | 59372958 | 8.74945576 | MYO1E |
| 15 | 59372805 | 59372958 | 8.74945576 | FAM81A |
| 1 | 44805787 | 44805984 | 1.379868451 | PLK3 |
| 1 | 44805787 | 44805984 | 1.379868451 | TCTEX1D4 |

| | | | | |
|---|---|---|---|---|
| 8 | 129352715 | 129352892 | -7.026060692 | CCDC26 |
| 8 | 129426268 | 129426440 | -8.400626828 | RP11-26E5.1 |
| 8 | 129426268 | 129426440 | -8.400626828 | CCDC26 |
| 8 | 129548322 | 129548523 | -5.389365227 | CCDC26 |
| 8 | 129628671 | 129628867 | -5.322253496 | CCDC26 |
| 8 | 129645972 | 129646115 | -2.866295718 | CCDC26 |
| 1 | 243126562 | 243126750 | 4.069166378 | CEP170 |
| 8 | 130054626 | 130054823 | 2.706466578 | ASAP1 |
| 8 | 130060751 | 130060954 | 1.986014136 | ASAP1 |
| 8 | 132123505 | 132123676 | -1.600562788 | KCNQ3 |
| 8 | 132988851 | 132989019 | 4.038407799 | TG |
| 8 | 133037906 | 133038232 | -3.493093365 | SLA |
| 8 | 133037906 | 133038232 | -3.493093365 | TG |
| 8 | 134573644 | 134573775 | -6.343687842 | ZFAT |
| 8 | 134602164 | 134602269 | -1.596857445 | ZFAT |
| 8 | 139742282 | 139742467 | -7.259326578 | TRAPPC9 |
| 8 | 139775788 | 139775971 | -1.885604137 | TRAPPC9 |
| 8 | 139851817 | 139851927 | 5.623370299 | TRAPPC9 |
| 8 | 139989440 | 139989554 | -4.511183907 | TRAPPC9 |
| 8 | 140097412 | 140097606 | -2.525403416 | PEG13 |
| 8 | 140097412 | 140097606 | -2.525403416 | TRAPPC9 |
| 8 | 140450976 | 140451185 | -1.221225528 | TRAPPC9 |
| 8 | 140634667 | 140634866 | 8.986388951 | AGO2 |
| 8 | 140636749 | 140636961 | 3.484934157 | ERICD |
| 8 | 140659424 | 140659559 | 3.93518236 | PTK2 |
| 8 | 141124858 | 141125058 | -1.617227688 | DENND3 |
| 8 | 141126135 | 141126320 | -1.19248249 | RP11-809O17.1 |
| 8 | 141126135 | 141126320 | -1.19248249 | DENND3 |
| 8 | 141127984 | 141128183 | -2.532402316 | DENND3 |
| 8 | 141127984 | 141128183 | -2.532402316 | RP11-809O17.1 |
| 8 | 141129013 | 141129237 | -8.57980265 | DENND3 |
| 8 | 141129013 | 141129237 | -8.57980265 | RP11-809O17.1 |
| 8 | 141129525 | 141129696 | -9.063468164 | DENND3 |
| 8 | 141129525 | 141129696 | -9.063468164 | RP11-809O17.1 |
| 8 | 141210877 | 141211079 | -1.061925403 | SLC45A4 |
| 8 | 141213160 | 141213299 | -8.499823868 | SLC45A4 |
| 8 | 141252383 | 141252581 | -8.540658495 | SLC45A4 |
| 8 | 141252383 | 141252581 | -8.540658495 | RP11-10J21.4 |
| 8 | 142308438 | 142308611 | -3.605872589 | TSNARE1 |
| 8 | 142663665 | 142663895 | -1.806768329 | JRK |
| 8 | 142664981 | 142665178 | -1.488014022 | JRK |
| 8 | 142666084 | 142666311 | -1.891235387 | JRK |
| 8 | 143008964 | 143009151 | -1.479089912 | RP11-273G15.2 |
| 8 | 143249259 | 143249453 | -1.884092162 | RP13-582O9.6 |
| 8 | 143249259 | 143249453 | -1.884092162 | ZFP41 |
| 8 | 143251083 | 143251275 | -1.517912412 | RP13-582O9.6 |
| 8 | 143251083 | 143251275 | -1.517912412 | ZFP41 |
| 8 | 143259654 | 143259852 | -2.47819235 | ZFP41 |
| 8 | 143259654 | 143259852 | -2.47819235 | RP13-582O9.6 |
| 8 | 143260369 | 143260567 | -2.237660158 | RP13-582O9.6 |
| 8 | 143260369 | 143260567 | -2.237660158 | ZFP41 |
| 8 | 143295910 | 143296098 | -1.980067919 | ZNF696 |
| 8 | 143366807 | 143366969 | -1.226105079 | RHPN1-AS1 |
| 8 | 143538650 | 143538841 | -1.479167377 | ZC3H3 |

| | | | | |
|---|---|---|---|---|
| 9 | 96388164 | 96388272 | -3.50188945 | ZNF367 |
| 9 | 96759155 | 96759404 | -2.627319402 | ZNF510 |
| 9 | 96926350 | 96926546 | 1.121986278 | MFSD14C |
| 9 | 97330473 | 97330630 | -5.483613129 | RP11-23J9.4 |
| 9 | 97330473 | 97330630 | -5.483613129 | CCDC180 |
| 1 | 247135015 | 247135212 | -3.742953348 | ZNF124 |
| 9 | 98015266 | 98015466 | -6.429476155 | ANP32B |
| 9 | 98124702 | 98124877 | -2.089156141 | CORO2A |
| 9 | 98255289 | 98255593 | -1.060536208 | TBC1D2 |
| 9 | 98732286 | 98732504 | -2.604890077 | ANKS6 |
| 9 | 98736249 | 98736436 | -3.934377733 | ANKS6 |
| 9 | 98783885 | 98784248 | -2.344864619 | ANKS6 |
| 9 | 98790068 | 98790168 | -2.278433022 | ANKS6 |
| 9 | 99149365 | 99149552 | -1.384249011 | TGFBR1 |
| 1 | 247165347 | 247165536 | -1.625166543 | ZNF124 |
| 9 | 100179414 | 100179543 | -1.981262647 | INVS |
| 9 | 100182033 | 100182232 | -1.75161473 | INVS |
| 9 | 100305809 | 100305915 | -4.993632006 | TEX10 |
| 9 | 100346904 | 100347106 | -1.459583941 | TEX10 |
| 9 | 101475896 | 101476142 | -1.928212764 | TMEM246-AS1 |
| 9 | 101475896 | 101476142 | -1.928212764 | TMEM246 |
| 9 | 101573177 | 101573370 | 10.18580857 | GRIN3A |
| 9 | 110137558 | 110137761 | 3.424636084 | AKAP2 |
| 9 | 110137558 | 110137761 | 3.424636084 | PALM2-AKAP2 |
| 9 | 110168553 | 110168769 | 3.757365135 | AKAP2 |
| 9 | 110168553 | 110168769 | 3.757365135 | PALM2-AKAP2 |
| 9 | 111026591 | 111026776 | -1.995889747 | LPAR1 |
| 9 | 111661245 | 111661373 | 3.451639671 | GNG10 |
| 9 | 111661245 | 111661373 | 3.451639671 | DNAJC25-GNG10 |
| 9 | 112574763 | 112574960 | -1.026800442 | KIAA1958 |
| 9 | 112659584 | 112659903 | -1.295505245 | KIAA1958 |
| 9 | 112868733 | 112868909 | -7.010186162 | SNX30 |
| 9 | 113373894 | 113374094 | -1.134536196 | HDHD3 |
| 9 | 113541487 | 113541674 | 3.726387507 | RGS3 |
| 9 | 113543683 | 113543867 | 3.8974289 | RGS3 |
| 9 | 114269724 | 114269927 | 3.948998337 | COL27A1 |
| 9 | 114377002 | 114377199 | -1.881554582 | AKNA |
| 9 | 114381155 | 114381358 | -1.709388676 | AKNA |
| 9 | 114789159 | 114789358 | 1.72557996 | TNFSF15 |
| 9 | 114790232 | 114790434 | 1.422778241 | TNFSF15 |
| 9 | 116698911 | 116699159 | -2.767637688 | ASTN2 |
| 9 | 116698911 | 116699159 | -2.767637688 | TRIM32 |
| 9 | 117712495 | 117712695 | 1.607348028 | TLR4 |
| 9 | 117713749 | 117713956 | 1.955695434 | TLR4 |
| 9 | 117714406 | 117714664 | 1.429894233 | TLR4 |
| 9 | 117715860 | 117716062 | 1.371103242 | TLR4 |
| 9 | 117716291 | 117716483 | 1.706589276 | TLR4 |
| 9 | 120778448 | 120778646 | -7.40839184 | FBXW2 |
| 9 | 120842930 | 120843128 | 8.762087192 | PSMD5 |
| 9 | 120842930 | 120843128 | 8.762087192 | PSMD5-AS1 |
| 9 | 120843957 | 120844157 | -5.312854422 | PSMD5-AS1 |
| 9 | 120845022 | 120845139 | -5.715226276 | PSMD5-AS1 |
| 9 | 120902797 | 120902951 | 7.441657215 | TRAF1 |
| 9 | 120904609 | 120904762 | 7.423624413 | TRAF1 |

| | | | | |
|---|---|---|---|---|
| 15 | 62017225 | 62017514 | -5.842593398 | VPS13C |
| 15 | 63305694 | 63305879 | -1.369412968 | APH1B |
| 1 | 45326954 | 45327192 | 1.845039603 | HPDL |
| 15 | 63591140 | 63591355 | -1.028605761 | USP3 |
| 15 | 63591140 | 63591355 | -1.028605761 | USP3-AS1 |
| 15 | 63620861 | 63621065 | -1.749027218 | HERC1 |
| 15 | 63908116 | 63908244 | -1.565963088 | DAPK2 |
| 15 | 64155929 | 64156130 | 1.243580418 | SNX22 |
| 15 | 64155929 | 64156130 | 1.243580418 | PPIB |
| 15 | 65615986 | 65616185 | -1.871136617 | SLC24A1 |
| 15 | 65616723 | 65616890 | -1.956921678 | SLC24A1 |
| 1 | 45345304 | 45345495 | -2.193027649 | TESK2 |
| 15 | 66702477 | 66702671 | 1.069993064 | SMAD6 |
| 15 | 67190796 | 67190985 | -1.051246212 | SMAD3 |
| 15 | 67407085 | 67407285 | -1.185964716 | IQCH-AS1 |
| 15 | 67407085 | 67407285 | -1.185964716 | IQCH |
| 15 | 67408144 | 67408352 | -1.154974471 | IQCH-AS1 |
| 15 | 67408144 | 67408352 | -1.154974471 | IQCH |
| 15 | 67422872 | 67423103 | -2.256578499 | IQCH |
| 15 | 67422872 | 67423103 | -2.256578499 | IQCH-AS1 |
| 15 | 67826449 | 67826609 | -2.270575971 | SKOR1 |
| 15 | 68184672 | 68184865 | -1.015393533 | PIAS1 |
| 15 | 68184672 | 68184865 | -1.015393533 | RP11-315D16.2 |
| 15 | 68193939 | 68194191 | 1.084108455 | PIAS1 |
| 15 | 68193939 | 68194191 | 1.084108455 | RP11-315D16.2 |
| 15 | 68193939 | 68194191 | 1.084108455 | CALML4 |
| 15 | 70049853 | 70050007 | 1.342413111 | TLE3 |
| 1 | 45608033 | 45608238 | 1.095127495 | NASP |
| 15 | 72046451 | 72046660 | 1.172342347 | MYO9A |
| 15 | 72140541 | 72140698 | -2.760884337 | RP11-2I17.4 |
| 15 | 72140541 | 72140698 | -2.760884337 | SENP8 |
| 15 | 72474581 | 72474783 | 9.133900838 | RP11-1007O24.3 |
| 15 | 72474581 | 72474783 | 9.133900838 | ARIH1 |
| 15 | 72751673 | 72751869 | 1.113107174 | ADPGK |
| 15 | 72783865 | 72784074 | 2.881770379 | ADPGK |
| 15 | 72783865 | 72784074 | 2.881770379 | ADPGK-AS1 |
| 15 | 73714197 | 73714397 | 1.783504245 | CD276 |
| 15 | 74175704 | 74175902 | 7.962545387 | RP11-665J16.1 |
| 15 | 74175704 | 74175902 | 7.962545387 | ISLR |
| 15 | 74361807 | 74361914 | 3.054630716 | CTD-2311M21.2 |
| 15 | 74361807 | 74361914 | 3.054630716 | CYP11A1 |
| 15 | 74410323 | 74410541 | 1.018084581 | SEMA7A |
| 15 | 74906982 | 74907162 | 3.179947973 | FAM219B |
| 15 | 75675990 | 75676211 | 1.631702069 | CSPG4 |
| 15 | 76259243 | 76259440 | -5.475642357 | ETFA |
| 15 | 76259243 | 76259440 | -5.475642357 | TYRO3P |
| 15 | 76259595 | 76259783 | -1.2410081 | TYRO3P |
| 15 | 76259595 | 76259783 | -1.2410081 | ETFA |
| 1 | 46608180 | 46608375 | 1.862149591 | MOB3C |
| 1 | 46608180 | 46608375 | 1.862149591 | MKNK1 |
| 15 | 77181310 | 77181429 | -6.724703307 | PEAK1 |
| 15 | 78280236 | 78280431 | -1.190185184 | WDR61 |
| 15 | 78280236 | 78280431 | -1.190185184 | DNAJA4 |
| 15 | 78981970 | 78982146 | -6.399337149 | RASGRF1 |

| | | | | |
|---|---|---|---|---|
| 8 | 143544496 | 143544663 | -1.163493316 | RP11-661A12.5 |
| 8 | 143559764 | 143559964 | -1.705539024 | GSDMD |
| 8 | 143560248 | 143560447 | -1.134813902 | GSDMD |
| 8 | 143573580 | 143573777 | -3.10080351 | RP11-661A12.9 |
| 8 | 143587928 | 143588087 | -10.42294408 | EEF1D |
| 8 | 143589780 | 143589979 | -2.19750899 | EEF1D |
| 8 | 143596800 | 143597035 | -3.073809543 | EEF1D |
| 8 | 143599505 | 143599703 | -1.409187078 | TIGD5 |
| 8 | 143599505 | 143599703 | -1.409187078 | EEF1D |
| 8 | 143632660 | 143632852 | -1.584280273 | RP11-661A12.12 |
| 1 | 243685247 | 243685380 | 5.488639962 | AKT3 |
| 8 | 143744738 | 143744856 | -1.796313458 | FAM83H-AS1 |
| 8 | 143770406 | 143770587 | -7.866322214 | FAM83H-AS1 |
| 8 | 143770406 | 143770587 | -7.866322214 | RP11-429J17.7 |
| 8 | 143922948 | 143923148 | -10.84059926 | PLEC |
| 1 | 243730449 | 243730649 | -3.266867883 | AKT3 |
| 1 | 243730449 | 243730649 | -3.266867883 | RP11-370K11.1 |
| 8 | 143985054 | 143985256 | 1.726675108 | PARP10 |
| 8 | 144102666 | 144102839 | -1.752602435 | SHARPIN |
| 8 | 144139683 | 144139905 | -1.673152513 | HGH1 |
| 8 | 144263475 | 144263696 | 7.695655364 | BOP1 |
| 1 | 244053903 | 244054100 | -1.949787084 | ZBTB18 |
| 8 | 144268114 | 144268314 | 6.754763804 | SCX |
| 8 | 144268114 | 144268314 | 6.754763804 | BOP1 |
| 8 | 144292687 | 144292886 | -9.375873357 | HSF1 |
| 8 | 144298636 | 144298828 | -1.6285671 | HSF1 |
| 8 | 144310522 | 144310709 | -6.093676037 | HSF1 |
| 8 | 144441171 | 144441392 | 7.120991575 | TONSL |
| 8 | 144450191 | 144450395 | -1.284037796 | CYHR1 |
| 1 | 244054559 | 244054759 | -2.250923138 | ZBTB18 |
| 8 | 144509120 | 144509318 | -2.489425594 | MFSD3 |
| 8 | 144520153 | 144520355 | -1.028145968 | LRRC14 |
| 8 | 144521175 | 144521376 | -1.020730522 | LRRC14 |
| 8 | 144524167 | 144524364 | -1.701837594 | LRRC24 |
| 8 | 144524167 | 144524364 | -1.701837594 | LRRC14 |
| 8 | 144548068 | 144548249 | -5.975642357 | ARHGAP39 |
| 8 | 144808311 | 144808560 | -2.016160644 | ZNF517 |
| 8 | 144810368 | 144810566 | 2.067090908 | ZNF517 |
| 8 | 144841416 | 144841628 | -1.033892762 | ZNF7 |
| 8 | 144841416 | 144841628 | -1.033892762 | COMMD5 |
| 9 | 14612 | 14813 | -1.407438244 | WASH1 |
| 9 | 112919 | 113117 | -1.101524109 | LINC01388 |
| 9 | 116629 | 116821 | -1.237645452 | FOXD4 |
| 1 | 244653692 | 244653895 | 7.872850291 | DESI2 |
| 9 | 214186 | 214387 | -1.568888433 | C9orf66 |
| 9 | 306981 | 307165 | 7.124037905 | DOCK8 |
| 9 | 710840 | 711042 | 1.621429007 | KANK1 |
| 9 | 712893 | 713094 | 1.720382636 | KANK1 |
| 1 | 244656943 | 244657125 | 7.91585155 | DESI2 |
| 9 | 2448325 | 2448431 | -6.007010235 | VLDLR-AS1 |
| 9 | 2655098 | 2655234 | -2.652661218 | VLDLR |
| 9 | 2719754 | 2719917 | -1.834811793 | KCNV2 |
| 9 | 3530417 | 3530641 | -1.533876915 | RFX3-AS1 |
| 1 | 244659575 | 244659779 | 8.919229582 | DESI2 |
| 9 | 120904788 | 120905009 | 7.593199132 | TRAF1 |
| 9 | 121100607 | 121100801 | -5.459431619 | CNTRL |
| 9 | 121377950 | 121378138 | -1.032065169 | RP11-162D16.2 |
| 9 | 121379029 | 121379228 | -1.215616331 | RP11-162D16.2 |
| 9 | 121379434 | 121379636 | -1.228682304 | RP11-162D16.2 |
| 9 | 122392268 | 122392475 | -1.316467969 | PTGS1 |
| 9 | 122820341 | 122820563 | -3.381582493 | PDCL |
| 9 | 123109651 | 123109850 | -6.296345308 | MIR600HG |
| 9 | 123109651 | 123109850 | -6.296345308 | STRBP |
| 9 | 123416587 | 123416744 | 7.387444994 | DENND1A |
| 9 | 124032571 | 124032768 | 2.545344858 | LHX2 |
| 9 | 124344587 | 124344787 | 6.928941914 | NEK6 |
| 9 | 125150476 | 125150604 | 3.011183907 | PPP6C |
| 1 | 12528250 | 12528408 | -4.829416199 | RP5-888M10.2 |
| 1 | 247451642 | 247451832 | 1.804285773 | OR2B11 |
| 9 | 127713818 | 127714014 | 1.780674972 | PTRH1 |
| 9 | 127713818 | 127714014 | 1.780674972 | CFAP157 |
| 9 | 127789587 | 127789787 | -1.376740565 | CDK9 |
| 9 | 127815257 | 127815459 | -1.836485151 | ENG |
| 9 | 127908294 | 127908492 | -2.29029203 | ST6GALNAC4 |
| 9 | 127921368 | 127921603 | -2.482017251 | PIP5KL1 |
| 9 | 128107581 | 128107852 | 1.010500114 | SLC25A25 |
| 9 | 128113415 | 128113601 | -1.719275043 | SLC25A25-AS1 |
| 9 | 128179135 | 128179324 | -2.021155402 | CIZ1 |
| 1 | 247876022 | 247876212 | -7.506465007 | TRIM58 |
| 9 | 128322482 | 128322672 | -6.60715956 | TRUB2 |
| 9 | 128322482 | 128322672 | -6.60715956 | COQ4 |
| 1 | 247895984 | 247896176 | -1.601891431 | OR2W3 |
| 9 | 129176763 | 129176962 | 11.98298085 | IER5L |
| 9 | 129176763 | 129176962 | 11.98298085 | RP11-247A12.8 |
| 9 | 129176763 | 129176962 | 11.98298085 | RP11-247A12.2 |
| 9 | 129488856 | 129489057 | -1.737693485 | LINC00963 |
| 9 | 129503375 | 129503672 | -1.474431102 | LINC00963 |
| 9 | 129635406 | 129635549 | -1.992791614 | NTMT1 |
| 9 | 129635406 | 129635549 | -1.992791614 | ASB6 |
| 9 | 131260691 | 131260891 | -2.23111704 | FAM78A |
| 9 | 131276367 | 131276596 | -2.331888793 | FAM78A |
| 1 | 248825151 | 248825342 | 1.96404388 | SH3BP5L |
| 9 | 131474792 | 131474996 | -2.436256658 | PRRC2B |
| 9 | 131495931 | 131496136 | -1.470292919 | PRRC2B |
| 1 | 248847259 | 248847474 | 1.948084017 | ZNF672 |
| 9 | 132905688 | 132905893 | -1.335215921 | TSC1 |
| 9 | 133128056 | 133128216 | 9.10490753 | RALGDS |
| 9 | 133417531 | 133417711 | -2.355485193 | REXO4 |
| 9 | 133417531 | 133417711 | -2.355485193 | ADAMTS13 |
| 9 | 133464443 | 133464585 | 6.678776002 | CACFD1 |
| 9 | 133465175 | 133465294 | 6.950056031 | CACFD1 |
| 9 | 133471615 | 133471830 | 4.633183937 | SLC2A6 |
| 9 | 133667759 | 133667935 | -2.737866715 | SARDH |
| 9 | 134026901 | 134027119 | -2.890786478 | LINC00094 |
| 9 | 134033314 | 134033513 | -4.020105714 | BRD3 |
| 9 | 134033314 | 134033513 | -4.020105714 | LINC00094 |
| 9 | 134034728 | 134034985 | -2.762711384 | BRD3 |
| 9 | 134036110 | 134036300 | -3.165231136 | BRD3 |
| 15 | 78981970 | 78982146 | -6.399337149 | RP11-16K12.1 |
| 15 | 78988381 | 78988577 | -1.084320542 | RASGRF1 |
| 15 | 79970801 | 79971002 | -1.861411513 | BCL2A1 |
| 15 | 81001132 | 81001312 | 8.990783641 | MESDC1 |
| 15 | 81001769 | 81001983 | 8.90482755 | MESDC1 |
| 15 | 81312877 | 81313066 | -5.058171981 | IL16 |
| 15 | 81312877 | 81313066 | -5.058171981 | STARD5 |
| 15 | 82043808 | 82043963 | 2.611196211 | MEX3B |
| 1 | 46718668 | 46718862 | 9.427689753 | EFCAB14 |
| 15 | 82786979 | 82787179 | -1.643107246 | FSD2 |
| 15 | 82846591 | 82846795 | 1.913050548 | HOMER2 |
| 15 | 83329149 | 83329322 | -3.745926548 | RP11-382A20.4 |
| 15 | 84605966 | 84606138 | 1.110849326 | ZSCAN2 |
| 15 | 84605966 | 84606138 | 1.110849326 | RP11-182J1.18 |
| 15 | 84639715 | 84639888 | -1.810307405 | WDR73 |
| 15 | 84639715 | 84639888 | -1.810307405 | SCAND2P |
| 1 | 841390 | 841584 | 1.539280361 | LINC01128 |
| 1 | 1598974 | 1599147 | 2.873077215 | FNDC10 |
| 1 | 47280811 | 47281009 | -1.528415041 | STIL |
| 15 | 88881396 | 88881588 | -5.704695468 | HAPLN3 |
| 15 | 89625069 | 89625256 | -1.317794287 | TICRR |
| 15 | 89625069 | 89625256 | -1.317794287 | KIF7 |
| 15 | 89629615 | 89629809 | 1.440677752 | KIF7 |
| 15 | 89629615 | 89629809 | 1.440677752 | TICRR |
| 15 | 89631808 | 89632001 | -1.732987232 | KIF7 |
| 15 | 89639827 | 89640017 | -4.72766361 | KIF7 |
| 15 | 89805974 | 89806196 | 1.499848291 | ANPEP |
| 15 | 90067181 | 90067379 | 1.65366702 | ZNF710 |
| 15 | 90240864 | 90241018 | -1.212855208 | GDPGP1 |
| 15 | 90961116 | 90961317 | 2.316066934 | RCCD1 |
| 15 | 92789105 | 92789377 | -1.422344536 | FAM174B |
| 1 | 50405737 | 50405841 | -1.475736094 | FCF1P6 |
| 15 | 97971355 | 97971565 | 3.598138759 | ARRDC4 |
| 15 | 97973299 | 97973485 | 3.695455558 | ARRDC4 |
| 15 | 99130878 | 99131057 | 1.364033572 | RP11-6O2.4 |
| 15 | 99130878 | 99131057 | 1.364033572 | SYNM |
| 15 | 99131583 | 99131752 | 1.715848252 | SYNM |
| 15 | 99131583 | 99131752 | 1.715848252 | RP11-6O2.4 |
| 15 | 99132580 | 99132773 | 2.41350354 | SYNM |
| 15 | 99133972 | 99134160 | 1.049191726 | SYNM |
| 15 | 99366224 | 99366502 | -1.002301139 | LRRC28 |
| 15 | 99368623 | 99368809 | -1.743675036 | LRRC28 |
| 1 | 50644666 | 50644861 | 6.175745705 | FAF1 |
| 15 | 99728019 | 99728152 | 1.605399896 | LYSMD4 |
| 15 | 99801907 | 99802012 | 1.788550191 | DNM1P46 |
| 15 | 100225581 | 100225840 | 1.213972492 | ADAMTS17 |
| 15 | 100349260 | 100349382 | 3.178776002 | SPATA41 |
| 15 | 100918404 | 100918602 | 1.766366271 | RP11-66B24.4 |
| 15 | 101068789 | 101068906 | -4.160964047 | LRRK1 |
| 15 | 101068789 | 101068906 | -4.160964047 | RP11-505E24.2 |
| 15 | 101072465 | 101072721 | -1.123899946 | LRRK1 |
| 15 | 101072465 | 101072721 | -1.123899946 | RP11-505E24.2 |
| 15 | 101170593 | 101170770 | 1.973433536 | RP11-424I19.2 |
| 1 | 50908674 | 50908990 | -1.297117089 | FAF1 |

| | | | | |
|---|---|---|---|---|
| 9 | 4662718 | 4662835 | -1.576858618 | PLPP6 |
| 9 | 4662718 | 4662835 | -1.576858618 | SPATA6L |
| 9 | 4726475 | 4726635 | -1.984524497 | AK3 |
| 9 | 4727056 | 4727267 | -1.330087565 | AK3 |
| 1 | 244701321 | 244701454 | -2.381676963 | DESI2 |
| 9 | 5034015 | 5034153 | -5.013952998 | JAK2 |
| 9 | 5035641 | 5035837 | -4.204695468 | JAK2 |
| 9 | 5037409 | 5037605 | -7.135074649 | JAK2 |
| 9 | 5113478 | 5113674 | -1.732931699 | JAK2 |
| 9 | 5113478 | 5113674 | -1.732931699 | IGHEP2 |
| 1 | 12208891 | 12209092 | 2.21570547 | TNFRSF1B |
| 9 | 5136535 | 5136719 | -1.46295949 | INSL6 |
| 9 | 5168511 | 5168705 | 1.380499643 | INSL6 |
| 9 | 5169562 | 5169803 | 1.942426162 | INSL6 |
| 9 | 5172239 | 5172437 | 2.532371382 | INSL6 |
| 9 | 5467826 | 5468026 | 3.35590939 | CD274 |
| 9 | 5539361 | 5539537 | 5.49540654 | PDCD1LG2 |
| 9 | 5569975 | 5570179 | 8.637407457 | PDCD1LG2 |
| 9 | 5779964 | 5780190 | -1.586794193 | ERMP1 |
| 9 | 5782783 | 5782975 | -1.060490196 | ERMP1 |
| 9 | 5895838 | 5895987 | -1.146864195 | MLANA |
| 9 | 5895838 | 5895987 | -1.146864195 | KIAA2026 |
| 9 | 5920503 | 5920670 | -1.043532648 | KIAA2026 |
| 9 | 5968169 | 5968370 | -1.079670009 | KIAA2026 |
| 1 | 245010214 | 245010367 | -4.02046928 | EFCAB2 |
| 9 | 5988462 | 5988626 | -6.162371555 | KIAA2026 |
| 9 | 6012411 | 6012584 | -1.438467536 | RANBP6 |
| 9 | 6013146 | 6013357 | -1.0450191 | RANBP6 |
| 1 | 245071155 | 245071354 | -7.769579406 | EFCAB2 |
| 9 | 12775342 | 12775543 | 1.71950197 | LURAP1L-AS1 |
| 9 | 12775342 | 12775543 | 1.71950197 | LURAP1L |
| 9 | 12821536 | 12821702 | 2.205232385 | LURAP1L |
| 9 | 15493245 | 15493367 | -4.669925001 | PSIP1 |
| 9 | 15744466 | 15744678 | -2.717569353 | CCDC171 |
| 9 | 17394653 | 17394799 | -1.335255794 | CNTLN |
| 9 | 19058144 | 19058410 | -1.910840048 | HAUS6 |
| 1 | 245091241 | 245091453 | -1.757372078 | EFCAB2 |
| 9 | 19372100 | 19372298 | 2.699873814 | DENND4C |
| 9 | 19451184 | 19451554 | -1.251743918 | ACER2 |
| 9 | 20345961 | 20346143 | -1.377842837 | MLLT3 |
| 1 | 245672071 | 245672239 | -1.983276758 | KIF26B |
| 9 | 32431347 | 32431547 | 7.616322884 | ACO1 |
| 9 | 32444532 | 32444758 | 5.119248302 | ACO1 |
| 9 | 32446367 | 32446565 | 7.564158301 | ACO1 |
| 9 | 32448674 | 32448867 | 6.239890132 | ACO1 |
| 9 | 32457123 | 32457325 | 2.998233874 | DDX58 |
| 1 | 245680643 | 245680745 | -2.482017251 | KIF26B |
| 9 | 33038678 | 33038883 | -1.343307688 | DNAJA1 |
| 9 | 33462603 | 33462804 | -1.462041018 | NOL6 |
| 1 | 245684733 | 245684930 | -5.134344096 | KIF26B |
| 9 | 33601296 | 33601496 | -1.543491504 | CYP4F26P |
| 1 | 245686296 | 245686464 | -2.611757106 | KIF26B |
| 9 | 34249062 | 34249253 | -8.907985561 | UBAP1 |
| 9 | 34256003 | 34256238 | -2.890005575 | KIF24 |
| 1 | 248918077 | 248918275 | -1.297289734 | PGBD2 |
| 9 | 135487933 | 135488052 | -1.943249403 | PPP1R26 |
| 9 | 136407173 | 136407372 | -1.339729005 | SDCCAG3 |
| 9 | 136438882 | 136439085 | -6.569775676 | INPP5E |
| 9 | 136613858 | 136614042 | -1.015577179 | LINC01451 |
| 9 | 136673495 | 136673637 | 6.13529367 | AGPAT2 |
| 9 | 136721215 | 136721422 | 1.401527392 | FAM69B |
| 9 | 136721215 | 136721422 | 1.401527392 | SNHG7 |
| 9 | 136723807 | 136723948 | -1.496455288 | SNHG7 |
| 9 | 136723807 | 136723948 | -1.496455288 | FAM69B |
| 9 | 136800719 | 136800956 | -1.572709896 | CCDC183 |
| 9 | 136800719 | 136800956 | -1.572709896 | RP11-216L13.18 |
| 9 | 136800719 | 136800956 | -1.572709896 | RP11-216L13.19 |
| 9 | 136839991 | 136840280 | 8.110850197 | RABL6 |
| 9 | 136925774 | 136925973 | 1.680034573 | TRAF2 |
| 9 | 136953261 | 136953458 | -1.312657727 | LCN12 |
| 9 | 137030804 | 137031006 | -2.040143894 | C9orf139 |
| 9 | 137030804 | 137031006 | -2.040143894 | FUT7 |
| 9 | 137062260 | 137062461 | -1.84734716 | RP11-229P13.22 |
| 9 | 137062260 | 137062461 | -1.84734716 | SAPCD2 |
| 9 | 137064107 | 137064277 | -2.128139039 | RP11-229P13.23 |
| 9 | 137064107 | 137064277 | -2.128139039 | SAPCD2 |
| 9 | 137082881 | 137083094 | -1.697380453 | UAP1L1 |
| 9 | 137085603 | 137085740 | -3.786113256 | MAN1B1-AS1 |
| 9 | 137164273 | 137164477 | -1.729550192 | GRIN1 |
| 9 | 137204493 | 137204730 | -1.020958532 | TMEM203 |
| 9 | 137486088 | 137486222 | -1.49943973 | PNPLA7 |
| 9 | 137555089 | 137555430 | -1.386824644 | DPH7 |
| 9 | 137582401 | 137582619 | -1.006041004 | ZMYND19 |
| 10 | 3122435 | 3122638 | 9.553074421 | PFKP |
| 9 | 137728232 | 137728446 | 9.925169736 | EHMT1 |
| X | 303021 | 303223 | -2.753418762 | PLCXD1 |
| X | 310152 | 310282 | -7.775463555 | GTPBP6 |
| X | 313480 | 313654 | -8.224106559 | GTPBP6 |
| X | 317088 | 317284 | -7.299607748 | GTPBP6 |
| 10 | 3123725 | 3123844 | 8.077804422 | PFKP |
| X | 350431 | 350614 | -5.350219859 | PPP2R3B |
| X | 1190725 | 1190921 | 3.06023858 | CRLF2 |
| X | 1309551 | 1309755 | -1.290084884 | CSF2RA |
| X | 1396395 | 1396528 | -1.447723126 | LINC00106 |
| 10 | 3124246 | 3124410 | 8.434604125 | PFKP |
| X | 1437018 | 1437226 | -1.252150027 | ASMTL |
| X | 1463169 | 1463370 | -2.399613413 | P2RY8 |
| X | 1600506 | 1600711 | -8.338639596 | AKAP17A |
| X | 1742581 | 1742818 | -1.863591065 | RP13-297E16.4 |
| 10 | 3781942 | 3782141 | 1.113596096 | KLF6 |
| X | 2220807 | 2221001 | -1.352376633 | DHRSX |
| X | 2489674 | 2489818 | 8.010652671 | DHRSX |
| X | 2489674 | 2489818 | 8.010652671 | ZBED1 |
| X | 2861933 | 2862105 | -7.807556163 | GYG2 |
| X | 2865549 | 2865711 | -2.554262228 | GYG2 |
| X | 2913981 | 2914143 | -1.396800883 | ARSD |
| X | 3821834 | 3822022 | -1.51994595 | RP11-706O15.1 |
| 10 | 5456784 | 5456982 | -1.635979167 | NET1 |
| 15 | 101188112 | 101188272 | -3.274032271 | CHSY1 |
| 15 | 101301958 | 101302145 | -1.569538073 | PCSK6 |
| 15 | 101301958 | 101302145 | -1.569538073 | RP11-299G20.2 |
| 15 | 101302867 | 101303177 | -1.122572909 | PCSK6 |
| 15 | 101302867 | 101303177 | -1.122572909 | RP11-299G20.2 |
| 15 | 101304756 | 101305036 | -1.296962369 | PCSK6 |
| 15 | 101304756 | 101305036 | -1.296962369 | RP11-299G20.2 |
| 15 | 101328856 | 101329086 | -1.892442372 | PCSK6 |
| 15 | 101329584 | 101329799 | -2.09855465 | PCSK6 |
| 1 | 50968370 | 50968562 | 7.488818845 | CDKN2C |
| 15 | 101751204 | 101751540 | -1.130718518 | DNM1P47 |
| 1 | 50969015 | 50969303 | 9.038658223 | CDKN2C |
| 15 | 101976319 | 101976471 | -1.209695989 | WASH3P |
| 16 | 57142 | 57371 | -1.074202118 | SNRNP25 |
| 16 | 112672 | 112822 | -6.175745705 | NPRL3 |
| 16 | 297961 | 298144 | 9.492937641 | AXIN1 |
| 16 | 520880 | 521066 | 1.340508062 | RAB11FIP3 |
| 16 | 547687 | 547913 | 1.8783982 | LA16c-366D1.3 |
| 16 | 547687 | 547913 | 1.8783982 | CAPN15 |
| 16 | 553454 | 553650 | 2.542998158 | LA16c-366D1.3 |
| 16 | 553454 | 553650 | 2.542998158 | CAPN15 |
| 16 | 589504 | 589677 | 3.944308212 | RAB40C |
| 16 | 614654 | 614815 | 1.019644566 | RAB40C |
| 16 | 627588 | 627761 | 1.382788127 | RAB40C |
| 16 | 720545 | 720758 | 2.666901186 | FAM173A |
| 16 | 785876 | 786065 | 3.618738172 | RPUSD1 |
| 16 | 888083 | 888416 | 1.287703862 | LMF1 |
| 16 | 968725 | 968902 | 5.86459793 | RP11-161M6.6 |
| 16 | 968725 | 968902 | 5.86459793 | LMF1 |
| 16 | 1257864 | 1257984 | -5.821251514 | RP11-616M22.5 |
| 16 | 1257864 | 1257984 | -5.821251514 | TPSD1 |
| 16 | 1349420 | 1349628 | 1.100768136 | BAIAP3 |
| 16 | 1349420 | 1349628 | 1.100768136 | TSR3 |
| 16 | 1434299 | 1434487 | 6.183161107 | CCDC154 |
| 16 | 1434861 | 1434963 | 6.090109991 | CCDC154 |
| 16 | 1453246 | 1453387 | 7.637061657 | CLCN7 |
| 16 | 1519866 | 1519980 | 6.84685237 | IFT140 |
| 16 | 1536787 | 1537033 | 2.762634242 | IFT140 |
| 16 | 1536787 | 1537033 | 2.762634242 | TMEM204 |
| 16 | 1732537 | 1732744 | 1.360323689 | MAPK8IP3 |
| 16 | 1776811 | 1777122 | 1.377424793 | SPSB3 |
| 16 | 1776811 | 1777122 | 1.377424793 | EME2 |
| 16 | 2089673 | 2089801 | 1.425002307 | PKD1 |
| 16 | 2108704 | 2108973 | 2.138298519 | PKD1 |
| 16 | 2111369 | 2111568 | 12.3072312 | PKD1 |
| 16 | 2177591 | 2177779 | 1.105691425 | CASKIN1 |
| 16 | 2177591 | 2177779 | 1.105691425 | TRAF7 |
| 1 | 51779389 | 51779586 | -1.536222818 | OSBPL9 |
| 16 | 2457514 | 2457704 | 4.863960227 | CCNF |
| 16 | 2457514 | 2457704 | 4.863960227 | RP11-715J22.4 |
| 16 | 2502552 | 2502778 | 1.102536186 | TBC1D24 |
| 16 | 2502552 | 2502778 | 1.102536186 | RP11-20I23.1 |
| 16 | 2571091 | 2571257 | 1.043731421 | RP11-20I23.6 |
| 16 | 2571091 | 2571257 | 1.043731421 | PDPK1 |

| | | | | | | | | | | | | | | |
|---|---|---|---|---|---|---|---|---|---|---|---|---|---|---|
| 9 | 34256831 | 34257042 | -2.994274592 | KIF24 | X | 7050089 | 7050238 | -1.747978747 | PUDP | 16 | 2597757 | 2598025 | 1.355177828 | CTD-3126B10.1 |
| 9 | 34257550 | 34257683 | -3.141648036 | KIF24 | X | 9502910 | 9503077 | -4.475642357 | TBL1X | 16 | 2597757 | 2598025 | 1.355177828 | PDPK1 |
| 9 | 34310886 | 34311092 | -2.285118193 | KIF24 | X | 9716250 | 9716510 | -1.201583038 | TBL1X | 16 | 2628885 | 2629071 | 2.986565083 | PDPK2P |
| 9 | 34367376 | 34367586 | -2.542609981 | KIAA1161 | 10 | 5639497 | 5639697 | -4.236043949 | ASB13 | 16 | 2628885 | 2629071 | 2.986565083 | AC141586.5 |
| 9 | 34370741 | 34370845 | -2.817586973 | KIAA1161 | X | 10122446 | 10122635 | 1.449535546 | WWC3 | 16 | 2637385 | 2637508 | 1.04600607 | PDPK2P |
| 9 | 34400582 | 34400798 | -1.209002417 | FAM219A | X | 11586420 | 11586562 | -2.24106982 | ARHGAP6 | 1 | 51840424 | 51840622 | -1.704904575 | NRDC |
| 9 | 34635466 | 34635670 | -2.066529653 | SIGMAR1 | X | 12885716 | 12885833 | 1.448418465 | TLR7 | 16 | 2971713 | 2971828 | 1.398071924 | PKMYT1 |
| 9 | 34680941 | 34681133 | -3.781989756 | RP11-195F19.9 | X | 12919410 | 12919597 | 5.038497235 | TLR8 | 16 | 2971713 | 2971828 | 1.398071924 | PAQR4 |
| 1 | 245702616 | 245702820 | 1.764018733 | KIF26B | X | 12920577 | 12920722 | 5.074168447 | TLR8 | 16 | 3013082 | 3013371 | 2.730101145 | CLDN9 |
| 9 | 35088878 | 35089080 | -1.133180309 | PIGO | X | 12921268 | 12921395 | 3.755419578 | TLR8 | 16 | 3013861 | 3014057 | 4.429425662 | CLDN9 |
| 9 | 35538064 | 35538256 | 2.222201301 | RUSC2 | 10 | 5640296 | 5640497 | -3.105568211 | ASB13 | 16 | 3084657 | 3084812 | 1.079348873 | RP11-473M20.9 |
| 9 | 35546876 | 35547069 | 1.860408548 | RUSC2 | X | 12921726 | 12921930 | 6.192253013 | TLR8 | 16 | 3140599 | 3140778 | -1.058765438 | ZNF213 |
| 9 | 35547610 | 35547813 | 1.674724699 | RUSC2 | X | 14020152 | 14020325 | -6.775373393 | GEMIN8 | 16 | 3242912 | 3243107 | -1.063256537 | MEFV |
| 9 | 35555290 | 35555480 | 1.671917044 | RUSC2 | X | 14682762 | 14682944 | -2.237866715 | GLRA2 | 16 | 3243294 | 3243492 | -1.990814564 | MEFV |
| 9 | 35609682 | 35609885 | 2.595392788 | CD72 | X | 14686640 | 14686832 | -2.132170302 | GLRA2 | 16 | 3282592 | 3282785 | -1.193846538 | ZNF263 |
| 9 | 35609682 | 35609885 | 2.595392788 | TESK1 | 10 | 5730608 | 5730804 | 1.377850838 | RP11-336A10.2 | 16 | 3412090 | 3412271 | -1.409226315 | LA16c-306E5.2 |
| 9 | 35649668 | 35649829 | 4.336212671 | SIT1 | 10 | 5730608 | 5730804 | 1.377850838 | FAM208B | 16 | 3486503 | 3486745 | 1.173043995 | NAA60 |
| 9 | 35660862 | 35661061 | 1.18367409 | ARHGEF39 | X | 15697329 | 15697439 | 5.902267201 | CA5B | 16 | 3486503 | 3486745 | 1.173043995 | LA16c-306E5.3 |
| 9 | 35660862 | 35661061 | 1.18367409 | CCDC107 | X | 15697329 | 15697439 | 5.902267201 | CA5BP1 | 16 | 3581382 | 3581589 | 1.128634164 | SLX4 |
| 9 | 35661688 | 35661885 | -5.431703844 | ARHGEF39 | X | 15784774 | 15784968 | -1.340347696 | CA5B | 16 | 3581382 | 3581589 | 1.128634164 | RP11-461A8.1 |
| 1 | 246147122 | 246147330 | -1.583636283 | SMYD3 | X | 19779786 | 19779891 | -3.52219706 | SH3KBP1 | 16 | 3582347 | 3582545 | 1.454046027 | SLX4 |
| 9 | 35724659 | 35724844 | -13.35997288 | TLN1 | X | 19914174 | 19914418 | -2.564316015 | CXorf23 | 16 | 3582347 | 3582545 | 1.454046027 | RP11-461A8.1 |
| 9 | 36293490 | 36293708 | 1.201049222 | CLTA | X | 20155168 | 20155370 | -1.446801351 | RPS6KA3 | 16 | 3661542 | 3661667 | 6.454665297 | DNASE1 |
| 1 | 246239415 | 246239581 | -1.338030817 | SMYD3 | X | 30559834 | 30560061 | 1.197925217 | CXorf21 | 16 | 3661542 | 3661667 | 6.454665297 | TRAP1 |
| 9 | 36413981 | 36414104 | -6.492476214 | RNF38 | X | 37727816 | 37727973 | -1.665627949 | XK | 16 | 3728427 | 3728630 | 1.070436933 | CREBBP |
| 9 | 36839531 | 36839722 | 1.647135733 | PAX5 | X | 37727816 | 37727973 | -1.665627949 | RP5-972B16.2 | 1 | 1615504 | 1615678 | 8.693684737 | RP11-345P4.9 |
| 9 | 36840212 | 36840430 | 2.674108718 | PAX5 | X | 38805197 | 38805396 | -2.335504592 | MID1IP1 | 1 | 1615504 | 1615678 | 8.693684737 | MIB2 |
| 9 | 37440978 | 37441177 | -1.804967589 | ZBTB5 | 10 | 5749119 | 5749369 | 1.105148069 | FAM208B | 16 | 3993653 | 3993755 | -1.21557161 | ADCY9 |
| 9 | 37503228 | 37503428 | -1.431739183 | POLR1E | X | 40074875 | 40075006 | -9.11777575 | BCOR | 16 | 4337374 | 4337544 | 2.363614485 | PAM16 |
| 9 | 37541374 | 37541572 | -1.133926355 | FBXO10 | X | 40172295 | 40172401 | -2.412214218 | BCOR | 16 | 4337374 | 4337544 | 2.363614485 | GLIS2 |
| 9 | 37541374 | 37541572 | -1.133926355 | RP11-613M10.8 | X | 40172770 | 40172959 | -5.962406252 | BCOR | 16 | 4337374 | 4337544 | 2.363614485 | RP11-295D4.1 |
| 1 | 246242156 | 246242331 | -1.463177398 | SMYD3 | X | 46472959 | 46473068 | -2.246527026 | KRBOX4 | 16 | 4381961 | 4382160 | 3.087366834 | CORO7-PAM16 |
| 9 | 37588810 | 37589008 | -2.823385672 | TOMM5 | X | 46600148 | 46600384 | -1.453279817 | SLC9A7 | 16 | 4381961 | 4382160 | 3.087366834 | CORO7 |
| 9 | 37588810 | 37589008 | -2.823385672 | RP11-613M10.8 | X | 46606441 | 46606645 | -1.591730729 | SLC9A7 | 16 | 4381961 | 4382160 | 3.087366834 | VASN |
| 9 | 37588810 | 37589008 | -2.823385672 | RP11-613M10.9 | 10 | 5784447 | 5784655 | -8.302189742 | GDI2 | 16 | 4383170 | 4383361 | 8.855672166 | CORO7 |
| 9 | 37765322 | 37765468 | 6.330889049 | RP11-613M10.9 | X | 47483550 | 47483836 | -2.179150242 | LINC01560 | 16 | 4383170 | 4383361 | 8.855672166 | CORO7-PAM16 |
| 9 | 37765322 | 37765468 | 6.330889049 | TRMT10B | X | 47904038 | 47904238 | -6.472904792 | ZNF81 | 16 | 4383170 | 4383361 | 8.855672166 | VASN |
| 9 | 38396117 | 38396315 | -3.901083277 | ALDH1B1 | X | 47915175 | 47915335 | -1.106366958 | ZNF81 | 16 | 4509284 | 4509475 | 1.529244758 | HMOX2 |
| 9 | 38396717 | 38396878 | -3.912144561 | ALDH1B1 | X | 47916341 | 47916510 | -1.253474077 | ZNF81 | 1 | 52389253 | 52389451 | 1.131441395 | ORC1 |
| 9 | 38397096 | 38397289 | -4.528386145 | ALDH1B1 | X | 47917955 | 47918151 | -1.245884911 | ZNF81 | 16 | 5042246 | 5042428 | 1.709576804 | RP11-165E7.1 |
| 9 | 38397938 | 38398135 | -4.030141898 | ALDH1B1 | X | 47921256 | 47921441 | -2.252764017 | ZNF81 | 16 | 5042246 | 5042428 | 1.709576804 | ALG1 |
| 9 | 38553897 | 38554095 | -2.479537664 | RP11-392E22.11 | X | 47976985 | 47977339 | -1.445457685 | ZNF182 | 16 | 5084695 | 5084882 | 1.288284019 | ALG1 |
| 9 | 38553897 | 38554095 | -2.479537664 | RP11-392E22.9 | X | 47976985 | 47977339 | -1.445457685 | ZNF81 | 16 | 5084695 | 5084882 | 1.288284019 | EEF2KMT |
| 9 | 38566467 | 38566704 | -2.533129579 | RP11-392E22.9 | X | 48485749 | 48485944 | -1.22341085 | FTSJ1 | 16 | 8783234 | 8783410 | -7.265081227 | TMEM186 |
| 9 | 38566467 | 38566704 | -2.533129579 | SNX18P3 | X | 48559703 | 48559827 | -1.609714673 | TBC1D25 | 16 | 8783234 | 8783410 | -7.265081227 | ABAT |
| 9 | 38566467 | 38566704 | -2.533129579 | RP11-392E22.11 | 10 | 5862212 | 5862409 | 1.749892308 | RPL12P28 | 16 | 8815737 | 8815842 | -4.704695468 | PMM2 |
| 9 | 38567530 | 38567698 | -2.971915713 | RP11-392E22.11 | 10 | 5862212 | 5862409 | 1.749892308 | ANKRD16 | 16 | 11549408 | 11549606 | 8.062868165 | LITAF |
| 9 | 38567530 | 38567698 | -2.971915713 | RP11-392E22.9 | X | 48700143 | 48700433 | -1.558620837 | SUV39H1 | 1 | 52525723 | 52525910 | -1.02700239 | ZCCHC11 |
| 9 | 38567530 | 38567698 | -2.971915713 | SNX18P3 | X | 48700143 | 48700433 | -1.558620837 | AF196970.3 | 16 | 11676467 | 11676670 | -1.86297501 | SNN |
| 9 | 38568712 | 38568919 | -2.691013557 | RP11-392E22.9 | X | 49001887 | 49002067 | 6.406088653 | GRIPAP1 | 16 | 15726703 | 15726874 | -3.997944131 | MYH11 |
| 9 | 38595678 | 38595878 | -2.814209882 | ANKRD18A | X | 51744207 | 51744353 | -1.954806882 | GSPT2 | 16 | 15726703 | 15726874 | -3.997944131 | AF001548.5 |
| 9 | 38620410 | 38620577 | -3.482067274 | FAM201A | X | 52947777 | 52947989 | -1.294286877 | FAM156A | 16 | 17236791 | 17236985 | -1.205838334 | XYLT1 |
| 9 | 38620410 | 38620577 | -3.482067274 | ANKRD18A | X | 53141928 | 53142160 | -2.276248783 | KANTR | 16 | 17237776 | 17237973 | -1.988922234 | XYLT1 |
| 9 | 38622192 | 38622509 | -2.933108903 | FAM201A | X | 53143006 | 53143206 | -2.646509361 | KANTR | 16 | 18385336 | 18385532 | 1.928461414 | PKD1P5 |

| | | | | | | | | | | | | | | |
|---|---|---|---|---|---|---|---|---|---|---|---|---|---|---|
| 9 | 39446877 | 39447060 | -1.068970299 | ZNF658B | X | 53143006 | 53143206 | -2.646509361 | ACTG1P10 | 16 | 18386554 | 18386822 | 1.524892088 | PKD1P5 |
| 9 | 39809440 | 39809656 | -2.293418198 | RP11-104G3.2 | X | 53145577 | 53145724 | -1.782392309 | KANTR | 16 | 19115513 | 19115715 | 1.360192952 | CTD-2349B8.1 |
| 9 | 39809440 | 39809656 | -2.293418198 | GLIDR | X | 53147160 | 53147360 | -2.964976474 | KANTR | 16 | 19115513 | 19115715 | 1.360192952 | ITPRIPL2 |
| 1 | 246271045 | 246271174 | -1.095765729 | SMYD3 | X | 53147931 | 53148082 | -5.94208526 | KANTR | 16 | 19183684 | 19183888 | 2.286311036 | SYT17 |
| 9 | 41135725 | 41135916 | -1.083817277 | CBWD7 | X | 53149790 | 53149975 | -3.840033221 | KANTR | 16 | 19267207 | 19267406 | 2.618907956 | SYT17 |
| 9 | 41135725 | 41135916 | -1.083817277 | FRG1HP | 10 | 5953049 | 5953245 | 8.542770179 | IL15RA | 1 | 1646380 | 1646579 | -6.28788779 | CDK11B |
| 9 | 41480221 | 41480421 | -1.282324492 | RP11-187C18.3 | X | 53236382 | 53236507 | -2.351329485 | IQSEC2 | 16 | 19714385 | 19714599 | 1.209339997 | AC002550.5 |
| 9 | 42950883 | 42951075 | -2.33371233 | CYP4F60P | X | 53250418 | 53250635 | -2.414793184 | IQSEC2 | 16 | 19714385 | 19714599 | 1.209339997 | KNOP1 |
| 9 | 61854589 | 61854771 | -2.685059857 | FAM27C | X | 53374456 | 53374657 | -1.361688345 | SMC1A | 16 | 20797825 | 20798004 | 1.033696422 | ERI2 |
| 9 | 61972662 | 61972843 | -1.315025195 | RP11-374M1.2 | X | 54173945 | 54174132 | 2.631517203 | FAM120C | 16 | 20859880 | 20860084 | 1.441024373 | ERI2 |
| 9 | 62801651 | 62801852 | -1.439094092 | LINC01410 | X | 55487195 | 55487411 | -3.005613628 | USP51 | 16 | 20859880 | 20860084 | 1.441024373 | DCUN1D3 |
| 9 | 62811048 | 62811154 | -1.009487358 | LINC01410 | X | 64190483 | 64190608 | 3.460006627 | AMER1 | 1 | 52873372 | 52873569 | -4.766664866 | ZYG11A |
| 9 | 62812986 | 62813246 | -2.362526136 | LINC01410 | X | 64191300 | 64191494 | 5.17501754 | AMER1 | 16 | 22325993 | 22326186 | 2.023001904 | POLR3E |
| 9 | 63581466 | 63581648 | 1.119166983 | RP11-452D2.1 | X | 64192180 | 64192388 | 2.095372691 | AMER1 | 16 | 22392808 | 22392978 | -5.254392581 | CDR2 |
| 9 | 63818002 | 63818170 | -2.304044792 | DUX4L50 | X | 64268624 | 64268789 | -5.112603718 | MTMR8 | 16 | 22392808 | 22392978 | -5.254392581 | MFSD13B |
| 9 | 66918214 | 66918381 | -1.121011799 | ZNF658 | X | 64381330 | 64381535 | -1.425845695 | MTMR8 | 16 | 22420909 | 22421116 | 1.612644806 | CDR2 |
| 9 | 66919568 | 66919739 | -8.199171444 | ZNF658 | X | 64381539 | 64381699 | 1.153919524 | MTMR8 | 16 | 22420909 | 22421116 | 1.612644806 | RRN3P3 |
| 9 | 67718526 | 67718733 | -2.408523233 | FAM27E3 | X | 68046720 | 68046992 | -2.987087777 | OPHN1 | 16 | 24570152 | 24570326 | -2.017104546 | RBBP6 |
| 9 | 68296132 | 68296309 | -1.04517924 | CBWD3 | 10 | 7163618 | 7163811 | -1.015905131 | SFMBT2 | 16 | 24570785 | 24570985 | -2.695766396 | RBBP6 |
| 9 | 68780838 | 68781038 | -1.445721898 | FAM122A | X | 68840509 | 68840746 | -1.75128928 | EFNB1 | 16 | 24571958 | 24572158 | -1.030045338 | RBBP6 |
| 9 | 68780838 | 68781038 | -1.445721898 | PIP5K1B | 10 | 7197233 | 7197418 | 3.078049074 | SFMBT2 | 16 | 25032178 | 25032338 | 1.377853757 | RP11-266L9.4 |
| 9 | 68782493 | 68782637 | -1.982241325 | PIP5K1B | X | 71043721 | 71043907 | -1.52219706 | SOCS5P4 | 1 | 53009595 | 53009823 | 6.073602462 | SCP2 |
| 9 | 68782493 | 68782637 | -1.982241325 | FAM122A | X | 71096404 | 71096608 | -1.940354358 | FOXO4 | 16 | 25240183 | 25240379 | -1.143066218 | ZKSCAN2 |
| 1 | 246849732 | 246849929 | -1.567539871 | AHCTF1 | X | 71463664 | 71463866 | -2.762892282 | TAF1 | 16 | 27448880 | 27449106 | 1.113286356 | IL21R-AS1 |
| 9 | 69254360 | 69254557 | -2.399474021 | TJP2 | X | 71573735 | 71573946 | -1.048244679 | OGT | 16 | 27448880 | 27449106 | 1.113286356 | IL21R |
| 9 | 70387620 | 70387824 | 1.111635101 | KLF9 | 10 | 7411383 | 7411501 | 5.921764267 | SFMBT2 | 1 | 53010316 | 53010422 | -4.165458439 | SCP2 |
| 9 | 71685167 | 71685340 | 3.288916253 | TMEM2 | X | 71616248 | 71616369 | 1.857862223 | CXCR3 | 16 | 27778834 | 27779028 | 1.422629548 | KIAA0556 |
| 9 | 71745276 | 71745475 | 1.858422706 | TMEM2 | X | 72205061 | 72205287 | -5.6077665 | ERCC6L | 16 | 29145350 | 29145509 | -2.559750998 | RP11-426C22.5 |
| 9 | 71874762 | 71874962 | -1.081043847 | ABHD17B | X | 72205061 | 72205287 | -5.6077665 | PIN4 | 16 | 29154411 | 29154597 | 2.918303672 | RP11-426C22.5 |
| 1 | 246850616 | 246850838 | -1.059210545 | AHCTF1 | X | 72208481 | 72208678 | -5.89190838 | PIN4 | 16 | 29272242 | 29272446 | -1.276045126 | RP11-426C22.8 |
| 9 | 77721352 | 77721554 | 1.09294207 | GNAQ | X | 72208481 | 72208678 | -5.89190838 | ERCC6L | 16 | 29364730 | 29364951 | -1.322904028 | SNX29P2 |
| 9 | 78271357 | 78271488 | -2.257943353 | CEP78 | X | 72263451 | 72263650 | -1.310508645 | PIN4 | 16 | 29364730 | 29364951 | -1.322904028 | RP11-231C14.7 |
| 9 | 81689158 | 81689372 | -1.21838246 | TLE1 | X | 72263451 | 72263650 | -1.310508645 | RPS4X | 16 | 29665879 | 29666077 | 1.365880941 | SPN |
| 9 | 83903493 | 83903735 | -1.579091181 | KIF27 | X | 73999761 | 73999953 | -1.329891305 | JPX | 16 | 29665879 | 29666077 | 1.365880941 | AC009133.22 |
| 9 | 83939536 | 83939723 | -1.572633877 | C9orf64 | X | 73999761 | 73999953 | -1.329891305 | FTX | 16 | 29816703 | 29816901 | 8.992106303 | PAGR1 |
| 9 | 84001023 | 84001210 | -1.360881594 | RMI1 | X | 74004242 | 74004433 | -1.322874208 | FTX | 16 | 29816703 | 29816901 | 8.992106303 | AC009133.23 |
| 9 | 84001987 | 84002186 | -2.950395838 | RMI1 | X | 74004242 | 74004433 | -1.322874208 | JPX | 16 | 29816703 | 29816901 | 8.992106303 | AC009133.12 |
| 1 | 246969514 | 246969673 | -8.654485 | ZNF695 | X | 74034224 | 74034422 | -1.330256767 | FTX | 16 | 29986525 | 29986728 | 1.267788321 | TAOK2 |
| 1 | 246969514 | 246969673 | -8.654485 | ZNF670-ZNF695 | X | 74034224 | 74034422 | -1.330256767 | JPX | 16 | 30355242 | 30355442 | 1.173590886 | CD2BP2 |
| 9 | 85632938 | 85633208 | -1.354520141 | AGTPBP1 | X | 74034863 | 74035063 | -1.530957583 | FTX | 16 | 30355242 | 30355442 | 1.173590886 | RP11-347C12.10 |
| 9 | 86048897 | 86049094 | -6.806549397 | GOLM1 | X | 74034863 | 74035063 | -1.530957583 | JPX | 16 | 30356808 | 30357024 | 1.110533023 | RP11-347C12.10 |
| 9 | 86301346 | 86301543 | 2.641935085 | ZCCHC6 | X | 74041637 | 74041840 | -1.316662761 | JPX | 16 | 30359345 | 30359538 | 1.118209606 | TBC1D10B |
| 9 | 86323489 | 86323671 | 1.826294382 | ZCCHC6 | X | 74041637 | 74041840 | -1.316662761 | FTX | 16 | 30524384 | 30524585 | 1.130323854 | ZNF768 |
| 9 | 86346393 | 86346610 | 1.527609698 | ZCCHC6 | X | 74044109 | 74044303 | -2.465990788 | FTX | 16 | 30532433 | 30532659 | 1.322787418 | ZNF747 |
| 9 | 86352778 | 86352979 | 1.418450719 | ZCCHC6 | X | 74044109 | 74044303 | -2.465990788 | JPX | 16 | 30532433 | 30532659 | 1.322787418 | AC002310.11 |
| 1 | 246986719 | 246986891 | -2.782463233 | ZNF695 | X | 74067847 | 74068057 | -1.90978748 | FTX | 16 | 30555478 | 30555671 | 9.103669247 | AC002310.13 |
| 1 | 246986719 | 246986891 | -2.782463233 | ZNF670-ZNF695 | X | 74067847 | 74068057 | -1.90978748 | JPX | 16 | 30555478 | 30555671 | 9.103669247 | ZNF764 |
| 9 | 87706709 | 87706931 | 3.525727501 | DAPK1 | X | 74072987 | 74073220 | -2.377188723 | FTX | 16 | 30569758 | 30569957 | 1.29174128 | ZNF688 |
| 9 | 87858039 | 87858364 | -2.524032554 | XXyac-YM21GA2.7 | X | 74075802 | 74075998 | -1.219338745 | FTX | 16 | 30775752 | 30775912 | 7.462545387 | RNF40 |
| 9 | 87921168 | 87921393 | -5.758475556 | SPATA31C1 | X | 74077280 | 74077456 | -2.329073424 | FTX | 16 | 30775752 | 30775912 | 7.462545387 | RP11-2C24.3 |
| 9 | 87922709 | 87922842 | -6.302486801 | SPATA31C1 | X | 74095049 | 74095245 | -1.318207648 | FTX | 16 | 30821249 | 30821380 | -1.098016786 | RP11-2C24.5 |
| 9 | 87923037 | 87923236 | -6.986932055 | SPATA31C1 | X | 74096647 | 74096847 | -2.672048721 | FTX | 16 | 30821249 | 30821380 | -1.098016786 | RP11-2C24.7 |
| 9 | 87967101 | 87967302 | -3.481330069 | CDK20 | X | 74097210 | 74097362 | -1.952589437 | FTX | 16 | 30851152 | 30851344 | -6.274410954 | BCL7C |
| 9 | 88129872 | 88130066 | -6.754514686 | SPATA31C2 | X | 74099656 | 74099807 | -1.837806106 | FTX | 1 | 54173409 | 54173623 | 1.018262938 | RP11-446E24.4 |

| | | | | |
|---|---|---|---|---|
| 9 | 88131620 | 88131819 | -6.566099234 | SPATA31C2 |
| 1 | 246987138 | 246987338 | -2.664431443 | ZNF670-ZNF695 |
| 1 | 246987138 | 246987338 | -2.664431443 | ZNF695 |
| 9 | 89361460 | 89361656 | -3.913735819 | SEMA4D |
| 9 | 89378757 | 89378952 | -2.106507597 | SEMA4D |
| 1 | 247038035 | 247038243 | -3.444622538 | ZNF670-ZNF695 |
| 1 | 247038035 | 247038243 | -3.444622538 | ZNF670 |
| 9 | 90896058 | 90896434 | -1.551965231 | SYK |
| 9 | 91229442 | 91229602 | 6.452317311 | AUH |
| 9 | 91409385 | 91409693 | 1.454170483 | NFIL3 |
| 9 | 92226237 | 92226383 | -6.825525846 | IARS |
| 9 | 92314430 | 92314624 | -1.353451448 | NOL8 |
| 9 | 92549034 | 92549153 | -2.632126192 | CENPP |
| 9 | 92574787 | 92574987 | -7.247427792 | CENPP |
| 9 | 92614118 | 92614318 | -4.429935516 | CENPP |
| 9 | 92614118 | 92614318 | -4.429935516 | IPPK |
| 9 | 92615779 | 92615969 | -3.130068015 | CENPP |
| 9 | 92615779 | 92615969 | -3.130068015 | IPPK |
| 9 | 92618865 | 92619065 | -6.496940945 | IPPK |
| 9 | 92618865 | 92619065 | -6.496940945 | CENPP |
| 9 | 92631987 | 92632162 | -8.15999479 | IPPK |
| 9 | 92719331 | 92719526 | -1.321609417 | BICD2 |
| 9 | 92832229 | 92832396 | 5.612603718 | ANKRD19P |
| 9 | 92836858 | 92837060 | 1.718114176 | ANKRD19P |
| 9 | 92836858 | 92837060 | 1.718114176 | EEF1DP2 |
| 9 | 92847235 | 92847499 | 1.466722007 | ZNF484 |
| 9 | 92847235 | 92847499 | 1.466722007 | ANKRD19P |
| 9 | 93121781 | 93121999 | 1.003757362 | NINJ1 |
| 9 | 93951931 | 93952136 | -2.281549209 | BARX1 |
| 9 | 93953680 | 93953921 | -2.468951462 | BARX1 |
| 9 | 93959071 | 93959248 | -8.002680694 | BARX1-AS1 |
| 9 | 94038000 | 94038126 | -2.663214744 | CYCSP24 |
| 9 | 94038000 | 94038126 | -2.663214744 | PTPDC1 |
| 9 | 94098294 | 94098494 | -1.142480007 | PTPDC1 |
| 9 | 94300884 | 94301086 | -1.111441219 | ZNF169 |
| 9 | 94459252 | 94459453 | -1.129498736 | MFSD14B |
| 9 | 94760280 | 94760536 | -1.340190484 | C9orf3 |
| 9 | 94772907 | 94773089 | -3.64604028 | C9orf3 |
| 9 | 95086815 | 95087008 | -3.005436087 | C9orf3 |
| 9 | 95881007 | 95881200 | -1.265956143 | ERCC6L2 |
| 9 | 95919807 | 95920005 | -1.02219706 | ERCC6L2 |
| 9 | 96320992 | 96321215 | 1.039486564 | SLC35D2 |
| 9 | 96388164 | 96388272 | -2.173816816 | ZNF367 |
| 9 | 96816829 | 96816989 | -2.979210448 | ZNF782 |
| 9 | 96817922 | 96818118 | -1.23164786 | ZNF782 |
| 9 | 96819318 | 96819501 | -1.249475614 | ZNF782 |
| 9 | 96926350 | 96926546 | -2.390151203 | MFSD14C |
| 9 | 97330473 | 97330630 | -2.060615372 | CCDC180 |
| 9 | 97330473 | 97330630 | -2.060615372 | RP11-23J9.4 |
| 9 | 98015266 | 98015466 | -4.27285037 | ANP32B |
| 9 | 98083348 | 98083546 | -1.322261508 | TRIM14 |
| 9 | 98213191 | 98213393 | 3.082332927 | TBC1D2 |
| 9 | 98220868 | 98221053 | 3.859322381 | TBC1D2 |
| 9 | 98255289 | 98255593 | 2.869419004 | TBC1D2 |

| | | | | |
|---|---|---|---|---|
| X | 74100394 | 74100553 | -1.854117709 | FTX |
| X | 74161235 | 74161517 | -1.278877149 | FTX |
| X | 74161235 | 74161517 | -1.278877149 | MKRN5P |
| X | 74193104 | 74193282 | -3.165234383 | FTX |
| X | 74194748 | 74194920 | -2.15532226 | FTX |
| X | 75741148 | 75741303 | -3.225230733 | TTC3P1 |
| X | 75746132 | 75746333 | -6.870733493 | TTC3P1 |
| X | 76177623 | 76177785 | -2.427438238 | PBDC1 |
| X | 77682879 | 77683101 | -1.050762288 | ATRX |
| X | 77843748 | 77843932 | 7.066049016 | MAGT1 |
| X | 77846837 | 77847094 | 7.531676473 | MAGT1 |
| 10 | 11462636 | 11462841 | 1.181782227 | USP6NL |
| X | 85255454 | 85255648 | -7.706946164 | ZNF711 |
| X | 85270851 | 85271072 | -1.809865614 | ZNF711 |
| X | 91835592 | 91835791 | -1.772873052 | PCDH11X |
| X | 91876691 | 91876860 | -1.337779624 | PCDH11X |
| X | 91877705 | 91877898 | -2.705916689 | PCDH11X |
| X | 91878863 | 91879054 | -1.583212802 | PCDH11X |
| X | 92022227 | 92022411 | -2.780457141 | PCDH11X |
| X | 100670938 | 100671039 | -4.403677461 | SRPX2 |
| X | 100914695 | 100914894 | -7.124112045 | XKRX |
| X | 101346541 | 101346736 | -1.017868163 | TIMM8A |
| X | 101473010 | 101473206 | -6.11860498 | ARMCX4 |
| 10 | 11870466 | 11870681 | -2.278811197 | PROSER2-AS1 |
| 10 | 11870466 | 11870681 | -2.278811197 | PROSER2 |
| X | 101883045 | 101883244 | -1.37153361 | ZMAT1 |
| X | 101884488 | 101884686 | -1.522649173 | ZMAT1 |
| X | 102884288 | 102884472 | -1.139140105 | LL0XNC01-237H1.2 |
| X | 102884288 | 102884472 | -1.139140105 | LINC00630 |
| X | 103253677 | 103253887 | -2.53631233 | TCEAL8 |
| X | 103609272 | 103609475 | -1.421136848 | TCEAL3 |
| X | 103630108 | 103630359 | -1.798731973 | TCEAL1 |
| X | 103825526 | 103825722 | 1.254811978 | RAB9B |
| X | 107066436 | 107066631 | -7.199839235 | RBM41 |
| X | 107714123 | 107714323 | -1.391928134 | TSC22D3 |
| X | 107841001 | 107841241 | -1.333952372 | MID2 |
| X | 109624043 | 109624241 | 1.736727093 | KCNE5 |
| X | 111755236 | 111755415 | -1.029465246 | ALG13 |
| X | 113692854 | 113693039 | -4.552954254 | XACT |
| X | 113729978 | 113730203 | -6.74467312 | XACT |
| X | 113820181 | 113820330 | -2.697595182 | XACT |
| X | 113823081 | 113823237 | -1.970690137 | XACT |
| X | 113825461 | 113825646 | -1.351486106 | XACT |
| X | 113857907 | 113858153 | -1.541907739 | XACT |
| X | 113858929 | 113859130 | -1.839698651 | XACT |
| X | 113861627 | 113861812 | -3.436209189 | XACT |
| X | 113863003 | 113863202 | -1.415600832 | XACT |
| X | 113863660 | 113863792 | -1.057899485 | XACT |
| X | 113925342 | 113925541 | -2.590886719 | XACT |
| X | 115931948 | 115932146 | -2.826519363 | DANT2 |
| X | 117909725 | 117909941 | -6.892317423 | KLHL13 |
| X | 118392694 | 118392904 | 1.157708218 | WDR44 |
| X | 118998211 | 118998376 | -1.479119146 | LONRF3 |
| X | 119452997 | 119453199 | -1.319933741 | SLC25A43 |

| | | | | |
|---|---|---|---|---|
| 1 | 54173409 | 54173623 | 1.018262938 | CYB5RL |
| 16 | 30979492 | 30979689 | 10.68937973 | SETD1A |
| 16 | 31061847 | 31062027 | 10.34299204 | RP11-196G11.6 |
| 16 | 31061847 | 31062027 | 10.34299204 | ZNF668 |
| 16 | 31064217 | 31064413 | 1.598974112 | ZNF668 |
| 16 | 31073064 | 31073284 | 1.848194362 | ZNF668 |
| 16 | 31112068 | 31112267 | 3.535094135 | BCKDK |
| 16 | 31203036 | 31203228 | 2.750887499 | PYCARD |
| 16 | 31203036 | 31203228 | 2.750887499 | PYCARD-AS1 |
| 16 | 31311866 | 31312030 | -6.108372929 | ITGAM |
| 16 | 31358073 | 31358290 | -1.185878208 | ITGAX |
| 16 | 31435875 | 31435977 | 6.116809838 | ZNF843 |
| 16 | 31450925 | 31451118 | 1.073864805 | RP11-452L6.1 |
| 16 | 31457860 | 31458043 | 1.14150884 | RP11-452L6.5 |
| 16 | 31708116 | 31708258 | 1.132170302 | CLUHP3 |
| 16 | 31764729 | 31764839 | -1.079739107 | ZNF720 |
| 16 | 31772880 | 31773064 | -6.949678461 | ZNF720 |
| 16 | 31794140 | 31794331 | -1.136001435 | ZNF720 |
| 16 | 46581774 | 46581974 | 1.330282593 | SHCBP1 |
| 16 | 46597980 | 46598172 | 1.629323544 | SHCBP1 |
| 16 | 47083314 | 47083539 | 1.00465189 | NETO2 |
| 16 | 47557223 | 47557397 | 5.845435505 | PHKB |
| 16 | 48542893 | 48543084 | -1.767929783 | N4BP1 |
| 16 | 50153970 | 50154153 | 7.130129 | PAPD5 |
| 16 | 50211188 | 50211381 | 7.34281242 | PAPD5 |
| 16 | 50633565 | 50633761 | 2.464505915 | NKD1 |
| 16 | 50736516 | 50736714 | 5.781121212 | RP11-327F22.1 |
| 16 | 53280372 | 53280579 | 2.3561618 | CHD9 |
| 16 | 57747498 | 57747695 | 2.024488266 | KATNB1 |
| 16 | 57816669 | 57816871 | 1.11939343 | KIFC3 |
| 16 | 57816669 | 57816871 | 1.11939343 | CTD-2600O9.1 |
| 16 | 57821151 | 57821260 | 5.96831897 | KIFC3 |
| 16 | 57996217 | 57996402 | 3.837028214 | ZNF319 |
| 16 | 58012854 | 58013013 | 3.26865543 | USB1 |
| 16 | 66566520 | 66566722 | 1.769904149 | CKLF-CMTM1 |
| 16 | 66566520 | 66566722 | 1.769904149 | CMTM1 |
| 16 | 66921930 | 66922065 | 5.693470123 | RRAD |
| 16 | 67160270 | 67160416 | -4.906890596 | FBXL8 |
| 16 | 67160270 | 67160416 | -4.906890596 | TRADD |
| 16 | 67164351 | 67164537 | 7.433494545 | RP11-5A19.5 |
| 16 | 67164351 | 67164537 | 7.433494545 | FBXL8 |
| 16 | 67183345 | 67183534 | 1.628361745 | KIAA0895L |
| 16 | 67675602 | 67675860 | 1.349796835 | GFOD2 |
| 16 | 67724950 | 67725149 | 1.21669055 | RANBP10 |
| 16 | 67841916 | 67842123 | 2.221931442 | THAP11 |
| 16 | 67841916 | 67842123 | 2.221931442 | CENPT |
| 16 | 67945797 | 67945976 | 2.438872125 | SLC12A4 |
| 16 | 68268134 | 68268237 | -4.408491812 | SLC7A6 |
| 1 | 59586147 | 59586312 | -2.627788514 | FGGY |
| 16 | 68563938 | 68564150 | -1.627009749 | ZFP90 |
| 16 | 69109738 | 69109882 | 4.791655971 | HAS3 |
| 16 | 69116749 | 69116947 | 3.389397327 | HAS3 |
| 16 | 69666797 | 69666956 | -5.430543453 | NFAT5 |
| 16 | 69693353 | 69693581 | -1.025816502 | NFAT5 |

| | | | | |
|---|---|---|---|---|
| 9 | 98732286 | 98732504 | 1.196570821 | ANKS6 |
| 9 | 99149365 | 99149552 | 1.15323438 | TGFBR1 |
| 9 | 99828185 | 99828384 | 6.528313085 | NR4A3 |
| 9 | 99863507 | 99863714 | 10.83736187 | NR4A3 |
| 9 | 100179414 | 100179543 | -1.681202644 | INVS |
| 9 | 100182033 | 100182232 | -1.119055558 | INVS |
| 9 | 100292694 | 100292935 | -1.854213679 | INVS |
| 9 | 100300500 | 100300698 | -1.460617089 | INVS |
| 9 | 100305809 | 100305915 | -2.708389926 | TEX10 |
| 9 | 100346904 | 100347106 | -1.010385102 | TEX10 |
| 9 | 101390441 | 101390645 | -1.378414967 | MRPL50 |
| 9 | 101475896 | 101476142 | -1.878201883 | TMEM246-AS1 |
| 9 | 101475896 | 101476142 | -1.878201883 | TMEM246 |
| 9 | 103204452 | 103204650 | -1.723968466 | LINC01492 |
| 9 | 104784212 | 104784425 | 1.505022508 | ABCA1 |
| 9 | 110137558 | 110137761 | -2.822222596 | PALM2-AKAP2 |
| 9 | 110137558 | 110137761 | -2.822222596 | AKAP2 |
| 9 | 110168553 | 110168769 | -2.055262343 | PALM2-AKAP2 |
| 9 | 110168553 | 110168769 | -2.055262343 | AKAP2 |
| 9 | 111026591 | 111026776 | 1.799170765 | LPAR1 |
| 9 | 112574318 | 112574486 | -1.548291996 | KIAA1958 |
| 9 | 112574763 | 112574960 | -1.980913083 | KIAA1958 |
| 9 | 112618342 | 112618546 | -1.924323418 | KIAA1958 |
| 9 | 112659584 | 112659903 | -1.181204142 | KIAA1958 |
| 9 | 112890425 | 112890624 | 2.21059112 | SLC46A2 |
| 9 | 113163133 | 113163332 | 3.430910792 | FKBP15 |
| 9 | 113163133 | 113163332 | 3.430910792 | SLC31A2 |
| 9 | 113291536 | 113291683 | -2.221555669 | PRPF4 |
| 9 | 113292218 | 113292410 | -1.822535471 | PRPF4 |
| 9 | 113408880 | 113409084 | -1.490820551 | POLE3 |
| 9 | 113584309 | 113584502 | -2.087588769 | RGS3 |
| 9 | 113584309 | 113584502 | -2.087588769 | RP11-168K11.2 |
| 9 | 114168342 | 114168542 | -1.251296424 | COL27A1 |
| 9 | 114269724 | 114269927 | -4.245926548 | COL27A1 |
| 9 | 114790232 | 114790434 | 10.40732898 | TNFSF15 |
| 9 | 116698911 | 116699159 | -1.157314608 | ASTN2 |
| 9 | 116698911 | 116699159 | -1.157314608 | TRIM32 |
| 9 | 120439600 | 120439799 | -1.424975361 | CDK5RAP2 |
| 9 | 120453604 | 120453818 | -1.006478499 | CDK5RAP2 |
| 9 | 120605284 | 120605484 | -1.998683796 | MEGF9 |
| 9 | 120778448 | 120778646 | 1.419751878 | FBXW2 |
| 9 | 120902797 | 120902951 | 3.734557086 | TRAF1 |
| 9 | 120904609 | 120904762 | 1.727571068 | TRAF1 |
| 9 | 120904788 | 120905009 | 2.055998339 | TRAF1 |
| 9 | 121100607 | 121100801 | -1.802349359 | CNTRL |
| 9 | 121379029 | 121379228 | -1.679671591 | RP11-162D16.2 |
| 9 | 121379434 | 121379636 | -1.104594812 | RP11-162D16.2 |
| 9 | 121402694 | 121402888 | 5.196158711 | RP11-162D16.2 |
| 9 | 121989442 | 121989652 | -1.236817657 | TTLL11 |
| 9 | 122392268 | 122392475 | 6.870197663 | PTGS1 |
| 9 | 122820341 | 122820563 | 1.551848328 | PDCL |
| 9 | 122919366 | 122919565 | -1.498769935 | ZBTB26 |
| 9 | 123416587 | 123416744 | -7.336212671 | DENND1A |
| 9 | 124329731 | 124329931 | 8.024903008 | NEK6 |

| | | | | |
|---|---|---|---|---|
| 10 | 13209168 | 13209367 | -5.222507423 | MCM10 |
| X | 119759515 | 119759794 | -2.054369799 | SOWAHD |
| X | 119871377 | 119871576 | -1.578840117 | NDUFA1 |
| X | 119871377 | 119871576 | -1.578840117 | RNF113A |
| X | 120254657 | 120254881 | -1.489098271 | ZBTB33 |
| X | 121049147 | 121049348 | -1.336905423 | GLUD2 |
| 10 | 13317517 | 13317728 | -2.850913129 | SEPHS1 |
| X | 126165162 | 126165342 | 1.1761592 | DCAF12L2 |
| X | 126551839 | 126552069 | 7.618903737 | DCAF12L1 |
| X | 129793913 | 129794112 | -2.445257442 | SASH3 |
| X | 129794899 | 129795097 | -2.267125469 | SASH3 |
| X | 129794899 | 129795097 | -2.267125469 | RP4-753P9.3 |
| 10 | 13646359 | 13646569 | -3.970699404 | RP11-295P9.3 |
| 10 | 13646359 | 13646569 | -3.970699404 | RP11-295P9.13 |
| 10 | 13646359 | 13646569 | -3.970699404 | FRMD4A |
| X | 130014998 | 130015201 | -1.293717793 | BCORL1 |
| X | 130056105 | 130056230 | -1.154807719 | BCORL1 |
| X | 130067268 | 130067464 | 1.17877471 | ELF4 |
| X | 133753656 | 133753848 | -1.135314819 | GPC3 |
| 10 | 13657294 | 13657502 | -5.220434584 | RP11-295P9.3 |
| 10 | 13657294 | 13657502 | -5.220434584 | FRMD4A |
| X | 134838760 | 134838883 | 1.788140129 | FAM122C |
| X | 134991597 | 134991796 | -1.711261375 | SMIM10 |
| X | 135021595 | 135021828 | -2.244098255 | FAM127C |
| X | 135286051 | 135286245 | -1.08277328 | ZNF75D |
| X | 135293965 | 135294177 | -1.291646045 | ZNF75D |
| X | 135360247 | 135360464 | -1.773792483 | ZNF449 |
| 10 | 13707253 | 13707452 | -4.48674619 | FRMD4A |
| X | 136046666 | 136046825 | -1.974108729 | SLC9A6 |
| X | 143628937 | 143629108 | -3.030969666 | SLITRK4 |
| X | 143629422 | 143629720 | -2.377496133 | SLITRK4 |
| X | 143630403 | 143630567 | -7.324852633 | SLITRK4 |
| X | 145824699 | 145824806 | 3.075751591 | SLITRK2 |
| X | 149524707 | 149524870 | -2.625406658 | AF011889.2 |
| X | 149524707 | 149524870 | -2.625406658 | AF011889.5 |
| 10 | 14520920 | 14521192 | 1.238802191 | FAM107B |
| X | 149597036 | 149597238 | -1.345580952 | TMEM185A |
| X | 150469885 | 150470082 | 1.029663459 | MAMLD1 |
| 10 | 14567803 | 14567998 | 1.980194072 | FAM107B |
| X | 153571875 | 153572043 | -1.072098918 | ATP2B3 |
| X | 153580309 | 153580515 | -8.43518236 | ATP2B3 |
| X | 153724911 | 153725145 | 1.752784988 | ABCD1 |
| X | 153905400 | 153905604 | -1.539999661 | AVPR2 |
| X | 153905400 | 153905604 | -1.539999661 | L1CAM |
| X | 153971091 | 153971209 | 10.5519922 | HCFC1 |
| X | 154023551 | 154023751 | -1.23164901 | MECP2 |
| X | 154484364 | 154484536 | -3.372011859 | UBL4A |
| X | 155122128 | 155122301 | -1.031870903 | MTCP1 |
| X | 155122128 | 155122301 | -1.031870903 | BRCC3 |
| X | 155125879 | 155126078 | -1.614250914 | MTCP1 |
| X | 155127069 | 155127270 | -1.6695017 | MTCP1 |
| X | 155541913 | 155542222 | -2.724434165 | TMLHE-AS1 |
| X | 155541913 | 155542222 | -2.724434165 | TMLHE |
| X | 155773768 | 155774058 | 1.089436149 | SPRY3 |

| | | | | |
|---|---|---|---|---|
| 16 | 69695062 | 69695256 | -1.544428977 | NFAT5 |
| 16 | 69696381 | 69696484 | -1.270237696 | NFAT5 |
| 1 | 62436807 | 62436986 | 7.985951373 | USP1 |
| 16 | 70333552 | 70333756 | -1.140678456 | RP11-529K1.2 |
| 16 | 70333552 | 70333756 | -1.140678456 | DDX19B |
| 16 | 70333552 | 70333756 | -1.140678456 | RP11-529K1.3 |
| 16 | 70371958 | 70372160 | 1.095032834 | DDX19A |
| 16 | 70371958 | 70372160 | 1.095032834 | RP11-529K1.3 |
| 16 | 70381519 | 70381719 | 2.207287114 | RP11-529K1.4 |
| 16 | 70381519 | 70381719 | 2.207287114 | ST3GAL2 |
| 16 | 70699149 | 70699346 | 7.757110455 | VAC14 |
| 16 | 71285287 | 71285446 | -7.326142406 | CMTR2 |
| 16 | 71808928 | 71809084 | 1.857039119 | AP1G1 |
| 1 | 63457537 | 63457781 | 1.362446381 | ITGB3BP |
| 1 | 64559236 | 64559433 | -7.480136749 | CACHD1 |
| 16 | 72116532 | 72116730 | -6.763738503 | PMFBP1 |
| 16 | 72788489 | 72788608 | -1.143449486 | ZFHX3 |
| 16 | 72788489 | 72788608 | -1.143449486 | RP5-991G20.1 |
| 16 | 72795826 | 72795998 | -2.061395513 | ZFHX3 |
| 16 | 72798153 | 72798414 | -1.234404335 | ZFHX3 |
| 16 | 72958681 | 72958878 | -1.073911503 | ZFHX3 |
| 1 | 64691980 | 64692174 | -6.899640811 | CACHD1 |
| 16 | 74961805 | 74962013 | -1.261827648 | WDR59 |
| 16 | 75229896 | 75230097 | 1.964315456 | BCAR1 |
| 16 | 75233867 | 75234081 | 1.00289384 | BCAR1 |
| 1 | 64777795 | 64777948 | -1.583378861 | RAVER2 |
| 16 | 75234994 | 75235206 | 2.368852165 | BCAR1 |
| 16 | 75264329 | 75264477 | -4.646160816 | BCAR1 |
| 16 | 75435421 | 75435639 | 1.584258284 | RP11-77K12.1 |
| 16 | 75435421 | 75435639 | 1.584258284 | RP11-77K12.9 |
| 16 | 75502439 | 75502603 | -5.22766361 | RP11-77K12.5 |
| 16 | 77213489 | 77213687 | -4.084962501 | RP11-538I12.2 |
| 16 | 79600467 | 79600660 | -1.270354631 | MAF |
| 16 | 81765056 | 81765255 | -4.803665157 | PLCG2 |
| 16 | 81765056 | 81765255 | -4.803665157 | RP11-960L18.1 |
| 1 | 65966831 | 65967017 | -1.178998675 | PDE4B |
| 16 | 83939535 | 83939670 | 6.333112001 | RP11-505K9.4 |
| 16 | 85088434 | 85088600 | -6.719136028 | KIAA0513 |
| 16 | 87430109 | 87430308 | -1.089118994 | ZCCHC14 |
| 16 | 87491959 | 87492152 | 9.822768531 | ZCCHC14 |
| 16 | 87701723 | 87701912 | 1.538440986 | KLHDC4 |
| 16 | 87831006 | 87831207 | -1.591424599 | SLC7A5 |
| 1 | 1917820 | 1918013 | 1.974183616 | TMEM52 |
| 16 | 87832433 | 87832634 | -1.266317467 | SLC7A5 |
| 16 | 88048894 | 88049107 | 1.044013789 | BANP |
| 16 | 88245771 | 88245965 | -1.904007673 | LA16c-444G7.1 |
| 16 | 88438847 | 88439001 | 1.07552166 | ZNF469 |
| 16 | 88677929 | 88678134 | 1.031608372 | SNAI3-AS1 |
| 16 | 88677929 | 88678134 | 1.031608372 | SNAI3 |
| 16 | 88697197 | 88697325 | 10.70436933 | RP5-1142A6.5 |
| 16 | 88697197 | 88697325 | 10.70436933 | RNF166 |
| 16 | 88816571 | 88816731 | 1.069662639 | GALNS |
| 16 | 88829087 | 88829294 | 7.162055105 | GALNS |
| 16 | 88876833 | 88876965 | 3.588769093 | CBFA2T3 |

| | | | | |
|---|---|---|---|---|
| 9 | 124344587 | 124344787 | -5.492920969 | NEK6 |
| 9 | 124351807 | 124352007 | 2.626527919 | NEK6 |
| 9 | 124354431 | 124354629 | 3.300987333 | PSMB7 |
| 9 | 124354849 | 124355046 | 4.519215748 | PSMB7 |
| 9 | 124355576 | 124355771 | 3.016232251 | PSMB7 |
| 9 | 124356058 | 124356243 | 3.488514153 | PSMB7 |
| 9 | 124358126 | 124358352 | 2.810134355 | PSMB7 |
| 9 | 124358798 | 124358984 | 3.310887521 | PSMB7 |
| 9 | 124942968 | 124943149 | -7.044228502 | GOLGA1 |
| 9 | 124942968 | 124943149 | -7.044228502 | SCAI |
| 9 | 125150476 | 125150604 | -7.165178358 | PPP6C |
| 1 | 12528250 | 12528408 | 3.151022773 | RP5-888M10.2 |
| 9 | 125236606 | 125236805 | -1.210300503 | HSPA5 |
| 9 | 126503830 | 126504032 | -1.263752173 | MVB12B |
| 9 | 126505295 | 126505502 | -1.968458562 | MVB12B |
| 9 | 126696446 | 126696611 | -2.980322781 | LMX1B |
| 9 | 126833258 | 126833484 | 1.659838722 | ZBTB43 |
| 9 | 127445071 | 127445272 | -1.346290287 | ZNF79 |
| 9 | 127506810 | 127507023 | 3.31142551 | FAM129B |
| 9 | 127713818 | 127714014 | -3.156277795 | PTRH1 |
| 9 | 127713818 | 127714014 | -3.156277795 | CFAP157 |
| 9 | 127731735 | 127731940 | -1.450333142 | TOR2A |
| 9 | 127908294 | 127908492 | 1.553025804 | ST6GALNAC4 |
| 9 | 128061452 | 128061719 | -1.69934852 | NAIF1 |
| 9 | 128063191 | 128063401 | -1.85656888 | NAIF1 |
| 9 | 128179135 | 128179324 | -1.751368798 | CIZ1 |
| 9 | 128193112 | 128193311 | -1.711020546 | CIZ1 |
| 9 | 128193112 | 128193311 | -1.711020546 | DNM1 |
| 9 | 128322482 | 128322672 | 1.368280427 | TRUB2 |
| 9 | 128322482 | 128322672 | 1.368280427 | COQ4 |
| 9 | 128333630 | 128333804 | -1.58172378 | COQ4 |
| 9 | 128633696 | 128633877 | -3.216740659 | WDR34 |
| 9 | 128821293 | 128821563 | -1.688955055 | SPOUT1 |
| 9 | 128821293 | 128821563 | -1.688955055 | ENDOG |
| 9 | 128916325 | 128916550 | 1.433660634 | LRRC8A |
| 9 | 128945806 | 128946020 | -1.028936971 | DOLK |
| 9 | 128945806 | 128946020 | -1.028936971 | RP11-101E3.5 |
| 9 | 128947001 | 128947174 | -1.132963411 | RP11-101E3.5 |
| 9 | 128947001 | 128947174 | -1.132963411 | DOLK |
| 9 | 129177179 | 129177347 | 3.67421615 | RP11-247A12.8 |
| 9 | 129177179 | 129177347 | 3.67421615 | RP11-247A12.2 |
| 9 | 129177179 | 129177347 | 3.67421615 | IER5L |
| 9 | 129488856 | 129489057 | -1.58855399 | LINC00963 |
| 9 | 129510923 | 129511117 | 5.024839205 | LINC00963 |
| 9 | 129809413 | 129809614 | 1.110656286 | TOR1B |
| 9 | 130233243 | 130233462 | 1.159880198 | NCS1 |
| 9 | 130701569 | 130701739 | -2.815032215 | EXOSC2 |
| 9 | 130703700 | 130703897 | -2.627680306 | EXOSC2 |
| 9 | 130884097 | 130884288 | -1.069642934 | ABL1 |
| 9 | 130885403 | 130885709 | -1.159269794 | ABL1 |
| 9 | 131197690 | 131197892 | -1.938394614 | NUP214 |
| 9 | 131276367 | 131276596 | -1.026591867 | FAM78A |
| 1 | 248825151 | 248825342 | -2.946078521 | SH3BP5L |
| 9 | 131474792 | 131474996 | -2.375788194 | PRRC2B |
| X | 155776411 | 155776570 | 2.337782525 | SPRY3 |
| X | 156009947 | 156010145 | -1.106025239 | AJ271736.10 |
| X | 156009947 | 156010145 | -1.106025239 | IL9R |
| Y | 2924675 | 2924858 | -1.640368204 | RPS4Y1 |
| Y | 3069859 | 3070050 | -1.065622267 | LINC00278 |
| Y | 5098094 | 5098294 | -1.903677461 | PCDH11Y |
| Y | 13915745 | 13915936 | -1.682286216 | ANOS2P |
| Y | 14622121 | 14622300 | 3.152056373 | NLGN4Y |
| Y | 14622344 | 14622489 | 8.795271597 | NLGN4Y |
| Y | 14774778 | 14774926 | 7.631964678 | NLGN4Y |
| Y | 14830156 | 14830286 | 2.398506489 | NLGN4Y |
| Y | 14840725 | 14840935 | 1.844681823 | NLGN4Y |
| Y | 19008567 | 19008777 | -2.147146608 | TTTY14 |
| Y | 19077388 | 19077506 | -5.351086343 | TTTY14 |
| 10 | 14909143 | 14909329 | -1.283064572 | DCLRE1C |
| Y | 19706241 | 19706441 | -1.195322145 | KDM5D |
| 10 | 14910748 | 14910982 | -1.614878655 | DCLRE1C |
| 10 | 14911656 | 14911877 | -7.22830555 | DCLRE1C |
| 10 | 15860215 | 15860320 | 4.814102843 | FAM188A |
| 10 | 16484464 | 16484677 | -2.610402886 | PTER |
| 10 | 17215071 | 17215268 | -7.586682415 | VIM-AS1 |
| 10 | 20050099 | 20050297 | -6.730739724 | PLXDC2 |
| 10 | 20263293 | 20263411 | -1.875082312 | PLXDC2 |
| 10 | 20279723 | 20279938 | -1.455142702 | PLXDC2 |
| 10 | 22252107 | 22252248 | -2.995358181 | RP11-573G6.4 |
| 10 | 23121089 | 23121288 | -2.759496423 | MSRB2 |
| 10 | 23121541 | 23121757 | -1.544026164 | MSRB2 |
| 10 | 23440542 | 23440744 | 1.635911738 | OTUD1 |
| 10 | 24063096 | 24063286 | -5.947408882 | KIAA1217 |
| 10 | 25023791 | 25023986 | -7.088203465 | THNSL1 |
| 10 | 25023791 | 25023986 | -7.088203465 | ENKUR |
| 10 | 25598288 | 25598484 | -6.509013133 | GPR158 |
| 10 | 25599308 | 25599508 | -1.251218808 | GPR158 |
| 10 | 27035456 | 27035567 | -1.689630635 | ANKRD26 |
| 10 | 27170141 | 27170336 | 1.243483684 | MASTL |
| 10 | 28523484 | 28523687 | -1.740852421 | WAC-AS1 |
| 1 | 15492757 | 15492960 | -1.277251094 | CASP9 |
| 10 | 29550893 | 29551009 | 1.666687936 | SVIL |
| 10 | 30026157 | 30026347 | -5.078410231 | KIAA1462 |
| 10 | 30028168 | 30028351 | -3.023067989 | KIAA1462 |
| 10 | 30029155 | 30029350 | -4.06251619 | KIAA1462 |
| 10 | 30313385 | 30313750 | -1.198843045 | MTPAP |
| 10 | 30849196 | 30849412 | -1.094246265 | ZNF438 |
| 10 | 30928688 | 30928869 | 1.649950894 | ZNF438 |
| 10 | 31408847 | 31408971 | -2.478597839 | ZEB1 |
| 10 | 31474673 | 31474827 | 6.928990498 | ZEB1 |
| 10 | 31521531 | 31521761 | -1.10408976 | ZEB1 |
| 10 | 31891650 | 31891821 | -1.973344529 | ARHGAP12 |
| 10 | 35211313 | 35211519 | 3.394840894 | CREM |
| 10 | 37832392 | 37832577 | -1.49073407 | ZNF248 |
| 10 | 37832392 | 37832577 | -1.49073407 | AL135791.1 |
| 10 | 38054591 | 38054813 | -1.771860239 | ZNF33A |
| 10 | 38064031 | 38064292 | 1.399749194 | ZNF33A |
| 1 | 15734152 | 15734326 | 1.803557757 | RP11-288I21.1 |
| 16 | 89160496 | 89160694 | -1.704552156 | LINC00304 |
| 16 | 89215640 | 89215763 | 2.720100264 | RP11-46C24.7 |
| 16 | 89282931 | 89283121 | -1.404942169 | ANKRD11 |
| 16 | 89283717 | 89283918 | -1.042162927 | ANKRD11 |
| 16 | 89521100 | 89521299 | -6.73456651 | RP11-104N10.2 |
| 16 | 89521100 | 89521299 | -6.73456651 | SPG7 |
| 16 | 89596029 | 89596148 | 2.777759361 | CPNE7 |
| 16 | 89660189 | 89660411 | 1.886569603 | SPATA33 |
| 16 | 89686286 | 89686529 | 1.098917745 | RP11-368I7.4 |
| 16 | 89686286 | 89686529 | 1.098917745 | CDK10 |
| 16 | 89697814 | 89698040 | 3.463055465 | SPATA2L |
| 16 | 89707418 | 89707615 | 2.294964377 | VPS9D1 |
| 16 | 89717431 | 89717631 | 1.411180399 | VPS9D1-AS1 |
| 16 | 89717431 | 89717631 | 1.411180399 | VPS9D1 |
| 16 | 89730467 | 89730677 | 7.171696971 | ZNF276 |
| 17 | 231094 | 231255 | -5.518086806 | RPH3AL |
| 17 | 263678 | 263865 | -4.656441478 | RPH3AL |
| 17 | 270773 | 270908 | -2.988639962 | RPH3AL |
| 17 | 368173 | 368370 | -6.246302152 | RPH3AL |
| 17 | 634680 | 634826 | -5.281121212 | VPS53 |
| 17 | 670549 | 670713 | -6.018086806 | VPS53 |
| 17 | 753523 | 753667 | 1.406793438 | DBIL5P |
| 17 | 753523 | 753667 | 1.406793438 | GEMIN4 |
| 17 | 800795 | 800985 | -1.03735338 | NXN |
| 17 | 1148409 | 1148598 | -1.949630317 | ABR |
| 17 | 1423051 | 1423231 | 3.019872808 | CRK |
| 17 | 1423256 | 1423446 | 7.351627367 | CRK |
| 17 | 1423577 | 1423758 | -3.468657169 | CRK |
| 17 | 1575155 | 1575355 | 1.836593052 | SLC43A2 |
| 1 | 69905526 | 69905664 | 1.190103293 | LRRC7 |
| 17 | 1628171 | 1628347 | 4.011183907 | SLC43A2 |
| 17 | 1733650 | 1733862 | 2.39450898 | WDR81 |
| 17 | 1733650 | 1733862 | 2.39450898 | RP11-961A15.1 |
| 17 | 2395045 | 2395239 | 7.377261095 | MNT |
| 17 | 2690523 | 2690715 | 9.90753839 | CLUH |
| 1 | 75758576 | 75758802 | -4.974183616 | ACADM |
| 17 | 3570882 | 3571037 | 1.4198882 | TRPV1 |
| 17 | 3655288 | 3655490 | 1.338591741 | CTNS |
| 17 | 3657908 | 3658104 | 2.026556599 | CTNS |
| 17 | 3657908 | 3658104 | 2.026556599 | RP11-235E17.6 |
| 17 | 3669367 | 3669583 | 1.789952976 | P2RX5-TAX1BP3 |
| 17 | 3669367 | 3669583 | 1.789952976 | EMC6 |
| 17 | 3677813 | 3677985 | 1.171696761 | P2RX5 |
| 17 | 3677813 | 3677985 | 1.171696761 | P2RX5-TAX1BP3 |
| 1 | 77107148 | 77107325 | -5.974183616 | PIGK |
| 17 | 3913595 | 3913827 | 1.399775243 | P2RX1 |
| 17 | 4510778 | 4510981 | 1.522458573 | SPNS2 |
| 17 | 4539712 | 4539888 | 2.036871382 | MYBBP1A |
| 17 | 4672651 | 4672849 | 1.210813536 | PELP1 |
| 17 | 4731488 | 4731670 | 2.507475171 | MED11 |
| 1 | 2190579 | 2190686 | 3.004437914 | FAAP20 |
| 17 | 5210684 | 5210866 | 3.743244217 | RP11-333E1.1 |
| 17 | 5210684 | 5210866 | 3.743244217 | SCIMP |
| 17 | 6782462 | 6782653 | -1.258921811 | FBXO39 |

| | | | | | | | | | | | | | | |
|---|---|---|---|---|---|---|---|---|---|---|---|---|---|---|
| 9 | 131475762 | 131475963 | -1.102770559 | PRRC2B | 1 | 15734152 | 15734326 | 1.803557757 | PLEKHM2 | 17 | 6803471 | 6803669 | -6.240899716 | TEKT1 |
| 9 | 131524920 | 131525064 | -1.204909169 | UCK1 | 10 | 42337214 | 42337340 | -2.188002384 | RP11-313J2.1 | 17 | 6884295 | 6884437 | -5.765934388 | ALOX12-AS1 |
| 9 | 131579443 | 131579643 | 2.401532135 | RAPGEF1 | 10 | 42513974 | 42514153 | -1.278464073 | ZNF37BP | 17 | 6884295 | 6884437 | -5.765934388 | ALOX12P2 |
| 9 | 131626155 | 131626328 | 2.36265974 | RAPGEF1 | 10 | 42515373 | 42515573 | -1.822394571 | ZNF37BP | 17 | 6934679 | 6934877 | -1.612770125 | ALOX12-AS1 |
| 1 | 248847259 | 248847474 | -2.512450377 | ZNF672 | 10 | 42534747 | 42534960 | -5.980000966 | ZNF37BP | 17 | 6934679 | 6934877 | -1.612770125 | ALOX12P2 |
| 9 | 132375835 | 132376024 | -1.147423766 | TTF1 | 10 | 42594273 | 42594459 | -1.66366836 | ZNF33B | 1 | 77565572 | 77565761 | -1.166802524 | ZZZ3 |
| 9 | 132679356 | 132679606 | -1.762147248 | GTF3C4 | 10 | 42796520 | 42796720 | -2.255097366 | BMS1 | 17 | 7353344 | 7353547 | 1.095331855 | RP11-542C16.1 |
| 9 | 132689040 | 132689224 | -1.136627124 | GTF3C4 | 10 | 42831029 | 42831220 | -1.315451959 | BMS1 | 17 | 7353344 | 7353547 | 1.095331855 | KCTD11 |
| 1 | 248850064 | 248850277 | -2.516975869 | ZNF692 | 1 | 15836078 | 15836362 | 1.033514248 | FLJ37453 | 17 | 7383622 | 7383790 | -3.26569073 | TNK1 |
| 9 | 133153433 | 133153646 | -1.65108883 | GBGT1 | 10 | 43557518 | 43557746 | -2.893082966 | ZNF239 | 17 | 7849086 | 7849294 | 2.483176915 | KDM6B |
| 9 | 133331463 | 133331654 | -1.603452789 | SURF6 | 10 | 43616717 | 43616911 | -7.18295615 | ZNF485 | 17 | 8174973 | 8175172 | 6.296695561 | TMEM107 |
| 9 | 133417531 | 133417711 | -1.674699016 | REXO4 | 10 | 43617116 | 43617234 | -2.938333337 | ZNF485 | 17 | 8269262 | 8269427 | 1.205375729 | PFAS |
| 9 | 133417531 | 133417711 | -1.674699016 | ADAMTS13 | 10 | 43649092 | 43649193 | -5.573602462 | ZNF32-AS3 | 17 | 8369795 | 8369996 | -1.539001256 | RP11-849F2.7 |
| 1 | 248916800 | 248916945 | -1.425030933 | PGBD2 | 10 | 44994180 | 44994363 | 1.286410249 | RASSF4 | 17 | 8369795 | 8369996 | -1.539001256 | KRBA2 |
| 9 | 133667759 | 133667935 | 5.03534001 | SARDH | 10 | 45000169 | 45000398 | -1.599959871 | C10orf25 | 17 | 8369795 | 8369996 | -1.539001256 | RP11-849F2.5 |
| 9 | 134026901 | 134027119 | -1.035598794 | LINC00094 | 10 | 45003332 | 45003530 | -1.389564773 | ZNF22 | 17 | 8476004 | 8476204 | 1.409986735 | MYH10 |
| 9 | 134033314 | 134033513 | -1.303564605 | LINC00094 | 10 | 45003332 | 45003530 | -1.389564773 | CEP164P1 | 17 | 8476004 | 8476204 | 1.409986735 | NDEL1 |
| 9 | 134033314 | 134033513 | -1.303564605 | BRD3 | 10 | 45004787 | 45004972 | -2.967729874 | ZNF22 | 17 | 8879205 | 8879405 | 1.00928923 | PIK3R5 |
| 9 | 134034728 | 134034985 | -1.427713527 | BRD3 | 10 | 45004787 | 45004972 | -2.967729874 | CEP164P1 | 17 | 8888360 | 8888565 | 1.519096391 | PIK3R5 |
| 9 | 134036110 | 134036300 | -1.422590164 | BRD3 | 10 | 45458352 | 45458554 | -1.107892332 | Mar-08 | 17 | 9243384 | 9243560 | 5.299491486 | NTN1 |
| 1 | 248918077 | 248918275 | -1.561529981 | PGBD2 | 10 | 45826751 | 45826946 | -7.894894168 | AGAP4 | 17 | 13496230 | 13496413 | 2.257963868 | HS3ST3A1 |
| 9 | 135468293 | 135468442 | 3.308868768 | PPP1R26-AS1 | 1 | 15873893 | 15874086 | -2.256803878 | SPEN | 17 | 13496572 | 13496785 | 1.69209303 | HS3ST3A1 |
| 9 | 135504169 | 135504368 | -1.355557647 | MRPS2 | 10 | 46827446 | 46827646 | -6.082139219 | AGAP13P | 17 | 13601627 | 13601818 | 3.831573602 | HS3ST3A1 |
| 9 | 135504169 | 135504368 | -1.355557647 | RP11-426A6.5 | 10 | 48446464 | 48446660 | 1.260691192 | ARHGAP22 | 1 | 78628015 | 78628264 | -1.119786949 | IFI44L |
| 9 | 135933141 | 135933337 | -1.292703238 | UBAC1 | 10 | 48450640 | 48450825 | 2.178886647 | ARHGAP22 | 17 | 13777032 | 13777186 | 1.716121804 | COX10-AS1 |
| 9 | 136115101 | 136115303 | -1.704429834 | C9orf69 | 10 | 49515725 | 49515948 | -1.185878183 | PGBD3 | 17 | 14034105 | 14034302 | 1.335737318 | COX10-AS1 |
| 9 | 136378320 | 136378520 | -1.312195913 | SNAPC4 | 10 | 49515725 | 49515948 | -1.185878183 | ERCC6 | 17 | 14303791 | 14304002 | 1.804994607 | HS3ST3B1 |
| 9 | 136407173 | 136407372 | -1.679038583 | SDCCAG3 | 10 | 49933763 | 49933963 | -1.702317952 | PARG | 17 | 14303791 | 14304002 | 1.804994607 | RP11-214O1.2 |
| 9 | 136423225 | 136423470 | -1.208468271 | PMPCA | 10 | 50307041 | 50307223 | 1.202360234 | SGMS1 | 1 | 78650255 | 78650555 | -1.12573903 | IFI44 |
| 9 | 136438882 | 136439085 | -1.331002727 | INPP5E | 10 | 52731337 | 52731535 | 1.430217052 | RP11-556E13.1 | 17 | 15786646 | 15786836 | 2.576514648 | MEIS3P1 |
| 9 | 136497114 | 136497312 | 1.336846611 | NOTCH1 | 10 | 52734233 | 52734422 | 1.001627699 | RP11-556E13.1 | 17 | 15952463 | 15952586 | -1.657732501 | ADORA2B |
| 9 | 136613858 | 136614042 | 1.832764526 | LINC01451 | 10 | 58828875 | 58829050 | -1.504577358 | BICC1 | 17 | 15974981 | 15975138 | -1.594410826 | ADORA2B |
| 9 | 136614588 | 136614788 | 7.312569022 | LINC01451 | 10 | 59788968 | 59789179 | -1.369147159 | CCDC6 | 1 | 2229430 | 2229658 | 1.935176255 | SKI |
| 9 | 136673495 | 136673637 | -9.651490114 | AGPAT2 | 10 | 62053058 | 62053267 | 2.338576804 | ARID5B | 1 | 78654066 | 78654264 | -1.03725939 | IFI44 |
| 9 | 136721215 | 136721422 | -2.611593188 | FAM69B | 10 | 62197969 | 62198191 | -2.755028846 | RTKN2 | 17 | 16349030 | 16349230 | 1.809405863 | PIGL |
| 9 | 136721215 | 136721422 | -2.611593188 | SNHG7 | 10 | 62376553 | 62376750 | -1.427775438 | ZNF365 | 17 | 16349030 | 16349230 | 1.809405863 | CENPV |
| 9 | 136723807 | 136723948 | -3.953797594 | FAM69B | 10 | 62805681 | 62805881 | -1.478864974 | RP11-436D10.3 | 17 | 16443629 | 16443729 | -2.314250693 | LRRC75A |
| 9 | 136723807 | 136723948 | -3.953797594 | SNHG7 | 10 | 62805681 | 62805881 | -1.478864974 | ADO | 17 | 16443629 | 16443729 | -2.314250693 | LRRC75A-AS1 |
| 9 | 136829525 | 136829810 | -1.72358253 | RABL6 | 10 | 62813929 | 62814107 | -1.755353822 | EGR2 | 17 | 16471688 | 16471882 | -1.365120862 | LRRC75A-AS1 |
| 9 | 136831244 | 136831444 | -1.808542806 | RABL6 | 10 | 63153705 | 63153902 | -1.358482285 | NRBF2 | 17 | 16471688 | 16471882 | -1.365120862 | LRRC75A |
| 9 | 136840681 | 136840881 | -2.748982864 | RABL6 | 10 | 63206729 | 63206926 | -1.284033086 | JMJD1C | 1 | 81801125 | 81801372 | 1.156433572 | ADGRL2 |
| 9 | 136925774 | 136925973 | -2.385106128 | TRAF2 | 10 | 63213707 | 63213891 | -3.565887706 | JMJD1C | 17 | 17167640 | 17167808 | 1.066501323 | MPRIP |
| 9 | 136940769 | 136940932 | -2.91869958 | FBXW5 | 10 | 63214912 | 63215117 | -1.593265483 | JMJD1C | 1 | 81802439 | 81802632 | 2.690185215 | ADGRL2 |
| 9 | 136953261 | 136953458 | 2.94417493 | LCN12 | 10 | 64169560 | 64169665 | -1.851759556 | DBF4P1 | 17 | 17491581 | 17491809 | 1.97656744 | MED9 |
| 9 | 137028852 | 137029049 | -7.394155252 | ABCA2 | 10 | 65653141 | 65653302 | -6.280405234 | LINC01515 | 17 | 17494755 | 17494947 | 2.464296514 | RASD1 |
| 9 | 137028852 | 137029049 | -7.394155252 | C9orf139 | 10 | 65657245 | 65657419 | -5.336212671 | LINC01515 | 17 | 17846881 | 17847020 | -7.007184621 | TOM1L2 |
| 9 | 137030804 | 137031006 | -3.114287705 | FUT7 | 10 | 67912738 | 67912942 | -1.940532996 | SIRT1 | 17 | 17955525 | 17955692 | -4.577409055 | TOM1L2 |
| 9 | 137030804 | 137031006 | -3.114287705 | C9orf139 | 10 | 67916351 | 67916516 | -1.151414515 | SIRT1 | 17 | 18183477 | 18183672 | 4.57487356 | RP11-258F1.1 |
| 9 | 137062260 | 137062461 | -4.443023783 | RP11-229P13.22 | 10 | 68644897 | 68645091 | -5.459431619 | TET1 | 17 | 18183477 | 18183672 | 4.57487356 | ALKBH5 |
| 9 | 137062260 | 137062461 | -4.443023783 | SAPCD2 | 10 | 68691451 | 68691670 | -3.76974098 | TET1 | 17 | 18264452 | 18264609 | 2.108744178 | MIEF2 |
| 9 | 137062844 | 137063117 | -4.872462191 | SAPCD2 | 1 | 16231785 | 16231958 | -2.206444516 | RSG1 | 17 | 18274863 | 18275057 | 1.592736579 | TOP3A |
| 9 | 137064107 | 137064277 | -2.560992449 | SAPCD2 | 10 | 69123588 | 69123734 | 1.991496787 | VPS26A | 17 | 18535288 | 18535484 | -1.155982739 | CTD-2303H24.2 |
| 9 | 137064107 | 137064277 | -2.560992449 | RP11-229P13.23 | 10 | 70098540 | 70098738 | -1.832515073 | H2AFY2 | 17 | 18538590 | 18538786 | -1.911465889 | CCDC144B |
| 9 | 137164273 | 137164477 | -4.149557998 | GRIN1 | 10 | 70098540 | 70098738 | -1.832515073 | AIFM2 | 17 | 18538590 | 18538786 | -1.911465889 | CTD-2303H24.2 |

| | | | | |
|---|---|---|---|---|
| 9 | 137187685 | 137187884 | -1.208576643 | ANAPC2 |
| 9 | 137242536 | 137242736 | -2.700984491 | TUBB4B |
| 9 | 137279800 | 137280002 | -1.231500247 | TOR4A |
| 9 | 137486088 | 137486222 | 6.957379419 | PNPLA7 |
| 9 | 137486741 | 137486934 | 1.143763916 | PNPLA7 |
| 9 | 137488203 | 137488401 | 2.409935766 | PNPLA7 |
| 9 | 137555089 | 137555430 | -2.189679639 | DPH7 |
| 9 | 137582401 | 137582619 | -2.114270231 | ZMYND19 |
| 9 | 137612155 | 137612256 | 6.26116816 | ARRDC1 |
| 9 | 137612155 | 137612256 | 6.26116816 | RP11-48C7.8 |
| 9 | 137616037 | 137616239 | -1.100404015 | RP11-48C7.8 |
| 9 | 137616037 | 137616239 | -1.100404015 | ARRDC1-AS1 |
| 9 | 137617968 | 137618164 | -1.656848345 | RP11-48C7.8 |
| 9 | 137617968 | 137618164 | -1.656848345 | ARRDC1-AS1 |
| 10 | 3122435 | 3122638 | -8.961216709 | PFKP |
| X | 310152 | 310282 | 7.775463555 | GTPBP6 |
| X | 313480 | 313654 | 8.224106559 | GTPBP6 |
| X | 317088 | 317284 | 7.299607748 | GTPBP6 |
| X | 342081 | 342281 | 7.743480343 | PPP2R3B |
| X | 350431 | 350614 | 5.350219859 | PPP2R3B |
| X | 1309551 | 1309755 | -1.597718886 | CSF2RA |
| X | 1389344 | 1389543 | -2.310118988 | SLC25A6 |
| X | 1396395 | 1396528 | 1.665595343 | LINC00106 |
| 10 | 3124246 | 3124410 | -8.273115652 | PFKP |
| X | 1463169 | 1463370 | -2.619490074 | P2RY8 |
| X | 1465496 | 1465692 | -3.84051622 | P2RY8 |
| X | 1466283 | 1466391 | -4.498441702 | P2RY8 |
| 10 | 3781942 | 3782141 | 1.873714109 | KLF6 |
| X | 1746293 | 1746419 | 1.349415233 | RP13-297E16.4 |
| 10 | 4235487 | 4235696 | 8.07903855 | LINC00702 |
| X | 2306830 | 2307030 | 5.284927804 | DHRSX |
| X | 2488656 | 2488856 | -1.382538288 | DHRSX |
| X | 2488656 | 2488856 | -1.382538288 | ZBED1 |
| X | 2489674 | 2489818 | -7.215846734 | DHRSX |
| X | 2489674 | 2489818 | -7.215846734 | ZBED1 |
| X | 3822085 | 3822193 | -1.563862404 | RP11-706O15.1 |
| X | 3828757 | 3829082 | -1.617683774 | RP11-706O15.1 |
| 10 | 5456784 | 5456982 | -2.185793209 | NET1 |
| X | 7050089 | 7050238 | -2.623778219 | PUDP |
| X | 7350140 | 7350349 | -1.132516681 | STS |
| X | 8532845 | 8533045 | 1.167618145 | ANOS1 |
| X | 9502910 | 9503077 | -1.556729025 | TBL1X |
| X | 9719104 | 9719247 | -2.503670988 | TBL1X |
| X | 9932253 | 9932427 | 5.88975709 | SHROOM2 |
| X | 9946772 | 9946941 | 2.627157271 | SHROOM2 |
| 10 | 5639497 | 5639697 | -1.329441765 | ASB13 |
| X | 10117452 | 10117641 | 2.070844019 | WWC3 |
| X | 12976827 | 12977029 | -8.359273127 | TMSB4X |
| X | 13760304 | 13760517 | -1.265502898 | OFD1 |
| X | 14682762 | 14682944 | -2.181012945 | GLRA2 |
| X | 14686640 | 14686832 | -2.489364934 | GLRA2 |
| 10 | 5730608 | 5730804 | -1.312658461 | FAM208B |
| 10 | 5730608 | 5730804 | -1.312658461 | RP11-336A10.2 |
| X | 14835051 | 14835236 | -5.586057466 | NPM1P9 |
| 10 | 70139679 | 70139886 | -2.584384266 | TYSND1 |
| 10 | 70566729 | 70566950 | -1.551054997 | PALD1 |
| 10 | 70760662 | 70760861 | -1.612841876 | ADAMTS14 |
| 10 | 71362537 | 71362741 | -1.67690258 | SLC29A3 |
| 10 | 72008533 | 72008732 | 1.34230152 | CHST3 |
| 10 | 72275123 | 72275327 | -2.061961545 | DDIT4 |
| 10 | 72275123 | 72275327 | -2.061961545 | RP11-442H21.2 |
| 10 | 72335506 | 72335633 | -1.362662375 | DNAJB12 |
| 10 | 72354677 | 72354821 | 1.133939767 | DNAJB12 |
| 10 | 73125063 | 73125198 | 3.321928095 | NUDT13 |
| 10 | 73125063 | 73125198 | 3.321928095 | RP11-152N13.16 |
| 10 | 73166846 | 73166988 | -7.085431975 | ECD |
| 10 | 73498973 | 73499194 | -1.159063262 | PPP3CB-AS1 |
| 10 | 73498973 | 73499194 | -1.159063262 | USP54 |
| 10 | 73516582 | 73516782 | -2.676861089 | PPP3CB-AS1 |
| 10 | 73516582 | 73516782 | -2.676861089 | USP54 |
| 10 | 73647436 | 73647687 | -4.639224729 | SYNPO2L |
| 10 | 73648515 | 73648688 | -1.046078413 | SYNPO2L |
| 10 | 73675160 | 73675348 | -2.307373134 | AGAP5 |
| 10 | 73675160 | 73675348 | -2.307373134 | RP11-464F9.1 |
| 1 | 16472269 | 16472476 | -2.095199376 | CROCCP3 |
| 10 | 73704003 | 73704257 | -1.506050831 | RP11-464F9.1 |
| 10 | 73704003 | 73704257 | -1.506050831 | RP11-574K11.29 |
| 10 | 73704003 | 73704257 | -1.506050831 | BMS1P4 |
| 10 | 73731095 | 73731266 | -5.320122468 | GLUD1P3 |
| 10 | 73783088 | 73783278 | -2.196283157 | CHCHD1 |
| 10 | 73813896 | 73814098 | -2.552812996 | RP11-574K11.5 |
| 10 | 73813896 | 73814098 | -2.552812996 | CAMK2G |
| 10 | 75028771 | 75028969 | -8.663557007 | KAT6B |
| 10 | 75555751 | 75555953 | -7.54806931 | C10orf11 |
| 10 | 75580892 | 75581094 | -1.645849082 | C10orf11 |
| 10 | 75688217 | 75688476 | -1.812669131 | C10orf11 |
| 1 | 16597064 | 16597253 | -6.699371846 | NBPF1 |
| 10 | 76219264 | 76219434 | -1.887029256 | C10orf11 |
| 10 | 76222129 | 76222297 | -1.092128646 | C10orf11 |
| 10 | 76475350 | 76475511 | -5.114409345 | C10orf11 |
| 10 | 77780949 | 77781109 | -3.505089202 | RP13-39P12.2 |
| 10 | 77792211 | 77792435 | -1.719703105 | RP13-39P12.3 |
| 10 | 77792211 | 77792435 | -1.719703105 | DLG5 |
| 10 | 78054505 | 78054724 | -1.502637471 | RPS24 |
| 10 | 79019029 | 79019175 | -1.869246532 | ZMIZ1-AS1 |
| 10 | 79313341 | 79313541 | -1.30461594 | ZMIZ1 |
| 1 | 16602346 | 16602516 | -2.36503195 | NBPF1 |
| 10 | 79315278 | 79315471 | -1.565659504 | ZMIZ1 |
| 10 | 79353843 | 79354041 | 9.269999675 | PPIF |
| 10 | 79711698 | 79711873 | -1.112097332 | NUTM2B |
| 10 | 79711698 | 79711873 | -1.112097332 | NUTM2B-AS1 |
| 1 | 17068851 | 17069073 | -1.742989506 | PADI2 |
| 10 | 79958364 | 79958534 | -1.844852891 | SFTPD |
| 10 | 80522215 | 80522412 | -1.315322306 | TSPAN14 |
| 10 | 84371415 | 84371525 | 1.038068068 | CCSER2 |
| 10 | 84371920 | 84372128 | 1.305001652 | CCSER2 |
| 1 | 17115541 | 17115727 | -5.938833787 | PADI2 |
| 10 | 84442956 | 84443144 | 1.53470578 | CCSER2 |
| 17 | 18661871 | 18662028 | -4.63339327 | ZNF286B |
| 17 | 18661871 | 18662028 | -4.63339327 | TBC1D28 |
| 17 | 19003648 | 19003787 | 2.361468524 | SLC5A10 |
| 17 | 19003648 | 19003787 | 2.361468524 | FAM83G |
| 17 | 19003796 | 19003996 | 1.571084178 | SLC5A10 |
| 17 | 19003796 | 19003996 | 1.571084178 | FAM83G |
| 17 | 20828900 | 20829073 | -1.481555705 | ABHD17AP6 |
| 17 | 20946728 | 20946934 | -1.138224392 | CCDC144NL-AS1 |
| 17 | 28326607 | 28326808 | -1.258714055 | TMEM97 |
| 17 | 28724448 | 28724585 | 1.963512576 | TLCD1 |
| 17 | 28759485 | 28759675 | 1.141370228 | FAM222B |
| 17 | 28921783 | 28921978 | -1.033104536 | PHF12 |
| 17 | 29077718 | 29077901 | 6.422745025 | TIAF1 |
| 17 | 29077718 | 29077901 | 6.422745025 | MYO18A |
| 17 | 29078415 | 29078605 | 3.705779032 | MYO18A |
| 17 | 29078415 | 29078605 | 3.705779032 | TIAF1 |
| 17 | 29166098 | 29166279 | 2.13008833 | MYO18A |
| 17 | 29166558 | 29166750 | 1.217920317 | MYO18A |
| 17 | 29560837 | 29561015 | 1.064430317 | ABHD15 |
| 17 | 29560837 | 29561015 | 1.064430317 | ABHD15-AS1 |
| 17 | 29567811 | 29568006 | 2.415497873 | TP53I13 |
| 17 | 29567811 | 29568006 | 2.415497873 | ABHD15-AS1 |
| 17 | 29614277 | 29614559 | 2.91979763 | ABHD15-AS1 |
| 17 | 29614277 | 29614559 | 2.91979763 | ANKRD13B |
| 17 | 29632570 | 29632767 | -1.150710679 | SSH2 |
| 17 | 29632570 | 29632767 | -1.150710679 | ABHD15-AS1 |
| 17 | 29636323 | 29636521 | -2.616031025 | ABHD15-AS1 |
| 17 | 29636323 | 29636521 | -2.616031025 | SSH2 |
| 17 | 29742589 | 29742727 | -6.994076049 | SSH2 |
| 17 | 30864925 | 30865128 | -1.893454199 | ATAD5 |
| 17 | 30864925 | 30865128 | -1.893454199 | RP13-753N3.1 |
| 17 | 30970742 | 30970937 | -1.861305922 | RNF135 |
| 17 | 31193610 | 31193810 | 1.205652941 | NF1 |
| 17 | 31531613 | 31531813 | 1.042237754 | RAB11FIP4 |
| 17 | 32160006 | 32160207 | -2.071964896 | CTD-2095E4.3 |
| 17 | 32160006 | 32160207 | -2.071964896 | RHOT1 |
| 1 | 86627760 | 86627952 | -1.031004831 | RP4-651E10.4 |
| 17 | 35258752 | 35258955 | -1.863132853 | SLFN5 |
| 17 | 35259311 | 35259601 | -2.445307746 | SLFN5 |
| 17 | 35265401 | 35265612 | -2.036776563 | SLFN5 |
| 17 | 35409819 | 35410016 | -1.548771271 | RP11-686D22.7 |
| 17 | 35422473 | 35422683 | -1.144651684 | SLFN12 |
| 17 | 35480004 | 35480203 | -4.830889049 | SLFN12L |
| 1 | 86631734 | 86631911 | -1.383865613 | RP4-651E10.4 |
| 17 | 36537448 | 36537648 | 1.01616235 | PIGW |
| 17 | 36537448 | 36537648 | 1.01616235 | MYO19 |
| 17 | 36599153 | 36599346 | 2.41487159 | DHRS11 |
| 17 | 36997811 | 36998021 | -6.530601204 | AATF |
| 17 | 37406349 | 37406554 | 1.305385412 | ACACA |
| 17 | 37512374 | 37512489 | 1.432741147 | DUSP14 |
| 17 | 38466413 | 38466567 | 2.736393166 | ARHGAP23 |
| 17 | 38466586 | 38466759 | 4.505655288 | ARHGAP23 |
| 1 | 87104551 | 87104750 | -1.095880349 | HS2ST1 |
| 1 | 87104551 | 87104750 | -1.095880349 | RP5-1052I5.2 |

| | | | | |
|---|---|---|---|---|
| X | 15697329 | 15697439 | -7.170898386 | CA5B |
| X | 15697329 | 15697439 | -7.170898386 | CA5BP1 |
| X | 15700615 | 15700820 | -2.052755209 | CA5BP1 |
| X | 15700615 | 15700820 | -2.052755209 | CA5B |
| X | 15702511 | 15702755 | -1.84659991 | CA5BP1 |
| X | 15702511 | 15702755 | -1.84659991 | AC112497.1 |
| X | 15702511 | 15702755 | -1.84659991 | CA5B |
| X | 16596477 | 16596656 | -7.247302808 | CTPS2 |
| X | 16597179 | 16597400 | -8.347804464 | CTPS2 |
| X | 19779786 | 19779891 | -2.401723118 | SH3KBP1 |
| X | 19780413 | 19780601 | -5.777294426 | SH3KBP1 |
| X | 20155168 | 20155370 | 1.166892633 | RPS6KA3 |
| X | 21856951 | 21857175 | -1.504538819 | MBTPS2 |
| X | 21856951 | 21857175 | -1.504538819 | YY2 |
| X | 30559834 | 30560061 | -4.481108473 | CXorf21 |
| X | 36963214 | 36963405 | -1.062577566 | BX842568.2 |
| X | 38796931 | 38797097 | -1.722779722 | RP4-646N3.1 |
| 10 | 5749119 | 5749369 | -1.091735692 | FAM208B |
| X | 40172295 | 40172401 | -1.004705204 | BCOR |
| X | 40172770 | 40172959 | -1.363455965 | BCOR |
| 10 | 5783351 | 5783545 | -6.44208526 | GDI2 |
| X | 46597785 | 46597941 | 1.475964757 | CHST7 |
| X | 46600148 | 46600384 | 2.993186988 | SLC9A7 |
| X | 46606441 | 46606645 | 2.959622055 | SLC9A7 |
| X | 46625358 | 46625493 | -2.660964047 | SLC9A7 |
| X | 46647075 | 46647298 | 2.811454328 | PGAM1P7 |
| X | 46647075 | 46647298 | 2.811454328 | SLC9A7 |
| X | 46648011 | 46648220 | 1.560219248 | SLC9A7 |
| X | 47447301 | 47447514 | -1.14521763 | ZNF41 |
| X | 47448049 | 47448249 | -1.133472778 | ZNF41 |
| 10 | 5784447 | 5784655 | 2.152585733 | GDI2 |
| X | 47483550 | 47483836 | -1.121514404 | LINC01560 |
| X | 47915175 | 47915335 | 1.020128986 | ZNF81 |
| X | 48485749 | 48485944 | -1.11340802 | FTSJ1 |
| X | 48700143 | 48700433 | -2.556414861 | AF196970.3 |
| X | 48700143 | 48700433 | -2.556414861 | SUV39H1 |
| X | 48823062 | 48823263 | -2.543874519 | HDAC6 |
| X | 49001887 | 49002067 | -7.388293623 | GRIPAP1 |
| X | 49175653 | 49175874 | -1.052084453 | PRICKLE3 |
| X | 49881509 | 49881718 | -1.90773632 | USP27X |
| X | 50065695 | 50065879 | -1.278083955 | CLCN5 |
| X | 51744207 | 51744353 | 1.647869285 | GSPT2 |
| X | 53141928 | 53142160 | -1.71377052 | KANTR |
| X | 53143006 | 53143206 | -1.376122264 | KANTR |
| X | 53143006 | 53143206 | -1.376122264 | ACTG1P10 |
| X | 53145577 | 53145724 | -2.605637746 | KANTR |
| X | 53147160 | 53147360 | -1.817093215 | KANTR |
| X | 53147931 | 53148082 | -2.152093256 | KANTR |
| X | 53149790 | 53149975 | -1.157267902 | KANTR |
| X | 53169039 | 53169228 | -1.654376353 | RP11-258C19.7 |
| 10 | 5953049 | 5953245 | -3.394445658 | IL15RA |
| X | 53236382 | 53236507 | 3.086286399 | IQSEC2 |
| X | 53250418 | 53250635 | 1.66800361 | IQSEC2 |
| X | 53374456 | 53374657 | -1.034267959 | SMC1A |
| 10 | 84513630 | 84513828 | 1.257734071 | CCSER2 |
| 10 | 86943532 | 86943722 | -2.139835204 | MMRN2 |
| 10 | 86965900 | 86966103 | -1.453678856 | MMRN2 |
| 10 | 86965900 | 86966103 | -1.453678856 | ADIRF-AS1 |
| 10 | 86969127 | 86969326 | -1.47539332 | MMRN2 |
| 10 | 86969127 | 86969326 | -1.47539332 | ADIRF |
| 10 | 86969127 | 86969326 | -1.47539332 | ADIRF-AS1 |
| 10 | 86970797 | 86970992 | -1.062122168 | ADIRF-AS1 |
| 10 | 86970797 | 86970992 | -1.062122168 | AGAP11 |
| 10 | 86970797 | 86970992 | -1.062122168 | RP11-96C23.15 |
| 10 | 86970797 | 86970992 | -1.062122168 | ADIRF |
| 1 | 17407853 | 17408053 | -1.423743154 | RCC2 |
| 10 | 87009549 | 87009692 | -2.356448917 | AGAP11 |
| 10 | 87009549 | 87009692 | -2.356448917 | RP11-96C23.11 |
| 10 | 87009549 | 87009692 | -2.356448917 | RP11-96C23.14 |
| 10 | 87009549 | 87009692 | -2.356448917 | RP11-96C23.13 |
| 10 | 87342591 | 87342752 | 1.283078181 | NUTM2A-AS1 |
| 10 | 87342591 | 87342752 | 1.283078181 | LINC00863 |
| 10 | 87356639 | 87356915 | -1.42465666 | LINC00863 |
| 10 | 87366168 | 87366348 | -1.832941248 | NUTM2D |
| 10 | 87862731 | 87862915 | 1.294767615 | KLLN |
| 10 | 89306244 | 89306435 | 4.638988319 | LIPA |
| 10 | 89306244 | 89306435 | 4.638988319 | IFIT2 |
| 10 | 89307352 | 89307554 | 4.386875302 | IFIT2 |
| 10 | 89307352 | 89307554 | 4.386875302 | LIPA |
| 10 | 89338913 | 89339125 | 2.549865551 | IFIT3 |
| 10 | 89338913 | 89339125 | 2.549865551 | LIPA |
| 10 | 89402578 | 89402726 | 1.860461689 | LIPA |
| 10 | 89402578 | 89402726 | 1.860461689 | IFIT1 |
| 10 | 89418155 | 89418352 | 1.784096173 | IFIT5 |
| 10 | 89584285 | 89584464 | -5.56592848 | PANK1 |
| 10 | 89611797 | 89612076 | -2.81248909 | PANK1 |
| 10 | 89634284 | 89634481 | -4.092942447 | PANK1 |
| 10 | 89737432 | 89737620 | -2.105268032 | KIF20B |
| 10 | 89738107 | 89738306 | -1.607045177 | KIF20B |
| 10 | 89765356 | 89765476 | 3.160964047 | KIF20B |
| 10 | 91222697 | 91222894 | 3.124183856 | PCGF5 |
| 10 | 91259791 | 91259988 | 4.265065592 | PCGF5 |
| 10 | 91260491 | 91260691 | 2.385487406 | PCGF5 |
| 10 | 92959457 | 92959644 | -7.414464914 | EXOC6 |
| 10 | 94532952 | 94533149 | -2.253355528 | HELLS |
| 10 | 94532952 | 94533149 | -2.253355528 | TBC1D12 |
| 10 | 95874149 | 95874349 | -2.035252585 | RP11-248J23.7 |
| 10 | 95874149 | 95874349 | -2.035252585 | ENTPD1-AS1 |
| 10 | 95875264 | 95875463 | -1.955716338 | ENTPD1-AS1 |
| 10 | 95875264 | 95875463 | -1.955716338 | RP11-248J23.7 |
| 10 | 96058078 | 96058276 | 1.055075761 | CCNJ |
| 10 | 96058078 | 96058276 | 1.055075761 | ENTPD1-AS1 |
| 10 | 96665366 | 96665563 | 7.719265989 | PIK3AP1 |
| 10 | 96955491 | 96955697 | 1.062097182 | LCOR |
| 1 | 1083551 | 1083758 | 2.101522957 | C1orf159 |
| 1 | 19328868 | 19329104 | -1.409093788 | PQLC2 |
| 10 | 97320213 | 97320430 | -2.254756404 | FRAT1 |
| 10 | 97333512 | 97333710 | -2.765455776 | FRAT2 |
| 17 | 39759516 | 39759773 | 1.111339303 | IKZF3 |
| 1 | 87134146 | 87134348 | -1.143083189 | RP5-1052I5.2 |
| 1 | 87134146 | 87134348 | -1.143083189 | LINC01140 |
| 17 | 40362569 | 40362741 | 1.688684867 | CTD-2267D19.3 |
| 1 | 87153567 | 87153765 | -7.30158134 | RP5-1052I5.2 |
| 1 | 87153567 | 87153765 | -7.30158134 | LINC01140 |
| 17 | 40363732 | 40363941 | 1.98323747 | GJD3 |
| 17 | 40363732 | 40363941 | 1.98323747 | CTD-2267D19.3 |
| 17 | 40554494 | 40554684 | -1.054262228 | CCR7 |
| 17 | 40555708 | 40555903 | -2.481207031 | CCR7 |
| 17 | 41802588 | 41802724 | 2.038843048 | P3H4 |
| 17 | 41968318 | 41968521 | 2.122565098 | CNP |
| 17 | 42043211 | 42043364 | -1.362244294 | ZNF385C |
| 17 | 42264540 | 42264757 | -1.605521994 | STAT5B |
| 17 | 42668103 | 42668230 | -1.245452133 | PLEKHH3 |
| 17 | 43138398 | 43138587 | -1.008148682 | NBR2 |
| 17 | 43138398 | 43138587 | -1.008148682 | BRCA1 |
| 17 | 43162770 | 43162959 | 1.59592642 | BRCA1 |
| 17 | 43162770 | 43162959 | 1.59592642 | RP11-242D8.1 |
| 17 | 43399996 | 43400112 | 1.409918586 | ARL4D |
| 17 | 44690920 | 44691084 | 1.366191591 | RP11-1072C15.6 |
| 17 | 44926328 | 44926430 | -5.31467831 | KIF18B |
| 17 | 44946606 | 44946922 | -1.375722546 | KIF18B |
| 17 | 45150021 | 45150223 | 2.079776759 | HEXIM1 |
| 17 | 45190846 | 45191047 | -6.218355771 | CTD-2020K17.1 |
| 17 | 45218905 | 45219155 | 1.092281555 | CTD-2020K17.1 |
| 17 | 45264350 | 45264547 | 1.705765696 | MAP3K14-AS1 |
| 17 | 45264350 | 45264547 | 1.705765696 | MAP3K14 |
| 17 | 45515322 | 45515520 | -2.477098155 | LRRC37A4P |
| 17 | 46512220 | 46512390 | 3.903677461 | LRRC37A2 |
| 17 | 46768470 | 46768673 | -1.232731481 | WNT3 |
| 17 | 46959940 | 46960146 | -1.27827387 | GOSR2 |
| 17 | 46959940 | 46960146 | -1.27827387 | RP11-156P1.2 |
| 17 | 46968608 | 46968771 | -6.265081227 | RP11-156P1.2 |
| 17 | 47009512 | 47009758 | 2.324365685 | RP11-156P1.3 |
| 17 | 47009512 | 47009758 | 2.324365685 | LRRC37A17P |
| 17 | 47009512 | 47009758 | 2.324365685 | RP11-156P1.2 |
| 17 | 47321235 | 47321378 | -1.158687588 | RP11-290H9.2 |
| 17 | 47321235 | 47321378 | -1.158687588 | THCAT158 |
| 17 | 47387475 | 47387640 | -2.723246281 | EFCAB13 |
| 17 | 47437484 | 47437679 | -7.150890981 | EFCAB13 |
| 17 | 47898913 | 47899075 | 6.596146407 | SP2-AS1 |
| 17 | 47898913 | 47899075 | 6.596146407 | SP2 |
| 17 | 49317008 | 49317382 | -1.262828652 | ZNF652 |
| 17 | 49404398 | 49404599 | 1.333230419 | RP11-81K2.1 |
| 17 | 49404398 | 49404599 | 1.333230419 | PHB |
| 17 | 49404398 | 49404599 | 1.333230419 | RP11-1079K10.4 |
| 1 | 89426084 | 89426270 | 2.821928095 | GBP1P1 |
| 17 | 50149248 | 50149488 | 1.046895135 | PPP1R9B |
| 17 | 50352973 | 50353163 | -2.145001664 | XYLT2 |
| 17 | 50381091 | 50381276 | 1.182698978 | LRRC59 |
| 17 | 50381091 | 50381276 | 1.182698978 | EME1 |
| 17 | 50526092 | 50526377 | -2.482892142 | MYCBPAP |
| 17 | 56891966 | 56892167 | -1.144597287 | TRIM25 |

| | | | | |
|---|---|---|---|---|
| X | 54173945 | 54174132 | -4.321051641 | FAM120C |
| X | 55007196 | 55007398 | -1.714736049 | APEX2 |
| X | 55486675 | 55486878 | -1.512322064 | USP51 |
| X | 56564560 | 56564758 | -1.151561688 | UBQLN2 |
| X | 63350254 | 63350453 | -2.462288454 | SPIN4-AS1 |
| X | 63350254 | 63350453 | -2.462288454 | LINC01278 |
| X | 63350254 | 63350453 | -2.462288454 | SPIN4 |
| X | 64190483 | 64190608 | -3.123464036 | AMER1 |
| X | 64191300 | 64191494 | -2.959973374 | AMER1 |
| X | 64192180 | 64192388 | -1.560471648 | AMER1 |
| X | 64268624 | 64268789 | -2.218952936 | MTMR8 |
| X | 64311504 | 64311701 | -2.040085174 | MTMR8 |
| 1 | 1055055 | 1055216 | 2.235710298 | AGRN |
| 1 | 1055055 | 1055216 | 2.235710298 | RP11-54O7.14 |
| X | 64381539 | 64381699 | -1.698329485 | MTMR8 |
| X | 66021887 | 66022083 | 12.24092884 | VSIG4 |
| X | 67544702 | 67544899 | -2.182777465 | AR |
| X | 67546051 | 67546265 | -1.801001611 | AR |
| X | 67546273 | 67546373 | -1.607070328 | AR |
| X | 68046720 | 68046992 | -1.020692344 | OPHN1 |
| X | 68378372 | 68378547 | -1.006575684 | OPHN1 |
| X | 68718396 | 68718593 | 1.071859359 | STARD8 |
| X | 68719112 | 68719294 | 1.159015726 | STARD8 |
| X | 68840509 | 68840746 | 1.352950595 | EFNB1 |
| X | 69162787 | 69163012 | 1.395777511 | PJA1 |
| X | 69505816 | 69506015 | -1.038855779 | FAM155B |
| 10 | 7197233 | 7197418 | -3.03002584 | SFMBT2 |
| X | 70502463 | 70502662 | 2.059025581 | DLG3 |
| X | 71167353 | 71167540 | 1.835500415 | NLGN3 |
| X | 71183471 | 71183652 | -3.91339336 | RP5-1091N2.9 |
| X | 71463664 | 71463866 | 2.043509365 | TAF1 |
| X | 71544645 | 71544846 | 1.78722589 | OGT |
| X | 71616248 | 71616369 | -3.681949437 | CXCR3 |
| X | 71617386 | 71617498 | -3.069510356 | CXCR3 |
| X | 72128318 | 72128506 | 1.279026525 | NHSL2 |
| X | 72128318 | 72128506 | 1.279026525 | RGAG4 |
| X | 72205061 | 72205287 | -4.891739397 | PIN4 |
| X | 72205061 | 72205287 | -4.891739397 | ERCC6L |
| X | 72208481 | 72208678 | -3.749047581 | ERCC6L |
| X | 72208481 | 72208678 | -3.749047581 | PIN4 |
| X | 72263451 | 72263650 | 2.817023619 | RPS4X |
| X | 72263451 | 72263650 | 2.817023619 | PIN4 |
| X | 73999761 | 73999953 | -1.07905825 | JPX |
| X | 73999761 | 73999953 | -1.07905825 | FTX |
| X | 74004242 | 74004433 | -1.147263096 | JPX |
| X | 74004242 | 74004433 | -1.147263096 | FTX |
| 10 | 7412296 | 7412473 | 4.160362674 | RP5-1031D4.3 |
| X | 74034863 | 74035063 | 6.987028011 | JPX |
| X | 74034863 | 74035063 | 6.987028011 | FTX |
| X | 74044109 | 74044303 | 1.004324439 | JPX |
| X | 74044109 | 74044303 | 1.004324439 | FTX |
| X | 74067847 | 74068057 | 1.57064329 | FTX |
| X | 74067847 | 74068057 | 1.57064329 | JPX |
| X | 74068660 | 74068892 | 1.017110133 | FTX |
| 10 | 97338178 | 97338364 | -4.319881376 | RP11-452K12.4 |
| 10 | 97570323 | 97570593 | -2.435213954 | UBTD1 |
| 10 | 97675443 | 97675603 | 6.615910589 | PI4K2A |
| 10 | 97677645 | 97677840 | -2.233740154 | AVPI1 |
| 10 | 97714635 | 97714853 | -1.511908467 | MARVELD1 |
| 10 | 97717829 | 97718025 | -2.267785911 | MARVELD1 |
| 10 | 98223078 | 98223249 | -1.987873904 | R3HCC1L |
| 10 | 98247999 | 98248220 | -2.299361879 | LOXL4 |
| 10 | 99611096 | 99611297 | 2.568791597 | SLC25A28 |
| 10 | 99714508 | 99714696 | -3.10940423 | COX15 |
| 10 | 99714508 | 99714696 | -3.10940423 | CUTC |
| 10 | 99929748 | 99929997 | -2.625589024 | DNMBP-AS1 |
| 10 | 99929748 | 99929997 | -2.625589024 | DNMBP |
| 10 | 99955346 | 99955500 | -2.221047551 | DNMBP-AS1 |
| 10 | 99955346 | 99955500 | -2.221047551 | DNMBP |
| 10 | 99956354 | 99956477 | -1.044045665 | DNMBP |
| 10 | 99956354 | 99956477 | -1.044045665 | DNMBP-AS1 |
| 10 | 100382630 | 100382867 | -4.415105231 | OLMALINC |
| 10 | 101007202 | 101007395 | -1.471385436 | LZTS2 |
| 10 | 101067708 | 101067910 | -1.933238629 | KAZALD1 |
| 10 | 101136889 | 101137116 | -2.10063314 | TLX1 |
| 10 | 101553191 | 101553434 | -1.076836932 | BTRC |
| 10 | 101786268 | 101786458 | -1.634771681 | MGEA5 |
| 10 | 101798062 | 101798262 | 7.023220974 | MGEA5 |
| 10 | 102108083 | 102108271 | -1.846943747 | LDB1 |
| 10 | 102630136 | 102630343 | -1.328195715 | SUFU |
| 10 | 102657053 | 102657252 | -1.344849057 | TRIM8 |
| 10 | 102815060 | 102815162 | -1.004994652 | WBP1L |
| 10 | 102919788 | 102919988 | -1.440769507 | CNNM2 |
| 10 | 102927245 | 102927445 | -1.450268295 | CNNM2 |
| 10 | 103446942 | 103447143 | -1.84552051 | CALHM2 |
| 10 | 103450028 | 103450228 | -4.575955352 | CALHM2 |
| 10 | 103450028 | 103450228 | -4.575955352 | RP11-225H22.7 |
| 10 | 103600539 | 103600723 | -7.296345308 | SH3PXD2A |
| 10 | 103600947 | 103601182 | -1.696638554 | SH3PXD2A |
| 10 | 103601541 | 103601670 | -1.994886528 | SH3PXD2A |
| 10 | 103601826 | 103602023 | -2.368011094 | SH3PXD2A |
| 1 | 20618573 | 20618772 | -4.353679566 | CDA |
| 10 | 103603282 | 103603481 | -2.06658961 | SH3PXD2A |
| 10 | 103677052 | 103677242 | -1.59980279 | SH3PXD2A |
| 10 | 104268144 | 104268342 | -5.406890596 | MIR4482 |
| 10 | 110510472 | 110510679 | 2.459898695 | DUSP5 |
| 10 | 112817593 | 112817752 | -1.429967685 | VTI1A |
| 10 | 113730221 | 113730431 | 1.786125342 | CASP7 |
| 10 | 113849148 | 113849332 | -6.729011024 | DCLRE1A |
| 10 | 113852876 | 113853075 | -2.282888332 | DCLRE1A |
| 1 | 20650942 | 20651116 | -2.572662376 | PINK1 |
| 1 | 20650942 | 20651116 | -2.572662376 | PINK1-AS |
| 10 | 114125112 | 114125306 | -1.764503047 | CCDC186 |
| 10 | 114861541 | 114861712 | -1.300701279 | FAM160B1 |
| 10 | 116886370 | 116886582 | -1.440344558 | SHTN1 |
| 10 | 116886370 | 116886582 | -1.440344558 | ENO4 |
| 10 | 117283948 | 117284147 | -1.050488016 | PDZD8 |
| 10 | 118040347 | 118040545 | -1.916818027 | RAB11FIP2 |
| 17 | 58302067 | 58302265 | 2.70521447 | TSPOAP1 |
| 17 | 58980157 | 58980356 | -1.244226606 | PPM1E |
| 1 | 2350903 | 2351147 | 1.054214707 | RP4-740C4.5 |
| 1 | 2350903 | 2351147 | 1.054214707 | MORN1 |
| 1 | 89584314 | 89584533 | 2.01847685 | FLJ27354 |
| 1 | 89584314 | 89584533 | 2.01847685 | LRRC8B |
| 17 | 60483719 | 60483910 | -1.847733419 | APPBP2 |
| 17 | 61405379 | 61405538 | 10.43161825 | TBX2 |
| 17 | 61405379 | 61405538 | 10.43161825 | TBX2-AS1 |
| 17 | 61408464 | 61408604 | 3.851887828 | TBX2 |
| 17 | 61408464 | 61408604 | 3.851887828 | TBX2-AS1 |
| 17 | 61412508 | 61412700 | 6.718355771 | C17orf82 |
| 1 | 89592908 | 89593107 | 2.363965562 | FLJ27354 |
| 1 | 89592908 | 89593107 | 2.363965562 | LRRC8B |
| 1 | 89713350 | 89713547 | 1.707253776 | RP11-302M6.4 |
| 1 | 89713350 | 89713547 | 1.707253776 | LRRC8C |
| 17 | 63420240 | 63420442 | -1.194119422 | TANC2 |
| 17 | 63420240 | 63420442 | -1.194119422 | RP11-269G24.3 |
| 17 | 63819728 | 63819929 | 1.279176862 | FTSJ3 |
| 1 | 89713704 | 89713904 | 1.26828344 | RP11-302M6.4 |
| 1 | 89713704 | 89713904 | 1.26828344 | LRRC8C |
| 17 | 64002549 | 64002786 | 1.036847164 | ICAM2 |
| 17 | 64002549 | 64002786 | 1.036847164 | PRR29 |
| 17 | 64148743 | 64148941 | -1.018311545 | TEX2 |
| 17 | 64188278 | 64188463 | -2.194366422 | TEX2 |
| 17 | 64360313 | 64360512 | -1.882732872 | PECAM1 |
| 1 | 89714396 | 89714603 | 1.519348109 | RP11-302M6.4 |
| 1 | 89714396 | 89714603 | 1.519348109 | LRRC8C |
| 17 | 64375213 | 64375411 | -1.608802671 | PECAM1 |
| 17 | 64778323 | 64778521 | 9.992322571 | RP13-104F24.2 |
| 1 | 89758062 | 89758254 | 2.260811016 | LRRC8C |
| 1 | 89758062 | 89758254 | 2.260811016 | RP11-302M6.4 |
| 1 | 89762027 | 89762240 | 3.645847555 | RP11-302M6.4 |
| 1 | 89762027 | 89762240 | 3.645847555 | LRRC8C |
| 17 | 68135439 | 68135551 | -1.380406168 | RP11-147L13.13 |
| 17 | 68135439 | 68135551 | -1.380406168 | LRRC37A16P |
| 17 | 68135439 | 68135551 | -1.380406168 | SH3GL1P3 |
| 1 | 89820827 | 89820984 | 7.300682284 | RP5-943J3.2 |
| 1 | 89820827 | 89820984 | 7.300682284 | RP11-302M6.4 |
| 17 | 72694811 | 72694993 | -1.336273113 | SLC39A11 |
| 17 | 73068029 | 73068267 | -1.038929484 | SLC39A11 |
| 17 | 73068029 | 73068267 | -1.038929484 | RP11-766H1.1 |
| 1 | 89924036 | 89924206 | -3.559891188 | RP11-302M6.4 |
| 1 | 89924036 | 89924206 | -3.559891188 | LRRC8D |
| 17 | 74522414 | 74522611 | -1.482212907 | CD300LB |
| 17 | 74771409 | 74771613 | 1.446559602 | NAT9 |
| 17 | 75003774 | 75003976 | 1.708579778 | CDR2L |
| 17 | 75502046 | 75502199 | 2.39979072 | CASKIN2 |
| 17 | 75522273 | 75522467 | 2.072915782 | TSEN54 |
| 17 | 75588685 | 75588891 | 13.10843912 | MYO15B |
| 17 | 75890982 | 75891182 | 1.6486477 | TRIM65 |
| 17 | 75937947 | 75938145 | 1.44770349 | FBF1 |
| 17 | 76077800 | 76077957 | -1.554158124 | ZACN |
| 17 | 76270359 | 76270562 | 2.28118576 | UBALD2 |

| | | | | |
|---|---|---|---|---|
| X | 74068660 | 74068892 | 1.017110133 | JPX |
| X | 74072987 | 74073220 | 1.855763649 | FTX |
| X | 74077280 | 74077456 | 1.004668932 | FTX |
| X | 74095049 | 74095245 | 1.916546512 | FTX |
| 10 | 7964185 | 7964389 | -1.068737296 | TAF3 |
| X | 74095528 | 74095688 | 1.565952781 | FTX |
| X | 74096647 | 74096847 | 1.70470722 | FTX |
| X | 74097210 | 74097362 | 1.592595879 | FTX |
| X | 74193104 | 74193282 | 2.909753424 | FTX |
| X | 74194748 | 74194920 | 1.509744952 | FTX |
| 10 | 7964989 | 7965198 | -1.283983798 | TAF3 |
| X | 74744057 | 74744200 | -7.678054019 | KIAA2022 |
| X | 75741148 | 75741303 | -1.612475163 | TTC3P1 |
| X | 76429786 | 76429904 | -1.895038465 | MAGEE1 |
| 10 | 8014739 | 8014949 | -1.001721865 | TAF3 |
| X | 77684118 | 77684317 | -1.251268226 | ATRX |
| X | 77843748 | 77843932 | -5.422745025 | MAGT1 |
| X | 77846837 | 77847094 | -8.041261654 | MAGT1 |
| X | 77988347 | 77988548 | 1.680917489 | ATP7A |
| 10 | 11462636 | 11462841 | 1.30875717 | USP6NL |
| X | 81279988 | 81280158 | -6.046048207 | SH3BGRL |
| X | 91876691 | 91876860 | 8.669503083 | PCDH11X |
| X | 92013811 | 92014035 | 2.827914416 | PCDH11X |
| 10 | 11862217 | 11862378 | -3.771691698 | PROSER2 |
| 10 | 11862217 | 11862378 | -3.771691698 | PROSER2-AS1 |
| X | 101163223 | 101163412 | -3.055604068 | CENPI |
| X | 101348325 | 101348659 | -1.26060335 | TIMM8A |
| X | 101489029 | 101489229 | -1.270700233 | ARMCX4 |
| 10 | 11870466 | 11870681 | -1.441608123 | PROSER2-AS1 |
| 10 | 11870466 | 11870681 | -1.441608123 | PROSER2 |
| X | 101552995 | 101553210 | 1.047410491 | ARMCX1 |
| X | 101883045 | 101883244 | -1.508410481 | ZMAT1 |
| X | 102602067 | 102602284 | -1.561241064 | ARMCX5 |
| X | 102602067 | 102602284 | -1.561241064 | ARMCX5-GPRASP2 |
| X | 102602067 | 102602284 | -1.561241064 | RP4-769N13.7 |
| X | 102714645 | 102714836 | -1.534194184 | GPRASP2 |
| X | 102714645 | 102714836 | -1.534194184 | ARMCX5-GPRASP2 |
| X | 102748904 | 102749102 | -1.751369551 | BHLHB9 |
| X | 102884288 | 102884472 | -1.121764626 | LL0XNC01-237H1.2 |
| X | 102884288 | 102884472 | -1.121764626 | LINC00630 |
| X | 103088928 | 103089123 | -6.406890596 | NXF3 |
| X | 103088928 | 103089123 | -6.406890596 | LL0XNC01-221F2.2 |
| X | 103609272 | 103609475 | -1.776363266 | TCEAL3 |
| 10 | 12014113 | 12014296 | -2.426433106 | UPF2 |
| X | 103825526 | 103825722 | -2.907536523 | RAB9B |
| X | 104130725 | 104130927 | -1.076816223 | SLC25A53 |
| X | 106829770 | 106829948 | -3.382435795 | MORC4 |
| X | 106829770 | 106829948 | -3.382435795 | TBC1D8B |
| X | 107065960 | 107066152 | -1.10526289 | RBM41 |
| X | 107066436 | 107066631 | -1.331779299 | RBM41 |
| X | 107088522 | 107088707 | -1.070757379 | RBM41 |
| X | 107714123 | 107714323 | -1.153121743 | TSC22D3 |
| X | 109624043 | 109624241 | -3.422615415 | KCNE5 |
| X | 111755236 | 111755415 | 1.915644996 | ALG13 |
| 10 | 118040347 | 118040545 | -1.916818027 | RP11-354M20.3 |
| 10 | 118301944 | 118302062 | -1.714024141 | FAM204A |
| 10 | 119036073 | 119036278 | -1.116458232 | EIF3A |
| 10 | 119037213 | 119037413 | -4.611003816 | EIF3A |
| 10 | 119596879 | 119597079 | -1.11219392 | TIAL1 |
| 1 | 20907753 | 20907923 | -1.186945725 | EIF4G3 |
| 10 | 121951534 | 121951730 | -1.126736726 | ATE1-AS1 |
| 10 | 122211131 | 122211329 | -2.41888502 | TACC2 |
| 10 | 122888272 | 122888512 | -2.369472979 | RP11-564D11.3 |
| 10 | 122970877 | 122971076 | -1.378027484 | PSTK |
| 1 | 20941856 | 20942056 | -1.013295192 | EIF4G3 |
| 10 | 123056188 | 123056382 | -1.533044595 | ACADSB |
| 10 | 123137053 | 123137237 | 7.838088738 | HMX3 |
| 10 | 124942938 | 124943139 | -1.297137273 | ZRANB1 |
| 10 | 124966469 | 124966666 | -1.748001961 | ZRANB1 |
| 10 | 124984898 | 124985102 | -2.097182624 | ZRANB1 |
| 10 | 124984898 | 124985102 | -2.097182624 | CTBP2 |
| 10 | 124989414 | 124989615 | -1.54798908 | CTBP2 |
| 10 | 125842483 | 125842651 | 1.310139307 | DHX32 |
| 10 | 125842483 | 125842651 | 1.310139307 | BCCIP |
| 10 | 125880825 | 125881012 | -2.837973935 | DHX32 |
| 10 | 127934348 | 127934531 | -5.265203169 | PTPRE |
| 10 | 127934348 | 127934531 | -5.265203169 | RP11-4C20.3 |
| 10 | 128098805 | 128099027 | -1.415598205 | MKI67 |
| 10 | 128101454 | 128101622 | -3.487207295 | MKI67 |
| 10 | 128104388 | 128104587 | -1.861024752 | MKI67 |
| 10 | 128106207 | 128106402 | -1.519983637 | MKI67 |
| 10 | 128108097 | 128108296 | -2.400330135 | MKI67 |
| 10 | 128108811 | 128109011 | -8.032939447 | MKI67 |
| 10 | 128109153 | 128109268 | -4.231694816 | MKI67 |
| 10 | 128115235 | 128115341 | -2.587667799 | MKI67 |
| 10 | 128115389 | 128115564 | -2.282987617 | MKI67 |
| 10 | 131970407 | 131970577 | -8.564691824 | BNIP3 |
| 10 | 131971157 | 131971356 | -1.241275762 | BNIP3 |
| 10 | 131971157 | 131971356 | -1.241275762 | RP11-45A17.2 |
| 1 | 22030402 | 22030610 | -1.311422122 | LINC00339 |
| 10 | 131996776 | 131997043 | -1.520059812 | RP11-45A17.3 |
| 10 | 132192389 | 132192549 | -6.41188264 | DPYSL4 |
| 10 | 132348083 | 132348284 | -2.881606183 | LRRC27 |
| 10 | 132406608 | 132406806 | -2.02310993 | PWWP2B |
| 10 | 132448181 | 132448381 | -3.661685035 | C10orf91 |
| 10 | 133241257 | 133241457 | -1.805337466 | VENTX |
| 10 | 133420064 | 133420251 | 1.504950929 | RP11-108K14.8 |
| 10 | 133420064 | 133420251 | 1.504950929 | MTG1 |
| 10 | 133476833 | 133476989 | -4.303665157 | SCART1 |
| 10 | 133481033 | 133481211 | -6.873362122 | SCART1 |
| 11 | 216437 | 216636 | -2.518058765 | SIRT3 |
| 11 | 281367 | 281509 | -4.921175172 | NLRP6 |
| 11 | 324831 | 325159 | -1.794169178 | IFITM3 |
| 11 | 324831 | 325159 | -1.794169178 | RP11-326C3.11 |
| 11 | 422131 | 422332 | -1.182483966 | ANO9 |
| 11 | 451520 | 451709 | -1.860087459 | PTDSS2 |
| 11 | 452138 | 452368 | -1.798467571 | PTDSS2 |
| 11 | 453922 | 454111 | -5.372416919 | PTDSS2 |
| 17 | 76386863 | 76387059 | 8.121782256 | SPHK1 |
| 17 | 78111815 | 78111986 | 1.008921188 | TMC6 |
| 17 | 78112600 | 78112801 | 1.066167155 | TMC6 |
| 17 | 78171914 | 78172079 | 1.448767052 | SYNGR2 |
| 17 | 78345120 | 78345318 | -3.745926548 | AC061992.2 |
| 17 | 78358008 | 78358205 | 1.569393941 | SOCS3 |
| 17 | 78488793 | 78488919 | 2.242119956 | DNAH17-AS1 |
| 17 | 78488793 | 78488919 | 2.242119956 | DNAH17 |
| 1 | 90892947 | 90893063 | -1.074692884 | PHKA1P1 |
| 17 | 79086538 | 79086792 | 1.039874384 | ENGASE |
| 17 | 79933590 | 79933785 | 1.108016128 | TBC1D16 |
| 17 | 80045652 | 80045857 | -1.420050456 | CCDC40 |
| 17 | 80210046 | 80210240 | -1.451554447 | SGSH |
| 17 | 80252831 | 80253029 | -1.194467823 | SLC26A11 |
| 17 | 80288042 | 80288235 | 2.22701812 | RNF213 |
| 17 | 80295498 | 80295698 | -1.290045248 | RNF213 |
| 17 | 80727249 | 80727378 | -4.563352236 | RPTOR |
| 17 | 80964719 | 80964915 | 1.187874061 | RPTOR |
| 17 | 80970698 | 80970865 | -5.392317423 | CTD-2561B21.5 |
| 17 | 81114684 | 81114871 | -5.879319819 | BAIAP2 |
| 17 | 81304898 | 81305097 | -5.082209063 | LINC00482 |
| 17 | 81324859 | 81325046 | -3.56592848 | TMEM105 |
| 17 | 81327252 | 81327448 | -5.964629204 | TMEM105 |
| 17 | 81451565 | 81451756 | 7.222507423 | BAHCC1 |
| 17 | 81461375 | 81461583 | 8.348660276 | RP11-1055B8.8 |
| 17 | 81461375 | 81461583 | 8.348660276 | BAHCC1 |
| 17 | 81541157 | 81541325 | 2.899884614 | FAAP100 |
| 17 | 81547119 | 81547327 | 1.376581758 | FAAP100 |
| 17 | 81665307 | 81665501 | 1.427190901 | OXLD1 |
| 17 | 81666098 | 81666302 | 1.874698825 | OXLD1 |
| 17 | 81707223 | 81707420 | 9.957234904 | RP13-1032I1.10 |
| 17 | 81707223 | 81707420 | 9.957234904 | MRPL12 |
| 17 | 81710693 | 81710892 | -1.100665403 | RP13-1032I1.10 |
| 17 | 81843258 | 81843454 | 10.2108393 | RP11-498C9.2 |
| 17 | 81843258 | 81843454 | 10.2108393 | P4HB |
| 17 | 81947325 | 81947446 | -1.005503975 | AC137723.5 |
| 17 | 82138815 | 82138984 | -6.076776016 | CCDC57 |
| 17 | 82246665 | 82246864 | -1.321580113 | CSNK1D |
| 17 | 82246665 | 82246864 | -1.321580113 | SLC16A3 |
| 17 | 82249820 | 82249996 | -1.342961694 | SLC16A3 |
| 17 | 82249820 | 82249996 | -1.342961694 | RP13-516M14.8 |
| 17 | 82249820 | 82249996 | -1.342961694 | CSNK1D |
| 17 | 82253317 | 82253473 | -8.707156807 | SLC16A3 |
| 17 | 82253317 | 82253473 | -8.707156807 | CSNK1D |
| 17 | 82254271 | 82254466 | 5.589021164 | SLC16A3 |
| 17 | 82254271 | 82254466 | 5.589021164 | CSNK1D |
| 17 | 82273462 | 82273641 | 8.415665994 | CSNK1D |
| 17 | 82321835 | 82322007 | 2.516931726 | SECTM1 |
| 17 | 82936117 | 82936299 | -5.609886775 | TBCD |
| 17 | 82978160 | 82978327 | -1.982892142 | B3GNTL1 |
| 17 | 83094358 | 83094558 | 1.607747607 | METRNL |
| 17 | 83104297 | 83104486 | 2.290675624 | AC144831.1 |
| 17 | 83105200 | 83105430 | 1.521991681 | AC144831.1 |
| 18 | 659603 | 659801 | -6.682887818 | TYMS |

| | | | | |
|---|---|---|---|---|
| X | 113692854 | 113693039 | -1.156092833 | XACT |
| X | 113700055 | 113700249 | 1.415404567 | XACT |
| X | 113729978 | 113730203 | -1.758802175 | XACT |
| X | 113820181 | 113820330 | -1.069756075 | XACT |
| X | 113823081 | 113823237 | -1.621408925 | XACT |
| X | 113823408 | 113823517 | -2.880395947 | XACT |
| X | 113825461 | 113825646 | -1.51330486 | XACT |
| X | 113857907 | 113858153 | -1.698978175 | XACT |
| X | 113858929 | 113859130 | -2.3194385 | XACT |
| X | 113861627 | 113861812 | -1.411501633 | XACT |
| X | 113863003 | 113863202 | -2.027077355 | XACT |
| X | 113863660 | 113863792 | -1.912052159 | XACT |
| X | 113925342 | 113925541 | -1.156953368 | XACT |
| X | 117909725 | 117909941 | -3.156299368 | KLHL13 |
| X | 118768538 | 118768724 | -6.93112495 | IL13RA1 |
| X | 118992269 | 118992469 | 1.339848869 | LONRF3 |
| X | 118998211 | 118998376 | 2.225313737 | LONRF3 |
| X | 119589521 | 119589720 | -2.162518036 | UBE2A |
| X | 119589521 | 119589720 | -2.162518036 | NKRF |
| 10 | 13209168 | 13209367 | -4.71774994 | MCM10 |
| X | 119591056 | 119591256 | -1.500877286 | NKRF |
| X | 119591056 | 119591256 | -1.500877286 | UBE2A |
| X | 119759515 | 119759794 | -2.006193513 | SOWAHD |
| X | 119871377 | 119871576 | -1.026012334 | RNF113A |
| X | 119871377 | 119871576 | -1.026012334 | NDUFA1 |
| X | 120255195 | 120255380 | -1.468867265 | ZBTB33 |
| X | 121049147 | 121049348 | 1.605820506 | GLUD2 |
| 10 | 13317517 | 13317728 | -2.760584922 | SEPHS1 |
| X | 126165162 | 126165342 | -2.297364164 | DCAF12L2 |
| X | 126551839 | 126552069 | -9.549591705 | DCAF12L1 |
| X | 129793913 | 129794112 | 1.327688862 | SASH3 |
| 10 | 13646359 | 13646569 | 12.59924598 | RP11-295P9.13 |
| 10 | 13646359 | 13646569 | 12.59924598 | FRMD4A |
| 10 | 13646359 | 13646569 | 12.59924598 | RP11-295P9.3 |
| X | 130412474 | 130412683 | -1.039648771 | RBMX2 |
| X | 132449215 | 132449374 | -6.07487356 | MBNL3 |
| X | 133753656 | 133753848 | 2.98282617 | GPC3 |
| 10 | 13657294 | 13657502 | 8.097878335 | FRMD4A |
| 10 | 13657294 | 13657502 | 8.097878335 | RP11-295P9.3 |
| X | 134991597 | 134991796 | -2.488702042 | SMIM10 |
| X | 135021595 | 135021828 | -1.086133985 | FAM127C |
| X | 135343350 | 135343547 | -2.403906047 | ZNF75D |
| 10 | 13707253 | 13707452 | 8.682904901 | FRMD4A |
| X | 136231796 | 136232011 | -1.897956991 | MAP7D3 |
| X | 136922481 | 136922707 | -1.742713414 | RP11-308D16.2 |
| X | 149483070 | 149483271 | 3.193396061 | IDS |
| X | 149483070 | 149483271 | 3.193396061 | AF011889.5 |
| X | 149525871 | 149526094 | -1.323165497 | AF011889.5 |
| X | 149525871 | 149526094 | -1.323165497 | AF011889.2 |
| X | 149525871 | 149526094 | -1.323165497 | IDSP1 |
| 10 | 14520920 | 14521192 | -1.485182214 | FAM107B |
| X | 149546747 | 149546847 | -3.860321644 | CXorf40A |
| X | 152869052 | 152869248 | -1.389086485 | NSDHL |
| 10 | 14567803 | 14567998 | -2.387837255 | FAM107B |
| 11 | 459272 | 459497 | -5.926654778 | PTDSS2 |
| 11 | 529039 | 529242 | 1.46289906 | RP13-46H24.1 |
| 1 | 23084178 | 23084388 | -1.889528983 | RP1-184J9.2 |
| 1 | 23084178 | 23084388 | -1.889528983 | LUZP1 |
| 11 | 532939 | 533150 | 1.17821474 | HRAS |
| 11 | 536254 | 536396 | 1.20583169 | HRAS |
| 11 | 538578 | 538702 | 1.352238871 | LRRC56 |
| 11 | 545321 | 545507 | 1.323034294 | LRRC56 |
| 11 | 546576 | 546760 | 3.270404849 | LRRC56 |
| 11 | 563720 | 563919 | -1.781679447 | RASSF7 |
| 11 | 566293 | 566488 | 5.593366645 | MIR210HG |
| 11 | 567272 | 567472 | 3.427473492 | MIR210HG |
| 11 | 644401 | 644602 | -1.042683257 | DEAF1 |
| 11 | 657648 | 657846 | 5.910089481 | DEAF1 |
| 1 | 23091279 | 23091482 | -1.343347789 | LUZP1 |
| 11 | 783816 | 783995 | -1.794067998 | AP006621.5 |
| 11 | 797256 | 797450 | -1.432406901 | SLC25A22 |
| 1 | 23092692 | 23092886 | -2.175401249 | LUZP1 |
| 11 | 1286696 | 1286925 | -2.161798986 | TOLLIP |
| 11 | 1453421 | 1453607 | -1.666711867 | BRSK2 |
| 11 | 1584513 | 1584677 | 1.545140303 | AP006285.2 |
| 11 | 1584513 | 1584677 | 1.545140303 | KRTAP5-AS1 |
| 11 | 1584513 | 1584677 | 1.545140303 | KRTAP5-1 |
| 11 | 2268748 | 2268939 | 2.119219817 | ASCL2 |
| 11 | 2328882 | 2329076 | -1.265778187 | CD81-AS1 |
| 11 | 2698468 | 2698632 | -5.237866715 | KCNQ1OT1 |
| 11 | 2698468 | 2698632 | -5.237866715 | KCNQ1 |
| 11 | 3132609 | 3132805 | 7.158988911 | OSBPL5 |
| 11 | 3133668 | 3133830 | 8.521780679 | OSBPL5 |
| 11 | 3359998 | 3360222 | -1.54873497 | ZNF195 |
| 1 | 23362354 | 23362511 | -7.692122061 | ZNF436 |
| 11 | 3712419 | 3712590 | 7.445155892 | NUP98 |
| 11 | 4052677 | 4052870 | -1.433139767 | STIM1 |
| 11 | 4385528 | 4385727 | 1.094347639 | TRIM21 |
| 1 | 1090210 | 1090350 | -1.098622372 | C1orf159 |
| 1 | 23509750 | 23509944 | -5.988639962 | E2F2 |
| 11 | 5714902 | 5715103 | 1.276954341 | TRIM5 |
| 11 | 5714902 | 5715103 | 1.276954341 | TRIM22 |
| 11 | 5715752 | 5715934 | 2.266878967 | TRIM22 |
| 11 | 5715752 | 5715934 | 2.266878967 | TRIM5 |
| 11 | 6390458 | 6390661 | 7.597301488 | SMPD1 |
| 11 | 6391713 | 6391907 | 1.103009512 | SMPD1 |
| 11 | 6557275 | 6557475 | -1.058325943 | DNHD1 |
| 11 | 6558220 | 6558469 | -1.35207679 | DNHD1 |
| 11 | 8668203 | 8668369 | -3.209863073 | TRIM66 |
| 11 | 8668713 | 8668902 | -1.598545491 | TRIM66 |
| 11 | 8671771 | 8671970 | -1.265633919 | TRIM66 |
| 1 | 23752257 | 23752456 | 1.402246647 | TCEB3 |
| 11 | 9139538 | 9139749 | 1.848984729 | DENND5A |
| 11 | 9203982 | 9204181 | 1.241896035 | DENND5A |
| 11 | 9280746 | 9280944 | -1.081917953 | TMEM41B |
| 11 | 9749816 | 9750017 | -1.240931334 | SWAP70 |
| 11 | 9779479 | 9779664 | -6.178501046 | SBF2 |
| 11 | 9779479 | 9779664 | -6.178501046 | SBF2-AS1 |
| 1 | 2384108 | 2384303 | 2.220804319 | MORN1 |
| 18 | 664661 | 664770 | -5.477098155 | TYMS |
| 18 | 667685 | 667852 | -7.157539392 | TYMS |
| 18 | 670603 | 670810 | -1.797753959 | TYMS |
| 18 | 670603 | 670810 | -1.797753959 | ENOSF1 |
| 18 | 2920855 | 2921033 | 4.789657969 | RP11-737O24.5 |
| 18 | 2920855 | 2921033 | 4.789657969 | LPIN2 |
| 18 | 3596614 | 3596812 | -2.2036621 | DLGAP1-AS1 |
| 18 | 3596614 | 3596812 | -2.2036621 | DLGAP1 |
| 18 | 8762411 | 8762612 | 1.851781927 | MTCL1 |
| 18 | 8783665 | 8783795 | 1.024806552 | MTCL1 |
| 18 | 8784173 | 8784369 | 8.461239938 | MTCL1 |
| 18 | 8824795 | 8825000 | 1.656760345 | MTCL1 |
| 18 | 8825434 | 8825743 | 2.453759542 | MTCL1 |
| 18 | 9136328 | 9136548 | 1.117461128 | ANKRD12 |
| 18 | 9136328 | 9136548 | 1.117461128 | RP11-21J18.1 |
| 18 | 9136328 | 9136548 | 1.117461128 | NDUFV2-AS1 |
| 18 | 9524470 | 9524666 | -6.39395128 | RALBP1 |
| 18 | 9887888 | 9888075 | -2.143497152 | TXNDC2 |
| 18 | 10471776 | 10471969 | 2.205009712 | APCDD1 |
| 18 | 10487999 | 10488198 | 5.889207074 | APCDD1 |
| 18 | 11859459 | 11859657 | 7.358409731 | GNAL |
| 18 | 12325037 | 12325236 | 1.989224305 | TUBB6 |
| 18 | 12326146 | 12326346 | 1.028032833 | TUBB6 |
| 18 | 12449233 | 12449561 | -1.330569637 | SPIRE1 |
| 18 | 13495621 | 13495794 | -2.701049222 | LDLRAD4 |
| 1 | 93166745 | 93166989 | -8.627864691 | TMED5 |
| 18 | 14114111 | 14114300 | -5.419995535 | ZNF519 |
| 18 | 21111145 | 21111334 | 9.16938995 | ROCK1 |
| 1 | 93180374 | 93180575 | 1.328840539 | TMED5 |
| 1 | 93180374 | 93180575 | 1.328840539 | CCDC18 |
| 18 | 23586676 | 23586857 | 1.206950536 | NPC1 |
| 18 | 23648885 | 23649030 | 1.488058933 | ANKRD29 |
| 18 | 25226973 | 25227172 | -1.335878651 | ZNF521 |
| 18 | 34714142 | 34714320 | -6.514643613 | DTNA |
| 18 | 34725045 | 34725216 | -6.842319528 | DTNA |
| 18 | 35374124 | 35374324 | -1.609024889 | ZNF396 |
| 18 | 35633241 | 35633424 | 1.787454418 | GALNT1 |
| 18 | 36786586 | 36786769 | -1.120516491 | TPGS2 |
| 18 | 36962707 | 36962860 | -1.861233012 | KIAA1328 |
| 18 | 44678467 | 44678774 | 2.205308402 | RP11-456K23.1 |
| 18 | 44701391 | 44701594 | 2.034046336 | SETBP1 |
| 18 | 44953013 | 44953224 | 2.069844496 | SETBP1 |
| 18 | 45038354 | 45038540 | 2.521381978 | SETBP1 |
| 18 | 45063651 | 45063793 | 3.801995546 | SETBP1 |
| 1 | 93266280 | 93266380 | 1.59216219 | CCDC18 |
| 1 | 93266280 | 93266380 | 1.59216219 | CCDC18-AS1 |
| 18 | 46253593 | 46253792 | -7.514665185 | C18orf25 |
| 18 | 46456602 | 46456738 | 6.868306665 | RNF165 |
| 18 | 46679412 | 46679620 | -1.169990938 | ST8SIA5 |
| 18 | 46812226 | 46812573 | -2.296118582 | PIAS2 |
| 18 | 46946057 | 46946211 | 1.233489214 | KATNAL2 |
| 18 | 47841713 | 47841908 | -1.482554717 | SMAD2 |
| 18 | 48949836 | 48950013 | 7.528445844 | SMAD7 |

| | | | | |
|---|---|---|---|---|
| X | 153459257 | 153459435 | -1.680031612 | HAUS7 |
| X | 153459257 | 153459435 | -1.680031612 | TREX2 |
| X | 153571875 | 153572043 | 2.548491599 | ATP2B3 |
| X | 153580309 | 153580515 | 2.003863681 | ATP2B3 |
| X | 153724911 | 153725145 | -1.472506204 | ABCD1 |
| X | 153905400 | 153905604 | -1.233940461 | L1CAM |
| X | 153905400 | 153905604 | -1.233940461 | AVPR2 |
| X | 153954478 | 153954684 | -1.753181744 | HCFC1 |
| X | 153955128 | 153955304 | -2.076802977 | HCFC1 |
| X | 153971091 | 153971209 | -11.19048811 | HCFC1 |
| X | 153971582 | 153971784 | -1.186213847 | HCFC1 |
| X | 153982872 | 153983099 | -1.041983867 | TMEM187 |
| 10 | 14842567 | 14842736 | -8.869732097 | HSPA14 |
| X | 154394300 | 154394563 | -1.555432374 | RPL10 |
| X | 154398569 | 154398766 | -7.779344879 | RPL10 |
| X | 154398569 | 154398766 | -7.779344879 | XX-FW83563B9.5 |
| X | 154460319 | 154460614 | 1.92145125 | PLXNA3 |
| X | 154484364 | 154484536 | -1.115221509 | UBL4A |
| 10 | 14843554 | 14843773 | -2.206996525 | HSPA14 |
| X | 155122128 | 155122301 | -2.233826606 | MTCP1 |
| X | 155122128 | 155122301 | -2.233826606 | BRCC3 |
| X | 155125879 | 155126078 | -1.49228661 | MTCP1 |
| X | 155773768 | 155774058 | -1.495678249 | SPRY3 |
| X | 156009947 | 156010145 | -2.454144762 | AJ271736.10 |
| X | 156009947 | 156010145 | -2.454144762 | IL9R |
| 10 | 14896925 | 14897116 | -2.425279026 | SUV39H2 |
| Y | 2924675 | 2924858 | 4.178776002 | RPS4Y1 |
| Y | 3069859 | 3070050 | 1.866178387 | LINC00278 |
| 10 | 14907848 | 14908043 | -2.144698685 | DCLRE1C |
| Y | 5099982 | 5100166 | 6.669925001 | PCDH11Y |
| 10 | 14908658 | 14908860 | -1.554247022 | DCLRE1C |
| Y | 14622121 | 14622300 | -1.626817854 | NLGN4Y |
| Y | 14622344 | 14622489 | -7.511357287 | NLGN4Y |
| Y | 14774778 | 14774926 | -3.863960227 | NLGN4Y |
| Y | 14830156 | 14830286 | -1.150965593 | NLGN4Y |
| 10 | 14909143 | 14909329 | -1.272717068 | DCLRE1C |
| Y | 19603739 | 19603970 | -1.171841324 | TXLNGY |
| 10 | 14910748 | 14910982 | -2.049430727 | DCLRE1C |
| 10 | 14911656 | 14911877 | -1.951886534 | DCLRE1C |
| 10 | 16591732 | 16591922 | -2.233268314 | RSU1 |
| 10 | 17142405 | 17142647 | -1.114479786 | TRDMT1 |
| 10 | 18513397 | 18513597 | -1.157180862 | CACNB2 |
| 10 | 18513397 | 18513597 | -1.157180862 | RP11-499P20.2 |
| 10 | 20050099 | 20050297 | 6.730739724 | PLXDC2 |
| 10 | 20263293 | 20263411 | 4.447573007 | PLXDC2 |
| 10 | 20279723 | 20279938 | 4.611283056 | PLXDC2 |
| 10 | 21494499 | 21494692 | -8.546811942 | CASC10 |
| 10 | 21515400 | 21515599 | -3.306505263 | SKIDA1 |
| 10 | 21796690 | 21796912 | -6.577250847 | DNAJC1 |
| 10 | 22252107 | 22252248 | 2.791459753 | RP11-573G6.4 |
| 10 | 22329574 | 22329790 | -1.836906846 | BMI1 |
| 10 | 23121089 | 23121288 | -1.973678558 | MSRB2 |
| 10 | 23121541 | 23121757 | -2.097336536 | MSRB2 |
| 10 | 23440542 | 23440744 | 1.99642857 | OTUD1 |
| 11 | 9807999 | 9808230 | -1.3080556 | SBF2-AS1 |
| 11 | 9807999 | 9808230 | -1.3080556 | SBF2 |
| 11 | 9809013 | 9809176 | -6.5691359 | SBF2-AS1 |
| 11 | 9809013 | 9809176 | -6.5691359 | SBF2 |
| 11 | 10170152 | 10170269 | 5.245926548 | SBF2 |
| 11 | 10471243 | 10471443 | 5.388627658 | AMPD3 |
| 11 | 10505643 | 10505818 | 1.436212277 | AMPD3 |
| 11 | 10853083 | 10853284 | -1.240148614 | ZBED5 |
| 1 | 23845560 | 23845758 | -1.444450551 | FUCA1 |
| 11 | 11956170 | 11956369 | -1.163951364 | USP47 |
| 11 | 12259875 | 12260066 | -1.6212478 | MICAL2 |
| 11 | 12358561 | 12358674 | -4.362256927 | MICALCL |
| 11 | 14265328 | 14265448 | -1.187683677 | RP11-21L19.1 |
| 11 | 14265328 | 14265448 | -1.187683677 | SPON1 |
| 11 | 14576287 | 14576497 | -3.166721041 | PSMA1 |
| 11 | 14595745 | 14595971 | -2.755596878 | PSMA1 |
| 11 | 14597787 | 14597988 | -2.127816901 | PSMA1 |
| 11 | 14600252 | 14600481 | -2.722926536 | PSMA1 |
| 11 | 14606153 | 14606337 | -4.775373393 | PSMA1 |
| 11 | 14663462 | 14663696 | -7.488684687 | PDE3B |
| 11 | 14666321 | 14666520 | -1.901096608 | PDE3B |
| 11 | 14666959 | 14667165 | -1.784729802 | PDE3B |
| 11 | 14717964 | 14718158 | -7.515074912 | PDE3B |
| 11 | 14721453 | 14721646 | -2.138535969 | PDE3B |
| 11 | 14777739 | 14777939 | -2.366857774 | PDE3B |
| 11 | 14779861 | 14780010 | -2.957313122 | PDE3B |
| 11 | 14783322 | 14783511 | -2.534946906 | PDE3B |
| 11 | 14846531 | 14846721 | -1.306061639 | PDE3B |
| 11 | 14863998 | 14864195 | -1.981892838 | PDE3B |
| 11 | 14869742 | 14869942 | -1.935577241 | PDE3B |
| 11 | 14883501 | 14883705 | -4.759599702 | CYP2R1 |
| 11 | 14884208 | 14884402 | -6.524754126 | CYP2R1 |
| 11 | 14891228 | 14891539 | -1.938600436 | CYP2R1 |
| 11 | 16826201 | 16826337 | 2.851803499 | PLEKHA7 |
| 1 | 24901636 | 24901835 | 1.119094368 | RUNX3 |
| 11 | 17169563 | 17169745 | -1.18485191 | PIK3C2A |
| 11 | 17353070 | 17353264 | -4.225605556 | NCR3LG1 |
| 11 | 17353678 | 17353884 | 1.39085755 | NCR3LG1 |
| 11 | 18702035 | 18702225 | -3.048714315 | TMEM86A |
| 1 | 24902097 | 24902296 | 9.38309621 | RUNX3 |
| 11 | 22624793 | 22625018 | -1.125853994 | FANCF |
| 11 | 27367948 | 27368218 | -1.686123561 | LGR4 |
| 11 | 27658359 | 27658531 | -2.547864002 | BDNF-AS |
| 11 | 27658359 | 27658531 | -2.547864002 | BDNF |
| 11 | 28036300 | 28036513 | -2.946448662 | KIF18A |
| 1 | 24931736 | 24931928 | 1.919655844 | RUNX3 |
| 11 | 28159004 | 28159204 | -2.096423166 | METTL15 |
| 11 | 30331076 | 30331276 | -1.310421446 | ARL14EP |
| 11 | 31495066 | 31495261 | -1.950002105 | IMMP1L |
| 11 | 31746563 | 31746757 | 1.660964047 | Z83001.1 |
| 11 | 31746563 | 31746757 | 1.660964047 | ELP4 |
| 11 | 32438117 | 32438320 | 1.124577905 | WT1-AS |
| 11 | 32438904 | 32439104 | 1.331482506 | WT1-AS_6 |
| 11 | 32438904 | 32439104 | 1.331482506 | WT1-AS |
| 18 | 48950320 | 48950527 | 1.100389357 | SMAD7 |
| 18 | 49294060 | 49294263 | -6.794298764 | DYM |
| 18 | 49590642 | 49590839 | -1.666000211 | LIPG |
| 18 | 51047007 | 51047137 | -1.495637984 | RP11-729L2.2 |
| 18 | 51047007 | 51047137 | -1.495637984 | SMAD4 |
| 18 | 57436743 | 57436934 | 1.727327239 | ONECUT2 |
| 18 | 57439335 | 57439533 | 6.902821561 | ONECUT2 |
| 1 | 93514828 | 93514981 | 1.045573944 | FNBP1L |
| 18 | 57440198 | 57440379 | 6.535982536 | ONECUT2 |
| 18 | 57483020 | 57483211 | -6.177124691 | ONECUT2 |
| 1 | 93550909 | 93551109 | -1.145831186 | FNBP1L |
| 18 | 58920336 | 58920545 | 1.132489706 | ZNF532 |
| 18 | 59051299 | 59051479 | 2.196158711 | OACYLP |
| 18 | 62523756 | 62523948 | 9.989517184 | ZCCHC2 |
| 18 | 62575274 | 62575443 | -1.0495752 | ZCCHC2 |
| 18 | 62576408 | 62576605 | -7.193739433 | ZCCHC2 |
| 18 | 62829903 | 62830122 | 7.258219367 | PHLPP1 |
| 18 | 63123875 | 63124160 | -1.490197721 | BCL2 |
| 18 | 63127417 | 63127617 | -1.364436331 | BCL2 |
| 18 | 63317599 | 63317773 | -2.176195975 | BCL2 |
| 18 | 63318659 | 63318788 | -1.492186189 | BCL2 |
| 18 | 63319424 | 63319613 | -1.108714089 | BCL2 |
| 18 | 70098931 | 70099131 | -4.65983606 | RTTN |
| 18 | 70099492 | 70099701 | -5.344125155 | RTTN |
| 18 | 74593571 | 74593772 | 1.054223528 | LINC00909 |
| 1 | 95063445 | 95063645 | -5.108751689 | ALG14 |
| 1 | 95063445 | 95063645 | -5.108751689 | RP11-313A24.1 |
| 18 | 75063577 | 75063720 | 1.070624504 | ZNF407 |
| 18 | 75064236 | 75064434 | 2.202327327 | ZNF407 |
| 18 | 75287359 | 75287529 | -1.279951902 | TSHZ1 |
| 18 | 76371537 | 76371724 | -2.183975111 | ZNF516 |
| 18 | 76441722 | 76441919 | 10.45963972 | ZNF516 |
| 18 | 77037453 | 77037649 | -2.558229195 | MBP |
| 1 | 99306980 | 99307185 | 2.44447904 | PLPPR4 |
| 18 | 79714939 | 79715208 | 1.361759549 | CTDP1 |
| 1 | 99969510 | 99969695 | -1.25265618 | RP5-884G6.2 |
| 18 | 80162058 | 80162283 | -1.474891538 | PARD6G |
| 18 | 80162058 | 80162283 | -1.474891538 | PARD6G-AS1 |
| 18 | 80162058 | 80162283 | -1.474891538 | AC139100.4 |
| 19 | 306562 | 306767 | 1.078200491 | MIER2 |
| 19 | 509101 | 509298 | 1.446660449 | TPGS1 |
| 19 | 643040 | 643194 | 3.821928095 | FGF22 |
| 19 | 1003220 | 1003412 | 7.852937668 | GRIN3B |
| 19 | 1005039 | 1005238 | 2.820949715 | GRIN3B |
| 19 | 1010503 | 1010672 | 10.81450218 | LLNLR-284B4.1 |
| 19 | 1010503 | 1010672 | 10.81450218 | TMEM259 |
| 19 | 1085561 | 1085742 | 1.095822218 | ARHGAP45 |
| 19 | 1157894 | 1158191 | 1.32633382 | SBNO2 |
| 19 | 1269936 | 1270145 | 9.303925211 | CIRBP-AS1 |
| 19 | 1269936 | 1270145 | 9.303925211 | CIRBP |
| 19 | 1278962 | 1279157 | 1.156108799 | C19orf24 |
| 19 | 1321925 | 1322088 | -1.884994035 | AC005330.2 |
| 19 | 1377160 | 1377323 | 2.813588661 | AC004623.2 |
| 19 | 1377160 | 1377323 | 2.813588661 | MUM1 |

| | | | | |
|---|---|---|---|---|
| 1 | 15215003 | 15215200 | 1.752874485 | TMEM51 |
| 10 | 24063096 | 24063286 | 2.043731421 | KIAA1217 |
| 10 | 25598288 | 25598484 | 1.339088131 | GPR158 |
| 10 | 27035148 | 27035345 | -1.836270324 | ANKRD26 |
| 1 | 15219533 | 15219733 | 2.026724169 | TMEM51 |
| 10 | 27035456 | 27035567 | -1.554417533 | ANKRD26 |
| 10 | 27170141 | 27170336 | -2.64356782 | MASTL |
| 10 | 28523484 | 28523687 | -1.047567864 | WAC-AS1 |
| 10 | 29467416 | 29467617 | -2.247313962 | SVIL |
| 10 | 29467416 | 29467617 | -2.247313962 | SVIL-AS1 |
| 10 | 29523733 | 29523846 | -1.494632427 | SVIL |
| 10 | 29550893 | 29551009 | -1.87454094 | SVIL |
| 10 | 30313385 | 30313750 | -1.160071182 | MTPAP |
| 10 | 30844934 | 30845140 | -1.148639698 | ZNF438 |
| 10 | 30849196 | 30849412 | -1.383970134 | ZNF438 |
| 10 | 30928688 | 30928869 | -4.805005078 | ZNF438 |
| 10 | 31318404 | 31318609 | -1.40525068 | ZEB1-AS1 |
| 10 | 31318404 | 31318609 | -1.40525068 | ZEB1 |
| 10 | 31474673 | 31474827 | -8.338612063 | ZEB1 |
| 10 | 31475938 | 31476042 | -7.633996824 | ZEB1 |
| 10 | 31861570 | 31861769 | 1.348085887 | ARHGAP12 |
| 1 | 1065725 | 1065931 | 5.582572578 | RP11-465B22.3 |
| 10 | 31891650 | 31891821 | 1.052977682 | ARHGAP12 |
| 10 | 33179990 | 33180203 | 6.242561651 | NRP1 |
| 10 | 34111110 | 34111283 | -2.558868598 | PARD3 |
| 10 | 37829136 | 37829357 | -2.333699954 | ZNF248 |
| 10 | 37829136 | 37829357 | -2.333699954 | AL135791.1 |
| 10 | 37832392 | 37832577 | -1.216756502 | ZNF248 |
| 10 | 37832392 | 37832577 | -1.216756502 | AL135791.1 |
| 10 | 38054591 | 38054813 | -1.513360122 | ZNF33A |
| 10 | 38064031 | 38064292 | -1.972816233 | ZNF33A |
| 10 | 38111051 | 38111240 | 6.312169273 | ZNF37A |
| 10 | 38124090 | 38124284 | 1.61177724 | ZNF37A |
| 10 | 42335543 | 42335741 | -4.088162963 | RP11-313J2.1 |
| 10 | 42337214 | 42337340 | -3.026071364 | RP11-313J2.1 |
| 10 | 42520016 | 42520201 | -1.34937874 | ZNF37BP |
| 10 | 42534747 | 42534960 | -1.683003073 | ZNF37BP |
| 10 | 42594273 | 42594459 | -2.504228777 | ZNF33B |
| 10 | 42796520 | 42796720 | -3.139049728 | BMS1 |
| 10 | 42831029 | 42831220 | -1.581948977 | BMS1 |
| 10 | 43155048 | 43155345 | 1.239820218 | CSGALNACT2 |
| 10 | 43183402 | 43183605 | 2.316886888 | CSGALNACT2 |
| 10 | 43557518 | 43557746 | -2.865106941 | ZNF239 |
| 10 | 43616717 | 43616911 | -1.488081184 | ZNF485 |
| 10 | 43617116 | 43617234 | -1.687422301 | ZNF485 |
| 10 | 44977471 | 44977710 | -2.761355386 | RASSF4 |
| 10 | 44977471 | 44977710 | -2.761355386 | C10orf10 |
| 10 | 44994180 | 44994363 | -1.7156806 | RASSF4 |
| 10 | 45003332 | 45003530 | -2.237839966 | CEP164P1 |
| 10 | 45003332 | 45003530 | -2.237839966 | ZNF22 |
| 10 | 45004787 | 45004972 | -2.890679857 | ZNF22 |
| 10 | 45004787 | 45004972 | -2.890679857 | CEP164P1 |
| 10 | 45429231 | 45429397 | -4.241403978 | ALOX5 |
| 10 | 45626531 | 45626720 | -1.630155619 | ZFAND4 |

| | | | | |
|---|---|---|---|---|
| 11 | 32853739 | 32853940 | 1.469494598 | PRRG4 |
| 1 | 25458582 | 25458785 | -1.251911635 | TMEM57 |
| 11 | 33353148 | 33353334 | -1.151753761 | HIPK3 |
| 11 | 33710148 | 33710346 | 1.599069948 | CD59 |
| 11 | 33859107 | 33859315 | -1.136066468 | LMO2 |
| 11 | 34159358 | 34159534 | -6.488103045 | ABTB2 |
| 11 | 35137798 | 35137952 | 9.560781935 | RP4-607I7.1 |
| 11 | 35200915 | 35201106 | 11.72076031 | CD44 |
| 11 | 35204584 | 35204784 | 11.04153333 | CD44 |
| 11 | 35229128 | 35229326 | 2.835199128 | CD44 |
| 11 | 43325521 | 43325707 | -6.324178791 | API5 |
| 11 | 43339437 | 43339633 | -1.277004786 | RP11-484D2.2 |
| 11 | 43339437 | 43339633 | -1.277004786 | API5 |
| 1 | 25566900 | 25567101 | -1.633334569 | LDLRAP1 |
| 11 | 45933913 | 45934135 | -2.233894548 | PHF21A |
| 11 | 46238413 | 46238575 | -1.688330414 | CTD-2589M5.5 |
| 11 | 46616086 | 46616262 | -1.130473177 | HARBI1 |
| 11 | 46743975 | 46744179 | -1.280536324 | CKAP5 |
| 11 | 47238155 | 47238353 | -2.125740634 | DDB2 |
| 11 | 47261563 | 47261762 | -1.363073185 | NR1H3 |
| 11 | 47577921 | 47578043 | -1.039689056 | NDUFS3 |
| 11 | 47577921 | 47578043 | -1.039689056 | KBTBD4 |
| 11 | 47577921 | 47578043 | -1.039689056 | RP11-76I23.7 |
| 11 | 57302758 | 57302932 | -6.703235384 | TNKS1BP1 |
| 11 | 57308985 | 57309274 | -1.664226464 | TNKS1BP1 |
| 11 | 57313568 | 57313798 | -1.376354269 | TNKS1BP1 |
| 11 | 57484943 | 57485154 | -1.814315349 | SLC43A1 |
| 11 | 57603562 | 57603674 | 6.478006539 | SERPING1 |
| 1 | 25783782 | 25783980 | -1.823518385 | MAN1C1 |
| 11 | 59152719 | 59152918 | 1.057416155 | FAM111A |
| 11 | 59182149 | 59182336 | 2.840548186 | DTX4 |
| 11 | 59204788 | 59204991 | 2.006825993 | DTX4 |
| 11 | 60365189 | 60365389 | 5.333112001 | MS4A6E |
| 11 | 60937011 | 60937146 | 5.123370299 | SLC15A3 |
| 11 | 60937011 | 60937146 | 5.123370299 | TMEM132A |
| 11 | 61004604 | 61004884 | -1.510745649 | CD6 |
| 11 | 61258488 | 61258686 | -4.219939411 | VWCE |
| 1 | 1093104 | 1093319 | 2.224619415 | C1orf159 |
| 11 | 61636213 | 61636340 | 1.433698539 | RPLP0P2 |
| 11 | 61636673 | 61637030 | 1.89612105 | RPLP0P2 |
| 11 | 61637785 | 61637984 | 1.925653712 | RPLP0P2 |
| 11 | 61786454 | 61786761 | -1.859826043 | MYRF |
| 11 | 61786454 | 61786761 | -1.859826043 | TMEM258 |
| 11 | 61898107 | 61898307 | -1.816193291 | RAB3IL1 |
| 11 | 61962573 | 61962790 | -1.527189939 | BEST1 |
| 11 | 61962573 | 61962790 | -1.527189939 | FTH1 |
| 1 | 25835197 | 25835404 | -4.686024413 | AUNIP |
| 11 | 62517473 | 62517664 | -1.668571731 | AHNAK |
| 11 | 62518397 | 62518599 | -1.422963301 | AHNAK |
| 11 | 62519325 | 62519516 | -1.056142083 | AHNAK |
| 11 | 62520607 | 62520808 | -1.208288225 | AHNAK |
| 11 | 62521963 | 62522232 | -1.616606863 | AHNAK |
| 11 | 62524548 | 62524745 | -1.246814198 | AHNAK |
| 11 | 62529547 | 62529754 | -1.358802177 | AHNAK |

| | | | | |
|---|---|---|---|---|
| 19 | 1392551 | 1392739 | -6.774471144 | NDUFS7 |
| 19 | 1392551 | 1392739 | -6.774471144 | AC005329.7 |
| 19 | 1474793 | 1475036 | 2.360031422 | C19orf25 |
| 19 | 1556788 | 1556979 | 8.190171342 | MEX3D |
| 19 | 1577655 | 1577856 | 1.136772867 | MBD3 |
| 19 | 1578421 | 1578608 | 9.742373103 | MBD3 |
| 19 | 1578421 | 1578608 | 9.742373103 | AC005943.2 |
| 1 | 100877595 | 100877800 | 1.027564764 | EXTL2 |
| 19 | 1782945 | 1783142 | -5.386569603 | ATP8B3 |
| 19 | 1827512 | 1827734 | 1.592297487 | REXO1 |
| 19 | 1854190 | 1854388 | 1.071433693 | KLF16 |
| 19 | 1877085 | 1877281 | 10.81052512 | ABHD17A |
| 19 | 1913142 | 1913322 | 8.362922757 | SCAMP4 |
| 19 | 1913142 | 1913322 | 8.362922757 | ADAT3 |
| 19 | 1980293 | 1980511 | 1.533439171 | CSNK1G2 |
| 19 | 2192026 | 2192224 | 1.591280087 | DOT1L |
| 19 | 2226499 | 2226724 | 12.69338203 | DOT1L |
| 1 | 103526083 | 103526289 | -7.272361499 | RNPC3 |
| 19 | 2353128 | 2353308 | 4.089413144 | SPPL2B |
| 19 | 2716334 | 2716535 | 1.053223731 | DIRAS1 |
| 19 | 2735390 | 2735504 | 6.483613129 | AC006538.4 |
| 19 | 2735390 | 2735504 | 6.483613129 | SLC39A3 |
| 19 | 2809265 | 2809475 | 1.736034354 | THOP1 |
| 19 | 2877951 | 2878151 | 1.903274381 | ZNF556 |
| 19 | 3118709 | 3118906 | 5.99540654 | AC005262.3 |
| 19 | 3118709 | 3118906 | 5.99540654 | GNA11 |
| 19 | 3121410 | 3121602 | 1.313617383 | AC005262.2 |
| 19 | 3121410 | 3121602 | 1.313617383 | GNA11 |
| 19 | 3207766 | 3207988 | 1.618598738 | NCLN |
| 19 | 3531354 | 3531528 | 8.184696434 | FZR1 |
| 19 | 3538415 | 3538654 | 1.014536549 | MFSD12 |
| 19 | 3562778 | 3562913 | 2.079935668 | MFSD12 |
| 19 | 3611264 | 3611460 | 3.029864828 | CACTIN |
| 19 | 3611264 | 3611460 | 3.029864828 | CACTIN-AS1 |
| 19 | 3657215 | 3657372 | -5.76080022 | PIP5K1C |
| 19 | 3958740 | 3958938 | 2.368169442 | DAPK3 |
| 19 | 3959021 | 3959263 | 12.00557847 | DAPK3 |
| 19 | 4047731 | 4047931 | 2.164080962 | ZBTB7A |
| 19 | 4054493 | 4054692 | 8.878528947 | ZBTB7A |
| 19 | 4216934 | 4217130 | 1.229674515 | ANKRD24 |
| 19 | 4538066 | 4538267 | 1.459007974 | CTB-50L17.14 |
| 19 | 4538066 | 4538267 | 1.459007974 | LRG1 |
| 19 | 4647344 | 4647542 | -5.088086575 | TNFAIP8L1 |
| 19 | 4647344 | 4647542 | -5.088086575 | MYDGF |
| 19 | 4652569 | 4652767 | -1.598737636 | TNFAIP8L1 |
| 19 | 4652569 | 4652767 | -1.598737636 | MYDGF |
| 19 | 4795035 | 4795235 | 2.006674427 | FEM1A |
| 19 | 4795035 | 4795235 | 2.006674427 | AC005523.2 |
| 19 | 5991198 | 5991326 | -1.69874456 | CTC-232P5.1 |
| 19 | 6057862 | 6058054 | -1.75763488 | RFX2 |
| 19 | 6189461 | 6189564 | 1.226708815 | RFX2 |
| 19 | 6189461 | 6189564 | 1.226708815 | ACSBG2 |
| 19 | 6414895 | 6415053 | 10.97675544 | KHSRP |
| 19 | 6583430 | 6583624 | -1.66688262 | CD70 |

| | | | | |
|---|---|---|---|---|
| 10 | 45826751 | 45826946 | -1.464268582 | AGAP4 |
| 10 | 45853862 | 45854014 | -1.545932913 | AGAP4 |
| 10 | 46809870 | 46810048 | -1.397590104 | BMS1P1 |
| 10 | 47502176 | 47502388 | -1.501057554 | AGAP9 |
| 10 | 47502176 | 47502388 | -1.501057554 | RP11-144G6.4 |
| 10 | 47529845 | 47530065 | -1.068344702 | RP11-144G6.12 |
| 10 | 49458864 | 49459066 | -1.005933659 | ERCC6 |
| 10 | 49458864 | 49459066 | -1.005933659 | RP11-123B3.2 |
| 10 | 50009080 | 50009349 | -1.615863841 | AGAP6 |
| 10 | 50009873 | 50010073 | -1.436592453 | AGAP6 |
| 10 | 51698233 | 51698439 | -1.324349677 | PRKG1 |
| 10 | 51698233 | 51698439 | -1.324349677 | CSTF2T |
| 10 | 52605336 | 52605496 | 1.816188346 | RP11-556E13.1 |
| 10 | 52730633 | 52730816 | 2.517453371 | RP11-556E13.1 |
| 10 | 52731337 | 52731535 | 1.469464777 | RP11-556E13.1 |
| 10 | 52734233 | 52734422 | 2.287622453 | RP11-556E13.1 |
| 10 | 58828875 | 58829050 | -2.728153884 | BICC1 |
| 10 | 59654177 | 59654377 | -2.161742276 | SLC16A9 |
| 10 | 60888337 | 60888544 | -1.217820231 | RHOBTB1 |
| 10 | 62053058 | 62053267 | 5.797628741 | ARID5B |
| 10 | 62092358 | 62092557 | 2.813628718 | ARID5B |
| 1 | 1082381 | 1082577 | -1.059012448 | C1orf159 |
| 1 | 15933852 | 15934049 | -1.565303787 | SPEN |
| 10 | 62093167 | 62093361 | 1.576554905 | ARID5B |
| 10 | 62197969 | 62198191 | -1.536679378 | RTKN2 |
| 10 | 62805681 | 62805881 | 1.580035193 | ADO |
| 10 | 62805681 | 62805881 | 1.580035193 | RP11-436D10.3 |
| 10 | 62813929 | 62814107 | 8.106534137 | EGR2 |
| 10 | 63465660 | 63465862 | -1.294499509 | JMJD1C-AS1 |
| 10 | 63465660 | 63465862 | -1.294499509 | JMJD1C |
| 10 | 64169560 | 64169665 | -1.686945033 | DBF4P1 |
| 10 | 65653141 | 65653302 | -1.809432704 | LINC01515 |
| 10 | 65654713 | 65654890 | -2.940672494 | LINC01515 |
| 10 | 65657245 | 65657419 | -2.622263536 | LINC01515 |
| 10 | 68644897 | 68645091 | -2.453594321 | TET1 |
| 10 | 68691451 | 68691670 | -3.174477058 | TET1 |
| 1 | 16231785 | 16231958 | -1.114767601 | RSG1 |
| 10 | 69123588 | 69123734 | -2.108372929 | VPS26A |
| 10 | 69208736 | 69208936 | -1.114829411 | SUPV3L1 |
| 10 | 70098540 | 70098738 | 6.909291089 | AIFM2 |
| 10 | 70098540 | 70098738 | 6.909291089 | H2AFY2 |
| 10 | 70108414 | 70108627 | 1.23641163 | H2AFY2 |
| 10 | 70108414 | 70108627 | 1.23641163 | AIFM2 |
| 10 | 70566729 | 70566950 | 1.725247083 | PALD1 |
| 10 | 70760662 | 70760861 | 4.949138733 | ADAMTS14 |
| 10 | 71362537 | 71362741 | 3.979723332 | SLC29A3 |
| 10 | 71713301 | 71713498 | 5.736852875 | C10orf105 |
| 10 | 71713301 | 71713498 | 5.736852875 | CDH23 |
| 10 | 71810198 | 71810383 | 3.929627422 | CDH23 |
| 10 | 73125063 | 73125198 | -6.487016805 | RP11-152N13.16 |
| 10 | 73125063 | 73125198 | -6.487016805 | NUDT13 |
| 10 | 73498455 | 73498643 | -7.188571366 | USP54 |
| 10 | 73498455 | 73498643 | -7.188571366 | PPP3CB-AS1 |
| 10 | 73498973 | 73499194 | -1.160392396 | PPP3CB-AS1 |
| 11 | 62531514 | 62531639 | -2.438924336 | AHNAK |
| 11 | 62531933 | 62532134 | -9.286749633 | AHNAK |
| 11 | 62532754 | 62533062 | -1.69461487 | AHNAK |
| 11 | 62533813 | 62534013 | -1.106754509 | AHNAK |
| 11 | 62578782 | 62578983 | -2.70430721 | RP11-864I4.1 |
| 11 | 62578782 | 62578983 | -2.70430721 | TUT1 |
| 11 | 62602205 | 62602489 | 1.104515616 | EML3 |
| 1 | 25863539 | 25863697 | -2.565622267 | PAQR7 |
| 11 | 62649444 | 62649643 | -1.045248451 | INTS5 |
| 11 | 62672055 | 62672285 | -1.261249207 | UQCC3 |
| 11 | 62672055 | 62672285 | -1.261249207 | LBHD1 |
| 11 | 62790434 | 62790667 | -1.596638421 | TMEM223 |
| 11 | 63763737 | 63763936 | -1.081135714 | C11orf95 |
| 11 | 63954844 | 63955044 | -1.936297132 | NAA40 |
| 11 | 64103420 | 64103578 | -5.363972657 | MACROD1 |
| 11 | 64103420 | 64103578 | -5.363972657 | FLRT1 |
| 11 | 64103420 | 64103578 | -5.363972657 | RP11-21A7A.2 |
| 11 | 64115712 | 64115912 | -2.687314637 | MACROD1 |
| 11 | 64115712 | 64115912 | -2.687314637 | FLRT1 |
| 11 | 64238488 | 64238687 | -1.34999067 | VEGFB |
| 11 | 64344461 | 64344661 | -1.212503721 | CCDC88B |
| 11 | 64349797 | 64349984 | -6.621140475 | CCDC88B |
| 1 | 26122195 | 26122394 | -1.560744496 | PDIK1L |
| 11 | 64450010 | 64450156 | -2.718702656 | AP003774.4 |
| 11 | 64810358 | 64810547 | -6.493454312 | MEN1 |
| 11 | 64854212 | 64854379 | 4.251243352 | EHD1 |
| 11 | 64854538 | 64854690 | 3.489191164 | EHD1 |
| 11 | 65040929 | 65041141 | -2.605690783 | SAC3D1 |
| 11 | 65044432 | 65044643 | -3.396470641 | SAC3D1 |
| 11 | 65079454 | 65079654 | -2.937234559 | CDCA5 |
| 11 | 65242162 | 65242349 | -1.890679857 | SLC22A20 |
| 11 | 65321329 | 65321575 | 1.717271798 | CDC42EP2 |
| 11 | 65424373 | 65424612 | 11.7027402 | NEAT1 |
| 1 | 26255365 | 26255564 | -5.210788237 | CEP85 |
| 11 | 65499486 | 65499627 | -13.67087184 | MALAT1 |
| 11 | 65654320 | 65654536 | 1.813135711 | RELA |
| 11 | 65726969 | 65727173 | -2.666305588 | KRT8P26 |
| 11 | 65794801 | 65794918 | -4.518086806 | OVOL1 |
| 11 | 65891381 | 65891593 | -1.281788599 | CCDC85B |
| 11 | 65998553 | 65998742 | 1.191311872 | EIF1AD |
| 11 | 66002120 | 66002304 | 6.513088798 | BANF1 |
| 11 | 66002120 | 66002304 | 6.513088798 | EIF1AD |
| 11 | 66345708 | 66345923 | -3.610692863 | B4GAT1 |
| 11 | 66346720 | 66346925 | -1.757548847 | B4GAT1 |
| 1 | 1204188 | 1204369 | 5.11136568 | TNFRSF18 |
| 1 | 26574346 | 26574547 | -1.104450863 | RPS6KA1 |
| 11 | 66669092 | 66669287 | -1.061388527 | RBM4B |
| 11 | 66677468 | 66677663 | -6.148314737 | RBM4B |
| 11 | 66686168 | 66686273 | -5.725605556 | SPTBN2 |
| 11 | 66700706 | 66700887 | 4.089954545 | SPTBN2 |
| 11 | 66779004 | 66779233 | -1.086436766 | C11orf80 |
| 11 | 66848962 | 66849161 | -2.285960871 | PC |
| 11 | 66857890 | 66858084 | -1.285139359 | LRFN4 |
| 11 | 66857890 | 66858084 | -1.285139359 | PC |
| 19 | 6593750 | 6593947 | -6.354865018 | RPL7P50 |
| 19 | 6593750 | 6593947 | -6.354865018 | CD70 |
| 1 | 107870353 | 107870541 | -6.658776968 | VAV3 |
| 19 | 7477586 | 7477729 | -6.273567766 | PEX11G |
| 19 | 7612378 | 7612698 | 3.008047843 | CAMSAP3 |
| 19 | 7741613 | 7741746 | 2.575037495 | CD209 |
| 19 | 7744260 | 7744455 | 2.562236374 | CD209 |
| 19 | 7745062 | 7745261 | 1.592909468 | CD209 |
| 19 | 7863435 | 7863595 | 7.126702164 | EVI5L |
| 1 | 108162060 | 108162261 | -5.02219706 | SLC25A24 |
| 19 | 8810260 | 8810437 | -7.375876626 | ZNF558 |
| 19 | 9674798 | 9674992 | 1.225855836 | ZNF562 |
| 1 | 108165575 | 108165767 | -1.578428385 | SLC25A24 |
| 19 | 10224141 | 10224338 | 7.956865456 | S1PR2 |
| 19 | 10224141 | 10224338 | 7.956865456 | DNMT1 |
| 19 | 10294540 | 10294743 | 2.256419061 | ICAM5 |
| 19 | 10334825 | 10335014 | 1.818495361 | ICAM3 |
| 19 | 10334825 | 10335014 | 1.818495361 | AC114271.2 |
| 19 | 10335222 | 10335422 | 2.231395315 | AC114271.2 |
| 19 | 10335222 | 10335422 | 2.231395315 | ICAM3 |
| 19 | 10643605 | 10643796 | -1.863510704 | SLC44A2 |
| 19 | 10653212 | 10653384 | -1.494963067 | ILF3-AS1 |
| 19 | 10869371 | 10869557 | 1.438511217 | C19orf38 |
| 19 | 11195717 | 11195910 | -1.502595765 | KANK2 |
| 19 | 11221085 | 11221255 | 5.658140766 | CTC-510F12.7 |
| 19 | 11221085 | 11221255 | 5.658140766 | DOCK6 |
| 19 | 11325061 | 11325258 | -1.295160519 | RAB3D |
| 19 | 11325061 | 11325258 | -1.295160519 | TSPAN16 |
| 1 | 108929691 | 108929889 | -1.159923964 | GPSM2 |
| 1 | 108929691 | 108929889 | -1.159923964 | RP11-475E11.9 |
| 1 | 108929691 | 108929889 | -1.159923964 | CLCC1 |
| 19 | 11680442 | 11680636 | -6.950669162 | CTC-499B15.5 |
| 19 | 11721618 | 11721818 | -1.7176615 | ZNF823 |
| 19 | 11904741 | 11904948 | -2.169346489 | ZNF69 |
| 19 | 11908656 | 11908762 | 2.38059604 | ZNF69 |
| 19 | 11909610 | 11909787 | -1.159059439 | ZNF69 |
| 1 | 109089183 | 109089414 | 1.820514282 | RP5-1065J22.8 |
| 19 | 11912530 | 11912719 | -1.558728791 | ZNF69 |
| 19 | 12043769 | 12043888 | 4.850219859 | CTD-2006C1.10 |
| 19 | 12043769 | 12043888 | 4.850219859 | CTD-2006C1.2 |
| 19 | 12043769 | 12043888 | 4.850219859 | ZNF878 |
| 1 | 109214608 | 109214803 | -1.19411374 | SARS |
| 19 | 12627911 | 12628033 | -1.000660887 | ZNF791 |
| 19 | 12627911 | 12628033 | -1.000660887 | ZNF490 |
| 19 | 12792045 | 12792247 | 2.415918206 | HOOK2 |
| 19 | 12792045 | 12792247 | 2.415918206 | JUNB |
| 1 | 109251295 | 109251477 | 3.22728823 | CELSR2 |
| 19 | 12792715 | 12792923 | 1.197006202 | JUNB |
| 19 | 12792715 | 12792923 | 1.197006202 | HOOK2 |
| 19 | 12894124 | 12894320 | -5.794825573 | RPS6P25 |
| 19 | 12894124 | 12894320 | -5.794825573 | GCDH |
| 1 | 109274548 | 109274765 | 1.440364017 | CELSR2 |
| 19 | 13762936 | 13763137 | 1.38319308 | CCDC130 |
| 19 | 14149787 | 14149970 | 7.549591705 | CTB-55O6.12 |

| | | | | |
|---|---|---|---|---|
| 10 | 73498973 | 73499194 | -1.160392396 | USP54 |
| 10 | 73516582 | 73516782 | -1.857103913 | PPP3CB-AS1 |
| 10 | 73516582 | 73516782 | -1.857103913 | USP54 |
| 10 | 73576218 | 73576402 | -1.193990834 | USP54 |
| 10 | 73626562 | 73626789 | -1.207908551 | RP11-464F9.20 |
| 10 | 73647436 | 73647687 | -2.556967526 | SYNPO2L |
| 10 | 73648515 | 73648688 | -1.452421383 | SYNPO2L |
| 10 | 73675431 | 73675631 | -1.382153667 | AGAP5 |
| 10 | 73675431 | 73675631 | -1.382153667 | RP11-464F9.1 |
| 10 | 73730321 | 73730528 | 1.346378127 | BMS1P4 |
| 10 | 73730321 | 73730528 | 1.346378127 | RP11-464F9.1 |
| 10 | 73733435 | 73733623 | -4.982892142 | GLUD1P3 |
| 10 | 73733435 | 73733623 | -4.982892142 | DUSP8P5 |
| 10 | 73779330 | 73779520 | -1.47747062 | FUT11 |
| 10 | 73783088 | 73783278 | -1.13512197 | CHCHD1 |
| 10 | 73792063 | 73792262 | 1.375613611 | ZSWIM8 |
| 1 | 16563988 | 16564183 | -6.791776465 | NBPF1 |
| 10 | 73813896 | 73814098 | -1.839356019 | CAMK2G |
| 10 | 73813896 | 73814098 | -1.839356019 | RP11-574K11.5 |
| 10 | 74352387 | 74352603 | -7.983613129 | ADK |
| 10 | 74363234 | 74363446 | -8.184517481 | ADK |
| 10 | 75150681 | 75150881 | 1.671507066 | SAMD8 |
| 10 | 75398674 | 75398879 | -2.164018757 | ZNF503 |
| 10 | 75398674 | 75398879 | -2.164018757 | RP11-399K21.11 |
| 10 | 75555751 | 75555953 | -1.266459558 | C10orf11 |
| 10 | 75578127 | 75578234 | -1.902390489 | C10orf11 |
| 10 | 75580892 | 75581094 | -1.712306879 | C10orf11 |
| 10 | 75688217 | 75688476 | -1.44806945 | C10orf11 |
| 1 | 1082669 | 1082824 | -1.84021191 | C1orf159 |
| 1 | 16597064 | 16597253 | 6.699371846 | NBPF1 |
| 10 | 76179385 | 76179588 | -1.483766879 | C10orf11 |
| 10 | 77780949 | 77781109 | 2.712607952 | RP13-39P12.2 |
| 10 | 77792211 | 77792435 | 1.309306445 | RP13-39P12.3 |
| 10 | 77792211 | 77792435 | 1.309306445 | DLG5 |
| 10 | 77821621 | 77821824 | 1.03951278 | DLG5 |
| 10 | 78054505 | 78054724 | 1.2086553 | RPS24 |
| 10 | 79313341 | 79313541 | 2.351097791 | ZMIZ1 |
| 10 | 79315278 | 79315471 | 1.7352842 | ZMIZ1 |
| 10 | 79353843 | 79354041 | -10.53256079 | PPIF |
| 10 | 79701170 | 79701383 | -1.978944182 | NUTM2B-AS1 |
| 10 | 79711698 | 79711873 | 6.226506677 | NUTM2B-AS1 |
| 10 | 79711698 | 79711873 | 6.226506677 | NUTM2B |
| 10 | 79826444 | 79826742 | -1.140591234 | NUTM2B-AS1 |
| 10 | 79826444 | 79826742 | -1.140591234 | RP11-182L21.6 |
| 1 | 17068851 | 17069073 | 3.189668226 | PADI2 |
| 10 | 79958364 | 79958534 | 6.896496796 | SFTPD |
| 1 | 17115541 | 17115727 | 5.938833787 | PADI2 |
| 10 | 86497253 | 86497456 | -1.025760846 | WAPL |
| 10 | 86709672 | 86709779 | -1.815025195 | LDB3 |
| 10 | 86943532 | 86943722 | 7.868782122 | MMRN2 |
| 10 | 86965900 | 86966103 | 2.003756217 | MMRN2 |
| 10 | 86965900 | 86966103 | 2.003756217 | ADIRF-AS1 |
| 10 | 86969127 | 86969326 | 1.163348384 | ADIRF |
| 10 | 86969127 | 86969326 | 1.163348384 | MMRN2 |
| 11 | 66860011 | 66860210 | 6.531360384 | LRFN4 |
| 11 | 66860011 | 66860210 | 6.531360384 | PC |
| 11 | 67049213 | 67049413 | -1.768041247 | SYT12 |
| 11 | 67397557 | 67397738 | 2.072127775 | RAD9A |
| 11 | 67398020 | 67398168 | 1.802909734 | RAD9A |
| 11 | 67409712 | 67409925 | -3.923298891 | PPP1CA |
| 11 | 67409712 | 67409925 | -3.923298891 | TBC1D10C |
| 11 | 67423803 | 67424011 | -1.289927868 | CARNS1 |
| 11 | 67425120 | 67425265 | -1.182930219 | CARNS1 |
| 11 | 67991356 | 67991604 | 2.2106902 | UNC93B1 |
| 11 | 68028036 | 68028221 | -1.083438827 | ALDH3B1 |
| 11 | 68028036 | 68028221 | -1.083438827 | RP5-901A4.1 |
| 11 | 68028544 | 68028772 | -2.243907698 | RP5-901A4.1 |
| 11 | 68028544 | 68028772 | -2.243907698 | ALDH3B1 |
| 11 | 68052150 | 68052347 | 6.166577675 | RP11-802E16.3 |
| 1 | 26731190 | 26731339 | -8.709156915 | ARID1A |
| 11 | 68980292 | 68980450 | -3.407165342 | MRGPRD |
| 11 | 69087926 | 69088138 | -1.103839228 | TPCN2 |
| 1 | 26774710 | 26774909 | -1.539579984 | ARID1A |
| 11 | 70206405 | 70206606 | -1.291594424 | FADD |
| 11 | 70206405 | 70206606 | -1.291594424 | RP11-805J14.5 |
| 11 | 72002577 | 72002773 | 2.075823458 | IL18BP |
| 11 | 72434167 | 72434362 | 1.258134732 | CLPB |
| 11 | 72838104 | 72838304 | 6.591119575 | ATG16L2 |
| 11 | 72838104 | 72838304 | 6.591119575 | FCHSD2 |
| 1 | 26787537 | 26787765 | 1.06595822 | PIGV |
| 11 | 73297570 | 73297785 | 2.156526614 | P2RY6 |
| 11 | 73310951 | 73311072 | -1.891939853 | ARHGEF17 |
| 11 | 73353126 | 73353248 | 4.406890596 | ARHGEF17 |
| 11 | 73367613 | 73367813 | -2.107496054 | ARHGEF17 |
| 11 | 73405084 | 73405289 | -1.603942274 | FAM168A |
| 11 | 74001182 | 74001306 | -5.967398296 | UCP3 |
| 11 | 74013202 | 74013334 | 1.393863058 | C2CD3 |
| 11 | 74128962 | 74129140 | -4.261780978 | C2CD3 |
| 11 | 74456995 | 74457206 | -2.482235928 | KCNE3 |
| 11 | 74456995 | 74457206 | -2.482235928 | RP11-702H23.4 |
| 11 | 74497759 | 74497926 | -1.132255532 | POLD3 |
| 11 | 74497759 | 74497926 | -1.132255532 | AP001372.2 |
| 11 | 75265807 | 75266026 | -2.32911442 | ARRB1 |
| 11 | 75571800 | 75572002 | -2.123445361 | SERPINH1 |
| 11 | 76141290 | 76141491 | -1.303984346 | UVRAG |
| 11 | 76381923 | 76382123 | -2.014678402 | RP11-111M22.2 |
| 11 | 76544300 | 76544502 | -1.158484892 | EMSY |
| 11 | 76796029 | 76796238 | -1.638235286 | TSKU |
| 11 | 76796556 | 76796686 | -1.696790246 | TSKU |
| 11 | 77701013 | 77701117 | -7.218355771 | RSF1 |
| 1 | 27534735 | 27534935 | -7.438642067 | AHDC1 |
| 11 | 78209739 | 78209946 | -2.211212622 | USP35 |
| 11 | 78218577 | 78218785 | -1.48740501 | GAB2 |
| 11 | 78226646 | 78226775 | -1.689769409 | GAB2 |
| 11 | 83165812 | 83166014 | -1.616022282 | PCF11 |
| 11 | 83166036 | 83166227 | -1.181199833 | PCF11 |
| 11 | 83167576 | 83167787 | -1.017271739 | PCF11 |
| 11 | 83213016 | 83213193 | -1.45817873 | RP11-727A23.1 |
| 19 | 14149787 | 14149970 | 7.549591705 | ADGRL1 |
| 19 | 14395211 | 14395410 | 1.710446851 | ADGRE5 |
| 19 | 14411264 | 14411456 | -1.699429962 | DDX39A |
| 19 | 14708898 | 14709064 | -1.906405944 | ZNF333 |
| 19 | 14767992 | 14768190 | -1.103031441 | ADGRE2 |
| 19 | 15398424 | 15398653 | 1.730413473 | AKAP8L |
| 19 | 16157381 | 16157505 | -1.883027792 | HSH2D |
| 19 | 16236751 | 16236881 | -4.930543453 | AP1M1 |
| 19 | 16501158 | 16501344 | -1.368294185 | C19orf44 |
| 19 | 16501158 | 16501344 | -1.368294185 | CTD-3222D19.2 |
| 19 | 16891614 | 16891857 | -1.293422593 | F2RL3 |
| 19 | 16893648 | 16893858 | -1.172164646 | CPAMD8 |
| 19 | 17305372 | 17305492 | 3.541497864 | ABHD8 |
| 19 | 17305372 | 17305492 | 3.541497864 | MRPL34 |
| 19 | 17319529 | 17319743 | 3.511603458 | DDA1 |
| 1 | 109665204 | 109665328 | -2.873616965 | GSTM4 |
| 19 | 17486385 | 17486588 | 1.684540976 | SLC27A1 |
| 19 | 17504586 | 17504785 | -2.443679669 | CTD-3131K8.2 |
| 19 | 17504586 | 17504785 | -2.443679669 | SLC27A1 |
| 19 | 18363468 | 18363669 | -1.211089576 | PGPEP1 |
| 19 | 18397035 | 18397154 | -4.123475063 | LRRC25 |
| 19 | 18434249 | 18434450 | 1.136460566 | SSBP4 |
| 19 | 18434249 | 18434450 | 1.136460566 | ISYNA1 |
| 19 | 18450547 | 18450745 | 10.69312361 | ELL |
| 19 | 18668885 | 18669076 | 1.73540421 | KLHL26 |
| 19 | 18919993 | 18920164 | -7.719477125 | DDX49 |
| 19 | 18919993 | 18920164 | -7.719477125 | AC002985.3 |
| 19 | 19506053 | 19506276 | 1.115609836 | GATAD2A |
| 1 | 109924904 | 109925099 | -1.269836177 | CSF1 |
| 19 | 19514171 | 19514291 | 2.229412148 | TSSK6 |
| 19 | 19724891 | 19725025 | -7.47891378 | ZNF14 |
| 19 | 19935060 | 19935231 | 1.282579077 | ZNF93 |
| 19 | 19935060 | 19935231 | 1.282579077 | AC007204.2 |
| 19 | 20227444 | 20227641 | -1.133746307 | CTC-260E6.6 |
| 1 | 110067851 | 110068050 | 1.785477821 | ALX3 |
| 1 | 110067851 | 110068050 | 1.785477821 | STRIP1 |
| 19 | 21057557 | 21057747 | -1.530700272 | ZNF430 |
| 19 | 21059196 | 21059395 | -1.04420143 | ZNF430 |
| 19 | 21116872 | 21117023 | -2.08164576 | ZNF714 |
| 19 | 21118567 | 21118762 | -1.515399656 | ZNF714 |
| 19 | 21182783 | 21182978 | -1.549604504 | ZNF431 |
| 1 | 110112761 | 110112961 | -2.59810039 | UBL4B |
| 19 | 21425819 | 21425923 | -1.675126682 | CTD-2561J22.3 |
| 19 | 21425819 | 21425923 | -1.675126682 | ZNF493 |
| 19 | 22677176 | 22677373 | -2.453445298 | CTC-457E21.9 |
| 19 | 22677919 | 22678112 | -2.29248125 | CTC-457E21.9 |
| 19 | 23101316 | 23101501 | 1.253743826 | ZNF730 |
| 1 | 110223499 | 110223686 | -2.667934417 | KCNC4 |
| 19 | 23259635 | 23259794 | -2.356359024 | RP11-15H20.8 |
| 19 | 23259635 | 23259794 | -2.356359024 | RP11-15H20.7 |
| 19 | 27699227 | 27699411 | -6.387187636 | LINC00662 |
| 19 | 27820272 | 27820381 | -4.56592848 | CTC-459F4.3 |
| 19 | 27820272 | 27820381 | -4.56592848 | LLNLF-65H9.1 |
| 19 | 29616814 | 29617013 | 2.588375823 | POP4 |

| | | | | |
|---|---|---|---|---|
| 10 | 86969127 | 86969326 | 1.163348384 | ADIRF-AS1 |
| 10 | 87009549 | 87009692 | 1.835968331 | RP11-96C23.14 |
| 10 | 87009549 | 87009692 | 1.835968331 | RP11-96C23.11 |
| 10 | 87009549 | 87009692 | 1.835968331 | AGAP11 |
| 10 | 87009549 | 87009692 | 1.835968331 | RP11-96C23.13 |
| 10 | 87151570 | 87151833 | -1.18423534 | FAM35A |
| 10 | 87342591 | 87342752 | -1.579610151 | LINC00863 |
| 10 | 87342591 | 87342752 | -1.579610151 | NUTM2A-AS1 |
| 10 | 87862731 | 87862915 | -3.434620128 | KLLN |
| 1 | 17697328 | 17697560 | 1.770003012 | ARHGEF10L |
| 10 | 88321819 | 88321919 | -1.544139307 | RNLS |
| 10 | 89306244 | 89306435 | 5.27266102 | LIPA |
| 10 | 89306244 | 89306435 | 5.27266102 | IFIT2 |
| 10 | 89307352 | 89307554 | 5.647580663 | IFIT2 |
| 10 | 89307352 | 89307554 | 5.647580663 | LIPA |
| 10 | 89338913 | 89339125 | 6.986166118 | IFIT3 |
| 10 | 89338913 | 89339125 | 6.986166118 | LIPA |
| 10 | 89402578 | 89402726 | 8.392173645 | LIPA |
| 10 | 89402578 | 89402726 | 8.392173645 | IFIT1 |
| 10 | 89418155 | 89418352 | 5.740538862 | IFIT5 |
| 10 | 89584285 | 89584464 | -2.411078774 | PANK1 |
| 10 | 89599403 | 89599623 | -3.110940607 | PANK1 |
| 10 | 89611797 | 89612076 | -1.990565266 | PANK1 |
| 10 | 89634284 | 89634481 | -1.144182803 | PANK1 |
| 10 | 89737432 | 89737620 | -3.424268976 | KIF20B |
| 1 | 19258783 | 19258983 | -1.184430728 | MRTO4 |
| 10 | 89738107 | 89738306 | -3.567714753 | KIF20B |
| 10 | 89765356 | 89765476 | -6.447976946 | KIF20B |
| 10 | 90748939 | 90749124 | 1.897646312 | HTR7 |
| 10 | 91259791 | 91259988 | -1.003459707 | PCGF5 |
| 10 | 91260491 | 91260691 | 3.937288485 | PCGF5 |
| 10 | 92959457 | 92959644 | 1.075825318 | EXOC6 |
| 10 | 93521440 | 93521636 | -7.966960564 | CEP55 |
| 10 | 93929919 | 93930039 | -3.47038559 | SLC35G1 |
| 10 | 93933395 | 93933586 | -1.25425522 | SLC35G1 |
| 10 | 93934746 | 93934946 | -6.838088738 | SLC35G1 |
| 10 | 95663563 | 95663756 | -1.335921691 | TCTN3 |
| 1 | 19327627 | 19327838 | -2.15237098 | PQLC2 |
| 10 | 96160003 | 96160263 | -1.042405572 | ZNF518A |
| 10 | 96665366 | 96665563 | -5.90024775 | PIK3AP1 |
| 1 | 789218 | 789408 | -1.321079696 | RP11-206L10.9 |
| 1 | 789218 | 789408 | -1.321079696 | RP5-857K21.4 |
| 1 | 1083551 | 1083758 | -1.61987456 | C1orf159 |
| 1 | 19328868 | 19329104 | -1.885595544 | PQLC2 |
| 10 | 97333512 | 97333710 | -1.185476735 | FRAT2 |
| 10 | 97338919 | 97339105 | 2.146741298 | RP11-452K12.4 |
| 10 | 97456918 | 97457140 | -1.324931661 | ZDHHC16 |
| 10 | 97570323 | 97570593 | 1.224511138 | UBTD1 |
| 10 | 97675443 | 97675603 | -7.087228802 | PI4K2A |
| 10 | 97677645 | 97677840 | 1.219960577 | AVPI1 |
| 10 | 98247999 | 98248220 | 3.291173276 | LOXL4 |
| 10 | 99704597 | 99704790 | -1.547854512 | CUTC |
| 10 | 99704597 | 99704790 | -1.547854512 | ENTPD7 |
| 10 | 99876858 | 99877057 | -1.513735581 | DNMBP |
| 11 | 83213016 | 83213193 | -1.45817873 | ANKRD42 |
| 11 | 83213016 | 83213193 | -1.45817873 | RP11-727A23.7 |
| 11 | 85757848 | 85758048 | -2.734924117 | SYTL2 |
| 11 | 86244778 | 86244979 | 1.036689906 | EED |
| 11 | 86951490 | 86951645 | 2.751428435 | PRSS23 |
| 11 | 86951490 | 86951645 | 2.751428435 | FZD4 |
| 11 | 93156881 | 93157077 | -1.75666405 | SLC36A4 |
| 11 | 93696968 | 93697124 | -1.293451978 | CEP295 |
| 11 | 93727329 | 93727542 | -1.016793682 | CEP295 |
| 11 | 93727329 | 93727542 | -1.016793682 | RP11-178H8.7 |
| 11 | 94180963 | 94181154 | 1.810392632 | PANX1 |
| 11 | 94181519 | 94181723 | 2.130271017 | PANX1 |
| 11 | 94543919 | 94544103 | -4.869890305 | FUT4 |
| 11 | 94543919 | 94544103 | -4.869890305 | PIWIL4 |
| 11 | 94799528 | 94799722 | -1.293245013 | AMOTL1 |
| 11 | 94870727 | 94870928 | -1.284505047 | AMOTL1 |
| 11 | 95070348 | 95070549 | -1.632526787 | SRSF8 |
| 11 | 95128551 | 95128810 | 4.716335865 | ENDOD1 |
| 11 | 95129398 | 95129597 | 3.949845904 | ENDOD1 |
| 11 | 95132320 | 95132518 | 3.837434946 | ENDOD1 |
| 11 | 95144828 | 95145027 | 5.766737873 | BUD13P1 |
| 11 | 95208892 | 95209058 | 5.736324268 | SESN3 |
| 11 | 95208892 | 95209058 | 5.736324268 | RP11-712B9.2 |
| 11 | 95811131 | 95811312 | -6.036233804 | CEP57 |
| 11 | 96074228 | 96074428 | 7.234312021 | MAML2 |
| 11 | 96341519 | 96341721 | 1.100729438 | MIR1260B |
| 11 | 96341519 | 96341721 | 1.100729438 | MAML2 |
| 11 | 96391890 | 96392091 | -1.133872431 | JRKL |
| 11 | 102324986 | 102325168 | 5.021091443 | BIRC3 |
| 11 | 107961994 | 107962197 | 3.40622487 | RAB39A |
| 11 | 108161498 | 108161621 | -7.076815597 | NPAT |
| 11 | 108173549 | 108173763 | -1.126878296 | NPAT |
| 11 | 108206663 | 108206832 | -1.009926528 | NPAT |
| 11 | 108207003 | 108207240 | -1.467485281 | NPAT |
| 11 | 108513237 | 108513356 | 2.624313586 | EXPH5 |
| 11 | 108514010 | 108514210 | 3.515626077 | EXPH5 |
| 11 | 111723863 | 111724064 | -1.669583003 | SIK2 |
| 11 | 111724725 | 111724915 | -1.357233719 | SIK2 |
| 11 | 111875439 | 111875638 | -8.134800778 | FDXACB1 |
| 11 | 111875439 | 111875638 | -8.134800778 | RP11-108O10.8 |
| 11 | 113224465 | 113224674 | 1.192170351 | NCAM1 |
| 11 | 113225547 | 113225685 | -5.955321365 | NCAM1 |
| 11 | 113226165 | 113226323 | 7.072010435 | NCAM1 |
| 11 | 113229724 | 113229922 | 1.313971187 | NCAM1 |
| 11 | 114400250 | 114400448 | -4.140716658 | C11orf71 |
| 11 | 114400250 | 114400448 | -4.140716658 | RBM7 |
| 11 | 114407528 | 114407728 | 1.184255171 | RBM7 |
| 11 | 114407528 | 114407728 | 1.184255171 | RP11-212D19.4 |
| 1 | 28466483 | 28466672 | 1.301305306 | PHACTR4 |
| 11 | 116849197 | 116849394 | 1.973326377 | APOA1-AS |
| 11 | 116849197 | 116849394 | 1.973326377 | SIK3 |
| 11 | 117995577 | 117995774 | 3.746624033 | IL10RA |
| 11 | 117998863 | 117999058 | 4.16009334 | IL10RA |
| 11 | 118502795 | 118502951 | -2.168987663 | KMT2A |
| 19 | 29674919 | 29675226 | 1.120421897 | PLEKHF1 |
| 19 | 31279485 | 31279627 | -3.825525846 | TSHZ3 |
| 19 | 32976860 | 32977015 | -1.064883937 | FAAP24 |
| 19 | 33205530 | 33205699 | 9.440795988 | LRP3 |
| 19 | 33207345 | 33207534 | 8.490292944 | LRP3 |
| 19 | 33207345 | 33207534 | 8.490292944 | CTD-2540B15.13 |
| 19 | 33301278 | 33301430 | 2.978885761 | CEBPA |
| 19 | 33301278 | 33301430 | 2.978885761 | CTD-2540B15.7 |
| 19 | 33399325 | 33399465 | 5.612603718 | PEPD |
| 19 | 34684663 | 34684783 | -1.318246401 | ZNF302 |
| 19 | 34741035 | 34741235 | -1.652097867 | ZNF181 |
| 19 | 34959290 | 34959501 | -1.025856935 | ZNF792 |
| 19 | 35347078 | 35347277 | -1.949257501 | CD22 |
| 19 | 35352766 | 35352948 | -4.46406322 | FFAR1 |
| 19 | 35543982 | 35544182 | 2.751280306 | GAPDHS |
| 19 | 35543982 | 35544182 | 2.751280306 | TMEM147-AS1 |
| 19 | 35721483 | 35721683 | 8.608297153 | KMT2B |
| 19 | 35940547 | 35940728 | 1.179275122 | LRFN3 |
| 19 | 36497411 | 36497608 | -2.39358852 | CTBP2P7 |
| 19 | 38396132 | 38396357 | -1.017429379 | SPRED3 |
| 19 | 38841877 | 38842076 | 6.806434249 | HNRNPL |
| 19 | 39307618 | 39307816 | 11.30031581 | LRFN1 |
| 19 | 39408026 | 39408226 | 1.57750109 | ZFP36 |
| 19 | 39532181 | 39532371 | -6.715879375 | EID2B |
| 19 | 40025225 | 40025420 | -2.982892142 | CTC-471F3.5 |
| 19 | 40036379 | 40036571 | 1.010782801 | ZNF780B |
| 19 | 40073705 | 40073887 | 1.061993636 | ZNF780A |
| 19 | 40073705 | 40073887 | 1.061993636 | AC005614.5 |
| 19 | 40074844 | 40075047 | 2.118766706 | AC005614.5 |
| 19 | 40074844 | 40075047 | 2.118766706 | ZNF780A |
| 19 | 40192214 | 40192483 | 2.972419929 | MAP3K10 |
| 19 | 40215114 | 40215329 | 1.159948961 | MAP3K10 |
| 19 | 40667396 | 40667620 | 1.633654497 | NUMBL |
| 19 | 40717443 | 40717608 | 1.069707474 | COQ8B |
| 19 | 40717443 | 40717608 | 1.069707474 | ITPKC |
| 19 | 40800884 | 40801083 | 10.90600363 | EGLN2 |
| 19 | 40800884 | 40801083 | 10.90600363 | RAB4B-EGLN2 |
| 19 | 41425503 | 41425703 | -1.939190868 | B3GNT8 |
| 19 | 41425503 | 41425703 | -1.939190868 | CTC-435M10.6 |
| 19 | 41426383 | 41426578 | -1.249184879 | B3GNT8 |
| 19 | 42089704 | 42089904 | 1.02092712 | POU2F2 |
| 19 | 42091013 | 42091199 | 1.41708552 | POU2F2 |
| 19 | 42137823 | 42137926 | -4.832667959 | POU2F2 |
| 19 | 42248743 | 42248948 | 11.1596875 | ERF |
| 19 | 42248743 | 42248948 | 11.1596875 | AC006486.9 |
| 19 | 42271926 | 42272056 | 7.928042523 | CIC |
| 19 | 42376600 | 42376753 | 2.628690828 | MEGF8 |
| 19 | 43594918 | 43595150 | 2.876267126 | IRGQ |
| 1 | 3770693 | 3770889 | -1.625714793 | CCDC27 |
| 1 | 111201849 | 111202025 | -6.18438963 | DENND2D |
| 1 | 111201849 | 111202025 | -6.18438963 | CHI3L2 |
| 19 | 43648852 | 43649052 | 9.303735598 | PLAUR |
| 19 | 43661671 | 43661868 | -1.142915471 | PLAUR |
| 1 | 111500252 | 111500446 | 4.094667623 | ADORA3 |

| | | | | |
|---|---|---|---|---|
| 10 | 99929748 | 99929997 | -1.589637252 | DNMBP-AS1 |
| 10 | 99929748 | 99929997 | -1.589637252 | DNMBP |
| 10 | 99956354 | 99956477 | -1.801879935 | DNMBP |
| 10 | 99956354 | 99956477 | -1.801879935 | DNMBP-AS1 |
| 10 | 100151784 | 100151989 | -2.939892246 | ERLIN1 |
| 10 | 100382630 | 100382867 | -3.272387717 | OLMALINC |
| 1 | 19682323 | 19682526 | -1.19021495 | TMCO4 |
| 10 | 100924144 | 100924339 | -1.131223802 | SLF2 |
| 10 | 100986426 | 100986624 | -1.495149093 | MRPL43 |
| 10 | 100987749 | 100987984 | -2.258682947 | C10orf2 |
| 10 | 100988427 | 100988616 | -2.366391658 | C10orf2 |
| 10 | 100988998 | 100989197 | -2.194887801 | C10orf2 |
| 1 | 19682931 | 19683048 | -1.074967517 | TMCO4 |
| 10 | 101060653 | 101060763 | -3.005413891 | RP11-108L7.15 |
| 10 | 101136889 | 101137116 | 1.895937672 | TLX1 |
| 10 | 101798062 | 101798262 | -7.740694758 | MGEA5 |
| 10 | 102139644 | 102139844 | -1.02695634 | PPRC1 |
| 10 | 102141701 | 102141900 | -2.222054586 | PPRC1 |
| 10 | 102643119 | 102643317 | 4.330889049 | RP11-47A8.5 |
| 10 | 102927245 | 102927445 | 1.285469967 | CNNM2 |
| 10 | 103388160 | 103388366 | -2.087008846 | TAF5 |
| 10 | 103425177 | 103425375 | -1.740514984 | PDCD11 |
| 10 | 103445408 | 103445620 | -1.616219211 | PDCD11 |
| 10 | 103446942 | 103447143 | 4.575236319 | CALHM2 |
| 10 | 103450028 | 103450228 | 5.124879014 | CALHM2 |
| 10 | 103450028 | 103450228 | 5.124879014 | RP11-225H22.7 |
| 10 | 103600539 | 103600723 | 7.296345308 | SH3PXD2A |
| 10 | 103600947 | 103601182 | 1.575445599 | SH3PXD2A |
| 10 | 103601541 | 103601670 | 2.097872906 | SH3PXD2A |
| 10 | 103601826 | 103602023 | 2.492019847 | SH3PXD2A |
| 1 | 20618573 | 20618772 | -3.73307117 | CDA |
| 10 | 103603282 | 103603481 | 2.725330705 | SH3PXD2A |
| 10 | 104268144 | 104268342 | -2.366556634 | MIR4482 |
| 10 | 104313402 | 104313594 | 1.120860993 | RP11-127L20.5 |
| 10 | 104313402 | 104313594 | 1.120860993 | ITPRIP |
| 10 | 110510472 | 110510679 | 4.32837484 | DUSP5 |
| 1 | 20648704 | 20648903 | -1.367575402 | PINK1 |
| 1 | 20648704 | 20648903 | -1.367575402 | PINK1-AS |
| 10 | 112595912 | 112596104 | 1.307813863 | VTI1A |
| 10 | 112808835 | 112809018 | -1.928012932 | VTI1A |
| 10 | 112810506 | 112810704 | -6.569775676 | VTI1A |
| 10 | 113849148 | 113849332 | -2.428006182 | DCLRE1A |
| 10 | 113852876 | 113853075 | -1.999279105 | DCLRE1A |
| 10 | 113876681 | 113876880 | -1.170781289 | NHLRC2 |
| 1 | 1087171 | 1087322 | 7.283978038 | C1orf159 |
| 10 | 114045484 | 114045744 | -2.209522516 | ADRB1 |
| 10 | 114125112 | 114125306 | 1.280357477 | CCDC186 |
| 10 | 114861541 | 114861712 | 1.034852604 | FAM160B1 |
| 10 | 118040347 | 118040545 | -1.019424262 | RAB11FIP2 |
| 10 | 118040347 | 118040545 | -1.019424262 | RP11-354M20.3 |
| 10 | 119042205 | 119042408 | -2.491561886 | EIF3A |
| 10 | 122211131 | 122211329 | 2.283352633 | TACC2 |
| 10 | 122888272 | 122888512 | -3.017888879 | RP11-564D11.3 |
| 10 | 122968631 | 122968760 | -1.955845791 | PSTK |

| | | | | |
|---|---|---|---|---|
| 11 | 118525274 | 118525430 | -1.167405714 | KMT2A |
| 11 | 118525274 | 118525430 | -1.167405714 | RP11-770J1.3 |
| 11 | 118628018 | 118628216 | -1.465322459 | PHLDB1 |
| 11 | 118628310 | 118628516 | -1.383593343 | PHLDB1 |
| 11 | 118751743 | 118751973 | -1.33251464 | DDX6 |
| 11 | 118786171 | 118786520 | -1.017657369 | DDX6 |
| 11 | 118898792 | 118898988 | 1.664724772 | BCL9L |
| 11 | 118899395 | 118899594 | 1.800452022 | BCL9L |
| 11 | 118902163 | 118902369 | 1.087317089 | BCL9L |
| 11 | 118996806 | 118996991 | -6.379319819 | RP11-110I1.12 |
| 11 | 119174612 | 119174795 | -4.024570797 | NLRX1 |
| 11 | 119334892 | 119335042 | -1.559676061 | RNF26 |
| 11 | 119356382 | 119356498 | 7.643567593 | USP2 |
| 11 | 119372875 | 119373023 | 6.218876036 | RP11-334E6.3 |
| 11 | 119372875 | 119373023 | 6.218876036 | USP2 |
| 11 | 119491332 | 119491527 | -1.56254761 | USP2-AS1 |
| 11 | 121086726 | 121087001 | -2.638598162 | TBCEL |
| 11 | 122809849 | 122810096 | -2.239499105 | UBASH3B |
| 11 | 122903976 | 122904175 | -2.858818053 | C11orf63 |
| 11 | 122934751 | 122934949 | -1.914290294 | C11orf63 |
| 11 | 123591038 | 123591204 | 2.585069836 | GRAMD1B |
| 11 | 123729712 | 123729912 | -7.340838828 | ZNF202 |
| 11 | 124746704 | 124746860 | 7.539367287 | NRGN |
| 11 | 124747325 | 124747520 | -1.78651256 | VSIG2 |
| 11 | 124748892 | 124749194 | -1.515005299 | VSIG2 |
| 11 | 124987751 | 124987968 | -1.911311969 | CCDC15 |
| 11 | 125086850 | 125087051 | -1.297930329 | SLC37A2 |
| 11 | 125088188 | 125088396 | -1.425881866 | SLC37A2 |
| 11 | 125895430 | 125895637 | 1.546718092 | HYLS1 |
| 11 | 125895430 | 125895637 | 1.546718092 | PUS3 |
| 11 | 125960877 | 125961060 | -4.07985159 | CDON |
| 11 | 126277812 | 126277976 | -1.000178883 | FOXRED1 |
| 11 | 126293955 | 126294141 | -1.875962997 | TIRAP |
| 11 | 126293955 | 126294141 | -1.875962997 | RP11-712L6.7 |
| 11 | 126341954 | 126342176 | -4.082273912 | DCPS |
| 11 | 126341954 | 126342176 | -4.082273912 | ST3GAL4-AS1 |
| 11 | 128810574 | 128810774 | -1.527458253 | FLI1 |
| 11 | 128969205 | 128969480 | -2.55275686 | ARHGAP32 |
| 11 | 128970609 | 128970772 | -1.922450073 | ARHGAP32 |
| 11 | 128972604 | 128972805 | -3.508848103 | ARHGAP32 |
| 11 | 129236434 | 129236628 | -1.982763891 | ARHGAP32 |
| 11 | 130108002 | 130108193 | -1.331158004 | APLP2 |
| 11 | 130108652 | 130108850 | -1.538401043 | APLP2 |
| 11 | 130261004 | 130261233 | -1.036714799 | ZBTB44 |
| 11 | 130405691 | 130405900 | -1.566895262 | ADAMTS8 |
| 1 | 29193345 | 29193555 | -1.19145429 | MECR |
| 11 | 130473891 | 130474093 | -2.967688638 | ADAMTS15 |
| 11 | 130476242 | 130476465 | -8.42146978 | ADAMTS15 |
| 11 | 130877643 | 130877926 | -1.401898551 | SNX19 |
| 11 | 134245590 | 134245787 | -1.802173087 | VPS26B |
| 11 | 134264948 | 134265156 | -1.411252058 | ACAD8 |
| 12 | 284599 | 284798 | -1.051686844 | KDM5A |
| 12 | 371489 | 371790 | -1.22308419 | KDM5A |
| 12 | 373612 | 373761 | -1.745520145 | KDM5A |

| | | | | |
|---|---|---|---|---|
| 1 | 111500252 | 111500446 | 4.094667623 | TMIGD3 |
| 1 | 111503406 | 111503568 | 3.330986905 | TMIGD3 |
| 1 | 111503406 | 111503568 | 3.330986905 | ADORA3 |
| 1 | 111596504 | 111596693 | 5.084537606 | RAP1A |
| 1 | 111599902 | 111600102 | 2.718842013 | LINC01160 |
| 1 | 111599902 | 111600102 | 2.718842013 | RAP1A |
| 19 | 45145594 | 45145760 | 7.889808385 | MARK4 |
| 19 | 45145594 | 45145760 | 7.889808385 | PPP1R37 |
| 19 | 45152860 | 45153000 | 1.355344907 | NKPD1 |
| 19 | 45152860 | 45153000 | 1.355344907 | MARK4 |
| 1 | 111608827 | 111609030 | 2.58925737 | RAP1A |
| 19 | 45529473 | 45529593 | 1.216775735 | OPA3 |
| 19 | 45712296 | 45712491 | 2.740273237 | FBXO46 |
| 19 | 45765498 | 45765678 | 1.031531588 | SIX5 |
| 19 | 45765498 | 45765678 | 1.031531588 | AC074212.5 |
| 1 | 111619297 | 111619497 | 10.37433311 | RAP1A |
| 19 | 45885600 | 45885777 | 7.402090725 | IRF2BP1 |
| 19 | 45890169 | 45890330 | 2.898273327 | MYPOP |
| 19 | 46620621 | 46620779 | 2.334442492 | PTGIR |
| 19 | 46755532 | 46755721 | 8.358409731 | FKRP |
| 19 | 46756460 | 46756669 | 1.679314302 | FKRP |
| 19 | 46775361 | 46775559 | 1.058211029 | SLC1A5 |
| 19 | 46775361 | 46775559 | 1.058211029 | FKRP |
| 1 | 111682386 | 111682566 | -6.569775676 | RAP1A |
| 1 | 111682386 | 111682566 | -6.569775676 | FAM212B |
| 19 | 46787803 | 46787979 | 1.935620349 | SLC1A5 |
| 19 | 46787803 | 46787979 | 1.935620349 | CTB-147N14.6 |
| 19 | 47066651 | 47066889 | 1.82553155 | ZC3H4 |
| 1 | 111724217 | 111724457 | 2.624847654 | FAM212B |
| 19 | 47320377 | 47320573 | -1.676518965 | C5AR1 |
| 19 | 47332134 | 47332244 | -1.023033425 | C5AR2 |
| 19 | 47341589 | 47341799 | -2.166933766 | C5AR2 |
| 19 | 47407156 | 47407346 | 1.360208874 | MEIS3 |
| 19 | 47680423 | 47680539 | 7.417235525 | CTD-2571L23.8 |
| 19 | 47680423 | 47680539 | 7.417235525 | GLTSCR1 |
| 1 | 111727272 | 111727545 | 2.876231309 | FAM212B |
| 19 | 47740969 | 47741182 | 2.390821885 | EHD2 |
| 19 | 47741517 | 47741648 | 2.236844162 | EHD2 |
| 19 | 48123881 | 48124079 | 5.86459793 | CTC-453G23.5 |
| 19 | 48123881 | 48124079 | 5.86459793 | LIG1 |
| 19 | 48195402 | 48195598 | -4.307354922 | ZNF114 |
| 19 | 48195402 | 48195598 | -4.307354922 | C19orf68 |
| 19 | 48195402 | 48195598 | -4.307354922 | CARD8 |
| 19 | 48286664 | 48286868 | -2.630303727 | ZNF114 |
| 19 | 48443921 | 48444118 | 1.983550785 | GRIN2D |
| 1 | 111766155 | 111766437 | 1.432478043 | DDX20 |
| 19 | 49150230 | 49150433 | 4.994152988 | PPFIA3 |
| 19 | 49470352 | 49470551 | 1.953763603 | ALDH16A1 |
| 19 | 49470352 | 49470551 | 1.953763603 | CTD-3148I10.9 |
| 19 | 49525037 | 49525237 | -1.141553539 | FCGRT |
| 19 | 49651303 | 49651492 | 11.11369868 | SCAF1 |
| 19 | 49654221 | 49654388 | 1.240243971 | SCAF1 |
| 19 | 49857848 | 49858031 | 8.483511784 | PTOV1 |
| 19 | 49857848 | 49858031 | 8.483511784 | PTOV1-AS2 |

| | | | | |
|---|---|---|---|---|
| 10 | 122970877 | 122971076 | -1.822762013 | PSTK |
| 1 | 20941856 | 20942056 | -1.056324283 | EIF4G3 |
| 10 | 122971416 | 122971564 | -8.630827108 | PSTK |
| 10 | 123056188 | 123056382 | -1.309673966 | ACADSB |
| 10 | 124989414 | 124989615 | -1.466834335 | CTBP2 |
| 10 | 125000992 | 125001118 | -6.959058926 | CTBP2 |
| 10 | 125000992 | 125001118 | -6.959058926 | RP11-59C5.3 |
| 10 | 125836025 | 125836227 | -1.697433437 | BCCIP |
| 10 | 125842483 | 125842651 | -3.475463894 | BCCIP |
| 10 | 125842483 | 125842651 | -3.475463894 | DHX32 |
| 10 | 125880825 | 125881012 | 1.469537981 | DHX32 |
| 10 | 127934348 | 127934531 | -4.701737231 | PTPRE |
| 10 | 127934348 | 127934531 | -4.701737231 | RP11-4C20.3 |
| 1 | 21219759 | 21220012 | 1.235550023 | ECE1 |
| 10 | 127935744 | 127935866 | -4.150933 | PTPRE |
| 10 | 127935744 | 127935866 | -4.150933 | RP11-4C20.4 |
| 10 | 128098805 | 128099027 | -3.805967884 | MKI67 |
| 10 | 128101454 | 128101622 | -3.953925381 | MKI67 |
| 10 | 128103969 | 128104150 | -5.073126777 | MKI67 |
| 10 | 128104388 | 128104587 | -4.429468447 | MKI67 |
| 10 | 128106207 | 128106402 | -4.478511404 | MKI67 |
| 10 | 128108097 | 128108296 | -3.814301728 | MKI67 |
| 10 | 128108811 | 128109011 | -4.296116298 | MKI67 |
| 10 | 128109153 | 128109268 | -3.842137631 | MKI67 |
| 10 | 128115235 | 128115341 | -3.202460022 | MKI67 |
| 10 | 128115389 | 128115564 | -3.407464439 | MKI67 |
| 10 | 129578292 | 129578474 | -6.450999093 | MGMT |
| 10 | 129837744 | 129837867 | -3.034034572 | RP11-234G16.4 |
| 10 | 129837744 | 129837867 | -3.034034572 | EBF3 |
| 10 | 131911674 | 131911875 | 7.80453164 | PPP2R2D |
| 10 | 131970407 | 131970577 | 3.968545417 | BNIP3 |
| 10 | 131971157 | 131971356 | 9.34216543 | BNIP3 |
| 10 | 131971157 | 131971356 | 9.34216543 | RP11-45A17.2 |
| 10 | 131996776 | 131997043 | 1.413008588 | RP11-45A17.3 |
| 10 | 132190111 | 132190291 | 4.307920907 | DPYSL4 |
| 10 | 132192389 | 132192549 | 5.061662781 | DPYSL4 |
| 10 | 132201745 | 132201931 | 4.034836764 | DPYSL4 |
| 10 | 132405524 | 132405698 | -1.663385556 | PWWP2B |
| 10 | 132406608 | 132406806 | -2.283309099 | PWWP2B |
| 10 | 132416931 | 132417124 | 5.772964885 | PWWP2B |
| 10 | 132434535 | 132434705 | -2.85786443 | RP11-432J24.3 |
| 10 | 132439046 | 132439235 | -3.695738096 | RP11-432J24.3 |
| 10 | 132448181 | 132448381 | -4.098484344 | C10orf91 |
| 10 | 132762567 | 132762764 | -7.380100076 | INPP5A |
| 10 | 132771555 | 132771765 | -6.40839184 | INPP5A |
| 1 | 1088576 | 1088766 | 6.821928095 | C1orf159 |
| 10 | 132772322 | 132772520 | -5.786113256 | INPP5A |
| 10 | 133420064 | 133420251 | -3.103490348 | MTG1 |
| 10 | 133420064 | 133420251 | -3.103490348 | RP11-108K14.8 |
| 10 | 133476833 | 133476989 | 4.303665157 | SCART1 |
| 10 | 133481033 | 133481211 | 6.873362122 | SCART1 |
| 11 | 207824 | 208031 | -1.117827999 | RIC8A |
| 11 | 214458 | 214700 | -1.051179341 | RIC8A |
| 11 | 295123 | 295336 | -1.664562086 | PGGHG |

| | | | | |
|---|---|---|---|---|
| 12 | 553503 | 553689 | -1.276940767 | B4GALNT3 |
| 12 | 561601 | 561799 | -1.891714541 | B4GALNT3 |
| 12 | 643967 | 644170 | -9.274948725 | NINJ2 |
| 12 | 643967 | 644170 | -9.274948725 | RP11-218M22.1 |
| 12 | 813513 | 813722 | 11.65512873 | WNK1 |
| 12 | 868043 | 868244 | -1.357114864 | WNK1 |
| 12 | 896233 | 896438 | -1.391239796 | WNK1 |
| 12 | 908633 | 908772 | -1.538708087 | WNK1 |
| 12 | 917855 | 918042 | -1.811663626 | RAD52 |
| 12 | 975822 | 976021 | -1.837558469 | RAD52 |
| 12 | 975822 | 976021 | -1.837558469 | RP11-359B12.2 |
| 12 | 976401 | 976587 | -3.304743289 | RAD52 |
| 12 | 976401 | 976587 | -3.304743289 | RP11-359B12.2 |
| 12 | 976840 | 977028 | -3.621348884 | RAD52 |
| 12 | 976840 | 977028 | -3.621348884 | RP11-359B12.2 |
| 12 | 1490512 | 1490716 | -1.30541193 | ERC1 |
| 12 | 1493132 | 1493361 | -1.269020727 | ERC1 |
| 12 | 1493704 | 1493865 | -1.559372767 | ERC1 |
| 12 | 1494222 | 1494441 | -1.406634268 | ERC1 |
| 12 | 1786105 | 1786306 | -1.050685054 | ADIPOR2 |
| 12 | 1787876 | 1788077 | -1.453126325 | ADIPOR2 |
| 12 | 1792948 | 1793116 | -4.048073248 | CACNA2D4 |
| 12 | 2859032 | 2859271 | -1.683422876 | FOXM1 |
| 12 | 2859032 | 2859271 | -1.683422876 | ITFG2 |
| 12 | 2874073 | 2874272 | -1.873982297 | FOXM1 |
| 12 | 2888173 | 2888392 | -1.091836896 | RHNO1 |
| 12 | 2888173 | 2888392 | -1.091836896 | TULP3 |
| 12 | 2889067 | 2889268 | -1.172234512 | TULP3 |
| 12 | 2889067 | 2889268 | -1.172234512 | RHNO1 |
| 12 | 2940714 | 2940843 | -2.269462758 | TULP3 |
| 12 | 3040379 | 3040584 | 1.418842266 | TEAD4 |
| 1 | 790194 | 790392 | -1.866919901 | RP5-857K21.4 |
| 1 | 790194 | 790392 | -1.866919901 | RP11-206L10.9 |
| 12 | 4520400 | 4520598 | -4.497176718 | C12orf4 |
| 12 | 4523499 | 4523685 | -6.118007096 | C12orf4 |
| 12 | 6018814 | 6019013 | -2.270522629 | VWF |
| 12 | 6329278 | 6329480 | 6.0691359 | TNFRSF1A |
| 12 | 6493575 | 6493869 | -2.371654414 | NCAPD2 |
| 12 | 6493575 | 6493869 | -2.371654414 | MRPL51 |
| 12 | 6619827 | 6620031 | -4.183648561 | LPAR5 |
| 12 | 6644352 | 6644543 | 5.04637857 | ACRBP |
| 12 | 6827113 | 6827313 | -1.917355106 | GPR162 |
| 12 | 6871204 | 6871411 | -2.609846965 | SPSB2 |
| 12 | 6872344 | 6872545 | -1.337720823 | SPSB2 |
| 12 | 7070468 | 7070698 | 1.772340998 | C1S |
| 12 | 7813947 | 7814207 | -1.179829242 | SLC2A14 |
| 12 | 8033089 | 8033288 | -1.2588819 | FOXJ2 |
| 12 | 8059954 | 8060130 | 2.790422395 | C3AR1 |
| 12 | 8235494 | 8235598 | -2.984903186 | RP11-266K4.9 |
| 12 | 8235494 | 8235598 | -2.984903186 | FAM86FP |
| 12 | 8238573 | 8238807 | -6.900854179 | RP11-266K4.9 |
| 12 | 8238573 | 8238807 | -6.900854179 | FAM86FP |
| 12 | 8776723 | 8776944 | -1.728675987 | A2ML1-AS1 |
| 12 | 8776723 | 8776944 | -1.728675987 | RIMKLB |

| | | | | |
|---|---|---|---|---|
| 19 | 49880533 | 49880725 | 1.471706538 | TBC1D17 |
| 19 | 49881205 | 49881403 | 7.513952998 | TBC1D17 |
| 19 | 49884434 | 49884639 | 1.312063411 | TBC1D17 |
| 19 | 49887903 | 49888097 | 1.548578119 | TBC1D17 |
| 19 | 49889763 | 49889972 | 2.672098491 | IL4I1 |
| 19 | 49940545 | 49940714 | -1.204695468 | CTC-326K19.6 |
| 19 | 50666627 | 50666898 | 2.026642397 | SHANK1 |
| 19 | 51413801 | 51414028 | 1.238347091 | SIGLEC10 |
| 1 | 112853846 | 112854045 | -7.461686958 | RP3-522D1.1 |
| 19 | 51992436 | 51992568 | -4.93518236 | ZNF615 |
| 1 | 112855282 | 112855514 | -1.126490371 | RP3-522D1.1 |
| 19 | 52035027 | 52035218 | -1.240395293 | ZNF432 |
| 19 | 52155382 | 52155483 | -1.189966849 | ZNF836 |
| 19 | 52155382 | 52155483 | -1.189966849 | CTC-471J1.8 |
| 19 | 52156788 | 52156987 | -7.476552575 | CTC-471J1.8 |
| 19 | 52156788 | 52156987 | -7.476552575 | ZNF836 |
| 19 | 52389143 | 52389430 | -1.174724579 | ZNF528-AS1 |
| 19 | 52416747 | 52416877 | -1.287035327 | ZNF528 |
| 1 | 113683119 | 113683272 | -1.385617748 | MAGI3 |
| 19 | 53875714 | 53875905 | 5.655874158 | MYADM |
| 19 | 53875714 | 53875905 | 5.655874158 | AC008440.5 |
| 19 | 54096378 | 54096559 | 4.916445007 | OSCAR |
| 1 | 113683710 | 113683916 | 1.339955452 | MAGI3 |
| 19 | 54354871 | 54355074 | -2.416874979 | LAIR1 |
| 19 | 54462847 | 54463008 | 1.388067493 | LENG9 |
| 19 | 55091177 | 55091376 | 1.102550444 | PPP1R12C |
| 19 | 55096869 | 55097062 | 6.352600189 | PPP1R12C |
| 19 | 55339068 | 55339205 | -1.085963677 | CTD-2105E13.16 |
| 19 | 55453062 | 55453262 | 1.088754368 | ISOC2 |
| 19 | 55500508 | 55500707 | 2.597261826 | SSC5D |
| 19 | 55602738 | 55602931 | 3.690025524 | ZNF524 |
| 19 | 55615572 | 55615755 | 9.004777644 | ZNF865 |
| 19 | 56087680 | 56087897 | 1.295389914 | ZNF787 |
| 19 | 57253185 | 57253382 | -1.016763379 | ZNF805 |
| 1 | 3779379 | 3779498 | -1.230948609 | LRRC47 |
| 1 | 113981188 | 113981417 | 2.105821966 | OLFML3 |
| 19 | 57378267 | 57378447 | -6.715879375 | AC003002.4 |
| 19 | 57378267 | 57378447 | -6.715879375 | ZNF547 |
| 19 | 57445176 | 57445315 | 1.270105929 | ZNF749 |
| 19 | 57445176 | 57445315 | 1.270105929 | AC004076.9 |
| 1 | 114397601 | 114397801 | -1.070674167 | TRIM33 |
| 19 | 57474078 | 57474280 | 1.309079905 | AC004076.9 |
| 19 | 57474078 | 57474280 | 1.309079905 | ZNF772 |
| 19 | 57492839 | 57493018 | -1.251446672 | ZNF419 |
| 19 | 57492839 | 57493018 | -1.251446672 | AC003005.4 |
| 19 | 57814686 | 57814870 | 1.822864302 | ZNF552 |
| 19 | 57814686 | 57814870 | 1.822864302 | ZNF586 |
| 19 | 57907500 | 57907681 | 1.267766396 | CTD-2583A14.9 |
| 19 | 57907500 | 57907681 | 1.267766396 | ZNF417 |
| 19 | 58009410 | 58009603 | -6.073602462 | CTD-2368P22.1 |
| 19 | 58349863 | 58350050 | 1.105160831 | CTD-2619J13.8 |
| 19 | 58349863 | 58350050 | 1.105160831 | A1BG |
| 19 | 58349863 | 58350050 | 1.105160831 | A1BG-AS1 |
| 19 | 58368807 | 58368967 | 9.126249564 | ZNF837 |

| | | | | |
|---|---|---|---|---|
| 11 | 324831 | 325159 | 4.005699651 | IFITM3 |
| 11 | 324831 | 325159 | 4.005699651 | RP11-326C3.11 |
| 11 | 434353 | 434626 | 3.886873265 | ANO9 |
| 11 | 451520 | 451709 | -2.811280238 | PTDSS2 |
| 11 | 452138 | 452368 | -2.347191582 | PTDSS2 |
| 11 | 453109 | 453353 | -2.24499087 | PTDSS2 |
| 11 | 453922 | 454111 | -2.75112034 | PTDSS2 |
| 11 | 459272 | 459497 | -1.684969003 | PTDSS2 |
| 11 | 529039 | 529242 | -2.484215517 | RP13-46H24.1 |
| 11 | 532370 | 532576 | -2.99899763 | HRAS |
| 11 | 532939 | 533150 | -2.310850972 | HRAS |
| 11 | 536254 | 536396 | -1.660575076 | HRAS |
| 11 | 536454 | 536720 | -1.348229934 | HRAS |
| 11 | 538578 | 538702 | -2.860258876 | LRRC56 |
| 11 | 545321 | 545507 | -1.306169027 | LRRC56 |
| 11 | 546576 | 546760 | -3.014311178 | LRRC56 |
| 11 | 563720 | 563919 | -1.059302379 | RASSF7 |
| 11 | 566293 | 566488 | -8.652777068 | MIR210HG |
| 11 | 567272 | 567472 | -4.873874824 | MIR210HG |
| 11 | 607369 | 607571 | -1.68333528 | PHRF1 |
| 11 | 608429 | 608730 | -2.102829398 | PHRF1 |
| 11 | 609196 | 609433 | -2.114178208 | PHRF1 |
| 11 | 780426 | 780598 | -2.309218327 | AP006621.5 |
| 11 | 780426 | 780598 | -2.309218327 | CMB9-55F22.1 |
| 11 | 797256 | 797450 | 2.112674833 | SLC25A22 |
| 11 | 804711 | 804909 | -1.202315342 | PIDD1 |
| 11 | 869094 | 869300 | -1.327678723 | CHID1 |
| 11 | 1010728 | 1010849 | -1.58586266 | AP2A2 |
| 11 | 1276968 | 1277163 | 1.153944694 | TOLLIP |
| 11 | 1453421 | 1453607 | 6.281121212 | BRSK2 |
| 1 | 23310566 | 23310790 | -2.30941879 | HNRNPR |
| 11 | 2313029 | 2313221 | 5.56592848 | TSPAN32 |
| 11 | 2328882 | 2329076 | 6.718095498 | CD81-AS1 |
| 11 | 2371813 | 2371945 | 2.293271768 | CD81-AS1 |
| 1 | 23361604 | 23361732 | -1.018954465 | ZNF436 |
| 11 | 2509383 | 2509561 | -5.485479655 | KCNQ1 |
| 11 | 2698468 | 2698632 | -2.545665688 | KCNQ1OT1 |
| 11 | 2698468 | 2698632 | -2.545665688 | KCNQ1 |
| 11 | 3132609 | 3132805 | -9.335805085 | OSBPL5 |
| 11 | 3133668 | 3133830 | -10.14458618 | OSBPL5 |
| 11 | 3359244 | 3359447 | -1.147185356 | ZNF195 |
| 11 | 3359998 | 3360222 | -1.146777557 | ZNF195 |
| 11 | 3665908 | 3666102 | -1.36677161 | CHRNA10 |
| 11 | 3676201 | 3676321 | -2.072364135 | NUP98 |
| 1 | 23362354 | 23362511 | -1.139986986 | ZNF436 |
| 11 | 3712419 | 3712590 | -6.485142838 | NUP98 |
| 11 | 3827865 | 3828065 | -1.243563087 | RHOG |
| 11 | 3855226 | 3855478 | -1.18089362 | AC090587.5 |
| 11 | 3855226 | 3855478 | -1.18089362 | STIM1 |
| 11 | 3855226 | 3855478 | -1.18089362 | AC090587.4 |
| 11 | 3856113 | 3856278 | -2.065475781 | STIM1 |
| 11 | 4475192 | 4475387 | 7.247802616 | OR52K3P |
| 1 | 1090210 | 1090350 | 8.19592386 | C1orf159 |
| 1 | 23509750 | 23509944 | -3.895523288 | E2F2 |
| 12 | 9753204 | 9753402 | 4.914045404 | CD69 |
| 1 | 32207789 | 32208050 | -1.524633684 | IQCC |
| 12 | 10847257 | 10847447 | -5.985592136 | PRH1 |
| 12 | 10847257 | 10847447 | -5.985592136 | PRR4 |
| 12 | 10847257 | 10847447 | -5.985592136 | PRH1-PRR4 |
| 12 | 11220100 | 11220238 | -2.550802362 | RP11-711K1.8 |
| 12 | 11379317 | 11379491 | -2.001540793 | PRB1 |
| 12 | 11382716 | 11383030 | -1.312854422 | PRB1 |
| 12 | 11500143 | 11500280 | -5.538407799 | PRB2 |
| 12 | 11720628 | 11720816 | -1.628458148 | ETV6 |
| 12 | 11824840 | 11824942 | 5.882435795 | ETV6 |
| 12 | 12165222 | 12165387 | -7.096069336 | BCL2L14 |
| 12 | 12165222 | 12165387 | -7.096069336 | LRP6 |
| 12 | 12181169 | 12181385 | -1.063063697 | LRP6 |
| 12 | 12181169 | 12181385 | -1.063063697 | BCL2L14 |
| 12 | 12203296 | 12203494 | -1.408144023 | BCL2L14 |
| 12 | 12203296 | 12203494 | -1.408144023 | LRP6 |
| 12 | 12465722 | 12465920 | -7.374934714 | BORCS5 |
| 12 | 12476959 | 12477146 | 1.384482787 | DUSP16 |
| 12 | 12477336 | 12477536 | 1.146808426 | DUSP16 |
| 12 | 12717877 | 12718080 | -1.030568373 | CDKN1B |
| 12 | 12774030 | 12774217 | 2.566234564 | APOLD1 |
| 12 | 12788787 | 12788986 | 1.097004085 | APOLD1 |
| 12 | 13002137 | 13002387 | -2.297491065 | HTR7P1 |
| 12 | 13002137 | 13002387 | -2.297491065 | RP11-377D9.3 |
| 12 | 13002711 | 13002906 | -2.676075214 | HTR7P1 |
| 12 | 13002711 | 13002906 | -2.676075214 | RP11-377D9.3 |
| 12 | 13004247 | 13004414 | -2.303729377 | HTR7P1 |
| 12 | 13004247 | 13004414 | -2.303729377 | RP11-377D9.3 |
| 12 | 14423994 | 14424182 | -1.406062466 | ATF7IP |
| 12 | 14424736 | 14424937 | -2.522265165 | ATF7IP |
| 12 | 14460635 | 14460899 | -2.118873372 | ATF7IP |
| 12 | 14497788 | 14497990 | -1.747070397 | ATF7IP |
| 12 | 19274690 | 19274887 | -2.120322063 | PLEKHA5 |
| 12 | 19283407 | 19283607 | -2.057298629 | PLEKHA5 |
| 12 | 19462412 | 19462615 | -1.607148368 | AEBP2 |
| 12 | 19582679 | 19582932 | -1.579767982 | AEBP2 |
| 1 | 1284551 | 1284709 | 2.785671845 | SCNN1D |
| 12 | 21515736 | 21515860 | -1.285687079 | GOLT1B |
| 12 | 23820795 | 23820931 | 1.072901639 | SOX5 |
| 12 | 25052386 | 25052579 | -3.114092895 | LRMP |
| 12 | 25107881 | 25108071 | -1.895795156 | LRMP |
| 12 | 26122758 | 26123090 | 1.254013206 | SSPN |
| 12 | 26122758 | 26123090 | 1.254013206 | BHLHE41 |
| 12 | 26974018 | 26974228 | -1.600551021 | RP11-421F16.3 |
| 12 | 26974018 | 26974228 | -1.600551021 | TM7SF3 |
| 12 | 27405173 | 27405327 | 2.572388993 | ARNTL2-AS1 |
| 12 | 27405173 | 27405327 | 2.572388993 | ARNTL2 |
| 12 | 27406787 | 27406990 | 8.215013891 | ARNTL2 |
| 12 | 27406787 | 27406990 | 8.215013891 | ARNTL2-AS1 |
| 12 | 27410293 | 27410496 | 2.711004554 | ARNTL2-AS1 |
| 12 | 27410293 | 27410496 | 2.711004554 | ARNTL2 |
| 12 | 27595376 | 27595567 | -4.906890596 | PPFIBP1 |
| 12 | 27797755 | 27797961 | -1.167724665 | KLHL42 |
| 19 | 58369799 | 58369994 | 3.245926548 | ZNF837 |
| 19 | 58440384 | 58440514 | 2.330125314 | CTD-2619J13.19 |
| 19 | 58472375 | 58472579 | 1.048607364 | ZNF324 |
| 19 | 58516510 | 58516707 | 1.846079982 | ZBTB45 |
| 19 | 58517400 | 58517529 | 1.151162979 | ZBTB45 |
| 19 | 58519146 | 58519350 | 1.538632334 | ZBTB45 |
| 19 | 58563096 | 58563264 | 8.230099948 | MZF1 |
| 19 | 58563096 | 58563264 | 8.230099948 | MZF1-AS1 |
| 19 | 58573603 | 58573871 | 1.66924481 | CENPBD1P1 |
| 19 | 58573603 | 58573871 | 1.66924481 | MZF1-AS1 |
| 2 | 257907 | 258114 | -1.190844997 | SH3YL1 |
| 2 | 3227267 | 3227372 | 6.554915327 | TSSC1 |
| 1 | 116046266 | 116046469 | -1.998078967 | SLC22A15 |
| 2 | 3461976 | 3462110 | 7.806650137 | TRAPPC12 |
| 2 | 6849514 | 6849716 | -1.257146597 | CMPK2 |
| 1 | 116067092 | 116067214 | -2.144394856 | SLC22A15 |
| 2 | 10039822 | 10040069 | -1.304160758 | RP11-254F7.2 |
| 2 | 10052407 | 10052529 | -1.054396414 | KLF11 |
| 2 | 10399894 | 10400097 | -1.073174319 | HPCAL1 |
| 2 | 10913818 | 10914010 | 1.276081566 | KCNF1 |
| 2 | 11446894 | 11447061 | -1.668524691 | E2F6 |
| 2 | 17781480 | 17781696 | -1.169100496 | GEN1 |
| 2 | 17781480 | 17781696 | -1.169100496 | SMC6 |
| 2 | 19278650 | 19278838 | -6.61920237 | LINC01376 |
| 2 | 20202484 | 20202680 | 2.716949178 | SDC1 |
| 2 | 20291515 | 20291714 | -3.260290137 | PUM2 |
| 2 | 23691514 | 23691710 | -6.412876916 | KLHL29 |
| 2 | 23927344 | 23927558 | 1.787955019 | UBXN2A |
| 2 | 24037873 | 24038020 | -4.642701109 | WDCP |
| 2 | 24037873 | 24038020 | -4.642701109 | MFSD2B |
| 2 | 24038852 | 24039050 | 1.228340641 | WDCP |
| 2 | 24038852 | 24039050 | 1.228340641 | MFSD2B |
| 2 | 24097294 | 24097514 | -1.210990489 | FAM228B |
| 2 | 24144600 | 24144811 | 1.086310562 | RP11-507M3.1 |
| 2 | 24144600 | 24144811 | 1.086310562 | FAM228B |
| 2 | 24819947 | 24820146 | 8.879290542 | ADCY3 |
| 2 | 24819947 | 24820146 | 8.879290542 | CENPO |
| 2 | 24826308 | 24826504 | 6.825862217 | RP11-443B20.1 |
| 2 | 24826308 | 24826504 | 6.825862217 | ADCY3 |
| 2 | 25326235 | 25326435 | -1.071599752 | DNMT3A |
| 2 | 25414025 | 25414179 | 5.899640811 | DTNB |
| 1 | 116526061 | 116526243 | -1.099693628 | CD58 |
| 2 | 26313905 | 26314102 | 1.386546049 | EPT1 |
| 2 | 26313905 | 26314102 | 1.386546049 | ADGRF3 |
| 2 | 27082819 | 27083051 | 2.167466087 | EMILIN1 |
| 1 | 116536053 | 116536252 | 2.229018961 | CD58 |
| 2 | 27357137 | 27357333 | 1.320077965 | AC074117.10 |
| 2 | 27358777 | 27359032 | 1.707494595 | AC074117.10 |
| 2 | 27381598 | 27381730 | 1.515917915 | PPM1G |
| 1 | 116564174 | 116564346 | -6.691447084 | CD58 |
| 2 | 27393006 | 27393198 | -6.380775616 | PPM1G |
| 2 | 27393006 | 27393198 | -6.380775616 | FTH1P3 |
| 2 | 27581328 | 27581522 | 1.687032859 | C2orf16 |
| 2 | 30606883 | 30607014 | 5.083709073 | LCLAT1 |

| | | | | |
|---|---|---|---|---|
| 11 | 5678303 | 5678501 | 1.06510424 | TRIM5 |
| 11 | 5679972 | 5680145 | 3.09607387 | TRIM5 |
| 11 | 6391713 | 6391907 | 1.395441903 | SMPD1 |
| 11 | 6557275 | 6557475 | 1.034404567 | DNHD1 |
| 11 | 6558220 | 6558469 | 1.656871268 | DNHD1 |
| 11 | 6563337 | 6563589 | 2.411495207 | DNHD1 |
| 11 | 6567119 | 6567228 | 1.387716604 | DNHD1 |
| 11 | 6600002 | 6600203 | -1.437661953 | RRP8 |
| 11 | 6601259 | 6601457 | -1.862419671 | RRP8 |
| 11 | 6682038 | 6682254 | -1.730006115 | MRPL17 |
| 11 | 7653344 | 7653590 | -1.347692617 | PPFIBP2 |
| 11 | 8000243 | 8000406 | -1.548317892 | EIF3F |
| 11 | 8730376 | 8730575 | 3.331508765 | ST5 |
| 11 | 10306789 | 10306983 | 2.79921238 | ADM |
| 11 | 10471243 | 10471443 | -3.877443751 | AMPD3 |
| 11 | 10505643 | 10505818 | 1.690164235 | AMPD3 |
| 11 | 10778795 | 10778991 | -2.296292269 | CTR9 |
| 1 | 23845560 | 23845758 | 1.731350007 | FUCA1 |
| 11 | 12259875 | 12260066 | 2.486571046 | MICAL2 |
| 11 | 14600252 | 14600481 | -1.215088525 | PSMA1 |
| 11 | 14606153 | 14606337 | -3.513812953 | PSMA1 |
| 11 | 14661161 | 14661364 | 1.345717736 | PDE3B |
| 11 | 14721453 | 14721646 | 1.068789416 | PDE3B |
| 11 | 14783322 | 14783511 | 1.085210344 | PDE3B |
| 1 | 24358286 | 24358493 | -2.123315975 | GRHL3 |
| 1 | 24358286 | 24358493 | -2.123315975 | STPG1 |
| 11 | 14883501 | 14883705 | 1.051904863 | CYP2R1 |
| 11 | 17168888 | 17169105 | -1.616501334 | PIK3C2A |
| 1 | 24901636 | 24901835 | 1.134583516 | RUNX3 |
| 11 | 17169563 | 17169745 | -1.019387154 | PIK3C2A |
| 11 | 17353070 | 17353264 | -4.817094365 | NCR3LG1 |
| 11 | 17353678 | 17353884 | -3.661367127 | NCR3LG1 |
| 11 | 17381147 | 17381362 | -3.433823238 | RP1-239B22.5 |
| 11 | 18702035 | 18702225 | 3.974924241 | TMEM86A |
| 11 | 22224571 | 22224727 | 1.899695001 | ANO5 |
| 11 | 22280056 | 22280191 | 1.650310068 | ANO5 |
| 11 | 22624793 | 22625018 | -1.926851768 | FANCF |
| 11 | 27367948 | 27368218 | -2.626303274 | LGR4 |
| 11 | 27675691 | 27675935 | 1.35172885 | BDNF |
| 11 | 27675691 | 27675935 | 1.35172885 | BDNF-AS |
| 11 | 28035457 | 28035604 | -3.682875933 | KIF18A |
| 11 | 28036300 | 28036513 | -2.91797451 | KIF18A |
| 11 | 31495066 | 31495261 | -2.108252393 | IMMP1L |
| 11 | 31746563 | 31746757 | -2.512767546 | Z83001.1 |
| 11 | 31746563 | 31746757 | -2.512767546 | ELP4 |
| 11 | 32438117 | 32438320 | -1.856214042 | WT1-AS |
| 1 | 1090795 | 1090974 | 7.844125155 | C1orf159 |
| 11 | 32438904 | 32439104 | -1.813391338 | WT1-AS |
| 11 | 32438904 | 32439104 | -1.813391338 | WT1-AS_6 |
| 11 | 32441056 | 32441233 | -1.398829524 | WT1-AS |
| 11 | 32453317 | 32453508 | 1.236537068 | WT1-AS |
| 11 | 32853739 | 32853940 | -2.927296505 | PRRG4 |
| 11 | 32931777 | 32931993 | -2.210031776 | QSER1 |
| 11 | 32933525 | 32933720 | -1.492351689 | QSER1 |
| 12 | 28275782 | 28276008 | -2.490950546 | CCDC91 |
| 12 | 29196603 | 29196878 | -1.398152952 | FAR2 |
| 12 | 29325620 | 29325755 | -9.668666129 | FAR2 |
| 12 | 29326554 | 29326762 | -9.224628741 | FAR2 |
| 12 | 29327848 | 29327980 | -5.81467831 | FAR2 |
| 12 | 29328437 | 29328637 | -7.238847517 | FAR2 |
| 12 | 29404181 | 29404323 | -1.198376931 | OVCH1-AS1 |
| 1 | 32826514 | 32826727 | -1.708974375 | S100PBP |
| 12 | 29406506 | 29406704 | -1.961312171 | OVCH1-AS1 |
| 12 | 29407291 | 29407446 | -3.316595743 | OVCH1-AS1 |
| 12 | 29410081 | 29410280 | -1.177187359 | OVCH1-AS1 |
| 12 | 29411736 | 29411882 | -1.284138584 | OVCH1-AS1 |
| 12 | 29441514 | 29441649 | -1.775871796 | OVCH1 |
| 12 | 29441514 | 29441649 | -1.775871796 | OVCH1-AS1 |
| 12 | 29442144 | 29442334 | -1.644760797 | OVCH1-AS1 |
| 12 | 29442144 | 29442334 | -1.644760797 | OVCH1 |
| 12 | 29449950 | 29450145 | -2.838243597 | OVCH1 |
| 12 | 29449950 | 29450145 | -2.838243597 | OVCH1-AS1 |
| 12 | 29465259 | 29465495 | -1.342618226 | OVCH1-AS1 |
| 12 | 29465259 | 29465495 | -1.342618226 | OVCH1 |
| 12 | 29584585 | 29584776 | 5.310568057 | TMTC1 |
| 12 | 29585423 | 29585613 | -1.253582612 | TMTC1 |
| 12 | 30630557 | 30630765 | -1.294849796 | IPO8 |
| 12 | 30710274 | 30710392 | -7.76492877 | CAPRIN2 |
| 12 | 30728779 | 30728978 | -2.406902359 | CAPRIN2 |
| 12 | 31387425 | 31387630 | -2.174948919 | DENND5B |
| 12 | 31447667 | 31447865 | -2.653157452 | DENND5B |
| 1 | 32936971 | 32937199 | 3.14605828 | RNF19B |
| 12 | 31452040 | 31452250 | -1.881761649 | DENND5B |
| 12 | 31479968 | 31480138 | -1.369267136 | DENND5B |
| 12 | 31550341 | 31550537 | -1.257602863 | DENND5B |
| 12 | 31551053 | 31551199 | -7.97635925 | DENND5B |
| 12 | 31982778 | 31982971 | -1.443136154 | KIAA1551 |
| 12 | 31983661 | 31983861 | -1.874520741 | KIAA1551 |
| 1 | 33008123 | 33008338 | 1.322836402 | AK2 |
| 1 | 33008123 | 33008338 | 1.322836402 | RP1-117O3.2 |
| 12 | 32107346 | 32107535 | -3.502495875 | RP11-843B15.2 |
| 12 | 32107346 | 32107535 | -3.502495875 | BICD1 |
| 12 | 32118881 | 32119080 | -2.914634849 | BICD1 |
| 12 | 32305830 | 32306030 | -2.710488898 | BICD1 |
| 12 | 32327517 | 32327720 | -2.470465388 | BICD1 |
| 12 | 32337431 | 32337716 | -2.819324931 | BICD1 |
| 12 | 32377653 | 32377852 | -3.403033113 | BICD1 |
| 12 | 32582066 | 32582325 | -3.426007888 | FGD4 |
| 12 | 32640444 | 32640652 | -3.827545623 | FGD4 |
| 1 | 33050779 | 33050997 | 2.986379006 | AK2 |
| 12 | 38320344 | 38320540 | -1.600312172 | ALG10B |
| 12 | 38906406 | 38906579 | 1.034372582 | RP11-396F22.1 |
| 12 | 38906406 | 38906579 | 1.034372582 | CPNE8 |
| 12 | 38909209 | 38909410 | 1.034419007 | RP11-396F22.1 |
| 12 | 39944614 | 39944832 | 1.367019122 | SLC2A13 |
| 12 | 42087740 | 42087998 | -1.040159582 | GXYLT1 |
| 1 | 33054328 | 33054509 | 1.460658453 | AK2 |
| 12 | 42225394 | 42225593 | -1.69742794 | YAF2 |
| 2 | 33234883 | 33235038 | -6.831556654 | LTBP1 |
| 2 | 33251072 | 33251255 | -2.608776932 | LTBP1 |
| 2 | 34594529 | 34594728 | -3.564641508 | AC073218.1 |
| 2 | 36578694 | 36578880 | -1.021131666 | FEZ2 |
| 2 | 38709822 | 38710089 | -5.899640811 | GALM |
| 2 | 38734383 | 38734586 | -1.2855716 | GALM |
| 1 | 117025579 | 117025778 | -1.661688799 | RP11-27K13.3 |
| 1 | 117025579 | 117025778 | -1.661688799 | CD101 |
| 2 | 39665209 | 39665408 | -1.212390752 | TMEM178A |
| 2 | 39665209 | 39665408 | -1.212390752 | AC007246.3 |
| 2 | 40177802 | 40178007 | 1.29220879 | SLC8A1-AS1 |
| 2 | 40177802 | 40178007 | 1.29220879 | SLC8A1 |
| 2 | 40235633 | 40235741 | -6.793478519 | SLC8A1-AS1 |
| 2 | 40235633 | 40235741 | -6.793478519 | SLC8A1 |
| 2 | 40428455 | 40428706 | 1.088418482 | SLC8A1 |
| 2 | 40430082 | 40430282 | 1.201245323 | SLC8A1 |
| 2 | 42168861 | 42169061 | 3.723422359 | AC083949.1 |
| 2 | 43675708 | 43675840 | -2.775879424 | C1GALT1C1L |
| 2 | 43675708 | 43675840 | -2.775879424 | PLEKHH2 |
| 2 | 44595708 | 44595866 | 1.261780978 | CAMKMT |
| 2 | 46184853 | 46185006 | -1.088266483 | PRKCE |
| 2 | 46526574 | 46526761 | -1.536879519 | ATP6V1E2 |
| 1 | 117402103 | 117402269 | 1.714922411 | MAN1A2 |
| 2 | 46603662 | 46603833 | -1.930872313 | PIGF |
| 2 | 53853882 | 53854006 | 6.23393012 | GPR75 |
| 2 | 53853882 | 53854006 | 6.23393012 | GPR75-ASB3 |
| 2 | 54631260 | 54631446 | -1.135435736 | SPTBN1 |
| 2 | 54649695 | 54649895 | -1.021038343 | SPTBN1 |
| 2 | 55050226 | 55050428 | 8.121679981 | RTN4 |
| 2 | 56173076 | 56173250 | 2.354229583 | RP11-481J13.1 |
| 2 | 56173076 | 56173250 | 2.354229583 | RP11-482H16.1 |
| 2 | 56193091 | 56193291 | 1.979963946 | RP11-482H16.1 |
| 2 | 56193091 | 56193291 | 1.979963946 | CCDC85A |
| 2 | 58173748 | 58173947 | -3.357122759 | FANCL |
| 2 | 60461832 | 60462034 | -3.074232815 | BCL11A |
| 2 | 60922567 | 60922743 | -1.509125852 | REL |
| 2 | 61040304 | 61040496 | -1.623271893 | PEX13 |
| 2 | 61224267 | 61224470 | -1.31601367 | USP34 |
| 2 | 61306063 | 61306266 | 6.540242229 | USP34 |
| 2 | 62146315 | 62146501 | 1.299624886 | RPSAP26 |
| 2 | 62146315 | 62146501 | 1.299624886 | COMMD1 |
| 2 | 62146315 | 62146501 | 1.299624886 | AC018462.2 |
| 2 | 63307698 | 63307897 | -5.312854422 | WDPCP |
| 2 | 63588439 | 63588643 | -6.922990007 | WDPCP |
| 2 | 63588439 | 63588643 | -6.922990007 | MDH1 |
| 2 | 63962119 | 63962321 | -1.345299192 | VPS54 |
| 2 | 64524426 | 64524627 | 2.598600336 | AFTPH |
| 2 | 65313775 | 65313978 | 1.379733487 | SPRED2 |
| 2 | 65500930 | 65501155 | 4.528168326 | AC007389.1 |
| 2 | 65500930 | 65501155 | 4.528168326 | AC074391.1 |
| 2 | 68130829 | 68131026 | 2.068560799 | WDR92 |
| 2 | 68395744 | 68395940 | 7.150355022 | PLEK |
| 2 | 68464979 | 68465180 | -1.892796411 | FBXO48 |
| 2 | 68826212 | 68826413 | 1.046577531 | ARHGAP25 |

| | | | | | | | | | | | | | | |
|---|---|---|---|---|---|---|---|---|---|---|---|---|---|---|
| 11 | 32935023 | 32935217 | -1.691794908 | QSER1 | 12 | 42427645 | 42427891 | -1.006890108 | PPHLN1 | 2 | 69741319 | 69741493 | 1.270590404 | ANXA4 |
| 11 | 32953908 | 32954120 | -1.790101325 | QSER1 | 12 | 43730572 | 43730754 | -2.322587311 | PUS7L | 2 | 69829852 | 69830050 | 9.032957849 | GMCL1 |
| 11 | 33237091 | 33237240 | -2.135523463 | RP1-89D4.1 | 12 | 43953341 | 43953541 | -1.262000062 | TMEM117 | 2 | 69937997 | 69938204 | -1.062872242 | MXD1 |
| 11 | 33286523 | 33286723 | -1.257409865 | HIPK3 | 12 | 44388016 | 44388215 | -1.823445125 | TMEM117 | 2 | 70927800 | 70928056 | 1.918470859 | VAX2 |
| 11 | 35200915 | 35201106 | -7.302768696 | CD44 | 12 | 45823090 | 45823222 | -2.732233604 | ARID2 | 2 | 70933053 | 70933252 | 1.531636065 | VAX2 |
| 11 | 35204584 | 35204784 | -5.417235525 | CD44 | 12 | 45922076 | 45922242 | -1.238200913 | SCAF11 | 1 | 119212400 | 119212677 | -1.38915963 | RP11-418J17.1 |
| 11 | 35229128 | 35229326 | 2.834290264 | CD44 | 1 | 33299423 | 33299636 | -2.799950115 | ZNF362 | 2 | 73462594 | 73462751 | -6.148027919 | ALMS1 |
| 11 | 36462446 | 36462653 | 1.404249732 | PRR5L | 12 | 46190112 | 46190308 | -2.72275994 | SLC38A1 | 2 | 74418205 | 74418357 | 1.477133041 | HMGA1P8 |
| 11 | 40792556 | 40792763 | 1.200439718 | LRRC4C | 12 | 46190962 | 46191149 | -3.618116491 | SLC38A1 | 2 | 74418205 | 74418357 | 1.477133041 | C2orf81 |
| 11 | 43339437 | 43339633 | -1.742745745 | RP11-484D2.2 | 12 | 46359142 | 46359322 | -1.563184747 | SLC38A2 | 2 | 74462615 | 74462810 | 1.174474969 | MOGS |
| 11 | 43339437 | 43339633 | -1.742745745 | API5 | 12 | 46399133 | 46399329 | -1.479534868 | RP11-96H19.1 | 2 | 74474260 | 74474468 | 1.070793716 | CCDC142 |
| 11 | 43898346 | 43898639 | -1.469084735 | ALKBH3 | 12 | 46399887 | 46400061 | -1.038937445 | RP11-96H19.1 | 2 | 74503135 | 74503386 | 2.33076523 | LBX2 |
| 11 | 43898346 | 43898639 | -1.469084735 | SEC14L1P1 | 12 | 46605153 | 46605336 | -1.241196384 | RP11-96H19.1 | 2 | 74503135 | 74503386 | 2.33076523 | LBX2-AS1 |
| 11 | 44244335 | 44244481 | -1.30784516 | EXT2 | 12 | 46605153 | 46605336 | -1.241196384 | RP11-446N19.1 | 2 | 74530109 | 74530304 | 1.143316163 | HTRA2 |
| 11 | 44264122 | 44264244 | -2.815729229 | ALX4 | 1 | 33324717 | 33324938 | -1.175556248 | PHC2 | 2 | 74832838 | 74833038 | -5.612000837 | RP11-259N19.1 |
| 11 | 45804454 | 45804647 | -5.498545625 | SLC35C1 | 1 | 33324717 | 33324938 | -1.175556248 | RP11-415J8.3 | 2 | 74833680 | 74833881 | 1.20329663 | RP11-259N19.1 |
| 11 | 45805620 | 45805817 | -1.001839539 | SLC35C1 | 12 | 46637207 | 46637405 | -5.332667959 | RP11-96H19.1 | 2 | 74834148 | 74834335 | 8.63189134 | HK2 |
| 11 | 45918053 | 45918246 | -3.630106943 | PEX16 | 12 | 46637207 | 46637405 | -5.332667959 | RP11-446N19.1 | 2 | 84870274 | 84870473 | 2.813766942 | TRABD2A |
| 11 | 46238413 | 46238575 | -2.398633052 | CTD-2589M5.5 | 12 | 47078680 | 47078913 | 1.349715537 | AMIGO2 | 2 | 86195752 | 86195912 | 3.543731421 | RP11-301O19.1 |
| 11 | 46603468 | 46603668 | -1.470349126 | HARBI1 | 12 | 47236250 | 47236467 | -1.256601227 | PCED1B | 2 | 86195752 | 86195912 | 3.543731421 | IMMT |
| 11 | 46615605 | 46615838 | -1.718491695 | HARBI1 | 1 | 33479211 | 33479443 | -1.17807918 | ZSCAN20 | 2 | 86212533 | 86212737 | 1.12266524 | MRPL35 |
| 11 | 46743975 | 46744179 | -2.572992338 | CKAP5 | 12 | 47738279 | 47738473 | 1.895198207 | RP1-197B17.3 | 1 | 119916460 | 119916657 | -2.449575579 | NOTCH2 |
| 11 | 47261563 | 47261762 | 1.609972031 | NR1H3 | 12 | 47738279 | 47738473 | 1.895198207 | RAPGEF3 | 2 | 86774695 | 86774903 | -1.012825454 | RMND5A |
| 11 | 47268624 | 47268806 | 1.037057666 | NR1H3 | 12 | 47783419 | 47783650 | -1.535581658 | HDAC7 | 2 | 87583420 | 87583548 | 2.05593986 | LINC00152 |
| 11 | 47355361 | 47355569 | -10.74026889 | SPI1 | 12 | 48768385 | 48768578 | -1.272343856 | ADCY6 | 2 | 87584752 | 87585031 | 2.111724864 | LINC00152 |
| 11 | 47723090 | 47723285 | 1.029532459 | FNBP4 | 12 | 48827666 | 48827858 | -2.82061536 | CACNB3 | 2 | 87782864 | 87783063 | -1.113143767 | RGPD2 |
| 11 | 47778948 | 47779167 | -1.738577731 | NUP160 | 12 | 48830033 | 48830233 | 1.050552903 | DDX23 | 1 | 120433844 | 120434027 | -6.627514285 | PFN1P2 |
| 11 | 57484943 | 57485154 | -1.581507824 | SLC43A1 | 12 | 48981544 | 48981715 | -4.96831897 | WNT1 | 2 | 91628528 | 91628684 | 1.151807108 | AC027612.6 |
| 11 | 57699057 | 57699290 | -1.400740825 | ZDHHC5 | 1 | 33494846 | 33494988 | -2.311053358 | ZSCAN20 | 2 | 95182159 | 95182373 | 1.459584463 | ZNF2 |
| 1 | 25783782 | 25783980 | 1.429427208 | MAN1C1 | 12 | 49330219 | 49330413 | -2.425279472 | TROAP | 1 | 120454378 | 120454537 | -3.142701109 | NBPF8 |
| 11 | 59135195 | 59135427 | -1.528901434 | AP001258.4 | 12 | 49351774 | 49351966 | -3.523423057 | DNAJC22 | 2 | 95838958 | 95839128 | -1.463702435 | ANKRD36C |
| 11 | 59152719 | 59152918 | -1.58149796 | FAM111A | 12 | 49625606 | 49625945 | 3.058459269 | PRPF40B | 2 | 95933196 | 95933391 | 6.164898169 | ANKRD36C |
| 11 | 59204788 | 59204991 | 1.591585776 | DTX4 | 12 | 49791719 | 49791904 | -3.156441478 | NCKAP5L | 2 | 96208514 | 96208692 | 8.810539431 | STARD7-AS1 |
| 11 | 59806411 | 59806611 | -1.615469467 | MRPL16 | 12 | 49795690 | 49795884 | -2.808486448 | NCKAP5L | 2 | 96208514 | 96208692 | 8.810539431 | STARD7 |
| 11 | 60172654 | 60172853 | -10.20242837 | MS4A6A | 12 | 50925124 | 50925280 | -1.744931634 | METTL7A | 2 | 96240218 | 96240387 | -2.227122586 | STARD7-AS1 |
| 11 | 60365189 | 60365389 | -7.232336686 | MS4A6E | 1 | 34976357 | 34976548 | -1.702088649 | RP11-244H3.1 | 1 | 6013810 | 6013969 | -4.281121212 | KCNAB2 |
| 11 | 60937011 | 60937146 | 4.794357318 | TMEM132A | 12 | 51245722 | 51245922 | -4.474745717 | DAZAP2 | 2 | 96824939 | 96825107 | -1.031778863 | ANKRD23 |
| 11 | 60937011 | 60937146 | 4.794357318 | SLC15A3 | 12 | 51245722 | 51245922 | -4.474745717 | SMAGP | 2 | 96824939 | 96825107 | -1.031778863 | CNNM3 |
| 11 | 61004604 | 61004884 | 8.43456797 | CD6 | 12 | 51352021 | 51352251 | -1.261254841 | GALNT6 | 2 | 97245384 | 97245558 | -1.143020375 | ANKRD36 |
| 11 | 61258488 | 61258686 | 2.777678019 | VWCE | 12 | 51354106 | 51354350 | -1.305510868 | GALNT6 | 2 | 97663317 | 97663518 | 8.18223834 | ACTR1B |
| 1 | 1093104 | 1093319 | -2.193625375 | C1orf159 | 1 | 34985387 | 34985547 | -1.300820621 | RP11-244H3.4 | 2 | 98582789 | 98582954 | -7.594506871 | INPP4A |
| 1 | 25832235 | 25832434 | -1.057441819 | MTFR1L | 12 | 53181700 | 53181834 | -1.457529415 | ZNF740 | 1 | 120802326 | 120802469 | 1.574049319 | NBPF26 |
| 1 | 25832235 | 25832434 | -1.057441819 | AUNIP | 12 | 53253396 | 53253598 | -1.079344022 | MFSD5 | 2 | 99125117 | 99125298 | -4.830889049 | TSGA10 |
| 11 | 61636673 | 61637030 | 8.400399405 | RPLP0P2 | 12 | 53411142 | 53411346 | -1.93114731 | SP1 | 2 | 99363525 | 99363721 | 5.083709073 | EIF5B |
| 11 | 61744013 | 61744219 | 1.353629544 | DAGLA | 12 | 53500889 | 53501068 | 1.872977189 | RP11-793H13.11 | 2 | 99384594 | 99384717 | 1.368843381 | EIF5B |
| 11 | 61786454 | 61786761 | 4.328458166 | TMEM258 | 12 | 53500889 | 53501068 | 1.872977189 | TARBP2 | 1 | 120840403 | 120840569 | -6.68758761 | NBPF26 |
| 11 | 61786454 | 61786761 | 4.328458166 | MYRF | 12 | 54362565 | 54362768 | 1.273829132 | RP11-753H16.5 | 2 | 99551307 | 99551490 | -1.533700783 | AFF3 |
| 11 | 61795849 | 61796166 | -2.980627998 | FADS2 | 12 | 54362565 | 54362768 | 1.273829132 | GPR84 | 2 | 101152477 | 101152649 | 1.092977912 | RP11-299H21.1 |
| 11 | 61795849 | 61796166 | -2.980627998 | FEN1 | 12 | 54362565 | 54362768 | 1.273829132 | RP11-753H16.3 | 2 | 101152477 | 101152649 | 1.092977912 | TBC1D8 |
| 11 | 61796842 | 61797034 | -4.28186943 | FADS2 | 12 | 54363011 | 54363227 | 1.156615831 | RP11-753H16.3 | 2 | 101381765 | 101381962 | 4.547152829 | CREG2 |
| 11 | 61796842 | 61797034 | -4.28186943 | FEN1 | 12 | 54363011 | 54363227 | 1.156615831 | GPR84 | 2 | 101750082 | 101750271 | -7.748863559 | MAP4K4 |
| 11 | 61962573 | 61962790 | 1.438745182 | BEST1 | 12 | 54363011 | 54363227 | 1.156615831 | RP11-753H16.5 | 1 | 120954377 | 120954578 | -1.347720203 | LINC00623 |
| 11 | 61962573 | 61962790 | 1.438745182 | FTH1 | 12 | 54395668 | 54395874 | -1.16678222 | ITGA5 | 2 | 102718749 | 102719065 | -1.210709723 | MFSD9 |
| 1 | 25835197 | 25835404 | -2.74418005 | AUNIP | 12 | 54395668 | 54395874 | -1.16678222 | RP11-753H16.3 | 2 | 105892979 | 105893102 | -1.324631229 | NCK2 |

| | | | | |
|---|---|---|---|---|
| 11 | 62519325 | 62519516 | -1.482262537 | AHNAK |
| 11 | 62521963 | 62522232 | -1.217149994 | AHNAK |
| 11 | 62523193 | 62523429 | -1.369461476 | AHNAK |
| 11 | 62525534 | 62525736 | -1.245485255 | AHNAK |
| 11 | 62526342 | 62526555 | -1.086690684 | AHNAK |
| 11 | 62529043 | 62529243 | -1.060167941 | AHNAK |
| 1 | 25862843 | 25862956 | -2.671384219 | PAQR7 |
| 1 | 25862843 | 25862956 | -2.671384219 | RP1-125I3.2 |
| 11 | 62531514 | 62531639 | -1.521081773 | AHNAK |
| 11 | 62531933 | 62532134 | -1.245773014 | AHNAK |
| 11 | 62602205 | 62602489 | -2.166575545 | EML3 |
| 11 | 62647848 | 62648060 | -1.907389004 | INTS5 |
| 11 | 62648553 | 62648757 | -1.311998874 | INTS5 |
| 1 | 25863539 | 25863697 | -1.586702785 | PAQR7 |
| 11 | 62649444 | 62649643 | -1.52974274 | INTS5 |
| 11 | 62672055 | 62672285 | -2.619337539 | LBHD1 |
| 11 | 62672055 | 62672285 | -2.619337539 | UQCC3 |
| 11 | 62752175 | 62752394 | -1.487081367 | ZBTB3 |
| 11 | 62786416 | 62786606 | -3.023636876 | TMEM223 |
| 11 | 62786416 | 62786606 | -3.023636876 | RP11-727F15.12 |
| 11 | 62786416 | 62786606 | -3.023636876 | TAF6L |
| 11 | 62787223 | 62787407 | -1.476290803 | TMEM223 |
| 11 | 62787223 | 62787407 | -1.476290803 | TAF6L |
| 11 | 62790434 | 62790667 | -1.740277106 | TMEM223 |
| 11 | 63719306 | 63719494 | -8.445891852 | RTN3 |
| 11 | 63720567 | 63720809 | -4.212346335 | RTN3 |
| 11 | 63954844 | 63955044 | -1.568658901 | NAA40 |
| 11 | 64115712 | 64115912 | -2.727136602 | FLRT1 |
| 11 | 64115712 | 64115912 | -2.727136602 | MACROD1 |
| 11 | 64246978 | 64247177 | 1.269547746 | RP11-783K16.13 |
| 11 | 64246978 | 64247177 | 1.269547746 | RP11-783K16.5 |
| 11 | 64344461 | 64344661 | -1.98763392 | CCDC88B |
| 11 | 64349797 | 64349984 | -2.382323866 | CCDC88B |
| 11 | 64450010 | 64450156 | -2.960210785 | AP003774.4 |
| 11 | 64765735 | 64765956 | -2.250699056 | SF1 |
| 11 | 64804252 | 64804612 | -1.832802694 | MEN1 |
| 11 | 64854212 | 64854379 | -2.029128276 | EHD1 |
| 11 | 64854538 | 64854690 | -1.581667945 | EHD1 |
| 11 | 64925523 | 64925731 | 1.384650788 | PPP2R5B |
| 11 | 65040929 | 65041141 | -2.148651777 | SAC3D1 |
| 11 | 65044432 | 65044643 | -2.377778632 | SAC3D1 |
| 1 | 26170595 | 26170789 | -1.481146347 | RP11-96L14.7 |
| 1 | 26170595 | 26170789 | -1.481146347 | ZNF593 |
| 11 | 65079454 | 65079654 | -3.253126238 | CDCA5 |
| 11 | 65116584 | 65116704 | -1.62493676 | ZNHIT2 |
| 11 | 65424373 | 65424612 | -11.56983633 | NEAT1 |
| 11 | 65455896 | 65456091 | 2.036375918 | CMB9-22P13.1 |
| 11 | 65466019 | 65466194 | 2.490064143 | CMB9-22P13.1 |
| 11 | 65499486 | 65499627 | 13.67087184 | MALAT1 |
| 11 | 65500360 | 65500579 | 2.743422606 | MALAT1 |
| 11 | 65581669 | 65581870 | -1.543112788 | EHBP1L1 |
| 11 | 65618432 | 65618607 | -1.034429635 | PCNX3 |
| 11 | 65654320 | 65654536 | -1.154609887 | RELA |
| 11 | 65726969 | 65727173 | 3.501349454 | KRT8P26 |
| 12 | 54395668 | 54395874 | -1.16678222 | RP11-753H16.5 |
| 12 | 55684011 | 55684205 | -8.523125044 | METTL7B |
| 12 | 55685376 | 55685566 | -1.715884747 | ITGA7 |
| 12 | 55690021 | 55690153 | -1.094665831 | ITGA7 |
| 1 | 35113745 | 35113965 | -1.528349084 | ZMYM1 |
| 12 | 55730482 | 55730669 | -1.038943803 | RP11-644F5.11 |
| 12 | 55749550 | 55749749 | -1.267002718 | GDF11 |
| 12 | 55751365 | 55751568 | -1.655687756 | GDF11 |
| 12 | 55752758 | 55753115 | -1.582529623 | GDF11 |
| 12 | 55752758 | 55753115 | -1.582529623 | SARNP |
| 12 | 55830187 | 55830401 | -1.527069824 | TMEM198B |
| 12 | 55830187 | 55830401 | -1.527069824 | DNAJC14 |
| 12 | 55901891 | 55902107 | -1.537834333 | PYM1 |
| 12 | 56004602 | 56004802 | -1.156202278 | SUOX |
| 1 | 35114985 | 35115271 | -2.181174751 | ZMYM1 |
| 12 | 56206416 | 56206630 | -1.579896235 | RNF41 |
| 12 | 56417571 | 56417715 | -2.667107234 | TIMELESS |
| 12 | 56511178 | 56511404 | -1.398575098 | RP11-153M3.1 |
| 12 | 57224318 | 57224462 | -7.120954345 | NXPH4 |
| 1 | 35381511 | 35381684 | -1.719966802 | ZMYM4 |
| 12 | 57626260 | 57626443 | 1.836526915 | B4GALNT1 |
| 12 | 57823398 | 57823598 | -1.516680745 | CTDSP2 |
| 12 | 58872591 | 58872789 | 1.862013269 | LRIG3 |
| 12 | 59779369 | 59779525 | -1.834255146 | SLC16A7 |
| 12 | 62191485 | 62191684 | 1.076475961 | FAM19A2 |
| 12 | 62592840 | 62593022 | -1.205218609 | MON2 |
| 12 | 62648366 | 62648569 | -2.176068597 | PPM1H |
| 12 | 64142461 | 64142645 | -1.816364833 | SRGAP1 |
| 12 | 64143226 | 64143329 | -1.630960393 | SRGAP1 |
| 12 | 64153130 | 64153320 | -3.239890132 | SRGAP1 |
| 12 | 64193964 | 64194171 | -1.051873962 | C12orf66 |
| 12 | 64451718 | 64451897 | 5.256863798 | RP11-114F3.4 |
| 12 | 64451718 | 64451897 | 5.256863798 | TBK1 |
| 12 | 64533415 | 64533591 | -6.274651684 | RASSF3 |
| 12 | 64533415 | 64533591 | -6.274651684 | RP11-439H13.2 |
| 12 | 65464216 | 65464415 | -4.660964047 | MSRB3 |
| 1 | 35577386 | 35577575 | -2.578680152 | TFAP2E |
| 1 | 35577386 | 35577575 | -2.578680152 | RP4-728D4.2 |
| 12 | 65613544 | 65613722 | -2.812559814 | RP11-230G5.2 |
| 12 | 65852936 | 65853130 | -1.875859702 | HMGA2 |
| 12 | 65852936 | 65853130 | -1.875859702 | RP11-366L20.2 |
| 12 | 65920911 | 65921109 | -2.038757585 | HMGA2 |
| 12 | 66165544 | 66165736 | 2.013031148 | RP11-745O10.4 |
| 12 | 66165544 | 66165736 | 2.013031148 | TMBIM4 |
| 12 | 66234632 | 66234770 | -6.010837021 | IRAK3 |
| 12 | 66234632 | 66234770 | -6.010837021 | RBMS1P1 |
| 12 | 66247765 | 66247969 | -1.370366714 | IRAK3 |
| 12 | 66310158 | 66310344 | 4.174558008 | HELB |
| 12 | 66331307 | 66331516 | 2.648562352 | HELB |
| 12 | 66338066 | 66338271 | 3.708632527 | HELB |
| 1 | 35594547 | 35594747 | -5.61860498 | TFAP2E |
| 12 | 67657141 | 67657365 | -1.817119872 | DYRK2 |
| 12 | 67658226 | 67658469 | -1.614376411 | DYRK2 |
| 12 | 68764739 | 68764939 | -1.274970668 | SLC35E3 |
| 2 | 108468791 | 108468897 | 7.133320078 | GCC2 |
| 2 | 108471034 | 108471234 | -1.187834114 | GCC2 |
| 2 | 108648809 | 108649000 | 8.832931642 | LIMS1 |
| 2 | 108649461 | 108649693 | 8.178148605 | LIMS1 |
| 1 | 121096108 | 121096309 | 1.238615531 | RP11-439A17.9 |
| 1 | 121096108 | 121096309 | 1.238615531 | FCGR1B |
| 1 | 121096108 | 121096309 | 1.238615531 | RP11-439A17.10 |
| 2 | 111119308 | 111119504 | -1.046554702 | BCL2L11 |
| 2 | 111123750 | 111123872 | -1.786813133 | BCL2L11 |
| 2 | 111164212 | 111164358 | -1.604900115 | BCL2L11 |
| 2 | 111223184 | 111223416 | 1.485296535 | MIR4435-2HG |
| 2 | 111224863 | 111225063 | 1.245737689 | MIR4435-2HG |
| 2 | 111246751 | 111246945 | 1.808316695 | MIR4435-2HG |
| 2 | 111292416 | 111292612 | 1.420268975 | MIR4435-2HG |
| 2 | 111367294 | 111367593 | 1.092109019 | MIR4435-2HG |
| 2 | 112028622 | 112028891 | -1.244984517 | MERTK |
| 1 | 6016571 | 6016765 | -2.219336529 | KCNAB2 |
| 1 | 121184041 | 121184252 | 1.933875712 | FAM72B |
| 2 | 112575148 | 112575350 | 1.101105044 | POLR1B |
| 2 | 112752381 | 112752586 | -1.10584837 | CKAP2L |
| 1 | 121184701 | 121184871 | 1.252793934 | SRGAP2C |
| 1 | 121184701 | 121184871 | 1.252793934 | FAM72B |
| 2 | 113113188 | 113113370 | 2.696158711 | IL1RN |
| 2 | 113132873 | 113133073 | 12.98251476 | IL1RN |
| 2 | 113598740 | 113598938 | -1.276458686 | WASH2P |
| 2 | 119263342 | 119263537 | -1.719185002 | STEAP3 |
| 2 | 126696079 | 126696278 | -1.128700423 | GYPC |
| 2 | 127428715 | 127428919 | -2.944671916 | PROC |
| 1 | 121406990 | 121407172 | -1.909713877 | CH17-437K3.1 |
| 1 | 143498486 | 143498650 | 6.243668808 | CH17-13I23.1 |
| 2 | 128268031 | 128268245 | 1.020630759 | HS6ST1 |
| 2 | 134333117 | 134333277 | 6.346525648 | MGAT5 |
| 2 | 134448817 | 134449019 | 1.3111045 | MGAT5 |
| 2 | 135202854 | 135203042 | -5.5 | ZRANB3 |
| 2 | 135774850 | 135775045 | -1.094006237 | UBXN4 |
| 2 | 138569007 | 138569192 | -1.618268302 | SPOPL |
| 2 | 138569007 | 138569192 | -1.618268302 | AC092620.2 |
| 2 | 143006244 | 143006432 | 6.689147428 | KYNU |
| 2 | 143008097 | 143008300 | -1.232818824 | KYNU |
| 2 | 143304138 | 143304337 | -6.777534196 | ARHGAP15 |
| 2 | 143304138 | 143304337 | -6.777534196 | AC096558.1 |
| 2 | 143304138 | 143304337 | -6.777534196 | RP11-570L15.2 |
| 2 | 143663607 | 143663807 | -6.792950725 | ARHGAP15 |
| 2 | 148024955 | 148025117 | 5.958872507 | MBD5 |
| 2 | 150469961 | 150470214 | -1.694117132 | RND3 |
| 2 | 151251351 | 151251509 | -1.643049722 | RBM43 |
| 2 | 151889823 | 151890012 | 7.139877828 | CTD-2021K4.1 |
| 2 | 151889823 | 151890012 | 7.139877828 | CACNB4 |
| 1 | 144423049 | 144423246 | -1.897284305 | NBPF15 |
| 2 | 152390176 | 152390325 | 5.577409055 | AC012066.1 |
| 2 | 152390176 | 152390325 | 5.577409055 | FMNL2 |
| 2 | 156325641 | 156325920 | 1.979300114 | NR4A2 |
| 2 | 156329793 | 156329993 | 2.200453325 | NR4A2 |
| 2 | 157321304 | 157321412 | -2.762346019 | ERMN |

| | | | | |
|---|---|---|---|---|
| 11 | 65751399 | 65751591 | 1.069965131 | RP11-770G2.2 |
| 11 | 65778775 | 65778962 | 1.620877325 | AP5B1 |
| 11 | 65787204 | 65787411 | 5.233006357 | OVOL1 |
| 11 | 65794801 | 65794918 | 4.518086806 | OVOL1 |
| 11 | 65891381 | 65891593 | -2.639045786 | CCDC85B |
| 11 | 65899626 | 65899794 | -2.953445298 | FOSL1 |
| 11 | 65998553 | 65998742 | -2.437906079 | EIF1AD |
| 11 | 66002120 | 66002304 | -7.968226955 | BANF1 |
| 11 | 66002120 | 66002304 | -7.968226955 | EIF1AD |
| 1 | 1204188 | 1204369 | 3.623963757 | TNFRSF18 |
| 11 | 66624221 | 66624420 | -2.105226181 | RBM14 |
| 11 | 66624221 | 66624420 | -2.105226181 | RBM14-RBM4 |
| 11 | 66643365 | 66643696 | -1.432415325 | RBM4 |
| 11 | 66643365 | 66643696 | -1.432415325 | RBM14-RBM4 |
| 11 | 66668709 | 66668901 | -1.059147953 | RBM4B |
| 11 | 66669092 | 66669287 | -2.192629715 | RBM4B |
| 11 | 66700706 | 66700887 | -5.549345048 | SPTBN2 |
| 11 | 66779004 | 66779233 | -1.887002914 | C11orf80 |
| 1 | 26692273 | 26692461 | -2.267553147 | RP5-968P14.2 |
| 11 | 66779701 | 66779895 | -1.840543801 | C11orf80 |
| 11 | 66848962 | 66849161 | 2.52462176 | PC |
| 11 | 66859520 | 66859725 | -1.3370029 | PC |
| 11 | 66859520 | 66859725 | -1.3370029 | LRFN4 |
| 1 | 26693011 | 26693208 | -1.363731559 | RP5-968P14.2 |
| 11 | 67290612 | 67290792 | -1.684442997 | ANKRD13D |
| 11 | 67333036 | 67333201 | -2.367746352 | RAD9A |
| 11 | 67365189 | 67365345 | 3.49547743 | CLCF1 |
| 11 | 67365189 | 67365345 | 3.49547743 | RAD9A |
| 11 | 67365189 | 67365345 | 3.49547743 | AP003419.11 |
| 11 | 67365415 | 67365539 | 4.595003894 | AP003419.11 |
| 11 | 67365415 | 67365539 | 4.595003894 | CLCF1 |
| 11 | 67365415 | 67365539 | 4.595003894 | RAD9A |
| 11 | 67397557 | 67397738 | -1.611160832 | RAD9A |
| 11 | 67398020 | 67398168 | -2.138535912 | RAD9A |
| 11 | 67409712 | 67409925 | 1.945167287 | PPP1CA |
| 11 | 67409712 | 67409925 | 1.945167287 | TBC1D10C |
| 11 | 67420483 | 67420677 | 6.417136758 | PPP1CA |
| 11 | 67420483 | 67420677 | 6.417136758 | CARNS1 |
| 11 | 67423803 | 67424011 | 10.61905628 | CARNS1 |
| 11 | 67424600 | 67424794 | 7.746927725 | CARNS1 |
| 11 | 67425120 | 67425265 | 5.524427511 | CARNS1 |
| 11 | 68052150 | 68052347 | -9.67985548 | RP11-802E16.3 |
| 11 | 68386414 | 68386530 | -3.45101242 | LRP5 |
| 11 | 68737163 | 68737345 | -4.142171079 | TESMIN |
| 11 | 69161933 | 69162048 | 5.72869044 | RP11-554A11.8 |
| 11 | 69161933 | 69162048 | 5.72869044 | TPCN2 |
| 11 | 69295848 | 69295984 | 6.741401449 | MYEOV |
| 11 | 71476379 | 71476595 | -2.247426221 | NADSYN1 |
| 11 | 71796666 | 71796859 | -1.150445384 | ALG1L9P |
| 11 | 71796666 | 71796859 | -1.150445384 | FAM86C1 |
| 11 | 71800330 | 71800529 | -1.477794705 | FAM86C1 |
| 11 | 71800330 | 71800529 | -1.477794705 | ALG1L9P |
| 11 | 71903382 | 71903524 | -3.203450855 | RP11-849H4.2 |
| 11 | 71903382 | 71903524 | -3.203450855 | OR7E126P |
| 12 | 68957447 | 68957637 | -1.197429809 | CPM |
| 12 | 68957447 | 68957637 | -1.197429809 | PRELID2P1 |
| 12 | 69284865 | 69285064 | -6.861937109 | C1GALT1P1 |
| 12 | 70182691 | 70182833 | -1.814906621 | LINC01481 |
| 12 | 70182691 | 70182833 | -1.814906621 | RP11-320P7.2 |
| 12 | 70183386 | 70183534 | -2.162160604 | RP11-320P7.2 |
| 12 | 70183386 | 70183534 | -2.162160604 | LINC01481 |
| 12 | 70235412 | 70235607 | -1.656225963 | LINC01481 |
| 12 | 70556387 | 70556579 | -2.31679331 | PTPRB |
| 12 | 70571188 | 70571369 | -5.826422487 | PTPRB |
| 12 | 70571188 | 70571369 | -5.826422487 | RP11-785D18.3 |
| 12 | 71663121 | 71663308 | 1.911550604 | ZFC3H1 |
| 12 | 71663121 | 71663308 | 1.911550604 | THAP2 |
| 12 | 71676729 | 71676839 | -1.339331043 | RP11-293I14.2 |
| 12 | 71676729 | 71676839 | -1.339331043 | THAP2 |
| 12 | 74539378 | 74539577 | -1.08464971 | ATXN7L3B |
| 1 | 35753831 | 35754031 | -1.795701858 | CLSPN |
| 12 | 76029934 | 76030143 | -1.127589629 | PHLDA1 |
| 12 | 76347542 | 76347812 | -1.29013566 | BBS10 |
| 1 | 35760438 | 35760639 | -1.295840517 | CLSPN |
| 12 | 82353209 | 82353409 | -1.304915687 | CCDC59 |
| 12 | 82423336 | 82423510 | -5.46831897 | METTL25 |
| 12 | 82857279 | 82857477 | -1.97932145 | TMTC2 |
| 12 | 85283570 | 85283734 | -3.271772687 | ALX1 |
| 1 | 35764360 | 35764525 | -1.245908102 | CLSPN |
| 12 | 89522802 | 89523037 | 1.28635889 | POC1B |
| 12 | 89522802 | 89523037 | 1.28635889 | POC1B-GALNT4 |
| 12 | 89522802 | 89523037 | 1.28635889 | GALNT4 |
| 1 | 35853720 | 35853918 | -1.408441042 | AGO4 |
| 12 | 89525562 | 89525742 | 1.840053209 | RP11-734K2.4 |
| 12 | 89525562 | 89525742 | 1.840053209 | POC1B |
| 12 | 89525562 | 89525742 | 1.840053209 | POC1B-GALNT4 |
| 12 | 89542108 | 89542334 | 2.205888868 | RP11-734K2.4 |
| 12 | 93398628 | 93398830 | -1.477946162 | NUDT4 |
| 12 | 94288575 | 94288745 | -2.045580636 | PLXNC1 |
| 1 | 35855946 | 35856141 | -2.671314106 | AGO4 |
| 12 | 98547891 | 98548112 | -2.30628224 | TMPO |
| 12 | 98613777 | 98613985 | -2.291962015 | IKBIP |
| 12 | 100059074 | 100059274 | -1.402320813 | UHRF1BP1L |
| 12 | 100586445 | 100586638 | -2.55706245 | GAS2L3 |
| 12 | 100624623 | 100624828 | -2.397317989 | GAS2L3 |
| 12 | 101921990 | 101922177 | 4.418034686 | DRAM1 |
| 1 | 36090215 | 36090382 | 1.501561268 | ADPRHL2 |
| 12 | 104348360 | 104348554 | 9.45277283 | TXNRD1 |
| 12 | 104573686 | 104573876 | -1.470497903 | CHST11 |
| 12 | 104805090 | 104805284 | 1.91316452 | SLC41A2 |
| 12 | 104928241 | 104928459 | 2.27437943 | SLC41A2 |
| 12 | 105024379 | 105024579 | -1.923502774 | ALDH1L2 |
| 12 | 105024379 | 105024579 | -1.923502774 | C12orf45 |
| 12 | 105026643 | 105026842 | -2.928872966 | ALDH1L2 |
| 12 | 105026643 | 105026842 | -2.928872966 | C12orf45 |
| 12 | 106066779 | 106066915 | -3.298094878 | NUAK1 |
| 1 | 36093530 | 36093732 | 1.59317878 | ADPRHL2 |
| 12 | 106346429 | 106346628 | -2.11969042 | TCP11L2 |
| 2 | 157737303 | 157737509 | 2.29900562 | ACVR1 |
| 2 | 159616109 | 159616271 | 1.062483691 | AC009506.1 |
| 2 | 159616109 | 159616271 | 1.062483691 | BAZ2B |
| 2 | 159902624 | 159902832 | 6.811140328 | LY75-CD302 |
| 2 | 159902624 | 159902832 | 6.811140328 | LY75 |
| 2 | 161235242 | 161235439 | -1.055100998 | AC009299.2 |
| 2 | 161235242 | 161235439 | -1.055100998 | TANK |
| 2 | 161501466 | 161501612 | 2.660964047 | AC009487.6 |
| 2 | 161501466 | 161501612 | 2.660964047 | SLC4A10 |
| 2 | 162308078 | 162308278 | 4.453445298 | IFIH1 |
| 2 | 162309529 | 162309663 | 1.190714553 | IFIH1 |
| 2 | 162310834 | 162311035 | -1.163698746 | IFIH1 |
| 2 | 168236459 | 168236640 | -4.038407799 | STK39 |
| 2 | 169095603 | 169095841 | 1.360234708 | DHRS9 |
| 2 | 169504894 | 169505120 | 1.043562051 | RP11-724O16.1 |
| 2 | 169504894 | 169505120 | 1.043562051 | BBS5 |
| 2 | 169688820 | 169688934 | 1.173072166 | CCDC173 |
| 2 | 171070523 | 171070666 | -1.829481541 | TLK1 |
| 2 | 171091928 | 171092129 | -8.407766701 | TLK1 |
| 2 | 172080444 | 172080589 | 1.460891305 | METAP1D |
| 2 | 172081914 | 172082070 | 1.027239353 | METAP1D |
| 2 | 172085468 | 172085669 | -2.307892565 | DLX1 |
| 2 | 172503966 | 172504096 | -8.615929335 | ITGA6 |
| 2 | 172503966 | 172504096 | -8.615929335 | AC093818.1 |
| 1 | 145291498 | 145291694 | -1.039127254 | NBPF20 |
| 2 | 172527271 | 172527467 | 5.108372929 | AC093818.1 |
| 2 | 174335556 | 174335662 | -5.033044595 | RP11-572N21.1 |
| 2 | 174335556 | 174335662 | -5.033044595 | SP9 |
| 2 | 174337187 | 174337391 | 2.690269075 | SP9 |
| 2 | 174337187 | 174337391 | 2.690269075 | RP11-572N21.1 |
| 1 | 145660334 | 145660482 | -5.642701109 | GPR89A |
| 2 | 176131468 | 176131660 | -4.406890596 | HOXD8 |
| 2 | 176131468 | 176131660 | -4.406890596 | HOXD-AS2 |
| 2 | 177221630 | 177221812 | 1.196267351 | HNRNPA3 |
| 2 | 177354738 | 177354913 | -6.240899716 | AC074286.1 |
| 2 | 177354738 | 177354913 | -6.240899716 | NFE2L2 |
| 2 | 177369243 | 177369394 | -1.522499753 | AC074286.1 |
| 2 | 177369243 | 177369394 | -1.522499753 | NFE2L2 |
| 2 | 177371884 | 177372062 | 4.320122468 | NFE2L2 |
| 2 | 177371884 | 177372062 | 4.320122468 | AC074286.1 |
| 2 | 177459862 | 177460036 | -4.89395128 | AGPS |
| 2 | 181033510 | 181033662 | -3.350724122 | UBE2E3 |
| 1 | 146058682 | 146058846 | 1.778068573 | RP11-458D21.1 |
| 2 | 181922123 | 181922319 | -1.135075282 | SSFA2 |
| 1 | 146066090 | 146066293 | -1.379225268 | NBPF10 |
| 2 | 184817919 | 184818052 | -7.044021426 | ZNF804A |
| 2 | 184937872 | 184938123 | 1.025388325 | ZNF804A |
| 2 | 185547076 | 185547276 | -1.749657205 | AC080125.1 |
| 2 | 187519403 | 187519565 | -5.94208526 | TFPI |
| 2 | 187519403 | 187519565 | -5.94208526 | AC007319.1 |
| 2 | 189563898 | 189564130 | 2.919428373 | SLC40A1 |
| 2 | 189713839 | 189714042 | -5.537739575 | ANKAR |
| 2 | 189854507 | 189854705 | 1.365470903 | PMS1 |
| 2 | 190476764 | 190476946 | -7.286882757 | MFSD6 |

| | | | | |
|---|---|---|---|---|
| 1 | 26780180 | 26780416 | -1.039012582 | ARID1A |
| 11 | 72013169 | 72013365 | -1.887133487 | NUMA1 |
| 11 | 72015511 | 72015696 | -2.823716431 | RP11-849H4.4 |
| 11 | 72015511 | 72015696 | -2.823716431 | NUMA1 |
| 11 | 72293306 | 72293601 | -2.250482245 | CLPB |
| 11 | 72434167 | 72434362 | -3.522990161 | CLPB |
| 11 | 72838104 | 72838304 | -7.954817651 | FCHSD2 |
| 11 | 72838104 | 72838304 | -7.954817651 | ATG16L2 |
| 1 | 26787537 | 26787765 | -2.737287552 | PIGV |
| 11 | 73235159 | 73235441 | -5.238628647 | P2RY2 |
| 11 | 73297570 | 73297785 | -2.750726771 | P2RY6 |
| 11 | 73309464 | 73309686 | -1.252383524 | ARHGEF17 |
| 11 | 73310951 | 73311072 | -1.535888721 | ARHGEF17 |
| 11 | 73353126 | 73353248 | -5.94208526 | ARHGEF17 |
| 11 | 73873096 | 73873327 | -1.237185094 | COA4 |
| 11 | 73927446 | 73927604 | -1.99725432 | PAAF1 |
| 11 | 74013202 | 74013334 | -3.77146364 | C2CD3 |
| 11 | 74033970 | 74034163 | -1.689145284 | C2CD3 |
| 11 | 74128962 | 74129140 | -3.816813529 | C2CD3 |
| 11 | 74456995 | 74457206 | -1.920294296 | KCNE3 |
| 11 | 74456995 | 74457206 | -1.920294296 | RP11-702H23.4 |
| 11 | 74640694 | 74640893 | -2.209059126 | POLD3 |
| 11 | 74842109 | 74842315 | -1.470023509 | XRRA1 |
| 11 | 74842109 | 74842315 | -1.470023509 | RNF169 |
| 11 | 74877761 | 74877930 | -6.519373778 | RP11-147I3.1 |
| 11 | 74877761 | 74877930 | -6.519373778 | XRRA1 |
| 11 | 74878464 | 74878663 | -7.033044595 | XRRA1 |
| 11 | 74878464 | 74878663 | -7.033044595 | RP11-147I3.1 |
| 11 | 75005929 | 75006138 | -1.466899198 | NEU3 |
| 11 | 75007196 | 75007365 | -2.476952364 | NEU3 |
| 11 | 75571800 | 75572002 | -1.580261784 | SERPINH1 |
| 11 | 76141290 | 76141491 | -1.568992048 | UVRAG |
| 11 | 76186757 | 76187053 | -1.190817824 | WNT11 |
| 11 | 76211235 | 76211434 | -2.857683029 | RP11-619A14.3 |
| 11 | 76351699 | 76351891 | -1.16920869 | THAP12 |
| 11 | 76381923 | 76382123 | -2.627375422 | RP11-111M22.2 |
| 1 | 27005799 | 27005967 | -4.619543513 | FAM46B |
| 11 | 76796029 | 76796238 | 2.385034942 | TSKU |
| 11 | 76796556 | 76796686 | 2.017433299 | TSKU |
| 11 | 76909070 | 76909254 | -6.141544177 | ACER3 |
| 11 | 77103530 | 77103700 | -1.595443346 | CAPN5 |
| 11 | 78226646 | 78226775 | 1.297579516 | GAB2 |
| 11 | 82732365 | 82732487 | -6.683843878 | FAM181B |
| 11 | 82931807 | 82931967 | -6.494822232 | PRCP |
| 11 | 82931807 | 82931967 | -6.494822232 | DDIAS |
| 1 | 27549151 | 27549336 | 1.350569459 | AHDC1 |
| 11 | 83286085 | 83286210 | -6.404482087 | CCDC90B |
| 11 | 83286085 | 83286210 | -6.404482087 | RP11-727A23.10 |
| 11 | 85627154 | 85627365 | -2.041026123 | DLG2 |
| 11 | 85663935 | 85664134 | -1.405914794 | CREBZF |
| 11 | 85757848 | 85758048 | 3.971889711 | SYTL2 |
| 1 | 27549846 | 27550038 | 1.05543848 | AHDC1 |
| 11 | 86244778 | 86244979 | -1.311462967 | EED |
| 11 | 86284371 | 86284540 | -1.37931773 | RP11-320L11.2 |
| 12 | 106509516 | 106509778 | -2.214310943 | RP11-144F15.1 |
| 12 | 106509516 | 106509778 | -2.214310943 | POLR3B |
| 12 | 106971167 | 106971321 | -1.025074505 | TMEM263 |
| 12 | 107734244 | 107734443 | -1.089532828 | PRDM4 |
| 1 | 36096988 | 36097186 | -1.161704547 | COL8A2 |
| 12 | 108291520 | 108291718 | 1.69694119 | CMKLR1 |
| 12 | 108623653 | 108623883 | -1.089811452 | SELPLG |
| 12 | 108647288 | 108647489 | -1.109047206 | CORO1C |
| 12 | 108788802 | 108789021 | -1.019869489 | SSH1 |
| 12 | 109053735 | 109053919 | 5.861702487 | USP30 |
| 12 | 109053735 | 109053919 | 5.861702487 | USP30-AS1 |
| 1 | 1315852 | 1316048 | -5.836212671 | CPSF3L |
| 12 | 109092650 | 109092895 | -2.853044542 | ALKBH2 |
| 12 | 109769177 | 109769287 | 2.671771409 | FAM222A-AS1 |
| 12 | 109769177 | 109769287 | 2.671771409 | FAM222A |
| 12 | 109769303 | 109769469 | 1.67258414 | FAM222A |
| 12 | 109769303 | 109769469 | 1.67258414 | FAM222A-AS1 |
| 12 | 109916521 | 109916712 | -1.179479413 | TCHP |
| 12 | 109930189 | 109930385 | -1.49584647 | GIT2 |
| 12 | 109930189 | 109930385 | -1.49584647 | TCHP |
| 12 | 109948567 | 109948773 | -1.377790636 | GIT2 |
| 12 | 109948567 | 109948773 | -1.377790636 | TCHP |
| 12 | 109948567 | 109948773 | -1.377790636 | RP11-837J7.4 |
| 12 | 110429953 | 110430144 | -6.643567593 | HMGA1P3 |
| 12 | 110429953 | 110430144 | -6.643567593 | RP11-478C19.2 |
| 12 | 110629035 | 110629218 | -2.192131708 | HVCN1 |
| 12 | 110629035 | 110629218 | -2.192131708 | TCTN1 |
| 12 | 110629775 | 110629990 | -1.158580176 | TCTN1 |
| 12 | 110629775 | 110629990 | -1.158580176 | HVCN1 |
| 12 | 111347738 | 111347863 | -1.984567476 | CUX2 |
| 12 | 111362513 | 111362714 | -1.570989085 | FAM109A |
| 1 | 36289196 | 36289396 | -1.075458844 | THRAP3 |
| 12 | 111840208 | 111840408 | -1.933232254 | MAPKAPK5-AS1 |
| 12 | 111898601 | 111898813 | -1.627726613 | MAPKAPK5 |
| 12 | 111931487 | 111931688 | -2.295590118 | TMEM116 |
| 12 | 112140809 | 112140975 | 3.699015537 | TRAFD1 |
| 12 | 112302019 | 112302152 | 6.440748192 | RPL7AP60 |
| 12 | 112302019 | 112302152 | 6.440748192 | HECTD4 |
| 12 | 112941875 | 112942070 | 8.144216773 | RP1-71H24.1 |
| 12 | 112941875 | 112942070 | 8.144216773 | OAS3 |
| 12 | 113009218 | 113009422 | 1.262011008 | RP1-71H24.1 |
| 12 | 113009218 | 113009422 | 1.262011008 | OAS2 |
| 12 | 113296082 | 113296231 | -1.835462956 | TPCN1 |
| 12 | 113299587 | 113299777 | -9.048434772 | SLC8B1 |
| 12 | 114671792 | 114672047 | 1.811048774 | TBX3 |
| 12 | 114683522 | 114683705 | 3.817949733 | TBX3 |
| 12 | 116237586 | 116237800 | -1.438516111 | MED13L |
| 12 | 116675777 | 116675977 | -8.51666848 | RP11-497G19.1 |
| 12 | 116712265 | 116712469 | -8.598051299 | C12orf49 |
| 12 | 116715957 | 116716085 | -7.937394641 | C12orf49 |
| 12 | 116718938 | 116719133 | -2.05265796 | C12orf49 |
| 12 | 116723589 | 116723792 | -8.242977531 | C12orf49 |
| 12 | 118152310 | 118152517 | -1.901341034 | TAOK3 |
| 12 | 118157456 | 118157670 | -1.326644124 | TAOK3 |
| 1 | 146794299 | 146794429 | 1.155714979 | HYDIN2 |
| 2 | 190955365 | 190955565 | -1.993727373 | GLS |
| 2 | 190956109 | 190956308 | -1.877650718 | GLS |
| 2 | 190959186 | 190959389 | -2.923307176 | GLS |
| 2 | 195749424 | 195749587 | 1.185530824 | DNAH7 |
| 2 | 196752852 | 196752956 | -5.586838568 | CCDC150 |
| 2 | 197260274 | 197260460 | -6.351627367 | ANKRD44-IT1 |
| 2 | 197260274 | 197260460 | -6.351627367 | AC013264.1 |
| 2 | 197260274 | 197260460 | -6.351627367 | ANKRD44 |
| 2 | 197260971 | 197261190 | -2.162471898 | AC013264.1 |
| 2 | 197260971 | 197261190 | -2.162471898 | ANKRD44-IT1 |
| 2 | 197260971 | 197261190 | -2.162471898 | ANKRD44 |
| 2 | 199459428 | 199459627 | 1.036083499 | SATB2 |
| 2 | 199459428 | 199459627 | 1.036083499 | SATB2-AS1 |
| 2 | 200892047 | 200892250 | -2.940100881 | NIF3L1 |
| 1 | 147177816 | 147178015 | -1.137106809 | RP11-337C18.10 |
| 1 | 147177816 | 147178015 | -1.137106809 | RP11-337C18.8 |
| 1 | 147177816 | 147178015 | -1.137106809 | FMO5 |
| 2 | 200981646 | 200981847 | -1.153740141 | FAM126B |
| 2 | 200981646 | 200981847 | -1.153740141 | AC005037.3 |
| 2 | 201034407 | 201034604 | -4.857122759 | FAM126B |
| 2 | 201138656 | 201138849 | 7.770215394 | IMPDH1P10 |
| 2 | 201138656 | 201138849 | 7.770215394 | CFLAR |
| 2 | 201246485 | 201246664 | 2.346315397 | CASP8 |
| 2 | 201296193 | 201296419 | 1.489911059 | ALS2CR12 |
| 2 | 202779453 | 202779615 | -1.897207933 | ICA1L |
| 2 | 202925001 | 202925195 | -1.63106622 | WDR12 |
| 2 | 202925001 | 202925195 | -1.63106622 | CARF |
| 2 | 202925001 | 202925195 | -1.63106622 | AC010900.2 |
| 2 | 203107739 | 203107925 | -6.105335672 | NBEAL1 |
| 2 | 207239835 | 207239978 | -2.353909624 | AC007879.7 |
| 2 | 207239835 | 207239978 | -2.353909624 | AC007879.5 |
| 2 | 207586488 | 207586687 | -3.158742832 | METTL21A |
| 2 | 207586488 | 207586687 | -3.158742832 | CREB1 |
| 2 | 207605096 | 207605242 | -4.614409345 | METTL21A |
| 2 | 207767023 | 207767151 | 3.064146205 | FZD5 |
| 2 | 207977100 | 207977327 | -1.183411833 | PLEKHM3 |
| 2 | 208001268 | 208001527 | -1.250236054 | PLEKHM3 |
| 2 | 210139650 | 210139843 | -5.830889049 | KANSL1L |
| 2 | 211382395 | 211382602 | -2.035797876 | ERBB4 |
| 2 | 211383722 | 211383959 | -2.343573074 | ERBB4 |
| 2 | 212237642 | 212237820 | -3.054262228 | ERBB4 |
| 2 | 214728842 | 214729019 | 1.119333495 | BARD1 |
| 2 | 214965327 | 214965479 | -1.321928095 | ABCA12 |
| 2 | 215367861 | 215368058 | -3.554262228 | FN1 |
| 2 | 215611818 | 215612018 | 1.956407715 | LINC00607 |
| 2 | 215611818 | 215612018 | 1.956407715 | AC012668.2 |
| 2 | 218035050 | 218035241 | 1.650584767 | RUFY4 |
| 2 | 218068701 | 218068913 | 1.057471195 | RUFY4 |
| 2 | 218388622 | 218388813 | 1.120338608 | SLC11A1 |
| 2 | 219250974 | 219251182 | 2.89527248 | TUBA4A |
| 2 | 219539363 | 219539538 | 1.957275618 | CHPF |
| 2 | 219539817 | 219539937 | 1.860897051 | CHPF |
| 2 | 219570680 | 219570872 | 9.179694627 | OBSL1 |

| | | | | |
|---|---|---|---|---|
| 11 | 86808389 | 86808595 | 2.011142803 | PRSS23 |
| 1 | 27551186 | 27551384 | 1.05474979 | AHDC1 |
| 11 | 93156881 | 93157077 | 2.145319031 | SLC36A4 |
| 11 | 93696968 | 93697124 | -2.012168234 | CEP295 |
| 11 | 93727329 | 93727542 | -1.546498203 | CEP295 |
| 11 | 93727329 | 93727542 | -1.546498203 | RP11-178H8.7 |
| 11 | 93778485 | 93778605 | -1.704042369 | TAF1D |
| 1 | 27660368 | 27660647 | 7.277309414 | RP11-288L9.1 |
| 11 | 94543919 | 94544103 | -3.97406064 | PIWIL4 |
| 11 | 94543919 | 94544103 | -3.97406064 | FUT4 |
| 11 | 94544990 | 94545173 | -2.584144267 | PIWIL4 |
| 11 | 94544990 | 94545173 | -2.584144267 | FUT4 |
| 11 | 94546089 | 94546289 | -2.948712369 | RP11-867G2.8 |
| 11 | 94546089 | 94546289 | -2.948712369 | FUT4 |
| 11 | 94546089 | 94546289 | -2.948712369 | PIWIL4 |
| 11 | 94549325 | 94549525 | -3.397272104 | RP11-867G2.8 |
| 11 | 94549325 | 94549525 | -3.397272104 | PIWIL4 |
| 11 | 94549325 | 94549525 | -3.397272104 | FUT4 |
| 11 | 94799528 | 94799722 | -1.689895372 | AMOTL1 |
| 11 | 94799729 | 94799919 | -1.767655284 | AMOTL1 |
| 11 | 94870727 | 94870928 | -1.536002885 | AMOTL1 |
| 11 | 95067962 | 95068161 | -1.211778351 | SRSF8 |
| 11 | 95129398 | 95129597 | -1.303522645 | ENDOD1 |
| 11 | 95144828 | 95145027 | 3.321928095 | BUD13P1 |
| 11 | 95208892 | 95209058 | 2.953445298 | RP11-712B9.2 |
| 11 | 95208892 | 95209058 | 2.953445298 | SESN3 |
| 11 | 95806715 | 95806889 | -4.872416919 | CEP57 |
| 11 | 95811131 | 95811312 | -1.790510641 | CEP57 |
| 11 | 95875933 | 95876117 | -5.652460835 | MTMR2 |
| 11 | 95978883 | 95979061 | 7.972070661 | MAML2 |
| 11 | 96093106 | 96093309 | 5.648923988 | MAML2 |
| 11 | 96341519 | 96341721 | 5.674766206 | MAML2 |
| 11 | 96341519 | 96341721 | 5.674766206 | MIR1260B |
| 11 | 96342875 | 96343079 | 5.366123149 | MAML2 |
| 11 | 96390474 | 96390673 | -1.202190115 | JRKL |
| 1 | 27881763 | 27881960 | 2.407691648 | THEMIS2 |
| 11 | 102229698 | 102229917 | 1.544348179 | RP11-864G5.3 |
| 11 | 102229698 | 102229917 | 1.544348179 | YAP1 |
| 1 | 1233866 | 1234064 | -1.206157378 | B3GALT6 |
| 1 | 27882738 | 27882938 | 2.086027781 | THEMIS2 |
| 11 | 106009608 | 106009811 | -1.059134907 | MSANTD4 |
| 11 | 107504914 | 107505112 | -1.542232851 | ALKBH8 |
| 11 | 107961994 | 107962197 | -1.081491451 | RAB39A |
| 11 | 108161498 | 108161621 | -1.346321359 | NPAT |
| 11 | 108173549 | 108173763 | -1.247780211 | NPAT |
| 11 | 108206663 | 108206832 | -1.898033614 | NPAT |
| 11 | 108207003 | 108207240 | -1.603720501 | NPAT |
| 11 | 108265520 | 108265674 | -6.649246109 | ATM |
| 11 | 108514010 | 108514210 | -3.243493671 | EXPH5 |
| 11 | 108723093 | 108723292 | -1.252761776 | DDX10 |
| 11 | 108778478 | 108778677 | -5.642701109 | DDX10 |
| 11 | 110102293 | 110102531 | 2.547995042 | ZC3H12C |
| 11 | 110136967 | 110137135 | 3.144087725 | ZC3H12C |
| 11 | 110165306 | 110165524 | 3.308303843 | ZC3H12C |
| 12 | 118160180 | 118160419 | -1.649247085 | TAOK3 |
| 12 | 120095553 | 120095684 | -6.05979481 | RAB35 |
| 12 | 120117446 | 120117785 | -1.329225817 | RP1-267D11.6 |
| 12 | 120117446 | 120117785 | -1.329225817 | RAB35 |
| 1 | 37475488 | 37475719 | 2.696274573 | ZC3H12A |
| 12 | 120444560 | 120444757 | -2.702131514 | TRIAP1 |
| 12 | 120444560 | 120444757 | -2.702131514 | AL021546.6 |
| 12 | 120579126 | 120579337 | -1.342414212 | POP5 |
| 12 | 120764632 | 120764850 | -1.506041412 | SPPL3 |
| 1 | 37482961 | 37483163 | 3.711306081 | ZC3H12A |
| 12 | 121020476 | 121020679 | 2.973634384 | OASL |
| 12 | 121185992 | 121186191 | 3.175084139 | P2RX7 |
| 12 | 121641480 | 121641763 | 1.313064691 | ORAI1 |
| 12 | 121779242 | 121779463 | -1.943489153 | RHOF |
| 12 | 121779242 | 121779463 | -1.943489153 | TMEM120B |
| 12 | 121803162 | 121803319 | 1.572598958 | RP11-347I19.7 |
| 12 | 121803162 | 121803319 | 1.572598958 | LINC01089 |
| 12 | 121803162 | 121803319 | 1.572598958 | RHOF |
| 12 | 122019712 | 122019859 | -3.19868648 | BCL7A |
| 12 | 122019712 | 122019859 | -3.19868648 | RP11-87C12.5 |
| 12 | 122021136 | 122021282 | -5.715226276 | BCL7A |
| 12 | 122059192 | 122059392 | -2.050764934 | BCL7A |
| 12 | 122060045 | 122060409 | -2.804778096 | BCL7A |
| 12 | 122061443 | 122061658 | -1.969419924 | BCL7A |
| 12 | 122701684 | 122701929 | 4.843872282 | RP11-324E6.6 |
| 12 | 122701684 | 122701929 | 4.843872282 | HCAR2 |
| 12 | 122702934 | 122703134 | 4.629404344 | RP11-324E6.6 |
| 12 | 122702934 | 122703134 | 4.629404344 | HCAR2 |
| 12 | 122715372 | 122715570 | 5.702662023 | RP11-324E6.6 |
| 12 | 122715372 | 122715570 | 5.702662023 | HCAR3 |
| 12 | 122716388 | 122716579 | 4.601601691 | HCAR3 |
| 12 | 122730035 | 122730179 | 6.993454312 | HCAR1 |
| 12 | 122865559 | 122865770 | -1.367214378 | RP11-463O12.3 |
| 12 | 122865559 | 122865770 | -1.367214378 | VPS37B |
| 12 | 122960031 | 122960150 | -1.838851724 | ABCB9 |
| 12 | 122974878 | 122975074 | -1.411983528 | OGFOD2 |
| 12 | 122974878 | 122975074 | -1.411983528 | ABCB9 |
| 1 | 37799814 | 37800025 | -1.720136809 | RP11-109P14.9 |
| 1 | 37799814 | 37800025 | -1.720136809 | MANEAL |
| 12 | 122980181 | 122980326 | -1.247116362 | ARL6IP4 |
| 12 | 122980181 | 122980326 | -1.247116362 | ABCB9 |
| 12 | 122980181 | 122980326 | -1.247116362 | RP11-197N18.2 |
| 12 | 122985603 | 122985804 | -1.654891935 | PITPNM2 |
| 12 | 123221558 | 123221718 | -1.628693921 | MPHOSPH9 |
| 12 | 123253676 | 123253885 | -1.74086444 | RP11-282O18.3 |
| 12 | 123253676 | 123253885 | -1.74086444 | C12orf65 |
| 12 | 123257082 | 123257293 | -1.555320089 | RP11-282O18.3 |
| 12 | 123257082 | 123257293 | -1.555320089 | C12orf65 |
| 12 | 123415604 | 123415815 | -1.666371467 | RILPL2 |
| 12 | 123619877 | 123620138 | -1.31435299 | DDX55 |
| 12 | 123933184 | 123933330 | -2.539110524 | DNAH10OS |
| 12 | 123933184 | 123933330 | -2.539110524 | CCDC92 |
| 12 | 123933184 | 123933330 | -2.539110524 | DNAH10 |
| 12 | 123946011 | 123946190 | -2.190410892 | CCDC92 |
| 2 | 219570680 | 219570872 | 9.179694627 | INHA |
| 2 | 222318595 | 222318815 | -1.821739804 | RP11-384O8.1 |
| 2 | 222318595 | 222318815 | -1.821739804 | AC010980.2 |
| 1 | 148103457 | 148103731 | -1.076364597 | NBPF11 |
| 2 | 223963643 | 223963829 | -1.350395991 | MRPL44 |
| 2 | 224000803 | 224000976 | -4.244834397 | SERPINE2 |
| 2 | 224812555 | 224812740 | 2.166041158 | DOCK10 |
| 2 | 226799074 | 226799273 | 1.291324317 | IRS1 |
| 1 | 148288638 | 148288844 | -1.729715809 | RP11-495P10.9 |
| 2 | 226931329 | 226931513 | 4.27344723 | RHBDD1 |
| 2 | 226986107 | 226986309 | -1.563185929 | RHBDD1 |
| 2 | 226986788 | 226986978 | -6.030854352 | RHBDD1 |
| 1 | 148317821 | 148318004 | -2.359631796 | RP11-495P10.4 |
| 2 | 230514902 | 230515024 | 6.429087655 | HMGB1P3 |
| 2 | 230514902 | 230515024 | 6.429087655 | SP100 |
| 2 | 230525607 | 230525720 | 1.256277003 | AC010149.4 |
| 2 | 230525607 | 230525720 | 1.256277003 | SP100 |
| 2 | 230527576 | 230527817 | -1.075073784 | SP100 |
| 2 | 230527576 | 230527817 | -1.075073784 | AC010149.4 |
| 2 | 230803019 | 230803198 | -4.857122759 | CAB39 |
| 2 | 230804183 | 230804379 | -2.346087317 | CAB39 |
| 2 | 231108663 | 231108786 | -4.114409345 | PSMD1 |
| 2 | 231108663 | 231108786 | -4.114409345 | HTR2B |
| 2 | 231373964 | 231374114 | -2.566275365 | ARMC9 |
| 2 | 231397793 | 231397970 | -6.226892528 | B3GNT7 |
| 2 | 231481726 | 231481924 | 1.767864912 | NCL |
| 2 | 231524879 | 231525098 | -1.877381137 | NMUR1 |
| 2 | 231559960 | 231560160 | -3.521318261 | AC104634.3 |
| 2 | 232344008 | 232344163 | -6.355833487 | DIS3L2 |
| 2 | 234495737 | 234495903 | 1.164997821 | ARL4C |
| 2 | 236125390 | 236125583 | 1.087439255 | AGAP1 |
| 2 | 237574501 | 237574682 | 1.675789952 | RAB17 |
| 2 | 237762707 | 237762907 | 1.005894656 | LRRFIP1 |
| 2 | 238141016 | 238141170 | 7.490761344 | KLHL30 |
| 2 | 238151206 | 238151386 | -1.424328665 | KLHL30 |
| 2 | 239124271 | 239124471 | 7.286113256 | HDAC4 |
| 2 | 239308705 | 239308894 | -4.453445298 | HDAC4 |
| 2 | 239358325 | 239358504 | -5.693739433 | HDAC4 |
| 2 | 240488014 | 240488211 | -1.241058357 | ANKMY1 |
| 2 | 240491092 | 240491286 | -1.219483758 | ANKMY1 |
| 2 | 240630847 | 240631040 | 1.596764346 | GPR35 |
| 2 | 241077577 | 241077719 | -6.243668808 | SNED1 |
| 2 | 241077577 | 241077719 | -6.243668808 | MTERF4 |
| 2 | 241231782 | 241231959 | 5.493632006 | HDLBP |
| 2 | 241572088 | 241572274 | -6.165318412 | BOK |
| 2 | 241672663 | 241672859 | 1.801018007 | ATG4B |
| 2 | 241686049 | 241686206 | -2.371356394 | DTYMK |
| 1 | 148992534 | 148992718 | -5.152460835 | PDE4DIP |
| 20 | 327901 | 328104 | 1.435455993 | NRSN2-AS1 |
| 20 | 327901 | 328104 | 1.435455993 | SOX12 |
| 20 | 396597 | 396838 | -1.46995923 | TRIB3 |
| 20 | 1155582 | 1155800 | -6.395276069 | PSMF1 |
| 20 | 1565159 | 1565352 | -2.843536059 | SIRPB1 |
| 20 | 1565159 | 1565352 | -2.843536059 | RP4-576H24.4 |

| | | | | |
|---|---|---|---|---|
| 1 | 28149652 | 28149862 | 1.064275757 | PTAFR |
| 11 | 110233352 | 110233503 | 1.427063166 | RDX |
| 11 | 113224465 | 113224674 | -2.588129388 | NCAM1 |
| 11 | 113225547 | 113225685 | -1.090809447 | NCAM1 |
| 11 | 113226165 | 113226323 | -8.306434249 | NCAM1 |
| 11 | 113227829 | 113228035 | -1.00156003 | NCAM1 |
| 11 | 113228907 | 113229108 | -1.038792459 | NCAM1 |
| 11 | 113229724 | 113229922 | -1.603545141 | NCAM1 |
| 11 | 116762711 | 116762915 | -2.099877915 | BUD13 |
| 11 | 116849197 | 116849394 | -1.860680221 | SIK3 |
| 11 | 116849197 | 116849394 | -1.860680221 | APOA1-AS |
| 11 | 116866875 | 116867043 | -1.697347338 | SIK3 |
| 11 | 117315812 | 117316011 | 7.384918922 | BACE1 |
| 11 | 117315812 | 117316011 | 7.384918922 | CEP164 |
| 11 | 117412218 | 117412447 | -2.003350813 | CEP164 |
| 11 | 117995577 | 117995774 | 3.406467775 | IL10RA |
| 11 | 117998863 | 117999058 | 3.577397718 | IL10RA |
| 11 | 118396210 | 118396403 | 1.330009753 | UBE4A |
| 11 | 118502795 | 118502951 | -1.764635688 | KMT2A |
| 11 | 118503985 | 118504169 | -2.025644556 | KMT2A |
| 11 | 118505461 | 118505680 | -1.17155832 | KMT2A |
| 11 | 118506103 | 118506306 | -2.326194255 | KMT2A |
| 11 | 118522211 | 118522410 | -2.003227672 | RP11-770J1.3 |
| 11 | 118522211 | 118522410 | -2.003227672 | KMT2A |
| 11 | 118525274 | 118525430 | -1.298447765 | KMT2A |
| 11 | 118525274 | 118525430 | -1.298447765 | RP11-770J1.3 |
| 11 | 118628018 | 118628216 | 1.838483303 | PHLDB1 |
| 11 | 118628310 | 118628516 | 1.962806407 | PHLDB1 |
| 11 | 118631151 | 118631317 | 1.264256611 | PHLDB1 |
| 11 | 118656755 | 118656953 | 2.276450646 | PHLDB1 |
| 11 | 118900079 | 118900234 | 1.200292104 | BCL9L |
| 11 | 118908488 | 118908678 | 1.968387772 | BCL9L |
| 11 | 118996806 | 118996991 | -3.002642956 | RP11-110I1.12 |
| 11 | 119094039 | 119094247 | -2.228143114 | H2AFX |
| 11 | 119174612 | 119174795 | -1.270515881 | NLRX1 |
| 11 | 119183522 | 119183680 | -2.467463988 | NLRX1 |
| 11 | 119334892 | 119335042 | -2.247032939 | RNF26 |
| 11 | 119335989 | 119336215 | -2.400473281 | RNF26 |
| 1 | 28715112 | 28715280 | -1.658680621 | GMEB1 |
| 11 | 119356382 | 119356498 | -6.243417511 | USP2 |
| 11 | 119372875 | 119373023 | -7.508882679 | USP2 |
| 11 | 119372875 | 119373023 | -7.508882679 | RP11-334E6.3 |
| 11 | 120407575 | 120407776 | -8.470569666 | ARHGEF12 |
| 11 | 123426457 | 123426637 | -1.273462 | GRAMD1B |
| 11 | 123591038 | 123591204 | -1.488149082 | GRAMD1B |
| 11 | 123724877 | 123725005 | -2.167932967 | ZNF202 |
| 11 | 123726246 | 123726393 | -1.079155824 | ZNF202 |
| 11 | 123730504 | 123730772 | -1.120159162 | ZNF202 |
| 11 | 124746704 | 124746860 | -7.191717151 | NRGN |
| 1 | 1234641 | 1234844 | -1.112242769 | B3GALT6 |
| 11 | 124747325 | 124747520 | 1.254780371 | VSIG2 |
| 11 | 124987751 | 124987968 | -2.492572242 | CCDC15 |
| 11 | 125086850 | 125087051 | 5.48808228 | SLC37A2 |
| 11 | 125088188 | 125088396 | 4.984866206 | SLC37A2 |
| 12 | 125025360 | 125025562 | -2.34150024 | BRI3BP |
| 12 | 125025360 | 125025562 | -2.34150024 | THRIL |
| 12 | 130971846 | 130972040 | -5.04637857 | ADGRD1 |
| 12 | 130972347 | 130972494 | -6.825862217 | ADGRD1 |
| 12 | 130973528 | 130973737 | -5.245926548 | ADGRD1 |
| 12 | 131089092 | 131089296 | -2.948418465 | ADGRD1 |
| 12 | 131139329 | 131139505 | -4.645550458 | ADGRD1 |
| 12 | 131943449 | 131943646 | 6.056219753 | PUS1 |
| 1 | 37990006 | 37990197 | -1.160120006 | SF3A3 |
| 12 | 132077659 | 132077848 | -1.311512584 | EP400 |
| 12 | 132077659 | 132077848 | -1.311512584 | RP13-820C6.2 |
| 12 | 132078455 | 132078655 | -1.442499931 | EP400 |
| 12 | 132078455 | 132078655 | -1.442499931 | RP13-820C6.2 |
| 12 | 132129207 | 132129442 | -1.057596317 | EP400NL |
| 12 | 132186909 | 132187151 | -1.059663392 | RP13-977J11.2 |
| 12 | 132189292 | 132189492 | -1.146179203 | RP13-977J11.2 |
| 1 | 38004596 | 38004840 | 4.539339171 | FHL3 |
| 12 | 132553114 | 132553299 | 6.037070731 | RP13-554M15.7 |
| 12 | 132553114 | 132553299 | 6.037070731 | FBRSL1 |
| 1 | 38860106 | 38860283 | 3.376660364 | RP5-864K19.4 |
| 1 | 38860106 | 38860283 | 3.376660364 | RP5-864K19.7 |
| 12 | 132926706 | 132926902 | -1.999471122 | ZNF605 |
| 12 | 132926706 | 132926902 | -1.999471122 | CHFR |
| 12 | 132942809 | 132943032 | -1.410275928 | ZNF605 |
| 12 | 132942809 | 132943032 | -1.410275928 | CHFR |
| 12 | 133058332 | 133058499 | -1.377602477 | ZNF84 |
| 12 | 133058933 | 133059136 | -1.25654612 | ZNF84 |
| 1 | 1327094 | 1327239 | 6.434025427 | CPTP |
| 1 | 39104819 | 39104960 | 8.781668903 | MACF1 |
| 12 | 133106498 | 133106646 | -1.064878844 | ZNF140 |
| 12 | 133121820 | 133122027 | -2.38441083 | CTD-2140B24.6 |
| 12 | 133121820 | 133122027 | -2.38441083 | ZNF891 |
| 12 | 133155568 | 133155774 | -2.953445298 | ZNF10 |
| 12 | 133155568 | 133155774 | -2.953445298 | CTD-2140B24.4 |
| 12 | 133156214 | 133156402 | -2.758680263 | ZNF10 |
| 12 | 133156214 | 133156402 | -2.758680263 | CTD-2140B24.4 |
| 12 | 133202471 | 133202586 | -7.218355771 | ZNF268 |
| 12 | 133204365 | 133204579 | -1.249552646 | ZNF268 |
| 13 | 18209000 | 18209198 | -2.079732203 | FAM230C |
| 1 | 39269118 | 39269303 | 7.488170182 | MACF1 |
| 13 | 19564368 | 19564515 | -4.708926257 | MRPS31P2 |
| 13 | 19564368 | 19564515 | -4.708926257 | CASC4P1 |
| 13 | 19824916 | 19825085 | -2.079152959 | ZMYM5 |
| 13 | 19864225 | 19864476 | -2.415788512 | RP11-61K9.3 |
| 13 | 19993579 | 19993784 | -1.245696243 | ZMYM2 |
| 13 | 20141824 | 20142122 | -1.844825391 | GJA3 |
| 13 | 20143091 | 20143288 | -1.62118596 | GJA3 |
| 13 | 20608076 | 20608249 | -1.532833459 | SLC35E1P1 |
| 13 | 20608076 | 20608249 | -1.532833459 | IFT88 |
| 13 | 20722933 | 20723068 | -4.706813965 | IL17D |
| 13 | 20981570 | 20981750 | -1.794118816 | LATS2 |
| 13 | 21358317 | 21358452 | -3.169925001 | GAPDHP52 |
| 13 | 23335479 | 23335679 | 1.256277003 | SACS |
| 13 | 24249396 | 24249593 | -3.285349522 | SPATA13 |
| 1 | 149554687 | 149554966 | -1.321836837 | NBPF19 |
| 20 | 3165037 | 3165237 | -3.37462047 | LZTS3 |
| 20 | 3866192 | 3866383 | 1.321077923 | MAVS |
| 1 | 149676703 | 149676876 | -3.4198882 | RP11-353N4.5 |
| 1 | 149676703 | 149676876 | -3.4198882 | LINC00869 |
| 20 | 4182176 | 4182368 | 1.274306195 | SMOX |
| 20 | 4699532 | 4699734 | -1.340971741 | PRNP |
| 20 | 4782835 | 4783042 | 2.691827081 | RASSF2 |
| 20 | 4784022 | 4784220 | 1.64890462 | RASSF2 |
| 20 | 5942503 | 5942720 | -1.477817337 | TRMT6 |
| 20 | 6768007 | 6768207 | 9.080479804 | BMP2 |
| 20 | 6768513 | 6768694 | 2.046208619 | BMP2 |
| 20 | 6778566 | 6778766 | 1.346870606 | BMP2 |
| 20 | 10639604 | 10639803 | 2.313757745 | JAG1 |
| 1 | 149787800 | 149787992 | 8.126101715 | HIST2H2BF |
| 1 | 149787800 | 149787992 | 8.126101715 | RP11-196G18.3 |
| 1 | 149787800 | 149787992 | 8.126101715 | FCGR1A |
| 20 | 17967968 | 17968191 | 6.923038993 | OVOL2 |
| 20 | 17967968 | 17968191 | 6.923038993 | SNX5 |
| 20 | 17990165 | 17990364 | 1.23719976 | MGME1 |
| 20 | 17990165 | 17990364 | 1.23719976 | OVOL2 |
| 20 | 18181567 | 18181761 | 1.720799163 | KAT14 |
| 1 | 149791320 | 149791522 | -2.552481199 | HIST2H2BF |
| 1 | 149791320 | 149791522 | -2.552481199 | RP11-196G18.3 |
| 1 | 149791320 | 149791522 | -2.552481199 | FCGR1A |
| 20 | 18384228 | 18384426 | -2.183638541 | DZANK1 |
| 20 | 20369623 | 20369790 | 4.145470201 | INSM1 |
| 20 | 20391442 | 20391617 | -5.882643049 | RALGAPA2 |
| 1 | 149792226 | 149792375 | 1.766664866 | RP11-196G18.3 |
| 1 | 149792226 | 149792375 | 1.766664866 | FCGR1A |
| 1 | 149792226 | 149792375 | 1.766664866 | HIST2H2BF |
| 20 | 21102441 | 21102657 | -2.087362994 | LINC00237 |
| 20 | 21161955 | 21162155 | -1.043181312 | KIZ-AS1 |
| 20 | 21161955 | 21162155 | -1.043181312 | RP5-872K7.7 |
| 20 | 21161955 | 21162155 | -1.043181312 | KIZ |
| 20 | 21168946 | 21169166 | -1.368482797 | KIZ |
| 20 | 21168946 | 21169166 | -1.368482797 | KIZ-AS1 |
| 20 | 21395773 | 21395977 | 2.157246642 | NKX2-4 |
| 20 | 21500076 | 21500274 | 1.523732387 | RP5-984P4.6 |
| 1 | 149859179 | 149859363 | 5.073602462 | HIST2H4B |
| 20 | 25221941 | 25222088 | 1.354482863 | ENTPD6 |
| 20 | 25240893 | 25241088 | 2.127544336 | RP5-965G21.4 |
| 20 | 25285817 | 25286001 | -5.099836172 | PYGB |
| 20 | 25476373 | 25476526 | 1.546669704 | NINL |
| 20 | 25683343 | 25683497 | -1.217193034 | ZNF337 |
| 20 | 25856460 | 25856659 | 2.903677461 | FAM182B |
| 20 | 25857042 | 25857210 | 5.642701109 | FAM182B |
| 20 | 31548487 | 31548676 | 8.268639539 | HM13 |
| 20 | 31665977 | 31666168 | 6.298094878 | BCL2L1 |
| 20 | 34550010 | 34550146 | 2.885744735 | MAP1LC3A |
| 20 | 35954150 | 35954335 | 7.940963245 | SCAND1 |
| 20 | 38089848 | 38090054 | 1.020131935 | RPRD1B |
| 20 | 40688517 | 40688619 | 9.055348699 | MAFB |
| 20 | 44224975 | 44225144 | -1.168141694 | OSER1-AS1 |

| | | | | |
|---|---|---|---|---|
| 11 | 125431537 | 125431717 | 5.986489893 | PKNOX2 |
| 11 | 125678320 | 125678430 | -2.625520368 | ACRV1 |
| 11 | 125680571 | 125680716 | -4.031867912 | ACRV1 |
| 11 | 125887117 | 125887305 | -6.819274522 | HYLS1 |
| 11 | 125899831 | 125900078 | -2.752946568 | PUS3 |
| 11 | 125899831 | 125900078 | -2.752946568 | HYLS1 |
| 11 | 125960877 | 125961060 | 1.203744906 | CDON |
| 11 | 126203194 | 126203440 | -1.470828726 | RPUSD4 |
| 1 | 28987531 | 28987732 | -1.501363219 | EPB41 |
| 11 | 126277812 | 126277976 | -2.626491621 | FOXRED1 |
| 11 | 126293955 | 126294141 | -1.551778644 | RP11-712L6.7 |
| 11 | 126293955 | 126294141 | -1.551778644 | TIRAP |
| 11 | 126317453 | 126317649 | -5.434411277 | DCPS |
| 11 | 126337236 | 126337435 | -7.94447862 | DCPS |
| 11 | 126341954 | 126342176 | -2.168689905 | DCPS |
| 11 | 126341954 | 126342176 | -2.168689905 | ST3GAL4-AS1 |
| 11 | 128462270 | 128462451 | 1.316128097 | ETS1 |
| 11 | 128810574 | 128810774 | -1.225012898 | FLI1 |
| 11 | 128970609 | 128970772 | -1.005568768 | ARHGAP32 |
| 1 | 29148902 | 29149106 | -3.172752177 | SRSF4 |
| 11 | 129869574 | 129869794 | -1.073112686 | NFRKB |
| 1 | 29193345 | 29193555 | -2.611079566 | MECR |
| 11 | 130449028 | 130449222 | -2.30710494 | ADAMTS15 |
| 11 | 130473891 | 130474093 | -1.386176934 | ADAMTS15 |
| 11 | 130476242 | 130476465 | -1.568512821 | ADAMTS15 |
| 11 | 130914576 | 130914777 | -1.120593816 | SNX19 |
| 11 | 130915096 | 130915301 | -1.294787199 | SNX19 |
| 11 | 134264948 | 134265156 | -1.018667044 | ACAD8 |
| 12 | 14615 | 14845 | -1.180977412 | WASH7P |
| 12 | 285489 | 285607 | -1.065513687 | KDM5A |
| 12 | 389303 | 389526 | -1.203803629 | KDM5A |
| 12 | 389303 | 389526 | -1.203803629 | CCDC77 |
| 12 | 643967 | 644170 | 2.437245837 | RP11-218M22.1 |
| 12 | 643967 | 644170 | 2.437245837 | NINJ2 |
| 1 | 30721559 | 30721741 | -1.72537558 | MATN1 |
| 1 | 30721559 | 30721741 | -1.72537558 | MATN1-AS1 |
| 12 | 868043 | 868244 | 10.92830927 | WNK1 |
| 12 | 868702 | 868904 | 3.317653238 | WNK1 |
| 12 | 908633 | 908772 | 1.62095317 | WNK1 |
| 12 | 916947 | 917154 | -1.231150598 | RAD52 |
| 12 | 975822 | 976021 | -1.928132261 | RAD52 |
| 12 | 975822 | 976021 | -1.928132261 | RP11-359B12.2 |
| 1 | 30872770 | 30873010 | 3.280419692 | SDC3 |
| 12 | 976401 | 976587 | -1.550121729 | RAD52 |
| 12 | 976401 | 976587 | -1.550121729 | RP11-359B12.2 |
| 1 | 30874408 | 30874603 | 2.74283383 | SDC3 |
| 12 | 1815766 | 1815936 | 6.817745703 | CACNA2D4 |
| 12 | 1817380 | 1817573 | 5.021779492 | CACNA2D4 |
| 12 | 1831220 | 1831401 | 5.729203307 | LRTM2 |
| 12 | 1831220 | 1831401 | 5.729203307 | CACNA2D4 |
| 12 | 2859032 | 2859271 | -3.865147233 | ITFG2 |
| 12 | 2859032 | 2859271 | -3.865147233 | FOXM1 |
| 12 | 2874073 | 2874272 | -4.419378968 | FOXM1 |
| 1 | 30876730 | 30876857 | 7.066892205 | SDC3 |

| | | | | |
|---|---|---|---|---|
| 13 | 24249396 | 24249593 | -3.285349522 | RP11-307N16.6 |
| 13 | 24297438 | 24297636 | -2.238107069 | RP11-307N16.6 |
| 13 | 24297438 | 24297636 | -2.238107069 | SPATA13 |
| 13 | 24303833 | 24303992 | -1.70243937 | SPATA13 |
| 13 | 24303833 | 24303992 | -1.70243937 | RP11-307N16.6 |
| 13 | 27430790 | 27431005 | -3.660975548 | GTF3A |
| 13 | 27440097 | 27440306 | -1.165237565 | MTIF3 |
| 13 | 27650456 | 27650654 | -1.454241341 | POLR1D |
| 13 | 27651064 | 27651170 | -2.34622128 | POLR1D |
| 13 | 27665643 | 27665828 | -2.880999461 | POLR1D |
| 13 | 28292436 | 28292637 | -1.06406302 | PAN3 |
| 13 | 28293668 | 28293868 | -1.573934201 | PAN3 |
| 13 | 28496765 | 28496968 | -1.649001128 | RP11-95G6.1 |
| 13 | 31329597 | 31329817 | -1.826627831 | B3GLCT |
| 13 | 31331166 | 31331313 | -2.139399742 | B3GLCT |
| 13 | 32298669 | 32298867 | -1.484207137 | FRY |
| 13 | 32332971 | 32333208 | 1.833758749 | BRCA2 |
| 13 | 32355014 | 32355192 | 1.334555519 | BRCA2 |
| 13 | 32770199 | 32770399 | -6.667695177 | PDS5B |
| 13 | 33105510 | 33105729 | -2.669773231 | STARD13 |
| 13 | 33129453 | 33129668 | -1.681579819 | STARD13 |
| 13 | 36227132 | 36227381 | -2.977873381 | CCDC169 |
| 13 | 36227132 | 36227381 | -2.977873381 | CCDC169-SOHLH2 |
| 13 | 36255111 | 36255286 | -1.931749 | CCDC169 |
| 13 | 36255111 | 36255286 | -1.931749 | CCDC169-SOHLH2 |
| 13 | 36819591 | 36819788 | 7.017485433 | RFXAP |
| 13 | 40559957 | 40560153 | -2.332981727 | FOXO1 |
| 13 | 40560480 | 40560671 | -1.331381144 | FOXO1 |
| 13 | 40864777 | 40865028 | -2.429021904 | TPTE2P5 |
| 13 | 40865558 | 40865729 | -2.25548096 | TPTE2P5 |
| 13 | 40875011 | 40875278 | -2.910697188 | TPTE2P5 |
| 13 | 40877905 | 40878103 | -1.112078439 | TPTE2P5 |
| 13 | 40941039 | 40941234 | -1.216795785 | ELF1 |
| 13 | 41043285 | 41043419 | -6.332778995 | ELF1 |
| 13 | 41193051 | 41193251 | -1.281501856 | KBTBD7 |
| 13 | 41193051 | 41193251 | -1.281501856 | RP11-74J13.8 |
| 13 | 41371217 | 41371418 | -1.982497451 | NAA16 |
| 13 | 41567947 | 41568156 | -1.666551101 | VWA8 |
| 13 | 41839476 | 41839681 | -1.199040174 | VWA8 |
| 13 | 42040017 | 42040137 | 2.502110233 | DGKH |
| 13 | 42041456 | 42041608 | -5.75042094 | DGKH |
| 13 | 42047634 | 42047820 | 1.231459948 | DGKH |
| 13 | 42048079 | 42048277 | 1.507999082 | DGKH |
| 1 | 39411514 | 39411709 | 2.324731167 | MACF1 |
| 13 | 42301891 | 42302085 | -6.475460176 | AKAP11 |
| 13 | 42303325 | 42303544 | -1.750335103 | AKAP11 |
| 13 | 45957049 | 45957231 | -1.986448821 | ZC3H13 |
| 1 | 39412034 | 39412212 | 2.545084774 | MACF1 |
| 13 | 45969644 | 45969844 | -1.294587917 | ZC3H13 |
| 13 | 46372197 | 46372411 | -2.101588558 | RUBCNL |
| 13 | 46437491 | 46437695 | -1.726163451 | PPP1R2P4 |
| 13 | 46437491 | 46437695 | -1.726163451 | RUBCNL |
| 1 | 39414183 | 39414381 | 3.314095018 | MACF1 |
| 13 | 48038815 | 48039003 | 1.030679544 | NUDT15 |

| | | | | |
|---|---|---|---|---|
| 1 | 150507137 | 150507339 | 1.512908549 | TARS2 |
| 20 | 44755949 | 44756135 | 6.654738177 | RIMS4 |
| 20 | 44958673 | 44958872 | -7.491051153 | TOMM34 |
| 20 | 44958673 | 44958872 | -7.491051153 | PABPC1L |
| 20 | 45706289 | 45706456 | 3.025886825 | WFDC13 |
| 20 | 46129071 | 46129271 | 3.560729756 | CD40 |
| 20 | 46541328 | 46541527 | 1.479553609 | OCSTAMP |
| 20 | 47636467 | 47636658 | -1.253804934 | NCOA3 |
| 20 | 47647025 | 47647227 | -1.141235182 | NCOA3 |
| 20 | 49144368 | 49144647 | 6.448922728 | STAU1 |
| 1 | 6415368 | 6415540 | -7.502790346 | HES2 |
| 20 | 49636181 | 49636421 | -1.514858662 | B4GALT5 |
| 20 | 49988332 | 49988485 | 1.235001408 | SNAI1 |
| 1 | 150579431 | 150579627 | 12.53189926 | MCL1 |
| 20 | 51003881 | 51004080 | -1.198008442 | KCNG1 |
| 20 | 51003881 | 51004080 | -1.198008442 | RP5-955M13.4 |
| 20 | 51598656 | 51598860 | -2.011732873 | ATP9A |
| 20 | 51600020 | 51600193 | -2.227943427 | ATP9A |
| 20 | 51601035 | 51601233 | -3.184964094 | ATP9A |
| 20 | 51784451 | 51784557 | -1.706182356 | SALL4 |
| 20 | 51791911 | 51792122 | -2.300467808 | SALL4 |
| 20 | 57371342 | 57371528 | 5.739890132 | RAE1 |
| 20 | 57652104 | 57652458 | 1.56329999 | PMEPA1 |
| 20 | 58709532 | 58709756 | -4.313363182 | STX16-NPEPL1 |
| 20 | 58709532 | 58709756 | -4.313363182 | NPEPL1 |
| 20 | 58991089 | 58991251 | 2.835538829 | NELFCD |
| 20 | 59774330 | 59774489 | -7.926898435 | PHACTR3 |
| 20 | 59805916 | 59806113 | -6.921764267 | PHACTR3 |
| 20 | 59938270 | 59938437 | -5.735837607 | FAM217B |
| 20 | 59938270 | 59938437 | -5.735837607 | PPP1R3D |
| 20 | 59939152 | 59939348 | 3.36827784 | FAM217B |
| 20 | 59939152 | 59939348 | 3.36827784 | PPP1R3D |
| 20 | 60058202 | 60058306 | 2.962595706 | C20orf197 |
| 20 | 60240145 | 60240314 | 1.223954874 | MIR646HG |
| 20 | 60323199 | 60323364 | -1.877443751 | MIR646HG |
| 1 | 151002283 | 151002493 | 1.255153082 | FAM63A |
| 20 | 62200877 | 62201079 | 2.291021513 | MTG2 |
| 20 | 62398430 | 62398574 | -1.921988194 | CABLES2 |
| 20 | 62399607 | 62399754 | -6.139805291 | CABLES2 |
| 20 | 62400903 | 62401237 | -1.192431897 | CABLES2 |
| 20 | 62401882 | 62402003 | -3.284927804 | CABLES2 |
| 20 | 62402599 | 62402783 | -5.079935668 | RP5-908M14.10 |
| 20 | 62402599 | 62402783 | -5.079935668 | CABLES2 |
| 20 | 62603094 | 62603274 | -1.488986847 | WI2-87327B8.2 |
| 20 | 62670165 | 62670287 | -5.724574323 | SLCO4A1 |
| 20 | 62714577 | 62714766 | -5.459431619 | NTSR1 |
| 20 | 62722573 | 62722712 | 5.897004823 | NTSR1 |
| 20 | 62751967 | 62752105 | 6.405084969 | NTSR1 |
| 1 | 6418626 | 6418837 | -1.826740311 | HES2 |
| 1 | 151059661 | 151059784 | 2.448630947 | MLLT11 |
| 1 | 151059661 | 151059784 | 2.448630947 | CDC42SE1 |
| 20 | 62879415 | 62879725 | 1.334629793 | DIDO1 |
| 20 | 62937620 | 62937821 | 2.775126736 | DIDO1 |
| 20 | 62967362 | 62967570 | 1.016571377 | SLC17A9 |

| | | | | |
|---|---|---|---|---|
| 12 | 2888173 | 2888392 | -2.269180537 | TULP3 |
| 12 | 2888173 | 2888392 | -2.269180537 | RHNO1 |
| 12 | 2889067 | 2889268 | -1.798193007 | TULP3 |
| 12 | 2889067 | 2889268 | -1.798193007 | RHNO1 |
| 12 | 3040379 | 3040584 | -2.357411592 | TEAD4 |
| 12 | 4520400 | 4520598 | 4.497176718 | C12orf4 |
| 12 | 6493575 | 6493869 | -1.615166914 | NCAPD2 |
| 12 | 6493575 | 6493869 | -1.615166914 | MRPL51 |
| 12 | 6557135 | 6557334 | -2.54135533 | NOP2 |
| 12 | 6644352 | 6644543 | -7.966245799 | ACRBP |
| 12 | 6666883 | 6667082 | -3.33297672 | RP4-761J14.8 |
| 12 | 6666883 | 6667082 | -3.33297672 | ZNF384 |
| 12 | 6672131 | 6672322 | -2.914960695 | ZNF384 |
| 12 | 6827113 | 6827313 | 6.24467312 | GPR162 |
| 12 | 6844971 | 6845276 | -2.965947318 | GNB3 |
| 12 | 6844971 | 6845276 | -2.965947318 | CDCA3 |
| 12 | 6884613 | 6884838 | -1.059468591 | DSTNP2 |
| 12 | 6884613 | 6884838 | -1.059468591 | LRRC23 |
| 12 | 6884613 | 6884838 | -1.059468591 | U47924.6 |
| 12 | 6884613 | 6884838 | -1.059468591 | SPSB2 |
| 12 | 6975761 | 6975929 | -1.948724543 | EMG1 |
| 12 | 6975761 | 6975929 | -1.948724543 | RP3-461F17.3 |
| 12 | 7070468 | 7070698 | -2.279871692 | C1S |
| 12 | 7080562 | 7080775 | 2.053300271 | C1R |
| 12 | 7096400 | 7096692 | -1.283946374 | C1RL |
| 12 | 7099512 | 7099721 | -1.111176406 | C1RL |
| 12 | 7110399 | 7110573 | -2.621788876 | C1RL-AS1 |
| 12 | 7110960 | 7111159 | -2.800978505 | C1RL-AS1 |
| 12 | 7115716 | 7115910 | -1.985835729 | C1RL-AS1 |
| 12 | 7115716 | 7115910 | -1.985835729 | ABC12-49244600F4.4 |
| 1 | 32181261 | 32181470 | -1.13472372 | TXLNA |
| 12 | 7813947 | 7814207 | 2.345378582 | SLC2A14 |
| 12 | 7921056 | 7921300 | 2.807495641 | SLC2A3 |
| 12 | 8033089 | 8033288 | -1.18665334 | FOXJ2 |
| 12 | 8221590 | 8221753 | -3.244050481 | FAM90A1 |
| 12 | 8235494 | 8235598 | -1.191993608 | FAM86FP |
| 12 | 8235494 | 8235598 | -1.191993608 | RP11-266K4.9 |
| 12 | 8238573 | 8238807 | -1.58158698 | RP11-266K4.9 |
| 12 | 8238573 | 8238807 | -1.58158698 | FAM86FP |
| 1 | 32194937 | 32195146 | -1.325941534 | TXLNA |
| 12 | 8242461 | 8242777 | -3.281757041 | RP11-266K4.9 |
| 12 | 8242461 | 8242777 | -3.281757041 | FAM86FP |
| 12 | 9695469 | 9695827 | 2.106627893 | RP11-75L1.2 |
| 12 | 9695469 | 9695827 | 2.106627893 | CLEC2D |
| 1 | 32207789 | 32208050 | -2.999248531 | IQCC |
| 12 | 11379317 | 11379491 | 1.487790724 | PRB1 |
| 12 | 11513987 | 11514185 | 5.690771476 | DDX55P1 |
| 12 | 12477336 | 12477536 | 1.024141097 | DUSP16 |
| 12 | 12717877 | 12718080 | -1.587239787 | CDKN1B |
| 12 | 12774030 | 12774217 | -4.383136272 | APOLD1 |
| 12 | 12787463 | 12787679 | -2.291107683 | APOLD1 |
| 12 | 12788787 | 12788986 | -2.252125259 | APOLD1 |
| 12 | 13002137 | 13002387 | 1.253411379 | HTR7P1 |
| 12 | 13002137 | 13002387 | 1.253411379 | RP11-377D9.3 |
| 13 | 48395405 | 48395612 | -2.687033313 | LPAR6 |
| 13 | 48395405 | 48395612 | -2.687033313 | RB1 |
| 13 | 48396446 | 48396634 | -2.414072486 | RB1 |
| 13 | 48396446 | 48396634 | -2.414072486 | LPAR6 |
| 13 | 48397192 | 48397392 | -1.465532257 | LPAR6 |
| 13 | 48397192 | 48397392 | -1.465532257 | RB1 |
| 13 | 48412266 | 48412415 | -1.653289517 | LPAR6 |
| 13 | 48412266 | 48412415 | -1.653289517 | RB1 |
| 13 | 48418527 | 48418683 | -3.043711798 | RB1 |
| 13 | 48418527 | 48418683 | -3.043711798 | LPAR6 |
| 13 | 48492519 | 48492715 | -1.829880142 | RCBTB2 |
| 13 | 49222191 | 49222417 | -4.623963757 | MLNR |
| 13 | 49274119 | 49274272 | 4.403677461 | CDADC1 |
| 13 | 49476533 | 49476763 | -3.778382804 | SETDB2 |
| 13 | 49630567 | 49630768 | -1.922965099 | ARL11 |
| 13 | 50715469 | 50715580 | -2.789091984 | DLEU7 |
| 13 | 50715469 | 50715580 | -2.789091984 | DLEU1 |
| 13 | 50807750 | 50807943 | -7.174087434 | DLEU7 |
| 13 | 50807750 | 50807943 | -7.174087434 | DLEU7-AS1 |
| 13 | 50834010 | 50834178 | -5.925874521 | DLEU7 |
| 13 | 50834010 | 50834178 | -5.925874521 | DLEU7-AS1 |
| 13 | 51369036 | 51369211 | -1.43541255 | INTS6 |
| 13 | 51460543 | 51460787 | -1.101411142 | INTS6-AS1 |
| 13 | 51460543 | 51460787 | -1.101411142 | RPS4XP16 |
| 1 | 39625397 | 39625606 | -1.623821298 | HEYL |
| 13 | 52461608 | 52461759 | -2.584173559 | CKAP2 |
| 13 | 57634846 | 57634997 | 6.910687696 | PCDH17 |
| 13 | 57725439 | 57725612 | 5.287484444 | PCDH17 |
| 13 | 67226738 | 67226919 | 2.823505168 | PCDH9 |
| 13 | 67228230 | 67228429 | 1.706885783 | PCDH9 |
| 13 | 71681840 | 71682045 | 7.198202048 | DACH1 |
| 13 | 72746751 | 72746922 | -1.683781758 | BORA |
| 1 | 1333701 | 1333879 | 6.869415578 | TAS1R3 |
| 13 | 73062278 | 73062492 | 2.018572504 | KLF5 |
| 13 | 73694451 | 73694644 | -1.811343566 | KLF12 |
| 13 | 73846143 | 73846354 | -1.363840762 | KLF12 |
| 13 | 75286754 | 75286981 | -1.385061008 | TBC1D4 |
| 13 | 76884861 | 76885063 | 2.146373797 | KCTD12 |
| 13 | 76991728 | 76991929 | 2.172927571 | CLN5 |
| 1 | 39758732 | 39758932 | -1.163997845 | BMP8B |
| 1 | 39758732 | 39758932 | -1.163997845 | PPIE |
| 13 | 78600866 | 78601051 | -5.480000966 | POU4F1 |
| 13 | 78600866 | 78601051 | -5.480000966 | RNF219-AS1 |
| 13 | 80337076 | 80337280 | -2.192446992 | SPRY2 |
| 1 | 39759656 | 39759857 | -2.038408629 | BMP8B |
| 1 | 39759656 | 39759857 | -2.038408629 | PPIE |
| 1 | 39769707 | 39769933 | -1.006106407 | BMP8B |
| 1 | 39769707 | 39769933 | -1.006106407 | OXCT2 |
| 13 | 87675603 | 87675803 | -1.164419335 | SLITRK5 |
| 13 | 87676421 | 87676624 | -1.855370369 | SLITRK5 |
| 13 | 87678108 | 87678309 | -1.816442468 | SLITRK5 |
| 13 | 87692729 | 87692923 | -2.694096327 | SLITRK5 |
| 1 | 39770334 | 39770528 | -2.20751875 | BMP8B |
| 1 | 39770334 | 39770528 | -2.20751875 | OXCT2 |
| 20 | 63196644 | 63196842 | -1.093337836 | YTHDF1 |
| 1 | 151159433 | 151159635 | 1.323964694 | TNFAIP8L2 |
| 1 | 151159433 | 151159635 | 1.323964694 | SCNM1 |
| 20 | 63590010 | 63590211 | 2.828402292 | GMEB2 |
| 20 | 63682552 | 63682748 | 2.29967092 | RTEL1-TNFRSF6B |
| 20 | 63682552 | 63682748 | 2.29967092 | RTEL1 |
| 20 | 63683740 | 63683920 | -2.709806486 | RTEL1 |
| 20 | 63683740 | 63683920 | -2.709806486 | RTEL1-TNFRSF6B |
| 20 | 63686356 | 63686578 | -2.782001441 | RTEL1 |
| 20 | 63686356 | 63686578 | -2.782001441 | RTEL1-TNFRSF6B |
| 20 | 63692617 | 63692802 | -8.152460835 | RTEL1-TNFRSF6B |
| 20 | 63692617 | 63692802 | -8.152460835 | RTEL1 |
| 20 | 63708746 | 63708973 | 1.209084827 | RP4-583P15.15 |
| 20 | 63708746 | 63708973 | 1.209084827 | ZGPAT |
| 1 | 151165524 | 151165662 | 1.075844919 | LYSMD1 |
| 1 | 151165524 | 151165662 | 1.075844919 | SCNM1 |
| 20 | 63735762 | 63735952 | 1.829724573 | RP4-583P15.15 |
| 20 | 63735762 | 63735952 | 1.829724573 | ZGPAT |
| 20 | 63743657 | 63743852 | 6.642628855 | ZBTB46 |
| 20 | 63746987 | 63747195 | 11.06635309 | ZBTB46 |
| 20 | 63770421 | 63770588 | -2.247098518 | ZBTB46 |
| 20 | 63774902 | 63775053 | 1.047354934 | ZBTB46 |
| 20 | 63861361 | 63861592 | 1.776616411 | RP4-591C20.9 |
| 20 | 63861361 | 63861592 | 1.776616411 | ABHD16B |
| 1 | 151281939 | 151282135 | 1.009678259 | ZNF687 |
| 20 | 63862153 | 63862257 | 10.43296554 | ABHD16B |
| 20 | 63862153 | 63862257 | 10.43296554 | RP4-591C20.9 |
| 20 | 63865527 | 63865728 | 9.206009917 | TPD52L2 |
| 20 | 63919980 | 63920182 | 1.690294208 | DNAJC5 |
| 20 | 63959983 | 63960176 | 4.460309078 | ZNF512B |
| 20 | 64063476 | 64063666 | 10.13730112 | TCEA2 |
| 21 | 5122387 | 5122574 | -2.469988112 | CH507-9B2.3 |
| 21 | 5122387 | 5122574 | -2.469988112 | CH507-9B2.4 |
| 21 | 5125654 | 5125814 | -1.376874208 | CH507-9B2.4 |
| 21 | 5125654 | 5125814 | -1.376874208 | CH507-9B2.3 |
| 21 | 5141734 | 5141914 | -2.836212671 | CH507-9B2.5 |
| 21 | 9809677 | 9809865 | -5.018086806 | MGC39584 |
| 21 | 10328797 | 10328992 | -1.111482549 | AP003900.6 |
| 21 | 14319936 | 14320131 | 2.57655804 | ABCC13 |
| 21 | 14329374 | 14329543 | 2.690410892 | ABCC13 |
| 21 | 24488894 | 24488994 | -1.46857634 | AP000476.1 |
| 21 | 25035245 | 25035406 | -2.136731303 | AP000235.2 |
| 21 | 26837627 | 26837824 | 2.879554428 | ADAMTS1 |
| 21 | 26844995 | 26845246 | 3.065896366 | ADAMTS1 |
| 21 | 26924417 | 26924543 | 4.669925001 | ADAMTS5 |
| 21 | 26924417 | 26924543 | 4.669925001 | AP001601.2 |
| 21 | 28875200 | 28875358 | -1.603496164 | N6AMT1 |
| 1 | 153626701 | 153626888 | 2.574324171 | S100A13 |
| 1 | 153626701 | 153626888 | 2.574324171 | RP1-178F15.5 |
| 21 | 29181951 | 29182151 | 1.084582938 | AF124730.4 |
| 21 | 31266314 | 31266434 | 3.316662761 | TIAM1 |
| 21 | 31998957 | 31999146 | 6.741403978 | HUNK |
| 21 | 32314602 | 32314802 | 1.466998461 | URB1 |
| 21 | 32314602 | 32314802 | 1.466998461 | MRAP |

| | | | | |
|---|---|---|---|---|
| 12 | 13002711 | 13002906 | 1.039841139 | RP11-377D9.3 |
| 12 | 13002711 | 13002906 | 1.039841139 | HTR7P1 |
| 12 | 13004247 | 13004414 | 1.880984351 | RP11-377D9.3 |
| 12 | 13004247 | 13004414 | 1.880984351 | HTR7P1 |
| 12 | 19462412 | 19462615 | -1.550483815 | AEBP2 |
| 12 | 20369052 | 20369203 | 4.488639962 | RP11-284H19.1 |
| 12 | 20680087 | 20680286 | 7.536568692 | PDE3A |
| 12 | 21515736 | 21515860 | 2.349188815 | GOLT1B |
| 12 | 23534053 | 23534439 | 1.015671894 | SOX5 |
| 12 | 23820795 | 23820931 | 5.093676037 | SOX5 |
| 12 | 24584079 | 24584225 | 3.853679566 | LINC00477 |
| 12 | 24584079 | 24584225 | 3.853679566 | KNOP1P1 |
| 12 | 25945831 | 25946035 | 3.060426395 | RASSF8-AS1 |
| 12 | 25958107 | 25958304 | 6.879944092 | RASSF8-AS1 |
| 12 | 26065057 | 26065251 | 2.110606669 | RASSF8 |
| 12 | 26068754 | 26068949 | 2.284498303 | RASSF8 |
| 12 | 26964557 | 26964755 | 1.224350323 | FGFR1OP2 |
| 12 | 27042902 | 27043022 | -1.770904649 | MED21 |
| 12 | 27160462 | 27160657 | -6.488639962 | RP11-582E3.2 |
| 12 | 27322342 | 27322542 | 2.182676116 | STK38L |
| 12 | 27406787 | 27406990 | -6.912876916 | ARNTL2 |
| 12 | 27406787 | 27406990 | -6.912876916 | ARNTL2-AS1 |
| 12 | 27648463 | 27648663 | -4.953445298 | PPFIBP1 |
| 12 | 27780895 | 27781099 | -1.661049107 | RP11-860B13.1 |
| 12 | 27780895 | 27781099 | -1.661049107 | KLHL42 |
| 12 | 29280936 | 29281106 | -6.164617871 | FAR2 |
| 12 | 29280936 | 29281106 | -6.164617871 | RP11-996F15.2 |
| 12 | 29325620 | 29325755 | 2.032059724 | FAR2 |
| 12 | 29326554 | 29326762 | 1.273076174 | FAR2 |
| 12 | 29404181 | 29404323 | 1.053623623 | OVCH1-AS1 |
| 1 | 32826514 | 32826727 | -1.293446601 | S100PBP |
| 12 | 29407291 | 29407446 | 1.211946123 | OVCH1-AS1 |
| 12 | 29410081 | 29410280 | 1.517922599 | OVCH1-AS1 |
| 12 | 29411736 | 29411882 | 1.548673248 | OVCH1-AS1 |
| 12 | 29506627 | 29506830 | 2.789444313 | TMTC1 |
| 12 | 29584585 | 29584776 | -5.471257253 | TMTC1 |
| 12 | 29585423 | 29585613 | 7.219136028 | TMTC1 |
| 12 | 30710274 | 30710392 | 4.679966269 | CAPRIN2 |
| 12 | 30728779 | 30728978 | 2.047824257 | CAPRIN2 |
| 12 | 31387425 | 31387630 | 2.088422793 | DENND5B |
| 12 | 31447667 | 31447865 | 1.593010336 | DENND5B |
| 1 | 32936971 | 32937199 | 1.409475409 | RNF19B |
| 12 | 31479968 | 31480138 | 1.303626426 | DENND5B |
| 12 | 31550341 | 31550537 | 3.857633737 | DENND5B |
| 12 | 31551053 | 31551199 | 2.137279271 | DENND5B |
| 12 | 31553177 | 31553376 | 1.967133517 | DENND5B |
| 12 | 31981166 | 31981359 | 1.463005874 | KIAA1551 |
| 12 | 31982778 | 31982971 | 1.599706257 | KIAA1551 |
| 12 | 31983661 | 31983861 | 1.190943164 | KIAA1551 |
| 12 | 31985559 | 31985859 | 1.463390537 | KIAA1551 |
| 1 | 33008123 | 33008338 | -2.733137726 | RP1-117O3.2 |
| 1 | 33008123 | 33008338 | -2.733137726 | AK2 |
| 12 | 32305830 | 32306030 | 1.269154413 | BICD1 |
| 12 | 32377653 | 32377852 | 2.990818895 | BICD1 |
| 13 | 93227188 | 93227311 | 3.318205689 | GPC6 |
| 13 | 93227188 | 93227311 | 3.318205689 | RP11-632L2.2 |
| 13 | 95676458 | 95676672 | -1.483845233 | DNAJC3-AS1 |
| 13 | 99255320 | 99255502 | -6.20918986 | UBAC2 |
| 13 | 99255320 | 99255502 | -6.20918986 | GPR18 |
| 13 | 99295643 | 99295972 | -1.172937177 | UBAC2 |
| 13 | 99295643 | 99295972 | -1.172937177 | GPR183 |
| 13 | 99501038 | 99501227 | -4.418742294 | RP11-214F16.8 |
| 13 | 99501038 | 99501227 | -4.418742294 | LINC00449 |
| 13 | 99700357 | 99700550 | -2.433758179 | CLYBL |
| 13 | 99965388 | 99965589 | -1.614180047 | ZIC5 |
| 13 | 100326383 | 100326582 | -1.083852657 | PCCA |
| 13 | 100358000 | 100358185 | -6.526284025 | PCCA |
| 1 | 39771404 | 39771619 | -1.170887907 | BMP8B |
| 13 | 102772510 | 102772734 | -1.715700197 | TEX30 |
| 13 | 102772992 | 102773165 | 6.097993649 | TEX30 |
| 13 | 102807384 | 102807575 | -2.073755112 | BIVM |
| 13 | 102807384 | 102807575 | -2.073755112 | BIVM-ERCC5 |
| 13 | 102839721 | 102839936 | -1.482144949 | BIVM-ERCC5 |
| 13 | 102839721 | 102839936 | -1.482144949 | BIVM |
| 13 | 102875587 | 102875787 | -1.145357129 | ERCC5 |
| 13 | 106492813 | 106492993 | 3.065243114 | EFNB2 |
| 13 | 109785219 | 109785420 | 10.76322216 | IRS2 |
| 13 | 110614133 | 110614303 | 5.635929499 | RP11-90L1.8 |
| 13 | 110892993 | 110893194 | -1.742524637 | ANKRD10 |
| 13 | 111303087 | 111303307 | -1.078696521 | ARHGEF7 |
| 13 | 112687308 | 112687487 | -5.518086806 | RP11-88E10.5 |
| 13 | 112688419 | 112688648 | -1.083857664 | RP11-88E10.5 |
| 1 | 799925 | 800198 | -1.549698887 | RP5-857K21.4 |
| 1 | 799925 | 800198 | -1.549698887 | RP11-206L10.9 |
| 1 | 39869852 | 39870046 | 9.061742451 | TRIT1 |
| 13 | 113413737 | 113413931 | -1.302512329 | ADPRHL1 |
| 13 | 113416237 | 113416403 | -3.205543613 | ADPRHL1 |
| 13 | 113456367 | 113456575 | -1.115327014 | DCUN1D2 |
| 13 | 113483931 | 113484116 | -1.097882503 | DCUN1D2 |
| 13 | 113816679 | 113816812 | -1.29248125 | GAS6-AS1 |
| 13 | 113816679 | 113816812 | -1.29248125 | TMEM255B |
| 1 | 39901403 | 39901540 | -7.097993649 | MYCL |
| 13 | 113817848 | 113818102 | -1.047359329 | GAS6-AS1 |
| 13 | 113818813 | 113818947 | -2.350219859 | GAS6-AS1 |
| 13 | 113832019 | 113832212 | -5.919995535 | GAS6-AS1 |
| 13 | 113832019 | 113832212 | -5.919995535 | GAS6 |
| 13 | 113842829 | 113843020 | -2.865255286 | GAS6-AS1 |
| 13 | 113842829 | 113843020 | -2.865255286 | GAS6 |
| 13 | 113844092 | 113844420 | -1.121302821 | GAS6-AS1 |
| 13 | 113844092 | 113844420 | -1.121302821 | GAS6 |
| 13 | 113844754 | 113844858 | -3.824263815 | GAS6-AS1 |
| 13 | 113844754 | 113844858 | -3.824263815 | GAS6 |
| 13 | 113927957 | 113928126 | -6.148314737 | LINC00565 |
| 1 | 39962665 | 39962862 | -8.439097319 | RP3-342P20.2 |
| 1 | 39962665 | 39962862 | -8.439097319 | MFSD2A |
| 13 | 113953900 | 113954105 | -1.068994515 | ABC13-47488600E17.1 |
| 13 | 113954447 | 113954642 | -1.44583044 | ABC13-47488600E17.1 |
| 13 | 113956509 | 113956703 | -1.982242054 | ABC13-47488600E17.1 |
| 21 | 32316630 | 32316867 | 1.648934345 | URB1 |
| 21 | 32347437 | 32347641 | 1.035358043 | URB1 |
| 1 | 153774482 | 153774807 | 2.190322397 | SLC27A3 |
| 1 | 153774482 | 153774807 | 2.190322397 | INTS3 |
| 21 | 32759739 | 32759946 | -1.313516888 | PAXBP1 |
| 21 | 33266600 | 33266800 | 1.222880499 | AP000295.9 |
| 21 | 33266600 | 33266800 | 1.222880499 | IL10RB |
| 21 | 33285796 | 33285993 | 1.582850208 | IL10RB |
| 21 | 33309108 | 33309297 | -1.328832496 | IL10RB |
| 1 | 153775705 | 153775899 | 9.605592435 | SLC27A3 |
| 21 | 33925886 | 33926091 | 1.384837676 | AP000304.12 |
| 21 | 33925886 | 33926091 | 1.384837676 | LINC00649 |
| 21 | 33976984 | 33977184 | 1.471972806 | LINC00649 |
| 21 | 34095019 | 34095218 | 1.831585734 | SLC5A3 |
| 21 | 34095019 | 34095218 | 1.831585734 | AP000320.7 |
| 21 | 34095019 | 34095218 | 1.831585734 | MRPS6 |
| 21 | 34096254 | 34096406 | 1.756810133 | AP000320.7 |
| 21 | 34096254 | 34096406 | 1.756810133 | MRPS6 |
| 21 | 34096254 | 34096406 | 1.756810133 | SLC5A3 |
| 21 | 34097117 | 34097317 | 1.456825629 | SLC5A3 |
| 21 | 34097117 | 34097317 | 1.456825629 | MRPS6 |
| 21 | 34097117 | 34097317 | 1.456825629 | AP000320.7 |
| 21 | 34182869 | 34183054 | 1.901834938 | LINC00310 |
| 21 | 34182869 | 34183054 | 1.901834938 | AP000320.7 |
| 21 | 34449265 | 34449398 | -1.711011352 | KCNE1 |
| 21 | 34496826 | 34497026 | -1.786991023 | KCNE1 |
| 21 | 34498411 | 34498536 | -1.782184936 | KCNE1 |
| 21 | 34520638 | 34520774 | 2.032449887 | RCAN1 |
| 21 | 36319973 | 36320094 | 4.672147954 | AP000692.10 |
| 1 | 153852057 | 153852259 | -1.640191473 | RP11-216N14.7 |
| 1 | 153852057 | 153852259 | -1.640191473 | GATAD2B |
| 21 | 37143865 | 37144065 | 5.552954254 | TTC3 |
| 21 | 37241223 | 37241519 | 6.473818469 | DSCR3 |
| 21 | 37267799 | 37267997 | 7.098761939 | DSCR3 |
| 21 | 37267799 | 37267997 | 7.098761939 | AP001412.1 |
| 1 | 153852936 | 153853136 | -8.708761457 | RP11-216N14.7 |
| 1 | 153852936 | 153853136 | -8.708761457 | GATAD2B |
| 21 | 38989170 | 38989364 | 1.576001547 | AF064858.8 |
| 21 | 39197348 | 39197507 | -1.163210251 | BRWD1 |
| 21 | 41377009 | 41377197 | 7.419995535 | MX2 |
| 21 | 41377542 | 41377730 | 2.138099433 | MX2 |
| 21 | 41390109 | 41390283 | -5.596146407 | MX2 |
| 1 | 153963561 | 153963759 | 2.761278422 | SLC39A1 |
| 21 | 42849327 | 42849530 | 1.483757776 | WDR4 |
| 1 | 154271037 | 154271236 | 1.792216421 | UBAP2L |
| 21 | 43724314 | 43724506 | -8.206813965 | PDXK |
| 21 | 43738563 | 43738762 | -7.469605941 | PDXK |
| 21 | 43809115 | 43809235 | -1.927707189 | AATBC |
| 21 | 43966755 | 43967022 | 5.957939689 | AGPAT3 |
| 21 | 44137708 | 44137883 | -3.660534546 | C21orf33 |
| 21 | 44233649 | 44233848 | 3.321928095 | ICOSLG |
| 21 | 44330310 | 44330557 | 1.756527286 | C21orf2 |
| 21 | 44351164 | 44351341 | 6.582139219 | TRPM2 |
| 21 | 44358733 | 44358866 | -1.518412298 | TRPM2 |

| | | | | |
|---|---|---|---|---|
| 12 | 32582066 | 32582325 | 2.730244185 | FGD4 |
| 12 | 32640444 | 32640652 | 2.970058906 | FGD4 |
| 12 | 34026033 | 34026232 | -1.314611901 | ALG10 |
| 12 | 34026033 | 34026232 | -1.314611901 | RP11-847H18.2 |
| 1 | 33050779 | 33050997 | 1.884356829 | AK2 |
| 12 | 38906108 | 38906294 | -1.102560705 | CPNE8 |
| 12 | 38906406 | 38906579 | -1.130437752 | CPNE8 |
| 12 | 38906406 | 38906579 | -1.130437752 | RP11-396F22.1 |
| 12 | 39619549 | 39619697 | -3.780484823 | ABCD2 |
| 1 | 33054328 | 33054509 | 2.684727137 | AK2 |
| 12 | 42460159 | 42460359 | -2.761936322 | PRICKLE1 |
| 12 | 42460159 | 42460359 | -2.761936322 | RP11-328C8.4 |
| 12 | 42464511 | 42464663 | -2.711312803 | RP11-328C8.4 |
| 12 | 42464511 | 42464663 | -2.711312803 | PRICKLE1 |
| 12 | 43956951 | 43957138 | -1.004508437 | TMEM117 |
| 12 | 44388016 | 44388215 | 1.493963584 | TMEM117 |
| 1 | 1313234 | 1313497 | -2.560599071 | CPSF3L |
| 1 | 33056541 | 33056740 | 2.750671007 | AK2 |
| 12 | 45394122 | 45394240 | -7.117595763 | ANO6 |
| 12 | 45726971 | 45727153 | -1.000851502 | RP11-352M15.2 |
| 12 | 46399133 | 46399329 | -1.973887076 | RP11-96H19.1 |
| 12 | 46399887 | 46400061 | -1.592579758 | RP11-96H19.1 |
| 12 | 46400445 | 46400644 | -3.774969563 | RP11-96H19.1 |
| 12 | 46604848 | 46604995 | -7.690298173 | RP11-446N19.1 |
| 12 | 46604848 | 46604995 | -7.690298173 | RP11-96H19.1 |
| 12 | 46605153 | 46605336 | -2.476240356 | RP11-446N19.1 |
| 12 | 46605153 | 46605336 | -2.476240356 | RP11-96H19.1 |
| 1 | 33324717 | 33324938 | 1.856993387 | PHC2 |
| 1 | 33324717 | 33324938 | 1.856993387 | RP11-415J8.3 |
| 12 | 46637207 | 46637405 | -2.501345164 | RP11-446N19.1 |
| 12 | 46637207 | 46637405 | -2.501345164 | RP11-96H19.1 |
| 12 | 46643600 | 46643708 | -7.917963517 | RP11-96H19.1 |
| 12 | 46643600 | 46643708 | -7.917963517 | RP11-446N19.1 |
| 12 | 47078075 | 47078224 | 3.109221074 | AMIGO2 |
| 12 | 47078680 | 47078913 | 2.747288687 | AMIGO2 |
| 12 | 47235108 | 47235318 | 2.229665901 | PCED1B |
| 12 | 47235674 | 47235920 | 1.490988481 | PCED1B |
| 12 | 47236250 | 47236467 | 1.36867875 | PCED1B |
| 12 | 47737010 | 47737208 | 2.556136207 | RP1-197B17.3 |
| 12 | 47737010 | 47737208 | 2.556136207 | RAPGEF3 |
| 12 | 47737341 | 47737463 | 2.225650522 | RAPGEF3 |
| 12 | 47737341 | 47737463 | 2.225650522 | RP1-197B17.3 |
| 12 | 48342929 | 48343108 | -1.199861262 | ZNF641 |
| 12 | 48815211 | 48815408 | 2.906534287 | CACNB3 |
| 12 | 48830033 | 48830233 | -1.744914387 | DDX23 |
| 12 | 48998009 | 48998224 | 3.941456199 | DDN |
| 12 | 49031365 | 49031656 | -1.241529072 | KMT2D |
| 12 | 49038177 | 49038376 | -1.520762003 | KMT2D |
| 12 | 49050066 | 49050229 | -1.024756978 | KMT2D |
| 12 | 49233311 | 49233520 | -5.302934678 | RP11-977B10.2 |
| 12 | 49233311 | 49233520 | -5.302934678 | TUBA1C |
| 12 | 49330219 | 49330413 | -2.926069704 | TROAP |
| 12 | 49625606 | 49625945 | 7.450291912 | PRPF40B |
| 12 | 49642519 | 49642767 | 2.968768174 | FMNL3 |
| 1 | 39963123 | 39963323 | 8.653951705 | MFSD2A |
| 1 | 39963123 | 39963323 | 8.653951705 | RP3-342P20.2 |
| 13 | 114116230 | 114116383 | 6.84821212 | RASA3 |
| 13 | 114304784 | 114304988 | -1.403131441 | UPF3A |
| 13 | 114323903 | 114324196 | -1.709960942 | CHAMP1 |
| 14 | 19097103 | 19097356 | -1.233997319 | DUXAP9 |
| 14 | 19300320 | 19300517 | -4.750918592 | AL589743.1 |
| 14 | 19300320 | 19300517 | -4.750918592 | LINC01296 |
| 14 | 19316403 | 19316587 | 1.550689425 | LINC01296 |
| 14 | 19316403 | 19316587 | 1.550689425 | BMS1P17 |
| 14 | 21074679 | 21074946 | -1.210380839 | ARHGEF40 |
| 14 | 22812995 | 22813195 | -1.824824487 | SLC7A7 |
| 14 | 22973218 | 22973471 | -2.666931146 | AJUBA |
| 14 | 22973218 | 22973471 | -2.666931146 | RP11-298I3.5 |
| 14 | 22982864 | 22983065 | -5.61928611 | RP11-298I3.4 |
| 14 | 22987276 | 22987496 | -1.078550481 | RP11-298I3.4 |
| 14 | 22987276 | 22987496 | -1.078550481 | C14orf93 |
| 14 | 23117383 | 23117579 | 4.281856906 | CEBPE |
| 14 | 23320487 | 23320686 | 1.525323222 | BCL2L2-PABPN1 |
| 14 | 23520941 | 23521131 | 1.027796293 | RP11-66N24.4 |
| 14 | 23520941 | 23521131 | 1.027796293 | ZFHX2 |
| 1 | 40257587 | 40257844 | -1.428001826 | RP1-39G22.7 |
| 14 | 23940418 | 23940750 | -2.450439395 | DHRS4-AS1 |
| 14 | 24103747 | 24103945 | -2.564232853 | PCK2 |
| 14 | 24103747 | 24103945 | -2.564232853 | NRL |
| 14 | 24213015 | 24213217 | -1.472243422 | RP11-468E2.1 |
| 14 | 24213015 | 24213217 | -1.472243422 | AL136419.6 |
| 14 | 24213015 | 24213217 | -1.472243422 | CHMP4A |
| 14 | 24240340 | 24240538 | 1.255923588 | TINF2 |
| 14 | 24427131 | 24427323 | 1.349305196 | CBLN3 |
| 14 | 30576222 | 30576422 | -1.540044557 | G2E3 |
| 1 | 40301008 | 40301218 | -1.313081773 | COL9A2 |
| 14 | 31026847 | 31027017 | 1.0801118 | AP4S1 |
| 14 | 31127902 | 31128080 | -6.91841968 | HECTD1 |
| 14 | 31690881 | 31691069 | -1.351834215 | NUBPL |
| 14 | 34518747 | 34518871 | -5.647023157 | EAPP |
| 14 | 34776045 | 34776246 | 1.226936723 | BAZ1A |
| 14 | 37591318 | 37591546 | 1.136083001 | FOXA1 |
| 14 | 39387175 | 39387364 | -1.536099499 | CTAGE5 |
| 14 | 41607789 | 41608031 | 8.314849063 | LRFN5 |
| 14 | 41886792 | 41886931 | 3.328947012 | LRFN5 |
| 14 | 41891661 | 41891895 | 2.950149211 | LRFN5 |
| 14 | 44964110 | 44964310 | -1.158370945 | KLHL28 |
| 14 | 44964110 | 44964310 | -1.158370945 | FAM179B |
| 14 | 45242195 | 45242398 | -1.989231037 | MIS18BP1 |
| 14 | 45246952 | 45247153 | -2.033284332 | MIS18BP1 |
| 14 | 49577524 | 49577748 | -2.162215301 | RPL32P29 |
| 14 | 49577524 | 49577748 | -2.162215301 | RPS29 |
| 14 | 49622154 | 49622355 | 12.66067372 | MGAT2 |
| 14 | 49622154 | 49622355 | 12.66067372 | RP11-649E7.5 |
| 14 | 50007029 | 50007225 | 3.548205539 | LINC01588 |
| 14 | 50118474 | 50118705 | -1.624251396 | SOS2 |
| 14 | 50159908 | 50160110 | -1.704194833 | SOS2 |
| 14 | 50329969 | 50330186 | -1.016167225 | CDKL1 |
| 21 | 44931985 | 44932182 | 3.123514758 | ITGB2 |
| 1 | 154502214 | 154502398 | 7.527895407 | TDRD10 |
| 21 | 44934375 | 44934550 | 1.662601223 | LINC01547 |
| 21 | 44976809 | 44976964 | 2.178922603 | FAM207A |
| 21 | 45072808 | 45073007 | 1.343662109 | SSR4P1 |
| 21 | 45073072 | 45073220 | 2.360802009 | SSR4P1 |
| 21 | 45162625 | 45162844 | 3.758064539 | ADARB1 |
| 21 | 45192603 | 45192802 | 1.465941886 | ADARB1 |
| 21 | 45222098 | 45222293 | 1.111643427 | ADARB1 |
| 21 | 45515838 | 45516037 | 3.769941158 | SLC19A1 |
| 21 | 45600792 | 45600962 | -4.974183616 | AL133493.2 |
| 21 | 45793389 | 45793583 | 3.196158711 | PCBP3 |
| 21 | 45852791 | 45852991 | 1.833557417 | PCBP3 |
| 21 | 46004663 | 46004867 | 2.904378265 | COL6A1 |
| 21 | 46129313 | 46129463 | 5.320371621 | COL6A2 |
| 21 | 46132383 | 46132557 | 11.47210026 | COL6A2 |
| 21 | 46291382 | 46291589 | 1.998658061 | YBEY |
| 21 | 46323065 | 46323268 | 1.006830408 | C21orf58 |
| 22 | 15821541 | 15821737 | 1.367077363 | DUXAP8 |
| 22 | 17548383 | 17548506 | -1.505912718 | CECR2 |
| 22 | 17552947 | 17553186 | -1.968197131 | CECR2 |
| 22 | 17638260 | 17638439 | -5.689959909 | BCL2L13 |
| 22 | 17777430 | 17777598 | 4.730739724 | LINC00528 |
| 22 | 17778514 | 17778680 | 4.033044595 | LINC00528 |
| 22 | 17818360 | 17818526 | 2.124123527 | MICAL3 |
| 22 | 17829962 | 17830166 | -6.520487299 | MICAL3 |
| 22 | 19432038 | 19432192 | 6.10715956 | C22orf39 |
| 22 | 19432038 | 19432192 | 6.10715956 | HIRA |
| 22 | 19432038 | 19432192 | 6.10715956 | MRPL40 |
| 22 | 19758701 | 19758899 | 4.451295229 | TBX1 |
| 22 | 19763916 | 19764103 | -3.857122759 | TBX1 |
| 22 | 19788550 | 19788710 | 1.03970673 | GNB1L |
| 22 | 19791927 | 19792110 | -1.422354168 | GNB1L |
| 22 | 19802417 | 19802615 | -1.5799887 | GNB1L |
| 1 | 155018235 | 155018414 | -8.241214921 | ZBTB7B |
| 22 | 19804163 | 19804356 | -1.285178084 | GNB1L |
| 22 | 20143248 | 20143419 | 2.309158859 | ZDHHC8 |
| 22 | 20145270 | 20145482 | 1.177427858 | ZDHHC8 |
| 1 | 155062476 | 155062677 | 2.144429584 | ADAM15 |
| 22 | 20241617 | 20241813 | 10.52109219 | RTN4R |
| 22 | 20405499 | 20405721 | 1.485483703 | ZNF74 |
| 22 | 21628210 | 21628409 | 1.253351163 | YDJC |
| 1 | 155338264 | 155338414 | -1.549897798 | ASH1L |
| 22 | 23743407 | 23743622 | -1.135881192 | ZNF70 |
| 22 | 23895845 | 23896033 | 1.428880878 | MIF-AS1 |
| 22 | 24178160 | 24178323 | 1.586519786 | CABIN1 |
| 22 | 25478239 | 25478366 | -4.161923536 | CTA-390C10.10 |
| 22 | 25478239 | 25478366 | -4.161923536 | CRYBB2P1 |
| 22 | 26434191 | 26434369 | -2.96831897 | ASPHD2 |
| 22 | 26464163 | 26464364 | -1.118364186 | HPS4 |
| 22 | 26514057 | 26514312 | -1.37045254 | CTA-445C9.14 |
| 22 | 27800436 | 27800592 | 1.799731035 | MN1 |
| 22 | 27983245 | 27983398 | -1.697904014 | TTC28 |
| 22 | 27983245 | 27983398 | -1.697904014 | TTC28-AS1 |

| | | | | |
|---|---|---|---|---|
| 12 | 49642519 | 49642767 | 2.968768174 | PRPF40B |
| 12 | 49643177 | 49643377 | 3.856676518 | PRPF40B |
| 12 | 49643177 | 49643377 | 3.856676518 | FMNL3 |
| 12 | 50744655 | 50744800 | 2.125946163 | DIP2B |
| 12 | 50925124 | 50925280 | 2.440068263 | METTL7A |
| 12 | 51172854 | 51173057 | -1.711539569 | TFCP2 |
| 12 | 51198518 | 51198713 | 1.537644064 | POU6F1 |
| 12 | 51245722 | 51245922 | 1.713646864 | DAZAP2 |
| 12 | 51245722 | 51245922 | 1.713646864 | SMAGP |
| 12 | 51352021 | 51352251 | 1.423378206 | GALNT6 |
| 12 | 53179798 | 53179980 | 1.000154118 | CSAD |
| 12 | 53500889 | 53501068 | -4.570610086 | TARBP2 |
| 12 | 53500889 | 53501068 | -4.570610086 | RP11-793H13.11 |
| 12 | 54181534 | 54181734 | -2.147392934 | SMUG1 |
| 12 | 54181960 | 54182144 | -1.056979353 | SMUG1 |
| 12 | 54292345 | 54292544 | -5.006052488 | NFE2 |
| 12 | 54292345 | 54292544 | -5.006052488 | RP11-968A15.8 |
| 12 | 54395668 | 54395874 | 1.300447887 | RP11-753H16.5 |
| 12 | 54395668 | 54395874 | 1.300447887 | ITGA5 |
| 12 | 54395668 | 54395874 | 1.300447887 | RP11-753H16.3 |
| 12 | 55724159 | 55724359 | -1.795597227 | RP11-644F5.10 |
| 12 | 55724159 | 55724359 | -1.795597227 | RDH5 |
| 1 | 1314475 | 1314612 | -2.495858776 | CPSF3L |
| 12 | 55730482 | 55730669 | -2.485417454 | RP11-644F5.11 |
| 12 | 55836964 | 55837164 | 7.243218505 | MMP19 |
| 12 | 56117481 | 56117704 | -1.040499975 | RPL41 |
| 12 | 56121349 | 56121596 | -1.510108475 | ZC3H10 |
| 12 | 56121349 | 56121596 | -1.510108475 | RP11-603J24.5 |
| 12 | 56121349 | 56121596 | -1.510108475 | ESYT1 |
| 12 | 56333014 | 56333186 | -1.650869348 | PAN2 |
| 12 | 56343905 | 56344098 | 1.851037966 | STAT2 |
| 12 | 56417571 | 56417715 | -2.967449055 | TIMELESS |
| 12 | 56469145 | 56469351 | -1.162759336 | MIP |
| 12 | 56469145 | 56469351 | -1.162759336 | SPRYD4 |
| 12 | 57091496 | 57091696 | 1.217553868 | NAB2 |
| 12 | 57094828 | 57095032 | 1.408075587 | NAB2 |
| 12 | 57516952 | 57517084 | -5.959990298 | DDIT3 |
| 12 | 57516952 | 57517084 | -5.959990298 | MARS |
| 12 | 57626260 | 57626443 | -2.864168133 | B4GALNT1 |
| 12 | 57626750 | 57626936 | -1.821928095 | B4GALNT1 |
| 12 | 57726324 | 57726534 | -1.742930164 | AGAP2-AS1 |
| 12 | 57726324 | 57726534 | -1.742930164 | AGAP2 |
| 12 | 57796390 | 57796603 | -1.293288007 | TSFM |
| 12 | 57956807 | 57956999 | -4.373242126 | ATP23 |
| 12 | 58872591 | 58872789 | -4.543920825 | LRIG3 |
| 12 | 59779369 | 59779525 | -1.665769845 | SLC16A7 |
| 12 | 62511746 | 62511942 | 1.252192492 | MON2 |
| 12 | 62648366 | 62648569 | 2.259632107 | PPM1H |
| 12 | 64142461 | 64142645 | 1.277605657 | SRGAP1 |
| 1 | 35560423 | 35560625 | -1.534812782 | NCDN |
| 12 | 64193964 | 64194171 | -1.065221276 | C12orf66 |
| 12 | 64716455 | 64716576 | 2.423026095 | GNS |
| 12 | 64876429 | 64876711 | -5.382376569 | TBC1D30 |
| 12 | 65464216 | 65464415 | -4.229049485 | MSRB3 |
| 14 | 50329969 | 50330186 | -1.016167225 | ATP5S |
| 14 | 50661385 | 50661590 | 1.356913052 | SAV1 |
| 14 | 50723552 | 50723759 | -2.444851635 | NIN |
| 14 | 50756663 | 50756863 | -1.308643719 | NIN |
| 14 | 50758124 | 50758326 | -1.59286829 | NIN |
| 14 | 50760077 | 50760274 | -1.872053029 | NIN |
| 1 | 40473208 | 40473425 | -7.853167063 | RP11-656D10.7 |
| 14 | 51066332 | 51066454 | 1.008009628 | TRIM9 |
| 14 | 53270118 | 53270300 | 1.008216145 | AL163953.3 |
| 1 | 40479005 | 40479204 | -3.341532116 | ZFP69 |
| 14 | 53323041 | 53323253 | -1.898063833 | AL163953.3 |
| 14 | 53950559 | 53950758 | -3.558900031 | BMP4 |
| 14 | 54405575 | 54405725 | -5.727149647 | CDKN3 |
| 14 | 54736892 | 54737104 | 11.14636541 | SAMD4A |
| 1 | 40486959 | 40487161 | -3.867728644 | ZFP69 |
| 14 | 54748660 | 54748856 | 1.672986784 | SAMD4A |
| 14 | 55298928 | 55299127 | -2.501851218 | CHMP4BP1 |
| 14 | 55298928 | 55299127 | -2.501851218 | FBXO34 |
| 1 | 40495532 | 40495772 | -1.967489665 | ZFP69 |
| 1 | 40495532 | 40495772 | -1.967489665 | RP11-656D10.3 |
| 14 | 55351432 | 55351626 | -2.13811295 | FBXO34 |
| 14 | 55352365 | 55352520 | -1.490463655 | FBXO34 |
| 14 | 55352619 | 55352801 | -1.282614299 | FBXO34 |
| 14 | 56296859 | 56297060 | -4.351748569 | PELI2 |
| 14 | 57409341 | 57409471 | -1.199892289 | NAA30 |
| 1 | 1354281 | 1354460 | 8.110340311 | MXRA8 |
| 1 | 40514725 | 40514966 | -1.41262735 | RP11-656D10.5 |
| 1 | 40514725 | 40514966 | -1.41262735 | EXO5 |
| 14 | 58266879 | 58267075 | -1.305628873 | PSMA3-AS1 |
| 14 | 58266879 | 58267075 | -1.305628873 | C14orf37 |
| 14 | 58266879 | 58267075 | -1.305628873 | PSMA3 |
| 14 | 58266879 | 58267075 | -1.305628873 | CTD-2002H8.2 |
| 14 | 58364245 | 58364446 | -2.963378469 | ARID4A |
| 14 | 58521660 | 58521854 | 5.609886775 | KIAA0586 |
| 14 | 58521660 | 58521854 | 5.609886775 | HNRNPCP1 |
| 14 | 58646742 | 58646871 | -1.158615654 | DACT1 |
| 14 | 59368636 | 59368881 | -1.211936899 | DAAM1 |
| 14 | 59463590 | 59463801 | 1.43373335 | L3HYPDH |
| 14 | 59463590 | 59463801 | 1.43373335 | GPR135 |
| 1 | 40516857 | 40517044 | -1.099666705 | RP11-656D10.6 |
| 14 | 60282941 | 60283146 | -1.394200241 | PPM1A |
| 14 | 60646250 | 60646448 | 1.060393197 | SIX1 |
| 14 | 60713331 | 60713476 | -1.476552696 | SIX4 |
| 14 | 60975793 | 60976000 | -2.01195971 | RP11-193F5.1 |
| 14 | 60975793 | 60976000 | -2.01195971 | TRMT5 |
| 14 | 60977600 | 60977766 | -2.027340229 | RP11-193F5.1 |
| 14 | 60977600 | 60977766 | -2.027340229 | TRMT5 |
| 14 | 60979578 | 60979798 | -1.897007908 | TRMT5 |
| 14 | 60979578 | 60979798 | -1.897007908 | RP11-193F5.1 |
| 14 | 61012438 | 61012645 | -2.824607302 | SLC38A6 |
| 14 | 61013153 | 61013351 | -4.099414854 | SLC38A6 |
| 14 | 61013805 | 61014005 | -3.061604251 | SLC38A6 |
| 14 | 61014856 | 61015043 | -5.800106777 | SLC38A6 |
| 14 | 61058062 | 61058259 | -6.661386588 | SLC38A6 |
| 22 | 28105602 | 28105792 | -2.028982541 | TTC28 |
| 22 | 28390613 | 28390811 | -7.400803663 | TTC28 |
| 22 | 29049305 | 29049503 | 1.974924787 | ZNRF3 |
| 22 | 29050554 | 29050769 | 1.296594959 | ZNRF3 |
| 22 | 29142187 | 29142314 | -6.257110455 | KREMEN1 |
| 22 | 29312232 | 29312460 | 1.577577894 | GAS2L1 |
| 22 | 29313042 | 29313242 | 1.059219516 | RASL10A |
| 22 | 29469816 | 29469947 | 1.121962791 | RFPL1S |
| 22 | 29490860 | 29491054 | 1.056926128 | NEFH |
| 22 | 30241071 | 30241261 | -4.897207933 | LIF |
| 22 | 30263910 | 30264115 | 1.435497875 | OSM |
| 22 | 30375771 | 30375962 | -2.809412976 | RNF215 |
| 22 | 30375771 | 30375962 | -2.809412976 | CCDC157 |
| 22 | 30668289 | 30668472 | 7.116285413 | SLC35E4 |
| 22 | 30903743 | 30903936 | -1.013740368 | OSBP2 |
| 22 | 31084474 | 31084659 | 2.469583499 | SMTN |
| 22 | 31291861 | 31292038 | 1.80967449 | PIK3IP1 |
| 22 | 31347128 | 31347276 | 3.398182069 | LINC01521 |
| 22 | 31347946 | 31348070 | 2.177630779 | LINC01521 |

| | | | | |
|---|---|---|---|---|
| 12 | 65852936 | 65853130 | 10.4208125 | HMGA2 |
| 12 | 65852936 | 65853130 | 10.4208125 | RP11-366L20.2 |
| 12 | 65920911 | 65921109 | 8.558131363 | HMGA2 |
| 12 | 66123851 | 66124051 | -1.201305176 | RP11-745O10.4 |
| 12 | 66123851 | 66124051 | -1.201305176 | LLPH |
| 12 | 66165544 | 66165736 | -3.806526369 | RP11-745O10.4 |
| 12 | 66165544 | 66165736 | -3.806526369 | TMBIM4 |
| 12 | 66234632 | 66234770 | -3.779410043 | IRAK3 |
| 12 | 66234632 | 66234770 | -3.779410043 | RBMS1P1 |
| 12 | 66247765 | 66247969 | -2.745594376 | IRAK3 |
| 12 | 66310158 | 66310344 | -2.865714495 | HELB |
| 12 | 66331307 | 66331516 | -1.355620099 | HELB |
| 12 | 66338066 | 66338271 | -2.49734832 | HELB |
| 1 | 35594547 | 35594747 | -2.188749942 | TFAP2E |
| 12 | 67305361 | 67305565 | -1.576980253 | CAND1 |
| 12 | 68957447 | 68957637 | -3.374293294 | PRELID2P1 |
| 12 | 68957447 | 68957637 | -3.374293294 | CPM |
| 1 | 35715768 | 35715988 | -1.427419382 | C1orf216 |
| 12 | 70182691 | 70182833 | 1.814906621 | LINC01481 |
| 12 | 70182691 | 70182833 | 1.814906621 | RP11-320P7.2 |
| 12 | 70183386 | 70183534 | 2.47588352 | RP11-320P7.2 |
| 12 | 70183386 | 70183534 | 2.47588352 | LINC01481 |
| 12 | 70556387 | 70556579 | 1.469034558 | PTPRB |
| 1 | 1315006 | 1315210 | -2.600183107 | CPSF3L |
| 1 | 35720766 | 35720966 | -4.704695468 | CLSPN |
| 12 | 70569827 | 70570030 | 1.864519339 | PTPRB |
| 12 | 70571188 | 70571369 | 1.086532354 | PTPRB |
| 12 | 70571188 | 70571369 | 1.086532354 | RP11-785D18.3 |
| 12 | 71632060 | 71632334 | -1.280714202 | ZFC3H1 |
| 12 | 71663121 | 71663308 | -1.511953576 | THAP2 |
| 12 | 71663121 | 71663308 | -1.511953576 | ZFC3H1 |
| 1 | 35753831 | 35754031 | -3.927274479 | CLSPN |
| 12 | 75360216 | 75360392 | 3.827318014 | GLIPR1L1 |
| 12 | 75360216 | 75360392 | 3.827318014 | CAPS2 |
| 12 | 76029934 | 76030143 | 7.455256541 | PHLDA1 |
| 12 | 76355767 | 76355964 | 2.440661111 | OSBPL8 |
| 1 | 35760438 | 35760639 | -4.208939179 | CLSPN |
| 12 | 78119517 | 78119715 | -3.476028325 | NAV3 |
| 12 | 82423336 | 82423510 | 1.878364424 | METTL25 |
| 12 | 82857279 | 82857477 | -1.330108383 | TMTC2 |
| 1 | 35764360 | 35764525 | -4.063785064 | CLSPN |
| 12 | 85805179 | 85805362 | -1.335219829 | RASSF9 |
| 12 | 89349291 | 89349513 | 3.97605321 | DUSP6 |
| 12 | 89350720 | 89350919 | 3.657115356 | DUSP6 |
| 12 | 89523737 | 89523843 | 1.071227476 | GALNT4 |
| 12 | 89523737 | 89523843 | 1.071227476 | POC1B |
| 12 | 89523737 | 89523843 | 1.071227476 | POC1B-GALNT4 |
| 12 | 92142710 | 92142903 | 1.365460418 | BTG1 |
| 12 | 92142710 | 92142903 | 1.365460418 | LINC01619 |
| 12 | 92144180 | 92144378 | 2.85294864 | BTG1 |
| 12 | 95209866 | 95209975 | 5.923920178 | FGD6 |
| 12 | 96249076 | 96249271 | 2.418124513 | ELK3 |
| 12 | 96259659 | 96259857 | 1.588885901 | ELK3 |
| 12 | 96280232 | 96280468 | 1.017918624 | CDK17 |
| 14 | 61061478 | 61061609 | -2.291144183 | SLC38A6 |
| 14 | 61549845 | 61550046 | -2.98045009 | RP11-47I22.4 |
| 14 | 61549845 | 61550046 | -2.98045009 | PRKCH |
| 14 | 61708301 | 61708485 | 7.299665871 | HIF1A |
| 14 | 64469693 | 64469889 | -1.11002424 | AKAP5 |
| 14 | 64469693 | 64469889 | -1.11002424 | ZBTB25 |
| 14 | 64487323 | 64487504 | -3.037477927 | ZBTB25 |
| 14 | 64522221 | 64522430 | -1.22384161 | ZBTB1 |
| 14 | 64522221 | 64522430 | -1.22384161 | RP11-973N13.4 |
| 1 | 40531588 | 40531838 | -1.955366689 | ZNF684 |
| 14 | 65061449 | 65061641 | -2.221391356 | CHURC1-FNTB |
| 14 | 65061449 | 65061641 | -2.221391356 | MAX |
| 14 | 65061449 | 65061641 | -2.221391356 | FNTB |
| 14 | 65561424 | 65561628 | -1.405088843 | FUT8 |
| 14 | 65615848 | 65616046 | -1.856226118 | FUT8 |
| 14 | 65633099 | 65633233 | -6.509013133 | FUT8 |
| 14 | 65742506 | 65742716 | -1.983557782 | FUT8 |
| 14 | 67273476 | 67273675 | -6.262148419 | MPP5 |
| 14 | 67274154 | 67274325 | -6.300188777 | MPP5 |
| 14 | 67327847 | 67328050 | -1.145019458 | MPP5 |
| 14 | 67327847 | 67328050 | -1.145019458 | ATP6V1D |
| 14 | 67328524 | 67328681 | -1.955666795 | ATP6V1D |
| 14 | 67328524 | 67328681 | -1.955666795 | MPP5 |
| 14 | 67678219 | 67678422 | -2.148473882 | RDH11 |
| 14 | 67678219 | 67678422 | -2.148473882 | RP11-1012A1.4 |
| 14 | 67727695 | 67727799 | -2.204695468 | RDH12 |
| 14 | 67727695 | 67727799 | -2.204695468 | ZFYVE26 |
| 14 | 68679714 | 68679908 | -3.497176718 | RAD51B |
| 14 | 68789524 | 68789729 | -1.398362073 | ZFP36L1 |
| 14 | 69241086 | 69241257 | -1.505613628 | EXD2 |
| 14 | 69241086 | 69241257 | -1.505613628 | RP11-363J20.2 |
| 14 | 69708803 | 69709004 | 1.571331154 | SUSD6 |
| 14 | 70730351 | 70730550 | 2.203228137 | MAP3K9 |
| 14 | 70988423 | 70988636 | 1.572824989 | PCNX1 |
| 14 | 71060987 | 71061225 | 1.069013642 | PCNX1 |
| 14 | 71061736 | 71061877 | 2.623582183 | PCNX1 |
| 14 | 72671063 | 72671182 | -2.072309083 | DPF3 |
| 14 | 73058402 | 73058534 | -4.594912279 | RBM25 |
| 14 | 73058402 | 73058534 | -4.594912279 | RP11-109N23.4 |
| 14 | 73239507 | 73239713 | -1.187589881 | PAPLN |
| 14 | 73245013 | 73245187 | -2.063877774 | PAPLN |
| 14 | 73245013 | 73245187 | -2.063877774 | RP4-647C14.2 |
| 14 | 73254178 | 73254375 | 8.032371382 | PAPLN |
| 14 | 73274279 | 73274450 | -1.732695203 | PAPLN |
| 14 | 73276504 | 73276703 | 1.223334302 | NUMB |
| 14 | 73543133 | 73543307 | -1.755335555 | HEATR4 |
| 14 | 73543133 | 73543307 | -1.755335555 | ACOT1 |
| 1 | 41165762 | 41165964 | -1.729586829 | SCMH1 |
| 14 | 73721377 | 73721553 | -8.822209816 | ELMSAN1 |
| 14 | 74802417 | 74802614 | -4.541264624 | YLPM1 |
| 14 | 75046924 | 75047129 | -1.779458237 | MLH3 |
| 14 | 75049235 | 75049424 | -1.421423757 | MLH3 |
| 14 | 75084444 | 75084644 | -1.183898873 | NEK9 |
| 14 | 75281058 | 75281248 | -1.973479836 | FOS |

| | | | | |
|---|---|---|---|---|
| 12 | 98547891 | 98548112 | -1.97224705 | TMPO |
| 12 | 98613777 | 98613985 | -1.557304586 | IKBIP |
| 12 | 98626398 | 98626696 | -2.708789098 | IKBIP |
| 12 | 100586445 | 100586638 | 3.028604627 | GAS2L3 |
| 12 | 101764930 | 101765136 | -1.053369425 | GNPTAB |
| 12 | 101921990 | 101922177 | -1.397860111 | DRAM1 |
| 12 | 102133971 | 102134167 | -7.815957976 | PARPBP |
| 12 | 103777748 | 103777937 | -2.411807148 | NT5DC3 |
| 12 | 104304216 | 104304413 | 1.605490199 | TXNRD1 |
| 12 | 104304216 | 104304413 | 1.605490199 | EID3 |
| 12 | 104348360 | 104348554 | -9.3234516 | TXNRD1 |
| 12 | 104757442 | 104757645 | 1.247089469 | CHST11 |
| 12 | 104805090 | 104805284 | 7.400803663 | SLC41A2 |
| 12 | 105166937 | 105167151 | 1.146916018 | KIAA1033 |
| 12 | 106066779 | 106066915 | -1.042963504 | NUAK1 |
| 12 | 106067284 | 106067479 | -1.197015249 | NUAK1 |
| 12 | 106346429 | 106346628 | 4.36690394 | TCP11L2 |
| 12 | 106509516 | 106509778 | -2.189809802 | RP11-144F15.1 |
| 12 | 106509516 | 106509778 | -2.189809802 | POLR3B |
| 12 | 106814988 | 106815192 | -2.035387814 | RIC8B |
| 12 | 106825598 | 106825791 | -2.12311539 | RIC8B |
| 12 | 106971167 | 106971321 | 1.225290385 | TMEM263 |
| 12 | 106978430 | 106978610 | -2.312983481 | TMEM263 |
| 12 | 106978430 | 106978610 | -2.312983481 | MTERF2 |
| 12 | 108523245 | 108523448 | -1.490711166 | SART3 |
| 12 | 108523245 | 108523448 | -1.490711166 | FICD |
| 12 | 108622478 | 108622626 | -3.558630602 | SELPLG |
| 12 | 108623653 | 108623883 | -2.866833072 | SELPLG |
| 12 | 109053735 | 109053919 | 3.660964047 | USP30-AS1 |
| 12 | 109053735 | 109053919 | 3.660964047 | USP30 |
| 1 | 1315852 | 1316048 | -2.819923041 | CPSF3L |
| 12 | 109092650 | 109092895 | -1.127771779 | ALKBH2 |
| 12 | 109267599 | 109267794 | -5.704515109 | ACACB |
| 12 | 109769177 | 109769287 | -2.503965301 | FAM222A |
| 12 | 109769177 | 109769287 | -2.503965301 | FAM222A-AS1 |
| 12 | 109769303 | 109769469 | -2.130102922 | FAM222A-AS1 |
| 12 | 109769303 | 109769469 | -2.130102922 | FAM222A |
| 1 | 36137038 | 36137244 | -1.326439022 | TRAPPC3 |
| 12 | 110373988 | 110374188 | -1.552269925 | ANAPC7 |
| 12 | 110470036 | 110470236 | 1.939346852 | FAM216A |
| 12 | 110502478 | 110502710 | -2.226201763 | RAD9B |
| 12 | 110502478 | 110502710 | -2.226201763 | RP3-424M6.4 |
| 12 | 111347738 | 111347863 | 2.049225964 | CUX2 |
| 12 | 111362513 | 111362714 | 1.983035347 | FAM109A |
| 12 | 111448167 | 111448373 | 1.57754986 | SH2B3 |
| 12 | 111813528 | 111813709 | -3.737285621 | ALDH2 |
| 12 | 111898601 | 111898813 | -1.466857452 | MAPKAPK5 |
| 12 | 111900015 | 111900210 | -1.346999508 | MAPKAPK5 |
| 12 | 111900015 | 111900210 | -1.346999508 | ADAM1A |
| 12 | 112010669 | 112010846 | -7.343851865 | TMEM116 |
| 12 | 112022339 | 112022540 | -2.079439308 | ERP29 |
| 12 | 112185015 | 112185228 | -1.626497379 | HECTD4 |
| 12 | 112505740 | 112505950 | -1.409297509 | PTPN11 |
| 12 | 113009218 | 113009422 | 11.61348858 | OAS2 |

| | | | | |
|---|---|---|---|---|
| 1 | 41510696 | 41510965 | 2.281416857 | HIVEP3 |
| 14 | 75469393 | 75469593 | -2.646362498 | JDP2 |
| 14 | 75546665 | 75546859 | 2.122742694 | BATF |
| 14 | 76206317 | 76206525 | -1.520959417 | GPATCH2L |
| 1 | 41513056 | 41513230 | 1.793520865 | HIVEP3 |
| 14 | 76782637 | 76782836 | -1.312520641 | RP11-488C13.5 |
| 14 | 76782637 | 76782836 | -1.312520641 | VASH1 |
| 14 | 76789026 | 76789259 | -1.373433135 | ANGEL1 |
| 14 | 77113646 | 77113831 | -1.29802479 | RP11-463C8.4 |
| 14 | 77113646 | 77113831 | -1.29802479 | CIPC |
| 14 | 77343264 | 77343439 | -1.473407003 | TMED8 |
| 1 | 41513573 | 41513761 | 1.134495021 | HIVEP3 |
| 14 | 80706659 | 80706858 | -4.5 | CEP128 |
| 14 | 80777963 | 80778156 | -5.664898169 | CEP128 |
| 14 | 80785082 | 80785253 | -2.255750169 | CEP128 |
| 14 | 81246202 | 81246401 | 1.191336472 | DYNLL1P1 |
| 14 | 85562749 | 85562916 | -7.0670527 | FLRT2 |
| 14 | 86038574 | 86038712 | 2.772160258 | CTD-2341M24.1 |
| 14 | 86038574 | 86038712 | 2.772160258 | RP11-1079H9.1 |
| 14 | 88010651 | 88010791 | -1.520178869 | GPR65 |
| 14 | 88010651 | 88010791 | -1.520178869 | RP11-300J18.2 |
| 14 | 88444472 | 88444646 | 1.707006247 | SPATA7 |
| 1 | 41543844 | 41544050 | 2.189282176 | RP11-486B10.4 |
| 1 | 41543844 | 41544050 | 2.189282176 | HIVEP3 |
| 14 | 89215554 | 89215725 | 3.094657557 | FOXN3 |
| 14 | 90304079 | 90304292 | -1.148563316 | NRDE2 |
| 1 | 41549152 | 41549349 | 3.052194984 | HIVEP3 |
| 14 | 90938133 | 90938251 | -1.497900006 | RPS6KA5 |
| 14 | 90939205 | 90939327 | -1.866200207 | RPS6KA5 |
| 14 | 91224933 | 91225133 | -1.2055546 | C14orf159 |
| 14 | 91233560 | 91233766 | 1.522424441 | GPR68 |
| 14 | 91258344 | 91258556 | 1.543416234 | CTD-2547L24.4 |
| 1 | 1361350 | 1361553 | -1.587623243 | MXRA8 |
| 1 | 41549558 | 41549758 | 2.844778868 | HIVEP3 |
| 14 | 91969587 | 91969785 | -1.720594378 | TRIP11 |
| 14 | 92014326 | 92014520 | -1.324440679 | TRIP11 |
| 14 | 92040260 | 92040469 | 2.350504091 | TRIP11 |
| 14 | 92040260 | 92040469 | 2.350504091 | ATXN3 |
| 14 | 92161680 | 92161879 | -4.777294426 | CPSF2 |
| 1 | 41551920 | 41552117 | 2.874794648 | HIVEP3 |
| 14 | 93184149 | 93184351 | -1.543887083 | MOAP1 |
| 14 | 93206776 | 93206929 | -1.76896748 | RP11-371E8.4 |
| 14 | 93206776 | 93206929 | -1.76896748 | GON7 |
| 14 | 93333431 | 93333593 | 1.030167832 | UNC79 |
| 1 | 41552654 | 41552852 | 2.533274707 | HIVEP3 |
| 14 | 94045875 | 94046072 | -3.841656829 | OTUB2 |
| 14 | 94114155 | 94114348 | 8.368392535 | IFI27 |
| 14 | 95417082 | 95417303 | -2.131665833 | SYNE3 |
| 14 | 95417753 | 95417896 | -2.426221406 | SYNE3 |
| 1 | 41555806 | 41556004 | 2.678644894 | HIVEP3 |
| 14 | 96354378 | 96354588 | -1.39542258 | ATG2B |
| 14 | 96355301 | 96355409 | -2.10268136 | ATG2B |
| 14 | 99398727 | 99398928 | -1.256431307 | SETD3 |
| 14 | 99515005 | 99515259 | -2.160784986 | CCNK |

| | | | | |
|---|---|---|---|---|
| 12 | 113009218 | 113009422 | 11.61348858 | RP1-71H24.1 |
| 12 | 113158627 | 113158849 | -1.908795233 | CFAP73 |
| 12 | 113158627 | 113158849 | -1.908795233 | DDX54 |
| 1 | 36322133 | 36322339 | -1.002145979 | SH3D21 |
| 1 | 36322133 | 36322339 | -1.002145979 | EVA1B |
| 12 | 114671792 | 114672047 | -1.195882879 | TBX3 |
| 12 | 114683522 | 114683705 | -2.305717356 | TBX3 |
| 12 | 116237586 | 116237800 | 1.208393487 | MED13L |
| 12 | 116665572 | 116665774 | -1.540759811 | RP11-497G19.1 |
| 12 | 116675777 | 116675977 | -2.060718946 | RP11-497G19.1 |
| 12 | 116693337 | 116693531 | -2.415663334 | RP11-497G19.1 |
| 1 | 37455600 | 37455819 | 2.973730262 | LINC01137 |
| 12 | 120095553 | 120095684 | -1.924403591 | RAB35 |
| 12 | 120127535 | 120127676 | -1.725280815 | GCN1 |
| 12 | 120303834 | 120304149 | -1.656651045 | SIRT4 |
| 1 | 37475488 | 37475719 | 1.816841034 | ZC3H12A |
| 12 | 120444560 | 120444757 | -1.37305403 | TRIAP1 |
| 12 | 120444560 | 120444757 | -1.37305403 | AL021546.6 |
| 12 | 120491129 | 120491326 | -2.698552177 | DYNLL1 |
| 12 | 120491129 | 120491326 | -2.698552177 | NRAV |
| 12 | 120498106 | 120498307 | -1.546966876 | DYNLL1 |
| 12 | 120503613 | 120503805 | -1.516501078 | COQ5 |
| 12 | 120579126 | 120579337 | -1.712414001 | POP5 |
| 12 | 120696439 | 120696639 | -2.294529849 | MLEC |
| 12 | 121003332 | 121003531 | -2.186811116 | C12orf43 |
| 1 | 37482961 | 37483163 | 1.125343007 | ZC3H12A |
| 12 | 121003741 | 121003941 | -1.317979364 | C12orf43 |
| 12 | 121020476 | 121020679 | 6.825051055 | OASL |
| 12 | 121185992 | 121186191 | 5.877578399 | P2RX7 |
| 12 | 121417608 | 121417811 | -1.101614931 | RNF34 |
| 12 | 121423485 | 121423688 | -1.524844828 | RNF34 |
| 12 | 121509696 | 121509884 | -1.868521323 | KDM2B |
| 12 | 121814009 | 121814205 | -1.319819797 | SETD1B |
| 1 | 1318944 | 1319161 | -2.545559222 | CPSF3L |
| 12 | 122142314 | 122142512 | -1.321273652 | MLXIP |
| 12 | 122701684 | 122701929 | 8.199597232 | RP11-324E6.6 |
| 12 | 122701684 | 122701929 | 8.199597232 | HCAR2 |
| 1 | 37708234 | 37708433 | -2.991398464 | CDCA8 |
| 12 | 122715372 | 122715570 | 8.02734455 | HCAR3 |
| 12 | 122715372 | 122715570 | 8.02734455 | RP11-324E6.6 |
| 12 | 122716388 | 122716579 | 5.620395666 | HCAR3 |
| 12 | 122777642 | 122777839 | -2.769350722 | CCDC62 |
| 12 | 122848515 | 122848681 | 7.753045295 | HIP1R |
| 12 | 122850859 | 122851031 | 6.469605941 | HIP1R |
| 12 | 122865559 | 122865770 | 1.139477006 | RP11-463O12.3 |
| 12 | 122865559 | 122865770 | 1.139477006 | VPS37B |
| 12 | 122960031 | 122960150 | 1.923169794 | ABCB9 |
| 12 | 122975723 | 122975921 | -1.083788394 | ABCB9 |
| 12 | 122975723 | 122975921 | -1.083788394 | OGFOD2 |
| 12 | 122975723 | 122975921 | -1.083788394 | RP11-197N18.2 |
| 12 | 122980181 | 122980326 | -2.452620917 | RP11-197N18.2 |
| 12 | 122980181 | 122980326 | -2.452620917 | ARL6IP4 |
| 12 | 122980181 | 122980326 | -2.452620917 | ABCB9 |
| 12 | 123221558 | 123221718 | -1.353993264 | MPHOSPH9 |
| 14 | 99515005 | 99515259 | -2.160784986 | CCDC85C |
| 14 | 99675903 | 99676107 | -1.495960175 | HHIPL1 |
| 1 | 41556506 | 41556894 | 2.33353182 | HIVEP3 |
| 14 | 99708816 | 99708963 | -5.464629204 | CYP46A1 |
| 14 | 99713409 | 99713574 | -4.696158711 | CYP46A1 |
| 14 | 100277802 | 100277982 | -1.108762282 | YY1 |
| 14 | 100291533 | 100291731 | -1.457966909 | SLC25A29 |
| 14 | 100291533 | 100291731 | -1.457966909 | RP11-638I2.6 |
| 14 | 100329186 | 100329348 | 2.735031501 | SLC25A47 |
| 1 | 41566517 | 41566714 | 3.184758202 | HIVEP3 |
| 14 | 100455211 | 100455379 | 1.473697958 | WDR25 |
| 14 | 101948756 | 101948911 | -3.383200745 | RP11-1017G21.5 |
| 14 | 102055130 | 102055276 | 1.165694212 | RP11-1017G21.4 |
| 14 | 102225648 | 102225845 | -2.450186805 | MOK |
| 1 | 41580406 | 41580590 | 1.644485335 | HIVEP3 |
| 14 | 102331518 | 102331668 | -7.015505954 | ZNF839 |
| 14 | 102726444 | 102726643 | -1.223544965 | RCOR1 |
| 14 | 102834688 | 102834883 | 1.582289035 | TRAF3 |
| 1 | 41581388 | 41581542 | 2.416013572 | HIVEP3 |
| 14 | 103135574 | 103135782 | 4.983879417 | TNFAIP2 |
| 14 | 103412270 | 103412427 | 6.735329437 | RPL10AP1 |
| 14 | 103412270 | 103412427 | 6.735329437 | MARK3 |
| 14 | 103534709 | 103534912 | 2.178561863 | TRMT61A |
| 1 | 41585030 | 41585241 | 2.236312335 | HIVEP3 |
| 14 | 103607167 | 103607282 | -4.642701109 | RP11-73M18.2 |
| 14 | 103607167 | 103607282 | -4.642701109 | KLC1 |
| 14 | 103607167 | 103607282 | -4.642701109 | APOPT1 |
| 14 | 103734789 | 103734989 | -1.680919967 | PPP1R13B |
| 14 | 104704130 | 104704319 | 7.132734271 | INF2 |
| 1 | 41847981 | 41848151 | 2.152325052 | HIVEP3 |
| 14 | 104801645 | 104801845 | 1.513392799 | ZBTB42 |
| 14 | 104884253 | 104884380 | -4.592437671 | CEP170B |
| 14 | 104938006 | 104938243 | -1.379935334 | AHNAK2 |
| 14 | 104938752 | 104938904 | -1.504210191 | AHNAK2 |
| 14 | 104940753 | 104940881 | -7.014471022 | AHNAK2 |
| 14 | 104941668 | 104942197 | -1.139254113 | AHNAK2 |
| 14 | 104946871 | 104947052 | -2.68839188 | AHNAK2 |
| 14 | 104947326 | 104947475 | -3.72491396 | AHNAK2 |
| 14 | 104948391 | 104948513 | -2.466061316 | AHNAK2 |
| 14 | 104952290 | 104952600 | -2.422507388 | AHNAK2 |
| 14 | 104953883 | 104954024 | -1.032059925 | AHNAK2 |
| 14 | 104956037 | 104956231 | -6.959058926 | AHNAK2 |
| 14 | 104957682 | 104957849 | -5.119732967 | AHNAK2 |
| 14 | 104994098 | 104994299 | 1.101629376 | C14orf79 |
| 14 | 105006391 | 105006578 | -2.744288497 | C14orf79 |
| 14 | 105050822 | 105051022 | 3.851620115 | GPR132 |
| 14 | 105054037 | 105054225 | 3.143672108 | GPR132 |
| 14 | 105055565 | 105055764 | 3.401036246 | GPR132 |
| 14 | 105142119 | 105142317 | -2.227223491 | JAG2 |
| 14 | 105247368 | 105247545 | -6.321477112 | BRF1 |
| 14 | 105250422 | 105250621 | -1.734046299 | BRF1 |
| 14 | 105250422 | 105250621 | -1.734046299 | BTBD6 |
| 14 | 105349788 | 105349989 | -1.624310286 | PACS2 |
| 14 | 105371268 | 105371467 | -5.353679566 | PACS2 |

| | | | | |
|---|---|---|---|---|
| 12 | 123251242 | 123251420 | -1.044631097 | C12orf65 |
| 12 | 123253676 | 123253885 | -1.887009069 | C12orf65 |
| 12 | 123253676 | 123253885 | -1.887009069 | RP11-282O18.3 |
| 12 | 123257082 | 123257293 | -1.977774689 | C12orf65 |
| 12 | 123257082 | 123257293 | -1.977774689 | RP11-282O18.3 |
| 12 | 123415604 | 123415815 | -1.016807027 | RILPL2 |
| 12 | 123472282 | 123472497 | 9.057821579 | SNRNP35 |
| 12 | 123472282 | 123472497 | 9.057821579 | RILPL1 |
| 12 | 123619877 | 123620138 | -2.287176383 | DDX55 |
| 12 | 123621281 | 123621481 | -1.255835898 | EIF2B1 |
| 12 | 124324844 | 124325043 | -3.117866651 | NCOR2 |
| 12 | 124777867 | 124778072 | -1.291836431 | SCARB1 |
| 12 | 125015291 | 125015562 | -3.826588409 | BRI3BP |
| 12 | 125025360 | 125025562 | -2.386235604 | THRIL |
| 12 | 125025360 | 125025562 | -2.386235604 | BRI3BP |
| 12 | 125162161 | 125162298 | -8.779561557 | RP11-158L12.6 |
| 12 | 130972347 | 130972494 | 1.213861379 | ADGRD1 |
| 12 | 131089092 | 131089296 | 7.062827811 | ADGRD1 |
| 12 | 131139329 | 131139505 | 7.667747518 | ADGRD1 |
| 12 | 131941333 | 131941536 | -1.899819477 | PUS1 |
| 12 | 131943449 | 131943646 | -8.202163329 | PUS1 |
| 1 | 37990006 | 37990197 | 8.766368198 | SF3A3 |
| 12 | 132104997 | 132105207 | -1.281577581 | EP400NL |
| 12 | 132108415 | 132108667 | -1.786316347 | EP400NL |
| 12 | 132129207 | 132129442 | -1.169508505 | EP400NL |
| 12 | 132130438 | 132130655 | -1.978357268 | EP400NL |
| 12 | 132553114 | 132553299 | -7.46721402 | RP13-554M15.7 |
| 12 | 132553114 | 132553299 | -7.46721402 | FBRSL1 |
| 12 | 132623999 | 132624180 | -6.836600282 | POLE |
| 12 | 132628813 | 132629007 | -4.817635352 | POLE |
| 12 | 132720842 | 132721117 | -1.876344646 | PGAM5 |
| 12 | 132738062 | 132738256 | -1.161082098 | ANKLE2 |
| 12 | 132754822 | 132755020 | -1.042429246 | ANKLE2 |
| 12 | 132808296 | 132808490 | -1.020330887 | AC127070.1 |
| 12 | 132808296 | 132808490 | -1.020330887 | GOLGA3 |
| 12 | 132941278 | 132941459 | -2.067993096 | ZNF605 |
| 12 | 132941278 | 132941459 | -2.067993096 | CHFR |
| 12 | 133010124 | 133010330 | -1.017660828 | ZNF26 |
| 12 | 133011393 | 133011596 | -1.206365539 | ZNF26 |
| 12 | 133121820 | 133122027 | -1.027500237 | CTD-2140B24.6 |
| 12 | 133121820 | 133122027 | -1.027500237 | ZNF891 |
| 13 | 18211133 | 18211373 | -1.225065314 | FAM230C |
| 1 | 39269118 | 39269303 | -6.02219706 | MACF1 |
| 13 | 19346161 | 19346361 | -1.352628367 | LINC00421 |
| 13 | 19564368 | 19564515 | -3.030347966 | CASC4P1 |
| 13 | 19564368 | 19564515 | -3.030347966 | MRPS31P2 |
| 13 | 19646828 | 19647029 | -1.060684926 | MPHOSPH8 |
| 13 | 19824916 | 19825085 | 1.39063385 | ZMYM5 |
| 13 | 20141824 | 20142122 | 1.004218933 | GJA3 |
| 13 | 20143091 | 20143288 | 1.28646301 | GJA3 |
| 13 | 20608076 | 20608249 | -2.307889107 | IFT88 |
| 13 | 20608076 | 20608249 | -2.307889107 | SLC35E1P1 |
| 13 | 20722933 | 20723068 | -2.84380457 | IL17D |
| 13 | 20773527 | 20773722 | -2.45103679 | EEF1AKMT1 |
| 14 | 105371268 | 105371467 | -5.353679566 | RP11-521B24.4 |
| 1 | 42653931 | 42654130 | -1.152062636 | CCDC30 |
| 14 | 105416831 | 105416959 | -6.142701109 | RP11-521B24.3 |
| 14 | 105416831 | 105416959 | -6.142701109 | TEX22 |
| 15 | 21294182 | 21294371 | -2.306718532 | RP11-32B5.1 |
| 15 | 21368721 | 21368917 | -1.575263218 | RP11-32B5.8 |
| 15 | 21370361 | 21370531 | -2.703613906 | RP11-32B5.8 |
| 15 | 22590049 | 22590250 | -2.321659022 | HERC2P2 |
| 15 | 24988423 | 24988589 | -4.014975405 | RP11-701H24.9 |
| 15 | 24988423 | 24988589 | -4.014975405 | SNHG14 |
| 15 | 24989742 | 24989969 | -2.98527268 | RP11-701H24.9 |
| 15 | 24989742 | 24989969 | -2.98527268 | SNHG14 |
| 15 | 24990731 | 24990926 | -4.930411859 | SNHG14 |
| 15 | 24990731 | 24990926 | -4.930411859 | RP11-701H24.9 |
| 15 | 24998192 | 24998412 | -1.787347083 | RP11-701H24.9 |
| 15 | 24998192 | 24998412 | -1.787347083 | SNHG14 |
| 15 | 25001437 | 25001626 | -3.433418673 | SNHG14 |
| 15 | 25709780 | 25709981 | -1.552447868 | RP11-345J18.2 |
| 15 | 25709780 | 25709981 | -1.552447868 | ATP10A |
| 15 | 25754139 | 25754307 | 8.764654267 | ATP10A |
| 15 | 28111687 | 28111891 | -2.449967714 | HERC2 |
| 15 | 28155038 | 28155231 | -1.878585393 | HERC2 |
| 1 | 42830530 | 42830683 | -1.780080887 | ERMAP |
| 15 | 28155678 | 28155829 | -3.072838728 | HERC2 |
| 15 | 28158214 | 28158325 | -4.827318014 | HERC2 |
| 15 | 28684499 | 28684696 | -2.523604689 | HERC2P9 |
| 15 | 28788564 | 28788702 | -3.004354365 | RP11-578F21.10 |
| 15 | 28789582 | 28789786 | -1.449298713 | PDCD6IPP2 |
| 15 | 28789582 | 28789786 | -1.449298713 | RP11-578F21.10 |
| 15 | 28847774 | 28848019 | -1.458613566 | PDCD6IPP2 |
| 15 | 28847774 | 28848019 | -1.458613566 | GOLGA6L7P |
| 15 | 28856174 | 28856398 | -1.528383888 | PDCD6IPP2 |
| 15 | 28858219 | 28858424 | -1.960577835 | PDCD6IPP2 |
| 15 | 28858219 | 28858424 | -1.960577835 | WI2-2334D6.1 |
| 15 | 30489408 | 30489612 | -1.78941889 | CTD-3092A11.2 |
| 15 | 30489408 | 30489612 | -1.78941889 | CTD-3092A11.1 |
| 15 | 30600242 | 30600406 | -2.403677461 | ULK4P2 |
| 15 | 30678252 | 30678448 | 3.745926548 | ARHGAP11B |
| 15 | 30741707 | 30741894 | -5.462406252 | ARHGAP11B |
| 15 | 30904781 | 30904934 | -3.042918863 | FAN1 |
| 15 | 30941538 | 30941809 | -1.034327122 | MTMR10 |
| 15 | 30941538 | 30941809 | -1.034327122 | FAN1 |
| 15 | 32499795 | 32499994 | -2.303199334 | RP11-632K20.2 |
| 15 | 32580739 | 32581034 | -2.298265501 | RP11-1000B6.3 |
| 15 | 32615058 | 32615258 | -3.3685633 | RP11-1000B6.8 |
| 15 | 32615058 | 32615258 | -3.3685633 | RP11-1000B6.5 |
| 15 | 32615058 | 32615258 | -3.3685633 | ARHGAP11A |
| 15 | 32636493 | 32636694 | -1.093520784 | ARHGAP11A |
| 15 | 34233338 | 34233570 | -1.688144697 | SLC12A6 |
| 15 | 34287976 | 34288117 | -7.41277755 | SLC12A6 |
| 15 | 34337032 | 34337232 | -1.62671001 | SLC12A6 |
| 15 | 34983123 | 34983323 | -1.285849465 | ZNF770 |
| 15 | 35084268 | 35084430 | -6.836654538 | NANOGP8 |
| 15 | 35298106 | 35298305 | -6.537070731 | RBM17P4 |

| | | | | |
|---|---|---|---|---|
| 13 | 21358317 | 21358452 | 1.449638706 | GAPDHP52 |
| 13 | 21701164 | 21701309 | 6.26569073 | FGF9 |
| 13 | 23330187 | 23330388 | -1.326195081 | SACS |
| 13 | 23335479 | 23335679 | -3.368102792 | SACS |
| 13 | 23337998 | 23338185 | -2.596510365 | SACS |
| 13 | 23339436 | 23339622 | -1.657893207 | SACS |
| 13 | 23341375 | 23341576 | -1.730907135 | SACS |
| 13 | 23354806 | 23355006 | -1.761298013 | SACS |
| 13 | 23355420 | 23355621 | -1.892296659 | SACS |
| 13 | 24223757 | 24223954 | -1.968546193 | RP11-307N16.6 |
| 13 | 24223757 | 24223954 | -1.968546193 | SPATA13 |
| 13 | 24249396 | 24249593 | -1.179526255 | SPATA13 |
| 13 | 24249396 | 24249593 | -1.179526255 | RP11-307N16.6 |
| 13 | 24303833 | 24303992 | -1.500367753 | RP11-307N16.6 |
| 13 | 24303833 | 24303992 | -1.500367753 | SPATA13 |
| 13 | 24905784 | 24905987 | -2.448207349 | CENPJ |
| 13 | 24912903 | 24913104 | -1.800805742 | CENPJ |
| 13 | 26214228 | 26214424 | -2.073773229 | RNF6 |
| 13 | 27069114 | 27069288 | 1.538498525 | USP12 |
| 13 | 27622935 | 27623133 | -2.198102901 | POLR1D |
| 13 | 28214712 | 28214914 | -1.278603116 | PAN3 |
| 13 | 28214712 | 28214914 | -1.278603116 | EEF1A1P3 |
| 13 | 28302569 | 28302765 | 2.456268579 | FLT1 |
| 13 | 31331166 | 31331313 | 1.191575792 | B3GLCT |
| 13 | 31953146 | 31953345 | -2.591886241 | EEF1DP3 |
| 13 | 32237542 | 32237743 | -2.614276859 | FRY |
| 13 | 32295263 | 32295462 | -1.709535319 | FRY |
| 13 | 32298669 | 32298867 | -2.42543736 | FRY |
| 13 | 32332971 | 32333208 | -4.421321312 | BRCA2 |
| 13 | 32337038 | 32337239 | -3.462724579 | BRCA2 |
| 13 | 32355014 | 32355192 | -3.406807877 | BRCA2 |
| 13 | 32398188 | 32398490 | -2.913172783 | BRCA2 |
| 13 | 32403002 | 32403203 | 1.915297773 | N4BP2L1 |
| 13 | 32442929 | 32443201 | 1.712280891 | N4BP2L2 |
| 13 | 32770199 | 32770399 | -1.491099455 | PDS5B |
| 13 | 33105510 | 33105729 | 3.207945735 | STARD13 |
| 13 | 33129453 | 33129668 | 2.94649764 | STARD13 |
| 13 | 33947734 | 33947915 | -3.556406321 | RFC3 |
| 13 | 36227132 | 36227381 | -1.882781397 | CCDC169 |
| 13 | 36227132 | 36227381 | -1.882781397 | CCDC169-SOHLH2 |
| 13 | 36335520 | 36335717 | -1.081528149 | SPG20 |
| 13 | 36819591 | 36819788 | -8.707837879 | RFXAP |
| 13 | 39048398 | 39048543 | -1.079592558 | NHLRC3 |
| 1 | 1328427 | 1328623 | -1.411861127 | CPTP |
| 13 | 40559957 | 40560153 | 1.370262623 | FOXO1 |
| 13 | 40864777 | 40865028 | -2.40493034 | TPTE2P5 |
| 13 | 40865558 | 40865729 | -1.131331073 | TPTE2P5 |
| 13 | 40875011 | 40875278 | -1.863237802 | TPTE2P5 |
| 13 | 40877905 | 40878103 | -2.214756254 | TPTE2P5 |
| 13 | 40933507 | 40933708 | -1.116642473 | ELF1 |
| 13 | 41043285 | 41043419 | -1.263540792 | ELF1 |
| 13 | 41046052 | 41046257 | -1.776122879 | ELF1 |
| 1 | 39410179 | 39410383 | -2.244002154 | MACF1 |
| 13 | 41131204 | 41131463 | -2.012412254 | KBTBD6 |
| 15 | 35298106 | 35298305 | -6.537070731 | DPH6 |
| 15 | 40089458 | 40089665 | -2.886378537 | BMF |
| 15 | 40090972 | 40091172 | -2.245457899 | BMF |
| 15 | 40335222 | 40335422 | -8.313317955 | C15orf52 |
| 15 | 40369742 | 40369884 | -4.890679857 | DISP2 |
| 15 | 40371687 | 40371972 | -2.716519459 | DISP2 |
| 15 | 40393726 | 40393932 | -1.988553553 | KNSTRN |
| 15 | 40420025 | 40420222 | -1.150891373 | IVD |
| 15 | 40471510 | 40471708 | -3.616809838 | CHST14 |
| 15 | 40472417 | 40472626 | -1.868990212 | CHST14 |
| 15 | 40569336 | 40569515 | -5.67987478 | RPUSD2 |
| 15 | 40621206 | 40621406 | -2.279046333 | KNL1 |
| 1 | 43413534 | 43413731 | 7.92061017 | SZT2 |
| 15 | 40623530 | 40623732 | -1.748429581 | KNL1 |
| 15 | 40856967 | 40857198 | -1.246605339 | SPINT1 |
| 15 | 40872914 | 40873104 | 2.005818981 | RHOV |
| 15 | 40873168 | 40873336 | 3.209347975 | RHOV |
| 15 | 40900355 | 40900558 | -1.037325784 | VPS18 |
| 15 | 40936630 | 40936811 | 6.388550586 | DLL4 |
| 15 | 40938006 | 40938206 | 5.835134803 | DLL4 |
| 15 | 40955860 | 40956077 | -4.500679956 | CHAC1 |
| 15 | 41298222 | 41298427 | -1.674302544 | OIP5-AS1 |
| 15 | 41503108 | 41503308 | -2.7550524 | ITPKA |
| 15 | 41669556 | 41669757 | -1.017606237 | MGA |
| 15 | 41696110 | 41696221 | -2.007968144 | MGA |
| 15 | 41696654 | 41696853 | -3.152342026 | MGA |
| 15 | 41713199 | 41713357 | -7.270427266 | MGA |
| 15 | 41749297 | 41749492 | -1.484618599 | MGA |
| 15 | 41750304 | 41750514 | -1.449060027 | MGA |
| 15 | 41766235 | 41766436 | -1.27140392 | MGA |
| 15 | 41823746 | 41823938 | -1.004971402 | MAPKBP1 |
| 15 | 41900450 | 41900647 | 2.040089246 | EHD4 |
| 15 | 41908471 | 41908650 | 8.125593252 | EHD4 |
| 15 | 41908471 | 41908650 | 8.125593252 | CTD-2382E5.6 |
| 15 | 42160267 | 42160519 | -1.00910758 | VPS39 |
| 15 | 42249995 | 42250154 | -5.064641508 | TMEM87A |
| 15 | 42299886 | 42300039 | -4.26569073 | GANC |
| 15 | 42304930 | 42305055 | -3.169925001 | GANC |
| 15 | 42336433 | 42336712 | -6.938162543 | GANC |
| 15 | 42439247 | 42439449 | -1.152575708 | ZNF106 |
| 15 | 42448270 | 42448485 | -1.099361677 | ZNF106 |
| 15 | 42451428 | 42451627 | -1.608149481 | ZNF106 |
| 15 | 42684544 | 42684746 | -1.321553272 | STARD9 |
| 15 | 42685002 | 42685190 | -1.567721435 | STARD9 |
| 15 | 42693877 | 42694066 | -6.636460663 | STARD9 |
| 15 | 42724021 | 42724204 | -1.420651127 | CTD-2036P10.5 |
| 15 | 42724021 | 42724204 | -1.420651127 | CDAN1 |
| 15 | 42745855 | 42746020 | -1.259908371 | TTBK2 |
| 15 | 42919875 | 42920082 | 1.062336072 | TTBK2 |
| 15 | 43409056 | 43409169 | -1.605912763 | TUBGCP4 |
| 15 | 43409056 | 43409169 | -1.605912763 | TP53BP1 |
| 15 | 43523096 | 43523296 | 4.669610091 | MAP1A |
| 15 | 43526380 | 43526559 | 3.625750309 | MAP1A |
| 1 | 43704541 | 43704742 | -1.580440348 | KDM4A |

| | | | | |
|---|---|---|---|---|
| 13 | 41193051 | 41193251 | -2.060915029 | KBTBD7 |
| 13 | 41193051 | 41193251 | -2.060915029 | RP11-74J13.8 |
| 13 | 42040017 | 42040137 | 4.160964047 | DGKH |
| 13 | 42041456 | 42041608 | -1.968227527 | DGKH |
| 13 | 42047634 | 42047820 | -2.072358854 | DGKH |
| 13 | 42048079 | 42048277 | -1.460282766 | DGKH |
| 1 | 39411514 | 39411709 | -3.600775839 | MACF1 |
| 13 | 42303325 | 42303544 | -1.517990875 | AKAP11 |
| 13 | 44434483 | 44434788 | 7.291847075 | TSC22D1 |
| 13 | 45391564 | 45391768 | 1.33422448 | TPT1-AS1 |
| 13 | 45395505 | 45395699 | -1.756404602 | SLC25A30 |
| 1 | 39412034 | 39412212 | -3.390679857 | MACF1 |
| 13 | 46296676 | 46296852 | -3.305415293 | LINC00563 |
| 13 | 46372197 | 46372411 | -1.194864753 | RUBCNL |
| 13 | 46750694 | 46750895 | 2.416671658 | LRCH1 |
| 1 | 39414183 | 39414381 | -4.141967556 | MACF1 |
| 13 | 48395405 | 48395612 | 2.873655657 | RB1 |
| 13 | 48395405 | 48395612 | 2.873655657 | LPAR6 |
| 13 | 48396446 | 48396634 | 9.282431995 | LPAR6 |
| 13 | 48396446 | 48396634 | 9.282431995 | RB1 |
| 13 | 48397192 | 48397392 | 3.073756063 | RB1 |
| 13 | 48397192 | 48397392 | 3.073756063 | LPAR6 |
| 13 | 48412266 | 48412415 | 11.21862256 | LPAR6 |
| 13 | 48412266 | 48412415 | 11.21862256 | RB1 |
| 13 | 48417190 | 48417394 | 2.968400916 | LPAR6 |
| 13 | 48417190 | 48417394 | 2.968400916 | RB1 |
| 13 | 48418527 | 48418683 | 8.743884696 | RB1 |
| 13 | 48418527 | 48418683 | 8.743884696 | LPAR6 |
| 13 | 48418926 | 48419110 | 2.022746658 | RB1 |
| 13 | 48418926 | 48419110 | 2.022746658 | LPAR6 |
| 13 | 48422065 | 48422282 | 2.331921236 | RB1 |
| 13 | 48422065 | 48422282 | 2.331921236 | LPAR6 |
| 13 | 48492519 | 48492715 | 9.201666236 | RCBTB2 |
| 13 | 49207061 | 49207302 | 3.256865005 | FNDC3A |
| 13 | 49222191 | 49222417 | -1.238586803 | MLNR |
| 13 | 49267801 | 49267987 | -1.592131331 | CDADC1 |
| 13 | 49274119 | 49274272 | -6.61920237 | CDADC1 |
| 13 | 50807750 | 50807943 | -1.744800438 | DLEU7 |
| 13 | 50807750 | 50807943 | -1.744800438 | DLEU7-AS1 |
| 13 | 50834010 | 50834178 | -1.427130624 | DLEU7 |
| 13 | 50834010 | 50834178 | -1.427130624 | DLEU7-AS1 |
| 13 | 50847831 | 50847971 | -3.136126945 | DLEU7-AS1 |
| 13 | 50867621 | 50867780 | -2.526855293 | RNASEH2B-AS1 |
| 13 | 50896644 | 50896773 | -1.982017251 | RNASEH2B-AS1 |
| 1 | 39527446 | 39527577 | -7.409690395 | RP11-69E11.4 |
| 13 | 50906523 | 50906664 | -1.465388279 | RNASEH2B-AS1 |
| 13 | 51251849 | 51252050 | -1.480335155 | FAM124A |
| 13 | 51281016 | 51281214 | -1.674917046 | FAM124A |
| 1 | 39625397 | 39625606 | -1.694555633 | HEYL |
| 13 | 52377320 | 52377512 | 2.871295565 | THSD1 |
| 13 | 52377808 | 52378000 | 2.785374321 | THSD1 |
| 13 | 52397322 | 52397545 | 1.672874993 | THSD1 |
| 13 | 52461608 | 52461759 | -2.184330457 | CKAP2 |
| 13 | 67226738 | 67226919 | -1.909679851 | PCDH9 |

| | | | | |
|---|---|---|---|---|
| 1 | 43704541 | 43704742 | -1.580440348 | KDM4A-AS1 |
| 15 | 44534999 | 44535212 | -1.974989964 | EIF3J-AS1 |
| 15 | 44536108 | 44536309 | -1.403524859 | EIF3J-AS1 |
| 15 | 44711123 | 44711313 | 1.135231395 | PATL2 |
| 15 | 44731551 | 44731758 | 1.846579379 | TRIM69 |
| 15 | 44767449 | 44767671 | 2.626160273 | TRIM69 |
| 15 | 45200794 | 45200908 | -6.768123608 | SHF |
| 15 | 45598823 | 45599019 | -4.198034779 | RP11-96O20.4 |
| 15 | 45598823 | 45599019 | -4.198034779 | BLOC1S6 |
| 15 | 48258570 | 48258761 | -3.302655573 | SLC12A1 |
| 15 | 48738525 | 48738643 | -1.238032819 | CEP152 |
| 15 | 48756164 | 48756413 | -1.351100088 | CEP152 |
| 15 | 48878737 | 48878971 | -1.223153191 | EID1 |
| 15 | 48878737 | 48878971 | -1.223153191 | SHC4 |
| 15 | 49860172 | 49860359 | -2.24948193 | ATP8B4 |
| 15 | 49936762 | 49936935 | -6.779509262 | ATP8B4 |
| 15 | 49948821 | 49949000 | -7.625149209 | ATP8B4 |
| 15 | 49998720 | 49998897 | -7.837606393 | ATP8B4 |
| 15 | 50000425 | 50000621 | -4.089954545 | ATP8B4 |
| 15 | 50235952 | 50236146 | -6.135147663 | SLC27A2 |
| 15 | 50354022 | 50354233 | -3.008379204 | GABPB1 |
| 1 | 43913723 | 43913918 | 4.441321525 | ST3GAL3 |
| 15 | 50481681 | 50481817 | -1.02338173 | USP8 |
| 15 | 50944970 | 50945164 | 1.182181681 | AP4E1 |
| 15 | 50944970 | 50945164 | 1.182181681 | DCAF13P3 |
| 15 | 51057983 | 51058183 | -1.432664541 | TNFAIP8L3 |
| 15 | 51057983 | 51058183 | -1.432664541 | RP11-108K3.1 |
| 15 | 51318654 | 51318807 | -3.911086231 | RP11-108K3.2 |
| 15 | 51318654 | 51318807 | -3.911086231 | CYP19A1 |
| 15 | 51384662 | 51384810 | -6.560604858 | GLDN |
| 15 | 51499379 | 51499578 | -2.315338408 | DMXL2 |
| 15 | 51549732 | 51549927 | -6.051643904 | DMXL2 |
| 15 | 51966184 | 51966374 | -1.449611967 | MAPK6 |
| 15 | 51966184 | 51966374 | -1.449611967 | LEO1 |
| 15 | 52381130 | 52381316 | -3.515563406 | MYO5A |
| 15 | 52476078 | 52476275 | -6.818538829 | MYO5A |
| 15 | 52608717 | 52608915 | -2.770492359 | FAM214A |
| 15 | 52609454 | 52609636 | -2.31727846 | FAM214A |
| 15 | 55317646 | 55317813 | -1.898544214 | PIGBOS1 |
| 15 | 55317646 | 55317813 | -1.898544214 | RAB27A |
| 15 | 55348472 | 55348671 | -3.039703705 | PIGB |
| 15 | 55348472 | 55348671 | -3.039703705 | CCPG1 |
| 15 | 55360419 | 55360642 | -1.714434783 | CCPG1 |
| 15 | 55360419 | 55360642 | -1.714434783 | DYX1C1-CCPG1 |
| 15 | 55544416 | 55544615 | -1.92008243 | PYGO1 |
| 1 | 43926544 | 43926733 | 5.513952998 | ST3GAL3 |
| 15 | 55546351 | 55546597 | -3.43878603 | PYGO1 |
| 15 | 55915731 | 55915930 | -6.589332426 | NEDD4 |
| 15 | 56093456 | 56093654 | -1.470013532 | RFX7 |
| 15 | 56093808 | 56094010 | -1.524579184 | RFX7 |
| 15 | 56095160 | 56095359 | -1.906818113 | RFX7 |
| 15 | 56095756 | 56095932 | -1.847625376 | RFX7 |
| 15 | 56123495 | 56123682 | -2.204481492 | RFX7 |
| 15 | 56436440 | 56436583 | -2.031071652 | TEX9 |

| | | | | | | | | | |
|---|---|---|---|---|---|---|---|---|---|
| 13 | 71440192 | 71440394 | -2.359122298 | DACH1 | 15 | 56436440 | 56436583 | -2.031071652 | MNS1 |
| 13 | 71681840 | 71682045 | -10.05638264 | DACH1 | 15 | 56439864 | 56440061 | -2.728213879 | MNS1 |
| 13 | 71866011 | 71866214 | -2.495481322 | DACH1 | 15 | 56439864 | 56440061 | -2.728213879 | TEX9 |
| 13 | 72746751 | 72746922 | -2.575716793 | BORA | 15 | 56442603 | 56442802 | -1.77041423 | TEX9 |
| 13 | 75286754 | 75286981 | 2.04009742 | TBC1D4 | 15 | 56442603 | 56442802 | -1.77041423 | MNS1 |
| 1 | 39758732 | 39758932 | -3.061907185 | PPIE | 15 | 56631551 | 56631805 | -2.505203313 | ZNF280D |
| 1 | 39758732 | 39758932 | -3.061907185 | BMP8B | 15 | 56820564 | 56820670 | -2.5 | ZNF280D |
| 13 | 77045044 | 77045253 | -1.310401749 | RP11-226E21.4 | 15 | 56954392 | 56954606 | -2.040938019 | TCF12 |
| 13 | 77045044 | 77045253 | -1.310401749 | MYCBP2-AS1 | 15 | 56955097 | 56955281 | -1.28528235 | TCF12 |
| 13 | 77045044 | 77045253 | -1.310401749 | MYCBP2 | 15 | 57286135 | 57286332 | -1.71557192 | TCF12 |
| 13 | 77098586 | 77098915 | -1.207714771 | RP11-226E21.4 | 15 | 57293449 | 57293607 | -2.279510549 | TCF12 |
| 13 | 77098586 | 77098915 | -1.207714771 | MYCBP2-AS1 | 15 | 57977254 | 57977374 | -3.836212671 | ALDH1A2 |
| 13 | 77098586 | 77098915 | -1.207714771 | MYCBP2 | 15 | 58316144 | 58316338 | -1.838742376 | ALDH1A2 |
| 13 | 78600866 | 78601051 | -2.847498817 | RNF219-AS1 | 15 | 58591956 | 58592156 | -2.845871191 | ADAM10 |
| 13 | 78600866 | 78601051 | -2.847498817 | POU4F1 | 15 | 58597355 | 58597554 | -1.464244897 | ADAM10 |
| 13 | 78601441 | 78601646 | -2.63139712 | RNF219-AS1 | 15 | 58771899 | 58772092 | -2.405324706 | FAM63B |
| 13 | 78601441 | 78601646 | -2.63139712 | POU4F1 | 15 | 58847342 | 58847540 | -1.864110664 | FAM63B |
| 13 | 78602400 | 78602589 | -7.149317703 | RNF219-AS1 | 15 | 58851654 | 58851844 | -2.119539239 | FAM63B |
| 13 | 78602400 | 78602589 | -7.149317703 | POU4F1 | 15 | 58854399 | 58854605 | -2.214288656 | FAM63B |
| 13 | 78615927 | 78616229 | -1.348053209 | RNF219-AS1 | 15 | 58864685 | 58864889 | -2.707546556 | RP11-30K9.5 |
| 13 | 78615927 | 78616229 | -1.348053209 | RNF219 | 15 | 58865384 | 58865578 | -4.008827077 | RNF111 |
| 13 | 78616665 | 78616883 | -1.018998095 | RNF219-AS1 | 15 | 59125935 | 59126088 | -1.261786766 | RP11-59H7.3 |
| 13 | 78616665 | 78616883 | -1.018998095 | RNF219 | 15 | 59126521 | 59126724 | -1.178049853 | RP11-59H7.3 |
| 13 | 80337076 | 80337280 | 3.472277319 | SPRY2 | 15 | 59127576 | 59127736 | -1.81903002 | RP11-59H7.3 |
| 1 | 39759656 | 39759857 | -2.990279744 | BMP8B | 1 | 44700979 | 44701137 | 6.54637857 | C1orf228 |
| 1 | 39759656 | 39759857 | -2.990279744 | PPIE | 15 | 59128483 | 59128685 | -2.032857722 | RP11-59H7.3 |
| 13 | 80340045 | 80340196 | 3.189483935 | SPRY2 | 15 | 59129183 | 59129385 | -1.267900796 | RP11-59H7.3 |
| 1 | 39769707 | 39769933 | -3.474578946 | BMP8B | 15 | 59136027 | 59136227 | -1.393443016 | MYO1E |
| 1 | 39769707 | 39769933 | -3.474578946 | OXCT2 | 15 | 59190397 | 59190597 | 5.269579406 | MYO1E |
| 13 | 87692729 | 87692923 | 2.145962347 | SLITRK5 | 1 | 44805787 | 44805984 | 3.115457816 | PLK3 |
| 1 | 39770334 | 39770528 | -3.987556818 | BMP8B | 1 | 44805787 | 44805984 | 3.115457816 | TCTEX1D4 |
| 1 | 39770334 | 39770528 | -3.987556818 | OXCT2 | 15 | 62049137 | 62049289 | -6.268123608 | VPS13C |
| 13 | 91693441 | 91693603 | 2.708222847 | GPC5 | 15 | 63141408 | 63141702 | 1.579336472 | LACTB |
| 13 | 93227188 | 93227311 | -1.494676378 | GPC6 | 15 | 63141408 | 63141702 | 1.579336472 | RPS27L |
| 13 | 93227188 | 93227311 | -1.494676378 | RP11-632L2.2 | 15 | 63305694 | 63305879 | -6.592437671 | APH1B |
| 13 | 99255320 | 99255502 | -2.975506574 | GPR18 | 15 | 63591140 | 63591355 | -2.456535069 | USP3-AS1 |
| 13 | 99255320 | 99255502 | -2.975506574 | UBAC2 | 15 | 63591140 | 63591355 | -2.456535069 | USP3 |
| 13 | 99295643 | 99295972 | 1.532896379 | UBAC2 | 15 | 63620861 | 63621065 | -1.577050017 | HERC1 |
| 13 | 99295643 | 99295972 | 1.532896379 | GPR183 | 15 | 64170586 | 64170843 | 1.185960892 | CSNK1G1 |
| 1 | 39770956 | 39771152 | -9.631710256 | OXCT2 | 15 | 64815787 | 64815985 | -5.55033117 | PIF1 |
| 1 | 39770956 | 39771152 | -9.631710256 | BMP8B | 15 | 64854797 | 64854988 | 1.563446191 | PLEKHO2 |
| 13 | 99700357 | 99700550 | -2.978889411 | CLYBL | 15 | 64854797 | 64854988 | 1.563446191 | AC069368.3 |
| 13 | 99982584 | 99982839 | -1.315637648 | ZIC2 | 1 | 45343237 | 45343441 | -1.354801685 | TOE1 |
| 13 | 100358000 | 100358185 | 1.973329771 | PCCA | 15 | 64951879 | 64951999 | -5.958872507 | ANKDD1A |
| 1 | 39771404 | 39771619 | -3.804797849 | BMP8B | 15 | 65624613 | 65624804 | -7.360041951 | SLC24A1 |
| 13 | 102772510 | 102772734 | -1.746304388 | TEX30 | 15 | 65624955 | 65625130 | -7.467490416 | SLC24A1 |
| 13 | 102772992 | 102773165 | -6.720694015 | TEX30 | 15 | 65625327 | 65625504 | 8.26080022 | SLC24A1 |
| 13 | 102862412 | 102862626 | -1.114642989 | ERCC5 | 15 | 65652134 | 65652314 | 1.966101331 | SLC24A1 |
| 13 | 102862412 | 102862626 | -1.114642989 | BIVM-ERCC5 | 15 | 66325981 | 66326181 | -2.321692358 | RP11-352G18.2 |
| 13 | 102875587 | 102875787 | -1.305375274 | ERCC5 | 15 | 66325981 | 66326181 | -2.321692358 | DIS3L |
| 13 | 109785219 | 109785420 | -8.909191228 | IRS2 | 15 | 66489815 | 66489996 | -6.405084969 | CTD-3185P2.1 |
| 13 | 110523551 | 110523760 | 1.534482106 | RAB20 | 15 | 66489815 | 66489996 | -6.405084969 | MAP2K1 |
| 13 | 110614133 | 110614303 | -1.487002396 | RP11-90L1.8 | 15 | 66702477 | 66702671 | -6.465738117 | SMAD6 |
| 13 | 110659909 | 110660185 | -1.068236575 | CARS2 | 15 | 67407085 | 67407285 | -2.421443652 | IQCH-AS1 |

| | | | | |
|---|---|---|---|---|
| 13 | 110879752 | 110879908 | 2.081361405 | ANKRD10 |
| 13 | 110883649 | 110883832 | 1.764674982 | ANKRD10 |
| 13 | 111228676 | 111228871 | -4.359180813 | ARHGEF7 |
| 1 | 39869548 | 39869745 | -3.885313467 | TRIT1 |
| 13 | 112689215 | 112689373 | -2.067843037 | RP11-88E10.5 |
| 13 | 112807226 | 112807434 | -2.011056274 | ATP11A |
| 13 | 112879543 | 112879764 | -2.02058444 | ATP11A |
| 13 | 112880505 | 112880778 | -1.972314619 | ATP11A |
| 1 | 39869852 | 39870046 | -10.51104504 | TRIT1 |
| 13 | 113251933 | 113252128 | -1.577662575 | CUL4A |
| 13 | 113252438 | 113252597 | -8.135786226 | CUL4A |
| 13 | 113263472 | 113263677 | -1.048772236 | CUL4A |
| 13 | 113402775 | 113402971 | 3.540012499 | ADPRHL1 |
| 13 | 113405638 | 113405835 | 1.591335115 | ADPRHL1 |
| 13 | 113406400 | 113406674 | 1.615624841 | ADPRHL1 |
| 13 | 113407792 | 113407967 | -4.584962501 | ADPRHL1 |
| 13 | 113410440 | 113410636 | -1.659750679 | ADPRHL1 |
| 13 | 113413737 | 113413931 | -1.231925961 | ADPRHL1 |
| 13 | 113416237 | 113416403 | -1.728404619 | ADPRHL1 |
| 13 | 113511732 | 113511960 | -1.580549962 | TMCO3 |
| 13 | 113511732 | 113511960 | -1.580549962 | RP11-230F18.6 |
| 13 | 113599899 | 113600097 | -2.651404017 | TFDP1 |
| 13 | 113816679 | 113816812 | -3.748529411 | GAS6-AS1 |
| 13 | 113816679 | 113816812 | -3.748529411 | TMEM255B |
| 1 | 39901403 | 39901540 | -1.317864085 | MYCL |
| 13 | 113817848 | 113818102 | -3.97034953 | GAS6-AS1 |
| 13 | 113818813 | 113818947 | -3.727383312 | GAS6-AS1 |
| 13 | 113953900 | 113954105 | 3.986859744 | ABC13-47488600E17.1 |
| 13 | 113954447 | 113954642 | 4.059064062 | ABC13-47488600E17.1 |
| 13 | 113956509 | 113956703 | 5.334557269 | ABC13-47488600E17.1 |
| 13 | 113973903 | 113974109 | 2.289618868 | ABC13-47488600E17.1 |
| 13 | 113979187 | 113979384 | -7.215226276 | RASA3 |
| 13 | 114052741 | 114052941 | 8.293185348 | RASA3 |
| 13 | 114058727 | 114058909 | -8.097253396 | RASA3 |
| 13 | 114059932 | 114060119 | -3.767334294 | RASA3 |
| 1 | 39963123 | 39963323 | -10.45155477 | MFSD2A |
| 1 | 39963123 | 39963323 | -10.45155477 | RP3-342P20.2 |
| 13 | 114062758 | 114062909 | -9.493965064 | RASA3 |
| 13 | 114076878 | 114077075 | 7.066049016 | RASA3 |
| 13 | 114304784 | 114304988 | -1.14731107 | UPF3A |
| 1 | 39969556 | 39969759 | -1.240702699 | MFSD2A |
| 13 | 114323903 | 114324196 | -1.272249401 | CHAMP1 |
| 13 | 114325595 | 114325794 | -1.526036804 | CHAMP1 |
| 14 | 19097103 | 19097356 | -1.10987695 | DUXAP9 |
| 14 | 19301249 | 19301422 | -5.428990498 | LINC01296 |
| 14 | 19301249 | 19301422 | -5.428990498 | AL589743.1 |
| 14 | 19316403 | 19316587 | -1.245926548 | LINC01296 |
| 14 | 19316403 | 19316587 | -1.245926548 | BMS1P17 |
| 14 | 20934758 | 20934957 | -7.989344059 | RP11-219E7.4 |
| 14 | 20983269 | 20983467 | -3.602873132 | RP11-84C10.1 |
| 14 | 21074679 | 21074946 | 2.012096295 | ARHGEF40 |
| 14 | 21081896 | 21082080 | 1.463947163 | ARHGEF40 |
| 14 | 21352713 | 21352905 | -2.50106173 | SUPT16H |
| 14 | 21385554 | 21385755 | -1.371267586 | CHD8 |
| 15 | 67407085 | 67407285 | -2.421443652 | IQCH |
| 15 | 67422872 | 67423103 | -2.008097421 | IQCH |
| 15 | 67422872 | 67423103 | -2.008097421 | IQCH-AS1 |
| 15 | 67448488 | 67448672 | -3.259630909 | IQCH |
| 15 | 67448488 | 67448672 | -3.259630909 | IQCH-AS1 |
| 15 | 67453357 | 67453565 | -1.459973374 | IQCH |
| 15 | 67453357 | 67453565 | -1.459973374 | IQCH-AS1 |
| 15 | 67526333 | 67526519 | -1.70025341 | C15orf61 |
| 15 | 67826449 | 67826609 | -6.83521748 | SKOR1 |
| 15 | 68193417 | 68193718 | -1.412214218 | RP11-315D16.2 |
| 15 | 68193417 | 68193718 | -1.412214218 | CALML4 |
| 15 | 68193417 | 68193718 | -1.412214218 | PIAS1 |
| 15 | 68193939 | 68194191 | -7.734486992 | PIAS1 |
| 15 | 68193939 | 68194191 | -7.734486992 | CALML4 |
| 15 | 68193939 | 68194191 | -7.734486992 | RP11-315D16.2 |
| 1 | 45607441 | 45607641 | -1.456674885 | NASP |
| 15 | 68197705 | 68197905 | -1.332273024 | RP11-315D16.2 |
| 15 | 68197705 | 68197905 | -1.332273024 | CALML4 |
| 15 | 68197705 | 68197905 | -1.332273024 | PIAS1 |
| 15 | 68205764 | 68206007 | -1.25337157 | RP11-315D16.2 |
| 15 | 68205764 | 68206007 | -1.25337157 | CALML4 |
| 15 | 68289691 | 68289880 | -1.00992408 | FEM1B |
| 15 | 68789331 | 68789541 | -2.184392095 | ANP32A |
| 15 | 69196440 | 69196645 | -1.464717643 | GLCE |
| 15 | 69255963 | 69256161 | -1.263293289 | GLCE |
| 15 | 69268860 | 69269054 | -1.92237525 | GLCE |
| 15 | 70832239 | 70832412 | -2.542988364 | LARP6 |
| 15 | 71826643 | 71826861 | -1.205476668 | MYO9A |
| 15 | 71897709 | 71897897 | -1.513901038 | MYO9A |
| 15 | 71898748 | 71898932 | -1.000129634 | MYO9A |
| 15 | 72140541 | 72140698 | -6.325525846 | RP11-2I17.4 |
| 15 | 72140541 | 72140698 | -6.325525846 | SENP8 |
| 15 | 72474581 | 72474783 | 9.372170569 | RP11-1007O24.3 |
| 15 | 72474581 | 72474783 | 9.372170569 | ARIH1 |
| 15 | 72733955 | 72734150 | -1.229180952 | BBS4 |
| 15 | 72783865 | 72784074 | 2.010590567 | ADPGK |
| 15 | 72783865 | 72784074 | 2.010590567 | ADPGK-AS1 |
| 15 | 73302659 | 73302856 | -1.459802138 | NEO1 |
| 15 | 73714197 | 73714397 | 1.212271638 | CD276 |
| 15 | 74035462 | 74035650 | 3.160372721 | PML |
| 15 | 74040937 | 74041113 | 9.549592989 | PML |
| 15 | 74044404 | 74044594 | 2.685253094 | PML |
| 15 | 74461623 | 74461754 | -6.266664866 | UBL7-AS1 |
| 15 | 74598662 | 74598860 | -7.709321625 | CLK3 |
| 15 | 74601694 | 74601876 | -3.953445298 | CLK3 |
| 15 | 74605361 | 74605575 | -2.631456847 | CLK3 |
| 15 | 74955608 | 74955823 | -2.027867506 | RPP25 |
| 15 | 75341462 | 75341745 | -1.046471184 | COMMD4 |
| 15 | 75371556 | 75371758 | -1.21285903 | SIN3A |
| 15 | 75468689 | 75468861 | -2.02246388 | PTPN9 |
| 15 | 75587274 | 75587405 | -2.379452431 | CTD-2323K18.1 |
| 15 | 75639293 | 75639493 | -1.51252632 | IMP3 |
| 15 | 75648857 | 75649056 | -2.206864228 | SNX33 |
| 15 | 75650234 | 75650439 | -2.316120484 | SNX33 |

| | | | | | | | | | |
|---|---|---|---|---|---|---|---|---|---|
| 14 | 21393646 | 21393914 | -1.085641826 | CHD8 | 15 | 75687398 | 75687613 | -2.193262734 | CSPG4 |
| 14 | 21431383 | 21431592 | -1.073146214 | CHD8 | 15 | 75689688 | 75689910 | -2.18408423 | CSPG4 |
| 14 | 22418617 | 22418813 | -2.490011149 | TRAC | 15 | 75727796 | 75727953 | -2.547100437 | DNM1P35 |
| 14 | 22418617 | 22418813 | -2.490011149 | TRDC | 15 | 76259243 | 76259440 | -5.475642357 | ETFA |
| 14 | 22418617 | 22418813 | -2.490011149 | AE000661.50 | 15 | 76259243 | 76259440 | -5.475642357 | TYRO3P |
| 14 | 22418617 | 22418813 | -2.490011149 | AE000661.37 | 15 | 77110899 | 77111095 | -6.671106798 | PEAK1 |
| 14 | 22473431 | 22473619 | 2.041884179 | AE000661.37 | 1 | 46608180 | 46608375 | 4.710841171 | MOB3C |
| 14 | 22473431 | 22473619 | 2.041884179 | TRAC | 1 | 46608180 | 46608375 | 4.710841171 | MKNK1 |
| 14 | 22812995 | 22813195 | 2.602732564 | SLC7A7 | 15 | 77479117 | 77479317 | -1.503901718 | HMG20A |
| 14 | 22845956 | 22846158 | 3.900075621 | MMP14 | 15 | 77896488 | 77896669 | -3.050054368 | CSPG4P13 |
| 14 | 22875974 | 22876183 | 1.623548342 | LRP10 | 15 | 77897254 | 77897405 | -1.750112747 | CSPG4P13 |
| 14 | 22987276 | 22987496 | -1.096627732 | C14orf93 | 15 | 77898894 | 77899097 | -1.72920341 | CSPG4P13 |
| 14 | 22987276 | 22987496 | -1.096627732 | RP11-298I3.4 | 1 | 46609491 | 46609699 | 2.442305915 | MOB3C |
| 14 | 23117383 | 23117579 | -6.258274768 | CEBPE | 1 | 46609491 | 46609699 | 2.442305915 | MKNK1 |
| 14 | 23275651 | 23275852 | -1.336519539 | HOMEZ | 15 | 78766537 | 78766735 | -1.196437435 | ADAMTS7 |
| 14 | 23320487 | 23320686 | -2.62181508 | BCL2L2-PABPN1 | 15 | 78988381 | 78988577 | -2.03124499 | RASGRF1 |
| 14 | 23520941 | 23521131 | 1.686591328 | ZFHX2 | 15 | 79458160 | 79458394 | -1.221799039 | KIAA1024 |
| 14 | 23520941 | 23521131 | 1.686591328 | RP11-66N24.4 | 1 | 46613191 | 46613393 | 2.735865756 | MOB3C |
| 14 | 23525980 | 23526198 | 1.205270289 | ZFHX2 | 1 | 46613191 | 46613393 | 2.735865756 | MKNK1 |
| 14 | 23525980 | 23526198 | 1.205270289 | RP11-66N24.4 | 15 | 79899022 | 79899218 | -8.443276979 | ST20 |
| 14 | 23534958 | 23535154 | 1.258920855 | ZFHX2 | 15 | 79899022 | 79899218 | -8.443276979 | ST20-MTHFS |
| 14 | 23534958 | 23535154 | 1.258920855 | RP11-66N24.4 | 15 | 81001132 | 81001312 | 6.661668145 | MESDC1 |
| 14 | 23940418 | 23940750 | -1.248071165 | DHRS4-AS1 | 15 | 81001769 | 81001983 | 9.096305341 | MESDC1 |
| 14 | 24240340 | 24240538 | -1.015185375 | TINF2 | 15 | 81299395 | 81299582 | -2.641691526 | IL16 |
| 14 | 24317682 | 24317825 | -7.659694938 | LTB4R | 15 | 81312877 | 81313066 | 3.314533882 | STARD5 |
| 14 | 24427131 | 24427323 | 1.754337569 | CBLN3 | 15 | 81312877 | 81313066 | 3.314533882 | IL16 |
| 14 | 24431815 | 24432044 | -1.906900768 | KHNYN | 15 | 82846591 | 82846795 | -7.54476662 | HOMER2 |
| 14 | 31690881 | 31691069 | -1.076310553 | NUBPL | 15 | 82849803 | 82850003 | -3.590948449 | HOMER2 |
| 14 | 34761926 | 34762130 | -2.239451382 | BAZ1A | 15 | 82920024 | 82920209 | 6.763738503 | HOMER2 |
| 14 | 34776045 | 34776246 | -1.798495661 | BAZ1A | 1 | 47097572 | 47097673 | -1.12165913 | CYP4Z1 |
| 14 | 44964110 | 44964310 | -1.168525036 | KLHL28 | 1 | 47097572 | 47097673 | -1.12165913 | CYP4A22-AS1 |
| 14 | 44964110 | 44964310 | -1.168525036 | FAM179B | 15 | 83017177 | 83017327 | -1.03332508 | RP11-382A20.7 |
| 14 | 45136188 | 45136385 | -2.894088045 | FANCM | 15 | 83017177 | 83017327 | -1.03332508 | BTBD1 |
| 14 | 45188849 | 45189122 | -2.405432828 | FANCM | 15 | 83017177 | 83017327 | -1.03332508 | RP11-382A20.5 |
| 14 | 45196056 | 45196203 | -4.253765193 | FANCM | 15 | 83329149 | 83329322 | 1.903677461 | RP11-382A20.4 |
| 14 | 45224318 | 45224475 | -2.181695379 | MIS18BP1 | 15 | 84512336 | 84512528 | -1.133196709 | GOLGA6L5P |
| 14 | 45242195 | 45242398 | -1.45728528 | MIS18BP1 | 15 | 84605966 | 84606138 | -6.414861368 | RP11-182J1.18 |
| 14 | 45246952 | 45247153 | -1.384516059 | MIS18BP1 | 15 | 84605966 | 84606138 | -6.414861368 | ZSCAN2 |
| 14 | 49577524 | 49577748 | 1.474838349 | RPS29 | 15 | 84620749 | 84620947 | -2.684616905 | ZSCAN2 |
| 14 | 49577524 | 49577748 | 1.474838349 | RPL32P29 | 15 | 84620749 | 84620947 | -2.684616905 | RP11-182J1.18 |
| 14 | 49618897 | 49619098 | -3.266994831 | RPL36AL | 15 | 84621894 | 84622144 | -2.243301734 | ZSCAN2 |
| 14 | 49622154 | 49622355 | -13.81766791 | MGAT2 | 15 | 84621894 | 84622144 | -2.243301734 | RP11-182J1.3 |
| 14 | 49622154 | 49622355 | -13.81766791 | RP11-649E7.5 | 15 | 84621894 | 84622144 | -2.243301734 | RP11-182J1.18 |
| 14 | 50007029 | 50007225 | 2.790429051 | LINC01588 | 15 | 84639715 | 84639888 | -1.567219816 | WDR73 |
| 14 | 50329969 | 50330186 | 1.221558024 | ATP5S | 15 | 84639715 | 84639888 | -1.567219816 | SCAND2P |
| 14 | 50329969 | 50330186 | 1.221558024 | CDKL1 | 1 | 841390 | 841584 | 1.440761938 | LINC01128 |
| 14 | 50756663 | 50756863 | -2.24012019 | NIN | 15 | 84643426 | 84643628 | -1.022494139 | WDR73 |
| 14 | 50757300 | 50757433 | -3.186652367 | NIN | 15 | 84643426 | 84643628 | -1.022494139 | SCAND2P |
| 14 | 50758124 | 50758326 | -1.977744223 | NIN | 15 | 84646112 | 84646307 | -1.123223959 | WDR73 |
| 14 | 50760077 | 50760274 | -1.544166656 | NIN | 15 | 84646112 | 84646307 | -1.123223959 | SCAND2P |
| 1 | 40473208 | 40473425 | 1.03406225 | RP11-656D10.7 | 15 | 84782910 | 84783109 | -1.61355048 | ZNF592 |
| 14 | 50943700 | 50943926 | -6.794298764 | PYGL | 15 | 84783915 | 84784195 | -1.169464657 | ZNF592 |
| 14 | 53045323 | 53045453 | 1.48414557 | DDHD1 | 15 | 84839971 | 84840128 | -2.286444834 | ALPK3 |
| 14 | 53270118 | 53270300 | 2.609956569 | AL163953.3 | 15 | 85580833 | 85581055 | 1.356109244 | AKAP13 |

| | | | | |
|---|---|---|---|---|
| 14 | 53320305 | 53320425 | -1.780703886 | AL163953.3 |
| 14 | 53950559 | 53950758 | 2.382253898 | BMP4 |
| 14 | 54405575 | 54405725 | 5.727149647 | CDKN3 |
| 14 | 54567734 | 54567887 | 6.098125414 | SAMD4A |
| 1 | 40486959 | 40487161 | 2.353106269 | ZFP69 |
| 14 | 55298928 | 55299127 | 7.573542825 | FBXO34 |
| 14 | 55298928 | 55299127 | 7.573542825 | CHMP4BP1 |
| 1 | 40495532 | 40495772 | -1.665248683 | RP11-656D10.3 |
| 1 | 40495532 | 40495772 | -1.665248683 | ZFP69 |
| 14 | 56647316 | 56647516 | -1.103357002 | RP11-1085N6.2 |
| 14 | 56647316 | 56647516 | -1.103357002 | TMEM260 |
| 14 | 57409341 | 57409471 | 1.506380727 | NAA30 |
| 1 | 40514725 | 40514966 | -1.118171973 | EXO5 |
| 1 | 40514725 | 40514966 | -1.118171973 | RP11-656D10.5 |
| 14 | 58521660 | 58521854 | -7.635950336 | HNRNPCP1 |
| 14 | 58521660 | 58521854 | -7.635950336 | KIAA0586 |
| 14 | 59368636 | 59368881 | 2.019061896 | DAAM1 |
| 14 | 59463590 | 59463801 | -1.574027576 | L3HYPDH |
| 14 | 59463590 | 59463801 | -1.574027576 | GPR135 |
| 14 | 60107682 | 60107884 | -1.143367779 | PCNX4 |
| 1 | 40516857 | 40517044 | -1.575843798 | RP11-656D10.6 |
| 14 | 60124421 | 60124658 | -1.050823808 | PCNX4 |
| 14 | 60248782 | 60248981 | 5.140965013 | PPM1A |
| 14 | 60889473 | 60889662 | -1.013885784 | MNAT1 |
| 14 | 60889473 | 60889662 | -1.013885784 | RP11-193F5.1 |
| 14 | 61012438 | 61012645 | 3.444141249 | SLC38A6 |
| 14 | 61013153 | 61013351 | 2.996184406 | SLC38A6 |
| 14 | 61013805 | 61014005 | 3.362622258 | SLC38A6 |
| 14 | 61014856 | 61015043 | 7.843838198 | SLC38A6 |
| 14 | 61058062 | 61058259 | 6.661386588 | SLC38A6 |
| 14 | 61061478 | 61061609 | 4.648266942 | SLC38A6 |
| 14 | 61708301 | 61708485 | -9.201526822 | HIF1A |
| 14 | 63598970 | 63599177 | -1.722311556 | CTD-2302E22.2 |
| 14 | 63598970 | 63599177 | -1.722311556 | WDR89 |
| 14 | 63599637 | 63599753 | -2.628020278 | WDR89 |
| 14 | 64531765 | 64531956 | 6.661245769 | ZBTB1 |
| 14 | 64531765 | 64531956 | 6.661245769 | RP11-973N13.4 |
| 14 | 64542411 | 64542574 | -2.256677083 | HSPA2 |
| 14 | 64741131 | 64741288 | -2.888330844 | PLEKHG3 |
| 14 | 64743452 | 64743657 | -2.896151965 | PLEKHG3 |
| 14 | 65061902 | 65062101 | -1.350030494 | FNTB |
| 14 | 65061902 | 65062101 | -1.350030494 | MAX |
| 14 | 65411798 | 65411999 | 7.588592961 | FUT8-AS1 |
| 14 | 65411798 | 65411999 | 7.588592961 | FUT8 |
| 14 | 65412122 | 65412310 | 2.554822364 | FUT8 |
| 14 | 65412122 | 65412310 | 2.554822364 | FUT8-AS1 |
| 14 | 65615848 | 65616046 | 1.349883488 | FUT8 |
| 14 | 65633099 | 65633233 | 6.509013133 | FUT8 |
| 14 | 67273476 | 67273675 | 6.262148419 | MPP5 |
| 14 | 67274154 | 67274325 | 3.604030065 | MPP5 |
| 14 | 67328524 | 67328681 | 1.059802451 | MPP5 |
| 14 | 67328524 | 67328681 | 1.059802451 | ATP6V1D |
| 14 | 67473277 | 67473478 | 1.284785801 | TMEM229B |
| 14 | 67562539 | 67562746 | -1.16601845 | PLEKHH1 |
| 15 | 88531105 | 88531296 | -2.04796221 | DET1 |
| 15 | 88626443 | 88626644 | 1.002112995 | AEN |
| 15 | 88881396 | 88881588 | 6.43820924 | HAPLN3 |
| 15 | 88910228 | 88910404 | -6.548604572 | MFGE8 |
| 15 | 89623864 | 89624056 | -6.284334347 | KIF7 |
| 15 | 89623864 | 89624056 | -6.284334347 | TICRR |
| 15 | 89626935 | 89627124 | -2.100256661 | KIF7 |
| 15 | 89626935 | 89627124 | -2.100256661 | TICRR |
| 15 | 89630776 | 89630963 | -3.543731421 | KIF7 |
| 15 | 89630776 | 89630963 | -3.543731421 | TICRR |
| 15 | 89631808 | 89632001 | -1.025468483 | KIF7 |
| 15 | 89683268 | 89683462 | -2.99491011 | PEX11A |
| 15 | 90067181 | 90067379 | 1.910832112 | ZNF710 |
| 15 | 90079858 | 90080067 | 1.985338047 | RP11-617F23.1 |
| 15 | 90079858 | 90080067 | 1.985338047 | ZNF710 |
| 1 | 47432887 | 47433096 | -1.695957639 | FOXD2-AS1 |
| 15 | 90089889 | 90090087 | 7.124890058 | IDH2 |
| 15 | 90229211 | 90229430 | 1.755909633 | SEMA4B |
| 15 | 90240864 | 90241018 | -1.944575689 | GDPGP1 |
| 15 | 90271635 | 90271841 | -1.154629219 | RP11-697E2.12 |
| 15 | 90271635 | 90271841 | -1.154629219 | NGRN |
| 15 | 90271635 | 90271841 | -1.154629219 | RP11-697E2.6 |
| 15 | 90275586 | 90275785 | -1.516327592 | RP11-697E2.12 |
| 15 | 90275586 | 90275785 | -1.516327592 | RP11-697E2.7 |
| 15 | 90275586 | 90275785 | -1.516327592 | NGRN |
| 15 | 90360482 | 90360671 | -1.885940184 | ZNF774 |
| 15 | 90749677 | 90749918 | -1.702819401 | BLM |
| 15 | 90760898 | 90761085 | -2.983886419 | BLM |
| 1 | 47433544 | 47433745 | -1.67797551 | FOXD2-AS1 |
| 15 | 90961116 | 90961317 | 1.285052514 | RCCD1 |
| 15 | 92789105 | 92789377 | -3.849419103 | FAM174B |
| 15 | 92897755 | 92897951 | -1.353435504 | RP11-437B10.1 |
| 15 | 92897755 | 92897951 | -1.353435504 | LINC01578 |
| 15 | 93024453 | 93024766 | -1.501628037 | CHD2 |
| 15 | 96331733 | 96331915 | -4.384092162 | NR2F2 |
| 15 | 96337569 | 96337762 | -1.611971251 | NR2F2 |
| 1 | 50405737 | 50405841 | -6.977143602 | FCF1P6 |
| 15 | 97674535 | 97674728 | -3.033941236 | LINC00923 |
| 15 | 97874219 | 97874428 | -2.483716069 | LINC00923 |
| 15 | 98707807 | 98708010 | -1.398926016 | IGF1R |
| 15 | 98958126 | 98958327 | -2.224233307 | IGF1R |
| 15 | 98958126 | 98958327 | -2.224233307 | RP11-654A16.3 |
| 15 | 98961649 | 98961827 | -2.309971806 | IGF1R |
| 15 | 98961649 | 98961827 | -2.309971806 | RP11-654A16.3 |
| 15 | 98962407 | 98962610 | -2.714561072 | RP11-654A16.3 |
| 15 | 98962407 | 98962610 | -2.714561072 | IGF1R |
| 15 | 99113972 | 99114154 | 5.320122468 | SYNM |
| 15 | 99132580 | 99132773 | -2.027433725 | SYNM |
| 15 | 99133972 | 99134160 | -1.240816627 | SYNM |
| 15 | 99368623 | 99368809 | -1.617587301 | LRRC28 |
| 15 | 99386099 | 99386389 | -1.054558133 | LRRC28 |
| 15 | 99678584 | 99678782 | -8.705656183 | MEF2A |
| 15 | 99717166 | 99717333 | -1.790702341 | LYSMD4 |
| 15 | 99799858 | 99800140 | -1.080675133 | DNM1P46 |

| | | | | |
|---|---|---|---|---|
| 14 | 67727695 | 67727799 | 5.608372929 | ZFYVE26 |
| 14 | 67727695 | 67727799 | 5.608372929 | RDH12 |
| 14 | 67729109 | 67729304 | 8.890964779 | ZFYVE26 |
| 14 | 67729109 | 67729304 | 8.890964779 | RDH12 |
| 14 | 68497220 | 68497379 | -1.785315105 | RAD51B |
| 14 | 68679714 | 68679908 | 3.115696317 | RAD51B |
| 14 | 68789524 | 68789729 | 6.127208215 | ZFP36L1 |
| 14 | 68887892 | 68888048 | -8.694681228 | ACTN1 |
| 14 | 68887892 | 68888048 | -8.694681228 | HMGN1P3 |
| 14 | 69708803 | 69709004 | 1.383700658 | SUSD6 |
| 1 | 40861262 | 40861464 | -1.433966381 | CITED4 |
| 14 | 70906987 | 70907195 | 1.748685051 | RP1-292L20.3 |
| 14 | 70978351 | 70978555 | 1.073535735 | PCNX1 |
| 14 | 71014759 | 71014971 | 1.842881924 | PCNX1 |
| 14 | 71061736 | 71061877 | 3.551643904 | PCNX1 |
| 14 | 71092596 | 71092797 | 3.538710902 | PCNX1 |
| 14 | 71109769 | 71109968 | 1.706401247 | PCNX1 |
| 14 | 71588402 | 71588580 | 1.383315688 | SIPA1L1 |
| 14 | 72671063 | 72671182 | 1.279827833 | DPF3 |
| 14 | 73058402 | 73058534 | -3.253381344 | RP11-109N23.4 |
| 14 | 73058402 | 73058534 | -3.253381344 | RBM25 |
| 14 | 73239507 | 73239713 | 7.067533973 | PAPLN |
| 1 | 41132575 | 41132723 | -2.831443285 | SCMH1 |
| 14 | 73276504 | 73276703 | 1.972200204 | NUMB |
| 14 | 73569221 | 73569439 | 1.99308709 | ACOT2 |
| 14 | 73595195 | 73595388 | -2.370225059 | ACOT4 |
| 1 | 41165762 | 41165964 | -1.847612862 | SCMH1 |
| 14 | 73718602 | 73718863 | 1.728613872 | ELMSAN1 |
| 14 | 74500295 | 74500487 | 3.174038847 | LTBP2 |
| 1 | 41246816 | 41246969 | -2.006708722 | RP11-399E6.1 |
| 14 | 74781390 | 74781594 | -1.783275519 | YLPM1 |
| 14 | 74840726 | 74840949 | -1.603041116 | YLPM1 |
| 14 | 74906591 | 74906781 | -1.957759336 | RPS6KL1 |
| 14 | 75049235 | 75049424 | -1.358616102 | MLH3 |
| 14 | 75071057 | 75071249 | -6.975460176 | ZC2HC1C |
| 1 | 41510696 | 41510965 | 1.294612052 | HIVEP3 |
| 14 | 75546665 | 75546859 | -2.792088253 | BATF |
| 14 | 75958898 | 75959049 | -1.228837915 | TGFB3 |
| 14 | 75958898 | 75959049 | -1.228837915 | IFT43 |
| 14 | 76166481 | 76166678 | -1.95752036 | GPATCH2L |
| 14 | 76206317 | 76206525 | 1.085442983 | GPATCH2L |
| 1 | 41513056 | 41513230 | 1.475618154 | HIVEP3 |
| 14 | 76782637 | 76782836 | 4.671211973 | RP11-488C13.5 |
| 14 | 76782637 | 76782836 | 4.671211973 | VASH1 |
| 14 | 77025365 | 77025564 | 3.161329242 | IRF2BPL |
| 14 | 77026328 | 77026520 | 2.269718429 | IRF2BPL |
| 14 | 77277203 | 77277403 | -1.347220489 | POMT2 |
| 14 | 77341724 | 77341922 | -1.78821277 | TMED8 |
| 14 | 77469059 | 77469328 | -1.599510513 | AHSA1 |
| 1 | 41513573 | 41513761 | 1.201488763 | HIVEP3 |
| 14 | 80480779 | 80480920 | -7.521107556 | CEP128 |
| 14 | 80706659 | 80706858 | -3.442943496 | CEP128 |
| 14 | 80777963 | 80778156 | -3.012149024 | CEP128 |
| 14 | 80785082 | 80785253 | -3.942183992 | CEP128 |
| 15 | 100225581 | 100225840 | 1.084787861 | ADAMTS17 |
| 15 | 100349260 | 100349382 | 4.914861368 | SPATA41 |
| 15 | 100623568 | 100623738 | -3.928990498 | ASB7 |
| 1 | 50905906 | 50906107 | -7.100142803 | FAF1 |
| 15 | 101068789 | 101068906 | 1.54637857 | RP11-505E24.2 |
| 15 | 101068789 | 101068906 | 1.54637857 | LRRK1 |
| 1 | 50908674 | 50908990 | -1.41219322 | FAF1 |
| 15 | 101188112 | 101188272 | -1.964154389 | CHSY1 |
| 15 | 101301958 | 101302145 | -1.696124041 | RP11-299G20.2 |
| 15 | 101301958 | 101302145 | -1.696124041 | PCSK6 |
| 15 | 101302867 | 101303177 | -1.66632726 | RP11-299G20.2 |
| 15 | 101302867 | 101303177 | -1.66632726 | PCSK6 |
| 15 | 101304756 | 101305036 | -1.529071387 | RP11-299G20.2 |
| 15 | 101304756 | 101305036 | -1.529071387 | PCSK6 |
| 15 | 101309832 | 101310032 | -2.623798835 | PCSK6 |
| 15 | 101328856 | 101329086 | -1.452736774 | PCSK6 |
| 15 | 101329584 | 101329799 | -2.284853937 | PCSK6 |
| 15 | 101490548 | 101490696 | -1.232474312 | PCSK6 |
| 15 | 101491390 | 101491596 | -1.23704066 | PCSK6 |
| 15 | 101492425 | 101492625 | -1.098472426 | PCSK6 |
| 15 | 101746314 | 101746519 | -1.116745065 | RP11-89K11.1 |
| 15 | 101751204 | 101751540 | -1.318620304 | DNM1P47 |
| 15 | 101764641 | 101764845 | -5.499295215 | DNM1P47 |
| 1 | 50969015 | 50969303 | 5.957379419 | CDKN2C |
| 15 | 101976319 | 101976471 | -1.14082225 | WASH3P |
| 16 | 112672 | 112822 | -6.175745705 | NPRL3 |
| 16 | 297961 | 298144 | 7.522431835 | AXIN1 |
| 1 | 50969873 | 50970072 | -1.284932576 | CDKN2C |
| 16 | 520631 | 520741 | 1.483872446 | RAB11FIP3 |
| 16 | 525442 | 525688 | -1.050836771 | LINC00235 |
| 16 | 553454 | 553650 | 1.879172115 | LA16c-366D1.3 |
| 16 | 553454 | 553650 | 1.879172115 | CAPN15 |
| 16 | 614654 | 614815 | -6.404079886 | RAB40C |
| 16 | 708799 | 708987 | -3.300204264 | LA16c-380A1.2 |
| 16 | 735232 | 735389 | -2.661701891 | HAGHL |
| 16 | 735232 | 735389 | -2.661701891 | NARFL |
| 16 | 865673 | 865874 | -1.660639482 | LMF1 |
| 16 | 888083 | 888416 | 1.282727666 | LMF1 |
| 16 | 1257864 | 1257984 | -1.081361382 | RP11-616M22.5 |
| 16 | 1257864 | 1257984 | -1.081361382 | TPSD1 |
| 16 | 1317529 | 1317653 | -7.483613129 | UBE2I |
| 16 | 1434299 | 1434487 | 5.93518236 | CCDC154 |
| 16 | 1434861 | 1434963 | 5.696158711 | CCDC154 |
| 16 | 1445897 | 1446100 | -1.437925696 | LA16c-390E6.5 |
| 16 | 1445897 | 1446100 | -1.437925696 | CLCN7 |
| 16 | 1938740 | 1938940 | 2.730464119 | MSRB1 |
| 16 | 2037980 | 2038180 | -1.911000849 | SLC9A3R2 |
| 16 | 2111369 | 2111568 | 8.64129085 | PKD1 |
| 16 | 2208787 | 2208892 | -4.973919917 | MLST8 |
| 16 | 2213295 | 2213515 | -1.368957628 | PGP |
| 16 | 2231151 | 2231324 | -5.745926548 | E4F1 |
| 16 | 2456764 | 2456968 | -1.946347026 | RP11-715J22.4 |
| 16 | 2456764 | 2456968 | -1.946347026 | CCNF |
| 16 | 2502552 | 2502778 | -1.881354244 | TBC1D24 |

| | | | | | | | | | |
|---|---|---|---|---|---|---|---|---|---|
| 14 | 81246202 | 81246401 | -2.15556346 | DYNLL1P1 | 16 | 2502552 | 2502778 | -1.881354244 | RP11-20I23.1 |
| 14 | 85562749 | 85562916 | 7.0670527 | FLRT2 | 16 | 2563681 | 2563851 | 4.52219706 | PDPK1 |
| 14 | 85621242 | 85621439 | 6.516378024 | FLRT2 | 16 | 2563681 | 2563851 | 4.52219706 | RP11-20I23.8 |
| 14 | 85623281 | 85623481 | 4.503819257 | FLRT2 | 16 | 2566803 | 2567040 | 1.832566425 | PDPK1 |
| 14 | 88010651 | 88010791 | 2.108217483 | RP11-300J18.2 | 16 | 2602294 | 2602483 | -1.211174947 | PDPK1 |
| 14 | 88010651 | 88010791 | 2.108217483 | GPR65 | 16 | 2948902 | 2949103 | -1.14339048 | LA16c-321D4.2 |
| 14 | 88480005 | 88480205 | 2.933105082 | PTPN21 | 16 | 2948902 | 2949103 | -1.14339048 | FLYWCH1 |
| 1 | 41543844 | 41544050 | 1.782581271 | HIVEP3 | 16 | 2968167 | 2968285 | -2.109023476 | PKMYT1 |
| 1 | 41543844 | 41544050 | 1.782581271 | RP11-486B10.4 | 16 | 2968167 | 2968285 | -2.109023476 | KREMEN2 |
| 14 | 89215554 | 89215725 | -2.973008463 | FOXN3 | 16 | 3013082 | 3013371 | -1.667351978 | CLDN9 |
| 14 | 89290373 | 89290616 | -1.019042232 | FOXN3 | 16 | 3053394 | 3053525 | 3.187658341 | MMP25-AS1 |
| 14 | 89290373 | 89290616 | -1.019042232 | CAP2P1 | 16 | 3053394 | 3053525 | 3.187658341 | MMP25 |
| 14 | 89963397 | 89963605 | -1.648088416 | TDP1 | 16 | 3072904 | 3073099 | 7.447077304 | IL32 |
| 14 | 90184761 | 90184935 | 5.134068286 | KCNK13 | 16 | 3084825 | 3085008 | -1.379894619 | RP11-473M20.9 |
| 1 | 41549152 | 41549349 | 1.454283403 | HIVEP3 | 16 | 3131951 | 3132150 | -1.553815179 | ZNF213 |
| 1 | 41549558 | 41549758 | 1.660522249 | HIVEP3 | 16 | 3131951 | 3132150 | -1.553815179 | ZNF213-AS1 |
| 14 | 92005720 | 92005920 | -1.053353653 | TRIP11 | 16 | 3140599 | 3140778 | -1.471579985 | ZNF213 |
| 14 | 92652241 | 92652444 | 1.246611037 | RIN3 | 16 | 3224159 | 3224360 | 1.094974716 | AJ003147.9 |
| 1 | 41551920 | 41552117 | 4.032372191 | HIVEP3 | 16 | 3224159 | 3224360 | 1.094974716 | ZNF200 |
| 14 | 92941491 | 92941666 | 6.077409055 | ITPK1 | 16 | 3242912 | 3243107 | 2.816282232 | MEFV |
| 14 | 93184149 | 93184351 | 2.211869519 | MOAP1 | 16 | 3243294 | 3243492 | 2.574880239 | MEFV |
| 14 | 93206776 | 93206929 | -1.964166159 | GON7 | 16 | 3282592 | 3282785 | -1.394233812 | ZNF263 |
| 14 | 93206776 | 93206929 | -1.964166159 | RP11-371E8.4 | 16 | 3282989 | 3283150 | -1.56154207 | ZNF263 |
| 14 | 93333431 | 93333593 | 1.884693536 | UNC79 | 1 | 52237692 | 52237814 | -2.95217388 | ZFYVE9 |
| 1 | 41552654 | 41552852 | 2.336280294 | HIVEP3 | 16 | 3299262 | 3299462 | -1.077130943 | ZNF263 |
| 14 | 94062313 | 94062522 | -1.253502028 | DDX24 | 16 | 3299262 | 3299462 | -1.077130943 | TIGD7 |
| 14 | 94079580 | 94079782 | -1.425600942 | DDX24 | 16 | 3300352 | 3300540 | -1.789450374 | TIGD7 |
| 14 | 95417082 | 95417303 | -1.06261639 | SYNE3 | 16 | 3300352 | 3300540 | -1.789450374 | ZNF263 |
| 1 | 41555806 | 41556004 | 2.522236619 | HIVEP3 | 16 | 3300925 | 3301125 | -2.215519824 | ZNF263 |
| 14 | 95543945 | 95544154 | -2.472625399 | GLRX5 | 16 | 3300925 | 3301125 | -2.215519824 | TIGD7 |
| 14 | 96354378 | 96354588 | 1.828950566 | ATG2B | 16 | 3308727 | 3308831 | -1.562139411 | ZNF75A |
| 14 | 96355301 | 96355409 | 1.221937404 | ATG2B | 16 | 3308727 | 3308831 | -1.562139411 | LA16c-360H6.2 |
| 14 | 96501731 | 96501886 | -7.640204156 | PAPOLA | 16 | 3310517 | 3310682 | 2.809804322 | ZNF75A |
| 14 | 96501731 | 96501886 | -7.640204156 | RP11-872J21.3 | 16 | 3318284 | 3318482 | -1.620170787 | ZNF75A |
| 14 | 96931339 | 96931528 | -7.186908375 | AL133168.3 | 1 | 52238551 | 52238731 | -2.199702877 | ZFYVE9 |
| 14 | 96931339 | 96931528 | -7.186908375 | VRK1 | 16 | 3401677 | 3401829 | -1.646813493 | ZNF174 |
| 1 | 41556506 | 41556894 | 2.052483451 | HIVEP3 | 16 | 3401677 | 3401829 | -1.646813493 | LA16c-306E5.2 |
| 14 | 99708816 | 99708963 | -2.351328772 | CYP46A1 | 16 | 3408922 | 3409121 | -1.496158583 | ZNF174 |
| 14 | 99713409 | 99713574 | -3.748486315 | CYP46A1 | 16 | 3408922 | 3409121 | -1.496158583 | LA16c-306E5.2 |
| 14 | 99713972 | 99714154 | -4.497637076 | CYP46A1 | 16 | 3412090 | 3412271 | -5.917630459 | LA16c-306E5.2 |
| 14 | 99714784 | 99714922 | -6.376294859 | CYP46A1 | 16 | 3436483 | 3436706 | -1.406693726 | ZNF597 |
| 14 | 100302681 | 100302880 | 3.380775616 | SLC25A29 | 16 | 3436483 | 3436706 | -1.406693726 | LA16c-306E5.2 |
| 14 | 100329186 | 100329348 | 2.331482506 | SLC25A47 | 16 | 3485034 | 3485218 | -1.336053333 | LA16c-306E5.3 |
| 1 | 41566517 | 41566714 | 2.070931509 | HIVEP3 | 16 | 3485034 | 3485218 | -1.336053333 | NAA60 |
| 14 | 101925331 | 101925478 | -1.129270918 | PPP2R5C | 16 | 3583230 | 3583419 | -1.017076757 | RP11-461A8.1 |
| 14 | 101948756 | 101948911 | -2.35536216 | RP11-1017G21.5 | 16 | 3583230 | 3583419 | -1.017076757 | SLX4 |
| 14 | 101986315 | 101986579 | -1.315315921 | DYNC1H1 | 16 | 3664899 | 3665099 | -7.79902625 | TRAP1 |
| 14 | 102225648 | 102225845 | 1.401571335 | MOK | 16 | 3664899 | 3665099 | -7.79902625 | DNASE1 |
| 1 | 41580406 | 41580590 | 1.107906177 | HIVEP3 | 16 | 3965882 | 3966092 | -1.749180576 | ADCY9 |
| 1 | 41581388 | 41581542 | 1.023501883 | HIVEP3 | 1 | 1615504 | 1615678 | 4.966845327 | MIB2 |
| 14 | 103534709 | 103534912 | -2.290855848 | TRMT61A | 1 | 1615504 | 1615678 | 4.966845327 | RP11-345P4.9 |
| 1 | 41585030 | 41585241 | 1.251484633 | HIVEP3 | 16 | 3993653 | 3993755 | -7.495140638 | ADCY9 |
| 14 | 103590504 | 103590710 | -1.274305723 | RP11-73M18.2 | 16 | 4113853 | 4114042 | -1.545819088 | ADCY9 |
| 14 | 103590504 | 103590710 | -1.274305723 | KLC1 | 16 | 4114345 | 4114533 | -1.400118466 | ADCY9 |

| | | | | | | | | | |
|---|---|---|---|---|---|---|---|---|---|
| 14 | 103590504 | 103590710 | -1.274305723 | APOPT1 | 16 | 4246261 | 4246456 | -2.549635663 | LINC01569 |
| 14 | 103705063 | 103705263 | -6.628529235 | XRCC3 | 16 | 4381961 | 4382160 | -6.04637857 | CORO7-PAM16 |
| 14 | 103705063 | 103705263 | -6.628529235 | KLC1 | 16 | 4381961 | 4382160 | -6.04637857 | VASN |
| 14 | 103712340 | 103712679 | 7.395912122 | XRCC3 | 16 | 4381961 | 4382160 | -6.04637857 | CORO7 |
| 14 | 103712340 | 103712679 | 7.395912122 | KLC1 | 16 | 4461786 | 4461989 | -1.135871169 | NMRAL1 |
| 14 | 103713418 | 103713598 | 6.826366479 | XRCC3 | 1 | 52389253 | 52389451 | -1.380406168 | ORC1 |
| 14 | 103713418 | 103713598 | 6.826366479 | KLC1 | 16 | 4609463 | 4609659 | -1.139300811 | UBALD1 |
| 14 | 103740004 | 103740199 | 1.924129619 | PPP1R13B | 16 | 4752166 | 4752390 | -1.010378039 | RP11-127I20.7 |
| 14 | 104704130 | 104704319 | -7.680149213 | INF2 | 16 | 4752166 | 4752390 | -1.010378039 | ZNF500 |
| 14 | 104719716 | 104719917 | -1.890925349 | INF2 | 16 | 4882664 | 4882784 | -1.404535697 | PPL |
| 14 | 104721545 | 104721776 | -1.872215145 | INF2 | 1 | 52393498 | 52393698 | -1.306394913 | ORC1 |
| 14 | 104889474 | 104889645 | -6.926118442 | CEP170B | 16 | 5025050 | 5025249 | -1.819793686 | NAGPA |
| 14 | 104938006 | 104938243 | 2.399172408 | AHNAK2 | 16 | 5025050 | 5025249 | -1.819793686 | RP11-165E7.1 |
| 14 | 104938752 | 104938904 | 3.862162181 | AHNAK2 | 16 | 5085111 | 5085302 | -1.927675285 | ALG1 |
| 1 | 1385962 | 1386107 | -1.50267117 | CCNL2 | 16 | 5085111 | 5085302 | -1.927675285 | EEF2KMT |
| 14 | 104941668 | 104942197 | 2.528318458 | AHNAK2 | 16 | 8714786 | 8714963 | -2.739195418 | ABAT |
| 14 | 104947326 | 104947475 | 1.430195971 | AHNAK2 | 16 | 8815737 | 8815842 | 2.479489553 | PMM2 |
| 14 | 104948391 | 104948513 | 1.21593499 | AHNAK2 | 16 | 8855031 | 8855237 | -1.101294594 | RP11-77H9.2 |
| 14 | 104951459 | 104951589 | 1.641170605 | AHNAK2 | 16 | 8855031 | 8855237 | -1.101294594 | CARHSP1 |
| 14 | 104952290 | 104952600 | 1.208845675 | AHNAK2 | 1 | 52417738 | 52417941 | -1.289525699 | PRPF38A |
| 14 | 104953883 | 104954024 | 3.658040816 | AHNAK2 | 1 | 52417738 | 52417941 | -1.289525699 | ZCCHC11 |
| 14 | 104994098 | 104994299 | -1.438833671 | C14orf79 | 16 | 8858515 | 8858715 | 1.136553716 | RP11-77H9.2 |
| 14 | 105006391 | 105006578 | -1.799788073 | C14orf79 | 16 | 8858515 | 8858715 | 1.136553716 | CARHSP1 |
| 14 | 105011258 | 105011463 | -1.551296237 | CDCA4 | 16 | 8859983 | 8860222 | 1.495646483 | CARHSP1 |
| 14 | 105050822 | 105051022 | -1.006997031 | GPR132 | 16 | 8859983 | 8860222 | 1.495646483 | RP11-77H9.2 |
| 14 | 105055565 | 105055764 | -1.044703707 | GPR132 | 16 | 8864264 | 8864442 | 3.490466461 | CARHSP1 |
| 14 | 105247368 | 105247545 | -2.55749674 | BRF1 | 16 | 10469160 | 10469282 | -1.902746198 | ATF7IP2 |
| 14 | 105250422 | 105250621 | -1.026459124 | BRF1 | 16 | 10906500 | 10906690 | 7.931875358 | CIITA |
| 14 | 105250422 | 105250621 | -1.026459124 | BTBD6 | 1 | 52475096 | 52475296 | -1.148395797 | ZCCHC11 |
| 14 | 105349788 | 105349989 | -1.050137285 | PACS2 | 16 | 10907853 | 10908006 | 8.641761243 | CIITA |
| 14 | 105371268 | 105371467 | -3.971157281 | RP11-521B24.4 | 16 | 10924593 | 10924778 | 9.252748787 | CIITA |
| 14 | 105371268 | 105371467 | -3.971157281 | PACS2 | 16 | 11549408 | 11549606 | 8.102763001 | LITAF |
| 14 | 105375118 | 105375335 | -7.316355632 | PACS2 | 1 | 52525723 | 52525910 | -1.679912583 | ZCCHC11 |
| 14 | 105387884 | 105388068 | -1.836283459 | PACS2 | 16 | 11676467 | 11676670 | 2.398943129 | SNN |
| 1 | 42653931 | 42654130 | 3.275459519 | CCDC30 | 16 | 11797904 | 11798106 | -2.235618747 | RP11-486I11.2 |
| 14 | 105401393 | 105401575 | -1.541970016 | TEX22 | 16 | 12051915 | 12052112 | -1.182581306 | SNX29 |
| 14 | 105416831 | 105416959 | -3.035305611 | TEX22 | 16 | 12569307 | 12569501 | -1.481425355 | CTD-3037G24.3 |
| 14 | 105416831 | 105416959 | -3.035305611 | RP11-521B24.3 | 16 | 12569307 | 12569501 | -1.481425355 | SNX29 |
| 14 | 105442660 | 105442858 | -3.638212034 | MTA1 | 16 | 13935170 | 13935416 | -1.640140669 | CTD-2135D7.2 |
| 14 | 105442660 | 105442858 | -3.638212034 | TEX22 | 16 | 13935170 | 13935416 | -1.640140669 | ERCC4 |
| 14 | 105498930 | 105499141 | -3.713898469 | C14orf80 | 1 | 52603309 | 52603606 | -5.754640084 | GPX7 |
| 15 | 21294182 | 21294371 | 2.061782166 | RP11-32B5.1 | 16 | 13947935 | 13948118 | -1.090375429 | ERCC4 |
| 1 | 42746479 | 42746692 | -1.169042752 | P3H1 | 16 | 14246714 | 14246897 | -8.258380941 | MKL2 |
| 15 | 25360454 | 25360655 | -1.133775539 | UBE3A | 16 | 14261139 | 14261476 | -2.78300625 | MKL2 |
| 15 | 25360454 | 25360655 | -1.133775539 | SNHG14 | 16 | 15125933 | 15126127 | -1.267290156 | PKD1P6-NPIPP1 |
| 15 | 25679353 | 25679554 | 1.435806297 | ATP10A | 16 | 15125933 | 15126127 | -1.267290156 | PDXDC1 |
| 15 | 28111687 | 28111891 | -1.070522174 | HERC2 | 16 | 15125933 | 15126127 | -1.267290156 | PKD1P6 |
| 15 | 28158214 | 28158325 | -1.273743898 | HERC2 | 16 | 15594752 | 15594951 | -1.866199492 | KIAA0430 |
| 15 | 28684499 | 28684696 | -1.026202216 | HERC2P9 | 16 | 15594752 | 15594951 | -1.866199492 | C16orf45 |
| 15 | 28702565 | 28702759 | 1.007959679 | GOLGA8M | 1 | 52687839 | 52688043 | -1.264631174 | COA7 |
| 15 | 28788564 | 28788702 | -1.655707434 | RP11-578F21.10 | 16 | 15635991 | 15636182 | -7.94807158 | KIAA0430 |
| 15 | 28789582 | 28789786 | -2.3363954 | PDCD6IPP2 | 16 | 15701449 | 15701650 | -1.633249389 | NDE1 |
| 15 | 28789582 | 28789786 | -2.3363954 | RP11-578F21.10 | 16 | 15701449 | 15701650 | -1.633249389 | CTB-193M12.5 |
| 15 | 29054278 | 29054481 | -1.03189494 | APBA2 | 16 | 15724369 | 15724588 | -2.361661844 | MYH11 |

| | | | | |
|---|---|---|---|---|
| 15 | 29268784 | 29268987 | -1.280863637 | FAM189A1 |
| 15 | 29268784 | 29268987 | -1.280863637 | NSMCE3 |
| 15 | 30489408 | 30489612 | -1.331702349 | CTD-3092A11.1 |
| 15 | 30489408 | 30489612 | -1.331702349 | CTD-3092A11.2 |
| 15 | 30626318 | 30626515 | -3.656399623 | ARHGAP11B |
| 15 | 30678252 | 30678448 | -6.226217154 | ARHGAP11B |
| 15 | 30736283 | 30736443 | -4.717389395 | ARHGAP11B |
| 15 | 30738216 | 30738428 | -3.281380244 | ARHGAP11B |
| 15 | 30741707 | 30741894 | -2.198135007 | ARHGAP11B |
| 15 | 32615058 | 32615258 | -1.000693966 | RP11-1000B6.8 |
| 15 | 32615058 | 32615258 | -1.000693966 | ARHGAP11A |
| 15 | 32615058 | 32615258 | -1.000693966 | RP11-1000B6.5 |
| 15 | 32636493 | 32636694 | -2.846054591 | ARHGAP11A |
| 15 | 34233338 | 34233570 | 3.000751589 | SLC12A6 |
| 15 | 34287976 | 34288117 | 3.143198145 | SLC12A6 |
| 15 | 34337032 | 34337232 | 1.799328379 | SLC12A6 |
| 15 | 35084268 | 35084430 | 4.836654538 | NANOGP8 |
| 15 | 35092377 | 35092570 | 2.830261148 | RP11-323I15.2 |
| 15 | 35391374 | 35391524 | -6.682682783 | DPH6 |
| 15 | 38351104 | 38351282 | 6.5746354 | SPRED1 |
| 15 | 40089458 | 40089665 | 2.030979178 | BMF |
| 15 | 40090972 | 40091172 | 3.727360332 | BMF |
| 15 | 40335222 | 40335422 | 2.201317118 | C15orf52 |
| 15 | 40369742 | 40369884 | -1.561637454 | DISP2 |
| 1 | 43409863 | 43410184 | -1.270290632 | SZT2 |
| 15 | 40393726 | 40393932 | -1.874592554 | KNSTRN |
| 15 | 40420025 | 40420222 | -1.437437859 | IVD |
| 15 | 40471510 | 40471708 | -2.294349433 | CHST14 |
| 15 | 40472417 | 40472626 | -2.353102127 | CHST14 |
| 15 | 40521147 | 40521272 | -6.143567593 | RP11-111A22.1 |
| 15 | 40569336 | 40569515 | -2.470373002 | RPUSD2 |
| 15 | 40573863 | 40574066 | -2.143223529 | RPUSD2 |
| 15 | 40621206 | 40621406 | -3.618625638 | KNL1 |
| 1 | 43413534 | 43413731 | -8.401737643 | SZT2 |
| 15 | 40621845 | 40622092 | -3.523530604 | KNL1 |
| 15 | 40622970 | 40623089 | -3.55087941 | KNL1 |
| 15 | 40623530 | 40623732 | -3.748404516 | KNL1 |
| 15 | 40856967 | 40857198 | 2.365093531 | SPINT1 |
| 15 | 40936630 | 40936811 | -5.061454431 | DLL4 |
| 15 | 40938006 | 40938206 | -4.403367394 | DLL4 |
| 15 | 41394953 | 41395145 | -1.801256649 | NDUFAF1 |
| 15 | 41396589 | 41396794 | -1.120662637 | NDUFAF1 |
| 15 | 41480616 | 41480816 | -1.207106524 | RTF1 |
| 15 | 41503108 | 41503308 | -1.267979069 | ITPKA |
| 15 | 41696110 | 41696221 | -1.211536784 | MGA |
| 15 | 41696654 | 41696853 | -1.849797473 | MGA |
| 1 | 43590957 | 43591160 | -2.107986267 | PTPRF |
| 15 | 41736404 | 41736603 | -1.38872112 | MGA |
| 15 | 41749297 | 41749492 | -1.357395645 | MGA |
| 15 | 41750304 | 41750514 | -1.295218609 | MGA |
| 15 | 41766235 | 41766436 | -1.150318354 | MGA |
| 15 | 41831424 | 41831630 | -2.49265498 | JMJD7-PLA2G4B |
| 15 | 41831424 | 41831630 | -2.49265498 | JMJD7 |
| 15 | 41900450 | 41900647 | -1.590439376 | EHD4 |

| | | | | |
|---|---|---|---|---|
| 16 | 15724369 | 15724588 | -2.361661844 | NDE1 |
| 16 | 15726703 | 15726874 | -2.516207069 | AF001548.5 |
| 16 | 15726703 | 15726874 | -2.516207069 | MYH11 |
| 16 | 16103467 | 16103680 | 1.815860369 | ABCC1 |
| 16 | 16149984 | 16150226 | -2.094099771 | ABCC6 |
| 16 | 16364884 | 16365040 | -3.5 | RP11-958N24.2 |
| 1 | 52771148 | 52771347 | -2.01057112 | ZYG11B |
| 16 | 17107503 | 17107699 | -1.563859574 | XYLT1 |
| 16 | 17108349 | 17108530 | -1.086972328 | XYLT1 |
| 16 | 17117711 | 17117927 | -2.242031461 | XYLT1 |
| 16 | 17236791 | 17236985 | -1.495825727 | XYLT1 |
| 16 | 17237776 | 17237973 | -7.648476022 | XYLT1 |
| 16 | 17259250 | 17259499 | -1.513176971 | XYLT1 |
| 16 | 18809221 | 18809421 | 1.212091263 | SMG1 |
| 16 | 18809221 | 18809421 | 1.212091263 | RP11-1035H13.2 |
| 16 | 18822213 | 18822417 | 1.098985196 | SMG1 |
| 16 | 19115513 | 19115715 | 2.097643557 | ITPRIPL2 |
| 16 | 19115513 | 19115715 | 2.097643557 | CTD-2349B8.1 |
| 16 | 19503379 | 19503644 | -1.195977166 | GDE1 |
| 1 | 1646380 | 1646579 | -6.28788779 | CDK11B |
| 16 | 19705274 | 19705445 | -1.743471483 | KNOP1 |
| 16 | 19705274 | 19705445 | -1.743471483 | C16orf62 |
| 16 | 19706948 | 19707147 | -2.601048934 | KNOP1 |
| 16 | 19706948 | 19707147 | -2.601048934 | AC002550.5 |
| 16 | 20723802 | 20723964 | -1.621727018 | THUMPD1 |
| 16 | 20723802 | 20723964 | -1.621727018 | ACSM3 |
| 16 | 20723802 | 20723964 | -1.621727018 | AC004381.8 |
| 16 | 20736956 | 20737198 | -1.244201479 | ACSM3 |
| 16 | 20736956 | 20737198 | -1.244201479 | THUMPD1 |
| 1 | 52873372 | 52873569 | -4.766664866 | ZYG11A |
| 16 | 21301112 | 21301308 | -6.781359714 | CRYM |
| 16 | 21301112 | 21301308 | -6.781359714 | CRYM-AS1 |
| 16 | 21641255 | 21641458 | -1.242990517 | IGSF6 |
| 16 | 21641255 | 21641458 | -1.242990517 | METTL9 |
| 16 | 22392808 | 22392978 | 2.613781957 | MFSD13B |
| 16 | 22392808 | 22392978 | 2.613781957 | CDR2 |
| 16 | 22393293 | 22393445 | 2.084962501 | MFSD13B |
| 16 | 22393293 | 22393445 | 2.084962501 | CDR2 |
| 1 | 52893057 | 52893299 | -2.340588908 | ZYG11A |
| 16 | 22419881 | 22420090 | 1.869956165 | RRN3P3 |
| 16 | 22419881 | 22420090 | 1.869956165 | CDR2 |
| 16 | 22420909 | 22421116 | 2.554696457 | CDR2 |
| 16 | 22420909 | 22421116 | 2.554696457 | RRN3P3 |
| 16 | 23064437 | 23064582 | -1.562530589 | USP31 |
| 16 | 23467472 | 23467672 | -1.042315942 | GGA2 |
| 1 | 52994900 | 52995006 | -1.506814818 | SCP2 |
| 1 | 52994900 | 52995006 | -1.506814818 | TUBBP10 |
| 16 | 23674101 | 23674334 | -1.414776103 | CTD-2196E14.9 |
| 16 | 23674101 | 23674334 | -1.414776103 | DCTN5 |
| 16 | 23893108 | 23893225 | -1.54397385 | PRKCB |
| 16 | 24540134 | 24540348 | 1.057332185 | RBBP6 |
| 16 | 24570785 | 24570985 | -1.071592915 | RBBP6 |
| 16 | 24791478 | 24791666 | -1.459388313 | TNRC6A |
| 16 | 24930955 | 24931153 | -2.876274441 | ARHGAP17 |

| | | | | |
|---|---|---|---|---|
| 15 | 41908471 | 41908650 | -7.797176883 | CTD-2382E5.6 |
| 15 | 41908471 | 41908650 | -7.797176883 | EHD4 |
| 1 | 43603809 | 43604018 | -4.013843769 | PTPRF |
| 15 | 42273655 | 42273851 | -7.850856284 | TMEM87A |
| 15 | 42273655 | 42273851 | -7.850856284 | GANC |
| 15 | 42299886 | 42300039 | -3.508780414 | GANC |
| 15 | 42304930 | 42305055 | -2.804258614 | GANC |
| 15 | 42336433 | 42336712 | -2.454076604 | GANC |
| 15 | 42684544 | 42684746 | -1.597107102 | STARD9 |
| 15 | 42691704 | 42691904 | -2.304010335 | STARD9 |
| 15 | 42724021 | 42724204 | -1.599622533 | CDAN1 |
| 15 | 42724021 | 42724204 | -1.599622533 | CTD-2036P10.5 |
| 15 | 42745855 | 42746020 | -1.272551007 | TTBK2 |
| 15 | 42752952 | 42753153 | -1.089425964 | TTBK2 |
| 15 | 42775367 | 42775574 | -2.116243628 | TTBK2 |
| 15 | 42919875 | 42920082 | -2.280315147 | TTBK2 |
| 1 | 43691005 | 43691187 | -9.275042699 | KDM4A |
| 15 | 43238915 | 43239059 | -3.871669215 | TGM5 |
| 15 | 43328936 | 43329288 | -2.479744799 | LCMT2 |
| 15 | 43409056 | 43409169 | -1.295092232 | TP53BP1 |
| 15 | 43409056 | 43409169 | -1.295092232 | TUBGCP4 |
| 15 | 43456118 | 43456317 | -1.24824346 | TP53BP1 |
| 15 | 43523096 | 43523296 | -2.007249785 | MAP1A |
| 15 | 43771155 | 43771304 | -9.194218893 | PDIA3 |
| 15 | 43804865 | 43805108 | -1.689135357 | MFAP1 |
| 15 | 43891770 | 43891963 | -7.785150225 | FRMD5 |
| 1 | 1400928 | 1401147 | -2.015220232 | RP4-758J18.2 |
| 1 | 43704541 | 43704742 | -1.503645524 | KDM4A |
| 1 | 43704541 | 43704742 | -1.503645524 | KDM4A-AS1 |
| 15 | 44352268 | 44352448 | -7.856370964 | CASC4 |
| 15 | 44503494 | 44503768 | -1.408934175 | CTDSPL2 |
| 15 | 44524127 | 44524314 | -2.478589359 | CTDSPL2 |
| 15 | 44536108 | 44536309 | -1.033339845 | EIF3J-AS1 |
| 15 | 44731551 | 44731758 | 2.289875653 | TRIM69 |
| 15 | 44767449 | 44767671 | 3.26721224 | TRIM69 |
| 1 | 43863326 | 43863530 | -4.782805104 | ST3GAL3 |
| 15 | 45598823 | 45599019 | 7.698034779 | RP11-96O20.4 |
| 15 | 45598823 | 45599019 | 7.698034779 | BLOC1S6 |
| 1 | 43910455 | 43910652 | -10.48737078 | ST3GAL3 |
| 15 | 48738525 | 48738643 | -2.490448939 | CEP152 |
| 15 | 48756164 | 48756413 | -3.222527574 | CEP152 |
| 15 | 48992480 | 48992682 | 1.544358371 | SECISBP2L |
| 1 | 43912870 | 43913068 | -10.99703772 | ST3GAL3 |
| 15 | 50235952 | 50236146 | -2.13212537 | SLC27A2 |
| 15 | 50355488 | 50355695 | -1.050869181 | GABPB1-AS1 |
| 1 | 43913723 | 43913918 | -10.82318135 | ST3GAL3 |
| 15 | 51057983 | 51058183 | 4.291176007 | RP11-108K3.1 |
| 15 | 51057983 | 51058183 | 4.291176007 | TNFAIP8L3 |
| 15 | 51318654 | 51318807 | 1.48521171 | CYP19A1 |
| 15 | 51318654 | 51318807 | 1.48521171 | RP11-108K3.2 |
| 15 | 51384662 | 51384810 | 6.560604858 | GLDN |
| 15 | 51966184 | 51966374 | -1.316555208 | MAPK6 |
| 15 | 51966184 | 51966374 | -1.316555208 | LEO1 |
| 1 | 43914964 | 43915187 | -5.644990548 | ST3GAL3 |
| 16 | 25240183 | 25240379 | -1.723747956 | ZKSCAN2 |
| 16 | 25256852 | 25257020 | -1.642545663 | ZKSCAN2 |
| 16 | 27210105 | 27210305 | -3.050356932 | KDM8 |
| 16 | 27448880 | 27449106 | -1.728413605 | IL21R |
| 16 | 27448880 | 27449106 | -1.728413605 | IL21R-AS1 |
| 16 | 27740592 | 27740793 | -1.246695184 | KIAA0556 |
| 1 | 53010316 | 53010422 | -4.165458439 | SCP2 |
| 1 | 53210407 | 53210611 | -8.559293137 | RP5-1024G6.2 |
| 1 | 53210407 | 53210611 | -8.559293137 | CPT2 |
| 16 | 29262699 | 29262940 | -1.966828187 | RP11-426C22.6 |
| 16 | 29272242 | 29272446 | -2.907709735 | RP11-426C22.8 |
| 16 | 29364730 | 29364951 | -1.440789754 | SNX29P2 |
| 16 | 29364730 | 29364951 | -1.440789754 | RP11-231C14.7 |
| 16 | 29665879 | 29666077 | -9.226723744 | SPN |
| 16 | 29665879 | 29666077 | -9.226723744 | AC009133.22 |
| 16 | 29743605 | 29743783 | -2.23712104 | C16orf54 |
| 16 | 29744012 | 29744209 | -2.67065397 | C16orf54 |
| 16 | 29819632 | 29819825 | -4.006673904 | PAGR1 |
| 16 | 29819632 | 29819825 | -4.006673904 | CTD-2574D22.6 |
| 16 | 29819632 | 29819825 | -4.006673904 | AC009133.12 |
| 16 | 29819632 | 29819825 | -4.006673904 | AC009133.23 |
| 16 | 29986525 | 29986728 | 1.431863537 | TAOK2 |
| 1 | 53244605 | 53244840 | -1.92704764 | LRP8 |
| 16 | 30397461 | 30397666 | -2.254674022 | ZNF48 |
| 16 | 30521626 | 30521902 | -1.972384322 | ITGAL |
| 1 | 1663428 | 1663629 | -2.354138724 | SLC35E2B |
| 1 | 53767852 | 53768052 | -1.77088284 | NDC1 |
| 16 | 30592397 | 30592579 | -6.01014275 | RP11-146F11.5 |
| 16 | 30779848 | 30780106 | -1.658913379 | ZNF629 |
| 16 | 30851152 | 30851344 | -6.274410954 | BCL7C |
| 1 | 54173409 | 54173623 | -1.332454754 | CYB5RL |
| 1 | 54173409 | 54173623 | -1.332454754 | RP11-446E24.4 |
| 16 | 30957309 | 30957493 | -1.889478463 | SETD1A |
| 16 | 30979492 | 30979689 | 7.524307293 | SETD1A |
| 16 | 31074098 | 31074259 | -2.747705458 | ZNF668 |
| 1 | 54174500 | 54174681 | -2.204399693 | CYB5RL |
| 1 | 54174500 | 54174681 | -2.204399693 | RP11-446E24.4 |
| 16 | 31080870 | 31081050 | -1.080443555 | ZNF646 |
| 16 | 31112068 | 31112267 | 2.672546221 | BCKDK |
| 16 | 31311866 | 31312030 | -6.108372929 | ITGAM |
| 16 | 31358073 | 31358290 | -1.429356638 | ITGAX |
| 16 | 31382017 | 31382216 | -1.073419148 | ITGAX |
| 16 | 31435875 | 31435977 | 3.422745025 | ZNF843 |
| 16 | 31436977 | 31437115 | -2.633056899 | ZNF843 |
| 1 | 54609116 | 54609277 | -2.194106512 | FAM151A |
| 1 | 54609116 | 54609277 | -2.194106512 | ACOT11 |
| 16 | 31450925 | 31451118 | -4.67918824 | RP11-452L6.1 |
| 16 | 31457041 | 31457245 | -1.187388322 | RP11-452L6.5 |
| 16 | 31708116 | 31708258 | 4.812195102 | CLUHP3 |
| 16 | 31764729 | 31764839 | -6.837096134 | ZNF720 |
| 16 | 31772880 | 31773064 | -4.099458602 | ZNF720 |
| 1 | 54609604 | 54609801 | -3.088017297 | FAM151A |
| 1 | 54609604 | 54609801 | -3.088017297 | ACOT11 |
| 16 | 31776810 | 31776970 | -2.500141872 | ZNF720 |

| | | | | | | | | | |
|---|---|---|---|---|---|---|---|---|---|
| 15 | 52381130 | 52381316 | 1.400353009 | MYO5A | 1 | 54653383 | 54653580 | -6.258650151 | MROH7-TTC4 |
| 15 | 52608717 | 52608915 | 1.311874811 | FAM214A | 1 | 54653383 | 54653580 | -6.258650151 | MROH7 |
| 15 | 52609454 | 52609636 | 1.174562882 | FAM214A | 16 | 46581774 | 46581974 | -6.884763977 | SHCBP1 |
| 15 | 55311813 | 55312009 | -2.314271567 | RAB27A | 16 | 46626447 | 46626636 | -4.251912869 | RAB43P1 |
| 15 | 55348472 | 55348671 | 8.546713941 | CCPG1 | 16 | 46802778 | 46802965 | -2.079612324 | C16orf87 |
| 15 | 55348472 | 55348671 | 8.546713941 | PIGB | 16 | 48542893 | 48543084 | -1.163924033 | N4BP1 |
| 15 | 55360419 | 55360642 | 1.982329697 | CCPG1 | 16 | 50211188 | 50211381 | 7.403325155 | PAPD5 |
| 15 | 55360419 | 55360642 | 1.982329697 | DYX1C1-CCPG1 | 16 | 50315371 | 50315620 | -1.432641641 | BRD7 |
| 1 | 43926544 | 43926733 | -10.33384511 | ST3GAL3 | 16 | 50315371 | 50315620 | -1.432641641 | ADCY7 |
| 15 | 56123495 | 56123682 | 1.985536407 | RFX7 | 16 | 50316653 | 50316852 | -1.554797661 | ADCY7 |
| 15 | 56134951 | 56135148 | 1.466144387 | RFX7 | 16 | 50316653 | 50316852 | -1.554797661 | BRD7 |
| 15 | 56399778 | 56399972 | 6.639515068 | TEX9 | 1 | 54757816 | 54758011 | -1.042326046 | PARS2 |
| 1 | 43930850 | 43931064 | -4.022299384 | ST3GAL3 | 16 | 50671785 | 50671940 | 1.422333347 | SNX20 |
| 15 | 56404792 | 56404969 | 3.491021435 | TEX9 | 16 | 50711297 | 50711426 | 6.655328578 | NOD2 |
| 15 | 56436440 | 56436583 | 9.306445045 | TEX9 | 16 | 50736516 | 50736714 | 9.067944806 | RP11-327F22.1 |
| 15 | 56436440 | 56436583 | 9.306445045 | MNS1 | 16 | 50749638 | 50749928 | 2.764619289 | CYLD |
| 15 | 56439864 | 56440061 | 9.585336638 | TEX9 | 16 | 50796385 | 50796581 | 2.318603445 | RP11-327F22.4 |
| 15 | 56439864 | 56440061 | 9.585336638 | MNS1 | 16 | 50796385 | 50796581 | 2.318603445 | CYLD |
| 15 | 56442603 | 56442802 | 8.511313945 | MNS1 | 16 | 53185180 | 53185375 | -1.242319492 | CHD9 |
| 15 | 56442603 | 56442802 | 8.511313945 | TEX9 | 16 | 53252485 | 53252768 | -1.059173637 | CHD9 |
| 15 | 56820564 | 56820670 | 2.035194664 | ZNF280D | 1 | 54758620 | 54758850 | -1.225559597 | PARS2 |
| 15 | 56917101 | 56917257 | -1.559681372 | ZNF280D | 16 | 53279575 | 53279771 | -2.004413659 | CHD9 |
| 1 | 43936845 | 43937047 | -1.869497844 | ARTN | 16 | 53280372 | 53280579 | 1.725330705 | CHD9 |
| 15 | 56917619 | 56917829 | -1.01720691 | ZNF280D | 16 | 53303823 | 53304012 | -1.178216142 | CHD9 |
| 15 | 57300580 | 57300784 | -4.045480322 | LINC00926 | 16 | 53324288 | 53324494 | -1.077732799 | CHD9 |
| 1 | 43947053 | 43947248 | -1.690071354 | IPO13 | 16 | 53844029 | 53844183 | -1.15908798 | FTO |
| 15 | 58184072 | 58184279 | 7.345128605 | AQP9 | 1 | 1665448 | 1665648 | -1.50962253 | SLC35E2B |
| 15 | 58184072 | 58184279 | 7.345128605 | ALDH1A2 | 16 | 54270604 | 54270789 | -2.531504899 | RP11-324D17.1 |
| 15 | 58316144 | 58316338 | 1.248456253 | ALDH1A2 | 16 | 54285181 | 54285381 | -1.490513322 | IRX3 |
| 15 | 58597355 | 58597554 | 1.529831192 | ADAM10 | 16 | 54286528 | 54286689 | -2.800563233 | IRX3 |
| 15 | 58771899 | 58772092 | 1.068354126 | FAM63B | 16 | 54286528 | 54286689 | -2.800563233 | RP11-434E6.5 |
| 15 | 58847342 | 58847540 | 3.930316485 | FAM63B | 16 | 54933667 | 54933869 | -1.164170087 | IRX5 |
| 1 | 43990414 | 43990611 | -2.715604175 | B4GALT2 | 1 | 54852043 | 54852249 | -1.094601275 | DHCR24 |
| 15 | 58851654 | 58851844 | 4.166071622 | FAM63B | 16 | 56990924 | 56991178 | 3.885219018 | NLRC5 |
| 15 | 58854399 | 58854605 | 2.732534705 | FAM63B | 16 | 56991926 | 56992129 | 3.77314365 | NLRC5 |
| 15 | 59125935 | 59126088 | 3.636080523 | RP11-59H7.3 | 16 | 57025466 | 57025770 | 2.286204974 | NLRC5 |
| 15 | 59126521 | 59126724 | 4.339765028 | RP11-59H7.3 | 16 | 57026478 | 57026686 | 2.190380063 | NLRC5 |
| 15 | 59127576 | 59127736 | 4.358838212 | RP11-59H7.3 | 16 | 57082574 | 57082773 | 1.966067603 | NLRC5 |
| 15 | 59128483 | 59128685 | 8.14158669 | RP11-59H7.3 | 16 | 57746375 | 57746534 | -4.669925001 | KATNB1 |
| 15 | 59129183 | 59129385 | 4.423062921 | RP11-59H7.3 | 16 | 57747498 | 57747695 | -4.94055698 | KATNB1 |
| 15 | 59135172 | 59135494 | 1.945931369 | MYO1E | 1 | 55068783 | 55069025 | -1.004632819 | USP24 |
| 15 | 59136027 | 59136227 | 3.436138761 | MYO1E | 16 | 57816669 | 57816871 | -5.725605556 | CTD-2600O9.1 |
| 15 | 59137144 | 59137344 | 2.443255095 | MYO1E | 16 | 57816669 | 57816871 | -5.725605556 | KIFC3 |
| 15 | 59158839 | 59159028 | -8.674588772 | MYO1E | 16 | 57996217 | 57996402 | 1.459584463 | ZNF319 |
| 15 | 59372661 | 59372788 | -6.238373102 | FAM81A | 16 | 58012854 | 58013013 | 2.873297148 | USB1 |
| 15 | 59372661 | 59372788 | -6.238373102 | MYO1E | 16 | 66510748 | 66510964 | -2.419017553 | TK2 |
| 15 | 59372805 | 59372958 | -7.914514482 | FAM81A | 16 | 66510748 | 66510964 | -2.419017553 | RP11-403P17.5 |
| 15 | 59372805 | 59372958 | -7.914514482 | MYO1E | 16 | 66566520 | 66566722 | -1.42926128 | CMTM1 |
| 1 | 44805787 | 44805984 | 5.586838568 | TCTEX1D4 | 16 | 66566520 | 66566722 | -1.42926128 | CKLF-CMTM1 |
| 1 | 44805787 | 44805984 | 5.586838568 | PLK3 | 16 | 66579588 | 66579711 | -6.801639305 | CMTM2 |
| 15 | 60449261 | 60449487 | -2.091046078 | ICE2 | 16 | 66885510 | 66885776 | -2.579888975 | PDP2 |
| 15 | 62019376 | 62019536 | -6.207210491 | VPS13C | 16 | 66921930 | 66922065 | 5.287296764 | RRAD |
| 15 | 63141408 | 63141702 | 1.95311841 | RPS27L | 16 | 67154435 | 67154639 | 1.960196806 | TRADD |
| 15 | 63141408 | 63141702 | 1.95311841 | LACTB | 16 | 67160270 | 67160416 | 3.128568185 | FBXL8 |

| | | | | | | | | | |
|---|---|---|---|---|---|---|---|---|---|
| 15 | 63305694 | 63305879 | -1.135290158 | APH1B | 16 | 67160270 | 67160416 | 3.128568185 | TRADD |
| 1 | 45326954 | 45327192 | -2.007408108 | HPDL | 16 | 67180178 | 67180332 | -6.945607053 | KIAA0895L |
| 15 | 63591140 | 63591355 | 1.161428783 | USP3-AS1 | 16 | 67265922 | 67266046 | -6.331890225 | SLC9A5 |
| 15 | 63591140 | 63591355 | 1.161428783 | USP3 | 16 | 67611190 | 67611439 | -1.219420889 | CTCF |
| 15 | 63601757 | 63601932 | -1.223631619 | FBXL22 | 16 | 67685543 | 67685721 | -3.015678298 | GFOD2 |
| 15 | 63908116 | 63908244 | 6.523902777 | DAPK2 | 16 | 67841916 | 67842123 | 1.342661404 | THAP11 |
| 15 | 64142830 | 64143069 | -2.149501494 | SNX1 | 16 | 67841916 | 67842123 | 1.342661404 | CENPT |
| 15 | 64155929 | 64156130 | -2.118405915 | SNX22 | 16 | 67843502 | 67843683 | -1.296362135 | THAP11 |
| 15 | 64155929 | 64156130 | -2.118405915 | PPIB | 16 | 67843502 | 67843683 | -1.296362135 | CENPT |
| 1 | 45327795 | 45327965 | -2.322569008 | HPDL | 16 | 67944485 | 67944679 | -1.03735387 | SLC12A4 |
| 15 | 64678084 | 64678282 | -1.029293795 | ZNF609 | 16 | 67945797 | 67945976 | 2.670992874 | SLC12A4 |
| 15 | 64815787 | 64815985 | -3.754858664 | PIF1 | 16 | 68021823 | 68022022 | -1.089995984 | DDX28 |
| 15 | 64865159 | 64865307 | 1.578717733 | PLEKHO2 | 16 | 68021823 | 68022022 | -1.089995984 | DUS2 |
| 15 | 64865159 | 64865307 | 1.578717733 | AC069368.3 | 16 | 68122460 | 68122681 | -1.388429385 | NFATC3 |
| 15 | 64865570 | 64865768 | 1.905490703 | AC069368.3 | 16 | 68191068 | 68191228 | -2.770908592 | NFATC3 |
| 15 | 64865570 | 64865768 | 1.905490703 | PLEKHO2 | 16 | 68259648 | 68259848 | -1.398170354 | PLA2G15 |
| 1 | 45343237 | 45343441 | -1.465224368 | TOE1 | 16 | 68259648 | 68259848 | -1.398170354 | RP11-96D1.7 |
| 15 | 64951879 | 64951999 | 1.272439977 | ANKDD1A | 16 | 68298671 | 68298881 | -1.170867 | SLC7A6OS |
| 15 | 65624613 | 65624804 | -1.00912295 | SLC24A1 | 16 | 68298671 | 68298881 | -1.170867 | SLC7A6 |
| 1 | 45345304 | 45345495 | 8.176280375 | TESK2 | 16 | 68563258 | 68563454 | -1.690689826 | ZFP90 |
| 15 | 65652134 | 65652314 | -2.711384036 | SLC24A1 | 16 | 68563938 | 68564150 | -1.569102356 | ZFP90 |
| 15 | 65690809 | 65691013 | -2.250526316 | DENND4A | 16 | 69033718 | 69033853 | -3.453445298 | RPS2P45 |
| 15 | 66325981 | 66326181 | -2.030578831 | RP11-352G18.2 | 16 | 69033718 | 69033853 | -3.453445298 | TANGO6 |
| 15 | 66325981 | 66326181 | -2.030578831 | DIS3L | 16 | 69083676 | 69083793 | -1.698617589 | TANGO6 |
| 15 | 66489815 | 66489996 | -2.388803789 | MAP2K1 | 16 | 69328908 | 69329060 | -1.283768772 | COG8 |
| 15 | 66489815 | 66489996 | -2.388803789 | CTD-3185P2.1 | 16 | 69328908 | 69329060 | -1.283768772 | PDF |
| 15 | 66702477 | 66702671 | -1.58336019 | SMAD6 | 16 | 69334837 | 69335039 | -1.047488316 | COG8 |
| 15 | 67448488 | 67448672 | -1.453484825 | IQCH-AS1 | 16 | 69341559 | 69341766 | 1.068821239 | RP11-343C2.9 |
| 15 | 67448488 | 67448672 | -1.453484825 | IQCH | 16 | 69341559 | 69341766 | 1.068821239 | NIP7 |
| 15 | 67526333 | 67526519 | -1.418794842 | C15orf61 | 16 | 69341559 | 69341766 | 1.068821239 | RP11-343C2.7 |
| 1 | 45607441 | 45607641 | -1.432118243 | NASP | 16 | 69666219 | 69666448 | -5.25042094 | NFAT5 |
| 15 | 69196440 | 69196645 | -1.022965711 | GLCE | 16 | 69666797 | 69666956 | 1.323787239 | NFAT5 |
| 15 | 69403628 | 69403846 | 7.346716253 | PAQR5 | 16 | 69695062 | 69695256 | -1.450247142 | NFAT5 |
| 15 | 69403628 | 69403846 | 7.346716253 | RP11-253M7.1 | 16 | 69742030 | 69742235 | -1.275995479 | NOB1 |
| 15 | 70667358 | 70667505 | -2.127176581 | UACA | 16 | 69742030 | 69742235 | -1.275995479 | CTD-2033A16.3 |
| 1 | 45608033 | 45608238 | -2.99506152 | NASP | 16 | 70160251 | 70160450 | -1.225204588 | RP11-296I10.3 |
| 15 | 70832239 | 70832412 | 1.968231593 | LARP6 | 16 | 70160251 | 70160450 | -1.225204588 | PDPR |
| 15 | 72046451 | 72046660 | -1.529558843 | MYO9A | 16 | 70250851 | 70251079 | -1.020119366 | EXOSC6 |
| 15 | 72474581 | 72474783 | -8.12881362 | ARIH1 | 16 | 70252513 | 70252712 | -2.282989419 | AARS |
| 15 | 72474581 | 72474783 | -8.12881362 | RP11-1007O24.3 | 1 | 62436807 | 62436986 | 5.5691359 | USP1 |
| 15 | 72751673 | 72751869 | -1.545894228 | ADPGK | 16 | 70333552 | 70333756 | -1.604290496 | RP11-529K1.2 |
| 15 | 73302659 | 73302856 | 1.305857817 | NEO1 | 16 | 70333552 | 70333756 | -1.604290496 | DDX19B |
| 15 | 73713102 | 73713305 | 2.498072952 | CD276 | 16 | 70333552 | 70333756 | -1.604290496 | RP11-529K1.3 |
| 15 | 74044404 | 74044594 | 1.782120959 | PML | 16 | 70663471 | 70663670 | -1.59483061 | MTSS1L |
| 1 | 46017260 | 46017453 | 5.867778012 | MAST2 | 16 | 70663471 | 70663670 | -1.59483061 | RP11-394B2.1 |
| 15 | 74410323 | 74410541 | 4.94319001 | SEMA7A | 16 | 70699149 | 70699346 | 8.625412892 | VAC14 |
| 15 | 74461623 | 74461754 | -2.910746065 | UBL7-AS1 | 1 | 1735485 | 1735620 | -2.95706527 | RP1-283E3.8 |
| 15 | 74598662 | 74598860 | -2.999510135 | CLK3 | 1 | 1735485 | 1735620 | -2.95706527 | RP1-283E3.4 |
| 15 | 74601694 | 74601876 | -4.3747392 | CLK3 | 1 | 1735485 | 1735620 | -2.95706527 | SLC35E2 |
| 15 | 74605361 | 74605575 | -2.373622851 | CLK3 | 1 | 62444821 | 62445024 | -1.592682315 | USP1 |
| 15 | 74629240 | 74629343 | -1.171170982 | CLK3 | 16 | 71285287 | 71285446 | -1.516720255 | CMTR2 |
| 15 | 74898949 | 74899165 | -1.741634625 | MPI | 16 | 71449440 | 71449621 | -3.403021869 | AC010547.9 |
| 15 | 74906982 | 74907162 | -4.828837865 | FAM219B | 16 | 71449440 | 71449621 | -3.403021869 | ZNF23 |
| 15 | 74955608 | 74955823 | -1.132548975 | RPP25 | 16 | 71449738 | 71449848 | -1.707781793 | AC010547.9 |

| | | | | | | | | | |
|---|---|---|---|---|---|---|---|---|---|
| 15 | 75341462 | 75341745 | -1.967329786 | COMMD4 | 16 | 71449738 | 71449848 | -1.707781793 | ZNF23 |
| 15 | 75371556 | 75371758 | -1.445019243 | SIN3A | 16 | 71808928 | 71809084 | 2.822504942 | AP1G1 |
| 1 | 46285629 | 46285854 | -1.906983696 | LRRC41 | 16 | 71860029 | 71860236 | -2.31460386 | ZNF821 |
| 15 | 75587274 | 75587405 | -2.727829027 | CTD-2323K18.1 | 16 | 71860029 | 71860236 | -2.31460386 | IST1 |
| 15 | 75639293 | 75639493 | -1.910419341 | IMP3 | 16 | 71860029 | 71860236 | -2.31460386 | ATXN1L |
| 15 | 75675990 | 75676211 | 4.158037154 | CSPG4 | 16 | 71860029 | 71860236 | -2.31460386 | RP11-417N10.4 |
| 15 | 75687398 | 75687613 | 3.053370784 | CSPG4 | 16 | 72024224 | 72024413 | -1.150731996 | DHODH |
| 15 | 75689688 | 75689910 | 3.761274881 | CSPG4 | 1 | 64559236 | 64559433 | 1.046911011 | CACHD1 |
| 15 | 75727796 | 75727953 | 1.649459393 | DNM1P35 | 16 | 72115381 | 72115567 | -1.176211538 | PMFBP1 |
| 15 | 75735200 | 75735400 | -2.557342155 | DNM1P35 | 16 | 72116532 | 72116730 | -2.602774456 | PMFBP1 |
| 1 | 46361785 | 46361904 | -1.429056558 | NSUN4 | 16 | 72795826 | 72795998 | -1.177303351 | ZFHX3 |
| 15 | 76259243 | 76259440 | -3.56540097 | TYRO3P | 16 | 72798153 | 72798414 | -1.792736459 | ZFHX3 |
| 15 | 76259243 | 76259440 | -3.56540097 | ETFA | 1 | 64691980 | 64692174 | 1.053874928 | CACHD1 |
| 15 | 76259595 | 76259783 | -2.188307812 | ETFA | 16 | 74452202 | 74452406 | -1.014358088 | GLG1 |
| 15 | 76259595 | 76259783 | -2.188307812 | TYRO3P | 16 | 74961805 | 74962013 | -1.34961343 | WDR59 |
| 15 | 77068832 | 77068986 | -4.957939689 | TSPAN3 | 16 | 75109470 | 75109700 | -2.119195208 | ZNRF1 |
| 15 | 77110899 | 77111095 | 2.856428488 | PEAK1 | 16 | 75109470 | 75109700 | -2.119195208 | RP11-252E2.1 |
| 15 | 77114235 | 77114400 | 1.104625002 | PEAK1 | 16 | 75169861 | 75170060 | -2.95908524 | ZFP1 |
| 15 | 77133068 | 77133247 | 1.061784289 | PEAK1 | 1 | 64777795 | 64777948 | -6.868306665 | RAVER2 |
| 15 | 77133300 | 77133435 | 1.165952982 | PEAK1 | 16 | 75264329 | 75264477 | 2.803541249 | BCAR1 |
| 1 | 46608180 | 46608375 | -3.285511655 | MOB3C | 16 | 75502439 | 75502603 | -5.22766361 | RP11-77K12.5 |
| 1 | 46608180 | 46608375 | -3.285511655 | MKNK1 | 1 | 65884855 | 65884962 | 1.182590646 | PDE4B |
| 15 | 77180174 | 77180411 | 1.178419147 | PEAK1 | 16 | 78119009 | 78119197 | -6.652460835 | WWOX |
| 15 | 77896488 | 77896669 | 6.909919897 | CSPG4P13 | 16 | 79600467 | 79600660 | -6.223799929 | MAF |
| 15 | 77897254 | 77897405 | 8.194008643 | CSPG4P13 | 16 | 80684675 | 80684842 | 1.075917801 | CDYL2 |
| 15 | 77898894 | 77899097 | 5.159892281 | CSPG4P13 | 16 | 83915457 | 83915647 | -1.531997266 | RP11-505K9.4 |
| 15 | 78012928 | 78013139 | -1.161783952 | TBC1D2B | 16 | 83915457 | 83915647 | -1.531997266 | MLYCD |
| 15 | 78766302 | 78766472 | 4.323264839 | ADAMTS7 | 1 | 65966831 | 65967017 | 1.956732839 | PDE4B |
| 15 | 78766537 | 78766735 | 3.441020323 | ADAMTS7 | 16 | 83932009 | 83932164 | -5.597993649 | RP11-505K9.4 |
| 15 | 79411517 | 79411733 | -1.613968421 | TMED3 | 1 | 65983116 | 65983416 | 2.391709419 | PDE4B |
| 15 | 79413015 | 79413217 | -1.425069998 | TMED3 | 16 | 84974731 | 84974946 | -1.124937787 | ZDHHC7 |
| 1 | 46613191 | 46613393 | 1.283670405 | MOB3C | 16 | 84976149 | 84976360 | -1.249939616 | ZDHHC7 |
| 1 | 46613191 | 46613393 | 1.283670405 | MKNK1 | 16 | 85170329 | 85170589 | -2.233811008 | GSE1 |
| 15 | 79899022 | 79899218 | 2.457325606 | ST20-MTHFS | 16 | 85672493 | 85672692 | -1.453050688 | GSE1 |
| 15 | 79899022 | 79899218 | 2.457325606 | ST20 | 16 | 85707669 | 85707861 | -4.43518236 | C16orf74 |
| 15 | 79923494 | 79923695 | 7.937730699 | ST20-MTHFS | 1 | 66322101 | 66322367 | 1.789469357 | PDE4B |
| 15 | 79923494 | 79923695 | 7.937730699 | ST20 | 16 | 86538287 | 86538405 | -1.190294065 | MTHFSD |
| 15 | 79923494 | 79923695 | 7.937730699 | ST20-AS1 | 16 | 87308875 | 87309073 | -1.834925699 | RP11-178L8.4 |
| 15 | 79970801 | 79971002 | 7.665333515 | BCL2A1 | 16 | 87308875 | 87309073 | -1.834925699 | C16orf95 |
| 15 | 81299395 | 81299582 | 6.532371382 | IL16 | 1 | 66372478 | 66372676 | 4.507754563 | PDE4B |
| 1 | 46718668 | 46718862 | -8.728758576 | EFCAB14 | 16 | 87700784 | 87700972 | -1.167610057 | KLHDC4 |
| 15 | 82786979 | 82787179 | 1.961923387 | FSD2 | 16 | 87831006 | 87831207 | -2.019494952 | SLC7A5 |
| 15 | 82833290 | 82833496 | 1.59711132 | WHAMM | 1 | 1917820 | 1918013 | -4.29248125 | TMEM52 |
| 15 | 82834245 | 82834469 | 1.095863949 | WHAMM | 16 | 87832433 | 87832634 | -1.979671559 | SLC7A5 |
| 15 | 82846591 | 82846795 | 2.210324128 | HOMER2 | 16 | 88048894 | 88049107 | 1.394639688 | BANP |
| 15 | 82849803 | 82850003 | 3.937053827 | HOMER2 | 16 | 88245771 | 88245965 | -6.950386243 | LA16c-444G7.1 |
| 15 | 82920024 | 82920209 | -6.189824559 | HOMER2 | 16 | 88432459 | 88432653 | -1.347733314 | ZNF469 |
| 1 | 47097572 | 47097673 | -1.050735396 | CYP4A22-AS1 | 16 | 88434581 | 88434771 | -1.064163185 | ZNF469 |
| 1 | 47097572 | 47097673 | -1.050735396 | CYP4Z1 | 16 | 88435531 | 88435722 | -1.556247307 | ZNF469 |
| 15 | 83012170 | 83012340 | -1.655885036 | RP11-382A20.5 | 16 | 88649270 | 88649475 | 9.262152245 | CYBA |
| 15 | 83017177 | 83017327 | 1.775914026 | BTBD1 | 16 | 88675999 | 88676152 | -1.449763604 | SNAI3-AS1 |
| 15 | 83017177 | 83017327 | 1.775914026 | RP11-382A20.7 | 16 | 88676614 | 88676774 | -1.074998975 | SNAI3-AS1 |
| 15 | 83017177 | 83017327 | 1.775914026 | RP11-382A20.5 | 16 | 88677929 | 88678134 | -1.453000006 | SNAI3 |
| 15 | 84605966 | 84606138 | -1.669171063 | RP11-182J1.18 | 16 | 88677929 | 88678134 | -1.453000006 | SNAI3-AS1 |

| | | | | |
|---|---|---|---|---|
| 15 | 84605966 | 84606138 | -1.669171063 | ZSCAN2 |
| 15 | 84641159 | 84641266 | -1.842288375 | WDR73 |
| 15 | 84641159 | 84641266 | -1.842288375 | SCAND2P |
| 1 | 841390 | 841584 | -2.789972213 | LINC01128 |
| 1 | 1598974 | 1599147 | -3.162911005 | FNDC10 |
| 1 | 47251512 | 47251739 | -3.206471609 | STIL |
| 15 | 84784569 | 84784750 | -1.057011102 | ZNF592 |
| 15 | 84839971 | 84840128 | 2.595136823 | ALPK3 |
| 15 | 84856488 | 84856774 | 1.356299664 | ALPK3 |
| 15 | 84868361 | 84868549 | 2.796008629 | ALPK3 |
| 1 | 47280322 | 47280531 | -3.232710485 | STIL |
| 15 | 85579358 | 85579618 | -1.739174122 | AKAP13 |
| 15 | 85579358 | 85579618 | -1.739174122 | RP11-815J21.2 |
| 15 | 85580833 | 85581055 | -1.608050282 | AKAP13 |
| 15 | 88459606 | 88459805 | -1.720279529 | MRPL46 |
| 15 | 88459606 | 88459805 | -1.720279529 | RP11-97O12.7 |
| 1 | 47280811 | 47281009 | -2.999429531 | STIL |
| 15 | 88531105 | 88531296 | -1.456852141 | DET1 |
| 15 | 88881396 | 88881588 | 5.704695468 | HAPLN3 |
| 15 | 88910228 | 88910404 | 6.548604572 | MFGE8 |
| 15 | 89623864 | 89624056 | -4.34877469 | TICRR |
| 15 | 89623864 | 89624056 | -4.34877469 | KIF7 |
| 15 | 89625069 | 89625256 | -4.233566644 | KIF7 |
| 15 | 89625069 | 89625256 | -4.233566644 | TICRR |
| 15 | 89626935 | 89627124 | -4.083833344 | TICRR |
| 15 | 89626935 | 89627124 | -4.083833344 | KIF7 |
| 15 | 89629615 | 89629809 | -5.552694744 | KIF7 |
| 15 | 89629615 | 89629809 | -5.552694744 | TICRR |
| 15 | 89630776 | 89630963 | -4.693691915 | KIF7 |
| 15 | 89630776 | 89630963 | -4.693691915 | TICRR |
| 15 | 89631808 | 89632001 | -5.212005011 | KIF7 |
| 15 | 89639827 | 89640017 | -4.332308476 | KIF7 |
| 15 | 89683268 | 89683462 | -1.272088443 | PEX11A |
| 15 | 90067181 | 90067379 | -1.925611274 | ZNF710 |
| 1 | 47432887 | 47433096 | -1.720254542 | FOXD2-AS1 |
| 15 | 90749677 | 90749918 | -3.630704187 | BLM |
| 15 | 90760898 | 90761085 | -3.770248568 | BLM |
| 1 | 47433544 | 47433745 | -1.180447675 | FOXD2-AS1 |
| 15 | 90881695 | 90881930 | 1.766875792 | FURIN |
| 15 | 90961116 | 90961317 | -4.63383253 | RCCD1 |
| 15 | 92162956 | 92163169 | -1.424627333 | SLCO3A1 |
| 15 | 92162956 | 92163169 | -1.424627333 | RP11-152L20.3 |
| 15 | 92162956 | 92163169 | -1.424627333 | RP11-24J19.1 |
| 15 | 92789105 | 92789377 | -2.58928196 | FAM174B |
| 15 | 96331733 | 96331915 | 1.422345881 | NR2F2 |
| 15 | 97630437 | 97630616 | -1.507673982 | LINC00923 |
| 15 | 97674535 | 97674728 | -1.192007737 | LINC00923 |
| 15 | 97874219 | 97874428 | -1.390522357 | LINC00923 |
| 15 | 97971355 | 97971565 | 1.593385058 | ARRDC4 |
| 15 | 97973299 | 97973485 | 2.013031148 | ARRDC4 |
| 15 | 99113972 | 99114154 | -7.315815903 | SYNM |
| 1 | 50417653 | 50417852 | -1.935493511 | DMRTA2 |
| 15 | 99130878 | 99131057 | -1.007249284 | RP11-6O2.4 |
| 15 | 99130878 | 99131057 | -1.007249284 | SYNM |

| | | | | |
|---|---|---|---|---|
| 16 | 88686759 | 88686937 | -1.996670332 | AC138028.1 |
| 16 | 88686759 | 88686937 | -1.996670332 | SNAI3-AS1 |
| 16 | 88829087 | 88829294 | 6.358409731 | GALNS |
| 16 | 88876376 | 88876598 | -3.132285158 | CBFA2T3 |
| 16 | 88876833 | 88876965 | -4.599836172 | CBFA2T3 |
| 16 | 89151166 | 89151365 | -1.053347226 | ACSF3 |
| 16 | 89152965 | 89153174 | -1.402460138 | ACSF3 |
| 16 | 89160496 | 89160694 | -1.244908055 | LINC00304 |
| 16 | 89200757 | 89200964 | -1.463176176 | SLC22A31 |
| 16 | 89215640 | 89215763 | -4.441321525 | RP11-46C24.7 |
| 16 | 89216825 | 89216955 | -1.545131238 | RP11-46C24.7 |
| 16 | 89235469 | 89235656 | -7.212607952 | ZNF778 |
| 16 | 89267847 | 89268053 | -1.483386824 | ANKRD11 |
| 16 | 89281835 | 89282035 | -1.087897926 | ANKRD11 |
| 16 | 89282931 | 89283121 | -1.725465313 | ANKRD11 |
| 16 | 89283717 | 89283918 | -1.645501022 | ANKRD11 |
| 16 | 89284706 | 89284906 | -1.63286723 | ANKRD11 |
| 16 | 89498822 | 89499006 | -4.846743479 | RP11-104N10.1 |
| 16 | 89498822 | 89499006 | -4.846743479 | SPG7 |
| 16 | 89686286 | 89686529 | -1.129259462 | RP11-368I7.4 |
| 16 | 89686286 | 89686529 | -1.129259462 | CDK10 |
| 16 | 89707418 | 89707615 | 4.498696822 | VPS9D1 |
| 16 | 89723370 | 89723580 | 1.013927127 | ZNF276 |
| 1 | 67033684 | 67033829 | -5.777294426 | SLC35D1 |
| 16 | 89730467 | 89730677 | 8.374397797 | ZNF276 |
| 16 | 89871239 | 89871429 | 1.066633265 | SPIRE2 |
| 16 | 89966134 | 89966369 | -1.27541709 | DEF8 |
| 1 | 67395727 | 67395927 | 2.846130411 | IL12RB2 |
| 16 | 89971299 | 89971529 | -1.360712372 | RP11-566K11.5 |
| 16 | 89971299 | 89971529 | -1.360712372 | CENPBD1 |
| 16 | 89971872 | 89972046 | -1.991233185 | RP11-566K11.5 |
| 16 | 89971872 | 89972046 | -1.991233185 | CENPBD1 |
| 16 | 90167310 | 90167533 | -1.124936579 | RP11-356C4.5 |
| 16 | 90167310 | 90167533 | -1.124936579 | FAM157C |
| 17 | 231094 | 231255 | -5.518086806 | RPH3AL |
| 17 | 270773 | 270908 | -2.988639962 | RPH3AL |
| 17 | 368173 | 368370 | -1.273580234 | RPH3AL |
| 17 | 634680 | 634826 | -5.281121212 | VPS53 |
| 17 | 753523 | 753667 | -5.02219706 | GEMIN4 |
| 17 | 753523 | 753667 | -5.02219706 | DBIL5P |
| 17 | 800795 | 800985 | -1.626339989 | NXN |
| 17 | 815353 | 815457 | -8.581451332 | NXN |
| 17 | 1001228 | 1001424 | -1.47588692 | TIMM22 |
| 17 | 1042480 | 1042678 | 6.206152602 | ABR |
| 17 | 1148409 | 1148598 | -7.364491685 | ABR |
| 17 | 1423051 | 1423231 | 2.763264003 | CRK |
| 17 | 1423256 | 1423446 | 7.769821753 | CRK |
| 17 | 1494976 | 1495155 | 1.103112098 | INPP5K |
| 17 | 1517682 | 1517934 | -1.176330611 | PITPNA |
| 17 | 1517682 | 1517934 | -1.176330611 | PITPNA-AS1 |
| 17 | 1533850 | 1533994 | 1.211042766 | PITPNA |
| 17 | 1575155 | 1575355 | 4.428866076 | SLC43A2 |
| 17 | 1628171 | 1628347 | 9.516726961 | SLC43A2 |
| 17 | 1634654 | 1634939 | 4.520707016 | SCARF1 |

| | | | | | | | | | |
|---|---|---|---|---|---|---|---|---|---|
| 15 | 99131583 | 99131752 | -1.011919543 | SYNM | 17 | 1712640 | 1712840 | 1.576901598 | MIR22HG |
| 15 | 99131583 | 99131752 | -1.011919543 | RP11-6O2.4 | 17 | 1725766 | 1725970 | -1.01501326 | RP11-961A15.1 |
| 15 | 99132580 | 99132773 | -2.106518176 | SYNM | 17 | 1725766 | 1725970 | -1.01501326 | WDR81 |
| 15 | 99137271 | 99137415 | -1.178733181 | TTC23 | 17 | 1777273 | 1777466 | 1.483558719 | SERPINF1 |
| 15 | 99249307 | 99249525 | -1.85687812 | TTC23 | 17 | 1800170 | 1800425 | -2.213752482 | SMYD4 |
| 15 | 99366224 | 99366502 | 8.582433827 | LRRC28 | 17 | 2305297 | 2305471 | 3.0605077 | SRR |
| 15 | 99368623 | 99368809 | 8.005700995 | LRRC28 | 17 | 2386596 | 2386796 | -1.045423418 | RP1-59D14.1 |
| 15 | 99376819 | 99377017 | 11.64034943 | LRRC28 | 17 | 2386596 | 2386796 | -1.045423418 | MNT |
| 15 | 99378576 | 99378773 | 9.565164618 | LRRC28 | 17 | 2395045 | 2395239 | 4.594912279 | MNT |
| 15 | 99380311 | 99380511 | 12.74170223 | LRRC28 | 17 | 2712646 | 2712844 | -2.053750297 | RP11-74E22.3 |
| 15 | 99380987 | 99381185 | 9.135033575 | LRRC28 | 17 | 2712646 | 2712844 | -2.053750297 | CLUH |
| 15 | 99381684 | 99381888 | 11.2607638 | LRRC28 | 17 | 3592184 | 3592382 | -2.637160416 | TRPV1 |
| 15 | 99386099 | 99386389 | 10.73039441 | LRRC28 | 17 | 3592184 | 3592382 | -2.637160416 | RP11-235E17.2 |
| 15 | 99386972 | 99387203 | 10.89609568 | LRRC28 | 17 | 3610322 | 3610523 | -1.521101248 | RP11-235E17.2 |
| 15 | 99678584 | 99678782 | 6.301978722 | MEF2A | 17 | 3610322 | 3610523 | -1.521101248 | SHPK |
| 15 | 99728019 | 99728152 | -2.39943139 | LYSMD4 | 17 | 3682291 | 3682479 | -1.037700404 | P2RX5-TAX1BP3 |
| 15 | 99792241 | 99792353 | -2.697588538 | CTD-2054N24.1 | 17 | 3682291 | 3682479 | -1.037700404 | P2RX5 |
| 15 | 99792241 | 99792353 | -2.697588538 | DNM1P46 | 17 | 3725792 | 3725976 | -3.77286528 | GSG2 |
| 15 | 99799858 | 99800140 | -3.651562006 | DNM1P46 | 17 | 3725792 | 3725976 | -3.77286528 | ITGAE |
| 15 | 99801053 | 99801235 | -2.077288519 | DNM1P46 | 17 | 3726069 | 3726277 | -2.789452175 | ITGAE |
| 15 | 99801907 | 99802012 | -3.798094878 | DNM1P46 | 17 | 3726069 | 3726277 | -2.789452175 | GSG2 |
| 15 | 100349260 | 100349382 | -6.886569603 | SPATA41 | 17 | 3861162 | 3861366 | -2.682051172 | CAMKK1 |
| 15 | 100623568 | 100623738 | -2.311783098 | ASB7 | 17 | 3861832 | 3862008 | -3.130548432 | CAMKK1 |
| 1 | 50905906 | 50906107 | -1.774660993 | FAF1 | 17 | 3913595 | 3913827 | -1.227914191 | P2RX1 |
| 15 | 100918404 | 100918602 | -3.639921347 | RP11-66B24.4 | 17 | 3915612 | 3915822 | -2.499872784 | P2RX1 |
| 15 | 101065750 | 101065957 | -1.176395804 | RP11-505E24.2 | 17 | 4510778 | 4510981 | -2.303665157 | SPNS2 |
| 15 | 101065750 | 101065957 | -1.176395804 | LRRK1 | 17 | 4537780 | 4538047 | -2.069002175 | SPNS2 |
| 15 | 101068789 | 101068906 | -3.119112793 | RP11-505E24.2 | 17 | 4731488 | 4731670 | 1.740563345 | MED11 |
| 15 | 101068789 | 101068906 | -3.119112793 | LRRK1 | 17 | 5092470 | 5092687 | -1.133806212 | ZFP3 |
| 15 | 101170593 | 101170770 | -3.657622506 | RP11-424I19.2 | 17 | 5106283 | 5106482 | -3.297600192 | ZNF232 |
| 1 | 50908674 | 50908990 | -1.019891698 | FAF1 | 1 | 2190579 | 2190686 | -3.947408882 | FAAP20 |
| 15 | 101301958 | 101302145 | 3.267225603 | RP11-299G20.2 | 17 | 5109367 | 5109562 | -2.275575505 | ZNF232 |
| 15 | 101301958 | 101302145 | 3.267225603 | PCSK6 | 17 | 5180914 | 5181115 | -1.59028356 | ZNF594 |
| 15 | 101302867 | 101303177 | 3.211256695 | RP11-299G20.2 | 17 | 5181864 | 5182045 | -1.389889607 | ZNF594 |
| 15 | 101302867 | 101303177 | 3.211256695 | PCSK6 | 17 | 5184033 | 5184233 | -8.317596895 | ZNF594 |
| 15 | 101304756 | 101305036 | 3.304526713 | RP11-299G20.2 | 17 | 5186758 | 5186960 | -1.202468149 | ZNF594 |
| 15 | 101304756 | 101305036 | 3.304526713 | PCSK6 | 17 | 5210684 | 5210866 | 2.59700531 | RP11-333E1.1 |
| 15 | 101309832 | 101310032 | 3.126212986 | PCSK6 | 17 | 5210684 | 5210866 | 2.59700531 | SCIMP |
| 15 | 101328856 | 101329086 | 3.308537343 | PCSK6 | 17 | 5486417 | 5486575 | -4.208926257 | DERL2 |
| 15 | 101329584 | 101329799 | 3.931554452 | PCSK6 | 17 | 5486417 | 5486575 | -4.208926257 | MIS12 |
| 15 | 101490548 | 101490696 | 4.45394606 | PCSK6 | 17 | 6652305 | 6652510 | -4.308055035 | C17orf100 |
| 15 | 101491390 | 101491596 | 3.137869456 | PCSK6 | 17 | 6775126 | 6775329 | 1.009737836 | XAF1 |
| 15 | 101492425 | 101492625 | 2.652333899 | PCSK6 | 17 | 6775126 | 6775329 | 1.009737836 | RP11-609D21.3 |
| 15 | 101764641 | 101764845 | -3.473492429 | DNM1P47 | 17 | 6782462 | 6782653 | 1.405681443 | FBXO39 |
| 1 | 50969015 | 50969303 | -10.68180111 | CDKN2C | 17 | 6785308 | 6785477 | 1.366613252 | FBXO39 |
| 16 | 29531 | 29718 | -1.943725798 | IL9RP3 | 17 | 6791233 | 6791432 | 1.471367615 | TEKT1 |
| 16 | 29531 | 29718 | -1.943725798 | Z84812.4 | 17 | 6791233 | 6791432 | 1.471367615 | FBXO39 |
| 16 | 57142 | 57371 | -2.893796443 | SNRNP25 | 17 | 7038566 | 7038760 | -1.416296107 | SLC16A13 |
| 16 | 112672 | 112822 | 6.175745705 | NPRL3 | 17 | 7039772 | 7039984 | -2.148857374 | SLC16A13 |
| 16 | 486621 | 486807 | -7.004433133 | RAB11FIP3 | 1 | 77565572 | 77565761 | -1.216547748 | ZZZ3 |
| 16 | 516978 | 517174 | -3.738535488 | RAB11FIP3 | 17 | 7383622 | 7383790 | -6.558171981 | TNK1 |
| 16 | 520631 | 520741 | 1.808074275 | RAB11FIP3 | 17 | 7415682 | 7415928 | -3.21414836 | NLGN2 |
| 16 | 520880 | 521066 | -2.671967901 | RAB11FIP3 | 17 | 7456828 | 7457005 | 1.002797731 | CHRNB1 |
| 16 | 525442 | 525688 | -2.209645332 | LINC00235 | 17 | 7463082 | 7463281 | -2.437072378 | ZBTB4 |

| | | | | |
|---|---|---|---|---|
| 16 | 526694 | 526889 | -2.640489233 | LINC00235 |
| 16 | 547687 | 547913 | -1.191742199 | LA16c-366D1.3 |
| 16 | 547687 | 547913 | -1.191742199 | CAPN15 |
| 16 | 589504 | 589677 | -2.621586992 | RAB40C |
| 16 | 614654 | 614815 | -1.819550533 | RAB40C |
| 1 | 50973928 | 50974122 | -2.07441702 | CDKN2C |
| 16 | 682518 | 682718 | -1.738763609 | JMJD8 |
| 16 | 682518 | 682718 | -1.738763609 | STUB1 |
| 16 | 708799 | 708987 | 6.546130812 | LA16c-380A1.2 |
| 16 | 720545 | 720758 | -2.200560709 | FAM173A |
| 16 | 730345 | 730460 | -1.142118213 | NARFL |
| 16 | 730345 | 730460 | -1.142118213 | HAGHL |
| 16 | 735232 | 735389 | 1.195711103 | NARFL |
| 16 | 735232 | 735389 | 1.195711103 | HAGHL |
| 16 | 785876 | 786065 | -2.43657697 | RPUSD1 |
| 16 | 887168 | 887316 | -1.111196211 | LMF1 |
| 16 | 888083 | 888416 | -1.315547811 | LMF1 |
| 16 | 966521 | 966723 | -2.82898979 | LMF1 |
| 16 | 967400 | 967616 | -1.727865295 | LMF1 |
| 16 | 968725 | 968902 | -8.048120795 | LMF1 |
| 16 | 968725 | 968902 | -8.048120795 | RP11-161M6.6 |
| 16 | 969788 | 969987 | -2.330409589 | LMF1 |
| 16 | 1257864 | 1257984 | 5.821251514 | RP11-616M22.5 |
| 16 | 1257864 | 1257984 | 5.821251514 | TPSD1 |
| 16 | 1349420 | 1349628 | -1.147352153 | BAIAP3 |
| 16 | 1349420 | 1349628 | -1.147352153 | TSR3 |
| 16 | 1420091 | 1420367 | -1.295727721 | C16orf91 |
| 16 | 1434299 | 1434487 | -7.257849919 | CCDC154 |
| 16 | 1434861 | 1434963 | -5.506661337 | CCDC154 |
| 16 | 1510053 | 1510256 | -1.857061867 | TELO2 |
| 16 | 1511133 | 1511361 | -2.048382633 | IFT140 |
| 16 | 1519866 | 1519980 | -5.341058382 | IFT140 |
| 16 | 1536787 | 1537033 | -4.28739136 | TMEM204 |
| 16 | 1536787 | 1537033 | -4.28739136 | IFT140 |
| 16 | 1699255 | 1699450 | -2.274540188 | HN1L |
| 16 | 1732537 | 1732744 | -1.530766634 | MAPK8IP3 |
| 16 | 1772231 | 1772450 | -1.836491821 | MRPS34 |
| 16 | 1776811 | 1777122 | -1.634984154 | EME2 |
| 16 | 1776811 | 1777122 | -1.634984154 | SPSB3 |
| 16 | 1938740 | 1938940 | -2.186274326 | MSRB1 |
| 16 | 2089673 | 2089801 | -1.090293556 | PKD1 |
| 16 | 2111369 | 2111568 | -6.651747688 | PKD1 |
| 16 | 2177591 | 2177779 | -1.519881713 | TRAF7 |
| 16 | 2177591 | 2177779 | -1.519881713 | CASKIN1 |
| 16 | 2213295 | 2213515 | -2.554038844 | PGP |
| 16 | 2231151 | 2231324 | 5.745926548 | E4F1 |
| 16 | 2235443 | 2235644 | -1.031709172 | E4F1 |
| 16 | 2456764 | 2456968 | -3.647542265 | RP11-715J22.4 |
| 16 | 2456764 | 2456968 | -3.647542265 | CCNF |
| 16 | 2457514 | 2457704 | -9.653331255 | CCNF |
| 16 | 2457514 | 2457704 | -9.653331255 | RP11-715J22.4 |
| 16 | 2496114 | 2496313 | -2.414393581 | RP11-20I23.1 |
| 16 | 2496114 | 2496313 | -2.414393581 | TBC1D24 |
| 16 | 2502552 | 2502778 | -2.599231975 | TBC1D24 |

| | | | | |
|---|---|---|---|---|
| 17 | 7466522 | 7466715 | -6.161668145 | ZBTB4 |
| 17 | 7849086 | 7849294 | 3.381450125 | KDM6B |
| 17 | 7915206 | 7915417 | -2.404022818 | AC025335.1 |
| 1 | 77889020 | 77889213 | -6.765203169 | NEXN-AS1 |
| 1 | 77889020 | 77889213 | -6.765203169 | NEXN |
| 17 | 8140923 | 8141131 | -1.718611672 | PER1 |
| 17 | 8227768 | 8227979 | -1.148243047 | CTC1 |
| 17 | 8227995 | 8228119 | -1.924108789 | CTC1 |
| 17 | 8369148 | 8369347 | -2.81754299 | RP11-849F2.5 |
| 17 | 8369148 | 8369347 | -2.81754299 | KRBA2 |
| 17 | 8369148 | 8369347 | -2.81754299 | RP11-849F2.7 |
| 17 | 8369795 | 8369996 | -2.105652858 | KRBA2 |
| 17 | 8369795 | 8369996 | -2.105652858 | RP11-849F2.7 |
| 17 | 8369795 | 8369996 | -2.105652858 | RP11-849F2.5 |
| 17 | 8879205 | 8879405 | 2.172185771 | PIK3R5 |
| 17 | 8880487 | 8880690 | 1.604417797 | PIK3R5 |
| 17 | 8888360 | 8888565 | 2.236960337 | PIK3R5 |
| 17 | 9239671 | 9239895 | 3.58353535 | NTN1 |
| 17 | 9243384 | 9243560 | 6.600993019 | NTN1 |
| 17 | 11978076 | 11978283 | -1.427756237 | ZNF18 |
| 17 | 13496230 | 13496413 | 1.590425421 | HS3ST3A1 |
| 17 | 13496572 | 13496785 | 2.138424442 | HS3ST3A1 |
| 17 | 13777032 | 13777186 | -2.320352526 | COX10-AS1 |
| 17 | 14302329 | 14302522 | 1.398997514 | HS3ST3B1 |
| 17 | 14303791 | 14304002 | 2.106521777 | RP11-214O1.2 |
| 17 | 14303791 | 14304002 | 2.106521777 | HS3ST3B1 |
| 17 | 14345492 | 14345697 | 1.49238616 | HS3ST3B1 |
| 17 | 15946098 | 15946298 | -4.150366936 | ADORA2B |
| 17 | 15952463 | 15952586 | -1.649283967 | ADORA2B |
| 17 | 15974981 | 15975138 | -3.222537414 | ADORA2B |
| 17 | 15976922 | 15977108 | -1.547988927 | ZSWIM7 |
| 17 | 16032135 | 16032335 | -1.287135336 | TTC19 |
| 17 | 16032135 | 16032335 | -1.287135336 | NCOR1 |
| 17 | 16216849 | 16217049 | -4.058171981 | NCOR1 |
| 17 | 16325965 | 16326086 | -1.459203308 | PIGL |
| 17 | 16349030 | 16349230 | -7.112603718 | PIGL |
| 17 | 16349030 | 16349230 | -7.112603718 | CENPV |
| 17 | 16443629 | 16443729 | -1.33492109 | LRRC75A |
| 17 | 16443629 | 16443729 | -1.33492109 | LRRC75A-AS1 |
| 17 | 16461543 | 16461737 | -1.163733139 | LRRC75A-AS1 |
| 17 | 16461543 | 16461737 | -1.163733139 | LRRC75A |
| 17 | 16471688 | 16471882 | -2.183266213 | LRRC75A |
| 17 | 16471688 | 16471882 | -2.183266213 | LRRC75A-AS1 |
| 17 | 16472159 | 16472311 | -1.737320837 | LRRC75A-AS1 |
| 17 | 16472159 | 16472311 | -1.737320837 | LRRC75A |
| 17 | 17167435 | 17167631 | -2.304754442 | MPRIP |
| 17 | 17167640 | 17167808 | -1.115946712 | MPRIP |
| 17 | 17187473 | 17187671 | -4.316770097 | MPRIP |
| 17 | 17188180 | 17188375 | -2.016390785 | MPRIP |
| 17 | 17202571 | 17202770 | -1.191584083 | RP11-45M22.4 |
| 17 | 17202571 | 17202770 | -1.191584083 | MPRIP |
| 17 | 17202571 | 17202770 | -1.191584083 | PLD6 |
| 1 | 81802439 | 81802632 | 2.467473278 | ADGRL2 |
| 17 | 17202861 | 17203043 | -3.153341569 | MPRIP |

| | | | | |
|---|---|---|---|---|
| 16 | 2502552 | 2502778 | -2.599231975 | RP11-20I23.1 |
| 16 | 2563681 | 2563851 | -6.465368669 | RP11-20I23.8 |
| 16 | 2563681 | 2563851 | -6.465368669 | PDPK1 |
| 1 | 51785646 | 51785792 | -5.689147428 | OSBPL9 |
| 16 | 2566803 | 2567040 | -2.012640983 | PDPK1 |
| 16 | 2571091 | 2571257 | -2.531084411 | RP11-20I23.6 |
| 16 | 2571091 | 2571257 | -2.531084411 | PDPK1 |
| 16 | 2628885 | 2629071 | -2.92292422 | AC141586.5 |
| 16 | 2628885 | 2629071 | -2.92292422 | PDPK2P |
| 16 | 2637385 | 2637508 | -2.681583601 | PDPK2P |
| 16 | 2637918 | 2638120 | -3.002246828 | PDPK2P |
| 16 | 2639206 | 2639333 | -1.619436178 | PDPK2P |
| 16 | 2640210 | 2640424 | -3.126046014 | PDPK2P |
| 16 | 2641614 | 2641818 | -1.393865452 | PDPK2P |
| 1 | 51840424 | 51840622 | -1.889912758 | NRDC |
| 16 | 2642800 | 2642999 | -1.944340845 | PDPK2P |
| 16 | 2708351 | 2708534 | 1.186644026 | KCTD5 |
| 16 | 2762476 | 2762674 | -11.60871508 | SRRM2 |
| 16 | 2767234 | 2767434 | -1.310050675 | SRRM2 |
| 16 | 2968167 | 2968285 | -1.221984304 | PKMYT1 |
| 16 | 2968167 | 2968285 | -1.221984304 | KREMEN2 |
| 16 | 2971713 | 2971828 | -2.813499697 | PKMYT1 |
| 16 | 2971713 | 2971828 | -2.813499697 | PAQR4 |
| 1 | 51878362 | 51878571 | -1.557024472 | NRDC |
| 16 | 3053394 | 3053525 | -1.315213266 | MMP25 |
| 16 | 3053394 | 3053525 | -1.315213266 | MMP25-AS1 |
| 16 | 3072904 | 3073099 | -4.975642357 | IL32 |
| 16 | 3084657 | 3084812 | -1.621041527 | RP11-473M20.9 |
| 16 | 3084825 | 3085008 | -1.295438324 | RP11-473M20.9 |
| 16 | 3124495 | 3124692 | -1.260965896 | ZNF213-AS1 |
| 1 | 52032411 | 52032612 | -1.430736948 | TXNDC12 |
| 1 | 52032411 | 52032612 | -1.430736948 | KTI12 |
| 16 | 3224159 | 3224360 | -1.369482286 | ZNF200 |
| 16 | 3224159 | 3224360 | -1.369482286 | AJ003147.9 |
| 16 | 3242912 | 3243107 | -2.129916361 | MEFV |
| 16 | 3243294 | 3243492 | -2.031773868 | MEFV |
| 16 | 3282592 | 3282785 | 3.202314744 | ZNF263 |
| 16 | 3282989 | 3283150 | 2.754750481 | ZNF263 |
| 16 | 3308727 | 3308831 | -1.322800697 | LA16c-360H6.2 |
| 16 | 3308727 | 3308831 | -1.322800697 | ZNF75A |
| 16 | 3309335 | 3309561 | -2.227365141 | ZNF75A |
| 16 | 3310517 | 3310682 | -4.438551514 | ZNF75A |
| 16 | 3318284 | 3318482 | -1.030928816 | ZNF75A |
| 16 | 3385787 | 3385970 | -1.382286505 | ZSCAN32 |
| 16 | 3385787 | 3385970 | -1.382286505 | LA16c-306E5.2 |
| 16 | 3412090 | 3412271 | 1.540186708 | LA16c-306E5.2 |
| 16 | 3412886 | 3413112 | -1.38948656 | LA16c-306E5.2 |
| 16 | 3486503 | 3486745 | -1.775975351 | LA16c-306E5.3 |
| 16 | 3486503 | 3486745 | -1.775975351 | NAA60 |
| 16 | 3581382 | 3581589 | -1.82342239 | RP11-461A8.1 |
| 16 | 3581382 | 3581589 | -1.82342239 | SLX4 |
| 16 | 3582347 | 3582545 | -1.742874988 | SLX4 |
| 16 | 3582347 | 3582545 | -1.742874988 | RP11-461A8.1 |
| 16 | 3583230 | 3583419 | -2.002248219 | SLX4 |
| 17 | 17202861 | 17203043 | -3.153341569 | PLD6 |
| 17 | 17202861 | 17203043 | -3.153341569 | RP11-45M22.4 |
| 17 | 17326361 | 17326512 | -2.103053669 | NT5M |
| 17 | 17494755 | 17494947 | -1.576475961 | RASD1 |
| 17 | 17793315 | 17793518 | -1.596939536 | RAI1 |
| 17 | 17794942 | 17795210 | -1.25505371 | RAI1 |
| 17 | 17797296 | 17797496 | -1.516287269 | RAI1 |
| 17 | 17797927 | 17798113 | -1.254858608 | RAI1 |
| 17 | 17955525 | 17955692 | -4.577409055 | TOM1L2 |
| 17 | 18065184 | 18065411 | -2.079192486 | GID4 |
| 17 | 18183477 | 18183672 | 5.464629204 | RP11-258F1.1 |
| 17 | 18183477 | 18183672 | 5.464629204 | ALKBH5 |
| 17 | 18316566 | 18316760 | -1.072319617 | SMCR8 |
| 17 | 18515536 | 18515744 | -1.627354917 | USP32P2 |
| 17 | 18515536 | 18515744 | -1.627354917 | CTD-2303H24.2 |
| 17 | 18534511 | 18534719 | -1.491454546 | CTD-2303H24.2 |
| 17 | 18535288 | 18535484 | -2.225401276 | CTD-2303H24.2 |
| 17 | 18536819 | 18537017 | -2.229947698 | CTD-2303H24.2 |
| 17 | 18538286 | 18538450 | -3.033487781 | CCDC144B |
| 17 | 18538286 | 18538450 | -3.033487781 | CTD-2303H24.2 |
| 17 | 18538590 | 18538786 | -1.059160778 | CCDC144B |
| 17 | 18538590 | 18538786 | -1.059160778 | CTD-2303H24.2 |
| 17 | 18580178 | 18580370 | -1.933366735 | CCDC144B |
| 17 | 18661871 | 18662028 | -4.63339327 | TBC1D28 |
| 17 | 18661871 | 18662028 | -4.63339327 | ZNF286B |
| 17 | 18663084 | 18663199 | -2.465526323 | ZNF286B |
| 17 | 18670673 | 18670860 | -1.044556598 | ZNF286B |
| 17 | 19283281 | 19283529 | -1.387055761 | EPN2 |
| 1 | 84926483 | 84926716 | 2.158030949 | MCOLN2 |
| 17 | 20828900 | 20829073 | -1.498731457 | ABHD17AP6 |
| 1 | 85100668 | 85100866 | 3.233190247 | WDR63 |
| 17 | 20922188 | 20922387 | -1.456388691 | CCDC144NL-AS1 |
| 17 | 20946728 | 20946934 | -2.043686078 | CCDC144NL-AS1 |
| 17 | 20987013 | 20987213 | -1.15878824 | CCDC144NL-AS1 |
| 17 | 21002570 | 21002772 | -1.392628334 | USP22 |
| 1 | 85158898 | 85159171 | -4.271398694 | SYDE2 |
| 17 | 28326607 | 28326808 | -1.581216206 | TMEM97 |
| 17 | 28345555 | 28345744 | 3.503909752 | TNFAIP1 |
| 17 | 28592803 | 28593000 | -1.866351625 | RP11-192H23.4 |
| 17 | 28592803 | 28593000 | -1.866351625 | SPAG5 |
| 1 | 85182194 | 85182393 | -1.992679399 | SYDE2 |
| 17 | 28658713 | 28658825 | -1.436359668 | SDF2 |
| 17 | 28724448 | 28724585 | 1.307911732 | TLCD1 |
| 17 | 28749229 | 28749585 | -1.038936944 | TRAF4 |
| 17 | 28758370 | 28758546 | 1.40540901 | FAM222B |
| 17 | 28759485 | 28759675 | 1.172515559 | FAM222B |
| 17 | 28906047 | 28906247 | -1.774070938 | PHF12 |
| 1 | 85267727 | 85267889 | -1.430546521 | BCL10 |
| 17 | 28912941 | 28913141 | -1.98946682 | PHF12 |
| 17 | 28921783 | 28921978 | -3.258120997 | PHF12 |
| 17 | 28924110 | 28924310 | -2.256026703 | PHF12 |
| 17 | 29077718 | 29077901 | 5.520144861 | TIAF1 |
| 17 | 29077718 | 29077901 | 5.520144861 | MYO18A |
| 1 | 86350437 | 86350634 | 1.061329624 | ODF2L |

| | | | | |
|---|---|---|---|---|
| 16 | 3583230 | 3583419 | -2.002248219 | RP11-461A8.1 |
| 16 | 3584837 | 3585041 | -2.15676748 | SLX4 |
| 16 | 3589112 | 3589372 | -2.132503812 | SLX4 |
| 16 | 3591079 | 3591199 | -1.738207102 | SLX4 |
| 16 | 3608518 | 3608856 | -1.818951864 | SLX4 |
| 16 | 3652957 | 3653169 | -1.26576166 | DNASE1 |
| 16 | 3652957 | 3653169 | -1.26576166 | TRAP1 |
| 16 | 3661542 | 3661667 | -7.228626306 | TRAP1 |
| 16 | 3661542 | 3661667 | -7.228626306 | DNASE1 |
| 16 | 3664899 | 3665099 | 1.398172071 | DNASE1 |
| 16 | 3664899 | 3665099 | 1.398172071 | TRAP1 |
| 16 | 3965882 | 3966092 | -1.343656987 | ADCY9 |
| 16 | 3993653 | 3993755 | -2.642792027 | ADCY9 |
| 16 | 4113853 | 4114042 | -1.798491843 | ADCY9 |
| 16 | 4114345 | 4114533 | -2.100644543 | ADCY9 |
| 16 | 4337374 | 4337544 | 5.44208526 | RP11-295D4.1 |
| 16 | 4337374 | 4337544 | 5.44208526 | PAM16 |
| 16 | 4337374 | 4337544 | 5.44208526 | GLIS2 |
| 16 | 4383170 | 4383361 | -7.576459288 | CORO7-PAM16 |
| 16 | 4383170 | 4383361 | -7.576459288 | CORO7 |
| 16 | 4383170 | 4383361 | -7.576459288 | VASN |
| 16 | 4461786 | 4461989 | -1.79954061 | NMRAL1 |
| 16 | 4509284 | 4509475 | -2.075470949 | HMOX2 |
| 1 | 52389253 | 52389451 | -4.346657783 | ORC1 |
| 16 | 4699381 | 4699577 | -1.068013873 | ANKS3 |
| 16 | 4752166 | 4752390 | -1.280659997 | ZNF500 |
| 16 | 4752166 | 4752390 | -1.280659997 | RP11-127I20.7 |
| 16 | 4860804 | 4861004 | -1.020761121 | UBN1 |
| 16 | 4874391 | 4874594 | -1.191483062 | UBN1 |
| 16 | 4882664 | 4882784 | -2.309417515 | PPL |
| 1 | 52393498 | 52393698 | -3.858147579 | ORC1 |
| 16 | 5025050 | 5025249 | -1.100845526 | RP11-165E7.1 |
| 16 | 5025050 | 5025249 | -1.100845526 | NAGPA |
| 16 | 5042246 | 5042428 | -1.886998163 | RP11-165E7.1 |
| 16 | 5042246 | 5042428 | -1.886998163 | ALG1 |
| 16 | 5084695 | 5084882 | -3.233779836 | ALG1 |
| 16 | 5084695 | 5084882 | -3.233779836 | EEF2KMT |
| 16 | 5085111 | 5085302 | -2.07691841 | ALG1 |
| 16 | 5085111 | 5085302 | -2.07691841 | EEF2KMT |
| 16 | 8714786 | 8714963 | 1.444484344 | ABAT |
| 16 | 8783234 | 8783410 | 2.776441265 | ABAT |
| 16 | 8783234 | 8783410 | 2.776441265 | TMEM186 |
| 16 | 8795957 | 8796160 | -1.117463604 | PMM2 |
| 16 | 8795957 | 8796160 | -1.117463604 | TMEM186 |
| 16 | 8815737 | 8815842 | -2.65897229 | PMM2 |
| 16 | 8855031 | 8855237 | -1.163649164 | RP11-77H9.2 |
| 16 | 8855031 | 8855237 | -1.163649164 | CARHSP1 |
| 1 | 52417738 | 52417941 | -1.11868518 | ZCCHC11 |
| 1 | 52417738 | 52417941 | -1.11868518 | PRPF38A |
| 16 | 10415395 | 10415595 | -1.143790511 | ATF7IP2 |
| 16 | 10763579 | 10763800 | -2.011080349 | NUBP1 |
| 16 | 10763579 | 10763800 | -2.011080349 | TVP23A |
| 16 | 10941683 | 10941869 | -1.824485381 | DEXI |
| 16 | 10941683 | 10941869 | -1.824485381 | RP11-876N24.4 |
| 17 | 29560837 | 29561015 | -1.388585958 | ABHD15 |
| 17 | 29560837 | 29561015 | -1.388585958 | ABHD15-AS1 |
| 17 | 29561632 | 29561827 | -2.509504057 | ABHD15-AS1 |
| 17 | 29561632 | 29561827 | -2.509504057 | ABHD15 |
| 17 | 29562501 | 29562751 | -2.888456678 | ABHD15-AS1 |
| 17 | 29562501 | 29562751 | -2.888456678 | ABHD15 |
| 17 | 29567811 | 29568006 | -1.030017983 | ABHD15-AS1 |
| 17 | 29567811 | 29568006 | -1.030017983 | TP53I13 |
| 17 | 29614277 | 29614559 | -1.342498645 | ANKRD13B |
| 17 | 29614277 | 29614559 | -1.342498645 | ABHD15-AS1 |
| 17 | 29630888 | 29631085 | -2.748126679 | ABHD15-AS1 |
| 17 | 29630888 | 29631085 | -2.748126679 | SSH2 |
| 17 | 29632570 | 29632767 | -2.681065657 | SSH2 |
| 17 | 29632570 | 29632767 | -2.681065657 | ABHD15-AS1 |
| 17 | 29636323 | 29636521 | -3.398438167 | ABHD15-AS1 |
| 17 | 29636323 | 29636521 | -3.398438167 | SSH2 |
| 17 | 29742589 | 29742727 | -6.994076049 | SSH2 |
| 1 | 86611457 | 86611637 | -1.530630008 | RP4-651E10.4 |
| 17 | 30624793 | 30624978 | -6.37462047 | SH3GL1P2 |
| 17 | 30624793 | 30624978 | -6.37462047 | AC005562.1 |
| 17 | 30632835 | 30633026 | -1.321063775 | AC005562.1 |
| 17 | 30632835 | 30633026 | -1.321063775 | LRRC37BP1 |
| 17 | 30834639 | 30834929 | -2.865901945 | CTD-2349P21.11 |
| 17 | 30834639 | 30834929 | -2.865901945 | ATAD5 |
| 17 | 30893851 | 30894045 | -2.288316035 | ATAD5 |
| 17 | 30904189 | 30904374 | -1.548994895 | TEFM |
| 1 | 86612146 | 86612340 | -2.150302163 | RP4-651E10.4 |
| 17 | 30958012 | 30958378 | -1.26084382 | ADAP2 |
| 17 | 30970742 | 30970937 | -2.740061902 | RNF135 |
| 17 | 30975398 | 30975579 | -1.202561015 | DPRXP4 |
| 17 | 30975398 | 30975579 | -1.202561015 | RNF135 |
| 17 | 31304365 | 31304567 | -2.087591758 | EVI2B |
| 17 | 31304365 | 31304567 | -2.087591758 | NF1 |
| 17 | 31318296 | 31318527 | -1.541892268 | NF1 |
| 17 | 31318296 | 31318527 | -1.541892268 | EVI2A |
| 17 | 31318296 | 31318527 | -1.541892268 | CTD-2370N5.3 |
| 17 | 31325950 | 31326123 | -1.944204154 | NF1 |
| 17 | 31531613 | 31531813 | -2.109144012 | RAB11FIP4 |
| 17 | 31998822 | 31998954 | -2.305225053 | SUZ12 |
| 1 | 86613880 | 86614040 | -1.509826714 | RP4-651E10.4 |
| 17 | 32160006 | 32160207 | -1.929439962 | CTD-2095E4.3 |
| 17 | 32160006 | 32160207 | -1.929439962 | RHOT1 |
| 17 | 32428845 | 32429039 | -1.081630636 | RP11-227G15.12 |
| 17 | 32430242 | 32430373 | -1.826989837 | RP11-227G15.12 |
| 17 | 32489797 | 32489963 | -3.038270777 | CDK5R1 |
| 1 | 86627760 | 86627952 | -1.125839188 | RP4-651E10.4 |
| 17 | 32490374 | 32490575 | -1.415277269 | CDK5R1 |
| 1 | 86630292 | 86630461 | -2.647408125 | RP4-651E10.4 |
| 17 | 35440789 | 35441008 | -1.915600832 | SLFN13 |
| 17 | 35470912 | 35471054 | 1.69591373 | SLFN12L |
| 17 | 35473260 | 35473441 | 1.170874768 | SLFN12L |
| 17 | 35480004 | 35480203 | 2.833588632 | SLFN12L |
| 1 | 86631734 | 86631911 | -2.211067903 | RP4-651E10.4 |
| 17 | 35489224 | 35489406 | -4.213132377 | SLFN12L |

| | | | | |
|---|---|---|---|---|
| 16 | 11179840 | 11179993 | -1.037374723 | CLEC16A |
| 16 | 11351258 | 11351458 | -2.373963326 | RMI2 |
| 16 | 11549408 | 11549606 | -6.936818178 | LITAF |
| 16 | 11797904 | 11798106 | -1.523881802 | RP11-486I11.2 |
| 16 | 11837809 | 11838009 | -1.588505588 | RSL1D1 |
| 16 | 11869692 | 11869893 | -1.746395803 | RP11-166B2.8 |
| 16 | 11869692 | 11869893 | -1.746395803 | GSPT1 |
| 1 | 52603309 | 52603606 | -1.624679734 | GPX7 |
| 16 | 14246714 | 14246897 | 1.380467716 | MKL2 |
| 16 | 14261139 | 14261476 | 1.679484022 | MKL2 |
| 16 | 14667918 | 14668028 | -1.41077729 | BFAR |
| 16 | 15036110 | 15036299 | 1.173994869 | PDXDC1 |
| 1 | 52687839 | 52688043 | -1.579858117 | COA7 |
| 16 | 15724369 | 15724588 | 1.055723147 | MYH11 |
| 16 | 15724369 | 15724588 | 1.055723147 | NDE1 |
| 16 | 15726703 | 15726874 | 1.570732543 | MYH11 |
| 16 | 15726703 | 15726874 | 1.570732543 | AF001548.5 |
| 16 | 16364884 | 16365040 | -1.183161107 | RP11-958N24.2 |
| 16 | 16376308 | 16376501 | -1.06535002 | RP11-958N24.2 |
| 16 | 17107503 | 17107699 | 1.813454025 | XYLT1 |
| 16 | 17108349 | 17108530 | 1.339983047 | XYLT1 |
| 16 | 17117711 | 17117927 | 2.92662165 | XYLT1 |
| 16 | 17123831 | 17124044 | 1.054423698 | XYLT1 |
| 16 | 17237776 | 17237973 | 1.300426937 | XYLT1 |
| 1 | 52857449 | 52857655 | -2.714861734 | ZYG11A |
| 16 | 18387193 | 18387330 | -1.154459494 | PKD1P5 |
| 16 | 19115513 | 19115715 | 1.812553332 | ITPRIPL2 |
| 16 | 19115513 | 19115715 | 1.812553332 | CTD-2349B8.1 |
| 16 | 19119027 | 19119228 | 1.506356796 | CTD-2349B8.1 |
| 16 | 19119027 | 19119228 | 1.506356796 | ITPRIPL2 |
| 16 | 19183684 | 19183888 | 1.570954713 | SYT17 |
| 16 | 19267207 | 19267406 | 1.511164282 | SYT17 |
| 16 | 19503379 | 19503644 | -1.393745628 | GDE1 |
| 1 | 52860627 | 52860785 | -3.232817331 | ZYG11A |
| 16 | 19537060 | 19537279 | -3.273187524 | CCP110 |
| 16 | 19704696 | 19704895 | -2.370886101 | C16orf62 |
| 16 | 19704696 | 19704895 | -2.370886101 | KNOP1 |
| 16 | 19705274 | 19705445 | -2.114750959 | C16orf62 |
| 16 | 19705274 | 19705445 | -2.114750959 | KNOP1 |
| 16 | 19706948 | 19707147 | -1.678926817 | AC002550.5 |
| 16 | 19706948 | 19707147 | -1.678926817 | KNOP1 |
| 16 | 19714385 | 19714599 | -2.606487787 | AC002550.5 |
| 16 | 19714385 | 19714599 | -2.606487787 | KNOP1 |
| 16 | 20723802 | 20723964 | 2.261780978 | THUMPD1 |
| 16 | 20723802 | 20723964 | 2.261780978 | AC004381.8 |
| 16 | 20723802 | 20723964 | 2.261780978 | ACSM3 |
| 16 | 20797825 | 20798004 | -2.570302214 | ERI2 |
| 1 | 52873372 | 52873569 | -3.366087719 | ZYG11A |
| 16 | 21301112 | 21301308 | -2.366349883 | CRYM |
| 16 | 21301112 | 21301308 | -2.366349883 | CRYM-AS1 |
| 16 | 21641255 | 21641458 | 9.273000695 | METTL9 |
| 16 | 21641255 | 21641458 | 9.273000695 | IGSF6 |
| 16 | 21647239 | 21647420 | 1.13795101 | METTL9 |
| 16 | 21647239 | 21647420 | 1.13795101 | IGSF6 |
| 17 | 35498461 | 35498644 | 6.963086214 | SLFN12L |
| 17 | 35498461 | 35498644 | 6.963086214 | RP11-1094M14.8 |
| 17 | 35514782 | 35514952 | 5.67616649 | SLFN12L |
| 17 | 35514782 | 35514952 | 5.67616649 | TOMM20P2 |
| 17 | 36496026 | 36496234 | -1.051290636 | MYO19 |
| 17 | 36496026 | 36496234 | -1.051290636 | ZNHIT3 |
| 17 | 36997811 | 36998021 | 1.750922434 | AATF |
| 17 | 37512374 | 37512489 | -1.239435943 | DUSP14 |
| 17 | 38399450 | 38399646 | -2.039001256 | SOCS7 |
| 1 | 2307137 | 2307331 | -1.432775018 | SKI |
| 17 | 38673067 | 38673387 | 2.572599779 | C17orf96 |
| 17 | 38702893 | 38703093 | 2.107182245 | CTB-58E17.1 |
| 17 | 38725753 | 38725972 | 1.502845381 | MLLT6 |
| 17 | 38767844 | 38768045 | -1.558418496 | PIP4K2B |
| 17 | 39175266 | 39175524 | -1.421584674 | CACNB1 |
| 17 | 39175266 | 39175524 | -1.421584674 | RP5-906A24.2 |
| 17 | 39347587 | 39347874 | -7.054384729 | FBXL20 |
| 1 | 87104551 | 87104750 | -1.741797086 | HS2ST1 |
| 1 | 87104551 | 87104750 | -1.741797086 | RP5-1052I5.2 |
| 1 | 87134146 | 87134348 | -1.907595019 | LINC01140 |
| 1 | 87134146 | 87134348 | -1.907595019 | RP5-1052I5.2 |
| 17 | 40123082 | 40123261 | -1.219735438 | MSL1 |
| 17 | 40126091 | 40126289 | -2.361039468 | MSL1 |
| 17 | 40129278 | 40129483 | -1.632243146 | MSL1 |
| 17 | 40163888 | 40164038 | 1.126776975 | CASC3 |
| 17 | 40356418 | 40356622 | -1.458730575 | RARA |
| 1 | 87153567 | 87153765 | -7.30158134 | LINC01140 |
| 1 | 87153567 | 87153765 | -7.30158134 | RP5-1052I5.2 |
| 17 | 40554494 | 40554684 | 6.090154042 | CCR7 |
| 17 | 40555708 | 40555903 | 6.069597459 | CCR7 |
| 17 | 40649026 | 40649213 | -1.542752864 | RP11-458J1.1 |
| 17 | 40649026 | 40649213 | -1.542752864 | RP5-1028K7.3 |
| 17 | 41810941 | 41811125 | -2.19482609 | P3H4 |
| 17 | 41968318 | 41968521 | 2.618541427 | CNP |
| 17 | 42121532 | 42121716 | 1.45548864 | CTD-2132N18.3 |
| 17 | 42121532 | 42121716 | 1.45548864 | HSPB9 |
| 17 | 42122437 | 42122609 | 1.192633734 | CTD-2132N18.3 |
| 17 | 42122437 | 42122609 | 1.192633734 | HSPB9 |
| 17 | 42192397 | 42192608 | -2.324354923 | GHDC |
| 17 | 42265148 | 42265343 | 6.313424882 | STAT5B |
| 17 | 42553832 | 42554031 | -1.821992324 | RP11-400F19.8 |
| 17 | 42553832 | 42554031 | -1.821992324 | RP11-400F19.6 |
| 17 | 42553832 | 42554031 | -1.821992324 | HSD17B1 |
| 17 | 42668103 | 42668230 | -1.542214391 | PLEKHH3 |
| 17 | 42980368 | 42980500 | -3.548577132 | PTGES3L |
| 17 | 42980368 | 42980500 | -3.548577132 | PTGES3L-AARSD1 |
| 17 | 43128451 | 43128554 | -2.079682431 | BRCA1 |
| 17 | 43128451 | 43128554 | -2.079682431 | NBR2 |
| 17 | 43136209 | 43136407 | -1.378896744 | NBR2 |
| 17 | 43136209 | 43136407 | -1.378896744 | BRCA1 |
| 17 | 43138398 | 43138587 | -3.192488519 | NBR2 |
| 17 | 43138398 | 43138587 | -3.192488519 | BRCA1 |
| 17 | 43144964 | 43145173 | -1.717497201 | NBR2 |
| 17 | 43144964 | 43145173 | -1.717497201 | BRCA1 |

| | | | | |
|---|---|---|---|---|
| 16 | 22325993 | 22326186 | -1.282237389 | POLR3E |
| 16 | 22333617 | 22333829 | -1.233429699 | POLR3E |
| 16 | 22392808 | 22392978 | -1.196606512 | MFSD13B |
| 16 | 22392808 | 22392978 | -1.196606512 | CDR2 |
| 1 | 52893057 | 52893299 | -2.271594359 | ZYG11A |
| 16 | 22419881 | 22420090 | -1.394726349 | CDR2 |
| 16 | 22419881 | 22420090 | -1.394726349 | RRN3P3 |
| 16 | 22420909 | 22421116 | -1.96614513 | RRN3P3 |
| 16 | 22420909 | 22421116 | -1.96614513 | CDR2 |
| 16 | 23064437 | 23064582 | -2.056418403 | USP31 |
| 16 | 23067573 | 23067772 | -2.538837226 | USP31 |
| 16 | 23068212 | 23068411 | -2.095680077 | USP31 |
| 16 | 23069123 | 23069386 | -1.570369858 | USP31 |
| 16 | 23467472 | 23467672 | -1.781948015 | GGA2 |
| 16 | 23524185 | 23524398 | -1.647943823 | EARS2 |
| 16 | 23629785 | 23629989 | -1.589837818 | PALB2 |
| 1 | 52994900 | 52995006 | -1.908206008 | SCP2 |
| 1 | 52994900 | 52995006 | -1.908206008 | TUBBP10 |
| 16 | 23635978 | 23636209 | -1.636609924 | PALB2 |
| 16 | 23669340 | 23669541 | -1.045426241 | DCTN5 |
| 16 | 24930955 | 24931153 | -1.392125273 | ARHGAP17 |
| 1 | 53009595 | 53009823 | -8.01666848 | SCP2 |
| 16 | 25247121 | 25247321 | -1.049900922 | ZKSCAN2 |
| 16 | 25256852 | 25257020 | -1.356437015 | ZKSCAN2 |
| 16 | 27220843 | 27221043 | -1.353146584 | KDM8 |
| 16 | 27221210 | 27221408 | -1.424474153 | KDM8 |
| 16 | 27448880 | 27449106 | 1.275505577 | IL21R |
| 16 | 27448880 | 27449106 | 1.275505577 | IL21R-AS1 |
| 1 | 53010316 | 53010422 | -3.077644828 | SCP2 |
| 16 | 27778834 | 27779028 | -1.758846864 | KIAA0556 |
| 16 | 28098436 | 28098640 | -1.13399963 | XPO6 |
| 16 | 28115059 | 28115257 | -1.255885744 | XPO6 |
| 16 | 28156144 | 28156343 | -1.066673239 | XPO6 |
| 16 | 28498667 | 28498872 | 1.119270046 | APOBR |
| 16 | 28934819 | 28934947 | -4.654272941 | RABEP2 |
| 16 | 28934819 | 28934947 | -4.654272941 | CD19 |
| 16 | 28935436 | 28935628 | -2.555860542 | RABEP2 |
| 16 | 28935436 | 28935628 | -2.555860542 | CD19 |
| 16 | 28963781 | 28963985 | -1.353688673 | NFATC2IP |
| 16 | 28963781 | 28963985 | -1.353688673 | RP11-264B17.2 |
| 16 | 29145350 | 29145509 | -2.234543019 | RP11-426C22.5 |
| 16 | 29154411 | 29154597 | -5.181426374 | RP11-426C22.5 |
| 16 | 29211061 | 29211262 | -1.594997185 | RP11-426C22.5 |
| 16 | 29364730 | 29364951 | 1.597844049 | SNX29P2 |
| 16 | 29364730 | 29364951 | 1.597844049 | RP11-231C14.7 |
| 16 | 29664224 | 29664427 | -1.877399565 | SPN |
| 16 | 29664224 | 29664427 | -1.877399565 | AC009133.22 |
| 16 | 29665879 | 29666077 | -2.907764952 | SPN |
| 16 | 29665879 | 29666077 | -2.907764952 | AC009133.22 |
| 16 | 29743605 | 29743783 | -1.512622866 | C16orf54 |
| 1 | 53210855 | 53211085 | -1.270736193 | RP5-1024G6.2 |
| 1 | 53210855 | 53211085 | -1.270736193 | CPT2 |
| 16 | 29744012 | 29744209 | -1.063153807 | C16orf54 |
| 16 | 29744551 | 29744705 | -2.063836882 | C16orf54 |
| 17 | 43144964 | 43145173 | -1.717497201 | CTD-3199J23.6 |
| 17 | 43162770 | 43162959 | -2.641726974 | RP11-242D8.1 |
| 17 | 43162770 | 43162959 | -2.641726974 | BRCA1 |
| 17 | 43230946 | 43231146 | 2.067598302 | LINC00854 |
| 17 | 43246576 | 43246774 | -8.323448085 | LINC00854 |
| 17 | 43399996 | 43400112 | -1.264866245 | ARL4D |
| 17 | 43400288 | 43400483 | -2.458069328 | ARL4D |
| 17 | 43766332 | 43766532 | -1.335459884 | DUSP3 |
| 17 | 44148050 | 44148241 | -2.896366887 | C17orf53 |
| 17 | 44206904 | 44207104 | -1.306475618 | UBTF |
| 17 | 44206904 | 44207104 | -1.306475618 | CTB-175E5.7 |
| 1 | 88983570 | 88983766 | 1.705338877 | KYAT3 |
| 1 | 88983570 | 88983766 | 1.705338877 | RBMXL1 |
| 17 | 44400357 | 44400506 | -1.458467585 | GPATCH8 |
| 17 | 44666570 | 44666687 | -3.307354922 | MEIOC |
| 17 | 44667627 | 44667811 | -3.160964047 | MEIOC |
| 17 | 44926328 | 44926430 | -5.31467831 | KIF18B |
| 17 | 44946606 | 44946922 | -5.959431619 | KIF18B |
| 17 | 44947252 | 44947451 | -7.115160358 | KIF18B |
| 17 | 45107344 | 45107555 | -1.614326627 | NMT1 |
| 17 | 45147754 | 45147951 | -1.573949348 | AC142472.6 |
| 17 | 45190846 | 45191047 | 1.505831459 | CTD-2020K17.1 |
| 17 | 45218905 | 45219155 | 2.543212834 | CTD-2020K17.1 |
| 1 | 2307708 | 2307907 | -1.899707734 | SKI |
| 17 | 45468378 | 45468553 | -7.055323214 | PLEKHM1 |
| 17 | 45475350 | 45475494 | -1.197401124 | PLEKHM1 |
| 17 | 45514612 | 45514813 | -2.829073998 | LRRC37A4P |
| 17 | 45515322 | 45515520 | -1.431084087 | LRRC37A4P |
| 1 | 89104414 | 89104552 | 7.804259672 | PTGES3P1 |
| 17 | 45549805 | 45549943 | -2.447506122 | RP11-798G7.7 |
| 17 | 45549805 | 45549943 | -2.447506122 | RP11-798G7.6 |
| 17 | 45549805 | 45549943 | -2.447506122 | LRRC37A4P |
| 17 | 46309988 | 46310179 | -3.428990498 | LRRC37A |
| 17 | 46309988 | 46310179 | -3.428990498 | ARL17B |
| 17 | 46548822 | 46549021 | -1.083279719 | ARL17A |
| 17 | 46548822 | 46549021 | -1.083279719 | AC138645.2 |
| 17 | 46548822 | 46549021 | -1.083279719 | AC138645.1 |
| 17 | 46548822 | 46549021 | -1.083279719 | LRRC37A2 |
| 17 | 46556082 | 46556286 | 1.204363682 | AC138645.2 |
| 17 | 46556082 | 46556286 | 1.204363682 | AC138645.1 |
| 17 | 46556082 | 46556286 | 1.204363682 | ARL17A |
| 17 | 46768470 | 46768673 | -2.145188127 | WNT3 |
| 17 | 46964384 | 46964588 | -1.540390467 | GOSR2 |
| 17 | 46964384 | 46964588 | -1.540390467 | RP11-156P1.2 |
| 17 | 47017938 | 47018135 | -1.831492516 | LRRC37A17P |
| 17 | 47017938 | 47018135 | -1.831492516 | RP11-156P1.2 |
| 17 | 47017938 | 47018135 | -1.831492516 | RP11-156P1.3 |
| 17 | 47056438 | 47056643 | -2.348964951 | RP11-156P1.3 |
| 17 | 47384121 | 47384354 | -1.167300962 | EFCAB13 |
| 17 | 47387475 | 47387640 | -2.136649924 | EFCAB13 |
| 17 | 47434891 | 47435066 | -6.293888758 | EFCAB13 |
| 17 | 47437484 | 47437679 | -2.596628753 | EFCAB13 |
| 17 | 47745134 | 47745287 | 8.074525164 | TBX21 |
| 1 | 89421458 | 89421658 | 10.2922441 | GBP1P1 |

| | | | | |
|---|---|---|---|---|
| 16 | 29819632 | 29819825 | -1.820241136 | CTD-2574D22.6 |
| 16 | 29819632 | 29819825 | -1.820241136 | AC009133.12 |
| 16 | 29819632 | 29819825 | -1.820241136 | AC009133.23 |
| 16 | 29819632 | 29819825 | -1.820241136 | PAGR1 |
| 16 | 29915138 | 29915328 | -2.036969197 | ASPHD1 |
| 16 | 29915138 | 29915328 | -2.036969197 | CTD-2574D22.3 |
| 16 | 29915138 | 29915328 | -2.036969197 | KCTD13 |
| 16 | 29986525 | 29986728 | -1.001453129 | TAOK2 |
| 16 | 30355242 | 30355442 | 8.654405031 | CD2BP2 |
| 16 | 30355242 | 30355442 | 8.654405031 | RP11-347C12.10 |
| 16 | 30356808 | 30357024 | 1.131554402 | RP11-347C12.10 |
| 1 | 53244605 | 53244840 | -1.770860051 | LRP8 |
| 16 | 30365536 | 30365742 | -1.515766792 | TBC1D10B |
| 16 | 30369325 | 30369525 | -1.024601869 | TBC1D10B |
| 16 | 30397461 | 30397666 | -1.087197755 | ZNF48 |
| 16 | 30521626 | 30521902 | -1.200542381 | ITGAL |
| 16 | 30524384 | 30524585 | -1.274507194 | ZNF768 |
| 16 | 30525687 | 30525887 | -1.309106958 | ZNF768 |
| 16 | 30532433 | 30532659 | -1.065194791 | AC002310.11 |
| 16 | 30532433 | 30532659 | -1.065194791 | ZNF747 |
| 16 | 30555091 | 30555322 | -1.082063097 | AC002310.13 |
| 16 | 30555091 | 30555322 | -1.082063097 | ZNF764 |
| 1 | 1663428 | 1663629 | -1.033425326 | SLC35E2B |
| 1 | 53767852 | 53768052 | -2.531417988 | NDC1 |
| 16 | 30569758 | 30569957 | -1.501776351 | ZNF688 |
| 16 | 30592397 | 30592579 | -1.364661045 | RP11-146F11.5 |
| 16 | 30604127 | 30604297 | -2.60394823 | RP11-146F11.5 |
| 16 | 30604127 | 30604297 | -2.60394823 | ZNF689 |
| 16 | 30604917 | 30605217 | -1.129361265 | RP11-146F11.5 |
| 16 | 30604917 | 30605217 | -1.129361265 | ZNF689 |
| 1 | 54096354 | 54096548 | -1.568024784 | TCEANC2 |
| 16 | 30757000 | 30757274 | -1.629620576 | PHKG2 |
| 16 | 30779848 | 30780106 | -1.033345795 | ZNF629 |
| 1 | 54173409 | 54173623 | -2.590305086 | RP11-446E24.4 |
| 1 | 54173409 | 54173623 | -2.590305086 | CYB5RL |
| 16 | 30948126 | 30948267 | -1.0876655 | FBXL19 |
| 16 | 30957309 | 30957493 | -1.501108764 | SETD1A |
| 16 | 30964927 | 30965142 | -1.005168735 | SETD1A |
| 16 | 31073064 | 31073284 | -1.064948949 | ZNF668 |
| 1 | 54174500 | 54174681 | -1.747205507 | RP11-446E24.4 |
| 1 | 54174500 | 54174681 | -1.747205507 | CYB5RL |
| 16 | 31112068 | 31112267 | -3.221578039 | BCKDK |
| 16 | 31203036 | 31203228 | -5.282260835 | PYCARD |
| 16 | 31203036 | 31203228 | -5.282260835 | PYCARD-AS1 |
| 16 | 31311866 | 31312030 | 6.108372929 | ITGAM |
| 16 | 31358073 | 31358290 | 5.119657986 | ITGAX |
| 16 | 31382017 | 31382216 | 5.634617464 | ITGAX |
| 16 | 31436977 | 31437115 | 6.436722056 | ZNF843 |
| 1 | 54609116 | 54609277 | -1.756326892 | FAM151A |
| 1 | 54609116 | 54609277 | -1.756326892 | ACOT11 |
| 16 | 31457860 | 31458043 | -1.007111576 | RP11-452L6.5 |
| 16 | 31462592 | 31462826 | -1.687646428 | ARMC5 |
| 16 | 31466808 | 31467010 | -1.890042643 | ARMC5 |
| 16 | 31708116 | 31708258 | -2.398055082 | CLUHP3 |
| 17 | 48050665 | 48050866 | -1.095674851 | NFE2L1 |
| 17 | 48058626 | 48058824 | -1.062526651 | NFE2L1 |
| 17 | 48059568 | 48059769 | -1.083241894 | NFE2L1 |
| 17 | 48596011 | 48596193 | -3.216306946 | HOXB-AS3 |
| 17 | 48596011 | 48596193 | -3.216306946 | HOXB6 |
| 17 | 48596011 | 48596193 | -3.216306946 | HOXB3 |
| 17 | 48607758 | 48607957 | -2.225910824 | HOXB7 |
| 17 | 49298116 | 49298318 | -2.547214276 | ZNF652 |
| 17 | 49317008 | 49317382 | -2.108276276 | ZNF652 |
| 17 | 49579233 | 49579425 | -4.185222621 | NXPH3 |
| 17 | 49969264 | 49969406 | -7.330137254 | DLX4 |
| 17 | 49974057 | 49974257 | -4.022976988 | DLX4 |
| 17 | 49991273 | 49991475 | -2.322829893 | DLX3 |
| 1 | 89426084 | 89426270 | 9.539437634 | GBP1P1 |
| 17 | 50360122 | 50360336 | -2.205865502 | XYLT2 |
| 17 | 50381091 | 50381276 | -5.406088653 | LRRC59 |
| 17 | 50381091 | 50381276 | -5.406088653 | EME1 |
| 17 | 50382811 | 50383006 | 1.993538364 | LRRC59 |
| 17 | 50485518 | 50485717 | -2.03547298 | RSAD1 |
| 17 | 50522853 | 50523053 | -1.700373421 | MYCBPAP |
| 17 | 50526092 | 50526377 | -2.006545 | MYCBPAP |
| 17 | 50863736 | 50863938 | -1.887630296 | TOB1 |
| 17 | 56832980 | 56833226 | 2.885045809 | C17orf67 |
| 17 | 56891966 | 56892167 | 1.690907002 | TRIM25 |
| 17 | 56950663 | 56950838 | -1.06260555 | COIL |
| 17 | 57106212 | 57106412 | -1.183229771 | AKAP1 |
| 17 | 57106920 | 57107083 | -1.230180074 | AKAP1 |
| 17 | 57109902 | 57110079 | -2.250149952 | AKAP1 |
| 17 | 57632140 | 57632344 | -6.569535827 | MSI2 |
| 17 | 58980157 | 58980356 | -1.525413606 | PPM1E |
| 17 | 59397052 | 59397228 | -2.005062626 | YPEL2 |
| 17 | 60043809 | 60043940 | -1.411384678 | HEATR6 |
| 17 | 60179227 | 60179428 | -1.699838065 | USP32 |
| 17 | 60181598 | 60181802 | -1.438833761 | USP32 |
| 17 | 60447738 | 60447927 | -3.217143904 | APPBP2 |
| 17 | 60483719 | 60483910 | -6.56608919 | APPBP2 |
| 17 | 60663177 | 60663380 | -1.133651815 | PPM1D |
| 17 | 61115134 | 61115243 | -3.928990498 | BCAS3 |
| 17 | 61683406 | 61683785 | 1.948585383 | BRIP1 |
| 17 | 62358396 | 62358552 | 6.399640811 | EFCAB3 |
| 17 | 62477243 | 62477441 | -1.256083584 | TLK2 |
| 17 | 62626123 | 62626324 | 1.492775749 | CTD-3035K23.6 |
| 17 | 62626586 | 62626784 | -1.658383603 | CTD-3035K23.7 |
| 17 | 63420240 | 63420442 | -2.218808963 | RP11-269G24.3 |
| 17 | 63420240 | 63420442 | -2.218808963 | TANC2 |
| 17 | 63421742 | 63421959 | -2.265694992 | RP11-269G24.3 |
| 17 | 63421742 | 63421959 | -2.265694992 | TANC2 |
| 17 | 63426536 | 63426725 | -2.146576526 | TANC2 |
| 17 | 63426536 | 63426725 | -2.146576526 | RP11-269G24.3 |
| 17 | 63608075 | 63608302 | -1.027922999 | TACO1 |
| 17 | 64043606 | 64043809 | -1.768911889 | ERN1 |
| 17 | 64129178 | 64129348 | -1.397697704 | ERN1 |
| 17 | 64148743 | 64148941 | -2.046630174 | TEX2 |
| 17 | 64193655 | 64193875 | -1.968426989 | TEX2 |

| | | | | |
|---|---|---|---|---|
| 16 | 31764729 | 31764839 | 3.222686789 | ZNF720 |
| 16 | 31774135 | 31774288 | 2.432222374 | ZNF720 |
| 1 | 54609604 | 54609801 | -3.012444773 | ACOT11 |
| 1 | 54609604 | 54609801 | -3.012444773 | FAM151A |
| 16 | 31776810 | 31776970 | 1.957178713 | ZNF720 |
| 16 | 31794140 | 31794331 | 1.652645285 | ZNF720 |
| 1 | 54653383 | 54653580 | 1.454984994 | MROH7-TTC4 |
| 1 | 54653383 | 54653580 | 1.454984994 | MROH7 |
| 16 | 46581774 | 46581974 | -3.896641379 | SHCBP1 |
| 16 | 46597980 | 46598172 | -3.898784822 | SHCBP1 |
| 16 | 47083314 | 47083539 | -1.378499314 | NETO2 |
| 16 | 47557223 | 47557397 | -7.280822974 | PHKB |
| 1 | 54741692 | 54742004 | -1.408077071 | MROH7-TTC4 |
| 1 | 54741692 | 54742004 | -1.408077071 | TTC4 |
| 16 | 48542893 | 48543084 | 1.657267706 | N4BP1 |
| 16 | 48561148 | 48561346 | 1.395479645 | N4BP1 |
| 16 | 48561148 | 48561346 | 1.395479645 | RP11-44I10.3 |
| 16 | 48561978 | 48562209 | 1.109112791 | RP11-44I10.3 |
| 16 | 48561978 | 48562209 | 1.109112791 | N4BP1 |
| 1 | 54757816 | 54758011 | -1.26223163 | PARS2 |
| 16 | 50671785 | 50671940 | -1.123089337 | SNX20 |
| 16 | 50796385 | 50796581 | 1.149996546 | CYLD |
| 16 | 50796385 | 50796581 | 1.149996546 | RP11-327F22.4 |
| 16 | 53185180 | 53185375 | 1.29354727 | CHD9 |
| 1 | 54758620 | 54758850 | -1.062966484 | PARS2 |
| 16 | 53280372 | 53280579 | -2.860703971 | CHD9 |
| 16 | 53826051 | 53826396 | -1.286674145 | FTO |
| 16 | 53844029 | 53844183 | -1.427077065 | FTO |
| 16 | 54239563 | 54239752 | -7.34818495 | RP11-324D17.2 |
| 1 | 54851193 | 54851392 | -1.914052838 | DHCR24 |
| 16 | 54264288 | 54264483 | -4.864219747 | RP11-324D17.1 |
| 16 | 54270604 | 54270789 | -3.380708135 | RP11-324D17.1 |
| 16 | 54285181 | 54285381 | -3.491341467 | IRX3 |
| 16 | 54286528 | 54286689 | -3.248412512 | IRX3 |
| 16 | 54286528 | 54286689 | -3.248412512 | RP11-434E6.5 |
| 16 | 54289098 | 54289291 | -2.928850742 | RP11-434E6.5 |
| 16 | 54291773 | 54291962 | -1.747769072 | RP11-434E6.4 |
| 16 | 54933667 | 54933869 | -1.44977431 | IRX5 |
| 16 | 56476011 | 56476256 | -1.090827576 | BBS2 |
| 16 | 56476011 | 56476256 | -1.090827576 | OGFOD1 |
| 1 | 54852043 | 54852249 | -1.546862953 | DHCR24 |
| 16 | 56991926 | 56992129 | -1.281097575 | NLRC5 |
| 16 | 57207902 | 57208103 | -1.143007229 | RSPRY1 |
| 16 | 57460486 | 57460677 | -1.232601112 | COQ9 |
| 16 | 57746375 | 57746534 | 4.669925001 | KATNB1 |
| 16 | 57747498 | 57747695 | -2.259816042 | KATNB1 |
| 1 | 55068783 | 55069025 | -1.047250138 | USP24 |
| 16 | 57996217 | 57996402 | -4.100087776 | ZNF319 |
| 16 | 58012854 | 58013013 | -3.261656511 | USB1 |
| 16 | 58392432 | 58392574 | -2.345767082 | RP11-481J2.3 |
| 16 | 58392432 | 58392574 | -2.345767082 | GINS3 |
| 16 | 58404443 | 58404657 | -1.455716841 | GINS3 |
| 16 | 58518661 | 58518864 | -1.345274538 | SETD6 |
| 16 | 66510748 | 66510964 | -1.171842432 | TK2 |
| 17 | 64212782 | 64213054 | -2.461692746 | TEX2 |
| 17 | 64322625 | 64322815 | -1.453692847 | PECAM1 |
| 17 | 64323502 | 64323703 | -1.562589917 | PECAM1 |
| 17 | 64369930 | 64370127 | 6.315588528 | PECAM1 |
| 17 | 64375213 | 64375411 | -1.494959771 | PECAM1 |
| 17 | 64412160 | 64412334 | -7.094178516 | PECAM1 |
| 17 | 64413707 | 64413838 | -3.484950197 | PECAM1 |
| 17 | 64496516 | 64496713 | -1.38405576 | POLG2 |
| 17 | 64500223 | 64500425 | -2.317911901 | DDX5 |
| 17 | 64778323 | 64778521 | 8.357686092 | RP13-104F24.2 |
| 17 | 64784052 | 64784249 | 2.014021519 | PLEKHM1P1 |
| 17 | 66757117 | 66757296 | 1.073454174 | PRKCA |
| 1 | 89762027 | 89762240 | 1.385757029 | RP11-302M6.4 |
| 1 | 89762027 | 89762240 | 1.385757029 | LRRC8C |
| 17 | 67991867 | 67992066 | -1.962914422 | C17orf58 |
| 17 | 68125711 | 68125875 | -1.057116963 | LRRC37A16P |
| 17 | 68135439 | 68135551 | 1.2865442 | LRRC37A16P |
| 17 | 68135439 | 68135551 | 1.2865442 | RP11-147L13.13 |
| 17 | 68135439 | 68135551 | 1.2865442 | SH3GL1P3 |
| 17 | 68269106 | 68269313 | -2.489720436 | SLC16A6 |
| 17 | 68269106 | 68269313 | -2.489720436 | ARSG |
| 17 | 68271414 | 68271612 | -2.970331506 | SLC16A6 |
| 17 | 68271414 | 68271612 | -2.970331506 | ARSG |
| 17 | 68307491 | 68307672 | -2.186856775 | ARSG |
| 17 | 68399296 | 68399400 | -5.44208526 | ARSG |
| 17 | 70174879 | 70175077 | 1.025934657 | KCNJ2 |
| 17 | 72646627 | 72646826 | -1.681632272 | SLC39A11 |
| 17 | 72694811 | 72694993 | -6.771455473 | SLC39A11 |
| 1 | 89869393 | 89869591 | -1.723087761 | RP11-302M6.4 |
| 1 | 89869393 | 89869591 | -1.723087761 | LRRC8D |
| 17 | 72818170 | 72818341 | -6.02882667 | SLC39A11 |
| 17 | 73068029 | 73068267 | -2.516112869 | RP11-766H1.1 |
| 17 | 73068029 | 73068267 | -2.516112869 | SLC39A11 |
| 17 | 73248504 | 73248700 | -1.169717407 | C17orf80 |
| 17 | 73248504 | 73248700 | -1.169717407 | CPSF4L |
| 17 | 73285309 | 73285456 | -1.184604049 | CDC42EP4 |
| 1 | 89924036 | 89924206 | -2.372136621 | RP11-302M6.4 |
| 1 | 89924036 | 89924206 | -2.372136621 | LRRC8D |
| 17 | 74431532 | 74431794 | -2.446790628 | GPRC5C |
| 17 | 74450349 | 74450490 | -3.605323249 | GPRC5C |
| 17 | 74484410 | 74484608 | -2.611375936 | CD300A |
| 17 | 74522414 | 74522611 | -3.710919133 | CD300LB |
| 17 | 74544675 | 74544853 | -7.022367813 | CD300C |
| 17 | 74838382 | 74838549 | -1.195515024 | TMEM104 |
| 17 | 75003774 | 75003976 | -1.486693199 | CDR2L |
| 1 | 2353558 | 2353725 | -1.208208484 | MORN1 |
| 1 | 89933836 | 89934035 | -1.321375958 | LRRC8D |
| 17 | 75100234 | 75100417 | -6.179462935 | SLC16A5 |
| 17 | 75492849 | 75493038 | -1.10049317 | TMEM94 |
| 17 | 75502046 | 75502199 | -1.415037499 | CASKIN2 |
| 1 | 89935304 | 89935515 | -1.578645177 | LRRC8D |
| 17 | 75524456 | 75524649 | -7.505089202 | TSEN54 |
| 17 | 75588685 | 75588891 | 10.53863605 | MYO15B |
| 17 | 75661030 | 75661227 | -1.543731421 | RECQL5 |

| | | | | |
|---|---|---|---|---|
| 16 | 66510748 | 66510964 | -1.171842432 | RP11-403P17.5 |
| 16 | 66579588 | 66579711 | 6.801639305 | CMTM2 |
| 16 | 66613216 | 66613416 | 1.269278845 | CMTM3 |
| 16 | 67160270 | 67160416 | -2.603946426 | FBXL8 |
| 16 | 67160270 | 67160416 | -2.603946426 | TRADD |
| 1 | 58781993 | 58782195 | 5.365063836 | JUN |
| 16 | 67164351 | 67164537 | -6.114409345 | FBXL8 |
| 16 | 67164351 | 67164537 | -6.114409345 | RP11-5A19.5 |
| 16 | 67180178 | 67180332 | -1.155579232 | KIAA0895L |
| 16 | 67183345 | 67183534 | -1.487294492 | KIAA0895L |
| 16 | 67484284 | 67484399 | -2.857509777 | RP11-297D21.4 |
| 16 | 67611190 | 67611439 | -1.256752795 | CTCF |
| 16 | 67675602 | 67675860 | -1.792619936 | GFOD2 |
| 16 | 67724950 | 67725149 | -2.414572649 | RANBP10 |
| 1 | 58783031 | 58783234 | 5.204338581 | JUN |
| 16 | 67843502 | 67843683 | -1.413761883 | CENPT |
| 16 | 67843502 | 67843683 | -1.413761883 | THAP11 |
| 16 | 67908859 | 67909067 | -1.437911953 | PSKH1 |
| 16 | 67927818 | 67928118 | -1.531865779 | PSKH1 |
| 16 | 67927818 | 67928118 | -1.531865779 | CTRL |
| 16 | 67945797 | 67945976 | -1.702023358 | SLC12A4 |
| 16 | 68021823 | 68022022 | -1.289470785 | DUS2 |
| 16 | 68021823 | 68022022 | -1.289470785 | DDX28 |
| 16 | 68259648 | 68259848 | 1.682291427 | RP11-96D1.7 |
| 16 | 68259648 | 68259848 | 1.682291427 | PLA2G15 |
| 16 | 68268134 | 68268237 | 4.408491812 | SLC7A6 |
| 16 | 68860058 | 68860258 | -2.441321525 | TANGO6 |
| 16 | 69033718 | 69033853 | -3.017261938 | RPS2P45 |
| 16 | 69033718 | 69033853 | -3.017261938 | TANGO6 |
| 16 | 69083676 | 69083793 | -2.354638521 | TANGO6 |
| 16 | 69109738 | 69109882 | 1.260976352 | HAS3 |
| 16 | 69118249 | 69118448 | -1.07709979 | HAS3 |
| 16 | 69118249 | 69118448 | -1.07709979 | CHTF8 |
| 16 | 69328908 | 69329060 | -1.423470916 | PDF |
| 16 | 69328908 | 69329060 | -1.423470916 | COG8 |
| 16 | 69334837 | 69335039 | -1.251069479 | COG8 |
| 16 | 69341559 | 69341766 | -1.869306226 | RP11-343C2.7 |
| 16 | 69341559 | 69341766 | -1.869306226 | RP11-343C2.9 |
| 16 | 69341559 | 69341766 | -1.869306226 | NIP7 |
| 16 | 69366892 | 69367096 | -1.303358321 | TERF2 |
| 16 | 69691937 | 69692200 | 1.166082025 | NFAT5 |
| 1 | 61088338 | 61088534 | -3.541463691 | NFIA |
| 16 | 69693353 | 69693581 | 1.482508076 | NFAT5 |
| 16 | 69695062 | 69695256 | 2.999736301 | NFAT5 |
| 16 | 69696381 | 69696484 | 2.21066116 | NFAT5 |
| 16 | 69742030 | 69742235 | -1.125611059 | NOB1 |
| 16 | 69742030 | 69742235 | -1.125611059 | CTD-2033A16.3 |
| 16 | 69743033 | 69743212 | -8.201247913 | NOB1 |
| 16 | 70250851 | 70251079 | -1.440149945 | EXOSC6 |
| 16 | 70371958 | 70372160 | -1.183349553 | DDX19A |
| 16 | 70371958 | 70372160 | -1.183349553 | RP11-529K1.3 |
| 16 | 70571841 | 70572041 | -2.120000128 | SF3B3 |
| 16 | 70699149 | 70699346 | -7.100525907 | VAC14 |
| 1 | 62444821 | 62445024 | -1.516618581 | USP1 |
| 17 | 75880270 | 75880458 | -2.451866856 | TRIM65 |
| 17 | 76077800 | 76077957 | -1.829403266 | ZACN |
| 17 | 76293188 | 76293434 | -1.163575601 | QRICH2 |
| 17 | 76311400 | 76311605 | -1.858279657 | PRPSAP1 |
| 17 | 77343971 | 77344192 | -2.067442515 | Sep-09 |
| 17 | 78105874 | 78106069 | -1.562123714 | TNRC6C |
| 17 | 78110626 | 78110827 | -1.782954668 | TMC6 |
| 17 | 78110626 | 78110827 | -1.782954668 | TNRC6C-AS1 |
| 17 | 78111815 | 78111986 | -1.768040105 | TMC6 |
| 17 | 78112600 | 78112801 | -1.313126048 | TMC6 |
| 17 | 78344128 | 78344316 | 6.843523781 | AC061992.2 |
| 17 | 78345120 | 78345318 | 6.309230989 | AC061992.2 |
| 17 | 78358008 | 78358205 | 4.077613703 | SOCS3 |
| 17 | 78891071 | 78891249 | -4.72766361 | TIMP2 |
| 17 | 78891071 | 78891249 | -4.72766361 | CEP295NL |
| 17 | 79784754 | 79784900 | -2.853679566 | CBX2 |
| 17 | 79794674 | 79794844 | -1.484838877 | CBX8 |
| 17 | 79834778 | 79834982 | -1.680590946 | CBX4 |
| 17 | 79932440 | 79932643 | -1.438587987 | TBC1D16 |
| 17 | 79939663 | 79939864 | -1.038935521 | TBC1D16 |
| 1 | 90939381 | 90939579 | -1.161219537 | ZNF644 |
| 17 | 80045652 | 80045857 | -4.293692175 | CCDC40 |
| 17 | 80210046 | 80210240 | -1.789974551 | SGSH |
| 17 | 80211530 | 80211702 | -1.562812668 | SGSH |
| 17 | 80295498 | 80295698 | 1.230218655 | RNF213 |
| 17 | 80319539 | 80319754 | 4.245573743 | RNF213 |
| 17 | 80332078 | 80332279 | 1.717922089 | RNF213 |
| 17 | 80339343 | 80339543 | 1.655338584 | RNF213 |
| 17 | 80345426 | 80345592 | 2.628789746 | RNF213 |
| 17 | 80347253 | 80347438 | 2.606278156 | RNF213 |
| 17 | 80348002 | 80348192 | 1.67527655 | RNF213 |
| 17 | 80365224 | 80365418 | 7.308159548 | CTD-2047H16.4 |
| 17 | 80365224 | 80365418 | 7.308159548 | RNF213 |
| 17 | 80384701 | 80384909 | 4.362365322 | RNF213 |
| 17 | 80384701 | 80384909 | 4.362365322 | CTD-2047H16.4 |
| 17 | 80394056 | 80394302 | 2.017672804 | CTD-2047H16.4 |
| 17 | 80394056 | 80394302 | 2.017672804 | RNF213 |
| 17 | 80395516 | 80395715 | 1.30426187 | RNF213 |
| 17 | 80395516 | 80395715 | 1.30426187 | CTD-2047H16.4 |
| 17 | 80581966 | 80582131 | 5.715226276 | RPTOR |
| 17 | 80727249 | 80727378 | -4.563352236 | RPTOR |
| 17 | 80891132 | 80891245 | -4.204695468 | RPTOR |
| 17 | 80930697 | 80930893 | 5.961849442 | RPTOR |
| 17 | 80957260 | 80957390 | 5.76569073 | RPTOR |
| 17 | 80970698 | 80970865 | 2.245926548 | CTD-2561B21.5 |
| 1 | 2364473 | 2364644 | -7.522645201 | MORN1 |
| 17 | 81030559 | 81030802 | -3.555925759 | BAIAP2-AS1 |
| 17 | 81032145 | 81032450 | -2.679424339 | BAIAP2-AS1 |
| 17 | 81114684 | 81114871 | -5.879319819 | BAIAP2 |
| 17 | 81304898 | 81305097 | -2.17472501 | LINC00482 |
| 17 | 81327252 | 81327448 | -5.011183907 | TMEM105 |
| 17 | 81461375 | 81461583 | 6.206813965 | BAHCC1 |
| 17 | 81461375 | 81461583 | 6.206813965 | RP11-1055B8.8 |
| 17 | 81541157 | 81541325 | 1.142872879 | FAAP100 |

| | | | | |
|---|---|---|---|---|
| 16 | 70740997 | 70741221 | -5.727920455 | VAC14 |
| 16 | 71284708 | 71284908 | -1.159737898 | CMTR2 |
| 1 | 63457537 | 63457781 | -3.40485217 | ITGB3BP |
| 16 | 72020688 | 72020836 | 7.047368753 | DHODH |
| 16 | 72024224 | 72024413 | -1.994813764 | DHODH |
| 16 | 72115381 | 72115567 | -1.308169084 | PMFBP1 |
| 16 | 72320471 | 72320622 | -2.506280281 | LINC01572 |
| 16 | 72786992 | 72787213 | 2.34335002 | ZFHX3 |
| 16 | 72786992 | 72787213 | 2.34335002 | RP5-991G20.1 |
| 16 | 72788489 | 72788608 | 2.876902856 | RP5-991G20.1 |
| 16 | 72788489 | 72788608 | 2.876902856 | ZFHX3 |
| 16 | 72795826 | 72795998 | 1.139248124 | ZFHX3 |
| 16 | 72798153 | 72798414 | 2.051775485 | ZFHX3 |
| 16 | 72958681 | 72958878 | 1.953549381 | ZFHX3 |
| 16 | 74325763 | 74325964 | -1.483561886 | AC009120.6 |
| 16 | 74661042 | 74661220 | -1.965770418 | RFWD3 |
| 16 | 75229896 | 75230097 | 1.24231696 | BCAR1 |
| 1 | 64777795 | 64777948 | -2.278518963 | RAVER2 |
| 16 | 75234994 | 75235206 | 1.063393755 | BCAR1 |
| 16 | 75264329 | 75264477 | -1.398895393 | BCAR1 |
| 16 | 75435421 | 75435639 | -1.168934819 | RP11-77K12.1 |
| 16 | 75435421 | 75435639 | -1.168934819 | RP11-77K12.9 |
| 16 | 75539926 | 75540045 | -1.903350245 | TMEM231 |
| 16 | 75539926 | 75540045 | -1.903350245 | RP11-77K12.7 |
| 16 | 75612329 | 75612599 | -1.061712347 | ADAT1 |
| 16 | 77198964 | 77199196 | -2.46351647 | MON1B |
| 16 | 77213489 | 77213687 | -2.327914416 | RP11-538I12.2 |
| 16 | 78119009 | 78119197 | -1.272534887 | WWOX |
| 16 | 81765056 | 81765255 | -3.02814169 | PLCG2 |
| 16 | 81765056 | 81765255 | -3.02814169 | RP11-960L18.1 |
| 16 | 83965878 | 83965986 | 1.545961245 | OSGIN1 |
| 16 | 84101814 | 84102014 | -1.343888478 | MBTPS1 |
| 16 | 84117551 | 84117705 | -1.489825677 | RP11-505K9.1 |
| 16 | 84179063 | 84179266 | -1.353094341 | TAF1C |
| 16 | 84745217 | 84745420 | -1.040145258 | USP10 |
| 16 | 85170329 | 85170589 | -2.441243376 | GSE1 |
| 16 | 85672493 | 85672692 | -1.780807806 | GSE1 |
| 16 | 85707669 | 85707861 | -4.420328449 | C16orf74 |
| 16 | 86538287 | 86538405 | -1.274431411 | MTHFSD |
| 16 | 87308875 | 87309073 | -2.79248125 | C16orf95 |
| 16 | 87308875 | 87309073 | -2.79248125 | RP11-178L8.4 |
| 16 | 87406598 | 87406764 | 1.258261533 | ZCCHC14 |
| 1 | 66372478 | 66372676 | 1.818468523 | PDE4B |
| 16 | 87410194 | 87410393 | 1.556571459 | ZCCHC14 |
| 16 | 87411551 | 87411795 | 1.450338941 | ZCCHC14 |
| 16 | 87430109 | 87430308 | 2.239703761 | ZCCHC14 |
| 16 | 87686795 | 87686985 | 2.671730628 | JPH3 |
| 16 | 87699824 | 87700025 | -1.609040334 | KLHDC4 |
| 16 | 87700784 | 87700972 | -2.060236291 | KLHDC4 |
| 16 | 87701723 | 87701912 | -2.970696983 | KLHDC4 |
| 16 | 87703171 | 87703377 | -1.610291218 | KLHDC4 |
| 1 | 1917820 | 1918013 | -1.94221247 | TMEM52 |
| 16 | 88048894 | 88049107 | -1.931442855 | BANP |
| 16 | 88432459 | 88432653 | -1.098777853 | ZNF469 |
| 1 | 92176968 | 92177169 | -2.286535022 | KIAA1107 |
| 1 | 92176968 | 92177169 | -2.286535022 | BTBD8 |
| 17 | 81707223 | 81707420 | 6.785935563 | RP13-1032I1.10 |
| 17 | 81707223 | 81707420 | 6.785935563 | MRPL12 |
| 17 | 81710218 | 81710423 | -7.238373102 | RP13-1032I1.10 |
| 17 | 81710693 | 81710892 | -6.608372929 | RP13-1032I1.10 |
| 1 | 92180532 | 92180714 | -1.632277311 | BTBD8 |
| 1 | 92180532 | 92180714 | -1.632277311 | KIAA1107 |
| 17 | 81901179 | 81901379 | -1.567082414 | NPB |
| 17 | 81901179 | 81901379 | -1.567082414 | PCYT2 |
| 17 | 81904551 | 81904778 | -1.352587414 | PCYT2 |
| 17 | 82056074 | 82056269 | 1.76308144 | GPS1 |
| 17 | 82108137 | 82108337 | 1.788162609 | CCDC57 |
| 1 | 92182136 | 92182335 | -2.845543176 | KIAA1107 |
| 1 | 92182136 | 92182335 | -2.845543176 | BTBD8 |
| 1 | 92184057 | 92184182 | -2.244746912 | BTBD8 |
| 1 | 92184057 | 92184182 | -2.244746912 | KIAA1107 |
| 17 | 82246665 | 82246864 | -1.538488949 | SLC16A3 |
| 17 | 82246665 | 82246864 | -1.538488949 | CSNK1D |
| 17 | 82247553 | 82247741 | -1.655811003 | SLC16A3 |
| 17 | 82247553 | 82247741 | -1.655811003 | CSNK1D |
| 17 | 82249820 | 82249996 | -1.335266523 | RP13-516M14.8 |
| 17 | 82249820 | 82249996 | -1.335266523 | CSNK1D |
| 17 | 82249820 | 82249996 | -1.335266523 | SLC16A3 |
| 17 | 82250449 | 82250606 | -7.914861368 | SLC16A3 |
| 17 | 82250449 | 82250606 | -7.914861368 | CSNK1D |
| 17 | 82250449 | 82250606 | -7.914861368 | RP13-516M14.8 |
| 17 | 82253317 | 82253473 | -1.696102963 | SLC16A3 |
| 17 | 82253317 | 82253473 | -1.696102963 | CSNK1D |
| 17 | 82255711 | 82255911 | -6.586838568 | CSNK1D |
| 17 | 82255711 | 82255911 | -6.586838568 | SLC16A3 |
| 17 | 82293875 | 82294074 | -1.625517726 | RP13-516M14.1 |
| 1 | 92387491 | 92387690 | -1.390500115 | RPAP2 |
| 17 | 82321835 | 82322007 | 7.633524266 | SECTM1 |
| 17 | 82420199 | 82420346 | 1.709524256 | HEXDC |
| 17 | 82443408 | 82443606 | 1.035501168 | C17orf62 |
| 17 | 82601737 | 82601937 | -1.016022724 | FOXK2 |
| 1 | 92388479 | 92388630 | -2.601936167 | RPAP2 |
| 17 | 82602997 | 82603277 | -1.919573769 | RP13-638C3.4 |
| 17 | 82602997 | 82603277 | -1.919573769 | FOXK2 |
| 17 | 82656314 | 82656506 | -1.663566627 | RAB40B |
| 17 | 83012277 | 83012472 | 1.335404234 | B3GNTL1 |
| 1 | 92390499 | 92390707 | -1.861769121 | RPAP2 |
| 17 | 83105200 | 83105430 | 1.009421895 | AC144831.1 |
| 18 | 659603 | 659801 | -1.087975538 | TYMS |
| 1 | 92391401 | 92391599 | -4.211603357 | RPAP2 |
| 18 | 662841 | 662998 | -1.478186534 | TYMS |
| 18 | 664661 | 664770 | -5.477098155 | TYMS |
| 18 | 670603 | 670810 | -2.508497084 | TYMS |
| 18 | 670603 | 670810 | -2.508497084 | ENOSF1 |
| 18 | 2652888 | 2653051 | -1.420690784 | CBX3P2 |
| 18 | 2802560 | 2802755 | -1.273122697 | SMCHD1 |
| 18 | 2802560 | 2802755 | -1.273122697 | RP11-703M24.5 |
| 18 | 2892090 | 2892296 | 1.162167591 | EMILIN2 |

| | | | | | | | | | |
|---|---|---|---|---|---|---|---|---|---|
| 16 | 88434581 | 88434771 | -1.004176858 | ZNF469 | 1 | 92393256 | 92393493 | -1.811363762 | RPAP2 |
| 16 | 88435531 | 88435722 | -1.109010797 | ZNF469 | 18 | 3457627 | 3457891 | -2.157038332 | TGIF1 |
| 16 | 88438847 | 88439001 | -1.586532354 | ZNF469 | 18 | 3596614 | 3596812 | -3.115967564 | DLGAP1 |
| 16 | 88570462 | 88570662 | -1.151342098 | ZC3H18 | 18 | 3596614 | 3596812 | -3.115967564 | DLGAP1-AS1 |
| 16 | 88577141 | 88577341 | -1.373992528 | ZC3H18 | 18 | 3602840 | 3603103 | -3.512216397 | DLGAP1 |
| 16 | 88631234 | 88631434 | -1.564180386 | ZC3H18 | 18 | 3602840 | 3603103 | -3.512216397 | DLGAP1-AS2 |
| 16 | 88649270 | 88649475 | -10.01460451 | CYBA | 18 | 5290877 | 5291079 | -2.993777164 | ZBTB14 |
| 16 | 88677929 | 88678134 | 1.270429479 | SNAI3 | 1 | 92475667 | 92475888 | -2.006265077 | GFI1 |
| 16 | 88677929 | 88678134 | 1.270429479 | SNAI3-AS1 | 18 | 8762411 | 8762612 | -2.338511527 | MTCL1 |
| 16 | 88686759 | 88686937 | 3.048610904 | SNAI3-AS1 | 18 | 8784173 | 8784369 | 6.471257253 | MTCL1 |
| 16 | 88686759 | 88686937 | 3.048610904 | AC138028.1 | 18 | 8825434 | 8825743 | 1.387879905 | MTCL1 |
| 16 | 88697197 | 88697325 | -7.117969712 | RP5-1142A6.5 | 18 | 9136328 | 9136548 | 1.687096838 | RP11-21J18.1 |
| 16 | 88697197 | 88697325 | -7.117969712 | RNF166 | 18 | 9136328 | 9136548 | 1.687096838 | ANKRD12 |
| 16 | 88751696 | 88751895 | -7.950622016 | PIEZO1 | 18 | 9136328 | 9136548 | 1.687096838 | NDUFV2-AS1 |
| 16 | 88764212 | 88764378 | -7.372206705 | PIEZO1 | 1 | 92484918 | 92485135 | -1.281329397 | GFI1 |
| 16 | 88829087 | 88829294 | -7.31365294 | GALNS | 18 | 9516906 | 9517103 | -1.498795002 | RALBP1 |
| 16 | 88876376 | 88876598 | -1.816302095 | CBFA2T3 | 18 | 9522030 | 9522258 | -1.658164599 | RALBP1 |
| 16 | 88876833 | 88876965 | -2.836212671 | CBFA2T3 | 18 | 9524470 | 9524666 | -1.797804872 | RALBP1 |
| 16 | 88890078 | 88890296 | -2.756295815 | CBFA2T3 | 18 | 9795327 | 9795527 | 8.715512079 | RAB31 |
| 16 | 88970328 | 88970527 | -7.336764962 | CBFA2T3 | 1 | 92513691 | 92513890 | -1.841377368 | EVI5 |
| 16 | 89119710 | 89119890 | -1.194044905 | ACSF3 | 18 | 11884382 | 11884584 | -1.265843322 | GNAL |
| 16 | 89160496 | 89160694 | 3.055610304 | LINC00304 | 18 | 11884382 | 11884584 | -1.265843322 | MPPE1 |
| 16 | 89200757 | 89200964 | -3.102010386 | SLC22A31 | 18 | 12325037 | 12325236 | 1.159533229 | TUBB6 |
| 16 | 89215640 | 89215763 | -2.359215645 | RP11-46C24.7 | 18 | 12449233 | 12449561 | -1.69993698 | SPIRE1 |
| 16 | 89498822 | 89499006 | -2.474508036 | SPG7 | 1 | 93136713 | 93136938 | -1.090244185 | MTF2 |
| 16 | 89498822 | 89499006 | -2.474508036 | RP11-104N10.1 | 18 | 12628082 | 12628278 | -1.346995954 | SPIRE1 |
| 16 | 89521100 | 89521299 | -1.250643741 | RP11-104N10.2 | 18 | 13049119 | 13049318 | -1.024543052 | CEP192 |
| 16 | 89521100 | 89521299 | -1.250643741 | SPG7 | 18 | 13056253 | 13056452 | -1.401005535 | CEP192 |
| 16 | 89536808 | 89537008 | -1.580854191 | SPG7 | 18 | 13479533 | 13479729 | -3.121145989 | LDLRAD4 |
| 16 | 89540827 | 89541016 | -1.435155208 | SPG7 | 18 | 13495621 | 13495794 | -1.6402345 | LDLRAD4 |
| 16 | 89557386 | 89557664 | -1.098064521 | SPG7 | 18 | 13498291 | 13498466 | -6.533801209 | LDLRAD4 |
| 16 | 89596029 | 89596148 | -1.366293577 | CPNE7 | 18 | 13499386 | 13499568 | -4.436741044 | LDLRAD4 |
| 16 | 89645673 | 89645873 | -1.206076144 | CHMP1A | 1 | 93166745 | 93166989 | -8.627864691 | TMED5 |
| 16 | 89660189 | 89660411 | -3.57220963 | SPATA33 | 18 | 13645310 | 13645511 | -3.739920369 | LDLRAD4 |
| 16 | 89670384 | 89670574 | -5.083403421 | SPATA33 | 18 | 13645310 | 13645511 | -3.739920369 | RP11-701H16.4 |
| 16 | 89696033 | 89696241 | -2.100086659 | CDK10 | 18 | 14074594 | 14074694 | -1.715922893 | ZNF519 |
| 16 | 89717431 | 89717631 | -2.547928075 | VPS9D1 | 18 | 21530322 | 21530513 | -1.267615616 | ESCO1 |
| 16 | 89717431 | 89717631 | -2.547928075 | VPS9D1-AS1 | 18 | 21573485 | 21573689 | -1.512948977 | ESCO1 |
| 16 | 89719129 | 89719329 | -1.662368039 | VPS9D1 | 18 | 22993195 | 22993393 | -1.71220187 | RBBP8 |
| 16 | 89730467 | 89730677 | -10.1748717 | ZNF276 | 18 | 23257416 | 23257651 | -1.462246112 | CABLES1 |
| 16 | 89737887 | 89738086 | -3.437836008 | ZNF276 | 18 | 23257416 | 23257651 | -1.462246112 | TMEM241 |
| 16 | 89737887 | 89738086 | -3.437836008 | FANCA | 18 | 23257416 | 23257651 | -1.462246112 | RP11-17J14.2 |
| 16 | 89740200 | 89740404 | -1.170768732 | FANCA | 18 | 24140794 | 24140967 | -2.678776002 | CABYR |
| 16 | 89740200 | 89740404 | -1.170768732 | ZNF276 | 18 | 25224628 | 25224823 | -1.42529004 | ZNF521 |
| 16 | 89871239 | 89871429 | -2.418048939 | SPIRE2 | 1 | 93219654 | 93219849 | -1.116935959 | CCDC18 |
| 16 | 89918990 | 89919270 | 2.123559754 | RP11-566K11.2 | 18 | 25226181 | 25226355 | -1.501208001 | ZNF521 |
| 16 | 89918990 | 89919270 | 2.123559754 | MC1R | 18 | 25226973 | 25227172 | -1.600871538 | ZNF521 |
| 16 | 89920306 | 89920514 | 2.16962809 | MC1R | 18 | 26389905 | 26390036 | -3 | TAF4B |
| 16 | 89920306 | 89920514 | 2.16962809 | RP11-566K11.4 | 18 | 32268037 | 32268242 | -1.215908194 | GAREM1 |
| 16 | 89920306 | 89920514 | 2.16962809 | RP11-566K11.2 | 18 | 32288025 | 32288228 | -1.456563566 | RP11-344B2.2 |
| 1 | 2142153 | 2142301 | -8.475050099 | RP5-892K4.1 | 18 | 32288025 | 32288228 | -1.456563566 | GAREM1 |
| 1 | 2142153 | 2142301 | -8.475050099 | PRKCZ | 18 | 34714142 | 34714320 | -6.514643613 | DTNA |
| 1 | 67395727 | 67395927 | -1.48713889 | IL12RB2 | 18 | 34725045 | 34725216 | -4.320122468 | DTNA |
| 16 | 89971299 | 89971529 | -1.197692779 | RP11-566K11.5 | 18 | 35242573 | 35242792 | -1.897217876 | ZNF397 |

| | | | | | | | | | |
|---|---|---|---|---|---|---|---|---|---|
| 16 | 89971299 | 89971529 | -1.197692779 | CENPBD1 | 18 | 35253858 | 35254061 | -2.018729794 | ZNF397 |
| 16 | 89971872 | 89972046 | -1.44718004 | RP11-566K11.5 | 18 | 35253858 | 35254061 | -2.018729794 | ZSCAN30 |
| 16 | 89971872 | 89972046 | -1.44718004 | CENPBD1 | 18 | 35260885 | 35261085 | -2.003547371 | ZSCAN30 |
| 16 | 89995480 | 89995679 | -1.543390333 | AFG3L1P | 18 | 35260885 | 35261085 | -2.003547371 | ZNF397 |
| 16 | 89995831 | 89996114 | -1.576193224 | AFG3L1P | 18 | 35262493 | 35262701 | -1.789437504 | ZSCAN30 |
| 16 | 90043169 | 90043369 | -1.77491899 | GAS8 | 18 | 35262493 | 35262701 | -1.789437504 | ZNF397 |
| 16 | 90043169 | 90043369 | -1.77491899 | URAHP | 18 | 35284120 | 35284326 | -1.849294461 | ZSCAN30 |
| 16 | 90167310 | 90167533 | -1.013622659 | RP11-356C4.5 | 18 | 35284120 | 35284326 | -1.849294461 | RP11-158H5.7 |
| 16 | 90167310 | 90167533 | -1.013622659 | FAM157C | 18 | 35306024 | 35306206 | -1.325112805 | ZNF271P |
| 17 | 147023 | 147222 | -2.300173126 | DOC2B | 18 | 35332479 | 35332684 | -1.134960396 | ZNF24 |
| 17 | 231094 | 231255 | -2.20312579 | RPH3AL | 1 | 2392780 | 2392989 | 1.004038257 | RER1 |
| 17 | 263678 | 263865 | -2.276908214 | RPH3AL | 1 | 93262285 | 93262481 | -1.745884151 | CCDC18 |
| 17 | 270773 | 270908 | -4.341051414 | RPH3AL | 1 | 93262285 | 93262481 | -1.745884151 | CCDC18-AS1 |
| 17 | 283503 | 283611 | -4.43518236 | RPH3AL | 18 | 35374124 | 35374324 | -7.856827504 | ZNF396 |
| 17 | 285934 | 286097 | -7.124037905 | RPH3AL | 18 | 35633241 | 35633424 | 2.862662839 | GALNT1 |
| 17 | 414024 | 414127 | -1.052124668 | C17orf97 | 18 | 35709625 | 35709909 | -1.083922163 | GALNT1 |
| 1 | 67413103 | 67413303 | -1.365657513 | SERBP1 | 18 | 36786586 | 36786769 | -2.652293841 | TPGS2 |
| 17 | 634680 | 634826 | 5.281121212 | VPS53 | 18 | 36937136 | 36937336 | -2.206279108 | KIAA1328 |
| 17 | 715664 | 715802 | -3.903677461 | VPS53 | 18 | 45063651 | 45063793 | 2.425954961 | SETBP1 |
| 17 | 740329 | 740508 | -8.144577177 | FAM57A | 18 | 45850218 | 45850386 | -1.059992725 | EPG5 |
| 17 | 741446 | 741723 | -2.454280246 | FAM57A | 18 | 46239889 | 46240080 | -7.731123317 | C18orf25 |
| 17 | 747187 | 747390 | -2.419932899 | GEMIN4 | 18 | 46253593 | 46253792 | -7.514665185 | C18orf25 |
| 17 | 753523 | 753667 | -2.811608212 | DBIL5P | 18 | 46483462 | 46483658 | -4.10715956 | LOXHD1 |
| 17 | 753523 | 753667 | -2.811608212 | GEMIN4 | 18 | 46560312 | 46560523 | -1.316962533 | LOXHD1 |
| 17 | 791910 | 792109 | -1.951480888 | MRM3 | 18 | 46561532 | 46561783 | -2.702897032 | LOXHD1 |
| 17 | 800795 | 800985 | 1.325552262 | NXN | 18 | 46679412 | 46679620 | -2.809932751 | ST8SIA5 |
| 17 | 815353 | 815457 | 2.521170384 | NXN | 18 | 46812226 | 46812573 | -1.308961904 | PIAS2 |
| 17 | 937905 | 938104 | -2.168189582 | NXN | 18 | 46946057 | 46946211 | -4.93518236 | KATNAL2 |
| 17 | 1001228 | 1001424 | -1.082327418 | TIMM22 | 18 | 47841713 | 47841908 | -2.794127564 | SMAD2 |
| 17 | 1005890 | 1006120 | 1.265362419 | ABR | 18 | 48027519 | 48027717 | -1.665379831 | ZBTB7C |
| 17 | 1042480 | 1042678 | -10.23484823 | ABR | 18 | 48040179 | 48040373 | -2.206212498 | ZBTB7C |
| 1 | 68161701 | 68161885 | -2.209156316 | CTBP2P8 | 18 | 48862846 | 48863044 | -6.788596383 | CTIF |
| 1 | 68161701 | 68161885 | -2.209156316 | WLS | 18 | 48949836 | 48950013 | 8.784412266 | SMAD7 |
| 1 | 68161701 | 68161885 | -2.209156316 | GNG12-AS1 | 18 | 48950320 | 48950527 | 1.946887648 | SMAD7 |
| 17 | 1148409 | 1148598 | 7.364491685 | ABR | 18 | 49590642 | 49590839 | -1.129705428 | LIPG |
| 17 | 1423051 | 1423231 | -2.848586002 | CRK | 1 | 93346517 | 93346716 | -1.156451563 | DR1 |
| 17 | 1423256 | 1423446 | -7.345762721 | CRK | 18 | 50984253 | 50984451 | -1.203688429 | ELAC1 |
| 17 | 1494976 | 1495155 | -1.705220985 | INPP5K | 18 | 50984253 | 50984451 | -1.203688429 | RP11-729L2.2 |
| 17 | 1517682 | 1517934 | -1.172345202 | PITPNA-AS1 | 18 | 51047007 | 51047137 | -1.221420802 | SMAD4 |
| 17 | 1517682 | 1517934 | -1.172345202 | PITPNA | 18 | 51047007 | 51047137 | -1.221420802 | RP11-729L2.2 |
| 17 | 1533850 | 1533994 | -1.715354467 | PITPNA | 18 | 51176723 | 51176922 | -1.355585975 | MEX3C |
| 17 | 1628171 | 1628347 | -2.169925001 | SLC43A2 | 18 | 51177200 | 51177379 | -1.133694637 | MEX3C |
| 17 | 1712640 | 1712840 | 2.955365934 | MIR22HG | 18 | 54290536 | 54290726 | -6.780046979 | POLI |
| 17 | 1777273 | 1777466 | -4.657230613 | SERPINF1 | 18 | 55227798 | 55227945 | 1.497356084 | TCF4 |
| 17 | 1800170 | 1800425 | -1.847268901 | SMYD4 | 18 | 57439335 | 57439533 | 4.414861368 | ONECUT2 |
| 17 | 1858067 | 1858258 | -7.580802152 | AC016292.3 | 1 | 93514828 | 93514981 | 1.119773963 | FNBP1L |
| 17 | 1858067 | 1858258 | -7.580802152 | RPA1 | 18 | 57440198 | 57440379 | 6.111246526 | ONECUT2 |
| 17 | 2305297 | 2305471 | -5.910036283 | SRR | 18 | 57483020 | 57483211 | -1.270234095 | ONECUT2 |
| 17 | 2395045 | 2395239 | -6.379319819 | MNT | 18 | 57483965 | 57484151 | 1.564112855 | ONECUT2 |
| 1 | 75758576 | 75758802 | -3.221119195 | ACADM | 18 | 59051299 | 59051479 | 1.803136503 | OACYLP |
| 17 | 2712646 | 2712844 | -1.66368044 | CLUH | 18 | 59902700 | 59902902 | 1.592849212 | PMAIP1 |
| 17 | 2712646 | 2712844 | -1.66368044 | RP11-74E22.3 | 18 | 62051724 | 62051878 | -1.114585477 | PIGN |
| 17 | 3669367 | 3669583 | -2.379736106 | P2RX5-TAX1BP3 | 18 | 62523756 | 62523948 | 8.109319455 | ZCCHC2 |
| 17 | 3669367 | 3669583 | -2.379736106 | EMC6 | 18 | 62539549 | 62539747 | 1.245065625 | ZCCHC2 |

| | | | | | | | | | |
|---|---|---|---|---|---|---|---|---|---|
| 17 | 3677813 | 3677985 | -2.215390329 | P2RX5 | 18 | 62829903 | 62830122 | 7.061979855 | PHLPP1 |
| 17 | 3677813 | 3677985 | -2.215390329 | P2RX5-TAX1BP3 | 1 | 93582432 | 93582639 | 2.175424163 | BCAR3 |
| 17 | 3682291 | 3682479 | 2.698189724 | P2RX5 | 18 | 63902935 | 63903133 | -3.229715809 | SERPINB2 |
| 17 | 3682291 | 3682479 | 2.698189724 | P2RX5-TAX1BP3 | 18 | 63902935 | 63903133 | -3.229715809 | SERPINB10 |
| 17 | 3725792 | 3725976 | -2.710916715 | ITGAE | 18 | 68836971 | 68837148 | -1.369993716 | CCDC102B |
| 17 | 3725792 | 3725976 | -2.710916715 | GSG2 | 1 | 844057 | 844256 | -5.756863798 | LINC01128 |
| 17 | 3726069 | 3726277 | -3.185731703 | ITGAE | 1 | 2403094 | 2403302 | -1.182187651 | RER1 |
| 17 | 3726069 | 3726277 | -3.185731703 | GSG2 | 18 | 70105153 | 70105266 | -4.25389732 | RTTN |
| 17 | 3861162 | 3861366 | 1.919627094 | CAMKK1 | 18 | 70490024 | 70490219 | 1.170878423 | RP11-644A7.2 |
| 17 | 3861832 | 3862008 | 2.217720563 | CAMKK1 | 18 | 74520576 | 74520776 | 1.719755764 | CNDP2 |
| 17 | 3913595 | 3913827 | -3.779790148 | P2RX1 | 18 | 74592969 | 74593216 | -2.020235681 | LINC00909 |
| 17 | 3915612 | 3915822 | -2.500969808 | P2RX1 | 1 | 95063445 | 95063645 | -1.271655555 | RP11-313A24.1 |
| 17 | 4006154 | 4006471 | -1.242297569 | ZZEF1 | 1 | 95063445 | 95063645 | -1.271655555 | ALG14 |
| 1 | 77107857 | 77108050 | -1.634493869 | PIGK | 18 | 74596926 | 74597167 | -1.175995109 | LINC00909 |
| 17 | 4510778 | 4510981 | 1.487531049 | SPNS2 | 18 | 74970360 | 74970539 | 1.049248882 | ZNF407 |
| 17 | 4537780 | 4538047 | 2.016669085 | SPNS2 | 18 | 75201570 | 75201770 | -1.988720358 | ZADH2 |
| 17 | 4539712 | 4539888 | -4.109621967 | MYBBP1A | 18 | 75211204 | 75211395 | -3.719071373 | TSHZ1 |
| 17 | 4672651 | 4672849 | -1.665536275 | PELP1 | 18 | 75285610 | 75285806 | -1.770140396 | TSHZ1 |
| 17 | 4731488 | 4731670 | -3.433528309 | MED11 | 18 | 75287359 | 75287529 | -2.263074102 | TSHZ1 |
| 17 | 4733149 | 4733412 | -1.015245864 | MED11 | 18 | 75288448 | 75288651 | -1.74492503 | TSHZ1 |
| 17 | 4789493 | 4789687 | 1.178923835 | GLTPD2 | 18 | 76362012 | 76362210 | -1.35485062 | ZNF516 |
| 17 | 5092470 | 5092687 | -1.334998324 | ZFP3 | 18 | 76371537 | 76371724 | -3.691852146 | ZNF516 |
| 1 | 2190579 | 2190686 | -2.826158884 | FAAP20 | 18 | 76380046 | 76380247 | -1.474554818 | ZNF516 |
| 1 | 77402568 | 77402766 | -7.253401719 | AK5 | 18 | 76441722 | 76441919 | 7.539158811 | ZNF516 |
| 17 | 5180914 | 5181115 | -1.460839597 | ZNF594 | 18 | 77016456 | 77016656 | -1.572344669 | MBP |
| 17 | 5181864 | 5182045 | -1.249290785 | ZNF594 | 18 | 77035260 | 77035462 | -1.549232931 | MBP |
| 17 | 5185498 | 5185784 | -1.22276705 | ZNF594 | 18 | 77037453 | 77037649 | -3.649245902 | MBP |
| 17 | 5186758 | 5186960 | -2.160974414 | ZNF594 | 18 | 77133571 | 77133776 | -2.539550891 | MBP |
| 17 | 5210684 | 5210866 | 5.52219706 | SCIMP | 18 | 79115341 | 79115554 | -1.657843969 | ATP9B |
| 17 | 5210684 | 5210866 | 5.52219706 | RP11-333E1.1 | 18 | 79365961 | 79366062 | -1.405869629 | ATP9B |
| 17 | 5444125 | 5444345 | -1.512012362 | DHX33 | 18 | 79527541 | 79527753 | -1.540386027 | NFATC1 |
| 17 | 5444125 | 5444345 | -1.512012362 | C1QBP | 1 | 99306980 | 99307185 | 3.773825315 | PLPPR4 |
| 1 | 77408869 | 77409061 | -7.924152533 | AK5 | 1 | 99969510 | 99969695 | -2.871820024 | RP5-884G6.2 |
| 17 | 5486417 | 5486575 | -3.209222225 | MIS12 | 18 | 80045944 | 80046149 | -2.193310346 | RBFA |
| 17 | 5486417 | 5486575 | -3.209222225 | DERL2 | 18 | 80045944 | 80046149 | -2.193310346 | RP11-795F19.5 |
| 17 | 5489199 | 5489411 | -1.223816239 | MIS12 | 18 | 80137534 | 80137736 | -1.146333655 | ADNP2 |
| 17 | 6652305 | 6652510 | 1.713079007 | C17orf100 | 18 | 80159602 | 80159761 | -6.032371382 | PARD6G |
| 17 | 6773287 | 6773490 | 2.104463136 | XAF1 | 18 | 80159602 | 80159761 | -6.032371382 | PARD6G-AS1 |
| 17 | 6773287 | 6773490 | 2.104463136 | RP11-609D21.3 | 18 | 80162058 | 80162283 | -1.5263092 | PARD6G |
| 17 | 6774260 | 6774462 | 2.213938678 | XAF1 | 18 | 80162058 | 80162283 | -1.5263092 | AC139100.4 |
| 17 | 6774260 | 6774462 | 2.213938678 | RP11-609D21.3 | 18 | 80162058 | 80162283 | -1.5263092 | PARD6G-AS1 |
| 17 | 6775126 | 6775329 | 2.306943916 | RP11-609D21.3 | 19 | 306562 | 306767 | -1.072918273 | MIER2 |
| 17 | 6775126 | 6775329 | 2.306943916 | XAF1 | 19 | 343850 | 344064 | -1.675914346 | MIER2 |
| 17 | 6782462 | 6782653 | 1.840500116 | FBXO39 | 19 | 509770 | 509929 | -6.014643613 | TPGS1 |
| 1 | 77412809 | 77412923 | -7.189959909 | AK5 | 19 | 643040 | 643194 | 3.453445298 | FGF22 |
| 17 | 7243913 | 7244119 | -1.282169235 | CTDNEP1 | 19 | 867449 | 867666 | -1.118326374 | MED16 |
| 17 | 7243913 | 7244119 | -1.282169235 | CTD-2545G14.7 | 19 | 958413 | 958613 | 3.761780978 | ARID3A |
| 17 | 7327697 | 7327880 | -1.015972518 | NEURL4 | 19 | 1010503 | 1010672 | 8.91909107 | TMEM259 |
| 17 | 7383622 | 7383790 | 6.058171981 | TNK1 | 19 | 1010503 | 1010672 | 8.91909107 | LLNLR-284B4.1 |
| 17 | 7456828 | 7457005 | -2.23765671 | CHRNB1 | 19 | 1157894 | 1158191 | 2.782377814 | SBNO2 |
| 1 | 77631943 | 77632172 | -1.014839362 | ZZZ3 | 19 | 1269936 | 1270145 | 9.34531276 | CIRBP |
| 17 | 7466522 | 7466715 | -1.573565971 | ZBTB4 | 19 | 1269936 | 1270145 | 9.34531276 | CIRBP-AS1 |
| 17 | 7579858 | 7580009 | -10.15418674 | CD68 | 19 | 1360522 | 1360723 | -2.121973606 | MUM1 |
| 17 | 7579858 | 7580009 | -10.15418674 | RP11-186B7.4 | 19 | 1370758 | 1370917 | -4.687853905 | MUM1 |

| | | | | | | | | | |
|---|---|---|---|---|---|---|---|---|---|
| 17 | 7583546 | 7583788 | 2.734438589 | AC113189.5 | 19 | 1377160 | 1377323 | -5.685888322 | MUM1 |
| 17 | 7583546 | 7583788 | 2.734438589 | MPDU1 | 19 | 1377160 | 1377323 | -5.685888322 | AC004623.2 |
| 17 | 7849086 | 7849294 | 1.256036622 | KDM6B | 19 | 1377746 | 1377953 | -2.654330705 | MUM1 |
| 1 | 77889020 | 77889213 | 6.765203169 | NEXN-AS1 | 19 | 1392551 | 1392739 | -6.774471144 | NDUFS7 |
| 1 | 77889020 | 77889213 | 6.765203169 | NEXN | 19 | 1392551 | 1392739 | -6.774471144 | AC005329.7 |
| 17 | 8140923 | 8141131 | 2.809336681 | PER1 | 19 | 1556788 | 1556979 | 5.101061912 | MEX3D |
| 17 | 8174973 | 8175172 | -8.79655694 | TMEM107 | 19 | 1574462 | 1574658 | -1.446847214 | MBD3 |
| 17 | 8227768 | 8227979 | -1.047152728 | CTC1 | 19 | 1827512 | 1827734 | 1.628978184 | REXO1 |
| 17 | 8269262 | 8269427 | -3.195427492 | PFAS | 19 | 1854190 | 1854388 | -1.051470038 | KLF16 |
| 17 | 8481679 | 8481792 | 2.032842122 | NDEL1 | 19 | 1913142 | 1913322 | 6.733420604 | SCAMP4 |
| 17 | 8481679 | 8481792 | 2.032842122 | MYH10 | 19 | 1913142 | 1913322 | 6.733420604 | ADAT3 |
| 17 | 8623149 | 8623283 | -3.605377962 | MYH10 | 19 | 2071248 | 2071492 | -1.729566516 | MOB3A |
| 17 | 9235570 | 9235705 | -3.669925001 | NTN1 | 19 | 2073198 | 2073409 | -1.560676974 | MOB3A |
| 17 | 9239671 | 9239895 | -2.346341278 | NTN1 | 19 | 2078248 | 2078453 | -1.673225224 | MOB3A |
| 17 | 9243384 | 9243560 | -5.561478689 | NTN1 | 19 | 2192026 | 2192224 | 1.632013516 | DOT1L |
| 17 | 9728353 | 9728538 | -1.942245724 | USP43 | 19 | 2292811 | 2293011 | -1.337905946 | LLfos-48D6.2 |
| 17 | 10658691 | 10658791 | -1.518161996 | CTC-297N7.1 | 19 | 2294218 | 2294503 | -1.941861369 | LLfos-48D6.2 |
| 17 | 12992507 | 12992696 | -2.822921539 | ELAC2 | 19 | 2295086 | 2295287 | -2.281320842 | LLfos-48D6.2 |
| 17 | 13496572 | 13496785 | 1.722531403 | HS3ST3A1 | 19 | 2513474 | 2513583 | -1.222760066 | GNG7 |
| 17 | 13777032 | 13777186 | 1.373936249 | COX10-AS1 | 19 | 2716334 | 2716535 | -1.385146714 | DIRAS1 |
| 17 | 14030033 | 14030235 | -1.85228158 | COX10-AS1 | 19 | 2717258 | 2717462 | -1.341847456 | DIRAS1 |
| 17 | 14034105 | 14034302 | -2.54120602 | COX10-AS1 | 19 | 2727853 | 2728014 | -1.002442291 | AC006538.1 |
| 17 | 14216792 | 14217006 | -1.309492613 | AC005224.2 | 19 | 2727853 | 2728014 | -1.002442291 | AC006538.4 |
| 17 | 14345492 | 14345697 | 2.735270685 | HS3ST3B1 | 19 | 2728649 | 2728855 | -1.749472698 | AC006538.4 |
| 17 | 15437991 | 15438212 | 2.131244904 | TVP23C-CDRT4 | 19 | 2728649 | 2728855 | -1.749472698 | AC006538.1 |
| 17 | 15437991 | 15438212 | 2.131244904 | CDRT4 | 19 | 2734568 | 2734820 | -1.384496608 | SLC39A3 |
| 17 | 15651452 | 15651651 | -1.447461986 | RP11-385D13.1 | 19 | 2734568 | 2734820 | -1.384496608 | AC006538.4 |
| 17 | 15651452 | 15651651 | -1.447461986 | TRIM16 | 19 | 2736838 | 2737036 | -1.746440146 | AC006538.4 |
| 17 | 15651452 | 15651651 | -1.447461986 | RP11-640I15.1 | 19 | 2736838 | 2737036 | -1.746440146 | SLC39A3 |
| 17 | 15715956 | 15716191 | -1.162776481 | ZNF286A | 19 | 2771876 | 2772073 | -5.013952998 | SGTA |
| 17 | 15715956 | 15716191 | -1.162776481 | TBC1D26 | 19 | 2834006 | 2834207 | -2.87318363 | ZNF554 |
| 17 | 15946098 | 15946298 | -1.37586318 | ADORA2B | 19 | 2852459 | 2852640 | -2.313455341 | ZNF555 |
| 1 | 2229430 | 2229658 | -1.054329079 | SKI | 19 | 2852459 | 2852640 | -2.313455341 | AC006130.3 |
| 17 | 16349030 | 16349230 | -3.110006786 | PIGL | 19 | 2853461 | 2853668 | -1.944858703 | AC006130.3 |
| 17 | 16349030 | 16349230 | -3.110006786 | CENPV | 19 | 2853461 | 2853668 | -1.944858703 | ZNF555 |
| 17 | 16443629 | 16443729 | 7.898273327 | LRRC75A-AS1 | 19 | 2859262 | 2859463 | -3.165470366 | ZNF555 |
| 17 | 16443629 | 16443729 | 7.898273327 | LRRC75A | 19 | 2877951 | 2878151 | -5.728946919 | ZNF556 |
| 17 | 16471688 | 16471882 | 1.927129319 | LRRC75A-AS1 | 19 | 2917181 | 2917375 | -2.30406916 | ZNF57 |
| 17 | 16471688 | 16471882 | 1.927129319 | LRRC75A | 19 | 2917181 | 2917375 | -2.30406916 | AC006277.2 |
| 17 | 16472159 | 16472311 | 1.974088539 | LRRC75A | 19 | 2934574 | 2934779 | -2.918911398 | ZNF77 |
| 17 | 16472159 | 16472311 | 1.974088539 | LRRC75A-AS1 | 19 | 3118709 | 3118906 | 6.299142587 | GNA11 |
| 1 | 81802439 | 81802632 | -1.854600033 | ADGRL2 | 19 | 3118709 | 3118906 | 6.299142587 | AC005262.3 |
| 17 | 17326361 | 17326512 | -1.770882971 | NT5M | 19 | 3122529 | 3122707 | -1.27640511 | GNA11 |
| 17 | 17491581 | 17491809 | -1.804866291 | MED9 | 19 | 3462797 | 3462998 | -1.663877173 | NFIC |
| 17 | 17494755 | 17494947 | 8.350043727 | RASD1 | 19 | 3509075 | 3509248 | 1.340026475 | FZR1 |
| 17 | 17793315 | 17793518 | -1.051959114 | RAI1 | 19 | 3531354 | 3531528 | 7.147023157 | FZR1 |
| 17 | 17794942 | 17795210 | -1.009622344 | RAI1 | 19 | 3562778 | 3562913 | -2.903677461 | MFSD12 |
| 17 | 17797927 | 17798113 | -1.001763483 | RAI1 | 19 | 3565314 | 3565503 | -7.171801069 | MFSD12 |
| 17 | 17955525 | 17955692 | -2.684310665 | TOM1L2 | 19 | 3959021 | 3959263 | 7.756709595 | DAPK3 |
| 17 | 18007601 | 18007752 | -2.484409172 | DRC3 | 19 | 4033014 | 4033118 | -5.245926548 | PIAS4 |
| 17 | 18007601 | 18007752 | -2.484409172 | ATPAF2 | 19 | 4409060 | 4409263 | -1.221605887 | CHAF1A |
| 17 | 18183477 | 18183672 | -7.232655654 | RP11-258F1.1 | 1 | 107057686 | 107057886 | -1.048526496 | PRMT6 |
| 17 | 18183477 | 18183672 | -7.232655654 | ALKBH5 | 19 | 4538066 | 4538267 | 1.671887038 | CTB-50L17.14 |
| 17 | 18274863 | 18275057 | -2.577645076 | TOP3A | 19 | 4538066 | 4538267 | 1.671887038 | LRG1 |

| | | | | |
|---|---|---|---|---|
| 1 | 84078809 | 84079007 | -2.255805379 | PRKACB |
| 17 | 18580178 | 18580370 | -1.072975159 | CCDC144B |
| 17 | 18661871 | 18662028 | -2.108577582 | ZNF286B |
| 17 | 18661871 | 18662028 | -2.108577582 | TBC1D28 |
| 17 | 18977958 | 18978176 | 1.301315211 | SLC5A10 |
| 17 | 18977958 | 18978176 | 1.301315211 | FAM83G |
| 17 | 19021424 | 19021624 | -1.379111607 | SLC5A10 |
| 17 | 19021424 | 19021624 | -1.379111607 | GRAP |
| 1 | 84926483 | 84926716 | -3.02559166 | MCOLN2 |
| 17 | 20204518 | 20204752 | -1.622887648 | AC004702.2 |
| 17 | 20204518 | 20204752 | -1.622887648 | SPECC1 |
| 17 | 20205479 | 20205681 | -2.109033635 | SPECC1 |
| 17 | 20205479 | 20205681 | -2.109033635 | AC004702.2 |
| 17 | 20238337 | 20238601 | -2.256665712 | SPECC1 |
| 17 | 20238337 | 20238601 | -2.256665712 | AC004702.2 |
| 17 | 20314169 | 20314308 | -2.591294694 | AC004702.2 |
| 17 | 20314169 | 20314308 | -2.591294694 | SPECC1 |
| 1 | 85100668 | 85100866 | -4.052640725 | WDR63 |
| 17 | 20895854 | 20896192 | -1.278183606 | CCDC144NL |
| 17 | 20895854 | 20896192 | -1.278183606 | CCDC144NL-AS1 |
| 17 | 20998404 | 20998601 | 1.095956275 | CCDC144NL-AS1 |
| 17 | 21147436 | 21147626 | -4.747207805 | DHRS7B |
| 17 | 21191117 | 21191294 | -1.36053665 | DHRS7B |
| 17 | 21198318 | 21198527 | -1.30288065 | TMEM11 |
| 17 | 21534831 | 21535045 | -1.591765086 | C17orf51 |
| 17 | 21534831 | 21535045 | -1.591765086 | RP11-822E23.2 |
| 17 | 21534831 | 21535045 | -1.591765086 | RP11-822E23.8 |
| 17 | 28326607 | 28326808 | -1.845087279 | TMEM97 |
| 17 | 28592803 | 28593000 | -2.789698387 | RP11-192H23.4 |
| 17 | 28592803 | 28593000 | -2.789698387 | SPAG5 |
| 1 | 85182194 | 85182393 | -1.193737064 | SYDE2 |
| 17 | 28634886 | 28635089 | -1.558002678 | RP11-192H23.7 |
| 17 | 28634886 | 28635089 | -1.558002678 | KIAA0100 |
| 17 | 28658713 | 28658825 | -1.148597267 | SDF2 |
| 17 | 28724448 | 28724585 | -4.035905101 | TLCD1 |
| 17 | 28860449 | 28860669 | -1.632551572 | ERAL1 |
| 17 | 28906047 | 28906247 | -1.242231029 | PHF12 |
| 17 | 28912941 | 28913141 | -1.162007128 | PHF12 |
| 17 | 29077718 | 29077901 | -8.879888839 | MYO18A |
| 17 | 29077718 | 29077901 | -8.879888839 | TIAF1 |
| 17 | 29078415 | 29078605 | -4.890497675 | MYO18A |
| 17 | 29078415 | 29078605 | -4.890497675 | TIAF1 |
| 17 | 29166098 | 29166279 | 1.714700081 | MYO18A |
| 17 | 29166558 | 29166750 | 1.22638136 | MYO18A |
| 17 | 29560837 | 29561015 | -3.047285552 | ABHD15 |
| 17 | 29560837 | 29561015 | -3.047285552 | ABHD15-AS1 |
| 17 | 29561632 | 29561827 | -2.054450354 | ABHD15-AS1 |
| 17 | 29561632 | 29561827 | -2.054450354 | ABHD15 |
| 17 | 29562501 | 29562751 | -1.93438159 | ABHD15 |
| 17 | 29562501 | 29562751 | -1.93438159 | ABHD15-AS1 |
| 17 | 29567811 | 29568006 | -2.11381085 | TP53I13 |
| 17 | 29567811 | 29568006 | -2.11381085 | ABHD15-AS1 |
| 17 | 29742589 | 29742727 | 6.994076049 | SSH2 |
| 17 | 30834639 | 30834929 | -3.571764218 | ATAD5 |
| 19 | 4652569 | 4652767 | -2.911225212 | MYDGF |
| 19 | 4652569 | 4652767 | -2.911225212 | TNFAIP8L1 |
| 19 | 4795035 | 4795235 | 1.454979704 | FEM1A |
| 19 | 4795035 | 4795235 | 1.454979704 | AC005523.2 |
| 19 | 4816060 | 4816256 | 1.649569438 | TICAM1 |
| 19 | 4816717 | 4816895 | 1.969829848 | TICAM1 |
| 19 | 4818113 | 4818240 | 1.998575533 | TICAM1 |
| 19 | 4961423 | 4961624 | -3.605960735 | UHRF1 |
| 19 | 5111630 | 5111823 | 6.178638537 | KDM4B |
| 19 | 5112188 | 5112355 | 6.143567593 | KDM4B |
| 19 | 5122998 | 5123165 | 7.237423336 | KDM4B |
| 19 | 5648446 | 5648643 | -1.028555296 | SAFB |
| 19 | 5711580 | 5711751 | 1.126037869 | LONP1 |
| 19 | 5909036 | 5909234 | -1.849065279 | AC104532.2 |
| 19 | 5909036 | 5909234 | -1.849065279 | VMAC |
| 19 | 5991198 | 5991326 | -6.88517709 | CTC-232P5.1 |
| 19 | 5994084 | 5994284 | -2.611942383 | RFX2 |
| 19 | 5994084 | 5994284 | -2.611942383 | CTC-232P5.1 |
| 19 | 6057862 | 6058054 | -6.648314737 | RFX2 |
| 19 | 6189461 | 6189564 | -1.512997604 | ACSBG2 |
| 19 | 6189461 | 6189564 | -1.512997604 | RFX2 |
| 1 | 107669185 | 107669372 | -5.486130924 | VAV3 |
| 19 | 6441695 | 6441896 | -1.222092927 | SLC25A23 |
| 19 | 6534936 | 6535134 | 1.50243502 | TNFSF9 |
| 19 | 6583430 | 6583624 | 2.321059679 | CD70 |
| 19 | 6593750 | 6593947 | -1.265532593 | RPL7P50 |
| 19 | 6593750 | 6593947 | -1.265532593 | CD70 |
| 19 | 6664459 | 6664661 | -1.71577189 | TNFSF14 |
| 19 | 6994768 | 6994925 | -6.204562855 | ADGRE4P |
| 19 | 7115303 | 7115505 | -1.753323417 | INSR |
| 19 | 7116027 | 7116227 | -2.110170841 | INSR |
| 19 | 7117123 | 7117314 | -1.725492098 | INSR |
| 19 | 7744260 | 7744455 | -3.155937215 | CD209 |
| 19 | 7863435 | 7863595 | 5.146160816 | EVI5L |
| 19 | 7982684 | 7982868 | 5.497176718 | ELAVL1 |
| 19 | 8498799 | 8498949 | -2.683313951 | PRAM1 |
| 19 | 8542800 | 8542968 | -1.182399872 | MYO1F |
| 19 | 8810260 | 8810437 | -4.853679566 | ZNF558 |
| 19 | 9341672 | 9341866 | -1.831737166 | ZNF559 |
| 19 | 9341672 | 9341866 | -1.831737166 | ZNF559-ZNF177 |
| 19 | 9381053 | 9381253 | -1.502364522 | ZNF559-ZNF177 |
| 19 | 9381053 | 9381253 | -1.502364522 | ZNF177 |
| 19 | 9414208 | 9414409 | -1.332969451 | ZNF266 |
| 19 | 9527598 | 9527799 | -1.268035347 | ZNF426 |
| 19 | 9529366 | 9529567 | -1.661088248 | ZNF426 |
| 19 | 9610569 | 9610775 | -1.446279375 | ZNF561 |
| 19 | 9653703 | 9653899 | -1.076707025 | ZNF562 |
| 19 | 9758285 | 9758450 | -6.728946919 | ZNF846 |
| 19 | 9857700 | 9857886 | -2.689689184 | OLFM2 |
| 19 | 10092136 | 10092383 | 2.462354714 | ANGPTL6 |
| 19 | 10092136 | 10092383 | 2.462354714 | C19orf66 |
| 19 | 10224141 | 10224338 | 8.089944829 | S1PR2 |
| 19 | 10224141 | 10224338 | 8.089944829 | DNMT1 |
| 19 | 10270577 | 10270771 | 3.597507991 | CTD-2369P2.5 |

| | | | | |
|---|---|---|---|---|
| 17 | 30834639 | 30834929 | -3.571764218 | CTD-2349P21.11 |
| 17 | 30864925 | 30865128 | 7.087462841 | ATAD5 |
| 17 | 30864925 | 30865128 | 7.087462841 | RP13-753N3.1 |
| 17 | 30893851 | 30894045 | -2.84243762 | ATAD5 |
| 17 | 30958012 | 30958378 | 2.191572871 | ADAP2 |
| 17 | 30970742 | 30970937 | 2.763381072 | RNF135 |
| 17 | 31318296 | 31318527 | 1.459226327 | EVI2A |
| 17 | 31318296 | 31318527 | 1.459226327 | CTD-2370N5.3 |
| 17 | 31318296 | 31318527 | 1.459226327 | NF1 |
| 17 | 31998822 | 31998954 | -1.483036914 | SUZ12 |
| 17 | 32021378 | 32021618 | -1.176159845 | LRRC37B |
| 17 | 32178109 | 32178297 | -1.749011507 | RHOT1 |
| 17 | 32489797 | 32489963 | -1.402668576 | CDK5R1 |
| 17 | 32490374 | 32490575 | -1.533892097 | CDK5R1 |
| 17 | 34962378 | 34962578 | -1.349491258 | ZNF830 |
| 17 | 34962378 | 34962578 | -1.349491258 | CCT6B |
| 17 | 35100765 | 35101027 | -2.006043582 | RAD51L3-RFFL |
| 17 | 35100765 | 35101027 | -2.006043582 | RAD51D |
| 17 | 35258752 | 35258955 | 5.427905982 | SLFN5 |
| 17 | 35259311 | 35259601 | 3.733904653 | SLFN5 |
| 17 | 35265401 | 35265612 | 5.733870089 | SLFN5 |
| 17 | 35352619 | 35352809 | -1.488971867 | SLFN11 |
| 17 | 35353805 | 35354011 | -1.036887316 | SLFN11 |
| 17 | 35362933 | 35363139 | -1.93368506 | SLFN11 |
| 17 | 35440789 | 35441008 | -1.075597199 | SLFN13 |
| 17 | 35441939 | 35442148 | -1.04362528 | SLFN13 |
| 17 | 35470912 | 35471054 | -1.309999417 | SLFN12L |
| 17 | 35480004 | 35480203 | -1.721082666 | SLFN12L |
| 17 | 35489224 | 35489406 | -2.352152904 | SLFN12L |
| 17 | 35498461 | 35498644 | 2.745926548 | RP11-1094M14.8 |
| 17 | 35498461 | 35498644 | 2.745926548 | SLFN12L |
| 17 | 35514782 | 35514952 | 4.169925001 | TOMM20P2 |
| 17 | 35514782 | 35514952 | 4.169925001 | SLFN12L |
| 17 | 35912872 | 35913046 | -4.421829693 | LRRC37A9P |
| 17 | 36496026 | 36496234 | -2.017245832 | MYO19 |
| 17 | 36496026 | 36496234 | -2.017245832 | ZNHIT3 |
| 17 | 36535138 | 36535335 | -2.069226716 | MYO19 |
| 17 | 36535138 | 36535335 | -2.069226716 | PIGW |
| 17 | 36537448 | 36537648 | -2.563121281 | PIGW |
| 17 | 36537448 | 36537648 | -2.563121281 | MYO19 |
| 17 | 36599153 | 36599346 | -2.582743007 | DHRS11 |
| 17 | 36602182 | 36602379 | -2.379100118 | MRM1 |
| 1 | 86703798 | 86704117 | 4.86919969 | RP4-612B15.3 |
| 17 | 36952879 | 36953082 | -1.236832785 | AATF |
| 17 | 36997811 | 36998021 | -2.020140452 | AATF |
| 17 | 37406349 | 37406554 | -1.473035811 | ACACA |
| 17 | 38322803 | 38323005 | -1.371316536 | MRPL45 |
| 17 | 38673067 | 38673387 | -1.645224569 | C17orf96 |
| 17 | 39175266 | 39175524 | -3.864220545 | RP5-906A24.2 |
| 17 | 39175266 | 39175524 | -3.864220545 | CACNB1 |
| 17 | 39409384 | 39409524 | -1.49093693 | MED1 |
| 1 | 87104551 | 87104750 | 3.327662965 | HS2ST1 |
| 1 | 87104551 | 87104750 | 3.327662965 | RP5-1052I5.2 |
| 17 | 39409914 | 39410095 | -1.027859014 | MED1 |
| 19 | 10285245 | 10285524 | 3.59032155 | ICAM1 |
| 19 | 10292081 | 10292254 | -5.951187557 | ICAM5 |
| 19 | 10294540 | 10294743 | 1.197148893 | ICAM5 |
| 19 | 10466990 | 10467190 | -1.760879498 | PDE4A |
| 19 | 10553479 | 10553679 | -1.181693776 | KRI1 |
| 19 | 10643605 | 10643796 | -2.73577865 | SLC44A2 |
| 1 | 108648692 | 108648885 | -3.021285692 | HENMT1 |
| 19 | 10653212 | 10653384 | -3.275696193 | ILF3-AS1 |
| 19 | 10869371 | 10869557 | 1.010596144 | C19orf38 |
| 19 | 11131434 | 11131634 | -1.140931938 | SPC24 |
| 19 | 11131434 | 11131634 | -1.140931938 | LDLR |
| 19 | 11146058 | 11146266 | -2.219383191 | SPC24 |
| 19 | 11193216 | 11193415 | -2.250499656 | KANK2 |
| 19 | 11193934 | 11194132 | -6.741970853 | KANK2 |
| 19 | 11195717 | 11195910 | -2.652471014 | KANK2 |
| 19 | 11325061 | 11325258 | -4.004411196 | TSPAN16 |
| 19 | 11325061 | 11325258 | -4.004411196 | RAB3D |
| 19 | 11375797 | 11375999 | -2.14367084 | SWSAP1 |
| 19 | 11617207 | 11617413 | -2.490537566 | ZNF627 |
| 19 | 11652686 | 11652887 | -2.493030405 | CTC-499B15.5 |
| 19 | 11652686 | 11652887 | -2.493030405 | ZNF887P |
| 1 | 108929691 | 108929889 | -3.491819022 | GPSM2 |
| 1 | 108929691 | 108929889 | -3.491819022 | CLCC1 |
| 1 | 108929691 | 108929889 | -3.491819022 | RP11-475E11.9 |
| 19 | 11680442 | 11680636 | -4.789705114 | CTC-499B15.5 |
| 19 | 11721618 | 11721818 | -2.911944144 | ZNF823 |
| 19 | 11722649 | 11722921 | -3.56901901 | ZNF823 |
| 19 | 11780618 | 11780816 | -1.813912185 | ZNF441 |
| 19 | 11781472 | 11781673 | -2.539513208 | ZNF441 |
| 19 | 11782877 | 11783084 | -1.987995253 | ZNF441 |
| 19 | 11832363 | 11832631 | -2.530233691 | ZNF440 |
| 19 | 11833568 | 11833763 | -2.383038711 | ZNF440 |
| 19 | 11867408 | 11867613 | -2.598135549 | ZNF439 |
| 19 | 11869028 | 11869228 | -2.384978488 | ZNF439 |
| 19 | 11908656 | 11908762 | -4.709980089 | ZNF69 |
| 19 | 11948438 | 11948619 | -1.718532427 | CTD-2006C1.12 |
| 19 | 11948438 | 11948619 | -1.718532427 | CTD-2006C1.13 |
| 19 | 11948438 | 11948619 | -1.718532427 | ZNF700 |
| 19 | 11950170 | 11950373 | -1.450328755 | ZNF700 |
| 19 | 11950170 | 11950373 | -1.450328755 | CTD-2006C1.12 |
| 19 | 11950170 | 11950373 | -1.450328755 | CTD-2006C1.13 |
| 19 | 11978145 | 11978344 | -4.855433959 | CTD-2006C1.13 |
| 19 | 11978145 | 11978344 | -4.855433959 | ZNF763 |
| 19 | 12005141 | 12005343 | -2.004604507 | CTD-2006C1.2 |
| 19 | 12016269 | 12016488 | -1.764297099 | ZNF433 |
| 19 | 12016269 | 12016488 | -1.764297099 | CTD-2006C1.2 |
| 19 | 12043769 | 12043888 | 3.336212671 | ZNF878 |
| 19 | 12043769 | 12043888 | 3.336212671 | CTD-2006C1.10 |
| 19 | 12043769 | 12043888 | 3.336212671 | CTD-2006C1.2 |
| 19 | 12111343 | 12111548 | -2.375320266 | ZNF20 |
| 19 | 12111343 | 12111548 | -2.375320266 | ZNF788 |
| 19 | 12113330 | 12113591 | -9.177034276 | ZNF788 |
| 19 | 12113330 | 12113591 | -9.177034276 | ZNF20 |
| 19 | 12132391 | 12132593 | -3.431525865 | ZNF788 |

| | | | | | | | | | |
|---|---|---|---|---|---|---|---|---|---|
| 17 | 39462366 | 39462567 | -1.157768427 | CDK12 | 19 | 12132391 | 12132593 | -3.431525865 | ZNF625-ZNF20 |
| 17 | 39471221 | 39471421 | -1.406029146 | CDK12 | 19 | 12132391 | 12132593 | -3.431525865 | ZNF20 |
| 17 | 39526031 | 39526231 | -1.789654413 | CDK12 | 19 | 12140322 | 12140495 | -4.604726683 | ZNF20 |
| 17 | 39531001 | 39531199 | -1.2728981 | CDK12 | 19 | 12140322 | 12140495 | -4.604726683 | ZNF625-ZNF20 |
| 17 | 39759516 | 39759773 | -1.519495852 | IKZF3 | 19 | 12186649 | 12186802 | -1.052356951 | ZNF136 |
| 17 | 39765789 | 39765981 | -1.539125723 | IKZF3 | 19 | 12247791 | 12247992 | -1.82564756 | ZNF44 |
| 1 | 87134146 | 87134348 | 3.543013468 | LINC01140 | 19 | 12272327 | 12272528 | -3.253489203 | ZNF44 |
| 1 | 87134146 | 87134348 | 3.543013468 | RP5-1052I5.2 | 19 | 12273009 | 12273125 | -2.318361964 | ZNF44 |
| 17 | 40163888 | 40164038 | -2.36223478 | CASC3 | 19 | 12273592 | 12273698 | -1.346492104 | ZNF44 |
| 17 | 40348159 | 40348347 | 7.137480236 | RARA | 1 | 109214608 | 109214803 | -1.121818192 | SARS |
| 17 | 40348159 | 40348347 | 7.137480236 | CTD-2267D19.1 | 19 | 12350635 | 12350836 | -2.310187378 | ZNF442 |
| 17 | 40356418 | 40356622 | -1.321474306 | RARA | 19 | 12380960 | 12381122 | -5.739890132 | CTD-3105H18.4 |
| 17 | 40362569 | 40362741 | -2.100610566 | CTD-2267D19.3 | 19 | 12380960 | 12381122 | -5.739890132 | CTD-3105H18.14 |
| 1 | 87153567 | 87153765 | 7.30158134 | RP5-1052I5.2 | 19 | 12390475 | 12390725 | -2.644807954 | ZNF799 |
| 1 | 87153567 | 87153765 | 7.30158134 | LINC01140 | 19 | 12390475 | 12390725 | -2.644807954 | CTD-3105H18.14 |
| 17 | 40363732 | 40363941 | -2.124122955 | GJD3 | 19 | 12430774 | 12430883 | -2.625537688 | ZNF443 |
| 17 | 40363732 | 40363941 | -2.124122955 | CTD-2267D19.3 | 19 | 12430774 | 12430883 | -2.625537688 | CTD-3105H18.16 |
| 17 | 40554494 | 40554684 | 8.242867494 | CCR7 | 19 | 12465262 | 12465464 | -4.800885394 | ZNF709 |
| 17 | 40555708 | 40555903 | 6.273688282 | CCR7 | 19 | 12465262 | 12465464 | -4.800885394 | CTD-3105H18.18 |
| 17 | 41802588 | 41802724 | -2.562347874 | P3H4 | 19 | 12465262 | 12465464 | -4.800885394 | CTD-2192J16.17 |
| 17 | 41810941 | 41811125 | -1.394115427 | P3H4 | 19 | 12527269 | 12527468 | -1.55278509 | CTD-2192J16.17 |
| 1 | 87328948 | 87329150 | 1.360541031 | LMO4 | 19 | 12527269 | 12527468 | -1.55278509 | CTD-2192J16.20 |
| 17 | 41968318 | 41968521 | -2.687344918 | CNP | 19 | 12527269 | 12527468 | -1.55278509 | ZNF564 |
| 17 | 42020481 | 42020733 | -8.124871543 | DNAJC7 | 1 | 109237828 | 109238027 | -2.675351873 | SARS |
| 17 | 42020481 | 42020733 | -8.124871543 | NKIRAS2 | 19 | 12580249 | 12580462 | -1.188610739 | CTD-2192J16.20 |
| 17 | 42113211 | 42113429 | -1.215539945 | KAT2A | 19 | 12580249 | 12580462 | -1.188610739 | ZNF490 |
| 17 | 42121532 | 42121716 | 1.397641045 | HSPB9 | 19 | 12581338 | 12581447 | -3.125614077 | ZNF490 |
| 17 | 42121532 | 42121716 | 1.397641045 | CTD-2132N18.3 | 19 | 12627911 | 12628033 | -1.53324875 | ZNF791 |
| 1 | 87344872 | 87345073 | 1.837826294 | LMO4 | 19 | 12627911 | 12628033 | -1.53324875 | ZNF490 |
| 17 | 42553832 | 42554031 | -1.200509327 | RP11-400F19.6 | 19 | 12628801 | 12629002 | -1.376775342 | ZNF490 |
| 17 | 42553832 | 42554031 | -1.200509327 | RP11-400F19.8 | 19 | 12628801 | 12629002 | -1.376775342 | ZNF791 |
| 17 | 42553832 | 42554031 | -1.200509327 | HSD17B1 | 19 | 12792715 | 12792923 | 1.896563772 | HOOK2 |
| 17 | 42668103 | 42668230 | -2.127153801 | PLEKHH3 | 19 | 12792715 | 12792923 | 1.896563772 | JUNB |
| 17 | 42797772 | 42798076 | -1.328819017 | COA3 | 19 | 12875722 | 12875938 | -1.113110699 | DNASE2 |
| 17 | 42991033 | 42991237 | -1.190601081 | RUNDC1 | 19 | 13099140 | 13099329 | -2.547821236 | LYL1 |
| 17 | 42993013 | 42993218 | -1.729217604 | RUNDC1 | 19 | 14052365 | 14052560 | -1.927018923 | IL27RA |
| 17 | 42993013 | 42993218 | -1.729217604 | AC055866.1 | 19 | 14149787 | 14149970 | 6.142701109 | ADGRL1 |
| 17 | 43094143 | 43094342 | -4.275921294 | BRCA1 | 19 | 14149787 | 14149970 | 6.142701109 | CTB-55O6.12 |
| 17 | 43128451 | 43128554 | -2.435517993 | BRCA1 | 19 | 14158114 | 14158326 | 1.048549388 | ADGRL1 |
| 17 | 43128451 | 43128554 | -2.435517993 | NBR2 | 19 | 14158114 | 14158326 | 1.048549388 | CTB-55O6.12 |
| 17 | 43136209 | 43136407 | -1.059656588 | BRCA1 | 19 | 14162996 | 14163235 | -1.129674193 | CTB-55O6.12 |
| 17 | 43136209 | 43136407 | -1.059656588 | NBR2 | 19 | 14162996 | 14163235 | -1.129674193 | ADGRL1 |
| 17 | 43162770 | 43162959 | -1.553414083 | BRCA1 | 1 | 109279889 | 109280088 | -4.414332714 | PSRC1 |
| 17 | 43162770 | 43162959 | -1.553414083 | RP11-242D8.1 | 19 | 14394244 | 14394436 | 4.32422324 | ADGRE5 |
| 17 | 43246576 | 43246774 | -2.014974661 | LINC00854 | 19 | 14395211 | 14395410 | 3.851581759 | ADGRE5 |
| 1 | 88833251 | 88833480 | -1.007887264 | PKN2 | 19 | 14408358 | 14408562 | 2.47016233 | ADGRE5 |
| 17 | 44023553 | 44023797 | -1.114165122 | TMEM101 | 19 | 14408358 | 14408562 | 2.47016233 | CTC-548K16.5 |
| 17 | 44148050 | 44148241 | -1.93558042 | C17orf53 | 19 | 14409184 | 14409287 | 9.070554637 | DDX39A |
| 17 | 44206904 | 44207104 | -1.473281493 | UBTF | 19 | 14411264 | 14411456 | 2.9476407 | DDX39A |
| 17 | 44206904 | 44207104 | -1.473281493 | CTB-175E5.7 | 19 | 14412645 | 14412843 | 9.538161202 | DDX39A |
| 1 | 88983570 | 88983766 | -1.086622785 | RBMXL1 | 19 | 14708898 | 14709064 | -8.489958186 | ZNF333 |
| 1 | 88983570 | 88983766 | -1.086622785 | KYAT3 | 1 | 3463377 | 3463528 | -3.213342041 | ARHGEF16 |
| 17 | 44666570 | 44666687 | -4.82844954 | MEIOC | 19 | 14735914 | 14736119 | -1.258332879 | ADGRE2 |
| 17 | 44667627 | 44667811 | -4.36310908 | MEIOC | 19 | 14767992 | 14768190 | -1.267072208 | ADGRE2 |

| | | | | |
|---|---|---|---|---|
| 17 | 44780102 | 44780253 | 1.5 | ADAM11 |
| 17 | 44926328 | 44926430 | -4.563764338 | KIF18B |
| 17 | 44946606 | 44946922 | -2.652719986 | KIF18B |
| 17 | 44947252 | 44947451 | -2.269041663 | KIF18B |
| 17 | 45105650 | 45105862 | -1.125760505 | NMT1 |
| 17 | 45107344 | 45107555 | -1.319159166 | NMT1 |
| 17 | 45108646 | 45108848 | -1.031203692 | NMT1 |
| 17 | 45169399 | 45169628 | -2.382930703 | RP13-890H12.2 |
| 17 | 45169399 | 45169628 | -2.382930703 | HEXIM2 |
| 17 | 45190846 | 45191047 | 6.218355771 | CTD-2020K17.1 |
| 17 | 45218905 | 45219155 | 4.325842434 | CTD-2020K17.1 |
| 17 | 45286755 | 45286955 | 1.220151078 | MAP3K14 |
| 17 | 45514612 | 45514813 | -1.661994352 | LRRC37A4P |
| 17 | 45515322 | 45515520 | -1.076856528 | LRRC37A4P |
| 1 | 89104414 | 89104552 | 5.869890305 | PTGES3P1 |
| 17 | 45518995 | 45519192 | -1.943117322 | LRRC37A4P |
| 17 | 45549805 | 45549943 | -2.005295977 | RP11-798G7.6 |
| 17 | 45549805 | 45549943 | -2.005295977 | RP11-798G7.7 |
| 17 | 45549805 | 45549943 | -2.005295977 | LRRC37A4P |
| 17 | 45550506 | 45550708 | -3.039032072 | RP11-798G7.6 |
| 17 | 45550506 | 45550708 | -3.039032072 | RP11-798G7.7 |
| 17 | 46309988 | 46310179 | -3.194379802 | LRRC37A |
| 17 | 46309988 | 46310179 | -3.194379802 | ARL17B |
| 17 | 46338222 | 46338427 | -1.22772639 | ARL17B |
| 17 | 46512220 | 46512390 | -5.669925001 | LRRC37A2 |
| 17 | 46548822 | 46549021 | -1.08332545 | AC138645.1 |
| 17 | 46548822 | 46549021 | -1.08332545 | LRRC37A2 |
| 17 | 46548822 | 46549021 | -1.08332545 | ARL17A |
| 17 | 46548822 | 46549021 | -1.08332545 | AC138645.2 |
| 17 | 46556082 | 46556286 | -2.211101683 | AC138645.1 |
| 17 | 46556082 | 46556286 | -2.211101683 | AC138645.2 |
| 17 | 46556082 | 46556286 | -2.211101683 | ARL17A |
| 17 | 46556634 | 46556836 | -8.141544177 | ARL17A |
| 17 | 46556634 | 46556836 | -8.141544177 | AC138645.1 |
| 17 | 46556634 | 46556836 | -8.141544177 | AC138645.2 |
| 17 | 46768470 | 46768673 | -2.533234715 | WNT3 |
| 17 | 46959940 | 46960146 | 1.668990288 | RP11-156P1.2 |
| 17 | 46959940 | 46960146 | 1.668990288 | GOSR2 |
| 17 | 46964384 | 46964588 | 1.009929235 | GOSR2 |
| 17 | 46964384 | 46964588 | 1.009929235 | RP11-156P1.2 |
| 17 | 46968608 | 46968771 | 2.311635929 | RP11-156P1.2 |
| 17 | 47009512 | 47009758 | -1.526717092 | LRRC37A17P |
| 17 | 47009512 | 47009758 | -1.526717092 | RP11-156P1.2 |
| 17 | 47009512 | 47009758 | -1.526717092 | RP11-156P1.3 |
| 17 | 47049860 | 47050045 | -2.368372785 | LRRC37A17P |
| 17 | 47049860 | 47050045 | -2.368372785 | RP11-156P1.3 |
| 17 | 47387475 | 47387640 | 1.3075765 | EFCAB13 |
| 17 | 47434891 | 47435066 | 2.793888758 | EFCAB13 |
| 17 | 47745134 | 47745287 | 2.068751762 | TBX21 |
| 17 | 47898913 | 47899075 | -6.993676423 | SP2-AS1 |
| 17 | 47898913 | 47899075 | -6.993676423 | SP2 |
| 17 | 48576475 | 48576651 | -1.602140007 | HOXB3 |
| 17 | 48576475 | 48576651 | -1.602140007 | HOXB-AS3 |
| 17 | 48576475 | 48576651 | -1.602140007 | HOXB4 |
| 19 | 15362202 | 15362400 | 7.837978516 | AKAP8 |
| 19 | 16157381 | 16157505 | 1.659192531 | HSH2D |
| 19 | 16236204 | 16236414 | 1.308081721 | AP1M1 |
| 19 | 16236751 | 16236881 | 2.023652857 | AP1M1 |
| 19 | 16327091 | 16327286 | 6.330332239 | KLF2 |
| 19 | 16579198 | 16579395 | -2.006245472 | CTC-429P9.4 |
| 19 | 16579198 | 16579395 | -2.006245472 | MED26 |
| 19 | 16579198 | 16579395 | -2.006245472 | CTD-3222D19.2 |
| 19 | 16630883 | 16631075 | -2.184075556 | CTC-429P9.1 |
| 19 | 16630883 | 16631075 | -2.184075556 | CTC-429P9.2 |
| 19 | 16630883 | 16631075 | -2.184075556 | CTC-429P9.4 |
| 19 | 16630883 | 16631075 | -2.184075556 | SMIM7 |
| 19 | 16879681 | 16879849 | 1.273390158 | SIN3B |
| 19 | 16890991 | 16891185 | -3.500585857 | F2RL3 |
| 19 | 16891614 | 16891857 | -4.496427713 | F2RL3 |
| 19 | 16893648 | 16893858 | -1.581687081 | CPAMD8 |
| 19 | 17231981 | 17232180 | -2.204157559 | NR2F6 |
| 19 | 17283435 | 17283635 | -6.555241655 | ANKLE1 |
| 19 | 17306400 | 17306592 | -2.861438524 | MRPL34 |
| 19 | 17306400 | 17306592 | -2.861438524 | ABHD8 |
| 1 | 109665204 | 109665328 | -4.569775676 | GSTM4 |
| 19 | 17341654 | 17341778 | -1.492873008 | GTPBP3 |
| 19 | 17486385 | 17486588 | -6.891601445 | SLC27A1 |
| 19 | 17504586 | 17504785 | -7.54290219 | SLC27A1 |
| 19 | 17504586 | 17504785 | -7.54290219 | CTD-3131K8.2 |
| 19 | 18012992 | 18013181 | -2.836040532 | ARRDC2 |
| 19 | 18013666 | 18013866 | -1.502128038 | ARRDC2 |
| 1 | 109896285 | 109896467 | -1.506036416 | RP11-195M16.3 |
| 19 | 18363468 | 18363669 | -2.513250138 | PGPEP1 |
| 19 | 18397035 | 18397154 | -7.959687734 | LRRC25 |
| 19 | 18450547 | 18450745 | 8.444989603 | ELL |
| 19 | 18777147 | 18777366 | -2.829163336 | CRTC1 |
| 19 | 18919993 | 18920164 | -7.719477125 | AC002985.3 |
| 19 | 18919993 | 18920164 | -7.719477125 | DDX49 |
| 19 | 19177022 | 19177222 | -1.876859245 | BORCS8 |
| 19 | 19177022 | 19177222 | -1.876859245 | BORCS8-MEF2B |
| 19 | 19257491 | 19257687 | -1.488639962 | HAPLN4 |
| 19 | 19257491 | 19257687 | -1.488639962 | AC138430.4 |
| 19 | 19508456 | 19508662 | -1.649663232 | GATAD2A |
| 19 | 19712872 | 19713071 | -1.062909581 | ZNF14 |
| 19 | 19724891 | 19725025 | -2.520974091 | ZNF14 |
| 19 | 19892429 | 19892606 | -3.699196372 | CTC-559E9.8 |
| 19 | 19892429 | 19892606 | -3.699196372 | ZNF253 |
| 19 | 19935060 | 19935231 | -5.936914287 | AC007204.2 |
| 19 | 19935060 | 19935231 | -5.936914287 | ZNF93 |
| 19 | 20006409 | 20006635 | -2.668934819 | ZNF682 |
| 19 | 20544399 | 20544614 | -1.903516732 | ZNF737 |
| 19 | 20545621 | 20545815 | -4.424337549 | ZNF737 |
| 19 | 20948982 | 20949097 | -2.506226694 | ZNF85 |
| 19 | 20950256 | 20950454 | -2.193740043 | ZNF85 |
| 1 | 3733412 | 3733612 | -1.413780225 | TP73 |
| 19 | 21057557 | 21057747 | -1.297854968 | ZNF430 |
| 19 | 21116872 | 21117023 | -1.611335034 | ZNF714 |
| 19 | 21118567 | 21118762 | -1.918510688 | ZNF714 |

| | | | | | | | | | |
|---|---|---|---|---|---|---|---|---|---|
| 17 | 48607758 | 48607957 | -1.584945545 | HOXB7 | 19 | 21182783 | 21182978 | -1.676950427 | ZNF431 |
| 17 | 49298116 | 49298318 | -1.381645535 | ZNF652 | 19 | 21183852 | 21184053 | -1.590944835 | ZNF431 |
| 17 | 49317008 | 49317382 | -1.506484991 | ZNF652 | 1 | 110112761 | 110112961 | -4.673792826 | UBL4B |
| 17 | 49404398 | 49404599 | -2.243210969 | RP11-1079K10.4 | 19 | 21293032 | 21293232 | -1.674394351 | ZNF708 |
| 17 | 49404398 | 49404599 | -2.243210969 | PHB | 19 | 21294402 | 21294603 | -1.386138695 | ZNF708 |
| 17 | 49404398 | 49404599 | -2.243210969 | RP11-81K2.1 | 19 | 21425819 | 21425923 | -4.285402219 | CTD-2561J22.3 |
| 17 | 49579233 | 49579425 | 7.062666372 | NXPH3 | 19 | 21425819 | 21425923 | -4.285402219 | ZNF493 |
| 17 | 49969264 | 49969406 | 1.952693503 | DLX4 | 19 | 21537534 | 21537647 | -5.894359166 | ZNF429 |
| 17 | 49974057 | 49974257 | 1.878256867 | DLX4 | 19 | 21726611 | 21726811 | -1.403896749 | ZNF100 |
| 17 | 49991273 | 49991475 | 7.824476181 | DLX3 | 19 | 21727635 | 21727834 | -1.679008494 | ZNF100 |
| 17 | 50089898 | 50090095 | 7.990576091 | ITGA3 | 19 | 21809593 | 21809819 | -2.403771136 | ZNF43 |
| 17 | 50367957 | 50368158 | -1.786854742 | MRPL27 | 19 | 23222307 | 23222511 | -1.060580641 | ZNF724 |
| 17 | 50381091 | 50381276 | -4.269874848 | LRRC59 | 19 | 23223238 | 23223434 | -2.959193117 | ZNF724 |
| 17 | 50381091 | 50381276 | -4.269874848 | EME1 | 19 | 23223238 | 23223434 | -2.959193117 | RP11-15H20.8 |
| 17 | 50382811 | 50383006 | -2.388496912 | LRRC59 | 1 | 110223499 | 110223686 | 1.050142486 | KCNC4 |
| 17 | 50755789 | 50756027 | -1.4963534 | LUC7L3 | 19 | 23223728 | 23223925 | -3.410693304 | RP11-15H20.8 |
| 17 | 50863736 | 50863938 | 1.042999046 | TOB1 | 19 | 23223728 | 23223925 | -3.410693304 | ZNF724 |
| 17 | 56832980 | 56833226 | -2.087933488 | C17orf67 | 19 | 23259635 | 23259794 | -5.233802775 | RP11-15H20.8 |
| 17 | 56950021 | 56950263 | -1.684008494 | COIL | 19 | 23259635 | 23259794 | -5.233802775 | RP11-15H20.7 |
| 17 | 56950663 | 56950838 | -1.950187812 | COIL | 19 | 23360829 | 23360963 | -2.089761323 | ZNF91 |
| 17 | 57106212 | 57106412 | -3.490977988 | AKAP1 | 19 | 23653279 | 23653493 | -1.312071706 | ZNF675 |
| 17 | 57106920 | 57107083 | -2.874486772 | AKAP1 | 19 | 24126327 | 24126525 | -1.790075735 | ZNF254 |
| 17 | 57109902 | 57110079 | -2.375220326 | AKAP1 | 19 | 24127823 | 24128131 | -1.030700272 | ZNF254 |
| 17 | 57632140 | 57632344 | -2.401351929 | MSI2 | 19 | 27696465 | 27696627 | -3.603225439 | LINC00662 |
| 17 | 58302067 | 58302265 | 6.851627367 | TSPOAP1 | 19 | 27699227 | 27699411 | -4.387187636 | LINC00662 |
| 17 | 58347372 | 58347474 | -8.786272403 | TSPOAP1-AS1 | 19 | 27820272 | 27820381 | -4.56592848 | CTC-459F4.3 |
| 17 | 58347372 | 58347474 | -8.786272403 | SUPT4H1 | 19 | 27820272 | 27820381 | -4.56592848 | LLNLF-65H9.1 |
| 17 | 58496035 | 58496237 | -1.087325547 | MTMR4 | 19 | 29702737 | 29702853 | 1.398449795 | C19orf12 |
| 17 | 59211007 | 59211211 | -1.323551168 | SMG8 | 19 | 31276686 | 31276886 | -1.237244068 | TSHZ3 |
| 17 | 59211007 | 59211211 | -1.323551168 | CTD-2510F5.6 | 19 | 32353318 | 32353512 | -2.536065611 | ZNF507 |
| 17 | 59213192 | 59213406 | -1.115102785 | SMG8 | 19 | 32353987 | 32354225 | -1.735975444 | ZNF507 |
| 17 | 59213192 | 59213406 | -1.115102785 | CTD-2510F5.6 | 1 | 110346210 | 110346410 | -2.022790517 | RBM15 |
| 17 | 59397052 | 59397228 | 1.401544016 | YPEL2 | 19 | 32382999 | 32383210 | -2.686192919 | ZNF507 |
| 17 | 59697446 | 59697647 | -1.152127244 | PTRH2 | 19 | 32691526 | 32691722 | -1.847135242 | CTD-2538C1.2 |
| 17 | 60043809 | 60043940 | -1.357685659 | HEATR6 | 19 | 32976860 | 32977015 | -5.562060656 | FAAP24 |
| 17 | 60447738 | 60447927 | 1.285493241 | APPBP2 | 19 | 33171655 | 33171833 | -1.645470201 | WDR88 |
| 17 | 60483719 | 60483910 | 1.744161096 | APPBP2 | 19 | 33172070 | 33172347 | -1.680826257 | WDR88 |
| 17 | 61408464 | 61408604 | 5.86735481 | TBX2-AS1 | 19 | 33174570 | 33174761 | -1.463650094 | WDR88 |
| 17 | 61408464 | 61408604 | 5.86735481 | TBX2 | 19 | 33175783 | 33176038 | -1.753569753 | WDR88 |
| 17 | 61483372 | 61483636 | -1.474662931 | TBX4 | 1 | 110347491 | 110347706 | -1.418604605 | LAMTOR5-AS1 |
| 17 | 61683406 | 61683785 | -3.651975219 | BRIP1 | 19 | 33205530 | 33205699 | 7.093676037 | LRP3 |
| 17 | 62477243 | 62477441 | -1.065030163 | TLK2 | 19 | 33207345 | 33207534 | 6.490516029 | LRP3 |
| 17 | 62626123 | 62626324 | -1.664320719 | CTD-3035K23.6 | 19 | 33207345 | 33207534 | 6.490516029 | CTD-2540B15.13 |
| 1 | 89713350 | 89713547 | -1.065481667 | RP11-302M6.4 | 19 | 33304624 | 33304827 | -3.095754279 | CEBPA-AS1 |
| 1 | 89713350 | 89713547 | -1.065481667 | LRRC8C | 19 | 33379539 | 33379738 | -1.905617198 | CEBPG |
| 17 | 63818031 | 63818249 | -1.126374758 | DDX42 | 19 | 33399325 | 33399465 | 6.077409055 | PEPD |
| 17 | 63819728 | 63819929 | -2.024695824 | FTSJ3 | 19 | 33812907 | 33813104 | -1.893732859 | KCTD15 |
| 17 | 64002549 | 64002786 | -2.822058738 | ICAM2 | 19 | 34341697 | 34341851 | -1.371409816 | KIAA0355 |
| 17 | 64002549 | 64002786 | -2.822058738 | PRR29 | 19 | 34684663 | 34684783 | -1.702195128 | ZNF302 |
| 17 | 64043606 | 64043809 | 1.879967652 | ERN1 | 1 | 110408006 | 110408158 | -5.796695561 | LAMTOR5-AS1 |
| 17 | 64129178 | 64129348 | 1.069604222 | ERN1 | 19 | 34733452 | 34733664 | -1.150799351 | CTD-2553L13.10 |
| 17 | 64148743 | 64148941 | 2.399240667 | TEX2 | 19 | 34741035 | 34741235 | -7.466776177 | ZNF181 |
| 17 | 64193655 | 64193875 | 1.279190668 | TEX2 | 19 | 34957878 | 34958050 | -6.552135564 | ZNF792 |
| 1 | 89714396 | 89714603 | -1.226840363 | LRRC8C | 19 | 34959290 | 34959501 | -2.523831333 | ZNF792 |

| | | | | | | | | | |
|---|---|---|---|---|---|---|---|---|---|
| 1 | 89714396 | 89714603 | -1.226840363 | RP11-302M6.4 | 19 | 35347078 | 35347277 | -1.35154996 | CD22 |
| 17 | 64369930 | 64370127 | -8.770623948 | PECAM1 | 19 | 35721483 | 35721683 | 8.535553085 | KMT2B |
| 17 | 64412160 | 64412334 | 7.094178516 | PECAM1 | 19 | 35937049 | 35937252 | -1.280859075 | LRFN3 |
| 17 | 64413707 | 64413838 | 3.429858738 | PECAM1 | 19 | 35939312 | 35939489 | -2.137101244 | LRFN3 |
| 17 | 64496516 | 64496713 | -1.347315515 | POLG2 | 19 | 35944957 | 35945172 | -1.249107646 | LRFN3 |
| 17 | 64778323 | 64778521 | -6.39659843 | RP13-104F24.2 | 19 | 35995701 | 35995959 | -1.034365982 | SDHAF1 |
| 17 | 64778995 | 64779201 | 1.370686887 | RP13-104F24.2 | 19 | 36337080 | 36337236 | -1.026581534 | ZFP14 |
| 17 | 64897420 | 64897609 | 1.220286296 | RP11-927P21.1 | 19 | 36449737 | 36449851 | -3.278888836 | ZNF566 |
| 17 | 64897420 | 64897609 | 1.220286296 | LRRC37A3 | 19 | 36496919 | 36497026 | -1.281794425 | CTBP2P7 |
| 17 | 64956406 | 64956528 | 3.736618709 | SLC16A6P1 | 1 | 3767168 | 3767377 | -2.596479064 | CCDC27 |
| 17 | 67912804 | 67912984 | -1.63412935 | BPTF | 1 | 110604463 | 110604660 | -3.752161224 | KCNA2 |
| 17 | 67945415 | 67945630 | -1.024564794 | BPTF | 19 | 36626069 | 36626293 | -1.379169051 | ZNF382 |
| 17 | 68135439 | 68135551 | -1.344537531 | SH3GL1P3 | 19 | 36626913 | 36627352 | -2.114487219 | ZNF382 |
| 17 | 68135439 | 68135551 | -1.344537531 | RP11-147L13.13 | 19 | 36638832 | 36638977 | -6.889654487 | ZNF461 |
| 17 | 68135439 | 68135551 | -1.344537531 | LRRC37A16P | 19 | 36719167 | 36719377 | -1.369118263 | ZNF567 |
| 17 | 68269106 | 68269313 | 5.047892417 | SLC16A6 | 19 | 36719167 | 36719377 | -1.369118263 | ZNF850 |
| 17 | 68269106 | 68269313 | 5.047892417 | ARSG | 19 | 36720101 | 36720396 | -1.488521151 | ZNF567 |
| 1 | 89820827 | 89820984 | -5.404884064 | RP5-943J3.2 | 19 | 36720101 | 36720396 | -1.488521151 | ZNF850 |
| 1 | 89820827 | 89820984 | -5.404884064 | RP11-302M6.4 | 19 | 36747983 | 36748182 | -3.584087208 | ZNF850 |
| 17 | 68271414 | 68271612 | 3.883477159 | SLC16A6 | 19 | 36750463 | 36750599 | -3.241410525 | ZNF850 |
| 17 | 68271414 | 68271612 | 3.883477159 | ARSG | 19 | 36819821 | 36820014 | -1.374028869 | ZNF790-AS1 |
| 17 | 68399296 | 68399400 | -1.119975396 | ARSG | 19 | 36819821 | 36820014 | -1.374028869 | ZNF790 |
| 17 | 70168614 | 70168821 | 8.626425096 | KCNJ2 | 19 | 37127102 | 37127293 | -7.828659225 | CTC-454I21.3 |
| 17 | 70168614 | 70168821 | 8.626425096 | KCNJ2-AS1 | 19 | 37127102 | 37127293 | -7.828659225 | ZNF420 |
| 17 | 70174879 | 70175077 | 10.60254104 | KCNJ2 | 19 | 37127102 | 37127293 | -7.828659225 | ZNF585A |
| 17 | 70176190 | 70176388 | 5.279329934 | KCNJ2 | 19 | 37151351 | 37151527 | -2.037873269 | ZNF585A |
| 17 | 72646627 | 72646826 | 1.074569869 | SLC39A11 | 19 | 37151351 | 37151527 | -2.037873269 | CTC-454I21.3 |
| 1 | 89869393 | 89869591 | -1.550234012 | LRRC8D | 19 | 37184964 | 37185164 | -1.167217339 | CTC-454I21.3 |
| 1 | 89869393 | 89869591 | -1.550234012 | RP11-302M6.4 | 19 | 37184964 | 37185164 | -1.167217339 | ZNF585B |
| 17 | 73236265 | 73236464 | -1.640788797 | FAM104A | 19 | 37186741 | 37186959 | -1.829346428 | ZNF585B |
| 17 | 73236265 | 73236464 | -1.640788797 | C17orf80 | 19 | 37186741 | 37186959 | -1.829346428 | CTC-454I21.3 |
| 17 | 73236795 | 73237000 | -2.015678298 | C17orf80 | 19 | 37389408 | 37389529 | -2.929600158 | ZNF527 |
| 17 | 74431532 | 74431794 | 3.599154305 | GPRC5C | 19 | 37484139 | 37484323 | -2.159416261 | ZNF570 |
| 17 | 74522414 | 74522611 | 3.844659288 | CD300LB | 19 | 37484914 | 37485118 | -1.105786257 | ZNF570 |
| 17 | 74771409 | 74771613 | -2.012840007 | NAT9 | 19 | 37635559 | 37635759 | -7.14587314 | ZFP30 |
| 17 | 75003774 | 75003976 | 3.772291613 | CDR2L | 19 | 37739652 | 37739876 | -2.640621922 | ZNF573 |
| 1 | 89933836 | 89934035 | -1.424269738 | LRRC8D | 19 | 38206660 | 38206891 | -1.49627049 | SIPA1L3 |
| 17 | 75100234 | 75100417 | -2.020968447 | SLC16A5 | 19 | 38327287 | 38327443 | -6.862256927 | KCNK6 |
| 17 | 75129087 | 75129297 | -1.193916845 | ARMC7 | 1 | 110848275 | 110848589 | -1.345114516 | NRBF2P3 |
| 17 | 75236998 | 75237180 | -1.08506993 | GGA3 | 19 | 38910658 | 38910858 | -1.846743479 | CCER2 |
| 17 | 75237581 | 75237699 | -1.0631796 | GGA3 | 19 | 39307618 | 39307816 | 6.267273034 | LRFN1 |
| 17 | 75238052 | 75238246 | -1.537929815 | GGA3 | 19 | 39314162 | 39314361 | -2.877443751 | LRFN1 |
| 17 | 75271881 | 75272223 | -1.947825966 | RP11-649A18.12 | 19 | 39314665 | 39314818 | -2.923563482 | LRFN1 |
| 17 | 75273222 | 75273420 | -3.283309345 | SLC25A19 | 19 | 39314665 | 39314818 | -2.923563482 | CTC-246B18.8 |
| 17 | 75273222 | 75273420 | -3.283309345 | RP11-649A18.12 | 19 | 39408026 | 39408226 | 1.109751572 | ZFP36 |
| 17 | 75502046 | 75502199 | -1.338761951 | CASKIN2 | 19 | 39532181 | 39532371 | -1.015706476 | EID2B |
| 17 | 75522273 | 75522467 | -2.454226318 | TSEN54 | 19 | 40014069 | 40014266 | -1.59587968 | ZNF546 |
| 1 | 89935304 | 89935515 | -1.093689739 | LRRC8D | 19 | 40023560 | 40023797 | -1.429013193 | CTC-471F3.5 |
| 17 | 75661030 | 75661227 | -1.677303808 | RECQL5 | 19 | 40025225 | 40025420 | -6.125593252 | CTC-471F3.5 |
| 17 | 75662574 | 75662777 | -1.611829095 | RECQL5 | 19 | 40034418 | 40034546 | -1.297371761 | ZNF780B |
| 17 | 75706451 | 75706660 | -1.098824615 | SAP30BP | 19 | 40035251 | 40035431 | -1.318698425 | ZNF780B |
| 17 | 75890982 | 75891182 | -2.124309564 | TRIM65 | 19 | 40038068 | 40038245 | -1.92340349 | ZNF780B |
| 17 | 75937947 | 75938145 | -3.109594066 | FBF1 | 19 | 40038899 | 40039101 | -1.621133754 | ZNF780B |
| 17 | 76039205 | 76039402 | -1.688790922 | SRP68 | 19 | 40041681 | 40041917 | -1.17367246 | ZNF780B |

| | | | | |
|---|---|---|---|---|
| 17 | 76077800 | 76077957 | 7.282848563 | ZACN |
| 17 | 76081151 | 76081305 | -1.231008993 | ZACN |
| 17 | 76081151 | 76081305 | -1.231008993 | EXOC7 |
| 17 | 76083446 | 76083625 | -1.402020438 | ZACN |
| 17 | 76083446 | 76083625 | -1.402020438 | EXOC7 |
| 17 | 76389698 | 76389898 | -1.004747549 | UBE2O |
| 17 | 76390020 | 76390131 | -1.124665459 | UBE2O |
| 17 | 76471430 | 76471668 | 3.906992664 | RHBDF2 |
| 17 | 76565508 | 76565698 | 1.824174011 | ST6GALNAC2 |
| 17 | 78050895 | 78051096 | -1.246987645 | TNRC6C |
| 17 | 78105874 | 78106069 | -1.516626781 | TNRC6C |
| 17 | 78171914 | 78172079 | -1.265098159 | SYNGR2 |
| 17 | 78345120 | 78345318 | -3.372678432 | AC061992.2 |
| 17 | 78358008 | 78358205 | 1.008818093 | SOCS3 |
| 17 | 78891071 | 78891249 | 4.72766361 | TIMP2 |
| 17 | 78891071 | 78891249 | 4.72766361 | CEP295NL |
| 17 | 78972285 | 78972479 | 5.95093626 | LGALS3BP |
| 17 | 79086538 | 79086792 | -1.31236815 | ENGASE |
| 17 | 79087937 | 79088137 | -1.313330647 | ENGASE |
| 17 | 79784754 | 79784900 | -3.266780388 | CBX2 |
| 17 | 79794674 | 79794844 | -1.766647251 | CBX8 |
| 17 | 79833905 | 79834082 | -1.314162421 | CBX4 |
| 17 | 79932440 | 79932643 | 2.885390439 | TBC1D16 |
| 17 | 80045652 | 80045857 | 1.871959567 | CCDC40 |
| 17 | 80252831 | 80253029 | 11.1403834 | SLC26A11 |
| 17 | 80288042 | 80288235 | 1.08926564 | RNF213 |
| 17 | 80295498 | 80295698 | 2.856744811 | RNF213 |
| 17 | 80332078 | 80332279 | 2.790028962 | RNF213 |
| 17 | 80339343 | 80339543 | 2.589237323 | RNF213 |
| 17 | 80347253 | 80347438 | 2.154935415 | RNF213 |
| 17 | 80348002 | 80348192 | 2.444089936 | RNF213 |
| 17 | 80365224 | 80365418 | -4.885744735 | CTD-2047H16.4 |
| 17 | 80365224 | 80365418 | -4.885744735 | RNF213 |
| 17 | 80384701 | 80384909 | 2.142947682 | CTD-2047H16.4 |
| 17 | 80384701 | 80384909 | 2.142947682 | RNF213 |
| 17 | 80388801 | 80388991 | -6.831890225 | RNF213 |
| 17 | 80388801 | 80388991 | -6.831890225 | CTD-2047H16.4 |
| 17 | 80394056 | 80394302 | 2.720820639 | RNF213 |
| 17 | 80394056 | 80394302 | 2.720820639 | CTD-2047H16.4 |
| 17 | 80395516 | 80395715 | 8.583395375 | RNF213 |
| 17 | 80395516 | 80395715 | 8.583395375 | CTD-2047H16.4 |
| 17 | 80581966 | 80582131 | -7.241970853 | RPTOR |
| 1 | 91020911 | 91021053 | -1.099339307 | ZNF644 |
| 17 | 80727249 | 80727378 | 4.563352236 | RPTOR |
| 17 | 80811656 | 80811817 | -6.614409345 | RPTOR |
| 17 | 80891132 | 80891245 | -3.595885239 | RPTOR |
| 17 | 80931364 | 80931561 | 6.658706307 | RPTOR |
| 17 | 80964719 | 80964915 | -1.283331175 | RPTOR |
| 17 | 80970698 | 80970865 | 1.258924152 | CTD-2561B21.5 |
| 1 | 91524045 | 91524298 | -3.41762698 | CDC7 |
| 17 | 81110106 | 81110302 | 1.006011566 | BAIAP2 |
| 17 | 81114684 | 81114871 | 5.879319819 | BAIAP2 |
| 17 | 81121460 | 81121657 | 2.36735481 | AATK |
| 17 | 81245916 | 81246106 | -1.3151691 | SLC38A10 |

| | | | | |
|---|---|---|---|---|
| 19 | 40043289 | 40043511 | -1.404222491 | ZNF780B |
| 19 | 40065792 | 40065990 | -1.404595181 | AC005614.5 |
| 19 | 40800884 | 40801083 | 8.437058427 | RAB4B-EGLN2 |
| 19 | 40800884 | 40801083 | 8.437058427 | EGLN2 |
| 19 | 40826657 | 40826800 | -1.993389796 | CYP2F2P |
| 19 | 40826657 | 40826800 | -1.993389796 | CTC-490E21.12 |
| 19 | 41425503 | 41425703 | -1.620542135 | B3GNT8 |
| 19 | 41425503 | 41425703 | -1.620542135 | CTC-435M10.6 |
| 19 | 41426383 | 41426578 | -2.032176983 | B3GNT8 |
| 1 | 111187283 | 111187470 | -1.837095167 | DENND2D |
| 19 | 42088557 | 42088758 | 2.341485122 | POU2F2 |
| 19 | 42089704 | 42089904 | 2.397404495 | POU2F2 |
| 19 | 42091013 | 42091199 | 2.260783559 | POU2F2 |
| 19 | 42137823 | 42137926 | 2.203230803 | POU2F2 |
| 19 | 42271926 | 42272056 | 8.858429821 | CIC |
| 19 | 43586104 | 43586312 | -1.284382719 | L34079.2 |
| 19 | 43586104 | 43586312 | -1.284382719 | IRGQ |
| 19 | 43592726 | 43592932 | -1.022618633 | L34079.2 |
| 19 | 43592726 | 43592932 | -1.022618633 | IRGQ |
| 1 | 3770693 | 3770889 | -1.45912672 | CCDC27 |
| 19 | 43659729 | 43659890 | -8.804509991 | PLAUR |
| 19 | 43847179 | 43847411 | -1.458574357 | ZNF283 |
| 19 | 43966871 | 43967107 | -2.540437723 | ZNF221 |
| 19 | 43970419 | 43970609 | -1.313042678 | ZNF155 |
| 19 | 43971454 | 43971694 | -1.517454299 | ZNF155 |
| 19 | 43973230 | 43973423 | -7.061979855 | ZNF155 |
| 1 | 111500252 | 111500446 | -4.997176718 | TMIGD3 |
| 1 | 111500252 | 111500446 | -4.997176718 | ADORA3 |
| 19 | 43974069 | 43974368 | -2.036309059 | ZNF155 |
| 19 | 43996162 | 43996358 | -1.629940055 | ZNF155 |
| 19 | 43997136 | 43997337 | -1.410047084 | RP11-15A1.7 |
| 19 | 43997136 | 43997337 | -1.410047084 | ZNF155 |
| 19 | 44032506 | 44032705 | -1.575085071 | ZNF222 |
| 19 | 44032506 | 44032705 | -1.575085071 | AC084219.2 |
| 19 | 44066902 | 44067107 | -2.064337277 | AC084219.2 |
| 19 | 44066902 | 44067107 | -2.064337277 | ZNF223 |
| 19 | 44086808 | 44087032 | -1.152061438 | AC084219.2 |
| 19 | 44086808 | 44087032 | -1.152061438 | ZNF284 |
| 19 | 44131767 | 44131976 | -1.330830228 | ZNF225 |
| 1 | 111503406 | 111503568 | -4.25389732 | ADORA3 |
| 1 | 111503406 | 111503568 | -4.25389732 | TMIGD3 |
| 19 | 44250995 | 44251188 | -6.006661337 | ZNF235 |
| 19 | 44252701 | 44252883 | -3.257849919 | ZNF235 |
| 19 | 44477017 | 44477209 | -1.20290993 | ZNF180 |
| 19 | 45145594 | 45145760 | 6.882228482 | MARK4 |
| 19 | 45145594 | 45145760 | 6.882228482 | PPP1R37 |
| 19 | 45152860 | 45153000 | -5.56592848 | MARK4 |
| 19 | 45152860 | 45153000 | -5.56592848 | NKPD1 |
| 19 | 45178413 | 45178602 | 2.083834018 | MARK4 |
| 19 | 45179805 | 45180025 | 1.877631109 | BLOC1S3 |
| 19 | 45179805 | 45180025 | 1.877631109 | AC005779.2 |
| 19 | 45179805 | 45180025 | 1.877631109 | MARK4 |
| 19 | 45408488 | 45408683 | -7.335964002 | CD3EAP |
| 19 | 45408488 | 45408683 | -7.335964002 | ERCC1 |

| | | | | |
|---|---|---|---|---|
| 17 | 81250617 | 81250812 | -1.133488374 | SLC38A10 |
| 17 | 81252335 | 81252509 | -2.085164871 | SLC38A10 |
| 17 | 81304898 | 81305097 | -2.3866975 | LINC00482 |
| 17 | 81323963 | 81324123 | -7.07487356 | TMEM105 |
| 17 | 81324859 | 81325046 | -3.234521607 | TMEM105 |
| 17 | 81327252 | 81327448 | -4.506620129 | TMEM105 |
| 17 | 81442533 | 81442907 | -1.85295384 | BAHCC1 |
| 17 | 81447302 | 81447509 | -1.488607374 | BAHCC1 |
| 17 | 81451565 | 81451756 | -6.128546572 | BAHCC1 |
| 17 | 81666098 | 81666302 | -2.018612197 | OXLD1 |
| 17 | 81673121 | 81673345 | -2.560062716 | CCDC137 |
| 17 | 81707223 | 81707420 | -8.65050245 | RP13-1032I1.10 |
| 17 | 81707223 | 81707420 | -8.65050245 | MRPL12 |
| 17 | 81710218 | 81710423 | -3.417027578 | RP13-1032I1.10 |
| 17 | 81710693 | 81710892 | -1.810773549 | RP13-1032I1.10 |
| 17 | 81843258 | 81843454 | -12.92533423 | RP11-498C9.2 |
| 17 | 81843258 | 81843454 | -12.92533423 | P4HB |
| 17 | 81868775 | 81868976 | -1.597696708 | ARHGDIA |
| 17 | 81901179 | 81901379 | -2.773107053 | PCYT2 |
| 17 | 81901179 | 81901379 | -2.773107053 | NPB |
| 17 | 81904551 | 81904778 | -1.9987133 | PCYT2 |
| 17 | 81912258 | 81912462 | -1.699012182 | SIRT7 |
| 17 | 81930277 | 81930477 | -1.734011975 | MAFG-AS1 |
| 17 | 81947325 | 81947446 | 1.991496787 | AC137723.5 |
| 17 | 82018984 | 82019184 | -3.250109505 | STRA13 |
| 17 | 82056074 | 82056269 | -1.762195016 | GPS1 |
| 17 | 82058009 | 82058228 | -1.818341365 | DUS1L |
| 17 | 82152880 | 82153070 | 1.043782444 | CCDC57 |
| 17 | 82154534 | 82154746 | 1.025909612 | CCDC57 |
| 17 | 82154534 | 82154746 | 1.025909612 | RP11-1376P16.2 |
| 17 | 82156871 | 82157071 | 1.018101281 | CCDC57 |
| 17 | 82195640 | 82195773 | -7.549838277 | CCDC57 |
| 17 | 82243151 | 82243394 | 1.262607419 | SLC16A3 |
| 17 | 82243151 | 82243394 | 1.262607419 | CSNK1D |
| 1 | 92184057 | 92184182 | 2.668321168 | KIAA1107 |
| 1 | 92184057 | 92184182 | 2.668321168 | BTBD8 |
| 17 | 82246665 | 82246864 | 1.856232498 | CSNK1D |
| 17 | 82246665 | 82246864 | 1.856232498 | SLC16A3 |
| 17 | 82247553 | 82247741 | 2.712544797 | CSNK1D |
| 17 | 82247553 | 82247741 | 2.712544797 | SLC16A3 |
| 17 | 82248966 | 82249165 | 2.716260542 | CSNK1D |
| 17 | 82248966 | 82249165 | 2.716260542 | RP13-516M14.8 |
| 17 | 82248966 | 82249165 | 2.716260542 | SLC16A3 |
| 17 | 82249820 | 82249996 | 3.298959953 | SLC16A3 |
| 17 | 82249820 | 82249996 | 3.298959953 | CSNK1D |
| 17 | 82249820 | 82249996 | 3.298959953 | RP13-516M14.8 |
| 17 | 82250449 | 82250606 | 7.914861368 | CSNK1D |
| 17 | 82250449 | 82250606 | 7.914861368 | RP13-516M14.8 |
| 17 | 82250449 | 82250606 | 7.914861368 | SLC16A3 |
| 17 | 82253317 | 82253473 | 8.707156807 | CSNK1D |
| 17 | 82253317 | 82253473 | 8.707156807 | SLC16A3 |
| 17 | 82255711 | 82255911 | 3.13339327 | CSNK1D |
| 17 | 82255711 | 82255911 | 3.13339327 | SLC16A3 |
| 17 | 82422578 | 82422686 | -1.455726415 | HEXDC |

| | | | | |
|---|---|---|---|---|
| 19 | 45409497 | 45409673 | -2.726446099 | CD3EAP |
| 19 | 45409497 | 45409673 | -2.726446099 | ERCC1 |
| 19 | 45468256 | 45468480 | -1.769595341 | ERCC1 |
| 19 | 45468256 | 45468480 | -1.769595341 | FOSB |
| 19 | 45472915 | 45473127 | -1.199205321 | ERCC1 |
| 19 | 45472915 | 45473127 | -1.199205321 | FOSB |
| 19 | 45529473 | 45529593 | 1.495682388 | OPA3 |
| 19 | 45681958 | 45682158 | -6.567854643 | GIPR |
| 19 | 45768980 | 45769209 | -2.462801339 | SIX5 |
| 19 | 45768980 | 45769209 | -2.462801339 | AC074212.6 |
| 19 | 45768980 | 45769209 | -2.462801339 | AC074212.5 |
| 1 | 111619297 | 111619497 | 7.718941043 | RAP1A |
| 19 | 45885600 | 45885777 | 3.729715809 | IRF2BP1 |
| 19 | 46620621 | 46620779 | 4.007850071 | PTGIR |
| 19 | 46621054 | 46621245 | 2.703255652 | PTGIR |
| 19 | 46755532 | 46755721 | 5.612603718 | FKRP |
| 19 | 46788099 | 46788294 | -1.829409925 | CTB-147N14.6 |
| 19 | 46788099 | 46788294 | -1.829409925 | SLC1A5 |
| 19 | 46918832 | 46919047 | -2.017936248 | ARHGAP35 |
| 19 | 46921231 | 46921459 | -1.550926055 | ARHGAP35 |
| 19 | 47001080 | 47001279 | -2.066942087 | ARHGAP35 |
| 19 | 47069153 | 47069383 | -9.641609756 | ZC3H4 |
| 1 | 111724217 | 111724457 | -4.041813947 | FAM212B |
| 19 | 47320377 | 47320573 | -1.825523565 | C5AR1 |
| 19 | 47332134 | 47332244 | -6.946366266 | C5AR2 |
| 19 | 47341589 | 47341799 | -2.218619166 | C5AR2 |
| 1 | 111727272 | 111727545 | -3.720767422 | FAM212B |
| 19 | 48093953 | 48094119 | 1.781612052 | PLA2G4C |
| 19 | 48170908 | 48171107 | -5.629283017 | C19orf68 |
| 19 | 48195402 | 48195598 | -4.307354922 | ZNF114 |
| 19 | 48195402 | 48195598 | -4.307354922 | CARD8 |
| 19 | 48195402 | 48195598 | -4.307354922 | C19orf68 |
| 19 | 48196553 | 48196756 | -1.378403271 | ZNF114 |
| 19 | 48196553 | 48196756 | -1.378403271 | C19orf68 |
| 19 | 48196553 | 48196756 | -1.378403271 | CARD8 |
| 19 | 48203843 | 48204058 | -1.773267894 | CARD8 |
| 19 | 48203843 | 48204058 | -1.773267894 | ZNF114 |
| 1 | 3775899 | 3776101 | -1.621109363 | SMIM1 |
| 19 | 48286153 | 48286292 | -3.627682923 | ZNF114 |
| 19 | 48286664 | 48286868 | -2.547898292 | ZNF114 |
| 19 | 48453067 | 48453268 | 1.133888452 | GRWD1 |
| 19 | 48498204 | 48498421 | -2.425982193 | LMTK3 |
| 19 | 49310638 | 49310885 | -2.189300272 | SLC6A16 |
| 19 | 49525037 | 49525237 | -1.020460821 | FCGRT |
| 19 | 49596221 | 49596422 | -3.36032382 | PRR12 |
| 19 | 49599274 | 49599473 | -8.087492091 | PRR12 |
| 19 | 49651303 | 49651492 | 9.07275216 | SCAF1 |
| 19 | 49857848 | 49858031 | 8.54322363 | PTOV1 |
| 19 | 49857848 | 49858031 | 8.54322363 | PTOV1-AS2 |
| 19 | 49877181 | 49877381 | 3.071722273 | AKT1S1 |
| 19 | 49877181 | 49877381 | 3.071722273 | PNKP |
| 19 | 49878972 | 49879148 | -9.409632697 | TBC1D17 |
| 19 | 49879531 | 49879713 | 3.50363046 | TBC1D17 |
| 19 | 49880533 | 49880725 | 4.048197737 | TBC1D17 |

| | | | | |
|---|---|---|---|---|
| 17 | 82443408 | 82443606 | -1.89296942 | C17orf62 |
| 17 | 82457874 | 82458085 | -1.119599852 | RP13-20L14.6 |
| 17 | 82602997 | 82603277 | -1.602373186 | RP13-638C3.4 |
| 17 | 82602997 | 82603277 | -1.602373186 | FOXK2 |
| 17 | 82656314 | 82656506 | 2.190122295 | RAB40B |
| 17 | 83012277 | 83012472 | -1.438650547 | B3GNTL1 |
| 17 | 83085006 | 83085207 | 3.545345631 | METRNL |
| 1 | 92390499 | 92390707 | 1.022475498 | RPAP2 |
| 17 | 83094358 | 83094558 | 3.439748231 | METRNL |
| 17 | 83105200 | 83105430 | 4.427378547 | AC144831.1 |
| 18 | 244437 | 244566 | -2.616788766 | THOC1 |
| 18 | 245539 | 245782 | -1.597800723 | THOC1 |
| 18 | 659603 | 659801 | 6.682887818 | TYMS |
| 1 | 2384108 | 2384303 | -4.052599965 | MORN1 |
| 18 | 662841 | 662998 | 8.905084969 | TYMS |
| 18 | 664661 | 664770 | 5.477098155 | TYMS |
| 18 | 667685 | 667852 | 7.157539392 | TYMS |
| 18 | 670603 | 670810 | 1.157866638 | ENOSF1 |
| 18 | 670603 | 670810 | 1.157866638 | TYMS |
| 18 | 2891080 | 2891281 | -1.303574371 | EMILIN2 |
| 18 | 2892090 | 2892296 | -1.34927098 | EMILIN2 |
| 18 | 2913583 | 2913784 | -1.204537314 | EMILIN2 |
| 18 | 2914533 | 2914881 | -2.12713702 | EMILIN2 |
| 18 | 2920855 | 2921033 | -6.235329437 | LPIN2 |
| 18 | 2920855 | 2921033 | -6.235329437 | RP11-737O24.5 |
| 18 | 3246851 | 3247065 | -1.147193412 | RP13-270P17.2 |
| 18 | 3246851 | 3247065 | -1.147193412 | RP13-270P17.3 |
| 18 | 3457627 | 3457891 | 1.885651242 | TGIF1 |
| 18 | 3596614 | 3596812 | 3.911363354 | DLGAP1-AS1 |
| 18 | 3596614 | 3596812 | 3.911363354 | DLGAP1 |
| 18 | 8762411 | 8762612 | -1.879803898 | MTCL1 |
| 18 | 8765671 | 8765868 | 2.030023692 | MTCL1 |
| 18 | 8784173 | 8784369 | -7.419257282 | MTCL1 |
| 18 | 8825434 | 8825743 | -1.002470975 | MTCL1 |
| 1 | 92484918 | 92485135 | -1.436862336 | GFI1 |
| 18 | 9270309 | 9270496 | 1.351065924 | ANKRD12 |
| 18 | 9524470 | 9524666 | -2.113004451 | RALBP1 |
| 18 | 9795327 | 9795527 | -5.780405234 | RAB31 |
| 18 | 9859329 | 9859528 | 1.413982021 | RAB31 |
| 18 | 9887888 | 9888075 | 2.599968785 | TXNDC2 |
| 1 | 92513691 | 92513890 | 1.133029388 | EVI5 |
| 18 | 11884382 | 11884584 | -1.07150533 | GNAL |
| 18 | 11884382 | 11884584 | -1.07150533 | MPPE1 |
| 18 | 12326146 | 12326346 | -2.596505284 | TUBB6 |
| 18 | 12329621 | 12329822 | -1.333183275 | AFG3L2 |
| 18 | 12329621 | 12329822 | -1.333183275 | TUBB6 |
| 18 | 12449233 | 12449561 | 1.898211943 | SPIRE1 |
| 18 | 12628082 | 12628278 | 1.26064089 | SPIRE1 |
| 18 | 13049119 | 13049318 | -1.782372104 | CEP192 |
| 18 | 13056253 | 13056452 | -1.559702661 | CEP192 |
| 18 | 13108281 | 13108421 | -2.044404633 | CEP192 |
| 18 | 13479533 | 13479729 | -1.142980751 | LDLRAD4 |
| 18 | 13498291 | 13498466 | -1.478984874 | LDLRAD4 |
| 18 | 13499386 | 13499568 | -1.568887685 | LDLRAD4 |

| | | | | |
|---|---|---|---|---|
| 19 | 49881205 | 49881403 | 10.23298942 | TBC1D17 |
| 19 | 49884434 | 49884639 | 3.880309804 | TBC1D17 |
| 19 | 49886523 | 49886720 | 3.34135991 | TBC1D17 |
| 19 | 49887251 | 49887451 | 3.197282942 | TBC1D17 |
| 19 | 49887903 | 49888097 | 3.075444111 | TBC1D17 |
| 19 | 49889763 | 49889972 | 3.386785568 | IL4I1 |
| 19 | 49932397 | 49932604 | 2.057313081 | ATF5 |
| 19 | 49932397 | 49932604 | 2.057313081 | CTC-326K19.6 |
| 19 | 49933601 | 49933775 | 1.559732499 | CTC-326K19.6 |
| 19 | 49933601 | 49933775 | 1.559732499 | ATF5 |
| 19 | 50045870 | 50045991 | -1.297358659 | ZNF473 |
| 19 | 50045870 | 50045991 | -1.297358659 | CTD-2126E3.3 |
| 19 | 50323278 | 50323480 | -1.385244836 | KCNC3 |
| 19 | 50509364 | 50509528 | -3.176212348 | JOSD2 |
| 19 | 50661777 | 50661952 | -1.090271143 | SHANK1 |
| 19 | 50798157 | 50798438 | -1.380740818 | C19orf48 |
| 19 | 51131081 | 51131280 | -1.483416568 | SIGLEC9 |
| 19 | 51587211 | 51587411 | -1.798766736 | ZNF175 |
| 19 | 51588138 | 51588323 | -1.29918067 | ZNF175 |
| 1 | 112853846 | 112854045 | -2.884277903 | RP3-522D1.1 |
| 19 | 51872758 | 51872943 | -1.516262907 | AC074141.1 |
| 19 | 51872758 | 51872943 | -1.516262907 | ZNF577 |
| 19 | 51944216 | 51944404 | -2.405735515 | ZNF613 |
| 19 | 51965255 | 51965363 | -1.057344519 | ZNF350-AS1 |
| 19 | 51965255 | 51965363 | -1.057344519 | ZNF350 |
| 19 | 51992436 | 51992568 | -4.93518236 | ZNF615 |
| 19 | 51993079 | 51993286 | -1.965497459 | ZNF615 |
| 1 | 112855282 | 112855514 | -7.134197897 | RP3-522D1.1 |
| 19 | 52035027 | 52035218 | -1.51310494 | ZNF432 |
| 19 | 52060424 | 52060617 | 2.254531193 | CTC-471J1.2 |
| 19 | 52060424 | 52060617 | 2.254531193 | ZNF432 |
| 19 | 52113725 | 52113915 | -4.106922567 | ZNF616 |
| 19 | 52116100 | 52116301 | -2.630951662 | ZNF616 |
| 1 | 112913975 | 112914176 | -1.153206573 | SLC16A1 |
| 19 | 52156788 | 52156987 | -2.171963206 | ZNF836 |
| 19 | 52156788 | 52156987 | -2.171963206 | CTC-471J1.8 |
| 19 | 52290213 | 52290412 | -1.193401465 | ZNF766 |
| 19 | 52290213 | 52290412 | -1.193401465 | CTD-2525I3.5 |
| 19 | 52324628 | 52324804 | -2.340835661 | CTD-2525I3.6 |
| 19 | 52324628 | 52324804 | -2.340835661 | ZNF480 |
| 19 | 52325482 | 52325681 | -1.199457476 | CTD-2525I3.6 |
| 19 | 52325482 | 52325681 | -1.199457476 | ZNF480 |
| 19 | 52366103 | 52366316 | -1.879301372 | ZNF610 |
| 19 | 52415991 | 52416108 | -1.7667161 | ZNF528 |
| 1 | 112917323 | 112917526 | -1.326708335 | SLC16A1 |
| 19 | 52416747 | 52416877 | -1.083722697 | ZNF528 |
| 19 | 52555328 | 52555534 | -1.539889558 | ZNF808 |
| 19 | 52563554 | 52563750 | -3.876214356 | ZNF808 |
| 19 | 52563554 | 52563750 | -3.876214356 | ZNF701 |
| 19 | 52583724 | 52583881 | -2.016247724 | CTD-3099C6.7 |
| 19 | 52583724 | 52583881 | -2.016247724 | ZNF701 |
| 19 | 52612777 | 52612983 | -1.302868695 | ZNF83 |
| 19 | 52614464 | 52614771 | -1.260644378 | ZNF83 |
| 1 | 112956787 | 112956921 | -1.577206001 | SLC16A1 |

| | | | | |
|---|---|---|---|---|
| 18 | 13500639 | 13500851 | -3.553981196 | RP11-53B2.5 |
| 18 | 13500639 | 13500851 | -3.553981196 | LDLRAD4 |
| 18 | 13525674 | 13525862 | -2.030809947 | LDLRAD4 |
| 1 | 93166745 | 93166989 | 1.828722104 | TMED5 |
| 18 | 13645310 | 13645511 | 1.545354015 | LDLRAD4 |
| 18 | 13645310 | 13645511 | 1.545354015 | RP11-701H16.4 |
| 18 | 13681856 | 13682056 | -1.289466272 | FAM210A |
| 18 | 14074594 | 14074694 | -3.169804646 | ZNF519 |
| 18 | 14114111 | 14114300 | -2.547402761 | ZNF519 |
| 1 | 93180374 | 93180575 | -2.172653125 | CCDC18 |
| 1 | 93180374 | 93180575 | -2.172653125 | TMED5 |
| 18 | 22993195 | 22993393 | -1.959340503 | RBBP8 |
| 18 | 23521964 | 23522162 | 3.386241757 | C18orf8 |
| 18 | 23521964 | 23522162 | 3.386241757 | NPC1 |
| 18 | 23531934 | 23532143 | 1.75879597 | NPC1 |
| 18 | 23648885 | 23649030 | 1.893298181 | ANKRD29 |
| 18 | 24140794 | 24140967 | -2.118919348 | CABYR |
| 18 | 25224628 | 25224823 | -1.722879513 | ZNF521 |
| 1 | 93219654 | 93219849 | -1.655489078 | CCDC18 |
| 18 | 25226973 | 25227172 | -1.393787212 | ZNF521 |
| 18 | 26389905 | 26390036 | -3.731826042 | TAF4B |
| 18 | 31625454 | 31625639 | -1.355776798 | B4GALT6 |
| 18 | 32268037 | 32268242 | 2.100259067 | GAREM1 |
| 18 | 32288025 | 32288228 | 1.634946762 | GAREM1 |
| 18 | 32288025 | 32288228 | 1.634946762 | RP11-344B2.2 |
| 18 | 34725045 | 34725216 | 2.951639671 | DTNA |
| 18 | 35242573 | 35242792 | -1.939131585 | ZNF397 |
| 1 | 93220875 | 93221068 | 1.18855604 | CCDC18 |
| 18 | 35260885 | 35261085 | 1.206058034 | ZSCAN30 |
| 18 | 35260885 | 35261085 | 1.206058034 | ZNF397 |
| 18 | 35332479 | 35332684 | -1.32391003 | ZNF24 |
| 18 | 35369550 | 35369659 | 1.937912623 | ZNF396 |
| 18 | 35374124 | 35374324 | 1.486937199 | ZNF396 |
| 18 | 35628094 | 35628262 | -1.139139681 | GALNT1 |
| 18 | 35633241 | 35633424 | -3.786733431 | GALNT1 |
| 18 | 36126424 | 36126624 | -1.51567708 | SLC39A6 |
| 18 | 36786586 | 36786769 | -1.976910627 | TPGS2 |
| 18 | 36962707 | 36962860 | -1.448221904 | KIAA1328 |
| 18 | 37027737 | 37027874 | -4.921175172 | KIAA1328 |
| 18 | 44678467 | 44678774 | 2.722093331 | RP11-456K23.1 |
| 18 | 44701391 | 44701594 | 3.635144918 | SETBP1 |
| 18 | 44953013 | 44953224 | 4.48681363 | SETBP1 |
| 18 | 45850218 | 45850386 | -1.099107658 | EPG5 |
| 1 | 93266280 | 93266380 | -3.04505621 | CCDC18 |
| 1 | 93266280 | 93266380 | -3.04505621 | CCDC18-AS1 |
| 18 | 46456602 | 46456738 | -5.77609438 | RNF165 |
| 18 | 46679412 | 46679620 | 1.375186176 | ST8SIA5 |
| 18 | 46946057 | 46946211 | -2.497321771 | KATNAL2 |
| 18 | 48027519 | 48027717 | 7.581824838 | ZBTB7C |
| 1 | 93268232 | 93268430 | -3.570736213 | CCDC18-AS1 |
| 1 | 93268232 | 93268430 | -3.570736213 | CCDC18 |
| 18 | 48862846 | 48863044 | 1.089224537 | CTIF |
| 18 | 48921454 | 48921653 | 2.699063359 | SMAD7 |
| 18 | 48949836 | 48950013 | -7.026835693 | SMAD7 |

| | | | | |
|---|---|---|---|---|
| 1 | 112956787 | 112956921 | -1.577206001 | SLC16A1-AS1 |
| 19 | 52704782 | 52704973 | -1.479498377 | ZNF611 |
| 19 | 52706356 | 52706556 | -1.828238555 | ZNF611 |
| 19 | 52800538 | 52800669 | -2.692613999 | ZNF28 |
| 19 | 52801455 | 52801655 | -1.721851072 | ZNF28 |
| 1 | 113123964 | 113124156 | -1.197155584 | LRIG2 |
| 19 | 52906847 | 52907038 | -1.182364971 | CTD-2331H12.8 |
| 19 | 52908012 | 52908220 | -1.307036803 | CTD-2331H12.8 |
| 19 | 52929247 | 52929451 | -1.273569295 | CTD-2331H12.8 |
| 19 | 52929247 | 52929451 | -1.273569295 | ZNF321P |
| 19 | 52929247 | 52929451 | -1.273569295 | ZNF816-ZNF321P |
| 19 | 52949576 | 52949774 | -1.394506914 | ZNF816 |
| 19 | 52949576 | 52949774 | -1.394506914 | ZNF816-ZNF321P |
| 19 | 52949576 | 52949774 | -1.394506914 | CTD-2331H12.8 |
| 19 | 52950164 | 52950304 | -2.587352688 | ZNF816 |
| 19 | 52950164 | 52950304 | -2.587352688 | ZNF816-ZNF321P |
| 19 | 52950164 | 52950304 | -2.587352688 | CTD-2331H12.8 |
| 1 | 113641988 | 113642210 | -1.07673724 | MAGI3 |
| 19 | 53352383 | 53352487 | -2.622290795 | ZNF525 |
| 19 | 53352383 | 53352487 | -2.622290795 | ZNF845 |
| 19 | 53386209 | 53386370 | -4.287454418 | ZNF525 |
| 19 | 53386209 | 53386370 | -4.287454418 | CTD-3141N22.1 |
| 19 | 53408059 | 53408229 | -1.171332533 | ZNF765 |
| 19 | 53426590 | 53426790 | -1.623066051 | ZNF765 |
| 1 | 113683119 | 113683272 | -6.396801655 | MAGI3 |
| 19 | 53456397 | 53456597 | -1.083161602 | ZNF761 |
| 19 | 53492211 | 53492360 | -1.157147214 | ZNF813 |
| 19 | 53810294 | 53810494 | -4.960176428 | NLRP12 |
| 19 | 53875714 | 53875905 | 3.660964047 | AC008440.5 |
| 19 | 53875714 | 53875905 | 3.660964047 | MYADM |
| 19 | 54096378 | 54096559 | 5.478551021 | OSCAR |
| 19 | 54160914 | 54161155 | -2.536399066 | TMC4 |
| 19 | 54165624 | 54165820 | -6.137771278 | TMC4 |
| 19 | 54179102 | 54179391 | 1.06971999 | MBOAT7 |
| 19 | 54216890 | 54217090 | 2.001987817 | LILRB3 |
| 19 | 54216890 | 54217090 | 2.001987817 | RPS9 |
| 19 | 54354871 | 54355074 | -1.176644759 | LAIR1 |
| 19 | 54444815 | 54444999 | -2.92633557 | LENG8-AS1 |
| 19 | 54498397 | 54498594 | -6.321477112 | LAIR2 |
| 19 | 54636915 | 54637117 | 2.033636309 | LILRB1 |
| 19 | 54636915 | 54637117 | 2.033636309 | AC009892.10 |
| 19 | 54813780 | 54813948 | 7.05979481 | KIR2DL4 |
| 19 | 55041607 | 55041834 | -1.327684966 | CTC-550B14.6 |
| 19 | 55041607 | 55041834 | -1.327684966 | CTC-550B14.7 |
| 19 | 55041607 | 55041834 | -1.327684966 | RDH13 |
| 19 | 55044714 | 55044911 | -1.276736107 | CTC-550B14.7 |
| 19 | 55044714 | 55044911 | -1.276736107 | RDH13 |
| 19 | 55044714 | 55044911 | -1.276736107 | CTC-550B14.6 |
| 19 | 55226953 | 55227152 | -2.772330279 | TMEM86B |
| 19 | 55312580 | 55312888 | 7.050125839 | TMEM150B |
| 19 | 55347521 | 55347726 | -1.949609373 | KMT5C |
| 19 | 55483913 | 55484116 | -1.035268012 | ZNF628 |
| 19 | 55500508 | 55500707 | 1.290911756 | SSC5D |
| 19 | 55502494 | 55502693 | -4.367467688 | SSC5D |

| | | | | | | | | | |
|---|---|---|---|---|---|---|---|---|---|
| 18 | 48950320 | 48950527 | 1.723996703 | SMAD7 | 1 | 113838017 | 113838140 | -2.483276758 | PTPN22 |
| 18 | 49294060 | 49294263 | 2.656236562 | DYM | 19 | 55540884 | 55541043 | 2.810899634 | SBK3 |
| 1 | 93346517 | 93346716 | -1.202121948 | DR1 | 19 | 55615572 | 55615755 | 7.611246526 | ZNF865 |
| 18 | 50269278 | 50269481 | -1.396512099 | MBD1 | 19 | 55616783 | 55616983 | -1.071701193 | ZNF865 |
| 18 | 50271364 | 50271562 | -1.353132681 | MBD1 | 19 | 55620963 | 55621153 | -1.8527202 | ZNF784 |
| 18 | 51002796 | 51002983 | 1.569893315 | RP11-729L2.2 | 19 | 55645152 | 55645359 | -1.281935959 | ZNF581 |
| 18 | 54289730 | 54289905 | -1.063911769 | POLI | 19 | 55645152 | 55645359 | -1.281935959 | CCDC106 |
| 18 | 54290536 | 54290726 | -1.055170501 | POLI | 1 | 113911640 | 113911875 | -1.353689899 | DCLRE1B |
| 18 | 54294276 | 54294491 | -1.382402915 | POLI | 19 | 56422915 | 56423067 | -3.470179274 | ZNF583 |
| 18 | 54378304 | 54378461 | -4.359731722 | C18orf54 | 19 | 56554766 | 56554960 | -1.721093283 | AC007228.11 |
| 18 | 57436743 | 57436934 | -6.027604237 | ONECUT2 | 19 | 56554766 | 56554960 | -1.721093283 | ZFP28 |
| 18 | 57439335 | 57439533 | -7.568715738 | ONECUT2 | 19 | 56577485 | 56577685 | -1.528819313 | ZNF470 |
| 1 | 93514828 | 93514981 | 5.431705979 | FNBP1L | 19 | 57290748 | 57291062 | -1.112258129 | ZNF460 |
| 18 | 57440198 | 57440379 | -8.245174753 | ONECUT2 | 1 | 3779379 | 3779498 | -2.582048697 | LRRC47 |
| 18 | 57476598 | 57476798 | -4.735297299 | ONECUT2 | 19 | 57329027 | 57329225 | -1.380936108 | ZNF543 |
| 18 | 57478341 | 57478442 | -4.205996671 | ONECUT2 | 19 | 57356150 | 57356360 | -1.478069418 | ZNF304 |
| 18 | 57483020 | 57483211 | -3.354157966 | ONECUT2 | 19 | 57377246 | 57377455 | -1.364371411 | ZNF547 |
| 18 | 57483965 | 57484151 | -5.129247785 | ONECUT2 | 19 | 57377246 | 57377455 | -1.364371411 | AC003002.4 |
| 18 | 57668090 | 57668291 | -1.766606489 | RP11-35G9.3 | 19 | 57410933 | 57411152 | -2.57127273 | AC003002.6 |
| 18 | 57668090 | 57668291 | -1.766606489 | RP11-35G9.5 | 19 | 57410933 | 57411152 | -2.57127273 | AC004076.7 |
| 18 | 57668090 | 57668291 | -1.766606489 | ATP8B1 | 19 | 57419846 | 57420141 | -1.103505148 | ZNF17 |
| 18 | 57737776 | 57737885 | -1.553901645 | ATP8B1 | 19 | 57419846 | 57420141 | -1.103505148 | AC003002.6 |
| 18 | 57737776 | 57737885 | -1.553901645 | RP11-35G9.5 | 19 | 57419846 | 57420141 | -1.103505148 | AC004076.7 |
| 18 | 57783982 | 57784146 | -2.683313951 | ATP8B1 | 19 | 57443517 | 57443740 | -1.027331227 | AC004076.9 |
| 18 | 58171313 | 58171502 | -4.258096239 | NEDD4L | 19 | 57443517 | 57443740 | -1.027331227 | ZNF749 |
| 18 | 58747419 | 58747621 | 1.620445761 | RP11-126O1.4 | 19 | 57445176 | 57445315 | -2.035060472 | ZNF749 |
| 18 | 58747419 | 58747621 | 1.620445761 | MALT1 | 19 | 57445176 | 57445315 | -2.035060472 | AC004076.9 |
| 1 | 93550909 | 93551109 | 1.774931497 | FNBP1L | 19 | 57473092 | 57473198 | -1.405276102 | AC004076.9 |
| 18 | 58919818 | 58919999 | 1.147423614 | ZNF532 | 19 | 57473092 | 57473198 | -1.405276102 | ZNF772 |
| 18 | 58984205 | 58984413 | 1.256332065 | ZNF532 | 19 | 57474078 | 57474280 | -1.38675159 | AC004076.9 |
| 18 | 62048989 | 62049159 | -1.345394739 | PIGN | 19 | 57474078 | 57474280 | -1.38675159 | ZNF772 |
| 18 | 62051724 | 62051878 | -1.620693682 | PIGN | 19 | 57492839 | 57493018 | -3.508082712 | ZNF419 |
| 18 | 62369093 | 62369361 | 1.550381488 | TNFRSF11A | 19 | 57492839 | 57493018 | -3.508082712 | AC003005.4 |
| 18 | 62539549 | 62539747 | 2.130908021 | ZCCHC2 | 19 | 57507646 | 57507971 | -1.591794718 | ZNF773 |
| 18 | 62574740 | 62574938 | 2.252492416 | ZCCHC2 | 19 | 57516124 | 57516308 | -1.808816887 | ZNF773 |
| 18 | 62575274 | 62575443 | 2.446886653 | ZCCHC2 | 19 | 57537356 | 57537554 | -1.157739422 | ZNF550 |
| 18 | 62576408 | 62576605 | 7.193739433 | ZCCHC2 | 19 | 57537356 | 57537554 | -1.157739422 | ZNF549 |
| 18 | 62717095 | 62717219 | -2.679229223 | PHLPP1 | 19 | 57538703 | 57538906 | -1.457899648 | ZNF550 |
| 18 | 62829903 | 62830122 | -8.467398296 | PHLPP1 | 19 | 57538703 | 57538906 | -1.457899648 | ZNF549 |
| 18 | 62978888 | 62979140 | -1.880678292 | PHLPP1 | 19 | 57547623 | 57547816 | -2.055157756 | ZNF549 |
| 18 | 63123875 | 63124160 | -2.759326578 | BCL2 | 19 | 57547623 | 57547816 | -2.055157756 | ZNF550 |
| 1 | 93582432 | 93582639 | 2.979922453 | BCAR3 | 19 | 57572233 | 57572430 | -1.215955659 | ZNF416 |
| 18 | 63127417 | 63127617 | -2.543203894 | BCL2 | 19 | 57573285 | 57573501 | -1.818356152 | ZNF416 |
| 18 | 63281162 | 63281309 | -2.871532806 | BCL2 | 19 | 57605933 | 57606153 | -1.800231797 | ZNF530 |
| 18 | 63317599 | 63317773 | -2.348856893 | BCL2 | 19 | 57620326 | 57620524 | -1.68392529 | ZNF134 |
| 18 | 63318109 | 63318317 | -2.830349627 | BCL2 | 19 | 57621285 | 57621521 | -1.615613104 | ZNF134 |
| 18 | 63318659 | 63318788 | -2.770957765 | BCL2 | 19 | 57640997 | 57641197 | -1.773277597 | ZNF211 |
| 18 | 63319424 | 63319613 | -2.833726776 | BCL2 | 19 | 57687610 | 57687848 | -2.152122084 | AC003006.7 |
| 18 | 63902935 | 63903133 | -8.856360508 | SERPINB10 | 19 | 57687610 | 57687848 | -2.152122084 | ZNF551 |
| 18 | 63902935 | 63903133 | -8.856360508 | SERPINB2 | 19 | 57721414 | 57721608 | -1.295317855 | ZNF671 |
| 18 | 64004938 | 64005121 | -1.652525293 | SERPINB8 | 19 | 57721414 | 57721608 | -1.295317855 | AC003006.7 |
| 18 | 68836971 | 68837148 | 1.947104176 | CCDC102B | 19 | 57729512 | 57729663 | -1.079906843 | AC003006.7 |
| 1 | 844057 | 844256 | -1.84767247 | LINC01128 | 19 | 57753854 | 57754054 | -1.65244513 | ZNF776 |
| 1 | 93876782 | 93876976 | -1.540574425 | DNTTIP2 | 1 | 114568081 | 114568284 | -1.75800497 | BCAS2 |

| | | | | |
|---|---|---|---|---|
| 18 | 70098931 | 70099131 | -3.426182328 | RTTN |
| 18 | 70099492 | 70099701 | -2.632597959 | RTTN |
| 18 | 70105153 | 70105266 | -2.438764268 | RTTN |
| 18 | 74520576 | 74520776 | 1.214332912 | CNDP2 |
| 18 | 74633523 | 74633743 | -1.083808563 | ZNF407 |
| 18 | 74970360 | 74970539 | 7.089176812 | ZNF407 |
| 18 | 74971442 | 74971625 | 6.177124691 | ZNF407 |
| 18 | 75063577 | 75063720 | -1.308715681 | ZNF407 |
| 18 | 75064236 | 75064434 | -1.436496454 | ZNF407 |
| 1 | 95244408 | 95244620 | -1.210110984 | RP11-57H12.5 |
| 1 | 95244408 | 95244620 | -1.210110984 | TMEM56-RWDD3 |
| 1 | 95244408 | 95244620 | -1.210110984 | RWDD3 |
| 18 | 76362012 | 76362210 | -2.750271417 | ZNF516 |
| 18 | 76371537 | 76371724 | -1.563757064 | ZNF516 |
| 18 | 76380046 | 76380247 | -1.356484531 | ZNF516 |
| 18 | 76492544 | 76492708 | -8.035228208 | ZNF516 |
| 18 | 76492544 | 76492708 | -8.035228208 | RP11-17M16.2 |
| 18 | 76495512 | 76495664 | -8.30453164 | C18orf65 |
| 18 | 76808677 | 76808867 | -2.067022981 | RP11-162A12.2 |
| 18 | 77016456 | 77016656 | 1.417077078 | MBP |
| 18 | 77035260 | 77035462 | 1.426486899 | MBP |
| 18 | 77037453 | 77037649 | 2.60010487 | MBP |
| 18 | 77133571 | 77133776 | 2.053740583 | MBP |
| 18 | 79115341 | 79115554 | 1.457757561 | ATP9B |
| 18 | 79527541 | 79527753 | 1.163803588 | NFATC1 |
| 18 | 79677686 | 79677910 | -1.404131949 | RP11-567M16.6 |
| 18 | 79677686 | 79677910 | -1.404131949 | RP11-567M16.3 |
| 18 | 79729327 | 79729526 | -2.081048782 | CTDP1 |
| 18 | 79732120 | 79732322 | -1.284634521 | CTDP1 |
| 18 | 79732585 | 79732689 | -1.65519547 | CTDP1 |
| 18 | 79736229 | 79736417 | -7.20087309 | CTDP1 |
| 18 | 79754062 | 79754222 | -1.343137499 | CTDP1 |
| 18 | 80045944 | 80046149 | -1.720987301 | RBFA |
| 18 | 80045944 | 80046149 | -1.720987301 | RP11-795F19.5 |
| 18 | 80137534 | 80137736 | -1.121579797 | ADNP2 |
| 18 | 80159602 | 80159761 | -1.888115903 | PARD6G-AS1 |
| 18 | 80159602 | 80159761 | -1.888115903 | PARD6G |
| 18 | 80160718 | 80160919 | -1.037264287 | PARD6G-AS1 |
| 18 | 80160718 | 80160919 | -1.037264287 | PARD6G |
| 19 | 509101 | 509298 | -2.934874622 | TPGS1 |
| 19 | 509770 | 509929 | -1.57501938 | TPGS1 |
| 19 | 603612 | 603812 | 3.000638666 | HCN2 |
| 19 | 643040 | 643194 | -6.698034779 | FGF22 |
| 19 | 851398 | 851520 | -10.10522153 | ELANE |
| 19 | 867449 | 867666 | -4.062821196 | MED16 |
| 19 | 958413 | 958613 | -4.577409055 | ARID3A |
| 19 | 1005039 | 1005238 | 1.76078828 | GRIN3B |
| 19 | 1085561 | 1085742 | -2.084433129 | ARHGAP45 |
| 19 | 1157894 | 1158191 | -2.395776749 | SBNO2 |
| 19 | 1269936 | 1270145 | -10.09192853 | CIRBP-AS1 |
| 19 | 1269936 | 1270145 | -10.09192853 | CIRBP |
| 19 | 1273729 | 1273999 | -2.107208506 | CIRBP |
| 19 | 1278962 | 1279157 | -1.759493483 | C19orf24 |
| 19 | 1321925 | 1322088 | 2.702495626 | AC005330.2 |

| | | | | |
|---|---|---|---|---|
| 19 | 57770374 | 57770568 | -2.086356794 | ZNF586 |
| 19 | 57814686 | 57814870 | 2.572891653 | ZNF552 |
| 19 | 57814686 | 57814870 | 2.572891653 | ZNF586 |
| 19 | 57841756 | 57841956 | 1.072819594 | CTD-2583A14.10 |
| 19 | 57841756 | 57841956 | 1.072819594 | ZNF587B |
| 19 | 57907500 | 57907681 | -5.582453463 | CTD-2583A14.9 |
| 19 | 57907500 | 57907681 | -5.582453463 | ZNF417 |
| 1 | 114625338 | 114625488 | -1.454782365 | DENND2C |
| 19 | 57909597 | 57909743 | -1.001423237 | CTD-2583A14.9 |
| 19 | 57909597 | 57909743 | -1.001423237 | ZNF417 |
| 19 | 57942265 | 57942570 | -1.764511314 | ZNF256 |
| 19 | 57978875 | 57979098 | -1.984623396 | ZNF606 |
| 19 | 57979886 | 57980085 | -2.171623147 | ZNF606 |
| 19 | 58009410 | 58009603 | -2.343886653 | CTD-2368P22.1 |
| 19 | 58128076 | 58128355 | -1.471976513 | ZNF329 |
| 19 | 58146447 | 58146627 | 2.074701339 | ZNF329 |
| 19 | 58212912 | 58213111 | -1.591498925 | ZNF274 |
| 19 | 58260853 | 58261071 | -1.537721982 | ZNF544 |
| 19 | 58260853 | 58261071 | -1.537721982 | CTD-3138B18.5 |
| 19 | 58260853 | 58261071 | -1.537721982 | CTD-3138B18.4 |
| 19 | 58262756 | 58262956 | -1.016305353 | CTD-3138B18.5 |
| 19 | 58262756 | 58262956 | -1.016305353 | CTD-3138B18.4 |
| 19 | 58262756 | 58262956 | -1.016305353 | ZNF544 |
| 1 | 114625822 | 114626011 | -1.542613918 | DENND2C |
| 19 | 58294358 | 58294553 | -1.791409634 | AC010642.1 |
| 19 | 58294358 | 58294553 | -1.791409634 | CTD-3138B18.4 |
| 19 | 58294358 | 58294553 | -1.791409634 | ZNF8 |
| 19 | 58349863 | 58350050 | -1.008146379 | CTD-2619J13.8 |
| 19 | 58349863 | 58350050 | -1.008146379 | A1BG |
| 19 | 58349863 | 58350050 | -1.008146379 | A1BG-AS1 |
| 19 | 58355696 | 58355873 | -1.17867122 | CTD-2619J13.8 |
| 19 | 58355696 | 58355873 | -1.17867122 | ZNF497 |
| 19 | 58357479 | 58357658 | -1.483465531 | CTD-2619J13.8 |
| 19 | 58357479 | 58357658 | -1.483465531 | ZNF497 |
| 19 | 58358599 | 58358724 | -1.23445154 | CTD-2619J13.9 |
| 19 | 58358599 | 58358724 | -1.23445154 | ZNF497 |
| 19 | 58358599 | 58358724 | -1.23445154 | CTD-2619J13.8 |
| 19 | 58359195 | 58359417 | -1.268030364 | CTD-2619J13.9 |
| 19 | 58359195 | 58359417 | -1.268030364 | ZNF497 |
| 19 | 58359195 | 58359417 | -1.268030364 | CTD-2619J13.8 |
| 19 | 58368807 | 58368967 | 6.031360384 | ZNF837 |
| 19 | 58369799 | 58369994 | -3.554262228 | ZNF837 |
| 19 | 58406084 | 58406284 | -5.293888758 | CTD-2619J13.14 |
| 19 | 58406084 | 58406284 | -5.293888758 | ZNF584 |
| 19 | 58407903 | 58408099 | -1.895364302 | CTD-2619J13.14 |
| 19 | 58407903 | 58408099 | -1.895364302 | ZNF584 |
| 19 | 58417903 | 58418070 | -2.361620417 | ZNF584 |
| 19 | 58452180 | 58452356 | 3.745926548 | ZNF324B |
| 1 | 115663708 | 115663837 | -7.135731514 | VANGL1 |
| 19 | 58475142 | 58475362 | -1.219680955 | ZNF446 |
| 19 | 58475142 | 58475362 | -1.219680955 | ZNF324 |
| 19 | 58475142 | 58475362 | -1.219680955 | CTD-2619J13.23 |
| 19 | 58479113 | 58479311 | -2.234445974 | ZNF446 |
| 19 | 58499695 | 58499879 | -1.201256271 | SLC27A5 |

| | | | | |
|---|---|---|---|---|
| 19 | 1370758 | 1370917 | 1.284275537 | MUM1 |
| 1 | 2508624 | 2508820 | -1.52130505 | PANK4 |
| 1 | 100874055 | 100874266 | -1.221705699 | EXTL2 |
| 19 | 1377160 | 1377323 | -1.69934039 | AC004623.2 |
| 19 | 1377160 | 1377323 | -1.69934039 | MUM1 |
| 19 | 1392551 | 1392739 | 6.774471144 | AC005329.7 |
| 19 | 1392551 | 1392739 | 6.774471144 | NDUFS7 |
| 19 | 1474793 | 1475036 | -3.278342748 | C19orf25 |
| 19 | 1505731 | 1505901 | -1.738227458 | ADAMTSL5 |
| 19 | 1556788 | 1556979 | -8.421126059 | MEX3D |
| 19 | 1574462 | 1574658 | -2.067146071 | MBD3 |
| 19 | 1577655 | 1577856 | -1.669139002 | MBD3 |
| 1 | 100877595 | 100877800 | -1.36261173 | EXTL2 |
| 19 | 1611344 | 1611542 | -2.055765303 | TCF3 |
| 19 | 1782945 | 1783142 | -1.871649764 | ATP8B3 |
| 19 | 1827512 | 1827734 | -1.308310454 | REXO1 |
| 19 | 1854190 | 1854388 | -1.211479063 | KLF16 |
| 19 | 1913142 | 1913322 | -7.572349035 | ADAT3 |
| 19 | 1913142 | 1913322 | -7.572349035 | SCAMP4 |
| 19 | 1986269 | 1986457 | -2.93886057 | BTBD2 |
| 19 | 1989618 | 1989848 | -1.072924671 | AC005306.3 |
| 19 | 1989618 | 1989848 | -1.072924671 | BTBD2 |
| 19 | 2076911 | 2077111 | -4.155468655 | MOB3A |
| 19 | 2192026 | 2192224 | 1.750581946 | DOT1L |
| 19 | 2229940 | 2230147 | 1.379498579 | PLEKHJ1 |
| 19 | 2229940 | 2230147 | 1.379498579 | DOT1L |
| 1 | 103526083 | 103526289 | -1.437947597 | RNPC3 |
| 19 | 2292811 | 2293011 | -2.239457829 | LLfos-48D6.2 |
| 19 | 2294218 | 2294503 | -1.786501569 | LLfos-48D6.2 |
| 19 | 2295086 | 2295287 | -1.771046642 | LLfos-48D6.2 |
| 19 | 2303742 | 2303929 | -6.346743479 | LLfos-48D6.2 |
| 19 | 2353627 | 2353769 | -1.331904527 | SPPL2B |
| 19 | 2426666 | 2426845 | -1.968922453 | TIMM13 |
| 19 | 2477897 | 2478099 | 1.367517962 | GADD45B |
| 19 | 2513474 | 2513583 | 1.123030664 | GNG7 |
| 19 | 2577035 | 2577234 | -6.895683207 | GNG7 |
| 19 | 2734568 | 2734820 | -1.328564599 | AC006538.4 |
| 19 | 2734568 | 2734820 | -1.328564599 | SLC39A3 |
| 19 | 2735390 | 2735504 | -7.583375765 | SLC39A3 |
| 19 | 2735390 | 2735504 | -7.583375765 | AC006538.4 |
| 19 | 2809265 | 2809475 | -3.604879203 | THOP1 |
| 19 | 2877951 | 2878151 | -4.390337087 | ZNF556 |
| 19 | 2917181 | 2917375 | -1.246441805 | AC006277.2 |
| 19 | 2917181 | 2917375 | -1.246441805 | ZNF57 |
| 19 | 2934574 | 2934779 | -1.284219475 | ZNF77 |
| 19 | 3163028 | 3163214 | -1.104000289 | GNA15 |
| 19 | 3207766 | 3207988 | -1.275146523 | NCLN |
| 19 | 3509075 | 3509248 | 5.753649562 | FZR1 |
| 19 | 3516222 | 3516394 | 7.279867784 | FZR1 |
| 19 | 3531354 | 3531528 | -5.667695177 | FZR1 |
| 19 | 3535112 | 3535313 | -1.295020627 | FZR1 |
| 19 | 3538415 | 3538654 | 1.293010819 | MFSD12 |
| 19 | 3562778 | 3562913 | -3.379944092 | MFSD12 |
| 19 | 3565314 | 3565503 | -2.406847724 | MFSD12 |

| | | | | |
|---|---|---|---|---|
| 19 | 58517400 | 58517529 | -6.927628914 | ZBTB45 |
| 19 | 58519146 | 58519350 | -2.883131933 | ZBTB45 |
| 19 | 58581586 | 58581811 | -2.396973409 | CENPBD1P1 |
| 19 | 58582874 | 58583086 | -1.08361629 | CENPBD1P1 |
| 2 | 257907 | 258114 | -1.809738856 | SH3YL1 |
| 2 | 259024 | 259231 | -1.014010833 | SH3YL1 |
| 2 | 3227267 | 3227372 | 6.614409345 | TSSC1 |
| 1 | 116046266 | 116046469 | -1.068559428 | SLC22A15 |
| 2 | 3461976 | 3462110 | 7.03371718 | TRAPPC12 |
| 2 | 6849514 | 6849716 | 2.488446223 | CMPK2 |
| 1 | 116067092 | 116067214 | -1.443031253 | SLC22A15 |
| 2 | 6877420 | 6877617 | 1.699521486 | RSAD2 |
| 2 | 6895789 | 6895994 | 3.479694629 | RSAD2 |
| 2 | 8667047 | 8667242 | -3.008332727 | ID2-AS1 |
| 2 | 8670893 | 8671073 | -3.479482373 | ID2-AS1 |
| 2 | 8683751 | 8683951 | -1.906567593 | ID2 |
| 2 | 8858664 | 8858835 | -5.696158711 | MBOAT2 |
| 2 | 9405863 | 9406060 | -3.802481725 | ITGB1BP1 |
| 1 | 116133272 | 116133431 | -2.582553993 | MAB21L3 |
| 2 | 10047803 | 10048004 | -1.508798693 | KLF11 |
| 2 | 10052407 | 10052529 | -1.178669461 | KLF11 |
| 2 | 10206543 | 10206855 | -2.092766371 | C2orf48 |
| 2 | 10210443 | 10210638 | -1.823215067 | C2orf48 |
| 2 | 10211101 | 10211304 | -1.989580441 | C2orf48 |
| 2 | 10399894 | 10400097 | -2.095593769 | HPCAL1 |
| 2 | 10449111 | 10449309 | -3.058506118 | RP11-320M2.1 |
| 2 | 10913818 | 10914010 | -2.951417615 | KCNF1 |
| 2 | 11318470 | 11318786 | -1.136356544 | ROCK2 |
| 2 | 12740700 | 12740888 | 2.653916633 | TRIB2 |
| 2 | 17655336 | 17655531 | 4.018361553 | VSNL1 |
| 2 | 20202484 | 20202680 | -4.660964047 | SDC1 |
| 2 | 20686711 | 20686974 | -1.687981405 | LDAH |
| 2 | 23691514 | 23691710 | -6.412876916 | KLHL29 |
| 2 | 23927344 | 23927558 | 1.357622408 | UBXN2A |
| 2 | 24077498 | 24077652 | -1.143643896 | FAM228B |
| 2 | 24077498 | 24077652 | -1.143643896 | TP53I3 |
| 1 | 116493134 | 116493334 | -1.660964047 | RP5-1086K13.1 |
| 2 | 24707579 | 24707765 | 1.146867278 | NCOA1 |
| 2 | 24768269 | 24768481 | 1.180822983 | NCOA1 |
| 2 | 24819947 | 24820146 | 8.060301197 | CENPO |
| 2 | 24819947 | 24820146 | 8.060301197 | ADCY3 |
| 2 | 24826308 | 24826504 | 4.027641218 | ADCY3 |
| 2 | 24826308 | 24826504 | 4.027641218 | RP11-443B20.1 |
| 1 | 116525191 | 116525392 | -1.439028611 | CD58 |
| 2 | 24830488 | 24830685 | 4.60715956 | ADCY3 |
| 2 | 25035465 | 25035629 | -7.401258183 | DNAJC27-AS1 |
| 2 | 25234177 | 25234386 | -1.379824556 | DNMT3A |
| 2 | 25326235 | 25326435 | -1.687686473 | DNMT3A |
| 2 | 25414025 | 25414179 | 3.857122759 | DTNB |
| 1 | 116526061 | 116526243 | -3.137396002 | CD58 |
| 2 | 25810051 | 25810306 | -1.635153626 | ASXL2 |
| 2 | 25810051 | 25810306 | -1.635153626 | TPM3P7 |
| 2 | 25856549 | 25856749 | -6.670064137 | AC010150.1 |
| 2 | 25856549 | 25856749 | -6.670064137 | ASXL2 |

| | | | | | | | | | |
|---|---|---|---|---|---|---|---|---|---|
| 19 | 3591396 | 3591603 | 1.114457799 | GIPC3 | 2 | 26311753 | 26311952 | 5.81467831 | EPT1 |
| 19 | 3610871 | 3611070 | -1.644873207 | CACTIN-AS1 | 2 | 26311753 | 26311952 | 5.81467831 | ADGRF3 |
| 19 | 3610871 | 3611070 | -1.644873207 | CACTIN | 2 | 27083445 | 27083722 | -1.217274402 | EMILIN1 |
| 19 | 3611264 | 3611460 | -2.834132509 | CACTIN | 2 | 27264200 | 27264398 | -3.040897046 | SLC30A3 |
| 19 | 3611264 | 3611460 | -2.834132509 | CACTIN-AS1 | 2 | 27326447 | 27326647 | -1.267585648 | GTF3C2 |
| 19 | 3958740 | 3958938 | -1.403727038 | DAPK3 | 1 | 116564174 | 116564346 | -2.355234413 | CD58 |
| 19 | 3976402 | 3976604 | -1.458879042 | EEF2 | 2 | 27627402 | 27627595 | -2.429565089 | RP11-158I13.2 |
| 19 | 4409060 | 4409263 | -2.688837255 | CHAF1A | 2 | 27627402 | 27627595 | -2.429565089 | CCDC121 |
| 19 | 4422499 | 4422698 | -3.074560413 | CHAF1A | 2 | 27653497 | 27653636 | -1.126097587 | SUPT7L |
| 1 | 107057686 | 107057886 | -2.286742368 | PRMT6 | 2 | 28331149 | 28331264 | -3.715922893 | BRE |
| 19 | 4450182 | 4450331 | -9.008240934 | CTB-50L17.8 | 2 | 28332920 | 28333087 | -5.55033117 | BRE |
| 19 | 4450182 | 4450331 | -9.008240934 | UBXN6 | 1 | 116941802 | 116941934 | -1.852224947 | PTGFRN |
| 19 | 4538066 | 4538267 | -4.078948342 | CTB-50L17.14 | 2 | 30158806 | 30159005 | -2.159612329 | YPEL5 |
| 19 | 4538066 | 4538267 | -4.078948342 | LRG1 | 2 | 30606883 | 30607014 | 5.404482087 | LCLAT1 |
| 19 | 4647344 | 4647542 | -2.48392386 | MYDGF | 2 | 32249920 | 32250119 | -3.769902518 | NLRC4 |
| 19 | 4647344 | 4647542 | -2.48392386 | TNFAIP8L1 | 2 | 32617779 | 32617980 | -2.861674389 | BIRC6 |
| 19 | 4649352 | 4649484 | -7.551643904 | MYDGF | 2 | 33234883 | 33235038 | -6.831556654 | LTBP1 |
| 19 | 4649352 | 4649484 | -7.551643904 | TNFAIP8L1 | 1 | 116961327 | 116961529 | -1.685793097 | PTGFRN |
| 19 | 4710770 | 4710969 | -1.972485935 | DPP9 | 2 | 33251072 | 33251255 | -5.693739433 | LTBP1 |
| 19 | 4713188 | 4713372 | -2.0029976 | DPP9 | 2 | 33558756 | 33558965 | -1.376418322 | AC020594.5 |
| 19 | 4795035 | 4795235 | -3.713751748 | FEM1A | 2 | 33558756 | 33558965 | -1.376418322 | RASGRP3 |
| 19 | 4795035 | 4795235 | -3.713751748 | AC005523.2 | 2 | 33826605 | 33826790 | 2.696946926 | LINC01317 |
| 19 | 4816060 | 4816256 | 1.971749127 | TICAM1 | 2 | 34594529 | 34594728 | 1.44844729 | AC073218.1 |
| 19 | 4961423 | 4961624 | -4.295712907 | UHRF1 | 2 | 36548583 | 36548804 | 1.746464814 | CRIM1 |
| 19 | 5080703 | 5080851 | 5.86735481 | KDM4B | 2 | 36548583 | 36548804 | 1.746464814 | FEZ2 |
| 19 | 5111630 | 5111823 | -7.28486848 | KDM4B | 1 | 116966968 | 116967136 | -3.428511992 | PTGFRN |
| 19 | 5112188 | 5112355 | -6.073602462 | KDM4B | 2 | 36814311 | 36814508 | 3.543731421 | VIT |
| 19 | 5122998 | 5123165 | -7.864491685 | KDM4B | 2 | 37316668 | 37316872 | 1.280779068 | PRKD3 |
| 19 | 5691922 | 5692124 | -1.091653182 | LONP1 | 2 | 37827671 | 37827776 | -5.307354922 | LINC00211 |
| 19 | 5917289 | 5917491 | -1.370821175 | RANBP3 | 2 | 38070907 | 38071150 | 1.707629028 | CYP1B1 |
| 19 | 5991198 | 5991326 | 6.88517709 | CTC-232P5.1 | 1 | 116986875 | 116987073 | -2.803705605 | PTGFRN |
| 19 | 5994084 | 5994284 | 2.253379475 | CTC-232P5.1 | 2 | 38074481 | 38074684 | 2.068540451 | CYP1B1-AS1 |
| 19 | 5994084 | 5994284 | 2.253379475 | RFX2 | 2 | 38074481 | 38074684 | 2.068540451 | CYP1B1 |
| 19 | 6057862 | 6058054 | 1.588195798 | RFX2 | 2 | 38492796 | 38493012 | -1.189733212 | AC016995.3 |
| 19 | 6189461 | 6189564 | -2.490516029 | RFX2 | 2 | 38581452 | 38581652 | 1.230674249 | HNRNPLL |
| 19 | 6189461 | 6189564 | -2.490516029 | ACSBG2 | 2 | 38709822 | 38710089 | 1.257013297 | GALM |
| 19 | 6210962 | 6211243 | -1.769012308 | MLLT1 | 2 | 38781508 | 38781711 | -1.80248508 | GEMIN6 |
| 19 | 6221341 | 6221522 | -4.397217351 | MLLT1 | 2 | 38861478 | 38861673 | -1.151164459 | DHX57 |
| 19 | 6222424 | 6222632 | -1.764616918 | MLLT1 | 1 | 117025579 | 117025778 | -2.08341596 | CD101 |
| 19 | 6441695 | 6441896 | -1.797798219 | SLC25A23 | 1 | 117025579 | 117025778 | -2.08341596 | RP11-27K13.3 |
| 19 | 6583430 | 6583624 | -2.082772791 | CD70 | 2 | 39498141 | 39498357 | -1.105186823 | AC007246.3 |
| 19 | 6593750 | 6593947 | -1.519231407 | CD70 | 2 | 39599434 | 39599654 | -1.999775158 | AC007246.3 |
| 19 | 6593750 | 6593947 | -1.519231407 | RPL7P50 | 2 | 39736679 | 39736879 | -1.450369025 | THUMPD2 |
| 19 | 6664459 | 6664661 | 3.510503498 | TNFSF14 | 2 | 42168861 | 42169061 | 3.183161107 | AC083949.1 |
| 19 | 6994768 | 6994925 | 4.412081605 | ADGRE4P | 2 | 42762682 | 42762882 | 1.275859708 | OXER1 |
| 19 | 7083680 | 7083841 | -1.195602393 | ZNF557 | 1 | 117118299 | 117118526 | -2.28491478 | TRIM45 |
| 19 | 7267500 | 7267694 | -1.694591286 | INSR | 2 | 43675708 | 43675840 | -2.907396627 | PLEKHH2 |
| 19 | 7612378 | 7612698 | -1.81596875 | CAMSAP3 | 2 | 43675708 | 43675840 | -2.907396627 | C1GALT1C1L |
| 19 | 7741613 | 7741746 | 3.050143441 | CD209 | 2 | 45006122 | 45006313 | -1.039147263 | SIX2 |
| 19 | 7744260 | 7744455 | 1.647966443 | CD209 | 2 | 46184853 | 46185006 | -1.64005396 | PRKCE |
| 19 | 7745062 | 7745261 | 2.689689184 | CD209 | 2 | 46512241 | 46512458 | -1.305717356 | ATP6V1E2 |
| 19 | 7982684 | 7982868 | -7.869679354 | ELAVL1 | 2 | 46523268 | 46523403 | -6.037070731 | ATP6V1E2 |
| 19 | 8542800 | 8542968 | -1.052760256 | MYO1F | 2 | 46526574 | 46526761 | -3.514335575 | ATP6V1E2 |
| 1 | 108162060 | 108162261 | -2.093488542 | SLC25A24 | 2 | 46905374 | 46905579 | -1.597901623 | AC016722.4 |

| | | | | |
|---|---|---|---|---|
| 19 | 9343094 | 9343294 | -1.841881549 | ZNF559 |
| 19 | 9343094 | 9343294 | -1.841881549 | ZNF559-ZNF177 |
| 1 | 108162817 | 108162987 | -6.198569903 | SLC25A24 |
| 19 | 9560500 | 9560722 | -1.920918264 | ZNF121 |
| 19 | 9566843 | 9567043 | -1.12576818 | ZNF121 |
| 19 | 9810470 | 9810679 | -1.114407066 | FBXL12 |
| 19 | 9815006 | 9815204 | -1.540844904 | FBXL12 |
| 19 | 10092136 | 10092383 | 2.235569283 | C19orf66 |
| 19 | 10092136 | 10092383 | 2.235569283 | ANGPTL6 |
| 19 | 10224141 | 10224338 | -5.844125155 | DNMT1 |
| 19 | 10224141 | 10224338 | -5.844125155 | S1PR2 |
| 19 | 10270577 | 10270771 | 2.564377906 | CTD-2369P2.5 |
| 19 | 10285245 | 10285524 | 7.632669767 | ICAM1 |
| 19 | 10292081 | 10292254 | 5.951187557 | ICAM5 |
| 19 | 10294540 | 10294743 | 6.25389732 | ICAM5 |
| 19 | 10334825 | 10335014 | -1.244761818 | AC114271.2 |
| 19 | 10334825 | 10335014 | -1.244761818 | ICAM3 |
| 19 | 10335222 | 10335422 | -1.171001609 | AC114271.2 |
| 19 | 10335222 | 10335422 | -1.171001609 | ICAM3 |
| 19 | 10486519 | 10486720 | -1.45325122 | KEAP1 |
| 19 | 10553479 | 10553679 | -3.014945453 | KRI1 |
| 19 | 10643605 | 10643796 | 5.584233346 | SLC44A2 |
| 1 | 108648692 | 108648885 | -1.885690243 | HENMT1 |
| 19 | 10691664 | 10691876 | -1.250952527 | ILF3 |
| 19 | 10833062 | 10833337 | -1.582187817 | DNM2 |
| 19 | 10833062 | 10833337 | -1.582187817 | TMED1 |
| 19 | 10869371 | 10869557 | -2.328349127 | C19orf38 |
| 19 | 10929662 | 10929872 | -1.069158721 | C19orf52 |
| 19 | 11146058 | 11146266 | -3.703898642 | SPC24 |
| 19 | 11176690 | 11176889 | -8.230643793 | KANK2 |
| 19 | 11193934 | 11194132 | -1.257853016 | KANK2 |
| 1 | 2563511 | 2563711 | 3.343333743 | TNFRSF14 |
| 19 | 11325061 | 11325258 | -2.563711272 | RAB3D |
| 19 | 11325061 | 11325258 | -2.563711272 | TSPAN16 |
| 19 | 11375797 | 11375999 | -1.316471265 | SWSAP1 |
| 19 | 11378026 | 11378167 | -1.581072746 | EPOR |
| 19 | 11617939 | 11618104 | -2.103831359 | ZNF627 |
| 19 | 11652686 | 11652887 | -3.402402283 | CTC-499B15.5 |
| 19 | 11652686 | 11652887 | -3.402402283 | ZNF887P |
| 1 | 108929691 | 108929889 | -2.71387775 | GPSM2 |
| 1 | 108929691 | 108929889 | -2.71387775 | RP11-475E11.9 |
| 1 | 108929691 | 108929889 | -2.71387775 | CLCC1 |
| 19 | 11655780 | 11655944 | -7.127735614 | CTC-499B15.5 |
| 19 | 11659608 | 11659738 | -3.756791083 | CTC-499B15.5 |
| 19 | 11667138 | 11667329 | -3.582381648 | CTC-499B15.5 |
| 19 | 11673783 | 11673931 | -4.062649752 | CTC-499B15.5 |
| 19 | 11680442 | 11680636 | -1.232440712 | CTC-499B15.5 |
| 19 | 11721618 | 11721818 | -1.398998541 | ZNF823 |
| 19 | 11722649 | 11722921 | -2.220468116 | ZNF823 |
| 19 | 11833568 | 11833763 | -1.17371914 | ZNF440 |
| 19 | 11867408 | 11867613 | -2.236766927 | ZNF439 |
| 19 | 11869028 | 11869228 | -2.379531046 | ZNF439 |
| 19 | 11904741 | 11904948 | -1.959622914 | ZNF69 |
| 19 | 11908656 | 11908762 | -3.764408841 | ZNF69 |
| 2 | 46905374 | 46905579 | -1.597901623 | MCFD2 |
| 2 | 47800789 | 47800990 | -1.01129308 | MSH6 |
| 2 | 47800789 | 47800990 | -1.01129308 | FBXO11 |
| 1 | 117522930 | 117523189 | -1.031720109 | MAN1A2 |
| 2 | 54051036 | 54051260 | -1.046549702 | ACYP2 |
| 2 | 56173076 | 56173250 | -1.554262228 | RP11-482H16.1 |
| 2 | 56173076 | 56173250 | -1.554262228 | RP11-481J13.1 |
| 2 | 58170650 | 58170816 | 3.148681227 | FANCL |
| 2 | 58173748 | 58173947 | 4.911000849 | FANCL |
| 2 | 58177245 | 58177429 | 7.914365766 | FANCL |
| 2 | 60922119 | 60922323 | 2.177112375 | REL |
| 2 | 60922567 | 60922743 | 1.805477759 | REL |
| 2 | 60929204 | 60929404 | 2.206647839 | RP11-373L24.1 |
| 2 | 60929204 | 60929404 | 2.206647839 | REL |
| 2 | 61031742 | 61031923 | -1.481455557 | PEX13 |
| 2 | 61039539 | 61039730 | -1.967657337 | PEX13 |
| 2 | 61040304 | 61040496 | -1.530951028 | PEX13 |
| 2 | 61048506 | 61048699 | -1.670945343 | PEX13 |
| 2 | 61188123 | 61188331 | -1.204550888 | USP34 |
| 2 | 61188123 | 61188331 | -1.204550888 | AHSA2 |
| 2 | 61190542 | 61190718 | -1.06911765 | AHSA2 |
| 2 | 61190542 | 61190718 | -1.06911765 | USP34 |
| 2 | 62840966 | 62841144 | -3.564641508 | AC007098.1 |
| 2 | 62840966 | 62841144 | -3.564641508 | EHBP1 |
| 2 | 62948338 | 62948511 | -6.877026184 | EHBP1 |
| 2 | 63212600 | 63212787 | -1.058058171 | WDPCP |
| 2 | 63404250 | 63404466 | -2.075606919 | WDPCP |
| 2 | 63564440 | 63564659 | -1.367014151 | WDPCP |
| 2 | 63588439 | 63588643 | -1.457621338 | MDH1 |
| 2 | 63588439 | 63588643 | -1.457621338 | WDPCP |
| 2 | 64094987 | 64095192 | 1.235418242 | PELI1 |
| 1 | 118883895 | 118884111 | -3.100773601 | TBX15 |
| 2 | 64458487 | 64458682 | -2.424881916 | LGALSL |
| 2 | 64523165 | 64523354 | -1.27366889 | RP11-568N6.1 |
| 2 | 65021210 | 65021411 | -1.129841554 | SLC1A4 |
| 2 | 65072265 | 65072468 | -2.232832476 | RAB1A |
| 2 | 65072265 | 65072468 | -2.232832476 | CEP68 |
| 2 | 67403519 | 67403792 | -1.10200247 | ETAA1 |
| 2 | 68130829 | 68131026 | -6.301929061 | WDR92 |
| 2 | 68464979 | 68465180 | -1.481869165 | FBXO48 |
| 2 | 68761973 | 68762173 | 1.889839557 | ARHGAP25 |
| 2 | 69245812 | 69245946 | -1.940208184 | ANTXR1 |
| 2 | 69458201 | 69458459 | -1.837568069 | AAK1 |
| 2 | 69465026 | 69465233 | -1.712083144 | AAK1 |
| 2 | 69466222 | 69466431 | -1.085102135 | AAK1 |
| 2 | 69654957 | 69655150 | -2.667317767 | ANXA4 |
| 2 | 69654957 | 69655150 | -2.667317767 | AAK1 |
| 2 | 69937997 | 69938204 | -1.057925791 | MXD1 |
| 2 | 70277228 | 70277425 | -2.027780441 | PCYOX1 |
| 2 | 71423239 | 71423460 | -1.08106519 | ZNF638 |
| 2 | 71426486 | 71426660 | -1.50928752 | ZNF638 |
| 2 | 72891387 | 72891528 | -1.034438889 | SPR |
| 2 | 73080847 | 73081046 | -1.54377761 | RAB11FIP5 |
| 1 | 119212400 | 119212677 | -6.174087434 | RP11-418J17.1 |

| | | | | |
|---|---|---|---|---|
| 19 | 11909610 | 11909787 | -1.345221702 | ZNF69 |
| 19 | 11910647 | 11910818 | -2.872291918 | ZNF69 |
| 19 | 11911435 | 11911659 | -3.307523804 | ZNF69 |
| 1 | 109089183 | 109089414 | -1.346459117 | RP5-1065J22.8 |
| 19 | 11912530 | 11912719 | -1.61059802 | ZNF69 |
| 19 | 11948438 | 11948619 | -1.157836847 | CTD-2006C1.13 |
| 19 | 11948438 | 11948619 | -1.157836847 | ZNF700 |
| 19 | 11948438 | 11948619 | -1.157836847 | CTD-2006C1.12 |
| 19 | 11950170 | 11950373 | -1.475926967 | ZNF700 |
| 19 | 11950170 | 11950373 | -1.475926967 | CTD-2006C1.12 |
| 19 | 11950170 | 11950373 | -1.475926967 | CTD-2006C1.13 |
| 19 | 11978145 | 11978344 | 1.778024904 | CTD-2006C1.13 |
| 19 | 11978145 | 11978344 | 1.778024904 | ZNF763 |
| 19 | 11979320 | 11979516 | -1.54482899 | ZNF763 |
| 19 | 12005141 | 12005343 | 2.093762643 | CTD-2006C1.2 |
| 19 | 12043769 | 12043888 | -8.188206855 | CTD-2006C1.10 |
| 19 | 12043769 | 12043888 | -8.188206855 | ZNF878 |
| 19 | 12043769 | 12043888 | -8.188206855 | CTD-2006C1.2 |
| 19 | 12111343 | 12111548 | -2.442878676 | ZNF20 |
| 19 | 12111343 | 12111548 | -2.442878676 | ZNF788 |
| 19 | 12113330 | 12113591 | -2.285694773 | ZNF788 |
| 19 | 12113330 | 12113591 | -2.285694773 | ZNF20 |
| 19 | 12132391 | 12132593 | -1.364896838 | ZNF625-ZNF20 |
| 19 | 12132391 | 12132593 | -1.364896838 | ZNF788 |
| 19 | 12132391 | 12132593 | -1.364896838 | ZNF20 |
| 19 | 12140322 | 12140495 | -2.444248222 | ZNF20 |
| 19 | 12140322 | 12140495 | -2.444248222 | ZNF625-ZNF20 |
| 19 | 12272327 | 12272528 | -1.777355224 | ZNF44 |
| 19 | 12273009 | 12273125 | -1.616305792 | ZNF44 |
| 19 | 12273592 | 12273698 | -2.294880496 | ZNF44 |
| 19 | 12349700 | 12349900 | -2.769907838 | ZNF442 |
| 19 | 12350635 | 12350836 | -1.912625644 | ZNF442 |
| 19 | 12380960 | 12381122 | -1.259933736 | CTD-3105H18.14 |
| 19 | 12380960 | 12381122 | -1.259933736 | CTD-3105H18.4 |
| 19 | 12390475 | 12390725 | -1.542668306 | CTD-3105H18.14 |
| 19 | 12390475 | 12390725 | -1.542668306 | ZNF799 |
| 19 | 12430165 | 12430361 | -2.760652042 | CTD-3105H18.16 |
| 19 | 12430165 | 12430361 | -2.760652042 | ZNF443 |
| 19 | 12430774 | 12430883 | -2.006845856 | CTD-3105H18.16 |
| 19 | 12430774 | 12430883 | -2.006845856 | ZNF443 |
| 19 | 12465262 | 12465464 | -2.812772156 | CTD-2192J16.17 |
| 19 | 12465262 | 12465464 | -2.812772156 | ZNF709 |
| 19 | 12465262 | 12465464 | -2.812772156 | CTD-3105H18.18 |
| 19 | 12526021 | 12526215 | -1.848512608 | ZNF564 |
| 19 | 12526021 | 12526215 | -1.848512608 | CTD-2192J16.20 |
| 19 | 12526021 | 12526215 | -1.848512608 | CTD-2192J16.17 |
| 19 | 12577453 | 12577675 | -2.155887693 | CTD-2192J16.20 |
| 19 | 12792045 | 12792247 | 1.316763654 | HOOK2 |
| 19 | 12792045 | 12792247 | 1.316763654 | JUNB |
| 1 | 109251295 | 109251477 | -2.222264398 | CELSR2 |
| 19 | 12792715 | 12792923 | 1.983753114 | HOOK2 |
| 19 | 12792715 | 12792923 | 1.983753114 | JUNB |
| 19 | 12875722 | 12875938 | 2.006972844 | DNASE2 |
| 19 | 12894124 | 12894320 | -2.732391796 | RPS6P25 |
| 2 | 73258729 | 73258934 | -1.828616827 | FBXO41 |
| 1 | 119596479 | 119596673 | 1.06306614 | GAPDHP33 |
| 1 | 119623793 | 119623970 | 1.149483704 | ZNF697 |
| 2 | 74418205 | 74418357 | -5.190771476 | C2orf81 |
| 2 | 74418205 | 74418357 | -5.190771476 | HMGA1P8 |
| 2 | 74461278 | 74461484 | -1.093300371 | MOGS |
| 2 | 74502718 | 74502918 | -1.088540012 | LBX2 |
| 2 | 74502718 | 74502918 | -1.088540012 | LBX2-AS1 |
| 2 | 74506814 | 74507003 | -8.105335672 | PCGF1 |
| 2 | 74556991 | 74557205 | -1.380003871 | DOK1 |
| 2 | 74930666 | 74930846 | -1.559897785 | LINC01291 |
| 2 | 84870274 | 84870473 | 2.642701109 | TRABD2A |
| 2 | 85607886 | 85608115 | -1.315206721 | C2orf68 |
| 2 | 85607886 | 85608115 | -1.315206721 | USP39 |
| 2 | 85840100 | 85840296 | -1.336994058 | ST3GAL5 |
| 2 | 85848106 | 85848213 | -1.689004793 | ST3GAL5 |
| 2 | 86195752 | 86195912 | 3.245926548 | RP11-301O19.1 |
| 2 | 86195752 | 86195912 | 3.245926548 | IMMT |
| 1 | 119916460 | 119916657 | -1.512996433 | NOTCH2 |
| 2 | 86604257 | 86604440 | -1.585012989 | AC015971.2 |
| 2 | 86604257 | 86604440 | -1.585012989 | RNF103-CHMP3 |
| 2 | 86604257 | 86604440 | -1.585012989 | RNF103 |
| 2 | 86774695 | 86774903 | -2.282212826 | RMND5A |
| 2 | 87583420 | 87583548 | 1.442826428 | LINC00152 |
| 2 | 87584752 | 87585031 | 1.741280252 | LINC00152 |
| 1 | 120433844 | 120434027 | 2.620645403 | PFN1P2 |
| 2 | 95149724 | 95149927 | -2.388660377 | ZNF514 |
| 2 | 95158931 | 95159086 | -4.082453463 | ZNF514 |
| 2 | 95181222 | 95181420 | -1.943066018 | ZNF2 |
| 2 | 95214478 | 95214710 | -3.451351399 | AC092835.2 |
| 2 | 95215154 | 95215386 | -1.92429502 | AC092835.2 |
| 2 | 95414095 | 95414291 | -1.46686105 | FAHD2A |
| 2 | 95414095 | 95414291 | -1.46686105 | AC009238.6 |
| 1 | 120454378 | 120454537 | 3.444137459 | NBPF8 |
| 2 | 95838958 | 95839128 | -7.002110233 | ANKRD36C |
| 1 | 120455825 | 120456007 | 7.15260338 | NBPF8 |
| 2 | 96022908 | 96023127 | -1.427044491 | FAHD2CP |
| 2 | 96022908 | 96023127 | -1.427044491 | GPAT2 |
| 2 | 96370814 | 96370992 | -1.629155998 | NCAPH |
| 2 | 96552339 | 96552538 | 1.052466123 | ARID5A |
| 1 | 120466197 | 120466368 | -1.967033928 | NBPF8 |
| 2 | 96810090 | 96810306 | -1.418208694 | CNNM4 |
| 2 | 96817188 | 96817404 | -1.256595387 | CNNM3 |
| 2 | 96824939 | 96825107 | -2.310649379 | CNNM3 |
| 2 | 96824939 | 96825107 | -2.310649379 | ANKRD23 |
| 2 | 96828458 | 96828665 | -3.040869738 | ANKRD23 |
| 2 | 96828458 | 96828665 | -3.040869738 | CNNM3 |
| 2 | 96860470 | 96860670 | -2.496469848 | SEMA4C |
| 2 | 97245384 | 97245558 | -2.211529708 | ANKRD36 |
| 2 | 97425925 | 97426123 | -2.146967953 | AC159540.2 |
| 2 | 97425925 | 97426123 | -2.146967953 | AC159540.1 |
| 2 | 97432736 | 97432932 | -4.874096425 | AC159540.2 |
| 2 | 97432736 | 97432932 | -4.874096425 | AC159540.1 |
| 2 | 97511675 | 97511874 | -2.847977028 | ANKRD36B |

| | | | | |
|---|---|---|---|---|
| 19 | 12894124 | 12894320 | -2.732391796 | GCDH |
| 19 | 12899522 | 12899723 | -2.432847646 | SYCE2 |
| 19 | 12899522 | 12899723 | -2.432847646 | GCDH |
| 19 | 12943614 | 12943814 | -14.29791205 | CALR |
| 19 | 13207007 | 13207215 | 3.635033959 | CACNA1A |
| 19 | 13242624 | 13242831 | 3.54518557 | CACNA1A |
| 19 | 13762936 | 13763137 | -1.233826792 | CCDC130 |
| 19 | 13830188 | 13830372 | 2.894527053 | ZSWIM4 |
| 19 | 13830890 | 13831090 | 2.604273243 | ZSWIM4 |
| 19 | 14149787 | 14149970 | -7.280942284 | ADGRL1 |
| 19 | 14149787 | 14149970 | -7.280942284 | CTB-55O6.12 |
| 19 | 14151133 | 14151339 | 1.146685242 | CTB-55O6.12 |
| 19 | 14151133 | 14151339 | 1.146685242 | ADGRL1 |
| 19 | 14158114 | 14158326 | -1.146719348 | ADGRL1 |
| 19 | 14158114 | 14158326 | -1.146719348 | CTB-55O6.12 |
| 1 | 109279889 | 109280088 | -3.573908666 | PSRC1 |
| 19 | 14394244 | 14394436 | -2.646868454 | ADGRE5 |
| 19 | 14395211 | 14395410 | -2.545616871 | ADGRE5 |
| 19 | 14411264 | 14411456 | 7.10712161 | DDX39A |
| 19 | 14412645 | 14412843 | -7.854595768 | DDX39A |
| 19 | 14515814 | 14516017 | -1.208821887 | DNAJB1 |
| 19 | 14708898 | 14709064 | 8.489958186 | ZNF333 |
| 1 | 3463377 | 3463528 | -1.333878271 | ARHGEF16 |
| 19 | 14735914 | 14736119 | 3.679551743 | ADGRE2 |
| 19 | 14767992 | 14768190 | 1.894757354 | ADGRE2 |
| 19 | 16124004 | 16124200 | -11.67087437 | RAB8A |
| 19 | 16124004 | 16124200 | -11.67087437 | CTD-2231E14.8 |
| 19 | 16133137 | 16133338 | -1.703859903 | RAB8A |
| 19 | 16133137 | 16133338 | -1.703859903 | CTD-2231E14.8 |
| 19 | 16157381 | 16157505 | -2.982550896 | HSH2D |
| 19 | 16191384 | 16191585 | -1.447276848 | FAM32A |
| 19 | 16236204 | 16236414 | -1.349790891 | AP1M1 |
| 19 | 16236751 | 16236881 | -2.178321825 | AP1M1 |
| 19 | 16327091 | 16327286 | -9.697698032 | KLF2 |
| 19 | 16509934 | 16510037 | -1.619314034 | CTD-3222D19.2 |
| 19 | 16509934 | 16510037 | -1.619314034 | C19orf44 |
| 19 | 16519088 | 16519288 | -1.693628629 | C19orf44 |
| 19 | 16519088 | 16519288 | -1.693628629 | CTD-3222D19.2 |
| 19 | 16519088 | 16519288 | -1.693628629 | CHERP |
| 1 | 109508006 | 109508201 | -1.209320277 | AMIGO1 |
| 19 | 16652532 | 16652737 | -1.635946018 | CTC-429P9.4 |
| 19 | 16652532 | 16652737 | -1.635946018 | SMIM7 |
| 19 | 16890991 | 16891185 | 1.178657762 | F2RL3 |
| 19 | 16891614 | 16891857 | 1.692351649 | F2RL3 |
| 19 | 16893648 | 16893858 | 5.307145408 | CPAMD8 |
| 19 | 17050043 | 17050220 | -2.084552261 | HAUS8 |
| 19 | 17231981 | 17232180 | -1.09007473 | NR2F6 |
| 19 | 17283435 | 17283635 | -4.521217633 | ANKLE1 |
| 19 | 17305372 | 17305492 | -1.63339327 | ABHD8 |
| 19 | 17305372 | 17305492 | -1.63339327 | MRPL34 |
| 19 | 17306400 | 17306592 | -1.583197718 | MRPL34 |
| 19 | 17306400 | 17306592 | -1.583197718 | ABHD8 |
| 19 | 17319529 | 17319743 | -2.358946647 | DDA1 |
| 1 | 109665204 | 109665328 | -1.361664056 | GSTM4 |

| | | | | |
|---|---|---|---|---|
| 1 | 120579636 | 120579825 | 7.294298764 | CH17-472G23.4 |
| 2 | 97663317 | 97663518 | 5.535731181 | ACTR1B |
| 2 | 97678200 | 97678308 | -2.672945669 | LINC01125 |
| 2 | 97702615 | 97702909 | -1.535037697 | LINC01125 |
| 2 | 97792623 | 97792822 | 1.017331195 | TMEM131 |
| 2 | 98570095 | 98570302 | -2.366745639 | INPP4A |
| 2 | 98582789 | 98582954 | -1.929608702 | INPP4A |
| 2 | 98590234 | 98590434 | -3.054132618 | INPP4A |
| 1 | 120802326 | 120802469 | 1.687849436 | NBPF26 |
| 2 | 98794302 | 98794522 | -6.544394119 | KIAA1211L |
| 2 | 99125117 | 99125298 | -1.842249087 | TSGA10 |
| 2 | 99126278 | 99126478 | -2.153800657 | TSGA10 |
| 2 | 99201500 | 99201728 | -6.604726683 | RP11-111H13.1 |
| 2 | 99551307 | 99551490 | -6.212476786 | AFF3 |
| 2 | 100392056 | 100392252 | -7.237866715 | CHST10 |
| 2 | 100392691 | 100392910 | -2.279220841 | CHST10 |
| 2 | 100393222 | 100393377 | -1.858457146 | CHST10 |
| 2 | 101007834 | 101008036 | -1.668871484 | RPL31 |
| 2 | 101007834 | 101008036 | -1.668871484 | TBC1D8 |
| 2 | 101068102 | 101068297 | -1.704393183 | TBC1D8 |
| 2 | 101152477 | 101152649 | -1.413375808 | TBC1D8 |
| 2 | 101152477 | 101152649 | -1.413375808 | RP11-299H21.1 |
| 2 | 101457266 | 101457476 | -2.161812193 | RFX8 |
| 2 | 102718749 | 102719065 | -1.928270595 | MFSD9 |
| 2 | 105881675 | 105881868 | -1.019063312 | NCK2 |
| 2 | 105892979 | 105893102 | -1.830808641 | NCK2 |
| 2 | 106160177 | 106160373 | -6.529334125 | UXS1 |
| 2 | 106424789 | 106424992 | -1.672222695 | RGPD3 |
| 2 | 108471034 | 108471234 | -1.582656448 | GCC2 |
| 2 | 108499151 | 108499346 | -5.357122759 | GCC2 |
| 2 | 108505479 | 108505593 | -6.613807972 | GCC2 |
| 2 | 108763602 | 108763798 | -1.680862953 | RANBP2 |
| 2 | 108765049 | 108765255 | -1.557265448 | RANBP2 |
| 2 | 108765759 | 108765959 | -1.513910728 | RANBP2 |
| 1 | 121096108 | 121096309 | 3.924460478 | RP11-439A17.10 |
| 1 | 121096108 | 121096309 | 3.924460478 | RP11-439A17.9 |
| 1 | 121096108 | 121096309 | 3.924460478 | FCGR1B |
| 2 | 108766455 | 108766662 | -1.493115869 | RANBP2 |
| 2 | 108767621 | 108767822 | -1.419518071 | RANBP2 |
| 2 | 108768108 | 108768297 | -1.607966284 | RANBP2 |
| 2 | 108771579 | 108771779 | -1.455845791 | RANBP2 |
| 2 | 108783519 | 108783715 | -2.510420754 | RANBP2 |
| 2 | 111123750 | 111123872 | -1.410666195 | BCL2L11 |
| 1 | 121097963 | 121098167 | 2.699256455 | RP11-439A17.10 |
| 2 | 111292416 | 111292612 | 1.769531227 | MIR4435-2HG |
| 2 | 111323487 | 111323682 | 2.308105034 | RP11-803D5.1 |
| 2 | 111323487 | 111323682 | 2.308105034 | MIR4435-2HG |
| 2 | 112028622 | 112028891 | -2.668124137 | MERTK |
| 1 | 6016571 | 6016765 | -6.076459288 | KCNAB2 |
| 1 | 121184041 | 121184252 | 2.043258083 | FAM72B |
| 2 | 112389852 | 112390004 | -6.377682252 | RGPD8 |
| 2 | 112538962 | 112539153 | -1.408190045 | TTL |
| 2 | 112659352 | 112659556 | -2.741815759 | SLC20A1 |
| 2 | 112736670 | 112736840 | -2.494676378 | NT5DC4 |

| | | | | | | | | | |
|---|---|---|---|---|---|---|---|---|---|
| 19 | 17341654 | 17341778 | -1.807215699 | GTPBP3 | 2 | 112736670 | 112736840 | -2.494676378 | CKAP2L |
| 19 | 17504586 | 17504785 | 7.54290219 | SLC27A1 | 2 | 112737116 | 112737311 | -5.667974283 | NT5DC4 |
| 19 | 17504586 | 17504785 | 7.54290219 | CTD-3131K8.2 | 2 | 112737116 | 112737311 | -5.667974283 | CKAP2L |
| 19 | 17581312 | 17581547 | -1.349353302 | COLGALT1 | 2 | 112752381 | 112752586 | -3.407714522 | CKAP2L |
| 19 | 17726789 | 17727028 | 1.218312172 | MAP1S | 2 | 112756055 | 112756279 | -2.04966844 | CKAP2L |
| 19 | 18013666 | 18013866 | -1.142698677 | ARRDC2 | 1 | 121184701 | 121184871 | 1.8148692 | SRGAP2C |
| 19 | 18059190 | 18059429 | 1.809222893 | IL12RB1 | 1 | 121184701 | 121184871 | 1.8148692 | FAM72B |
| 19 | 18264412 | 18264571 | 1.821986383 | KIAA1683 | 2 | 113113188 | 113113370 | 6.245258324 | IL1RN |
| 1 | 109896285 | 109896467 | -1.416445007 | RP11-195M16.3 | 2 | 113132873 | 113133073 | 11.90925798 | IL1RN |
| 19 | 18265051 | 18265211 | 3.839533034 | KIAA1683 | 2 | 113201508 | 113201708 | -1.213915792 | PSD4 |
| 19 | 18266360 | 18266563 | 2.828563727 | KIAA1683 | 2 | 113218176 | 113218385 | -2.303428595 | PAX8-AS1 |
| 19 | 18363468 | 18363669 | 1.709562827 | PGPEP1 | 2 | 113218176 | 113218385 | -2.303428595 | PAX8 |
| 19 | 18388750 | 18388951 | 9.678951978 | GDF15 | 2 | 113489561 | 113489761 | 3.56156266 | CBWD2 |
| 19 | 18397035 | 18397154 | 7.959687734 | LRRC25 | 2 | 113498909 | 113499048 | 3.740712862 | FOXD4L1 |
| 1 | 109923427 | 109923619 | 11.35754974 | CSF1 | 2 | 113598740 | 113598938 | -1.088407069 | WASH2P |
| 19 | 18919993 | 18920164 | 7.719477125 | DDX49 | 2 | 114000281 | 114000479 | 1.959548969 | LINC01191 |
| 19 | 18919993 | 18920164 | 7.719477125 | AC002985.3 | 2 | 119245499 | 119245727 | -2.583020783 | STEAP3-AS1 |
| 19 | 19009979 | 19010175 | -1.964905626 | SUGP2 | 2 | 119245499 | 119245727 | -2.583020783 | STEAP3 |
| 19 | 19024758 | 19024959 | -2.506607933 | SUGP2 | 2 | 119263342 | 119263537 | -2.826904832 | STEAP3 |
| 19 | 19025720 | 19025920 | -2.394995917 | SUGP2 | 2 | 119681779 | 119681983 | -2.137436171 | TMEM177 |
| 19 | 19177022 | 19177222 | -1.404958936 | BORCS8-MEF2B | 1 | 121406335 | 121406489 | 1.718678265 | CH17-437K3.1 |
| 19 | 19177022 | 19177222 | -1.404958936 | BORCS8 | 2 | 121650888 | 121651081 | -1.307485271 | NIFK-AS1 |
| 19 | 19257491 | 19257687 | 2.129933563 | AC138430.4 | 2 | 127307626 | 127307810 | -1.815479854 | MAP3K2 |
| 19 | 19257491 | 19257687 | 2.129933563 | HAPLN4 | 2 | 127428715 | 127428919 | -3.982315979 | PROC |
| 19 | 19333403 | 19333655 | -1.152495756 | MAU2 | 2 | 127702293 | 127702494 | -1.05558071 | SFT2D3 |
| 19 | 19508456 | 19508662 | -1.705531752 | GATAD2A | 2 | 127702293 | 127702494 | -1.05558071 | WDR33 |
| 19 | 19514171 | 19514291 | -3.332755692 | TSSK6 | 2 | 127708667 | 127708859 | -1.324979216 | WDR33 |
| 19 | 19679273 | 19679481 | -1.318869579 | ZNF101 | 2 | 127717208 | 127717408 | -1.116401102 | WDR33 |
| 19 | 19681184 | 19681380 | -2.672704123 | ZNF101 | 2 | 128265614 | 128265815 | -1.555083391 | HS6ST1 |
| 1 | 109929120 | 109929336 | 9.525488063 | CSF1 | 2 | 128268031 | 128268245 | -1.063620559 | HS6ST1 |
| 19 | 19935060 | 19935231 | -3.662051591 | AC007204.2 | 2 | 130373464 | 130373738 | -1.00954342 | PTPN18 |
| 19 | 19935060 | 19935231 | -3.662051591 | ZNF93 | 2 | 131520765 | 131521061 | -1.462718153 | AC093838.4 |
| 19 | 20006409 | 20006635 | -2.023473041 | ZNF682 | 2 | 135202854 | 135203042 | -5.5 | ZRANB3 |
| 19 | 20164722 | 20164890 | -7.100449303 | CTC-260E6.6 | 2 | 135207553 | 135207698 | -2.405309796 | ZRANB3 |
| 19 | 20226263 | 20226484 | -4.362615536 | CTC-260E6.6 | 2 | 135481571 | 135481797 | -3.052897332 | ZRANB3 |
| 19 | 20227444 | 20227641 | -3.356364654 | CTC-260E6.6 | 1 | 143883527 | 143883717 | 3.297569916 | FCGR1CP |
| 19 | 20805896 | 20806162 | -1.809918595 | ZNF66 | 1 | 143883527 | 143883717 | 3.297569916 | RP5-998N21.4 |
| 19 | 20843182 | 20843382 | -1.990614385 | CTD-2542C24.3 | 1 | 143883527 | 143883717 | 3.297569916 | HIST2H2BB |
| 19 | 20950256 | 20950454 | -2.324605314 | ZNF85 | 2 | 136115539 | 136115744 | -2.177142977 | CXCR4 |
| 19 | 21116872 | 21117023 | 1.057336408 | ZNF714 | 2 | 138013842 | 138014041 | -2.256084237 | HNMT |
| 19 | 21183852 | 21184053 | -1.413531402 | ZNF431 | 1 | 143904447 | 143904623 | -8.504126764 | RP5-998N21.10 |
| 19 | 21536204 | 21536403 | -1.315674079 | ZNF429 | 1 | 143904447 | 143904623 | -8.504126764 | HIST2H2BB |
| 19 | 21537534 | 21537647 | -1.2637816 | ZNF429 | 2 | 143006244 | 143006432 | 8.336571708 | KYNU |
| 19 | 21726611 | 21726811 | -1.449556034 | ZNF100 | 2 | 143008097 | 143008300 | 1.592723853 | KYNU |
| 19 | 21727635 | 21727834 | -1.260657296 | ZNF100 | 2 | 143010996 | 143011096 | 1.759773435 | KYNU |
| 19 | 21809593 | 21809819 | -1.544871082 | ZNF43 | 1 | 6065562 | 6065819 | -10.22427999 | KCNAB2 |
| 19 | 22391611 | 22391756 | -1.211360358 | ZNF98 | 1 | 143905566 | 143905756 | -3.225932306 | RP5-998N21.10 |
| 19 | 22677176 | 22677373 | -4.660964047 | CTC-457E21.9 | 1 | 143905566 | 143905756 | -3.225932306 | HIST2H3PS2 |
| 19 | 22677919 | 22678112 | -4.412876916 | CTC-457E21.9 | 1 | 143905566 | 143905756 | -3.225932306 | RP5-998N21.7 |
| 19 | 23101316 | 23101501 | -3.978294644 | ZNF730 | 2 | 143663607 | 143663807 | -3.792950725 | ARHGAP15 |
| 19 | 23146773 | 23146971 | -2.754679799 | ZNF730 | 2 | 143666998 | 143667247 | -1.006689515 | ARHGAP15 |
| 19 | 23222307 | 23222511 | -3.398002393 | ZNF724 | 2 | 144389555 | 144389763 | -1.109194661 | ZEB2 |
| 19 | 23223238 | 23223434 | -3.366390296 | RP11-15H20.8 | 2 | 144399985 | 144400174 | -1.311179172 | ZEB2 |
| 19 | 23223238 | 23223434 | -3.366390296 | ZNF724 | 2 | 144790234 | 144790420 | -2.982892142 | TEX41 |

| | | | | |
|---|---|---|---|---|
| 19 | 23223728 | 23223925 | -2.387569596 | ZNF724 |
| 19 | 23223728 | 23223925 | -2.387569596 | RP11-15H20.8 |
| 19 | 23259635 | 23259794 | -2.546348713 | RP11-15H20.7 |
| 19 | 23259635 | 23259794 | -2.546348713 | RP11-15H20.8 |
| 19 | 23653279 | 23653493 | -1.335616286 | ZNF675 |
| 19 | 24126327 | 24126525 | -1.054459363 | ZNF254 |
| 19 | 24127823 | 24128131 | -1.077481366 | ZNF254 |
| 1 | 110341007 | 110341207 | -1.059814413 | RBM15 |
| 19 | 27797797 | 27797997 | -2.165931017 | LLNLF-65H9.1 |
| 19 | 27797797 | 27797997 | -2.165931017 | CTC-459F4.3 |
| 19 | 27805696 | 27805943 | -1.917410884 | LLNLF-65H9.1 |
| 19 | 27805696 | 27805943 | -1.917410884 | CTC-459F4.3 |
| 19 | 27805696 | 27805943 | -1.917410884 | SLC25A1P5 |
| 19 | 27820272 | 27820381 | -2.059664771 | LLNLF-65H9.1 |
| 19 | 27820272 | 27820381 | -2.059664771 | CTC-459F4.3 |
| 19 | 29616814 | 29617013 | -1.469572397 | POP4 |
| 19 | 29674919 | 29675226 | 2.131987929 | PLEKHF1 |
| 19 | 31279485 | 31279627 | -3.705834538 | TSHZ3 |
| 19 | 32353987 | 32354225 | -1.666656838 | ZNF507 |
| 19 | 32598164 | 32598368 | -1.485292316 | ANKRD27 |
| 19 | 32969170 | 32969344 | -1.374552471 | CEP89 |
| 19 | 32970105 | 32970296 | -1.842032977 | CEP89 |
| 19 | 32976860 | 32977015 | -2.48927579 | FAAP24 |
| 1 | 110347491 | 110347706 | -1.85390021 | LAMTOR5-AS1 |
| 19 | 33301278 | 33301430 | -1.279376114 | CTD-2540B15.7 |
| 19 | 33301278 | 33301430 | -1.279376114 | CEBPA |
| 19 | 33304624 | 33304827 | -1.247434736 | CEBPA-AS1 |
| 19 | 33399325 | 33399465 | -6.972081493 | PEPD |
| 19 | 33812907 | 33813104 | -1.764944824 | KCTD15 |
| 19 | 34341697 | 34341851 | 1.832461 | KIAA0355 |
| 19 | 34684663 | 34684783 | -1.173799677 | ZNF302 |
| 19 | 35543177 | 35543369 | -7.455906637 | TMEM147-AS1 |
| 19 | 35543177 | 35543369 | -7.455906637 | GAPDHS |
| 19 | 35543982 | 35544182 | -4.690298173 | GAPDHS |
| 19 | 35543982 | 35544182 | -4.690298173 | TMEM147-AS1 |
| 19 | 35545032 | 35545228 | -1.199412335 | GAPDHS |
| 19 | 35545032 | 35545228 | -1.199412335 | TMEM147-AS1 |
| 19 | 35708162 | 35708398 | 7.364863473 | ZBTB32 |
| 19 | 35721483 | 35721683 | -8.548604572 | KMT2B |
| 19 | 36337080 | 36337236 | -1.838192424 | ZFP14 |
| 19 | 36497411 | 36497608 | 2.617318009 | CTBP2P7 |
| 1 | 110604463 | 110604660 | 6.544642475 | KCNA2 |
| 19 | 36626069 | 36626293 | 1.092846159 | ZNF382 |
| 19 | 36626913 | 36627352 | 1.189299939 | ZNF382 |
| 19 | 36638832 | 36638977 | -1.349240454 | ZNF461 |
| 19 | 36747983 | 36748182 | -2.80756521 | ZNF850 |
| 19 | 36750463 | 36750599 | -2.879391496 | ZNF850 |
| 19 | 37242641 | 37242814 | -1.367457756 | ZNF383 |
| 19 | 37362663 | 37362803 | -1.4654597 | HKR1 |
| 19 | 37389408 | 37389529 | -1.09383095 | ZNF527 |
| 19 | 38396132 | 38396357 | 2.697818891 | SPRED3 |
| 19 | 38910658 | 38910858 | -2.323281703 | CCER2 |
| 19 | 38932735 | 38932952 | -2.618599958 | CTC-360G5.8 |
| 19 | 38932735 | 38932952 | -2.618599958 | MRPS12 |
| 2 | 147927190 | 147927392 | 1.41801611 | ACVR2A |
| 2 | 148088997 | 148089165 | 1.162570971 | MBD5 |
| 2 | 148469086 | 148469350 | -1.355326797 | MBD5 |
| 1 | 144068225 | 144068424 | -1.672416227 | SRGAP2D |
| 2 | 151251351 | 151251509 | -1.02383169 | RBM43 |
| 2 | 152389689 | 152389892 | 1.391878308 | FMNL2 |
| 1 | 144423049 | 144423246 | -1.652546443 | NBPF15 |
| 2 | 152390176 | 152390325 | 8.323750293 | FMNL2 |
| 2 | 152390176 | 152390325 | 8.323750293 | AC012066.1 |
| 2 | 156325641 | 156325920 | -1.242162363 | NR4A2 |
| 2 | 157321304 | 157321412 | -3.598034645 | ERMN |
| 2 | 158625119 | 158625320 | -1.480270921 | PKP4 |
| 2 | 158682686 | 158682848 | -4.064641508 | PKP4 |
| 2 | 158682686 | 158682848 | -4.064641508 | AC005042.4 |
| 2 | 159474328 | 159474514 | -2.758363882 | BAZ2B |
| 2 | 159499270 | 159499427 | -5.725605556 | BAZ2B |
| 2 | 159500185 | 159500376 | -2.112989369 | BAZ2B |
| 2 | 159797285 | 159797484 | -8.394868701 | CD302 |
| 2 | 159797285 | 159797484 | -8.394868701 | LY75-CD302 |
| 2 | 162308078 | 162308278 | 10.09606813 | IFIH1 |
| 2 | 162309529 | 162309663 | 4.529437353 | IFIH1 |
| 2 | 162310834 | 162311035 | 3.076818934 | IFIH1 |
| 2 | 162318079 | 162318290 | 2.538111142 | IFIH1 |
| 2 | 165770345 | 165770544 | 1.251994211 | GALNT3 |
| 2 | 168236459 | 168236640 | -4.038407799 | STK39 |
| 2 | 169081594 | 169081795 | -1.09442142 | DHRS9 |
| 2 | 169083251 | 169083441 | -2.538858416 | DHRS9 |
| 2 | 169095603 | 169095841 | -1.165792702 | DHRS9 |
| 1 | 145168001 | 145168125 | 2.092699107 | RP11-403I13.8 |
| 2 | 169546386 | 169546572 | -1.320945647 | FASTKD1 |
| 2 | 169636995 | 169637195 | -2.186879674 | PPIG |
| 2 | 169688820 | 169688934 | -4.79248125 | CCDC173 |
| 2 | 169749393 | 169749563 | -1.564641508 | KLHL23 |
| 2 | 170993473 | 170993689 | -2.170339819 | TLK1 |
| 2 | 171091928 | 171092129 | -2.161840153 | TLK1 |
| 2 | 172088737 | 172088939 | -2.416157609 | DLX1 |
| 2 | 172503966 | 172504096 | -1.00543791 | AC093818.1 |
| 2 | 172503966 | 172504096 | -1.00543791 | ITGA6 |
| 1 | 145291498 | 145291694 | -1.190577658 | NBPF20 |
| 2 | 172527271 | 172527467 | 5.474183616 | AC093818.1 |
| 2 | 172529119 | 172529220 | 6.797628741 | AC093818.1 |
| 2 | 172530057 | 172530240 | 3.661999023 | AC093818.1 |
| 2 | 173910041 | 173910236 | -7.380100076 | SP3 |
| 2 | 173955061 | 173955261 | -1.160403608 | SP3 |
| 2 | 174335556 | 174335662 | -5.033044595 | RP11-572N21.1 |
| 2 | 174335556 | 174335662 | -5.033044595 | SP9 |
| 1 | 6098604 | 6098805 | -1.000001064 | KCNAB2 |
| 1 | 145660334 | 145660482 | 1.35190743 | GPR89A |
| 2 | 174395969 | 174396161 | -2.569626653 | SCRN3 |
| 2 | 175930005 | 175930217 | -1.143519603 | LNPK |
| 2 | 176131468 | 176131660 | 3.200420748 | HOXD-AS2 |
| 2 | 176131468 | 176131660 | 3.200420748 | HOXD8 |
| 2 | 177369243 | 177369394 | -3.040711457 | AC074286.1 |
| 2 | 177369243 | 177369394 | -3.040711457 | NFE2L2 |

| | | | | | | | | | |
|---|---|---|---|---|---|---|---|---|---|
| 19 | 39386080 | 39386283 | -1.513016802 | PAF1 | 2 | 177459111 | 177459304 | -4.010174323 | AGPS |
| 19 | 39532181 | 39532371 | -1.594316462 | EID2B | 2 | 178542172 | 178542333 | -5.753104195 | TTN |
| 19 | 40014069 | 40014266 | -1.410905884 | ZNF546 | 2 | 178542172 | 178542333 | -5.753104195 | TTN-AS1 |
| 19 | 40033477 | 40033659 | -1.945989117 | ZNF780B | 2 | 178542671 | 178542866 | -6.13339327 | TTN |
| 19 | 40034418 | 40034546 | -1.791449872 | ZNF780B | 2 | 178542671 | 178542866 | -6.13339327 | TTN-AS1 |
| 19 | 40034678 | 40034863 | -2.029871042 | ZNF780B | 2 | 178548240 | 178548437 | -5.369890305 | TTN-AS1 |
| 19 | 40035251 | 40035431 | -1.656957028 | ZNF780B | 2 | 178548240 | 178548437 | -5.369890305 | TTN |
| 19 | 40036379 | 40036571 | -1.875262501 | ZNF780B | 2 | 179945480 | 179945677 | -1.106829481 | CWC22 |
| 1 | 110951617 | 110951840 | -1.382659699 | LRIF1 | 2 | 181032950 | 181033113 | -7.520401705 | UBE2E3 |
| 1 | 110951617 | 110951840 | -1.382659699 | RP11-96K19.2 | 2 | 181033510 | 181033662 | -3.735790351 | UBE2E3 |
| 19 | 40038068 | 40038245 | -1.256108561 | ZNF780B | 1 | 146058682 | 146058846 | -2.078252243 | RP11-458D21.1 |
| 19 | 40038899 | 40039101 | -1.037690956 | ZNF780B | 2 | 181891323 | 181891519 | -1.254348303 | RP11-366L5.1 |
| 19 | 40041681 | 40041917 | -1.03374883 | ZNF780B | 2 | 181902153 | 181902353 | -1.030598454 | SSFA2 |
| 19 | 40073705 | 40073887 | -2.250214131 | ZNF780A | 2 | 181915398 | 181915505 | -1.404727029 | SSFA2 |
| 19 | 40073705 | 40073887 | -2.250214131 | AC005614.5 | 2 | 181916118 | 181916317 | -1.21443042 | SSFA2 |
| 19 | 40074844 | 40075047 | -2.121455872 | AC005614.5 | 2 | 181922123 | 181922319 | -9.110319057 | SSFA2 |
| 19 | 40074844 | 40075047 | -2.121455872 | ZNF780A | 2 | 184649737 | 184649928 | -7.573582584 | ZNF804A |
| 19 | 40075964 | 40076164 | -1.620169604 | ZNF780A | 2 | 185547076 | 185547276 | -2.638460994 | AC080125.1 |
| 19 | 40075964 | 40076164 | -1.620169604 | AC005614.5 | 2 | 187519403 | 187519565 | -5.94208526 | AC007319.1 |
| 19 | 40192214 | 40192483 | -1.211633006 | MAP3K10 | 2 | 187519403 | 187519565 | -5.94208526 | TFPI |
| 19 | 40215114 | 40215329 | 1.579199374 | MAP3K10 | 2 | 189563898 | 189564130 | -1.752133633 | SLC40A1 |
| 19 | 40422811 | 40423109 | 2.533438794 | SERTAD1 | 2 | 189665989 | 189666227 | -1.178674606 | ASNSD1 |
| 19 | 40667396 | 40667620 | 2.193357608 | NUMBL | 2 | 189713839 | 189714042 | -1.180616816 | ANKAR |
| 19 | 40807907 | 40808110 | -1.010727092 | RAB4B-EGLN2 | 2 | 189854507 | 189854705 | -2.68128504 | PMS1 |
| 19 | 40807907 | 40808110 | -1.010727092 | CTC-490E21.12 | 2 | 189863760 | 189863950 | -1.311952849 | PMS1 |
| 19 | 40807907 | 40808110 | -1.010727092 | EGLN2 | 2 | 190200042 | 190200234 | -1.357924073 | HIBCH |
| 19 | 40826657 | 40826800 | 1.474805735 | CTC-490E21.12 | 2 | 190200042 | 190200234 | -1.357924073 | C2orf88 |
| 19 | 40826657 | 40826800 | 1.474805735 | CYP2F2P | 2 | 190242103 | 190242224 | -2.406166299 | HIBCH |
| 19 | 41353360 | 41353568 | 2.893727817 | TMEM91 | 2 | 190371127 | 190371325 | -1.365529208 | INPP1 |
| 19 | 41353360 | 41353568 | 2.893727817 | CTC-435M10.3 | 2 | 190476764 | 190476946 | -7.286882757 | MFSD6 |
| 19 | 41353360 | 41353568 | 2.893727817 | TGFB1 | 2 | 190500163 | 190500300 | -2.06280507 | MFSD6 |
| 19 | 41426383 | 41426578 | 1.093694679 | B3GNT8 | 2 | 190509044 | 190509194 | -1.456003073 | NEMP2 |
| 1 | 111187283 | 111187470 | -1.505036221 | DENND2D | 2 | 190509044 | 190509194 | -1.456003073 | MFSD6 |
| 19 | 41889498 | 41889729 | -1.903061157 | ARHGEF1 | 1 | 146794299 | 146794429 | -6.049345048 | HYDIN2 |
| 19 | 42079644 | 42079838 | -1.482236467 | ZNF574 | 2 | 190557328 | 190557489 | -3.982892142 | AC093388.3 |
| 19 | 42081129 | 42081328 | -1.413824578 | ZNF574 | 2 | 190955365 | 190955565 | 1.564132402 | GLS |
| 19 | 42088557 | 42088758 | 1.247564114 | POU2F2 | 2 | 190956109 | 190956308 | 1.663526564 | GLS |
| 19 | 42089704 | 42089904 | 1.221759152 | POU2F2 | 2 | 190959186 | 190959389 | 1.336165206 | GLS |
| 19 | 42091013 | 42091199 | 1.258961108 | POU2F2 | 2 | 195680549 | 195680748 | -1.76867084 | SLC39A10 |
| 19 | 42137823 | 42137926 | 4.832667959 | POU2F2 | 2 | 195744397 | 195744628 | -1.966426545 | DNAH7 |
| 19 | 42226285 | 42226484 | -1.698831498 | ZNF526 | 2 | 195746275 | 195746467 | -7.209980089 | DNAH7 |
| 1 | 111200826 | 111200989 | 6.9027208 | DENND2D | 2 | 196752852 | 196752956 | -2.501876067 | CCDC150 |
| 1 | 111200826 | 111200989 | 6.9027208 | CHI3L2 | 1 | 147172812 | 147173136 | -1.219309468 | RP11-337C18.8 |
| 19 | 42271926 | 42272056 | -7.320517835 | CIC | 2 | 199965398 | 199965612 | 1.413588386 | C2orf47 |
| 19 | 42376600 | 42376753 | -2.204646463 | MEGF8 | 2 | 200571382 | 200571585 | -2.628791713 | SGO2 |
| 19 | 43586104 | 43586312 | 1.379114953 | IRGQ | 2 | 200573329 | 200573527 | -5.866507661 | SGO2 |
| 19 | 43586104 | 43586312 | 1.379114953 | L34079.2 | 2 | 200892047 | 200892250 | -1.988790802 | NIF3L1 |
| 19 | 43592726 | 43592932 | 1.608498882 | L34079.2 | 1 | 6222964 | 6223168 | -1.204079066 | ICMT |
| 19 | 43592726 | 43592932 | 1.608498882 | IRGQ | 2 | 201116734 | 201116935 | 2.189544423 | CFLAR |
| 19 | 43594918 | 43595150 | 1.554467186 | IRGQ | 2 | 201117337 | 201117540 | 2.123434033 | CFLAR |
| 1 | 3770693 | 3770889 | 1.47477975 | CCDC27 | 2 | 201137497 | 201137700 | 4.174427809 | IMPDH1P10 |
| 19 | 43648852 | 43649052 | -7.887007431 | PLAUR | 2 | 201137497 | 201137700 | 4.174427809 | CFLAR |
| 19 | 43659729 | 43659890 | 3.014852022 | PLAUR | 2 | 201138656 | 201138849 | 10.67606914 | IMPDH1P10 |
| 19 | 43661671 | 43661868 | 4.297789218 | PLAUR | 2 | 201138656 | 201138849 | 10.67606914 | CFLAR |

| | | | | | | | | | |
|---|---|---|---|---|---|---|---|---|---|
| 19 | 43766682 | 43766882 | 1.979652578 | KCNN4 | 2 | 201160584 | 201160781 | 2.164998331 | CFLAR |
| 19 | 43975288 | 43975493 | -1.484261159 | ZNF155 | 2 | 201163742 | 201163942 | 2.553186099 | CFLAR |
| 19 | 44010934 | 44011143 | -2.287569437 | ZNF230 | 2 | 201167248 | 201167439 | 1.939488003 | AC007283.5 |
| 19 | 44235206 | 44235483 | -1.059163557 | ZNF235 | 2 | 201167248 | 201167439 | 1.939488003 | CFLAR |
| 19 | 44235206 | 44235483 | -1.059163557 | ZNF227 | 2 | 201169393 | 201169517 | 1.345371126 | CFLAR |
| 19 | 44236810 | 44237005 | -2.494730789 | ZNF235 | 2 | 201219334 | 201219543 | 1.41238354 | CASP10 |
| 19 | 44236810 | 44237005 | -2.494730789 | ZNF227 | 2 | 201240464 | 201240664 | 2.039699854 | CASP8 |
| 19 | 44252701 | 44252883 | 1.019246759 | ZNF235 | 2 | 201246485 | 201246664 | 5.020403207 | CASP8 |
| 19 | 44289030 | 44289176 | -1.150355804 | ZNF235 | 2 | 201296193 | 201296419 | 1.945385465 | ALS2CR12 |
| 19 | 44476420 | 44476618 | -2.122161675 | ZNF180 | 2 | 201380721 | 201380925 | -1.10841497 | TRAK2 |
| 19 | 44477017 | 44477209 | -1.259956128 | ZNF180 | 2 | 201761162 | 201761411 | -1.293735425 | ALS2 |
| 19 | 44661832 | 44662039 | -1.240952268 | CTB-171A8.1 | 2 | 201761569 | 201761769 | -1.529496963 | ALS2 |
| 19 | 44661832 | 44662039 | -1.240952268 | PVR | 2 | 202377093 | 202377297 | 1.046156074 | BMPR2 |
| 19 | 44663169 | 44663370 | -1.010065351 | PVR | 1 | 147203716 | 147203898 | -5.477098155 | CCT8P1 |
| 19 | 44663169 | 44663370 | -1.010065351 | CTB-171A8.1 | 1 | 147203716 | 147203898 | -5.477098155 | RP11-337C18.8 |
| 19 | 44663169 | 44663370 | -1.010065351 | CEACAM19 | 1 | 147203716 | 147203898 | -5.477098155 | RP11-337C18.10 |
| 19 | 44878636 | 44878837 | 1.393528763 | NECTIN2 | 1 | 147203716 | 147203898 | -5.477098155 | FMO5 |
| 19 | 45090550 | 45090750 | -1.115740372 | MARK4 | 2 | 202765963 | 202766162 | -2.260857948 | FAM117B |
| 19 | 45090550 | 45090750 | -1.115740372 | GEMIN7 | 2 | 202779453 | 202779615 | -2.277474966 | ICA1L |
| 19 | 45145594 | 45145760 | -6.93518236 | PPP1R37 | 2 | 202925001 | 202925195 | -2.591761302 | WDR12 |
| 19 | 45145594 | 45145760 | -6.93518236 | MARK4 | 2 | 202925001 | 202925195 | -2.591761302 | CARF |
| 19 | 45152860 | 45153000 | -2.085551596 | NKPD1 | 2 | 202925001 | 202925195 | -2.591761302 | AC010900.2 |
| 19 | 45152860 | 45153000 | -2.085551596 | MARK4 | 2 | 202947292 | 202947393 | -5.738373102 | CARF |
| 19 | 45408488 | 45408683 | -1.659686195 | ERCC1 | 2 | 202947292 | 202947393 | -5.738373102 | WDR12 |
| 19 | 45408488 | 45408683 | -1.659686195 | CD3EAP | 1 | 147205809 | 147206052 | 1.498710186 | RP11-337C18.8 |
| 19 | 45409497 | 45409673 | -1.405634664 | CD3EAP | 1 | 147205809 | 147206052 | 1.498710186 | RP11-337C18.10 |
| 19 | 45409497 | 45409673 | -1.405634664 | ERCC1 | 1 | 147205809 | 147206052 | 1.498710186 | FMO5 |
| 19 | 45412665 | 45412845 | -8.309558322 | ERCC1 | 2 | 206086159 | 206086371 | -1.471725402 | AC007383.3 |
| 1 | 111608827 | 111609030 | 7.193544193 | RAP1A | 2 | 206086159 | 206086371 | -1.471725402 | INO80D |
| 19 | 45468256 | 45468480 | 2.654078955 | ERCC1 | 2 | 206766683 | 206766891 | -1.064730627 | FASTKD2 |
| 19 | 45468256 | 45468480 | 2.654078955 | FOSB | 2 | 207239835 | 207239978 | -1.777547494 | AC007879.5 |
| 19 | 45472915 | 45473127 | 2.175665981 | FOSB | 2 | 207239835 | 207239978 | -1.777547494 | AC007879.7 |
| 19 | 45472915 | 45473127 | 2.175665981 | ERCC1 | 2 | 207404410 | 207404598 | 3.73113029 | AC007879.5 |
| 19 | 45529473 | 45529593 | -1.054153488 | OPA3 | 1 | 147206596 | 147206697 | 1.843089513 | RP11-337C18.10 |
| 19 | 45553327 | 45553528 | -1.269187096 | OPA3 | 1 | 147206596 | 147206697 | 1.843089513 | FMO5 |
| 19 | 45681958 | 45682158 | -1.771624201 | GIPR | 1 | 147206596 | 147206697 | 1.843089513 | RP11-337C18.8 |
| 19 | 45712296 | 45712491 | -1.53300452 | FBXO46 | 2 | 207605096 | 207605242 | -4.614409345 | METTL21A |
| 19 | 45765498 | 45765678 | -2.069329436 | AC074212.5 | 2 | 207613149 | 207613349 | -1.344796576 | METTL21A |
| 19 | 45765498 | 45765678 | -2.069329436 | SIX5 | 2 | 207977100 | 207977327 | -1.221656891 | PLEKHM3 |
| 19 | 45768980 | 45769209 | -1.232051285 | AC074212.6 | 2 | 208001268 | 208001527 | -1.200756435 | PLEKHM3 |
| 19 | 45768980 | 45769209 | -1.232051285 | AC074212.5 | 2 | 210055585 | 210055722 | -1.087284713 | KANSL1L |
| 19 | 45768980 | 45769209 | -1.232051285 | SIX5 | 2 | 210055585 | 210055722 | -1.087284713 | AC007038.7 |
| 19 | 45883985 | 45884185 | -1.82236198 | IRF2BP1 | 2 | 211382395 | 211382602 | -1.162360306 | ERBB4 |
| 19 | 45885103 | 45885293 | -1.336672592 | IRF2BP1 | 2 | 211383722 | 211383959 | -1.887025125 | ERBB4 |
| 19 | 45890169 | 45890330 | -3.656335139 | MYPOP | 2 | 212237642 | 212237820 | -1.675248624 | ERBB4 |
| 19 | 46609600 | 46609843 | -1.922004031 | CTB-12A17.3 | 2 | 213675310 | 213675499 | -1.858706398 | SPAG16 |
| 19 | 46609600 | 46609843 | -1.922004031 | CALM3 | 2 | 215611818 | 215612018 | -4.169925001 | AC012668.2 |
| 19 | 46621054 | 46621245 | 1.154163896 | PTGIR | 2 | 215611818 | 215612018 | -4.169925001 | LINC00607 |
| 19 | 46775361 | 46775559 | -3.039574927 | SLC1A5 | 2 | 215944722 | 215944909 | 1.042720422 | MREG |
| 19 | 46775361 | 46775559 | -3.039574927 | FKRP | 2 | 218068701 | 218068913 | 1.876303178 | RUFY4 |
| 1 | 111682386 | 111682566 | 6.569775676 | FAM212B | 2 | 218069294 | 218069432 | 3.046410405 | RUFY4 |
| 1 | 111682386 | 111682566 | 6.569775676 | RAP1A | 2 | 218090224 | 218090408 | 2.435330313 | RUFY4 |
| 19 | 46787803 | 46787979 | -2.045283713 | SLC1A5 | 2 | 218345325 | 218345512 | -3.430729583 | PNKD |
| 19 | 46787803 | 46787979 | -2.045283713 | CTB-147N14.6 | 2 | 218345325 | 218345512 | -3.430729583 | CATIP-AS2 |

| | | | | |
|---|---|---|---|---|
| 19 | 46788099 | 46788294 | -1.624566081 | CTB-147N14.6 |
| 19 | 46788099 | 46788294 | -1.624566081 | SLC1A5 |
| 19 | 47065889 | 47066107 | -1.590884593 | ZC3H4 |
| 19 | 47066651 | 47066889 | -1.45616998 | ZC3H4 |
| 19 | 47069153 | 47069383 | -1.406668849 | ZC3H4 |
| 19 | 47085109 | 47085308 | -2.8290797 | ZC3H4 |
| 19 | 47221527 | 47221834 | 1.562065856 | BBC3 |
| 19 | 47320377 | 47320573 | 12.3537346 | C5AR1 |
| 19 | 47332134 | 47332244 | 6.946366266 | C5AR2 |
| 19 | 47341589 | 47341799 | 7.453809949 | C5AR2 |
| 19 | 47407156 | 47407346 | -2.006680189 | MEIS3 |
| 19 | 48123881 | 48124079 | -7.826044352 | CTC-453G23.5 |
| 19 | 48123881 | 48124079 | -7.826044352 | LIG1 |
| 19 | 48170908 | 48171107 | -2.05663933 | C19orf68 |
| 19 | 48195402 | 48195598 | -3.31590405 | CARD8 |
| 19 | 48195402 | 48195598 | -3.31590405 | ZNF114 |
| 19 | 48195402 | 48195598 | -3.31590405 | C19orf68 |
| 19 | 48196553 | 48196756 | -1.390391342 | CARD8 |
| 19 | 48196553 | 48196756 | -1.390391342 | ZNF114 |
| 19 | 48196553 | 48196756 | -1.390391342 | C19orf68 |
| 19 | 48203843 | 48204058 | 1.166718508 | CARD8 |
| 19 | 48203843 | 48204058 | 1.166718508 | ZNF114 |
| 1 | 3775899 | 3776101 | 1.067399295 | SMIM1 |
| 19 | 48286153 | 48286292 | -3.294574226 | ZNF114 |
| 19 | 48286664 | 48286868 | -2.722004226 | ZNF114 |
| 19 | 48453067 | 48453268 | -1.640110281 | GRWD1 |
| 1 | 111766155 | 111766437 | -1.116869997 | DDX20 |
| 19 | 48873453 | 48873656 | 1.793774839 | PPP1R15A |
| 19 | 48874436 | 48874634 | 1.172845363 | PPP1R15A |
| 19 | 49150230 | 49150433 | -3.08880943 | PPFIA3 |
| 19 | 49310638 | 49310885 | 1.104337771 | SLC6A16 |
| 19 | 49470352 | 49470551 | -3.116759972 | CTD-3148I10.9 |
| 19 | 49470352 | 49470551 | -3.116759972 | ALDH16A1 |
| 19 | 49525037 | 49525237 | 8.201672478 | FCGRT |
| 19 | 49596221 | 49596422 | -1.1775762 | PRR12 |
| 1 | 112456055 | 112456253 | 1.892056771 | CTTNBP2NL |
| 19 | 49599274 | 49599473 | -1.760122055 | PRR12 |
| 19 | 49651303 | 49651492 | -10.13698201 | SCAF1 |
| 19 | 49653256 | 49653440 | -1.501928283 | SCAF1 |
| 19 | 49654221 | 49654388 | -1.453532612 | SCAF1 |
| 19 | 49654544 | 49654740 | -1.34610881 | SCAF1 |
| 19 | 49658356 | 49658557 | -1.557580641 | SCAF1 |
| 19 | 49857848 | 49858031 | -9.316016573 | PTOV1 |
| 19 | 49857848 | 49858031 | -9.316016573 | PTOV1-AS2 |
| 19 | 49877181 | 49877381 | 2.346509429 | PNKP |
| 19 | 49877181 | 49877381 | 2.346509429 | AKT1S1 |
| 1 | 112457139 | 112457329 | 1.055837134 | CTTNBP2NL |
| 19 | 49879531 | 49879713 | 6.497176718 | TBC1D17 |
| 19 | 49880533 | 49880725 | 8.187383907 | TBC1D17 |
| 19 | 49884434 | 49884639 | 10.57385338 | TBC1D17 |
| 19 | 49886523 | 49886720 | 8.501352579 | TBC1D17 |
| 19 | 49887251 | 49887451 | 9.680003434 | TBC1D17 |
| 19 | 49887903 | 49888097 | 7.594989474 | TBC1D17 |
| 19 | 49889763 | 49889972 | 7.21210248 | IL4I1 |

| | | | | |
|---|---|---|---|---|
| 2 | 218594339 | 218594541 | -1.167391 | CNOT9 |
| 2 | 218633415 | 218633614 | -1.178956429 | PLCD4 |
| 2 | 218633415 | 218633614 | -1.178956429 | RP11-548H3.1 |
| 2 | 218638154 | 218638380 | -1.322033767 | ZNF142 |
| 2 | 218644114 | 218644296 | -1.474207886 | ZNF142 |
| 2 | 218873906 | 218874104 | -2.264441914 | WNT6 |
| 2 | 219234804 | 219235000 | -1.253059455 | ANKZF1 |
| 2 | 219250974 | 219251182 | -1.612329248 | TUBA4A |
| 2 | 219498275 | 219498514 | -2.220708013 | SPEG |
| 2 | 219498275 | 219498514 | -2.220708013 | AC053503.11 |
| 1 | 6223811 | 6224010 | -7.828868564 | ICMT |
| 2 | 219570680 | 219570872 | 5.231251136 | INHA |
| 2 | 219570680 | 219570872 | 5.231251136 | OBSL1 |
| 2 | 222318595 | 222318815 | 1.362839968 | AC010980.2 |
| 2 | 222318595 | 222318815 | 1.362839968 | RP11-384O8.1 |
| 2 | 223963643 | 223963829 | -2.32919477 | MRPL44 |
| 2 | 224000803 | 224000976 | 1.627352284 | SERPINE2 |
| 2 | 224811471 | 224811670 | -1.112247482 | DOCK10 |
| 1 | 148288638 | 148288844 | -9.345626399 | RP11-495P10.9 |
| 2 | 226931329 | 226931513 | 6.143567593 | RHBDD1 |
| 2 | 230169075 | 230169274 | 1.276440651 | SP110 |
| 2 | 230169075 | 230169274 | 1.276440651 | AC009950.2 |
| 1 | 148317821 | 148318004 | -6.038407799 | RP11-495P10.4 |
| 2 | 230177339 | 230177535 | 1.738466524 | SP110 |
| 2 | 230177339 | 230177535 | 1.738466524 | AC009950.1 |
| 2 | 230196800 | 230197001 | 2.24467832 | SP110 |
| 2 | 230196800 | 230197001 | 2.24467832 | AC009950.1 |
| 2 | 230197582 | 230197716 | 9.313326866 | SP110 |
| 2 | 230197582 | 230197716 | 9.313326866 | AC009950.1 |
| 2 | 230402843 | 230403044 | 1.190530646 | SP140L |
| 2 | 230412036 | 230412223 | 1.437979368 | RP11-69J7.1 |
| 2 | 230514902 | 230515024 | 7.969605941 | HMGB1P3 |
| 2 | 230514902 | 230515024 | 7.969605941 | SP100 |
| 2 | 230525607 | 230525720 | 3.086660128 | SP100 |
| 2 | 230525607 | 230525720 | 3.086660128 | AC010149.4 |
| 1 | 148334772 | 148334956 | -1.52882512 | RP11-495P10.3 |
| 2 | 230526059 | 230526259 | 1.822206783 | SP100 |
| 2 | 230526059 | 230526259 | 1.822206783 | AC010149.4 |
| 2 | 230527576 | 230527817 | 1.048338542 | AC010149.4 |
| 2 | 230527576 | 230527817 | 1.048338542 | SP100 |
| 2 | 230530412 | 230530568 | 1.814849063 | AC010149.4 |
| 2 | 230530412 | 230530568 | 1.814849063 | SP100 |
| 2 | 230531115 | 230531323 | 1.766519551 | SP100 |
| 2 | 230531115 | 230531323 | 1.766519551 | AC010149.4 |
| 2 | 230804183 | 230804379 | -1.451226239 | CAB39 |
| 2 | 231108663 | 231108786 | -4.114409345 | PSMD1 |
| 2 | 231108663 | 231108786 | -4.114409345 | HTR2B |
| 2 | 231373964 | 231374114 | -2.1797014 | ARMC9 |
| 2 | 231397793 | 231397970 | -6.226892528 | B3GNT7 |
| 2 | 231524879 | 231525098 | -3.65505133 | NMUR1 |
| 2 | 232340480 | 232340679 | 1.128663106 | DIS3L2 |
| 2 | 232344008 | 232344163 | -3.355833487 | DIS3L2 |
| 2 | 232548585 | 232548783 | -2.345377145 | TIGD1 |
| 2 | 232549173 | 232549282 | -2.531030484 | TIGD1 |

| | | | | | | | | | |
|---|---|---|---|---|---|---|---|---|---|
| 19 | 49908313 | 49908514 | -1.283796088 | IL4I1 | 2 | 232549772 | 232549965 | -2.134730337 | TIGD1 |
| 19 | 49908313 | 49908514 | -1.283796088 | NUP62 | 2 | 232581015 | 232581223 | 1.088915896 | AC073254.1 |
| 1 | 112457693 | 112457881 | 3.812346966 | CTTNBP2NL | 2 | 232581015 | 232581223 | 1.088915896 | EIF4E2 |
| 19 | 49940545 | 49940714 | 3.890679857 | CTC-326K19.6 | 2 | 233088651 | 233088857 | 1.784305792 | INPP5D |
| 19 | 50045224 | 50045343 | -1.57456698 | CTD-2126E3.3 | 2 | 233206898 | 233207102 | -1.679901763 | INPP5D |
| 19 | 50045224 | 50045343 | -1.57456698 | ZNF473 | 2 | 233212851 | 233212994 | -1.456353961 | ATG16L1 |
| 19 | 50509364 | 50509528 | 2.033175024 | JOSD2 | 2 | 233840664 | 233840862 | -2.529592852 | HJURP |
| 1 | 112493500 | 112493777 | 2.443890886 | WNT2B | 2 | 233841707 | 233841945 | -3.223318554 | HJURP |
| 19 | 50798157 | 50798438 | -2.217379831 | C19orf48 | 2 | 233866866 | 233867066 | -2.459873395 | AC005538.3 |
| 19 | 51413801 | 51414028 | -2.219929811 | SIGLEC10 | 2 | 233866866 | 233867066 | -2.459873395 | MSL3P1 |
| 19 | 51992436 | 51992568 | -1.993467983 | ZNF615 | 2 | 236124302 | 236124503 | -1.988897127 | AGAP1 |
| 1 | 112855282 | 112855514 | -1.664726543 | RP3-522D1.1 | 2 | 236165694 | 236165864 | -1.357721398 | GBX2 |
| 19 | 52034060 | 52034339 | 1.087041118 | ZNF432 | 2 | 237084406 | 237084622 | -3.134775876 | AC105760.2 |
| 19 | 52035027 | 52035218 | 1.284847872 | ZNF432 | 2 | 237529384 | 237529571 | -6.654596053 | MLPH |
| 19 | 52060424 | 52060617 | -2.130480432 | ZNF432 | 2 | 237779775 | 237780017 | -1.16195833 | LRRFIP1 |
| 19 | 52060424 | 52060617 | -2.130480432 | CTC-471J1.2 | 2 | 238151206 | 238151386 | -6.470707235 | KLHL30 |
| 19 | 52115102 | 52115292 | -1.41940157 | ZNF616 | 2 | 238166996 | 238167239 | -1.614038898 | ERFE |
| 19 | 52323241 | 52323437 | -3.178769812 | CTD-2525I3.6 | 2 | 238446499 | 238446713 | -1.463652386 | ASB1 |
| 19 | 52323241 | 52323437 | -3.178769812 | ZNF480 | 2 | 239160434 | 239160601 | 2.032477721 | HDAC4 |
| 19 | 52324628 | 52324804 | -3.138237267 | CTD-2525I3.6 | 2 | 239358325 | 239358504 | -2.182555526 | HDAC4 |
| 19 | 52324628 | 52324804 | -3.138237267 | ZNF480 | 1 | 6224826 | 6225026 | -1.156309582 | ICMT |
| 19 | 52325482 | 52325681 | -3.767197808 | ZNF480 | 2 | 240488014 | 240488211 | -4.824995488 | ANKMY1 |
| 19 | 52325482 | 52325681 | -3.767197808 | CTD-2525I3.6 | 2 | 240491092 | 240491286 | -1.903677461 | ANKMY1 |
| 19 | 52415991 | 52416108 | 1.111196211 | ZNF528 | 2 | 240496558 | 240496881 | -1.079922312 | ANKMY1 |
| 19 | 52583724 | 52583881 | -1.021051801 | ZNF701 | 2 | 240524014 | 240524297 | -1.131477292 | ANKMY1 |
| 19 | 52583724 | 52583881 | -1.021051801 | CTD-3099C6.7 | 2 | 240529240 | 240529444 | -1.981048224 | ANKMY1 |
| 19 | 52584145 | 52584385 | -1.736995921 | ZNF701 | 2 | 241077577 | 241077719 | -6.243668808 | SNED1 |
| 19 | 52584145 | 52584385 | -1.736995921 | CTD-3099C6.7 | 2 | 241077577 | 241077719 | -6.243668808 | MTERF4 |
| 19 | 52653840 | 52654040 | -1.67163522 | ZNF83 | 2 | 241096049 | 241096259 | -1.06637999 | MTERF4 |
| 1 | 112956787 | 112956921 | -1.476021235 | SLC16A1 | 2 | 241126910 | 241127121 | -1.851119108 | PASK |
| 1 | 112956787 | 112956921 | -1.476021235 | SLC16A1-AS1 | 2 | 241227599 | 241227794 | -3.223618161 | HDLBP |
| 19 | 52950164 | 52950304 | 1.128051899 | CTD-2331H12.8 | 2 | 241396174 | 241396373 | 8.09622347 | FARP2 |
| 19 | 52950164 | 52950304 | 1.128051899 | ZNF816-ZNF321P | 2 | 241569586 | 241569781 | -6.16152738 | BOK |
| 19 | 52950164 | 52950304 | 1.128051899 | ZNF816 | 2 | 241572088 | 241572274 | -2.495393411 | BOK |
| 19 | 53068352 | 53068708 | -1.072315152 | ZNF160 | 2 | 241686049 | 241686206 | -6.318765276 | DTYMK |
| 1 | 113641988 | 113642210 | -1.548845802 | MAGI3 | 2 | 241689897 | 241690131 | -1.014276444 | ING5 |
| 19 | 53386209 | 53386370 | -1.002203551 | CTD-3141N22.1 | 2 | 241725043 | 241725275 | -1.379029813 | ING5 |
| 19 | 53386209 | 53386370 | -1.002203551 | ZNF525 | 2 | 241725043 | 241725275 | -1.379029813 | AC114730.11 |
| 19 | 53410660 | 53410836 | -1.071014041 | ZNF765 | 1 | 148992534 | 148992718 | 2.975763306 | PDE4DIP |
| 19 | 53442291 | 53442494 | -1.672230177 | TPM3P9 | 20 | 326900 | 327100 | -3.163397118 | SOX12 |
| 19 | 53442291 | 53442494 | -1.672230177 | ZNF761 | 20 | 326900 | 327100 | -3.163397118 | NRSN2-AS1 |
| 19 | 53479901 | 53480125 | -1.554516556 | TPM3P6 | 20 | 327901 | 328104 | -2.891053853 | SOX12 |
| 19 | 53479901 | 53480125 | -1.554516556 | ZNF813 | 20 | 327901 | 328104 | -2.891053853 | NRSN2-AS1 |
| 19 | 53492922 | 53493149 | -1.089669304 | ZNF813 | 20 | 396597 | 396838 | -2.707272276 | TRIB3 |
| 19 | 53577127 | 53577354 | -1.419776819 | ZNF331 | 20 | 420163 | 420363 | 2.82070724 | RBCK1 |
| 19 | 53810294 | 53810494 | -1.575722333 | NLRP12 | 20 | 435648 | 435849 | -1.244806016 | TBC1D20 |
| 19 | 53875714 | 53875905 | -10.80135487 | AC008440.5 | 20 | 438710 | 438875 | -1.066471717 | TBC1D20 |
| 19 | 53875714 | 53875905 | -10.80135487 | MYADM | 20 | 1565159 | 1565352 | -2.697445683 | SIRPB1 |
| 19 | 54094830 | 54095030 | 2.751787502 | OSCAR | 20 | 1565159 | 1565352 | -2.697445683 | RP4-576H24.4 |
| 19 | 54096378 | 54096559 | -5.649604009 | OSCAR | 20 | 2794192 | 2794342 | 9.385693149 | CPXM1 |
| 1 | 113683710 | 113683916 | -1.990494658 | MAGI3 | 20 | 3047730 | 3047987 | -1.985151375 | MRPS26 |
| 19 | 54111046 | 54111245 | -1.699279093 | TFPT | 20 | 3121899 | 3122089 | -1.823416016 | UBOX5 |
| 19 | 54160914 | 54161155 | -1.008546621 | TMC4 | 20 | 3121899 | 3122089 | -1.823416016 | UBOX5-AS1 |
| 19 | 54165624 | 54165820 | -1.051206768 | TMC4 | 20 | 3148873 | 3149071 | 1.337009767 | FASTKD5 |

| | | | | |
|---|---|---|---|---|
| 19 | 54179102 | 54179391 | -1.858445301 | MBOAT7 |
| 19 | 54216890 | 54217090 | 8.993729164 | RPS9 |
| 19 | 54216890 | 54217090 | 8.993729164 | LILRB3 |
| 19 | 54460900 | 54461094 | 1.425959681 | LENG8 |
| 19 | 54461997 | 54462196 | -3.324397575 | LENG9 |
| 19 | 54461997 | 54462196 | -3.324397575 | LENG8 |
| 19 | 54462847 | 54463008 | -2.17737951 | LENG9 |
| 19 | 54498397 | 54498594 | -1.208554182 | LAIR2 |
| 19 | 54636915 | 54637117 | 2.161539711 | LILRB1 |
| 19 | 54636915 | 54637117 | 2.161539711 | AC009892.10 |
| 19 | 55041607 | 55041834 | -1.095743027 | CTC-550B14.7 |
| 19 | 55041607 | 55041834 | -1.095743027 | RDH13 |
| 19 | 55041607 | 55041834 | -1.095743027 | CTC-550B14.6 |
| 19 | 55044714 | 55044911 | -1.890874471 | CTC-550B14.7 |
| 19 | 55044714 | 55044911 | -1.890874471 | CTC-550B14.6 |
| 19 | 55044714 | 55044911 | -1.890874471 | RDH13 |
| 19 | 55079379 | 55079530 | -7.297103976 | EPS8L1 |
| 19 | 55091177 | 55091376 | -2.154935694 | PPP1R12C |
| 19 | 55226953 | 55227152 | -1.548741278 | TMEM86B |
| 19 | 55312580 | 55312888 | -6.906890596 | TMEM150B |
| 19 | 55339068 | 55339205 | -1.939232753 | CTD-2105E13.16 |
| 19 | 55347521 | 55347726 | -1.767181723 | KMT5C |
| 19 | 55453062 | 55453262 | -2.761049951 | ISOC2 |
| 19 | 55500508 | 55500707 | 6.046873831 | SSC5D |
| 19 | 55502494 | 55502693 | -2.19208393 | SSC5D |
| 1 | 113838017 | 113838140 | -4.245364512 | PTPN22 |
| 19 | 55592264 | 55592482 | -1.037647102 | FIZ1 |
| 19 | 55602738 | 55602931 | -2.553628355 | ZNF524 |
| 19 | 55615572 | 55615755 | -7.105107361 | ZNF865 |
| 1 | 113911640 | 113911875 | -1.699649531 | DCLRE1B |
| 19 | 55704382 | 55704622 | -1.074134805 | EPN1 |
| 19 | 55705124 | 55705327 | -1.295871408 | EPN1 |
| 19 | 56087680 | 56087897 | -1.479623 | ZNF787 |
| 19 | 56576833 | 56576938 | -1.553274252 | ZNF470 |
| 1 | 113940815 | 113941015 | -1.598732507 | HIPK1 |
| 1 | 853915 | 854108 | 1.521319241 | LINC01128 |
| 1 | 3779379 | 3779498 | 1.884826262 | LRRC47 |
| 19 | 57410933 | 57411152 | 1.161634756 | AC004076.7 |
| 19 | 57410933 | 57411152 | 1.161634756 | AC003002.6 |
| 19 | 57443517 | 57443740 | -1.922180561 | AC004076.9 |
| 19 | 57443517 | 57443740 | -1.922180561 | ZNF749 |
| 19 | 57445176 | 57445315 | -3.430619931 | ZNF749 |
| 19 | 57445176 | 57445315 | -3.430619931 | AC004076.9 |
| 19 | 57473092 | 57473198 | -1.355240803 | ZNF772 |
| 19 | 57473092 | 57473198 | -1.355240803 | AC004076.9 |
| 19 | 57474078 | 57474280 | -1.8033857 | ZNF772 |
| 19 | 57474078 | 57474280 | -1.8033857 | AC004076.9 |
| 19 | 57507646 | 57507971 | -1.925273595 | ZNF773 |
| 19 | 57538703 | 57538906 | -1.113071647 | ZNF550 |
| 19 | 57538703 | 57538906 | -1.113071647 | ZNF549 |
| 19 | 57547623 | 57547816 | -1.157821495 | ZNF550 |
| 19 | 57547623 | 57547816 | -1.157821495 | ZNF549 |
| 19 | 57572233 | 57572430 | -1.360730265 | ZNF416 |
| 19 | 57605933 | 57606153 | -2.045827713 | ZNF530 |

| | | | | |
|---|---|---|---|---|
| 20 | 3148873 | 3149071 | 1.337009767 | UBOX5-AS1 |
| 20 | 3148873 | 3149071 | 1.337009767 | UBOX5 |
| 20 | 3646800 | 3647017 | -2.118548692 | ATRN |
| 20 | 3648381 | 3648537 | -1.518846681 | ATRN |
| 20 | 3784875 | 3785074 | -1.042210699 | CENPB |
| 20 | 4182176 | 4182368 | -1.66270854 | SMOX |
| 20 | 4699532 | 4699734 | -1.237423251 | PRNP |
| 20 | 4782835 | 4783042 | 2.071781658 | RASSF2 |
| 20 | 5942503 | 5942720 | -1.839537651 | TRMT6 |
| 20 | 6768513 | 6768694 | 1.86050389 | BMP2 |
| 20 | 6778566 | 6778766 | 1.725220538 | BMP2 |
| 20 | 10622769 | 10622940 | -2.546280701 | SLX4IP |
| 20 | 10639604 | 10639803 | 2.569309091 | JAG1 |
| 20 | 11923264 | 11923553 | -1.926343441 | BTBD3 |
| 20 | 13299278 | 13299485 | -1.505768804 | ISM1 |
| 20 | 16379696 | 16379892 | -1.452852643 | KIF16B |
| 20 | 17967968 | 17968191 | 5.967029697 | SNX5 |
| 20 | 17967968 | 17968191 | 5.967029697 | OVOL2 |
| 20 | 17970048 | 17970246 | -2.228348325 | MGME1 |
| 20 | 17970048 | 17970246 | -2.228348325 | OVOL2 |
| 20 | 18162651 | 18162852 | -1.705281457 | KAT14 |
| 20 | 18181567 | 18181761 | -1.548489318 | KAT14 |
| 1 | 149791320 | 149791522 | 2.61551346 | RP11-196G18.3 |
| 1 | 149791320 | 149791522 | 2.61551346 | HIST2H2BF |
| 1 | 149791320 | 149791522 | 2.61551346 | FCGR1A |
| 20 | 18384228 | 18384426 | -4.865790064 | DZANK1 |
| 20 | 18794634 | 18794814 | -2.316760617 | RP5-1068E13.7 |
| 20 | 19996920 | 19997190 | -2.366010638 | RIN2 |
| 20 | 20000632 | 20000853 | -2.255743495 | RIN2 |
| 20 | 20391442 | 20391617 | -1.475752454 | RALGAPA2 |
| 20 | 20392890 | 20393082 | -1.941798488 | RALGAPA2 |
| 20 | 20512787 | 20512987 | -1.549886786 | RALGAPA2 |
| 20 | 21093412 | 21093553 | -4.321928095 | LINC00237 |
| 1 | 149792226 | 149792375 | 2.247382346 | FCGR1A |
| 1 | 149792226 | 149792375 | 2.247382346 | HIST2H2BF |
| 1 | 149792226 | 149792375 | 2.247382346 | RP11-196G18.3 |
| 20 | 21101054 | 21101227 | 3.204695468 | LINC00237 |
| 20 | 21102441 | 21102657 | -1.845413315 | LINC00237 |
| 20 | 21103356 | 21103563 | -5.281757041 | LINC00237 |
| 20 | 21161955 | 21162155 | -2.029058411 | RP5-872K7.7 |
| 20 | 21161955 | 21162155 | -2.029058411 | KIZ |
| 20 | 21161955 | 21162155 | -2.029058411 | KIZ-AS1 |
| 20 | 21167005 | 21167154 | -5.761780978 | KIZ |
| 20 | 21167005 | 21167154 | -5.761780978 | KIZ-AS1 |
| 20 | 21168946 | 21169166 | -1.99848702 | KIZ |
| 20 | 21168946 | 21169166 | -1.99848702 | KIZ-AS1 |
| 20 | 21500076 | 21500274 | -1.803841289 | RP5-984P4.6 |
| 20 | 23047585 | 23047786 | -1.362986859 | THBD |
| 1 | 149859179 | 149859363 | 3.669925001 | HIST2H4B |
| 20 | 23628441 | 23628597 | -1.855084193 | CST3 |
| 20 | 25221941 | 25222088 | -1.431533911 | ENTPD6 |
| 20 | 25264822 | 25265010 | -6.473818469 | PYGB |
| 1 | 149900359 | 149900561 | -1.199811112 | BOLA1 |
| 20 | 25285817 | 25286001 | 3.885938411 | PYGB |

| | | | | | | | | | |
|---|---|---|---|---|---|---|---|---|---|
| 19 | 57687610 | 57687848 | -1.421393115 | AC003006.7 | 20 | 25289959 | 25290139 | 8.083983446 | PYGB |
| 19 | 57687610 | 57687848 | -1.421393115 | ZNF551 | 20 | 25291542 | 25291653 | 7.770926432 | PYGB |
| 19 | 57729512 | 57729663 | -1.2325196 | AC003006.7 | 20 | 25507172 | 25507371 | -1.942083129 | NINL |
| 19 | 57770374 | 57770568 | -1.053613048 | ZNF586 | 20 | 25509303 | 25509507 | -1.268087714 | NINL |
| 19 | 57780132 | 57780481 | -1.666518978 | ZNF586 | 20 | 25616257 | 25616452 | -1.392293228 | NANP |
| 19 | 57786665 | 57786866 | -1.199505868 | ZNF586 | 1 | 149926743 | 149926943 | -9.56885582 | SF3B4 |
| 19 | 57841756 | 57841956 | -3.085906883 | ZNF587B | 20 | 25683343 | 25683497 | -1.441608323 | ZNF337 |
| 19 | 57841756 | 57841956 | -3.085906883 | CTD-2583A14.10 | 20 | 25690935 | 25691173 | -1.563339814 | ZNF337 |
| 19 | 57868218 | 57868470 | -1.348223015 | ZNF814 | 20 | 25857675 | 25857846 | -5.890474899 | FAM182B |
| 19 | 57868218 | 57868470 | -1.348223015 | CTD-2583A14.11 | 20 | 25865190 | 25865339 | 4.579935668 | FAM182B |
| 19 | 57907500 | 57907681 | -2.174841929 | ZNF417 | 20 | 31548487 | 31548676 | 5.042404194 | HM13 |
| 19 | 57907500 | 57907681 | -2.174841929 | CTD-2583A14.9 | 20 | 31665977 | 31666168 | 6.358409731 | BCL2L1 |
| 1 | 114625338 | 114625488 | 3.210731884 | DENND2C | 20 | 31801173 | 31801303 | -6.943943483 | TPX2 |
| 19 | 58146447 | 58146627 | 1.003558312 | ZNF329 | 1 | 149955801 | 149956005 | -1.033165037 | OTUD7B |
| 19 | 58260853 | 58261071 | -1.451697749 | ZNF544 | 20 | 32195422 | 32195682 | 1.221294174 | PLAGL2 |
| 19 | 58260853 | 58261071 | -1.451697749 | CTD-3138B18.4 | 20 | 32197177 | 32197386 | 1.086234472 | PLAGL2 |
| 19 | 58260853 | 58261071 | -1.451697749 | CTD-3138B18.5 | 20 | 32333458 | 32333589 | 1.053532844 | KIF3B |
| 19 | 58262756 | 58262956 | -1.106547082 | CTD-3138B18.5 | 20 | 32435555 | 32435748 | -1.365267181 | ASXL1 |
| 19 | 58262756 | 58262956 | -1.106547082 | CTD-3138B18.4 | 20 | 32436616 | 32436829 | -1.085246847 | ASXL1 |
| 19 | 58262756 | 58262956 | -1.106547082 | ZNF544 | 20 | 32438654 | 32438820 | -1.583202232 | ASXL1 |
| 1 | 114625822 | 114626011 | 2.309454916 | DENND2C | 20 | 33645032 | 33645231 | -1.106292185 | CBFA2T2 |
| 19 | 58359195 | 58359417 | 1.207773169 | CTD-2619J13.9 | 20 | 33707659 | 33707864 | -3.759228228 | PXMP4 |
| 19 | 58359195 | 58359417 | 1.207773169 | ZNF497 | 20 | 33989703 | 33989959 | -2.75632197 | RALY-AS1 |
| 19 | 58359195 | 58359417 | 1.207773169 | CTD-2619J13.8 | 20 | 33989703 | 33989959 | -2.75632197 | RP5-1125A11.6 |
| 19 | 58368807 | 58368967 | -6.391396749 | ZNF837 | 1 | 150101056 | 150101249 | -2.473885021 | VPS45 |
| 19 | 58406084 | 58406284 | -2.109612571 | CTD-2619J13.14 | 20 | 34089601 | 34089802 | -1.377361303 | EIF2S2 |
| 19 | 58406084 | 58406284 | -2.109612571 | ZNF584 | 20 | 34089601 | 34089802 | -1.377361303 | RALY |
| 19 | 58407903 | 58408099 | -2.100042238 | ZNF584 | 20 | 34560656 | 34560844 | -1.578422699 | PIGU |
| 19 | 58407903 | 58408099 | -2.100042238 | CTD-2619J13.14 | 1 | 150106095 | 150106207 | -1.691369478 | VPS45 |
| 19 | 58440384 | 58440514 | -3.183161107 | CTD-2619J13.19 | 20 | 35224697 | 35224869 | -2.091740662 | MMP24-AS1 |
| 19 | 58452180 | 58452356 | -7.621810146 | ZNF324B | 20 | 35503004 | 35503184 | -2.149702759 | CEP250 |
| 1 | 115663708 | 115663837 | -1.229994299 | VANGL1 | 20 | 35503829 | 35504130 | -1.965435059 | CEP250 |
| 19 | 58455442 | 58455670 | -1.702126867 | ZNF324B | 20 | 35507592 | 35507789 | -1.594236832 | CEP250 |
| 19 | 58456719 | 58456998 | -2.191814755 | ZNF324B | 1 | 150159262 | 150159488 | 1.728850657 | PLEKHO1 |
| 19 | 58470988 | 58471190 | -1.265181107 | ZNF324 | 20 | 35948115 | 35948316 | -1.981770069 | PHF20 |
| 19 | 58472375 | 58472579 | -1.906102005 | ZNF324 | 20 | 36229473 | 36229635 | -2.304518318 | EPB41L1 |
| 19 | 58479113 | 58479311 | -1.341379441 | ZNF446 | 20 | 36612033 | 36612235 | 1.233940699 | TGIF2-C20orf24 |
| 19 | 58516510 | 58516707 | -1.992551641 | ZBTB45 | 20 | 36612033 | 36612235 | 1.233940699 | C20orf24 |
| 19 | 58517400 | 58517529 | -1.359342579 | ZBTB45 | 1 | 150364352 | 150364549 | -5.249572794 | RPRD2 |
| 1 | 115691511 | 115691712 | -1.442446747 | VANGL1 | 20 | 36784474 | 36784622 | -1.530778513 | SOGA1 |
| 19 | 58519146 | 58519350 | -1.394335135 | ZBTB45 | 20 | 36785695 | 36785903 | -1.601985687 | SOGA1 |
| 19 | 58582874 | 58583086 | -1.065612956 | CENPBD1P1 | 20 | 36786254 | 36786390 | -1.922848046 | SOGA1 |
| 2 | 259024 | 259231 | -1.125737947 | SH3YL1 | 20 | 36793429 | 36793628 | -1.879180524 | SOGA1 |
| 2 | 3461030 | 3461228 | -1.028260939 | TRAPPC12 | 20 | 37404047 | 37404251 | 3.344399567 | SRC |
| 1 | 116046266 | 116046469 | -1.568291933 | SLC22A15 | 1 | 150470990 | 150471260 | -2.08659871 | RPRD2 |
| 2 | 3461976 | 3462110 | -8.953048941 | TRAPPC12 | 20 | 38951841 | 38952017 | -1.310102479 | FAM83D |
| 2 | 3467267 | 3467465 | -1.029091884 | TRAPPC12 | 20 | 40687418 | 40687649 | 1.159523552 | MAFB |
| 2 | 3523135 | 3523248 | -2.093178909 | AC142528.1 | 20 | 41174619 | 41174803 | -2.068702788 | PLCG1 |
| 2 | 3545750 | 3545949 | -1.126670505 | RNASEH1 | 20 | 41185101 | 41185299 | -7.493409882 | PLCG1 |
| 2 | 3545750 | 3545949 | -1.126670505 | RP13-512J5.1 | 20 | 41185101 | 41185299 | -7.493409882 | ZHX3 |
| 2 | 6849514 | 6849716 | 3.946565102 | CMPK2 | 20 | 41202159 | 41202356 | -2.057799009 | ZHX3 |
| 2 | 6895789 | 6895994 | 7.97647591 | RSAD2 | 1 | 150472909 | 150473109 | -1.040388867 | RPRD2 |
| 2 | 6948488 | 6948681 | -5.729715809 | RNF144A | 20 | 41202443 | 41202544 | -1.295940957 | ZHX3 |
| 2 | 8667047 | 8667242 | 7.811997884 | ID2-AS1 | 20 | 41204429 | 41204602 | -2.547954484 | ZHX3 |

| | | | | | | | | | |
|---|---|---|---|---|---|---|---|---|---|
| 2 | 8683751 | 8683951 | 3.906833412 | ID2 | 20 | 41405132 | 41405334 | -1.851319859 | CHD6 |
| 2 | 8858664 | 8858835 | -2.155847131 | MBOAT2 | 20 | 41415272 | 41415466 | -1.006109774 | CHD6 |
| 2 | 9405863 | 9406060 | 2.796736957 | ITGB1BP1 | 20 | 41421127 | 41421327 | -2.54548739 | CHD6 |
| 2 | 9490274 | 9490474 | 2.542892096 | ADAM17 | 20 | 41421731 | 41421979 | -2.067732248 | CHD6 |
| 2 | 9490274 | 9490474 | 2.542892096 | IAH1 | 20 | 41533236 | 41533442 | -1.454266536 | CHD6 |
| 2 | 10039822 | 10040069 | -1.775643359 | RP11-254F7.2 | 20 | 44068932 | 44069137 | -2.513549082 | TOX2 |
| 2 | 10200379 | 10200551 | -6.901258183 | C2orf48 | 20 | 45797376 | 45797575 | -4.221912642 | DNTTIP1 |
| 2 | 10206543 | 10206855 | -2.093844799 | C2orf48 | 1 | 150548624 | 150548758 | 1.833661778 | RP11-54A4.2 |
| 2 | 10210443 | 10210638 | -2.468410283 | C2orf48 | 20 | 45947684 | 45947917 | -1.47248067 | PCIF1 |
| 2 | 10211101 | 10211304 | -1.849684124 | C2orf48 | 20 | 46129071 | 46129271 | 9.587506106 | CD40 |
| 2 | 10399894 | 10400097 | 2.428425631 | HPCAL1 | 20 | 46541328 | 46541527 | 1.05975036 | OCSTAMP |
| 2 | 11640435 | 11640558 | 5.707342618 | GREB1 | 20 | 46686816 | 46687017 | -1.138726346 | TP53RK |
| 2 | 12740700 | 12740888 | -1.089032695 | TRIB2 | 20 | 47210499 | 47210835 | -1.372629752 | ZMYND8 |
| 2 | 17781480 | 17781696 | -2.506555644 | GEN1 | 20 | 47276608 | 47276806 | -1.925324765 | ZMYND8 |
| 2 | 17781480 | 17781696 | -2.506555644 | SMC6 | 20 | 47635909 | 47636109 | -1.232582498 | NCOA3 |
| 2 | 17783926 | 17784132 | -2.212134242 | SMC6 | 20 | 47636467 | 47636658 | -1.76682412 | NCOA3 |
| 2 | 17783926 | 17784132 | -2.212134242 | GEN1 | 20 | 48625322 | 48625527 | -2.877496135 | PREX1 |
| 2 | 19901443 | 19901589 | 1.142701109 | TTC32 | 20 | 49144368 | 49144647 | 6.656654108 | STAU1 |
| 2 | 20202484 | 20202680 | 4.660964047 | SDC1 | 20 | 49238174 | 49238374 | 5.97473099 | ZNFX1 |
| 2 | 20684299 | 20684497 | -1.041695374 | LDAH | 20 | 49238174 | 49238374 | 5.97473099 | DDX27 |
| 1 | 116405569 | 116405763 | -2.263290615 | ATP1A1-AS1 | 20 | 49247193 | 49247390 | 2.286005424 | ZNFX1 |
| 1 | 116405569 | 116405763 | -2.263290615 | ATP1A1 | 20 | 49248718 | 49248950 | 2.393288623 | ZNFX1 |
| 2 | 23927344 | 23927558 | -2.134730337 | UBXN2A | 20 | 49270103 | 49270367 | 2.085313859 | ZNFX1 |
| 2 | 23999678 | 23999896 | -1.023711866 | UBXN2A | 20 | 49271481 | 49271684 | 1.613098636 | ZNFX1 |
| 2 | 24037873 | 24038020 | -2.533458568 | MFSD2B | 20 | 49636181 | 49636421 | 1.41332437 | B4GALT5 |
| 2 | 24037873 | 24038020 | -2.533458568 | WDCP | 20 | 50192219 | 50192414 | 1.022180118 | CEBPB |
| 2 | 24038852 | 24039050 | -2.957028675 | WDCP | 20 | 50750321 | 50750511 | -1.825765059 | PARD6B |
| 2 | 24038852 | 24039050 | -2.957028675 | MFSD2B | 20 | 50891240 | 50891439 | -1.269239416 | ADNP |
| 2 | 24077498 | 24077652 | -1.064400272 | TP53I3 | 20 | 50894210 | 50894411 | -1.661480957 | ADNP |
| 2 | 24077498 | 24077652 | -1.064400272 | FAM228B | 20 | 50959758 | 50959957 | -1.016803736 | MOCS3 |
| 2 | 24100053 | 24100261 | -1.069280415 | FAM228B | 20 | 51003881 | 51004080 | -2.069567204 | RP5-955M13.4 |
| 2 | 24144600 | 24144811 | -1.931623842 | RP11-507M3.1 | 20 | 51003881 | 51004080 | -2.069567204 | KCNG1 |
| 2 | 24144600 | 24144811 | -1.931623842 | FAM228B | 20 | 51390958 | 51391179 | -1.636177762 | NFATC2 |
| 2 | 24819570 | 24819768 | -2.700587443 | CENPO | 20 | 51598656 | 51598860 | -1.42967234 | ATP9A |
| 2 | 24819570 | 24819768 | -2.700587443 | ADCY3 | 20 | 51600020 | 51600193 | -1.426107022 | ATP9A |
| 2 | 24819947 | 24820146 | -11.47421124 | CENPO | 20 | 51784451 | 51784557 | -3.560562247 | SALL4 |
| 2 | 24819947 | 24820146 | -11.47421124 | ADCY3 | 20 | 51791911 | 51792122 | -2.786763103 | SALL4 |
| 2 | 24824400 | 24824597 | -9.227350457 | ADCY3 | 20 | 52084471 | 52084656 | -2.192387937 | ZFP64 |
| 2 | 24826308 | 24826504 | -9.299258905 | ADCY3 | 20 | 52152765 | 52152986 | -1.742266713 | ZFP64 |
| 2 | 24826308 | 24826504 | -9.299258905 | RP11-443B20.1 | 20 | 53569048 | 53569336 | -1.1683248 | ZNF217 |
| 1 | 116525191 | 116525392 | 1.404390757 | CD58 | 20 | 53569048 | 53569336 | -1.1683248 | RP4-724E16.2 |
| 2 | 24830488 | 24830685 | -8.971394691 | ADCY3 | 20 | 53571783 | 53571979 | -1.363510354 | RP4-724E16.2 |
| 2 | 25234177 | 25234386 | 1.838985458 | DNMT3A | 20 | 53571783 | 53571979 | -1.363510354 | ZNF217 |
| 2 | 25326235 | 25326435 | 7.70937103 | DNMT3A | 20 | 53575934 | 53576132 | -1.613696127 | ZNF217 |
| 2 | 25414025 | 25414179 | -7.727470917 | DTNB | 20 | 53582826 | 53583010 | -1.555938029 | ZNF217 |
| 2 | 25744044 | 25744300 | -1.077368666 | ASXL2 | 20 | 56369971 | 56370170 | -3.778940193 | AURKA |
| 1 | 116526061 | 116526243 | 2.302601379 | CD58 | 20 | 56399061 | 56399262 | -2.434686759 | CSTF1 |
| 2 | 26188194 | 26188398 | -1.595537657 | GAREM2 | 20 | 56403480 | 56403677 | -1.70524072 | CSTF1 |
| 2 | 26311753 | 26311952 | -6.123963757 | EPT1 | 20 | 56452835 | 56453035 | -1.929787256 | CASS4 |
| 2 | 26311753 | 26311952 | -6.123963757 | ADGRF3 | 20 | 56458697 | 56458894 | -1.456224685 | CASS4 |
| 2 | 27082819 | 27083051 | -1.977172732 | EMILIN1 | 20 | 57371342 | 57371528 | 6.71090093 | RAE1 |
| 2 | 27083445 | 27083722 | -1.576376918 | EMILIN1 | 20 | 57407898 | 57408098 | -1.852171802 | RBM38 |
| 2 | 27085467 | 27085646 | -1.164338607 | EMILIN1 | 20 | 58703564 | 58703839 | -1.04245201 | STX16-NPEPL1 |
| 1 | 116536053 | 116536252 | -1.316902231 | CD58 | 20 | 58703564 | 58703839 | -1.04245201 | NPEPL1 |

| | | | | |
|---|---|---|---|---|
| 2 | 27152797 | 27153000 | -4.515971189 | TCF23 |
| 2 | 27264200 | 27264398 | 6.494342343 | SLC30A3 |
| 2 | 27264947 | 27265108 | 6.149604009 | SLC30A3 |
| 2 | 27326447 | 27326647 | -1.314475371 | GTF3C2 |
| 2 | 27357137 | 27357333 | -2.607981071 | AC074117.10 |
| 2 | 27358777 | 27359032 | -1.055334454 | AC074117.10 |
| 2 | 27359741 | 27359937 | -1.331924767 | AC074117.10 |
| 2 | 27381598 | 27381730 | -2.942897944 | PPM1G |
| 2 | 27393006 | 27393198 | -2.390800726 | PPM1G |
| 2 | 27393006 | 27393198 | -2.390800726 | FTH1P3 |
| 2 | 27581328 | 27581522 | -1.479514109 | C2orf16 |
| 2 | 27627402 | 27627595 | -3.082470019 | RP11-158I13.2 |
| 2 | 27627402 | 27627595 | -3.082470019 | CCDC121 |
| 2 | 27653497 | 27653636 | -1.017412965 | SUPT7L |
| 2 | 27657486 | 27657710 | -1.762472556 | SUPT7L |
| 2 | 27664019 | 27664218 | -1.353767855 | SLC4A1AP |
| 2 | 28331149 | 28331264 | -1.135704473 | BRE |
| 2 | 28332027 | 28332156 | -1.391638613 | BRE |
| 2 | 28332920 | 28333087 | -2.122821809 | BRE |
| 1 | 3862441 | 3862639 | -1.468194201 | DFFB |
| 2 | 28869800 | 28869996 | -1.40158689 | TRMT61B |
| 2 | 30158806 | 30159005 | 2.535855407 | YPEL5 |
| 2 | 30606883 | 30607014 | -6.743668808 | LCLAT1 |
| 2 | 32251423 | 32251620 | 1.362446381 | NLRC4 |
| 2 | 33527177 | 33527374 | 2.702212298 | RASGRP3 |
| 2 | 33558756 | 33558965 | 2.807646454 | RASGRP3 |
| 2 | 33558756 | 33558965 | 2.807646454 | AC020594.5 |
| 2 | 33562745 | 33562948 | 2.07344176 | AC020594.5 |
| 2 | 33562745 | 33562948 | 2.07344176 | RASGRP3 |
| 2 | 36548583 | 36548804 | 2.492446554 | FEZ2 |
| 2 | 36548583 | 36548804 | 2.492446554 | CRIM1 |
| 2 | 36808039 | 36808238 | -9.702437728 | VIT |
| 2 | 36814311 | 36814508 | -10.80129507 | VIT |
| 2 | 37228114 | 37228315 | -1.112255178 | CEBPZ |
| 2 | 37826425 | 37826625 | -3.548195007 | LINC00211 |
| 2 | 37827671 | 37827776 | -3.311595484 | LINC00211 |
| 1 | 116986875 | 116987073 | 1.478763958 | PTGFRN |
| 2 | 38492796 | 38493012 | 1.886024027 | AC016995.3 |
| 2 | 38709822 | 38710089 | 5.899640811 | GALM |
| 2 | 38734383 | 38734586 | 1.548023386 | GALM |
| 2 | 38781508 | 38781711 | -1.85613318 | GEMIN6 |
| 2 | 38861478 | 38861673 | -1.039332947 | DHX57 |
| 1 | 117025579 | 117025778 | 3.802118616 | CD101 |
| 1 | 117025579 | 117025778 | 3.802118616 | RP11-27K13.3 |
| 2 | 39498141 | 39498357 | 3.898150789 | AC007246.3 |
| 2 | 39599434 | 39599654 | 3.593277078 | AC007246.3 |
| 2 | 40164956 | 40165123 | 1.287388674 | SLC8A1-AS1 |
| 2 | 40164956 | 40165123 | 1.287388674 | SLC8A1 |
| 2 | 40177802 | 40178007 | 1.192074023 | SLC8A1-AS1 |
| 2 | 40177802 | 40178007 | 1.192074023 | SLC8A1 |
| 1 | 117075044 | 117075242 | -3.070080684 | TTF2 |
| 2 | 40428455 | 40428706 | 1.547644718 | SLC8A1 |
| 2 | 40428914 | 40429109 | 1.449213256 | SLC8A1 |
| 2 | 40430082 | 40430282 | 1.166897563 | SLC8A1 |
| 20 | 58991089 | 58991251 | 5.732443024 | NELFCD |
| 20 | 58995394 | 58995593 | 1.028002517 | CTSZ |
| 20 | 59774330 | 59774489 | -3.549454684 | PHACTR3 |
| 20 | 59805916 | 59806113 | -6.921764267 | PHACTR3 |
| 20 | 59944424 | 59944524 | -1.671336592 | FAM217B |
| 20 | 59944864 | 59945061 | -1.691084841 | FAM217B |
| 20 | 60057387 | 60057554 | -3.936914287 | C20orf197 |
| 20 | 60058202 | 60058306 | -3.277294426 | C20orf197 |
| 20 | 60097094 | 60097299 | -2.765495745 | MIR646HG |
| 20 | 60279158 | 60279365 | -2.972429223 | MIR646HG |
| 20 | 60322491 | 60322610 | -6.510576748 | MIR646HG |
| 20 | 62067076 | 62067275 | -1.999590957 | RP11-11M20.4 |
| 20 | 62286015 | 62286215 | -1.638013939 | OSBPL2 |
| 20 | 62295543 | 62295744 | -1.292033658 | OSBPL2 |
| 1 | 151006420 | 151006620 | -3.54342827 | FAM63A |
| 20 | 62399607 | 62399754 | -1.112164073 | CABLES2 |
| 20 | 62401882 | 62402003 | -3.284927804 | CABLES2 |
| 1 | 151033937 | 151034161 | -1.007207795 | PRUNE1 |
| 20 | 62402599 | 62402783 | 1.781681482 | CABLES2 |
| 20 | 62402599 | 62402783 | 1.781681482 | RP5-908M14.10 |
| 20 | 62602167 | 62602371 | -1.752220488 | WI2-87327B8.2 |
| 20 | 62603094 | 62603274 | -7.538825124 | WI2-87327B8.2 |
| 20 | 62656444 | 62656544 | -6.444466733 | SLCO4A1 |
| 20 | 62670165 | 62670287 | -5.724574323 | SLCO4A1 |
| 20 | 62710207 | 62710475 | -1.139525557 | NTSR1 |
| 20 | 62716459 | 62716684 | -1.74471532 | NTSR1 |
| 20 | 62722573 | 62722712 | 5.218355771 | NTSR1 |
| 20 | 62724925 | 62725107 | -1.237840141 | NTSR1 |
| 20 | 62746593 | 62746789 | 4.39395128 | NTSR1 |
| 1 | 6418626 | 6418837 | -1.025907621 | HES2 |
| 20 | 62760212 | 62760406 | -6.062383768 | NTSR1 |
| 20 | 62762161 | 62762362 | -2.455748842 | NTSR1 |
| 20 | 62799886 | 62800088 | -1.526261804 | MRGBP |
| 20 | 62812984 | 62813182 | 1.843460675 | OGFR |
| 20 | 62841893 | 62842026 | -6.833112001 | TCFL5 |
| 20 | 62844309 | 62844508 | -1.946955226 | TCFL5 |
| 20 | 62852674 | 62852908 | -1.256800184 | TCFL5 |
| 20 | 62909853 | 62910054 | -1.335279587 | DIDO1 |
| 20 | 62944864 | 62945062 | -1.092767733 | GID8 |
| 20 | 63564817 | 63564985 | 2.7983492 | HELZ2 |
| 20 | 63565713 | 63565961 | 2.983628415 | HELZ2 |
| 20 | 63669386 | 63669583 | 5.94208526 | RTEL1-TNFRSF6B |
| 20 | 63669386 | 63669583 | 5.94208526 | RTEL1 |
| 20 | 63682552 | 63682748 | 2.680373672 | RTEL1 |
| 20 | 63682552 | 63682748 | 2.680373672 | RTEL1-TNFRSF6B |
| 20 | 63686356 | 63686578 | 1.622109392 | RTEL1-TNFRSF6B |
| 20 | 63686356 | 63686578 | 1.622109392 | RTEL1 |
| 20 | 63708746 | 63708973 | 1.255487805 | RP4-583P15.15 |
| 20 | 63708746 | 63708973 | 1.255487805 | ZGPAT |
| 1 | 151165524 | 151165662 | -1.6202668 | SCNM1 |
| 1 | 151165524 | 151165662 | -1.6202668 | LYSMD1 |
| 20 | 63743657 | 63743852 | 6.444371624 | ZBTB46 |
| 20 | 63746987 | 63747195 | 7.595606736 | ZBTB46 |
| 20 | 63770421 | 63770588 | -6.218355771 | ZBTB46 |

| | | | | |
|---|---|---|---|---|
| 1 | 117118299 | 117118526 | -3.483147377 | TRIM45 |
| 2 | 43574557 | 43574675 | -1.245443029 | THADA |
| 2 | 44595708 | 44595866 | -2.61913951 | CAMKMT |
| 2 | 44944962 | 44945181 | -2.891050187 | SIX3 |
| 2 | 45006122 | 45006313 | -2.805596405 | SIX2 |
| 2 | 45389355 | 45389581 | -1.040075664 | SRBD1 |
| 2 | 45599601 | 45599855 | -2.290534964 | SRBD1 |
| 2 | 46184853 | 46185006 | 1.705538656 | PRKCE |
| 2 | 46512241 | 46512458 | -1.995438763 | ATP6V1E2 |
| 2 | 46519094 | 46519206 | -2.334935384 | ATP6V1E2 |
| 2 | 46526574 | 46526761 | -2.004823208 | ATP6V1E2 |
| 2 | 46541652 | 46541860 | -2.065293427 | RHOQ |
| 2 | 46541652 | 46541860 | -2.065293427 | ATP6V1E2 |
| 2 | 46580923 | 46581127 | 1.74113913 | PIGF |
| 2 | 46580923 | 46581127 | 1.74113913 | RHOQ |
| 1 | 117402103 | 117402269 | -2.802657972 | MAN1A2 |
| 2 | 46603662 | 46603833 | 7.411355947 | PIGF |
| 2 | 46604443 | 46604642 | 7.448922728 | PIGF |
| 2 | 47799224 | 47799423 | -2.174394516 | MSH6 |
| 2 | 47799224 | 47799423 | -2.174394516 | FBXO11 |
| 2 | 47800789 | 47800990 | -2.189978261 | FBXO11 |
| 2 | 47800789 | 47800990 | -2.189978261 | MSH6 |
| 2 | 47803320 | 47803510 | -8.554078438 | FBXO11 |
| 2 | 47803320 | 47803510 | -8.554078438 | MSH6 |
| 2 | 48346298 | 48346491 | -1.164711113 | FOXN2 |
| 2 | 53853240 | 53853547 | -1.054324353 | GPR75-ASB3 |
| 2 | 53853240 | 53853547 | -1.054324353 | GPR75 |
| 2 | 53853882 | 53854006 | -6.410089481 | GPR75 |
| 2 | 53853882 | 53854006 | -6.410089481 | GPR75-ASB3 |
| 2 | 54558587 | 54558792 | -1.229876216 | SPTBN1 |
| 1 | 3890684 | 3890890 | -1.307613671 | C1orf174 |
| 1 | 117623660 | 117623918 | 1.818400716 | FAM46C |
| 2 | 55224387 | 55224572 | -1.128509061 | CLHC1 |
| 2 | 55224387 | 55224572 | -1.128509061 | AC012358.4 |
| 2 | 55617531 | 55617755 | -1.111175763 | PPP4R3B |
| 2 | 56193091 | 56193291 | 1.614468835 | CCDC85A |
| 2 | 56193091 | 56193291 | 1.614468835 | RP11-482H16.1 |
| 2 | 58170650 | 58170816 | 4.95419631 | FANCL |
| 2 | 58173748 | 58173947 | 3.357122759 | FANCL |
| 2 | 60461832 | 60462034 | -2.729595449 | BCL11A |
| 2 | 61040304 | 61040496 | 1.12251218 | PEX13 |
| 2 | 61048506 | 61048699 | 1.208487566 | PEX13 |
| 2 | 61190542 | 61190718 | -1.072704572 | USP34 |
| 2 | 61190542 | 61190718 | -1.072704572 | AHSA2 |
| 2 | 61306063 | 61306266 | -7.691244468 | USP34 |
| 2 | 61537995 | 61538195 | -1.304054507 | XPO1 |
| 2 | 62146315 | 62146501 | -1.59965108 | RPSAP26 |
| 2 | 62146315 | 62146501 | -1.59965108 | AC018462.2 |
| 2 | 62146315 | 62146501 | -1.59965108 | COMMD1 |
| 2 | 62146890 | 62147064 | -2.216055506 | COMMD1 |
| 2 | 62146890 | 62147064 | -2.216055506 | RPSAP26 |
| 2 | 62706905 | 62707104 | -1.985192005 | EHBP1 |
| 2 | 62840966 | 62841144 | -2.31841587 | AC007098.1 |
| 2 | 62840966 | 62841144 | -2.31841587 | EHBP1 |
| 20 | 63774902 | 63775053 | 2.739621407 | ZBTB46 |
| 20 | 63797874 | 63798148 | -1.234875697 | ZBTB46 |
| 1 | 151281939 | 151282135 | 1.116086721 | ZNF687 |
| 20 | 63865527 | 63865728 | 8.831733883 | TPD52L2 |
| 20 | 63919980 | 63920182 | 2.081752914 | DNAJC5 |
| 20 | 64063476 | 64063666 | 6.806261491 | TCEA2 |
| 20 | 64098888 | 64099081 | -4.398218123 | OPRL1 |
| 20 | 64273237 | 64273524 | -1.665474824 | PCMTD2 |
| 21 | 5097341 | 5097570 | -1.120650456 | CH507-9B2.3 |
| 21 | 5103991 | 5104198 | 1.515612164 | CH507-9B2.3 |
| 21 | 5120794 | 5120993 | 1.48438443 | CH507-9B2.3 |
| 21 | 5120794 | 5120993 | 1.48438443 | CH507-9B2.4 |
| 21 | 9790533 | 9790715 | -3.63339327 | MGC39584 |
| 21 | 9809677 | 9809865 | -5.018086806 | MGC39584 |
| 21 | 14076128 | 14076320 | 5.564641508 | AP001347.6 |
| 21 | 14143700 | 14143902 | -1.697964338 | AP001347.6 |
| 21 | 14143700 | 14143902 | -1.697964338 | LIPI |
| 21 | 14143700 | 14143902 | -1.697964338 | ERLEC1P1 |
| 21 | 14319936 | 14320131 | -4.474183616 | ABCC13 |
| 21 | 14842287 | 14842428 | -1.04843077 | AF127936.7 |
| 21 | 14842287 | 14842428 | -1.04843077 | AF127936.9 |
| 21 | 17796665 | 17796887 | 2.309895333 | AL109761.5 |
| 21 | 17796665 | 17796887 | 2.309895333 | C21orf91 |
| 21 | 24477217 | 24477383 | 1.0103788 | AP000476.1 |
| 21 | 24489571 | 24489738 | 2.335264805 | AP000476.1 |
| 21 | 25035245 | 25035406 | -1.310692954 | AP000235.2 |
| 1 | 151874315 | 151874518 | -1.108482652 | THEM4 |
| 21 | 25768942 | 25769184 | -1.174318651 | GABPA |
| 21 | 26924417 | 26924543 | 8.114409345 | ADAMTS5 |
| 21 | 26924417 | 26924543 | 8.114409345 | AP001601.2 |
| 21 | 26965390 | 26965534 | 4.372480501 | ADAMTS5 |
| 21 | 28874792 | 28875004 | 1.369975962 | N6AMT1 |
| 21 | 29298394 | 29298592 | 3.678304585 | BACH1 |
| 21 | 31266314 | 31266434 | 3.114409345 | TIAM1 |
| 21 | 32314602 | 32314802 | 1.073347484 | MRAP |
| 21 | 32314602 | 32314802 | 1.073347484 | URB1 |
| 21 | 32316630 | 32316867 | 1.043249079 | URB1 |
| 21 | 32393491 | 32393680 | -3.541146458 | URB1-AS1 |
| 21 | 32759184 | 32759376 | -1.757306781 | PAXBP1 |
| 1 | 153774482 | 153774807 | -1.955353318 | INTS3 |
| 1 | 153774482 | 153774807 | -1.955353318 | SLC27A3 |
| 21 | 32759739 | 32759946 | -2.071693989 | PAXBP1 |
| 21 | 33027792 | 33028102 | -1.722748405 | AP000282.2 |
| 21 | 33027792 | 33028102 | -1.722748405 | OLIG2 |
| 21 | 33309108 | 33309297 | -7.247646463 | IL10RB |
| 21 | 33969216 | 33969412 | -1.981465589 | LINC00649 |
| 21 | 33976984 | 33977184 | -1.363269111 | LINC00649 |
| 1 | 153819655 | 153819852 | -2.691259037 | GATAD2B |
| 21 | 34182869 | 34183054 | -2.860812524 | AP000320.7 |
| 21 | 34182869 | 34183054 | -2.860812524 | LINC00310 |
| 21 | 34449265 | 34449398 | -1.505467739 | KCNE1 |
| 21 | 34496826 | 34497026 | -2.372885561 | KCNE1 |
| 21 | 34498411 | 34498536 | -3.221057061 | KCNE1 |
| 21 | 34520638 | 34520774 | -6.239005586 | RCAN1 |

| | | | | |
|---|---|---|---|---|
| 2 | 62948338 | 62948511 | -2.192929327 | EHBP1 |
| 2 | 63056322 | 63056519 | -1.888725172 | OTX1 |
| 2 | 63196686 | 63196919 | -1.293464435 | WDPCP |
| 2 | 63212600 | 63212787 | -1.956442282 | WDPCP |
| 2 | 63215537 | 63215727 | -2.376505718 | WDPCP |
| 1 | 117959271 | 117959472 | -1.927015261 | WDR3 |
| 1 | 117959271 | 117959472 | -1.927015261 | SPAG17 |
| 2 | 63275802 | 63276009 | -1.575381372 | WDPCP |
| 2 | 63307698 | 63307897 | -2.408179955 | WDPCP |
| 2 | 63308425 | 63308620 | -1.652653029 | WDPCP |
| 2 | 63588439 | 63588643 | -1.71466305 | WDPCP |
| 2 | 63588439 | 63588643 | -1.71466305 | MDH1 |
| 2 | 64094987 | 64095192 | 2.677537233 | PELI1 |
| 1 | 118883895 | 118884111 | 3.383373224 | TBX15 |
| 2 | 64458487 | 64458682 | 2.278924153 | LGALSL |
| 2 | 64523165 | 64523354 | 2.101189358 | RP11-568N6.1 |
| 2 | 64634358 | 64634577 | 1.94573332 | SERTAD2 |
| 2 | 64635938 | 64636139 | 2.117670241 | SERTAD2 |
| 2 | 64999920 | 65000100 | -7.452740513 | SLC1A4 |
| 2 | 65021210 | 65021411 | 1.246424868 | SLC1A4 |
| 1 | 119032982 | 119033185 | -1.251935892 | WARS2 |
| 2 | 65070790 | 65070989 | 2.257267801 | CEP68 |
| 2 | 65070790 | 65070989 | 2.257267801 | RAB1A |
| 2 | 65313775 | 65313978 | 1.079700589 | SPRED2 |
| 2 | 65500930 | 65501155 | -4.269148364 | AC074391.1 |
| 2 | 65500930 | 65501155 | -4.269148364 | AC007389.1 |
| 2 | 67403519 | 67403792 | -1.424705106 | ETAA1 |
| 2 | 68130829 | 68131026 | -2.556021779 | WDR92 |
| 2 | 68131100 | 68131286 | -1.547859405 | RP11-474G23.1 |
| 2 | 68131100 | 68131286 | -1.547859405 | WDR92 |
| 1 | 119179417 | 119179599 | -2.740147611 | RP11-418J17.1 |
| 2 | 68734943 | 68735152 | 1.600052643 | ARHGAP25 |
| 2 | 68822566 | 68822776 | 1.644268366 | ARHGAP25 |
| 2 | 68826212 | 68826413 | 1.316833906 | ARHGAP25 |
| 2 | 69741319 | 69741493 | -3.459377617 | ANXA4 |
| 1 | 119180228 | 119180446 | -2.324461116 | RP11-418J17.1 |
| 2 | 69937997 | 69938204 | 2.762700273 | MXD1 |
| 2 | 70087687 | 70087887 | -1.027169087 | PCBP1-AS1 |
| 2 | 70087687 | 70087887 | -1.027169087 | PCBP1 |
| 2 | 70181456 | 70181558 | -1.02552169 | C2orf42 |
| 2 | 70267285 | 70267388 | -6.086838568 | PCYOX1 |
| 2 | 70277228 | 70277425 | -1.24873349 | PCYOX1 |
| 2 | 70900991 | 70901190 | -2.439149218 | VAX2 |
| 2 | 70902082 | 70902281 | -1.117565767 | VAX2 |
| 2 | 70927800 | 70928056 | -2.570104762 | VAX2 |
| 1 | 119182968 | 119183190 | -1.453098315 | RP11-418J17.1 |
| 2 | 70933053 | 70933252 | -2.285089342 | VAX2 |
| 2 | 71133252 | 71133454 | -1.418014823 | MPHOSPH10 |
| 2 | 72891387 | 72891528 | -2.37172777 | SPR |
| 1 | 6013397 | 6013584 | -4.378185497 | KCNAB2 |
| 1 | 119212400 | 119212677 | -1.280859108 | RP11-418J17.1 |
| 2 | 73258729 | 73258934 | -1.165760078 | FBXO41 |
| 2 | 73424598 | 73424796 | -1.743204479 | ALMS1 |
| 2 | 73448103 | 73448271 | -1.684677433 | ALMS1 |
| 21 | 34846925 | 34847116 | 1.655992646 | RUNX1 |
| 21 | 34968020 | 34968215 | 1.206579037 | RUNX1 |
| 1 | 153828062 | 153828263 | -1.611891134 | GATAD2B |
| 21 | 36054483 | 36054686 | -1.464423772 | SETD4 |
| 21 | 36072626 | 36072838 | -1.834463029 | SETD4 |
| 21 | 36072626 | 36072838 | -1.834463029 | AP000688.14 |
| 21 | 36072626 | 36072838 | -1.834463029 | CBR1 |
| 21 | 36230598 | 36230801 | -1.234603405 | DOPEY2 |
| 21 | 36936549 | 36936749 | -1.759337635 | HLCS |
| 21 | 37114747 | 37114963 | -1.443674811 | TTC3 |
| 1 | 153852057 | 153852259 | -1.580801275 | RP11-216N14.7 |
| 1 | 153852057 | 153852259 | -1.580801275 | GATAD2B |
| 21 | 37143190 | 37143387 | -6.162371555 | TTC3 |
| 21 | 37241223 | 37241519 | 5.451187557 | DSCR3 |
| 1 | 153852936 | 153853136 | -3.193772784 | RP11-216N14.7 |
| 1 | 153852936 | 153853136 | -3.193772784 | GATAD2B |
| 21 | 38822824 | 38823026 | 2.394778224 | ETS2 |
| 21 | 39196378 | 39196638 | -1.67529254 | BRWD1 |
| 21 | 39197348 | 39197507 | -2.200195362 | BRWD1 |
| 21 | 39199005 | 39199209 | -2.177491013 | BRWD1 |
| 21 | 41377009 | 41377197 | 10.63085322 | MX2 |
| 21 | 41377542 | 41377730 | 5.522545354 | MX2 |
| 21 | 41390109 | 41390283 | 2.570291458 | MX2 |
| 21 | 41888854 | 41889172 | -1.979890194 | C2CD2 |
| 21 | 41952326 | 41952525 | -1.988639962 | C2CD2 |
| 21 | 42580361 | 42580565 | 2.222497237 | SLC37A1 |
| 21 | 42849327 | 42849530 | 1.052866218 | WDR4 |
| 21 | 43724314 | 43724506 | -8.206813965 | PDXK |
| 1 | 154402607 | 154402799 | 3.059742994 | RP11-350G8.5 |
| 21 | 43954751 | 43954935 | 8.062666372 | AGPAT3 |
| 21 | 43966755 | 43967022 | 8.123370299 | AGPAT3 |
| 21 | 44082870 | 44083068 | -1.020357509 | TRAPPC10 |
| 21 | 44227251 | 44227453 | 1.171854101 | ICOSLG |
| 21 | 44233649 | 44233848 | 4.573602462 | ICOSLG |
| 21 | 44329366 | 44329549 | 1.091825119 | C21orf2 |
| 21 | 44329366 | 44329549 | 1.091825119 | AP001062.8 |
| 21 | 44330310 | 44330557 | 2.534772635 | C21orf2 |
| 21 | 44351164 | 44351341 | 5.979638753 | TRPM2 |
| 21 | 44353513 | 44353713 | 5.826198431 | TRPM2 |
| 21 | 44441749 | 44441955 | -1.336697661 | TRPM2 |
| 1 | 154484128 | 154484295 | -7.30158134 | SHE |
| 21 | 44457556 | 44457760 | 1.433937141 | LRRC3 |
| 21 | 44934375 | 44934550 | 1.221642398 | LINC01547 |
| 21 | 44938891 | 44939089 | -1.209873959 | LINC01547 |
| 21 | 45144630 | 45144741 | -1.229720221 | ADARB1 |
| 21 | 45162625 | 45162844 | 1.898557869 | ADARB1 |
| 21 | 45176188 | 45176346 | -1.402752761 | ADARB1 |
| 21 | 45192603 | 45192802 | -6.287769623 | ADARB1 |
| 21 | 45222098 | 45222293 | -1.201891795 | ADARB1 |
| 21 | 45226110 | 45226304 | -2.005213526 | ADARB1 |
| 21 | 45296047 | 45296279 | -1.158694468 | LINC00205 |
| 21 | 45793389 | 45793583 | 3.183161107 | PCBP3 |
| 21 | 45852791 | 45852991 | 1.339049893 | PCBP3 |
| 21 | 46129313 | 46129463 | 1.906179729 | COL6A2 |

| | | | | | | | | | |
|---|---|---|---|---|---|---|---|---|---|
| 2 | 73448574 | 73448898 | -1.931682625 | ALMS1 | 21 | 46321353 | 46321550 | -6.84821212 | C21orf58 |
| 2 | 73451288 | 73451486 | -1.603835458 | ALMS1 | 21 | 46322512 | 46322720 | -2.035039218 | C21orf58 |
| 2 | 73453054 | 73453235 | -1.106633418 | ALMS1 | 21 | 46323065 | 46323268 | -2.102361666 | C21orf58 |
| 2 | 73453777 | 73453973 | -2.276700446 | ALMS1 | 21 | 46363446 | 46363644 | -1.613011396 | PCNT |
| 2 | 73491220 | 73491415 | -1.416957469 | ALMS1 | 21 | 46416146 | 46416453 | -1.479403601 | PCNT |
| 2 | 73519642 | 73519831 | -1.303268298 | ALMS1 | 21 | 46567789 | 46567985 | -1.339357904 | DIP2A |
| 2 | 73572404 | 73572617 | -1.069724868 | ALMS1 | 21 | 46658914 | 46659113 | -1.646677339 | PRMT2 |
| 2 | 74046525 | 74046724 | -1.069336613 | TET3 | 22 | 15854579 | 15854764 | -1.276216383 | NBEAP3 |
| 2 | 74048136 | 74048336 | -1.931106196 | TET3 | 22 | 15854579 | 15854764 | -1.276216383 | LA16c-60H5.7 |
| 2 | 74120680 | 74120816 | -1.247057496 | FNBP1P1 | 22 | 16648730 | 16648919 | -2.057910379 | TPTEP1 |
| 1 | 119623793 | 119623970 | 2.89324146 | ZNF697 | 22 | 16650728 | 16650885 | -2.233719599 | TPTEP1 |
| 2 | 74130762 | 74130909 | -1.935064915 | RP11-287D1.4 | 22 | 16665306 | 16665518 | -1.940536527 | ANKRD62P1-PARP4P3 |
| 2 | 74418205 | 74418357 | -2.850292647 | C2orf81 | 22 | 16665306 | 16665518 | -1.940536527 | TPTEP1 |
| 2 | 74418205 | 74418357 | -2.850292647 | HMGA1P8 | 22 | 17058301 | 17058559 | -1.767494122 | CECR7 |
| 2 | 74474260 | 74474468 | -1.960901153 | CCDC142 | 22 | 17070078 | 17070191 | -2.458972892 | AC006946.16 |
| 2 | 74481891 | 74482094 | -1.3990618 | CCDC142 | 22 | 17114771 | 17114987 | -2.434256239 | IL17RA |
| 2 | 74506814 | 74507003 | 2.222278293 | PCGF1 | 22 | 17119362 | 17119595 | -1.978973435 | CECR6 |
| 1 | 119624026 | 119624226 | 2.606662028 | ZNF697 | 22 | 17552947 | 17553186 | -1.671625932 | CECR2 |
| 2 | 74832838 | 74833038 | -2.747533113 | RP11-259N19.1 | 22 | 17638260 | 17638439 | 1.107668832 | BCL2L13 |
| 2 | 74834148 | 74834335 | -8.427781004 | HK2 | 22 | 17777430 | 17777598 | 3.696158711 | LINC00528 |
| 2 | 74930666 | 74930846 | -2.009245646 | LINC01291 | 22 | 17790074 | 17790273 | -1.411308482 | XXbac-B461K10.4 |
| 1 | 119625931 | 119626156 | 2.648553604 | ZNF697 | 22 | 17790074 | 17790273 | -1.411308482 | MICAL3 |
| 2 | 85326608 | 85326806 | -10.35379163 | TGOLN2 | 22 | 17790768 | 17791048 | -1.274895472 | XXbac-B461K10.4 |
| 2 | 85327222 | 85327428 | -1.290826172 | TGOLN2 | 22 | 17790768 | 17791048 | -1.274895472 | MICAL3 |
| 2 | 85544828 | 85544986 | -1.116477967 | MAT2A | 22 | 17818360 | 17818526 | 2.026380072 | MICAL3 |
| 2 | 85544828 | 85544986 | -1.116477967 | GGCX | 22 | 18606695 | 18606895 | -4.974913355 | RIMBP3 |
| 2 | 85839400 | 85839599 | 2.831750287 | ST3GAL5 | 22 | 18607188 | 18607366 | -6.826198431 | RIMBP3 |
| 2 | 85840100 | 85840296 | 3.607409281 | ST3GAL5 | 22 | 18608655 | 18608853 | -3.580430047 | RIMBP3 |
| 2 | 85848106 | 85848213 | 4.096488846 | ST3GAL5 | 22 | 19030555 | 19030655 | 1.64815849 | DGCR5 |
| 2 | 85851245 | 85851451 | 2.396534378 | ST3GAL5 | 22 | 19131299 | 19131468 | -1.398026623 | DGCR14 |
| 2 | 85889528 | 85889729 | 3.83974005 | ST3GAL5-AS1 | 22 | 19131299 | 19131468 | -1.398026623 | TSSK2 |
| 2 | 86020937 | 86021228 | -1.467993207 | POLR1A | 22 | 19179651 | 19179775 | -2.516147828 | CLTCL1 |
| 2 | 86021744 | 86021943 | -1.728915131 | POLR1A | 22 | 19189488 | 19189638 | -4.564641508 | CLTCL1 |
| 2 | 86027229 | 86027450 | -2.031007422 | POLR1A | 22 | 19432038 | 19432192 | 7.586174685 | MRPL40 |
| 2 | 86195752 | 86195912 | -6.812626111 | RP11-301O19.1 | 22 | 19432038 | 19432192 | 7.586174685 | C22orf39 |
| 2 | 86195752 | 86195912 | -6.812626111 | IMMT | 22 | 19432038 | 19432192 | 7.586174685 | HIRA |
| 2 | 86212533 | 86212737 | -1.831201648 | MRPL35 | 22 | 19444046 | 19444244 | -1.527236988 | HIRA |
| 2 | 86604257 | 86604440 | 1.12462811 | RNF103 | 22 | 19444046 | 19444244 | -1.527236988 | C22orf39 |
| 2 | 86604257 | 86604440 | 1.12462811 | AC015971.2 | 22 | 19722515 | 19722715 | -1.207617239 | Sep-05 |
| 2 | 86604257 | 86604440 | 1.12462811 | RNF103-CHMP3 | 22 | 19724485 | 19724688 | -1.295868381 | GP1BB |
| 2 | 87584752 | 87585031 | -1.425831533 | LINC00152 | 22 | 19724485 | 19724688 | -1.295868381 | Sep-05 |
| 2 | 88574948 | 88575058 | 1.208384209 | AC104134.2 | 22 | 19758701 | 19758899 | 2.801018007 | TBX1 |
| 2 | 88574948 | 88575058 | 1.208384209 | EIF2AK3 | 22 | 19782968 | 19783165 | -1.474183616 | TBX1 |
| 1 | 120433844 | 120434027 | 6.627514285 | PFN1P2 | 22 | 19784801 | 19784999 | -1.502411428 | GNB1L |
| 2 | 91628528 | 91628684 | -2.581574382 | AC027612.6 | 22 | 19786335 | 19786473 | -2.374575566 | GNB1L |
| 2 | 95158931 | 95159086 | -2.441449314 | ZNF514 | 22 | 19791927 | 19792110 | -2.044364837 | GNB1L |
| 2 | 95182159 | 95182373 | -2.248112566 | ZNF2 | 22 | 19796704 | 19796898 | -1.231198997 | GNB1L |
| 1 | 120454378 | 120454537 | 3.142701109 | NBPF8 | 22 | 19801137 | 19801381 | -1.060656187 | GNB1L |
| 2 | 95816163 | 95816351 | -1.600941376 | LINC00342 | 22 | 19801748 | 19801946 | -1.550880169 | GNB1L |
| 2 | 95838958 | 95839128 | -1.093245804 | ANKRD36C | 1 | 155018235 | 155018414 | -8.241214921 | ZBTB7B |
| 2 | 95855737 | 95856030 | -1.970859786 | ANKRD36C | 22 | 19803061 | 19803299 | -1.787846082 | GNB1L |
| 2 | 95933196 | 95933391 | -7.767485771 | ANKRD36C | 22 | 19804163 | 19804356 | -3.809359595 | GNB1L |
| 2 | 96186632 | 96186835 | -1.118905797 | STARD7 | 22 | 19851987 | 19852301 | -1.223762276 | GNB1L |
| 2 | 96240218 | 96240387 | 1.271447464 | STARD7-AS1 | 22 | 19851987 | 19852301 | -1.223762276 | C22orf29 |

| | | | | |
|---|---|---|---|---|
| 2 | 96370814 | 96370992 | -1.7190346 | NCAPH |
| 2 | 96552339 | 96552538 | 1.442385015 | ARID5A |
| 2 | 96593606 | 96593811 | -1.554359427 | KANSL3 |
| 2 | 96706972 | 96707213 | -1.80610634 | LMAN2L |
| 2 | 97245384 | 97245558 | -1.626932237 | ANKRD36 |
| 2 | 97427578 | 97427762 | 1.480147263 | AC159540.2 |
| 2 | 97427578 | 97427762 | 1.480147263 | AC159540.1 |
| 2 | 97427764 | 97427911 | 1.251912869 | AC159540.2 |
| 2 | 97427764 | 97427911 | 1.251912869 | AC159540.1 |
| 2 | 97429511 | 97429710 | 1.723236243 | AC159540.2 |
| 2 | 97429511 | 97429710 | 1.723236243 | AC159540.1 |
| 2 | 97432736 | 97432932 | 1.351899365 | AC159540.2 |
| 2 | 97432736 | 97432932 | 1.351899365 | AC159540.1 |
| 2 | 97702615 | 97702909 | 2.95289549 | LINC01125 |
| 2 | 98570095 | 98570302 | 1.222343176 | INPP4A |
| 2 | 98582789 | 98582954 | 1.414632091 | INPP4A |
| 2 | 99125117 | 99125298 | -2.232866333 | TSGA10 |
| 2 | 99162605 | 99162805 | -1.325917872 | LIPT1 |
| 2 | 99162605 | 99162805 | -1.325917872 | MITD1 |
| 2 | 99162605 | 99162805 | -1.325917872 | RP11-111H13.1 |
| 2 | 99162605 | 99162805 | -1.325917872 | RP11-38C17.1 |
| 2 | 99201024 | 99201200 | -2.575796589 | RP11-111H13.1 |
| 2 | 99201500 | 99201728 | -1.346778797 | RP11-111H13.1 |
| 2 | 99361093 | 99361302 | -1.482936619 | EIF5B |
| 2 | 99363525 | 99363721 | -8.622339862 | EIF5B |
| 2 | 99384594 | 99384717 | 4.114409345 | EIF5B |
| 2 | 99399319 | 99399522 | -1.310383996 | EIF5B |
| 2 | 99438871 | 99439092 | -1.139439425 | REV1 |
| 2 | 99551307 | 99551490 | -2.914225379 | AFF3 |
| 2 | 100392056 | 100392252 | -1.829957871 | CHST10 |
| 2 | 100392691 | 100392910 | -1.137731399 | CHST10 |
| 2 | 100393222 | 100393377 | -1.299023237 | CHST10 |
| 2 | 101147574 | 101147758 | -7.361403766 | TBC1D8 |
| 2 | 101152477 | 101152649 | -1.467920763 | TBC1D8 |
| 2 | 101152477 | 101152649 | -1.467920763 | RP11-299H21.1 |
| 2 | 101274129 | 101274327 | 1.087826366 | RNF149 |
| 2 | 101381765 | 101381962 | 4.890679857 | CREG2 |
| 2 | 101457983 | 101458184 | -1.092088856 | RFX8 |
| 2 | 105307768 | 105308012 | -1.140882554 | TGFBRAP1 |
| 2 | 105342928 | 105343128 | -1.275904429 | C2orf49 |
| 2 | 105892979 | 105893102 | 1.245641985 | NCK2 |
| 2 | 106160177 | 106160373 | 2.135382845 | UXS1 |
| 1 | 120985257 | 120985451 | -1.373179728 | LINC00623 |
| 2 | 108468791 | 108468897 | -6.961431552 | GCC2 |
| 2 | 108499151 | 108499346 | -2.290079922 | GCC2 |
| 2 | 108765049 | 108765255 | 1.413004717 | RANBP2 |
| 2 | 108765759 | 108765959 | 1.250751158 | RANBP2 |
| 1 | 121096108 | 121096309 | -2.36086554 | RP11-439A17.9 |
| 1 | 121096108 | 121096309 | -2.36086554 | RP11-439A17.10 |
| 1 | 121096108 | 121096309 | -2.36086554 | FCGR1B |
| 2 | 108767621 | 108767822 | 1.01906949 | RANBP2 |
| 2 | 108783519 | 108783715 | 2.745816494 | RANBP2 |
| 2 | 108877503 | 108877699 | -1.695125774 | CCDC138 |
| 2 | 111119308 | 111119504 | 3.590998917 | BCL2L11 |

| | | | | |
|---|---|---|---|---|
| 22 | 20086249 | 20086428 | 1.20773681 | DGCR8 |
| 22 | 20241617 | 20241813 | 6.6862285 | RTN4R |
| 22 | 20308727 | 20308880 | -5.840399517 | XXbac-B444P24.10 |
| 22 | 20405499 | 20405721 | -1.352720556 | ZNF74 |
| 22 | 20406820 | 20406973 | -2.104866425 | ZNF74 |
| 22 | 20953244 | 20953398 | -1.146758019 | CRKL |
| 22 | 20953727 | 20953921 | -1.737839425 | CRKL |
| 1 | 155069065 | 155069268 | -1.132763647 | EFNA4 |
| 1 | 155069065 | 155069268 | -1.132763647 | RP11-540D14.8 |
| 22 | 21388428 | 21388632 | -2.734905785 | RIMBP3B |
| 1 | 155338264 | 155338414 | -1.650424913 | ASH1L |
| 22 | 22488443 | 22488648 | -2.824817156 | ZNF280B |
| 22 | 22547902 | 22548089 | -9.140333286 | PRAME |
| 22 | 22549943 | 22550142 | -1.844474759 | PRAME |
| 22 | 23181824 | 23182021 | -1.040231679 | BCR |
| 22 | 23315644 | 23315816 | -1.341551796 | BCR |
| 22 | 23743407 | 23743622 | -2.090683645 | ZNF70 |
| 22 | 24166743 | 24167013 | -1.155580904 | CABIN1 |
| 22 | 24373565 | 24373760 | -2.34447204 | SPECC1L-ADORA2A |
| 22 | 24373565 | 24373760 | -2.34447204 | SPECC1L |
| 22 | 24401175 | 24401335 | 9.977656805 | SPECC1L-ADORA2A |
| 22 | 24401175 | 24401335 | 9.977656805 | SPECC1L |
| 22 | 24422657 | 24422856 | 7.881113149 | SPECC1L-ADORA2A |
| 22 | 24422657 | 24422856 | 7.881113149 | ADORA2A |
| 22 | 24441683 | 24441880 | 4.489951993 | ADORA2A-AS1 |
| 22 | 24441683 | 24441880 | 4.489951993 | ADORA2A |
| 22 | 24441683 | 24441880 | 4.489951993 | SPECC1L-ADORA2A |
| 1 | 155459834 | 155460034 | -1.567848593 | ASH1L |
| 22 | 25478239 | 25478366 | -1.92736645 | CTA-390C10.10 |
| 22 | 25478239 | 25478366 | -1.92736645 | CRYBB2P1 |
| 22 | 26433684 | 26433851 | 3.997644139 | ASPHD2 |
| 22 | 26434191 | 26434369 | 2.781354847 | ASPHD2 |
| 22 | 26442939 | 26443114 | -3.362721862 | ASPHD2 |
| 1 | 155468176 | 155468348 | -3.107085957 | ASH1L |
| 22 | 26453128 | 26453327 | -2.551643904 | HPS4 |
| 22 | 26464163 | 26464364 | -2.921522057 | HPS4 |
| 22 | 26464419 | 26464547 | -1.765733577 | HPS4 |
| 22 | 26481901 | 26482101 | -1.528509583 | HPS4 |
| 22 | 26514057 | 26514312 | -7.066611251 | CTA-445C9.14 |
| 22 | 27921998 | 27922170 | -2.186804523 | TTC28-AS1 |
| 22 | 27921998 | 27922170 | -2.186804523 | TTC28-AS1_2 |
| 22 | 27924319 | 27924521 | -1.552685319 | TTC28-AS1 |
| 22 | 27999416 | 27999611 | -1.323972041 | TTC28-AS1 |
| 22 | 27999416 | 27999611 | -1.323972041 | TTC28 |
| 22 | 28000500 | 28000668 | -4.080007727 | TTC28-AS1 |
| 22 | 28000500 | 28000668 | -4.080007727 | TTC28 |
| 22 | 28001066 | 28001254 | -6.390679857 | TTC28 |
| 22 | 28001066 | 28001254 | -6.390679857 | TTC28-AS1 |
| 22 | 28105602 | 28105792 | -1.118601376 | TTC28 |
| 22 | 28387193 | 28387330 | -1.253092694 | TTC28 |
| 22 | 28390613 | 28390811 | -2.854425093 | TTC28 |
| 22 | 28506481 | 28506623 | -2.164400486 | TTC28 |
| 22 | 29142187 | 29142314 | 1.597754564 | KREMEN1 |
| 22 | 29694827 | 29695023 | -8.118847729 | NF2 |

| | | | | |
|---|---|---|---|---|
| 2 | 111246751 | 111246945 | -1.130752659 | MIR4435-2HG |
| 2 | 111323487 | 111323682 | -3.390808428 | RP11-803D5.1 |
| 2 | 111323487 | 111323682 | -3.390808428 | MIR4435-2HG |
| 2 | 111767933 | 111768122 | -2.582128037 | ANAPC1 |
| 2 | 112028622 | 112028891 | 11.54562007 | MERTK |
| 1 | 121184041 | 121184252 | -3.008428422 | FAM72B |
| 2 | 112575148 | 112575350 | -2.245471774 | POLR1B |
| 2 | 112659352 | 112659556 | 2.39638755 | SLC20A1 |
| 2 | 112736670 | 112736840 | -2.800190497 | NT5DC4 |
| 2 | 112736670 | 112736840 | -2.800190497 | CKAP2L |
| 2 | 112737116 | 112737311 | -3.051989845 | NT5DC4 |
| 2 | 112737116 | 112737311 | -3.051989845 | CKAP2L |
| 2 | 112752381 | 112752586 | -2.579712975 | CKAP2L |
| 2 | 112756055 | 112756279 | -2.59118689 | CKAP2L |
| 2 | 113132873 | 113133073 | -7.807124864 | IL1RN |
| 2 | 113218176 | 113218385 | 1.701215988 | PAX8-AS1 |
| 2 | 113218176 | 113218385 | 1.701215988 | PAX8 |
| 2 | 113489561 | 113489761 | -2.972531043 | CBWD2 |
| 2 | 113598740 | 113598938 | 1.338555284 | WASH2P |
| 2 | 114000281 | 114000479 | 2.912935205 | LINC01191 |
| 2 | 119245499 | 119245727 | -1.81418715 | STEAP3 |
| 2 | 119245499 | 119245727 | -1.81418715 | STEAP3-AS1 |
| 2 | 119263342 | 119263537 | -1.549231878 | STEAP3 |
| 2 | 119681779 | 119681983 | -1.96124306 | TMEM177 |
| 1 | 121406335 | 121406489 | -1.606823189 | CH17-437K3.1 |
| 2 | 120350042 | 120350244 | 1.102473797 | INHBB |
| 2 | 121650888 | 121651081 | -1.890638741 | NIFK-AS1 |
| 2 | 127307626 | 127307810 | 2.067919689 | MAP3K2 |
| 2 | 127504826 | 127505029 | -1.090660518 | IWS1 |
| 2 | 127504826 | 127505029 | -1.090660518 | AC010976.2 |
| 2 | 127717208 | 127717408 | -1.302542684 | WDR33 |
| 2 | 127720027 | 127720230 | -1.023342171 | WDR33 |
| 2 | 127846781 | 127846985 | -1.427444441 | POLR2D |
| 2 | 127846781 | 127846985 | -1.427444441 | AC006011.4 |
| 2 | 127847890 | 127848131 | -1.571447012 | POLR2D |
| 2 | 128189802 | 128190002 | -1.137608175 | UGGT1 |
| 2 | 130152355 | 130152558 | -2.372054129 | SMPD4 |
| 2 | 130373464 | 130373738 | -2.498032398 | PTPN18 |
| 2 | 130374019 | 130374190 | -1.827479663 | PTPN18 |
| 2 | 131723688 | 131723879 | 1.654013639 | C2orf27A |
| 2 | 131723688 | 131723879 | 1.654013639 | AC093724.2 |
| 2 | 134333117 | 134333277 | -5.178225985 | MGAT5 |
| 2 | 134866479 | 134866689 | -1.784039218 | CCNT2-AS1 |
| 2 | 134866479 | 134866689 | -1.784039218 | ACMSD |
| 2 | 134867307 | 134867660 | -2.350362948 | ACMSD |
| 2 | 134867307 | 134867660 | -2.350362948 | CCNT2-AS1 |
| 2 | 135207553 | 135207698 | -1.363176226 | ZRANB3 |
| 2 | 135480996 | 135481164 | -3.207564196 | ZRANB3 |
| 2 | 135481571 | 135481797 | -2.014455329 | ZRANB3 |
| 2 | 135483641 | 135483787 | -6.968962742 | ZRANB3 |
| 2 | 135774850 | 135775045 | 1.144375496 | UBXN4 |
| 1 | 143883527 | 143883717 | -2.142167698 | HIST2H2BB |
| 1 | 143883527 | 143883717 | -2.142167698 | FCGR1CP |
| 1 | 143883527 | 143883717 | -2.142167698 | RP5-998N21.4 |

| | | | | |
|---|---|---|---|---|
| 22 | 30019779 | 30019980 | -1.441835149 | RP3-394A18.1 |
| 22 | 30019779 | 30019980 | -1.441835149 | MTMR3 |
| 22 | 30019779 | 30019980 | -1.441835149 | HORMAD2-AS1 |
| 22 | 30025728 | 30025924 | -1.092443398 | HORMAD2-AS1 |
| 22 | 30025728 | 30025924 | -1.092443398 | MTMR3 |
| 22 | 30025728 | 30025924 | -1.092443398 | RP3-394A18.1 |
| 22 | 30241071 | 30241261 | 4.139606592 | LIF |
| 22 | 30263910 | 30264115 | -4.204579858 | OSM |
| 22 | 30375451 | 30375651 | -2.088940863 | CCDC157 |
| 22 | 30375451 | 30375651 | -2.088940863 | RNF215 |
| 22 | 30375771 | 30375962 | -2.838859821 | CCDC157 |
| 22 | 30375771 | 30375962 | -2.838859821 | RNF215 |
| 22 | 30377944 | 30378145 | -1.651241972 | CCDC157 |
| 22 | 30377944 | 30378145 | -1.651241972 | RNF215 |
| 22 | 30903743 | 30903936 | -1.194172247 | OSBP2 |
| 22 | 30925967 | 30926120 | -2.071302198 | MORC2-AS1 |
| 22 | 30925967 | 30926120 | -2.071302198 | MORC2 |
| 22 | 30967924 | 30968091 | -1.029152574 | MORC2 |
| 1 | 155563227 | 155563449 | 1.290566349 | ASH1L-AS1 |
| 22 | 31276907 | 31277139 | 4.76564908 | LIMK2 |
| 22 | 31278726 | 31278870 | 4.044693491 | LIMK2 |

| | | | | |
|---|---|---|---|---|
| 2 | 136115539 | 136115744 | -1.302207077 | CXCR4 |
| 2 | 143006244 | 143006432 | -5.623370299 | KYNU |
| 2 | 143008097 | 143008300 | 2.134244418 | KYNU |
| 2 | 143010996 | 143011096 | 2.831585667 | KYNU |
| 1 | 6065562 | 6065819 | -1.846842499 | KCNAB2 |
| 2 | 143328355 | 143328569 | -1.385051763 | ARHGAP15 |
| 2 | 143328355 | 143328569 | -1.385051763 | RP11-570L15.2 |
| 2 | 143328355 | 143328569 | -1.385051763 | AC096558.1 |
| 2 | 147927190 | 147927392 | 2.983260673 | ACVR2A |
| 2 | 148092196 | 148092333 | -4.79248125 | MBD5 |
| 2 | 149587437 | 149587596 | -1.093845144 | MMADHC |
| 2 | 149587437 | 149587596 | -1.093845144 | AC144449.1 |
| 2 | 150469961 | 150470214 | 5.373023676 | RND3 |
| 2 | 151251351 | 151251509 | 1.371007727 | RBM43 |
| 2 | 151463178 | 151463377 | -1.884731072 | RIF1 |
| 2 | 151463972 | 151464176 | -1.403529605 | RIF1 |
| 2 | 151464542 | 151464752 | -1.45378361 | RIF1 |
| 2 | 151474914 | 151475246 | -1.495108298 | RIF1 |
| 2 | 151486190 | 151486386 | -1.599689238 | NEB |
| 2 | 151486190 | 151486386 | -1.599689238 | RIF1 |
| 2 | 152389689 | 152389892 | 8.151480077 | FMNL2 |
| 1 | 144423049 | 144423246 | 1.987089363 | NBPF15 |
| 2 | 152647809 | 152648009 | 8.028397006 | FMNL2 |
| 2 | 156325641 | 156325920 | 3.439175595 | NR4A2 |
| 2 | 156329793 | 156329993 | 2.869984041 | NR4A2 |
| 2 | 157415845 | 157416055 | 5.125653722 | CYTIP |
| 2 | 157737303 | 157737509 | 1.375639228 | ACVR1 |
| 2 | 158625119 | 158625320 | -1.839676739 | PKP4 |
| 2 | 158682686 | 158682848 | -3.022314152 | AC005042.4 |
| 2 | 158682686 | 158682848 | -3.022314152 | PKP4 |
| 2 | 159349863 | 159350037 | -1.787960122 | BAZ2B |
| 2 | 159499270 | 159499427 | -1.461234861 | BAZ2B |
| 2 | 159616109 | 159616271 | -1.224430844 | AC009506.1 |
| 2 | 159616109 | 159616271 | -1.224430844 | BAZ2B |
| 1 | 145096124 | 145096264 | -2.534836764 | FAM72D |
| 2 | 159797285 | 159797484 | -1.463149149 | CD302 |
| 2 | 159797285 | 159797484 | -1.463149149 | LY75-CD302 |
| 2 | 159902624 | 159902832 | -9.312037164 | LY75 |
| 2 | 159902624 | 159902832 | -9.312037164 | LY75-CD302 |
| 2 | 161501466 | 161501612 | -3.282755069 | SLC4A10 |
| 2 | 161501466 | 161501612 | -3.282755069 | AC009487.6 |
| 2 | 162308078 | 162308278 | -6.991853096 | IFIH1 |
| 2 | 162309529 | 162309663 | -1.489290508 | IFIH1 |
| 2 | 162310834 | 162311035 | 1.14621205 | IFIH1 |
| 2 | 162318079 | 162318290 | 1.92672655 | IFIH1 |
| 2 | 163610870 | 163611045 | -1.656268793 | FIGN |
| 2 | 168236459 | 168236640 | -2.82480773 | STK39 |
| 2 | 169081594 | 169081795 | -2.760505563 | DHRS9 |
| 2 | 169083251 | 169083441 | -3.042745284 | DHRS9 |
| 2 | 169095603 | 169095841 | -4.291020121 | DHRS9 |
| 2 | 169546386 | 169546572 | -1.471250292 | FASTKD1 |
| 2 | 169700921 | 169701131 | -1.547496233 | PHOSPHO2 |
| 2 | 169700921 | 169701131 | -1.547496233 | KLHL23 |
| 2 | 169735821 | 169736015 | -4.73931192 | KLHL23 |

| | | | | |
|---|---|---|---|---|
| 2 | 169749393 | 169749563 | -3.062798985 | KLHL23 |
| 1 | 145168729 | 145168928 | -1.243118679 | RP11-403I13.8 |
| 2 | 171091928 | 171092129 | 1.504441546 | TLK1 |
| 2 | 171324130 | 171324332 | -1.658656159 | METTL8 |
| 2 | 172080444 | 172080589 | -2.662913587 | METAP1D |
| 2 | 172081914 | 172082070 | -3.34591214 | METAP1D |
| 2 | 172085468 | 172085669 | -2.786006897 | DLX1 |
| 2 | 172088737 | 172088939 | -2.782222461 | DLX1 |
| 2 | 172504243 | 172504461 | 5.036156044 | ITGA6 |
| 2 | 172504243 | 172504461 | 5.036156044 | AC093818.1 |
| 2 | 172527271 | 172527467 | -6.386157287 | AC093818.1 |
| 2 | 172529119 | 172529220 | -5.291541384 | AC093818.1 |
| 2 | 172530057 | 172530240 | -4.065870558 | AC093818.1 |
| 2 | 173266496 | 173266705 | -2.402533822 | AC013461.1 |
| 2 | 173266496 | 173266705 | -2.402533822 | MLK7-AS1 |
| 2 | 173910041 | 173910236 | -1.54419917 | SP3 |
| 2 | 174335556 | 174335662 | 5.033044595 | SP9 |
| 2 | 174335556 | 174335662 | 5.033044595 | RP11-572N21.1 |
| 2 | 174337187 | 174337391 | -1.767818208 | RP11-572N21.1 |
| 2 | 174337187 | 174337391 | -1.767818208 | SP9 |
| 1 | 145660334 | 145660482 | -2.058205306 | GPR89A |
| 2 | 175930005 | 175930217 | -1.060056233 | LNPK |
| 2 | 177221630 | 177221812 | -3.414918393 | HNRNPA3 |
| 2 | 177227516 | 177227729 | 1.615234957 | NFE2L2 |
| 2 | 177354738 | 177354913 | -1.028437261 | AC074286.1 |
| 2 | 177354738 | 177354913 | -1.028437261 | NFE2L2 |
| 1 | 145660874 | 145661111 | -2.146390875 | GPR89A |
| 2 | 177369243 | 177369394 | -2.187559093 | NFE2L2 |
| 2 | 177369243 | 177369394 | -2.187559093 | AC074286.1 |
| 2 | 177371884 | 177372062 | -7.153671568 | AC074286.1 |
| 2 | 177371884 | 177372062 | -7.153671568 | NFE2L2 |
| 2 | 178539026 | 178539243 | -1.654473257 | TTN-AS1 |
| 2 | 178539026 | 178539243 | -1.654473257 | TTN |
| 2 | 178542172 | 178542333 | -1.796265523 | TTN-AS1 |
| 2 | 178542172 | 178542333 | -1.796265523 | TTN |
| 2 | 178542671 | 178542866 | -1.505831459 | TTN |
| 2 | 178542671 | 178542866 | -1.505831459 | TTN-AS1 |
| 2 | 178546130 | 178546323 | -2.460968979 | TTN-AS1 |
| 2 | 178546130 | 178546323 | -2.460968979 | TTN |
| 2 | 178548240 | 178548437 | -2.986151502 | TTN-AS1 |
| 2 | 178548240 | 178548437 | -2.986151502 | TTN |
| 2 | 178576862 | 178577061 | -1.079542492 | TTN-AS1 |
| 2 | 178576862 | 178577061 | -1.079542492 | TTN |
| 1 | 146058682 | 146058846 | 1.285770992 | RP11-458D21.1 |
| 2 | 181535533 | 181535664 | -1.38774053 | ITGA4 |
| 2 | 181915398 | 181915505 | -1.039308308 | SSFA2 |
| 2 | 184752195 | 184752409 | 1.36219432 | ZNF804A |
| 2 | 187519403 | 187519565 | 5.94208526 | AC007319.1 |
| 2 | 187519403 | 187519565 | 5.94208526 | TFPI |
| 2 | 189563898 | 189564130 | -2.639255534 | SLC40A1 |
| 2 | 189665989 | 189666227 | -1.137843879 | ASNSD1 |
| 2 | 189667233 | 189667472 | -1.20784067 | ASNSD1 |
| 2 | 189670398 | 189670599 | -1.373387652 | ASNSD1 |
| 2 | 189854507 | 189854705 | -3.164898169 | PMS1 |

| | | | | |
|---|---|---|---|---|
| 2 | 189863760 | 189863950 | -1.639558626 | PMS1 |
| 2 | 190200042 | 190200234 | 2.951187557 | HIBCH |
| 2 | 190200042 | 190200234 | 2.951187557 | C2orf88 |
| 2 | 190473542 | 190473737 | 5.903677461 | MFSD6 |
| 2 | 190476764 | 190476946 | 7.286882757 | MFSD6 |
| 2 | 190500163 | 190500300 | 8.42110764 | MFSD6 |
| 2 | 190509044 | 190509194 | -2.106594044 | MFSD6 |
| 2 | 190509044 | 190509194 | -2.106594044 | NEMP2 |
| 2 | 190557328 | 190557489 | -2.353762395 | AC093388.3 |
| 2 | 190659682 | 190659885 | -1.359979821 | NAB1 |
| 2 | 190955365 | 190955565 | 8.615760605 | GLS |
| 2 | 190956109 | 190956308 | 5.689951177 | GLS |
| 2 | 190959186 | 190959389 | 4.023741531 | GLS |
| 2 | 191696245 | 191696444 | 6.087930401 | NABP1 |
| 2 | 196752852 | 196752956 | 2.857122759 | CCDC150 |
| 2 | 197260274 | 197260460 | 6.351627367 | ANKRD44 |
| 2 | 197260274 | 197260460 | 6.351627367 | ANKRD44-IT1 |
| 2 | 197260274 | 197260460 | 6.351627367 | AC013264.1 |
| 2 | 197260971 | 197261190 | 1.668054494 | ANKRD44-IT1 |
| 2 | 197260971 | 197261190 | 1.668054494 | ANKRD44 |
| 2 | 197260971 | 197261190 | 1.668054494 | AC013264.1 |
| 1 | 147172812 | 147173136 | -1.316262975 | RP11-337C18.8 |
| 2 | 199459428 | 199459627 | -2.01874657 | SATB2-AS1 |
| 2 | 199459428 | 199459627 | -2.01874657 | SATB2 |
| 2 | 199925510 | 199925669 | -1.770268536 | C2orf69 |
| 2 | 199965398 | 199965612 | -1.797579781 | C2orf47 |
| 2 | 200571382 | 200571585 | -3.111214654 | SGO2 |
| 2 | 200573329 | 200573527 | -3.304670765 | SGO2 |
| 2 | 200892047 | 200892250 | -1.445812561 | NIF3L1 |
| 1 | 6222964 | 6223168 | -1.748582837 | ICMT |
| 2 | 201116734 | 201116935 | -1.216878348 | CFLAR |
| 2 | 201117337 | 201117540 | -1.613422755 | CFLAR |
| 2 | 201138656 | 201138849 | -8.092437671 | IMPDH1P10 |
| 2 | 201138656 | 201138849 | -8.092437671 | CFLAR |
| 2 | 201169393 | 201169517 | 1.024263664 | CFLAR |
| 2 | 201246485 | 201246664 | -2.826972504 | CASP8 |
| 2 | 202036384 | 202036584 | 1.124655786 | FZD7 |
| 2 | 202555801 | 202555902 | 1.326648778 | BMPR2 |
| 2 | 202559847 | 202560012 | 1.448640461 | BMPR2 |
| 1 | 147203716 | 147203898 | -2.669853179 | RP11-337C18.10 |
| 1 | 147203716 | 147203898 | -2.669853179 | FMO5 |
| 1 | 147203716 | 147203898 | -2.669853179 | CCT8P1 |
| 1 | 147203716 | 147203898 | -2.669853179 | RP11-337C18.8 |
| 2 | 202765963 | 202766162 | -1.884965703 | FAM117B |
| 2 | 202925001 | 202925195 | -2.044606601 | CARF |
| 2 | 202925001 | 202925195 | -2.044606601 | AC010900.2 |
| 2 | 202925001 | 202925195 | -2.044606601 | WDR12 |
| 2 | 203107739 | 203107925 | -1.036063757 | NBEAL1 |
| 2 | 203439474 | 203439723 | 4.415036972 | ABI2 |
| 2 | 203439474 | 203439723 | 4.415036972 | RAPH1 |
| 2 | 203440177 | 203440484 | 8.816902051 | RAPH1 |
| 2 | 203440177 | 203440484 | 8.816902051 | ABI2 |
| 1 | 147205809 | 147206052 | -1.586275235 | RP11-337C18.8 |
| 1 | 147205809 | 147206052 | -1.586275235 | RP11-337C18.10 |

| | | | | |
|---|---|---|---|---|
| 1 | 147205809 | 147206052 | -1.586275235 | FMO5 |
| 2 | 206766683 | 206766891 | -1.423621481 | FASTKD2 |
| 2 | 207123871 | 207124089 | -1.085162479 | KLF7 |
| 2 | 207186747 | 207186935 | 1.73057815 | AC007879.2 |
| 2 | 207186747 | 207186935 | 1.73057815 | AC007879.1 |
| 2 | 207239835 | 207239978 | 1.994135354 | AC007879.7 |
| 2 | 207239835 | 207239978 | 1.994135354 | AC007879.5 |
| 2 | 207605096 | 207605242 | -2.541464812 | METTL21A |
| 2 | 207613149 | 207613349 | -2.228181741 | METTL21A |
| 2 | 210153688 | 210153885 | -1.074917026 | KANSL1L |
| 2 | 211383722 | 211383959 | 5.917644096 | ERBB4 |
| 2 | 212237642 | 212237820 | 6.50167111 | ERBB4 |
| 2 | 214728842 | 214729019 | -2.87177702 | BARD1 |
| 2 | 214780753 | 214780953 | -1.846680731 | BARD1 |
| 1 | 147619317 | 147619597 | -1.027718195 | BCL9 |
| 2 | 214963210 | 214963406 | -2.778950793 | ABCA12 |
| 2 | 214963210 | 214963406 | -2.778950793 | AC072062.1 |
| 2 | 214965327 | 214965479 | -1.969103683 | ABCA12 |
| 2 | 214965969 | 214966164 | -2.485822194 | ABCA12 |
| 2 | 215367861 | 215368058 | -3.494342343 | FN1 |
| 2 | 215944722 | 215944909 | 1.871612292 | MREG |
| 2 | 218035050 | 218035241 | -3.877595279 | RUFY4 |
| 2 | 218068701 | 218068913 | -1.978400084 | RUFY4 |
| 2 | 218069294 | 218069432 | -2.163566627 | RUFY4 |
| 2 | 218090224 | 218090408 | -1.690364724 | RUFY4 |
| 2 | 218135159 | 218135266 | -8.487033615 | CXCR2 |
| 2 | 218136683 | 218136880 | -5.539931129 | CXCR2 |
| 2 | 218345325 | 218345512 | 5.363853889 | CATIP-AS2 |
| 2 | 218345325 | 218345512 | 5.363853889 | PNKD |
| 2 | 218388622 | 218388813 | 1.789938372 | SLC11A1 |
| 2 | 218454743 | 218454907 | -1.081688863 | USP37 |
| 1 | 147983136 | 147983353 | -1.047513657 | GPR89B |
| 2 | 218638154 | 218638380 | -1.12650861 | ZNF142 |
| 2 | 218873906 | 218874104 | -2.264326739 | WNT6 |
| 2 | 219234804 | 219235000 | -1.215561883 | ANKZF1 |
| 2 | 219250974 | 219251182 | -1.852927531 | TUBA4A |
| 1 | 856737 | 856914 | 2.14179094 | LINC01128 |
| 1 | 6223811 | 6224010 | -1.463832791 | ICMT |
| 2 | 222318595 | 222318815 | 1.288350156 | AC010980.2 |
| 2 | 222318595 | 222318815 | 1.288350156 | RP11-384O8.1 |
| 2 | 222646777 | 222646985 | -1.050973825 | FARSB |
| 2 | 223598540 | 223598756 | 4.575245909 | SCG2 |
| 2 | 224000803 | 224000976 | 5.537315648 | SERPINE2 |
| 2 | 224810757 | 224810955 | 2.465164015 | DOCK10 |
| 2 | 224811471 | 224811670 | 1.557935159 | DOCK10 |
| 2 | 226837076 | 226837277 | -1.259562776 | RHBDD1 |
| 2 | 226931329 | 226931513 | -6.37462047 | RHBDD1 |
| 2 | 226939930 | 226940191 | -1.027354587 | RHBDD1 |
| 2 | 230169075 | 230169274 | 3.126608435 | SP110 |
| 2 | 230169075 | 230169274 | 3.126608435 | AC009950.2 |
| 2 | 230177339 | 230177535 | 2.146792897 | AC009950.1 |
| 2 | 230177339 | 230177535 | 2.146792897 | SP110 |
| 2 | 230196800 | 230197001 | 3.992931035 | AC009950.1 |
| 2 | 230196800 | 230197001 | 3.992931035 | SP110 |

| | | | | |
|---|---|---|---|---|
| 2 | 230514902 | 230515024 | -7.04290219 | SP100 |
| 2 | 230514902 | 230515024 | -7.04290219 | HMGB1P3 |
| 1 | 148334772 | 148334956 | 2.411818161 | RP11-495P10.3 |
| 2 | 230544072 | 230544273 | 2.317581779 | SP100 |
| 2 | 230544072 | 230544273 | 2.317581779 | AC010149.4 |
| 2 | 230713788 | 230713947 | -8.340536831 | CAB39 |
| 2 | 230803019 | 230803198 | 4.857122759 | CAB39 |
| 2 | 230804183 | 230804379 | 1.913851514 | CAB39 |
| 2 | 231108663 | 231108786 | 4.114409345 | HTR2B |
| 2 | 231108663 | 231108786 | 4.114409345 | PSMD1 |
| 2 | 231371882 | 231372074 | 8.412522863 | ARMC9 |
| 2 | 231373964 | 231374114 | 6.847396577 | ARMC9 |
| 2 | 231397793 | 231397970 | -3.29020873 | B3GNT7 |
| 2 | 231481726 | 231481924 | -2.228401653 | NCL |
| 2 | 231524879 | 231525098 | 2.315095877 | NMUR1 |
| 2 | 231559960 | 231560160 | 6.924995722 | AC104634.3 |
| 2 | 231779195 | 231779382 | -5.998060182 | PDE6D |
| 2 | 232260010 | 232260265 | -1.396290214 | DIS3L2 |
| 2 | 232340480 | 232340679 | -2.548485874 | DIS3L2 |
| 2 | 232344008 | 232344163 | -1.844039121 | DIS3L2 |
| 2 | 232549772 | 232549965 | -1.133888109 | TIGD1 |
| 2 | 232581015 | 232581223 | -1.396411019 | AC073254.1 |
| 2 | 232581015 | 232581223 | -1.396411019 | EIF4E2 |
| 2 | 233063378 | 233063571 | -8.005679154 | INPP5D |
| 2 | 233088651 | 233088857 | -1.869984041 | INPP5D |
| 2 | 233840664 | 233840862 | -3.785928693 | HJURP |
| 2 | 233841707 | 233841945 | -3.141919771 | HJURP |
| 2 | 234495737 | 234495903 | 1.384024443 | ARL4C |
| 2 | 236125390 | 236125583 | -1.551364362 | AGAP1 |
| 2 | 236165694 | 236165864 | 1.603185085 | GBX2 |
| 2 | 237084406 | 237084622 | -1.32025587 | AC105760.2 |
| 1 | 148362837 | 148363059 | 1.241459986 | LINC01138 |
| 2 | 237574501 | 237574682 | 3.502344323 | RAB17 |
| 2 | 237581793 | 237582037 | 3.823369349 | RAB17 |
| 2 | 238151206 | 238151386 | 1.01538587 | KLHL30 |
| 1 | 148459432 | 148459605 | 3.005095765 | AC245100.1 |
| 1 | 148459432 | 148459605 | 3.005095765 | LINC01138 |
| 2 | 238329022 | 238329227 | -1.437506058 | TRAF3IP1 |
| 2 | 239052409 | 239052607 | -1.151365507 | HDAC4 |
| 2 | 239124271 | 239124471 | -10.12541116 | HDAC4 |
| 2 | 239160434 | 239160601 | -3.108819446 | HDAC4 |
| 2 | 239308705 | 239308894 | -1.079935668 | HDAC4 |
| 1 | 6224826 | 6225026 | -1.516166504 | ICMT |
| 2 | 240466224 | 240466420 | 1.36377269 | GPC1 |
| 2 | 240496558 | 240496881 | -1.03755471 | ANKMY1 |
| 2 | 240561692 | 240561888 | -1.26494405 | DUSP28 |
| 2 | 240561692 | 240561888 | -1.26494405 | ANKMY1 |
| 1 | 148953347 | 148953533 | -2.124850452 | PDE4DIP |
| 2 | 241126910 | 241127121 | -3.620523072 | PASK |
| 2 | 241227599 | 241227794 | -1.260075034 | HDLBP |
| 2 | 241231782 | 241231959 | -9.213304371 | HDLBP |
| 2 | 241396174 | 241396373 | -6.625371265 | FARP2 |
| 1 | 148981374 | 148981566 | -1.344518808 | PDE4DIP |
| 2 | 241563190 | 241563398 | -1.019521519 | BOK |

| | | | | |
|---|---|---|---|---|
| 2 | 241672663 | 241672859 | -1.224295801 | ATG4B |
| 2 | 241677706 | 241677896 | 1.071739511 | DTYMK |
| 2 | 241686049 | 241686206 | 6.318765276 | DTYMK |
| 2 | 241689897 | 241690131 | -1.353739934 | ING5 |
| 2 | 241721124 | 241721321 | -1.456233747 | ING5 |
| 2 | 241725043 | 241725275 | -1.203765185 | AC114730.11 |
| 2 | 241725043 | 241725275 | -1.203765185 | ING5 |
| 2 | 241728244 | 241728443 | -1.153033736 | ING5 |
| 2 | 241975290 | 241975434 | -5.012965257 | LINC01237 |
| 2 | 241975290 | 241975434 | -5.012965257 | AC093642.3 |
| 20 | 298205 | 298418 | -1.143479021 | ZCCHC3 |
| 20 | 326900 | 327100 | -1.310290947 | NRSN2-AS1 |
| 20 | 326900 | 327100 | -1.310290947 | SOX12 |
| 20 | 327901 | 328104 | -2.101903702 | NRSN2-AS1 |
| 20 | 327901 | 328104 | -2.101903702 | SOX12 |
| 20 | 511808 | 511966 | -1.568878675 | CSNK2A1 |
| 1 | 149176229 | 149176400 | 3.844125155 | CH17-189H20.1 |
| 20 | 845442 | 845807 | -1.641645488 | FAM110A |
| 20 | 1155582 | 1155800 | 1.181619647 | PSMF1 |
| 20 | 1540886 | 1541085 | -5.415319221 | RP4-576H24.4 |
| 20 | 1540886 | 1541085 | -5.415319221 | SIRPD |
| 20 | 1540886 | 1541085 | -5.415319221 | RP4-576H24.2 |
| 20 | 1541663 | 1541813 | -4.421309865 | SIRPD |
| 20 | 1541663 | 1541813 | -4.421309865 | RP4-576H24.2 |
| 20 | 1541663 | 1541813 | -4.421309865 | RP4-576H24.4 |
| 20 | 1565159 | 1565352 | -4.519835855 | RP4-576H24.4 |
| 20 | 1565159 | 1565352 | -4.519835855 | SIRPB1 |
| 20 | 1937736 | 1937942 | 1.306550449 | SIRPA |
| 20 | 2144277 | 2144454 | -1.215902576 | STK35 |
| 20 | 2483587 | 2483872 | -1.119417052 | RP4-734P14.4 |
| 20 | 2483587 | 2483872 | -1.119417052 | ZNF343 |
| 20 | 3047730 | 3047987 | -1.524297098 | MRPS26 |
| 20 | 3148873 | 3149071 | -1.168636017 | UBOX5 |
| 20 | 3148873 | 3149071 | -1.168636017 | FASTKD5 |
| 20 | 3148873 | 3149071 | -1.168636017 | UBOX5-AS1 |
| 1 | 149648579 | 149648779 | -1.89777549 | LINC00869 |
| 20 | 3165037 | 3165237 | 3.602460211 | LZTS3 |
| 20 | 3647867 | 3648041 | -1.602945522 | ATRN |
| 20 | 3648381 | 3648537 | -1.227364917 | ATRN |
| 20 | 3687654 | 3687894 | 8.400892515 | SIGLEC1 |
| 20 | 3866192 | 3866383 | -2.526219908 | MAVS |
| 20 | 3917000 | 3917196 | -1.224742123 | PANK2 |
| 1 | 149676703 | 149676876 | -1.929483862 | LINC00869 |
| 1 | 149676703 | 149676876 | -1.929483862 | RP11-353N4.5 |
| 20 | 4182176 | 4182368 | 2.064834339 | SMOX |
| 20 | 4699532 | 4699734 | 1.561633259 | PRNP |
| 20 | 4782835 | 4783042 | -5.226940444 | RASSF2 |
| 20 | 4784022 | 4784220 | -3.855539119 | RASSF2 |
| 20 | 4856944 | 4857149 | 1.17902961 | SLC23A2 |
| 20 | 6768007 | 6768207 | -4.321928095 | BMP2 |
| 20 | 6768513 | 6768694 | 1.280555928 | BMP2 |
| 20 | 6778566 | 6778766 | 1.805150857 | BMP2 |
| 20 | 10639604 | 10639803 | 1.881565055 | JAG1 |
| 20 | 17967968 | 17968191 | -8.380100076 | OVOL2 |

| | | | | |
|---|---|---|---|---|
| 20 | 17967968 | 17968191 | -8.380100076 | SNX5 |
| 20 | 17970048 | 17970246 | -2.334528806 | MGME1 |
| 20 | 17970048 | 17970246 | -2.334528806 | OVOL2 |
| 20 | 17990165 | 17990364 | -1.092212286 | MGME1 |
| 20 | 17990165 | 17990364 | -1.092212286 | OVOL2 |
| 20 | 18024673 | 18024842 | -5.938200917 | OVOL2 |
| 20 | 18142131 | 18142327 | -1.996880944 | KAT14 |
| 20 | 18142131 | 18142327 | -1.996880944 | PET117 |
| 20 | 18142644 | 18142869 | -2.332493008 | PET117 |
| 20 | 18142644 | 18142869 | -2.332493008 | KAT14 |
| 20 | 18162651 | 18162852 | -1.306442682 | KAT14 |
| 20 | 18181567 | 18181761 | -2.044567802 | KAT14 |
| 1 | 149791320 | 149791522 | -1.662392626 | RP11-196G18.3 |
| 1 | 149791320 | 149791522 | -1.662392626 | FCGR1A |
| 1 | 149791320 | 149791522 | -1.662392626 | HIST2H2BF |
| 20 | 18384228 | 18384426 | -5.734121925 | DZANK1 |
| 20 | 20000632 | 20000853 | 10.58217298 | RIN2 |
| 20 | 20391442 | 20391617 | -3.264568607 | RALGAPA2 |
| 20 | 20392890 | 20393082 | -1.311394735 | RALGAPA2 |
| 20 | 20512787 | 20512987 | -1.74570549 | RALGAPA2 |
| 20 | 21093412 | 21093553 | -3.363739025 | LINC00237 |
| 1 | 149792226 | 149792375 | 5.229715809 | HIST2H2BF |
| 1 | 149792226 | 149792375 | 5.229715809 | FCGR1A |
| 1 | 149792226 | 149792375 | 5.229715809 | RP11-196G18.3 |
| 20 | 21101054 | 21101227 | -7.512786084 | LINC00237 |
| 20 | 21102441 | 21102657 | -2.780664277 | LINC00237 |
| 20 | 21103356 | 21103563 | -1.940584612 | LINC00237 |
| 20 | 21167005 | 21167154 | 5.761780978 | KIZ |
| 20 | 21167005 | 21167154 | 5.761780978 | KIZ-AS1 |
| 20 | 21168946 | 21169166 | -1.032955401 | KIZ |
| 20 | 21168946 | 21169166 | -1.032955401 | KIZ-AS1 |
| 20 | 21395773 | 21395977 | -3.510971873 | NKX2-4 |
| 20 | 21500076 | 21500274 | -3.204446799 | RP5-984P4.6 |
| 20 | 23047585 | 23047786 | -1.716804335 | THBD |
| 20 | 23354243 | 23354444 | -2.340880048 | NXT1 |
| 20 | 23365072 | 23365276 | -1.673723708 | GZF1 |
| 20 | 25221941 | 25222088 | -1.136205434 | ENTPD6 |
| 20 | 25264822 | 25265010 | 6.473818469 | PYGB |
| 1 | 149900359 | 149900561 | -1.129750585 | BOLA1 |
| 20 | 25285817 | 25286001 | 5.099836172 | PYGB |
| 20 | 25289959 | 25290139 | -5.637771278 | PYGB |
| 20 | 25291542 | 25291653 | -6.018086806 | PYGB |
| 20 | 25300499 | 25300703 | 1.065843653 | ABHD12 |
| 20 | 25476373 | 25476526 | -2.234630689 | NINL |
| 20 | 25616257 | 25616452 | -1.113648404 | NANP |
| 20 | 25683343 | 25683497 | 2.243991706 | ZNF337 |
| 20 | 25690935 | 25691173 | 1.32065795 | ZNF337 |
| 20 | 25834002 | 25834165 | -1.477175715 | FAM182B |
| 20 | 25856460 | 25856659 | -6.928212764 | FAM182B |
| 20 | 25857042 | 25857210 | -8.620940383 | FAM182B |
| 20 | 25857675 | 25857846 | -2.692076737 | FAM182B |
| 20 | 25865190 | 25865339 | -5.664617871 | FAM182B |
| 1 | 149953041 | 149953240 | 3.007953078 | OTUD7B |
| 20 | 31548487 | 31548676 | -10.18781035 | HM13 |

| | | | | |
|---|---|---|---|---|
| 20 | 31564666 | 31564857 | -5.406890596 | HM13 |
| 20 | 31665977 | 31666168 | -6.414861368 | BCL2L1 |
| 20 | 31721775 | 31721977 | 1.387493071 | BCL2L1 |
| 20 | 31721775 | 31721977 | 1.387493071 | ABALON |
| 20 | 31801173 | 31801303 | -3.434424255 | TPX2 |
| 20 | 31945497 | 31945705 | -1.511038222 | PDRG1 |
| 1 | 149955801 | 149956005 | 3.291637075 | OTUD7B |
| 20 | 32195422 | 32195682 | -2.106630432 | PLAGL2 |
| 20 | 32197177 | 32197386 | -2.203082127 | PLAGL2 |
| 20 | 32234627 | 32234871 | -1.739711294 | POFUT1 |
| 20 | 32333977 | 32334177 | -1.029656166 | KIF3B |
| 20 | 32433479 | 32433700 | -1.371804635 | ASXL1 |
| 20 | 32436616 | 32436829 | -1.47718668 | ASXL1 |
| 1 | 149957533 | 149957804 | 2.434115277 | OTUD7B |
| 20 | 32437667 | 32437805 | -3.077369516 | ASXL1 |
| 20 | 32438654 | 32438820 | -2.121159176 | ASXL1 |
| 20 | 32447511 | 32447716 | -1.357050036 | AL034550.1 |
| 20 | 32447511 | 32447716 | -1.357050036 | NOL4L |
| 20 | 33408200 | 33408378 | -1.840735241 | SNTA1 |
| 20 | 33648144 | 33648307 | -2.080178858 | CBFA2T2 |
| 20 | 33649101 | 33649202 | -1.622264925 | CBFA2T2 |
| 20 | 33707659 | 33707864 | -1.279179571 | PXMP4 |
| 20 | 34550010 | 34550146 | -3.707342618 | MAP1LC3A |
| 20 | 34560656 | 34560844 | -1.614788543 | PIGU |
| 20 | 34741276 | 34741431 | -1.389535995 | NCOA6 |
| 20 | 35224697 | 35224869 | 2.501731928 | MMP24-AS1 |
| 20 | 35267670 | 35267853 | 1.5551756 | MMP24 |
| 20 | 35267670 | 35267853 | 1.5551756 | MMP24-AS1 |
| 20 | 35503004 | 35503184 | -1.176625612 | CEP250 |
| 20 | 35503829 | 35504130 | -1.197888893 | CEP250 |
| 20 | 35507592 | 35507789 | -1.896937577 | CEP250 |
| 1 | 150159262 | 150159488 | 5.122606531 | PLEKHO1 |
| 20 | 35669182 | 35669285 | -2.075952037 | NFS1 |
| 20 | 35669182 | 35669285 | -2.075952037 | RP1-309K20.6 |
| 20 | 35954150 | 35954335 | -6.603964723 | SCAND1 |
| 20 | 36229473 | 36229635 | 7.108183475 | EPB41L1 |
| 20 | 36240036 | 36240236 | -1.134442501 | AAR2 |
| 20 | 36526880 | 36527078 | 1.570234273 | DLGAP4 |
| 20 | 36526880 | 36527078 | 1.570234273 | DLGAP4-AS1 |
| 20 | 36612033 | 36612235 | -2.249285006 | TGIF2-C20orf24 |
| 20 | 36612033 | 36612235 | -2.249285006 | C20orf24 |
| 20 | 36785695 | 36785903 | -1.387661803 | SOGA1 |
| 20 | 36793429 | 36793628 | -1.134869224 | SOGA1 |
| 20 | 36815965 | 36816164 | -1.760958388 | SOGA1 |
| 20 | 37404047 | 37404251 | 3.475083699 | SRC |
| 20 | 38011724 | 38011910 | -1.632904462 | TTI1 |
| 20 | 38013352 | 38013576 | -1.349708239 | TTI1 |
| 20 | 38089848 | 38090054 | -1.161779457 | RPRD1B |
| 20 | 38951841 | 38952017 | -3.676095744 | FAM83D |
| 20 | 40687418 | 40687649 | 8.80155388 | MAFB |
| 20 | 41174619 | 41174803 | -1.236999499 | PLCG1 |
| 20 | 41182843 | 41183041 | -1.224485307 | PLCG1 |
| 20 | 41182843 | 41183041 | -1.224485307 | ZHX3 |
| 20 | 41185101 | 41185299 | -1.478079283 | PLCG1 |

| | | | | |
|---|---|---|---|---|
| 20 | 41185101 | 41185299 | -1.478079283 | ZHX3 |
| 20 | 41202159 | 41202356 | -1.711612276 | ZHX3 |
| 1 | 150472909 | 150473109 | -1.66657651 | RPRD2 |
| 20 | 41202443 | 41202544 | -1.325733931 | ZHX3 |
| 20 | 44224975 | 44225144 | 1.177553631 | OSER1-AS1 |
| 1 | 150507137 | 150507339 | -1.112084799 | TARS2 |
| 20 | 44489641 | 44489837 | -1.253877955 | TTPAL |
| 20 | 44958673 | 44958872 | 7.491051153 | TOMM34 |
| 20 | 44958673 | 44958872 | 7.491051153 | PABPC1L |
| 20 | 44976503 | 44976812 | -1.653825375 | STK4 |
| 20 | 45367115 | 45367301 | -1.237006169 | SYS1-DBNDD2 |
| 20 | 45367115 | 45367301 | -1.237006169 | SYS1 |
| 20 | 45367710 | 45367812 | -1.954810944 | SYS1-DBNDD2 |
| 20 | 45367710 | 45367812 | -1.954810944 | SYS1 |
| 20 | 45374813 | 45375011 | -1.748019579 | SYS1 |
| 20 | 45374813 | 45375011 | -1.748019579 | TP53TG5 |
| 20 | 45374813 | 45375011 | -1.748019579 | SYS1-DBNDD2 |
| 20 | 45797376 | 45797575 | -1.541274543 | DNTTIP1 |
| 1 | 150548624 | 150548758 | 6.592670027 | RP11-54A4.2 |
| 20 | 45876704 | 45876905 | -1.919699394 | ZSWIM3 |
| 20 | 45877942 | 45878247 | -1.819830891 | ZSWIM3 |
| 20 | 45883851 | 45884053 | -1.189745749 | ZSWIM1 |
| 20 | 45947684 | 45947917 | -1.138744164 | PCIF1 |
| 20 | 46062346 | 46062547 | -1.317746979 | NCOA5 |
| 20 | 46367267 | 46367470 | 1.057870439 | ELMO2 |
| 20 | 46367267 | 46367470 | 1.057870439 | RP11-394O2.3 |
| 20 | 46541328 | 46541527 | 10.88793854 | OCSTAMP |
| 20 | 47635909 | 47636109 | 1.671336508 | NCOA3 |
| 20 | 47636467 | 47636658 | 2.537501543 | NCOA3 |
| 20 | 47647025 | 47647227 | 3.014759624 | NCOA3 |
| 20 | 48625322 | 48625527 | -2.561143061 | PREX1 |
| 20 | 48639337 | 48639528 | -7.62884124 | PREX1 |
| 20 | 49144368 | 49144647 | -6.805742972 | STAU1 |
| 20 | 49270103 | 49270367 | 1.039541223 | ZNFX1 |
| 20 | 49271481 | 49271684 | 1.066520708 | ZNFX1 |
| 1 | 6415368 | 6415540 | 7.502790346 | HES2 |
| 20 | 49904969 | 49905105 | -1.927735808 | SPATA2 |
| 20 | 49906531 | 49906697 | -1.228005998 | SPATA2 |
| 20 | 49988332 | 49988485 | 2.219051049 | SNAI1 |
| 20 | 50192219 | 50192414 | 1.447727267 | CEBPB |
| 20 | 50894210 | 50894411 | -1.131417619 | ADNP |
| 20 | 50959758 | 50959957 | -1.1925544 | MOCS3 |
| 20 | 51598656 | 51598860 | 4.110993679 | ATP9A |
| 20 | 51600020 | 51600193 | 5.400005691 | ATP9A |
| 20 | 51601035 | 51601233 | 4.17491203 | ATP9A |
| 1 | 150648353 | 150648554 | -1.218245489 | GOLPH3L |
| 20 | 52152765 | 52152986 | -1.410647267 | ZFP64 |
| 20 | 53569048 | 53569336 | -1.714704453 | RP4-724E16.2 |
| 20 | 53569048 | 53569336 | -1.714704453 | ZNF217 |
| 20 | 53571783 | 53571979 | -1.340476332 | ZNF217 |
| 20 | 53571783 | 53571979 | -1.340476332 | RP4-724E16.2 |
| 20 | 56369971 | 56370170 | -2.12190813 | AURKA |
| 20 | 56452835 | 56453035 | 5.482673953 | CASS4 |
| 20 | 56458697 | 56458894 | 6.457345596 | CASS4 |

| | | | | |
|---|---|---|---|---|
| 20 | 57371342 | 57371528 | -6.68438963 | RAE1 |
| 20 | 57652104 | 57652458 | 3.511786788 | PMEPA1 |
| 20 | 58991089 | 58991251 | 2.988639962 | NELFCD |
| 20 | 59774330 | 59774489 | -3.831988994 | PHACTR3 |
| 20 | 59805916 | 59806113 | -3.983796057 | PHACTR3 |
| 20 | 59847190 | 59847453 | -4.348648923 | PHACTR3 |
| 20 | 59904356 | 59904546 | -7.977143602 | SYCP2 |
| 20 | 59938270 | 59938437 | -2.391565988 | FAM217B |
| 20 | 59938270 | 59938437 | -2.391565988 | PPP1R3D |
| 20 | 59939152 | 59939348 | -3.471257253 | FAM217B |
| 20 | 59939152 | 59939348 | -3.471257253 | PPP1R3D |
| 20 | 59944424 | 59944524 | -1.70646429 | FAM217B |
| 20 | 59944864 | 59945061 | -1.577634063 | FAM217B |
| 20 | 60057387 | 60057554 | -1.71612546 | C20orf197 |
| 20 | 60058202 | 60058306 | -5.414270825 | C20orf197 |
| 20 | 60240145 | 60240314 | 5.614409345 | MIR646HG |
| 20 | 60279158 | 60279365 | 7.13339327 | MIR646HG |
| 20 | 60322491 | 60322610 | 1.799544365 | MIR646HG |
| 1 | 151002283 | 151002493 | -1.087797393 | FAM63A |
| 20 | 62200877 | 62201079 | -2.060069505 | MTG2 |
| 20 | 62202374 | 62202574 | -1.723232258 | MTG2 |
| 20 | 62389257 | 62389456 | -1.369230718 | CABLES2 |
| 20 | 62397333 | 62397478 | -2.332919616 | CABLES2 |
| 20 | 62398430 | 62398574 | -1.051718458 | CABLES2 |
| 20 | 62399607 | 62399754 | -1.247176394 | CABLES2 |
| 20 | 62401882 | 62402003 | -3.545961245 | CABLES2 |
| 1 | 151033937 | 151034161 | -1.131118791 | PRUNE1 |
| 20 | 62402599 | 62402783 | -2.785684066 | RP5-908M14.10 |
| 20 | 62402599 | 62402783 | -2.785684066 | CABLES2 |
| 20 | 62602167 | 62602371 | 2.796883256 | WI2-87327B8.2 |
| 20 | 62664923 | 62665121 | -10.35662571 | SLCO4A1-AS1 |
| 20 | 62664923 | 62665121 | -10.35662571 | SLCO4A1 |
| 1 | 151053034 | 151053234 | 1.007309255 | CDC42SE1 |
| 20 | 62669134 | 62669331 | -8.870499739 | SLCO4A1 |
| 20 | 62710207 | 62710475 | -2.152981233 | NTSR1 |
| 20 | 62714577 | 62714766 | -3.411584912 | NTSR1 |
| 20 | 62722573 | 62722712 | -7.335328125 | NTSR1 |
| 20 | 62724925 | 62725107 | -1.863116024 | NTSR1 |
| 20 | 62746593 | 62746789 | -7.493632006 | NTSR1 |
| 1 | 6418626 | 6418837 | 7.229234347 | HES2 |
| 1 | 151059661 | 151059784 | 2.2123331 | MLLT11 |
| 1 | 151059661 | 151059784 | 2.2123331 | CDC42SE1 |
| 20 | 62799886 | 62800088 | -1.087665227 | MRGBP |
| 20 | 62827816 | 62827949 | -8.212304594 | COL9A3 |
| 20 | 62841893 | 62842026 | -1.669349338 | TCFL5 |
| 20 | 62844309 | 62844508 | -1.406847984 | TCFL5 |
| 20 | 62852674 | 62852908 | -2.499050827 | TCFL5 |
| 20 | 62881568 | 62881766 | -1.226323922 | DIDO1 |
| 20 | 62937620 | 62937821 | 6.464629204 | DIDO1 |
| 20 | 62967362 | 62967570 | -1.205082422 | SLC17A9 |
| 1 | 151159433 | 151159635 | -1.121853179 | TNFAIP8L2 |
| 1 | 151159433 | 151159635 | -1.121853179 | SCNM1 |
| 20 | 63564817 | 63564985 | 3.143122324 | HELZ2 |
| 20 | 63565713 | 63565961 | 3.65601919 | HELZ2 |

| | | | | |
|---|---|---|---|---|
| 20 | 63669386 | 63669583 | -7.249268801 | RTEL1 |
| 20 | 63669386 | 63669583 | -7.249268801 | RTEL1-TNFRSF6B |
| 20 | 63682552 | 63682748 | -3.392822828 | RTEL1 |
| 20 | 63682552 | 63682748 | -3.392822828 | RTEL1-TNFRSF6B |
| 20 | 63683740 | 63683920 | 4.443939624 | RTEL1 |
| 20 | 63683740 | 63683920 | 4.443939624 | RTEL1-TNFRSF6B |
| 20 | 63686356 | 63686578 | 7.328380012 | RTEL1-TNFRSF6B |
| 20 | 63686356 | 63686578 | 7.328380012 | RTEL1 |
| 20 | 63692617 | 63692802 | 2.630263775 | RTEL1-TNFRSF6B |
| 20 | 63692617 | 63692802 | 2.630263775 | RTEL1 |
| 20 | 63708746 | 63708973 | -1.283986649 | ZGPAT |
| 20 | 63708746 | 63708973 | -1.283986649 | RP4-583P15.15 |
| 1 | 151165524 | 151165662 | -1.812837769 | SCNM1 |
| 1 | 151165524 | 151165662 | -1.812837769 | LYSMD1 |
| 20 | 63735762 | 63735952 | -1.195260211 | RP4-583P15.15 |
| 20 | 63735762 | 63735952 | -1.195260211 | ZGPAT |
| 20 | 63743657 | 63743852 | -10.64292121 | ZBTB46 |
| 20 | 63767566 | 63767769 | 1.370371967 | ZBTB46 |
| 20 | 63769569 | 63769736 | 1.624679734 | ZBTB46 |
| 20 | 63797874 | 63798148 | 1.679999663 | ZBTB46 |
| 1 | 151281939 | 151282135 | -2.575096818 | ZNF687 |
| 20 | 63919980 | 63920182 | -1.172055864 | DNAJC5 |
| 20 | 63931473 | 63931642 | -2.706892012 | DNAJC5 |
| 20 | 63954293 | 63954618 | 1.943867477 | UCKL1 |
| 20 | 63954293 | 63954618 | 1.943867477 | UCKL1-AS1 |
| 20 | 63955171 | 63955369 | 6.559146117 | UCKL1 |
| 20 | 63955171 | 63955369 | 6.559146117 | UCKL1-AS1 |
| 20 | 63959983 | 63960176 | -4.163239643 | ZNF512B |
| 20 | 64063476 | 64063666 | -5.597993649 | TCEA2 |
| 20 | 64259926 | 64260107 | 2.153298379 | PCMTD2 |
| 21 | 5036159 | 5036361 | 1.61628499 | CH507-9B2.1 |
| 21 | 5037828 | 5038022 | 1.216479704 | CH507-9B2.1 |
| 21 | 5103991 | 5104198 | 2.611162312 | CH507-9B2.3 |
| 1 | 151292513 | 151292844 | -1.123455207 | PI4KB |
| 21 | 5104610 | 5104719 | 4.0629571 | CH507-9B2.3 |
| 21 | 5106045 | 5106282 | 2.920860791 | CH507-9B2.3 |
| 21 | 5120794 | 5120993 | 6.723799929 | CH507-9B2.3 |
| 21 | 5120794 | 5120993 | 6.723799929 | CH507-9B2.4 |
| 21 | 5122387 | 5122574 | 4.110360442 | CH507-9B2.3 |
| 21 | 5122387 | 5122574 | 4.110360442 | CH507-9B2.4 |
| 21 | 5125654 | 5125814 | 7.543731421 | CH507-9B2.4 |
| 21 | 5125654 | 5125814 | 7.543731421 | CH507-9B2.3 |
| 21 | 5130927 | 5131129 | -1.786342085 | CH507-9B2.5 |
| 21 | 5130927 | 5131129 | -1.786342085 | CH507-9B2.4 |
| 21 | 5141734 | 5141914 | 1.599247077 | CH507-9B2.5 |
| 21 | 7626305 | 7626470 | -3.90894865 | CH507-338C24.2 |
| 21 | 9069550 | 9069748 | -1.155216228 | TEKT4P2 |
| 21 | 9790533 | 9790715 | -1.435315797 | MGC39584 |
| 21 | 9803863 | 9804052 | -8.312797351 | MGC39584 |
| 21 | 9809677 | 9809865 | -3.061691208 | MGC39584 |
| 21 | 14076128 | 14076320 | -6.845762721 | AP001347.6 |
| 1 | 151405940 | 151406140 | -1.424346826 | POGZ |
| 21 | 14319936 | 14320131 | -3.104410891 | ABCC13 |
| 21 | 14329374 | 14329543 | -4.624927085 | ABCC13 |

| | | | | |
|---|---|---|---|---|
| 21 | 14332050 | 14332172 | -4.599780407 | ABCC13 |
| 1 | 6441253 | 6441436 | -6.655874158 | ESPN |
| 21 | 25735132 | 25735328 | -2.52894774 | GABPA |
| 21 | 25735132 | 25735328 | -2.52894774 | ATP5J |
| 21 | 26837627 | 26837824 | -4.601028573 | ADAMTS1 |
| 21 | 26844995 | 26845246 | -4.011499805 | ADAMTS1 |
| 21 | 26924417 | 26924543 | -3.31467831 | ADAMTS5 |
| 21 | 26924417 | 26924543 | -3.31467831 | AP001601.2 |
| 21 | 26965390 | 26965534 | -2.484103473 | ADAMTS5 |
| 21 | 28874792 | 28875004 | -2.223410546 | N6AMT1 |
| 21 | 28875200 | 28875358 | -1.640586753 | N6AMT1 |
| 21 | 29047010 | 29047213 | -1.229484761 | USP16 |
| 21 | 29342428 | 29342638 | 1.446157206 | BACH1 |
| 21 | 32314602 | 32314802 | -2.481134128 | MRAP |
| 21 | 32314602 | 32314802 | -2.481134128 | URB1 |
| 21 | 32316630 | 32316867 | -2.165887826 | URB1 |
| 21 | 32347437 | 32347641 | -2.233032924 | URB1 |
| 1 | 153774482 | 153774807 | -1.602440737 | SLC27A3 |
| 1 | 153774482 | 153774807 | -1.602440737 | INTS3 |
| 21 | 33027792 | 33028102 | 2.787447385 | OLIG2 |
| 21 | 33027792 | 33028102 | 2.787447385 | AP000282.2 |
| 21 | 33285796 | 33285993 | -3.349459549 | IL10RB |
| 21 | 33309108 | 33309297 | 2.577721461 | IL10RB |
| 21 | 33436959 | 33437132 | 1.194928802 | TMEM50B |
| 21 | 33436959 | 33437132 | 1.194928802 | IFNGR2 |
| 21 | 33551389 | 33551587 | -1.362847205 | SON |
| 21 | 33976984 | 33977184 | 1.272721109 | LINC00649 |
| 21 | 34096254 | 34096406 | 1.063066937 | SLC5A3 |
| 21 | 34096254 | 34096406 | 1.063066937 | AP000320.7 |
| 21 | 34096254 | 34096406 | 1.063066937 | MRPS6 |
| 21 | 34097117 | 34097317 | 1.276535119 | MRPS6 |
| 21 | 34097117 | 34097317 | 1.276535119 | SLC5A3 |
| 21 | 34097117 | 34097317 | 1.276535119 | AP000320.7 |
| 21 | 34449265 | 34449398 | 3.947976946 | KCNE1 |
| 21 | 34496826 | 34497026 | 2.923766545 | KCNE1 |
| 21 | 34498411 | 34498536 | 5.382021109 | KCNE1 |
| 21 | 34517729 | 34517946 | 1.018076545 | RCAN1 |
| 21 | 34520638 | 34520774 | 6.239005586 | RCAN1 |
| 21 | 34846925 | 34847116 | 8.292305119 | RUNX1 |
| 21 | 34887333 | 34887533 | -1.37525843 | RUNX1 |
| 21 | 34968020 | 34968215 | -1.839095106 | RUNX1 |
| 1 | 153852057 | 153852259 | 1.679303238 | GATAD2B |
| 1 | 153852057 | 153852259 | 1.679303238 | RP11-216N14.7 |
| 21 | 37142075 | 37142190 | -6.401863763 | TTC3 |
| 21 | 37143190 | 37143387 | -1.475926937 | TTC3 |
| 21 | 37143865 | 37144065 | -6.479729323 | TTC3 |
| 21 | 37222063 | 37222263 | -1.09237056 | AP001432.14 |
| 21 | 37241223 | 37241519 | -7.818184862 | DSCR3 |
| 21 | 37261268 | 37261456 | -1.062112909 | DSCR3 |
| 21 | 37267799 | 37267997 | -5.196158711 | DSCR3 |
| 21 | 37267799 | 37267997 | -5.196158711 | AP001412.1 |
| 1 | 153852936 | 153853136 | 4.052319979 | RP11-216N14.7 |
| 1 | 153852936 | 153853136 | 4.052319979 | GATAD2B |
| 21 | 38822824 | 38823026 | -1.400121541 | ETS2 |

| | | | | |
|---|---|---|---|---|
| 21 | 38989170 | 38989364 | -5.833469504 | AF064858.8 |
| 21 | 41390109 | 41390283 | 5.596146407 | MX2 |
| 21 | 41801167 | 41801367 | -1.206372844 | PRDM15 |
| 21 | 41952173 | 41952303 | -2.424859939 | C2CD2 |
| 21 | 41993624 | 41993824 | 1.025543334 | ZBTB21 |
| 21 | 42580361 | 42580565 | -1.056800169 | SLC37A1 |
| 21 | 42849327 | 42849530 | -3.144482288 | WDR4 |
| 21 | 42903977 | 42904165 | -1.229653305 | NDUFV3 |
| 1 | 6524459 | 6524664 | -1.887920241 | NOL9 |
| 1 | 154271037 | 154271236 | -1.973543618 | UBAP2L |
| 21 | 43416672 | 43416822 | 1.261780978 | SIK1 |
| 21 | 43694784 | 43694984 | -1.27387179 | RRP1B |
| 21 | 43724314 | 43724506 | 8.206813965 | PDXK |
| 21 | 43744787 | 43745016 | 1.468201512 | PDXK |
| 1 | 154349122 | 154349356 | 2.236411893 | ATP8B2 |
| 21 | 43803789 | 43803971 | -1.05885588 | RRP1 |
| 21 | 43809115 | 43809235 | 2.063217589 | AATBC |
| 21 | 43809767 | 43809955 | -6.61162378 | AATBC |
| 21 | 43810509 | 43810650 | -5.803665157 | AATBC |
| 21 | 43954751 | 43954935 | -7.180594945 | AGPAT3 |
| 21 | 43966755 | 43967022 | -7.799011707 | AGPAT3 |
| 21 | 44137708 | 44137883 | 5.856693257 | C21orf33 |
| 21 | 44226686 | 44226855 | 2.765257358 | ICOSLG |
| 21 | 44227251 | 44227453 | 2.234320146 | ICOSLG |
| 21 | 44330310 | 44330557 | -1.193480666 | C21orf2 |
| 21 | 44333500 | 44333700 | 1.673256867 | C21orf2 |
| 21 | 44333500 | 44333700 | 1.673256867 | AP001062.7 |
| 21 | 44351164 | 44351341 | -10.49825674 | TRPM2 |
| 21 | 44353513 | 44353713 | -9.446595949 | TRPM2 |
| 21 | 44357248 | 44357530 | -3.512089799 | TRPM2 |
| 21 | 44358733 | 44358866 | -2.839991121 | TRPM2 |
| 21 | 44441749 | 44441955 | -1.520989391 | TRPM2 |
| 1 | 154484128 | 154484295 | 2.755202769 | SHE |
| 21 | 44769410 | 44769620 | -1.951766617 | UBE2G2 |
| 21 | 44897832 | 44898027 | -7.459408331 | ITGB2 |
| 21 | 44931481 | 44931674 | 2.288260065 | ITGB2 |
| 21 | 44931985 | 44932182 | 5.544394119 | ITGB2 |
| 21 | 44934375 | 44934550 | 2.385373659 | LINC01547 |
| 21 | 44976809 | 44976964 | -3.259626588 | FAM207A |
| 21 | 45073072 | 45073220 | -1.171977201 | SSR4P1 |
| 21 | 45145542 | 45145753 | 1.10183826 | ADARB1 |
| 21 | 45162625 | 45162844 | 4.329105741 | ADARB1 |
| 21 | 45176188 | 45176346 | 1.373481526 | ADARB1 |
| 21 | 45192603 | 45192802 | 6.287769623 | ADARB1 |
| 21 | 45222098 | 45222293 | 1.813868262 | ADARB1 |
| 21 | 45294578 | 45294775 | -1.180631246 | LINC00205 |
| 21 | 45515838 | 45516037 | -3.70051732 | SLC19A1 |
| 21 | 45596998 | 45597118 | -2.292515797 | AL133493.2 |
| 21 | 45600792 | 45600962 | -1.623502488 | AL133493.2 |
| 1 | 154584813 | 154585016 | 1.030301947 | ADAR |
| 21 | 45793389 | 45793583 | -4.503653823 | PCBP3 |
| 21 | 45852791 | 45852991 | -3.621252706 | PCBP3 |
| 21 | 45860579 | 45860750 | -7.015678298 | PCBP3 |
| 21 | 45918923 | 45919120 | -6.771757608 | PCBP3-OT1 |

| | | | | |
|---|---|---|---|---|
| 21 | 45918923 | 45919120 | -6.771757608 | PCBP3 |
| 21 | 46004663 | 46004867 | 1.907836327 | COL6A1 |
| 21 | 46129313 | 46129463 | 3.26569073 | COL6A2 |
| 21 | 46244846 | 46245006 | -10.76946933 | MCM3AP-AS1 |
| 21 | 46244846 | 46245006 | -10.76946933 | MCM3AP |
| 21 | 46249882 | 46250071 | -10.27997139 | MCM3AP-AS1 |
| 21 | 46249882 | 46250071 | -10.27997139 | MCM3AP |
| 21 | 46250732 | 46250884 | -7.750358709 | MCM3AP |
| 21 | 46250732 | 46250884 | -7.750358709 | MCM3AP-AS1 |
| 21 | 46291382 | 46291589 | -1.873653162 | YBEY |
| 21 | 46321353 | 46321550 | -4.09735543 | C21orf58 |
| 21 | 46322512 | 46322720 | -4.017482975 | C21orf58 |
| 21 | 46323065 | 46323268 | -3.361729135 | C21orf58 |
| 21 | 46363446 | 46363644 | -1.356692655 | PCNT |
| 21 | 46366840 | 46367052 | -1.831569697 | PCNT |
| 21 | 46416146 | 46416453 | -1.727918871 | PCNT |
| 21 | 46431675 | 46431879 | -1.768659522 | PCNT |
| 22 | 15821541 | 15821737 | -1.450243779 | DUXAP8 |
| 22 | 16665306 | 16665518 | 1.019975114 | ANKRD62P1-PARP4P3 |
| 22 | 16665306 | 16665518 | 1.019975114 | TPTEP1 |
| 22 | 17057286 | 17057493 | -1.742713414 | CECR7 |
| 1 | 154958548 | 154958752 | -1.442171773 | PYGO2 |
| 22 | 17777430 | 17777598 | -7.496668237 | LINC00528 |
| 22 | 17778514 | 17778680 | -6.775193033 | LINC00528 |
| 1 | 154959560 | 154959708 | -1.202680024 | PYGO2 |
| 22 | 17807760 | 17807938 | -1.952292399 | MICAL3 |
| 22 | 17807760 | 17807938 | -1.952292399 | XXbac-B461K10.4 |
| 22 | 17818360 | 17818526 | -1.711435887 | MICAL3 |
| 22 | 17818842 | 17819021 | -2.102138253 | MICAL3 |
| 22 | 18145535 | 18145789 | 6.603049307 | TUBA8 |
| 1 | 154988577 | 154988781 | -1.331046548 | FLAD1 |
| 22 | 18607188 | 18607366 | -1.014805069 | RIMBP3 |
| 22 | 18608655 | 18608853 | -1.87438906 | RIMBP3 |
| 22 | 18989886 | 18990113 | 4.478369573 | DGCR5 |
| 22 | 19030555 | 19030655 | 1.222655255 | DGCR5 |
| 22 | 19036850 | 19037096 | -2.256467584 | DGCR2 |
| 22 | 19038491 | 19038746 | -1.060137239 | DGCR2 |
| 22 | 19131299 | 19131468 | -1.453416205 | DGCR14 |
| 22 | 19131299 | 19131468 | -1.453416205 | TSSK2 |
| 22 | 19134036 | 19134236 | -1.05788341 | DGCR14 |
| 22 | 19179651 | 19179775 | -4.185969197 | CLTCL1 |
| 22 | 19189488 | 19189638 | -6.002073859 | CLTCL1 |
| 22 | 19432038 | 19432192 | -7.266969014 | MRPL40 |
| 22 | 19432038 | 19432192 | -7.266969014 | HIRA |
| 22 | 19432038 | 19432192 | -7.266969014 | C22orf39 |
| 22 | 19444046 | 19444244 | -1.875057892 | C22orf39 |
| 22 | 19444046 | 19444244 | -1.875057892 | HIRA |
| 22 | 19722515 | 19722715 | -2.036731652 | Sep-05 |
| 22 | 19724485 | 19724688 | -1.785826909 | Sep-05 |
| 22 | 19724485 | 19724688 | -1.785826909 | GP1BB |
| 22 | 19763916 | 19764103 | -3.419661891 | TBX1 |
| 22 | 19782968 | 19783165 | -2.765192167 | TBX1 |
| 22 | 19784801 | 19784999 | -1.336922456 | GNB1L |
| 22 | 19786335 | 19786473 | -2.30498746 | GNB1L |

| | | | | |
|---|---|---|---|---|
| 22 | 19791927 | 19792110 | -1.251206967 | GNB1L |
| 22 | 19796704 | 19796898 | -1.879592062 | GNB1L |
| 22 | 19801137 | 19801381 | -1.777490571 | GNB1L |
| 22 | 19801748 | 19801946 | -1.296626237 | GNB1L |
| 22 | 19802417 | 19802615 | -1.451262601 | GNB1L |
| 22 | 19803061 | 19803299 | -1.208373235 | GNB1L |
| 22 | 19804163 | 19804356 | -1.673558778 | GNB1L |
| 22 | 19851987 | 19852301 | -1.483685776 | GNB1L |
| 22 | 19851987 | 19852301 | -1.483685776 | C22orf29 |
| 22 | 19886667 | 19886843 | -1.14861699 | TXNRD2 |
| 22 | 20062923 | 20063130 | -1.835236558 | TANGO2 |
| 22 | 20062923 | 20063130 | -1.835236558 | AC006547.13 |
| 22 | 20086249 | 20086428 | -2.491457064 | DGCR8 |
| 22 | 20110252 | 20110450 | -2.590197334 | DGCR8 |
| 22 | 20405499 | 20405721 | -1.380958711 | ZNF74 |
| 22 | 20465063 | 20465276 | -1.757712299 | KLHL22 |
| 22 | 20953727 | 20953921 | -3.430103557 | CRKL |
| 22 | 20999873 | 21000073 | -1.817035639 | THAP7 |
| 22 | 21388428 | 21388632 | -1.836642212 | RIMBP3B |
| 22 | 21450121 | 21450301 | -2.299301544 | HIC2 |
| 22 | 21545847 | 21546031 | -1.413266144 | RIMBP3C |
| 22 | 21622651 | 21622777 | -1.721997487 | UBE2L3 |
| 22 | 21628210 | 21628409 | -3.209621749 | YDJC |
| 1 | 155338264 | 155338414 | 2.289948275 | ASH1L |
| 22 | 21697160 | 21697364 | -1.285322269 | PPIL2 |
| 22 | 21942931 | 21943129 | -8.489427834 | PPM1F |
| 22 | 21942931 | 21943129 | -8.489427834 | LL22NC03-86G7.1 |
| 22 | 22321583 | 22321815 | -1.063775025 | BMS1P20 |
| 22 | 22488443 | 22488648 | -1.091646164 | ZNF280B |
| 22 | 23263685 | 23263898 | -1.10369862 | BCR |
| 22 | 23263685 | 23263898 | -1.10369862 | FBXW4P1 |
| 22 | 23895845 | 23896033 | -1.800448958 | MIF-AS1 |
| 22 | 24178160 | 24178323 | -1.383335425 | CABIN1 |
| 22 | 24321761 | 24321964 | -1.12994925 | SPECC1L-ADORA2A |
| 22 | 24321761 | 24321964 | -1.12994925 | SPECC1L |
| 22 | 24373565 | 24373760 | -1.257347621 | SPECC1L-ADORA2A |
| 22 | 24373565 | 24373760 | -1.257347621 | SPECC1L |
| 22 | 24441683 | 24441880 | 2.650955311 | SPECC1L-ADORA2A |
| 22 | 24441683 | 24441880 | 2.650955311 | ADORA2A |
| 22 | 24441683 | 24441880 | 2.650955311 | ADORA2A-AS1 |
| 22 | 25466221 | 25466400 | -6.771455473 | CRYBB2P1 |
| 22 | 26433684 | 26433851 | -1.15737822 | ASPHD2 |
| 22 | 26434191 | 26434369 | -1.066741503 | ASPHD2 |
| 22 | 26442939 | 26443114 | 2.838540351 | ASPHD2 |
| 22 | 26450784 | 26450951 | -2.786657496 | HPS4 |
| 22 | 26453128 | 26453327 | -1.965837436 | HPS4 |
| 22 | 26464163 | 26464364 | -1.8320778 | HPS4 |
| 22 | 26464419 | 26464547 | -2.063955654 | HPS4 |
| 22 | 26481901 | 26482101 | -1.582696158 | HPS4 |
| 22 | 26491537 | 26491798 | -1.3183039 | SRRD |
| 22 | 26491537 | 26491798 | -1.3183039 | TFIP11 |
| 22 | 27921998 | 27922170 | -1.011293982 | TTC28-AS1 |
| 22 | 27921998 | 27922170 | -1.011293982 | TTC28-AS1_2 |
| 22 | 27924319 | 27924521 | -1.283128398 | TTC28-AS1 |

| | | | | |
|---|---|---|---|---|
| 22 | 27999416 | 27999611 | -1.155500603 | TTC28-AS1 |
| 22 | 27999416 | 27999611 | -1.155500603 | TTC28 |
| 22 | 28001066 | 28001254 | -1.074134285 | TTC28-AS1 |
| 22 | 28001066 | 28001254 | -1.074134285 | TTC28 |
| 22 | 28387193 | 28387330 | 2.438872125 | TTC28 |
| 22 | 28506130 | 28506325 | -1.376783781 | TTC28 |
| 22 | 28786129 | 28786335 | -1.04246708 | CCDC117 |
| 22 | 29049305 | 29049503 | -2.436669018 | ZNRF3 |
| 22 | 29050554 | 29050769 | -1.573166409 | ZNRF3 |
| 22 | 29053475 | 29053681 | -1.989735813 | ZNRF3 |
| 22 | 29313042 | 29313242 | 3.328741308 | RASL10A |
| 22 | 29469816 | 29469947 | -4.241563146 | RFPL1S |
| 22 | 29490860 | 29491054 | -3.037668913 | NEFH |
| 22 | 30241071 | 30241261 | 4.897207933 | LIF |
| 22 | 30263910 | 30264115 | 2.954963545 | OSM |
| 22 | 30375771 | 30375962 | 1.033347422 | CCDC157 |
| 22 | 30375771 | 30375962 | 1.033347422 | RNF215 |
| 22 | 30428726 | 30428926 | -2.577503988 | MTFP1 |
| 22 | 30428726 | 30428926 | -2.577503988 | RP4-539M6.19 |
| 22 | 30903743 | 30903936 | 1.900960518 | OSBP2 |
| 1 | 155563227 | 155563449 | -1.785385375 | ASH1L-AS1 |
| 22 | 31084474 | 31084659 | -2.742062867 | SMTN |
| 22 | 31276907 | 31277139 | -2.307930405 | LIMK2 |
| 22 | 31278726 | 31278870 | -2.186662975 | LIMK2 |
| 22 | 31291861 | 31292038 | -1.568751762 | PIK3IP1 |
| 1 | 155614380 | 155614583 | -2.157772151 | MSTO1 |
| 22 | 31344940 | 31345112 | -2.199739495 | PATZ1 |
| 22 | 31347128 | 31347276 | -3.325525846 | LINC01521 |
| 22 | 31347946 | 31348070 | -4.047730675 | LINC01521 |
| 22 | 31618936 | 31619105 | -1.370841043 | PISD |
| 22 | 31619568 | 31619765 | -1.349037334 | PISD |
| 22 | 31712315 | 31712517 | -1.202707748 | PRR14L |

**Additional file 6**

**Table S2:** Lists of gene ontology terms representing genes with enriched m6A peaks in Mϕ, M1 and M2 populations used to generate enrichment

| Mϕ | | | | | | | |
|---|---|---|---|---|---|---|---|
| **Term** | **Overlap** | **P.value** | **Adjusted.P.value** | **Odds Ratio** | **Combined Score** | **Genes** | **Count** |
| glycosaminoglycan bio | 26/97 | 8.23E-07 | 0.003643586 | 3.51268655 | 49.21525397 | HS3ST3A1;SDC3;SLC | 26 |
| glycosaminoglycan met | 17/54 | 6.20E-06 | 0.01340643 | 4.394554512 | 52.6943186 | HS3ST3B1;HS3ST3A | 17 |
| artery morphogenesis ( | Dec-30 | 9.08E-06 | 0.01340643 | 6.366261666 | 73.90808113 | BMP4;NRP1;TGFB2; | 12 |
| positive regulation of gl | Sep-18 | 1.39E-05 | 0.014603616 | 9.539029536 | 106.7087154 | EGR2;BMP2;NOTCH | 9 |
| positive regulation of as | 6-Aug | 1.75E-05 | 0.014603616 | 28.58293839 | 313.0448771 | BMP2;NOTCH1;SERI | 6 |
| chondrocyte differentiat | Dec-32 | 1.98E-05 | 0.014603616 | 5.729001585 | 62.04884404 | BMP4;BMP2;BMPR2; | 12 |
| regulation of cell migrat | 65/408 | 2.49E-05 | 0.015763332 | 1.828305235 | 19.38019592 | FLT1;SERPINE2;CSF | 65 |
| aminoglycan biosynthet | 15/49 | 3.14E-05 | 0.017380046 | 4.215919701 | 43.71453436 | HS3ST3B1;HS3ST3A | 15 |
| establishment of endoth | Nov-29 | 3.88E-05 | 0.019104896 | 5.832658688 | 59.23948649 | RAB1A;AFDN;RAP1 | 11 |
| negative regulation of T | Dec-35 | 5.58E-05 | 0.024699353 | 4.980913664 | 48.78467357 | CD274;IL4I1;PRNP;C | 12 |
| heart development (GO | 32/164 | 6.76E-05 | 0.026300665 | 2.324968856 | 22.32448261 | NOTCH1;BMPR2;CH | 32 |
| regulation of pathway-r | 16/58 | 7.13E-05 | 0.026300665 | 3.640727016 | 34.76597042 | ACVR1;TGFB2;BMP | 16 |
| positive regulation of ep | 13/42 | 9.24E-05 | 0.031484273 | 4.280418459 | 39.76186376 | TGFB2;NOTCH1;JAG | 13 |
| positive regulation of pr | 58/371 | 0.000112981 | 0.0341924 | 1.784010337 | 16.21360641 | BCAR3;NRP1;BMPR | 58 |
| positive regulation of ce | 45/269 | 0.000135247 | 0.0341924 | 1.930191532 | 17.19493074 | NRP1;DOCK5;SEMA | 45 |
| positive regulation of pe | 27/134 | 0.000138853 | 0.0341924 | 2.416947837 | 21.46755724 | NRP1;SRC;PDGFB;N | 27 |
| positive regulation of ca | 5-Jul | 0.000138962 | 0.0341924 | 23.80657895 | 211.4335891 | ACVR1;TGFB2;JAG1 | 5 |
| regulation of lens fiber | 5-Jul | 0.000138962 | 0.0341924 | 23.80657895 | 211.4335891 | SPRED3;SPRED2;SPI | 5 |
| ventricular septum mor | Oct-28 | 0.000157714 | 0.03676403 | 5.29961888 | 46.39670838 | ACVR1;BMP4;TGFB | 10 |
| regulation of GTPase ac | 34/189 | 0.000214085 | 0.044532165 | 2.103274081 | 17.77085223 | USP6NL;BCAR3;PLX | 34 |
| cardiac muscle cell diffe | Sep-24 | 0.000219589 | 0.044532165 | 5.721518987 | 48.19667177 | ACVR1;KDM6B;BMI | 9 |
| regulation of epithelial t | 18/76 | 0.000221203 | 0.044532165 | 2.966449208 | 24.96691268 | PHLDB1;TGFB2;TGF | 18 |
| positive regulation of ap | 49/310 | 0.000284778 | 0.053923736 | 1.803957755 | 14.72714849 | TRIO;BTG1;CCAR2;I | 49 |
| cellular response to typ | 16/65 | 0.000304379 | 0.053923736 | 3.119413144 | 25.25863005 | IFITM3;EGR1;SP100; | 16 |
| type I interferon signali | 16/65 | 0.000304379 | 0.053923736 | 3.119413144 | 25.25863005 | IFITM3;EGR1;SP100; | 16 |
| positive regulation of pa | 13/47 | 0.000324221 | 0.055229787 | 3.64993471 | 29.32388721 | ACVR1;TGFB2;BMP | 13 |
| coronary vasculature m | 5-Aug | 0.000341367 | 0.055996856 | 15.87017544 | 126.6845001 | NRP1;SPRED1;NOTC | 5 |
| aminoglycan catabolic p | Oct-31 | 0.000408602 | 0.056840914 | 4.541776605 | 35.43843439 | GPC1;FUCA1;SDC3; | 10 |
| positive regulation of ce | 23/114 | 0.000411746 | 0.056840914 | 2.417886046 | 18.84767115 | RB1;PPP1R15A;TGF | 23 |
| regulation of astrocyte | 6-Dec | 0.000412615 | 0.056840914 | 9.525539758 | 74.23248836 | BMP2;NOTCH1;SERI | 6 |
| cellular response to gro | 29/158 | 0.000427688 | 0.056840914 | 2.152914828 | 16.70040991 | NRP1;NOTCH1;BMP | 29 |
| endocardial cushion dev | Aug-21 | 0.000436349 | 0.056840914 | 5.865779976 | 45.38393546 | ACVR1;BMP4;TGFB | 8 |
| regulation of cardiac mu | Aug-21 | 0.000436349 | 0.056840914 | 5.865779976 | 45.38393546 | NOTCH1;ERBB4;PIM | 8 |
| regulation of fibroblast | Aug-21 | 0.000436349 | 0.056840914 | 5.865779976 | 45.38393546 | TGFB1;PRKCE;GNA | 8 |
| negative regulation of c | 32/182 | 0.000490927 | 0.062123268 | 2.043922406 | 15.57308573 | CD274;CD84;C5AR2; | 32 |
| cell surface receptor sig | Jul-17 | 0.000578806 | 0.066305504 | 6.669915701 | 49.72116959 | ACVR1;BMP4;BMP2; | 7 |
| glycosaminoglycan cata | 14/56 | 0.000601476 | 0.066305504 | 3.182266878 | 23.60008729 | TGFB1;FUCA1;SDC3 | 14 |
| chondroitin sulfate prot | Sep-27 | 0.000604997 | 0.066305504 | 4.76714135 | 35.32588539 | CSGALNACT1;BCAN | 9 |
| positive regulation of pr | 28/154 | 0.000622172 | 0.066305504 | 2.127390043 | 15.70502004 | SLC27A1;CSF1;CAB | 28 |
| regulation of cellular co | 13/50 | 0.00062411 | 0.066305504 | 3.353436946 | 24.74562588 | SP100;ARHGAP18;N | 13 |
| osteoclast differentiatio | Aug-22 | 0.000628772 | 0.066305504 | 5.446494465 | 40.15015118 | CSF1;SRC;GPR183;O | 8 |
| peptidyl-threonine deph | Aug-22 | 0.000628772 | 0.066305504 | 5.446494465 | 40.15015118 | DUSP4;DUSP5;PPM1 | 8 |
| cardiac cell developmen | Jun-13 | 0.000704608 | 0.069249838 | 8.164296998 | 59.255397 | JAG1;NOTCH1;ALPK | 6 |
| pathway-restricted SMA | Jun-13 | 0.000704608 | 0.069249838 | 8.164296998 | 59.255397 | ACVR1;TGFB2;BMP | 6 |
| plasma membrane tubul | Jun-13 | 0.000704608 | 0.069249838 | 8.164296998 | 59.255397 | BIN3;MICALL1;PAC | 6 |
| transmembrane recepto | 25/133 | 0.000719235 | 0.069249838 | 2.214711387 | 16.02857986 | NRROS;BMPR2;SRC; | 25 |
| pulmonary valve develo | Jul-18 | 0.000869644 | 0.08195006 | 6.063224447 | 42.73013013 | BMP4;TGFB2;BMPR | 7 |
| cartilage development ( | 13/52 | 0.000932947 | 0.085018438 | 3.18111346 | 22.19514302 | WNT2B;TGFB1;SCX; | 13 |
| regulation of peptidyl-ty | 19/92 | 0.000940597 | 0.085018438 | 2.487093072 | 17.33254149 | ACVR1;NRP1;PRNP; | 19 |
| positive regulation of ce | 36/221 | 0.000991085 | 0.087790342 | 1.864734719 | 12.89782906 | SEMA7A;NOTCH1;F | 36 |
| heart morphogenesis (C | 13/53 | 0.001129935 | 0.09131921 | 3.101413848 | 21.04493861 | ACVR1;RYR2;TGFB | 13 |
| positive regulation of ca | Jun-14 | 0.001134016 | 0.09131921 | 7.143364929 | 48.44622974 | NOTCH1;ERBB4;PIM | 6 |
| response to chemokine | Jun-14 | 0.001134016 | 0.09131921 | 7.143364929 | 48.44622974 | LRCH1;DUSP1;WNK | 6 |
| regulation of Rho prote | 16/73 | 0.00119446 | 0.094469015 | 2.680411988 | 18.03933563 | ABCA1;NRP1;KANK | 16 |
| chondroitin sulfate prot | 5-Oct | 0.001304448 | 0.094711472 | 9.521052632 | 63.2385972 | CSGALNACT1;CSGA | 5 |
| Rap protein signal trans | 5-Oct | 0.001304448 | 0.094711472 | 9.521052632 | 63.2385972 | RAP1A;RAP2B;RAP | 5 |
| regulation of axon guid | 5-Oct | 0.001304448 | 0.094711472 | 9.521052632 | 63.2385972 | NRP1;ZSWIM8;ZSWI | 5 |
| regulation of T-helper 1 | 5-Oct | 0.001304448 | 0.094711472 | 9.521052632 | 63.2385972 | TBX21;ZC3H12A;RC | 5 |
| secretory granule organ | 5-Oct | 0.001304448 | 0.094711472 | 9.521052632 | 63.2385972 | ABCA1;F2R;DTNBP1 | 5 |
| protein phosphorylatio | 68/496 | 0.001395849 | 0.099713155 | 1.527983964 | 10.04535222 | DYRK3;BMPR2;FLT1 | 68 |
| chondroitin sulfate meta | Sep-30 | 0.001420867 | 0.099889234 | 4.085443038 | 26.78615722 | CSGALNACT1;BCAN | 9 |
| aortic valve developme | Oct-36 | 0.001506372 | 0.102648161 | 3.667343211 | 23.83058379 | RB1;BMP4;BMP2;BM | 10 |
| positive regulation of pr | Nov-42 | 0.001506464 | 0.102648161 | 3.38426951 | 21.99094904 | PRNP;ADAM17;DOC | 11 |
| outflow tract septum m | Aug-25 | 0.001641084 | 0.10905472 | 4.484604174 | 28.75706849 | ACVR1;BMP4;NRP1; | 8 |
| peptidyl-serine phosph | 27/156 | 0.001649733 | 0.10905472 | 2.002303291 | 12.82904149 | DYRK3;MAST4;CAB | 27 |
| positive regulation of E | 29/172 | 0.001720722 | 0.109998467 | 1.940626538 | 12.3521099 | NRP1;SEMA7A;NOT | 29 |
| circulatory system deve | 23/126 | 0.001726258 | 0.109998467 | 2.134766774 | 13.58095703 | ACVR1;MEF2A;KDM | 23 |
| dermatan sulfate proteo | Jun-15 | 0.001738517 | 0.109998467 | 6.349306653 | 40.34808317 | CSGALNACT1;BCAN | 6 |
| regulation of T cell prol | 16/76 | 0.001868575 | 0.116291663 | 2.545967884 | 15.99524448 | CD274;CEBPB;SFTP | 16 |
| regulation of p38MAP | Oct-37 | 0.001890494 | 0.116291663 | 3.531320238 | 22.14461573 | CYLD;PER1;BMP2;D | 10 |
| endothelial cell develop | Aug-26 | 0.002174783 | 0.123995352 | 4.235225209 | 25.96543035 | RAP1A;RDX;PECAM | 8 |
| establishment of endoth | 5-Nov | 0.002204488 | 0.123995352 | 7.93377193 | 48.53294612 | AFDN;RAB1A;RAP2 | 5 |
| negative regulation of T | 5-Nov | 0.002204488 | 0.123995352 | 7.93377193 | 48.53294612 | TBX21;TNFSF4;ZC3H | 5 |
| positive regulation of m | 5-Nov | 0.002204488 | 0.123995352 | 7.93377193 | 48.53294612 | EGR2;MYRF;ITGAX; | 5 |
| regulation of extrinsic a | 14/64 | 0.002409994 | 0.127351668 | 2.671919619 | 16.10668162 | ACVR1;SP100;SRC;G | 14 |
| chondroitin sulfate bios | Jul-21 | 0.002461868 | 0.127351668 | 4.76317176 | 28.61158567 | CSGALNACT1;BCAN | 7 |
| cellular response to BM | 15/71 | 0.002465614 | 0.127351668 | 2.556547619 | 15.35287182 | ACVR1;TGFB2;BMP | 15 |
| mesenchymal cell differ | Dec-51 | 0.00252973 | 0.127351668 | 2.93486123 | 17.54942163 | TGFB2;BMP2;NOTC | 12 |
| mesenchyme developm | Jun-16 | 0.00255911 | 0.127351668 | 5.714060032 | 34.10205716 | BMP4;BMP2;BMPR2; | 6 |
| platelet-derived growth | Jun-16 | 0.00255911 | 0.127351668 | 5.714060032 | 34.10205716 | NRP1;NR4A3;PDGF | 6 |
| positive regulation of co | Jun-16 | 0.00255911 | 0.127351668 | 5.714060032 | 34.10205716 | BMP4;TGFB1;F2R;SC | 6 |
| positive regulation of m | Jun-16 | 0.00255911 | 0.127351668 | 5.714060032 | 34.10205716 | SEMA7A;NR4A3;DD | 6 |
| negative regulation of tr | 20/108 | 0.00280524 | 0.136516592 | 2.171087533 | 12.75788786 | ACVR1;NOTCH1;TG | 20 |
| BMP signaling pathway | 14/65 | 0.002807499 | 0.136516592 | 2.619383872 | 15.39008827 | ACVR1;TGFB2;BMP | 14 |
| entrainment of circadia | Aug-27 | 0.002835748 | 0.136516592 | 4.012096662 | 23.53275095 | PER1;PER3;ID2;BHL | 8 |
| negative regulation of ly | Oct-39 | 0.002894738 | 0.137858023 | 3.287416978 | 19.2144938 | CD274;IL4I1;CEBPB; | 10 |
| protein dephosphorylati | 24/139 | 0.002992857 | 0.141014485 | 1.994868594 | 11.59323266 | DUSP4;PPP1R15A;D | 24 |

| M1 | | | | | | | |
|---|---|---|---|---|---|---|---|
| **Term** | **Overlap** | **P.value** | **Adjusted.P.value** | **Odds Ratio** | **Combined Score** | **Genes** | **Count** |
| positive regulation of I- | 44/171 | 5.71E-13 | 2.30E-09 | 4.242648026 | 119.6025018 | SLC35B2;CD40;GPR | 44 |
| regulation of I-kappaB | 51/224 | 1.43E-12 | 2.88E-09 | 3.617899678 | 98.65991521 | SLC35B2;CD40;GPR | 51 |
| cytokine-mediated sign | 94/621 | 1.68E-10 | 2.25E-07 | 2.210433417 | 49.75103921 | CIITA;IL1RN;CD40;T | 94 |
| I-kappaB kinase/NF-ka | 22/62 | 4.73E-10 | 4.76E-07 | 6.66915734 | 143.1929653 | TICAM2;RIPK2;TRA | 22 |
| defense response to vir | 31/133 | 1.91E-08 | 1.36E-05 | 3.694760421 | 65.67663153 | RTP4;CD40;CLPB;IFI | 31 |
| interferon-gamma-medi | 21/68 | 2.03E-08 | 1.36E-05 | 5.412262038 | 95.8762125 | CIITA;CAMK2D;SP1 | 21 |
| negative regulation of v | 21/70 | 3.58E-08 | 2.06E-05 | 5.190789474 | 88.99488043 | APOBEC3F;RSAD2; | 21 |
| defense response to sy | 29/124 | 5.01E-08 | 2.52E-05 | 3.707574492 | 62.31958545 | RTP4;CD40;IFIT5;NL | 29 |
| response to cytokine (C | 32/150 | 1.05E-07 | 4.70E-05 | 3.296110344 | 52.95923145 | HDAC4;CD274;CSF1 | 32 |
| death-inducing signalin | 6-Jul | 1.36E-06 | 0.00054507 | 72.1485342 | 974.8023214 | CASP8;TRADD;RIPK | 6 |
| negative regulation of v | 16/54 | 1.81E-06 | 0.000660516 | 5.086039689 | 67.25590904 | APOBEC3F;RSAD2; | 16 |
| regulation of viral geno | 18/67 | 2.02E-06 | 0.000678199 | 4.440484007 | 58.21563376 | APOBEC3F;RSAD2;S | 18 |
| cytoplasmic pattern rec | Nov-27 | 2.25E-06 | 0.000689065 | 8.287295752 | 107.7795742 | IFIH1;CYLD;CASP8; | 11 |
| regulation of interferon | 15/49 | 2.40E-06 | 0.000689065 | 5.326786678 | 68.92944801 | DDX58;RIPK2;LILRE | 15 |
| regulation of tumor nec | 16/58 | 5.10E-06 | 0.001367206 | 4.600655738 | 56.06346717 | TRADD;TNFAIP3;TR | 16 |
| innate immune respons | 46/302 | 6.67E-06 | 0.001586076 | 2.187860577 | 26.07342864 | CD40;SP100;CIITA;C | 46 |
| positive regulation of N | 29/155 | 6.71E-06 | 0.001586076 | 2.790674603 | 33.24339864 | CD40;TRADD;NOD1 | 29 |
| toll-like receptor 3 sign | 7-Dec | 8.91E-06 | 0.001939672 | 16.84198175 | 195.8366424 | CASP8;TNIP2;UNC9 | 7 |
| positive regulation of in | Dec-36 | 9.17E-06 | 0.001939672 | 6.028449967 | 69.92902116 | IFIH1;TBK1;DDX58; | 12 |
| regulation of interleukin | 19/83 | 1.37E-05 | 0.002749621 | 3.588052316 | 40.18474048 | GSDMD;SERPINB1; | 19 |
| cellular response to inte | 24/121 | 1.48E-05 | 0.002835976 | 2.994841963 | 33.30229435 | CIITA;CAMK2D;SP1 | 24 |
| toll-like receptor signali | 20/91 | 1.56E-05 | 0.002855608 | 3.405468974 | 37.68650455 | TICAM2;RIPK2;UNC | 20 |
| positive regulation of in | 70/546 | 1.73E-05 | 0.003024896 | 1.797796617 | 19.71179972 | BCAR3;SLC35B2;GP | 70 |
| regulation of interleukin | 14/51 | 2.09E-05 | 0.00350909 | 4.564824156 | 49.17859452 | CD40;RIPK2;IL16;LI | 14 |
| positive regulation of cy | 48/335 | 2.24E-05 | 0.003594795 | 2.035646023 | 21.79857227 | PTGER4;CD274;CSF | 48 |
| cellular response to typ | 16/65 | 2.47E-05 | 0.003678224 | 3.941920375 | 41.817963 | SP100;RSAD2;MX2;I | 16 |
| type I interferon signali | 16/65 | 2.47E-05 | 0.003678224 | 3.941920375 | 41.817963 | SP100;RSAD2;MX2;I | 16 |
| positive regulation of D | 38/246 | 2.98E-05 | 0.004284355 | 2.218441322 | 23.11531088 | FOXA1;HDAC4;CD4 | 38 |
| inflammatory response | 36/230 | 3.59E-05 | 0.004769169 | 2.252080693 | 23.04678971 | PTGER4;HDAC4;CS | 36 |
| regulation of cytokine-r | 17/74 | 3.65E-05 | 0.004769169 | 3.60125708 | 36.79859999 | TRADD;TNFAIP3;TR | 17 |
| positive regulation of ex | Oct-29 | 3.68E-05 | 0.004769169 | 6.339166008 | 64.72689813 | CYLD;TNFSF10;PM | 10 |
| positive regulation of tr | 128/1183 | 4.84E-05 | 0.006073979 | 1.494391617 | 14.84983325 | FOXA1;ICE1;MAML | 128 |
| TRIF-dependent toll-lik | Oct-30 | 5.14E-05 | 0.006258018 | 6.021881123 | 59.4746353 | IKBKB;CD40;CASP8 | 10 |
| regulation of defense re | Nov-36 | 5.42E-05 | 0.006411206 | 5.301281046 | 52.07122127 | APOBEC3F;TRAF3IP | 11 |
| regulation of macropha | Jul-15 | 5.89E-05 | 0.006763264 | 10.52452738 | 102.5082526 | CASP8;HSF1;LIF;PR | 7 |
| positive regulation of in | Aug-20 | 6.62E-05 | 0.007360081 | 8.022178735 | 77.19113649 | IFIH1;TBK1;DDX58; | 8 |
| positive regulation of ax | Dec-43 | 6.77E-05 | 0.007360081 | 4.665414882 | 44.78706675 | TIAM2;MACF1;TIAM | 12 |
| MyD88-independent to | Oct-31 | 7.06E-05 | 0.007471156 | 5.734813847 | 54.81424855 | IKBKB;CD40;CASP8 | 10 |
| regulation of extrinsic a | 15/64 | 8.09E-05 | 0.008342567 | 3.693128628 | 34.79614031 | SP100;SRC;TRADD; | 15 |
| positive regulation of pe | 24/134 | 8.37E-05 | 0.008412002 | 2.639036376 | 24.77594244 | LYN;CSF1R;CD40;S | 24 |
| neural crest cell develop | Dec-45 | 0.000109964 | 0.010781881 | 4.382186813 | 39.94518123 | TBX1;SEMA4A;SEM | 12 |
| negative chemotaxis (G | Oct-33 | 0.000127846 | 0.012236655 | 5.23556641 | 46.93521361 | SEMA4A;EPHA7;SE | 10 |
| positive regulation of de | Sep-28 | 0.000167988 | 0.015704895 | 5.701525354 | 49.55549105 | APOBEC3F;TRAF3IP | 9 |
| positive regulation of ty | 16/77 | 0.000216231 | 0.019755675 | 3.164396668 | 26.70485614 | DDX58;RIPK2;RELA | 16 |
| regulation of extrinsic a | Nov-42 | 0.000251437 | 0.022461689 | 4.273835125 | 35.42290807 | SP100;CASP8;TRAD | 11 |
| positive regulation of cy | Oct-36 | 0.000283748 | 0.024269497 | 4.630708938 | 37.82096587 | TICAM2;NLRC5;RIP | 10 |
| positive regulation of m | Oct-36 | 0.000283748 | 0.024269497 | 4.630708938 | 37.82096587 | CCR1;CASP8;OCSTA | 10 |
| extrinsic apoptotic sign | 15/72 | 0.000327113 | 0.027395722 | 3.173415189 | 25.46730606 | DAXX;PDPK1;CD70; | 15 |
| regulation of erythrocyt | Oct-37 | 0.000362238 | 0.029125808 | 4.458959286 | 35.32926815 | LYN;STAT5B;ZFP36; | 10 |
| negative regulation of in | Jun-14 | 0.000362261 | 0.029125808 | 9.01514658 | 71.42831189 | TLR8;LILRB1;CCR7; | 6 |
| activation of cysteine-ty | 16/81 | 0.000396694 | 0.030635392 | 2.969018916 | 23.25438292 | TNFSF15;TRADD;RP | 16 |
| regulation of interferon | Aug-25 | 0.000401536 | 0.030635392 | 5.66117954 | 44.27163256 | IFIH1;TBK1;DDX58; | 8 |
| positive regulation of tr | 98/908 | 0.000403899 | 0.030635392 | 1.47977208 | 11.56344895 | FOXA1;MAML2;NR | 98 |
| cellular response to gro | 25/158 | 0.000447485 | 0.033231862 | 2.272253854 | 17.52332266 | FOXC1;BMPR2;PDG | 25 |
| positive regulation of ty | 13/59 | 0.000454665 | 0.033231862 | 3.405545755 | 26.20891035 | LYN;CSF1R;CD40;T | 13 |
| cellular response to cyt | 58/482 | 0.000472316 | 0.033905568 | 1.66355488 | 12.73927289 | IL1RN;CD40;TRADD | 58 |
| cytoplasmic pattern rec | 5-Oct | 0.000489288 | 0.034507696 | 12.01432292 | 91.57988309 | IFIH1;CLPB;DDX58; | 5 |
| regulation of programm | Jul-20 | 0.00050325 | 0.034880453 | 6.474877144 | 49.17295509 | CYLD;PDCD6IP;CAS | 7 |
| positive regulation of in | 13/61 | 0.000638611 | 0.043512155 | 3.263293521 | 24.00548773 | DDX58;NOD1;NOD2 | 13 |
| positive regulation of in | 13/62 | 0.000751903 | 0.049365635 | 3.196522064 | 22.99227528 | GSDMD;RIPK2;PYH | 13 |
| stress-activated MAPK | 13/62 | 0.000751903 | 0.049365635 | 3.196522064 | 22.99227528 | PTGER4;RIPK2;GPS1 | 13 |
| negative regulation of c | 27/182 | 0.000761361 | 0.049365635 | 2.105970086 | 15.12171504 | PTGER4;CD274;NLR | 27 |
| negative regulation of e | Sep-34 | 0.000837218 | 0.052773285 | 4.331749347 | 30.69229153 | CASP8;TRADD;TNF | 9 |
| positive regulation of m | 5-Nov | 0.000840172 | 0.052773285 | 10.01139323 | 70.8997286 | CASP8;HSF1;LIF;PR | 5 |
| nucleotide-binding olig | Aug-28 | 0.000937235 | 0.057300805 | 4.81121983 | 33.54659991 | CYLD;CASP8;RIPK2 | 8 |
| positive regulation of re | 22/139 | 0.000953275 | 0.057300805 | 2.270522105 | 15.79285972 | PTGER4;ABCC1;FEM | 22 |
| osteoclast differentiatio | Jul-22 | 0.000965718 | 0.057300805 | 5.61095176 | 38.95481112 | FARP2;CSF1R;SBNO | 7 |
| positive regulation of in | Dec-56 | 0.000969267 | 0.057300805 | 3.284678043 | 22.79228265 | PANX1;GSDMD;P2R | 12 |
| negative regulation of e | 15/80 | 0.001050539 | 0.059481245 | 2.781631213 | 19.07768287 | SRC;TRADD;TNFAI | 15 |
| regulation of small GT | 22/141 | 0.00115605 | 0.06320403 | 2.232118654 | 15.09525163 | ARHGAP11B;SRC;A | 22 |
| regulation of intracellul | 52/437 | 0.00116072 | 0.06320403 | 1.639465169 | 11.08067771 | ARHGAP11B;EHMT | 52 |
| regulation of type I inte | 16/89 | 0.001163457 | 0.06320403 | 2.642497193 | 17.8536605 | DDX58;NLRC5;TNF | 16 |
| positive regulation of cy | Jun-17 | 0.001220687 | 0.065428797 | 6.555404205 | 43.97589251 | NOD2;TICAM1;HIF1 | 6 |
| regulation of necroptoti | Jul-23 | 0.001296705 | 0.068213834 | 5.259982073 | 34.96798815 | CYLD;PDCD6IP;CAS | 7 |
| regulation of osteoclast | Sep-36 | 0.001306583 | 0.068213834 | 4.010443864 | 26.63070942 | CCR1;CEBPB;MAFB; | 9 |
| tumor necrosis factor-m | 19/116 | 0.001361197 | 0.070154003 | 2.363127735 | 15.59520335 | EDARADD;CD40;TN | 19 |
| protein-containing com | 35/267 | 0.001403969 | 0.07144249 | 1.8258718 | 11.99315096 | AMER1;CD40;TRAD | 35 |
| neural crest cell migrati | Sep-37 | 0.001610124 | 0.080908732 | 3.867003916 | 24.87041942 | TBX1;SEMA4A;SEM | 9 |
| nucleotide-binding dom | Jul-24 | 0.001710125 | 0.083957266 | 4.950302937 | 31.53931614 | CYLD;CASP8;RIPK2 | 7 |
| programmed necrotic ce | Jun-18 | 0.001712561 | 0.083957266 | 6.008794788 | 38.27461246 | CYLD;TICAM2;PPIF; | 6 |
| positive regulation of in | 14/76 | 0.001816498 | 0.087979774 | 2.720472358 | 17.16847913 | DDX58;CYBA;IL16; | 14 |
| regulation of tyrosine p | 13/68 | 0.001844703 | 0.088282232 | 2.846882437 | 17.92236846 | LYN;CSF1R;CD40;T | 13 |
| cellular response to lipo | 19/120 | 0.002042896 | 0.096014827 | 2.269044119 | 14.05306776 | LYN;CD274;TICAM2 | 19 |
| toll-like receptor 9 sign | May-13 | 0.002054049 | 0.096014827 | 7.50773112 | 46.45740875 | UNC93B1;TNIP2;TL | 5 |
| pattern recognition rece | Dec-61 | 0.002116 | 0.097773789 | 2.94870597 | 18.1588031 | TRAF3;UNC93B1;TI | 12 |
| mesenchymal cell devel | Jun-19 | 0.002341301 | 0.105753152 | 5.546279128 | 33.59408149 | FOXC1;FAM172A;T | 6 |
| positive regulation of o | Jun-19 | 0.002341301 | 0.105753152 | 5.546279128 | 33.59408149 | PDP1;GDNF;ABL2;N | 6 |

| M2 | | | | | | | |
|---|---|---|---|---|---|---|---|
| **Term** | **Overlap** | **P.value** | **Adjusted.P.value** | **Odds Ratio** | **Combined Score** | **Genes** | **Count** |
| positive regulation of a | 19/43 | 8.30E-05 | 0.421623745 | 3.538092628 | 33.24433504 | ROBO2;MACF1;SEM | 19 |
| regulation of GTPase ac | 54/189 | 0.000367342 | 0.856957887 | 1.792695075 | 14.17881702 | USP6NL;ARHGAP11 | 54 |
| negative regulation of t | 24/68 | 0.000670737 | 0.856957887 | 2.438080032 | 17.81537683 | FBN2;SMAD2;ZNF4 | 24 |
| negative regulation of t | 34/108 | 0.000675169 | 0.856957887 | 2.055556382 | 15.00668794 | ZNF451;DLX1;UBE2 | 34 |
| camera-type eye morph | 7-Nov | 0.001144021 | 0.999996202 | 7.804971319 | 52.86467608 | SKI;TENM3;NF1;SH | 7 |
| regulation of transform | 31/100 | 0.001504919 | 0.999996202 | 2.008953868 | 13.05622379 | ZNF451;CITED2;LDL | 31 |
| positive regulation of n | Nov-24 | 0.001833225 | 0.999996202 | 3.775877663 | 23.79436737 | BACE1;PRNP;EPHA | 11 |
| embryonic limb morph | 14/36 | 0.003102445 | 0.999996202 | 2.840473703 | 16.40534022 | OSR2;HDAC2;CREB | 14 |
| death-inducing signalin | 5-Jul | 0.003124298 | 0.999996202 | 11.14523615 | 64.29180211 | CASP8;TRADD;RIPK | 5 |
| negative regulation of p | Oct-22 | 0.003178568 | 0.999996202 | 3.717878622 | 21.38272658 | BMP4;CDA;CDKN1A | 10 |
| positive regulation of a | Oct-22 | 0.003178568 | 0.999996202 | 3.717878622 | 21.38272658 | MACF1;SEMA7A;GC | 10 |
| negative regulation of i | 52/198 | 0.00352867 | 0.999996202 | 1.594269002 | 9.002572839 | DGKG;MAPKBP1;SF | 52 |
| regulation of small GT | 39/141 | 0.004156117 | 0.999996202 | 1.709999514 | 9.376224849 | ARHGAP11A;ARHG | 39 |
| neural crest cell migrati | 14/37 | 0.00416051 | 0.999996202 | 2.716808263 | 14.89386253 | TBX1;SEMA4A;SEM | 14 |
| negative regulation of C | Oct-23 | 0.004709377 | 0.999996202 | 3.431677672 | 18.38761408 | ARFGEF1;GPSM1;K | 10 |
| BMP signaling pathway | 21/65 | 0.004798199 | 0.999996202 | 2.131565172 | 11.38152336 | ACVR1;SMAD2;SM | 21 |
| negative regulation of t | 152/684 | 0.005130161 | 0.999996202 | 1.28392654 | 6.769654551 | RB1;ZNF451;SPI1;UE | 152 |
| regulation of plasma me | 22/70 | 0.005714099 | 0.999996202 | 2.04703785 | 10.57257937 | USP6NL;RABGAP1; | 22 |
| regulation of pathway-r | 19/58 | 0.006018025 | 0.999996202 | 2.175285115 | 11.12222438 | ACVR1;GDF11;SMA | 19 |
| regulation of establishm | Aug-17 | 0.006365448 | 0.999996202 | 3.964298725 | 20.04694615 | PPP1R12A;PROC;RO | 8 |
| cellular response to gro | 42/158 | 0.006490368 | 0.999996202 | 1.619225136 | 8.156743193 | DLX1;FOXC1;FLT1; | 42 |
| activation of GTPase ac | 30/105 | 0.006745914 | 0.999996202 | 1.78746564 | 8.935215993 | USP6NL;TBC1D9B;R | 30 |
| cellular response to BM | 22/71 | 0.006874245 | 0.999996202 | 2.005138424 | 9.985536273 | ACVR1;SMAD2;DLX | 22 |
| positive regulation of S | Jul-14 | 0.00711084 | 0.999996202 | 4.459164163 | 22.05562776 | BMP4;SMAD4;RBPM | 7 |
| response to chemokine | Jul-14 | 0.00711084 | 0.999996202 | 4.459164163 | 22.05562776 | FOXC1;LRCH1;DUS | 7 |
| smooth muscle contract | Jul-14 | 0.00711084 | 0.999996202 | 4.459164163 | 22.05562776 | SMTN;ROCK1;MYH | 7 |
| beta-catenin destruction | Sep-21 | 0.00811749 | 0.999996202 | 3.345176278 | 16.10278977 | AMER1;FZD1;FZD5; | 9 |
| regulation of cilium ass | 20/64 | 0.008755738 | 0.999996202 | 2.029505582 | 9.615890771 | USP6NL;RABGAP1; | 20 |
| regulation of organelle | 23/77 | 0.009384208 | 0.999996202 | 1.902118579 | 8.880472419 | USP6NL;STIL;PDCD | 23 |
| embryonic axis specific | Aug-18 | 0.009638613 | 0.999996202 | 3.567650273 | 16.5609544 | SMAD2;STIL;FZD5;P | 8 |
| positive regulation of hi | Jul-15 | 0.011245783 | 0.999996202 | 3.901529637 | 17.50913654 | DNMT1;SMAD4;KM | 7 |
| regulation of macropha | Jul-15 | 0.011245783 | 0.999996202 | 3.901529637 | 17.50913654 | CASP8;CSF1;HSF1;L | 7 |
| regulation of non-canon | Jul-15 | 0.011245783 | 0.999996202 | 3.901529637 | 17.50913654 | TIAM1;LBX2;RNF21 | 7 |
| negative regulation of c | 14/41 | 0.011629962 | 0.999996202 | 2.313750532 | 10.30583947 | IL1RN;AKNA;JAG1; | 14 |
| neural crest cell develop | 15/45 | 0.011664578 | 0.999996202 | 2.231316726 | 9.932033836 | TBX1;SEMA4A;SEM | 15 |
| ganglion development ( | 6-Dec | 0.012719497 | 0.999996202 | 4.458219552 | 19.45843085 | FZD3;SEMA3A;CTN | 6 |
| mesoderm morphogene | 6-Dec | 0.012719497 | 0.999996202 | 4.458219552 | 19.45843085 | CRB2;EOMES;SMAD | 6 |
| regulation of actin filam | Oct-26 | 0.012876084 | 0.999996202 | 2.787725533 | 12.1332509 | PTGER4;ROCK1;SYN | 10 |
| negative regulation of p | 5-Sep | 0.013522897 | 0.999996202 | 5.571935572 | 23.97810575 | PMEPA1;EMILIN1;L | 5 |
| semaphorin-plexin sign | 5-Sep | 0.013522897 | 0.999996202 | 5.571935572 | 23.97810575 | PLXNA2;PLXNA1;P | 5 |
| positive regulation of B | Nov-30 | 0.013759528 | 0.999996202 | 2.582545371 | 11.06885073 | VAV3;CD40;MEF2C; | 11 |
| regulation of histone H | Aug-19 | 0.01401366 | 0.999996202 | 3.243119722 | 13.84073568 | DNMT1;SMAD4;KM | 8 |
| regulation of mitochon | Aug-19 | 0.01401366 | 0.999996202 | 3.243119722 | 13.84073568 | ARHGAP11B;HEBP2 | 8 |
| regulation of B cell pro | 15/46 | 0.014439997 | 0.999996202 | 2.159206309 | 9.150183848 | VAV3;CD40;MEF2C; | 15 |
| positive regulation of R | 19/63 | 0.01551253 | 0.999996202 | 1.927501931 | 8.030179715 | NOTCH2;RTN4R;LR | 19 |
| regulation of cellular co | 36/139 | 0.016675724 | 0.999996202 | 1.561748856 | 6.393489415 | CAMK2D;TFRC;RO | 36 |
| positive regulation of G | 52/214 | 0.017131457 | 0.999996202 | 1.435390036 | 5.837500078 | USP6NL;ARHGAP11 | 52 |
| chondroitin sulfate prot | Oct-27 | 0.017157737 | 0.999996202 | 2.62358087 | 10.66565927 | BCAN;VCAN;CHPF; | 10 |
| positive regulation of R | Nov-31 | 0.017869877 | 0.999996202 | 2.453267706 | 9.873516473 | RTN4R;P2RY8;SYNP | 11 |
| odontogenesis (GO:00 | 13/39 | 0.018222449 | 0.999996202 | 2.230642955 | 8.933950299 | TBX1;OSR2;FOXC1; | 13 |
| regulation of peptidyl-t | 13/39 | 0.018222449 | 0.999996202 | 2.230642955 | 8.933950299 | CAB39;SPHK1;AXIN | 13 |
| negative regulation of t | 199/948 | 0.018228241 | 0.999996202 | 1.193487925 | 4.779660382 | RB1;SPI1;EHMT1;NC | 199 |
| negative regulation of d | 22/77 | 0.018492706 | 0.999996202 | 1.785737795 | 7.12577038 | TBX1;FOXC1;SPI1;T | 22 |
| eye morphogenesis (G | Aug-20 | 0.019671949 | 0.999996202 | 2.972677596 | 11.67834689 | FBN2;SKI;NIPBL;TE | 8 |
| protein localization to p | 35/136 | 0.019835452 | 0.999996202 | 1.548198172 | 6.069377207 | PLEKHF1;ROCK1;R | 35 |
| dermatan sulfate metab | Jun-13 | 0.020006391 | 0.999996202 | 3.821096981 | 14.94699854 | BCAN;VCAN;IDUA; | 6 |
| positive regulation of n | Jun-13 | 0.020006391 | 0.999996202 | 3.821096981 | 14.94699854 | SRGAP2C;NIPBL;SE | 6 |
| semaphorin-plexin sign | Jun-13 | 0.020006391 | 0.999996202 | 3.821096981 | 14.94699854 | SEMA3A;PLXNA2;P | 6 |
| ameboidal-type cell mig | 16/52 | 0.020780127 | 0.999996202 | 1.983205549 | 7.682458731 | TBX1;SEMA4A;SEM | 16 |
| mammary gland develo | Sep-24 | 0.021422997 | 0.999996202 | 2.675649084 | 10.2832962 | TGFB3;ERBB4;GPAT | 9 |
| regulation of axon exte | Sep-24 | 0.021422997 | 0.999996202 | 2.675649084 | 10.2832962 | SEMA4A;SEMA7A;S | 9 |
| cellular response to inte | Oct-28 | 0.022403961 | 0.999996202 | 2.477674503 | 9.411490015 | CEBPA;SOCS3;SBN | 10 |
| positive regulation of cy | Oct-28 | 0.022403961 | 0.999996202 | 2.477674503 | 9.411490015 | P2RY8;GPR35;LPAR | 10 |
| positive regulation of o | 13/40 | 0.022578274 | 0.999996202 | 2.147894817 | 8.14216915 | FBN2;ACVR1;OSR2; | 13 |
| regulation of cytokine-r | 21/74 | 0.022652841 | 0.999996202 | 1.768623474 | 6.698608383 | RNASEL;SPPL2B;TR | 21 |
| chromatin remodeling a | Nov-32 | 0.022826345 | 0.999996202 | 2.336302199 | 8.830848329 | CENPT;HELLS;CENP | 11 |
| positive regulation of b | Dec-36 | 0.022846329 | 0.999996202 | 2.230306346 | 8.42824927 | FBN2;ACVR1;BMP4; | 12 |
| regulation of ossificatio | Dec-36 | 0.022846329 | 0.999996202 | 2.230306346 | 8.42824927 | ACVR1;BMP4;MEF2 | 12 |
| peptidyl-serine phosph | 39/156 | 0.023005036 | 0.999996202 | 1.489391017 | 5.618045641 | CAMK2D;MAST4;CA | 39 |
| adherens junction asse | 5-Oct | 0.023005731 | 0.999996202 | 4.457275457 | 16.81289606 | TBCD;CTNNB1;PIP5 | 5 |
| chloride ion homeostas | 5-Oct | 0.023005731 | 0.999996202 | 4.457275457 | 16.81289606 | SLC12A4;SLC12A6;S | 5 |
| cytoplasmic pattern rec | 5-Oct | 0.023005731 | 0.999996202 | 4.457275457 | 16.81289606 | IFIH1;RNF135;CLPB; | 5 |
| ectoderm development ( | 5-Oct | 0.023005731 | 0.999996202 | 4.457275457 | 16.81289606 | ITGAM;SMURF1;ITG | 5 |
| monovalent inorganic a | 5-Oct | 0.023005731 | 0.999996202 | 4.457275457 | 16.81289606 | SLC12A4;SLC12A6;S | 5 |
| negative regulation of h | 5-Oct | 0.023005731 | 0.999996202 | 4.457275457 | 16.81289606 | GJA1;RAB11FIP1;LI | 5 |
| negative regulation of s | 5-Oct | 0.023005731 | 0.999996202 | 4.457275457 | 16.81289606 | CAMK2D;NEDD4;PR | 5 |
| negative regulation of t | 5-Oct | 0.023005731 | 0.999996202 | 4.457275457 | 16.81289606 | CREB1;DACH1;BHL | 5 |
| nerve growth factor sig | 5-Oct | 0.023005731 | 0.999996202 | 4.457275457 | 16.81289606 | RAP1A;BDNF;MAGI | 5 |
| primary miRNA proces | 5-Oct | 0.023005731 | 0.999996202 | 4.457275457 | 16.81289606 | SMAD2;SMAD1;SM | 5 |
| Rap protein signal trans | 5-Oct | 0.023005731 | 0.999996202 | 4.457275457 | 16.81289606 | RAP2C;RAP1A;SGS | 5 |
| regulation of macropha | 5-Oct | 0.023005731 | 0.999996202 | 4.457275457 | 16.81289606 | CX3CR1;CSF1;MST1 | 5 |
| secretory granule organ | 5-Oct | 0.023005731 | 0.999996202 | 4.457275457 | 16.81289606 | ABCA1;F2R;DTNBP1 | 5 |
| Wnt signaling pathway | 5-Oct | 0.023005731 | 0.999996202 | 4.457275457 | 16.81289606 | FZD1;CTNNB1;WNT | 5 |
| positive regulation of ce | 25/92 | 0.023538986 | 0.999996202 | 1.665942863 | 6.245781801 | ICE1;TFRC;FAF1;SY | 25 |
| positive regulation of tr | 25/92 | 0.023538986 | 0.999996202 | 1.665942863 | 6.245781801 | CRB2;NUMA1;CITE | 25 |
| regulation of bone min | 17/57 | 0.023769492 | 0.999996202 | 1.896494111 | 7.091659786 | ACVR1;FBN2;OSR2; | 17 |
| regulation of cell cycle | 68/296 | 0.025096257 | 0.999996202 | 1.334152047 | 4.916399073 | RB1;CDKN1A;PPP1R | 68 |

| | | | | | | | |
|---|---|---|---|---|---|---|---|
| negative regulation of M | 18/94 | 0.003060973 | 0.142705779 | 2.261609907 | 13.09251057 | DUSP4;DUSP1;SPRY | 18 |
| positive regulation of pı | 42/286 | 0.003129323 | 0.144372616 | 1.649336079 | 9.51161994 | TRIO;PSEN2;NLRC4; | 42 |
| negative regulation of c | 15/73 | 0.003272693 | 0.149007995 | 2.468117132 | 14.12291666 | CD164;NOTCH1;JAG | 15 |
| intrinsic apoptotic signa | 19/102 | 0.003297084 | 0.149007995 | 2.186229542 | 12.49368299 | PPP1R15A;MOAP1;B | 19 |
| regulation of cell cycle ( | 43/296 | 0.003450004 | 0.149258609 | 1.628590109 | 9.233096203 | RB1;PPP1R15A;NOT | 43 |
| cardiocyte differentiatio | 5-Dec | 0.003484402 | 0.149258609 | 6.8 | 38.48431966 | KDM6B;BMP2;TGFB | 5 |
| positive regulation of m | 5-Dec | 0.003484402 | 0.149258609 | 6.8 | 38.48431966 | ITGAM;DDX58;CAM | 5 |
| regulation of CD4-posi | 5-Dec | 0.003484402 | 0.149258609 | 6.8 | 38.48431966 | IRF4;TNFSF4;RUNX | 5 |
| regulation of protein kir | 20/110 | 0.003496353 | 0.149258609 | 2.122605364 | 12.00552999 | RB1;CSF1;C5AR1;DT | 20 |
| phosphorylation (GO:0 | 55/400 | 0.003554492 | 0.149258609 | 1.529573051 | 8.626093104 | DYRK3;BMPR2;MAS | 55 |
| positive regulation of ca | Jun-17 | 0.003639279 | 0.149258609 | 5.194312796 | 29.17110259 | NOTCH1;ERBB4;PIM | 6 |
| positive regulation of co | Jun-17 | 0.003639279 | 0.149258609 | 5.194312796 | 29.17110259 | BMP4;TGFB1;F2R;SC | 6 |
| regulation of ERK1 anc | 36/238 | 0.003641506 | 0.149258609 | 1.70618057 | 9.580814514 | NRP1;SEMA7A;NOT | 36 |
| photoperiodism (GO:0( | Aug-28 | 0.003643302 | 0.149258609 | 3.81128097 | 21.39982804 | PER1;PER3;ID2;BHLI | 8 |
| cellular response to vas | Sep-34 | 0.003677965 | 0.149258609 | 3.431012658 | 19.23218385 | ERN1;NRP1;FLT1;NC | 9 |
| Notch signaling pathwa | 13/60 | 0.003707033 | 0.149258609 | 2.638477801 | 14.76894172 | NOTCH1;JAG1;MAM | 13 |
| positive regulation of G | 33/214 | 0.003856406 | 0.153515466 | 1.744705341 | 9.697106356 | USP6NL;BCAR3;GPR | 33 |
| cellular response to org | 25/150 | 0.004023654 | 0.153515466 | 1.911702128 | 10.54411711 | KCNE1;AQP9;PDGFI | 25 |
| negative regulation of c | Oct-41 | 0.004284412 | 0.155538219 | 3.074985105 | 16.76719223 | BMP2;NOTCH1;JAG | 10 |
| negative regulation of tı | 14/68 | 0.004333489 | 0.156040825 | 2.473451241 | 13.45899395 | NRROS;TGFB1;LDLI | 14 |
| artery development (GC | Jul-23 | 0.00438181 | 0.156508352 | 4.167314278 | 22.62973939 | NRP1;BMPR2;NF1;HI | 7 |
| positive regulation of nı | 67/511 | 0.004472709 | 0.158477028 | 1.449157672 | 7.839596658 | SCX;FGF2;ETS1;ZMI | 67 |
| positive regulation of ce | 38/258 | 0.004540702 | 0.159609285 | 1.65372255 | 8.921293404 | YAP1;CEBPB;BTG1;I | 38 |
| negative regulation of ir | Aug-29 | 0.004617769 | 0.161040162 | 3.629590582 | 19.51937014 | CD274;PRNP;TNFSF | 8 |
| positive regulation of ex | Jun-18 | 0.005024102 | 0.169860655 | 4.7611901 | 25.20340092 | RB1;BMP2;TGFB1;M | 6 |
| positive regulation of ne | Jun-18 | 0.005024102 | 0.169860655 | 4.7611901 | 25.20340092 | RIMS2;EGR2;MYRF; | 6 |
| regulation of B cell diff | Jun-18 | 0.005024102 | 0.169860655 | 4.7611901 | 25.20340092 | IL4I1;MMP14;ID2;SL | 6 |
| regulation of cartilage d | Jun-18 | 0.005024102 | 0.169860655 | 4.7611901 | 25.20340092 | BMP4;BMP2;BMPR2; | 6 |
| peptidyl-serine modifica | 27/169 | 0.005209878 | 0.17388919 | 1.817677631 | 9.555892825 | DYRK3;MAST4;CAB | 27 |
| cardiac right ventricle m | May-13 | 0.00522178 | 0.17388919 | 5.949671053 | 31.26502755 | BMP4;TGFB2;JAG1;Z | 5 |
| endoderm formation (G | Sep-36 | 0.005551018 | 0.182114499 | 3.176511955 | 16.4980852 | DUSP4;DUSP5;MMP | 9 |
| regulation of ossificatio | Sep-36 | 0.005551018 | 0.182114499 | 3.176511955 | 16.4980852 | ACVR1;BMP4;TGFB2 | 9 |
| regulation of endothelia | 17/92 | 0.005726775 | 0.186499157 | 2.163418079 | 11.16886818 | NRP1;BMPR2;EGR3; | 17 |
| plasma membrane invag | Aug-30 | 0.005780282 | 0.186867659 | 3.464417501 | 17.8531924 | ITGAM;ITGB2;SPIRE | 8 |
| regulation of anatomica | 21/123 | 0.00600687 | 0.192387274 | 1.966263894 | 10.05714763 | PHLDB1;BRSK2;SP1 | 21 |
| regulation of transformi | 18/100 | 0.006037894 | 0.192387274 | 2.095428283 | 10.70701001 | TGFB1;ITGA3;LDLR | 18 |
| myeloid leukocyte diffe | 14/71 | 0.006470859 | 0.195108576 | 2.342879939 | 11.80916081 | CSF1;LEF1;TREM2;G | 14 |
| negative regulation of tı | 85/684 | 0.00651798 | 0.195108576 | 1.364141701 | 6.86598546 | RB1;WWC2;ZBTB21; | 85 |
| positive regulation of eı | 16/86 | 0.006750801 | 0.195108576 | 2.181048174 | 10.90108405 | NRP1;BMPR2;TGFB1 | 16 |
| atrioventricular valve m | Jun-19 | 0.006759324 | 0.195108576 | 4.394701665 | 21.95958751 | ACVR1;TGFB2;BMP2 | 6 |
| endocardial cushion mo | Jun-19 | 0.006759324 | 0.195108576 | 4.394701665 | 21.95958751 | ACVR1;BMP4;TGFB2 | 6 |
| heparan sulfate proteog | Jun-19 | 0.006759324 | 0.195108576 | 4.394701665 | 21.95958751 | CSGALNACT1;HS3S | 6 |
| regulation of collagen b | Jun-19 | 0.006759324 | 0.195108576 | 4.394701665 | 21.95958751 | BMP4;TGFB1;F2R;SC | 6 |
| regulation of oligodend | Jun-19 | 0.006759324 | 0.195108576 | 4.394701665 | 21.95958751 | NOTCH1;SLC45A3;N | 6 |
| cellular response to cAI | Aug-31 | 0.007152565 | 0.195108576 | 3.313607298 | 16.37016188 | KCNE1;RAP1A;AQP | 8 |
| proteoglycan biosynthe | Aug-31 | 0.007152565 | 0.195108576 | 3.313607298 | 16.37016188 | CSGALNACT1;HS3S | 8 |
| cellular polysaccharide | Jul-25 | 0.007257883 | 0.195108576 | 3.70386957 | 18.24402826 | CSGALNACT1;EXT1 | 7 |
| positive regulation of re | Jul-25 | 0.007257883 | 0.195108576 | 3.70386957 | 18.24402826 | MLLT11;MOAP1;BCI | 7 |
| atrioventricular valve de | May-14 | 0.007492766 | 0.195208603 | 5.288304094 | 25.87999353 | ACVR1;TGFB2;BMP2 | 5 |
| regulation of protein mo | Nov-51 | 0.007575303 | 0.196204785 | 2.621502112 | 12.80043276 | SPRED3;SPRED2;SPI | 11 |
| extracellular structure o | 32/216 | 0.007965725 | 0.20511743 | 1.663084101 | 8.037032307 | ITGAM;PAPLN;ITGB | 32 |
| ventricular septum deve | Sep-38 | 0.008086221 | 0.207016607 | 2.957114797 | 14.24617783 | ACVR1;TGFB2;BMPI | 9 |
| cellular response to org | 18/103 | 0.008235467 | 0.209625772 | 2.021135322 | 9.700045224 | EGR1;KCNE1;AQP9; | 18 |
| negative regulation of T | Oct-45 | 0.008599319 | 0.216399915 | 2.722955145 | 12.95057137 | CD274;IL4I1;CEBPB;I | 10 |
| regulation of pri-miRN | Oct-45 | 0.008599319 | 0.216399915 | 2.722955145 | 12.95057137 | BMP4;EGR1;JUN;TGI | 10 |
| regulation of cell migrat | Aug-32 | 0.008756713 | 0.218754631 | 3.175364611 | 15.04467011 | STARD13;NOTCH1; | 8 |
| cardiac epithelial to mes | Jun-20 | 0.008890457 | 0.218754631 | 4.08056872 | 19.27161544 | BMP4;TGFB2;BMP2; | 6 |
| mesoderm formation (C | Jun-20 | 0.008890457 | 0.218754631 | 4.08056872 | 19.27161544 | KDM6B;BMPR2;ITG | 6 |
| toll-like receptor 4 signa | Jun-20 | 0.008890457 | 0.218754631 | 4.08056872 | 19.27161544 | ITGAM;TICAM2;PRK | 6 |
| negative regulation of p | Jul-26 | 0.009130609 | 0.222194869 | 3.508734956 | 16.47745062 | KCNE1;DAB2;TGFB1 | 7 |
| regulation of myelinatio | Jul-26 | 0.009130609 | 0.222194869 | 3.508734956 | 16.47745062 | EGR2;MYRF;ITGAX; | 7 |
| regulation of endothelia | 16/89 | 0.009408544 | 0.22770733 | 2.091067971 | 9.757209769 | NRP1;SP100;BMPR2; | 16 |
| regulation of peptidyl-tl | Sep-39 | 0.009643669 | 0.230874644 | 2.858386076 | 13.26706651 | RPTOR;SPRED3;SPR | 9 |
| regulation of protein tyı | Sep-39 | 0.009643669 | 0.230874644 | 2.858386076 | 13.26706651 | PRNP;RAP2B;DLG3;I | 9 |
| regulation of MAP kina | 17/97 | 0.009770079 | 0.232643442 | 2.027641684 | 9.384799058 | TGFB1;FLT1;DUSP1; | 17 |
| positive regulation of pı | 25/161 | 0.009979588 | 0.236361462 | 1.756003598 | 8.090283509 | SRC;PDGFB;PDGFA; | 25 |
| negative regulation of p | 16/90 | 0.010458922 | 0.24639662 | 2.062695835 | 9.406511623 | RB1;TESK1;DTNBP1; | 16 |
| interferon-gamma-medi | 13/68 | 0.011045192 | 0.252160591 | 2.253699789 | 10.15463053 | SP100;IFNGR1;IFNG | 13 |
| regulation of cell junctio | Jul-27 | 0.011333815 | 0.253823733 | 3.333113804 | 14.93223177 | RAP1A;RAPGEF1;SN | 7 |
| regulation of vasculatur | Jul-27 | 0.011333815 | 0.253823733 | 3.333113804 | 14.93223177 | SP100;PLXND1;NF1; | 7 |
| positive regulation of os | Sep-40 | 0.011416929 | 0.253823733 | 2.76602695 | 12.37149274 | ACVR1;BMP4;P2RX7 | 9 |
| positive regulation of va | Jun-21 | 0.011461898 | 0.253823733 | 3.808320169 | 17.01834299 | ERN1;JUN;NR4A3;FC | 6 |
| regulation of lipopolysa | Jun-21 | 0.011461898 | 0.253823733 | 3.808320169 | 17.01834299 | TRIM5;CARD8;CD18 | 6 |
| response to progesteror | Jun-21 | 0.011461898 | 0.253823733 | 3.808320169 | 17.01834299 | YAP1;TGFB2;TGFB1 | 6 |
| epidermal growth factoı | Oct-47 | 0.011730213 | 0.257193633 | 2.575483135 | 11.4495355 | ADAM17;TGFB1;GA | 10 |
| epithelial to mesenchym | Oct-47 | 0.011730213 | 0.257193633 | 2.575483135 | 11.4495355 | TGFB2;BMP2;NOTCI | 10 |
| extracellular matrix org | 41/300 | 0.011796628 | 0.257375692 | 1.51473147 | 6.725319188 | ITGAM;PAPLN;ITGB | 41 |
| positive regulation of M | 38/274 | 0.011936382 | 0.259148216 | 1.540225868 | 6.820373093 | BCAR3;NRP1;SEMA | 38 |
| positive regulation of pı | Aug-34 | 0.012749418 | 0.274112477 | 2.930781396 | 12.78485885 | EGR1;JUN;TGFB2;TC | 8 |
| ventricular cardiac musc | Aug-34 | 0.012749418 | 0.274112477 | 2.930781396 | 12.78485885 | RYR2;TGFB2;NOTCI | 8 |
| positive regulation of tr | 16/92 | 0.012836213 | 0.274645348 | 2.008191469 | 8.746647751 | ACVR1;TGFB2;BMPI | 16 |
| positive regulation of eı | 14/77 | 0.013221992 | 0.276261875 | 2.119043422 | 9.166714444 | NRP1;BMPR2;EGR3;I | 14 |
| defense response to syr | 20/124 | 0.013343067 | 0.276261875 | 1.835441747 | 7.923158449 | IFITM3;RSAD2;DDX | 20 |
| positive regulation of bi | Sep-41 | 0.013422952 | 0.276261875 | 2.679440269 | 11.55050217 | ACVR1;BMP4;P2RX7 | 9 |
| regulation of interleukir | Oct-48 | 0.01358575 | 0.276261875 | 2.507568393 | 10.77936902 | PRNP;HOMER2;IRF4 | 10 |
| positive regulation of pl | Jul-28 | 0.013897324 | 0.276261875 | 3.174218476 | 13.57314545 | FLT1;TGFB1;PDGFB; | 7 |
| necroptotic process (GC | May-16 | 0.013920265 | 0.276261875 | 4.326315789 | 18.49244561 | CYLD;TICAM2;ITPK | 5 |
| positive regulation of sı | May-16 | 0.013920265 | 0.276261875 | 4.326315789 | 18.49244561 | SLC45A3;PRKCE;AD | 5 |
| regulation of interleukir | May-16 | 0.013920265 | 0.276261875 | 4.326315789 | 18.49244561 | IRF4;TNFSF4;LEF1;R | 5 |
| positive regulation of hy | 23/149 | 0.013972152 | 0.276261875 | 1.742856299 | 7.443197389 | USP6NL;BCAR3;MT | 23 |
| nervous system develop | 57/447 | 0.014136614 | 0.278271396 | 1.400245588 | 5.963627891 | CHRM3;DAGLA;PLX | 57 |
| defense response to vir | 21/133 | 0.014379354 | 0.281797167 | 1.789709395 | 7.591878954 | IFITM3;RSAD2;DDX | 21 |

| | | | | | | | |
|---|---|---|---|---|---|---|---|
| cellular response to top | Aug-32 | 0.002406396 | 0.107485709 | 4.008480104 | 24.16963143 | ATXN3;DAXX;HSPA | 8 |
| response to lipopolysac | 23/159 | 0.002515625 | 0.111129803 | 2.041332442 | 12.21785241 | CD274;TICAM2;TNF | 23 |
| myeloid leukocyte diffe | 13/71 | 0.002756124 | 0.120430639 | 2.699189836 | 15.90883569 | FARP2;CSF1R;CEBP | 13 |
| regulation of viral life c | Jul-26 | 0.002837202 | 0.122640329 | 4.428738077 | 25.97427003 | TRIM5;LAMP3;TRIM | 7 |
| response to chemokine | May-14 | 0.002993934 | 0.127155199 | 6.673177083 | 38.77894638 | FOXC1;DUSP1;WNK | 5 |
| cellular response to mol | 18/115 | 0.003004911 | 0.127155199 | 2.237282629 | 12.99303495 | CD274;TICAM2;TNF | 18 |
| positive regulation of nı | 57/511 | 0.003062357 | 0.12823622 | 1.523274695 | 8.817582566 | CIITA;CITED2;SIX1; | 57 |
| positive regulation of ir | 15/89 | 0.003150709 | 0.129396634 | 2.44212922 | 14.06697621 | PTGER4;ABCC1;FEM | 15 |
| positive regulation of d | 16/98 | 0.00325101 | 0.129396634 | 2.351315474 | 13.47019157 | PTGER4;ABCC1;FEM | 16 |
| regulation of interleukir | 14/81 | 0.003363101 | 0.132545745 | 2.516767831 | 14.33272053 | DDX58;NOD1;SSC5I | 14 |
| positive regulation of st | 16/99 | 0.003606889 | 0.139923739 | 2.322859964 | 13.06587738 | AXIN1;TRAF2;TRAF | 16 |
| positive regulation of ir | Aug-34 | 0.003619918 | 0.139923739 | 3.699734056 | 20.79732988 | CD40;RIPK2;CCR7;II | 8 |
| regulation of neuron pr | May-15 | 0.004208649 | 0.149723606 | 6.005533854 | 32.85395562 | MACF1;FZD4;SPRY | 5 |
| regulation of interleukir | 17/110 | 0.004414202 | 0.1556587 | 2.202904072 | 11.94619066 | DDX58;TNFAIP3;CY | 17 |
| cellular response to inte | Jul-28 | 0.004459747 | 0.155897253 | 4.006518905 | 21.68593735 | SOCS3;SBNO2;SRC; | 7 |
| negative regulation of ty | Sep-43 | 0.00480819 | 0.16520448 | 3.183554753 | 16.99201512 | IFIH1;CYLD;TBK1;D | 9 |
| regulation of cytokine p | Sep-43 | 0.00480819 | 0.16520448 | 3.183554753 | 16.99201512 | ZC3H12A;NOD2;MEI | 9 |
| aortic valve developmer | Aug-36 | 0.005259558 | 0.176663131 | 3.435094586 | 18.02637408 | DLL4;BMP2;BMPR2; | 8 |
| positive regulation of a: | Jun-22 | 0.005273526 | 0.176663131 | 4.505618893 | 23.63222346 | MACF1;DBNL;RUFY | 6 |
| peptidyl-serine modifica | 23/169 | 0.005433632 | 0.179575166 | 1.900477377 | 9.911269862 | SMG1;CAMK2D;GA | 23 |
| negative regulation of ir | Jul-29 | 0.005494464 | 0.179575166 | 3.824196989 | 19.90117556 | CD274;ZC3H12A;LIL | 7 |
| negative regulation of n | Jul-29 | 0.005494464 | 0.179575166 | 3.824196989 | 19.90117556 | CDK6;MAFB;INHBA | 7 |
| protein K63-linked ubi | Sep-44 | 0.005639883 | 0.18284136 | 3.092428198 | 16.01225924 | TRAF3IP2;TRIM5;TR | 9 |
| necroptotic process (GC | May-16 | 0.00573777 | 0.184526693 | 5.459280303 | 28.17362374 | CYLD;TICAM2;RIPK | 5 |
| activation of GTPase ac | 16/105 | 0.006474737 | 0.206574937 | 2.165555351 | 10.91406832 | USP6NL;RABGAP1L | 16 |
| regulation of macropha | Jul-30 | 0.006698896 | 0.21204381 | 3.657729154 | 18.3099062 | LRFN5;TNIP2;ZC3H1 | 7 |
| regulation of MAP kina | 15/97 | 0.007181961 | 0.221149918 | 2.202913723 | 10.87398471 | LYN;CD40;DUSP1;PI | 15 |
| regulation of establishn | May-17 | 0.007619894 | 0.221149918 | 5.00406901 | 24.40480866 | IKBKB;S1PR3;S1PR2 | 5 |
| positive regulation of G | 27/214 | 0.007714589 | 0.221165541 | 1.742538447 | 8.47682594 | USP6NL;RTN4R;BCA | 27 |
| negative regulation of n | 25/194 | 0.00777571 | 0.221165541 | 1.784710621 | 8.667894137 | DNMT1;CITED2;PDC | 25 |
| positive regulation of pı | 42/371 | 0.007812315 | 0.221165541 | 1.544008062 | 7.491610382 | BCAR3;CSF1R;CD40 | 42 |
| dendritic cell differentia | Jun-24 | 0.008332782 | 0.234250221 | 4.004560261 | 19.17206437 | LYN;TRPM2;UBD;LI | 6 |
| peptidyl-serine phosph | 21/156 | 0.008650941 | 0.24147275 | 1.875263158 | 8.907663435 | SMG1;CAMK2D;MA | 21 |
| regulation of viral entry | Aug-39 | 0.008709838 | 0.24147275 | 3.102161059 | 14.71448703 | CIITA;TRIM5;TRIM2 | 8 |
| regulation of inflammat | 26/206 | 0.008774081 | 0.241587698 | 1.742772277 | 8.253688102 | PTGER4;TRADD;TNI | 26 |
| regulation of cilium ass | Nov-64 | 0.009193786 | 0.2514219 | 2.496806018 | 11.70809137 | USP6NL;CYLD;ODF | 11 |
| positive regulation of JI | Dec-73 | 0.009530798 | 0.254001788 | 2.367088743 | 11.0146008 | DUSP22;RASSF2;TR | 12 |
| negative regulation of c | Jul-32 | 0.009679863 | 0.254001788 | 3.364745763 | 15.60470671 | SEMA4A;SEMA6A;S | 7 |
| positive regulation of cy | 17/119 | 0.009717734 | 0.254001788 | 2.007545932 | 9.302572058 | TNFSF15;TRADD;RP | 17 |
| positive regulation of ca | May-18 | 0.009891951 | 0.254001788 | 4.618890224 | 21.32095382 | BMP2;BMPR2;BMP1 | 5 |
| positive regulation of st | May-18 | 0.009891951 | 0.254001788 | 4.618890224 | 21.32095382 | LYN;RNF13;TAOK2; | 5 |
| pulmonary valve develc | May-18 | 0.009891951 | 0.254001788 | 4.618890224 | 21.32095382 | BMPR2;JAG1;SLIT2; | 5 |
| regulation of cartilage d | May-18 | 0.009891951 | 0.254001788 | 4.618890224 | 21.32095382 | BMP2;BMPR2;BMP1 | 5 |
| regulation of presynaps | May-18 | 0.009891951 | 0.254001788 | 4.618890224 | 21.32095382 | LRFN5;LRFN4;SLITR | 5 |
| regulation of presynaps | May-18 | 0.009891951 | 0.254001788 | 4.618890224 | 21.32095382 | LRFN5;LRFN4;SLITR | 5 |
| skeletal system develop | 21/158 | 0.00995256 | 0.254001788 | 1.847685363 | 8.517691732 | TEAD4;PTGER4;SLC | 21 |
| negative regulation of iı | 25/198 | 0.009983155 | 0.254001788 | 1.743064346 | 8.030046655 | PHLPP1;SRC;TNFAI | 25 |
| regulation of transcripti | 198/2206 | 0.010955819 | 0.271866623 | 1.20786326 | 5.452155292 | MAML2;BACH1;IL4I | 198 |
| positive regulation of ce | Nov-66 | 0.011542383 | 0.283647217 | 2.405751634 | 10.73381305 | FOXA1;CITED2;CCD | 11 |
| cellular response to mec | Sep-49 | 0.011571678 | 0.283647217 | 2.705140339 | 12.06274763 | PTGER4;CD40;CASP | 9 |
| cellular response to inte | May-19 | 0.012588618 | 0.293036606 | 4.288736979 | 18.76306227 | STAT5B;CISH;IRS2;I | 5 |
| interleukin-7-mediated : | May-19 | 0.012588618 | 0.293036606 | 4.288736979 | 18.76306227 | STAT5B;CISH;IRS2;I | 5 |
| interleukin-1-mediated : | 14/94 | 0.012683674 | 0.293036606 | 2.106303209 | 9.199152096 | IL1RN;RIPK2;CUL1; | 14 |
| axon guidance (GO:00( | 25/203 | 0.01343079 | 0.308524424 | 1.693638641 | 7.299930547 | SEMA3C;SRC;IRS2;N | 25 |
| positive regulation of ap | Oct-59 | 0.013963465 | 0.311905929 | 2.454044975 | 10.48198925 | CYLD;TNFSF10;PMA | 10 |
| regulation of interferon | 13/86 | 0.014190458 | 0.311905929 | 2.142813598 | 9.118069406 | CD274;LILRB1;PDCL | 13 |
| protein phosphorylatior | 52/496 | 0.014486233 | 0.31475633 | 1.416967673 | 6.000229734 | SMG1;BMPR2;MAST | 52 |
| regulation of JNK casca | 15/105 | 0.014563353 | 0.31475633 | 2.006225426 | 8.484822869 | PHLPP1;AXIN1;TRA | 15 |
| cellular response to unf | Jun-27 | 0.015037675 | 0.319848959 | 3.431921824 | 14.40445048 | DAXX;HSPA1L;RHE | 6 |
| negative regulation of p | Jun-27 | 0.015037675 | 0.319848959 | 3.431921824 | 14.40445048 | SRGAP2C;ODF2L;LI | 6 |
| positive regulation of eı | Jun-27 | 0.015037675 | 0.319848959 | 3.431921824 | 14.40445048 | STAT5B;INHBA;HIF | 6 |
| negative regulation of g | 36/322 | 0.015584892 | 0.324523612 | 1.519940525 | 6.325161529 | PTGER4;BTG2;DNM | 36 |
| homophilic cell adhesio | Oct-60 | 0.015633536 | 0.324523612 | 2.404833442 | 10.00010771 | SDK1;TENM3;PALLI | 10 |
| adaptive immune respoı | May-20 | 0.015741817 | 0.324523612 | 4.002604167 | 16.61654945 | JAG1;CD70;TLR4;IL1 | 5 |
| modulation by symbior | May-20 | 0.015741817 | 0.324523612 | 4.002604167 | 16.61654945 | TRIM5;TRIM25;TRIM | 5 |
| negative regulation of a | May-20 | 0.015741817 | 0.324523612 | 4.002604167 | 16.61654945 | SEMA4A;SEMA6A;S | 5 |
| negative regulation of o | May-20 | 0.015741817 | 0.324523612 | 4.002604167 | 16.61654945 | MAFB;LILRB1;LILRI | 5 |
| ameboidal-type cell mig | Sep-52 | 0.016861031 | 0.345823194 | 2.515999757 | 10.2721984 | TBX1;SEMA4A;SEM | 9 |
| protein polyubiquitinati | 35/314 | 0.017620322 | 0.355061609 | 1.514372617 | 6.116100265 | TRAF3IP2;CUL1;RNI | 35 |
| cellular response to epi | Jun-28 | 0.017922569 | 0.355061609 | 3.275747705 | 13.17405653 | DUSP22;STAT5B;FO | 6 |
| regulation of extrinsic a | Jun-28 | 0.017922569 | 0.355061609 | 3.275747705 | 13.17405653 | GDNF;INHBA;TNF;H | 6 |
| regulation of apoptotic | 73/742 | 0.018228554 | 0.355061609 | 1.322350656 | 5.295704948 | FOXA1;TNFAIP8;CI | 73 |
| axonogenesis (GO:000 | 28/240 | 0.018309853 | 0.355061609 | 1.592849393 | 6.371900875 | SEMA3C;SRC;PLPPR | 28 |
| cellular response to lipi | 26/219 | 0.018326116 | 0.355061609 | 1.6242275 | 6.495981171 | CD274;ADCY3;TNFA | 26 |
| cellular response to hea | Jul-36 | 0.018361693 | 0.355061609 | 2.900013487 | 11.59277117 | ATXN3;DAXX;CLPE | 7 |
| positive regulation of N | Sep-53 | 0.01896398 | 0.355061609 | 2.458684429 | 9.749209698 | NFAT5;TRADD;TLR | 9 |
| regulation of NIK/NF-l | Dec-80 | 0.019077937 | 0.355061609 | 2.122609933 | 8.403885497 | NFAT5;CYLD;RASSI | 12 |
| beta-catenin destruction | May-21 | 0.019380308 | 0.359026908 | 3.752237956 | 14.79694201 | AMER1;FZD1;CSNK | 5 |
| regulation of cellular co | 18/139 | 0.020613791 | 0.380125872 | 1.79117987 | 6.952992986 | OGFR;CAMK2D;EGI | 18 |
| negative regulation of tı | Jun-29 | 0.021164689 | 0.387088628 | 3.133153944 | 12.07962781 | ZFP36;DTX3L;SRC;II | 6 |
| positive regulation of m | Jul-37 | 0.021183955 | 0.387088628 | 2.803194263 | 10.80494378 | STAT5B;INHBA;RAI | 7 |
| cell-cell adhesion via pl | 21/170 | 0.021412345 | 0.389491519 | 1.697765807 | 6.525851274 | TENM3;AMIGO3;PC | 21 |
| regulation of cytokine p | 19/150 | 0.021900218 | 0.396571508 | 1.746556861 | 6.674045611 | LYN;PTGER4;DDX5 | 19 |
| peptidyl-threonine deph | May-22 | 0.023529369 | 0.420949782 | 3.531326593 | 13.2407299 | PDP1;DUSP5;DUSP1 | 5 |
| regulation of viral relea | May-22 | 0.023529369 | 0.420949782 | 3.531326593 | 13.2407299 | TRIM5;TRIM25;TRIM | 5 |
| connective tissue develc | Jul-38 | 0.024293619 | 0.420949782 | 2.712621441 | 10.08428291 | ZNF516;ITGB8;ARID | 7 |
| negative regulation of iı | Jul-38 | 0.024293619 | 0.420949782 | 2.712621441 | 10.08428291 | OAS3;NLRC5;SLAM | 7 |
| regulation of axon exteı | Jul-38 | 0.024293619 | 0.420949782 | 2.712621441 | 10.08428291 | RTN4R;MACF1;DBN | 7 |
| regulation of osteoblast | Dec-83 | 0.024852578 | 0.42878696 | 2.032590573 | 7.510003063 | BMP2;BMPR2;CDK6 | 12 |
| cellular response to tur | 23/194 | 0.025701392 | 0.435170947 | 1.620414673 | 5.932678596 | EDARADD;CD40;TN | 23 |
| negative regulation of I | 18/143 | 0.026640852 | 0.435170947 | 1.733483913 | 6.284415641 | EOMES;HDAC4;SP1 | 18 |
| cellular response to org | 16/123 | 0.026898519 | 0.435170947 | 1.799491344 | 6.506392143 | NCOA1;CSF1R;STAT | 16 |

| | | | | | | | |
|---|---|---|---|---|---|---|---|
| protein localization to n | 47/195 | 0.025764344 | 0.999996202 | 1.419363025 | 5.193113983 | PLEKHF1;ROCK1;RA | 47 |
| intrinsic apoptotic signa | 27/102 | 0.026597331 | 0.999996202 | 1.607393573 | 5.829927147 | BRSK2;CDKN1A;PPI | 27 |
| chondroitin sulfate bios | Aug-21 | 0.026782396 | 0.999996202 | 2.74384195 | 9.93273659 | BCAN;VCAN;CHPF; | 8 |
| positive regulation of tr | Aug-21 | 0.026782396 | 0.999996202 | 2.74384195 | 9.93273659 | SMAD1;SMAD4;BMI | 8 |
| in utero embryonic dev | Sep-25 | 0.028158437 | 0.999996202 | 2.508267286 | 8.954284095 | FUT8;MBNL1;STIL;B | 9 |
| negative regulation of c | 22/80 | 0.028321623 | 0.999996202 | 1.693059943 | 6.034285239 | DLX1;SMAD1;SKOR | 22 |
| centromere complex ass | Dec-37 | 0.028321827 | 0.999996202 | 2.140962801 | 7.630653658 | CENPT;HELLS;CENP | 12 |
| negative chemotaxis (G | Nov-33 | 0.028715405 | 0.999996202 | 2.229969921 | 7.917110283 | SEMA4A;SEMA7A;E | 11 |
| regulation of intracellul | 96/437 | 0.029488529 | 0.999996202 | 1.260320577 | 4.441059612 | ARHGAP11A;ARHG | 96 |
| regulation of cell projec | Jun-14 | 0.029678069 | 0.999996202 | 3.343255052 | 11.75938783 | MACF1;SEMA4D;SP | 6 |
| protein localization to c | 35/140 | 0.029976046 | 0.999996202 | 1.488852188 | 5.221935686 | PLEKHF1;ROCK1;RA | 35 |
| regulation of BMP sign | 21/76 | 0.030107682 | 0.999996202 | 1.704100506 | 5.969421366 | CRB2;SMAD2;DLX1; | 21 |
| regulation of epithelial t | 21/76 | 0.030107682 | 0.999996202 | 1.704100506 | 5.969421366 | FOXA1;CRB2;SMAD | 21 |
| positive regulation of n | 20/72 | 0.031994712 | 0.999996202 | 1.716430499 | 5.908270716 | ROBO2;CX3CR1;CDI | 20 |
| regulation of fibroblast | 14/46 | 0.032238481 | 0.999996202 | 1.951628352 | 6.703051943 | CDKN1A;SPHK1;PD | 14 |
| phospholipase C-activa | 22/81 | 0.032350543 | 0.999996202 | 1.66426174 | 5.710289209 | CX3CR1;HOMER1;G | 22 |
| negative regulation of a | 17/59 | 0.032865297 | 0.999996202 | 1.805963141 | 6.167974481 | KDM4A;IL10RA;HTR | 17 |
| interleukin-6-mediated | Jul-18 | 0.033517985 | 0.999996202 | 2.836954632 | 9.633370574 | CEBPA;SOCS3;SMAI | 7 |
| negative regulation of c | Jul-18 | 0.033517985 | 0.999996202 | 2.836954632 | 9.633370574 | CAMK2D;CAB39;SL | 7 |
| negative regulation of le | Jul-18 | 0.033517985 | 0.999996202 | 2.836954632 | 9.633370574 | IL4I1;WNK1;FGL1;M | 7 |
| pulmonary valve develc | Jul-18 | 0.033517985 | 0.999996202 | 2.836954632 | 9.633370574 | NOTCH2;ROBO2;BM | 7 |
| positive regulation of ep | 13/42 | 0.033531915 | 0.999996202 | 1.999518845 | 6.788881579 | CRB2;SMAD2;FOXC | 13 |
| negative regulation of p | 23/86 | 0.034476002 | 0.999996202 | 1.629485923 | 5.487280452 | HDAC4;PRNP;CDKN | 23 |
| regulation of apoptotic | Dec-38 | 0.034686859 | 0.999996202 | 2.058491836 | 6.919402849 | CX3CR1;RB1;PYCAF | 12 |
| regulation of axon exteı | Dec-38 | 0.034686859 | 0.999996202 | 2.058491836 | 6.919402849 | RTN4R;MACF1;CDK | 12 |
| cellular response to ner | Aug-22 | 0.035494204 | 0.999996202 | 2.547697112 | 8.505196032 | EHD1;RAP1A;BDNF; | 8 |
| limb morphogenesis (G | Aug-22 | 0.035494204 | 0.999996202 | 2.547697112 | 8.505196032 | SKI;MBNL1;NIPBL;Z | 8 |
| osteoclast differentiatioı | Aug-22 | 0.035494204 | 0.999996202 | 2.547697112 | 8.505196032 | CSF1R;SBNO2;CSF1; | 8 |
| peptidyl-threonine deph | Aug-22 | 0.035494204 | 0.999996202 | 2.547697112 | 8.505196032 | DUSP4;PPM1A;DUSI | 8 |
| epithelial tube morphog | Nov-34 | 0.035617381 | 0.999996202 | 2.132883927 | 7.113000549 | ACVR1;BMP4;CSF1F | 11 |
| cell proliferation in fore | 5-Nov | 0.035914389 | 0.999996202 | 3.714168714 | 12.35561777 | NUMBL;PCM1;AKN | 5 |
| midbrain dopaminergic | 5-Nov | 0.035914389 | 0.999996202 | 3.714168714 | 12.35561777 | FZD1;NR4A2;CTNNE | 5 |
| positive regulation of m | 5-Nov | 0.035914389 | 0.999996202 | 3.714168714 | 12.35561777 | CASP8;CSF1;HSF1;L | 5 |
| positive regulation of ne | 5-Nov | 0.035914389 | 0.999996202 | 3.714168714 | 12.35561777 | CX3CR1;FZD3;CTNN | 5 |
| positive regulation of ne | 5-Nov | 0.035914389 | 0.999996202 | 3.714168714 | 12.35561777 | LBX2;WNT5B;CSNK | 5 |
| regulation of inclusion l | 5-Nov | 0.035914389 | 0.999996202 | 3.714168714 | 12.35561777 | DNAJA4;SACS;HSPA | 5 |
| negative regulation of iı | Oct-30 | 0.036199058 | 0.999996202 | 2.22963368 | 7.399534751 | CX3CR1;SERPINB1; | 10 |
| negative regulation of iı | Sep-26 | 0.036249103 | 0.999996202 | 2.360577464 | 7.830839566 | CX3CR1;SERPINB1; | 9 |
| regulation of interleukir | 15/51 | 0.036541382 | 0.999996202 | 1.858746236 | 6.151167356 | CD40;IL16;MEFV;MA | 15 |
| regulation of hydrolase | 20/73 | 0.036708542 | 0.999996202 | 1.683941576 | 5.564998829 | VAV3;RALBP1;RAB | 20 |
| regulation of I-kappaB | 52/224 | 0.038093632 | 0.999996202 | 1.35110105 | 4.415003903 | MAPKBP1;CD40;GPI | 52 |
| positive regulation of pa | 14/47 | 0.038406636 | 0.999996202 | 1.892371996 | 6.168233864 | ACVR1;GDF11;SMA | 14 |
| negative regulation of a | 21/78 | 0.039270131 | 0.999996202 | 1.644105465 | 5.322447927 | RB1;ACVR1;CX3CR1 | 21 |
| regulation of cell-matrix | 18/65 | 0.041618023 | 0.999996202 | 1.70871466 | 5.432383158 | MACF1;JAG1;CSF1;I | 18 |
| mitotic intra-S DNA da | Jun-15 | 0.041966276 | 0.999996202 | 2.971600218 | 9.422614281 | MDC1;EME1;EME2;F | 6 |
| myoblast differentiatior | Jun-15 | 0.041966276 | 0.999996202 | 2.971600218 | 9.422614281 | RB1;MBNL1;JAG1;SI | 6 |
| negative regulation of c | 43/182 | 0.042792238 | 0.999996202 | 1.381888365 | 4.354880977 | PTGER4;MAPKBP1; | 43 |
| endocytic recycling (GC | 15/52 | 0.042931084 | 0.999996202 | 1.808398872 | 5.693127445 | DENND1A;LMTK2;V | 15 |
| negative regulation of E | 15/52 | 0.042931084 | 0.999996202 | 1.808398872 | 5.693127445 | DLX1;SMAD1;SKOR | 15 |
| positive regulation of d | 15/52 | 0.042931084 | 0.999996202 | 1.808398872 | 5.693127445 | MACF1;SEMA7A;DE | 15 |
| glycosaminoglycan bios | 25/97 | 0.042995998 | 0.999996202 | 1.549775826 | 4.876599365 | HS3ST3A1;CHPF;SD | 25 |
| semaphorin-plexin sign | Nov-35 | 0.043604269 | 0.999996202 | 2.043888433 | 6.402685335 | SEMA4A;SEMA7A;S | 11 |
| positive regulation of ul | 21/79 | 0.044551101 | 0.999996202 | 1.615659541 | 5.02650813 | CEBPA;FBXW7;CSN | 21 |
| N-glycan processing (C | Jul-19 | 0.044895521 | 0.999996202 | 2.600382409 | 8.070071635 | FUT8;GNPTAB;MAN | 7 |
| neural crest cell differer | Jul-19 | 0.044895521 | 0.999996202 | 2.600382409 | 8.070071635 | MEF2C;FAM172A;PI | 7 |
| regulation of collagen b | Jul-19 | 0.044895521 | 0.999996202 | 2.600382409 | 8.070071635 | BMP4;HDAC2;TGFB | 7 |
| negative regulation of a | Oct-31 | 0.044927247 | 0.999996202 | 2.123330469 | 6.588080434 | RTN4R;CDKL3;SEM | 10 |
| cytosolic transport (GO | 29/116 | 0.044934231 | 0.999996202 | 1.488046167 | 4.61674567 | TMEM87A;GOSR2;G | 29 |
| positive regulation of ap | 69/310 | 0.044946827 | 0.999996202 | 1.280068576 | 3.971124892 | FOXA1;FAF1;TRADI | 69 |
| phosphorylation (GO:0 | 87/400 | 0.045355387 | 0.999996202 | 1.243386064 | 3.846074501 | DGKG;MAML1;HTR | 87 |
| DNA methylation (GO | Aug-23 | 0.045930705 | 0.999996202 | 2.377704918 | 7.324808737 | N6AMT1;HELLS;DNI | 8 |
| regulation of mononucl | Aug-23 | 0.045930705 | 0.999996202 | 2.377704918 | 7.324808737 | CSF1R;STK10;CSF1; | 8 |
| regulation of SMAD pı | Aug-23 | 0.045930705 | 0.999996202 | 2.377704918 | 7.324808737 | BMP4;SMAD4;RBPM | 8 |
| chromatin remodeling ( | 26/103 | 0.049843139 | 0.999996202 | 1.507056919 | 4.51947444 | FOXA1;RB1;KDM5A | 26 |

| | | | | | | | |
|---|---|---|---|---|---|---|---|
| female gonad developm | Jun-22 | 0.014516123 | 0.281982047 | 3.570102686 | 15.11044301 | CEBPB;INHBB;INHB | 6 |
| negative regulation of ir | Jun-22 | 0.014516123 | 0.281982047 | 3.570102686 | 15.11044301 | PRNP;HOMER2;TBX | 6 |
| external encapsulating s | 31/217 | 0.01463225 | 0.282675204 | 1.592760583 | 6.728660502 | ITGAM;PAPLN;ITGB | 31 |
| negative regulation of a | 14/78 | 0.014743277 | 0.282675204 | 2.085817689 | 8.795826703 | RB1;ACVR1;SRC;BD | 14 |
| negative regulation of p | 14/78 | 0.014743277 | 0.282675204 | 2.085817689 | 8.795826703 | RB1;DUSP1;HEG1;SF | 14 |
| negative regulation of R | Aug-35 | 0.015181885 | 0.285476227 | 2.822077745 | 11.81788052 | KANK1;TGFB2;ITGA | 8 |
| semaphorin-plexin sign | Aug-35 | 0.015181885 | 0.285476227 | 2.822077745 | 11.81788052 | NRP1;SEMA7A;PLXN | 8 |
| regulation of interferon- | 15/86 | 0.015270045 | 0.285476227 | 2.014755198 | 8.425428689 | CD274;PRNP;SLC11A | 15 |
| positive regulation of ep | 16/94 | 0.015616297 | 0.285476227 | 1.956482198 | 8.137870787 | DOCK5;NRP1;TGFB2 | 16 |
| response to lipopolysac | 24/159 | 0.015854086 | 0.287777652 | 1.697442259 | 7.03475751 | CD274;TICAM2;SLC1 | 24 |
| negative regulation of D | 22/143 | 0.016545915 | 0.297893731 | 1.735528412 | 7.118471156 | RB1;PRNP;SP100;PA | 22 |
| receptor-mediated endo | 22/143 | 0.016545915 | 0.297893731 | 1.735528412 | 7.118471156 | NLGN3;CD163;ITGAI | 22 |
| intrinsic apoptotic signa | Jul-29 | 0.016849914 | 0.301968523 | 3.029768177 | 12.37178478 | PPP1R15A;ERN1;BRS | 7 |
| carboxylic acid transpo | Nov-57 | 0.017360591 | 0.301968523 | 2.278809513 | 9.237274107 | SLC23A2;SLC7A6;SL | 11 |
| regulation of bone mine | Nov-57 | 0.017360591 | 0.301968523 | 2.278809513 | 9.237274107 | ACVR1;BMP4;P2RX7 | 11 |
| actin filament organizati | 23/152 | 0.017439999 | 0.301968523 | 1.702040547 | 6.891543317 | GAS2L3;SHROOM2;I | 23 |
| negative regulation of p | 23/152 | 0.017439999 | 0.301968523 | 1.702040547 | 6.891543317 | RB1;PRNP;TGFB1;IG | 23 |
| cyclic-nucleotide-media | Aug-36 | 0.017933565 | 0.301968523 | 2.72113864 | 10.94191934 | RIMS2;HTR2B;PDE3A | 8 |
| positive regulation of bo | Aug-36 | 0.017933565 | 0.301968523 | 2.72113864 | 10.94191934 | ACVR1;BMP4;P2RX7 | 8 |
| chloride transmembrane | Oct-50 | 0.017943974 | 0.301968523 | 2.381926121 | 9.576536204 | GABRR2;GABRA4;A | 10 |
| neuron migration (GO:0 | Oct-50 | 0.017943974 | 0.301968523 | 2.381926121 | 9.576536204 | NR4A2;NRP1;RAPGE | 10 |
| hexose transmembrane | Jun-23 | 0.018092945 | 0.301968523 | 3.359910789 | 13.48074556 | SLC2A14;SLC45A3;S | 6 |
| positive regulation of m | Jun-23 | 0.018092945 | 0.301968523 | 3.359910789 | 13.48074556 | BMP4;PAX8;LIF;ITG | 6 |
| regulation of interferon- | Jun-23 | 0.018092945 | 0.301968523 | 3.359910789 | 13.48074556 | SOCS3;IFNGR1;IFNG | 6 |
| regulation of toll-like re | Jun-23 | 0.018092945 | 0.301968523 | 3.359910789 | 13.48074556 | TICAM2;NINJ1;NR1H | 6 |
| negative regulation of c | 14/80 | 0.018185513 | 0.301968523 | 2.022386744 | 8.103966584 | RB1;BTG2;BTG1;PPP | 14 |
| positive regulation of pl | Sep-43 | 0.01820013 | 0.301968523 | 2.52154691 | 10.10214034 | FLT1;BDNF;GNAQ;C | 9 |
| positive regulation of T | Sep-43 | 0.01820013 | 0.301968523 | 2.52154691 | 10.10214034 | IL4I1;DUSP10;BCL6; | 9 |
| mesenchyme morphoge | May-17 | 0.018204018 | 0.301968523 | 3.965570175 | 15.886522 | ACVR1;TGFB2;BMP2 | 5 |
| negative regulation of p | May-17 | 0.018204018 | 0.301968523 | 3.965570175 | 15.886522 | SPRED3;SPRED2;SPF | 5 |
| pharyngeal system deve | May-17 | 0.018204018 | 0.301968523 | 3.965570175 | 15.886522 | BMP4;TGFB2;BMPR2 | 5 |
| positive regulation of sr | May-17 | 0.018204018 | 0.301968523 | 3.965570175 | 15.886522 | DOCK5;NRP1;NR4A1 | 5 |
| pulmonary valve morph | May-17 | 0.018204018 | 0.301968523 | 3.965570175 | 15.886522 | BMP4;TGFB2;JAG1;N | 5 |
| inorganic anion transme | Dec-65 | 0.018625089 | 0.306827728 | 2.157940376 | 8.595606835 | GABRR2;SLC37A2;S | 12 |
| positive regulation of in | 67/546 | 0.018635507 | 0.306827728 | 1.340605769 | 5.339212577 | BCAR3;SLC35B2;FLT | 67 |
| dephosphorylation (GC | 23/153 | 0.018735297 | 0.307328253 | 1.68885392 | 6.717156414 | DUSP4;DUSP5;PPM1 | 23 |
| regulation of cytoskelet | 18/112 | 0.018864627 | 0.308307869 | 1.826709062 | 7.25288751 | PHLDB1;TESK1;HIP1 | 18 |
| cell-cell adhesion via pl | 25/170 | 0.018993159 | 0.30926729 | 1.646184886 | 6.524944178 | CD84;ITGAM;PCDHG | 25 |
| regulation of actin filam | 13/73 | 0.019522443 | 0.316721238 | 2.065318887 | 8.129488739 | KANK1;DBNL;PRKC | 13 |
| positive regulation of m | Nov-58 | 0.019636044 | 0.317401609 | 2.230200634 | 8.765554699 | MLLT11;MOAP1;BCI | 11 |
| positive regulation of lip | Jul-30 | 0.020218938 | 0.325635189 | 2.897878774 | 11.305018 | FLT1;TGFB1;PDGFB; | 7 |
| positive regulation of fa | Oct-51 | 0.020472921 | 0.326203599 | 2.323701654 | 9.036067512 | CEBPB;BMP2;ID2;ZC | 10 |
| regulation of substrate a | Oct-51 | 0.020472921 | 0.326203599 | 2.323701654 | 9.036067512 | DOCK5;NRP1;KANK | 10 |
| negative regulation of n | 30/214 | 0.020475186 | 0.326203599 | 1.557478261 | 6.056318924 | RB1;NOTCH1;BMPR | 30 |
| ERBB signaling pathwa | Dec-66 | 0.020853267 | 0.329625714 | 2.117861126 | 8.196640736 | ADAM17;GAREM1;T | 12 |
| positive regulation of bi | Sep-44 | 0.02100378 | 0.329625714 | 2.449367089 | 9.462034532 | BMP4;TGFB1;F2R;SC | 9 |
| phagocytosis, engulfme | Aug-37 | 0.021024866 | 0.329625714 | 2.627160853 | 10.14622515 | ITGAM;ITGB2;CD36; | 8 |
| negative regulation of in | 28/198 | 0.022095026 | 0.329625714 | 1.572910464 | 5.996568228 | SRC;NPRL3;WWC2;P | 28 |
| positive regulation of ne | Jun-24 | 0.022228875 | 0.329625714 | 3.173073547 | 12.07787028 | BACE1;PRNP;TGFB2 | 6 |
| cellular response to tran | 18/114 | 0.022256243 | 0.329625714 | 1.788453895 | 6.805304482 | ACVR1;JUN;TGFB2;I | 18 |
| regulation of peptidyl-s | 16/98 | 0.022550596 | 0.329625714 | 1.860630867 | 7.055500647 | BCAR3;TGFB1;BDNF | 16 |
| transmembrane recepto | 51/404 | 0.022980724 | 0.332506827 | 1.382573778 | 5.216588451 | BCAR3;NRP1;FLT1;C | 51 |
| regulation of cell differe | 23/156 | 0.023076255 | 0.332506827 | 1.650483808 | 6.220592771 | PHLDB1;CASZ1;ZFH | 23 |
| regulation of intrinsic ap | Oct-52 | 0.023252079 | 0.332506827 | 2.26824978 | 8.531705691 | CYLD;BCL2L11;SRC; | 10 |
| negative regulation of R | May-18 | 0.023273226 | 0.332506827 | 3.660323887 | 13.76447116 | KANK1;ITGA3;HEG1 | 5 |
| positive chemotaxis (GO | May-18 | 0.023273226 | 0.332506827 | 3.660323887 | 13.76447116 | NRP1;ANGPT1;MET; | 5 |
| positive regulation of ca | May-18 | 0.023273226 | 0.332506827 | 3.660323887 | 13.76447116 | BMP4;BMP2;BMPR2; | 5 |
| programmed necrotic ce | May-18 | 0.023273226 | 0.332506827 | 3.660323887 | 13.76447116 | CYLD;TICAM2;ITPK | 5 |
| positive regulation of ep | 19/123 | 0.023718754 | 0.337782514 | 1.742745126 | 6.520462125 | YAP1;NRP1;EGR3;BM | 19 |
| integrin-mediated signa | 13/75 | 0.024020276 | 0.337849993 | 1.99847405 | 7.452023884 | NRP1;SEMA7A;SRC; | 13 |
| positive regulation of T | 13/75 | 0.024020276 | 0.337849993 | 1.99847405 | 7.452023884 | CD274;DOCK8;ZBTB | 13 |
| positive regulation of R | Jul-31 | 0.024029969 | 0.337849993 | 2.776980155 | 10.35384142 | LPAR6;GPR65;F2R;LI | 7 |
| positive regulation of N | Sep-45 | 0.024104899 | 0.337849993 | 2.381197257 | 8.870769818 | YAP1;TGFB2;NOTCH | 9 |
| regulation of receptor si | Sep-45 | 0.024104899 | 0.337849993 | 2.381197257 | 8.870769818 | SOCS3;NOTCH1;ERE | 9 |
| enzyme linked receptor | 21/140 | 0.024411623 | 0.340882241 | 1.683776695 | 6.251350866 | TRIO;BMPR2;FLT1;C | 21 |
| response to cAMP (GC | Aug-38 | 0.024475176 | 0.340882241 | 2.539448252 | 9.421596592 | KCNE1;RAP1A;AQP9 | 8 |
| negative regulation of c | 48/379 | 0.025383491 | 0.35242471 | 1.38720966 | 5.096131462 | BTG2;BTG1;NOTCH1 | 48 |
| cytokine-mediated signa | 74/621 | 0.02581405 | 0.357282587 | 1.296533298 | 4.741210103 | IFITM3;SLC27A1;ITG | 74 |
| skeletal system develop | 23/158 | 0.026375921 | 0.363921976 | 1.625851143 | 5.910462787 | TGFB2;CHD7;EIF2AK | 23 |
| cAMP-mediated signali | Jun-25 | 0.026956563 | 0.368823834 | 3.005903384 | 10.86191748 | RIMS2;PDE3A;RAPG | 6 |
| in utero embryonic deve | Jun-25 | 0.026956563 | 0.368823834 | 3.005903384 | 10.86191748 | FUT8;BMP2;CHD7;A | 6 |
| negative regulation of c | 68/566 | 0.027033226 | 0.368823834 | 1.308003926 | 4.722794859 | RB1;BTG2;BTG1;BMI | 68 |
| cellular response to cyto | 59/482 | 0.027064291 | 0.368823834 | 1.33525942 | 4.819672398 | ITGAM;CSF1;DOCK8 | 59 |
| response to cytokine (G | 22/150 | 0.027237515 | 0.37004587 | 1.639977762 | 5.909102321 | IFITM3;CD274;ITIH4; | 22 |
| regulation of fibroblast | Sep-46 | 0.027518175 | 0.370196591 | 2.316712282 | 8.323735466 | PARP10;FTH1;PDGFC | 9 |
| positive regulation of m | 44/345 | 0.02815337 | 0.370196591 | 1.397698162 | 4.989905731 | SLC27A1;CEBPB;PA | 44 |
| negative regulation of s | Aug-39 | 0.028302666 | 0.370196591 | 2.457394528 | 8.760118245 | PNKD;NOTCH1;RSA | 8 |
| regulation of cellular co | Aug-39 | 0.028302666 | 0.370196591 | 2.457394528 | 8.760118245 | PARP10;NOTCH1;RA | 8 |
| aortic valve morphogen | Jul-32 | 0.028306486 | 0.370196591 | 2.665753425 | 9.502516067 | RB1;BMP4;NOTCH1; | 7 |
| endodermal cell differen | Jul-32 | 0.028306486 | 0.370196591 | 2.665753425 | 9.502516067 | MMP14;ITGB2;COL6 | 7 |
| regulation of chemotaxi | Jul-32 | 0.028306486 | 0.370196591 | 2.665753425 | 9.502516067 | TGFB1;GPR183;PDGI | 7 |
| regulation of intracellula | Jul-32 | 0.028306486 | 0.370196591 | 2.665753425 | 9.502516067 | BMP4;PLK3;PCM1;TC | 7 |
| positive regulation of M | Dec-69 | 0.028722521 | 0.370196591 | 2.006061112 | 7.121664881 | FLT1;TGFB1;PDGFC; | 12 |
| activin receptor signalin | May-19 | 0.029171057 | 0.370196591 | 3.398684211 | 12.01291535 | ACVR1;INHBB;INHE | 5 |
| establishment of melano | May-19 | 0.029171057 | 0.370196591 | 3.398684211 | 12.01291535 | BLOC1S6;MREG;SHF | 5 |
| striated muscle cell diffe | May-19 | 0.029171057 | 0.370196591 | 3.398684211 | 12.01291535 | KDM6B;BMP2;SDC1 | 5 |
| regulation of epithelial c | 15/93 | 0.029371876 | 0.371680113 | 1.833231583 | 6.467123422 | YAP1;TGFB2;TGFB1 | 15 |
| negative regulation of v | Oct-54 | 0.029609306 | 0.373617144 | 2.164907652 | 7.619753082 | IFITM3;IFIH1;PARP1 | 10 |
| regulation of protein bin | 18/118 | 0.030409899 | 0.382629096 | 1.716534181 | 5.995831762 | DTX3L;PDGFB;HIP1I | 18 |
| regulation of interleukin | 17/110 | 0.031054149 | 0.389628408 | 1.742949244 | 6.051559586 | DDX58;C5AR2;F2R;T | 17 |
| positive regulation of ne | Sep-47 | 0.031257433 | 0.389969493 | 2.255621252 | 7.816851121 | BACE1;EGR1;PRNP;T | 9 |
| regulation of protein loc | Sep-47 | 0.031257433 | 0.389969493 | 2.255621252 | 7.816851121 | YAP1;BMP4;TGFB1;I | 9 |
| regulation of peptidyl-th | Jul-39 | 0.027702576 | 0.441270204 | 2.62770942 | 9.423570041 | RPTOR;RIPK2;AXIN | 7 |
| regulation of protein tyr | Jul-39 | 0.027702576 | 0.441270204 | 2.62770942 | 9.423570041 | CSF1R;SH3BP5L;DO | 7 |
| negative regulation of e | May-23 | 0.028210558 | 0.441270204 | 3.334960938 | 11.89933732 | GDNF;TNF;HSPA1B; | 5 |
| negative regulation of s | May-23 | 0.028210558 | 0.441270204 | 3.334960938 | 11.89933732 | GDNF;TNF;HSPA1B; | 5 |
| regulation of B cell rece | May-23 | 0.028210558 | 0.441270204 | 3.334960938 | 11.89933732 | LYN;ELF2;PAX5;FO | 5 |
| regulation of interferon- | May-23 | 0.028210558 | 0.441270204 | 3.334960938 | 11.89933732 | SOCS3;NLRC5;PARP | 5 |
| regulation of mononucl | May-23 | 0.028210558 | 0.441270204 | 3.334960938 | 11.89933732 | LYN;CSF1R;SLIT2;TI | 5 |
| regulation of toll-like re | May-23 | 0.028210558 | 0.441270204 | 3.334960938 | 11.89933732 | LYN;TICAM2;NINJ1; | 5 |
| positive regulation of ki | 15/114 | 0.028721759 | 0.445096145 | 1.822947695 | 6.471647142 | CSF1R;VAC14;DAX | 15 |
| regulation of tumor nec | 16/124 | 0.028757378 | 0.445096145 | 1.78273224 | 6.326668792 | DDX58;TNFAIP3;CY | 16 |
| negative regulation of a | Jun-31 | 0.028787313 | 0.445096145 | 2.882188925 | 10.22548899 | RTN4R;SEMA4A;SEI | 6 |
| negative regulation of M | Aug-48 | 0.029012735 | 0.445157235 | 2.403000652 | 8.506671328 | LYN;DNAJA1;DUSP | 8 |
| regulation of interleukin | Aug-48 | 0.029012735 | 0.445157235 | 2.403000652 | 8.506671328 | ZFP36;HOMER2;IRF4 | 8 |
| positive regulation of si | 28/252 | 0.03222691 | 0.488876147 | 1.506526768 | 5.174849327 | MACF1;MAML2;CIT | 28 |
| RNA modification (GC | Sep-58 | 0.032385678 | 0.489437687 | 2.207198806 | 7.570777981 | PUS3;ANKRD16;AP | 9 |
| regulation of cell cycle ( | 32/296 | 0.032704145 | 0.492399493 | 1.461533827 | 4.998816097 | CITED2;ADARB1;PK | 32 |
| small GTPase mediated | 14/106 | 0.03285551 | 0.492832643 | 1.830372142 | 6.251884673 | DNMT1;RALA;RRAI | 14 |
| regulation of intracellul | Jun-32 | 0.03319673 | 0.49609983 | 2.771185167 | 9.436727669 | PLK3;ICE1;IFI27;RU | 6 |
| regulation of axon exter | May-24 | 0.033441559 | 0.497907655 | 3.159265351 | 10.73504426 | SEMA4A;SEMA6A;S | 5 |
| T cell receptor signaling | 19/158 | 0.035168169 | 0.506537354 | 1.645317123 | 5.507886466 | PDPK1;RIPK2;RFTN | 19 |
| positive regulation of ce | 15/117 | 0.035173702 | 0.506537354 | 1.769042479 | 5.921792886 | FZD1;LYN;TENM3;F | 15 |
| JNK cascade (GO:000 | Jul-41 | 0.035460943 | 0.506537354 | 2.472869852 | 8.257712113 | PTGER4;RIPK2;GPS1 | 7 |
| protein autophosphoryl | 19/159 | 0.037171273 | 0.506537354 | 1.63347569 | 5.377759758 | LYN;CSF1R;SMG1;C | 19 |
| phosphorylation (GO:0 | 41/400 | 0.037422736 | 0.508241215 | 1.378087279 | 4.527673838 | CAMK2D;BMPR2;PN | 41 |
| regulation of peptidyl-s | 13/98 | 0.037621808 | 0.509224477 | 1.839097629 | 6.032555419 | BCAR3;RIPK2;LIF;A | 13 |
| antigen processing and | Jun-33 | 0.038020971 | 0.512900346 | 2.668403909 | 8.724659864 | SEC24A;ERAP2;TAP | 6 |
| negative regulation of c | May-25 | 0.039236094 | 0.517924705 | 3.001139323 | 9.718163863 | IKBKB;DUSP22;SRC | 5 |
| outflow tract septum m | May-25 | 0.039236094 | 0.517924705 | 3.001139323 | 9.718163863 | TBX1;BMPR2;SEMA | 5 |
| positive regulation of p | May-25 | 0.039236094 | 0.517924705 | 3.001139323 | 9.718163863 | RPTOR;RIPK2;AXIN | 5 |
| regulation of heterotypi | May-25 | 0.039236094 | 0.517924705 | 3.001139323 | 9.718163863 | IL1RN;WNK1;MYAD | 5 |
| response to organic cyc | Sep-60 | 0.03924312 | 0.517924705 | 2.120411611 | 6.86584858 | FOXA1;TRPM2;STA | 9 |
| positive regulation of pr | Nov-79 | 0.039295282 | 0.517924705 | 1.944454056 | 6.293518793 | RNF144B;PLK3;FZR1 | 11 |
| positive regulation of ul | Nov-79 | 0.039295282 | 0.517924705 | 1.944454056 | 6.293518793 | RNF144B;CSNK1A1; | 11 |
| heterophilic cell-cell adl | Jul-42 | 0.039828927 | 0.521538394 | 2.40208605 | 7.742312041 | TENM3;AMIGO3;AM | 7 |
| regulation of chemokine | Jul-42 | 0.039828927 | 0.521538394 | 2.40208605 | 7.742312041 | CSF1R;TLR7;HIF1A; | 7 |
| positive regulation of fa | Aug-51 | 0.040015557 | 0.522280965 | 2.234985361 | 7.1932713 | ZFP36;CEBPB;BMP2 | 8 |
| cellular response to org | 18/150 | 0.040245268 | 0.52357921 | 1.640929983 | 5.271918899 | PDGFB;ADCY3;NR3 | 18 |
| negative regulation of p | 39/381 | 0.042238342 | 0.547735925 | 1.375481814 | 4.352611627 | TNFAIP8;CITED2;SR | 39 |
| negative regulation of c | Nov-80 | 0.04251834 | 0.548050703 | 1.916169366 | 6.050917503 | RUBCN;RTEL1;SRC; | 11 |
| MAPK cascade (GO:0 | 32/303 | 0.043016224 | 0.548050703 | 1.423234272 | 4.477748259 | PTGER4;PDGFB;CUI | 32 |
| positive regulation of au | Dec-90 | 0.043127537 | 0.548050703 | 1.849474267 | 5.813995433 | PLK3;TBK1;DHRSX; | 12 |
| positive regulation of pr | Jun-34 | 0.043269586 | 0.548050703 | 2.572964169 | 8.079893026 | KLF5;PDGFB;NR3C1 | 6 |
| cartilage development ( | Aug-52 | 0.044225474 | 0.548050703 | 2.184071636 | 6.810927631 | BMP2;ITGB8;TRIP11 | 8 |
| positive regulation of de | Aug-52 | 0.044225474 | 0.548050703 | 2.184071636 | 6.810927631 | RIMS2;MACF1;DBN | 8 |
| regulation of cell-cell ad | Jul-43 | 0.044533225 | 0.548050703 | 2.335234681 | 7.266128785 | EPHA7;TENM3;JAG1 | 7 |
| positive regulation of ce | Oct-71 | 0.044882185 | 0.548050703 | 1.969997109 | 6.114308283 | DTX3L;LIMK2;PYHI | 10 |
| chaperone cofactor-dep | May-26 | 0.045603904 | 0.548050703 | 2.858072917 | 8.825048831 | GAK;HSPA1L;DNAJ | 5 |
| endothelial cell develop | May-26 | 0.045603904 | 0.548050703 | 2.858072917 | 8.825048831 | RAP1A;MYADM;PE | 5 |
| metanephros developme | May-26 | 0.045603904 | 0.548050703 | 2.858072917 | 8.825048831 | NIPBL;GDNF;BASP1 | 5 |
| mononuclear cell differe | May-26 | 0.045603904 | 0.548050703 | 2.858072917 | 8.825048831 | LYN;CSF1R;TRPM2; | 5 |
| negative regulation of in | May-26 | 0.045603904 | 0.548050703 | 2.858072917 | 8.825048831 | SERPINB1;CPTP;ZC3 | 5 |
| regulation of monocyte | May-26 | 0.045603904 | 0.548050703 | 2.858072917 | 8.825048831 | CCR1;LYN;DUSP1;S | 5 |
| regulation of synapse o | May-26 | 0.045603904 | 0.548050703 | 2.858072917 | 8.825048831 | ARHGAP22;GRIPAP | 5 |
| regulation of telomere m | May-26 | 0.045603904 | 0.548050703 | 2.858072917 | 8.825048831 | RTEL1;SMG1;TINF2; | 5 |
| negative regulation of tr | 65/684 | 0.045727312 | 0.548050703 | 1.269203322 | 3.9155678 | CIITA;CITED2;BHLH | 65 |
| positive regulation of ne | Oct-72 | 0.048676169 | 0.548050703 | 1.938117612 | 5.858087843 | TIAM2;TIAM1;SERP | 10 |
| cellular response to hyd | Jun-35 | 0.048950046 | 0.548050703 | 2.484106481 | 7.494437379 | TRPM2;PPIF;TNFAIF | 6 |
| semaphorin-plexin sign | Jun-35 | 0.048950046 | 0.548050703 | 2.484106481 | 7.494437379 | FARP2;SEMA4A;SEM | 6 |
| branching morphogenes | Jul-44 | 0.04958015 | 0.548050703 | 2.271996899 | 6.825452962 | DLL4;BMP2;GDNF;S | 7 |
| positive regulation of os | Jul-44 | 0.04958015 | 0.548050703 | 2.271996899 | 6.825452962 | BMP2;BMPR2;SUCO | 7 |
| regulation of peptidyl-ty | Dec-92 | 0.049720625 | 0.548050703 | 1.803041203 | 5.411531462 | EPHA7;SEMA4D;RIP | 12 |

| | | | | | | | |
|---|---|---|---|---|---|---|---|
| regulation of plasma me | Dec-70 | 0.031768484 | 0.394124969 | 1.971364555 | 6.799789443 | USP6NL;CYLD;TBC1 | 12 |
| wound healing (GO:00 | Dec-70 | 0.031768484 | 0.394124969 | 1.971364555 | 6.799789443 | TGFB2;SMPD1;SLC1 | 12 |
| regulation of biominera | Jun-26 | 0.032304357 | 0.395777804 | 2.855450237 | 9.801484782 | CEBPB;NOTCH1;TGI | 6 |
| regulation of regulatory | Jun-26 | 0.032304357 | 0.395777804 | 2.855450237 | 9.801484782 | IL4I1;DUSP10;BCL6; | 6 |
| negative regulation of c | Aug-40 | 0.032524105 | 0.395777804 | 2.380469162 | 8.15494883 | KANK1;NOTCH1;JA | 8 |
| apoptotic mitochondrial | Jul-33 | 0.033069597 | 0.395777804 | 2.563082597 | 8.73790984 | AIFM2;ERBB4;BNIP | 7 |
| regulation of blood ves | Oct-55 | 0.033210083 | 0.395777804 | 2.116681325 | 7.207091961 | ADAM17;TGFB1;AN | 10 |
| positive regulation of tr | 103/908 | 0.034280783 | 0.403802094 | 1.227669739 | 4.141139161 | MAML2;PLXND1;SC | 103 |
| actin cytoskeleton reorg | Nov-63 | 0.034408326 | 0.404229384 | 2.015199821 | 6.790128534 | NRP1;BRSK2;SIPA1L | 11 |
| regulation of cell popul | 88/764 | 0.034641341 | 0.405890211 | 1.247970352 | 4.196559239 | BTG2;VIPR1;BTG1;B | 88 |
| cellular protein modific | 115/1025 | 0.034867523 | 0.407462431 | 1.213257413 | 4.071933855 | CHRM3;CSF1;TESK2 | 115 |
| cellular response to lipo | 18/120 | 0.035235813 | 0.40909917 | 1.68268961 | 5.629761664 | CD274;TICAM2;TGFI | 18 |
| negative regulation of M | Sep-48 | 0.035335518 | 0.40909917 | 2.197663096 | 7.346494693 | BMP4;RGS2;DUSP10 | 9 |
| positive regulation of bl | Sep-48 | 0.035335518 | 0.40909917 | 2.197663096 | 7.346494693 | ADAM17;TGFB1;AN | 9 |
| fatty acid transport (GC | May-20 | 0.035931266 | 0.40909917 | 3.171929825 | 10.55030628 | SLC27A1;LCN12;CD3 | 5 |
| lipopolysaccharide-med | May-20 | 0.035931266 | 0.40909917 | 3.171929825 | 10.55030628 | TGFB1;CD6;PRKCE; | 5 |
| microtubule nucleation | May-20 | 0.035931266 | 0.40909917 | 3.171929825 | 10.55030628 | CCDC57;NDE1;CSNK | 5 |
| negative regulation of li | May-20 | 0.035931266 | 0.40909917 | 3.171929825 | 10.55030628 | ABCA1;OSBPL8;NR1 | 5 |
| regulation of lymphocy | May-20 | 0.035931266 | 0.40909917 | 3.171929825 | 10.55030628 | IL4I1;TBX21;SLAMF | 5 |
| regulation of programm | May-20 | 0.035931266 | 0.40909917 | 3.171929825 | 10.55030628 | CYLD;SDCBP;PELI1; | 5 |
| regulation of smooth m | May-20 | 0.035931266 | 0.40909917 | 3.171929825 | 10.55030628 | NRP1;IGFBP3;PDGFI | 5 |
| regulation of protein kin | 28/207 | 0.036806013 | 0.413753501 | 1.493075543 | 4.930275853 | SRC;PDGFB;PDGFA; | 28 |
| blood vessel morphoge | Oct-56 | 0.03710631 | 0.413753501 | 2.070551795 | 6.820331855 | NRP1;FLT1;TGFB1;R | 10 |
| regulation of cell develo | Aug-41 | 0.037154692 | 0.413753501 | 2.308205939 | 7.6001494 | YAP1;BHLHE40;SLA | 8 |
| regulation of release of | Aug-41 | 0.037154692 | 0.413753501 | 2.308205939 | 7.6001494 | MLLT11;MOAP1;BCI | 8 |
| regulation of cilium ass | Nov-64 | 0.038108597 | 0.415154285 | 1.977067602 | 6.459703397 | USP6NL;YAP1;CYLD | 11 |
| epithelial tube morphog | Jul-34 | 0.038337797 | 0.415154285 | 2.468017016 | 8.048990798 | ACVR1;BMP4;RYR2; | 7 |
| negative regulation of b | Jul-34 | 0.038337797 | 0.415154285 | 2.468017016 | 8.048990798 | STARD13;NOTCH1;T | 7 |
| organelle disassembly ( | Jul-34 | 0.038337797 | 0.415154285 | 2.468017016 | 8.048990798 | PLK3;DYRK3;UBA5; | 7 |
| regulation of myoblast | Jul-34 | 0.038337797 | 0.415154285 | 2.468017016 | 8.048990798 | BMP4;BTG1;NOTCH | 7 |
| negative regulation of c | 13/80 | 0.038551867 | 0.416454195 | 1.848821432 | 6.019301751 | BMPR2;NOTCH1;SPI | 13 |
| sulfur compound biosy | 17/113 | 0.038934901 | 0.419568559 | 1.688200697 | 5.479670249 | CSGALNACT1;SLC3 | 17 |
| positive regulation of ac | Sep-49 | 0.039764181 | 0.426429927 | 2.142602848 | 6.909441527 | BAIAP2L2;BAIAP2L | 9 |
| regulation of smooth m | Sep-49 | 0.039764181 | 0.426429927 | 2.142602848 | 6.909441527 | BMP4;PTGIR;NR4A3 | 9 |
| transforming growth fa | 14/89 | 0.041565155 | 0.444666835 | 1.778811916 | 5.657499012 | ACVR1;JUN;TGFB2; | 14 |
| activation of protein kin | 17/114 | 0.041857967 | 0.446720329 | 1.670703739 | 5.30193342 | OSBPL8;TGFB2;ANG | 17 |
| cellular response to pep | 16/106 | 0.043047955 | 0.450411454 | 1.694488559 | 5.329913048 | SRC;DENND4C;PDE | 16 |
| positive regulation of pr | 16/106 | 0.043047955 | 0.450411454 | 1.694488559 | 5.329913048 | SLC27A1;TGFB1;FLT | 16 |
| cellular response to ster | May-21 | 0.043577953 | 0.450411454 | 2.973519737 | 9.316643722 | BMP4;SRC;NR3C1;TI | 5 |
| focal adhesion assembly | May-21 | 0.043577953 | 0.450411454 | 2.973519737 | 9.316643722 | SRC;PEAK1;TESK2;V | 5 |
| heparan sulfate proteog | May-21 | 0.043577953 | 0.450411454 | 2.973519737 | 9.316643722 | HS3ST3B1;EXT1;GPC | 5 |
| melanosome localizatio | May-21 | 0.043577953 | 0.450411454 | 2.973519737 | 9.316643722 | BLOC1S6;MREG;SHI | 5 |
| negative regulation of c | May-21 | 0.043577953 | 0.450411454 | 2.973519737 | 9.316643722 | ACVR1;IGFBP3;LIF; | 5 |
| negative regulation of p | May-21 | 0.043577953 | 0.450411454 | 2.973519737 | 9.316643722 | DAB2;TGFB1;NUMB | 5 |
| negative regulation of p | May-21 | 0.043577953 | 0.450411454 | 2.973519737 | 9.316643722 | DAB2;TGFB1;NUMB | 5 |
| negative regulation of s | May-21 | 0.043577953 | 0.450411454 | 2.973519737 | 9.316643722 | TGFB2;SPRY4;SPRY | 5 |
| neurotrophin signaling | May-21 | 0.043577953 | 0.450411454 | 2.973519737 | 9.316643722 | RAP1A;BDNF;RAPG | 5 |
| positive regulation of in | May-21 | 0.043577953 | 0.450411454 | 2.973519737 | 9.316643722 | HCAR2;TGFB2;RSAI | 5 |
| regulation of anoikis (G | May-21 | 0.043577953 | 0.450411454 | 2.973519737 | 9.316643722 | NOTCH1;SRC;SIK1;I | 5 |
| cellular response to amy | Jul-35 | 0.044126781 | 0.450411454 | 2.379741834 | 7.426432755 | PRNP;CACNA1A;TR | 7 |
| peptidyl-tyrosine depho | Jul-35 | 0.044126781 | 0.450411454 | 2.379741834 | 7.426432755 | DUSP4;DUSP5;PTPR | 7 |
| regulation of phosphati | Jul-35 | 0.044126781 | 0.450411454 | 2.379741834 | 7.426432755 | FLT1;TGFB1;PDGFB; | 7 |
| respiratory system deve | Jul-35 | 0.044126781 | 0.450411454 | 2.379741834 | 7.426432755 | BMP4;FGF9;ITGA3;C | 7 |
| negative regulation of E | Sep-50 | 0.044553987 | 0.450411454 | 2.090228466 | 6.50281285 | SPRED3;DUSP4;SPR | 9 |
| negative regulation of tr | 106/948 | 0.04471811 | 0.450411454 | 1.207333449 | 3.751639851 | RB1;NAB2;WWC2;ZE | 106 |
| positive regulation of de | Jun-28 | 0.044950124 | 0.450411454 | 2.595576619 | 8.052002338 | DTX3L;DDX58;ZC3H | 6 |
| regulation of B cell acti | Jun-28 | 0.044950124 | 0.450411454 | 2.595576619 | 8.052002338 | IL4I1;THEMIS2;TNFS | 6 |
| regulation of plasma me | Jun-28 | 0.044950124 | 0.450411454 | 2.595576619 | 8.052002338 | BRSK2;HOMER1;NC | 6 |
| positive regulation of tr | 130/1183 | 0.044993823 | 0.450411454 | 1.185324291 | 3.675963337 | MAML2;SCX;NR3C1 | 130 |
| positive regulation of bi | 14/90 | 0.045081346 | 0.450411454 | 1.755309082 | 5.440206135 | DTX3L;PLAUR;HIP1 | 14 |
| chordate embryonic dev | Oct-58 | 0.045822427 | 0.450411454 | 1.984058927 | 6.116817227 | KIAA1217;FUT8;BMI | 10 |
| positive regulation of pr | Oct-58 | 0.045822427 | 0.450411454 | 1.984058927 | 6.116817227 | BMP4;BMP2;DTX3L; | 10 |
| positive regulation of ce | 20/141 | 0.046060486 | 0.450411454 | 1.576079093 | 4.850855955 | PFKFB4;PLEKHF1;D | 20 |
| regulation of protein ph | 34/266 | 0.047375489 | 0.450411454 | 1.399173409 | 4.266989597 | CEBPB;ITGB2;TESK1 | 34 |
| positive regulation of st | 15/99 | 0.047626192 | 0.450411454 | 1.701719577 | 5.180668146 | TGFB2;GADD45B;FZ | 15 |
| negative regulation of ty | Aug-43 | 0.04769541 | 0.450411454 | 2.176067475 | 6.621599474 | IFIH1;RNF135;CYLD | 8 |
| regulation of cell-cell ac | Aug-43 | 0.04769541 | 0.450411454 | 2.176067475 | 6.621599474 | AFDN;JAG1;TGFB1; | 8 |
| regulation of cell motilit | 19/133 | 0.047882467 | 0.451215844 | 1.588989042 | 4.828947031 | TGFB1;PLXND1;SER | 19 |
| intrinsic apoptotic signa | Sep-51 | 0.04971422 | 0.466492118 | 2.040348101 | 6.124031929 | MOAP1;BCL2L11;BC | 9 |
| negative regulation of p | Sep-51 | 0.04971422 | 0.466492118 | 2.040348101 | 6.124031929 | DTX3L;NXN;TNFAIF | 9 |

**Additional file 9**
**Table S3: Lists of RNA 5hmC-enriched peaks in monocytes and macrophages.**

| mo | | | |
|---|---|---|---|
| **Chr** | **Start** | **End** | **Gene name** |
| 13 | 109786086 | 109786469 | IRS2 |
| 13 | 109786086 | 109786469 | RP11-40E6.1 |
| 3 | 133573830 | 133574052 | CDV3 |
| 3 | 170038002 | 170038207 | GPR160 |
| 3 | 196568665 | 196568917 | WDR53 |
| 3 | 196568665 | 196568917 | FBXO45 |
| 4 | 26321262 | 26321476 | RBPJ |
| 4 | 39977588 | 39977870 | PDS5A |
| 4 | 55396124 | 55396324 | TMEM165 |
| 5 | 344006 | 344255 | PDCD6 |
| 5 | 344006 | 344255 | AHRR |
| 5 | 473406 | 473645 | CTD-2228K2.7 |
| 5 | 473406 | 473645 | SLC9A3 |
| 5 | 10564349 | 10564553 | ANKRD33B |
| 5 | 17275438 | 17275732 | BASP1 |
| 5 | 36241696 | 36241917 | NADK2 |
| 5 | 154755837 | 154756038 | LARP1 |
| 1 | 212285570 | 212285771 | PPP2R5A |
| 1 | 220879960 | 220880179 | HLX-AS1 |
| 1 | 220879960 | 220880179 | HLX |
| 1 | 229558286 | 229558517 | ABCB10 |
| 6 | 7910574 | 7910774 | TXNDC5 |
| 6 | 7910574 | 7910774 | BLOC1S5-TXNDC5 |
| 10 | 70404179 | 70404409 | EIF4EBP2 |
| 10 | 91798627 | 91798849 | TNKS2 |
| 10 | 96586758 | 96586970 | TM9SF3 |
| 6 | 45422668 | 45422874 | RUNX2 |
| 6 | 89352551 | 89352754 | UBE2J1 |
| 6 | 170584355 | 170584579 | PDCD2 |
| 7 | 192658 | 192861 | AC093627.12 |
| 7 | 3301340 | 3301565 | SDK1 |
| 7 | 3301340 | 3301565 | AC073316.1 |
| 7 | 5388479 | 5388680 | TNRC18 |
| 19 | 929378 | 929591 | ARID3A |
| 11 | 3855689 | 3855917 | STIM1 |
| 11 | 32829802 | 32830005 | PRRG4 |
| 11 | 63839248 | 63839473 | MARK2 |
| 11 | 64246507 | 64246710 | PPP1R14B |
| 11 | 64246507 | 64246710 | RP11-783K16.5 |
| 11 | 94544460 | 94544663 | FUT4 |
| 11 | 94544460 | 94544663 | PIWIL4 |
| 11 | 120211039 | 120211248 | OAF |
| 12 | 752962 | 753166 | WNK1 |
| 8 | 30812559 | 30812813 | PPP2CB |
| 8 | 38787225 | 38787437 | TACC1 |
| 8 | 65841499 | 65841705 | PDE7A |
| 8 | 67343433 | 67343653 | ARFGEF1 |
| 12 | 71754926 | 71755157 | RAB21 |
| 12 | 92145517 | 92145739 | BTG1 |
| 12 | 92145517 | 92145739 | RP11-796E2.4 |
| 12 | 104137604 | 104137958 | NFYB |
| 12 | 104457024 | 104457227 | CHST11 |
| 12 | 111418413 | 111418622 | SH2B3 |

| Mϕ | | | |
|---|---|---|---|
| **Chr** | **Start** | **End** | **Gene name** |
| 21 | 8216762 | 8217006 | CH507-513H4.1 |
| 21 | 8400315 | 8400516 | CH507-528H12.1 |
| 21 | 39349229 | 39349430 | HMGN1 |
| 22 | 28800294 | 28800511 | XBP1 |
| 22 | 42615150 | 42615397 | RNU12 |
| 3 | 62372384 | 62372585 | FEZF2 |
| 3 | 71753625 | 71753825 | EIF4E3 |
| 3 | 129315971 | 129316184 | H1FX |
| 3 | 129315971 | 129316184 | H1FX-AS1 |
| 4 | 15003756 | 15003958 | CPEB2 |
| 5 | 443097 | 443297 | EXOC3 |
| 5 | 443097 | 443297 | EXOC3-AS1 |
| 5 | 168579365 | 168579568 | PANK3 |
| 6 | 33788818 | 33789022 | LEMD2 |
| 6 | 36639471 | 36639678 | RNU1-88P |
| 6 | 52995664 | 52995867 | 7SK |
| 6 | 52995664 | 52995867 | RN7SK |
| 7 | 193297 | 193569 | AC093627.12 |
| 7 | 193297 | 193569 | FAM20C |
| 7 | 3301234 | 3301472 | SDK1 |
| 7 | 3301234 | 3301472 | AC073316.1 |
| 7 | 65873155 | 65873360 | VKORC1L1 |
| 8 | 143977227 | 143977443 | PARP10 |
| 9 | 35657771 | 35657976 | RMRP |
| 1 | 10694452 | 10694652 | CASZ1 |
| 1 | 28508499 | 28508713 | SNORA73B |
| 1 | 28508499 | 28508713 | RCC1 |
| 1 | 28508499 | 28508713 | SNHG3 |
| 1 | 206685047 | 206685252 | MAPKAPK2 |
| X | 47586596 | 47586799 | TIMP1 |
| X | 47586596 | 47586799 | SYN1 |
| 10 | 130157454 | 130157655 | GLRX3 |
| 11 | 27700481 | 27700682 | BDNF |
| 11 | 27700481 | 27700682 | RP11-587D21.4 |
| X | 84187841 | 84188041 | RPS6KA6 |
| X | 150983268 | 150983515 | HMGB3 |
| 11 | 47355000 | 47355219 | SPI1 |
| 11 | 65499543 | 65499803 | MALAT1 |
| 11 | 93732336 | 93732550 | TAF1D |
| 11 | 93732336 | 93732550 | SNORA8 |
| 12 | 752924 | 753158 | WNK1 |
| 12 | 117370191 | 117370392 | NOS1 |
| 12 | 120291720 | 120291934 | RNU4-2 |
| 12 | 120293052 | 120293267 | RNU4-1 |
| 14 | 99420812 | 99421013 | SETD3 |
| 15 | 40358106 | 40358351 | DISP2 |
| 1 | 40861928 | 40862133 | CITED4 |
| 16 | 28063165 | 28063366 | GSG1L |
| 16 | 87492563 | 87492797 | RP11-482M8.1 |
| 16 | 88643298 | 88643507 | CYBA |
| 17 | 45221961 | 45222181 | CTD-2020K17.1 |
| 17 | 45221961 | 45222181 | FMNL1 |
| 19 | 519411 | 519646 | TPGS1 |

| M1 | | | |
|---|---|---|---|
| **Chr** | **Start** | **End** | **Gene name** |
| 8 | 11842483 | 11842686 | CTSB |
| X | 32866586 | 32866790 | DMD |
| 11 | 65499634 | 65499858 | MALAT1 |
| 19 | 48966514 | 48966783 | FTL |
| 2 | 169636706 | 169636907 | PPIG |
| 3 | 194486697 | 194486909 | ATP13A3 |
| 17 | 80281254 | 80281505 | RNF213 |
| 10 | 22714315 | 22714556 | PIP4K2A |
| 10 | 70404172 | 70404404 | EIF4EBP2 |
| 10 | 130157451 | 130157655 | GLRX3 |
| 1 | 24929938 | 24930143 | RUNX3 |
| 11 | 1956360 | 1956581 | MRPL23 |
| 11 | 27700479 | 27700682 | BDNF |
| 11 | 27700479 | 27700682 | RP11-587D21.4 |
| 11 | 47355038 | 47355244 | SPI1 |
| 11 | 67266439 | 67266685 | GRK2 |
| 11 | 67991464 | 67991703 | UNC93B1 |
| 11 | 68003825 | 68004027 | UNC93B1 |
| 12 | 752828 | 753141 | WNK1 |
| 12 | 27332854 | 27333095 | ARNTL2 |
| 12 | 104137677 | 104137897 | NFYB |
| 13 | 109786274 | 109786475 | IRS2 |
| 13 | 109786274 | 109786475 | RP11-40E6.1 |
| 1 | 28508496 | 28508704 | SNORA73B |
| 1 | 28508496 | 28508704 | RCC1 |
| 1 | 28508496 | 28508704 | SNHG3 |
| 1 | 31938088 | 31938306 | PTP4A2 |
| 1 | 32964580 | 32964793 | RNF19B |
| 1 | 35192306 | 35192609 | SFPQ |
| 1 | 206685031 | 206685244 | MAPKAPK2 |
| 1 | 212285560 | 212285806 | PPP2R5A |
| 1 | 229558252 | 229558586 | ABCB10 |
| 13 | 91348714 | 91348930 | MIR17HG |
| 14 | 23321497 | 23321738 | PABPN1 |
| 14 | 23321497 | 23321738 | BCL2L2-PABPN1 |
| 14 | 54902522 | 54902736 | GCH1 |
| 14 | 92719186 | 92719390 | LGMN |
| 15 | 25863087 | 25863348 | ATP10A |
| 15 | 101251650 | 101251864 | CHSY1 |
| 16 | 381679 | 381927 | TMEM8A |
| 16 | 680413 | 680636 | STUB1 |
| 16 | 19114139 | 19114342 | ITPRIPL2 |
| 16 | 19114139 | 19114342 | CTD-2349B8.1 |
| 16 | 28063162 | 28063365 | GSG1L |
| 16 | 28211657 | 28211910 | XPO6 |
| 16 | 48609919 | 48610139 | N4BP1 |
| 16 | 72787572 | 72787819 | RP5-991G20.1 |
| 16 | 72787572 | 72787819 | ZFHX3 |
| 16 | 87602399 | 87602649 | JPH3 |
| 17 | 40122279 | 40122480 | MSL1 |
| 17 | 45221937 | 45222141 | CTD-2020K17.1 |
| 17 | 45221937 | 45222141 | FMNL1 |
| 17 | 62065229 | 62065433 | MED13 |

| M2 | | | |
|---|---|---|---|
| **Chr** | **Start** | **End** | **Gene name** |
| 11 | 65855129 | 65855337 | CFL1 |
| 11 | 65855129 | 65855337 | SNX32 |
| 14 | 20343109 | 20343359 | RPPH1 |
| 19 | 2273037 | 2273241 | OAZ1 |
| 19 | 2273037 | 2273241 | LLfos-48D6.2 |
| 19 | 17403046 | 17403255 | BST2 |
| 19 | 40378176 | 40378380 | PLD3 |
| 19 | 49889719 | 49889947 | IL4I1 |
| 2 | 215435872 | 215436079 | FN1 |
| 22 | 36289110 | 36289311 | MYH9 |
| 3 | 131502717 | 131502948 | MRPL3 |
| 6 | 31356174 | 31356404 | HLA-B |
| 6 | 44250220 | 44250436 | HSP90AB1 |
| 9 | 127506096 | 127506312 | FAM129B |
| 1 | 27668218 | 27668421 | IFI6 |
| 1 | 42456621 | 42456844 | PPCS |
| 10 | 17229378 | 17229601 | VIM-AS1 |
| 10 | 17229378 | 17229601 | VIM |
| 11 | 60951079 | 60951326 | SLC15A3 |
| 19 | 49490575 | 49490801 | RPL13A |
| 19 | 49490575 | 49490801 | SNORD33 |
| 21 | 8397590 | 8397796 | CH507-528H12.1 |
| 21 | 8400314 | 8400516 | CH507-528H12.1 |
| 21 | 34073656 | 34073873 | MRPS6 |
| 21 | 34073656 | 34073873 | SLC5A3 |
| 21 | 34073656 | 34073873 | AP000320.7 |
| 21 | 34074007 | 34074227 | MRPS6 |
| 21 | 34074007 | 34074227 | SLC5A3 |
| 21 | 34074007 | 34074227 | AP000320.7 |
| 21 | 41830164 | 41830395 | PRDM15 |
| 22 | 28800303 | 28800513 | XBP1 |
| 5 | 154755380 | 154755600 | LARP1 |
| 6 | 52995636 | 52995864 | 7SK |
| 6 | 52995636 | 52995864 | RN7SK |
| 7 | 237638 | 237871 | FAM20C |
| 7 | 47474552 | 47474882 | TNS3 |
| 8 | 38787229 | 38787440 | TACC1 |
| 9 | 35657753 | 35657965 | RMRP |
| 9 | 111896842 | 111897042 | UGCG |
| 9 | 136672468 | 136672673 | EGFL7 |
| X | 150983268 | 150983477 | HMGB3 |
| 7 | 50334649 | 50334972 | IKZF1 |
| 1 | 44721738 | 44721950 | RNU5F-1 |
| 1 | 44721738 | 44721950 | C1orf228 |
| 11 | 93732330 | 93732553 | TAF1D |
| 11 | 93732330 | 93732553 | SNORA8 |
| 12 | 120291719 | 120291924 | RNU4-2 |
| 12 | 120293053 | 120293258 | RNU4-1 |
| 14 | 49862502 | 49862702 | RP11-596C23.6 |
| 14 | 49862502 | 49862702 | RN7SL2 |
| 14 | 92719183 | 92719386 | LGMN |
| 14 | 105168146 | 105168357 | JAG2 |
| 14 | 105168146 | 105168357 | RP11-44N21.4 |

| | | | |
|---|---|---|---|
| 12 | 124993724 | 124993947 | BRI3BP |
| 8 | 143792308 | 143792521 | SCRIB |
| 8 | 144358169 | 144358403 | SLC52A2 |
| 8 | 144358169 | 144358403 | FBXL6 |
| 8 | 144517522 | 144517751 | RECQL4 |
| 13 | 91348775 | 91348986 | MIR17HG |
| 13 | 99981881 | 99982123 | ZIC2 |
| 9 | 2621603 | 2621835 | VLDLR |
| 9 | 2621603 | 2621835 | VLDLR-AS1 |
| 9 | 33817469 | 33817784 | UBE2R2 |
| 9 | 33817469 | 33817784 | UBE2R2-AS1 |
| 9 | 37800862 | 37801063 | DCAF10 |
| 9 | 37800862 | 37801063 | EXOSC3 |
| 9 | 37800862 | 37801063 | RP11-613M10.9 |
| 14 | 49862534 | 49862745 | RP11-596C23.6 |
| 14 | 49862534 | 49862745 | RN7SL2 |
| 9 | 137070018 | 137070392 | SAPCD2 |
| 9 | 137278718 | 137278937 | TOR4A |
| 9 | 137459055 | 137459279 | NSMF |
| 14 | 91509663 | 91509884 | PPP4R3A |
| 14 | 99420807 | 99421016 | SETD3 |
| 15 | 31326936 | 31327143 | KLF13 |
| 15 | 74782220 | 74782440 | CSK |
| 15 | 83066944 | 83067169 | BTBD1 |
| 15 | 83066944 | 83067169 | RP11-382A20.6 |
| 15 | 83066944 | 83067169 | RP11-382A20.5 |
| 16 | 4116048 | 4116305 | ADCY9 |
| 16 | 11915586 | 11915794 | GSPT1 |
| 16 | 54285998 | 54286208 | IRX3 |
| 16 | 54285998 | 54286208 | RP11-434E6.5 |
| 16 | 72787572 | 72787814 | RP5-991G20.1 |
| 16 | 72787572 | 72787814 | ZFHX3 |
| 16 | 81444998 | 81445224 | CMIP |
| 16 | 88803855 | 88804057 | CDT1 |
| 17 | 16353167 | 16353379 | CENPV |
| 17 | 40122878 | 40123089 | MSL1 |
| 17 | 45221952 | 45222187 | CTD-2020K17.1 |
| 17 | 45221952 | 45222187 | FMNL1 |
| 17 | 67825523 | 67825753 | BPTF |
| 18 | 2906838 | 2907092 | EMILIN2 |
| 18 | 54224346 | 54224586 | MBD2 |
| 18 | 62715649 | 62715896 | PHLPP1 |
| 19 | 797569 | 797770 | PTBP1 |
| 19 | 821659 | 821868 | PLPPR3 |
| 19 | 1206175 | 1206416 | STK11 |
| 19 | 12734474 | 12734678 | C19orf43 |
| 19 | 13140608 | 13140835 | NACC1 |
| 19 | 13140608 | 13140835 | CTC-250I14.3 |
| 19 | 14090341 | 14090590 | SAMD1 |
| 19 | 17245704 | 17245904 | NR2F6 |
| 19 | 17245704 | 17245904 | AC010646.3 |
| 19 | 18940967 | 18941168 | HOMER3 |
| 19 | 18940967 | 18941168 | AC005932.1 |
| 19 | 33302234 | 33302442 | CEBPA |
| 19 | 35000327 | 35000538 | GRAMD1A |
| 19 | 35718170 | 35718373 | KMT2B |
| 19 | 38849669 | 38849870 | HNRNPL |
| 19 | 55407538 | 55407759 | UBE2S |
| X | 40097648 | 40097936 | BCOR |
| 19 | 2248201 | 2248443 | SF3A2 |
| 19 | 3544185 | 3544396 | MFSD12 |
| 19 | 3544185 | 3544396 | AC005786.7 |
| 19 | 18419437 | 18419663 | SSBP4 |
| 19 | 48966590 | 48966840 | FTL |
| 19 | 55258473 | 55258727 | PPP6R1 |
| 2 | 39120825 | 39121033 | SOS1 |
| 13 | 109786157 | 109786461 | IRS2 |
| 13 | 109786157 | 109786461 | RP11-40E6.1 |
| 2 | 109794459 | 109794703 | RGPD5 |
| 20 | 408078 | 408281 | RBCK1 |
| 8 | 22693231 | 22693440 | EGR3 |
| 8 | 22693231 | 22693440 | RP11-459E5.1 |
| 17 | 39921788 | 39922003 | ORMDL3 |
| 7 | 50334946 | 50335207 | IKZF1 |
| 1 | 26696445 | 26696673 | ARID1A |
| 1 | 31938092 | 31938327 | PTP4A2 |
| 1 | 40373830 | 40374062 | SMAP2 |
| 1 | 41620621 | 41620858 | HIVEP3 |
| 1 | 45637292 | 45637565 | GPBP1L1 |
| 1 | 92275306 | 92275512 | GLMN |
| 19 | 19628255 | 19628460 | LPAR2 |
| 9 | 135836653 | 135836866 | CAMSAP1 |
| MT | 14065 | 14344 | MT-ND5 |
| MT | 14065 | 14344 | MT-ND6 |
| 5 | 154446242 | 154446507 | SAP30L |
| 1 | 108699464 | 108699760 | PRPF38B |
| 1 | 113929660 | 113929889 | HIPK1 |
| 1 | 156670775 | 156671029 | NES |
| 6 | 43171448 | 43171847 | SRF |
| 6 | 43628991 | 43629197 | GTPBP2 |
| 6 | 44250219 | 44250431 | HSP90AB1 |
| 6 | 108561381 | 108561590 | FOXO3 |
| 6 | 113860216 | 113860424 | MARCKS |
| 6 | 136792156 | 136792362 | MAP3K5 |
| 6 | 149317680 | 149317885 | TAB2 |
| 6 | 159710663 | 159710964 | SOD2 |
| 6 | 169724266 | 169724473 | PHF10 |
| 7 | 240242 | 240480 | FAM20C |
| 10 | 70404205 | 70404411 | EIF4EBP2 |
| 7 | 6706689 | 6706894 | ZNF12 |
| 7 | 6706689 | 6706894 | AC073343.13 |
| 7 | 47351986 | 47352238 | TNS3 |
| 11 | 47356735 | 47356951 | SPI1 |
| 7 | 47515447 | 47515683 | TNS3 |
| 11 | 47358085 | 47358297 | SPI1 |
| 11 | 58579228 | 58579429 | ZFP91 |
| 11 | 58579228 | 58579429 | ZFP91-CNTF |
| 7 | 99438924 | 99439139 | CPSF4 |
| 7 | 99438924 | 99439139 | PTCD1 |
| 7 | 99438924 | 99439139 | ATP5J2-PTCD1 |
| 8 | 80486462 | 80486664 | ZBTB10 |
| 1 | 40210451 | 40210677 | RLF |
| 8 | 144326513 | 144326734 | DGAT1 |
| 8 | 144528792 | 144529043 | C8orf82 |
| 12 | 76031292 | 76031498 | PHLDA1 |
| 12 | 76031292 | 76031498 | RP11-290L1.3 |
| 12 | 104137600 | 104137872 | NFYB |
| 12 | 110281467 | 110281681 | ATP2A2 |
| 17 | 76501070 | 76501278 | RHBDF2 |
| 17 | 78925110 | 78925344 | TIMP2 |
| 17 | 83079604 | 83079810 | METRNL |
| 18 | 51030367 | 51030569 | SMAD4 |
| 18 | 51030367 | 51030569 | RP11-729L2.2 |
| 19 | 10283526 | 10283754 | ICAM1 |
| 19 | 10283526 | 10283754 | CTD-2369P2.5 |
| 19 | 18419461 | 18419672 | SSBP4 |
| 19 | 55117367 | 55117649 | PPP1R12C |
| 2 | 109794454 | 109794662 | RGPD5 |
| 2 | 148644811 | 148645026 | EPC2 |
| 2 | 201166121 | 201166326 | CFLAR |
| 2 | 224585027 | 224585229 | CUL3 |
| 2 | 239345799 | 239346047 | HDAC4 |
| 20 | 49713608 | 49713809 | B4GALT5 |
| 20 | 58891371 | 58891583 | GNAS |
| 21 | 8216762 | 8217009 | CH507-513H4.1 |
| 21 | 8397572 | 8397793 | CH507-528H12.1 |
| 21 | 41830143 | 41830534 | PRDM15 |
| 3 | 71389098 | 71389302 | FOXP1 |
| 3 | 129605150 | 129605353 | PLXND1 |
| 4 | 15003723 | 15003981 | CPEB2 |
| 4 | 176792313 | 176792526 | VEGFC |
| 5 | 443097 | 443336 | EXOC3 |
| 5 | 443097 | 443336 | EXOC3-AS1 |
| 5 | 856739 | 856957 | BRD9 |
| 5 | 10564343 | 10564563 | ANKRD33B |
| 5 | 45695731 | 45695936 | HCN1 |
| 5 | 154755800 | 154756045 | LARP1 |
| 5 | 168579340 | 168579564 | PANK3 |
| 6 | 71288860 | 71289101 | OGFRL1 |
| 6 | 71288860 | 71289101 | RP11-154D6.1 |
| 15 | 74782224 | 74782435 | CSK |
| 6 | 113859797 | 113860032 | MARCKS |
| 6 | 149317702 | 149317920 | TAB2 |
| 6 | 154733504 | 154733730 | SCAF8 |
| 6 | 163414737 | 163415062 | QKI |
| 7 | 192690 | 192903 | AC093627.12 |
| 7 | 193302 | 193532 | AC093627.12 |
| 7 | 193302 | 193532 | FAM20C |
| 7 | 237588 | 237833 | FAM20C |
| 7 | 519414 | 519635 | PDGFA |
| 7 | 30285156 | 30285363 | ZNRF2 |
| 7 | 65873155 | 65873356 | VKORC1L1 |
| 7 | 98281710 | 98281931 | BRI3 |
| 8 | 30812564 | 30812779 | PPP2CB |
| 8 | 38787230 | 38787445 | TACC1 |
| 8 | 130358376 | 130358593 | ASAP1 |
| 8 | 143733535 | 143733744 | FAM83H |
| 9 | 113250 | 113458 | LINC01388 |
| 9 | 35605456 | 35605675 | TESK1 |
| 9 | 77020192 | 77020479 | FOXB2 |
| 9 | 128689334 | 128689547 | SET |
| X | 84187840 | 84188044 | RPS6KA6 |
| 16 | 9091730 | 9091966 | C16orf72 |
| 11 | 6611859 | 6612067 | TAF10 |
| X | 139932404 | 139932638 | ATP11C |
| X | 147911980 | 147912225 | FMR1 |
| 22 | 50525836 | 50526206 | SCO2 |
| 15 | 63121881 | 63122088 | LACTB |
| 16 | 28063157 | 28063366 | GSG1L |
| 16 | 72787578 | 72787821 | RP5-991G20.1 |
| 16 | 72787578 | 72787821 | ZFHX3 |
| 16 | 87602395 | 87602604 | JPH3 |
| 16 | 88643315 | 88643516 | CYBA |
| 17 | 8173426 | 8173632 | TMEM107 |
| 17 | 8173426 | 8173632 | SNORD118 |
| 1 | 6030693 | 6030949 | KCNAB2 |
| 1 | 10694454 | 10694661 | CASZ1 |
| 17 | 42980586 | 42980815 | RUNDC1 |
| 17 | 50056080 | 50056329 | ITGA3 |
| 17 | 62065229 | 62065438 | MED13 |
| 19 | 10928856 | 10929061 | C19orf52 |
| 19 | 10960888 | 10961092 | SMARCA4 |
| 19 | 16076561 | 16076766 | TPM4 |
| 19 | 16076561 | 16076766 | CTD-2231E14.2 |
| 19 | 18281311 | 18281627 | JUND |
| 19 | 42242288 | 42242508 | GSK3A |
| 19 | 42242288 | 42242508 | AC006486.9 |
| 19 | 49107795 | 49108028 | SNRNP70 |
| 19 | 55117424 | 55117641 | PPP1R12C |
| 2 | 101252899 | 101253104 | CNOT11 |
| 2 | 148644786 | 148644991 | EPC2 |
| 2 | 177217566 | 177217767 | HNRNPA3 |
| 2 | 231711907 | 231712131 | PTMA |
| 2 | 234496817 | 234497033 | ARL4C |
| 20 | 25390599 | 25390809 | ABHD12 |
| 20 | 59007031 | 59007235 | CTSZ |
| 21 | 45992609 | 45992869 | COL6A1 |
| 22 | 38700712 | 38700914 | JOSD1 |
| 22 | 38735778 | 38736015 | GTPBP1 |
| 22 | 38735778 | 38736015 | SUN2 |
| 22 | 38735778 | 38736015 | RP3-508I15.18 |
| 22 | 38735778 | 38736015 | RP3-508I15.19 |
| 5 | 68215773 | 68215975 | PIK3R1 |
| 5 | 79513844 | 79514109 | HOMER1 |
| 5 | 87268328 | 87268591 | RASA1 |
| 6 | 62285966 | 62286207 | KHDRBS2 |
| 6 | 149317699 | 149317912 | TAB2 |
| 7 | 519424 | 519702 | PDGFA |
| 7 | 99438939 | 99439160 | CPSF4 |
| 7 | 99438939 | 99439160 | PTCD1 |
| 7 | 99438939 | 99439160 | ATP5J2-PTCD1 |
| 7 | 158829302 | 158829515 | ESYT2 |
| 8 | 30812569 | 30812784 | PPP2CB |
| 8 | 42896960 | 42897161 | HOOK3 |
| 8 | 42896960 | 42897161 | RNF170 |
| 8 | 80486469 | 80486730 | ZBTB10 |
| 8 | 143733523 | 143733728 | FAM83H |
| 8 | 144137856 | 144138057 | HGH1 |
| 9 | 38068923 | 38069163 | SHB |
| 9 | 127568854 | 127569066 | FAM129B |
| 9 | 128689337 | 128689543 | SET |
| 9 | 137205444 | 137205647 | TMEM203 |
| 11 | 47356728 | 47356993 | SPI1 |
| 1 | 154558469 | 154558671 | UBE2Q1 |
| 1 | 212285526 | 212285735 | PPP2R5A |
| 1 | 229558314 | 229558547 | ABCB10 |

| | | | | | | | | | | | | | | | |
|---|---|---|---|---|---|---|---|---|---|---|---|---|---|---|---|
| 2 | 180980616 | 180980831 | UBE2E3 | 9 | 5832803 | 5833042 | ERMP1 | 22 | 50525836 | 50526206 | TYMP | 11 | 6611853 | 6612143 | TAF10 |
| 2 | 230712922 | 230713125 | CAB39 | 9 | 35605455 | 35605721 | TESK1 | 19 | 18281317 | 18281518 | JUND | 11 | 65572587 | 65572797 | SSSCA1 |
| 20 | 408078 | 408285 | RBCK1 | 9 | 35689889 | 35690104 | TPM2 | 10 | 79069016 | 79069256 | ZMIZ1 | 11 | 65572587 | 65572797 | FAM89B |
| 20 | 36050724 | 36050935 | NORAD | 9 | 125786497 | 125786712 | PBX3 | 11 | 60951409 | 60951623 | SLC15A3 | 12 | 71754948 | 71755165 | RAB21 |
| 20 | 58891359 | 58891570 | GNAS | 9 | 127568850 | 127569069 | FAM129B | 1 | 150150766 | 150150967 | PLEKHO1 | 12 | 104137620 | 104137887 | NFYB |
| 21 | 34791973 | 34792178 | RUNX1 | 9 | 128689343 | 128689550 | SET | 1 | 167553516 | 167553759 | CREG1 | 12 | 120469518 | 120469741 | SRSF9 |
| 22 | 17109204 | 17109415 | IL17RA | MT | 11763 | 12031 | MT-ND4 | 16 | 1964360 | 1964570 | RPS2 | 14 | 22871877 | 22872110 | LRP10 |
| 22 | 23767458 | 23767668 | CHCHD10 | 14 | 23321517 | 23321739 | PABPN1 | 19 | 519419 | 519646 | TPGS1 | 14 | 23321465 | 23321739 | PABPN1 |
| 22 | 38755721 | 38755956 | SUN2 | 14 | 23321517 | 23321739 | BCL2L2-PABPN1 | 19 | 44846340 | 44846564 | NECTIN2 | 14 | 23321465 | 23321739 | BCL2L2-PABPN1 |
| 7 | 2354891 | 2355333 | EIF3B | X | 41216131 | 41216390 | USP9X | 21 | 8401664 | 8401864 | CH507-528H12.1 | 15 | 99105064 | 99105340 | SYNM |
| 22 | 42270423 | 42270649 | OGFRP1 | 15 | 50481730 | 50481963 | USP8 | 5 | 79514194 | 79514404 | HOMER1 | 15 | 99105064 | 99105340 | RP11-654A16.3 |
| 22 | 42270423 | 42270649 | TCF20 | X | 136511693 | 136511966 | HTATSF1 | 7 | 30633809 | 30634043 | GARS | 16 | 680431 | 680650 | STUB1 |
| 7 | 5388789 | 5389137 | TNRC18 | 15 | 90452296 | 90452536 | IQGAP1 | 7 | 157138985 | 157139203 | UBE3C | 16 | 1964350 | 1964568 | RPS2 |
| 19 | 18281307 | 18281518 | JUND | 15 | 101251660 | 101251873 | CHSY1 | 9 | 93134273 | 93134476 | NINJ1 | 16 | 19114168 | 19114370 | ITPRIPL2 |
| 1 | 15409977 | 15410178 | EFHD2 | 16 | 381713 | 381920 | TMEM8A | 7 | 47474619 | 47474940 | TNS3 | 16 | 19114168 | 19114370 | CTD-2349B8.1 |
| 1 | 21176550 | 21176767 | EIF4G3 | 16 | 1964365 | 1964572 | RPS2 | 10 | 28533262 | 28533508 | WAC | 16 | 48609954 | 48610159 | N4BP1 |
| 1 | 39576449 | 39576650 | PABPC4 | 16 | 9091737 | 9091951 | C16orf72 | 10 | 96586763 | 96586968 | TM9SF3 | 17 | 4498887 | 4499140 | AC118754.4 |
| 1 | 40373880 | 40374086 | SMAP2 | 16 | 19114089 | 19114321 | ITPRIPL2 | 11 | 93732336 | 93732552 | TAF1D | 17 | 4498887 | 4499140 | SPNS2 |
| 1 | 51236439 | 51236646 | RNF11 | 16 | 19114089 | 19114321 | CTD-2349B8.1 | 11 | 93732336 | 93732552 | SNORA8 | 1 | 11926199 | 11926408 | KIAA2013 |
| 1 | 92480724 | 92480953 | GFI1 | 16 | 22008118 | 22008330 | C16orf52 | 12 | 92145469 | 92145718 | BTG1 | 2 | 177216786 | 177216992 | HNRNPA3 |
| 1 | 113929715 | 113929920 | HIPK1 | 16 | 48609921 | 48610123 | N4BP1 | 12 | 92145469 | 92145718 | RP11-796E2.4 | 20 | 297695 | 297908 | ZCCHC3 |
| 1 | 150974621 | 150974844 | CERS2 | 17 | 4498918 | 4499159 | AC118754.4 | 1 | 32964202 | 32964417 | RNF19B | 6 | 43171434 | 43171866 | SRF |
| 1 | 150974621 | 150974844 | RP11-316M1.3 | 17 | 4498918 | 4499159 | SPNS2 | 1 | 40373851 | 40374068 | SMAP2 | 17 | 40122306 | 40122536 | MSL1 |
| 1 | 155250338 | 155250575 | FAM189B | 17 | 80281261 | 80281654 | RNF213 | 1 | 150579271 | 150579481 | MCL1 | 17 | 83079598 | 83079807 | METRNL |
| 1 | 156500536 | 156500739 | MEF2D | 17 | 82078443 | 82078681 | FASN | 12 | 120291727 | 120291932 | RNU4-2 | 18 | 54224353 | 54224571 | MBD2 |
| 8 | 130358240 | 130358471 | ASAP1 | 18 | 54224362 | 54224570 | MBD2 | 12 | 120293061 | 120293267 | RNU4-1 | 19 | 3544190 | 3544390 | MFSD12 |
| 3 | 52056145 | 52056350 | DUSP7 | 19 | 4035735 | 4036008 | PIAS4 | 12 | 124567495 | 124567764 | NCOR2 | 19 | 3544190 | 3544390 | AC005786.7 |
| 8 | 144527036 | 144527301 | C8orf82 | 19 | 39435690 | 39435898 | RPS16 | 14 | 49853717 | 49853921 | Metazoa_SRP | 19 | 3557344 | 3557557 | MFSD12 |
| 5 | 178204742 | 178204943 | HNRNPAB | 19 | 42242276 | 42242480 | GSK3A | 14 | 49862501 | 49862731 | RP11-596C23.6 | 19 | 3557344 | 3557557 | AC005786.7 |
| 11 | 6611884 | 6612171 | TAF10 | 19 | 42242276 | 42242480 | AC006486.9 | 14 | 49862501 | 49862731 | RN7SL2 | 19 | 12734442 | 12734671 | C19orf43 |
| 16 | 28063161 | 28063369 | GSG1L | 19 | 42284096 | 42284317 | CIC | 14 | 103127111 | 103127391 | **TNFAIP2** | 19 | 18280174 | 18280398 | JUND |
| 17 | 62065227 | 62065433 | MED13 | 19 | 55117439 | 55117642 | PPP1R12C | 15 | 63121907 | 63122129 | LACTB | 19 | 44908821 | 44909087 | APOE |
| 19 | 1244450 | 1244684 | ATP5D | 19 | 56088358 | 56088564 | ZNF787 | 14 | 49586598 | 49586862 | RN7SL1 | 2 | 230712911 | 230713145 | CAB39 |
| 11 | 93732334 | 93732552 | TAF1D | 2 | 36597953 | 36598157 | FEZ2 | 14 | 49586598 | 49586862 | RPS29 | 3 | 136195763 | 136196024 | MSL2 |
| 11 | 93732334 | 93732552 | SNORA8 | 5 | 473411 | 473662 | CTD-2228K2.7 | 16 | 88643297 | 88643528 | CYBA | 5 | 168579343 | 168579600 | PANK3 |
| 1 | 224434495 | 224434698 | CNIH3 | 5 | 473411 | 473662 | SLC9A3 | 1 | 40861925 | 40862136 | CITED4 | 6 | 20421681 | 20421905 | RN7SL128P |
| 1 | 224434495 | 224434698 | WDR26 | 2 | 148644823 | 148645032 | EPC2 | 1 | 41662575 | 41662791 | HIVEP3 | 6 | 20421681 | 20421905 | E2F3 |
| 19 | 49658264 | 49658510 | SCAF1 | 2 | 177217569 | 177217816 | HNRNPA3 | 18 | 54224354 | 54224558 | MBD2 | 6 | 166956199 | 166956475 | RNASET2 |
| 2 | 32916109 | 32916315 | LINC00486 | 2 | 230712916 | 230713144 | CAB39 | 19 | 3557312 | 3557548 | MFSD12 | 7 | 240239 | 240470 | FAM20C |
| 9 | 125747169 | 125747369 | PBX3 | 2 | 234496785 | 234497052 | ARL4C | 19 | 3557312 | 3557548 | AC005786.7 | 7 | 50335108 | 50335442 | IKZF1 |
| 15 | 40358095 | 40358302 | DISP2 | 2 | 239345795 | 239346063 | HDAC4 | 19 | 12734473 | 12734682 | C19orf43 | 8 | 144326493 | 144326766 | DGAT1 |
| 6 | 26156765 | 26156988 | HIST1H1E | 20 | 57660654 | 57660958 | PMEPA1 | 19 | 18280175 | 18280385 | JUND | 9 | 5832792 | 5832997 | ERMP1 |
| 16 | 492307 | 492517 | RAB11FIP3 | 20 | 59007016 | 59007241 | CTSZ | 19 | 44908825 | 44909072 | APOE | 9 | 127505514 | 127505746 | FAM129B |
| 6 | 52995630 | 52995866 | 7SK | 21 | 41830126 | 41830353 | PRDM15 | 2 | 230712933 | 230713169 | CAB39 | 9 | 133356189 | 133356430 | SURF1 |
| 6 | 52995630 | 52995866 | RN7SK | 22 | 41091886 | 41092107 | EP300 | 20 | 59007018 | 59007253 | CTSZ | X | 47586589 | 47586798 | TIMP1 |
| 21 | 39349226 | 39349433 | HMGN1 | 22 | 50525833 | 50526189 | SCO2 | 21 | 8400314 | 8400517 | CH507-528H12.1 | X | 47586589 | 47586798 | SYN1 |
| 16 | 88645952 | 88646156 | CYBA | 22 | 50525833 | 50526189 | TYMP | 22 | 28800301 | 28800505 | XBP1 | 10 | 28533266 | 28533489 | WAC |
| 17 | 50133767 | 50133971 | PPP1R9B | 3 | 23235529 | 23235754 | UBE2E2 | 6 | 16439207 | 16439455 | ATXN1 | 11 | 65499558 | 65499805 | MALAT1 |
| 17 | 82078469 | 82078681 | FASN | 3 | 25664312 | 25664551 | TOP2B | 6 | 52995664 | 52995866 | 7SK | 6 | 163414722 | 163414967 | QKI |
| 19 | 18280188 | 18280389 | JUND | 3 | 33439988 | 33440222 | UBP1 | 6 | 52995664 | 52995866 | RN7SK | 12 | 110281472 | 110281796 | ATP2A2 |
| 10 | 27154163 | 27154408 | YME1L1 | 8 | 143733470 | 143733710 | FAM83H | 6 | 113860210 | 113860429 | MARCKS | 14 | 49586572 | 49586834 | RN7SL1 |
| 9 | 35689902 | 35690165 | TPM2 | 4 | 40594571 | 40594795 | RBM47 | 6 | 170584337 | 170584572 | PDCD2 | 14 | 49586572 | 49586834 | RPS29 |
| 11 | 67991476 | 67991709 | UNC93B1 | 5 | 856605 | 856912 | BRD9 | 7 | 240210 | 240446 | FAM20C | 15 | 101251656 | 101251867 | CHSY1 |
| 16 | 1445428 | 1445680 | CLCN7 | 5 | 17275394 | 17275717 | BASP1 | 7 | 50334700 | 50335016 | IKZF1 | 1 | 108699439 | 108699686 | PRPF38B |
| 16 | 1445428 | 1445680 | LA16c-390E6.5 | 5 | 154755646 | 154755859 | LARP1 | 8 | 143977213 | 143977447 | PARP10 | 1 | 150579248 | 150579476 | MCL1 |
| 5 | 178547316 | 178547559 | COL23A1 | 9 | 78031450 | 78031716 | GNAQ | 9 | 35657761 | 35657978 | RMRP | 1 | 156670850 | 156671077 | NES |
| 6 | 16439202 | 16439405 | ATXN1 | 19 | 44908835 | 44909119 | APOE | 9 | 111896835 | 111897049 | UGCG | 10 | 96586741 | 96586967 | TM9SF3 |
| 16 | 397313 | 397555 | NME4 | 11 | 6611859 | 6612080 | TAF10 | MT | 13999 | 14352 | MT-ND5 | 11 | 60951421 | 60951671 | SLC15A3 |
| 7 | 30285144 | 30285358 | ZNRF2 | 3 | 194486720 | 194486995 | ATP13A3 | MT | 13999 | 14352 | MT-ND6 | 11 | 67266479 | 67266734 | GRK2 |

| | | | |
|---|---|---|---|
| 7 | 65873216 | 65873448 | VKORC1L1 |
| 22 | 28800302 | 28800508 | XBP1 |
| 3 | 71753623 | 71753834 | EIF4E3 |
| 16 | 87602390 | 87602599 | JPH3 |
| 16 | 88643318 | 88643523 | CYBA |
| 3 | 127672468 | 127672715 | PODXL2 |
| 3 | 129315959 | 129316184 | H1FX |
| 3 | 129315959 | 129316184 | H1FX-AS1 |
| 3 | 129605142 | 129605353 | PLXND1 |
| 17 | 40122203 | 40122568 | MSL1 |
| 1 | 40861339 | 40861581 | CITED4 |
| 17 | 81833586 | 81833799 | PPP1R27 |
| 1 | 150579256 | 150579486 | MCL1 |
| 19 | 812566 | 812880 | PLPPR3 |
| 1 | 206685017 | 206685265 | MAPKAPK2 |
| 19 | 39435651 | 39435898 | RPS16 |
| 19 | 55675242 | 55675468 | EPN1 |
| 1 | 40861921 | 40862139 | CITED4 |
| 12 | 110281482 | 110281812 | ATP2A2 |
| 10 | 130157451 | 130157660 | GLRX3 |
| 11 | 1956371 | 1956593 | MRPL23 |
| 11 | 27700478 | 27700687 | BDNF |
| 11 | 27700478 | 27700687 | RP11-587D21.4 |
| 9 | 35657758 | 35657976 | RMRP |
| 11 | 47355010 | 47355242 | SPI1 |
| 11 | 67266450 | 67266707 | GRK2 |
| 9 | 78031449 | 78031706 | GNAQ |
| 9 | 128689313 | 128689551 | SET |
| MT | 11773 | 12036 | MT-ND4 |
| 14 | 49586576 | 49586831 | RN7SL1 |
| 14 | 49586576 | 49586831 | RPS29 |
| X | 32866581 | 32866790 | DMD |
| 14 | 102507034 | 102507402 | ANKRD9 |
| 12 | 124567342 | 124567773 | NCOR2 |
| X | 147912022 | 147912233 | FMR1 |
| X | 150983265 | 150983516 | HMGB3 |
| 5 | 154755387 | 154755619 | LARP1 |
| 5 | 168579347 | 168579585 | PANK3 |
| 1 | 10694447 | 10694656 | CASZ1 |
| 6 | 163414754 | 163414977 | QKI |
| 21 | 8216757 | 8216966 | CH507-513H4.1 |
| 21 | 8397569 | 8397789 | CH507-528H12.1 |
| 21 | 8400315 | 8400517 | CH507-528H12.1 |
| 1 | 35192304 | 35192731 | SFPQ |

| | | | |
|---|---|---|---|
| 1 | 150579258 | 150579477 | MCL1 |
| 1 | 212285533 | 212285876 | PPP2R5A |
| 10 | 28533263 | 28533479 | WAC |
| 6 | 163414719 | 163414989 | QKI |
| 7 | 192647 | 192892 | AC093627.12 |
| 7 | 519373 | 519713 | PDGFA |
| 10 | 96586736 | 96586964 | TM9SF3 |
| 11 | 67266446 | 67266695 | GRK2 |
| 8 | 30812507 | 30812754 | PPP2CB |
| 12 | 124567568 | 124567770 | NCOR2 |
| 9 | 111896841 | 111897057 | UGCG |
| 14 | 49586628 | 49586831 | RN7SL1 |
| 14 | 49586628 | 49586831 | RPS29 |
| 14 | 49853717 | 49853919 | Metazoa_SRP |
| 14 | 49862498 | 49862740 | RP11-596C23.6 |
| 14 | 49862498 | 49862740 | RN7SL2 |
| 15 | 63121846 | 63122120 | LACTB |
| 16 | 1445425 | 1445680 | CLCN7 |
| 16 | 1445425 | 1445680 | LA16c-390E6.5 |
| 16 | 87602393 | 87602649 | JPH3 |
| 17 | 62065227 | 62065430 | MED13 |
| 17 | 83079607 | 83079807 | METRNL |
| 18 | 62715681 | 62715988 | PHLPP1 |
| 19 | 3557347 | 3557564 | MFSD12 |
| 19 | 3557347 | 3557564 | AC005786.7 |
| 19 | 12734454 | 12734682 | C19orf43 |
| 19 | 18280177 | 18280401 | JUND |
| 19 | 18281309 | 18281611 | JUND |
| 17 | 40122214 | 40122522 | MSL1 |

| | | | |
|---|---|---|---|
| 11 | 47356699 | 47356942 | SPI1 |
| 1 | 10694451 | 10694657 | CASZ1 |
| 12 | 110281467 | 110281798 | ATP2A2 |
| 17 | 7853393 | 7853666 | KDM6B |

| | | | |
|---|---|---|---|
| 12 | 76031290 | 76031527 | PHLDA1 |
| 12 | 76031290 | 76031527 | RP11-290L1.3 |
| 12 | 124567560 | 124567769 | NCOR2 |
| 16 | 1445443 | 1445667 | CLCN7 |
| 16 | 1445443 | 1445667 | LA16c-390E6.5 |
| 16 | 9091702 | 9091953 | C16orf72 |
| 17 | 17042498 | 17042707 | MPRIP |
| 17 | 20868961 | 20869183 | CCDC144NL |
| 17 | 20868961 | 20869183 | CCDC144NL-AS1 |
| 17 | 28041767 | 28042045 | NLK |
| MT | 13992 | 14374 | MT-ND5 |
| MT | 13992 | 14374 | MT-ND6 |
| 1 | 40161378 | 40161589 | RLF |
| 21 | 45992089 | 45992320 | COL6A1 |
| 3 | 194486704 | 194486908 | ATP13A3 |
| 5 | 154755624 | 154756140 | LARP1 |

**Additional file 12**
**Table S4:** Oligonucleotide sequences

**RT-qPCR Primers**

| cDNA target | Primer | Sequence (5'-3') |
|---|---|---|
| *hCXCL10* | Forward | AAGTGGCATTCAAGGAGTACC |
| | Reverse | ATGCAGGTACAGCGTACAGT |
| *hIL-6* | Forward | CCTGAACCTTCCAAAGATGGC |
| | Reverse | CACCAGGCAAGTCTCCTCATT |
| *hCD80* | Forward | ACGCCCTGTATAACAGTGTCC |
| | Reverse | ACGCCCTGTATAACAGTGTCC |
| *hTNFa* | Forward | CTGCACTTTGGAGTGATCGG |
| | Reverse | TCAGCTTGAGGGTTTGCTAC |
| *hCD209* | Forward | AAGTAACCGCTTCACCTGGA |
| | Reverse | CTTCCTCCCCAACGTTGTTG |
| *hCD206* | Forward | GCTAAACCTACTCATGAATTACTTACAACAA |
| | Reverse | GAAGACGGTTTAGAAGGGTCCAT |
| *hb2m* | Forward | TGCCGTGTGAACCATGTGAC |
| | Reverse | CGGCATCTTCAAACCTCCAT |
| *h18S* | Forward | ACCCGTTGAACCCCATTCGTGA |
| | Reverse | GCCTCACTAAACCATCCAATCGG |
| *hGAPDH* | Forward | TCAAGGCTGAGAACGGGAAG |
| | Reverse | TCAAGGCTGAGAACGGGAAG |
| *hCSF1* | Forward | CGCTGAGGAGTGAAAGAACC |
| | Reverse | TTCAGGCTCCTCTCTCTGGA |
| *hCD36* | Forward | ACTCAGTGTTGGTGTGGTGA |
| | Reverse | ATGCAGGGCCTAGGATTTGT |
| *hMSR1* | Forward | CAGCCCTTCCCTTCAAGAGA |
| | Reverse | TCTTCGTTTCCCACTTCAGGA |
| *hYAP1* | Forward | GAACTGCTTCGGCAGGCAAT |
| | Reverse | TTGGTAACTGGCTACGCAGG |
| *hICAM1* | Forward | GGCCTCAGTCAGTGTGA |
| | Reverse | AACCCCATTCAGCGTCA |

| | | |
|---|---|---|
| *hCOL6A2* | Forward | GAGTGTGACGTCATGACCTAC |
| | Reverse | TGTACCCAATGCTCTCGGAG |
| *hCD163* | Forward | TCGTCGCATTATTCTTCTTGACTAA |
| | Reverse | GGTGGACTAAGTTCTCTCCTCTTGA |
| *hGRK2* | Forward | TCCAGCCATACATCGAAGAGA |
| | Reverse | CAAAACCGTGTGAACTTATCGC |
| *hARID1A* | Forward | GGCGGGACTAACCCATACTC |
| | Reverse | GGCCCTGTTGACCATACCC |
| *hMYC* | Forward | AAACACAAACTTGAACAGCTAC |
| | Reverse | ATTTGAGGCAGTTTACATTATGG |

**shRNA Sequence (5'-3')**

| | |
|---|---|
| Sh Con_F | CCGGTTTGGATTGAAGGGAGCTCTTCAAGAGAGAGCTCCCTTCAATCCAAACTTTTTC |
| Sh Con_R | AATTGAAAAAAGTTTGGATTGAAGGGAGCTCTCTCTTGAAGAGCTCCCTTCAATCCAAA |
| Mettl3_Sh1_F | CCGGTCACCAAGGAGACTGCTATTTCTCGAGAAATAGCAGTCTCCTTGGTGATTTTTG |
| Mettl3_Sh1_R | AATTCAAAAAGCTGCACTTCAGACGAATTATCTCGAGTAATTCGTCTGAAGTGCAGC |
| Mettl3_Sh2_F | CCGGGCAAGTATGTTCACTATGAAACTCGAGTTTCATAGTGAACATACTTGCTTTTTG |
| Mettl3_Sh2_R | AATTCAAAAAGCAAGTATGTTCACTATGAAACTCGAGTTTCATAGTGAACATACTTGC |
